# Minor pilin genes are involved in motility and natural competence in *Synechocystis* sp. PCC 6803

**DOI:** 10.1101/2020.12.15.422378

**Authors:** Sabrina Oeser, Thomas Wallner, Lenka Bučinská, Heike Bähre, Nils Schürgers, Annegret Wilde

## Abstract

Cyanobacteria synthesize type IV pili, which are known to be essential for motility, adhesion and natural competence. They consist of long flexible fibres that are primarily composed of the major pilin PilA1 in *Synechocystis* sp. PCC 6803. In addition, *Synechocystis* encodes less abundant pilin-like proteins, which are known as minor pilins. The transcription of the minor pilin genes *pilA5, pilA6* and *pilA9*-*pilA11* is inversely regulated in response to different conditions. In this study, we show that the minor pilin PilA5 is essential for natural transformation but is dispensable for motility and flocculation. In contrast, a set of minor pilins encoded by the *pilA9*-*slr2019* transcriptional unit are necessary for motility but are dispensable for natural transformation. Neither *pilA5*-*pilA6* nor *pilA9*-*slr2019* are essential for pilus assembly as mutant strains showed type IV pili on the cell surface. Microarray analysis demonstrated that the transcription levels of known and newly predicted minor pilin genes change in response to surface contact. A total of 120 genes were determined to have altered transcription between planktonic and surface growth. Among these genes, 13 are located on the pSYSM plasmid. The results of our study indicate that different minor pilins facilitate distinct pilus functions.

## Introduction

The unicellular, coccoidal cyanobacterium *Synechocystis* sp. PCC 6803 (hereinafter designated *Synechocystis*) carries two types of cell appendages – thin pili with diameters of 2-3 nm and lengths of 0.5-1 µm and thick pili with diameters of 6-8 nm and possible lengths well beyond 1 µm (Bhaya *et al*., 2000). Whereas the composition and role of thin pili has not been elucidated, genetic analyses have demonstrated that the thick pili are type IV pili (T4P) (Bhaya *et al*., 2000; Yoshihara *et al*., 2001). In *Synechocystis*, T4P are involved in cell adhesion (Nakane and Nishizaka, 2017), cell-cell aggregation (Conradi *et al*., 2019), natural transformation (Yoshihara *et al*., 2001) and twitching motility – a jerky motion on surfaces (Bhaya *et al*., 2000; Mattick, 2002).

The core components of T4P are widely conserved across different prokaryotic phyla (Denise *et al*., 2019). T4P subunits of *Synechocystis* have been identified by homology to *Pseudomonas aeruginosa* (hereinafter designated *Pseudomonas*) and *Myxococcus xanthus* (hereinafter designated *Myxococcus*) pilus proteins (Bhaya *et al*., 1999; Bhaya *et al*., 2000; Yoshihara *et al*., 2001). PilQ is an integral outer membrane secretin and enables the emergence of the extending pilus fibre. The pilus platform is formed by the integral inner membrane protein PilC and traffic ATPases that facilitate the extension (PilB) or retraction (PilT) of the fibre. The PilMNO proteins align the pilus components throughout the periplasm. The most abundant structural protein of the pilus fibre is the major pilin PilA. PilA prepilins are inserted into the inner membrane, where they are processed by the bifunctional leader peptidase/methylase PilD and are subsequently inserted into the growing fibre (reviewed in Pelicic, 2008). In addition to the highly abundant major pilin, T4P-forming bacteria usually encode several, less abundant minor pilins with different functions. Minor pilins from *Pseudomonas* and *Myxococcus* can prime pilus assembly and influence pilus retraction (Nguyen *et al*., 2015; Treuner-Lange *et al*., 2020). Incorporation of minor pilins into the pilus filament was shown for *Pseudomonas* (Giltner *et al*., 2010) and *Vibrio cholerae* (Ng *et al*., 2016; Ellison *et al*., 2018), and location at the pilus tip was demonstrated for *Myxococcus* (Treuner-Lange *et al*., 2020). Furthermore, specific minor pilins are involved in pilus functions, such as adhesion or natural transformation, in various bacteria (Wolfgang *et al*., 1999; Winther-Larsen *et al*., 2001; Ng *et al*., 2016), including the cyanobacterium *Synechococcus elongatus* PCC 7942 (Taton *et al*., 2020). The involvement of minor pilins in DNA binding at the T4P tip was demonstrated for *Vibrio cholerae* (Ellison *et al*., 2018). Nevertheless, the role and mechanism of function of many minor pilins have not been completely elucidated to date.

*Synechocystis* encodes at least nine minor pilins, PilA2-PilA11. PilA3 was misidentified and is now considered a Tat protein that has characteristics of both TatA and TatB of the Tat protein secretion system (Aldridge *et al*., 2008). All major and minor pilins contain a conserved PilD cleavage site (G|XXXE) followed by a hydrophobic stretch (Linhartová *et al*., 2014). Most minor pilin genes of *Synechocystis* are organized in operons (Fig. 1A). Based on the transcriptomic data by Kopf *et al*. (2014), the transcriptional unit (TU) *TU763* comprises a polycistronic mRNA encoding the minor pilins PilA9, PilA10 and PilA11 together with the open reading frames *slr2018* and *slr2019*. The minor pilin genes *pilA5* and *pilA6* constitute *TU2300*. The downstream *pilA7* and *pilA8* genes are most likely transcribed from a different promotor. To date, knowledge regarding minor pilin function in *Synechocystis* is scarce. Deletion mutants of *pilA1, pilA10* and *pilA11* are non-motile on agar plates (Yoshihara *et al*., 2001; Bhaya *et al*., 2001). Furthermore, it was demonstrated that deletion of the whole *pilA9-slr2019* genomic region not only leads to loss of motility (Wallner *et al*., 2020) but also impairs flocculation (Conradi *et al*., 2019), which describes the aggregation of cells into floating assemblages in liquid culture. Hu *et al*. (2018) showed, that the asRNA PilR negatively regulates the amount of *pilA11* mRNA and the amount of the corresponding protein. Thus, overexpression of the asRNA led to inhibition of motility in this study. In another study it was shown that PilA4 is located within the pilus fibre if the major pilin PilA1 is present (Cengic *et al*., 2018).

**Figure 1:**
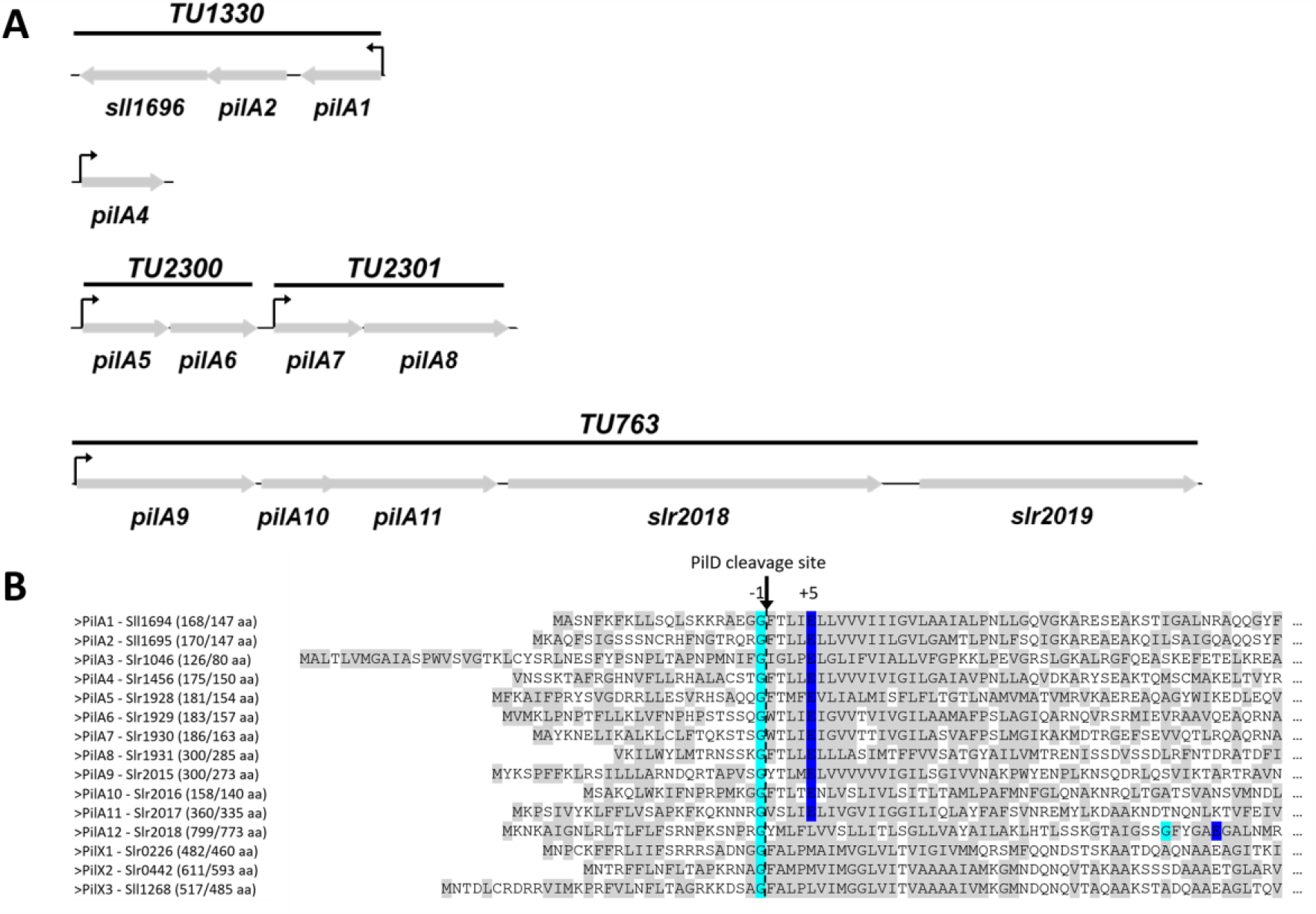
Major and minor pilins of *Synechocystis*. (**A**) Schematic representation of *Synechocystis* gene clusters encoding PilA1 and minor pilins. PilX homologues *slr0226, slr0442* and *sll1268* are not encoded in an operon or in proximity to other pilins. (**B**) Multiple alignment of the major pilin PilA1 and the proposed minor pilins. The potential cleavage site by PilD is marked with the dotted line and an arrow. The conserved glycine and glutamate at positions −1 and +5 in relation to the cleavage site are highlighted in turquois and blue, respectively. The newly proposed minor pilins PilA12 (Slr2018), Slr0226, Slr0442 and Sll1268 contain a hydrophobic amino acid substitution at position +5. The length of the full-length protein/mature protein is given next to the protein name in brackets. Hydrophobic amino acid residues are highlighted in grey. PilA3 was previously shown to belong to the Tat system and is no minor pilin (see introduction).

As in many other bacteria, a functional T4P system is crucial for natural competence of *Synechocystis* (Yoshihara *et al*., 2001). The uptake mechanism has not been fully elucidated, but it is known that DNA is processed to a single-stranded form during DNA uptake (Barten and Lill, 1995) and that ComA (Slr0197) (Yura *et al*., 1999; Yoshihara *et al*., 2001) and ComF (Slr0388) (Nakasugi *et al*., 2006) homologues are important for competence of *Synechocystis* cells.

T4P are also involved in surface attachment, which is an important part in the life of prokaryotes, enabling biofilm formation. Thus, biofilms confer a fitness advantage over planktonic solitary cells by protection from diverse environmental stresses, better nutrient availability and drug and predator resistance (Laventie and Jenal, 2020). Surface recognition by different appendages, including T4P, triggers signal transduction cascades involving second messengers, quorum-sensing systems, two-component systems and small regulatory RNAs (Laventie and Jenal, 2020). For *Pseudomonas*, which possesses T4P and a single polar flagellum, it was suggested that pilus tension leads to conformational change of the T4P, which subsequently leads to 3’,5’-cyclic adenosine monophosphate (cAMP) production and activation of virulence programmes (Persat *et al*., 2015). Additionally, surface contact of the flagellum (Laventie *et al*., 2019), as well as surface adhesion of the minor pilin PilY1 and a functional T4P (Rodesney *et al*., 2017), trigger the production of c-di-GMP. Increased c-di-GMP levels promote surface acclimation by inhibiting flagella motility, thereby promoting faster T4P-dependent surface attachment and production of extracellular polymeric substances (EPS) (Rodesney *et al*., 2017).

For *Synechocystis*, little is known regarding surface sensing and the transition from planktonic to sessile lifestyle. However, *Synechocystis* encodes cAMP- and c-di-GMP-dependent signalling components (Ohmori and Okamoto, 2004; Agostoni *et al*., 2013). Intracellular c-di-GMP levels can alter cellular deposition (Agostoni *et al*., 2016) and control flocculation (Conradi *et al*., 2019) and motility (Savakis *et al*., 2012). Motility is also dependent on cAMP, as inactivation of the adenylate cyclase gene *cya1* leads to inhibition of phototaxis on agar plates (Terauchi and Ohmori, 1999).

In this study, we investigate the role played by minor pilins in *Synechocystis* and show that the minor pilin PilA5 is involved in natural competence. Furthermore, our data imply that among others, genes encoding minor pilins, cell envelope structures and genes located on the plasmid pSYSM are major targets of a putative surface sensing system. We also investigated changes in second messenger production upon surface contact.

## Results

### Minor pilins play roles in motility and surface attachment

In addition to the major pilin PilA1, *Synechocystis* encodes at least nine different minor pilins (Linhartová *et al*., 2014). Considering that the two TUs encoding PilA5-PilA6 and PilA9-Slr2019 are inversely regulated in response to blue light (Wallner *et al*., 2020), we questioned if these minor pilins are involved in different pilus functions. In contrast to the Δ*pilA9*-*slr2019* mutant, which is non-motile, a Δ*pilA5*-*pilA6* mutant strain exhibited a wild-type (WT) phototaxis response (Wallner *et al*., 2020). To identify other phenotypes, we first tested whether the Δ*pilA5*-*pilA6* strain is impaired in flocculation. Fig. 2 shows that the Δ*pilA5*-*pilA6* strain exhibits a WT flocculation response and clearly differs from the non-flocculating and non-motile Δ*pilA9*-*slr2019* and Δ*hfq* mutant strains (Conradi *et al*., 2019; Wallner *et al*., 2020). The Δ*hfq* strain was employed as a control because deletion of the *hfq* gene, which encodes the cyanobacterial homologue of the RNA chaperone Hfq, that was shown to bind to the pilus base (Schuergers *et al*., 2014), leads to non-motile and non-piliated cells (Dienst *et al*., 2008). To determine whether the attachment of T4P to surfaces and their dynamics is impaired in the Δ*pilA5*-*pilA6* and Δ*pilA9*-*slr2019* strains, we used fluorescent beads and monitored the retraction of the beads to the cell surface, as described by Nakane & Nishizaka (2017). To that end, a glass surface was coated with 4% collodion; thus, cells were not able to move, though they were still able to assemble and retract T4P. Our bead assays suggest that the WT pili attach to nearby beads and transport them towards the cell (Fig. 3A and video S1). Interestingly, Δ*pilA9-slr2019* mutant cells were not able to adhere to the collodion surface. However, in some cases, single cells were trapped on the glass slide, but they were not able to attach to the beads (Fig. 3B and video S2). In contrast, Δ*pilA5*-*pilA6* mutant cells behave similar to the WT and retract beads towards the cell (Fig. 3C and video S3).

**Figure 2:**
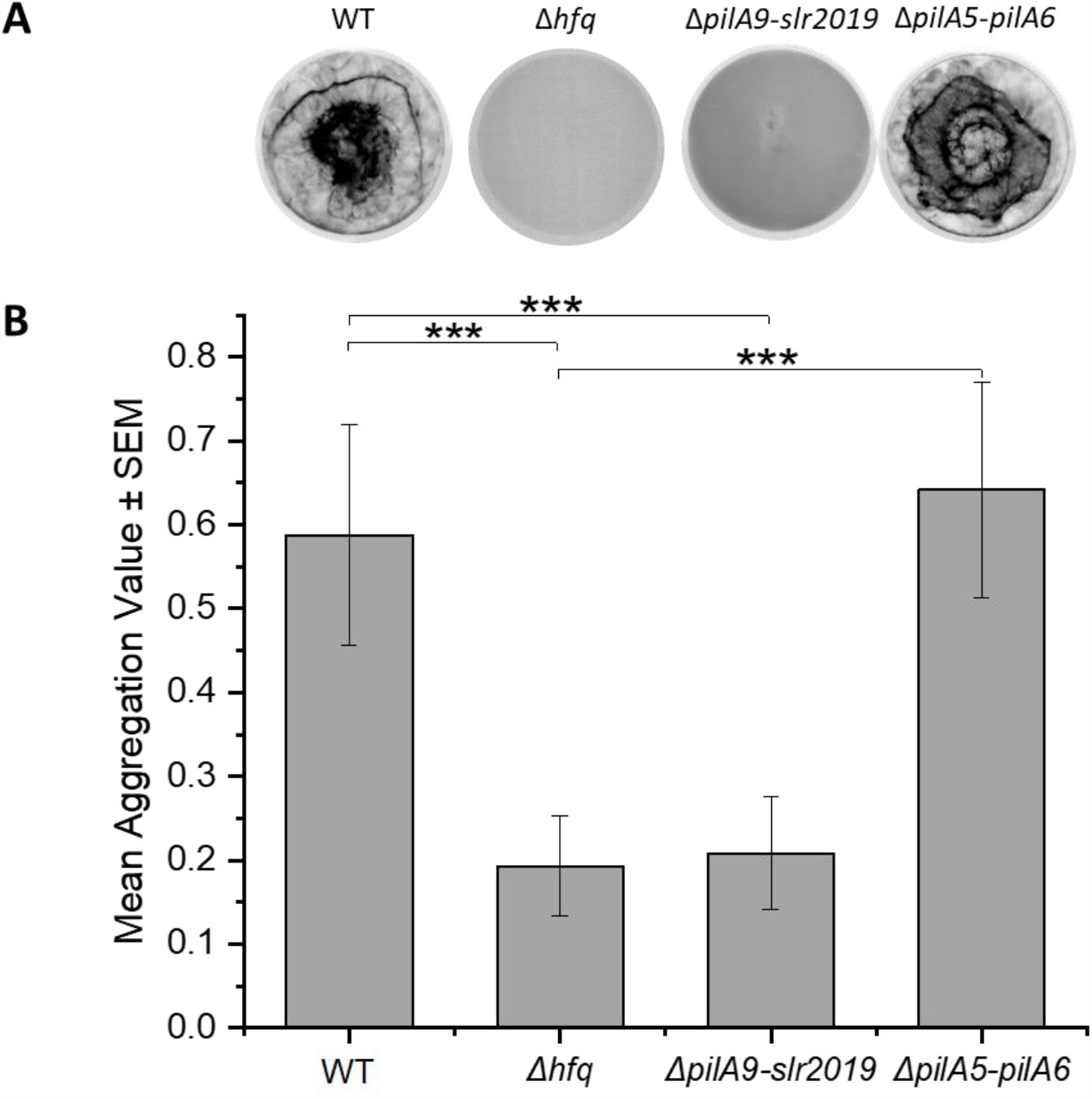
Involvement of minor pilins in flocculation. **(A)** Representative flocculation assays for WT *Synechocystis* cells and deletion mutants Δ*hfq*, Δ*pilA9-slr2019* and Δ*pilA5-pilA6*. Chlorophyll fluorescence is shown in inverted greyscale; thus, areas with more chlorophyll appear darker. **(B)** Mean aggregation values ± standard errors of WT (n=19), Δ*pilA5-pilA6* (n=19), Δ*hfq* (n=12) and Δ*pilA9-slr2019* (n=7). Three asterisks indicate statistical significance with *p*-values ≤ 0.001, calculated via Student’s t-test.

**Figure 3:**
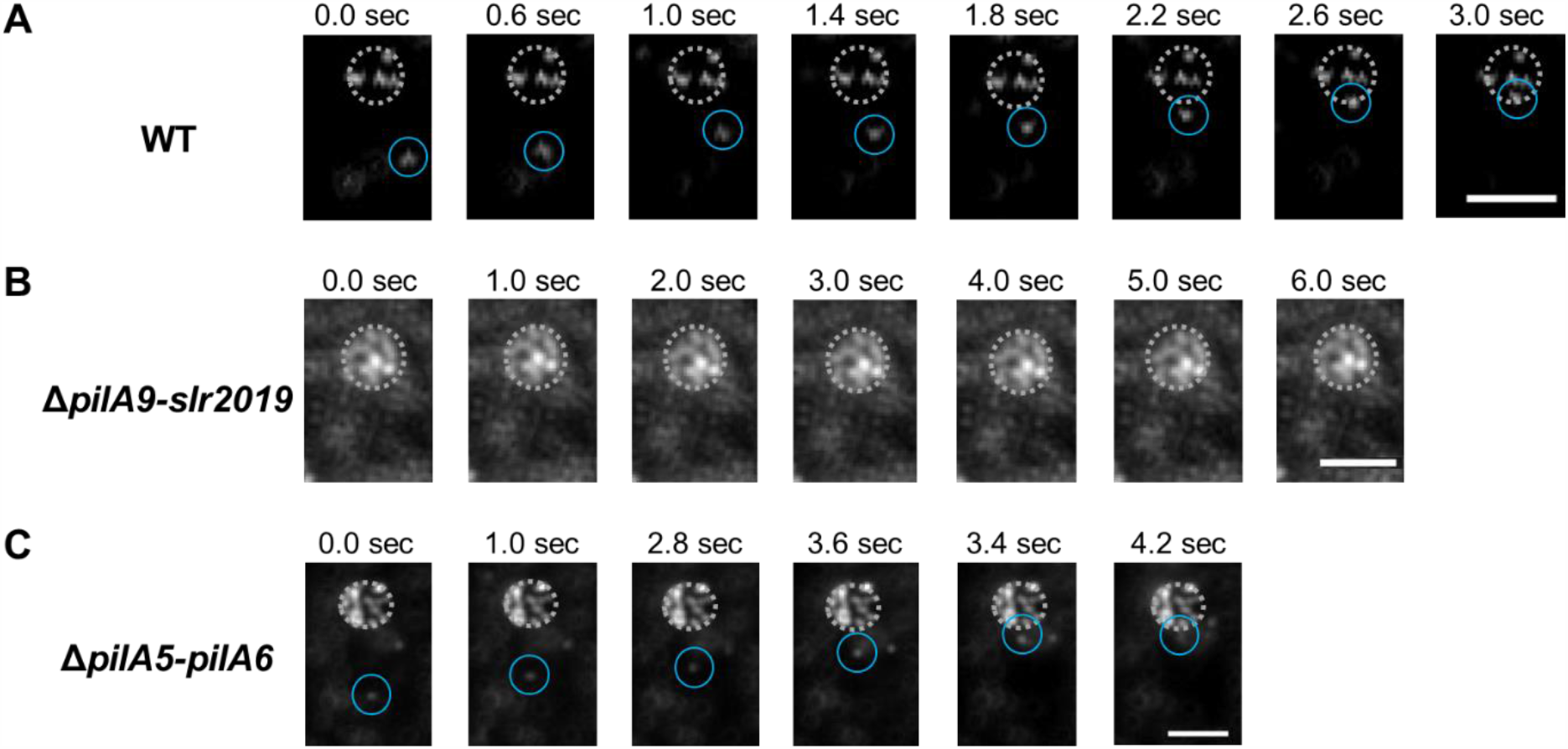
Pili can bind and retract fluorescent beads. Picture series of WT **(A)**, Δ*pilA9-slr2019* **(B)** and Δ*pilA5-pilA6* **(C)** cells binding fluorescent beads in the medium. Cells are immobilized on a collodion surface and indicated by dotted circles. Representative pilus-bound beads are indicated by a blue circle. Scale bars: 3 µm. Original videos can be found in the supplementary data (videos S1-S3).

These results indicate that the lack of the *pilA9*-*slr2019* gene cluster causes a defect in T4P function related to motility and attachment to biotic and abiotic substances, whereas the minor pilins PilA5 and PilA6 are dispensable for these processes.

### Role of minor pilins in natural competence

T4P are also known to be important for natural competence (reviewed in Piepenbrink, 2019). Therefore, we investigated the ability of the minor pilin mutants described above to be transformed by exogenous DNA. We used a plasmid that enables integration via homologous recombination of a streptomycin resistance cassette into the chromosomal region of the gene encoding the small RNA PsrR1 (Georg *et al*., 2014). The Δ*pilA9*-*slr2019* mutant strain was transformable with a transformation efficiency similar to that of the WT (Fig. 4A). The Δ*hfq* strain was employed as a negative control, as this strain is not transformable due to the lack of T4P (Dienst *et al*., 2008). Notably, the Δ*pilA5*-*pilA6* mutant strain was not transformable (Fig. 4A). To discriminate the function of PilA5 and PilA6, we complemented the Δ*pilA5*-*pilA6* strain with plasmids for the expression of *pilA5, pilA6* or the whole *pilA5*-*pilA6* operon (Fig. S1). To each coding sequence, the putative native promoter region (extending 450 bp upstream of the *pilA5* start codon) was fused. Though the standard deviations of transformation efficiency were very high in these experiments, it is clear that only in the strains that contained the *pilA5* gene, either the *pilA5* gene alone or in combination with *pilA6*, streptomycin-resistant clones were detectable (Fig. 4B). Exconjugants containing the plasmid with only the *pilA6* gene were not transformable (Fig. 4B). Therefore, we conclude that the minor pilin PilA5 is essential for DNA uptake.

**Figure 4:**
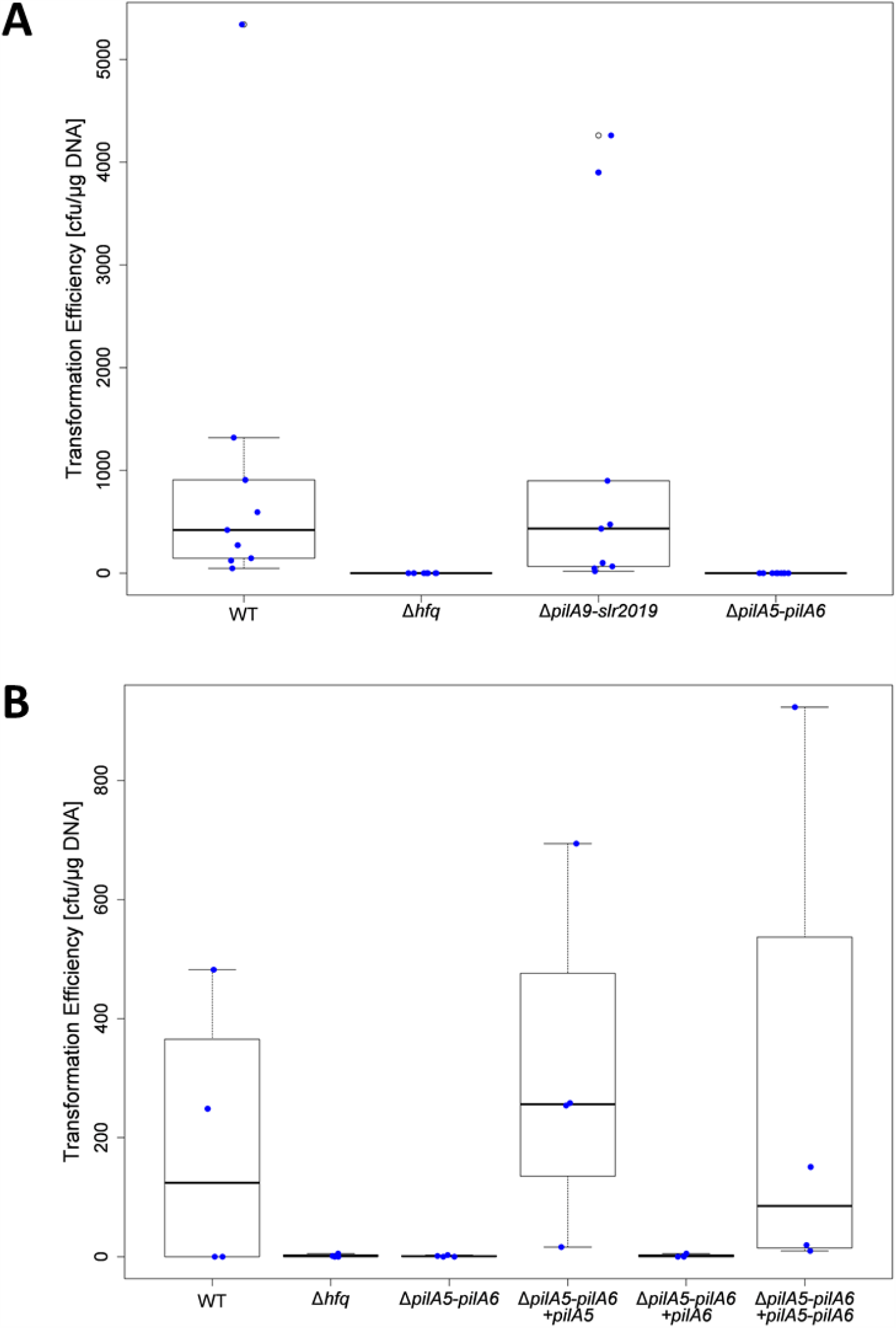
Transformation efficiency of different minor pilin mutants. **(A)** The transformation efficiency of WT, Δ*hfq*, Δ*pilA9*-*slr2019* and Δ*pilA5*-*pilA6* cells was determined (n=9). The deletion mutant Δ*hfq* without T4P was used as a negative control. Transformation was performed with different amounts of DNA ranging from 500 ng to 1 µg. **(B)** The transformation efficiency of WT, Δ*pilA5*-*pilA6*/+*pilA5*, Δ*pilA5*-*pilA6*/+*pilA6 and* Δ*pilA5*-*pilA6*/+*pilA5-pilA6* cells was determined (n=4). Natural transformation in the deletion mutant was restored by complementation with *pilA5* and *pilA5*-*pilA6*.

### Minor pilin mutants assemble thick pili on the surface of Synechocystis *cells*

To determine whether the lack of specific pilus functions is related to a defect in pilus assembly, we examined negatively stained cells by transmission electron microscopy (TEM). Notably, we were able to detect thick pili in the WT and the two minor pilin operon deletion mutants (Fig. 5). In all strains, we measured a diameter of the thick pili between 5.5 and 9 nm (Fig. S2), which is comparable to the 6 to 8 nm described by Bhaya *et al*. (2000). This result suggests that all minor pilin deletion mutants can assemble T4P. In addition, *Synechocystis* also possesses thin pili with unknown molecular composition and function. All mutant strains analysed in our study were shown to produce this kind of appendage (Fig. 5). Therefore, the minor pilins PilA5, PilA6, PilA9, PilA10 and PilA11 and proteins Slr2018 and Slr2019 are not involved in general formation of thin or thick pili. Notably, we could also visualize thin pili on the Δ*hfq* strain (Fig. 5F). This is in contrast to previous observations by Dienst *et al*. (2008), which described the mutant non-piliated.

**Figure 5:**
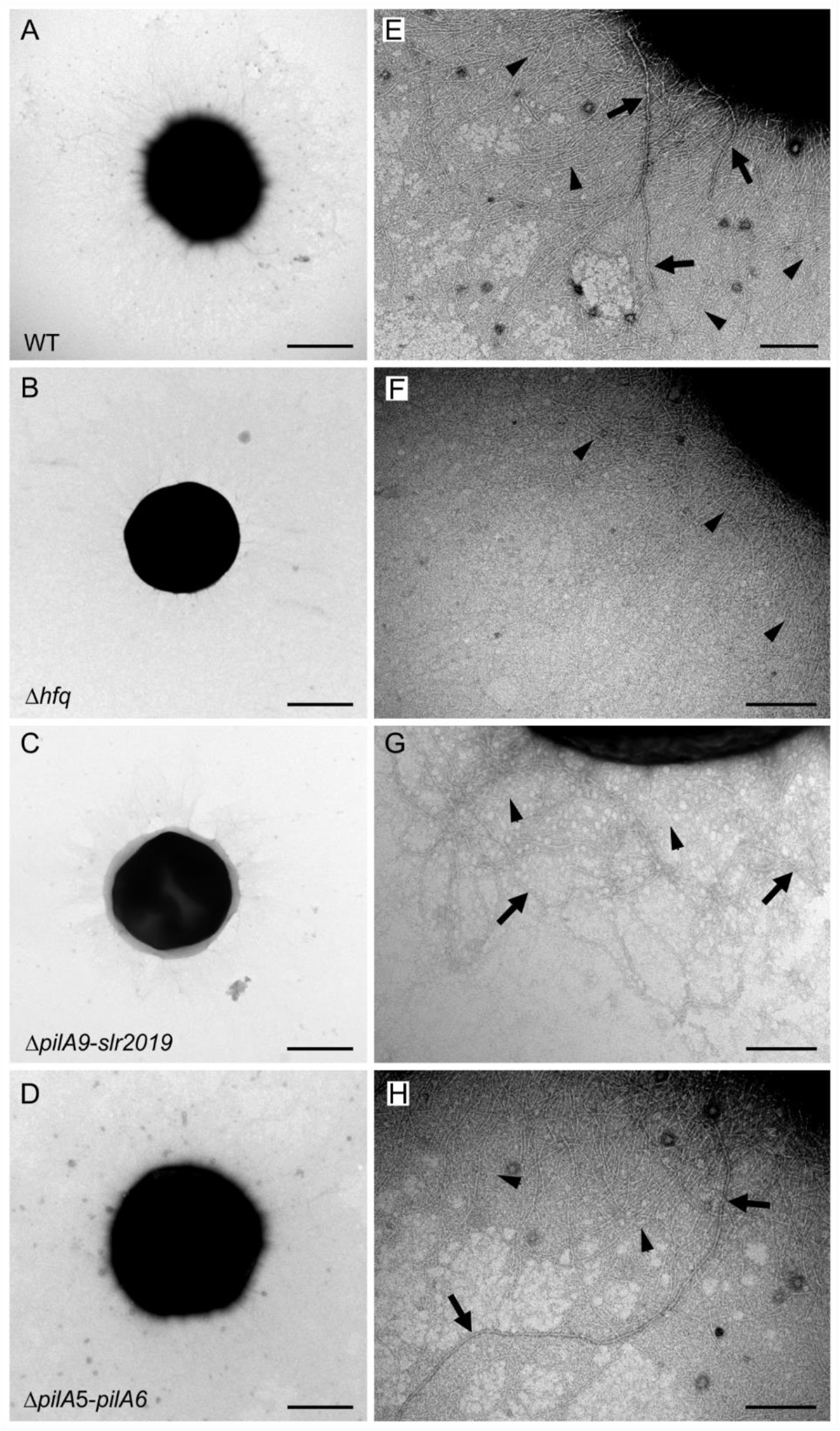
Electron micrographs of different negatively stained *Synechocystis* strains. WT **(A, E)**, T4P lacking deletion mutant Δ*hfq* **(B, F)** and deletion mutants Δ*pilA9-slr2019* **(C, G)** and Δ*pilA5-pilA6* **(D, H)** were negatively stained with 1% uranyl acetate. Shown are whole cells **(A-D)** and close-up views **(E-H)**. Scale bars are 1 µm in pictures A and B; 2 µm in C and D and 200 nm in E and H. Arrows depict representative thick pili, and arrowheads representative thin pili.

In summary, minor pilin deletion mutants assemble thick pili, and the presence of thin pili was not impaired in any of the constructed mutants. Therefore, loss of motility and flocculation in Δ*pilA9*-*slr2019* and impaired natural competence in Δ*pilA5*-*pilA6* did not correlate with loss of T4P or thin pili.

### Transcriptional changes in response to surface acclimation

As shown above, pili and minor pilins are involved in attachment and surface motility. Therefore, we attempted to determine if their transcription is regulated upon surface contact and whether *Synechocystis* is able to respond to surface contact in general. In *Pseudomonas*, mechanosensing of a surface with T4P leads to upregulation of a cAMP-dependent operon with significant transcriptional changes within 3 h of surface contact (Persat *et al*., 2015). To evaluate the acclimation capacity of the slower growing *Synechocystis*, researchers usually analyse gene expression changes within 1 to 24 h after transfer to new conditions (Hernández-Prieto *et al*., 2016). Therefore, we analysed genome-wide transcription changes between *Synechocystis* planktonic cell cultures and cells on a surface (sessile culture) after 4 and 8 h of surface incubation. We employed a microarray design that enables identification of all coding and non-coding RNA transcripts identified by RNA-seq (Mitschke *et al*., 2011). We defined a log_2_ fold change (FC) ≥ |−0.8| and an adjusted *p*-value < 0.05 as the threshold for a transcriptional change (comparable to Wallner et al., 2020). Based on these criteria, the transcript accumulation of 120 genes changed after surface acclimation (Fig. 6, Fig. S3 (with labels), Tables 1 and 2). As depicted in the volcano plot in blue, genes on the 120 kbp pSYSM plasmid showed the highest differential transcription upon surface contact (Fig. 6, Fig. S3). The transcript levels of 10 genes on the pSYSM plasmid were downregulated, and three were upregulated, together representing approximately 10 % of all annotated genes on this plasmid (Kaneko *et al*., 2003). One of the most upregulated TU at both time points after transfer of the cells to a surface is located on the pSYSM plasmid and encodes the two hypothetical proteins Slr5087 and Slr5088.

**Table 1:**
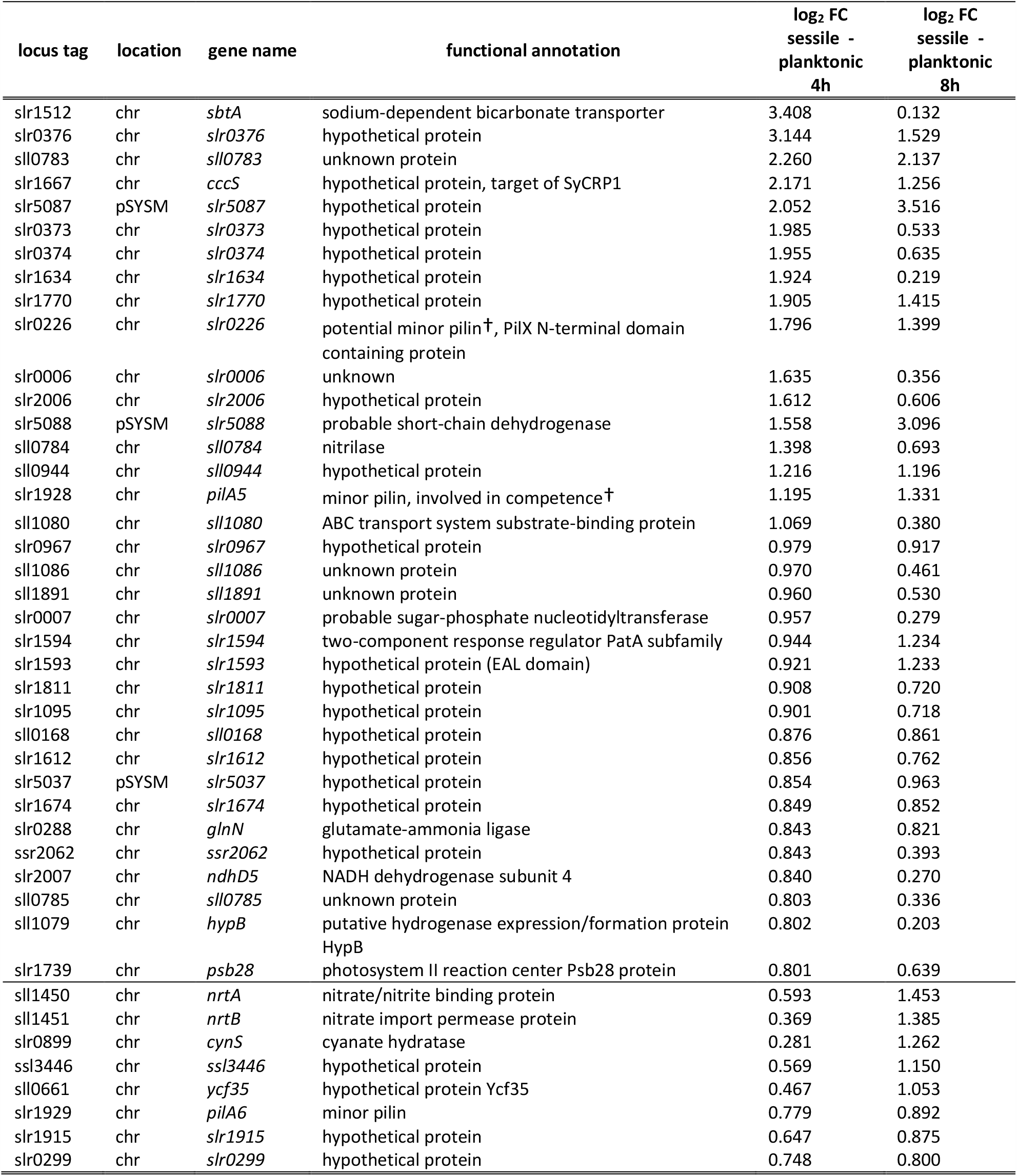
Upregulated genes in sessile conditions compared to planktonic conditions. Listed are differentially transcribed genes after growth for 4 or 8 h either under planktonic or sessile conditions. Shown are only protein encoding genes. Hits underneath the horizontal solid line indicate genes that were only considered significant after 8 h. Fold changes (FC) were considered significant with a log_2_ FC ≤ −0.8 or ≥ +0.8 and adjusted p-value < 0.05. Adjusted p-values were calculated using the Benjamini-Hochberg test. Functional annotation was derived from the CyanoBase and UniProt databases (Jan 2020). †shown in this study. chr – chromosome

| locus tag | location | gene name | functional annotation | $\log_2$ FC<br>sessile -<br>planktonic<br>4h | $\log_2$ FC<br>sessile -<br>planktonic<br>8h |
| --- | --- | --- | --- | --- | --- |
| slr1512 | chr | *sbtA* | sodium-dependent bicarbonate transporter | 3.408 | 0.132 |
| slr0376 | chr | *slr0376* | hypothetical protein | 3.144 | 1.529 |
| sll0783 | chr | *sll0783* | unknown protein | 2.260 | 2.137 |
| slr1667 | chr | *cccS* | hypothetical protein, target of SyCRP1 | 2.171 | 1.256 |
| slr5087 | pSYSM | *slr5087* | hypothetical protein | 2.052 | 3.516 |
| slr0373 | chr | *slr0373* | hypothetical protein | 1.985 | 0.533 |
| slr0374 | chr | *slr0374* | hypothetical protein | 1.955 | 0.635 |
| slr1634 | chr | *slr1634* | hypothetical protein | 1.924 | 0.219 |
| slr1770 | chr | *slr1770* | hypothetical protein | 1.905 | 1.415 |
| slr0226 | chr | *slr0226* | potential minor pilin†, PilX N-terminal domain containing protein | 1.796 | 1.399 |
| slr0006 | chr | *slr0006* | unknown | 1.635 | 0.356 |
| slr2006 | chr | *slr2006* | hypothetical protein | 1.612 | 0.606 |
| slr5088 | pSYSM | *slr5088* | probable short-chain dehydrogenase | 1.558 | 3.096 |
| sll0784 | chr | *sll0784* | nitrilase | 1.398 | 0.693 |
| sll0944 | chr | *sll0944* | hypothetical protein | 1.216 | 1.196 |
| slr1928 | chr | *pilA5* | minor pilin, involved in competence† | 1.195 | 1.331 |
| sll1080 | chr | *sll1080* | ABC transport system substrate-binding protein | 1.069 | 0.380 |
| slr0967 | chr | *slr0967* | hypothetical protein | 0.979 | 0.917 |
| sll1086 | chr | *sll1086* | unknown protein | 0.970 | 0.461 |
| sll1891 | chr | *sll1891* | unknown protein | 0.960 | 0.530 |
| slr0007 | chr | *slr0007* | probable sugar-phosphate nucleotidyltransferase | 0.957 | 0.279 |
| slr1594 | chr | *slr1594* | two-component response regulator PatA subfamily | 0.944 | 1.234 |
| slr1593 | chr | *slr1593* | hypothetical protein (EAL domain) | 0.921 | 1.233 |
| slr1811 | chr | *slr1811* | hypothetical protein | 0.908 | 0.720 |
| slr1095 | chr | *slr1095* | hypothetical protein | 0.901 | 0.718 |
| sll0168 | chr | *sll0168* | hypothetical protein | 0.876 | 0.861 |
| slr1612 | chr | *slr1612* | hypothetical protein | 0.856 | 0.762 |
| slr5037 | pSYSM | *slr5037* | hypothetical protein | 0.854 | 0.963 |
| slr1674 | chr | *slr1674* | hypothetical protein | 0.849 | 0.852 |
| slr0288 | chr | *glnN* | glutamate-ammonia ligase | 0.843 | 0.821 |
| ssr2062 | chr | *ssr2062* | hypothetical protein | 0.843 | 0.393 |
| slr2007 | chr | *ndhD5* | NADH dehydrogenase subunit 4 | 0.840 | 0.270 |
| sll0785 | chr | *sll0785* | unknown protein | 0.803 | 0.336 |
| sll1079 | chr | *hypB* | putative hydrogenase expression/formation protein HypB | 0.802 | 0.203 |
| slr1739 | chr | *psb28* | photosystem II reaction center Psb28 protein | 0.801 | 0.639 |
| sll1450 | chr | *nrtA* | nitrate/nitrite binding protein | 0.593 | 1.453 |
| sll1451 | chr | *nrtB* | nitrate import permease protein | 0.369 | 1.385 |
| slr0899 | chr | *cynS* | cyanate hydratase | 0.281 | 1.262 |
| ssl3446 | chr | *ssl3446* | hypothetical protein | 0.569 | 1.150 |
| sll0661 | chr | *ycf35* | hypothetical protein Ycf35 | 0.467 | 1.053 |
| slr1929 | chr | *pilA6* | minor pilin | 0.779 | 0.892 |
| slr1915 | chr | *slr1915* | hypothetical protein | 0.647 | 0.875 |
| slr0299 | chr | *slr0299* | hypothetical protein | 0.748 | 0.800 |

**Table 2:**
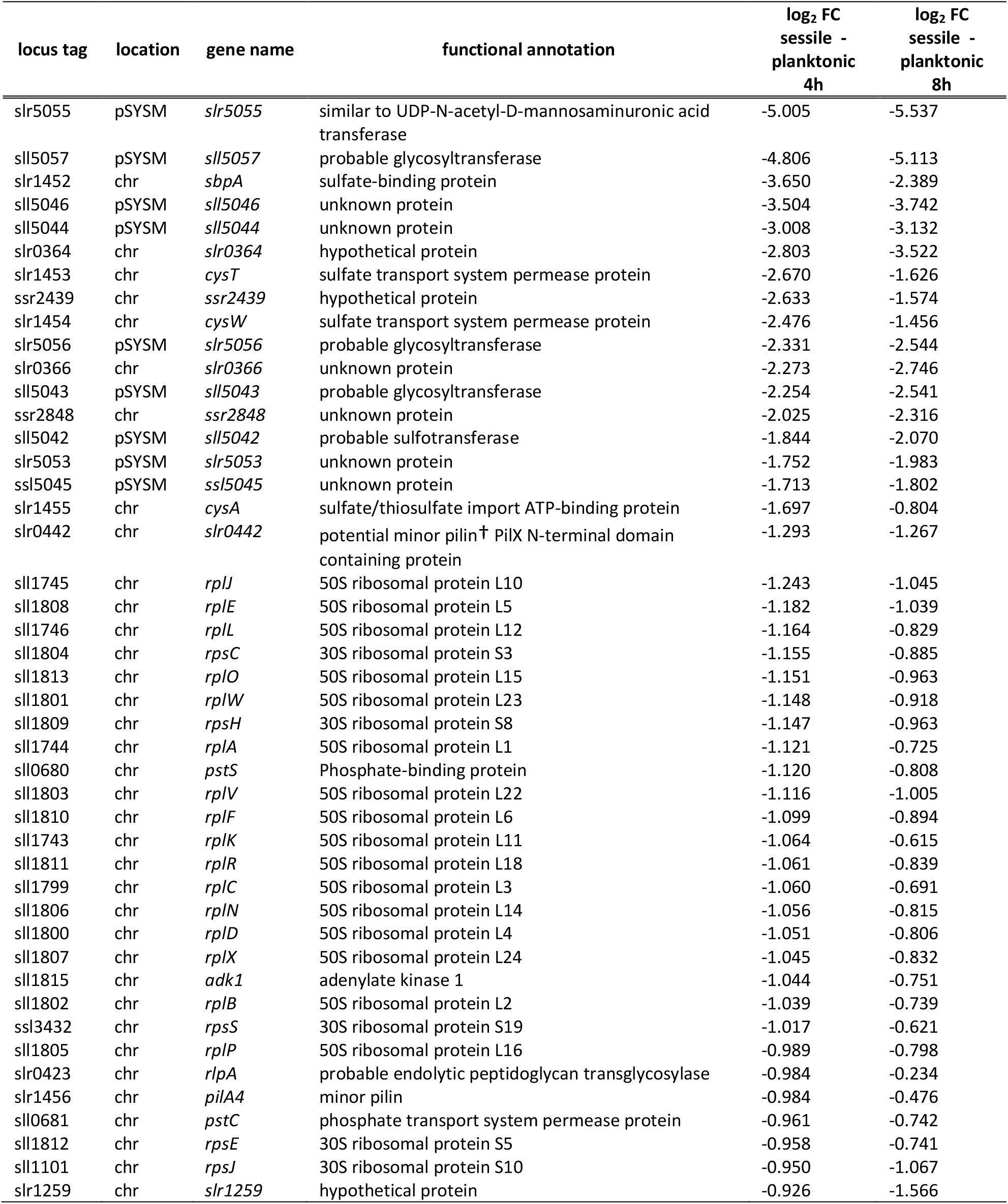
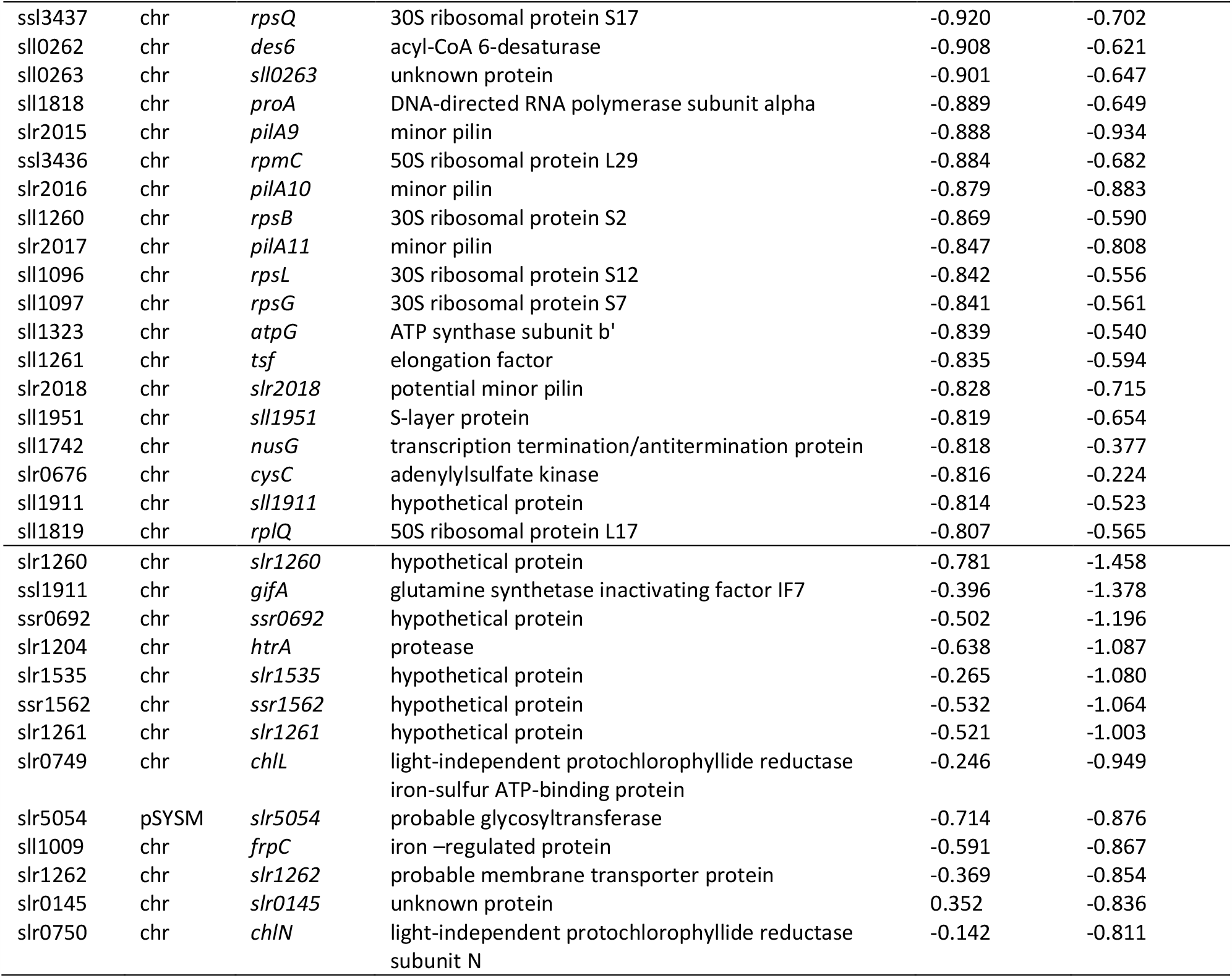
Downregulated genes in sessile conditions compared to planktonic conditions. Listed are differentially transcribed genes after growth for 4 or 8 h either under planktonic or sessile conditions. Shown are only protein encoding genes. Hits underneath the horizontal solid line indicate genes that were only considered significant after 8 h. Fold changes (FC) were considered significant with a log_2_ FC ≤ −0.8 or ≥ +0.8 and adjusted p-value < 0.05. Adjusted p-values were calculated using the Benjamini-Hochberg test. Functional annotation was derived from the CyanoBase and UniProt databases (Jan 2020). †shown in this study. chr – chromosome

**Figure 6:**
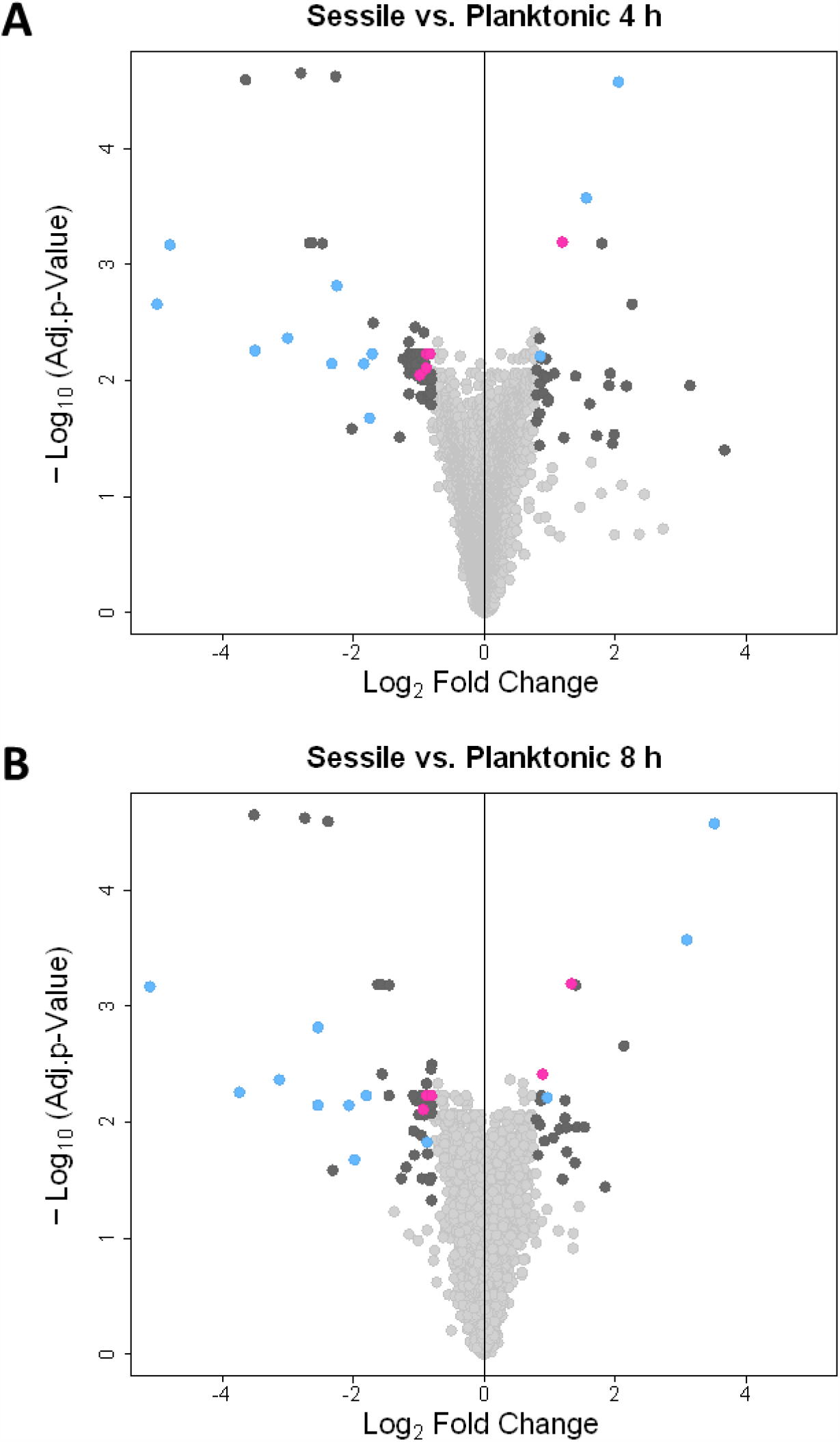
Volcano plots of differentially transcribed genes after incubation for 4 h (A) or 8 h (B) either in planktonic or sessile conditions. Negative log_10_-transformed adjusted p-values are plotted against the log_2_ fold changes of the data derived from a transcriptome microarray (corresponding data in Tables 1 and 2). Fold changes (FC) were considered significant with a log_2_ FC ≤ −0.8 or ≥ +0.8 and adjusted p-value < 0.05 and are represented by dark grey points. Adjusted p-values were calculated using the Benjamini-Hochberg test. FCs were calculated for sessile versus planktonic conditions; thus, positive log_2_ FC represents upregulation under sessile conditions (right side of plot), and negative log_2_ FC represents downregulation under sessile conditions (left side of plot). Genes located on the pSYSM plasmid are depicted in blue, and minor pilin genes are depicted in pink. For a detailed view and labelling, see Figure S3.

Notably, the mRNA of the minor pilin gene operon *pilA5*-*pilA6* accumulated under sessile conditions, whereas the genes *pilA9, pilA10, pilA11* and *slr2018* showed higher transcript accumulation during the planktonic lifestyle. The downstream *slr2019* open reading frame does not appear to be differentially transcribed based on our significance criteria, implying that this gene may not be part of *TU763* or that there is posttranscriptional gene regulation. Interestingly, the most strongly upregulated mRNA after 4 h of incubation on an agar plate encodes the high affinity bicarbonate uptake system SbtA (Shibata *et al*., 2002), implying that cells suffer from inorganic carbon (C_i_) limitation during surface acclimation. However, after 8 h on a surface, the mRNA from this gene no longer accumulates (Table 1). Three other genes (*slr0373, slr0374*, and *slr0376*), which are known to accumulate under several stress conditions (Singh and Sherman, 2002), including low C_i_ conditions (Eisenhut *et al*., 2007), show a similar response (Table 1). Further mRNAs that were previously found to accumulate under C_i_ limitation, including the hypothetical genes *slr1634, slr1770, slr0226, slr0006*, and *slr2006* (Orf *et al*., 2016), behave similar to the *sbtA* transcript in our study (Table 1). Notably, we did not identify components of the inducible bicarbonate/CO_2_ uptake systems (e.g., *ndhF3/D3, cmp*, and *cupA*), which are also typical low C_i_-responsive transcripts.

Furthermore, most upregulated genes 4 h after transfer to agar plates are *sll0783*, which is involved in polyhydroxybutyrate synthesis (Schlebusch and Forchhammer, 2010), and *slr1667*, a component of a putative chaperone usher pathway (Schuergers and Wilde, 2015) (Table 1). Loci that show increasing transcript accumulation in sessile cultures over the 8-h acclimation period are either located on pSYSM or are involved in nitrate or cyanate metabolic processes (e.g. *nrtAB* and *cynS*) or pilus function (such as the minor pilin genes *pilA5* and *pilA6*) (Table 1). A further operon with a similar transcription profile is *slr1593*/*slr1594*, which encodes an EAL domain protein and a CheY-like response regulator with unknown functions (Table 1).

The most downregulated genes in response to surface exposure are again transcribed from the pSYSM plasmid; among these genes, several encode putative glycosyl transferases (Table 2). Other genes that show higher transcription in planktonic cultures than in sessile cultures encode sulphate and phosphate transport systems, proteins related to transcription/translation and hypothetical proteins (Table 2). The transcription levels of some genes remain low or decrease further in surface-grown cells after 8 h. These genes encode pSYSM-based transcripts, the polycistronic mRNA for minor pilins (*pilA9-pilA11*), *gifA*, which encodes the glutamine synthetase inactivating factor IF7, enzymes of chlorophyll biosynthesis (*chlL, chlN*), a Deg protease (*htrA*) and a number of hypothetical proteins (Table 2).

In addition to the transcriptional changes in mRNAs, we detected 16 non-coding RNAs (ncRNA) which were upregulated after 4h on surface (e.g. PmgR1 (*ncr0700*) and IsaR1 (*ncl1600*)) and 6 ncRNAs which were downregulated under sessile conditions (e.g. SyR6 (*ncl0880*) and CsiR1 (*ncr038*) (Supplementary Data Table). These ncRNAs are known to be involved in acclimation to nutrient starvation (Kopf *et al*., 2014; De Porcellinis *et al*., 2016; Giner-Lamia *et al*., 2017; Rübsam *et al*., 2018).

Generally, differences in transcript accumulation between the two time points indicate a dynamic response of the cells with short upregulation of several stress-related functions and a more stable or even increasing expression change (e.g., transcription from the pSYSM plasmid), which suggests a more specific function of these genes in surface acclimation.

Additionally, we performed Northern blot hybridizations with RNA probes against *slr5055* and *pilA9* to verify the results of the microarray analysis (Fig. 7). In keeping with the microarray results, *slr5055* mRNA showed a strong signal under planktonic conditions and was not detectable under sessile conditions after 4 and 8 h of surface contact (Fig. 7A). Additionally, *pilA9* mRNA also accumulated under planktonic conditions and exhibited lower signal intensity under sessile conditions at 10 min, 1, 4 and 8 h after surface contact (Fig. 7B).

**Figure 7:**
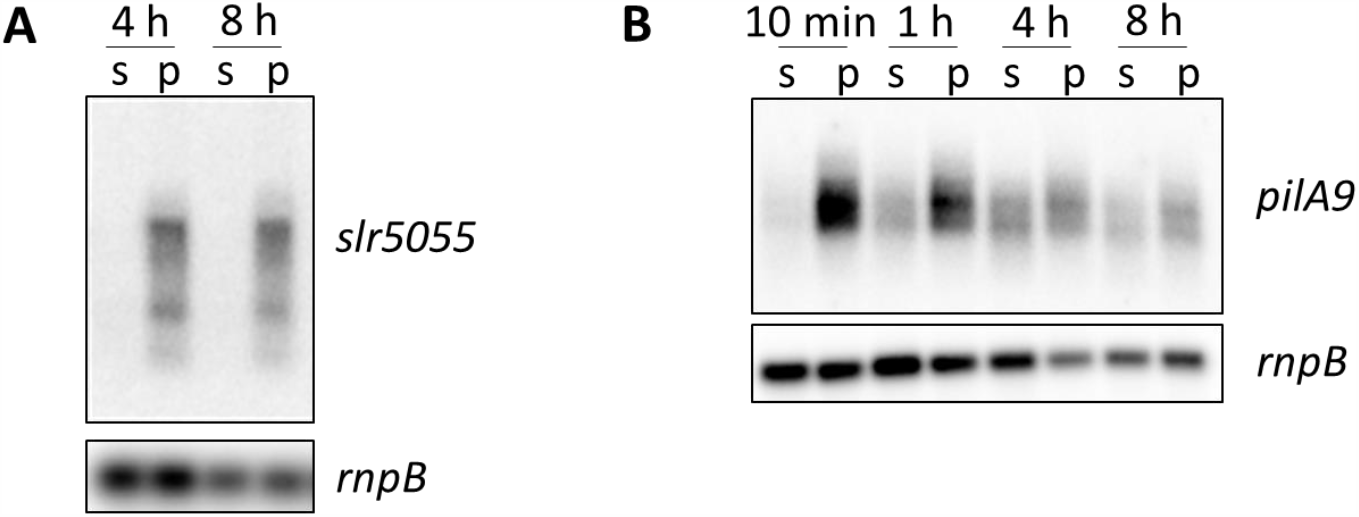
Northern blot analyses verify the microarray results. (**A**) RNAs sampled after 4 h or 8 h of acclimation to sessile (s) or planktonic (p) conditions were hybridized with *slr5055*. (**B**) Transcript accumulation of *pilA9* at timepoints of 10 min, 1 h, 4 h and 8 h. The *rnpB* probe was used for normalization. At least duplicates were performed, and only representative pictures are presented here.

In summary, the reported transcriptional changes occurring after transfer of a liquid culture to an agar surface suggest that the *Synechocystis* acclimation response includes different cellular processes. Downregulation of protein synthesis might relate to a slower growth of cultures on a surface, whereas upregulation of a subset of C_i_-responsive genes suggests CO_2_ limitation, at least in the first 4 h after transfer from planktonic cultures to sessile conditions. Genes which show a stable or increasing transcriptional change after 8 h of surface contact might be stronger candidates for a function in surface growth.

### Identification of putative new minor pilins of Synechocystis

Our microarray analysis identified several chromosomal genes that also showed a comparably stable transcriptional change during the 8 h of surface acclimation. Among these genes, *slr0226*, as well as *pilA5* and *pilA6*, exhibited increased transcription under sessile conditions. Constant downregulation upon sessile conditions was identified for *slr0442*, as well as the minor pilins from operon *pilA9*-*slr2019* (Table 1). Notably, these transcripts were also coregulated in other transcriptomic studies (Yoshimura *et al*., 2002a; Dienst *et al*., 2008; Kizawa *et al*., 2016; Wallner *et al*., 2020).

Synteny analysis using the tool FlaGs (Table S6) demonstrated that in many cyanobacteria, a hypothetical gene (listed as 2 in the analysis output; Fig. S4; Table S1) is located directly upstream or downstream of *pilA5* homologues from other cyanobacteria (Fig. S4A, Table S1). Although such a gene was not found in proximity to *pilA5* in *Synechocystis*, blastp analysis identified three *Synechocystis* homologues of the predicted hypothetical genes - *slr0226, slr0442* and *sll1268*. Further analysis showed that pilins are often encoded in genetic proximity to the homologues of these three genes (Fig. S4B-D, Table S1). Slr0226, Slr0442 and Sll1268 are annotated as hypothetical proteins, but all contain a conserved PilX-N-terminal domain (HHsearch probability > 95%; Fig. S5). PilX is a minor pilin from *Pseudomonas* that is essential for motility and is implicated as a key promotor of pilus assembly. Furthermore, PilX stimulates opening of the PilQ secretin pore in the outer membrane (Giltner *et al*., 2010; Giltner *et al*., 2011).

Notably, we also found a PilX-N domain in Slr2018 (Fig. 3; CDvist; HHsearch probability > 95%). The protein Slr2018 encoded within the *pilA9*-*slr2019* operon might therefore be a PilX-like protein or an additional minor pilin (proposed by Chandra *et al*., 2017) of unusually large size (773 aa). Nevertheless, a multiple sequence alignment of the potential minor pilins and PilX homologues shows that the hydrophobic regions of Slr0226, Slr0442 and Sll1268 are much more conserved to each other than to the hydrophobic region of Slr2018 (Fig. 1B).

These results suggest that in addition to the nine previously identified minor pilins, *Synechocystis* encodes three PilX-like minor pilins (Slr0442, Slr0226 and Sll1268), which are in other cyanobacteria in synteny with putative PilA5 homologues. The transcription levels of two of them change in response to surface acclimation.

### Possible functions of pSYSM-encoded genes

Some of the most differentially transcribed genes between planktonic and surface-grown cells were encoded on the pSYSM plasmid. The physiological roles of the corresponding proteins have not been fully elucidated to date; thus, we performed blastp homology searches and NCBI conserved domain searches. The three genes with upregulated transcription in sessile lifestyle were ambiguous and could not be classified into a specific biological process (*slr5087, slr5088, slr5037*; Table 1). However, of the ten pSYSM genes with downregulated transcription upon surface contact, six showed homology to glycosyltransferases, two for sulphotransferases, and one for a cyanoexosortase, while one was ambiguous. One of the potential glycosyltransferases (Sll5057) also harboured a conserved domain for an anti-anti-sigma regulatory factor.

As the genes *slr5087* and *slr5088*, encoded on the pSYSM plasmid, showed the highest upregulation upon surface contact, we sought to test the phenotypes of their respective deletion mutant, Δ*slr5087*-*5088* (Fig. S1). We performed phototaxis experiments and flocculation assays. However, we could not detect any difference between the mutant and the WT (Fig. 8).

**Figure 8:**
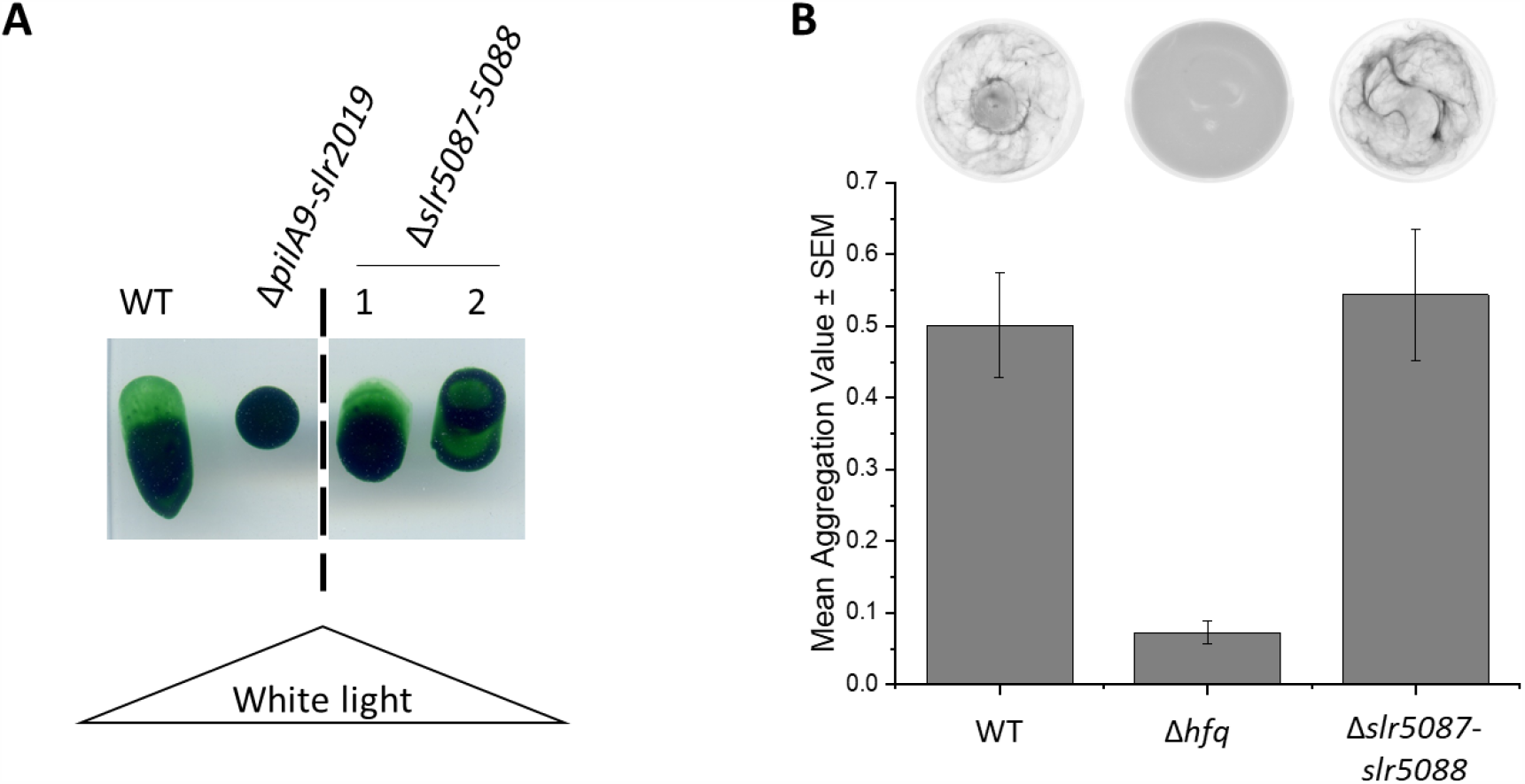
Phenotypes of the Δ*slr5087*-*5088* deletion mutant. (**A**) Phototaxis assay of the Δ*slr5087*-*5088* deletion mutant. Picture was taken after 6 days under unidirectional illumination with white light. Shown are two different clones. Δ*pilA9*-*slr2019* was used as negative control. (**B**) Flocculation of the Δ*slr5087*-*5088* deletion mutant. Chlorophyll fluorescence is shown in inverted greyscale; thus, areas with more chlorophyll appear darker. Presented are mean aggregation values ± standard errors of WT, Δ*hfq* and Δ*slr5078*-*slr5088* (n=8).

### Concentrations of the second messenger nucleotides cAMP and c-di-GMP change upon acclimation to a surface

Previously, it was shown that the two sets of minor pilins analysed in this study are differentially transcribed in a c-di-GMP-dependent manner (Wallner *et al*., 2020). Therefore, we analysed the intracellular accumulation of c-di-GMP upon the transfer of cells from a planktonic lifestyle to a surface. Cells were treated as for RNA sampling. We were not able to detect significant changes (*p*-values ≤ 0.05) in the cellular c-di-GMP level after planktonic cells were incubated for 4 and 8 h on an agar plate (Fig. 9). Therefore, an altered c-di-GMP content appears not to be directly responsible for the transcriptional changes detected in the microarray analysis at these time points. However, elevated c-di-GMP concentrations were detected after 10 min and 1 h under sessile conditions (Fig. 9). This result suggests a very fast response of the second messenger to surface contact. Comparison of c-di-GMP-dependent transcriptional changes (Wallner *et al*., 2020) and altered gene transcript levels upon surface contact indicates only a small overlap or even opposite changes between these two analyses (see discussion). Further, we revealed a rapid response of cAMP accumulation upon surface contact and an approximately 3.6-fold elevated cAMP level after 4 h on agar compared to planktonically grown cells.

**Figure 9:**
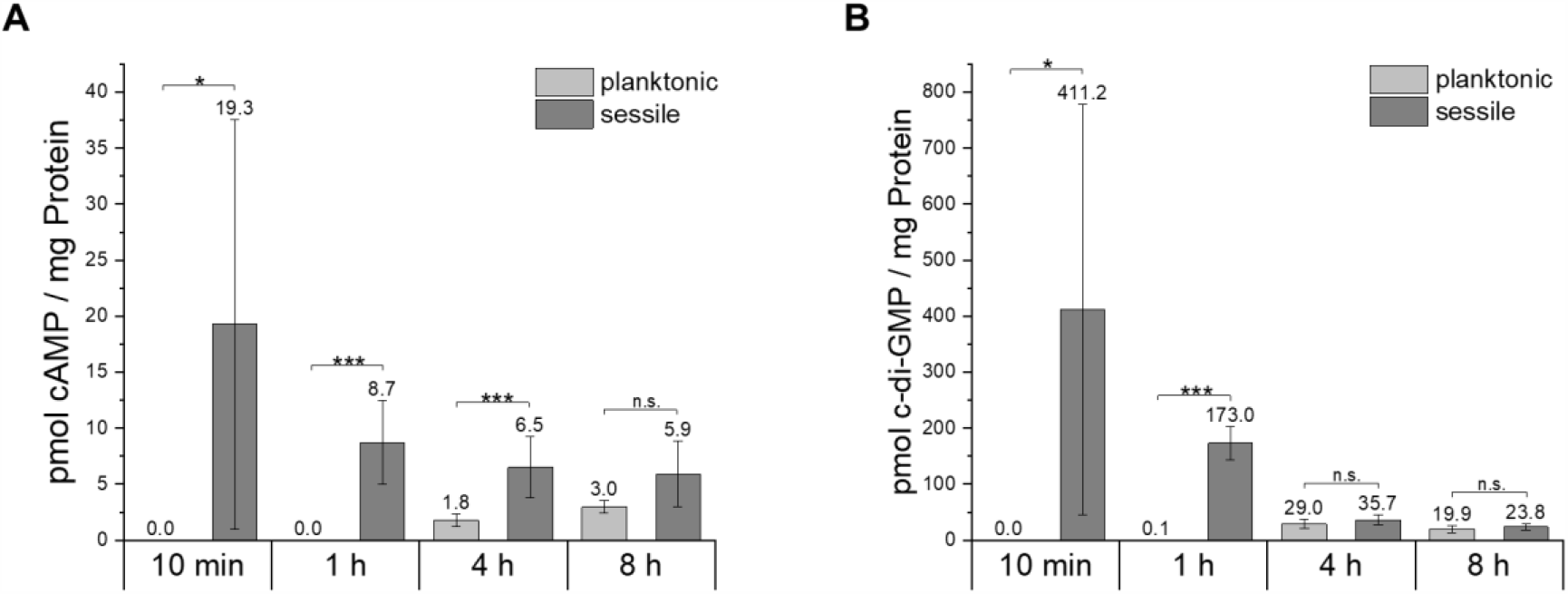
Intracellular cyclic nucleotide messenger concentrations in planktonic or sessile-grown *Synechocystis* WT cells. A planktonic preculture was cultivated either under planktonic or sessile conditions, and cAMP **(A)** and c-di-GMP **(B)** concentrations were determined after 10 min, 1 h, 4 h and 8 h. The bars display the mean concentration of cyclic nucleotide concentration normalized to whole cell protein amount. Error bars represent standard deviation. Experiments were performed as biological and extraction technical replicates. Significance was tested using a two-tailed Student’s t-test. * p ≤ 0.05, ** p ≤ 0.01, *** p ≤ 0.001, n.s. not significant

Taken together, we detected a very rapid increasing response of the second messengers cAMP and c-di-GMP within the first minutes after surface contact. However, the intracellular second messenger levels after 4 and 8 h were almost indistinguishable between sessile and planktonic cells.

## Discussion

### Function of minor pilins in natural competence, motility and flocculation

We showed that two known minor pilin operons, *pilA9*-*slr2019* and *pilA5*-*pilA6*, are involved in different pilus functions (Figure 10). We demonstrated that the *pilA5* mutant is not impaired in motility, flocculation or attachment to beads, but this gene is indispensable for the natural competence of *Synechocystis*. Furthermore, we showed that *pilA5* mRNA accumulated in sessile cells. Within a biofilm it might be beneficial for the cell to take up DNA from adjacent cells, either as a nutrient or to gain new traits (Vorkapic *et al*., 2016). In contrast, it is unlikely for planktonic, single cells to be adjoin free-floating DNA; therefore, the demand for DNA uptake and PilA5 might be lower.

**Figure 10:**
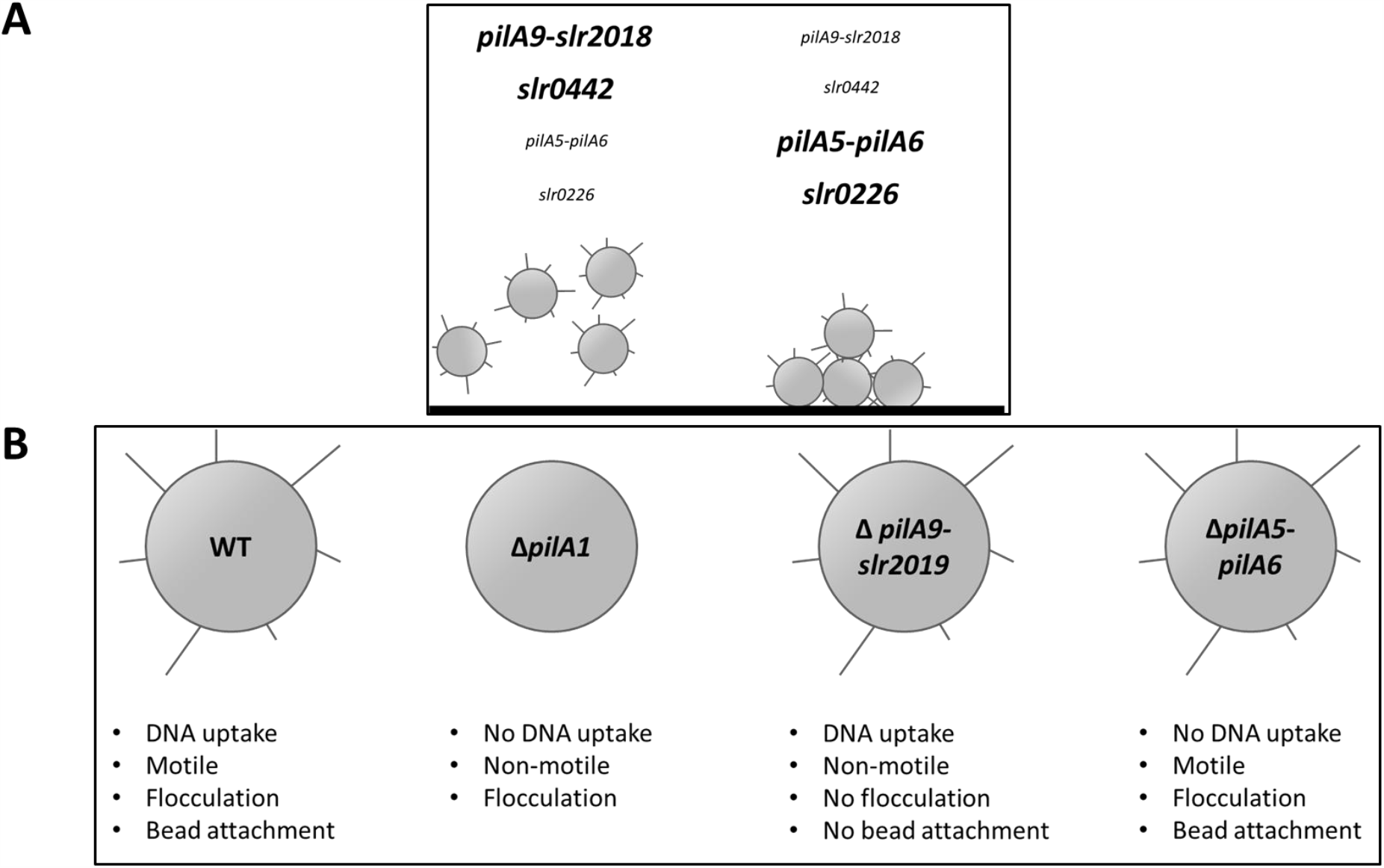
Summary of minor pilin functions. (**A**) Transcriptional change of minor pilin genes upon surface contact after 4 h and 8 h. Increased transcript names represent higher transcript accumulation. (**B**) Observed phenotypes for *Synechocystis* WT, Δ*pilA1*, Δ*pilA9*-*slr2019* and Δ*pilA5*-*6* mutant cells. Cells shown with or without schematic T4P. Data integrated from Yoshihara *et al*. (2001), Conradi *et al*. (2019) and Wallner *et al*. (2020).

In contrast to *pilA5*, the *pilA9*-*slr2019* operon is dispensable for natural competence but is involved in motility, bead attachment and flocculation. During floc formation, cells presumably bind to other cells or cellular components excreted by adjacent cells, such as T4P or EPS (Trunk *et al*., 2018). We assume that the non-motile, non-flocculating *pilA9*-*slr2019* mutant can still retract its pili, as they are still transformable. It seems that this mutant assembles pili that are not able to attach to a surface, other cells or beads. Recently, Treuner-Lange *et al*. (2020) suggested that in *Myxococcus*, several minor pilins form a complex that is part of the pilus fibre, specifically at the pilus tip. This complex of four minor pilins and PilY1 is involved in priming pilus extension and adhesion. These minor pilins show only limited similarity to the *Synechocystis* minor pilins. Furthermore, in contrast to *Myxococcus*, the minor pilin mutants of our study still formed T4P (Fig. 5). Therefore, we surmised that these mutants do not form a comparable priming complex.

Indeed, the minor pilin PilA4 from *Synechocystis* was detected within the T4P fibre (Cengic *et al*., 2018) and PilA2 and PilA11 localize at the cell surface (Hu *et al*., 2018; Cengic *et al*., 2018). Hu *et al*. (2018) reported that suppression or overexpression of *pilA11* expression via the antisense RNA PilR leads to smaller or thicker pili diameters, respectively. However, we were not able to measure significantly smaller diameters of the T4P in our Δ*pilA9*-*slr2019* mutant (Fig. S2). Eventually, the deletion of the complete minor pilin operon repeals the phenotype arising from the suppression of only *pilA11*. More likely, the negative stain used for sample preparation influences pilus visualization. Whereas, Hu *et al*. (2018) used phosphotungstic acid for staining, uronic acid was employed in our study. Additionally, TEM measurements are dependent of several parameters like noise, defocus or sample concentration, which might have added to the measured discrepancy. Whether the other minor pilins analysed in our study can be inserted into a pilus fibre remains unclear. Therefore, we attempted to model their structure. Due to poor inter- and intraspecies conservation of the amino acid sequences, we were only able to model PilA1, PilA5, PilA6 and PilA10 with acceptable sequence coverage (Fig. S6). These proteins all seem to form the hydrophobic N-terminal helix (Fig. S7), which is the most conserved trait within pilins (Giltner *et al*., 2012) and thus is assumed to be essential for the insertion of pilins into the fibre. For *Pseudomonas*, it was shown that the addition of hydrophobic substances to N-terminally truncated pilins reenables the formation of a pilus-like fimbrial structure (Audette *et al*., 2004). We surmise, that at least PilA5, PilA6 and PilA10 can be incorporated into the pilus fibre due to the conservation of the hydrophobic handle. Moreover, secondary structure prediction (Fig. S7) indicated hydrophobic handles for all known pilins and thus, all might be incorporated.

In addition to the assumption that PilA5 is incorporated into the pilus fibre, our results suggested that this protein facilitates DNA binding. However, we were not able to detect typical DNA binding motifs (Luscombe *et al*., 2000) or a region with enriched positively charged amino acids in the PilA5 sequence, as was shown for the minor pilin ComP of *Neisseria* (Berry *et al*., 2016). In this study, the researchers identified a region with a highly positively charged surface, which is probably involved in the electrostatic attraction of negatively charged DNA. Further studies should demonstrate by which mechanism PilA5 is involved in natural transformation. For *Thermus thermophilus*, recent studies demonstrated that this naturally competent, gram-negative bacterium assembles two distinct T4P filaments composed of two different pilins, that is, a wide pilus required for natural competence and a narrow pilus for twitching motility (Neuhaus *et al*., 2020). For *Synechocystis*, we did not obtain evidence from electron microscopy indicating that the different minor pilins form distinct pili (Fig. S2). This finding is supported by exoproteome analyses, which showed that the major pilin PilA1 is considerably more abundant in the exoproteome than other minor pilins (Sergeyenko and Los, 2000). Therefore, we suggest that the minor pilins might be incorporated within a filament that is primarily composed of PilA1.

### Identification of further minor pilins in Synechocystis

Taton *et al*. (2020) recently discovered indispensable proteins (minor pilin PilA3_S.e._, RntA_S.e._ and RntB_S.e_) for natural transformation of the cyanobacterium *Synechococcus elongatus* PCC 7942. RntB_S.e._ and PilA3_S.e._ exhibit modest homology to the PilA5 and PilA6 proteins of *Synechocystis* (Table S2). Notably, RntA_S.e._ shows homology to the PilX domain containing minor pilin Slr0226 (Table S2). Slr0226 together with Slr0442 and Sll1268 are newly proposed as minor pilins of *Synechocystis*. We denote them PilX1, PilX2 and PilX3, as they carry a PilX domain (Fig. 1B). Although PilX from *Pseudomonas* lacks glutamate at position +5 (E_+5_) of the PilD cleavage site, it was shown to be incorporated into the pilus fibre (Giltner *et al*., 2010). Pilins with substitution of E_+5_ to V_+5_ can still be cleaved and methylated (Strom and Lory, 1991). Moreover, it was hypothesized that pilins with a non-polar substitution have an improved ability to leave the membrane during assembly (Giltner *et al*., 2012). Thus, these proteins were proposed to form the pilus tip complex. This property was shown for the minor pilin GspK of the type II secretion complex of *E. coli*, which also lacks E_+5_ (Korotkov and Hol, 2008), and a similar PilX minor pilin found in *Acinetobacter baumannii* T4P (Piepenbrink, 2019). The three *Synechocystis* PilX homologues also have been determined to have different hydrophobic amino acid substitutions at position E_+5_ (Fig. 1B). Additionally, secondary structure predictions indicated the existence of hydrophobic N-terminal helices, which are characteristic of minor pilins and seem to be vital for insertion into the pilus (Fig. S7). Generally, PilX amino acid sequences are longer than those of the major pilin which is also true for the *Synechocystis* PilX-like proteins (Fig. 1). It has been further postulated that larger proteins locate at the pilus tip, as they sterically hinder the binding of previous pilins (discussed in Giltner *et al*., 2012). Indeed, minor pilins, such as PilX from *Pseudomonas*, have been determined to be incorporated at the pilus tip (Giltner *et al*., 2010). Thus, we surmised that *Synechocystis* minor pilins can also form minor pilin tip complexes for different T4P functions.

### Response of Synechocystis *to surface contact*

Our transcriptome analyses demonstrated that minor pilins might be major targets of a surface response in *Synechocystis* cells. In addition to *pilA5, pilA6* and *pilA9-pilA11*, at least two additional putative minor pilin genes (*slr0442* and *slr0226*) were observed to respond to surface attachment. Importantly, some transcripts were determined to accumulate in surface-grown cells, while others accumulated only in planktonic culture, suggesting distinct functions. Our microarray analysis demonstrated that *slr0226* mRNA accumulates under sessile conditions along with minor pilin mRNA *pilA5* and *pilA6*. In contrast, the transcription of *slr0442* was downregulated under sessile conditions along with the minor pilin operons *pilA9, pilA10, pilA11*, and *slr2018* (Table 1). This contrasting behaviour in the transcriptomic response of minor pilin genes was also detected in other microarray studies. The cAMP receptor protein SyCRP1 activated the transcription of *slr0442*, as well as the *pilA9, pilA10, pilA11*, and *slr2018* genes (Yoshimura *et al*., 2002a; Hedger *et al*., 2009). Similar transcription profiles for *slr0442* and *pilA9*-*slr2018* were observed in a microarray study that compared gene expression between WT and an *hfq* mutant (Dienst *et al*., 2008) and the transcriptional regulator LexA (Kizawa *et al*., 2016). The transcripts of *slr0226* and *pilA5*-*pilA9* accumulated under low c-di-GMP conditions (Wallner *et al*., 2020). Thus, in *Synechocystis* cells specific sets of minor pilins are coregulated, suggesting that they belong to the same regulon.

Laventie *et al*. (2019) showed for *Pseudomonas* that the c-di-GMP concentration increased within seconds upon surface contact, thereby leading to pilus assembly and enhanced surface attachment. *Pseudomonas* appeared to maintain these high c-di-GMP levels until the cells obtained the signal to detach from the surface (Laventie *et al*., 2019). However, we were not able to detect major changes in second messenger accumulation after 4 and 8 h of acclimation to a surface. At shorter time points, such as 10 min or 1 h after transfer to sessile conditions, an increase in c-di-GMP and cAMP levels was detected, suggesting that there is a short-term surface response. It is possible that these fast changes in second messenger levels were responsible for the measured transcriptional changes observed after 4 and 8 h. Indeed, 11 out of 17 genes that are known to be regulated by c-di-GMP under blue-light illumination also responded to surface contact in our analysis (Wallner *et al*., 2020). However, we identified 120 surface-responding genes in our microarray study, which suggested a considerably larger modulation for this signal. In addition, the accumulation of minor pilin-encoding transcripts in our study was inverse to the c-di-GMP response. For example, the *pilA9*-*slr2019* operon was downregulated on a surface (Tables 1 and 2), whereas a high c-di-GMP content has been observed to lead to upregulation of this mRNA (Wallner *et al*., 2020). Several other genes also responded in a contrasting way in both microarray studies. Interestingly, the accumulation of an mRNA encoding the EAL-domain protein Slr1593 was elevated in cells on the surface but also did not lead to a measurably lower c-di-GMP content in the cells (Fig. 9). It has been proposed that local pools of second messenger molecules could function to control cellular behaviour (Sarenko *et al*., 2017). Therefore, we could not exclude the possibility that localized changes in c-di-GMP control the transcription of several of the targets identified in this study. Furthermore, we detected 8 out of 18 genes that have been determined to be differentially regulated upon disruption of the cAMP receptor protein SyCRP1 (*Synechocystis*’ carbon regulated protein 1) (Yoshimura *et al*., 2002a). However, in this study, gene transcription appeared to operate again in the opposite way, as minor pilin operons *pilA9*-*pilA11* and *slr0442* were upregulated under higher cAMP levels (Yoshimura *et al*., 2002a), and in our study, they appeared to be downregulated under sessile conditions, where we eventually measured elevated cAMP levels.

We could not exclude the possibility that the different growth conditions in the microarray studies contributed to the differences in mRNA accumulation. Wallner *et al*. (2020) used mixotrophic conditions combined with green and blue light, and Yoshimura *et al*. (2002) used high CO_2_ conditions, whereas the current microarray study was performed with cells grown under photoautotrophic low-CO_2_ conditions. The two CRP-like transcription factors SyCRP1 and SyCRP2 have been determined to be involved in second messenger signalling and are known to control at least the *pilA9-slr2019* operon (Yoshimura *et al*., 2002b; Song *et al*., 2018; Wallner *et al*., 2020). CRPs are often used by proteobacteria to coordinate responses to carbon sources, and binding of the second messenger cAMP has been observed to lead to either activation or repression of gene transcription as part of the catabolite-repression response (Botsford and Harman, 1992). In cyanobacteria, it has been shown that cAMP is a high-carbon signal (Selim *et al*., 2018). It appears that glucose, high-CO_2_ conditions, different stages of surface attachment and the circadian clock (Taton *et al*., 2020) could present further signals to the regulatory network.

## Materials and Methods

### Bacterial strains and culture conditions

The WT of *Synechocystis* sp. PCC 6803 employed in this study is motile and can grow photoautotrophically, mixotrophically and chemoheterotrophically on glucose. This strain was originally obtained from S. Shestakov (Moscow State University, Russia) in 1993 and was re-sequenced in 2012 (Trautmann *et al*., 2012).

Cyanobacteria were cultivated under either sessile or planktonic conditions. Planktonic cultures were grown in modified 1 x BG11 (Rippka *et al*., 1979) substituted with 0.3 % (w/v) sodium thiosulphate and 10 mM (w/v) (N-[tris-(hydroxymethyl)-methyl]-2-aminoethane sulphonic acid) buffer (TES) pH 8.0. Cultures were grown photoautotrophically with orbital shakers at 140 rpm at 30°C under continuous white light illumination (Philips TLD Super 80/840) of 50 µmol photons m^-2^ s^-1^. Sessile cultures were grown on 0.75% agar plates (Bacto-Agar) using the same medium and growth conditions without shaking as described for planktonic cultures.

Cyanobacterial strains were cultivated on BG11 agar plates as described above, and antibiotics, if necessary, were added at the following concentrations: chloramphenicol, 14 µg ml^-1^; kanamycin, 40 µg ml^-1^; and gentamycin, 10 µg ml^-1^. For cloning of plasmids, *E. coli* DH5α was used, which was grown in LB medium supplemented with antibiotics at the following concentrations: ampicillin, 100 µg ml^-1^; streptomycin, 25 µg ml^-1^; and gentamycin, 10 µg ml^-1^.

### Construction of mutant strains

The deletion mutant Δ*pilA5-pilA6* was created as described in Wallner *et al*. (2020). Three plasmids were constructed bearing either the single gene *pilA5* or *pilA6* alone or the operon *pilA5*-*pilA6* under the control of the native promoter (P_pilA5-pilA6_). For a list of oligonucleotides used to generate all mutant constructs, see Table S3. The gene fragments P_pilA5-pilA6_-*pilA5* and P_pilA5-pilA6_-*pilA5*-*pilA6* were amplified from WT chromosomal DNA with an SdaI recognition site at the 5’-end (P1-P3) and inserted into the pJET 1.2 vector (CloneJET PCR Cloning Kit, Thermo Scientific™, Germany). For the third construct, assembly cloning (AQUA cloning) (Beyer *et al*., 2015) was used to fuse the pJET 1.2 vector with the P_pilA5-pilA6_ fragment containing the SdaI recognition site at the 5’-end and the *pilA6* gene fragment (P4-P9). The subcloning vectors were cleaved with SdaI and HindIII restriction enzymes, and fragments were ligated into the pVZ322 vector (NCBI accession number AF100175), which was cleaved with the same restriction enzymes. The final plasmids were transferred into the *Synechocystis* deletion mutant Δ*pilA5-pilA6* via triparental mating (Elhai and Wolk, 1988) with *E. coli* J53 (NCBI:txid1144303) harbouring the conjugative helper plasmid RP4 (NCBI:txid2503) and *E. coli* DH5α with the constructed plasmid. Verification of correct mutant strains was validated by PCR using primer pair P10-P11 (Fig. S1). A list of all plasmids and strains used in this study can be found in Tables S4 and S5.

A *Synechocystis* knockout mutant of the *slr5087*-*slr5088* operon located on the pSYSM plasmid was generated (Fig. S1D). First, we amplified chromosomal regions to be used as homologous recombination sites and added a NsiI restriction site to homologous region 1 (HR1) and overlapping regions using primer pairs P16-P17 and P18-P19. A combined homologous region sequence was obtained by overlap PCR using the two amplified polynucleotides and primer pair P16-P19. The construct was inserted into the pJET1.2 vector according to the manufacturer’s instructions (CloneJET PCR Cloning Kit, Thermo Scientific™, Germany).

Furthermore, a chloramphenicol resistance cassette was amplified from the pACYC184 vector (GenBank: X06403.1), and NsiI restriction sites were added (P20-P21). This amplification product and the vector were cleaved by NsiI and ligated using T4 DNA ligase (New England BioLabs, Inc.). Insert orientation was checked by sequencing. Finally, *Synechocystis* WT was transformed with the plasmid. DNA of the mutants was extracted from cells by three repeated cycles of shock freezing (−80°C) and heating (60°C) for 10 minutes each. DNA was separated from cell debris by centrifugation at 3000 x *g* for 3 minutes at 4°C. Correct construct insertion and full segregation were tested by colony PCR using the primer pair P22-P23 (Fig. S4E).

### Flocculation assay

Flocculation assays were mainly performed as described in Conradi *et al*. (2019). Briefly, cultures were diluted to an OD_750nm_ 0.25, and 6 ml were transferred to 6-well plates (Corning Costar®, non-treated, 392-0213, VWR, Germany). Cultures were incubated for 48 h at 30°C and 40 μmol photons m^−2^ s^−1^ of white light at 95 rpm on an orbital shaker. Images were taken by measuring chlorophyll autofluorescence using a Typhoon FLA4500 imaging system (GE Healthcare) with laser excitation at 473 nm and fluorescence detection at 665 nm. The aggregation score was calculated by dividing the standard deviation by the mean intensity as described in Conradi *et al*. (2019).

### Transformation assays

Natural transformation competence was tested with a suicide plasmid encoding streptomycin resistance. Cells were grown to log phase (OD_750nm_ 0.6 – 0.8) in planktonic culture, and 15 ml were harvested by centrifugation (3200 x *g*, RT). The cell pellet was resuspended in 300 µl BG11 medium. Then, 500 or 1000 ng of plasmid pDrive-Δ*psrR1*-Strep (Georg *et al*., 2014) were added to the suspension, and cultures were incubated at 30°C under white light (≈ 50 µmol photons m^-2^ s^-1^) without shaking. After three hours, cells were plated on sterile filters (0.2 µm pore size, MN615, Macherey-Nagel), placed on BG11 agar plates and incubated under standard conditions. After two days, the filters were transferred to new BG11 agar plates supplemented with 5 µg ml^-1^ streptomycin and incubated further until colonies appeared. Integration of the construct used led to deletion of the sRNA gene *psrR1*, which had no phenotypical effect under normal cultivation conditions (Georg *et al*., 2014).

### Transmission electron microscopy

Strains were grown in BG-11 medium under continuous white light conditions (40-50 µmol photons m^-2^ s^-1^) at 28°C for 2 days to an OD_730nm_ 0.7-0.8. Cells were dropped onto formvar-coated grids, negatively stained with 1% aqueous uranyl acetate (w/v) according to Harris (1997) and examined by a Jeol 1010 transmission electron microscope operated at 80 kV equipped with a Mega View III camera (SIS). Acquired pictures were analysed with ImageJ (Abramoff *et al*., 2004).

### Fluorescent bead assay

Fluorescent bead assays were performed as described in Nakane & Nishizaka (2017) with slight changes. The coverslip was prepared as described by Nakane & Nishizaka (2017) but coated with 4% (v/v) collodion in isoamyl acetate. Cells were added and incubated for 10 min, and then unattached cells were removed by rinsing with BG11. Coverslips were transferred to the microscope stage and illuminated for 2 min with red light (λ_max_ = 640 nm). Then, 0.2 µm fluorescent polystyrene beads (F8848, Thermo Fisher Scientific, Germany, final concentration 0.02% (w/v) in BG11) were added, and the fluorescent signal was detected by excitation at 426-450 nm, cut off at 458 nm and detected at 467 - 499 nm with an upright microscope (Nikon Instruments, Japan).

### Phototaxis experiments

Phototactic movement of cyanobacterial cells was analysed on 0.5% agar plates containing BG11, 10 mM (w/v) TES buffer (pH 8.0), 0.3 % (w/v) sodium thiosulphate and 11 mM glucose as previously described by Jakob *et al*. (2017). Cells were concentrated in a small volume of BG11, and 7 µl of cell suspension were spotted as droplets in triplicate on the phototaxis plate. Plates were then illuminated with unidirectional white light (≈ 5 µmol photons m^-2^ s^-1^) and incubated for 6 days at 30°C. Pictures were taken with a flatbed scanner.

### RNA extraction

A planktonic preculture was grown in BG11 medium to OD_750nm_ 0.6-0.8. Then, 35 ml were centrifuged at 4000 x *g* for 10 min at room temperature. For planktonic cultures, pellets were resuspended in 35 ml BG11 medium and incubated as described above in the culture conditions section. For sessile cultures, pellets were resuspended in 300 µl BG11 medium, plated onto BG11 agar plates and incubated as described above in the culture conditions section. After 10 min, 1 h, 4 h or 8 h RNA was harvested. For planktonic cultures, cells were collected by vacuum filtration through 0.8 µm polyethersulphone filter disks (Pall, Germany). Filters were immediately transferred to 1.6 ml PGTX solution (Pinto *et al*., 2009), vortexed, frozen in liquid nitrogen and stored at −80°C until further use. For sessile cultures, cells were scratched from the agar plates and directly suspended in 1.6 ml PGTX solution, vortexed, frozen in liquid nitrogen and stored at −80°C. RNA was then extracted according to Pinto *et al*. (2009) with modifications as described in Wallner *et al*. (2020).

### Microarray analysis

Removal of potential DNA from the RNA samples was performed using Ambion TurboDNase (Thermo Fisher Scientific, Germany) according to the manufacturer’s instructions, but addition and incubation with Turbo DNase were performed twice. RNA was precipitated overnight with 3 M sodium acetate (pH 5.2) in 97 % (v/v) ethanol. Furthermore, RNA was collected by centrifugation at 20800 x *g* for 30 min, washed with 70% ethanol and eluted in nuclease-free water. The concentration was checked on a NanoDrop ND2000 spectrophotometer (Thermo Fisher Scientific, Germany), and integrity was verified using a fragment analyser (Advanced Analytical Technologies, Germany). Furthermore, RNA was labelled with Cy3 (ULS™ Fluorescent Labeling Kit for Agilent Arrays (EA-023), Kreatech, Germany), fragmented, and 600 ng were hybridized with a high-resolution custom-made microarray (Agilent Technologies, Germany, Design ID 075764, format 8 × 60 K; slide layout = IS-62976-8-V2). The full dataset is available in the database NCBI’s Gene Expression Omnibus and accessible through GEO Series accession number GSE161586 (https://www.ncbi.nlm.nih.gov/geo/query/acc.cgi?acc=GSE161586). Further, the analysed dataset, containing all coding and non-coding RNAs can be found in the Supplementary Data Table. Graphical visualizations of the microarray results for the chromosome and all large plasmids are available in data sets S1-S5.

### Northern blot hybridization

To verify the microarray results, Northern blot hybridizations were performed. RNA was extracted as described above. Seven micrograms of RNA was separated on a denaturing 1.3% (w/v) agarose-formaldehyde electrophoresis gel and blotted onto a Roti-Nylon plus membrane (Carl Roth, Germany). The Northern blot was hybridized with probes generated by *in vitro* transcription of PCR fragments (P12-P15, Table S3) using radioactively labelled [α-^32^P]-UTP and the Ambion T7 polymerase Maxiscript kit (Thermo Fisher Scientific, Germany). Signals were detected by phosphoimaging on a Typhoon FLA4500 imaging system (GE Healthcare).

### Extraction of nucleotide second messengers

Cells for planktonic and sessile cultures were grown the same way as described for RNA extraction. To obtain measurable amounts of second messengers for sessile cultures, cells from two plates were scratched and suspended in 250 µl BG11 medium. Cells from 50 µl of this suspension were pelleted at 11000 x *g* at 4°C for 2 min for nucleotide extraction, and 50 µl were pelleted and used for determination of protein concentration. For planktonic samples, 10 ml of a liquid cell culture were pelleted for nucleotide and 1 ml for protein quantification. Cell pellets were resuspended in 300 µl extraction solution (acetonitrile/methanol/water 2:2:1 (v/v/v)), incubated at 4°C for 15 min and heated to 95°C for 10 min. Samples were snap-cooled on ice and stored for further use at −20°C or centrifuged at 21000 x *g* at 4°C for 10 min, and the supernatant was collected. The extraction was repeated twice from the pellets with 200 µl extraction solution each, without heat treatment. Supernatants were combined, and proteins precipitated by incubation at −20°C overnight. The samples were centrifuged at 21000 x *g* at 4°C for 10 min, and the supernatant containing the extracted second messengers was air dried in a SpeedVac at 42°C. Quantification of c-di-GMP and cAMP was performed by HPLC/MS/MS analysis, as previously described (Burhenne and Kaever, 2013). Second messengers were normalized to the total protein amount. For protein quantification, cell pellets were resuspended in 50 µl phosphate-buffered saline (PBS), supplemented with 0.7 volume of glass bead mix (0.1-0.11 & 0.25-0.5 mm) and vortexed for 60 sec. Then, the samples were snap-cooled at −80°C twice and heated to 40°C for 10 min each. After sedimentation of the beads, 2 µl of the supernatant were used for protein quantification using the Direct Detect system (Merck Millipore).

### Bioinformatic analysis

All bioinformatic tools used are listed in Table S6.

## Supporting information

Supplementary data set (microarray genome plot)

Supplementary data set (raw microarray data)

## Acknowledgements

We thank D. Nakane (Gakushuin University), who established the bead assay in our lab, and A. Jakob (University of Freiburg) for performing the assays together with S. Oeser. We acknowledge V. Reimann and W. Bigott (both University of Freiburg) for help with microarray analysis and Northern Blots hybridizations. We are grateful to the students S. Klostermayer and A. Moellering for experimental contributions. We thank R. Sobotka (Institute of Microbiology of the Czech Academy of Sciences) for scientific exchange. We acknowledge the Laboratory of Electron Microscopy, the core facility of Biology Centre of CAS supported by the MEYS CR (LM2015062 Czech-BioImaging). This work was supported by grants to AW of the German Science Foundation (DFG) as part of DFG Priority Programme SPP1879 (Wi 2014/7-1) and the SFB1381 (A2).

## Author contributions

SO performed experiments, analyzed and interpreted the data, wrote the manuscript and contributed to the design of the study.

TW performed experiments, interpreted the data and contributed to the design of the study. LB performed experiments and analyzed data.

NS interpreted the data and contributed to the design of the study.

AW contributed to the design of the study, interpreted data and wrote the manuscript.

