## Supplementary data set (microarray genome plot) for "Minor pilin genes are involved in motility and natural competence in *Synechocystis* sp. PCC 6803"

### Supplementary Data Sets

Data Set S1: **Graphical visualization of the microarray results of the chromosome.** This microarray data presents changes between Synechocystis cells of a sessile and planktonic lifestyle after 4 or 8 h, that are located on the chromosome. The transcription values for planktonic cells after 4 h (liquid_4h, blue) or 8 h (liquid_8h, ice blue) and sessile cells after 4 h (solid_4h, red) or 8 h (solid_8h, orange) are given in log2 scale. The read numbers of exponential (dark grey) or stationary phase (light grey) grown cells are shown in log2 scale and are extracted from Kopf *et al.* (2014). Supporting data can be found in the Supplementary Data Table.

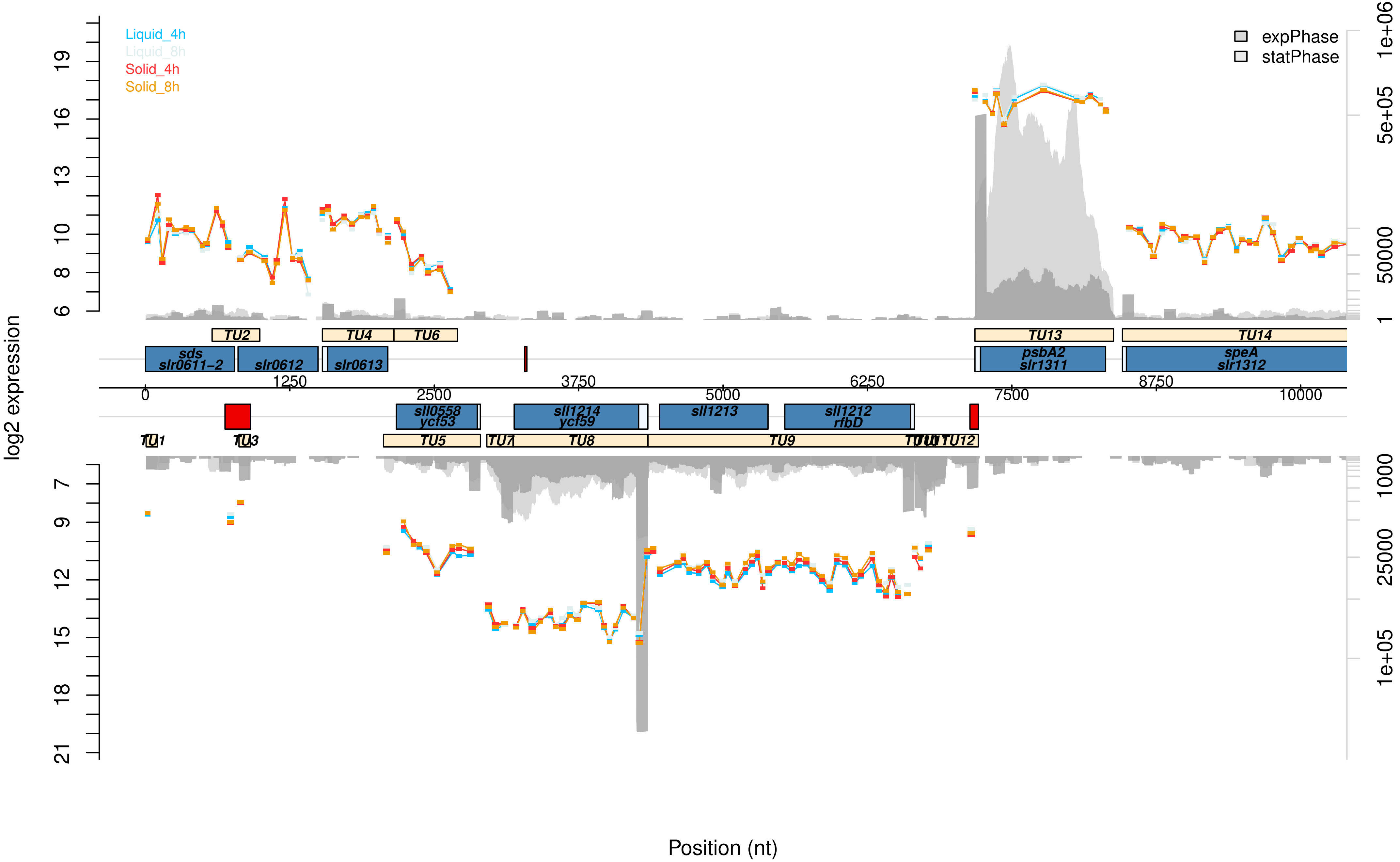

Liquid_4h
Liquid_8h
Solid_4h
Solid_8h
expPhase
statPhase
log2 expression
Position (nt)
TU1
TU2
TU3
TU4
TU5
TU6
TU7
TU8
TU9
TU12
TU13
TU14
sds
slr0611−2
slr0612
slr0613
sll0558
ycf53
sll1214
ycf59
sll1213
sll1212
rfbD
psbA2
slr1311
speA
slr1312

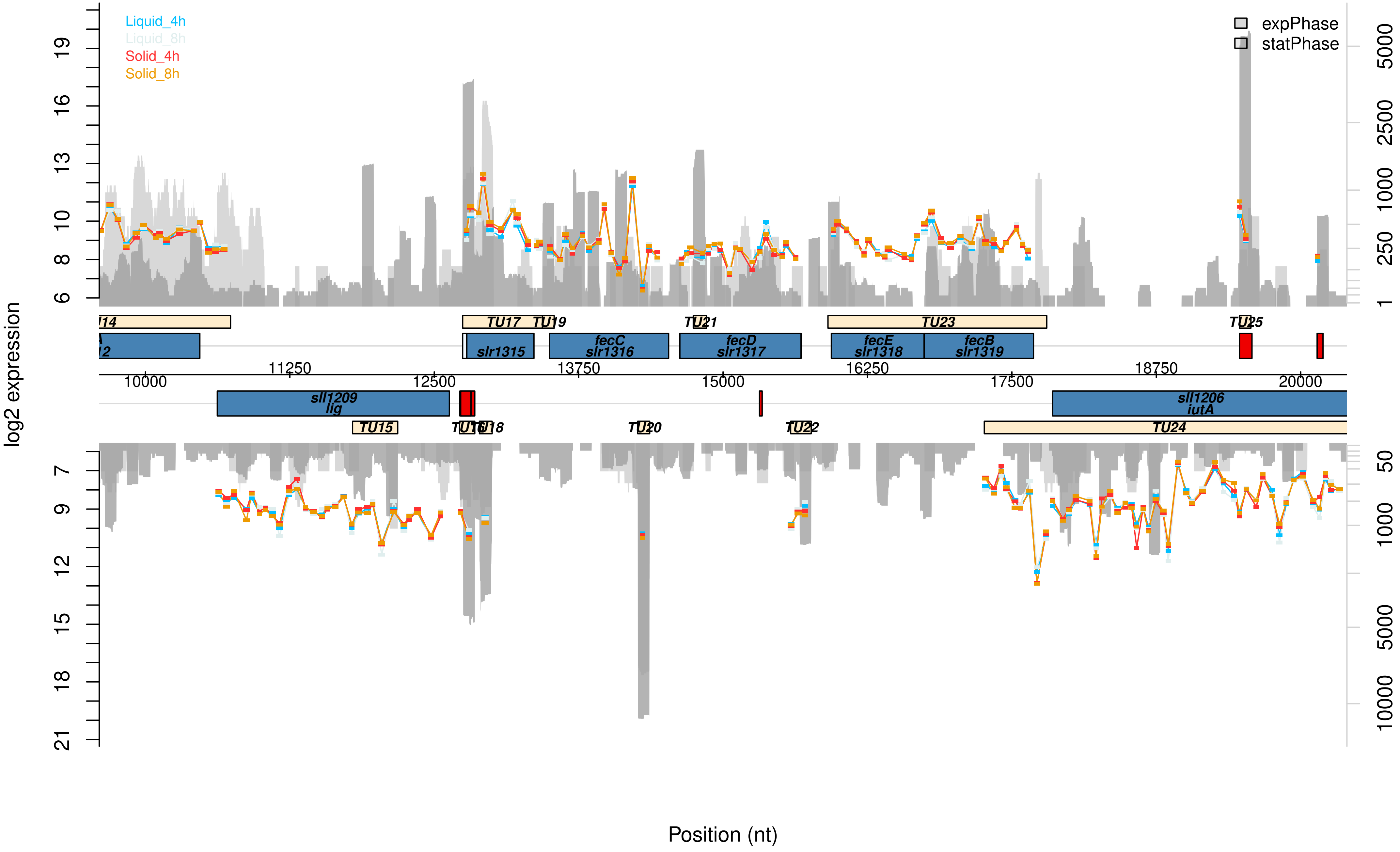

Liquid_4h
Liquid_8h
Solid_4h
Solid_8h
expPhase
statPhase
log2 expression
TU14
TU17
TU19
TU21
TU23
TU25
slr1315
fecC
slr1316
fecD
slr1317
fecE
slr1318
fecB
slr1319
10000
11250
12500
13750
15000
16250
17500
18750
20000
sll1209
lig
TU15
TU16
TU18
TU20
TU22
sll1206
iutA
TU24
Position (nt)

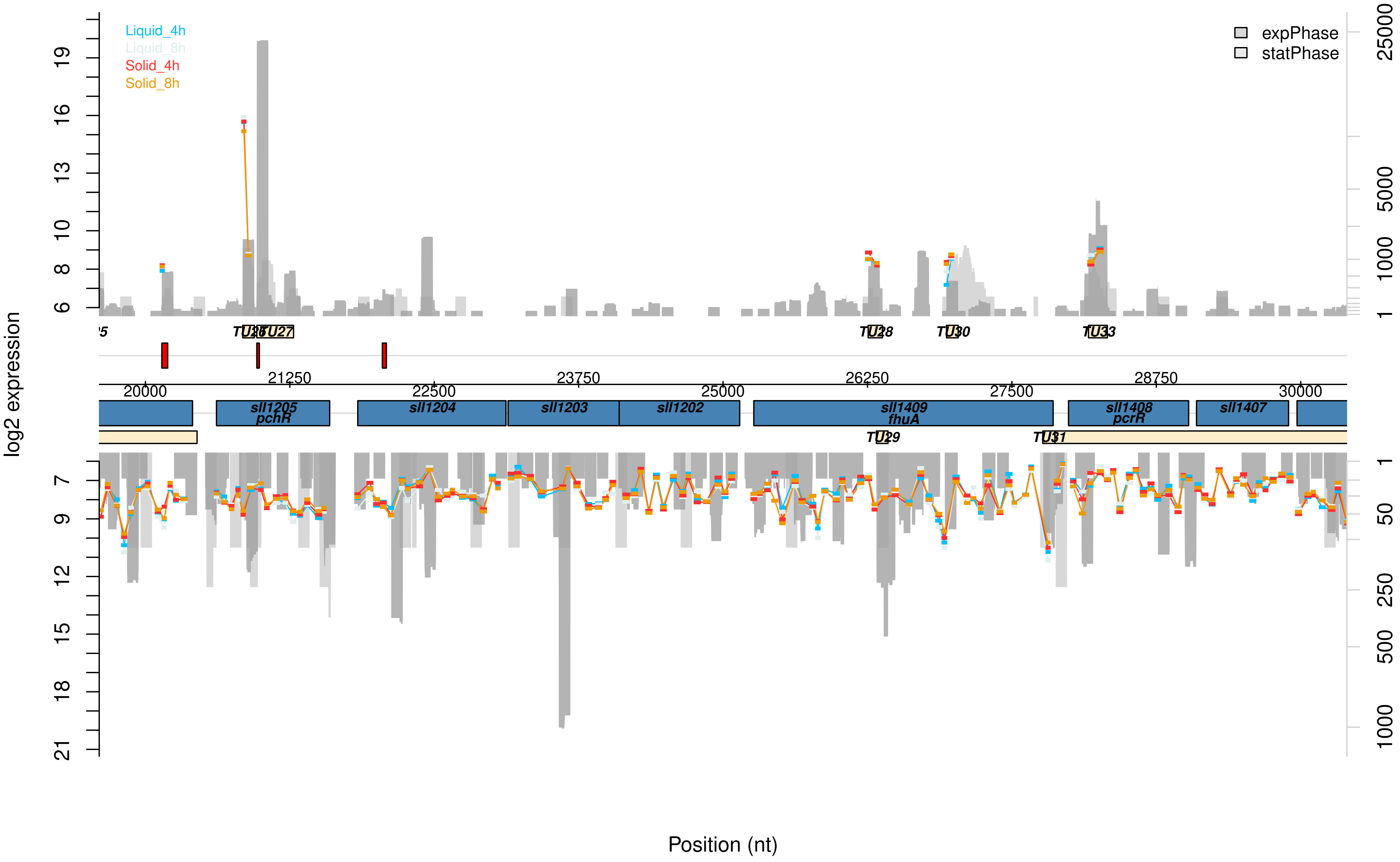

Liquid_4h
Liquid_8h
Solid_4h
Solid_8h
expPhase
statPhase
log2 expression
TU26
TU27
TU28
TU30
TU33
21250
23750
26250
28750
20000
22500
25000
27500
30000
sll1205
pchR
sll1204
sll1203
sll1202
sll1409
fhuA
sll1408
pcrR
sll1407
TU29
TU31
Position (nt)

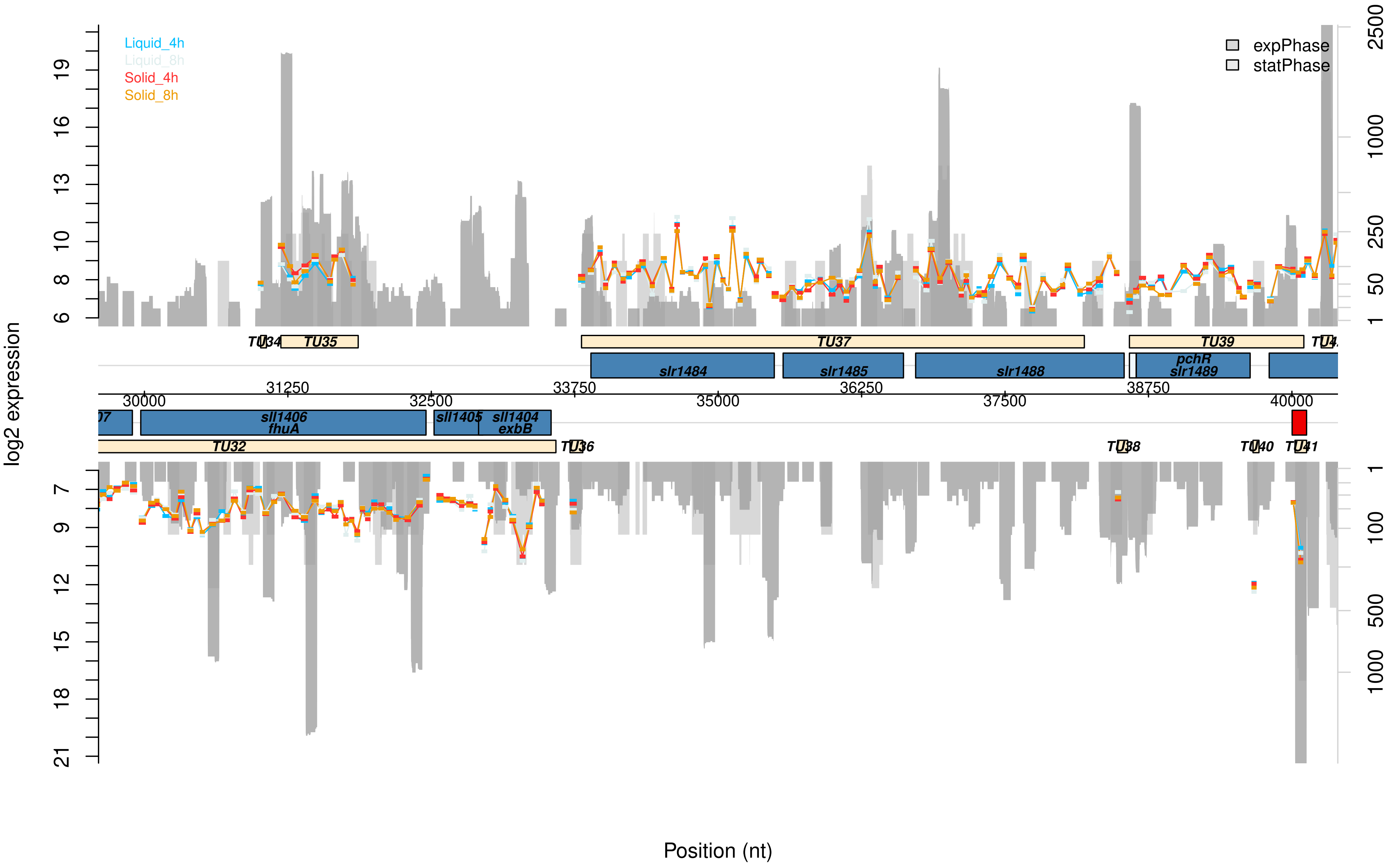
Liquid_4h
Liquid_8h
Solid_4h
Solid_8h
expPhase
statPhase
log2 expression
Position (nt)
TU34
TU35
TU37
TU39
TU32
TU36
TU38
TU40
TU41
slr1484
slr1485
slr1488
pchR
slr1489
sll1406
fhuA
sll1405
sll1404
exbB
31250
33750
36250
38750
30000
32500
35000
37500
40000
6
8
10
13
16
19
7
9
12
15
18
21
2500
1000
250
50
1
100
500

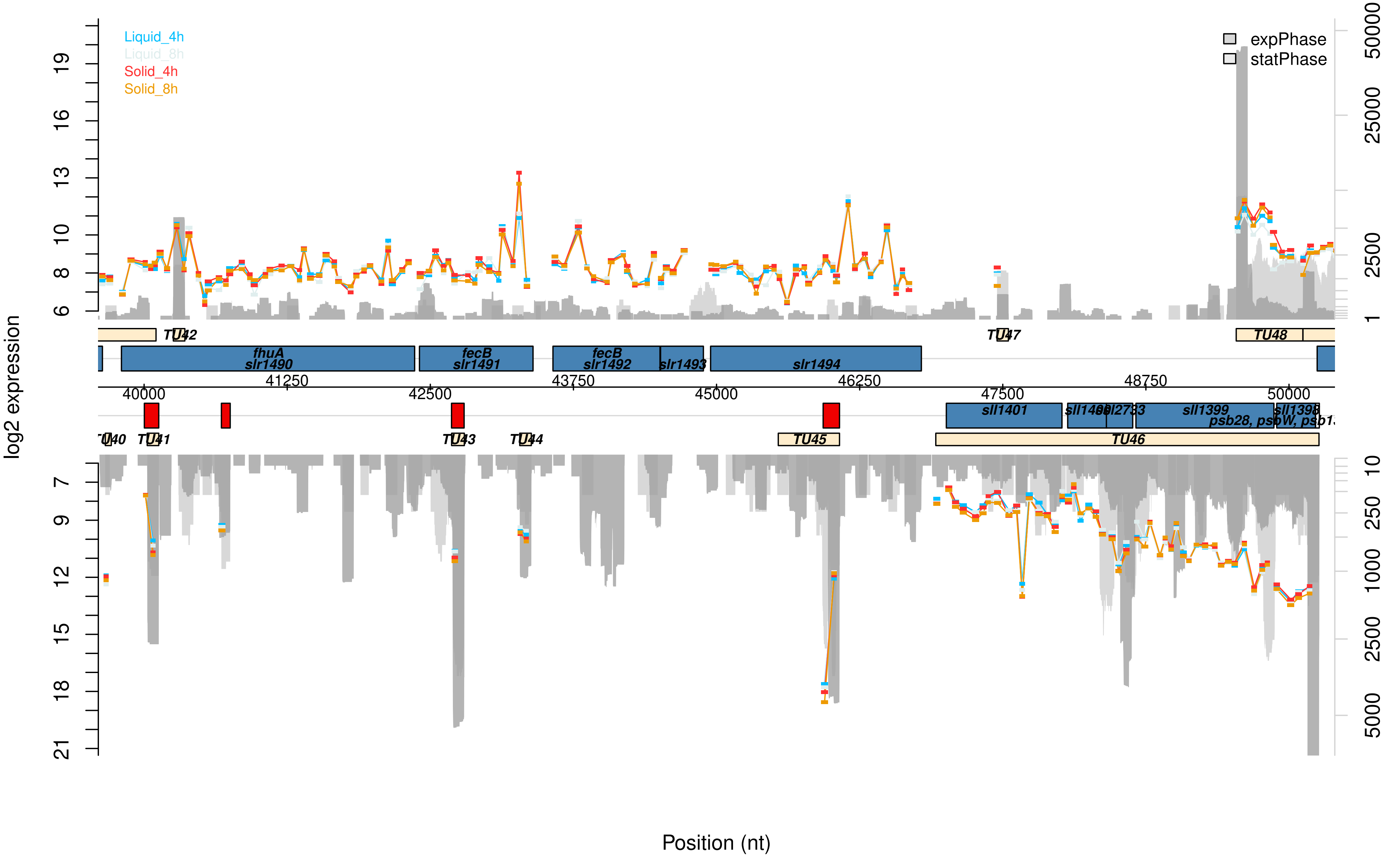

Liquid_4h
Liquid_8h
Solid_4h
Solid_8h
expPhase
statPhase
log2 expression
Position (nt)
TU40
TU41
TU42
TU43
TU44
TU45
TU46
TU47
TU48
fhuA
slr1490
fecB
slr1491
fecB
slr1492
slr1493
slr1494
sll1401
sll1400
sll2733
sll1399
sll1398
psb28, psbW, psb1
40000
41250
42500
43750
45000
46250
47500
48750
50000

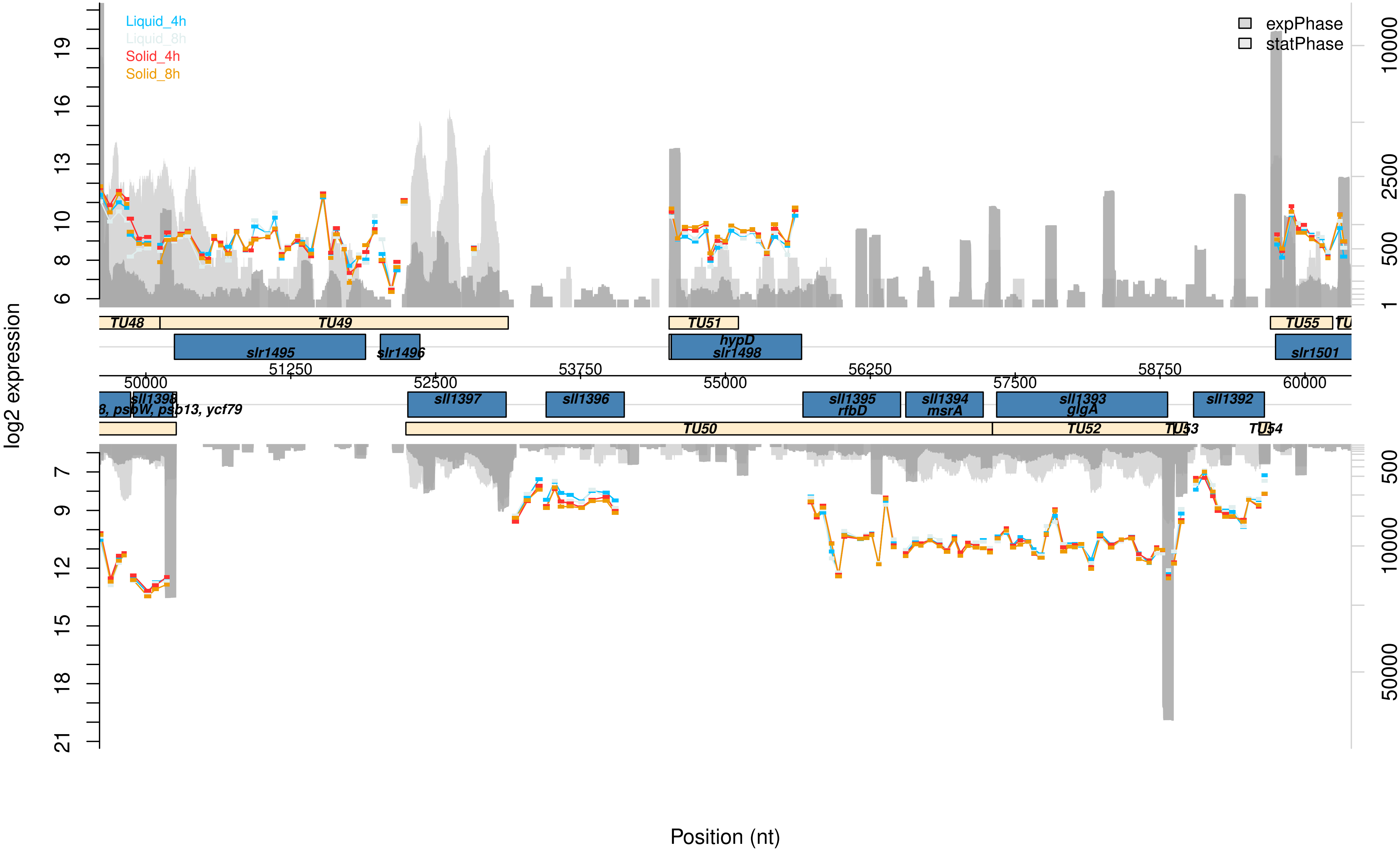

Liquid_4h
Liquid_8h
Solid_4h
Solid_8h
expPhase
statPhase
log2 expression
Position (nt)
TU48
TU49
TU51
TU55
slr1495
slr1496
hypD
slr1498
slr1501
sll1398
psbW, psb13, ycf79
sll1397
sll1396
sll1395
rfbD
sll1394
msrA
sll1393
glgA
sll1392
TU50
TU52
TU53
TU54
50000
51250
52500
53750
55000
56250
57500
58750
60000

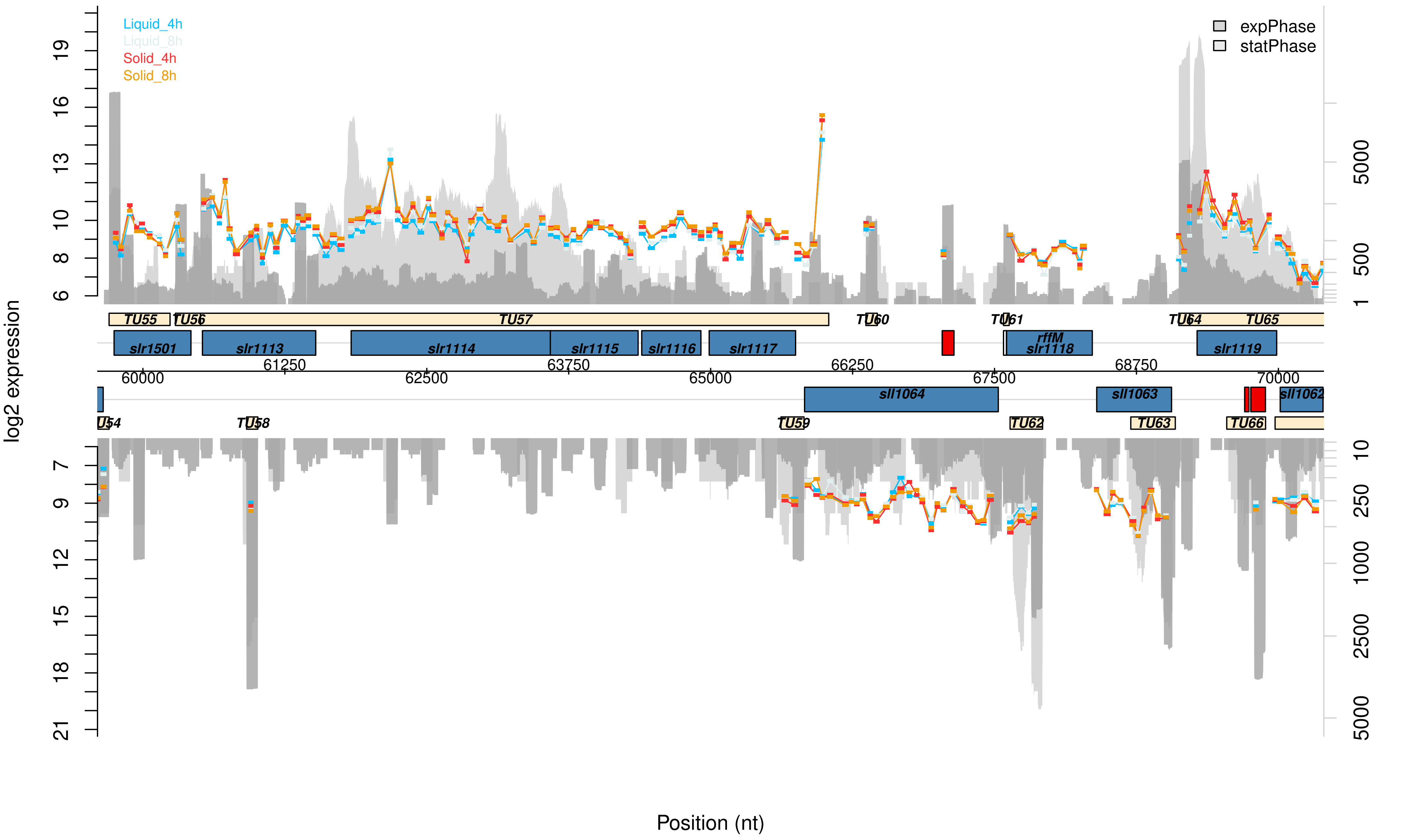

Liquid_4h
Liquid_8h
Solid_4h
Solid_8h
expPhase
statPhase
log2 expression
Position (nt)
TU55
TU56
TU57
TU60
TU61
TU64
TU65
slr1501
slr1113
slr1114
slr1115
slr1116
slr1117
rffM
slr1118
slr1119
sll1064
sll1063
sll1062
TU58
TU59
TU62
TU63
TU66
60000
61250
62500
63750
65000
66250
67500
68750
70000

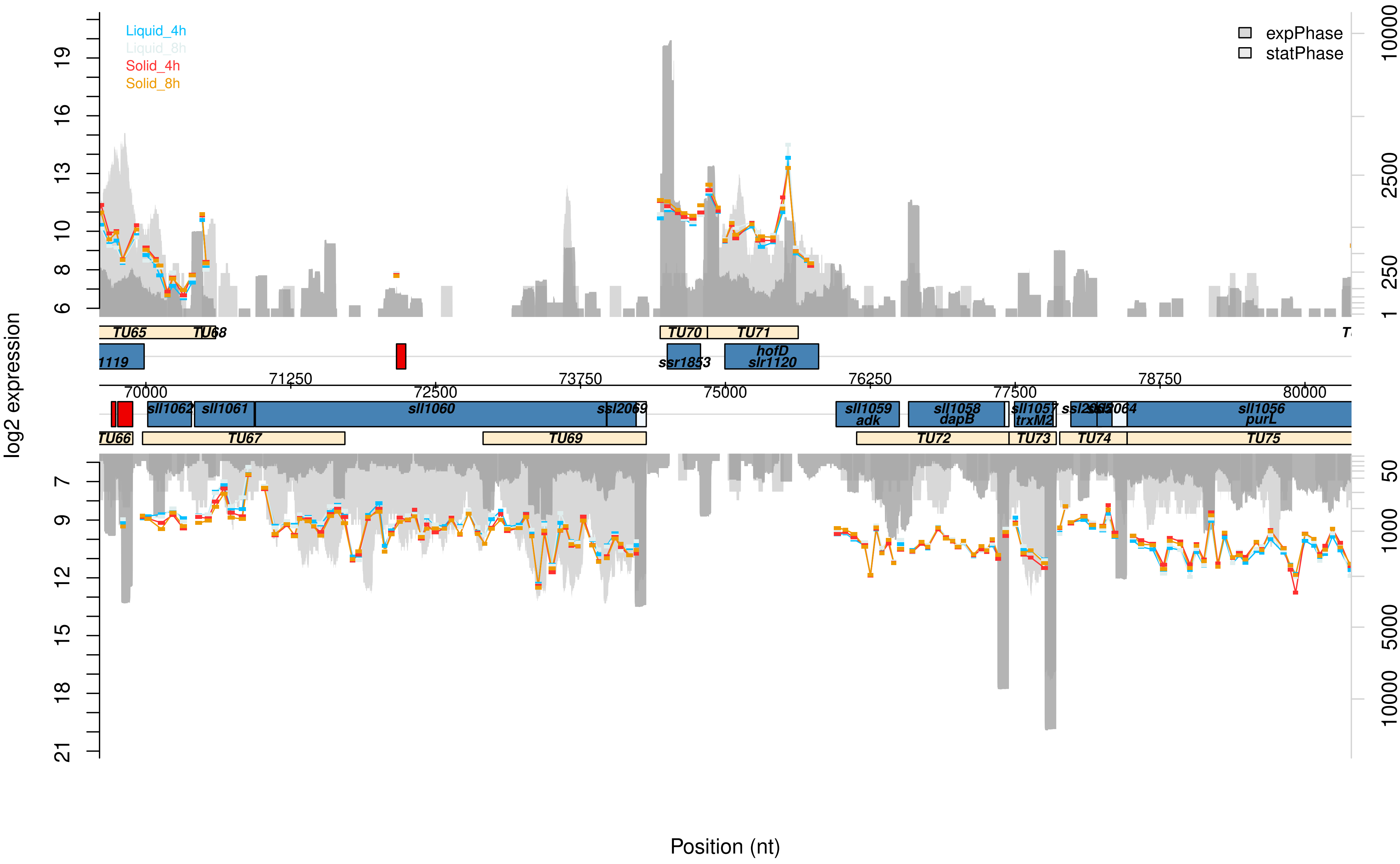

Liquid_4h
Liquid_8h
Solid_4h
Solid_8h
expPhase
statPhase
log2 expression
Position (nt)
TU65
TU68
TU70
TU71
TU66
TU67
TU69
TU72
TU73
TU74
TU75
1119
ssr1853
hofD
slr1120
sll1062
sll1061
sll1060
ssl2069
sll1059
adk
sll1058
dapB
sll1057
trxM2
sll1056
purL
70000
71250
72500
73750
75000
76250
77500
78750
80000

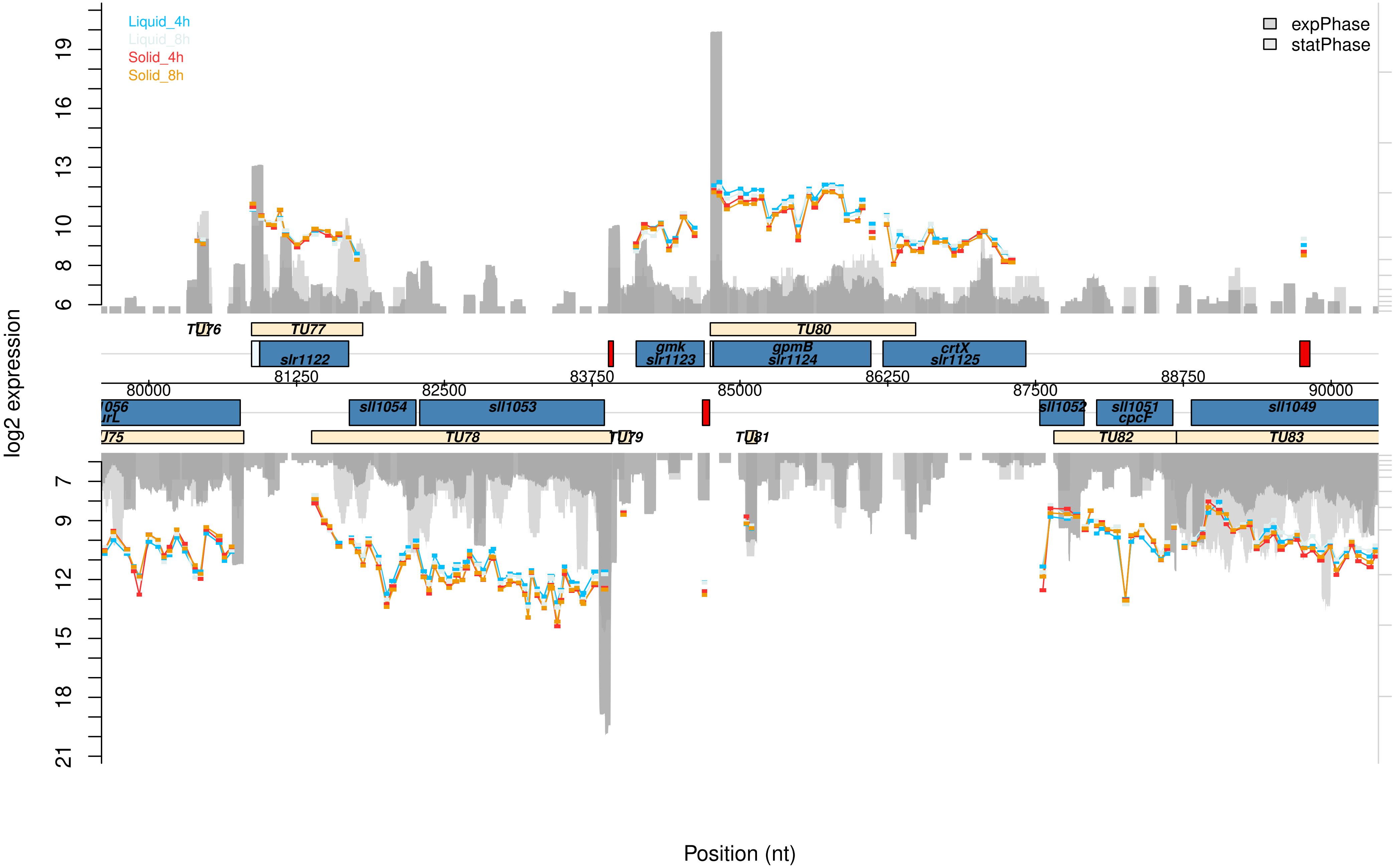

Liquid_4h
Liquid_8h
Solid_4h
Solid_8h
expPhase
statPhase
log2 expression
Position (nt)
TU76
TU77
TU80
slr1122
gmk
slr1123
gpmB
slr1124
crtX
slr1125
81250
83750
86250
88750
80000
82500
85000
87500
90000
sll1054
sll1053
sll1052
sll1051
cpcF
sll1049
TU78
TU79
TU81
TU82
TU83

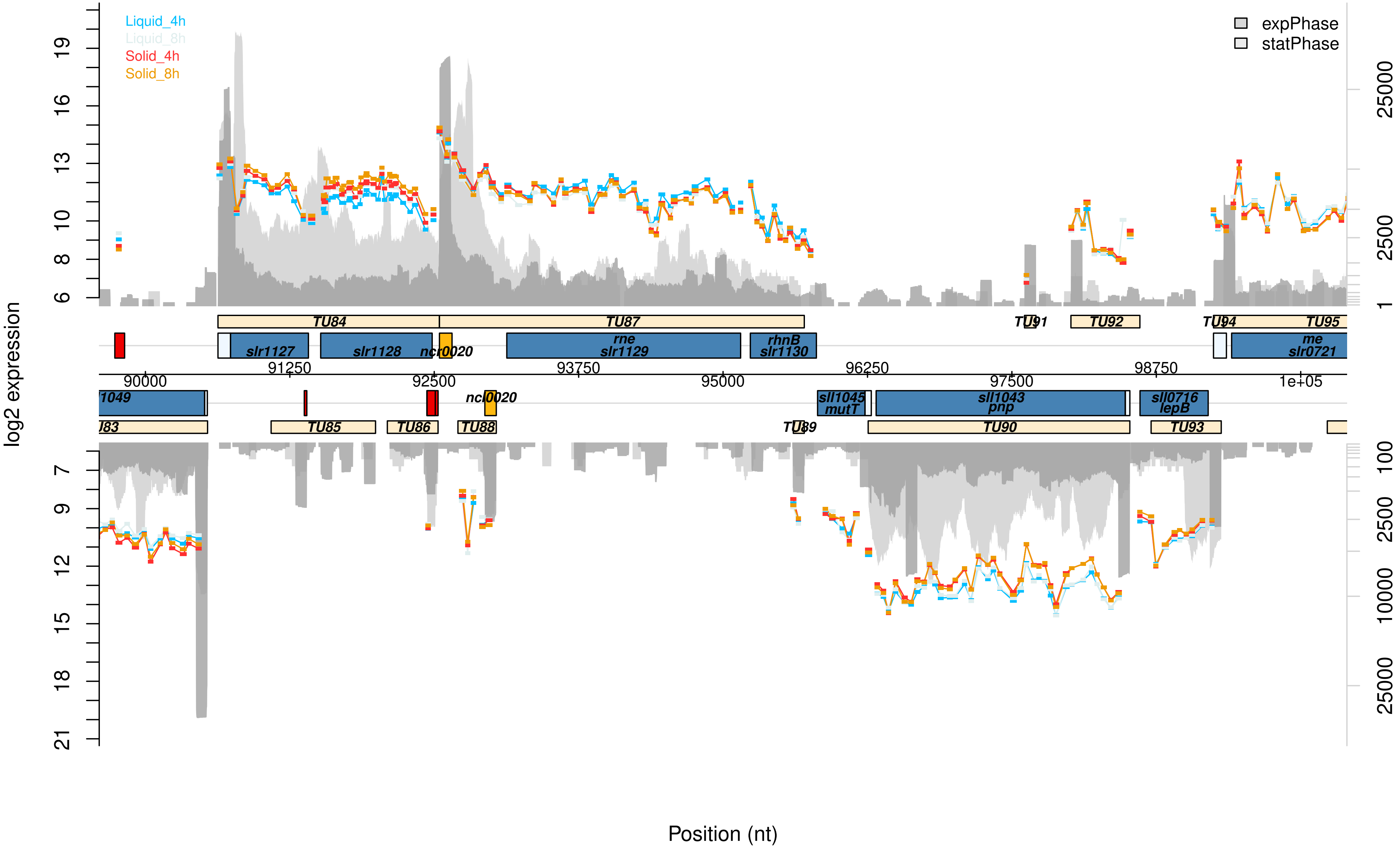

Liquid_4h
Liquid_8h
Solid_4h
Solid_8h
expPhase
statPhase
log2 expression
Position (nt)
TU84
TU87
TU91
TU92
TU94
TU95
slr1127
slr1128
ncr0020
rne
slr1129
rhnB
slr1130
me
slr0721
90000
91250
92500
93750
95000
96250
97500
98750
1e+05
sll1045
mutT
sll1043
pnp
sll0716
lepB
TU85
TU86
TU88
TU89
TU90
TU93

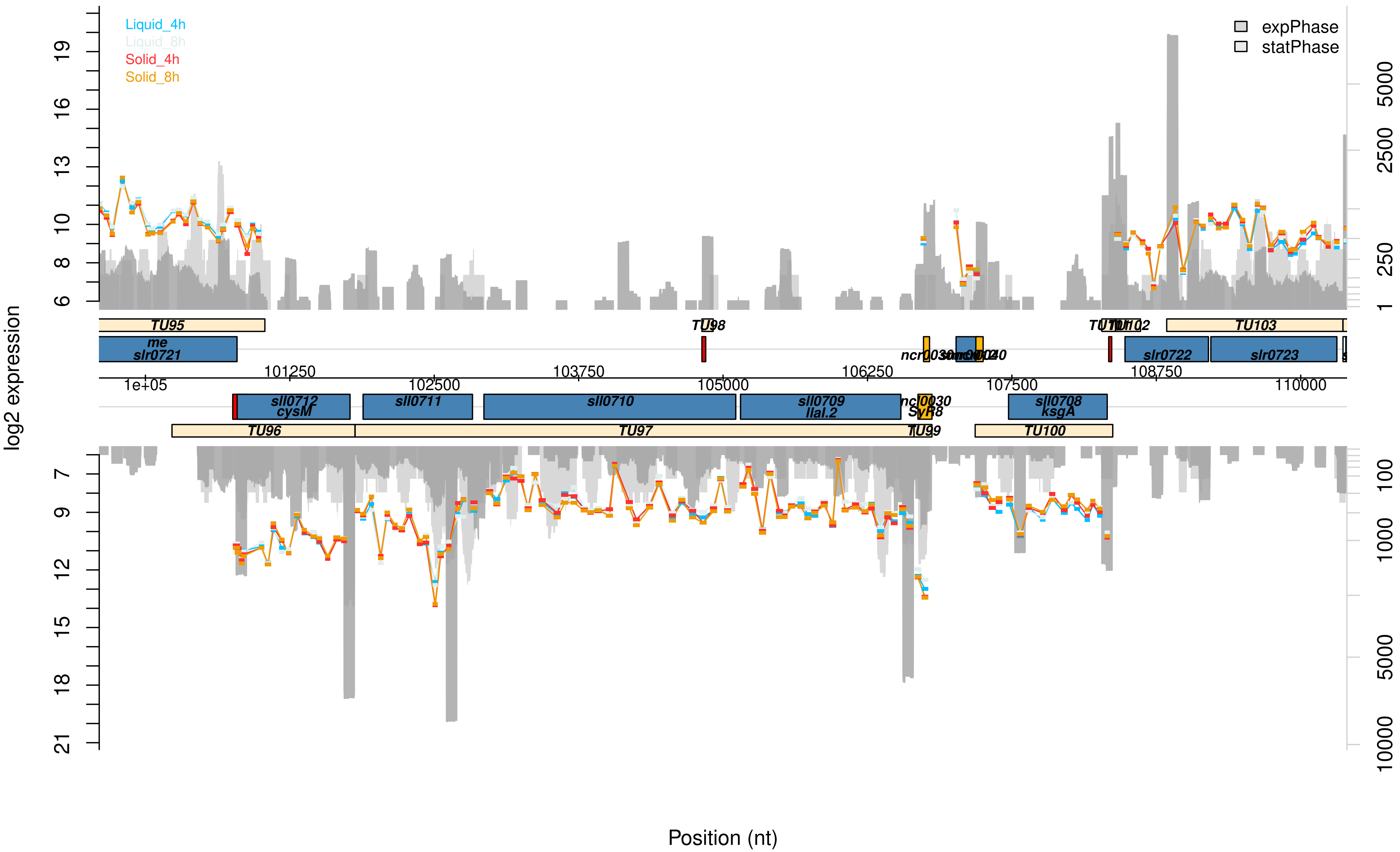

Liquid_4h
Liquid_8h
Solid_4h
Solid_8h
expPhase
statPhase
log2 expression
Position (nt)
6
8
10
13
16
19
7
9
12
15
18
21
1
250
2500
5000
100
1000
10000
TU95
TU98
TU102
TU103
me
slr0721
slr0722
slr0723
101250
103750
106250
108750
1e+05
102500
105000
107500
110000
sll0712
cysM
sll0711
sll0710
sll0709
llaL.2
ncl0030
SyR8
sll0708
ksgA
TU96
TU97
TU99
TU100

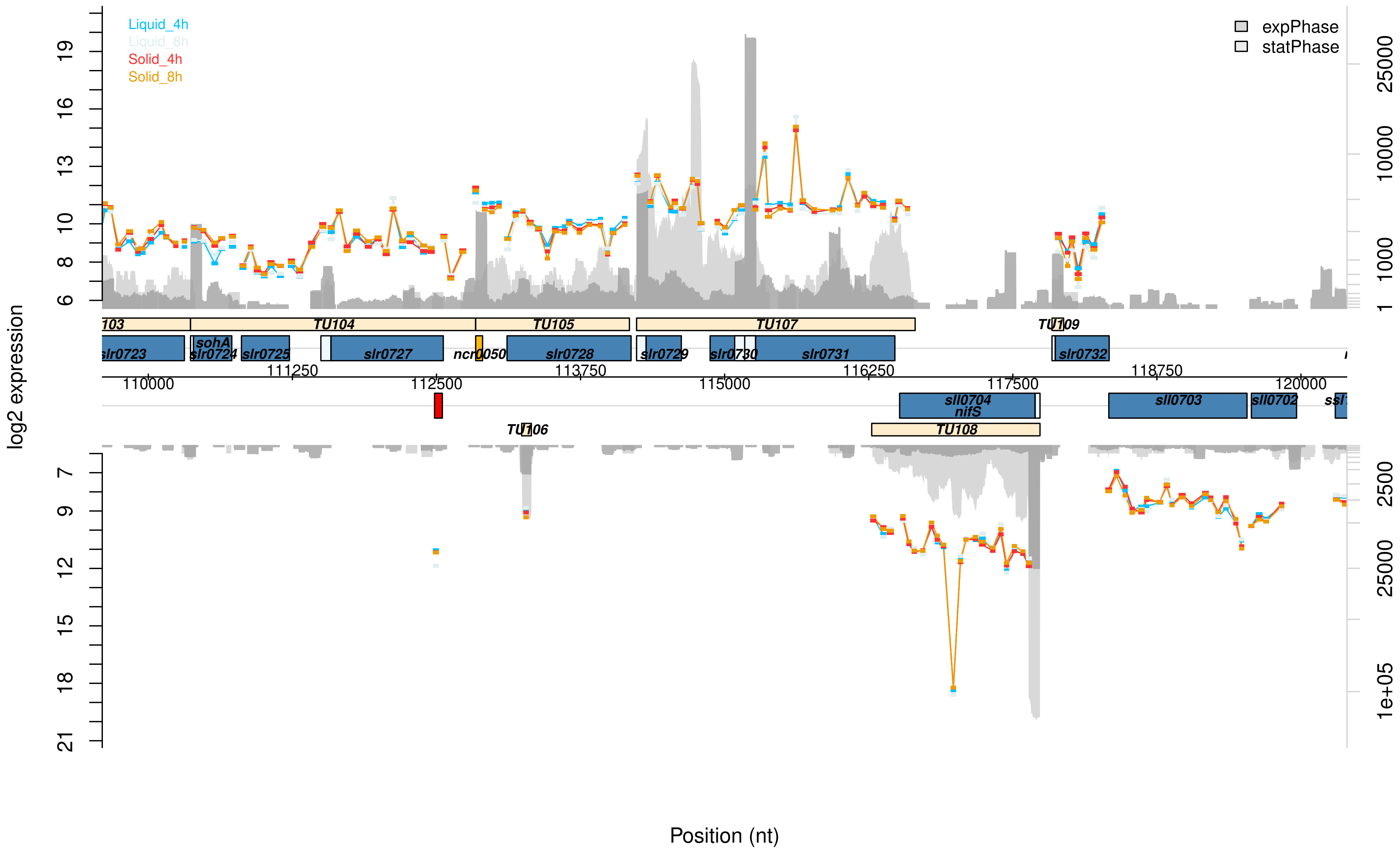

Liquid_4h
Liquid_8h
Solid_4h
Solid_8h
expPhase
statPhase
log2 expression
Position (nt)
TU104
TU105
TU107
TU109
TU106
TU108
sohA
slr0723
slr0724
slr0725
slr0727
ncr0050
slr0728
slr0729
slr0730
slr0731
slr0732
sll0704
nifS
sll0703
sll0702
110000
111250
112500
113750
115000
116250
117500
118750
120000

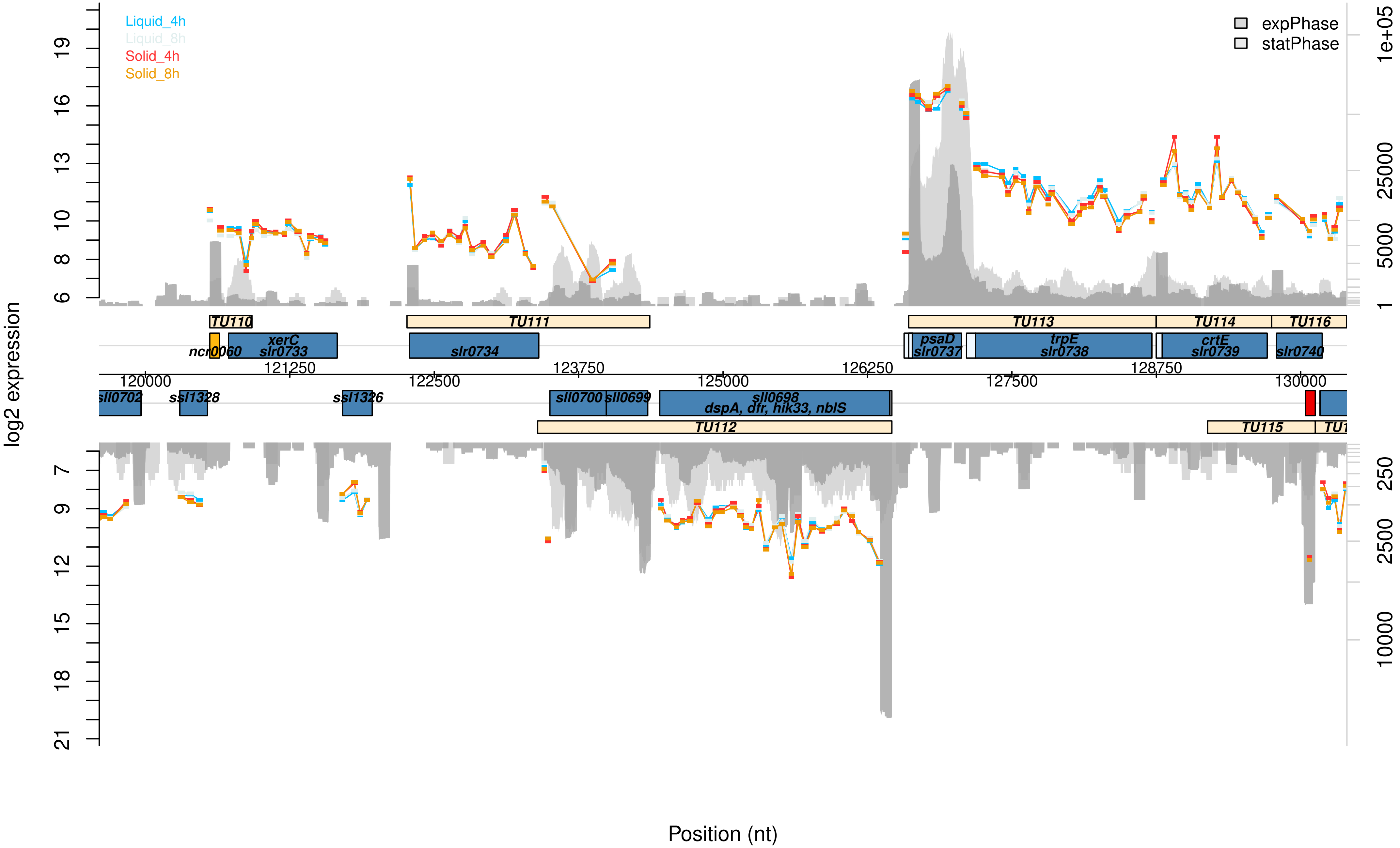

Liquid_4h
Liquid_8h
Solid_4h
Solid_8h
expPhase
statPhase
log2 expression
Position (nt)
TU110
TU111
TU112
TU113
TU114
TU115
TU116
ncr0060
xerC
slr0733
slr0734
psaD
slr0737
trpE
slr0738
crtE
slr0739
slr0740
sll0702
ssl1328
ssl1326
sll0700
sll0699
sll0698
dspA, dfr, hik33, nblS

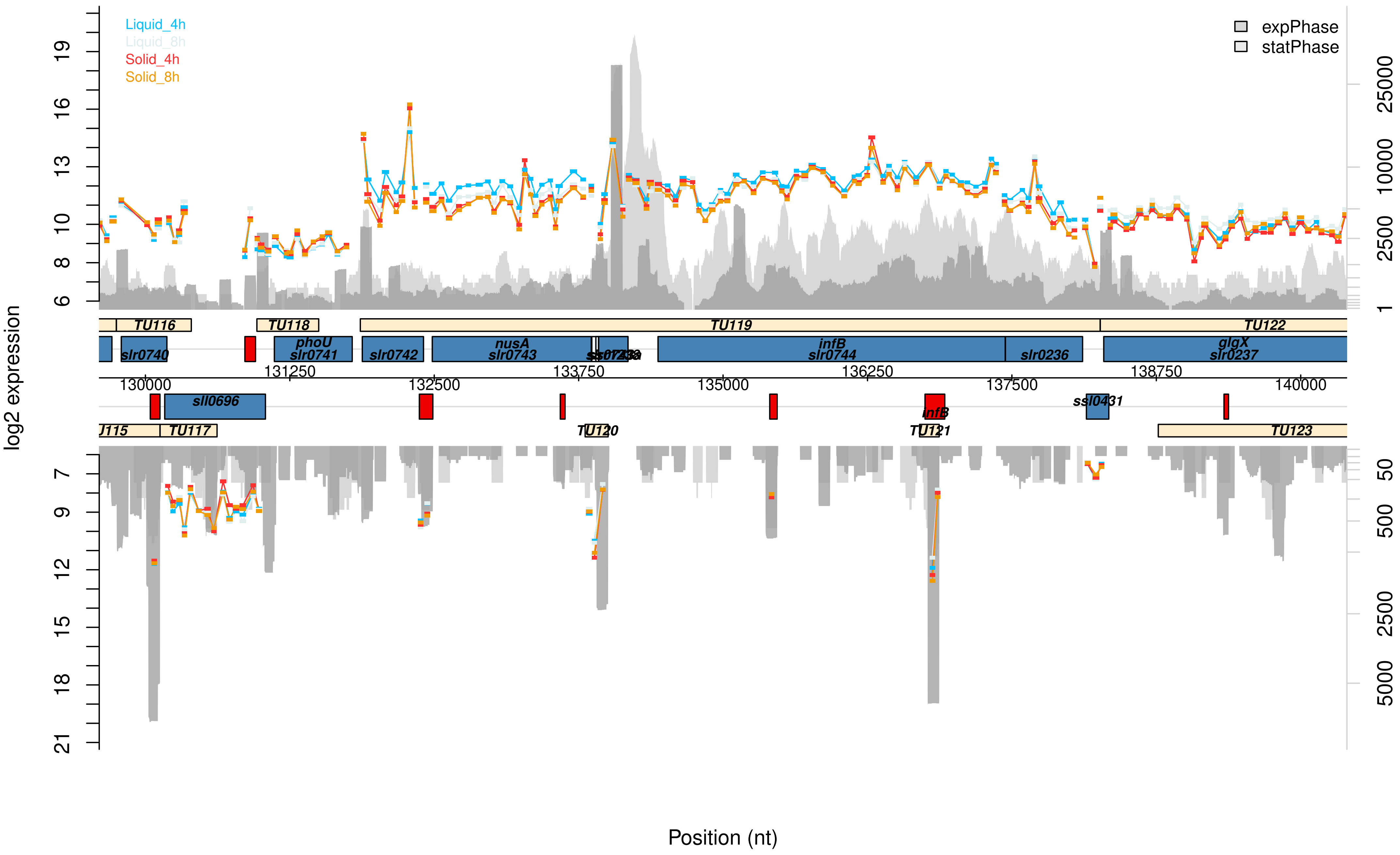

Liquid_4h
Liquid_8h
Solid_4h
Solid_8h
expPhase
statPhase
log2 expression
Position (nt)
TU116
TU118
TU119
TU122
slr0740
phoU
slr0741
slr0742
nusA
slr0743
infB
slr0744
slr0236
glgX
slr0237
sll0696
ssl0431
TU117
TU120
TU121
TU123
130000
131250
132500
133750
135000
136250
137500
138750
140000

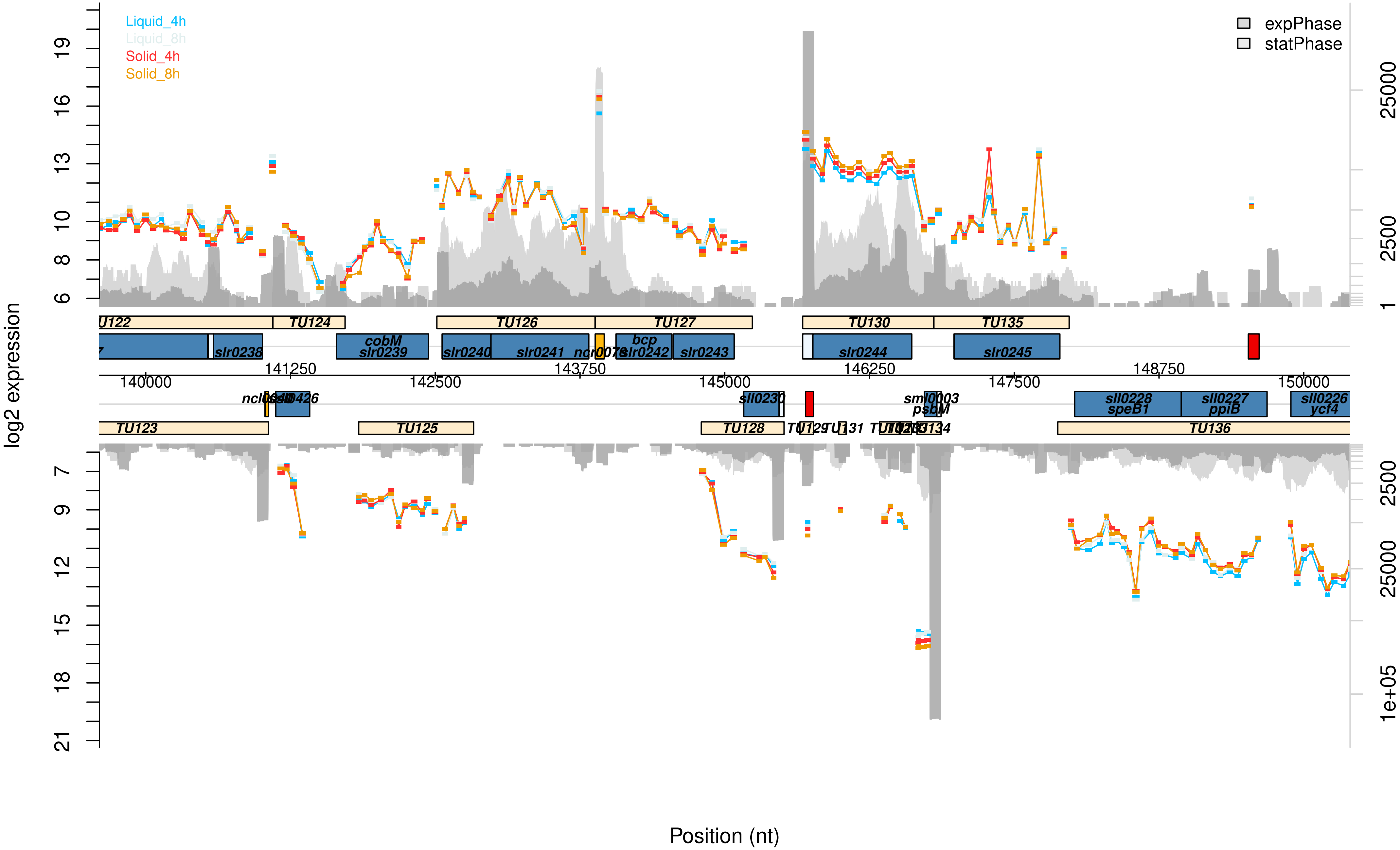
Liquid_4h
Liquid_8h
Solid_4h
Solid_8h
expPhase
statPhase
log2 expression
Position (nt)
TU122
TU124
TU126
TU127
TU130
TU135
cobM
slr0239
slr0238
slr0240
slr0241
ncr0078
bcp
slr0242
slr0243
slr0244
slr0245
140000
141250
142500
143750
145000
146250
147500
148750
150000
ncl0350
sll0426
sll0230
sml0003
psbM
sll0228
speB1
sll0227
ppiB
sll0226
ycf4
TU123
TU125
TU128
TU136

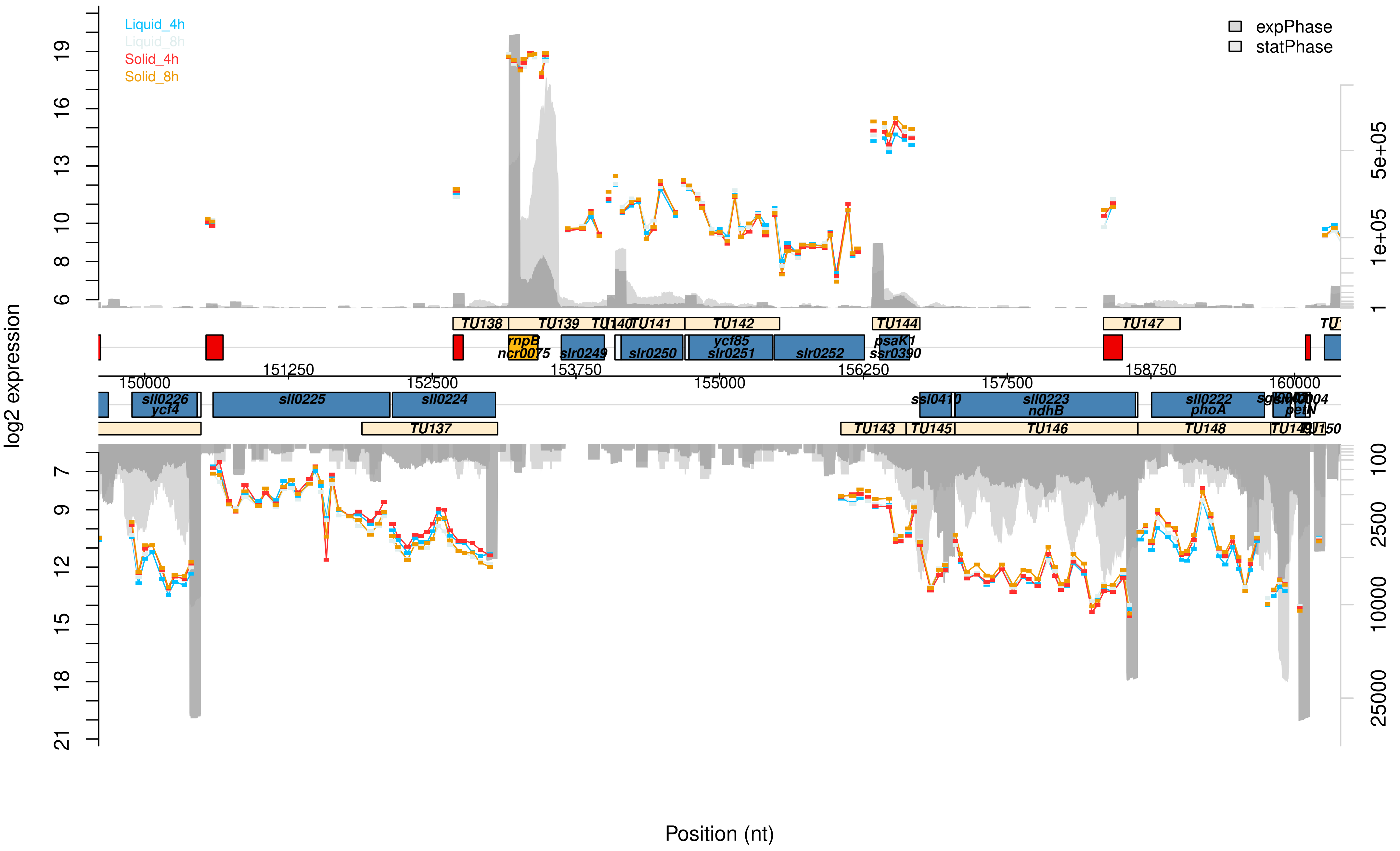

Liquid_4h
Liquid_8h
Solid_4h
Solid_8h
expPhase
statPhase
log2 expression
Position (nt)
TU137
TU138
TU139
TU140
TU141
TU142
TU143
TU144
TU145
TU146
TU147
TU148
sll0226
ycf4
sll0225
sll0224
rnpB
ncr0075
slr0249
slr0250
ycf85
slr0251
slr0252
psaK1
ssr0390
ssl0410
sll0223
ndhB
sll0222
phoA
petN
150000
151250
152500
153750
155000
156250
157500
158750
160000

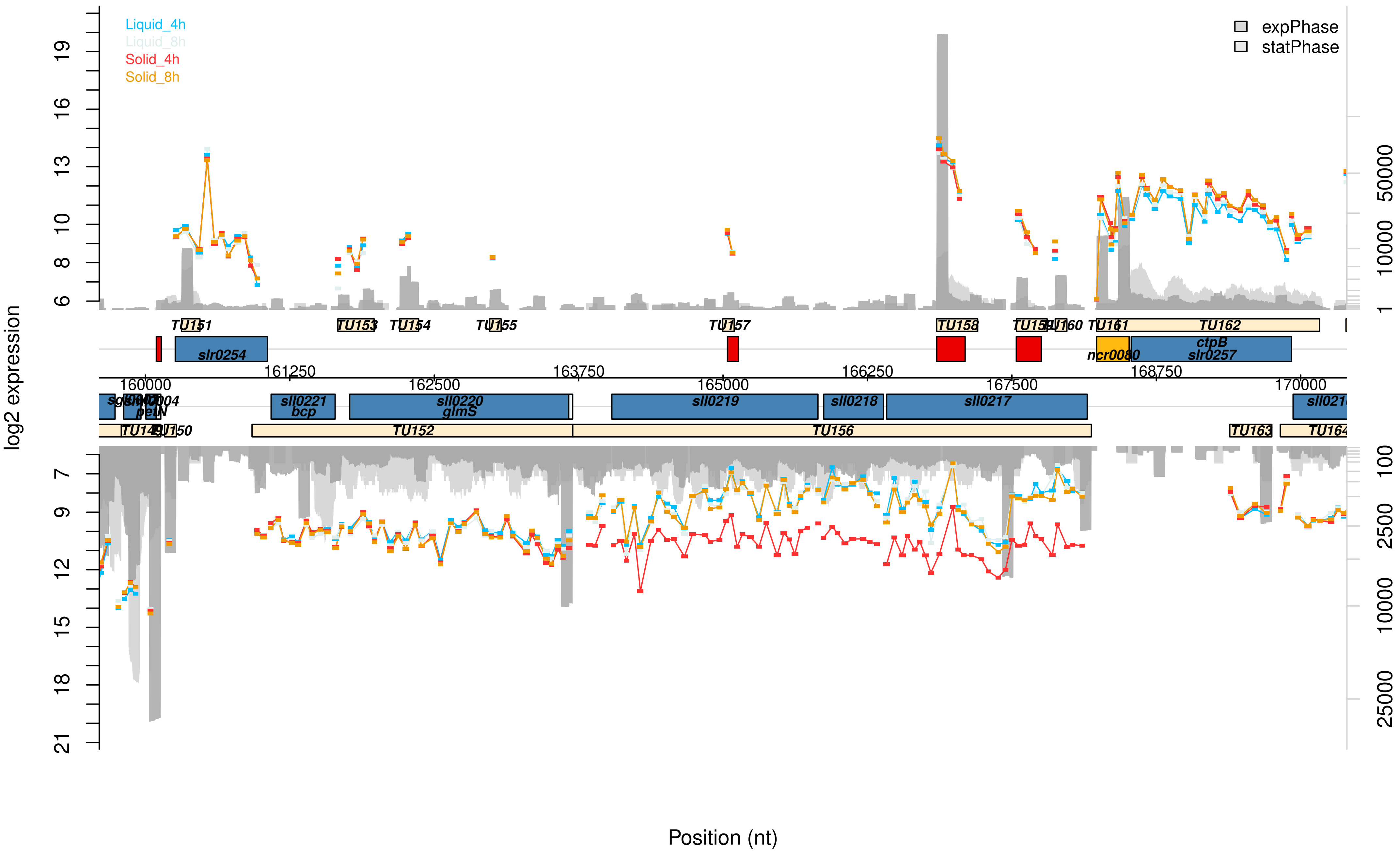

Liquid_4h
Liquid_8h
Solid_4h
Solid_8h
expPhase
statPhase
log2 expression
Position (nt)
TU151
TU153
TU154
TU155
TU157
TU158
TU160
TU162
slr0254
ncr0080
ctpB
slr0257
160000
161250
162500
163750
165000
166250
167500
168750
170000
sll0221
bcp
sll0220
glmS
sll0219
sll0218
sll0217
petN
TU150
TU152
TU156
TU163

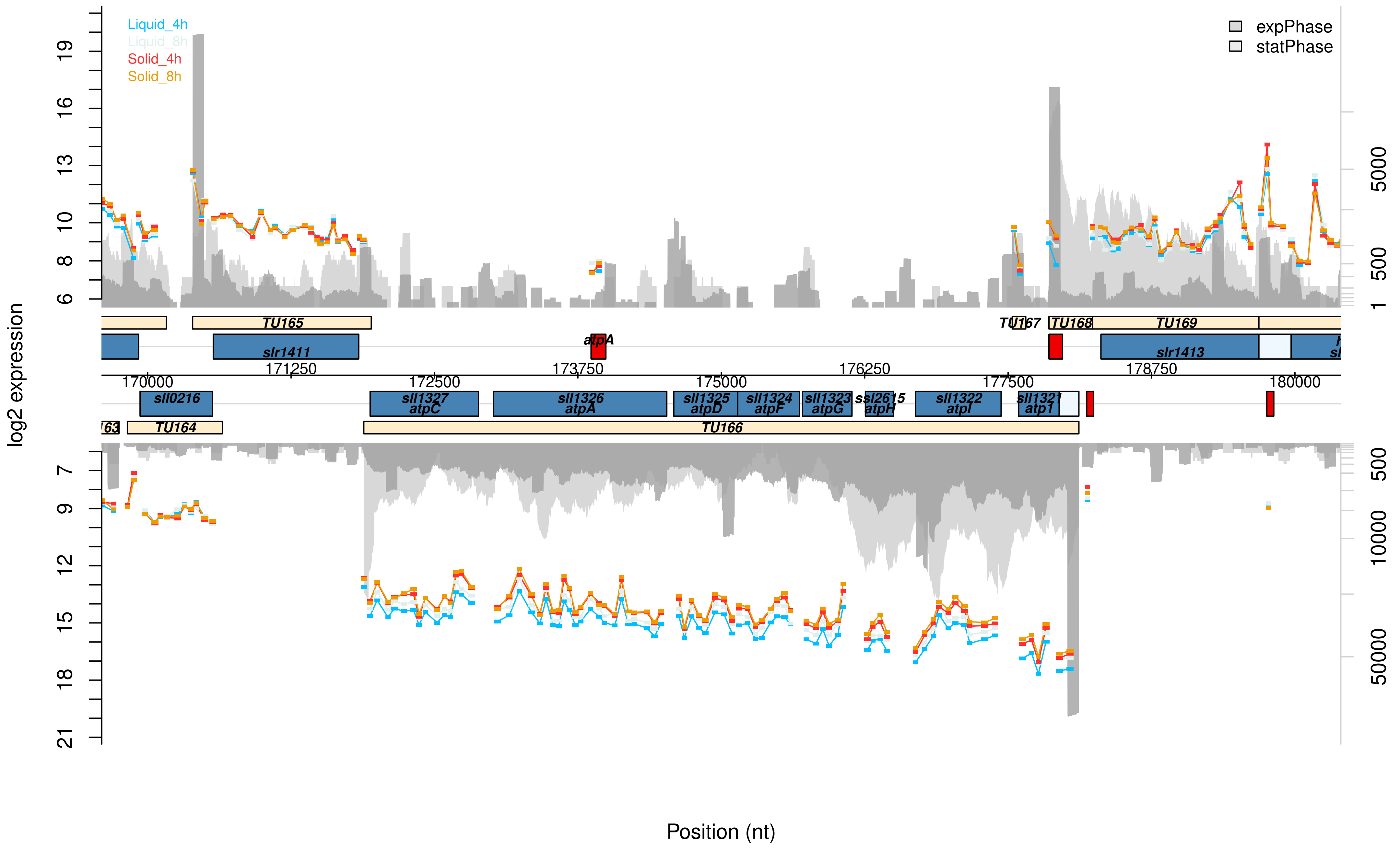

Liquid_4h
Liquid_8h
Solid_4h
Solid_8h
expPhase
statPhase
log2 expression
Position (nt)
TU165
TU167
TU168
TU169
TU164
TU166
slr1411
atpA
slr1413
sll0216
sll1327 atpC
sll1326 atpA
sll1325 atpD
sll1324 atpF
sll1323 atpG
ssl2615 atpH
sll1322 atpI
sll1321 atp1
170000
171250
172500
173750
175000
176250
177500
178750
180000

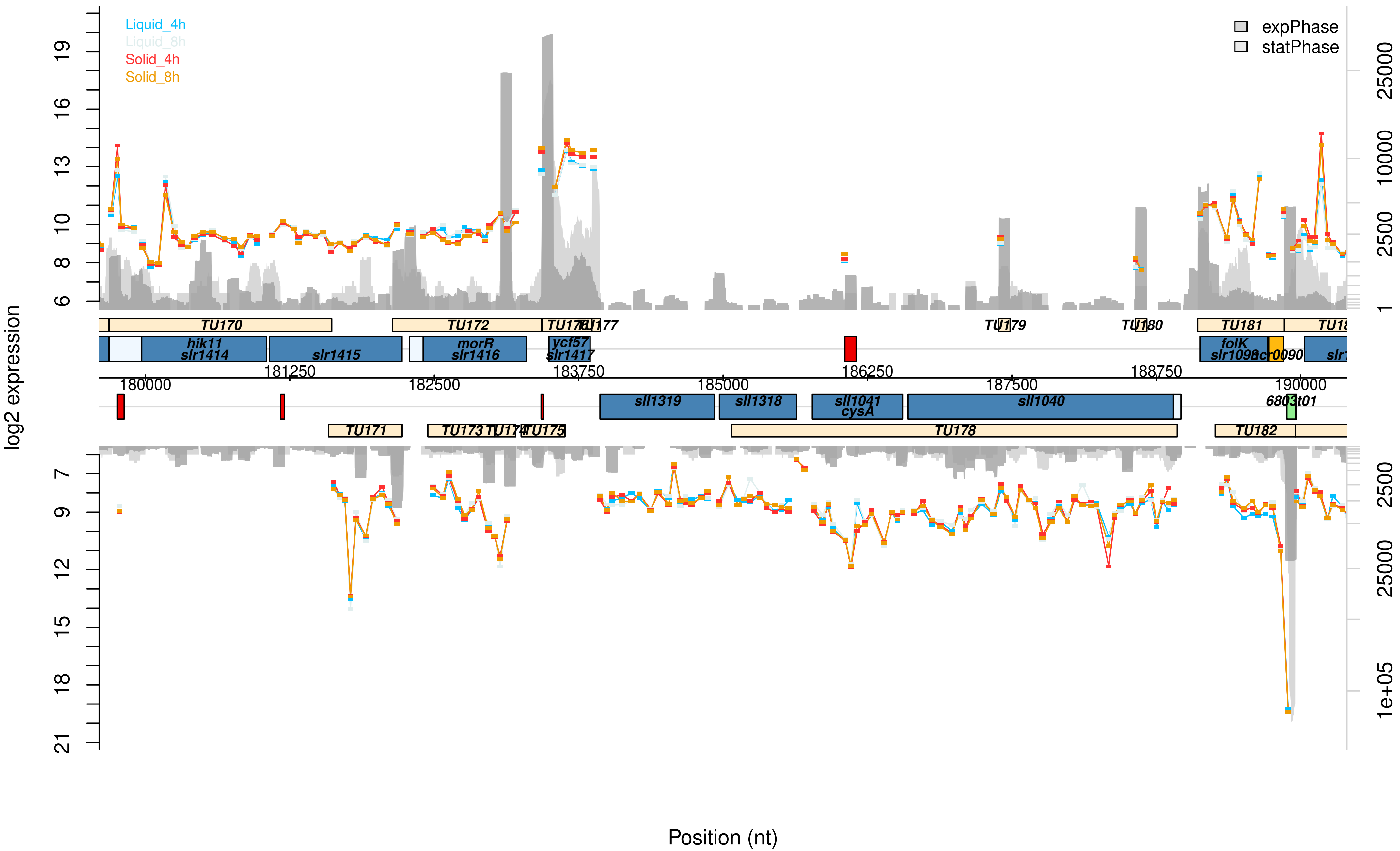
Liquid_4h
Liquid_8h
Solid_4h
Solid_8h
expPhase
statPhase
log2 expression
Position (nt)
TU170
TU172
TU176
TU177
TU179
TU180
TU181
TU171
TU173
TU174
TU175
TU178
TU182
hik11
slr1414
slr1415
morR
slr1416
ycf57
slr1417
folK
slr1098
ssr0090
sll1319
sll1318
sll1041
cysA
sll1040
6803t01
180000
181250
182500
183750
185000
186250
187500
188750
190000

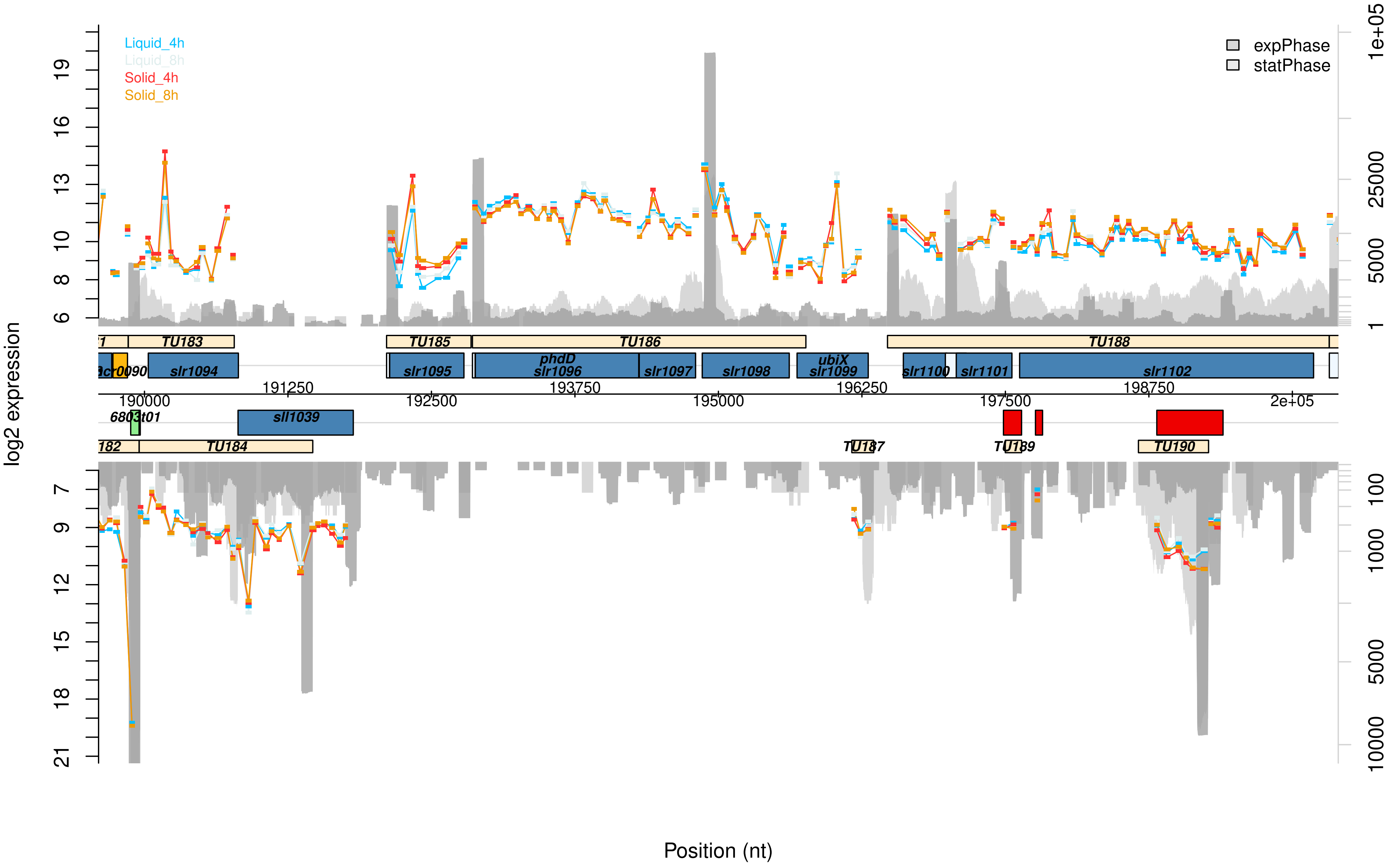

Liquid_4h
Liquid_8h
Solid_4h
Solid_8h
expPhase
statPhase
log2 expression
Position (nt)
TU183
TU185
TU186
TU188
slr1094
slr1095
phdD
slr1096
slr1097
slr1098
ubiX
slr1099
slr1100
slr1101
slr1102
sll1039
TU184
TU187
TU189
TU190
190000
191250
192500
193750
195000
196250
197500
198750
2e+05
1e+05
25000
5000
1
100
1000
10000

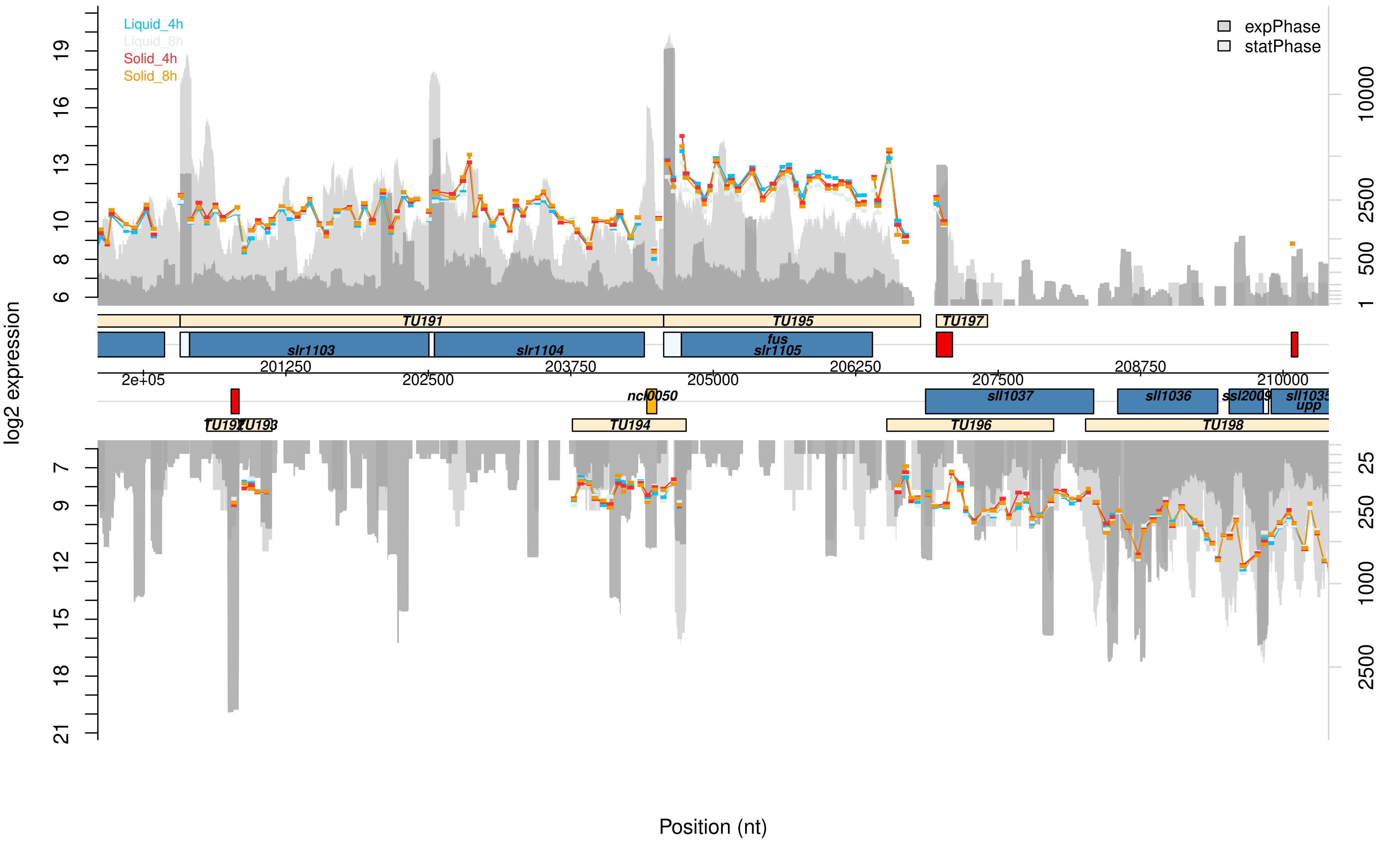

Liquid_4h
Liquid_8h
Solid_4h
Solid_8h
expPhase
statPhase
log2 expression
TU191
TU195
TU197
slr1103
slr1104
fus
slr1105
201250
203750
206250
208750
2e+05
202500
205000
207500
210000
ncl0050
sll1037
sll1036
ssl2009
sll1035
upp
TU192
TU193
TU194
TU196
TU198
Position (nt)

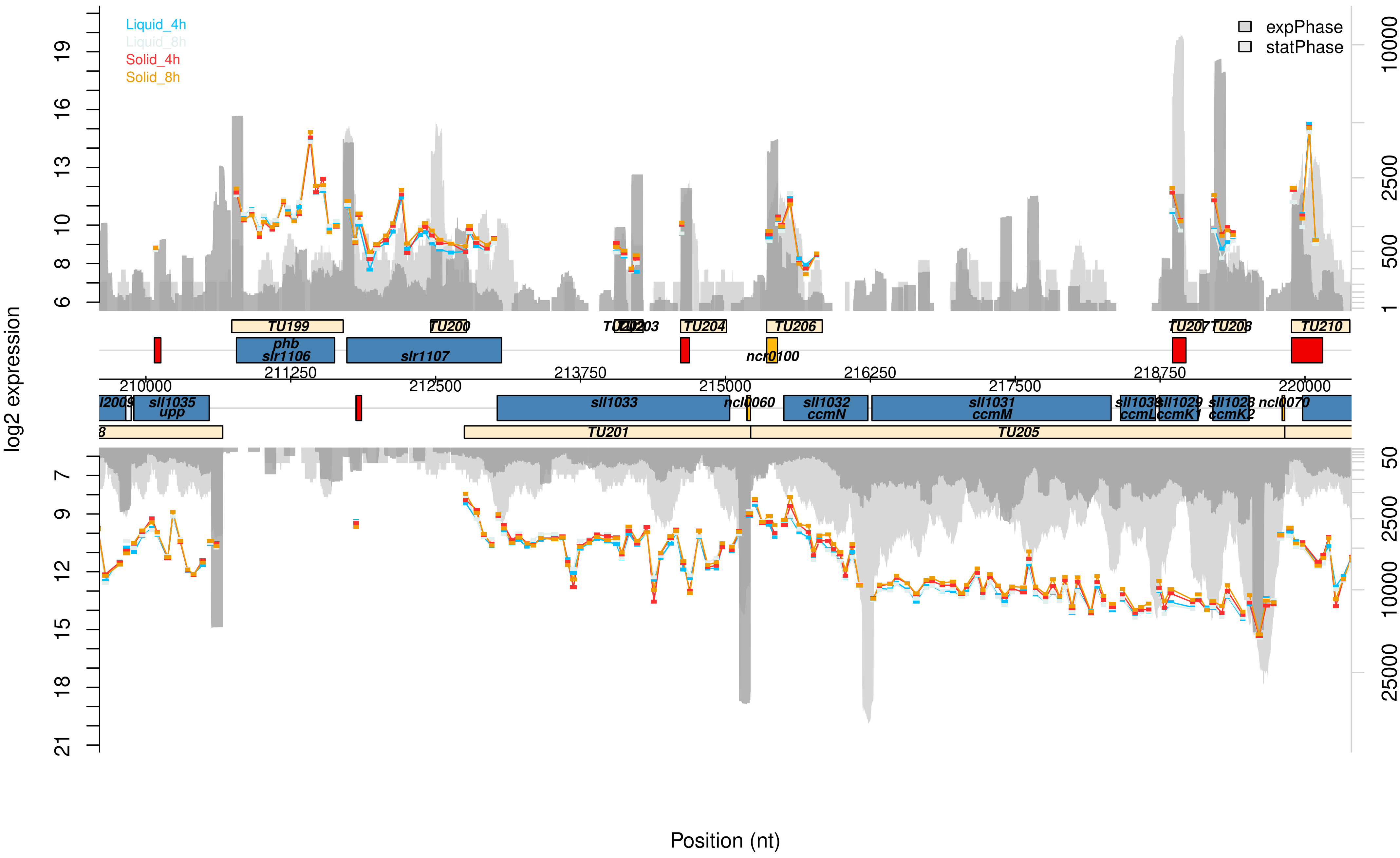

Liquid_4h
Liquid_8h
Solid_4h
Solid_8h
expPhase
statPhase
log2 expression
Position (nt)
TU199
TU200
TU202
TU203
TU204
TU206
TU207
TU208
TU210
phb
slr1106
slr1107
ncr0100
210000
211250
212500
213750
215000
216250
217500
218750
220000
sll1035
upp
sll1033
ncl0060
sll1032
ccmN
sll1031
ccmM
ccmL
sll1029
ccmK1
sll1028
ccmK2
ncl0070
TU201
TU205

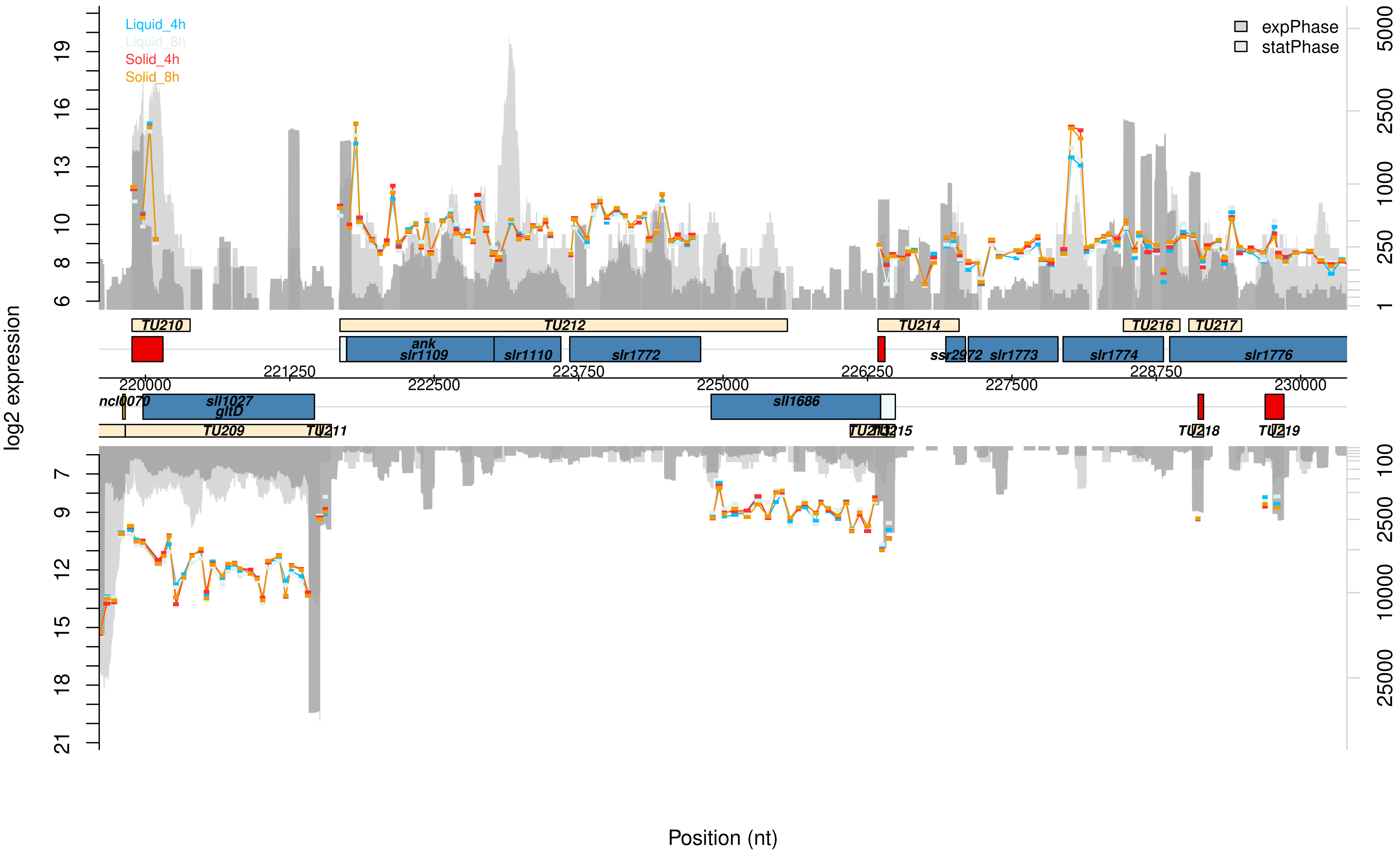

Liquid_4h
Liquid_8h
Solid_4h
Solid_8h
expPhase
statPhase
log2 expression
Position (nt)
TU210
TU212
TU214
TU216
TU217
ank
slr1109
slr1110
slr1772
ssr2972
slr1773
slr1774
slr1776
ncl0070
sll1027
gltD
sll1686
TU209
TU211
TU213
TU215
TU218
TU219
220000
221250
222500
223750
225000
226250
227500
228750
230000

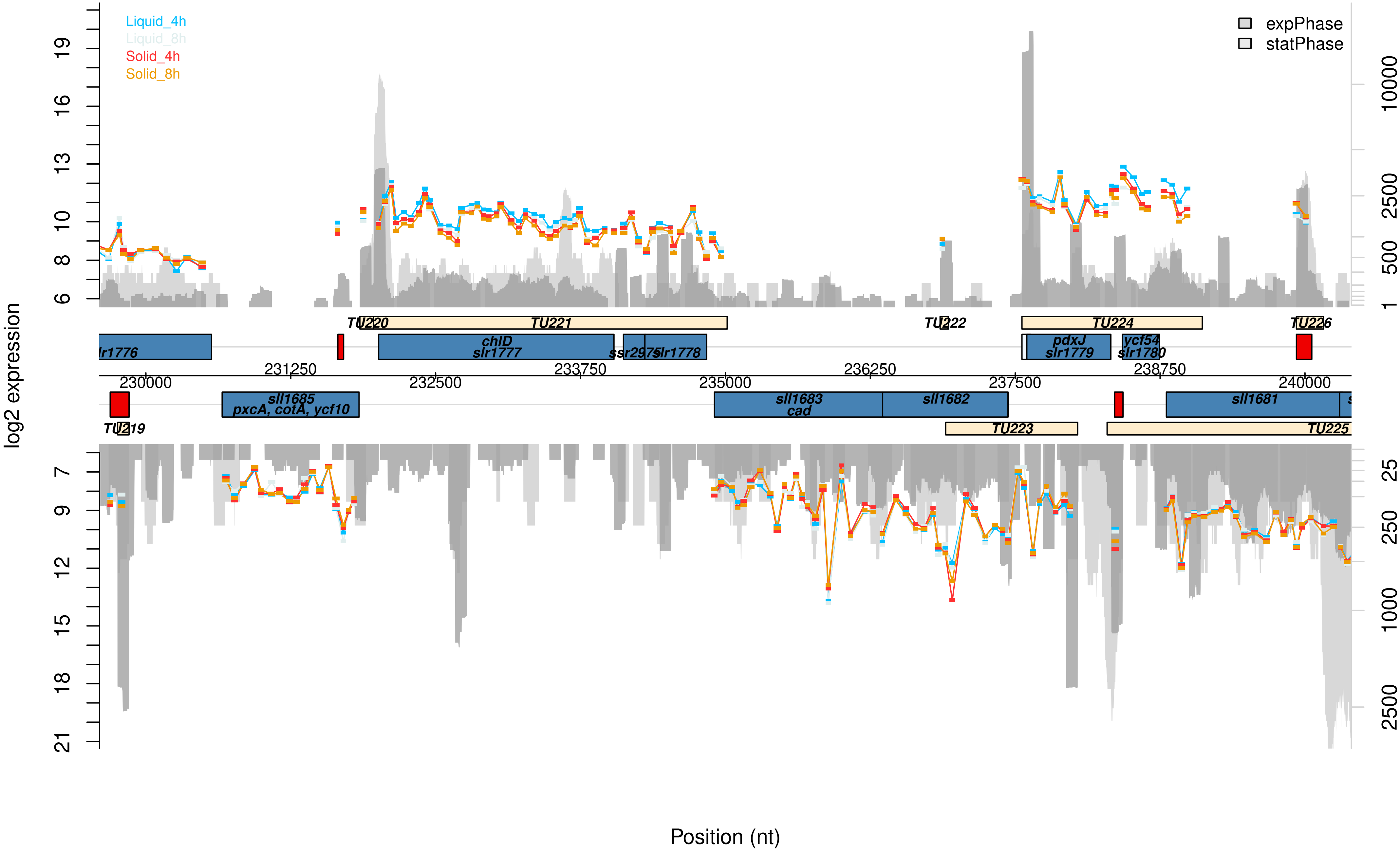

Liquid_4h
Liquid_8h
Solid_4h
Solid_8h
expPhase
statPhase
log2 expression
Position (nt)
TU219
TU220
TU221
TU222
TU223
TU224
TU225
TU226
chlD
slr1777
slr1778
pdxJ
slr1779
ycf54
slr1780
sll1685
pxcA, cotA, ycf10
sll1683
cad
sll1682
sll1681
230000
231250
232500
233750
235000
236250
237500
238750
240000

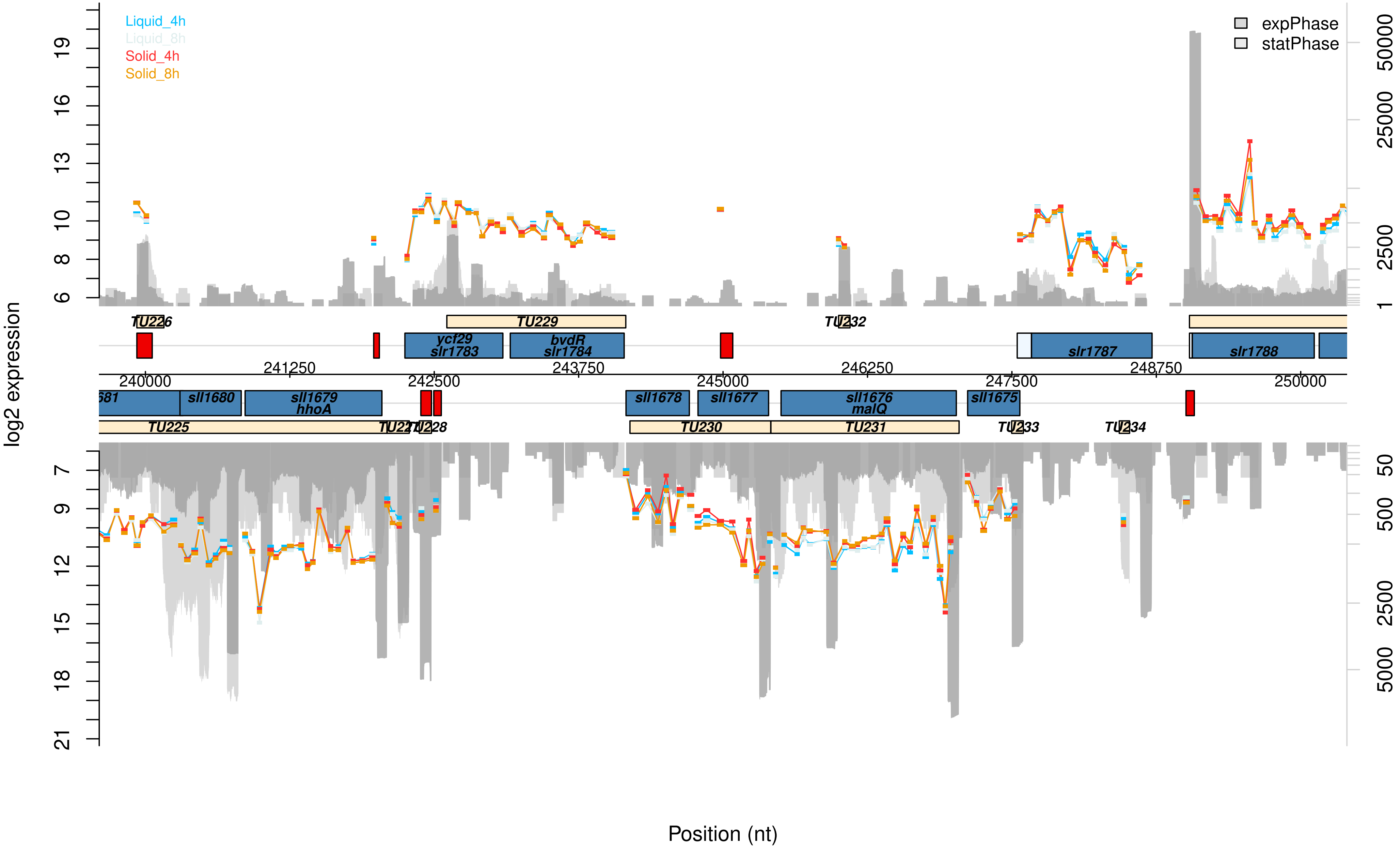

Liquid_4h
Liquid_8h
Solid_4h
Solid_8h
expPhase
statPhase
log2 expression
Position (nt)
TU226
TU229
TU232
ycf29
slr1783
bvdR
slr1784
slr1787
slr1788
240000
241250
242500
243750
245000
246250
247500
248750
250000
sll1680
sll1679
hhoA
sll1678
sll1677
sll1676
malQ
sll1675
TU225
TU230
TU231
TU233
TU234

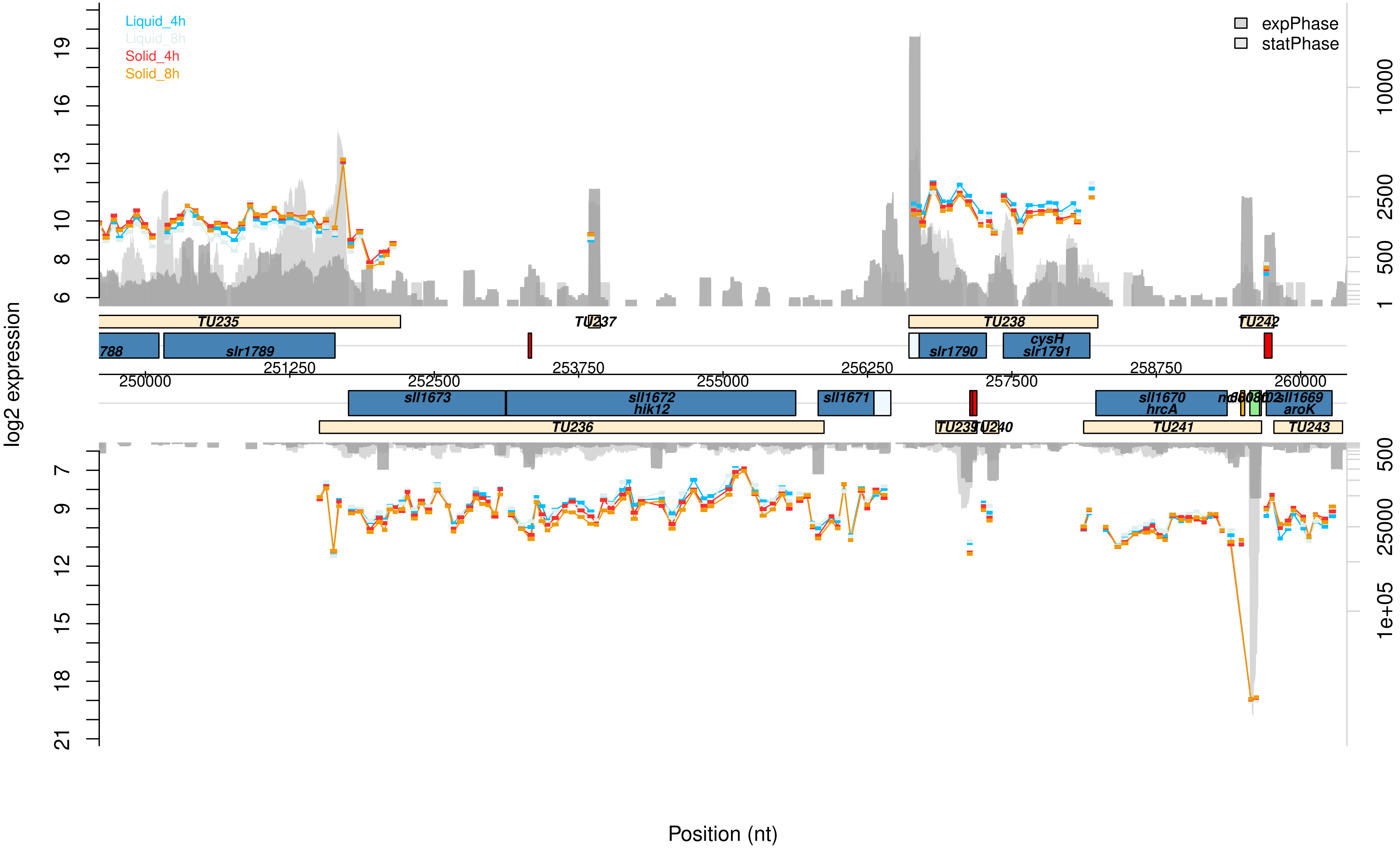

Liquid_4h
Liquid_8h
Solid_4h
Solid_8h
expPhase
statPhase
log2 expression
Position (nt)
TU235
TU237
TU238
TU242
788
slr1789
slr1790
cysH
slr1791
sll1673
sll1672
hik12
sll1671
sll1670
hrcA
sll1669
aroK
TU236
TU241
TU243
250000
251250
252500
253750
255000
256250
257500
258750
260000

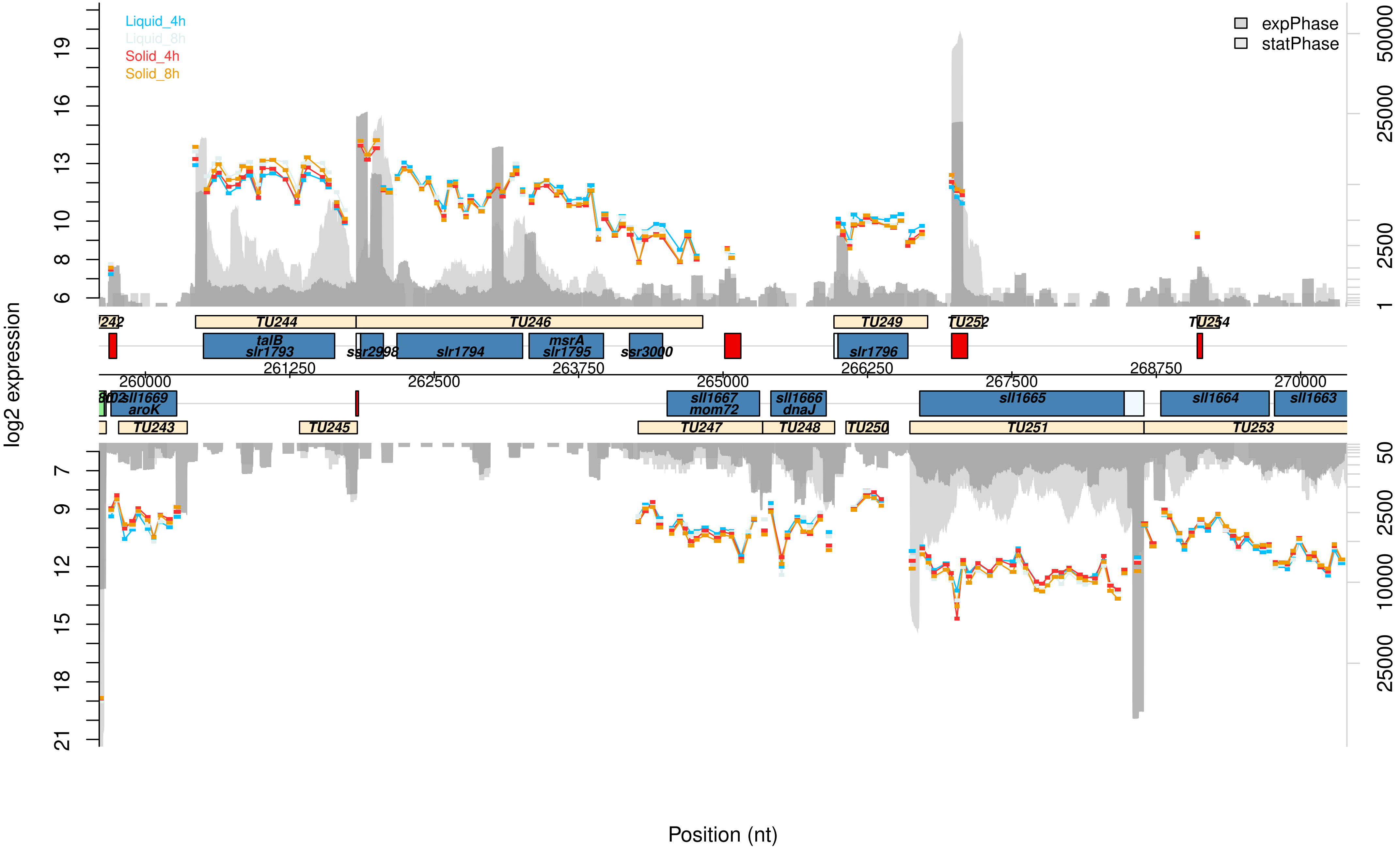

Liquid_4h
Liquid_8h
Solid_4h
Solid_8h
expPhase
statPhase
log2 expression
Position (nt)
TU244
TU246
TU249
TU252
TU254
talB
slr1793
ssr2998
slr1794
msrA
slr1795
ssr3000
slr1796
261250
263750
266250
268750
260000
262500
265000
267500
270000
sll1669
aroK
sll1667
mom72
sll1666
dnaJ
sll1665
sll1664
sll1663
TU243
TU245
TU247
TU248
TU250
TU251
TU253

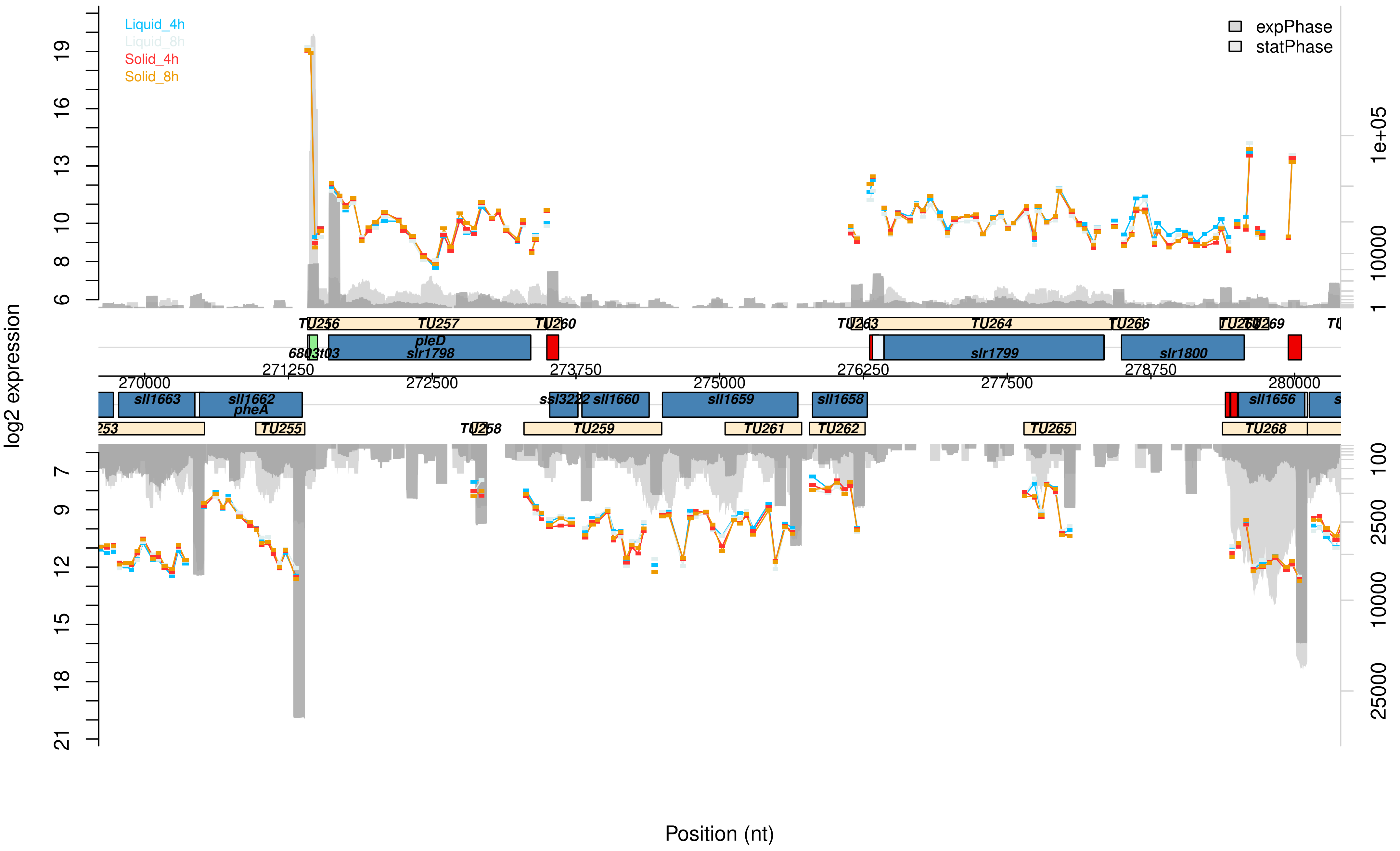

Liquid_4h
Liquid_8h
Solid_4h
Solid_8h
expPhase
statPhase
log2 expression
Position (nt)
TU256
TU257
TU260
TU263
TU264
TU266
TU269
6803t03
pleD
slr1798
slr1799
slr1800
sll1663
sll1662
pheA
ssl3222
sll1660
sll1659
sll1658
sll1656
TU253
TU255
TU258
TU259
TU261
TU262
TU265
TU268
270000
271250
272500
273750
275000
276250
277500
278750
280000

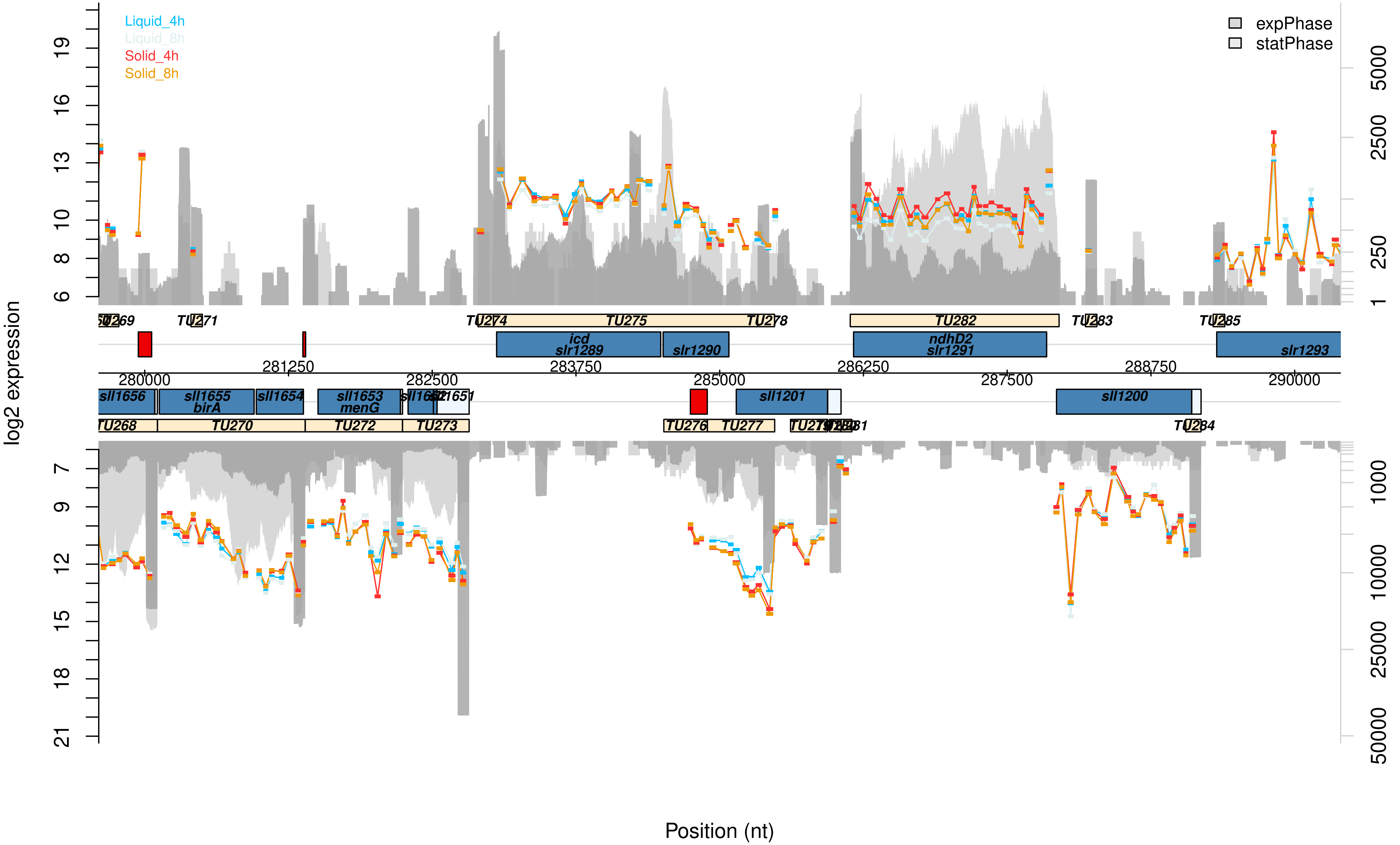

Liquid_4h
Liquid_8h
Solid_4h
Solid_8h
expPhase
statPhase
log2 expression
Position (nt)
sll1656
sll1655
birA
sll1654
sll1653
menG
icd
slr1289
slr1290
ndhD2
slr1291
slr1293
sll1201
sll1200
TU268
TU269
TU270
TU271
TU272
TU273
TU274
TU275
TU276
TU277
TU278
TU282
TU283
TU284
TU285
280000
281250
282500
283750
285000
286250
287500
288750
290000

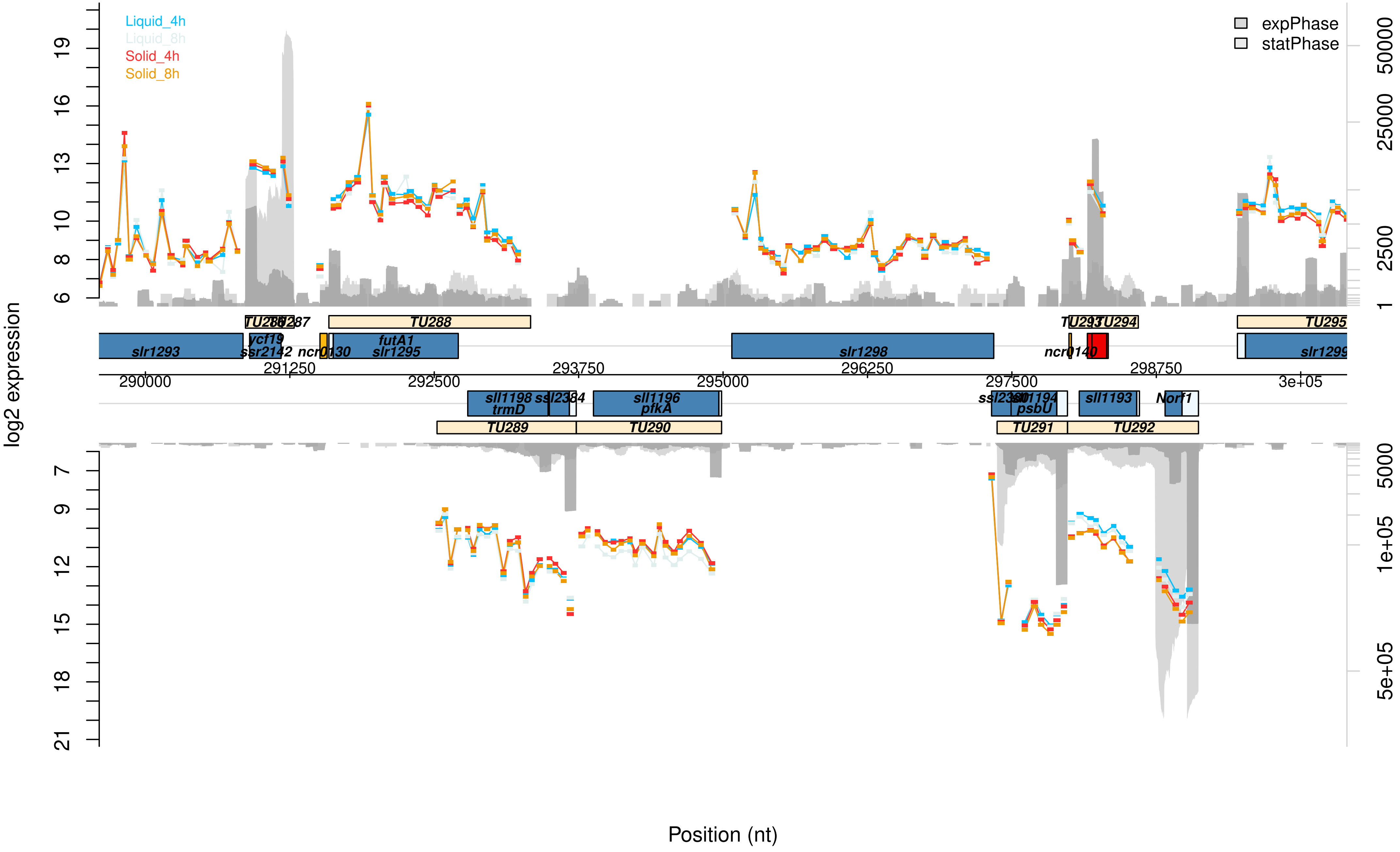

Liquid_4h
Liquid_8h
Solid_4h
Solid_8h
expPhase
statPhase
log2 expression
Position (nt)
TU288
TU289
TU290
TU291
TU292
TU294
TU295
slr1293
ycf19
ssr2142
futA1
slr1295
slr1298
slr1299
sll1198
trmD
sll1196
pfkA
psbU
sll1193
Norf1
290000
291250
292500
293750
295000
296250
297500
298750
3e+05
50000
25000
2500
5000
1e+05
5e+05

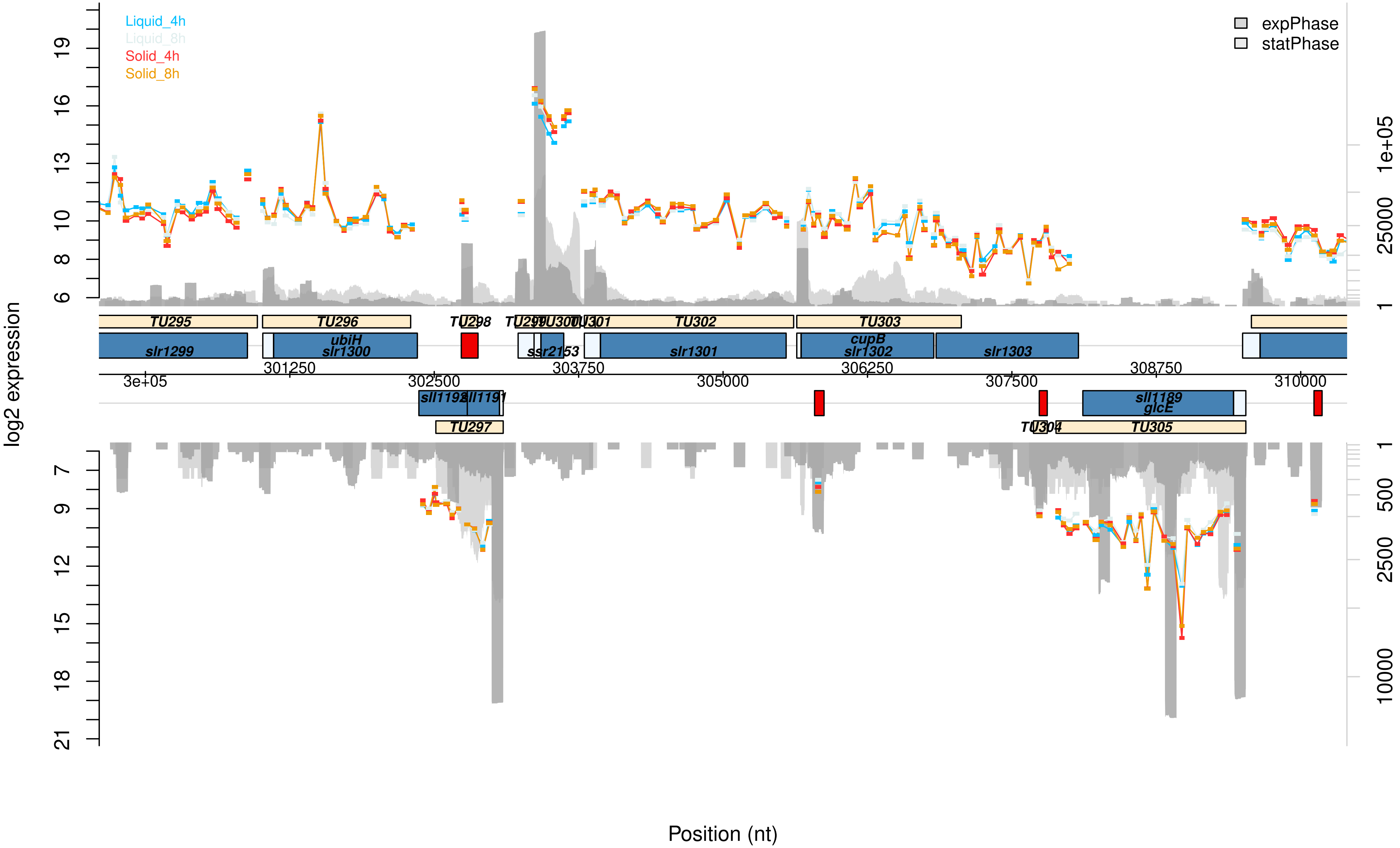

Liquid_4h
Liquid_8h
Solid_4h
Solid_8h
expPhase
statPhase
log2 expression
Position (nt)
TU295
TU296
TU298
TU299
TU300
TU301
TU302
TU303
TU297
TU304
TU305
slr1299
ubiH
slr1300
ssr2153
slr1301
cupB
slr1302
slr1303
sll1192
sll1191
sll1189
glcE
3e+05
301250
302500
303750
305000
306250
307500
308750
310000

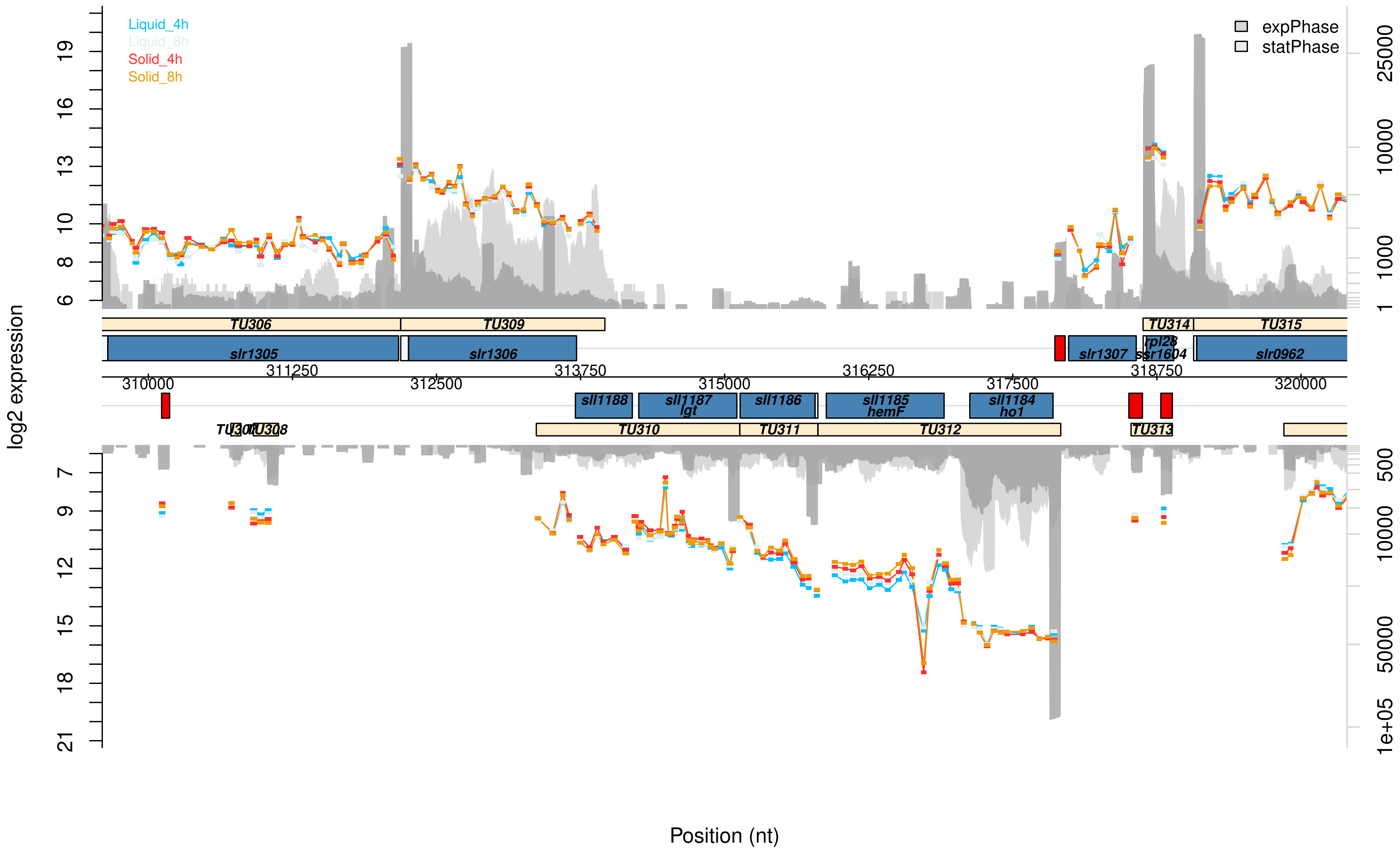
Liquid_4h
Liquid_8h
Solid_4h
Solid_8h
expPhase
statPhase
log2 expression
Position (nt)
TU306
TU309
TU314
TU315
slr1305
slr1306
slr1307
rpl28
ssr1604
slr0962
sll1188
sll1187
lgt
sll1186
sll1185
hemF
sll1184
ho1
TU307
TU308
TU310
TU311
TU312
TU313
310000
311250
312500
313750
315000
316250
317500
318750
320000
6
8
10
13
16
19
7
9
12
15
18
21
1
1000
10000
25000
500
50000
1e+05

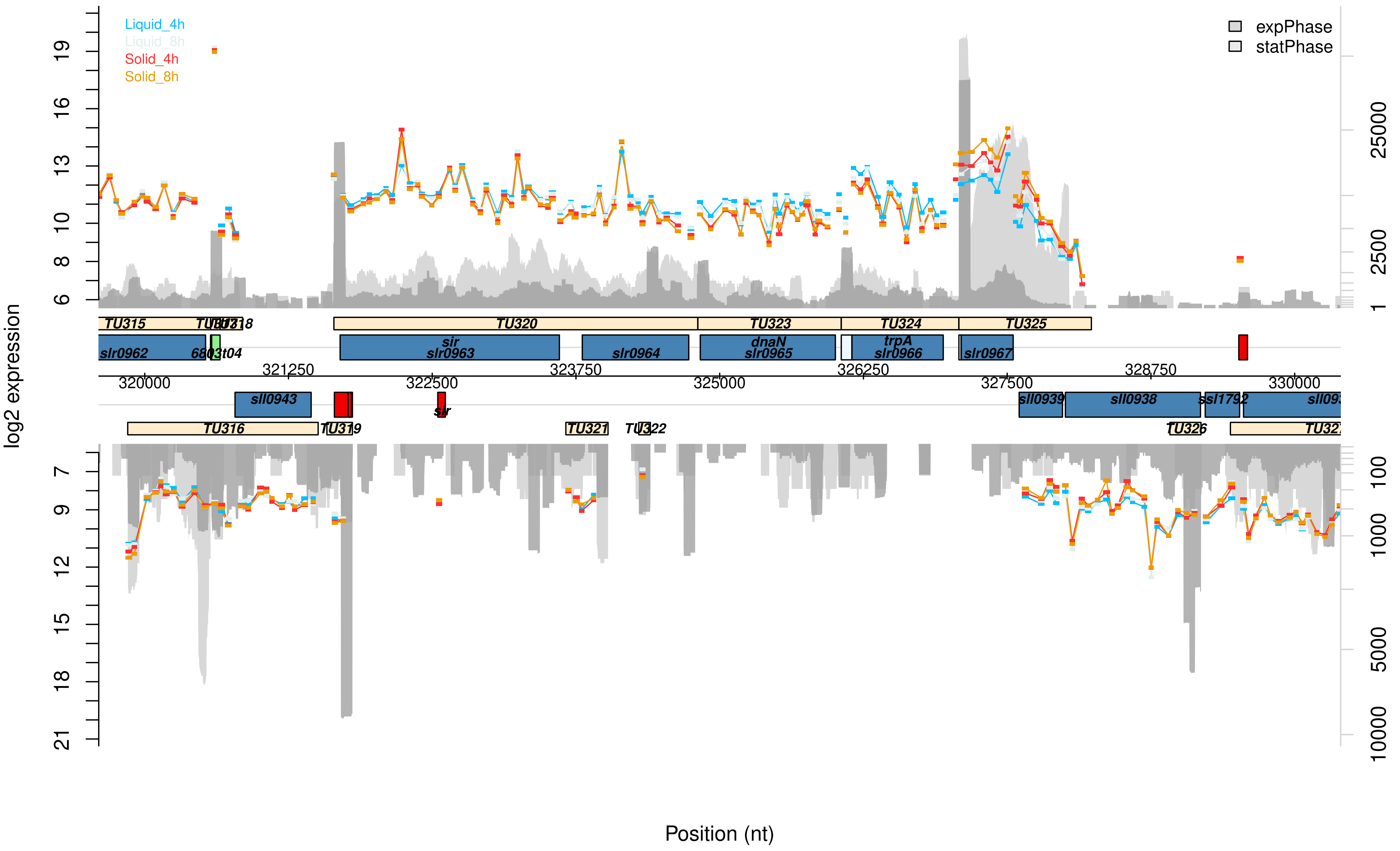

Liquid_4h
Liquid_8h
Solid_4h
Solid_8h
expPhase
statPhase
log2 expression
Position (nt)
TU315
TU320
TU323
TU324
TU325
TU316
TU319
TU321
TU326
slr0962
sir
slr0963
slr0964
dnaN
slr0965
trpA
slr0966
slr0967
sll0943
sll0939
sll0938
ssl1792
320000
321250
322500
323750
325000
326250
327500
328750
330000

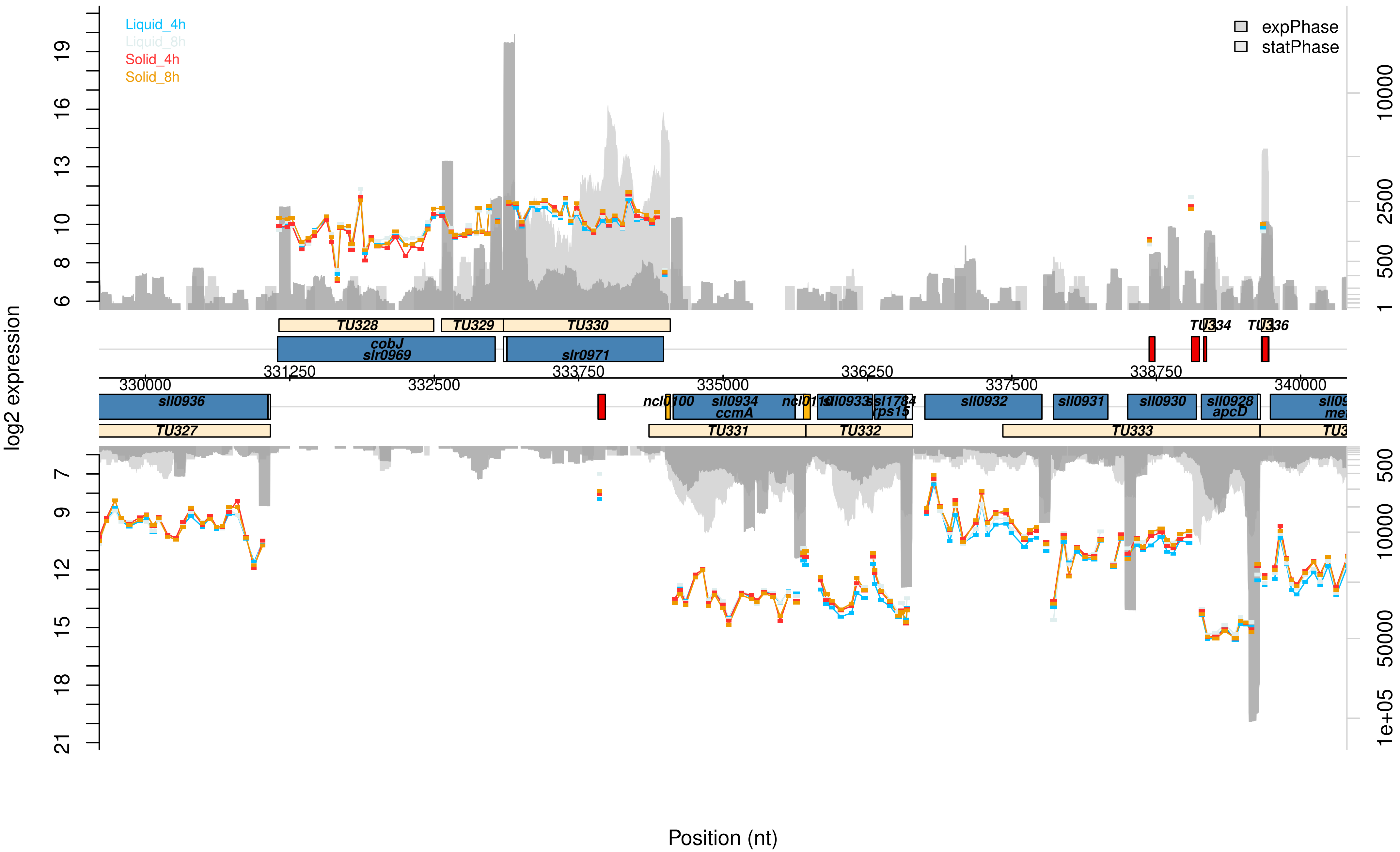

Liquid_4h
Liquid_8h
Solid_4h
Solid_8h
expPhase
statPhase
log2 expression
Position (nt)
TU328
TU329
TU330
TU334
TU336
cobJ
slr0969
slr0971
331250
333750
336250
338750
330000
332500
335000
337500
340000
sll0936
ncl0100
sll0934
ccmA
sll0933
ssl1784
rps15
sll0932
sll0931
sll0930
sll0928
apcD
TU327
TU331
TU332
TU333

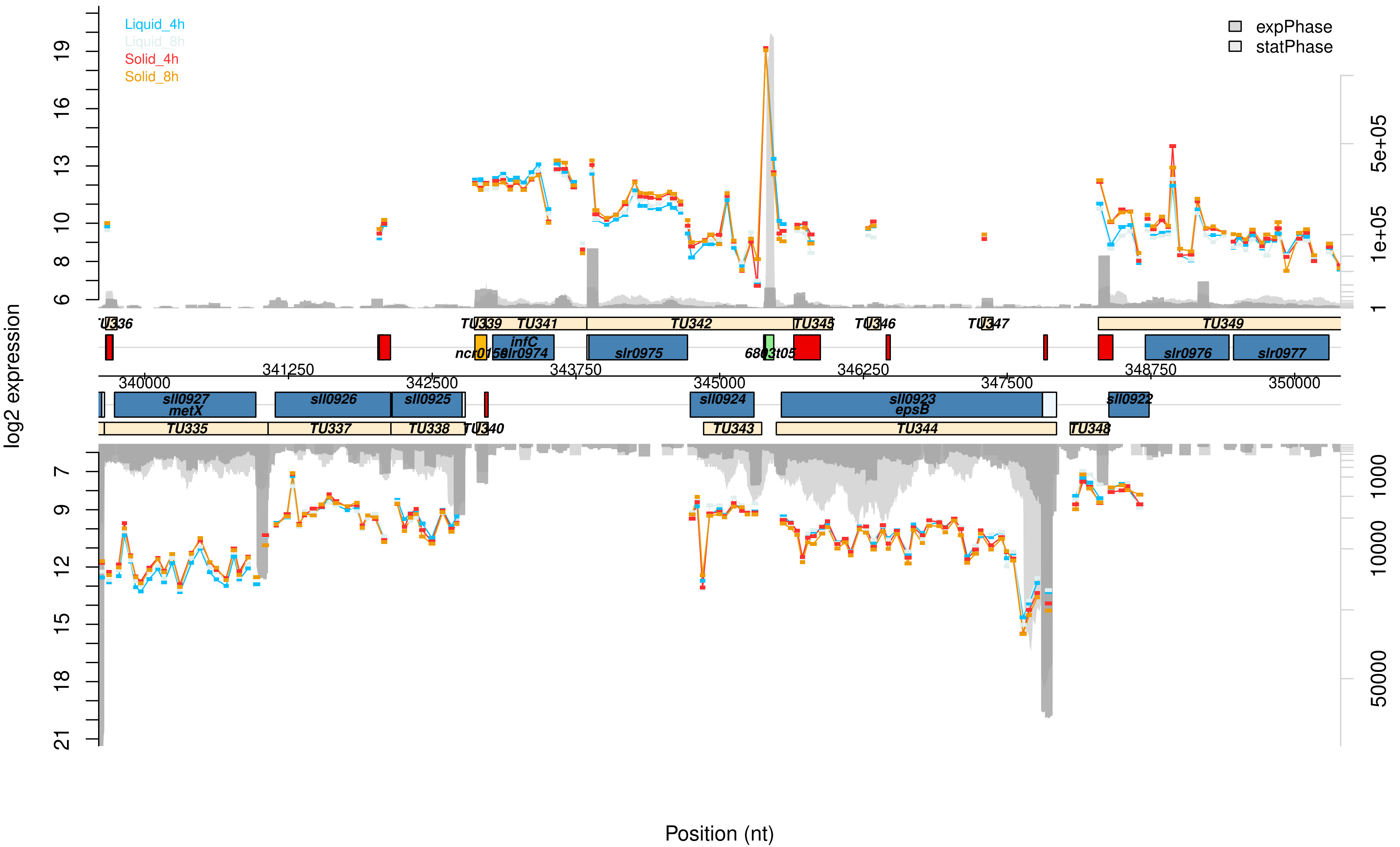

Liquid_4h
Liquid_8h
Solid_4h
Solid_8h
expPhase
statPhase
log2 expression
Position (nt)
TU335
TU336
TU337
TU338
TU339
TU340
TU341
TU342
TU343
TU344
TU345
TU346
TU347
TU348
TU349
sll0927
metX
sll0926
sll0925
sll0924
sll0923
epsB
sll0922
infC
slr0974
slr0975
slr0976
slr0977
340000
341250
342500
343750
345000
346250
347500
348750
350000
5e+05
1e+05
1000
10000
50000

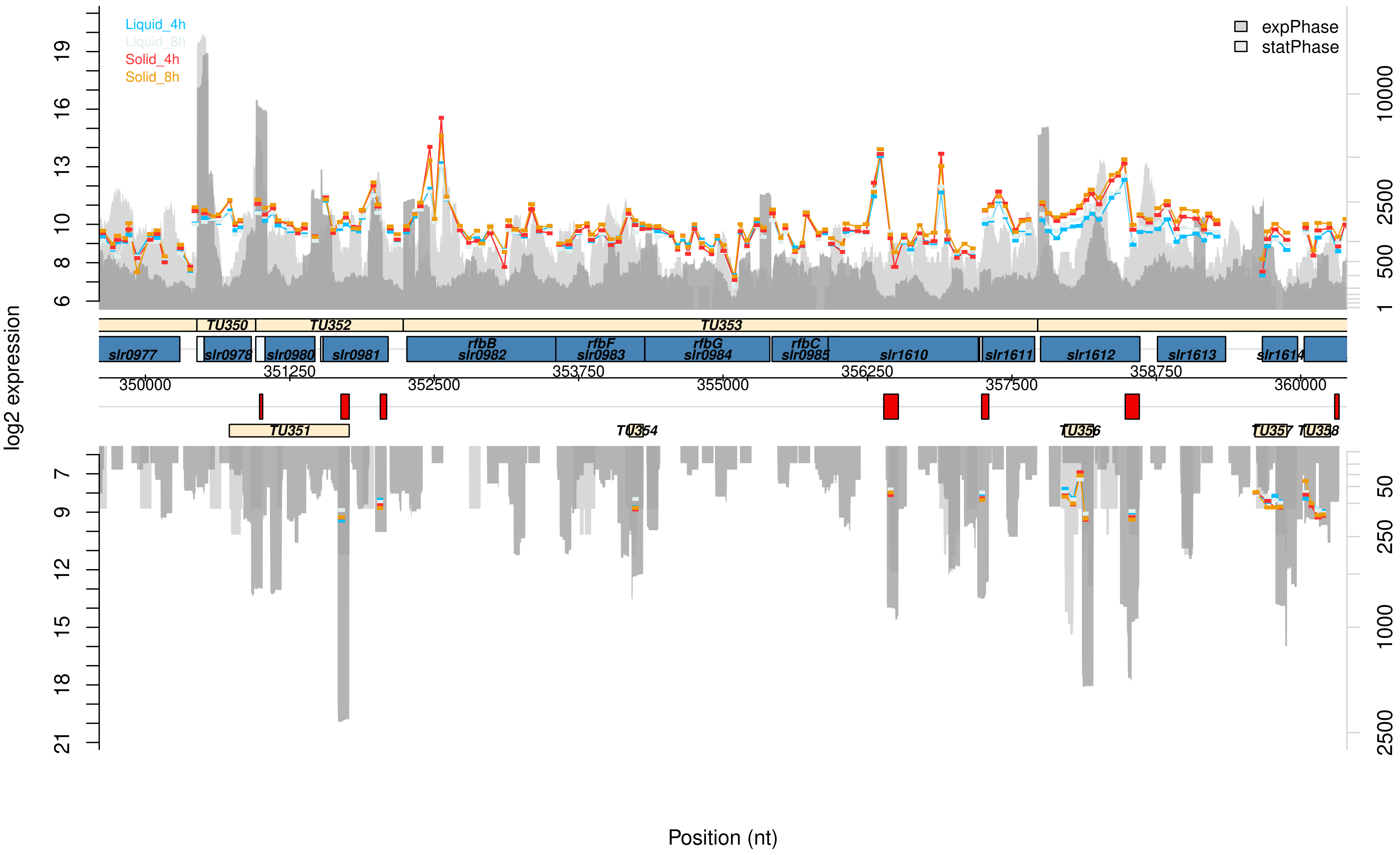

Liquid_4h
Liquid_8h
Solid_4h
Solid_8h
expPhase
statPhase
log2 expression
6
8
10
13
16
19
7
9
12
15
18
21
1
500
2500
10000
50
250
1000
2500
TU350
TU352
TU353
TU351
TU354
TU356
TU357
TU358
slr0977
slr0978
slr0980
slr0981
rfbB
slr0982
rfbF
slr0983
rfbG
slr0984
rfbC
slr0985
slr1610
slr1611
slr1612
slr1613
slr1614
350000
351250
352500
353750
355000
356250
357500
358750
360000
Position (nt)

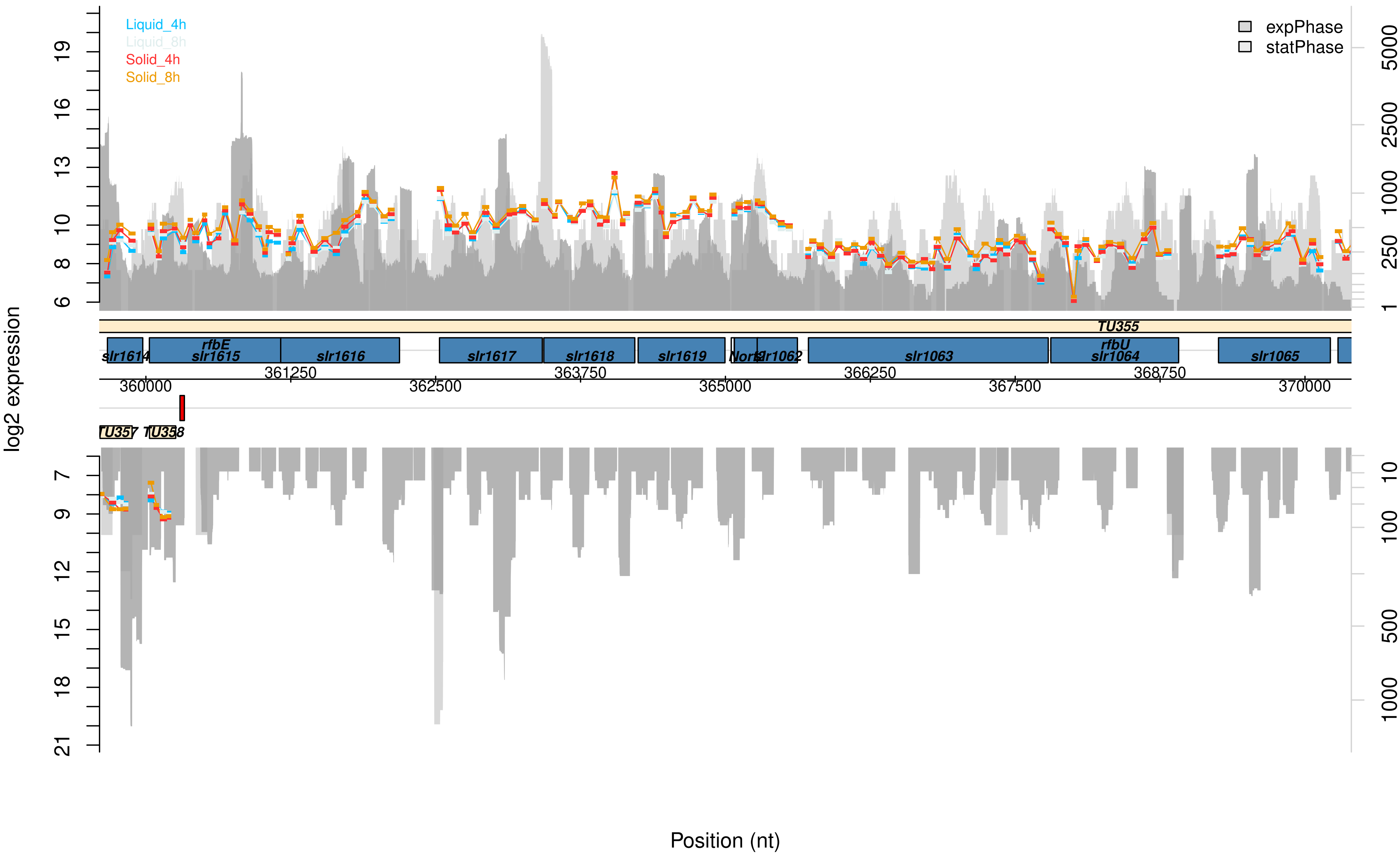

Liquid_4h
Liquid_8h
Solid_4h
Solid_8h
expPhase
statPhase
log2 expression
TU355
TU357
TU358
slr1614
rfbE
slr1615
slr1616
slr1617
slr1618
slr1619
slr1062
slr1063
rfbU
slr1064
slr1065
360000
361250
362500
363750
365000
366250
367500
368750
370000
Position (nt)

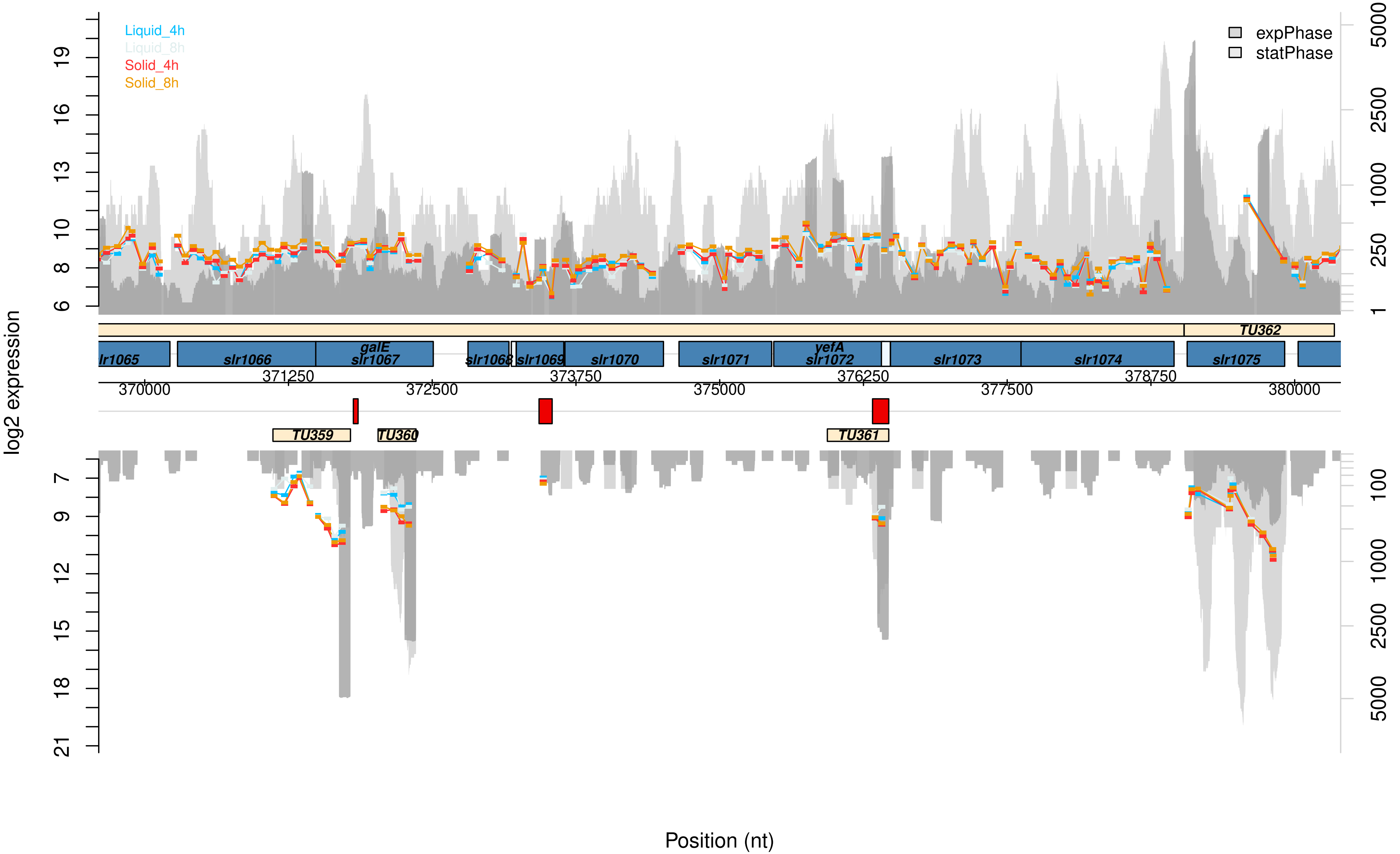

Liquid_4h
Liquid_8h
Solid_4h
Solid_8h
expPhase
statPhase
log2 expression
lr1065
slr1066
galE
slr1067
slr1068
slr1069
slr1070
slr1071
yefA
slr1072
slr1073
slr1074
slr1075
TU359
TU360
TU361
TU362
370000
371250
372500
373750
375000
376250
377500
378750
380000
Position (nt)

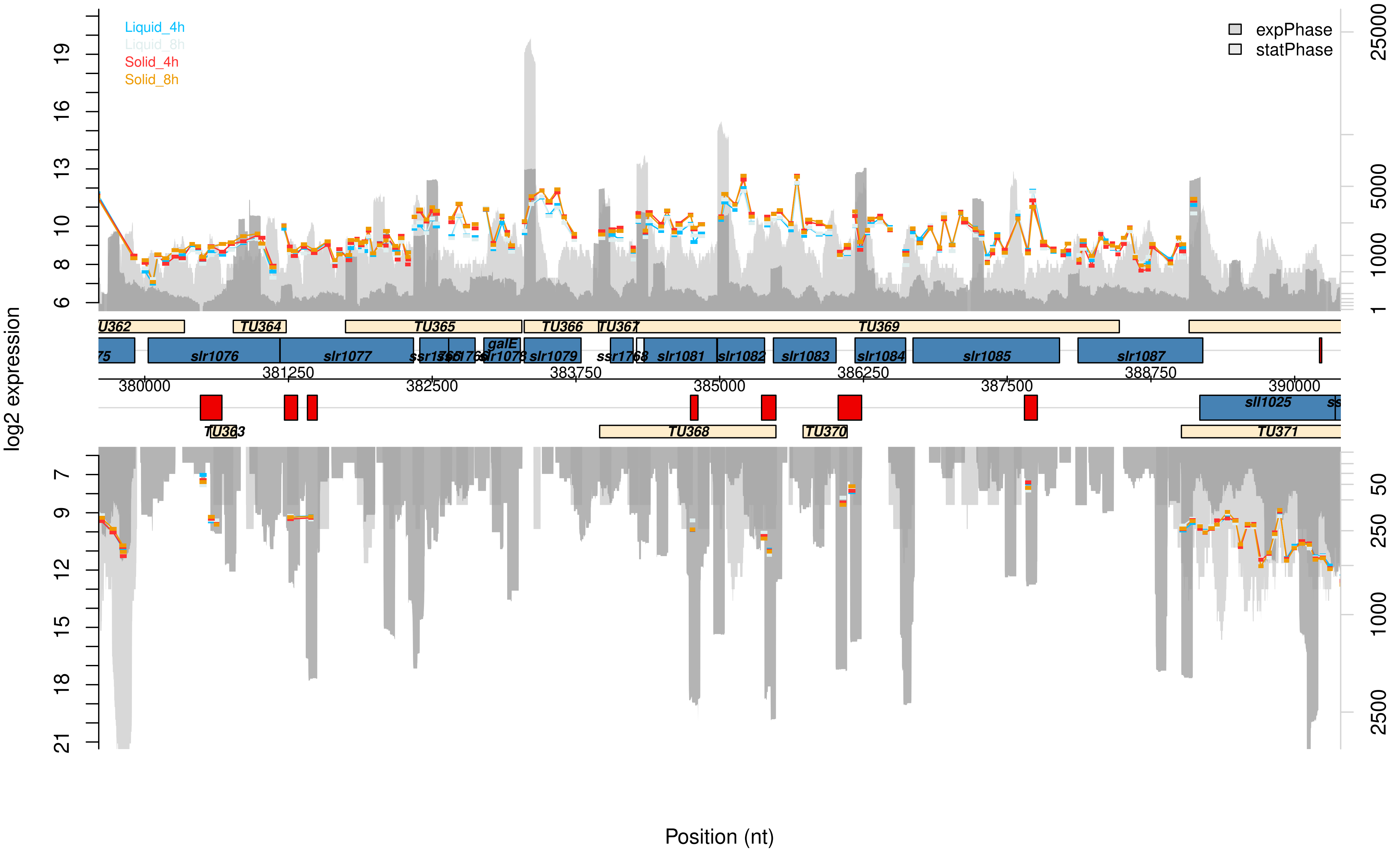

Liquid_4h
Liquid_8h
Solid_4h
Solid_8h
expPhase
statPhase
log2 expression
Position (nt)
TU362
TU363
TU364
TU365
TU366
TU367
TU368
TU369
TU370
TU371
slr1076
slr1077
galE
slr1079
ssr1768
slr1081
slr1082
slr1083
slr1084
slr1085
slr1087
sll1025
380000
381250
382500
383750
385000
386250
387500
388750
390000

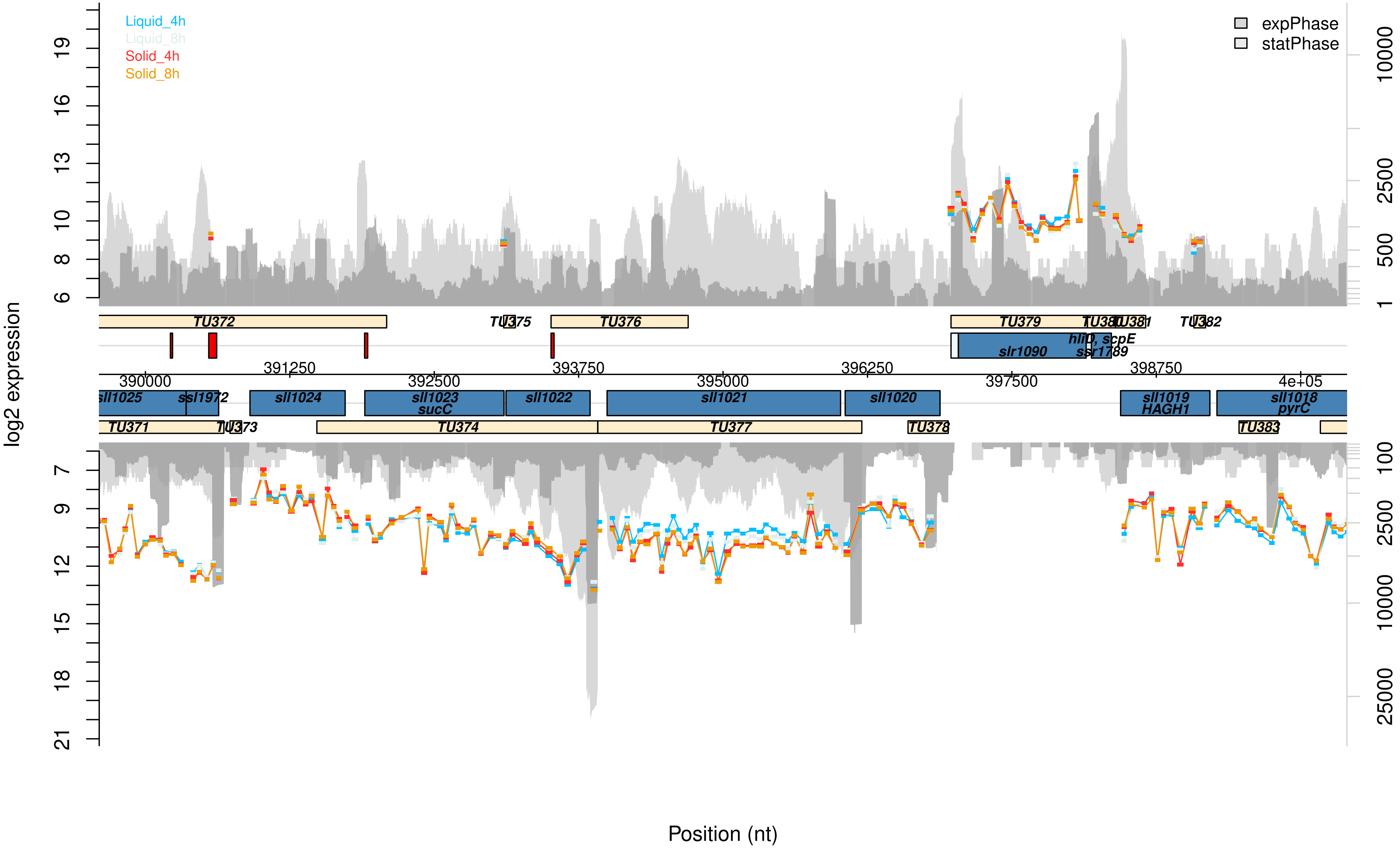

Liquid_4h
Liquid_8h
Solid_4h
Solid_8h
expPhase
statPhase
log2 expression
Position (nt)
TU372
TU375
TU376
TU379
TU380
TU381
TU382
slr1090
hliD, scpE
ssr1789
sll1025
ssl1972
sll1024
sll1023
sucC
sll1022
sll1021
sll1020
sll1019
HAGH1
sll1018
pyrC
TU371
TU373
TU374
TU377
TU378
TU383
390000
391250
392500
393750
395000
396250
397500
398750
4e+05

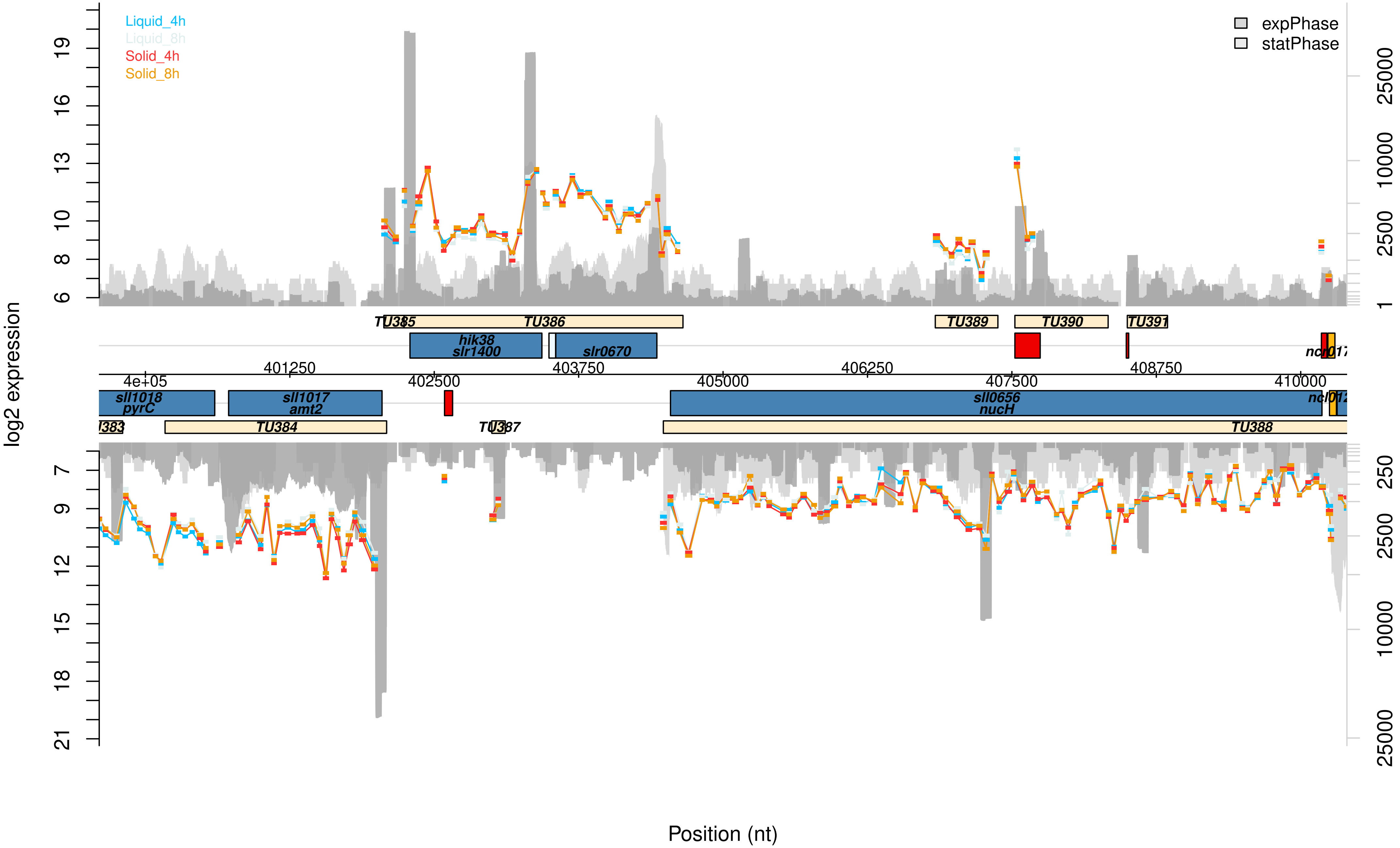

Liquid_4h
Liquid_8h
Solid_4h
Solid_8h
expPhase
statPhase
log2 expression
Position (nt)
TU385
TU386
TU389
TU390
TU391
hik38
slr1400
slr0670
sll1018
pyrC
sll1017
amt2
sll0656
nucH
TU384
TU387
TU388
4e+05
401250
402500
403750
405000
406250
407500
408750
410000

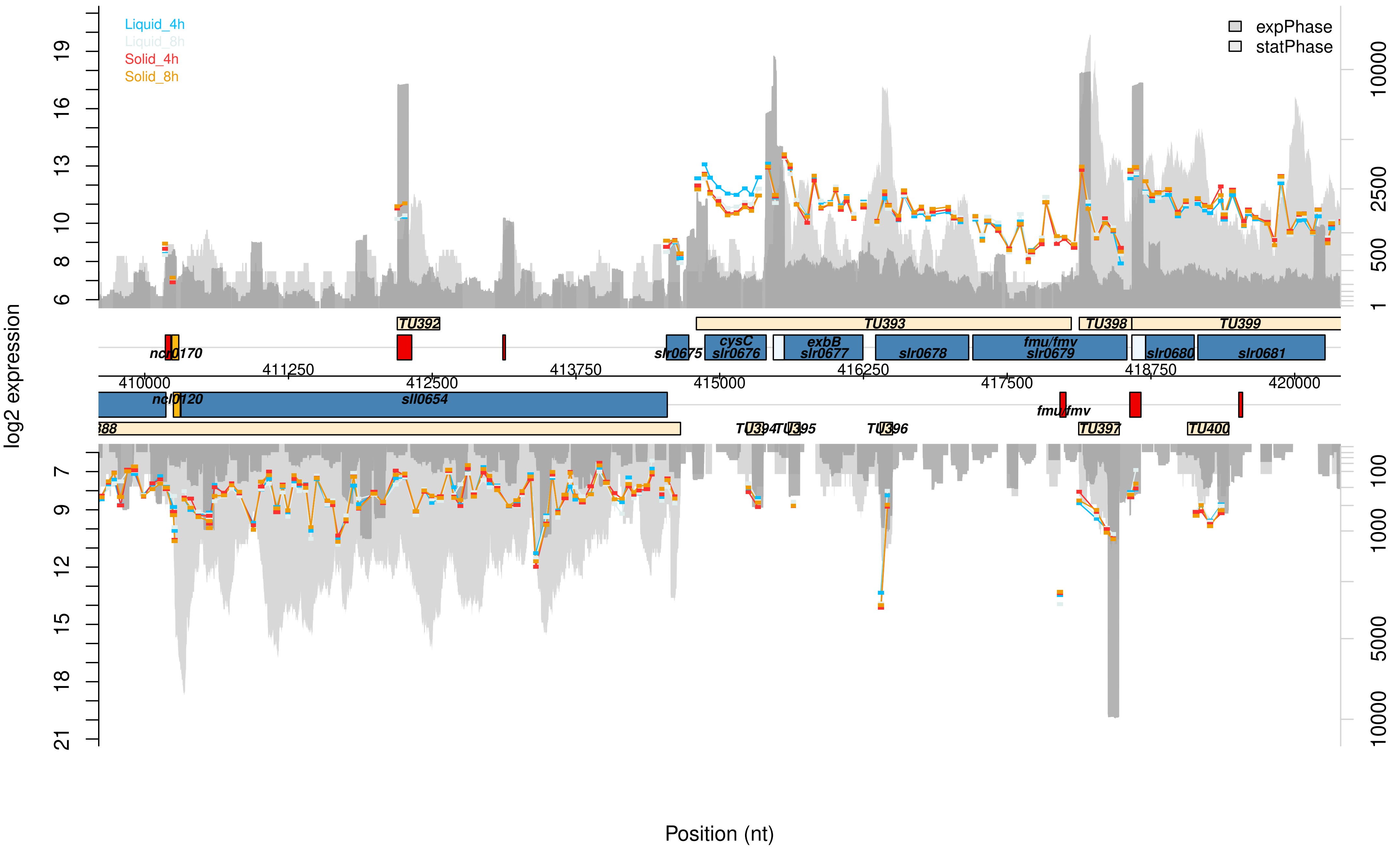

Liquid_4h
Liquid_8h
Solid_4h
Solid_8h
expPhase
statPhase
log2 expression
Position (nt)
TU392
TU393
TU398
TU399
ncl0170
slr0675
cysC slr0676
exbB slr0677
slr0678
fmu/fmv slr0679
slr0680
slr0681
411250
413750
416250
418750
410000
412500
415000
417500
420000
ncl0120
sll0654
fmu/fmv
TU394
TU395
TU396
TU397
TU400

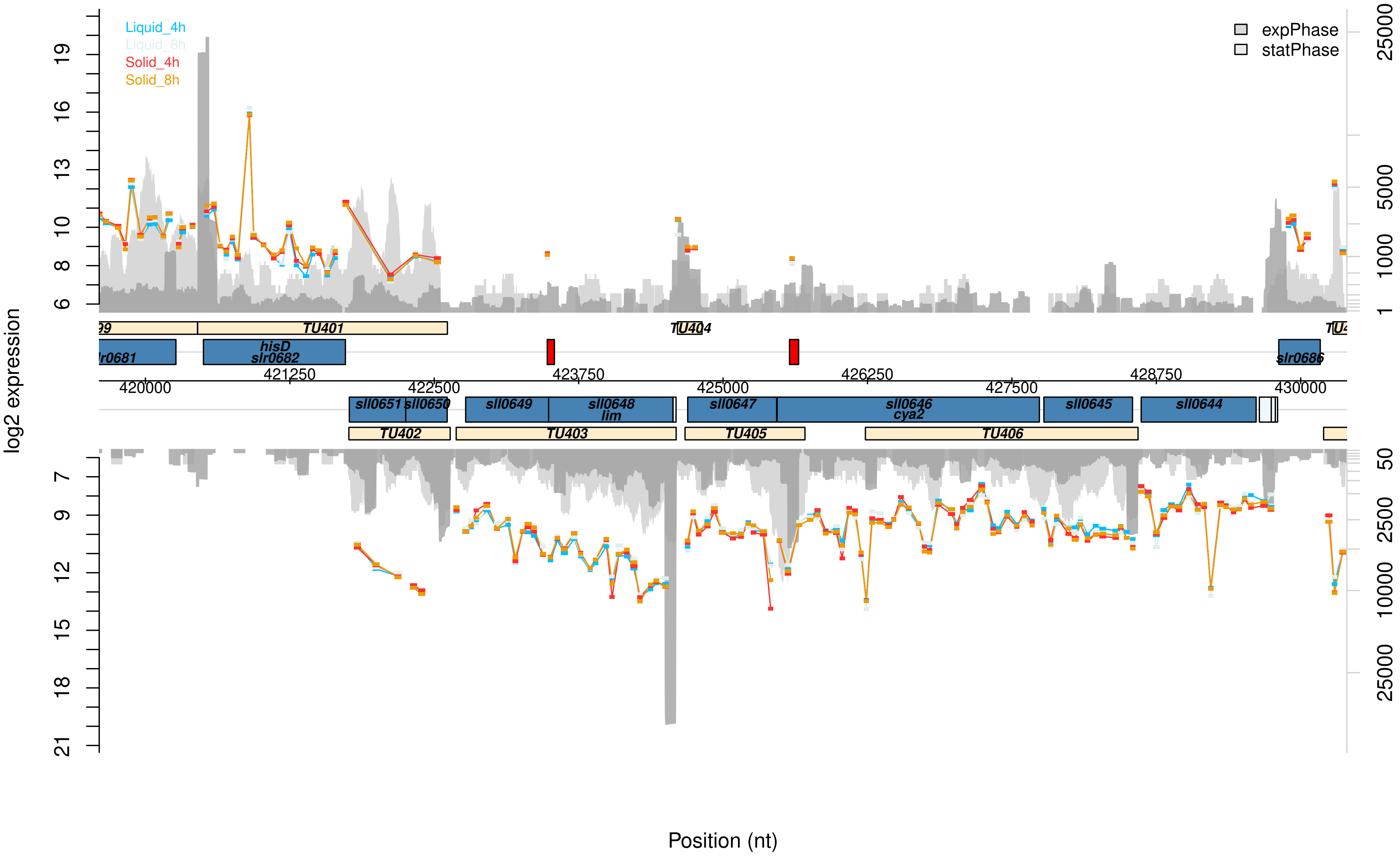

Liquid_4h
Liquid_8h
Solid_4h
Solid_8h
expPhase
statPhase
log2 expression
Position (nt)
TU401
TU402
TU403
TU404
TU405
TU406
hisD
slr0682
slr0686
sll0651
sll0650
sll0649
sll0648
lim
sll0647
sll0646
cya2
sll0645
sll0644
420000
421250
422500
423750
425000
426250
427500
428750
430000

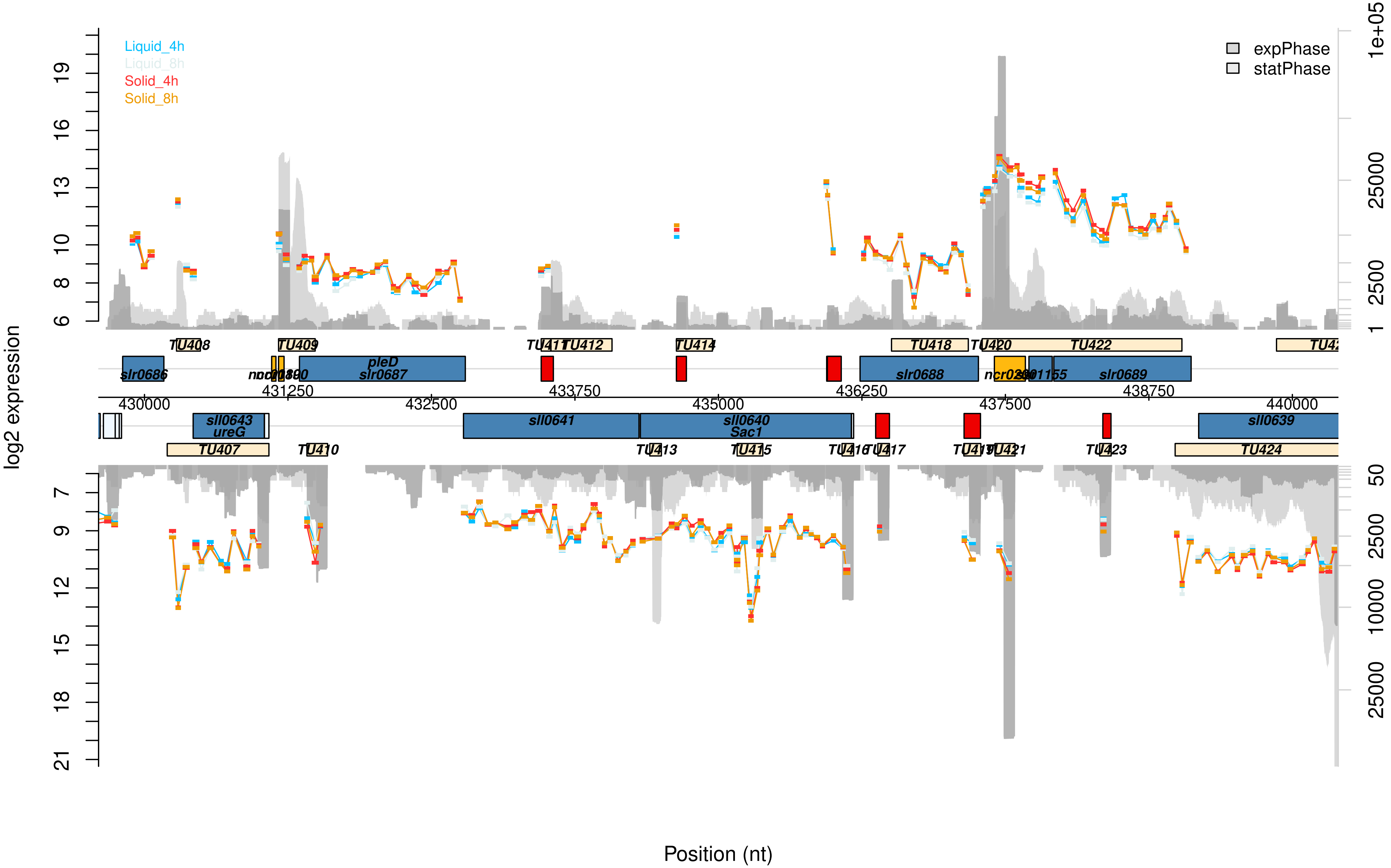

Liquid_4h
Liquid_8h
Solid_4h
Solid_8h
expPhase
statPhase
log2 expression
Position (nt)
TU407
TU408
TU409
TU410
TU412
TU413
TU414
TU415
TU417
TU418
TU420
TU422
TU423
TU424
slr0686
pleD
slr0687
slr0688
slr0689
sll0643
ureG
sll0641
sll0640
Sac1
sll0639
430000
431250
432500
433750
435000
436250
437500
438750
440000

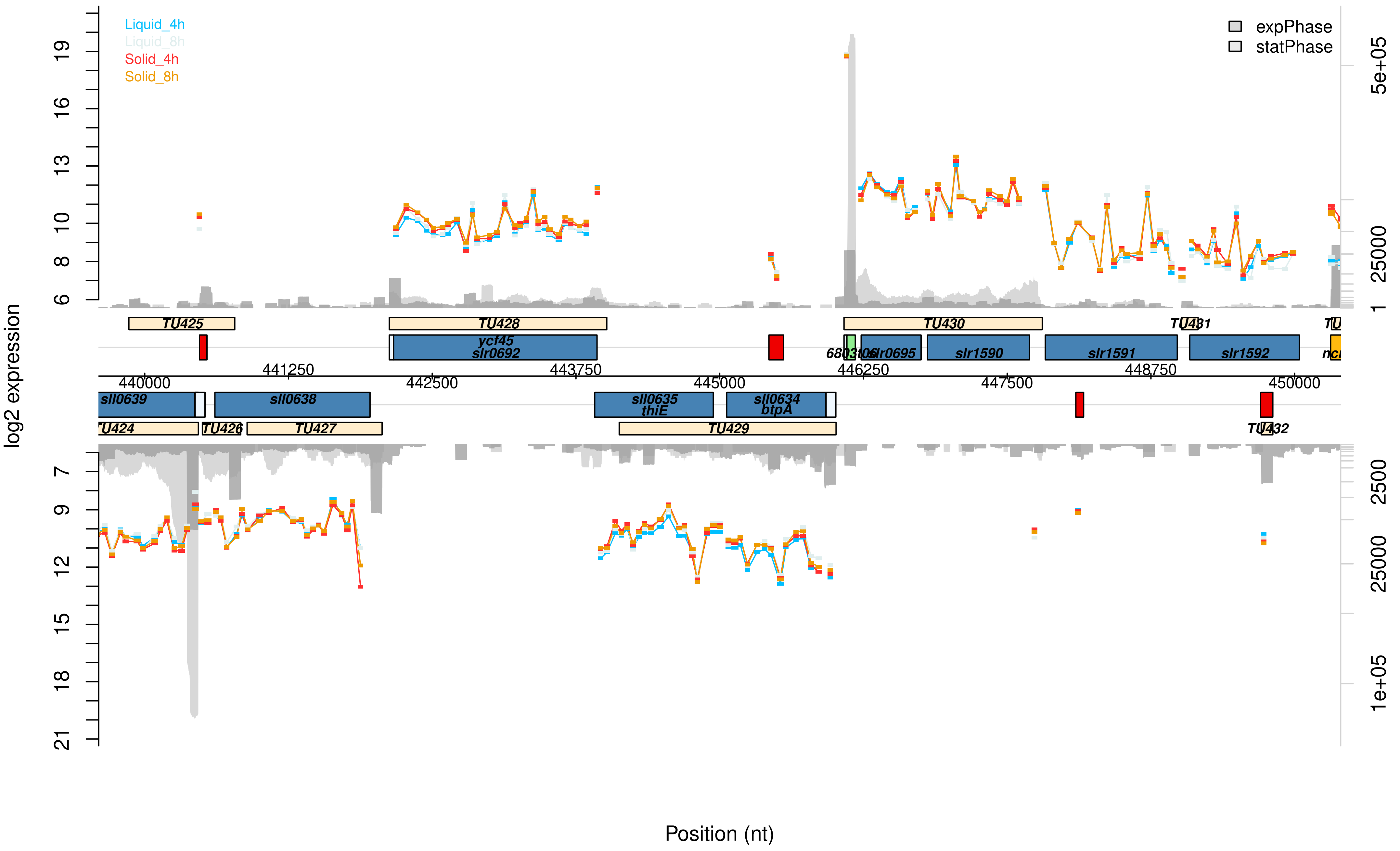

Liquid_4h
Liquid_8h
Solid_4h
Solid_8h
expPhase
statPhase
log2 expression
Position (nt)
TU424
TU425
TU426
TU427
TU428
TU429
TU430
TU431
TU432
sll0639
sll0638
ycf45
slr0692
sll0635
thiE
sll0634
btpA
slr0695
slr1590
slr1591
slr1592
440000
441250
442500
443750
445000
446250
447500
448750
450000
5e+05
25000
2500
1e+05

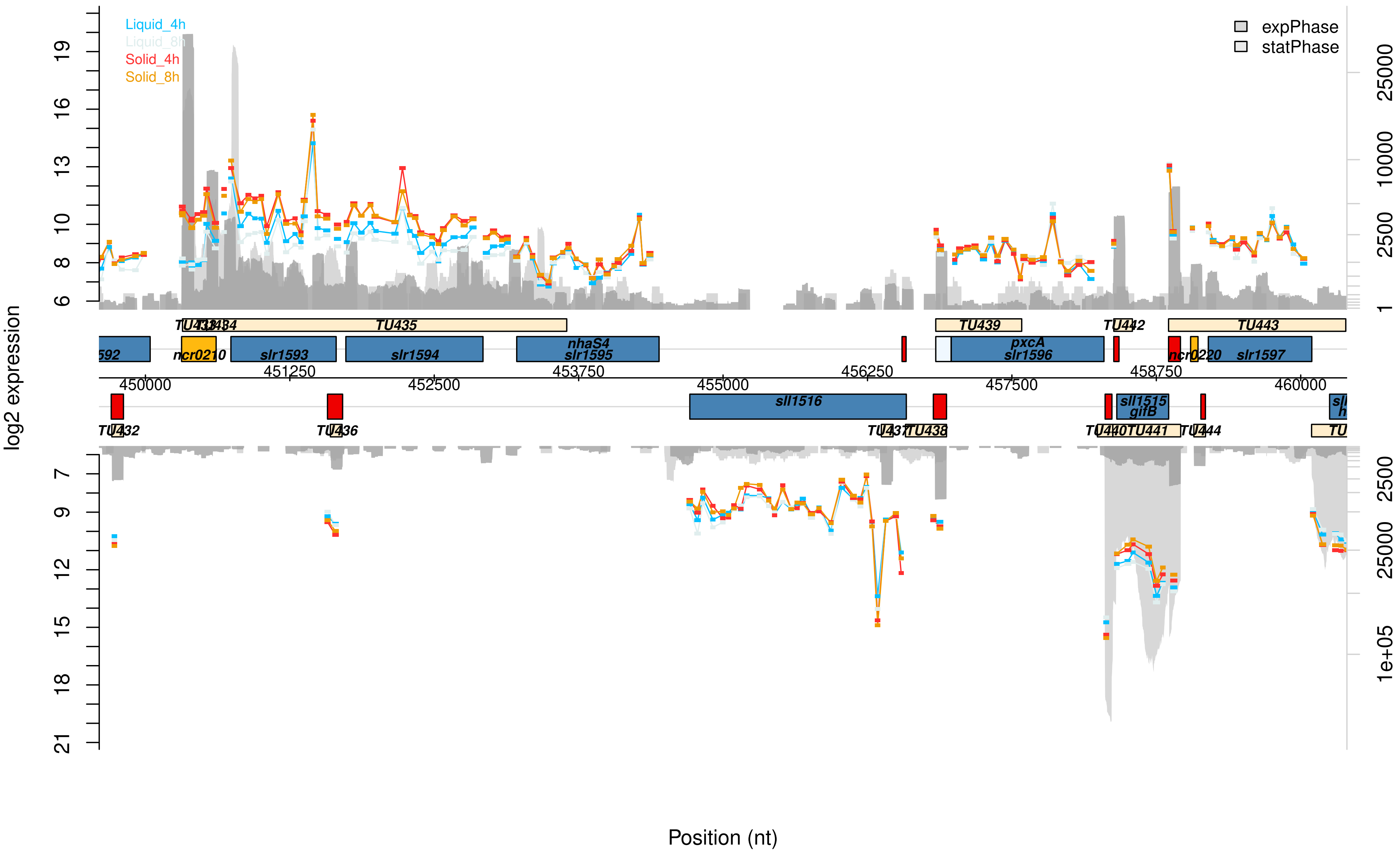
Liquid_4h
Liquid_8h
Solid_4h
Solid_8h
expPhase
statPhase
log2 expression
Position (nt)
TU433
TU434
TU435
TU439
TU442
TU443
ncr0210
slr1593
slr1594
nhaS4
slr1595
pxcA
slr1596
ncr0220
slr1597
sll1516
sll1515
gifB
TU432
TU436
TU437
TU438
TU440
TU441
TU444
450000
451250
452500
453750
455000
456250
457500
458750
460000

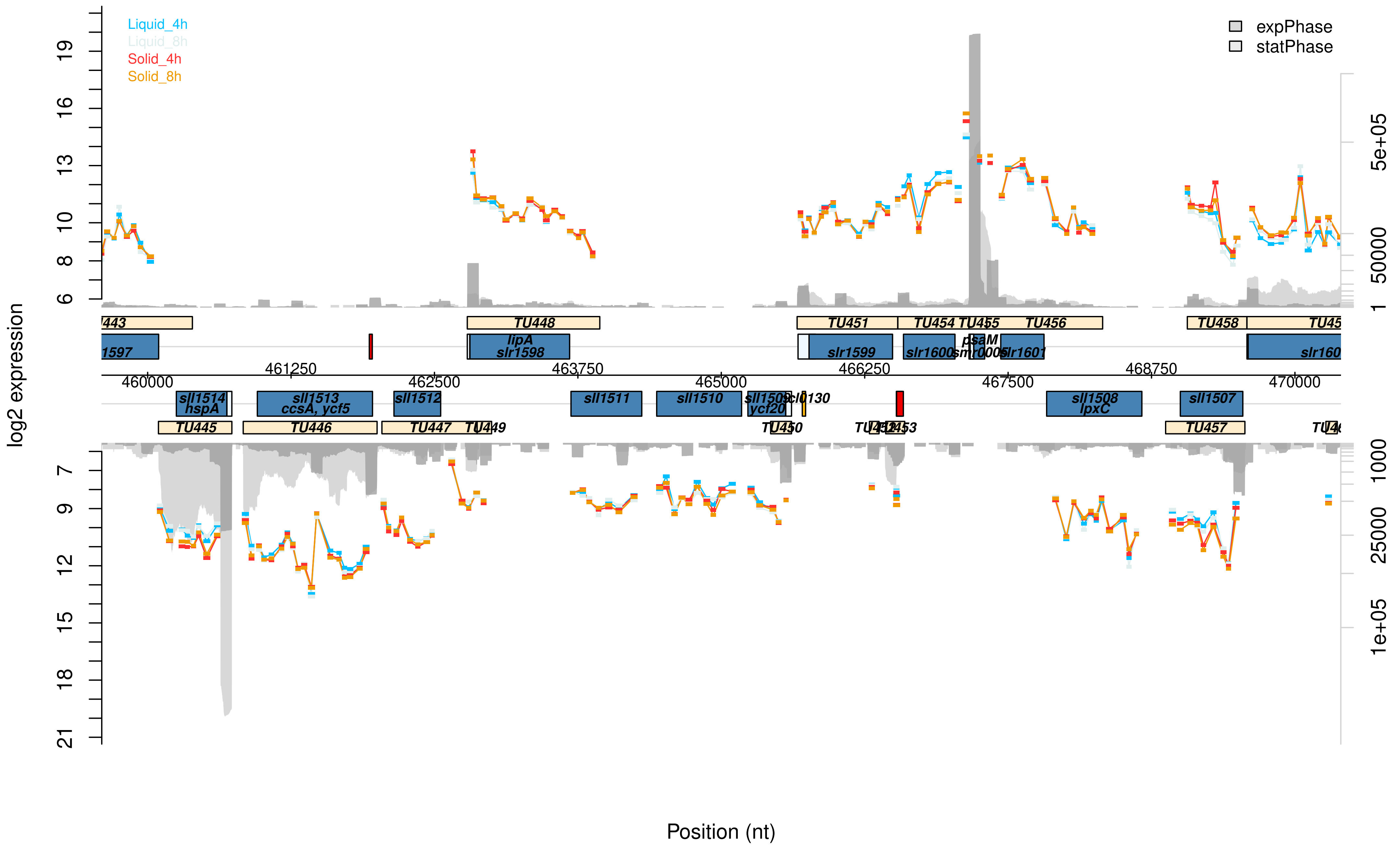

Liquid_4h
Liquid_8h
Solid_4h
Solid_8h
expPhase
statPhase
log2 expression
Position (nt)
TU448
TU451
TU454
TU455
TU456
TU458
lipA
slr1598
slr1599
slr1600
psaM
smr0005
slr1601
sll1514
hspA
sll1513
ccsA, ycf5
sll1512
sll1511
sll1510
sll1508
lpxC
sll1507
ycf20
TU445
TU446
TU447
TU449
TU450
TU457
460000
461250
462500
463750
465000
466250
467500
468750
470000

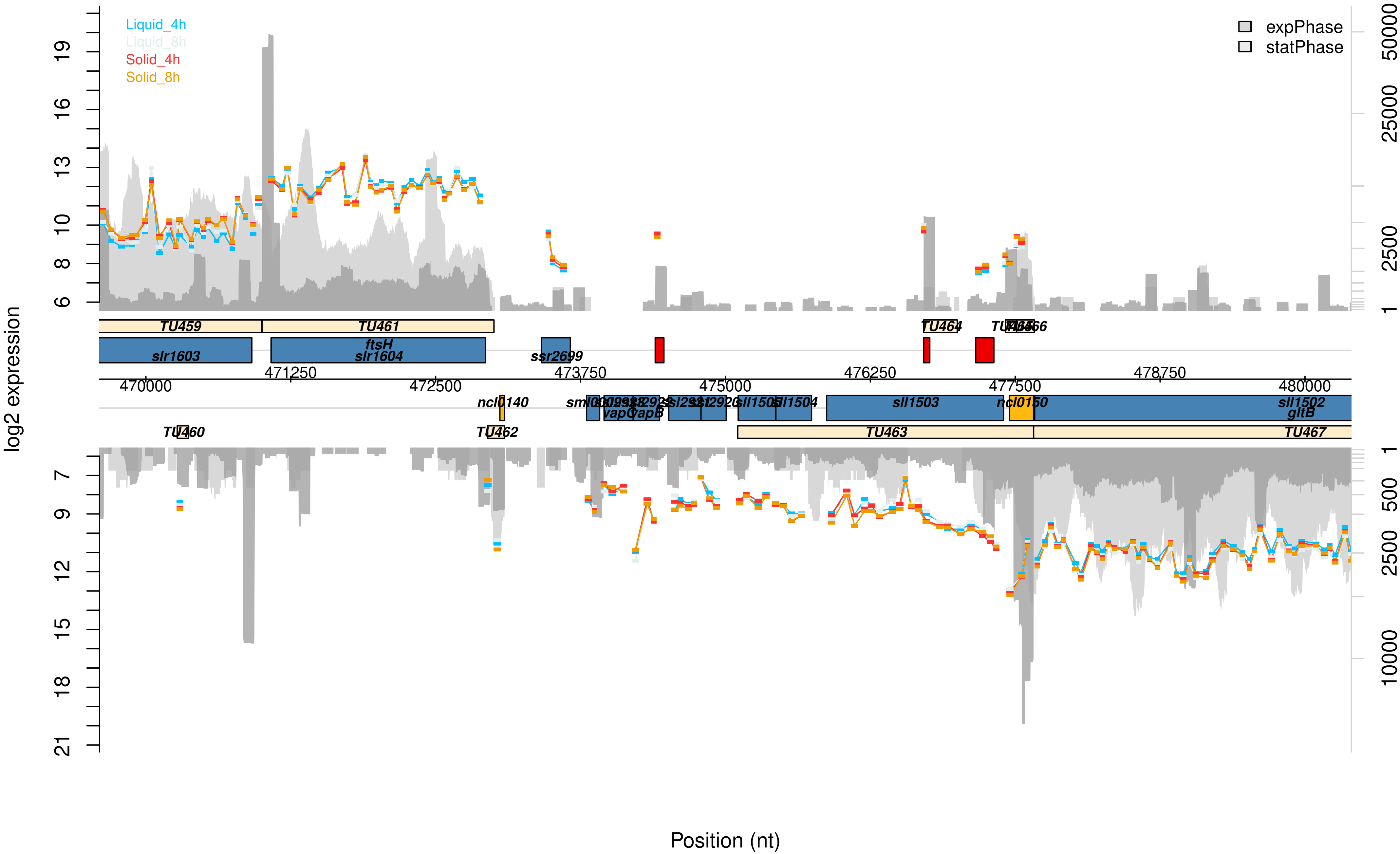

Liquid_4h
Liquid_8h
Solid_4h
Solid_8h
expPhase
statPhase
log2 expression
Position (nt)
TU459
TU461
TU464
TU460
TU462
TU463
TU467
slr1603
ftsH
slr1604
ssr2699
ncl0140
ncl0150
sll1503
sll1502
gltB
vapC
sll1504
470000
471250
472500
473750
475000
476250
477500
478750
480000

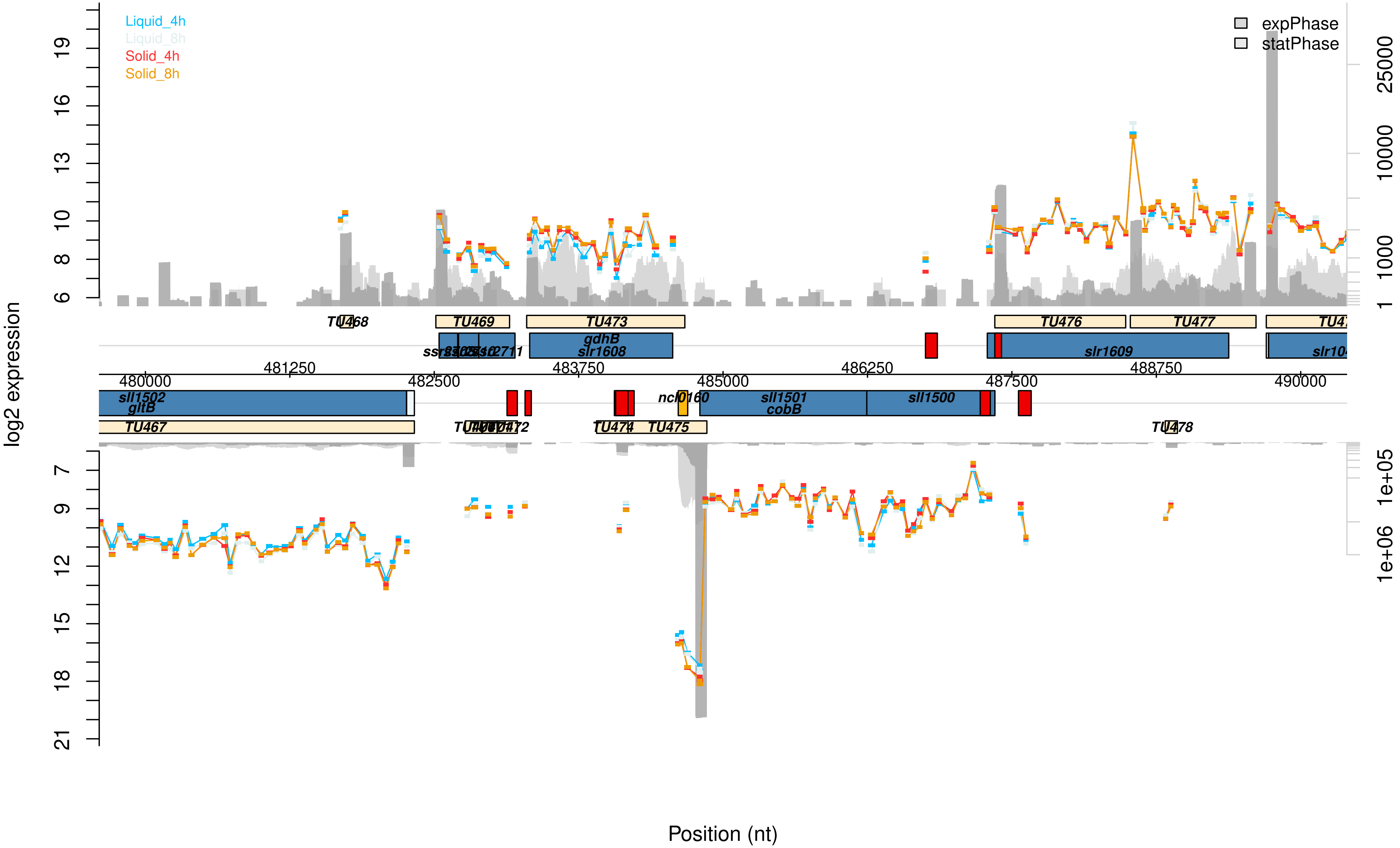

Liquid_4h
Liquid_8h
Solid_4h
Solid_8h
expPhase
statPhase
log2 expression
Position (nt)
TU468
TU469
TU473
TU476
TU477
gdhB
slr1608
slr1609
481250
483750
486250
488750
480000
482500
485000
487500
490000
sll1502
gltB
ncl0160
sll1501
cobB
sll1500
TU467
TU474
TU475
TU478

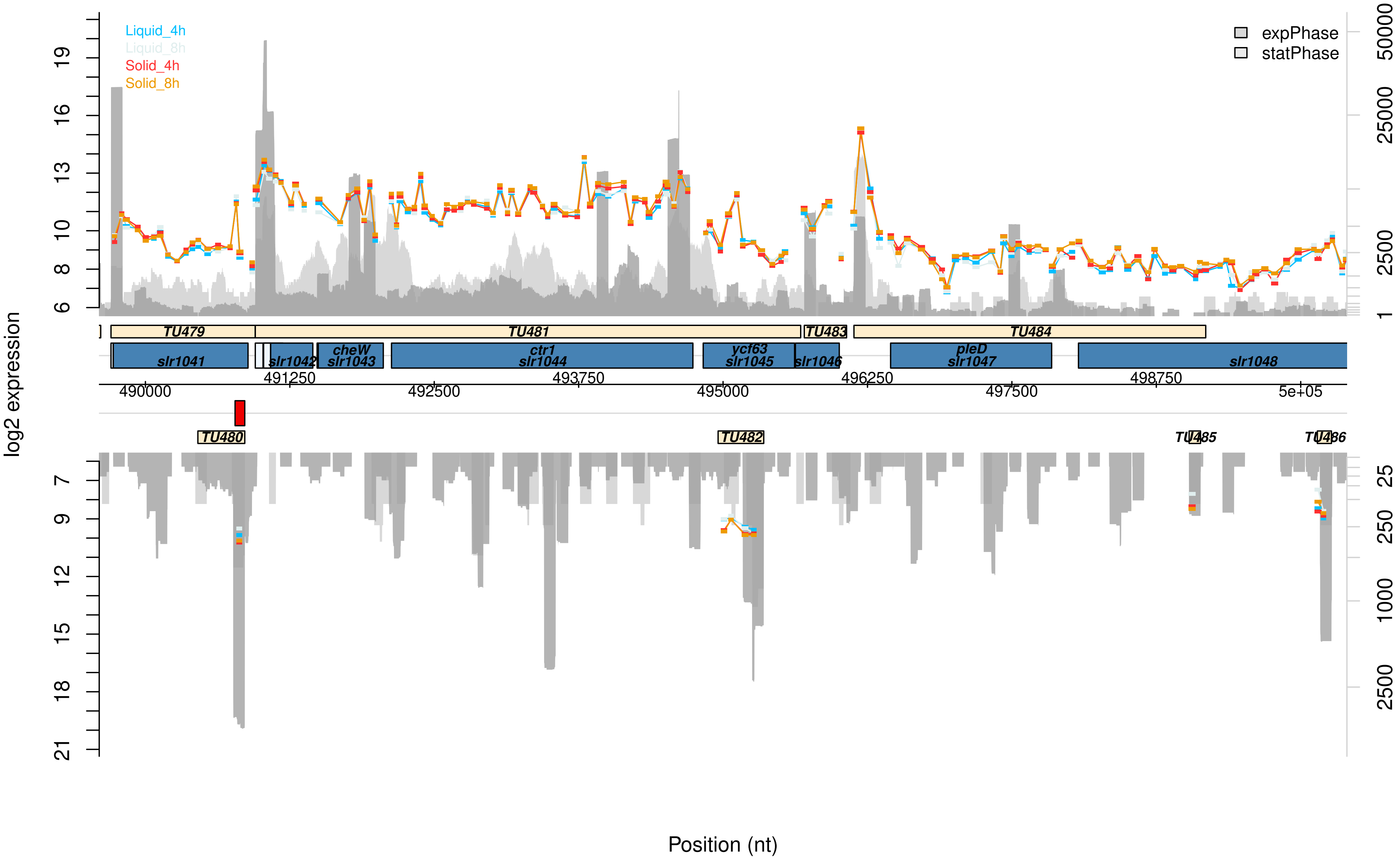

Liquid_4h
Liquid_8h
Solid_4h
Solid_8h
expPhase
statPhase
log2 expression
TU479
TU481
TU483
TU484
slr1041
slr1042
cheW
slr1043
ctr1
slr1044
ycf63
slr1045
slr1046
pleD
slr1047
slr1048
490000
491250
492500
493750
495000
496250
497500
498750
5e+05
TU480
TU482
TU485
TU486
Position (nt)

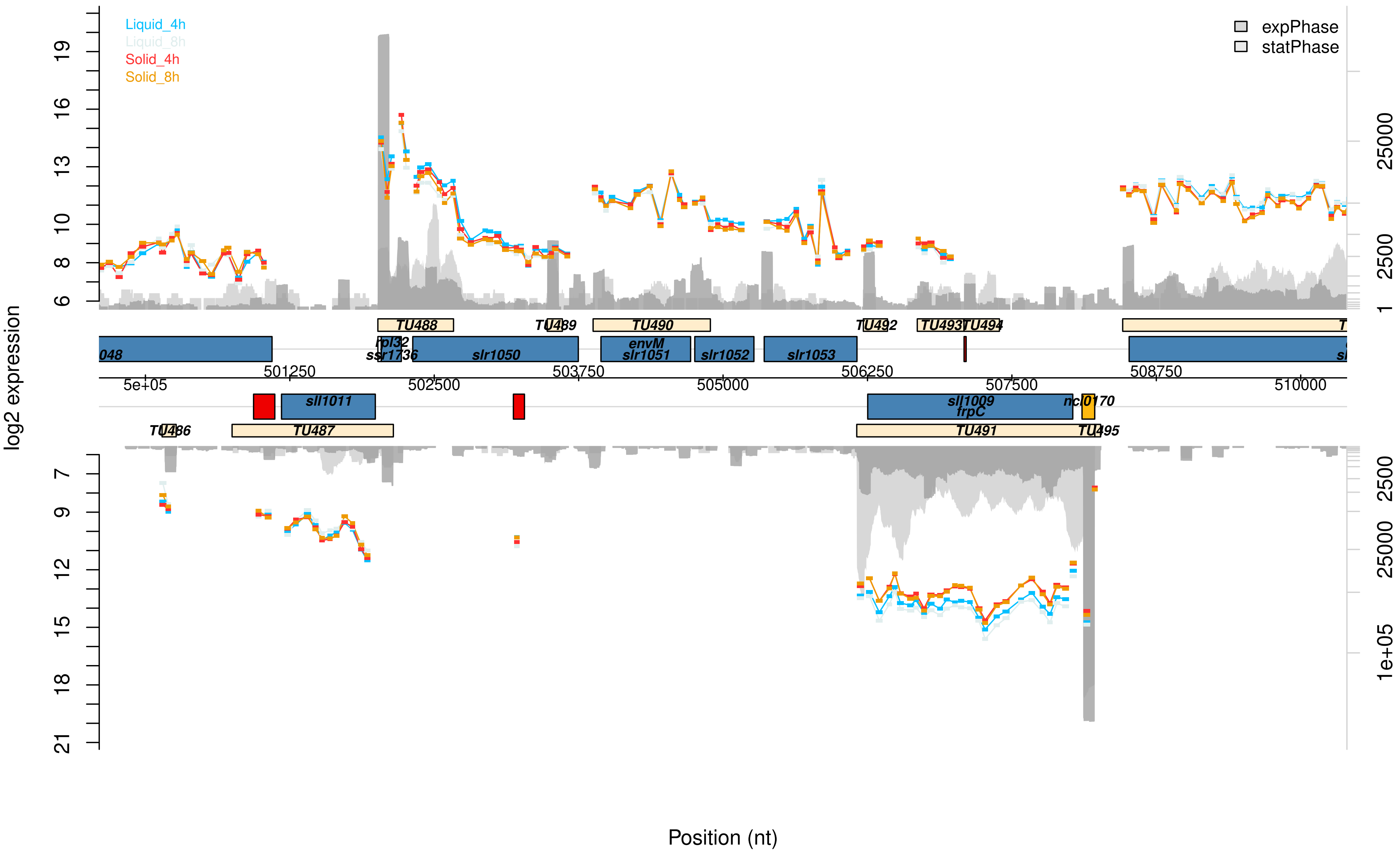

Liquid_4h
Liquid_8h
Solid_4h
Solid_8h
expPhase
statPhase
log2 expression
TU486
TU487
TU488
TU489
TU490
TU491
TU492
TU493
TU494
TU495
rpl32
ssr1736
slr1050
envM
slr1051
slr1052
slr1053
sll1011
sll1009
frpC
ncl0170
Position (nt)

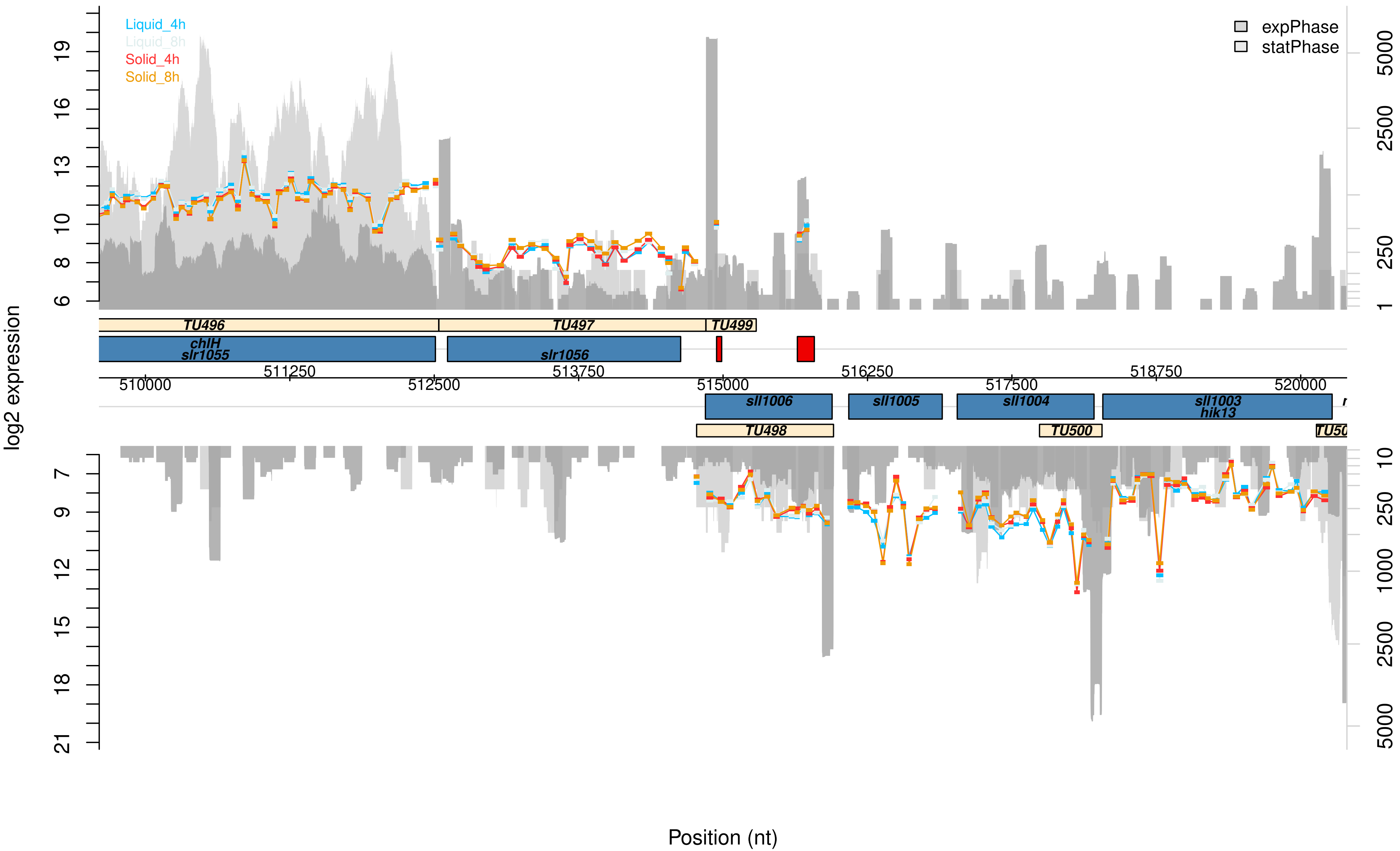
Liquid_4h
Liquid_8h
Solid_4h
Solid_8h
expPhase
statPhase
log2 expression
TU496
TU497
TU499
chlH
slr1055
slr1056
511250
513750
516250
518750
510000
512500
515000
517500
520000
sll1006
sll1005
sll1004
sll1003
hik13
TU498
TU500
Position (nt)

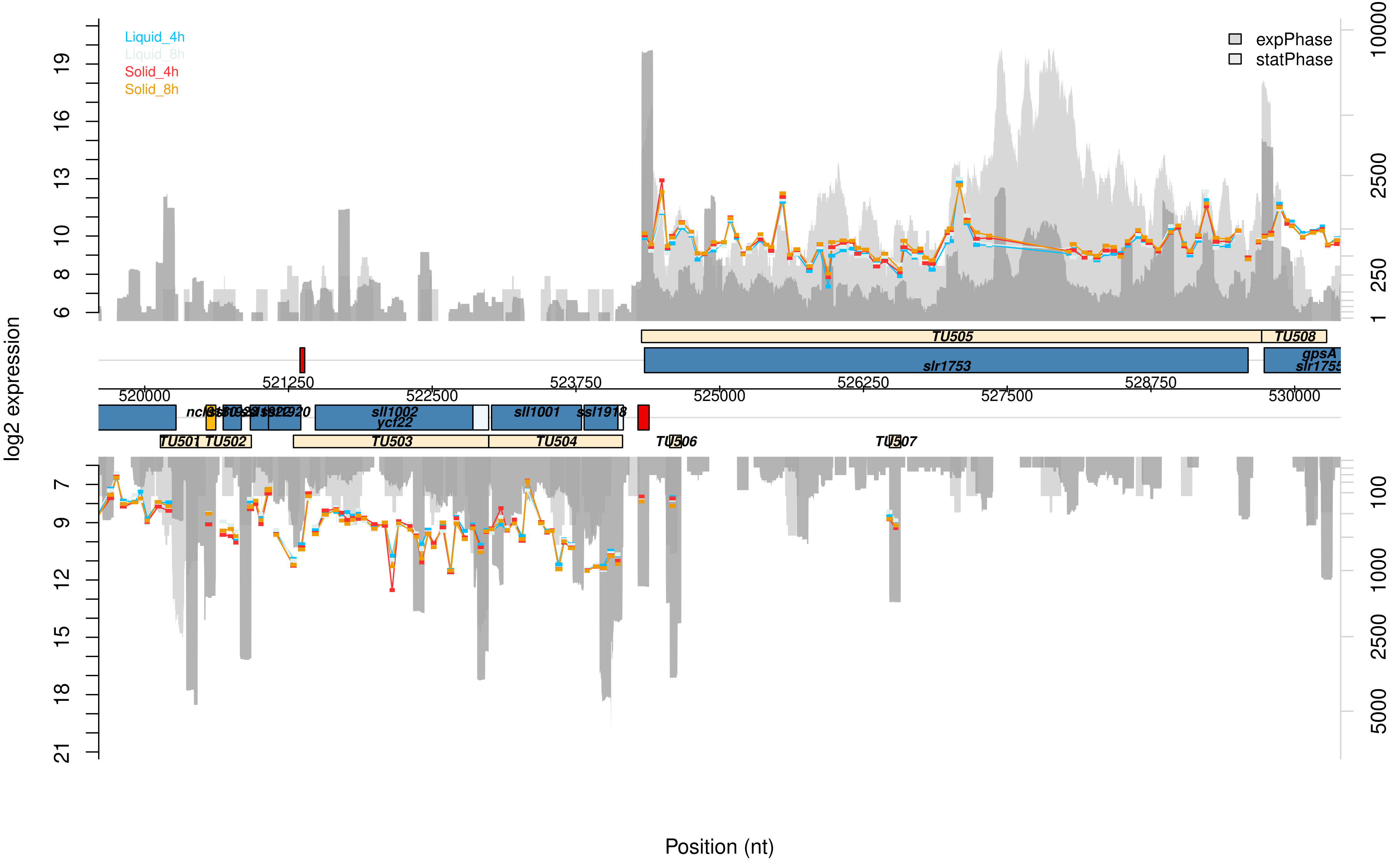

Liquid_4h
Liquid_8h
Solid_4h
Solid_8h
expPhase
statPhase
log2 expression
TU505
TU508
slr1753
gpsA
slr1755
ncl
sll1002
ycf22
sll1001
ssl1918
TU501
TU502
TU503
TU504
TU506
TU507
Position (nt)

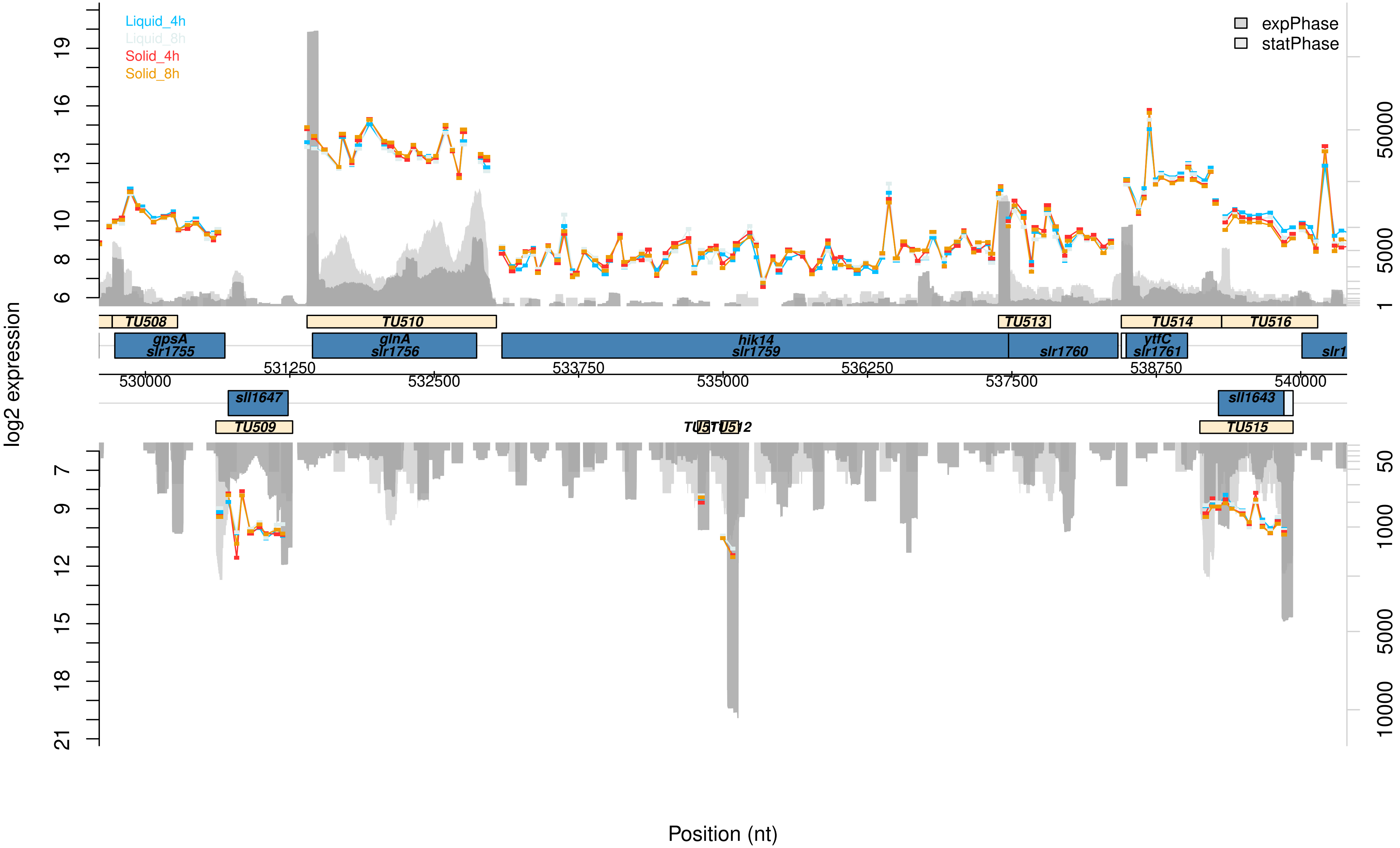

Liquid_4h
Liquid_8h
Solid_4h
Solid_8h
expPhase
statPhase
log2 expression
19
16
13
10
8
6
7
9
12
15
18
21
50000
5000
1
50
1000
5000
10000
TU508
TU510
TU513
TU514
TU516
gpsA
slr1755
glnA
slr1756
hik14
slr1759
slr1760
ytfC
slr1761
slr1
531250
533750
536250
538750
530000
532500
535000
537500
540000
sll1647
sll1643
TU509
TU515
Position (nt)

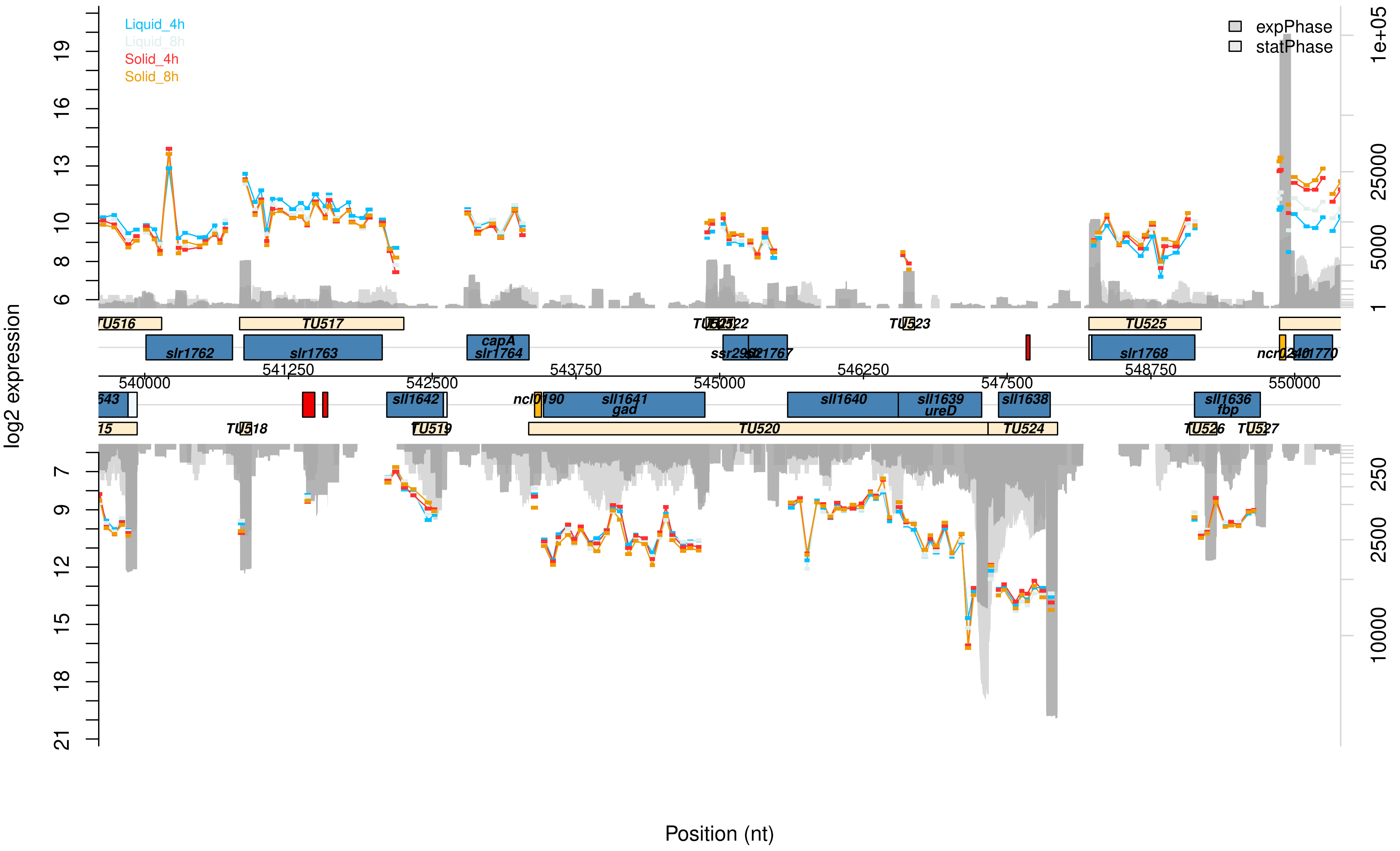

Liquid_4h
Liquid_8h
Solid_4h
Solid_8h
expPhase
statPhase
log2 expression
Position (nt)
TU516
TU517
TU525
TU523
slr1762
slr1763
capA
slr1764
ssr2962
slr1767
slr1768
ncr0240
slr1770
541250
543750
546250
548750
540000
542500
545000
547500
550000
sll1642
ncl0190
sll1641
gad
sll1640
sll1639
ureD
sll1638
sll1636
fbp
TU518
TU519
TU520
TU524
TU526
TU527

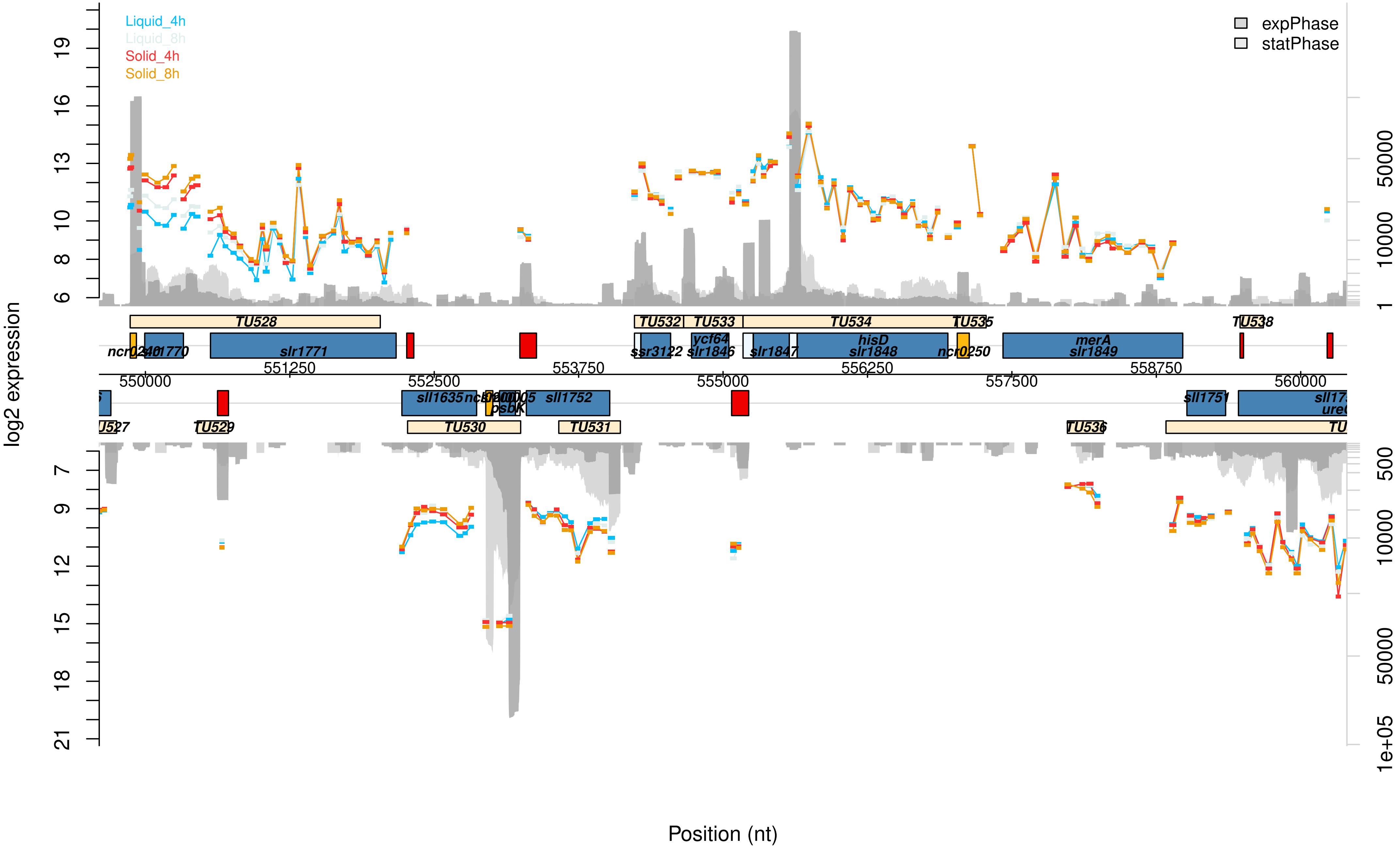

Liquid_4h
Liquid_8h
Solid_4h
Solid_8h
expPhase
statPhase
log2 expression
Position (nt)
TU528
TU532
TU533
TU534
TU535
TU538
ncr0240
slr1770
slr1771
ssr3122
ycf64
slr1846
slr1847
hisD
slr1848
ncr0250
merA
slr1849
550000
551250
552500
553750
555000
556250
557500
558750
560000
sll1635
psbK
sll1752
sll1751
TU529
TU530
TU531
TU536

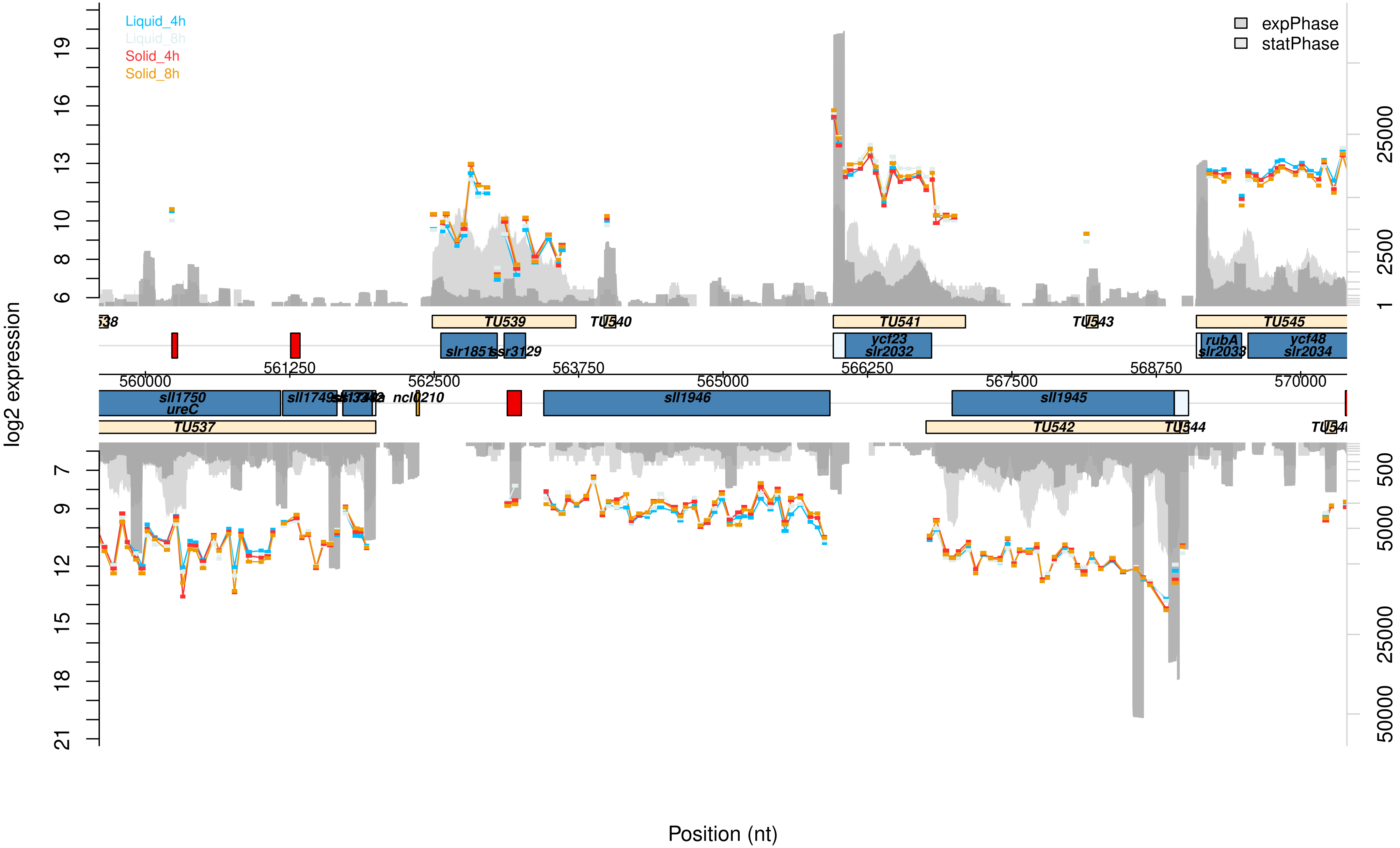

Liquid_4h
Liquid_8h
Solid_4h
Solid_8h
expPhase
statPhase
log2 expression
Position (nt)
TU537
TU539
TU540
TU541
TU542
TU543
TU544
TU545
sll1750
ureC
sll1749
ncl0210
sll1946
sll1945
slr1851
ssr3129
ycf23
slr2032
rubA
slr2033
ycf48
slr2034
560000
561250
562500
563750
565000
566250
567500
568750
570000

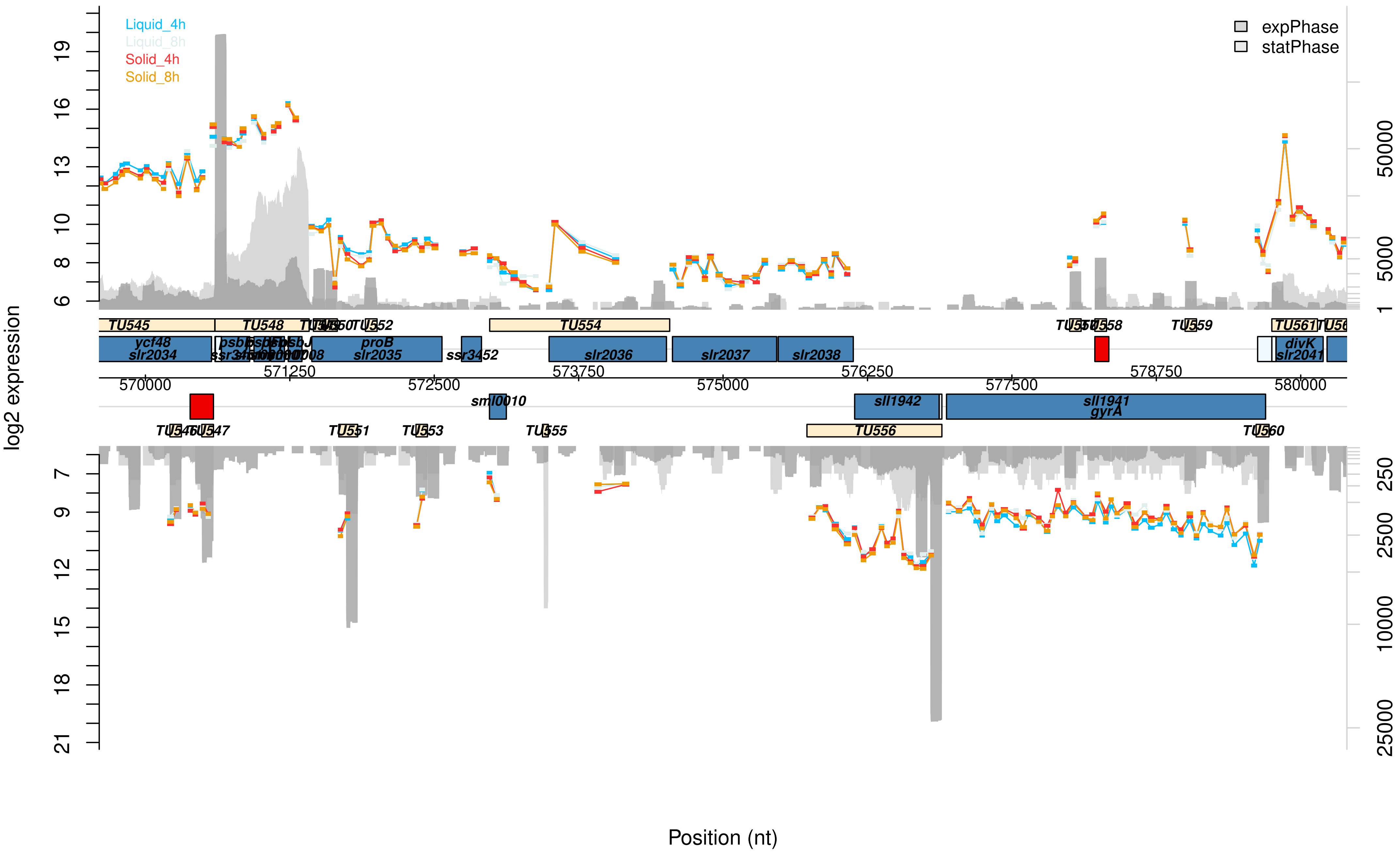

Liquid_4h
Liquid_8h
Solid_4h
Solid_8h
expPhase
statPhase
log2 expression
Position (nt)
TU545
TU548
TU552
TU554
TU558
TU559
ycf48
slr2034
proB
slr2035
ssr3452
slr2036
slr2037
slr2038
divK
slr2041
sml0010
sll1942
sll1941
gyrA
TU551
TU553
TU555
TU556
TU560
570000
571250
572500
573750
575000
576250
577500
578750
580000

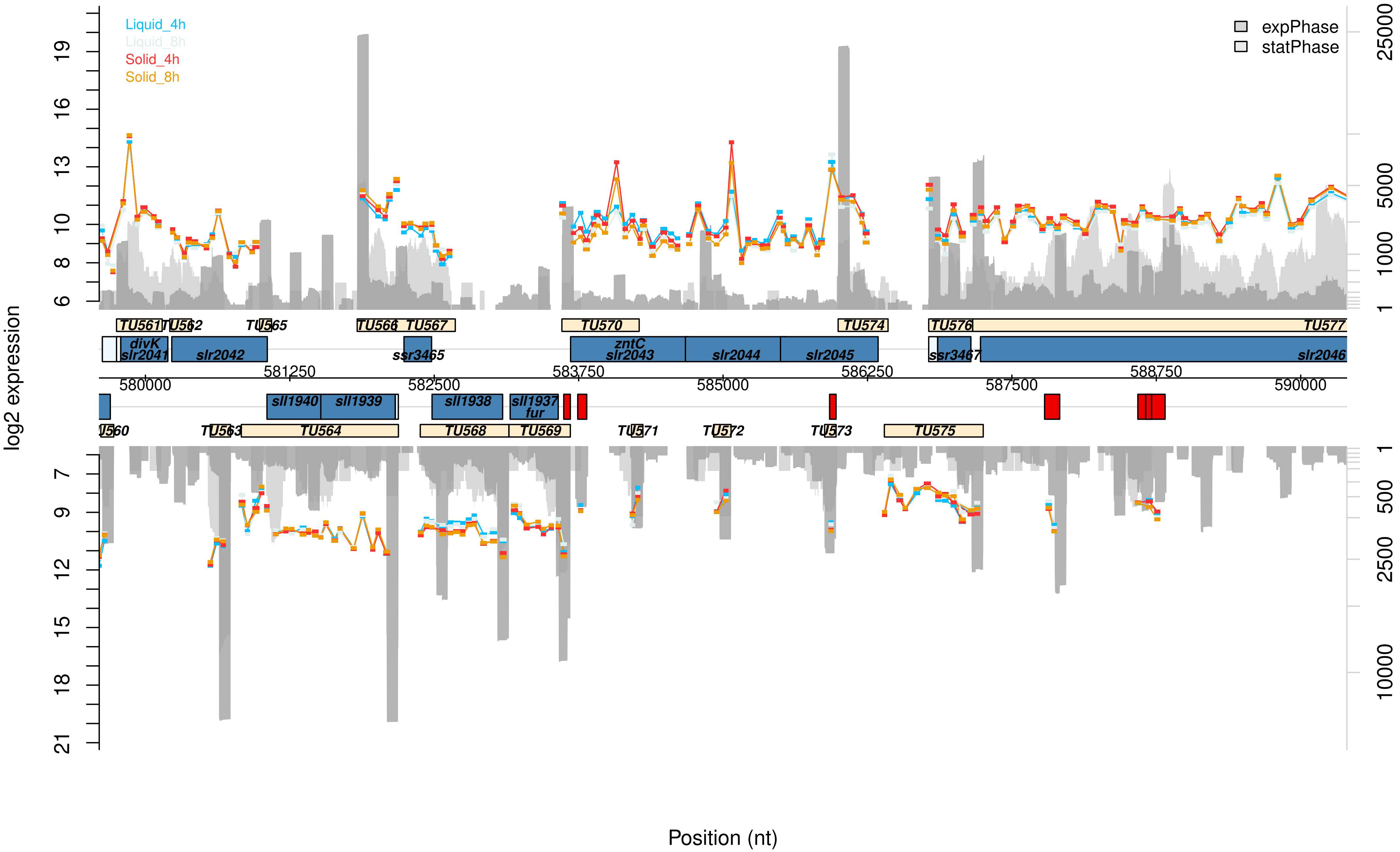
Liquid_4h
Liquid_8h
Solid_4h
Solid_8h
expPhase
statPhase
log2 expression
Position (nt)
TU561
TU562
TU565
TU566
TU567
TU570
TU574
TU576
TU577
divK
slr2041
slr2042
ssr3465
zntC
slr2043
slr2044
slr2045
ssr3467
slr2046
580000
581250
582500
583750
585000
586250
587500
588750
590000
sll1940
sll1939
sll1938
sll1937
fur
TU560
TU563
TU564
TU568
TU569
TU571
TU572
TU573
TU575

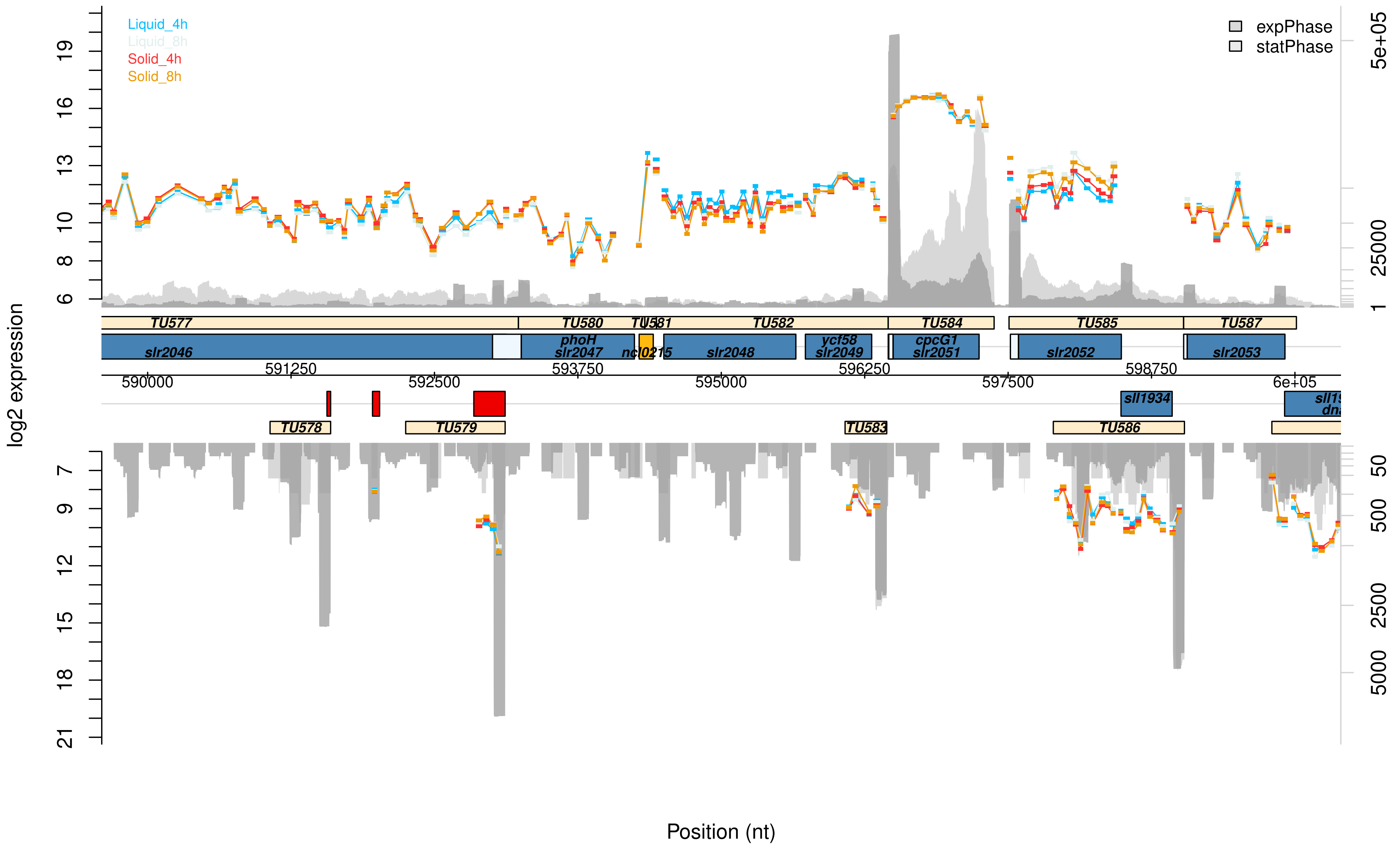

Liquid_4h
Liquid_8h
Solid_4h
Solid_8h
expPhase
statPhase
log2 expression
Position (nt)
TU577
TU578
TU579
TU580
TU581
TU582
TU583
TU584
TU585
TU586
TU587
slr2046
phoH
slr2047
ncl0215
slr2048
ycf58
slr2049
cpcG1
slr2051
slr2052
slr2053
sll1934
590000
591250
592500
593750
595000
596250
597500
598750
6e+05
5e+05
25000
2500
5000

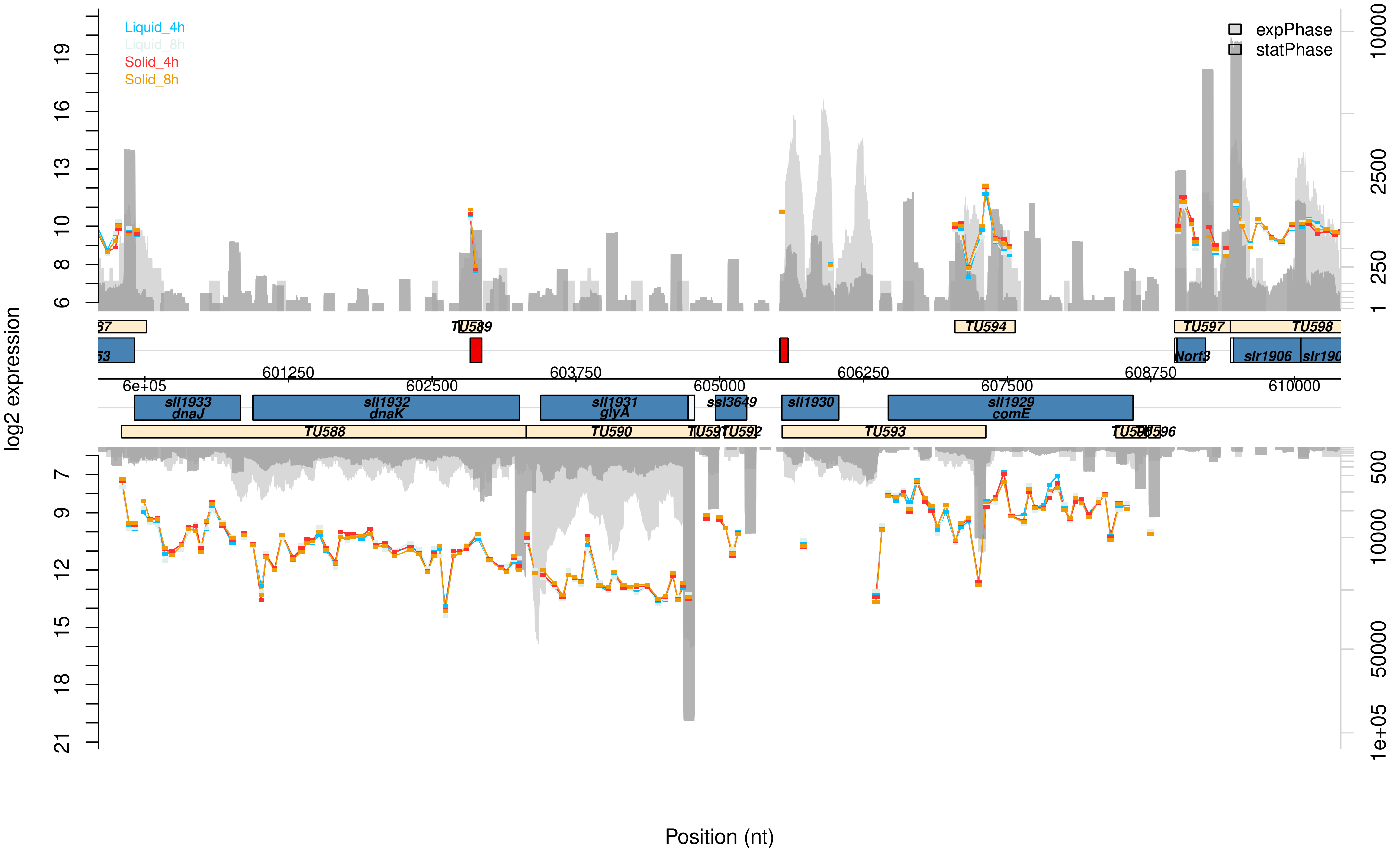

Liquid_4h
Liquid_8h
Solid_4h
Solid_8h
expPhase
statPhase
log2 expression
Position (nt)
TU589
TU594
TU597
TU598
Norf3
slr1906
6e+05
601250
602500
603750
605000
606250
607500
608750
610000
sll1933 dnaJ
sll1932 dnaK
sll1931 glyA
ssl3649
sll1930
sll1929 comE
TU588
TU590
TU591
TU592
TU593
TU596

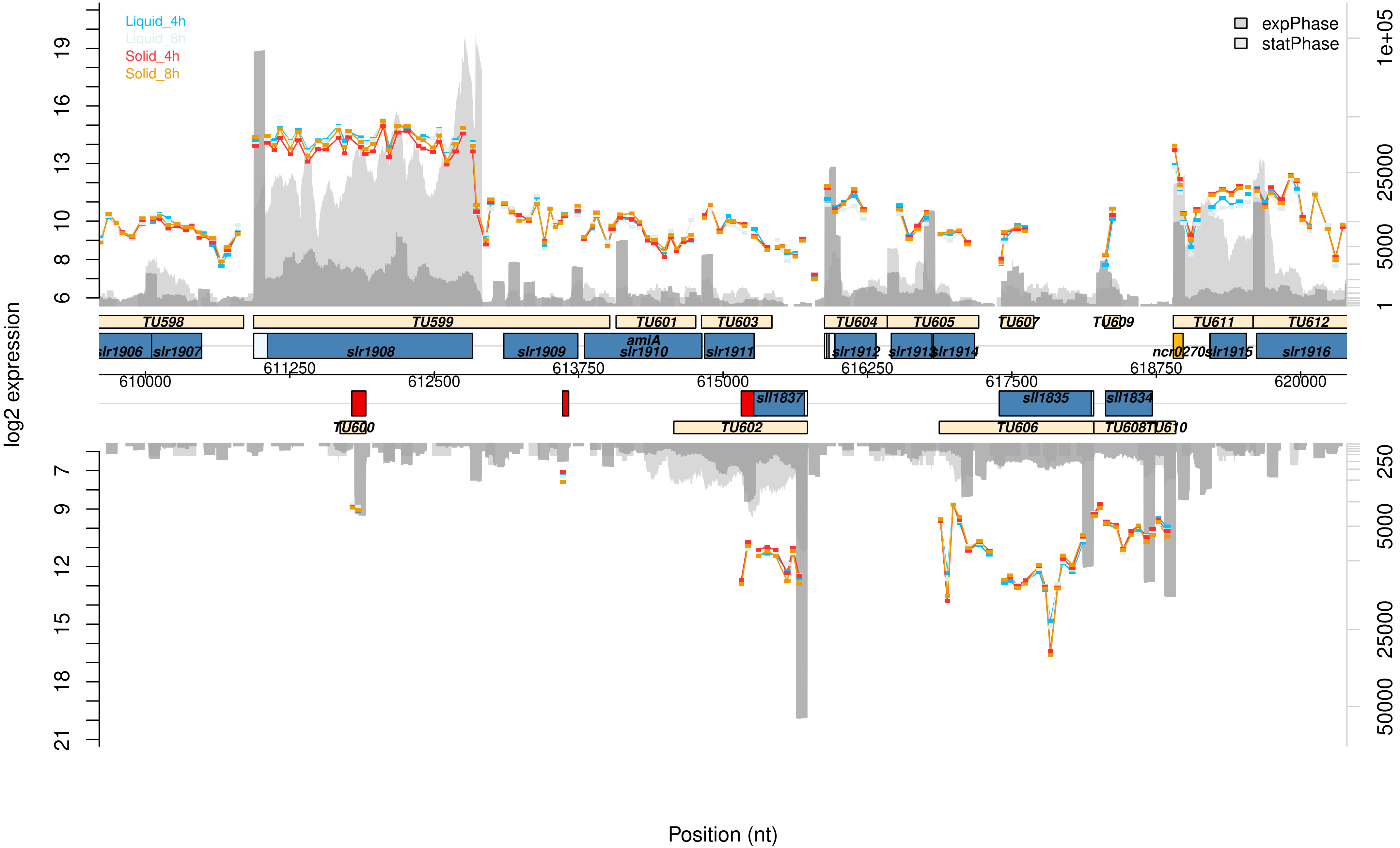

Liquid_4h
Liquid_8h
Solid_4h
Solid_8h
expPhase
statPhase
log2 expression
Position (nt)
TU598
TU599
TU601
TU603
TU604
TU605
TU607
TU609
TU611
TU612
slr1906
slr1907
slr1908
slr1909
amiA
slr1910
slr1911
slr1912
slr1913
slr1914
ncr0270
slr1915
slr1916
610000
611250
612500
613750
615000
616250
617500
618750
620000
sll1837
sll1835
sll1834
TU600
TU602
TU606
TU608
TU610

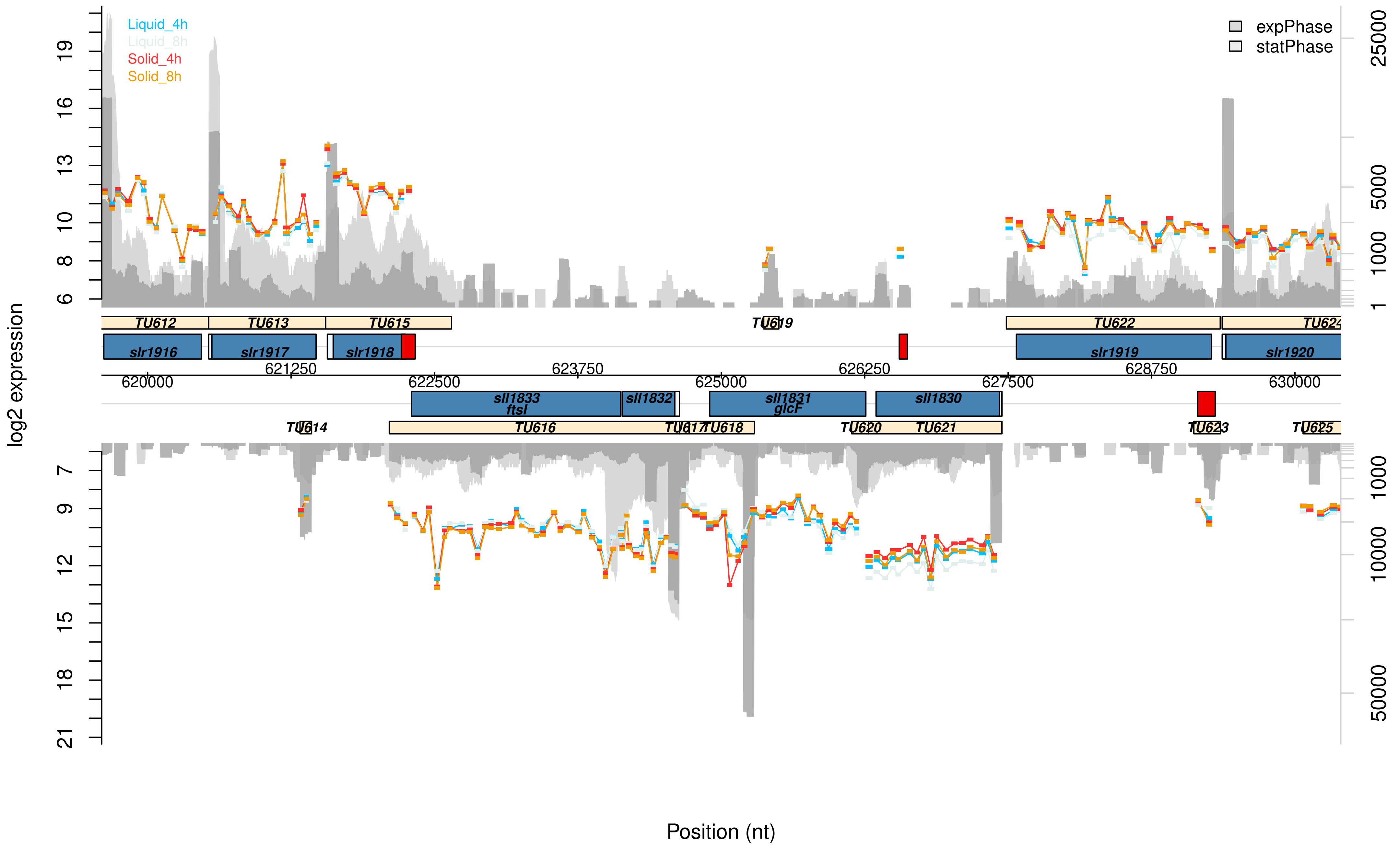
Liquid_4h
Liquid_8h
Solid_4h
Solid_8h
expPhase
statPhase
log2 expression
Position (nt)
TU612
TU613
TU615
TU619
TU622
TU624
slr1916
slr1917
slr1918
slr1919
slr1920
620000
621250
622500
623750
625000
626250
627500
628750
630000
sll1833
ftsI
sll1832
sll1831
glcF
sll1830
TU614
TU616
TU617
TU618
TU620
TU621
TU623
TU625

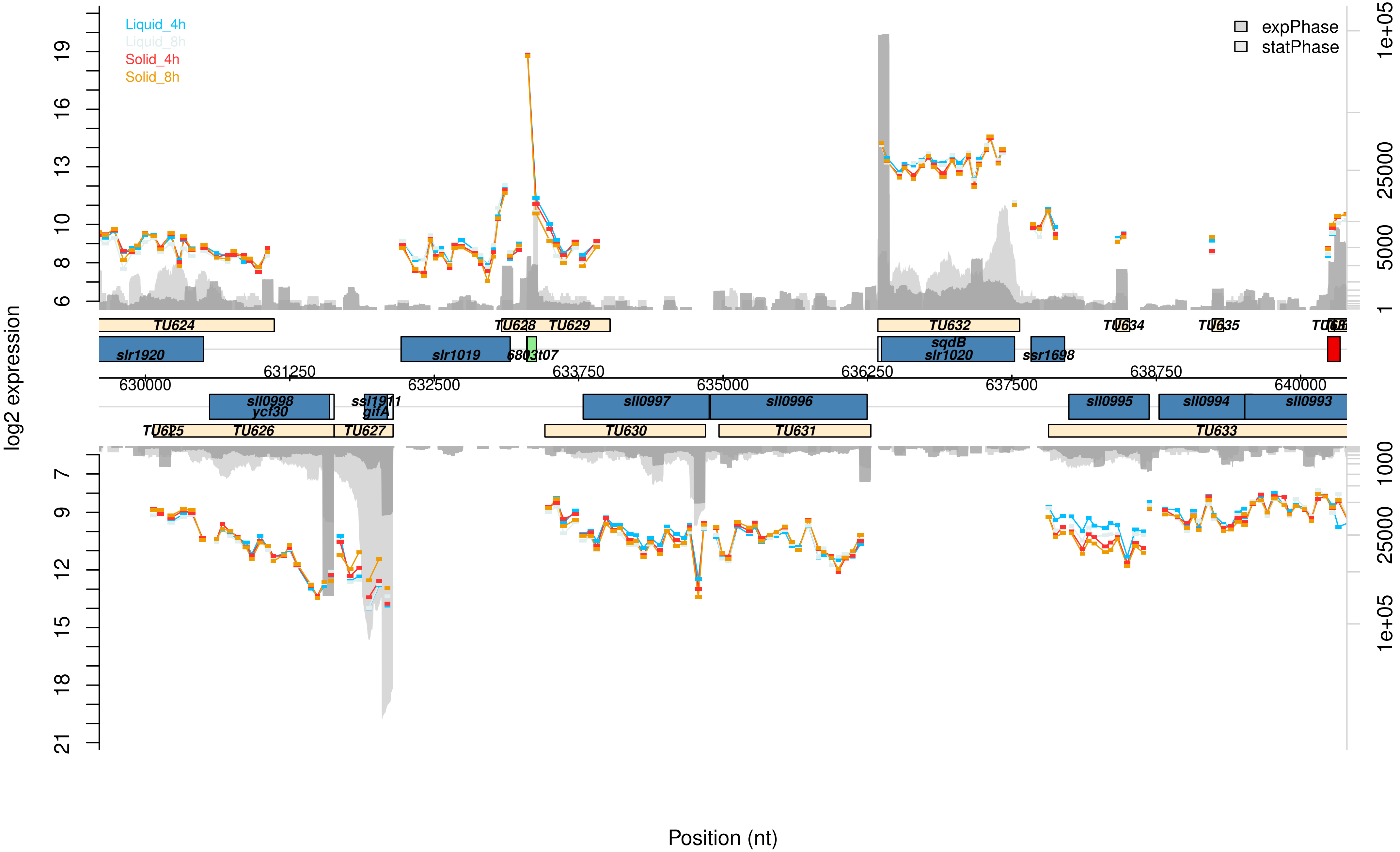

Liquid_4h
Liquid_8h
Solid_4h
Solid_8h
expPhase
statPhase
log2 expression
Position (nt)
TU624
TU628
TU629
TU632
TU634
TU635
slr1920
slr1019
6803t07
sqdB
slr1020
ssr1698
630000
631250
632500
633750
635000
636250
637500
638750
640000
sll0998
ycf30
ssl1911
gifA
sll0997
sll0996
sll0995
sll0994
sll0993
TU625
TU626
TU627
TU630
TU631
TU633
6
8
10
13
16
19
7
9
12
15
18
21
1
5000
25000
1e+05
1000

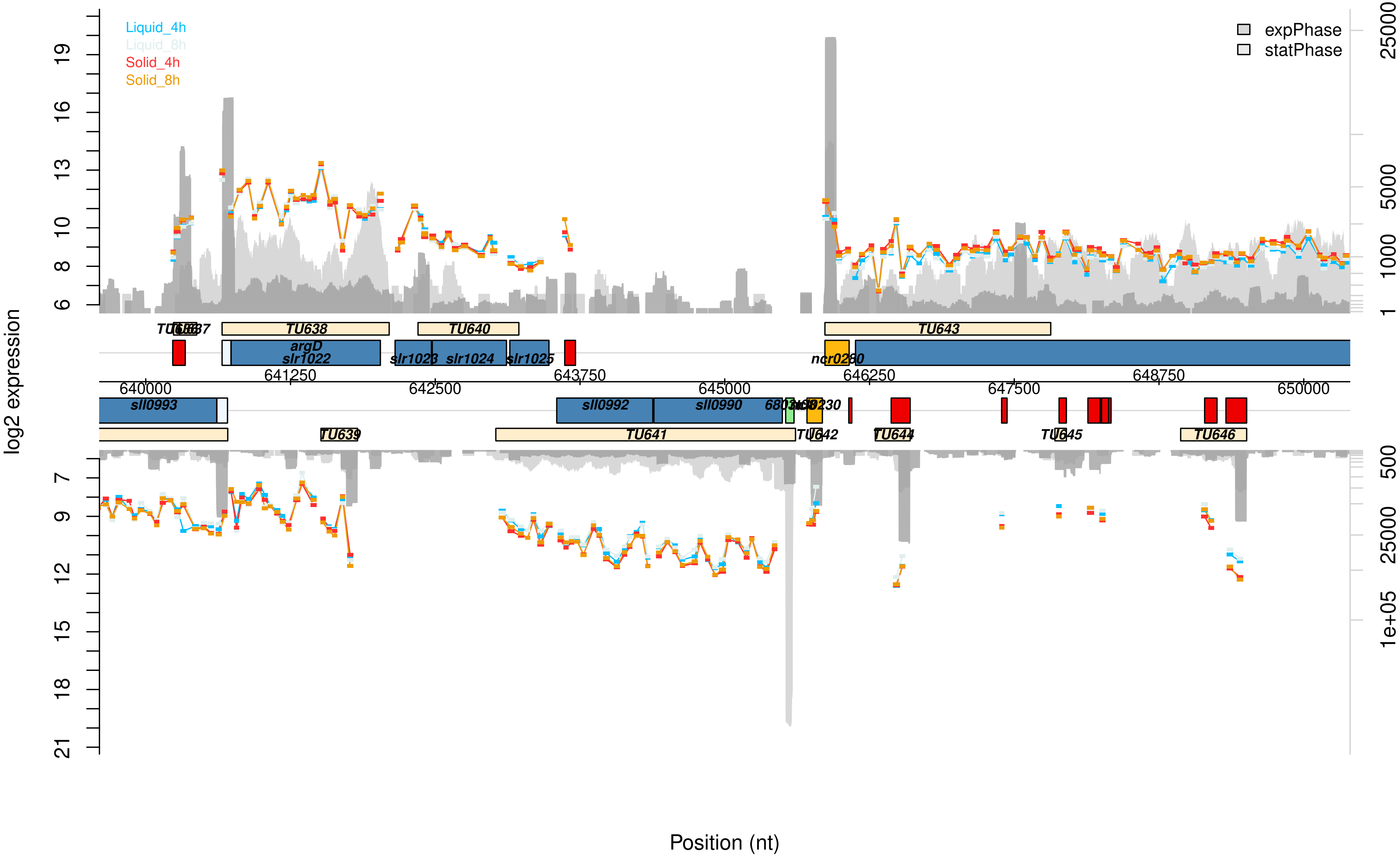

Liquid_4h
Liquid_8h
Solid_4h
Solid_8h
expPhase
statPhase
log2 expression
Position (nt)
TU638
TU640
TU643
argD
slr1022
slr1023
slr1024
slr1025
ncr0260
sll0993
sll0992
sll0990
TU639
TU641
TU642
TU644
TU645
TU646
640000
641250
642500
643750
645000
646250
647500
648750
650000

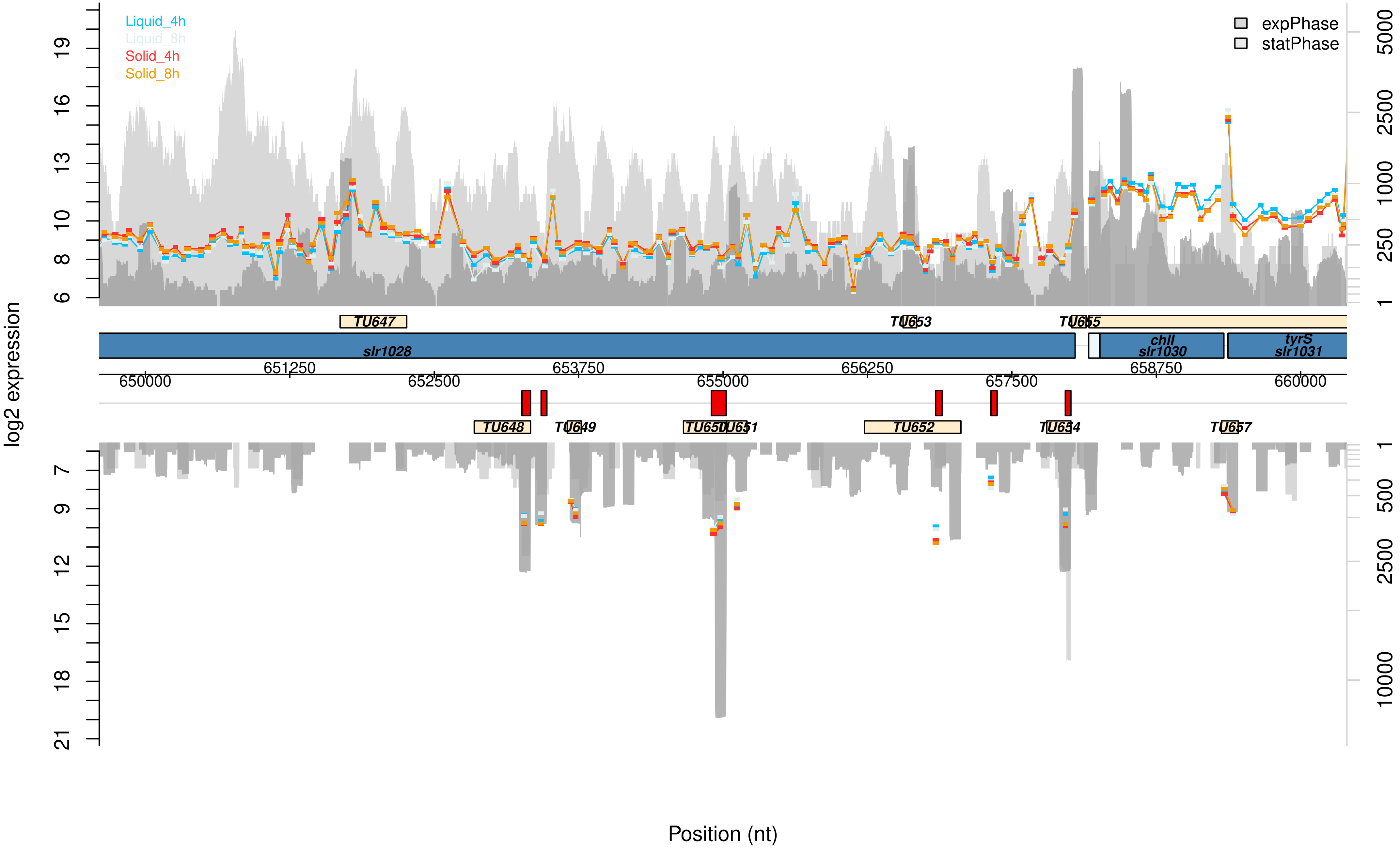

Liquid_4h
Liquid_8h
Solid_4h
Solid_8h
expPhase
statPhase
log2 expression
TU647
TU653
TU655
slr1028
chlI
slr1030
tyrS
slr1031
650000
651250
652500
653750
655000
656250
657500
658750
660000
TU648
TU649
TU650
TU651
TU652
TU654
TU657
Position (nt)

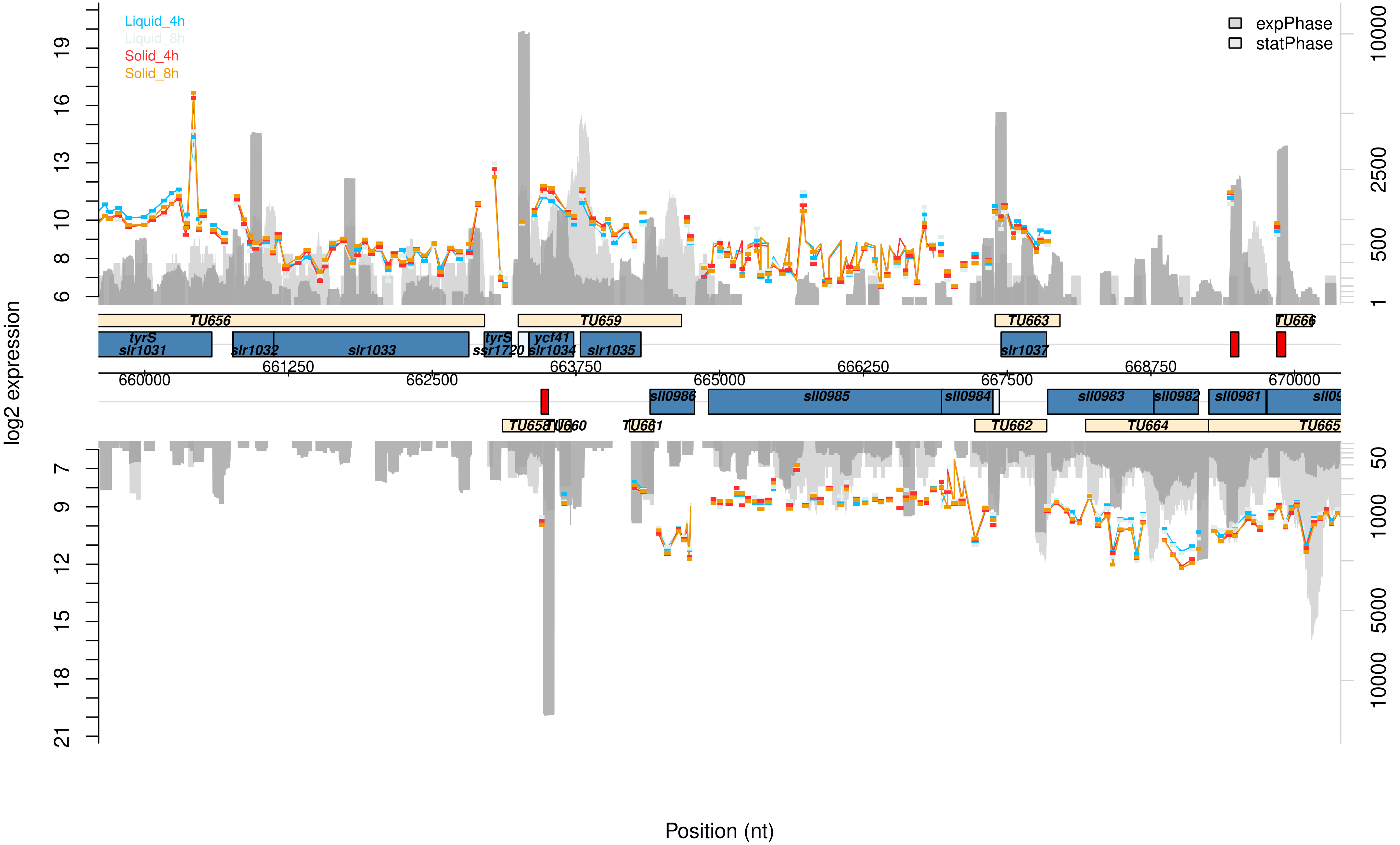

Liquid_4h
Liquid_8h
Solid_4h
Solid_8h
expPhase
statPhase
log2 expression
19
16
13
10
8
6
7
9
12
15
18
21
10000
2500
500
1
50
1000
5000
10000
TU656
TU659
TU663
TU666
tyrS
slr1031
slr1032
slr1033
tyrS
ssr1720
ycf41
slr1034
slr1035
slr1037
661250
663750
666250
668750
660000
662500
665000
667500
670000
sll0986
sll0985
sll0984
sll0983
sll0982
sll0981
TU658
TU660
TU661
TU662
TU664
TU665
Position (nt)

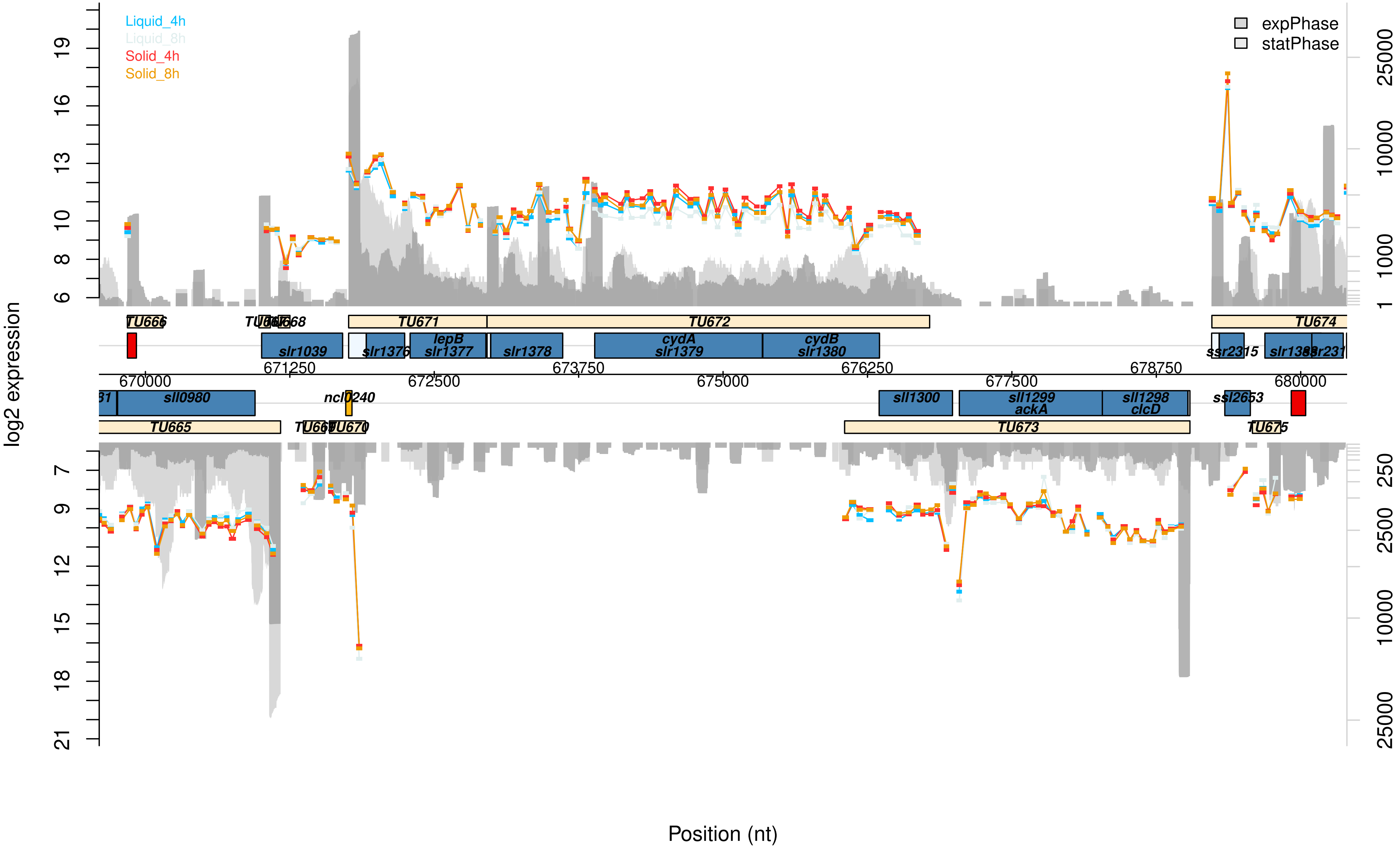

Liquid_4h
Liquid_8h
Solid_4h
Solid_8h
expPhase
statPhase
log2 expression
Position (nt)
670000
671250
672500
673750
675000
676250
677500
678750
680000
TU665
TU666
TU671
TU672
TU673
TU674
sll0980
slr1039
slr1376
lepB
slr1377
slr1378
cydA
slr1379
cydB
slr1380
ncl0240
sll1300
sll1299
ackA
sll1298
clcD
ssl2653

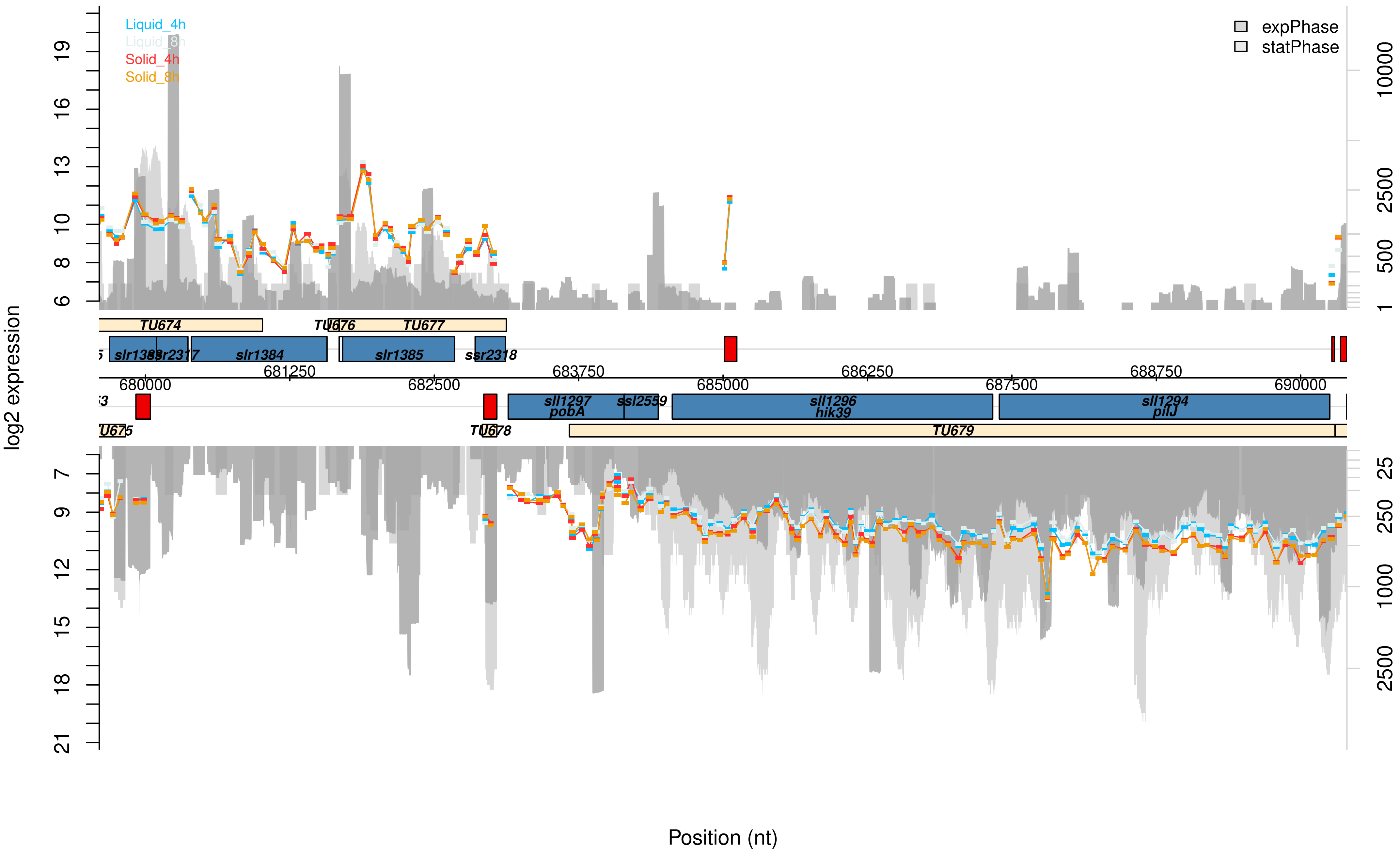

Liquid_4h
Liquid_8h
Solid_4h
Solid_8h
expPhase
statPhase
log2 expression
19
16
13
10
8
6
10000
2500
500
1
TU674
TU676
TU677
slr1383
ssr2317
slr1384
slr1385
ssr2318
681250
683750
686250
688750
680000
682500
685000
687500
690000
sll1297
pobA
ssl2559
sll1296
hik39
sll1294
pilJ
TU675
TU678
TU679
7
9
12
15
18
21
25
250
1000
2500
Position (nt)

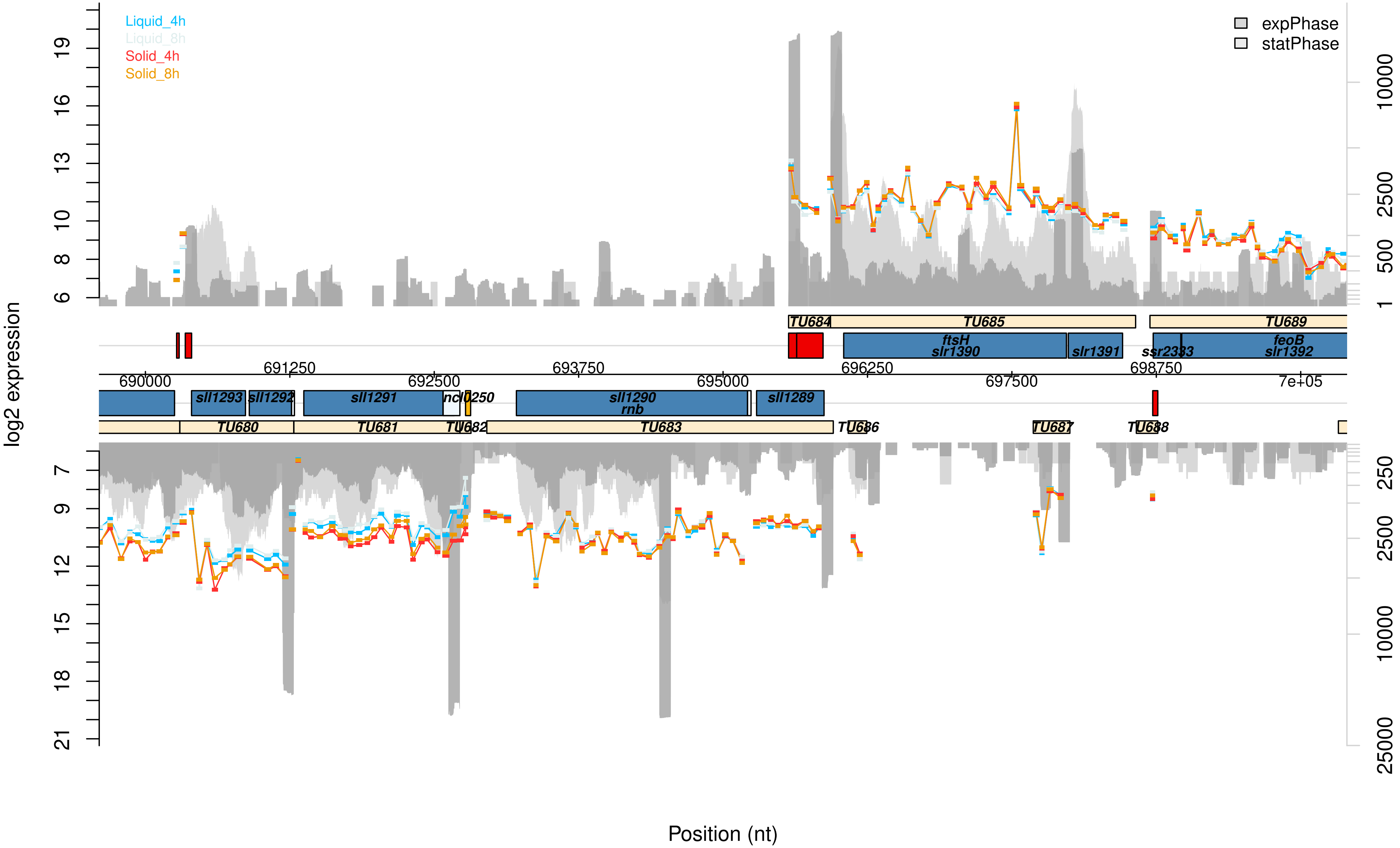

Liquid_4h
Liquid_8h
Solid_4h
Solid_8h
expPhase
statPhase
log2 expression
Position (nt)
TU684
TU685
TU689
ftsH
slr1390
slr1391
ssr2333
feoB
slr1392
690000
691250
692500
693750
695000
696250
697500
698750
7e+05
sll1293
sll1292
sll1291
ncl0250
sll1290
rnb
sll1289
TU680
TU681
TU682
TU683
TU686
TU687
TU688

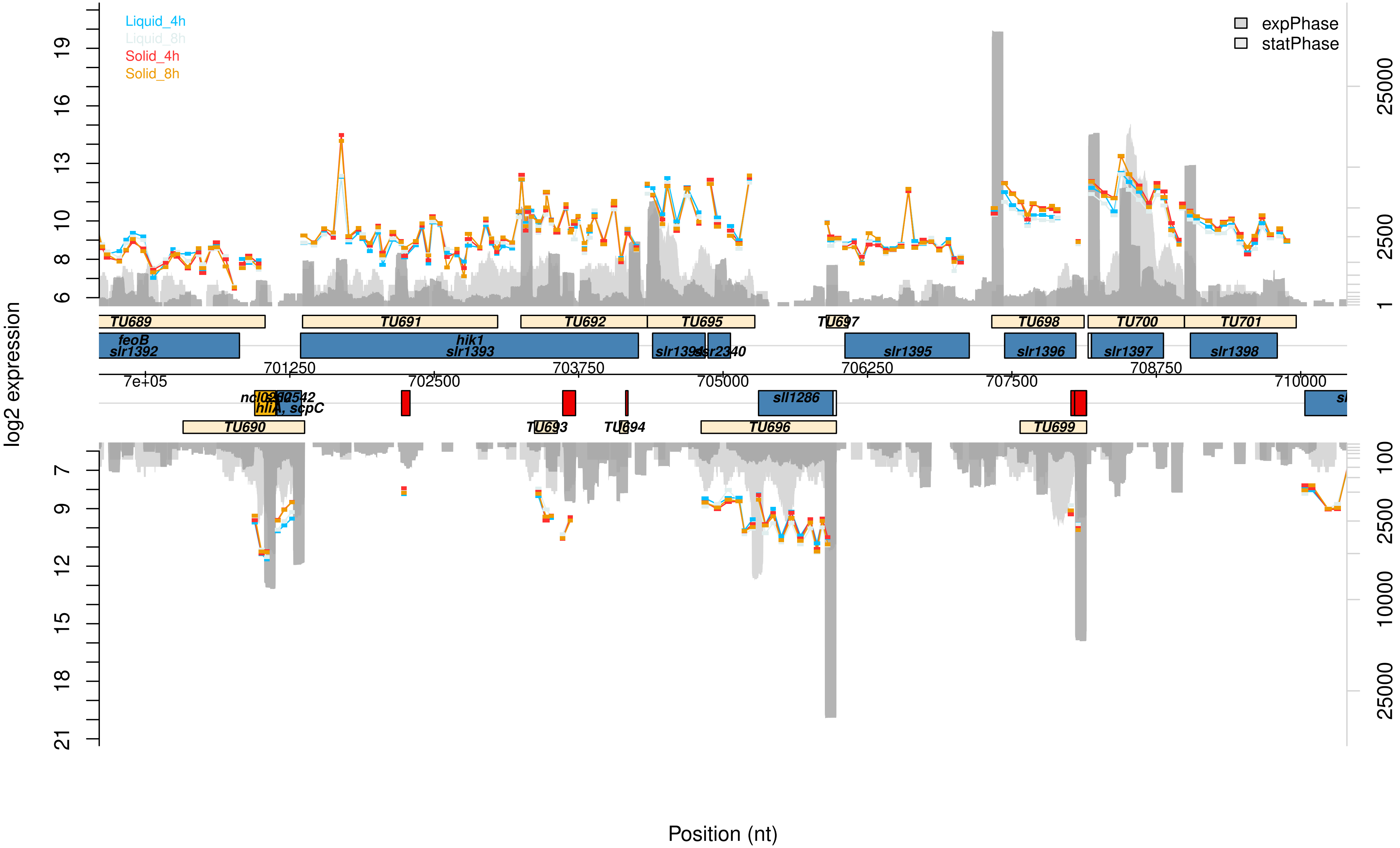

Liquid_4h
Liquid_8h
Solid_4h
Solid_8h
expPhase
statPhase
log2 expression
Position (nt)
TU689
TU691
TU692
TU695
TU697
TU698
TU700
TU701
feoB
slr1392
hik1
slr1393
slr1395
slr1396
slr1397
slr1398
sll1286
TU690
TU693
TU694
TU696
TU699
7e+05
701250
702500
703750
705000
706250
707500
708750
710000

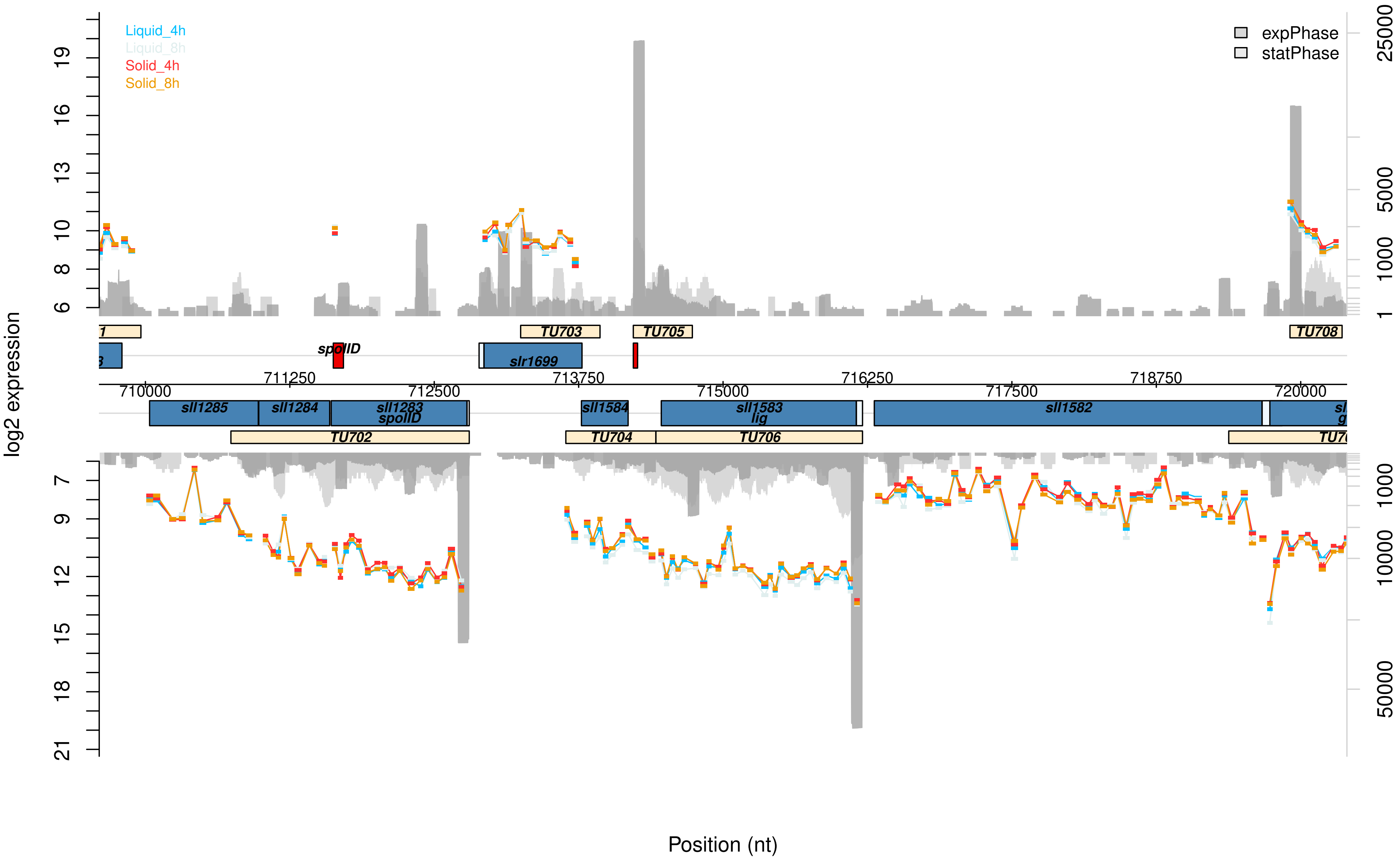

Liquid_4h
Liquid_8h
Solid_4h
Solid_8h
expPhase
statPhase
log2 expression
TU703
TU705
TU708
spoIID
slr1699
710000
711250
712500
713750
715000
716250
717500
718750
720000
sll1285
sll1284
sll1283
spoIID
sll1584
sll1583
lig
sll1582
TU702
TU704
TU706
Position (nt)

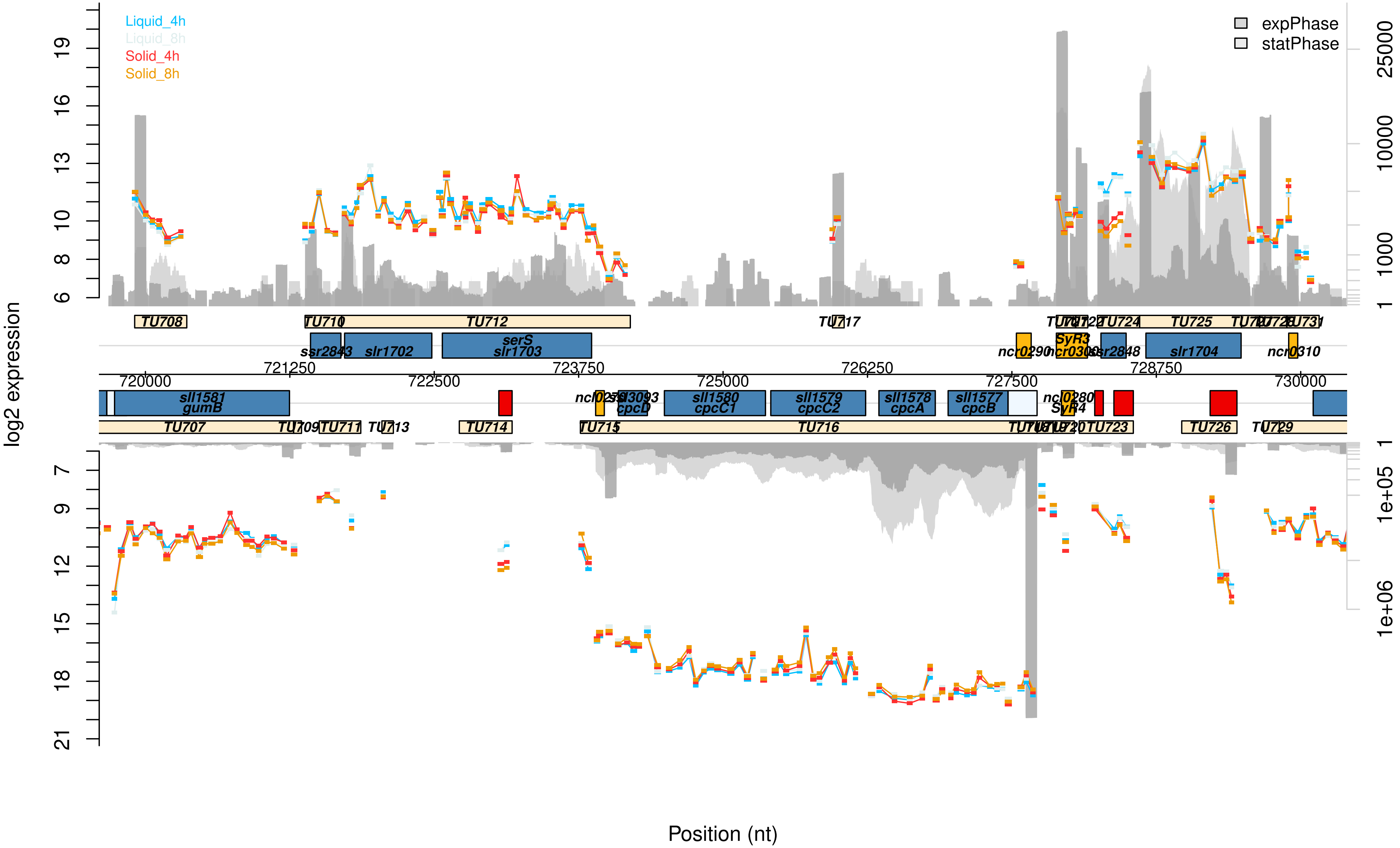

Liquid_4h
Liquid_8h
Solid_4h
Solid_8h
expPhase
statPhase
log2 expression
Position (nt)
TU708
TU710
TU712
TU717
TU725
TU731
ssr2843
slr1702
serS
slr1703
ncr0290
SyR3
slr1704
ncr0310
sll1581
gumB
TU707
TU711
TU714
TU715
TU716
TU723
TU726
sll1580
cpcC1
sll1579
cpcC2
sll1578
cpcA
sll1577
cpcB
cpcD
SyR4
720000
721250
722500
723750
725000
726250
727500
728750
730000

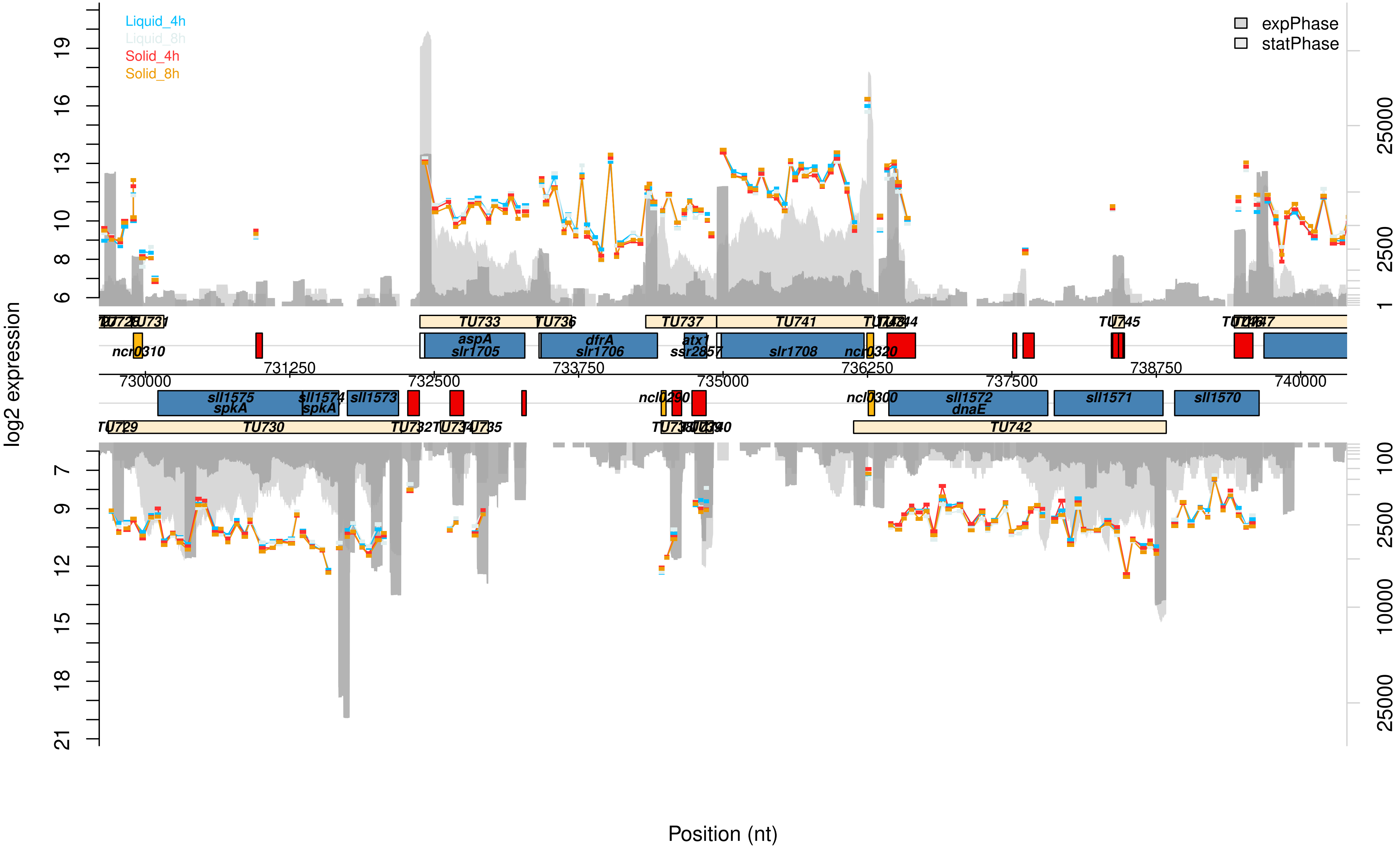

Liquid_4h
Liquid_8h
Solid_4h
Solid_8h
expPhase
statPhase
log2 expression
Position (nt)
aspA
slr1705
dfrA
slr1706
atx1
ssr2857
slr1708
ncr0310
ncr0320
sll1575
spkA
sll1574
sll1573
ncl0290
ncl0300
sll1572
dnaE
sll1571
sll1570
730000
731250
732500
733750
735000
736250
737500
738750
740000

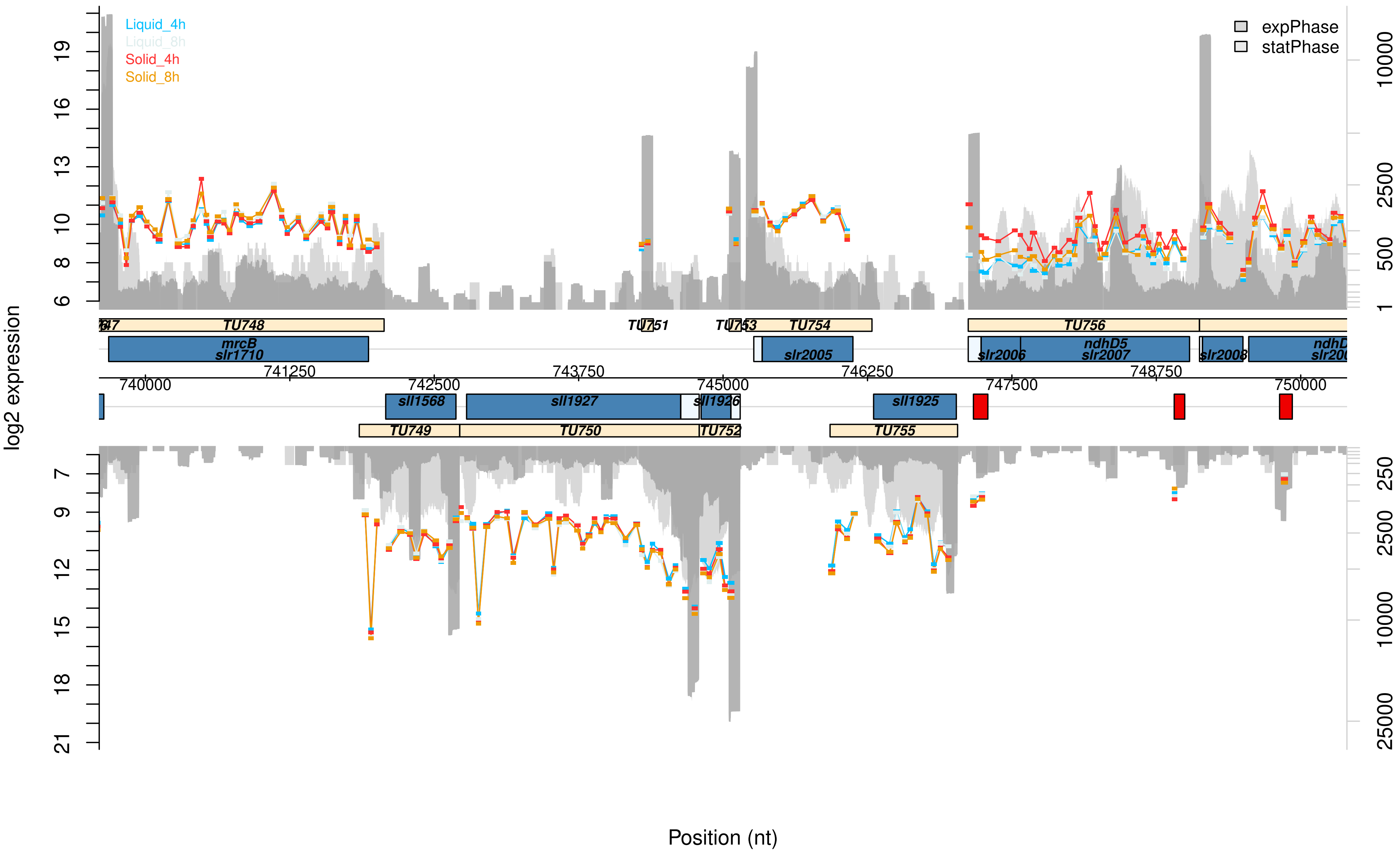

Liquid_4h
Liquid_8h
Solid_4h
Solid_8h
expPhase
statPhase
log2 expression
Position (nt)
TU748
TU751
TU753
TU754
TU756
mrcB
slr1710
slr2005
slr2006
ndhD5
slr2007
slr2008
740000
741250
742500
743750
745000
746250
747500
748750
750000
sll1568
sll1927
sll1926
sll1925
TU749
TU750
TU752
TU755

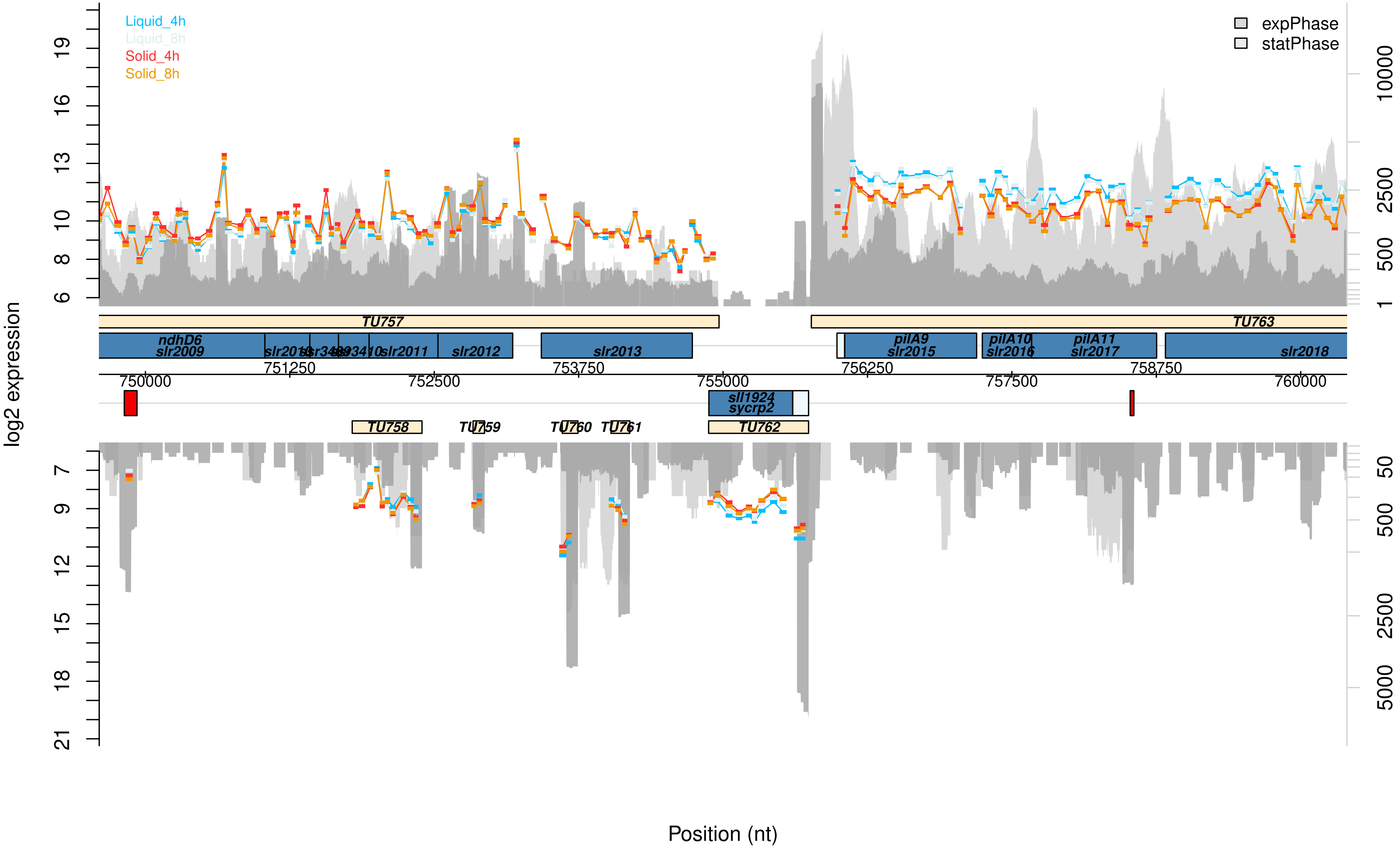
Liquid_4h
Liquid_8h
Solid_4h
Solid_8h
expPhase
statPhase
log2 expression
19
16
13
10
8
6
10000
2500
500
1
TU757
TU763
ndhD6
slr2009
slr2010
ssr3409
ssr3410
slr2011
slr2012
slr2013
pilA9
slr2015
pilA10
slr2016
pilA11
slr2017
slr2018
751250
753750
756250
758750
750000
752500
755000
757500
760000
sll1924
sycrp2
TU758
TU759
TU760
TU761
TU762
7
9
12
15
18
21
50
500
2500
5000
Position (nt)

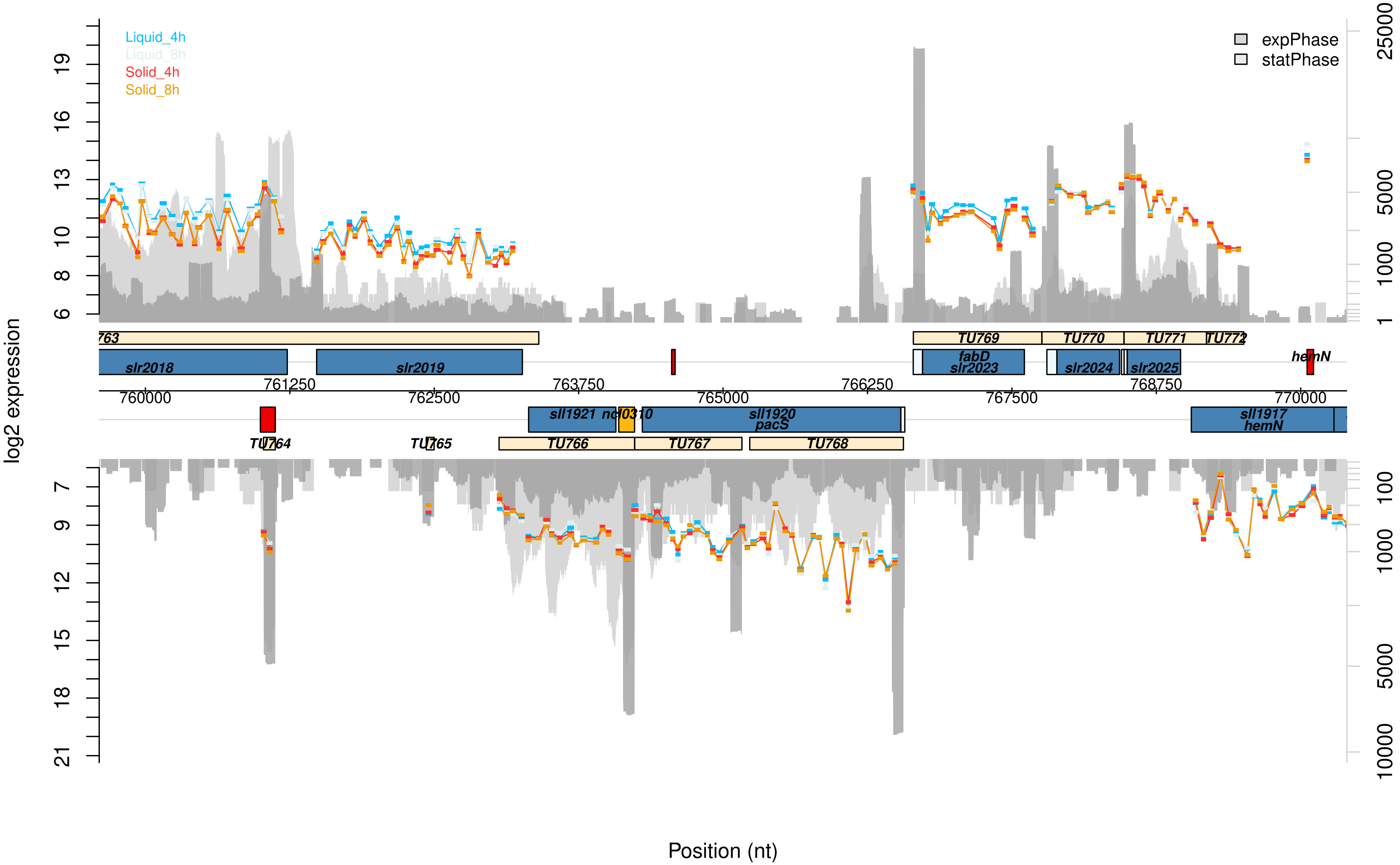

Liquid_4h
Liquid_8h
Solid_4h
Solid_8h
expPhase
statPhase
log2 expression
Position (nt)
19
16
13
10
8
6
7
9
12
15
18
21
25000
5000
1000
1
100
1000
5000
10000
760000
761250
762500
763750
765000
766250
767500
768750
770000
763
TU769
TU770
TU771
TU772
slr2018
slr2019
fabD
slr2023
slr2024
slr2025
hemN
sll1921
ncl0310
sll1920
pacS
sll1917
hemN
TU764
TU765
TU766
TU767
TU768

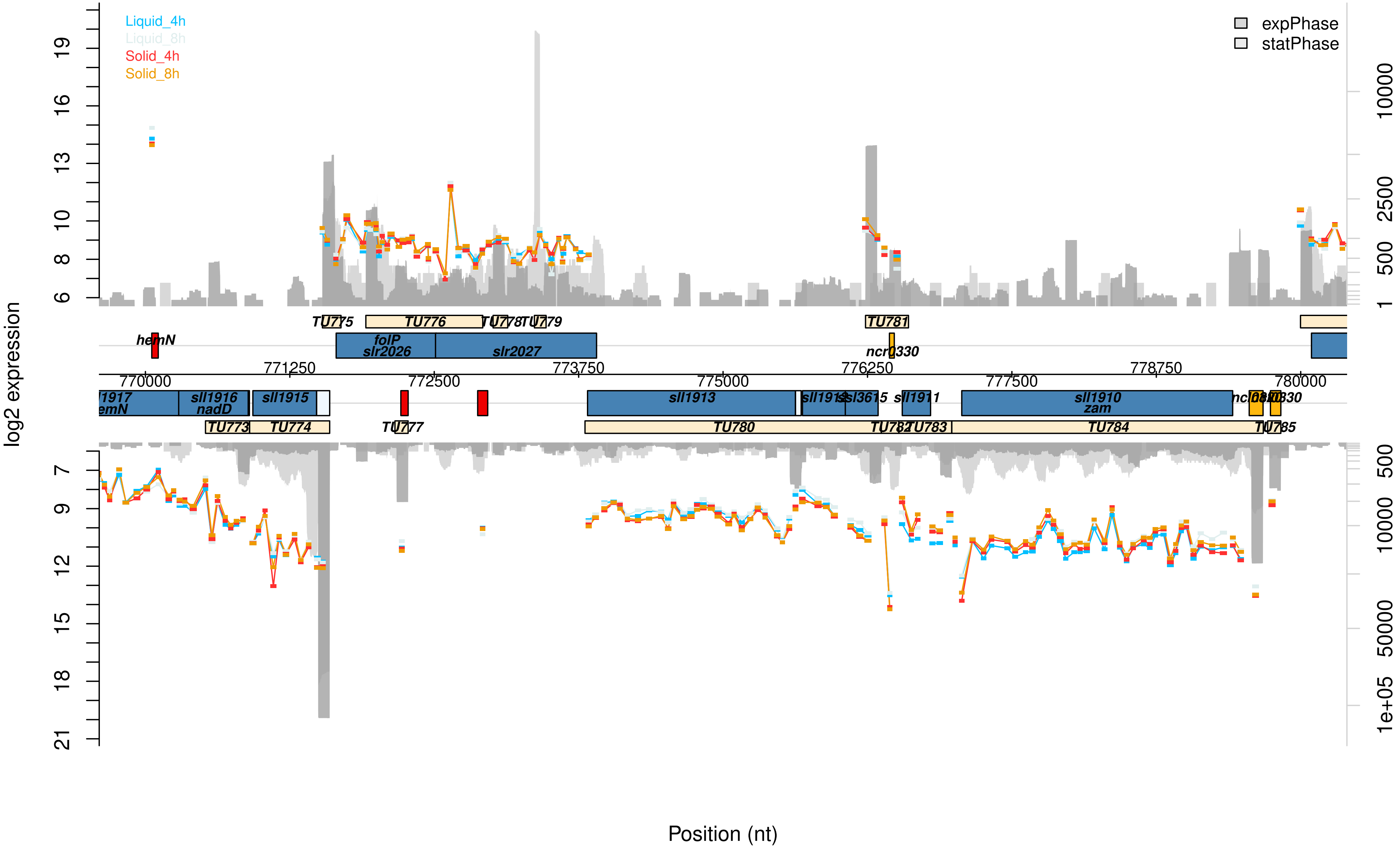

Liquid_4h
Liquid_8h
Solid_4h
Solid_8h
expPhase
statPhase
log2 expression
Position (nt)
TU775
TU776
TU778
TU779
TU781
hemN
folP
slr2026
slr2027
ncr0330
771250
772500
773750
775000
776250
777500
778750
780000
770000
sll1916
nadD
sll1915
sll1913
sll3615
sll1911
sll1910
zam
TU773
TU774
TU777
TU780
TU783
TU784
TU785

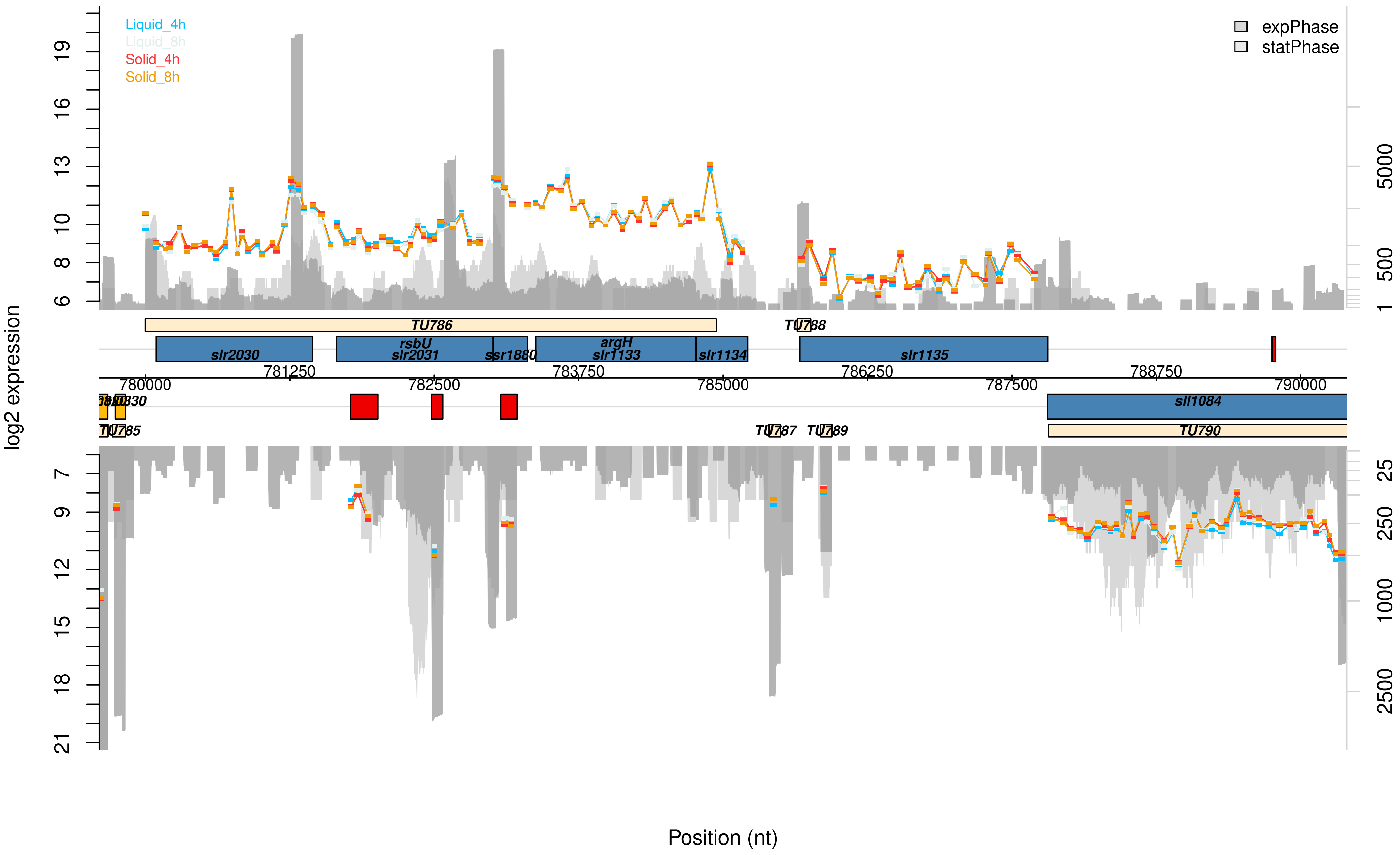

Liquid_4h
Liquid_8h
Solid_4h
Solid_8h
expPhase
statPhase
log2 expression
19
16
13
10
8
6
5000
500
1
TU786
TU788
slr2030
rsbU
slr2031
ssr1880
argH
slr1133
slr1134
slr1135
780000
781250
782500
783750
785000
786250
787500
788750
790000
sll1084
TU785
TU787
TU789
TU790
7
9
12
15
18
21
25
250
1000
2500
Position (nt)

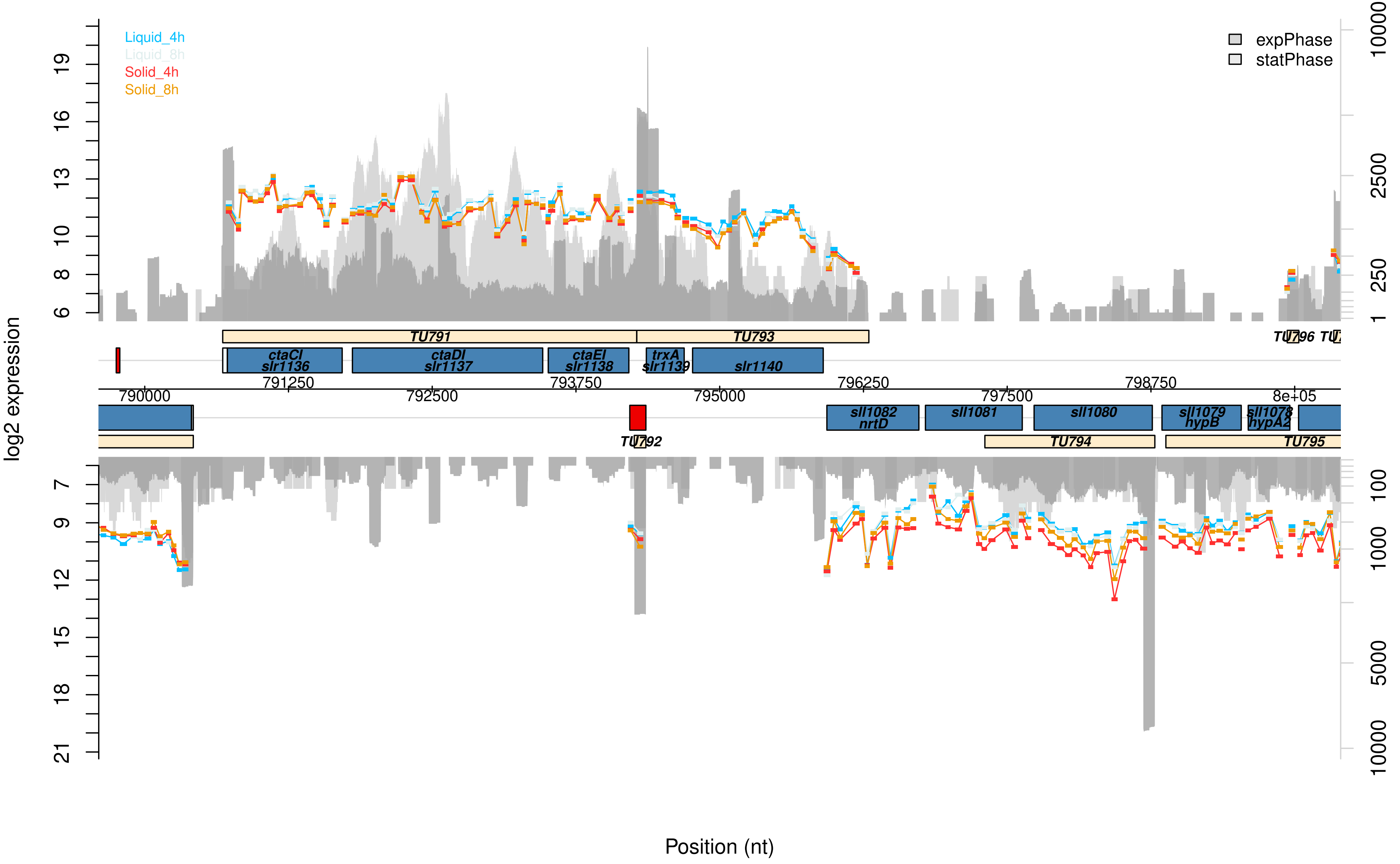

Liquid_4h
Liquid_8h
Solid_4h
Solid_8h
expPhase
statPhase
log2 expression
TU791
TU793
TU796
ctaCI
slr1136
ctaDI
slr1137
ctaEI
slr1138
trxA
slr1139
slr1140
790000
791250
792500
793750
795000
796250
797500
798750
8e+05
sll1082
nrtD
sll1081
sll1080
sll1079
hypB
sll1078
hypA2
TU792
TU794
TU795
Position (nt)

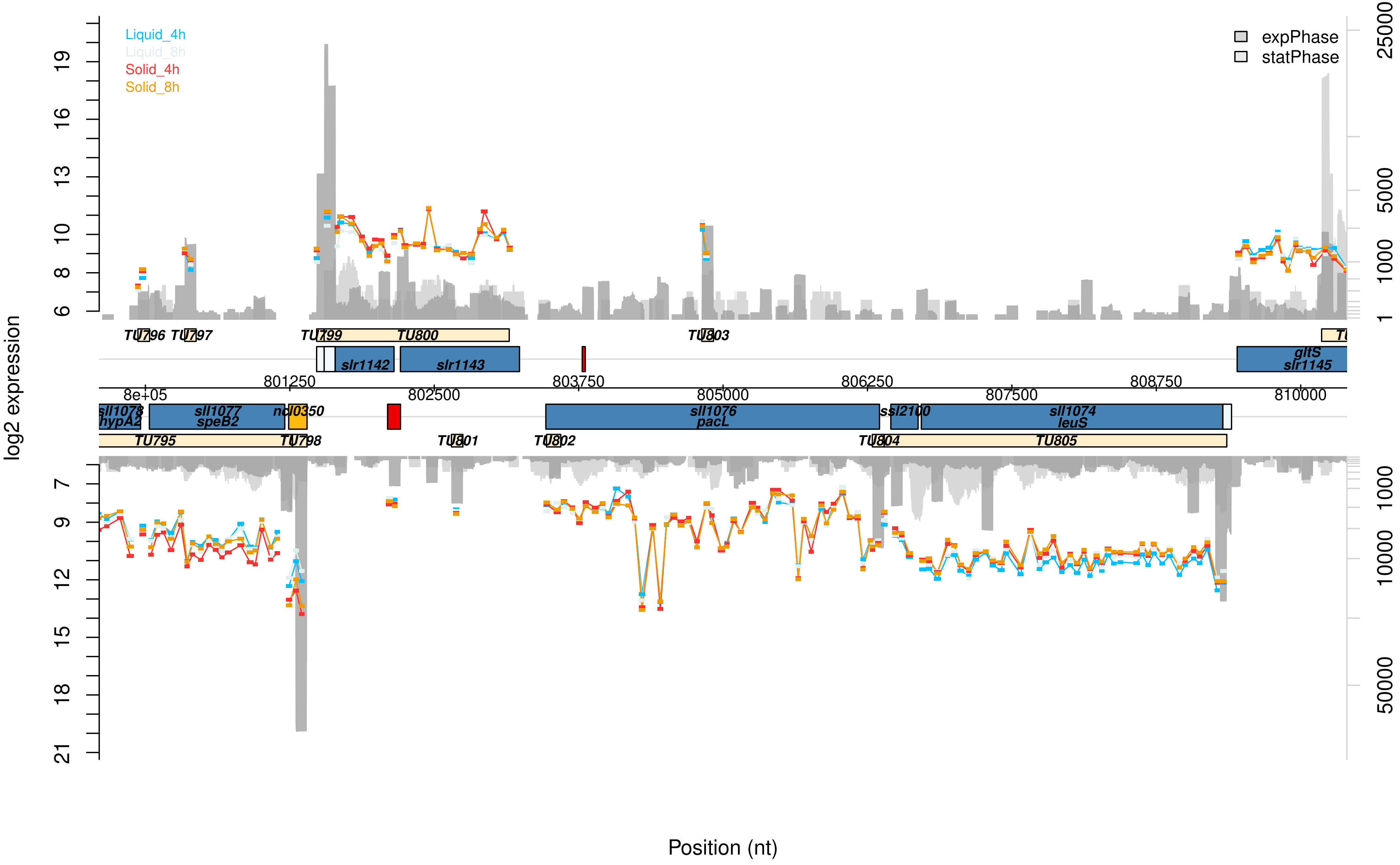

Liquid_4h
Liquid_8h
Solid_4h
Solid_8h
expPhase
statPhase
log2 expression
Position (nt)
TU796 TU797 TU799 TU800 TU803
slr1142 slr1143
gltS slr1145
801250 803750 806250 808750
8e+05 802500 805000 807500 810000
sll1078 hypA2
sll1077 speB2
ncl0350
sll1076 pacL
ssl2100
sll1074 leuS
TU795 TU798 TU801 TU802 TU804 TU805
6 8 10 13 16 19
7 9 12 15 18 21
1 1000 5000 25000
1000 10000 50000

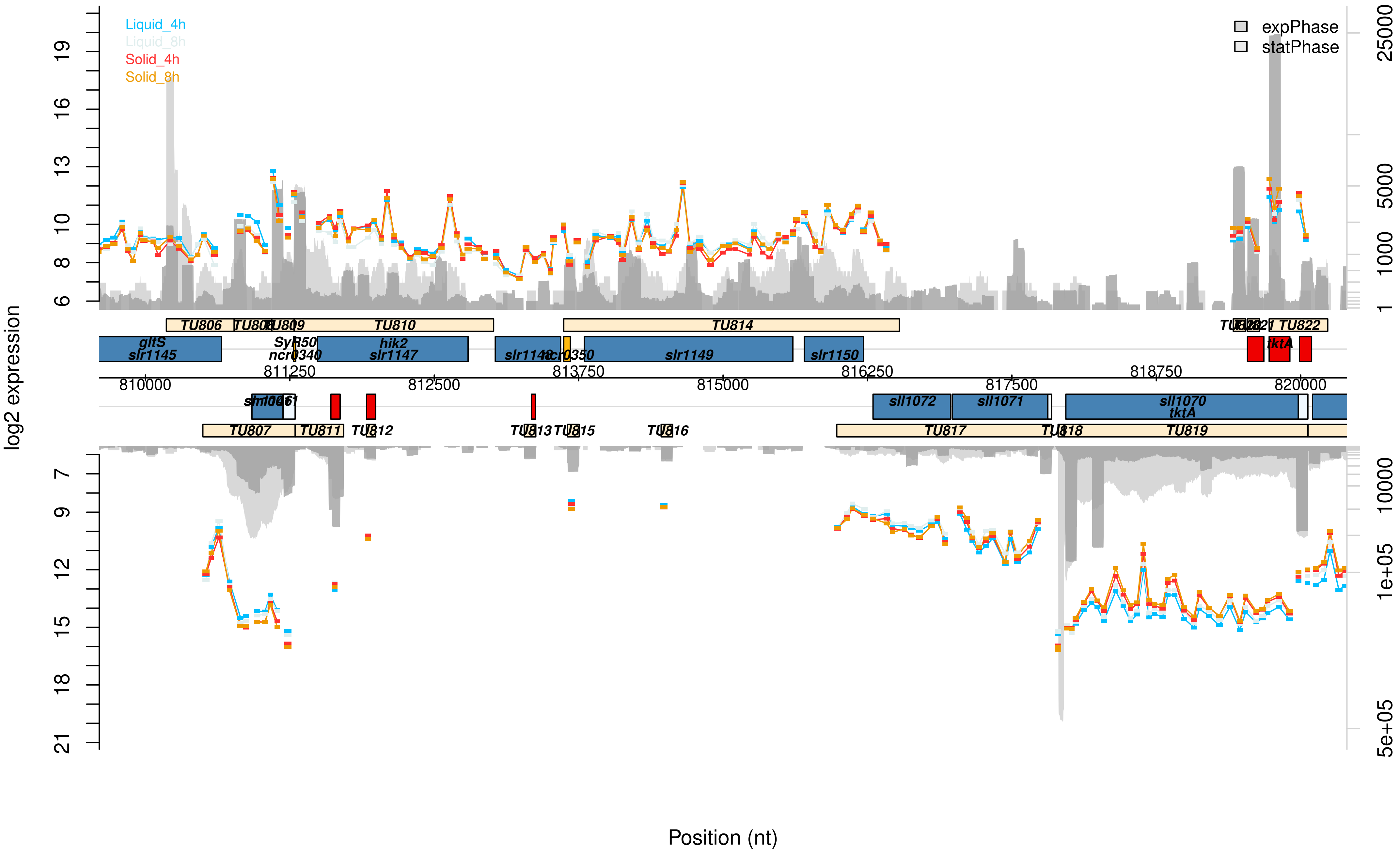

Liquid_4h
Liquid_8h
Solid_4h
Solid_8h
expPhase
statPhase
log2 expression
Position (nt)
TU806
TU808
TU809
TU810
TU814
TU822
gltS
slr1145
SyR50
ncr0340
hik2
slr1147
slr1148
ncr0350
slr1149
slr1150
tktA
810000
811250
812500
813750
815000
816250
817500
818750
820000
sll1072
sll1071
sll1070
tktA
TU807
TU811
TU812
TU813
TU815
TU816
TU817
TU818
TU819

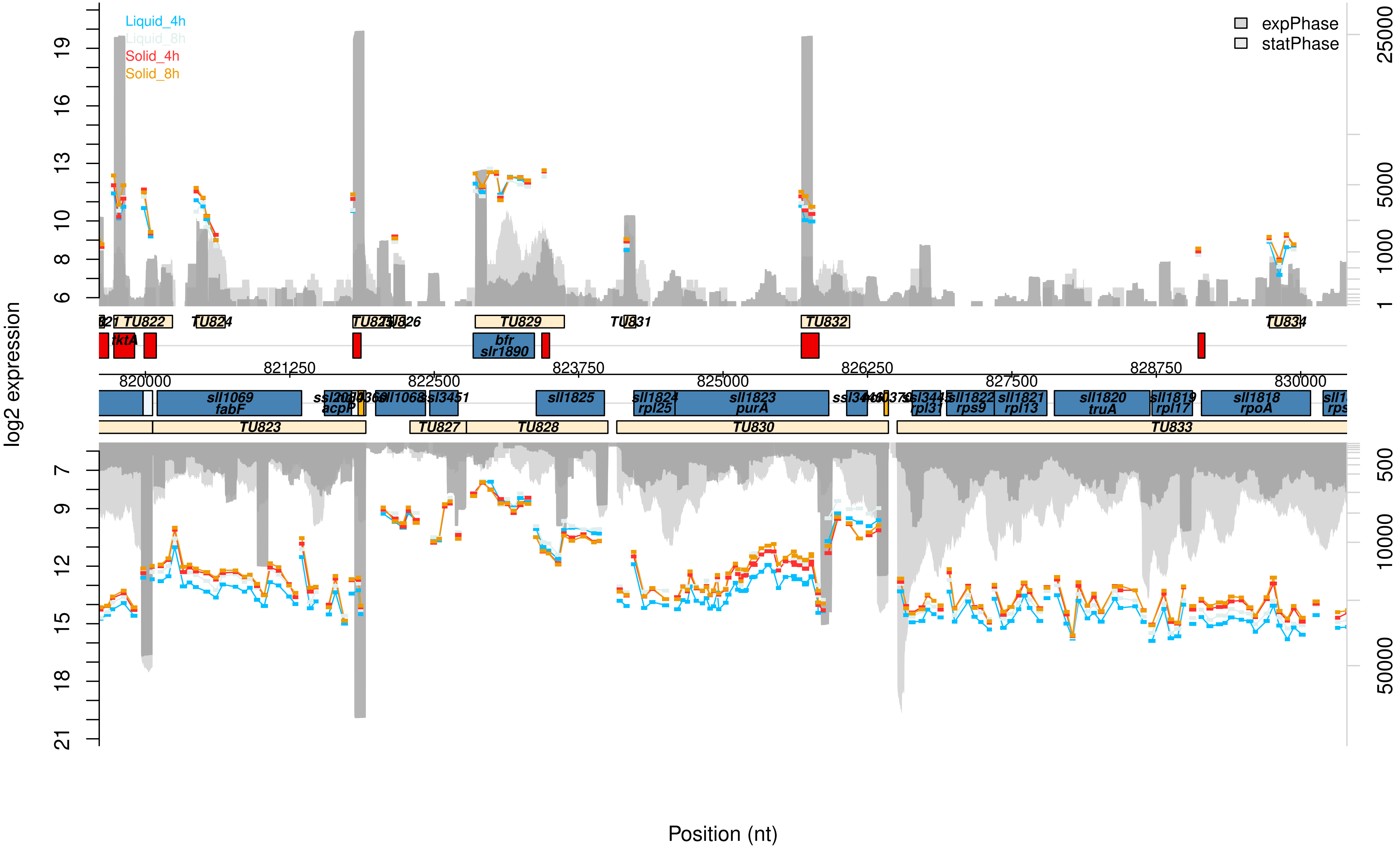

Liquid_4h
Liquid_8h
Solid_4h
Solid_8h
expPhase
statPhase
log2 expression
Position (nt)
TU822
TU824
TU829
TU831
TU832
TU834
tktA
bfr
slr1890
sll1069
fabF
acpP
sll1068
ssl3451
sll1825
sll1824
rpl25
sll1823
purA
ssl3445
rpl31
sll1822
rps9
sll1821
rpl13
sll1820
truA
sll1819
rpl17
sll1818
rpoA
TU823
TU827
TU828
TU830
TU833
820000
821250
822500
823750
825000
826250
827500
828750
830000

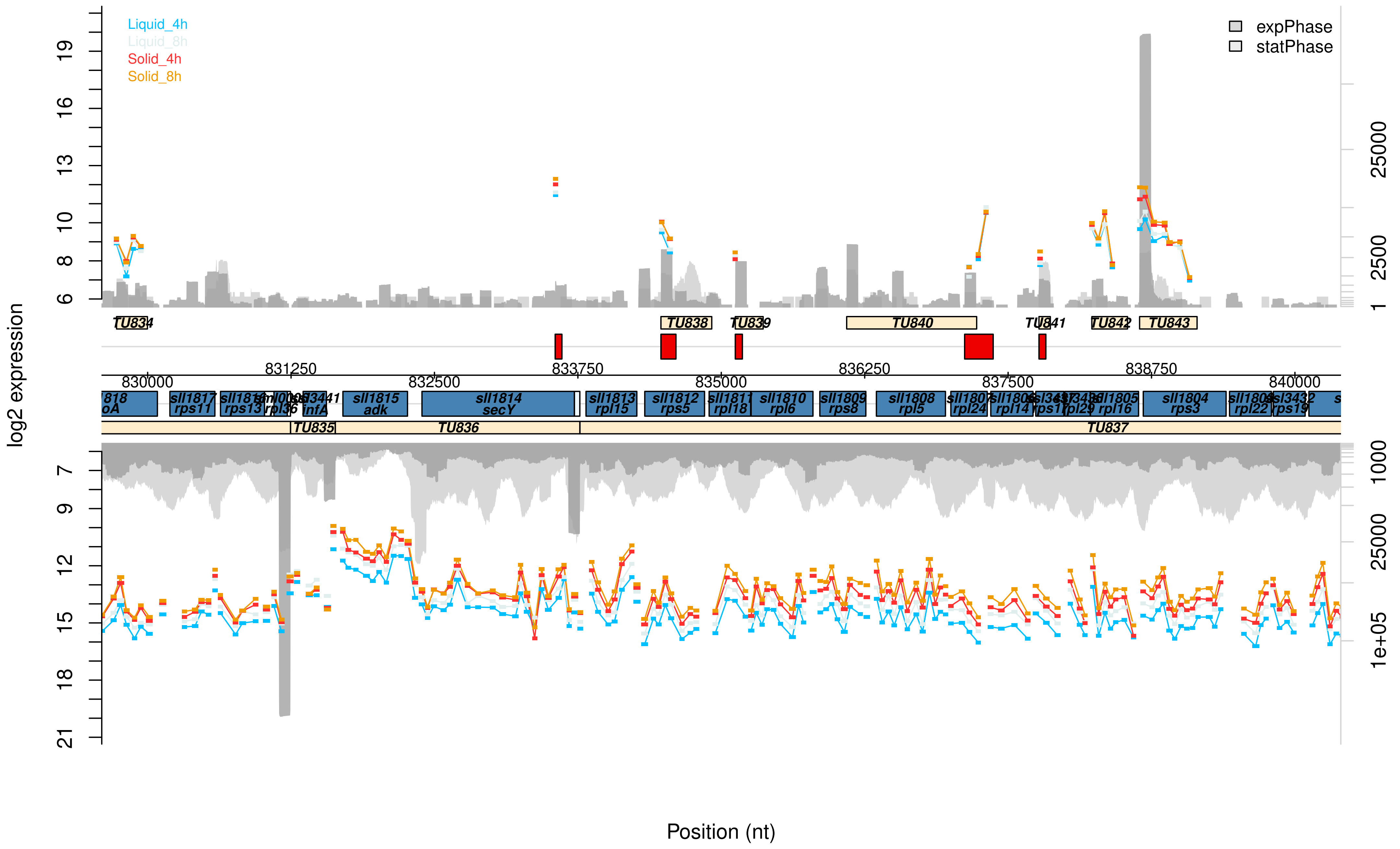
Liquid_4h
Liquid_8h
Solid_4h
Solid_8h
expPhase
statPhase
log2 expression
Position (nt)
TU834
TU835
TU836
TU837
TU838
TU839
TU840
TU841
TU842
TU843
830000
831250
832500
833750
835000
836250
837500
838750
840000
sll1817 rps11
sll1815 adk
sll1814 secY
sll1813 rpl15
sll1812 rps5
sll1811 rpl18
sll1810 rpl6
sll1809 rps8
sll1808 rpl5
rpl24
rpl14
rpl29
sll1805 rpl16
sll1804 rps3
rpl22
rps19
infA
rps13
rpl36

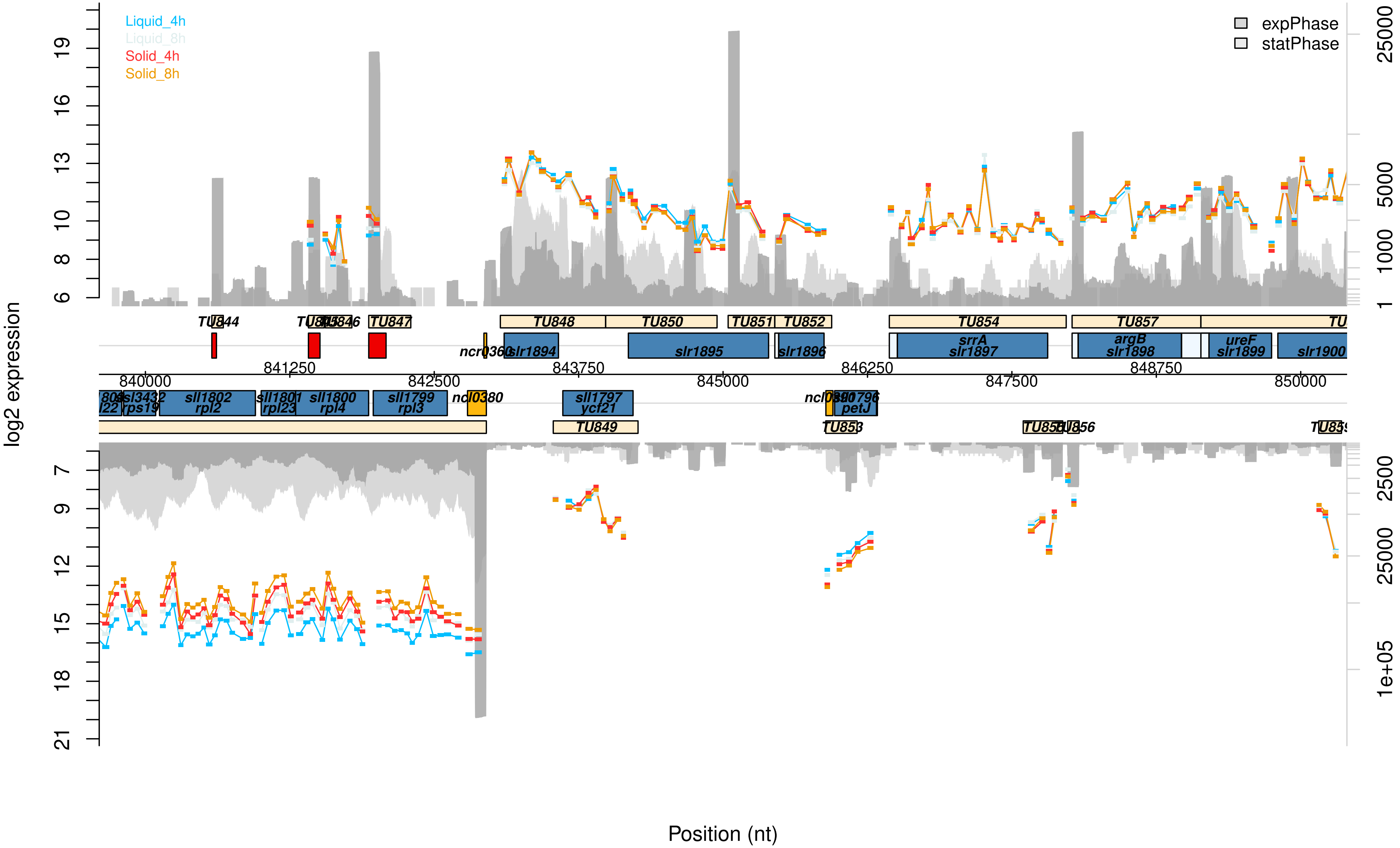

Liquid_4h
Liquid_8h
Solid_4h
Solid_8h
expPhase
statPhase
log2 expression
Position (nt)
TU844
TU846
TU847
TU848
TU850
TU851
TU852
TU854
TU857
TU849
TU853
TU856
ncr0360
slr1894
slr1895
slr1896
srrA
slr1897
argB
slr1898
ureF
slr1899
slr1900
sll1802
rpl2
sll1801
rpl23
sll1800
rpl4
sll1799
rpl3
ncl0380
sll1797
ycf21
petJ
ssl3432
rps19
840000
841250
842500
843750
845000
846250
847500
848750
850000

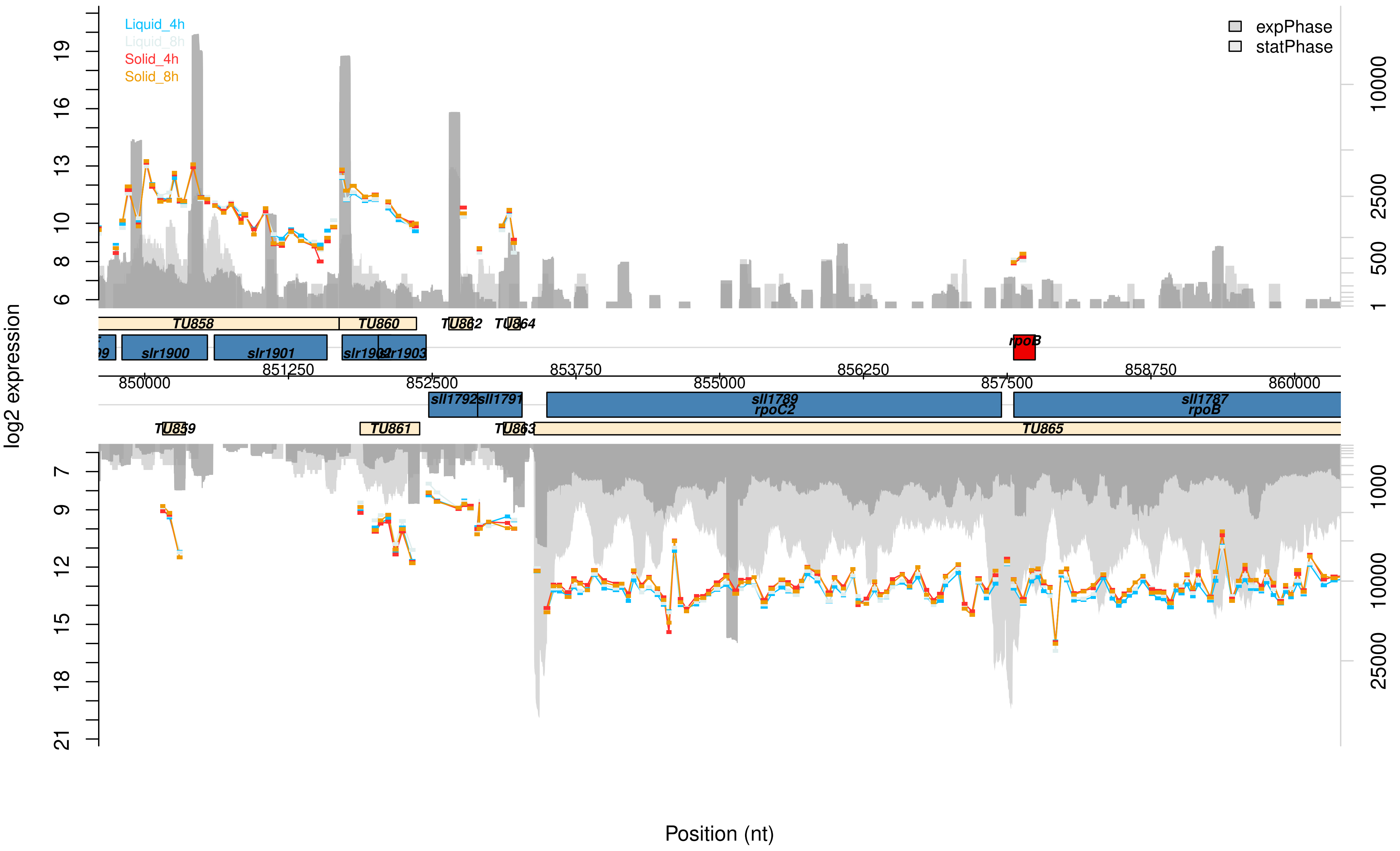

Liquid_4h
Liquid_8h
Solid_4h
Solid_8h
expPhase
statPhase
log2 expression
TU858
TU860
TU862
TU864
slr1900
slr1901
slr1902
slr1903
rpoB
sll1792
sll1791
sll1789
rpoC2
sll1787
rpoB
TU859
TU861
TU863
TU865
850000
851250
852500
853750
855000
856250
857500
858750
860000
Position (nt)

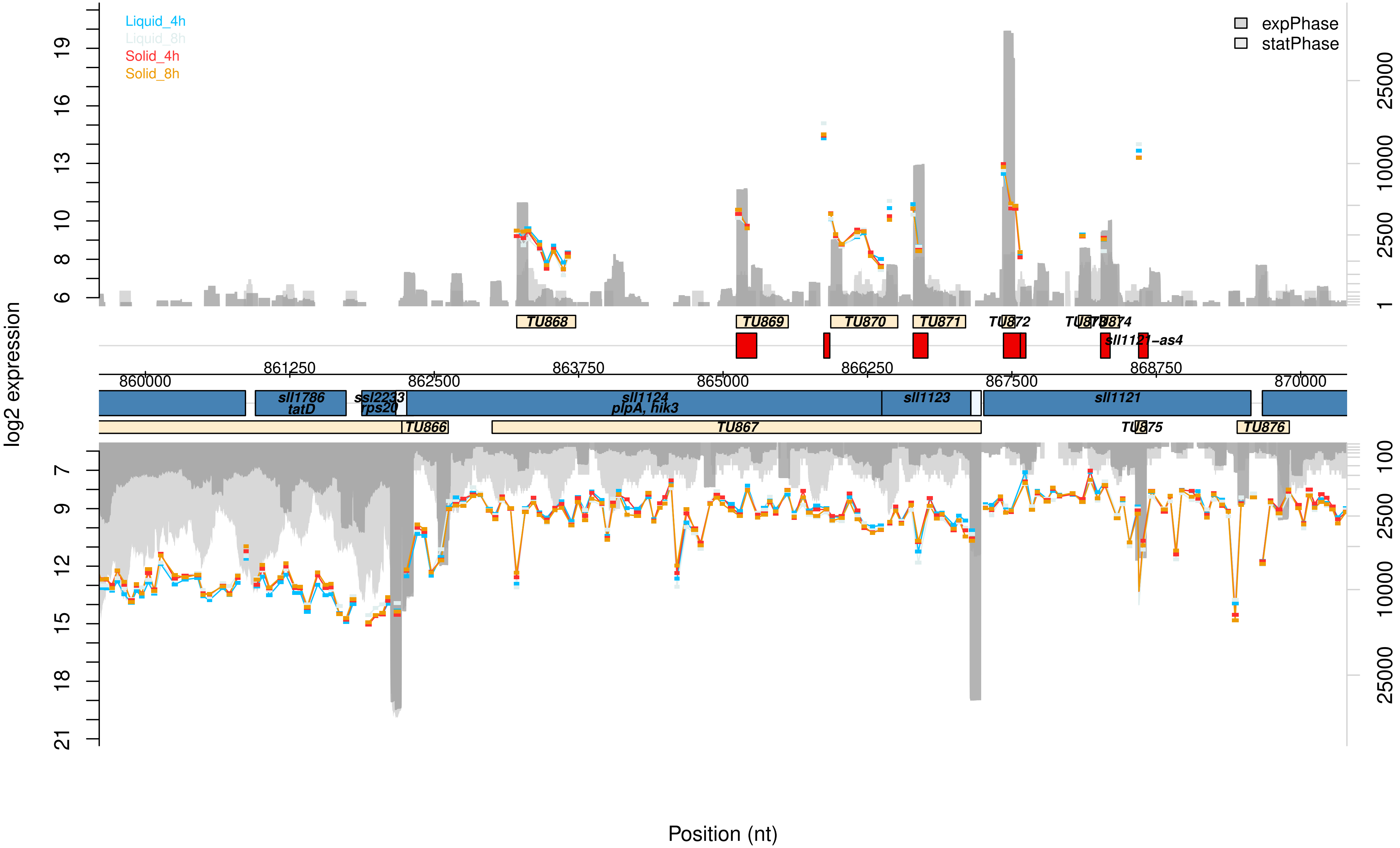

Liquid_4h
Liquid_8h
Solid_4h
Solid_8h
expPhase
statPhase
log2 expression
TU868
TU869
TU870
TU871
TU872
TU873
TU874
sll1121-as4
860000
861250
862500
863750
865000
866250
867500
868750
870000
sll1786
tatD
ssl2233
rps20
sll1124
plpA, hik3
sll1123
sll1121
TU866
TU867
TU875
TU876
Position (nt)

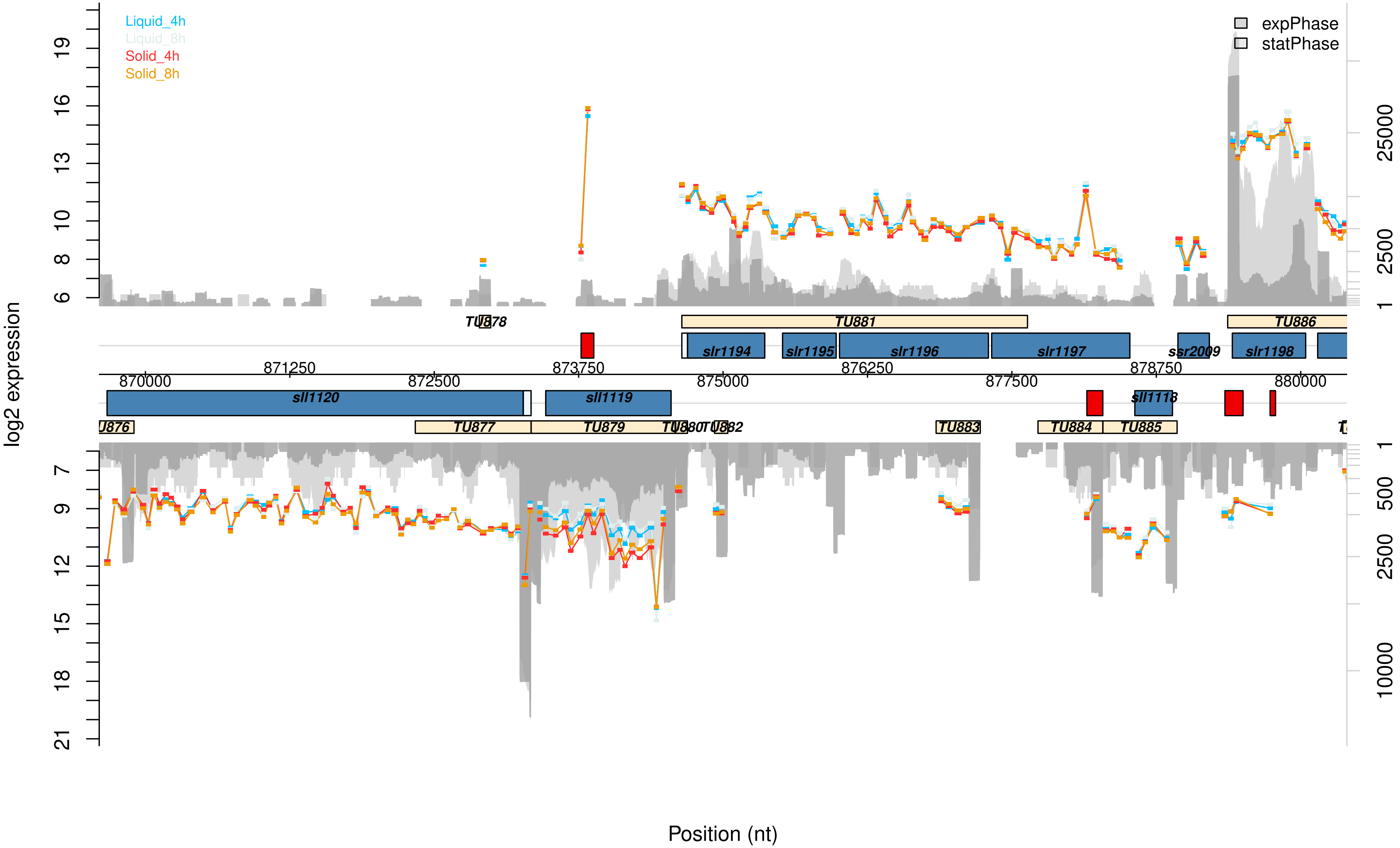
Liquid_4h
Liquid_8h
Solid_4h
Solid_8h
expPhase
statPhase
log2 expression
Position (nt)
6
8
10
13
16
19
7
9
12
15
18
21
1
2500
25000
1
500
2500
10000
870000
871250
872500
873750
875000
876250
877500
878750
880000
TU878
TU881
TU886
slr1194
slr1195
slr1196
slr1197
ssr2009
slr1198
sll1120
sll1119
sll1118
TU876
TU877
TU879
TU880
TU882
TU883
TU884
TU885

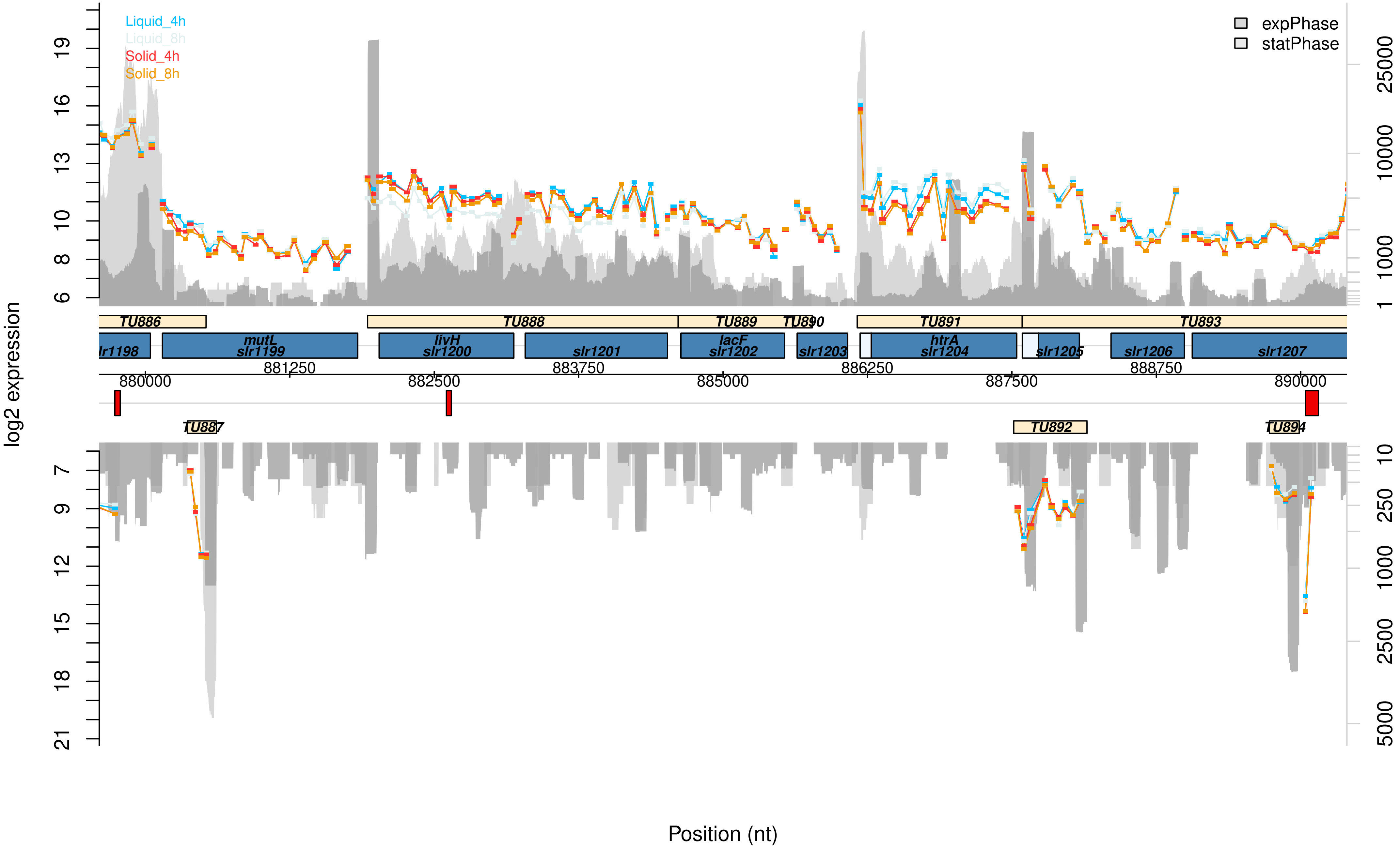

Liquid_4h
Liquid_8h
Solid_4h
Solid_8h
expPhase
statPhase
log2 expression
TU886
TU888
TU889
TU890
TU891
TU893
TU887
TU892
TU894
lr1198
mutL
slr1199
livH
slr1200
slr1201
lacF
slr1202
slr1203
htrA
slr1204
slr1205
slr1206
slr1207
880000
881250
882500
883750
885000
886250
887500
888750
890000
Position (nt)

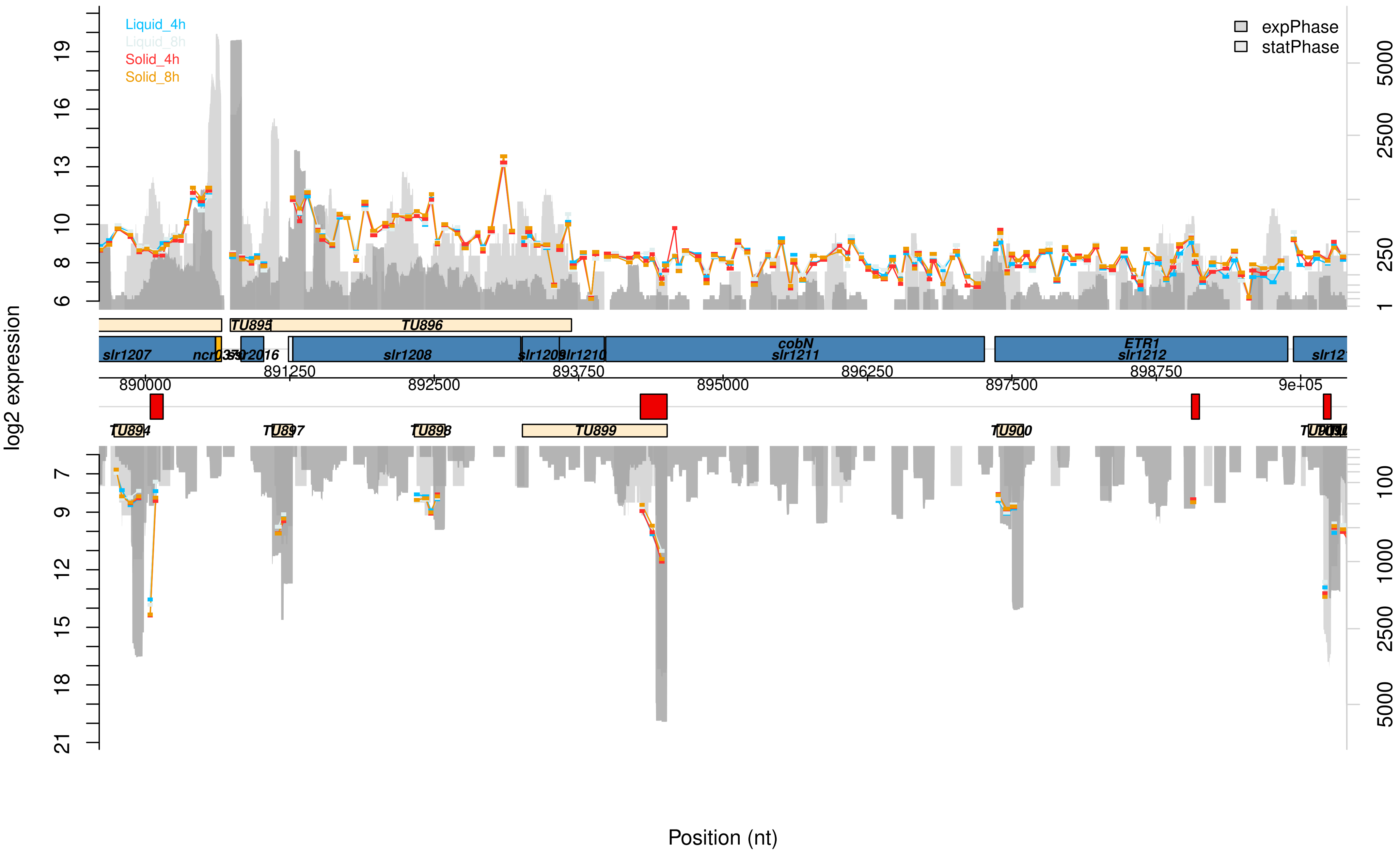

Liquid_4h
Liquid_8h
Solid_4h
Solid_8h
expPhase
statPhase
log2 expression
TU895
TU896
slr1207
slr1208
slr1210
cobN
slr1211
ETR1
slr1212
890000
891250
892500
893750
895000
896250
897500
898750
9e+05
TU894
TU897
TU898
TU899
TU900
Position (nt)

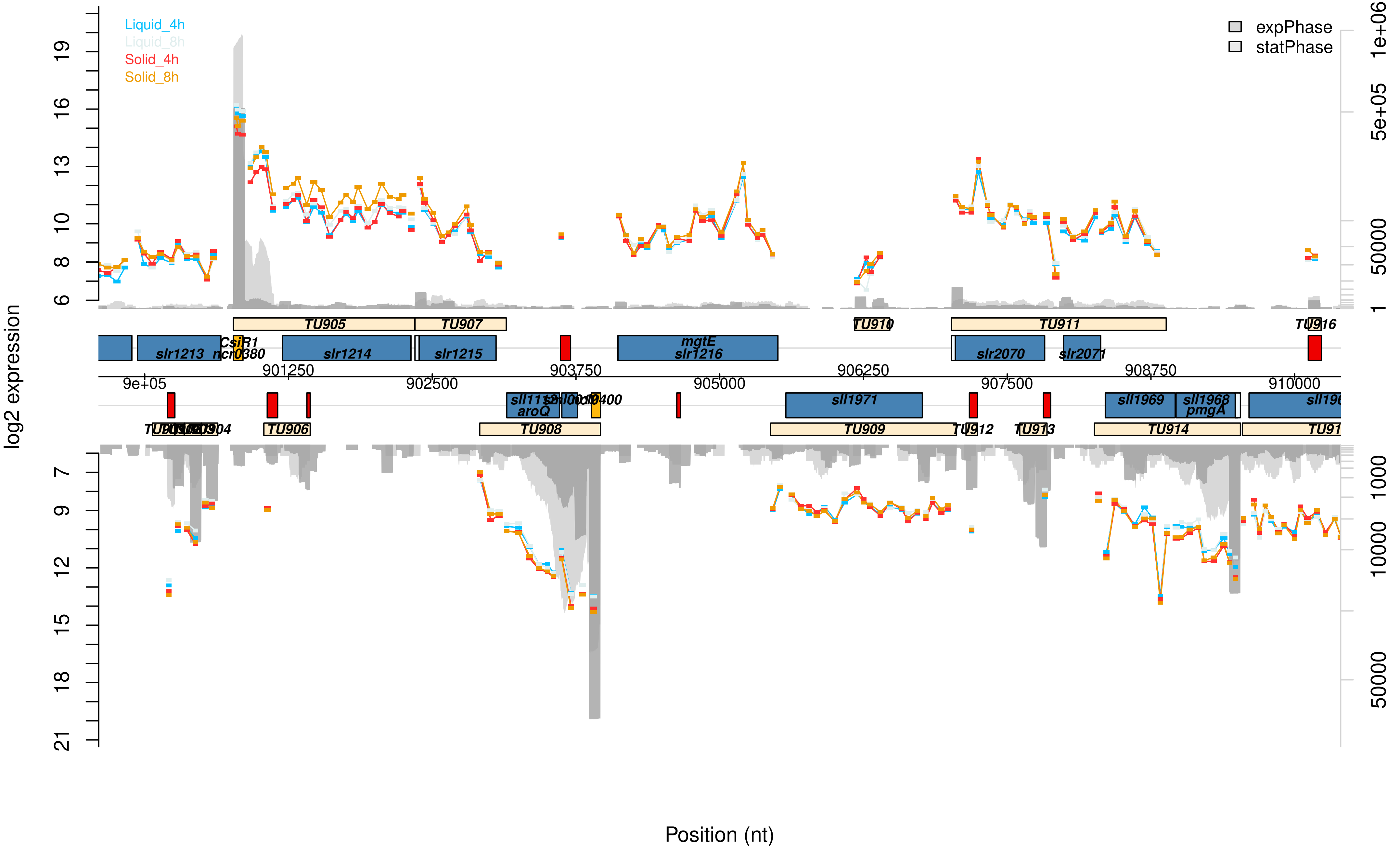

Liquid_4h
Liquid_8h
Solid_4h
Solid_8h
expPhase
statPhase
log2 expression
Position (nt)
TU905
TU907
TU910
TU911
TU916
slr1213
CsiR1
ncl0380
slr1214
slr1215
mgtE
slr1216
slr2070
slr2071
9e+05
901250
902500
903750
905000
906250
907500
908750
910000
aroQ
sll1971
sll1969
sll1968
pmgA
TU906
TU908
TU909
TU913
TU914

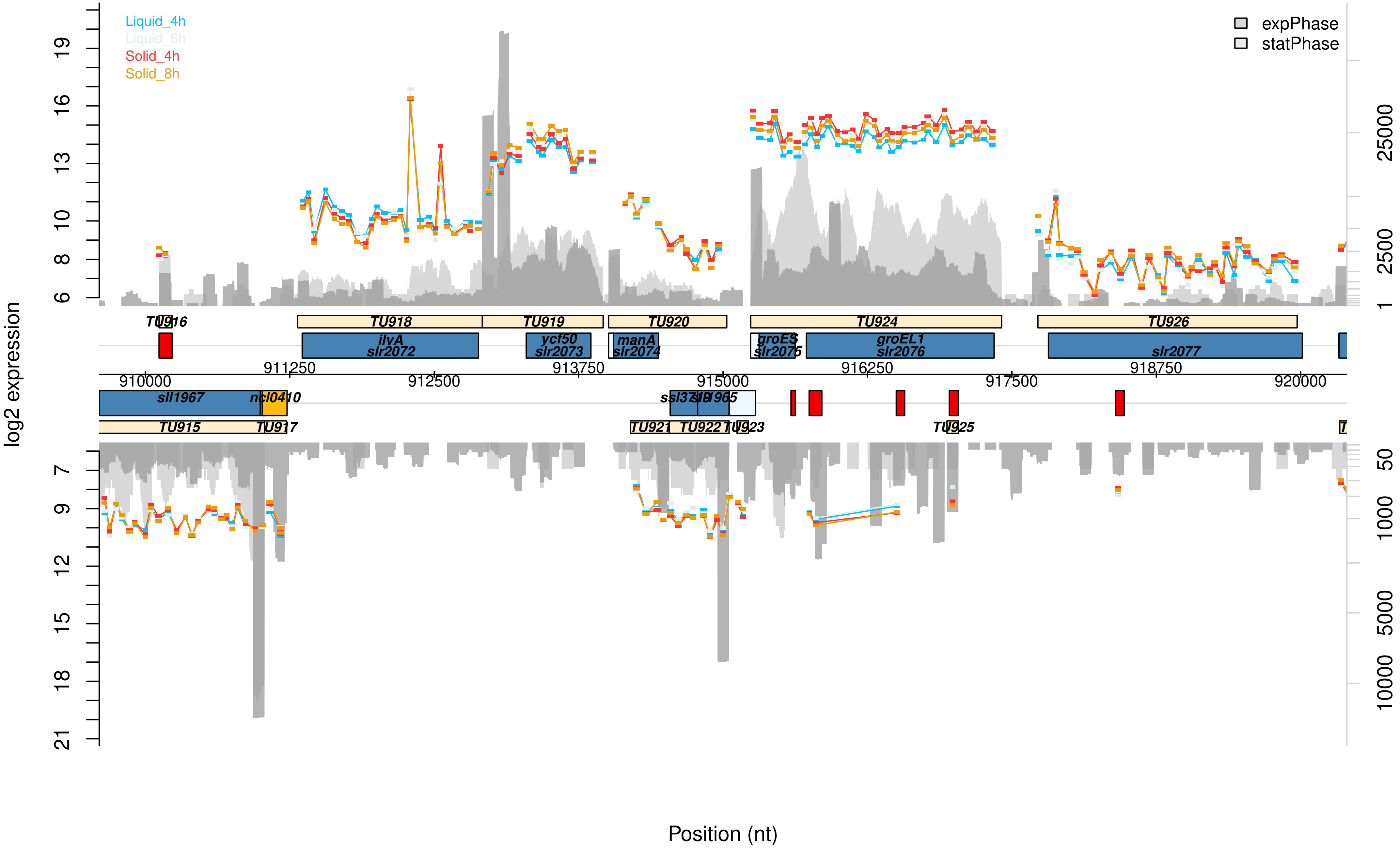

Liquid_4h
Liquid_8h
Solid_4h
Solid_8h
expPhase
statPhase
log2 expression
Position (nt)
TU916
TU918
TU919
TU920
TU924
TU926
ilvA
slr2072
ycf50
slr2073
manA
slr2074
groES
slr2075
groEL1
slr2076
slr2077
910000
911250
912500
913750
915000
916250
917500
918750
920000
sll1967
ncl0410
TU915
TU917
TU921
TU922
TU923
TU925

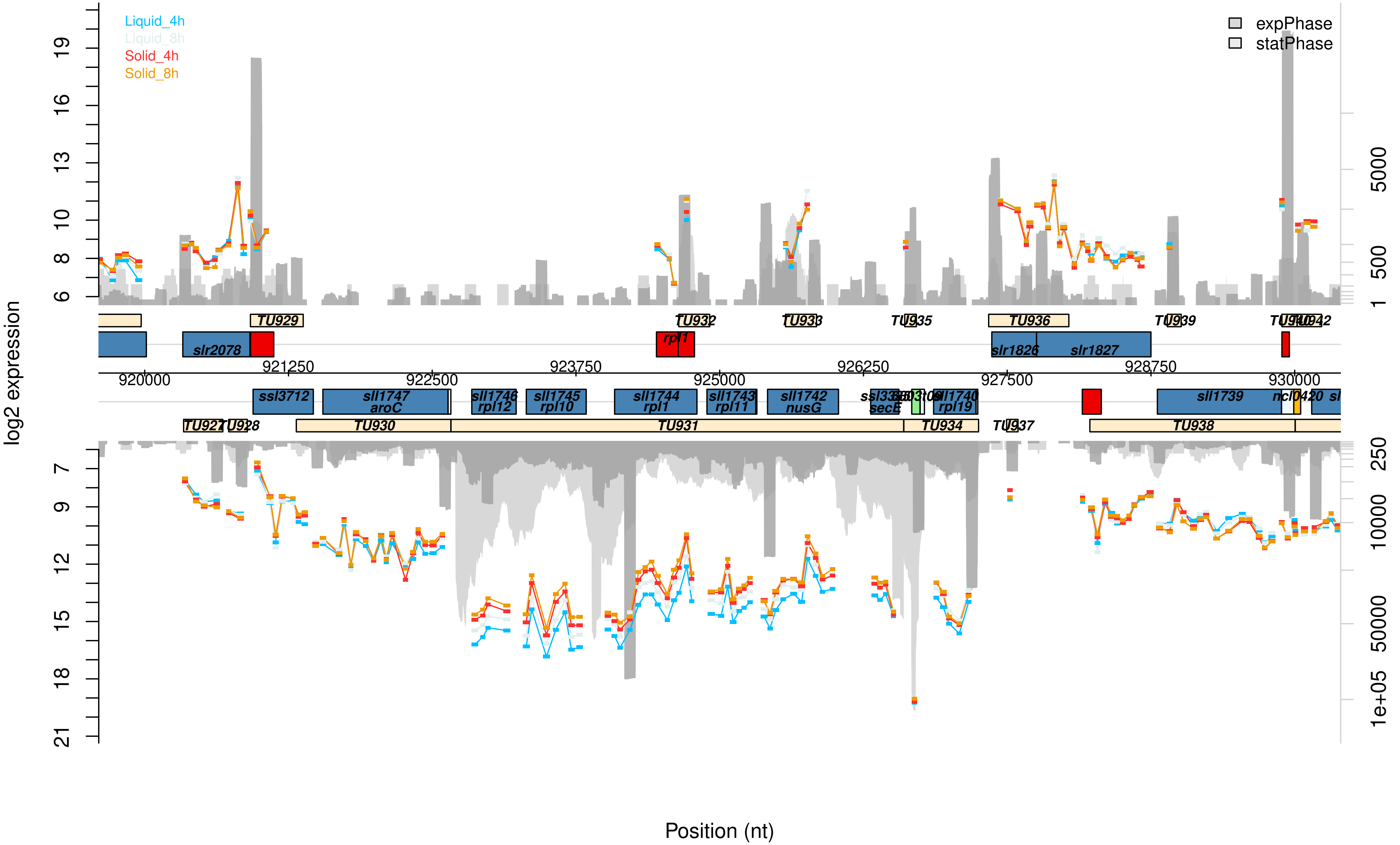

Liquid_4h
Liquid_8h
Solid_4h
Solid_8h
expPhase
statPhase
log2 expression
Position (nt)
TU929
TU932
TU933
TU935
TU936
TU939
TU942
slr2078
rpl1
slr1826
slr1827
920000
921250
922500
923750
925000
926250
927500
928750
930000
ssl3712
sll1747
aroC
sll1746
rpl12
sll1745
rpl10
sll1744
rpl1
sll1743
rpl11
sll1742
nusG
secE
sll1740
rpl19
sll1739
ncl0420
TU927
TU928
TU930
TU931
TU934
TU937
TU938

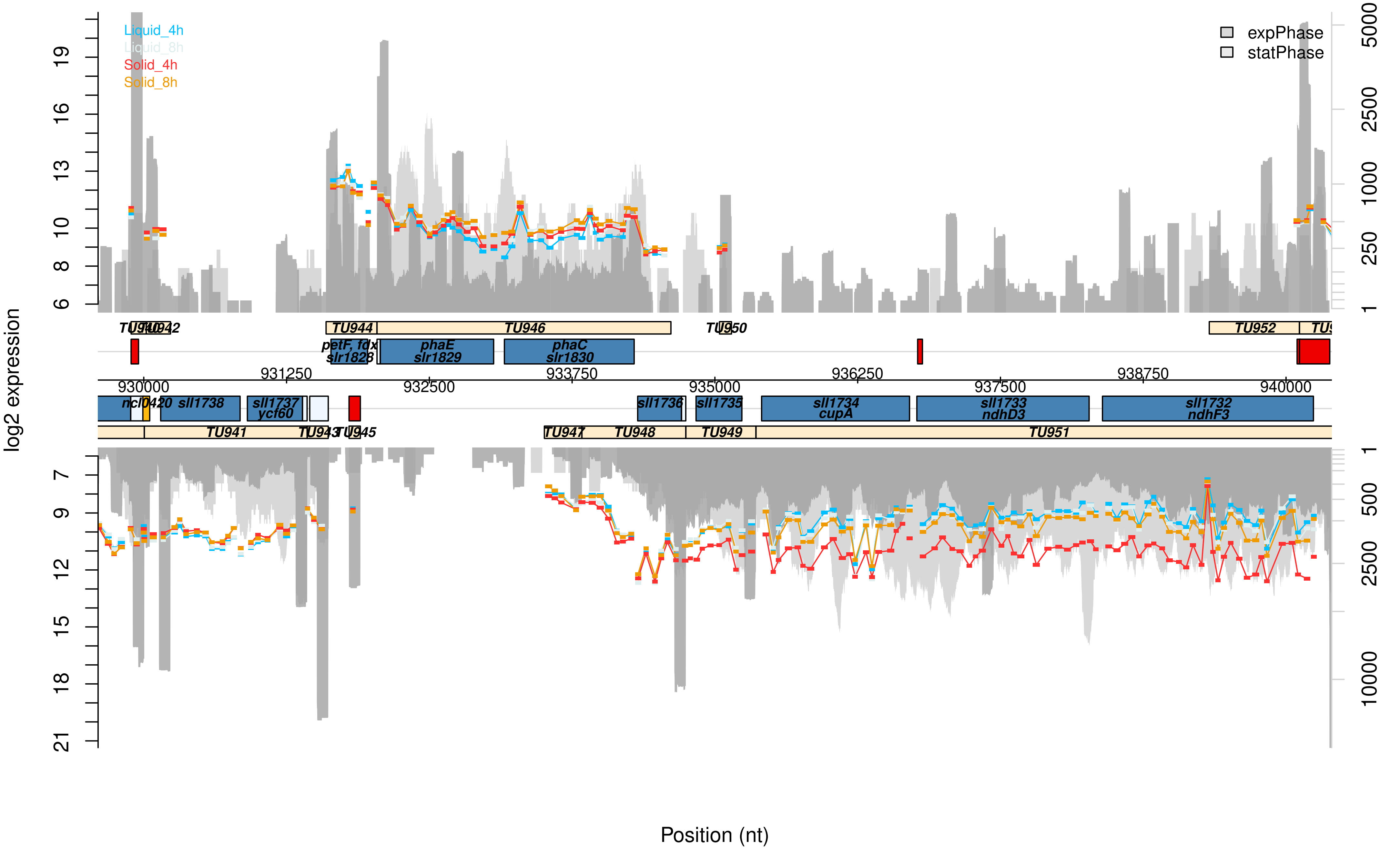

Liquid_4h
Liquid_8h
Solid_4h
Solid_8h
expPhase
statPhase
log2 expression
Position (nt)
TU944
TU946
TU950
TU952
petF, fdx
slr1828
phaE
slr1829
phaC
slr1830
930000
931250
932500
933750
935000
936250
937500
938750
940000
ncl0420
sll1738
sll1737
ycf60
sll1736
sll1735
sll1734
cupA
sll1733
ndhD3
sll1732
ndhF3
TU941
TU947
TU948
TU949
TU951

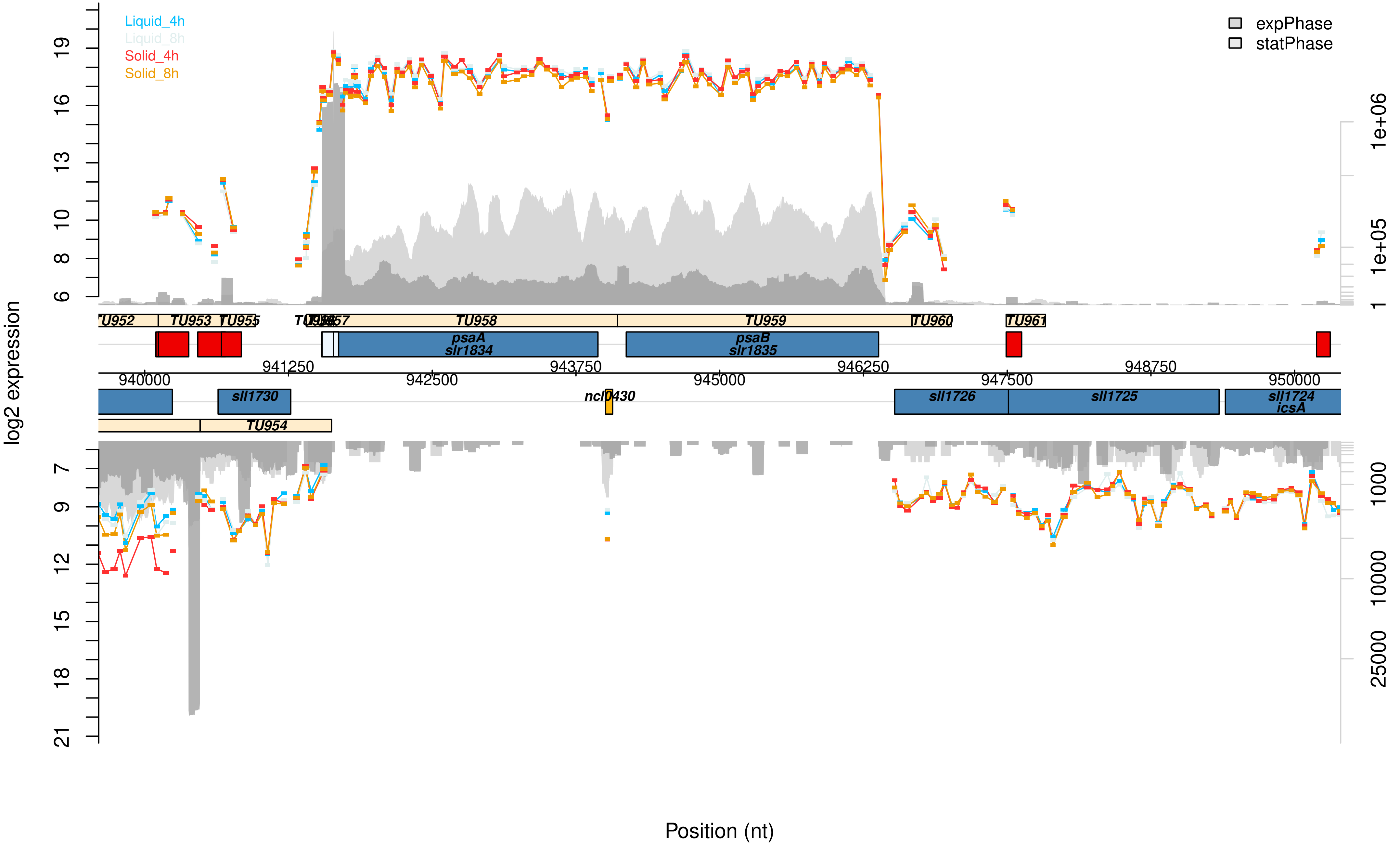

Liquid_4h
Liquid_8h
Solid_4h
Solid_8h
expPhase
statPhase
log2 expression
Position (nt)
TU952
TU953
TU955
TU958
TU959
TU960
TU961
TU954
psaA
slr1834
psaB
slr1835
sll1730
ncl0430
sll1726
sll1725
sll1724
icsA
940000
941250
942500
943750
945000
946250
947500
948750
950000
1e+06
1e+05
1
1000
10000
25000

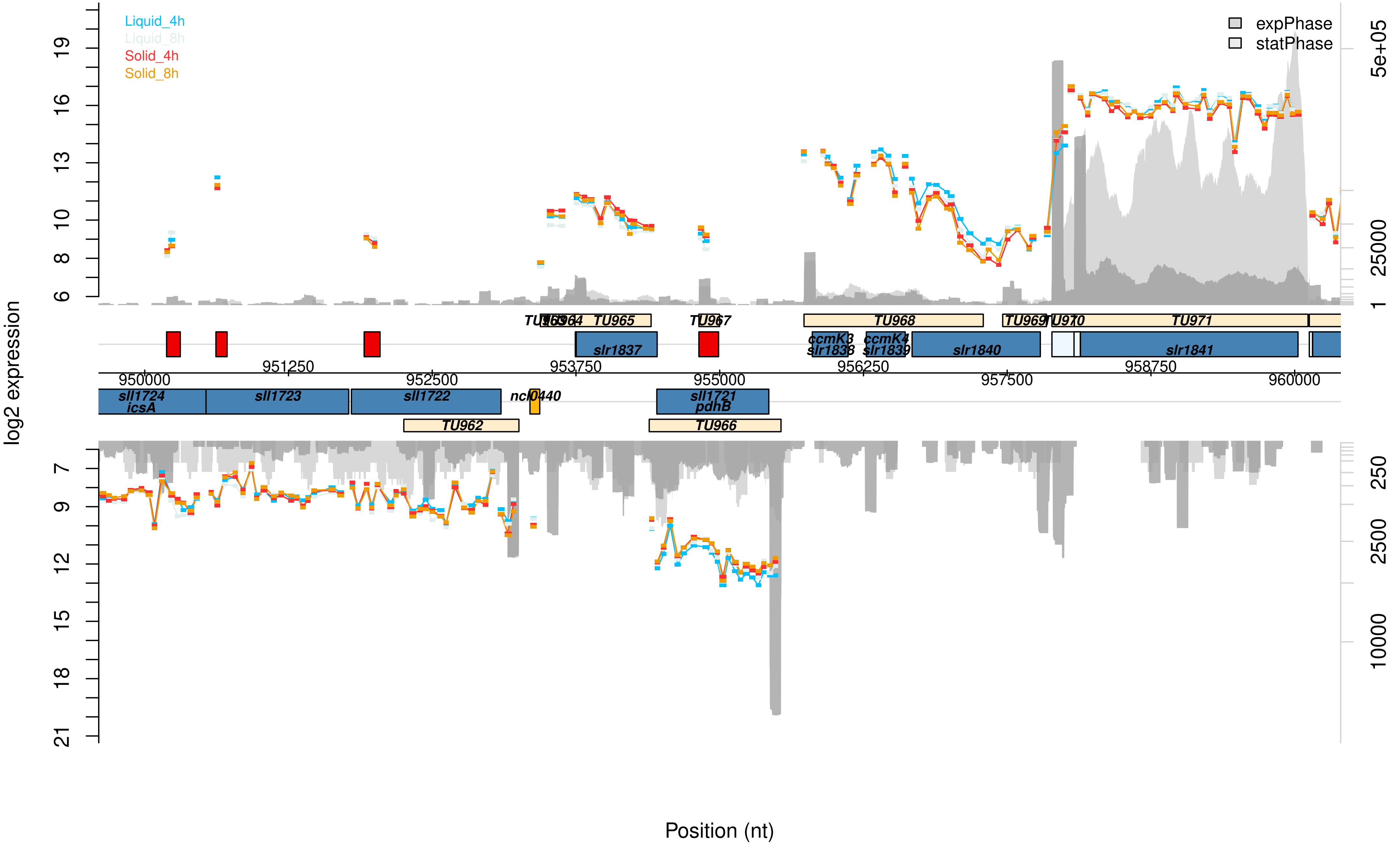

Liquid_4h
Liquid_8h
Solid_4h
Solid_8h
expPhase
statPhase
log2 expression
Position (nt)
TU963 TU964 TU965 TU967 TU968 TU969 TU970 TU971
slr1837 ccmK3 slr1838 ccmK4 slr1839 slr1840 slr1841
950000 951250 952500 953750 955000 956250 957500 958750 960000
sll1724 icsA sll1723 sll1722 ncl0440 sll1721 pdhB
TU962 TU966

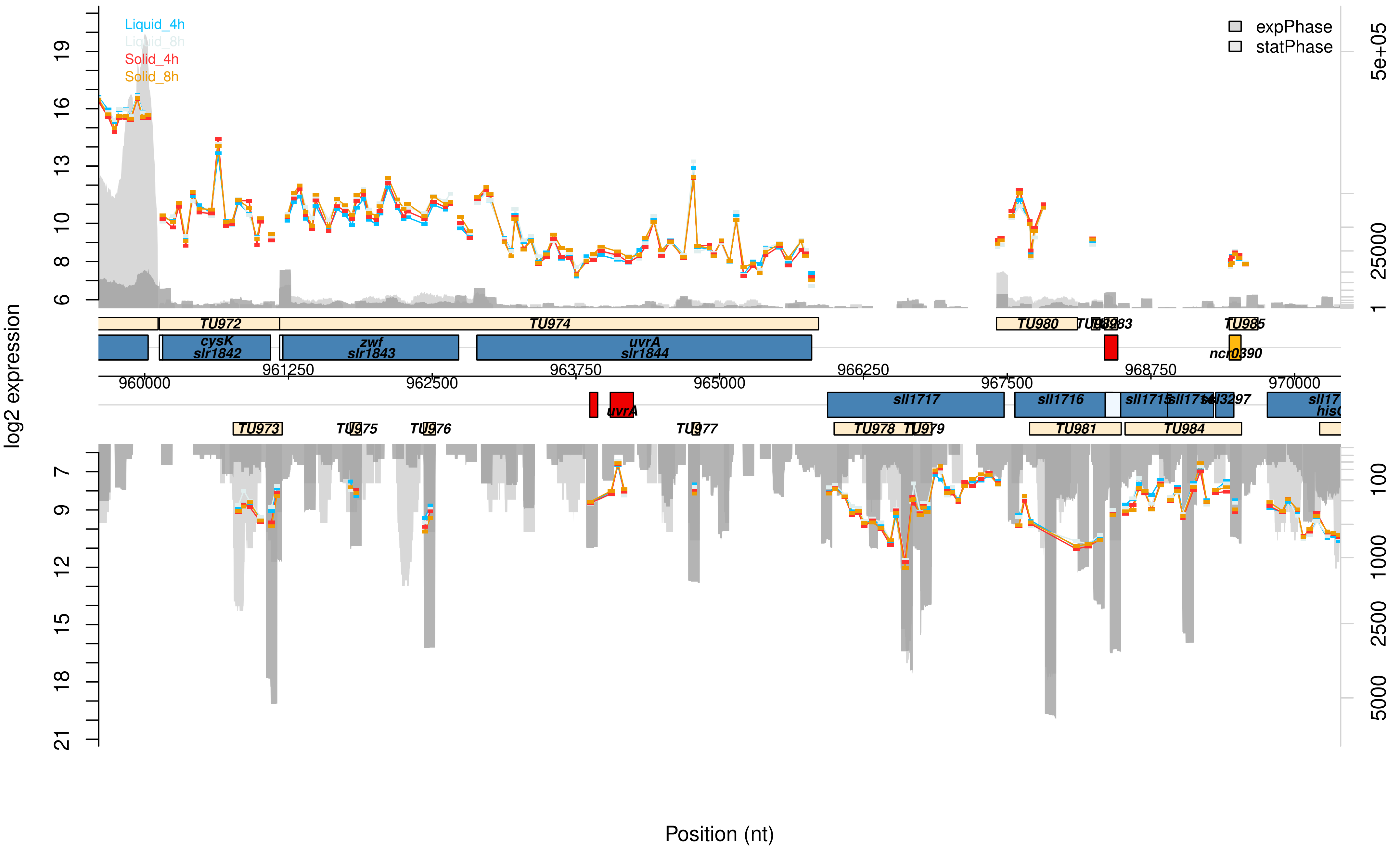

Liquid_4h
Liquid_8h
Solid_4h
Solid_8h
expPhase
statPhase
log2 expression
19
16
13
10
8
6
7
9
12
15
18
21
5e+05
25000
1
100
1000
2500
5000
TU972
TU974
TU980
TU985
cysK
slr1842
zwf
slr1843
uvrA
slr1844
ncr0390
960000
961250
962500
963750
965000
966250
967500
968750
970000
uvrA
sll1717
sll1716
TU973
TU975
TU976
TU977
TU978
TU979
TU981
TU984
Position (nt)

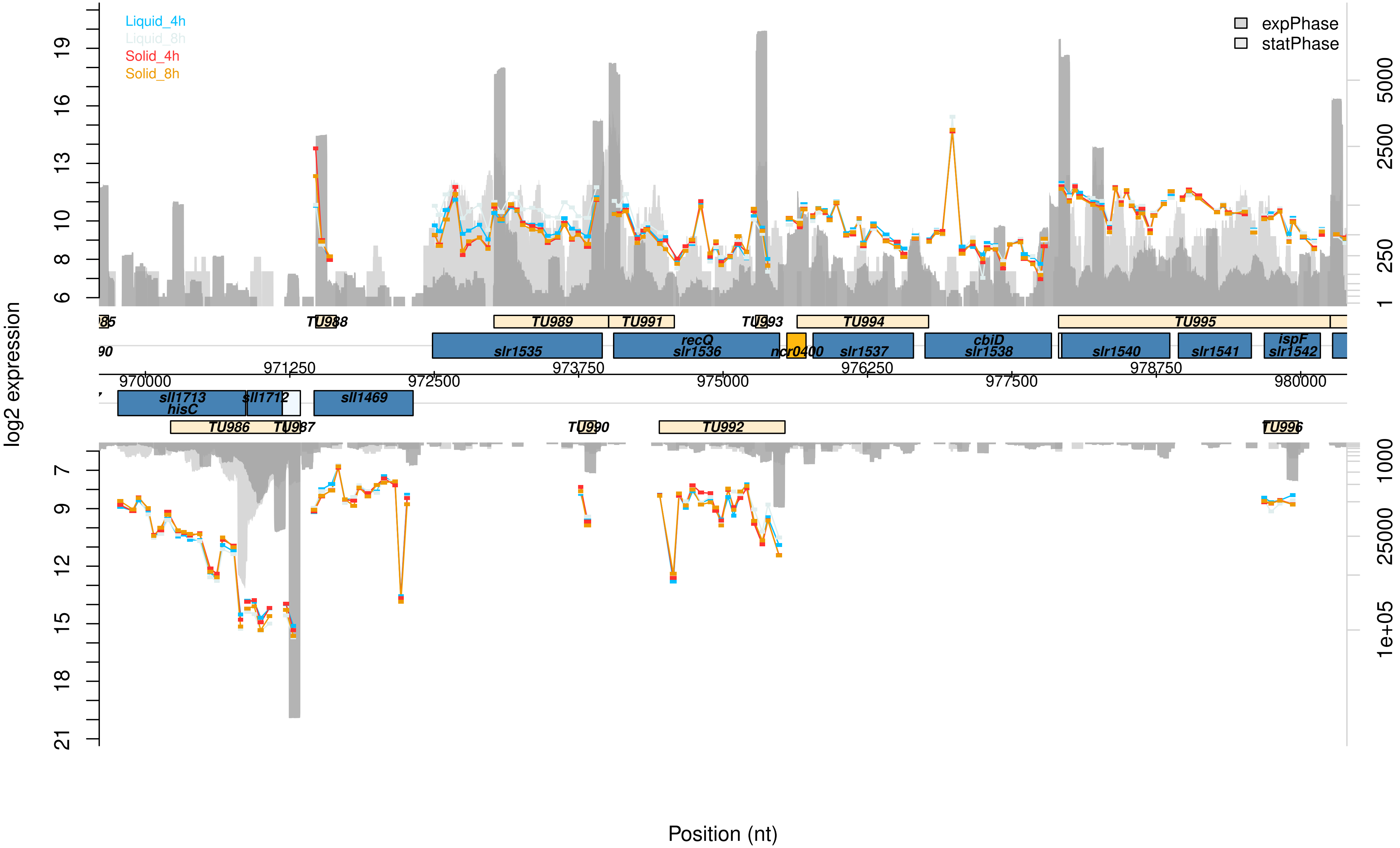

Liquid_4h
Liquid_8h
Solid_4h
Solid_8h
expPhase
statPhase
log2 expression
Position (nt)
TU988
TU989
TU991
TU993
TU994
TU995
slr1535
recQ
slr1536
ncr0400
slr1537
cbiD
slr1538
slr1540
slr1541
ispF
slr1542
970000
971250
972500
973750
975000
976250
977500
978750
980000
sll1713
hisC
sll1712
sll1469
TU986
TU987
TU990
TU992
TU996

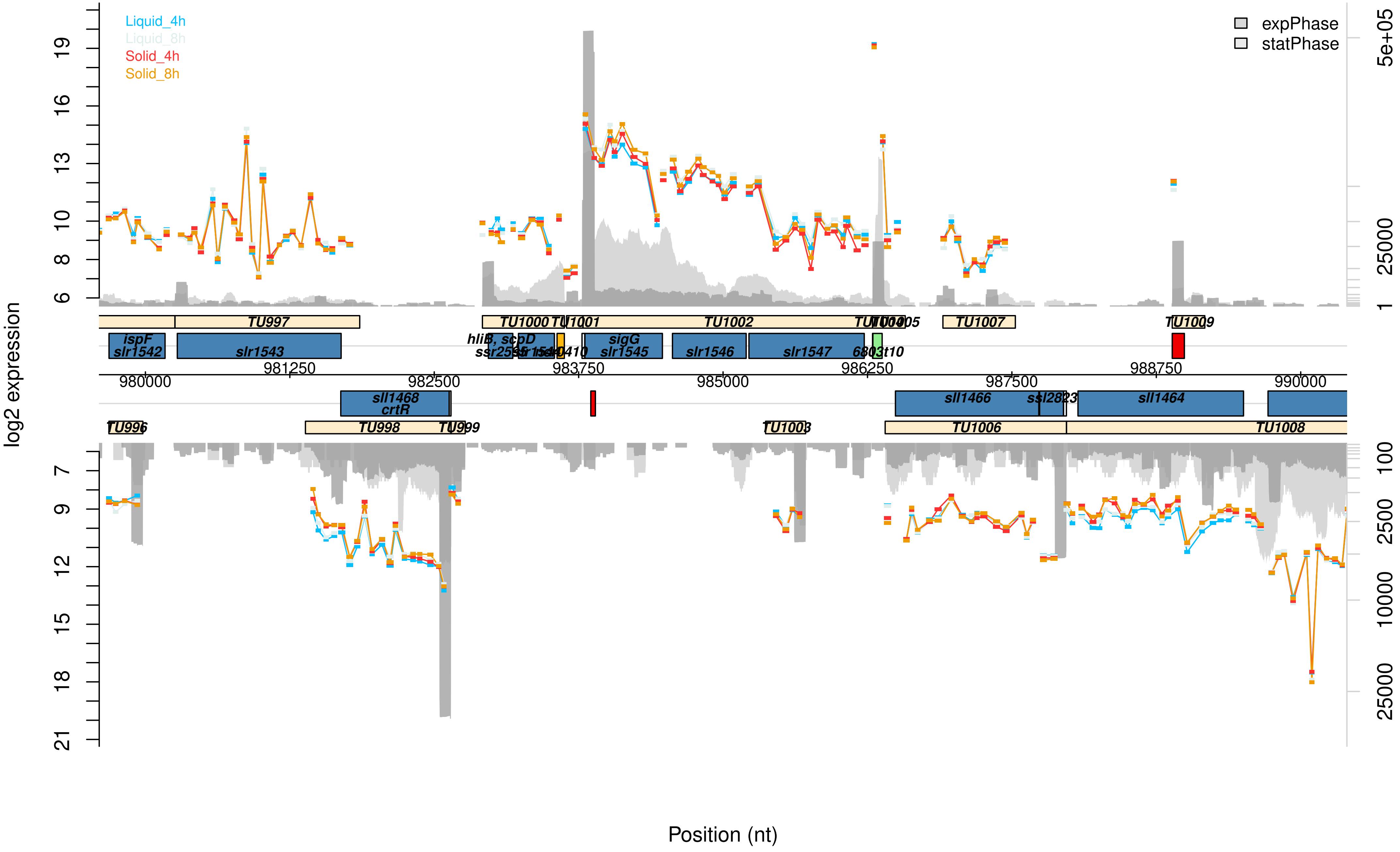

Liquid_4h
Liquid_8h
Solid_4h
Solid_8h
expPhase
statPhase
log2 expression
Position (nt)
TU997
TU1000
TU1001
TU1002
TU1004
TU1005
TU1007
TU1009
ispF
slr1542
slr1543
hliB, scpD
ssr2595
slr1544
sigG
slr1545
slr1546
slr1547
6803t10
980000
981250
982500
983750
985000
986250
987500
988750
990000
sll1468
crtR
sll1466
ssl2823
sll1464
TU996
TU998
TU999
TU1003
TU1006
TU1008

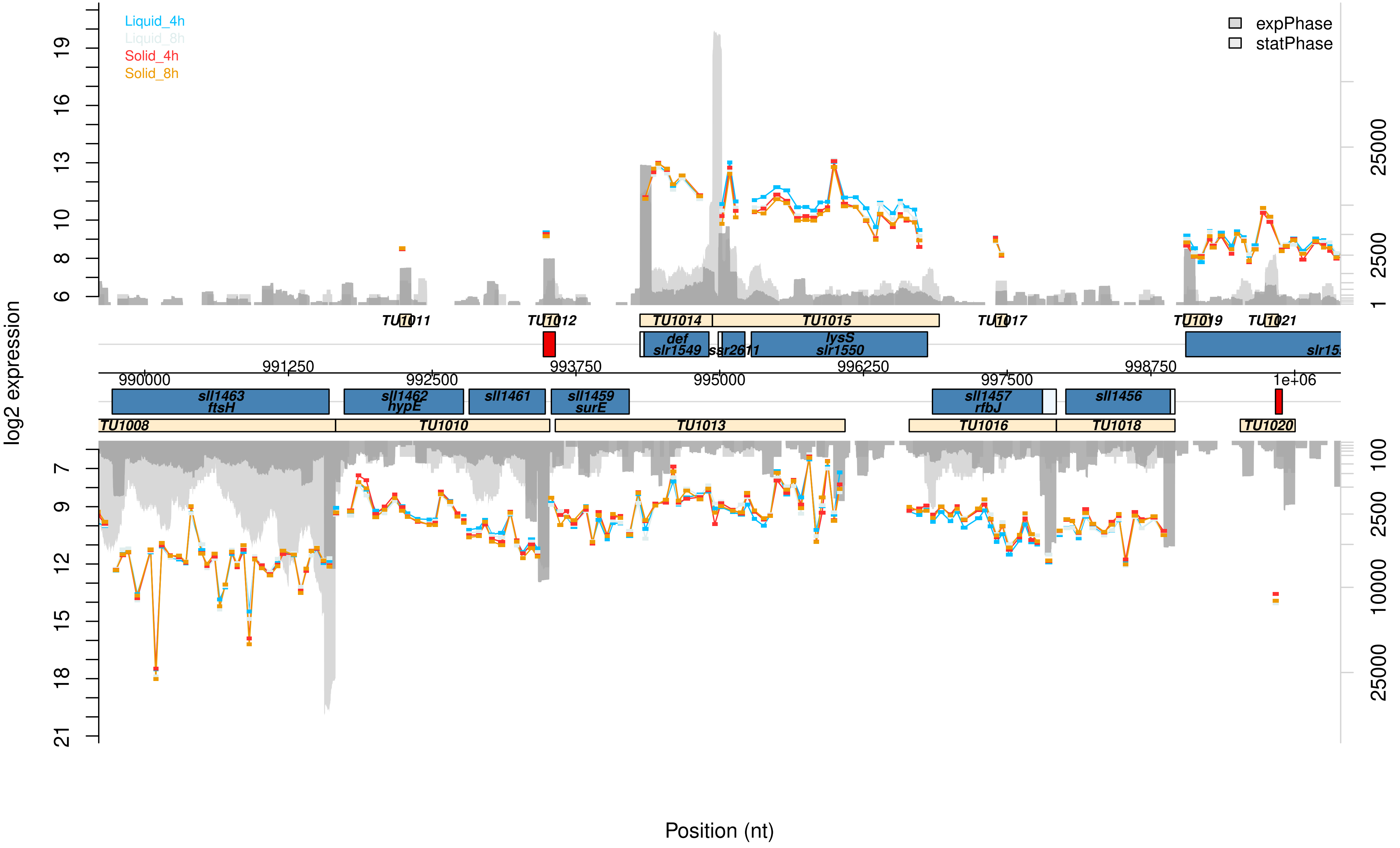
Liquid_4h
Liquid_8h
Solid_4h
Solid_8h
expPhase
statPhase
log2 expression
Position (nt)
990000
991250
992500
993750
995000
996250
997500
998750
1e+06
TU1008
TU1010
TU1011
TU1012
TU1013
TU1014
TU1015
TU1016
TU1017
TU1018
TU1019
TU1020
TU1021
sll1463
ftsH
sll1462
hypE
sll1461
sll1459
surE
def
slr1549
ssr2611
lysS
slr1550
sll1457
rfbJ
sll1456
6
8
10
13
16
19
7
9
12
15
18
21
1
2500
25000
100
10000

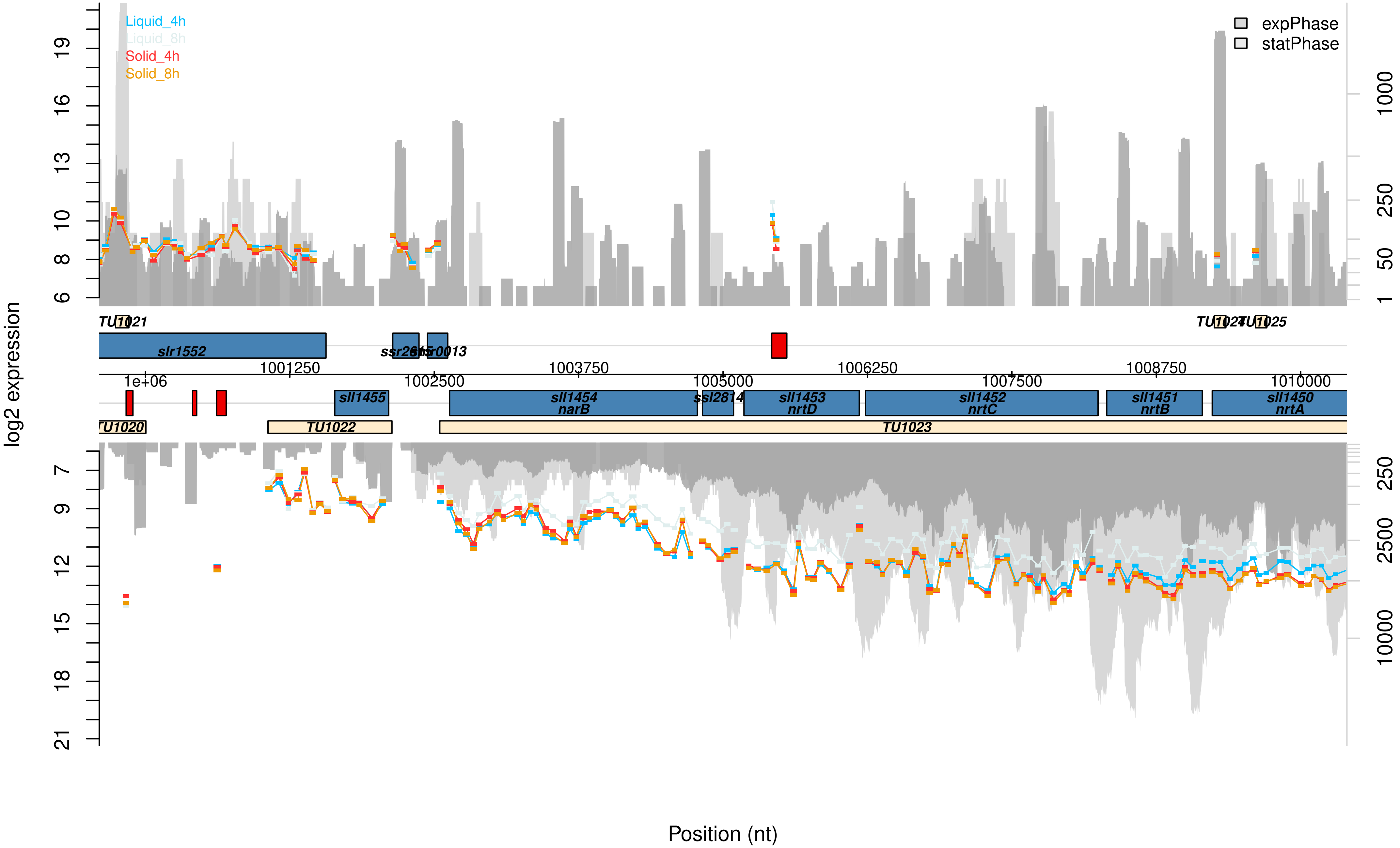

Liquid_4h
Liquid_8h
Solid_4h
Solid_8h
expPhase
statPhase
log2 expression
Position (nt)
TU1021
TU1020
TU1022
TU1023
TU1024
TU1025
slr1552
ssr2843
smr0013
sll1455
sll1454 narB
ssl2814
sll1453 nrtD
sll1452 nrtC
sll1451 nrtB
sll1450 nrtA
1e+06
1001250
1002500
1003750
1005000
1006250
1007500
1008750
1010000

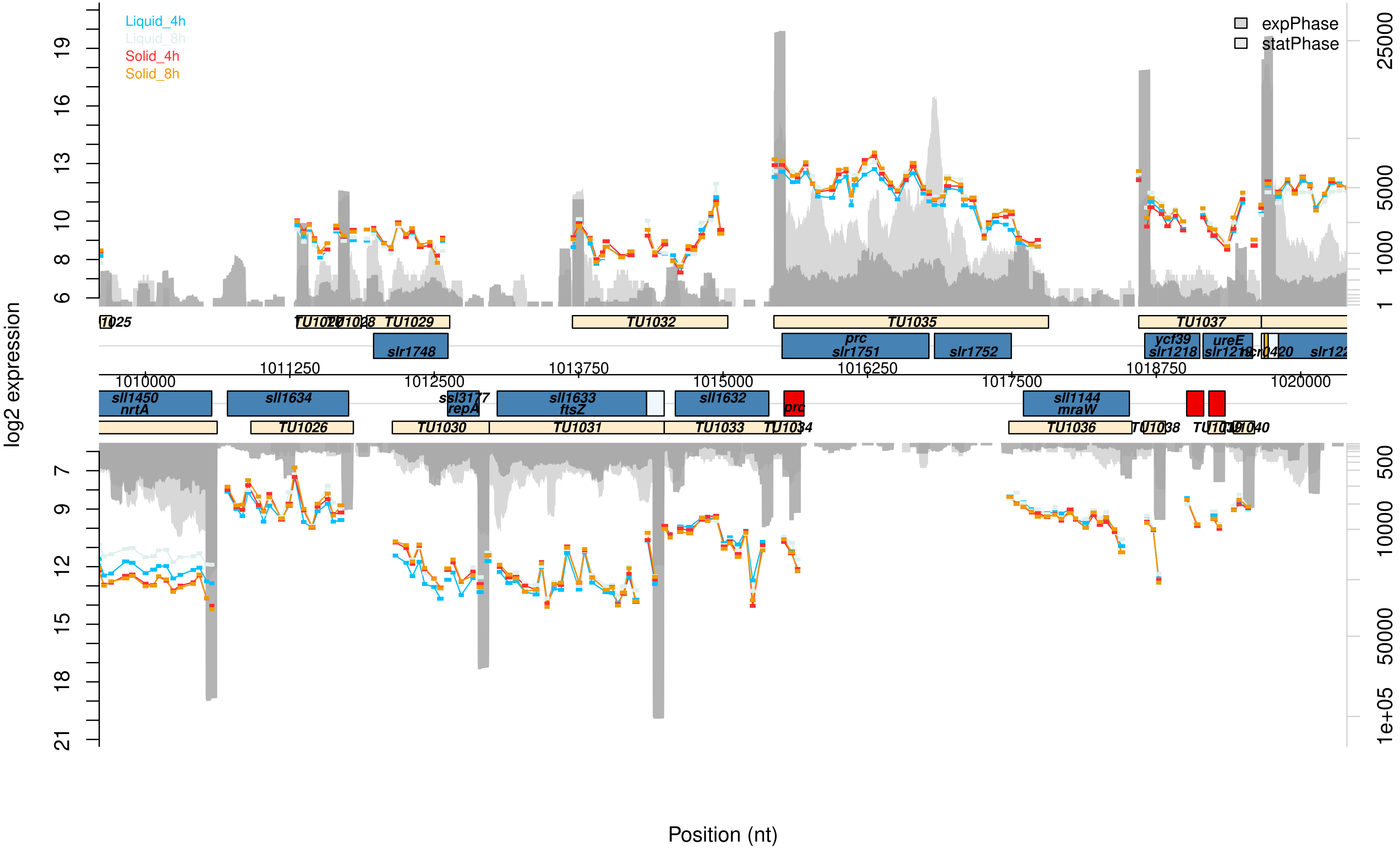

Liquid_4h
Liquid_8h
Solid_4h
Solid_8h
expPhase
statPhase
log2 expression
Position (nt)
TU1026
TU1029
TU1032
TU1035
TU1037
slr1748
prc
slr1751
slr1752
ycf39
slr1218
ureE
slr1219
slr122
1010000
1011250
1012500
1013750
1015000
1016250
1017500
1018750
1020000
sll1450
nrtA
sll1634
ssl3177
repA
sll1633
ftsZ
sll1632
prc
sll1144
mraW
TU1030
TU1031
TU1033
TU1034
TU1036

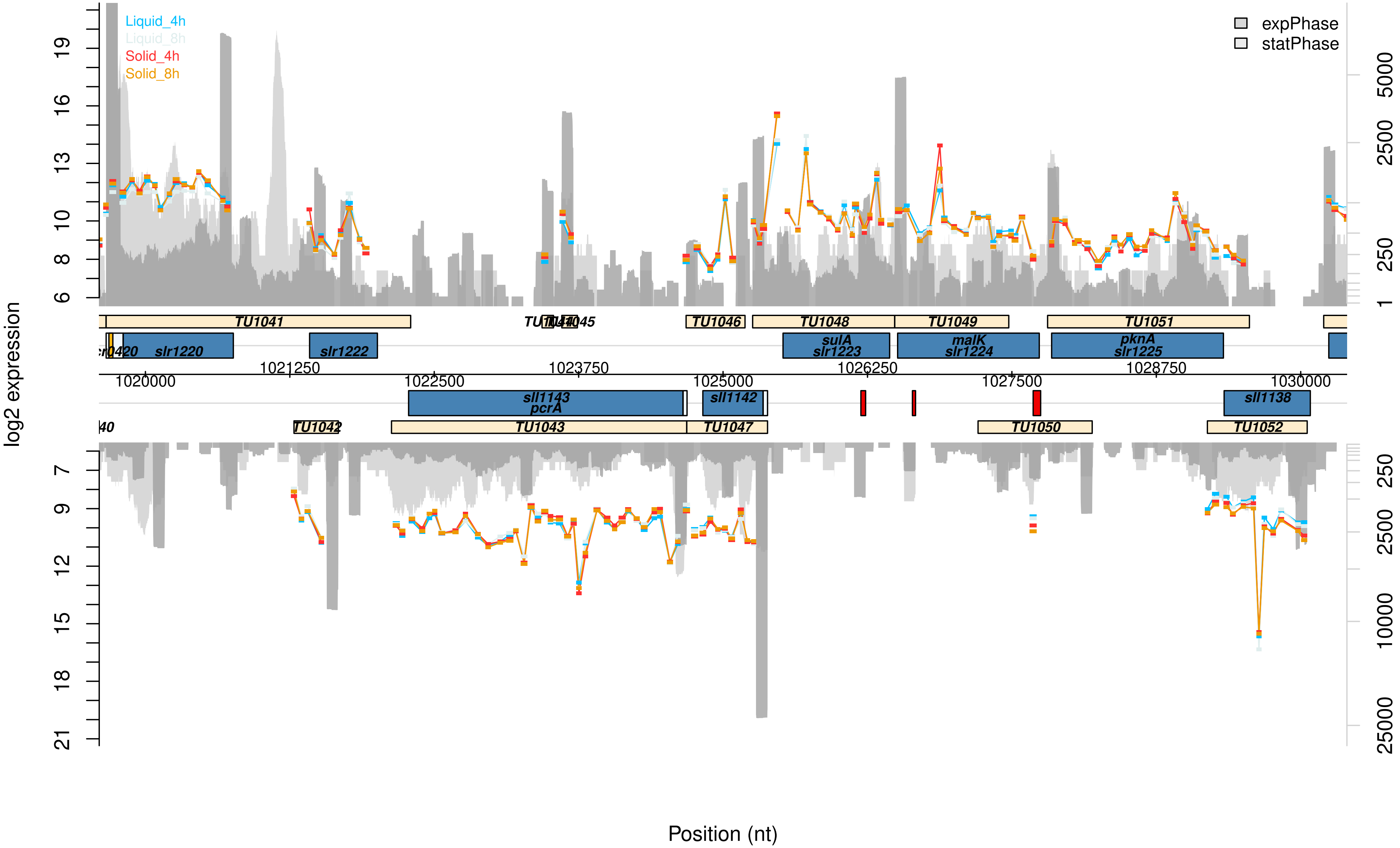

Liquid_4h
Liquid_8h
Solid_4h
Solid_8h
expPhase
statPhase
log2 expression
Position (nt)
TU1041
TU1045
TU1046
TU1048
TU1049
TU1051
slr1220
slr1222
sulA
slr1223
malK
slr1224
pknA
slr1225
1020000
1021250
1022500
1023750
1025000
1026250
1027500
1028750
1030000
sll1143
pcrA
sll1142
sll1138
TU1042
TU1043
TU1047
TU1050
TU1052

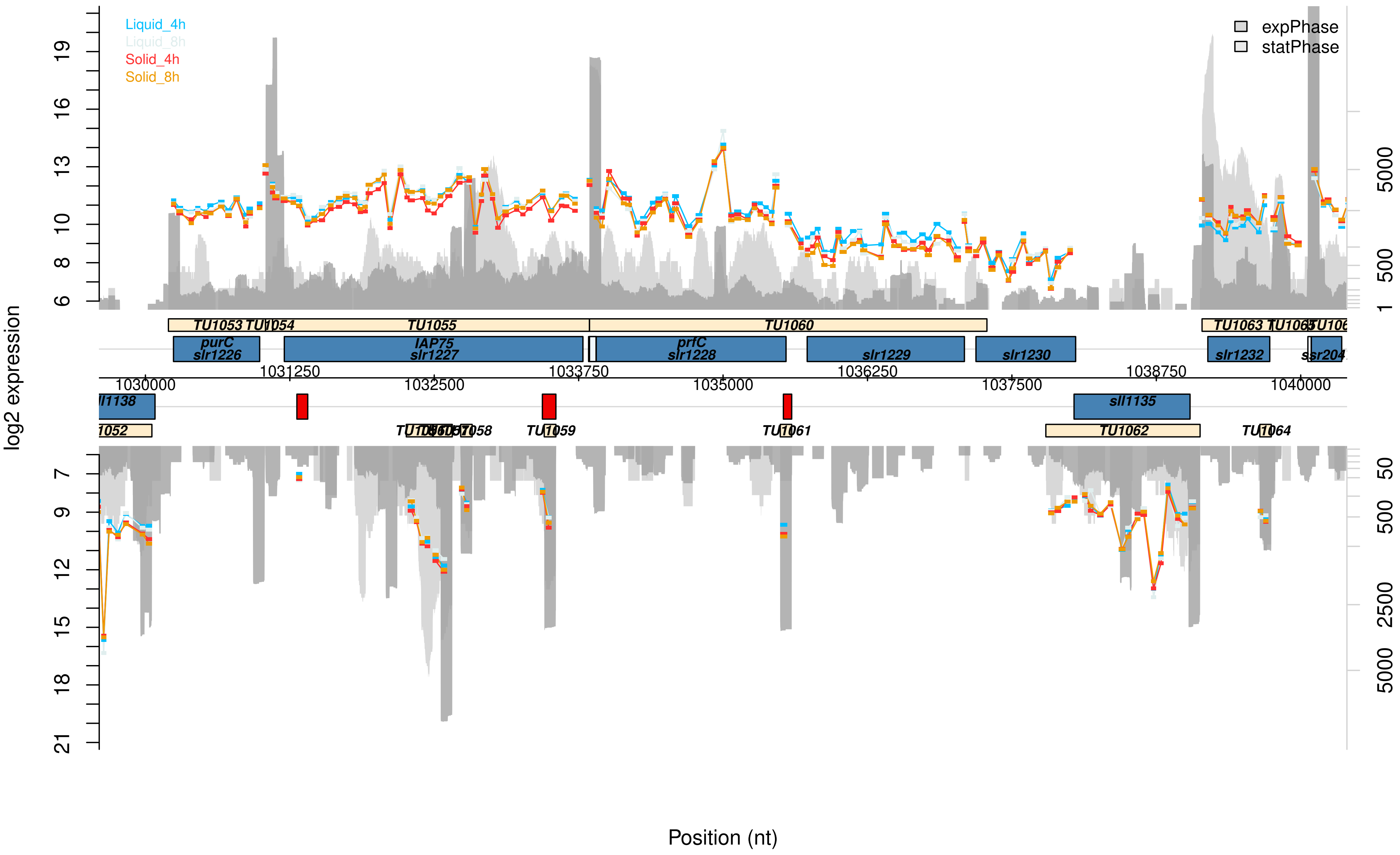

Liquid_4h
Liquid_8h
Solid_4h
Solid_8h
expPhase
statPhase
log2 expression
Position (nt)
TU1053
TU1054
TU1055
TU1060
TU1063
purC
slr1226
IAP75
slr1227
prfC
slr1228
slr1229
slr1230
slr1232
1031250
1033750
1036250
1038750
1030000
1032500
1035000
1037500
1040000
sll1135
TU1059
TU1061
TU1062
TU1064

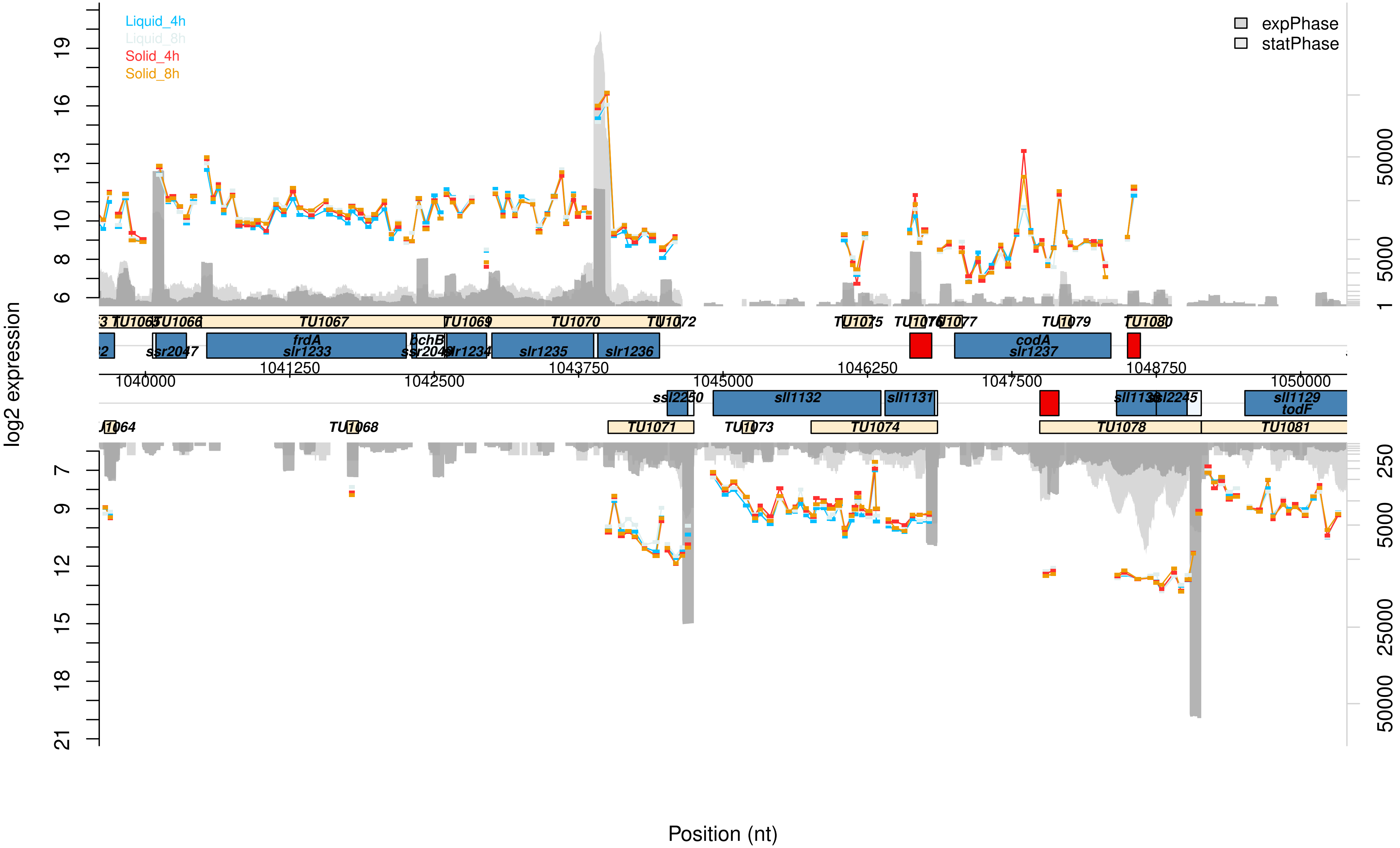

Liquid_4h
Liquid_8h
Solid_4h
Solid_8h
expPhase
statPhase
log2 expression
Position (nt)
TU1065
TU1066
TU1067
TU1069
TU1070
TU1072
TU1075
TU1077
TU1079
TU1080
ssr2047
frdA
slr1233
pchB
slr1234
slr1235
slr1236
codA
slr1237
1040000
1041250
1042500
1043750
1045000
1046250
1047500
1048750
1050000
sll1132
sll1131
sll1129
todF
ssl2245
TU1068
TU1071
TU1073
TU1074
TU1078
TU1081

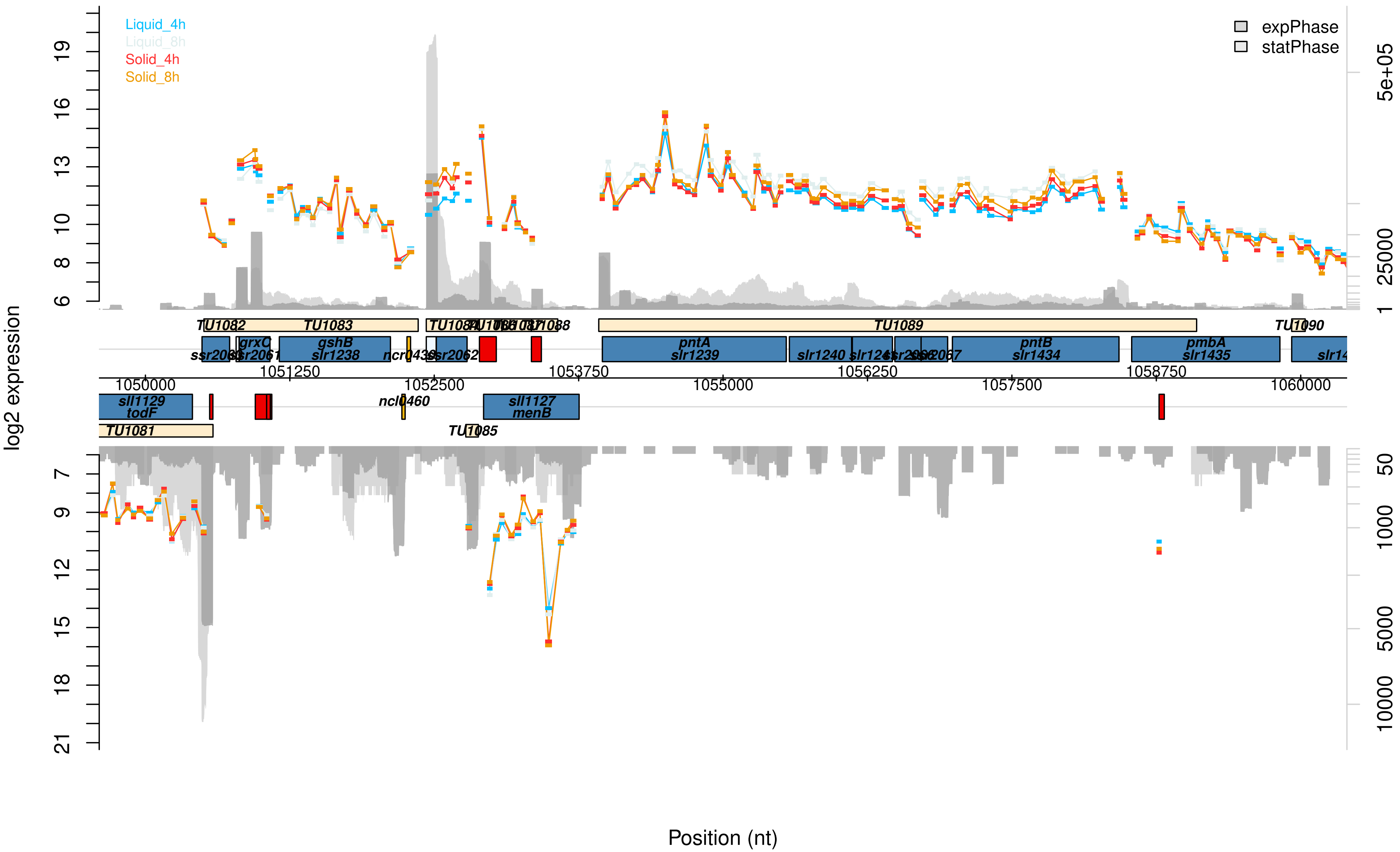
Liquid_4h
Liquid_8h
Solid_4h
Solid_8h
expPhase
statPhase
log2 expression
Position (nt)
TU1081
TU1082
TU1083
TU1085
TU1089
TU1090
sll1129
todF
grxC
gshB
slr1238
ncl0460
sll1127
menB
pntA
slr1239
slr1240
pntB
slr1434
pmbA
slr1435
1050000
1051250
1052500
1053750
1055000
1056250
1057500
1058750
1060000

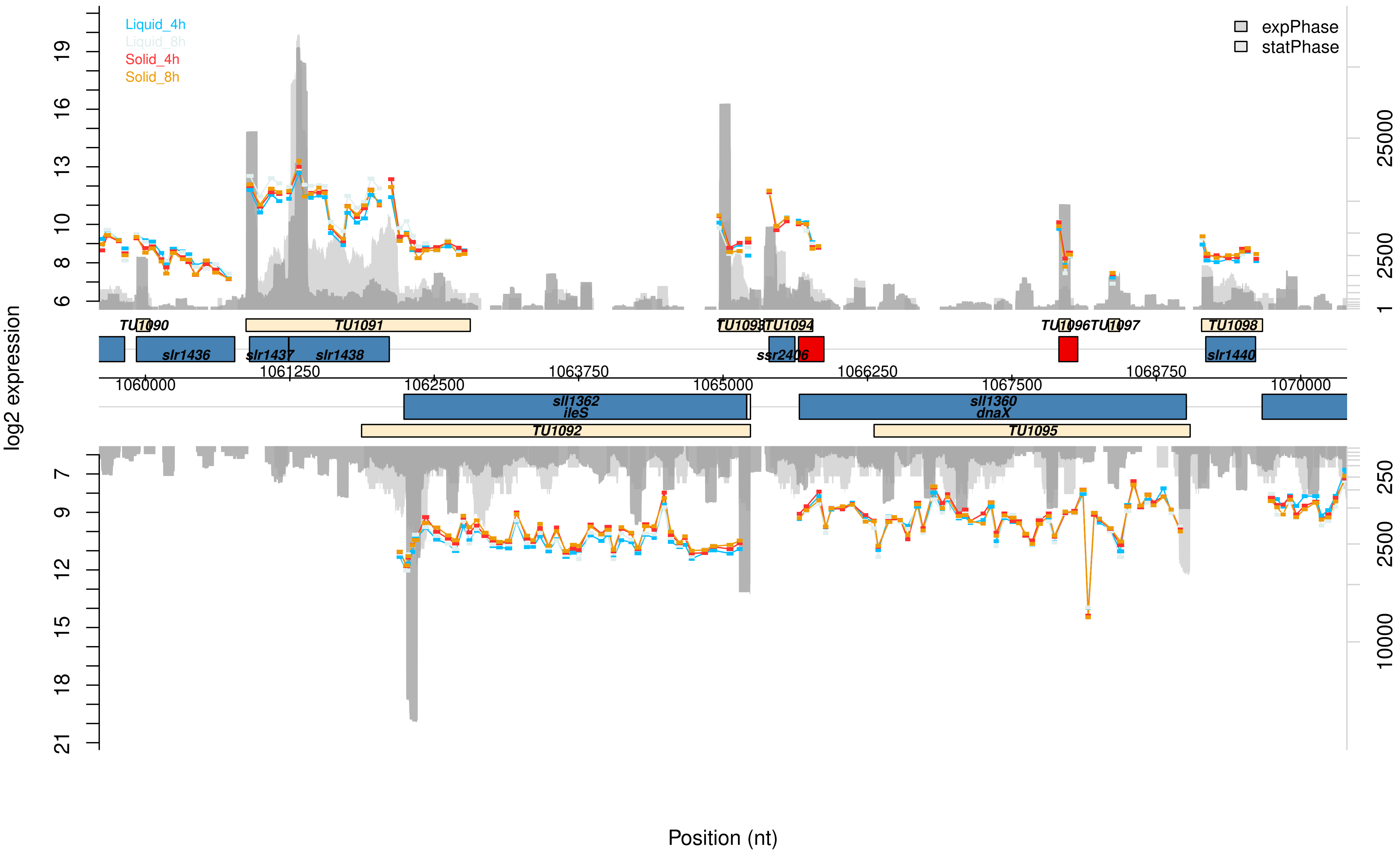

Liquid_4h
Liquid_8h
Solid_4h
Solid_8h
expPhase
statPhase
log2 expression
Position (nt)
TU1090
TU1091
TU1093
TU1094
TU1096
TU1097
TU1098
slr1436
slr1437
slr1438
ssr2406
slr1440
sll1362
ileS
sll1360
dnaX
TU1092
TU1095
1060000
1061250
1062500
1063750
1065000
1066250
1067500
1068750
1070000

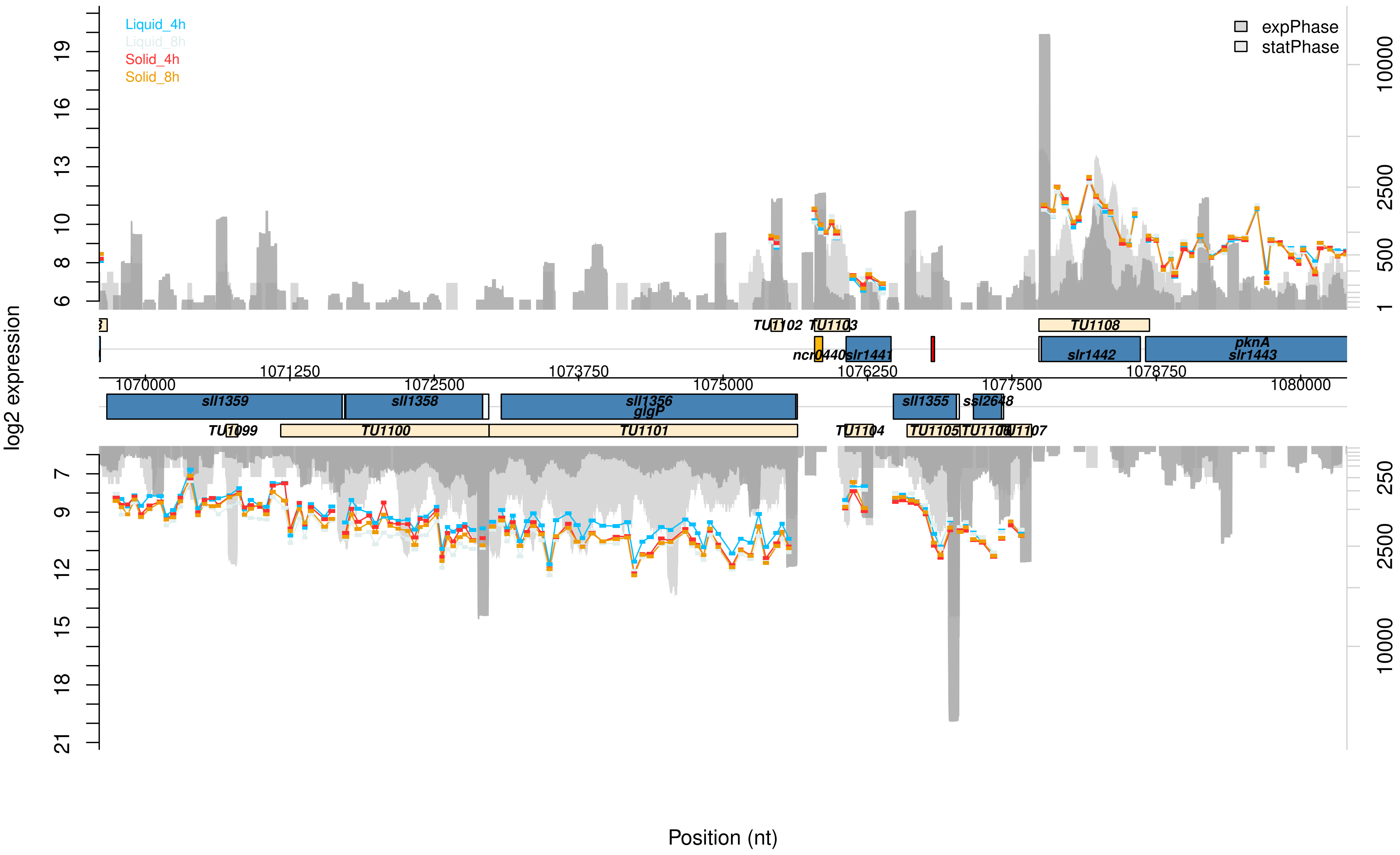

Liquid_4h
Liquid_8h
Solid_4h
Solid_8h
expPhase
statPhase
log2 expression
Position (nt)
TU1102
TU1103
TU1108
ncr0440
slr1441
slr1442
pknA
slr1443
1070000
1071250
1072500
1073750
1075000
1076250
1077500
1078750
1080000
sll1359
sll1358
sll1356
glgP
sll1355
ssl2648
TU1099
TU1100
TU1101
TU1104
TU1105
TU1106
TU1107

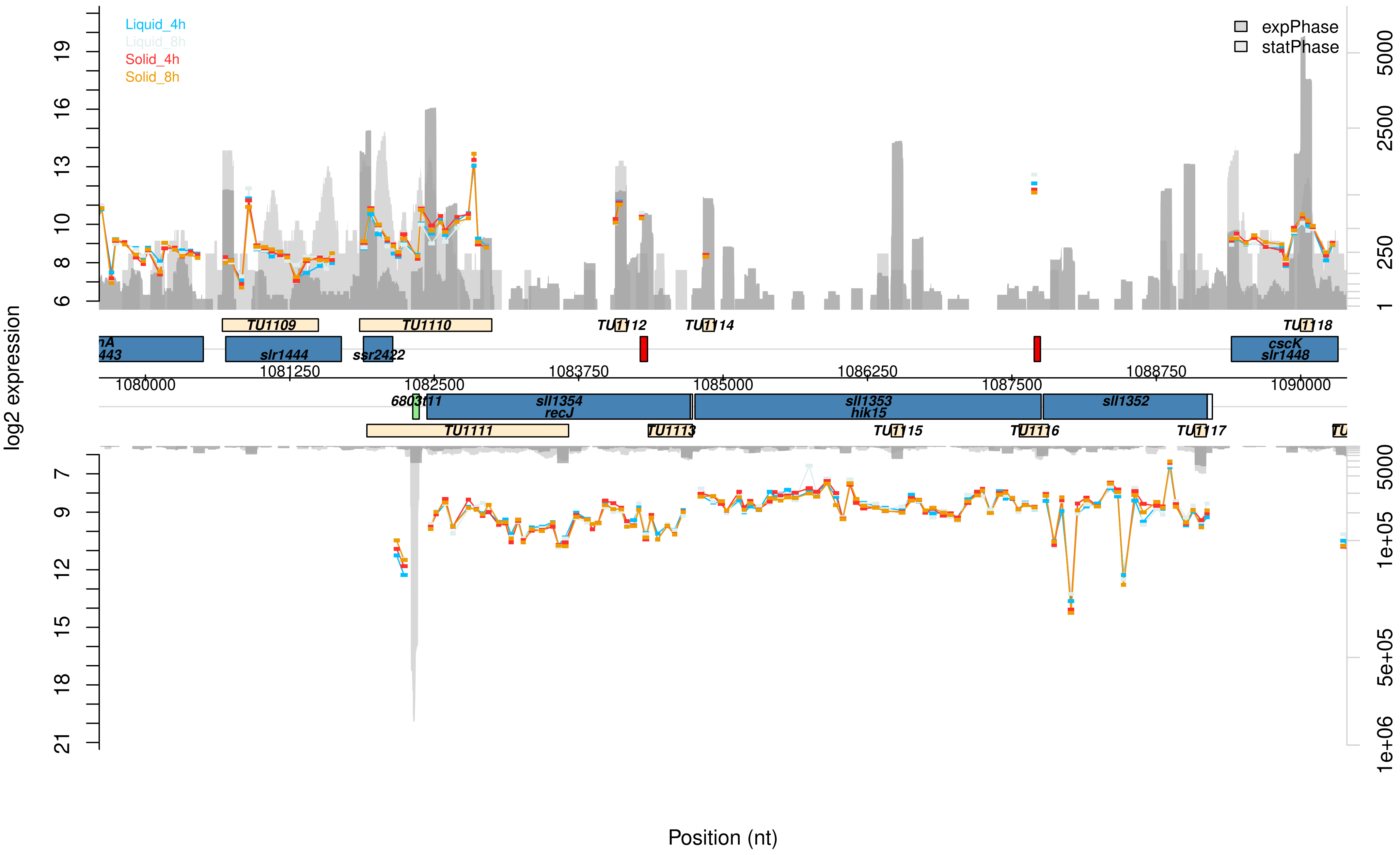

Liquid_4h
Liquid_8h
Solid_4h
Solid_8h
expPhase
statPhase
log2 expression
TU1109
TU1110
TU1112
TU1114
TU1118
nA
443
slr1444
ssr2422
cscK
slr1448
1080000
1081250
1082500
1083750
1085000
1086250
1087500
1088750
1090000
6803t11
sll1354
recJ
sll1353
hik15
sll1352
TU1111
TU1113
TU1115
TU1116
TU1117
Position (nt)

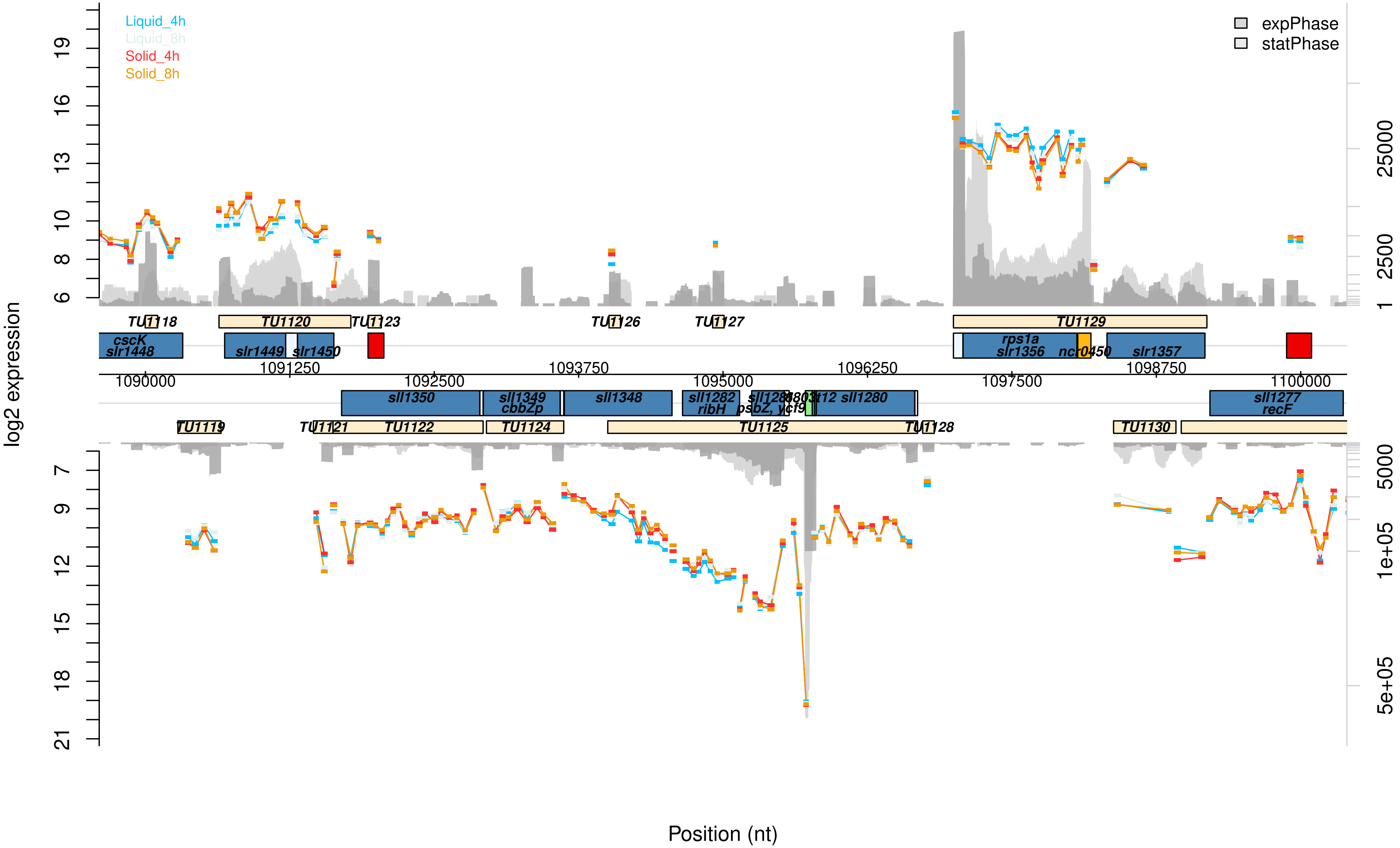

Liquid_4h
Liquid_8h
Solid_4h
Solid_8h
expPhase
statPhase
log2 expression
Position (nt)
TU1118
TU1120
TU1123
TU1126
TU1127
TU1129
cscK
slr1448
slr1449
slr1450
rps1a
slr1356
ncr0450
slr1357
1090000
1091250
1092500
1093750
1095000
1096250
1097500
1098750
1100000
sll1350
sll1349
cbbZp
sll1348
sll1282
ribH
psbZ, ycf9
sll1280
sll1277
recF
TU1119
TU1121
TU1122
TU1124
TU1125
TU1128
TU1130

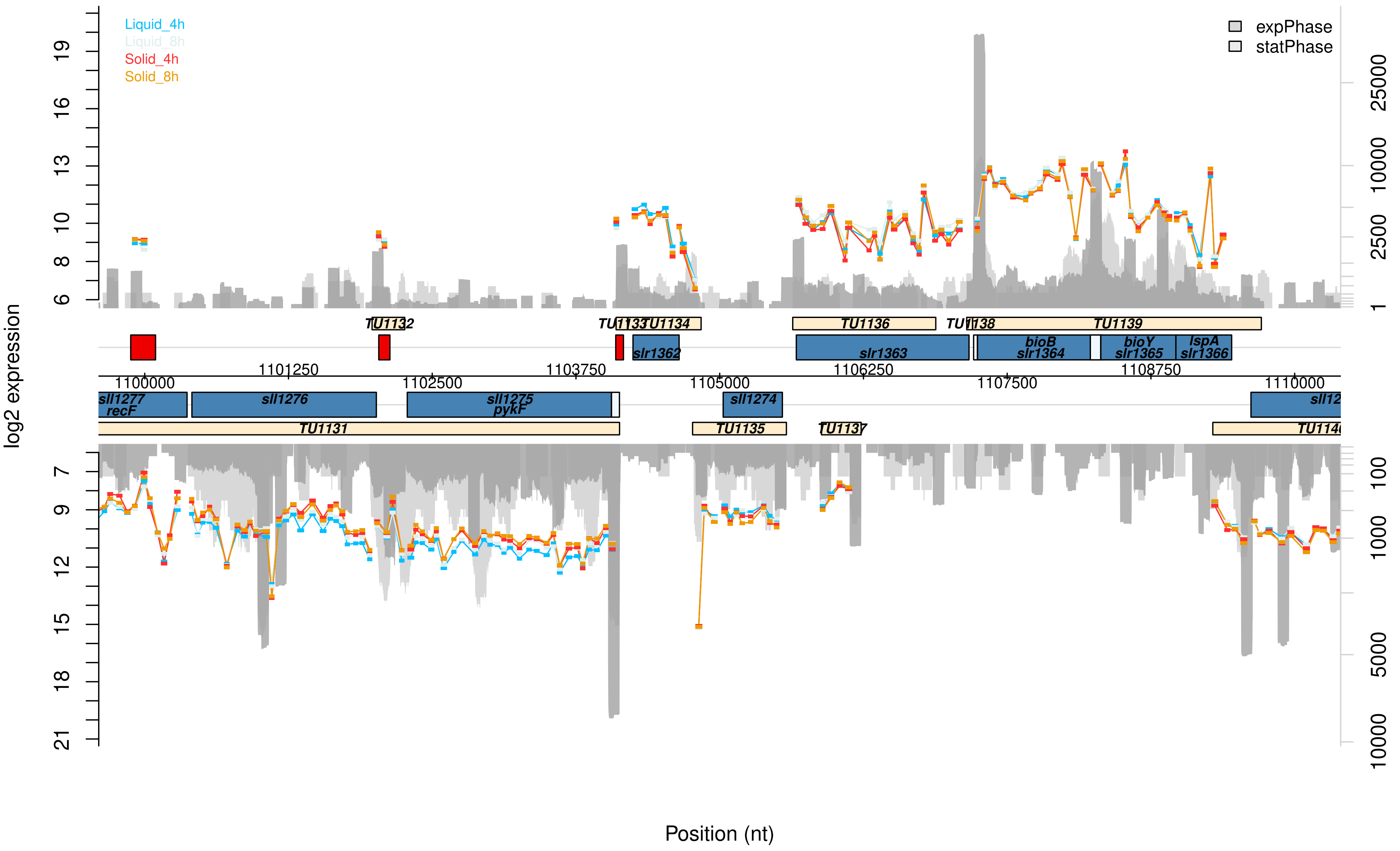

Liquid_4h
Liquid_8h
Solid_4h
Solid_8h
expPhase
statPhase
log2 expression
Position (nt)
TU1131
TU1132
TU1133
TU1134
TU1135
TU1136
TU1137
TU1138
TU1139
sll1277
recF
sll1276
sll1275
pykF
sll1274
slr1362
slr1363
bioB
slr1364
bioY
slr1365
lspA
slr1366
1100000
1101250
1102500
1103750
1105000
1106250
1107500
1108750
1110000

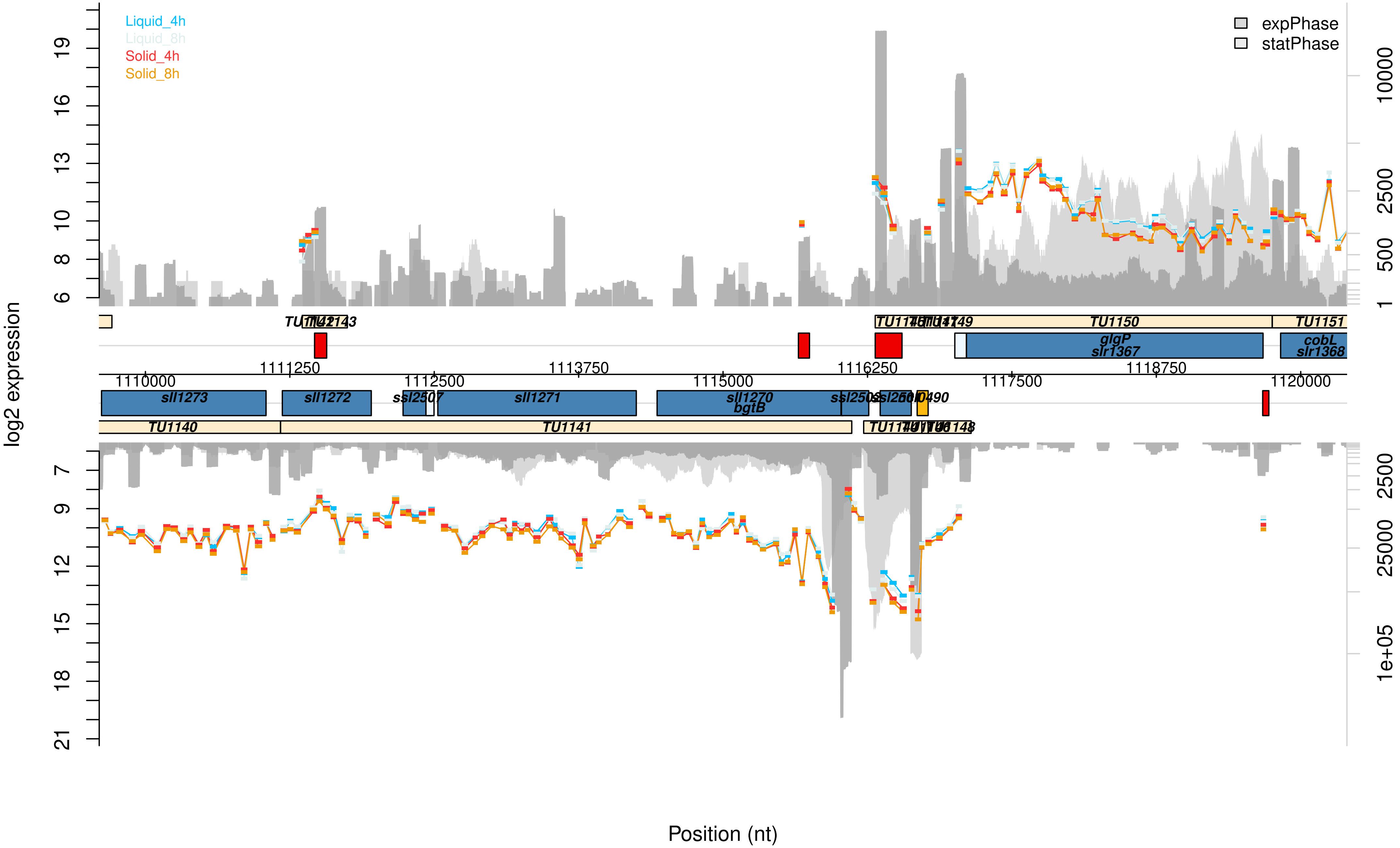

Liquid_4h
Liquid_8h
Solid_4h
Solid_8h
expPhase
statPhase
log2 expression
Position (nt)
TU1150
TU1151
glgP
slr1367
cobL
slr1368
1110000
1111250
1112500
1113750
1115000
1116250
1117500
1118750
1120000
sll1273
sll1272
ssl2507
sll1271
sll1270
bgtB
TU1140
TU1141

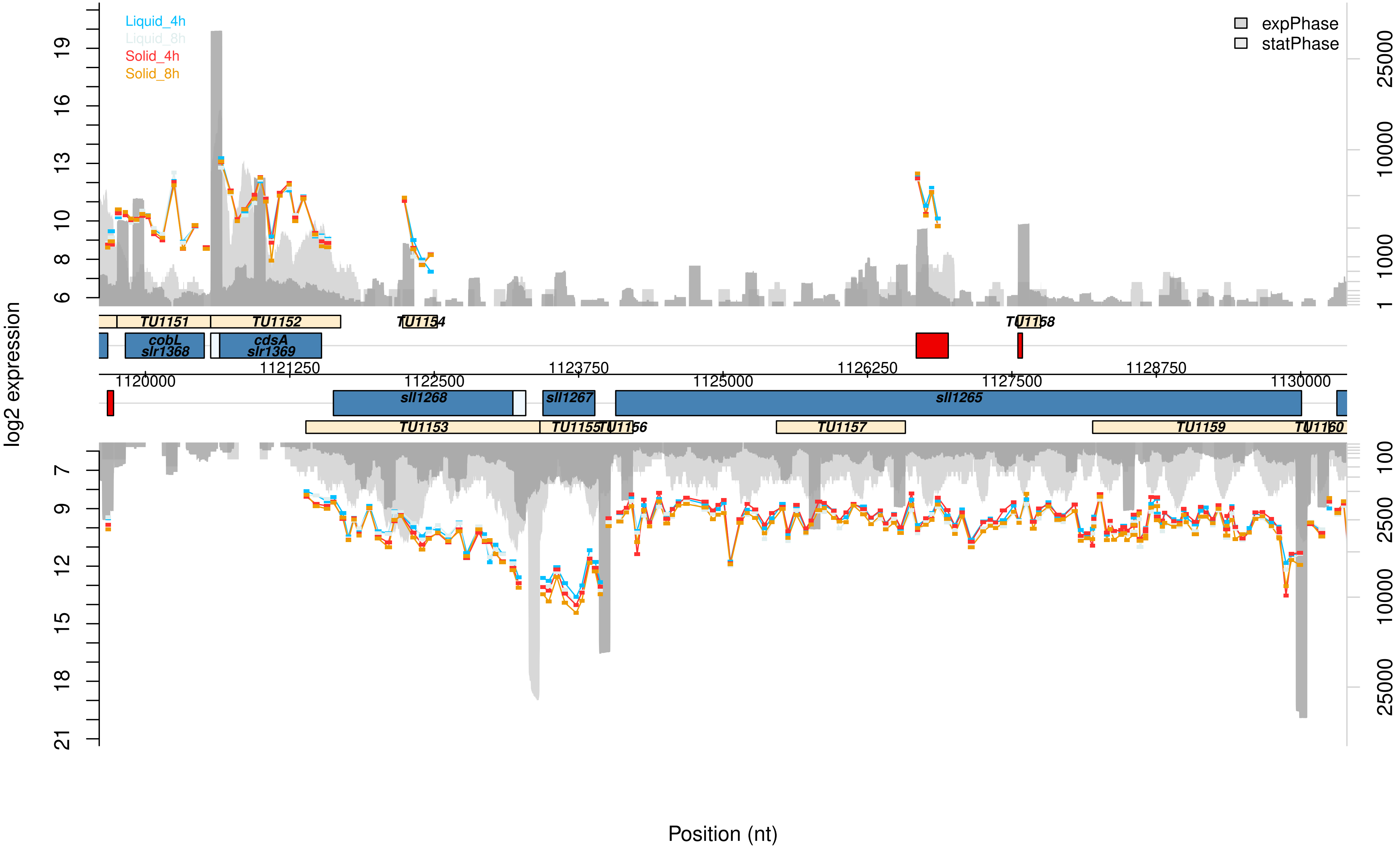

Liquid_4h
Liquid_8h
Solid_4h
Solid_8h
expPhase
statPhase
log2 expression
TU1151
TU1152
TU1154
TU1158
cobL
slr1368
cdsA
slr1369
1120000
1121250
1122500
1123750
1125000
1126250
1127500
1128750
1130000
sll1268
sll1267
sll1265
TU1153
TU1155
TU1156
TU1157
TU1159
TU1160
Position (nt)

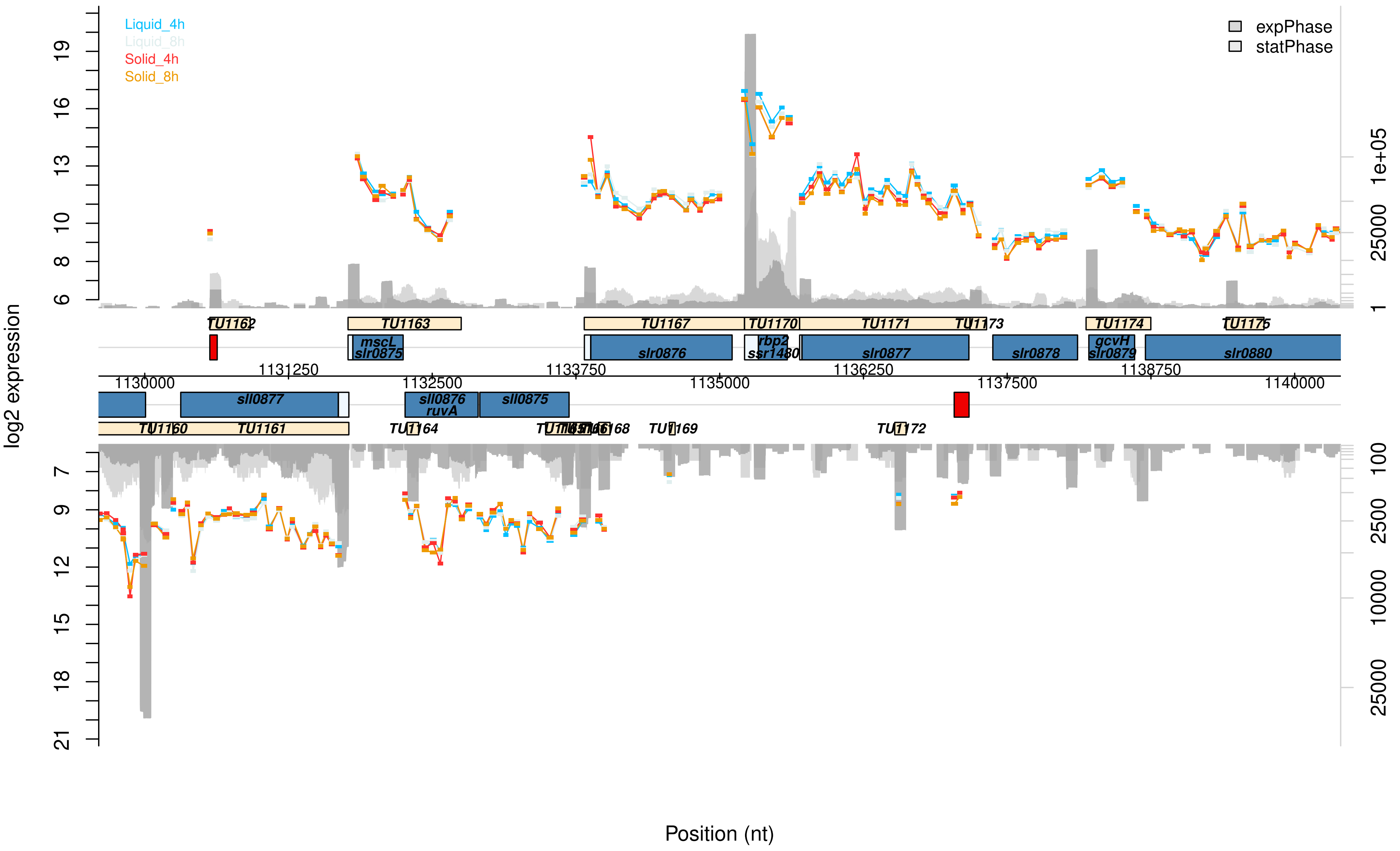

Liquid_4h
Liquid_8h
Solid_4h
Solid_8h
expPhase
statPhase
log2 expression
Position (nt)
TU1162
TU1163
TU1167
TU1170
TU1171
TU1173
TU1174
TU1175
mscL
slr0875
slr0876
rbp2
ssr1480
slr0877
slr0878
gcvH
slr0879
slr0880
1130000
1131250
1132500
1133750
1135000
1136250
1137500
1138750
1140000
sll0877
sll0876
ruvA
sll0875
TU1160
TU1161
TU1164
TU1168
TU1169
TU1172

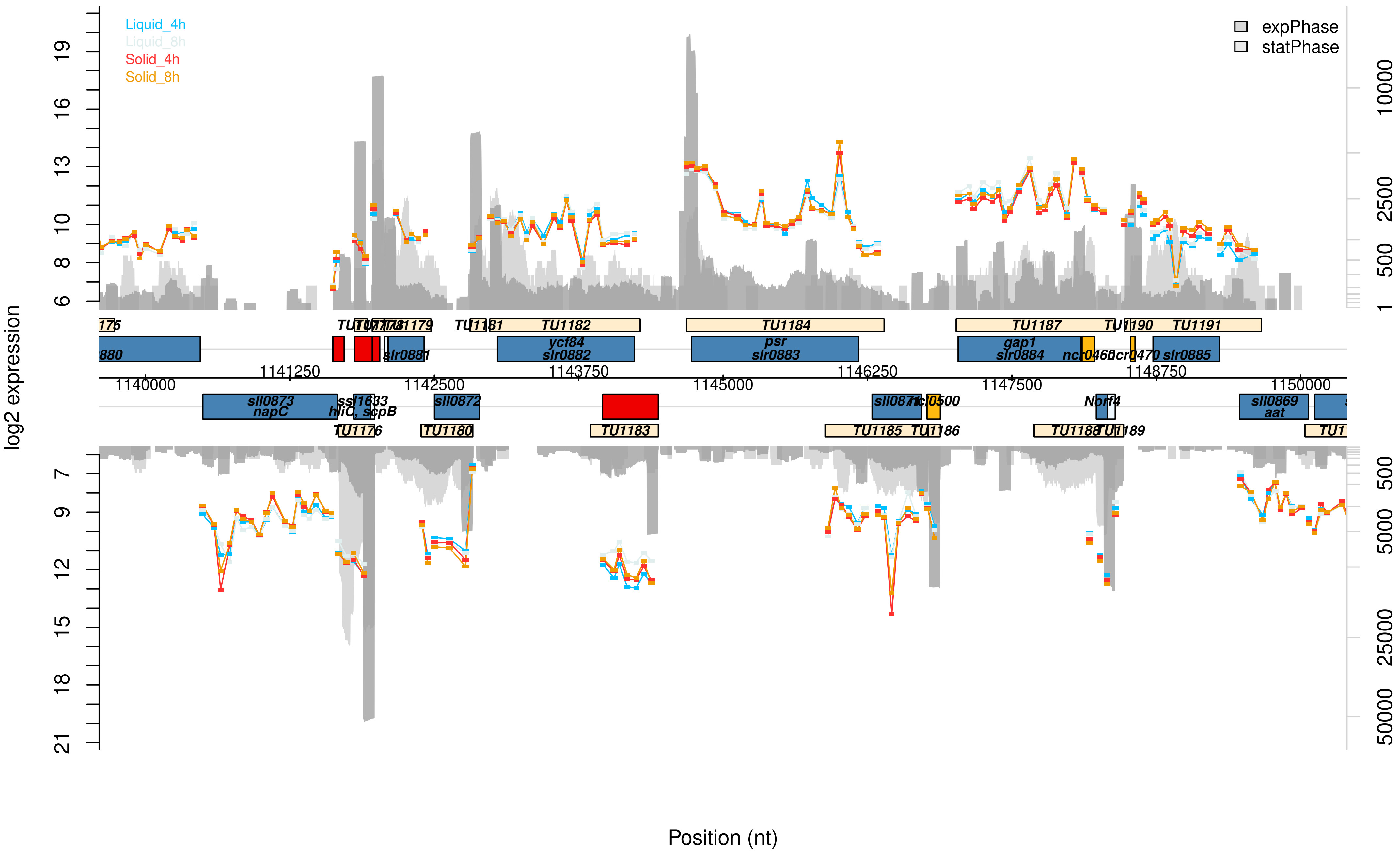

Liquid_4h
Liquid_8h
Solid_4h
Solid_8h
expPhase
statPhase
log2 expression
Position (nt)
TU1182
TU1184
TU1187
TU1191
ycf84
slr0882
psr
slr0883
gap1
slr0884
slr0885
slr0881
sll0873
napC
hliC, scpB
sll0872
TU1176
TU1180
TU1183
TU1185
sll0869
aat
Norf4
1140000
1141250
1142500
1143750
1145000
1146250
1147500
1148750
1150000

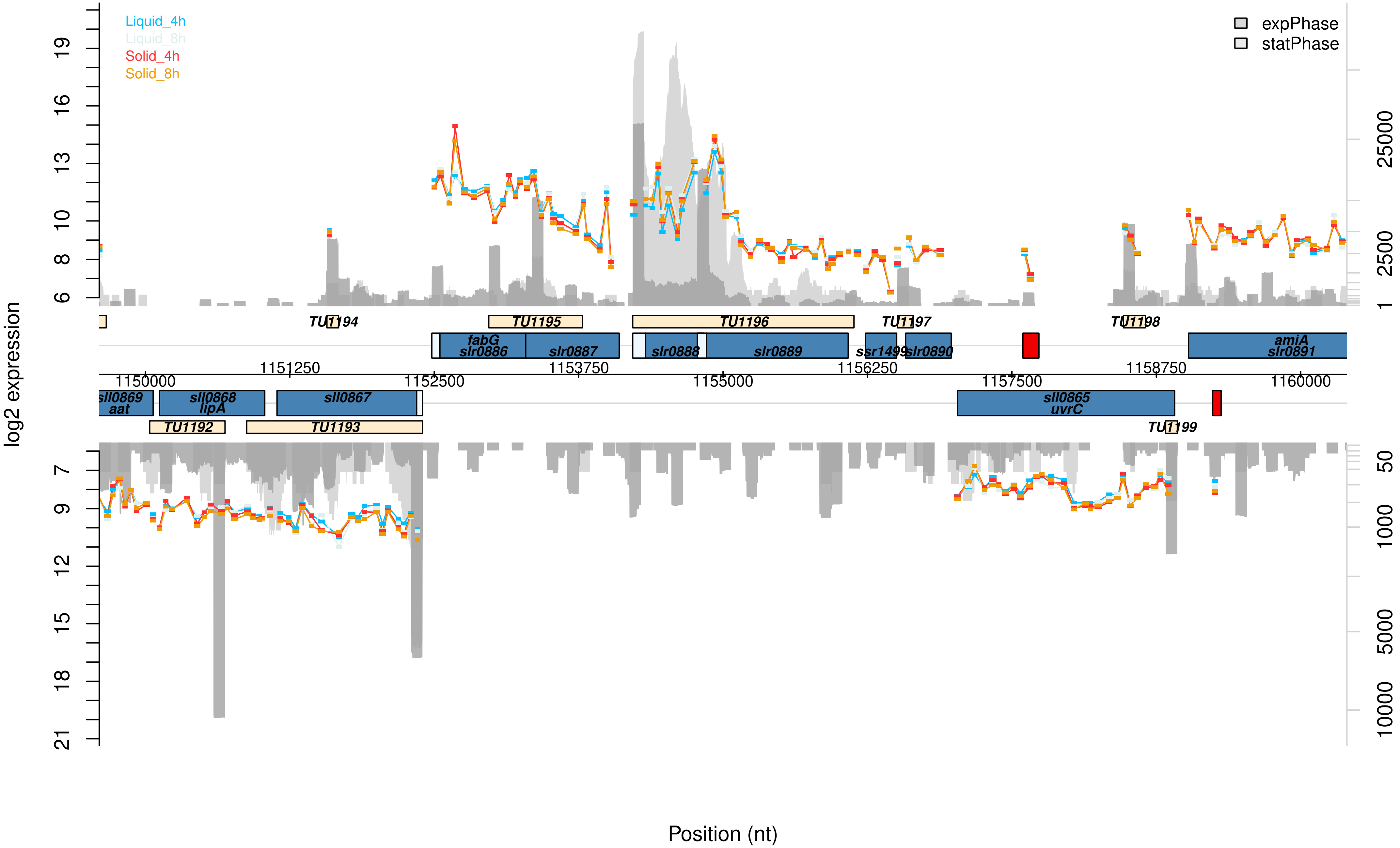

Liquid_4h
Liquid_8h
Solid_4h
Solid_8h
expPhase
statPhase
log2 expression
Position (nt)
TU1194
TU1195
TU1196
TU1197
TU1198
fabG
slr0886
slr0887
slr0888
slr0889
ssr1499
slr0890
amiA
slr0891
1150000
1151250
1152500
1153750
1155000
1156250
1157500
1158750
1160000
sll0869
aat
sll0868
lipA
sll0867
sll0865
uvrC
TU1192
TU1193
TU1199

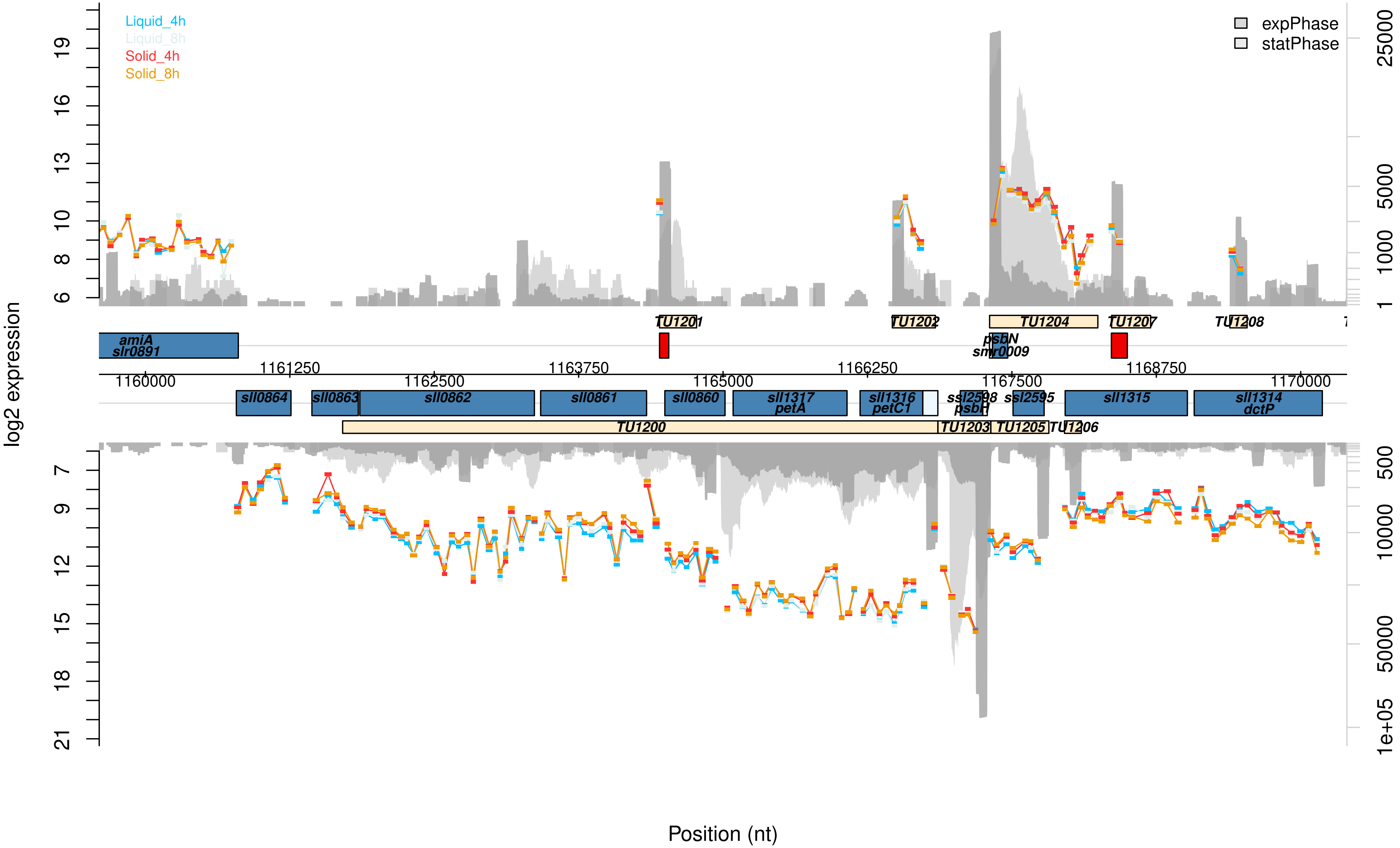
Liquid_4h
Liquid_8h
Solid_4h
Solid_8h
expPhase
statPhase
log2 expression
Position (nt)
amiA
slr0891
TU1201
TU1202
TU1204
TU1207
TU1208
psbN
smr0009
sll0864
sll0863
sll0862
sll0861
sll0860
sll1317
petA
sll1316
petC1
ssl2598
psbH
ssl2595
sll1315
sll1314
dctP
TU1200
TU1203
TU1205
TU1206
1160000
1161250
1162500
1163750
1165000
1166250
1167500
1168750
1170000
25000
5000
1000
1
500
10000
50000
1e+05

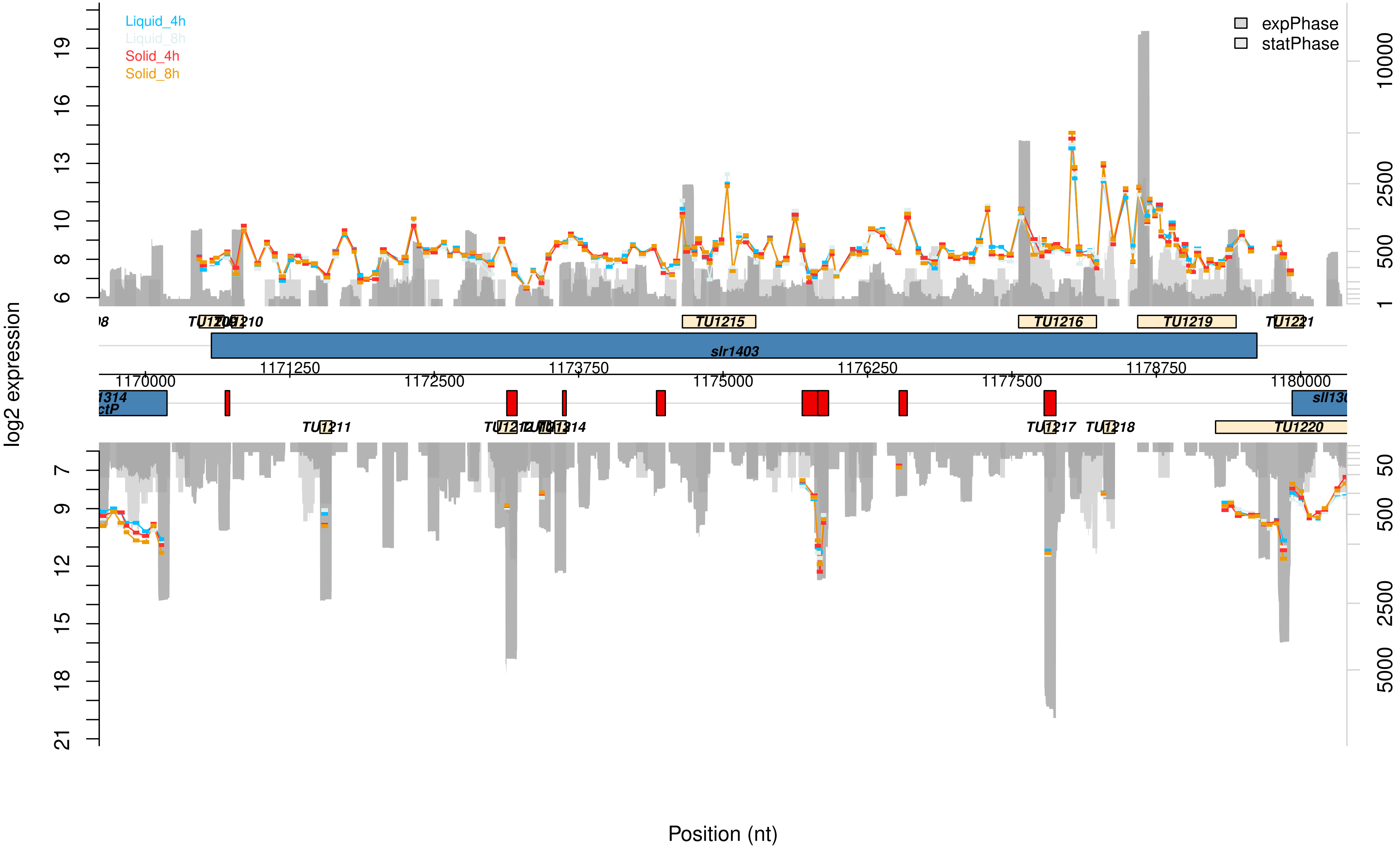

Liquid_4h
Liquid_8h
Solid_4h
Solid_8h
expPhase
statPhase
log2 expression
Position (nt)
TU1215
TU1216
TU1219
TU1221
slr1403
1171250
1173750
1176250
1178750
1170000
1172500
1175000
1177500
1180000
TU1211
TU1217
TU1218
TU1220

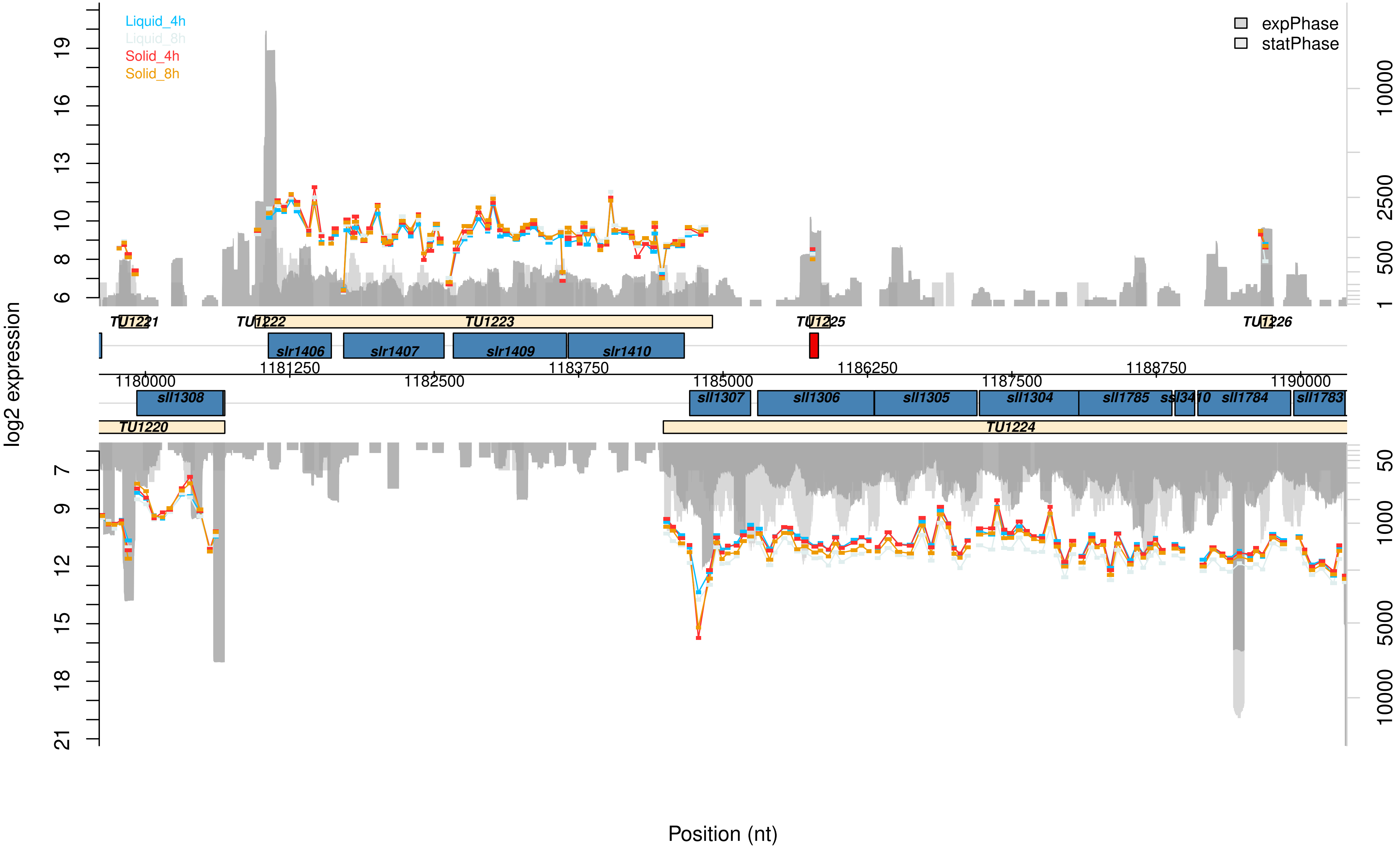

Liquid_4h
Liquid_8h
Solid_4h
Solid_8h
expPhase
statPhase
log2 expression
Position (nt)
TU1221
TU1222
TU1223
TU1225
TU1226
slr1406
slr1407
slr1409
slr1410
1180000
1181250
1182500
1183750
1185000
1186250
1187500
1188750
1190000
sll1308
sll1307
sll1306
sll1305
sll1304
sll1785
ssl3410
sll1784
sll1783
TU1220
TU1224

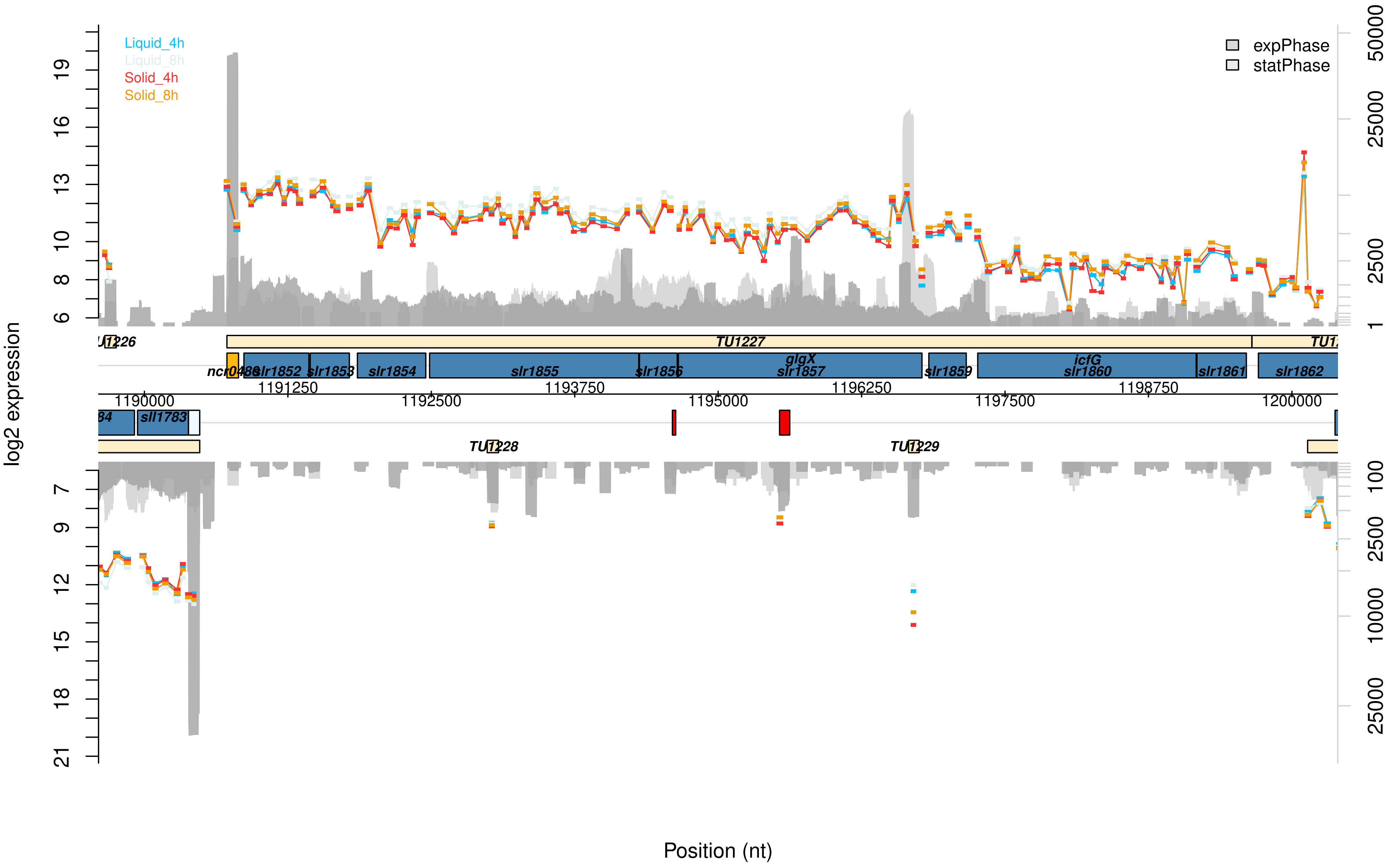

Liquid_4h
Liquid_8h
Solid_4h
Solid_8h
expPhase
statPhase
log2 expression
Position (nt)
TU1227
ncr0488
slr1852
slr1853
slr1854
slr1855
slr1856
glgX
slr1857
slr1859
icfG
slr1860
slr1861
slr1862
sll1783
TU1228
TU1229
1190000
1191250
1192500
1193750
1195000
1196250
1197500
1198750
1200000

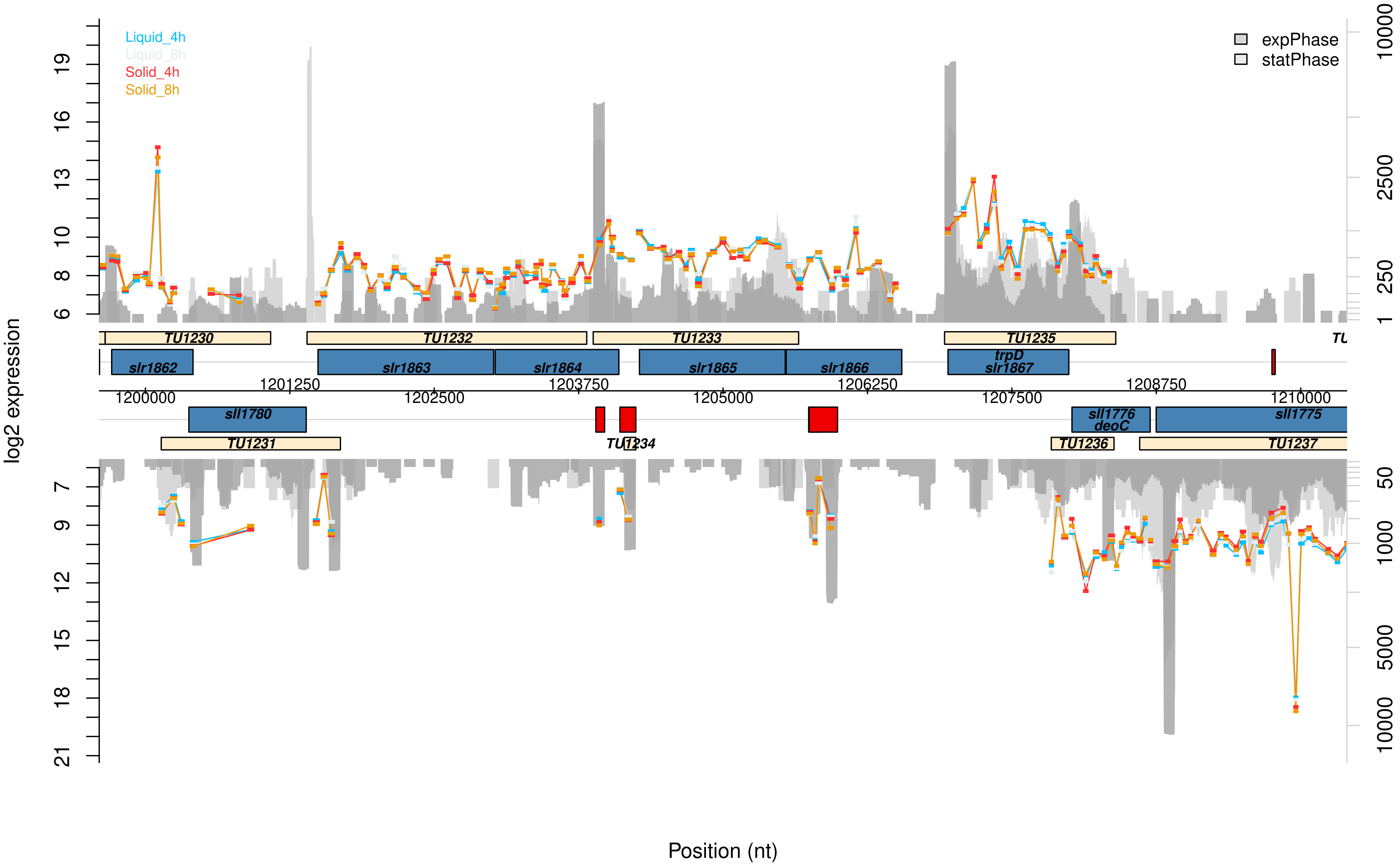

Liquid_4h
Liquid_8h
Solid_4h
Solid_8h
expPhase
statPhase
log2 expression
TU1230
TU1232
TU1233
TU1235
slr1862
slr1863
slr1864
slr1865
slr1866
trpD
slr1867
1200000
1201250
1202500
1203750
1205000
1206250
1207500
1208750
1210000
sll1780
sll1776
deoC
sll1775
TU1231
TU1234
TU1236
TU1237
Position (nt)

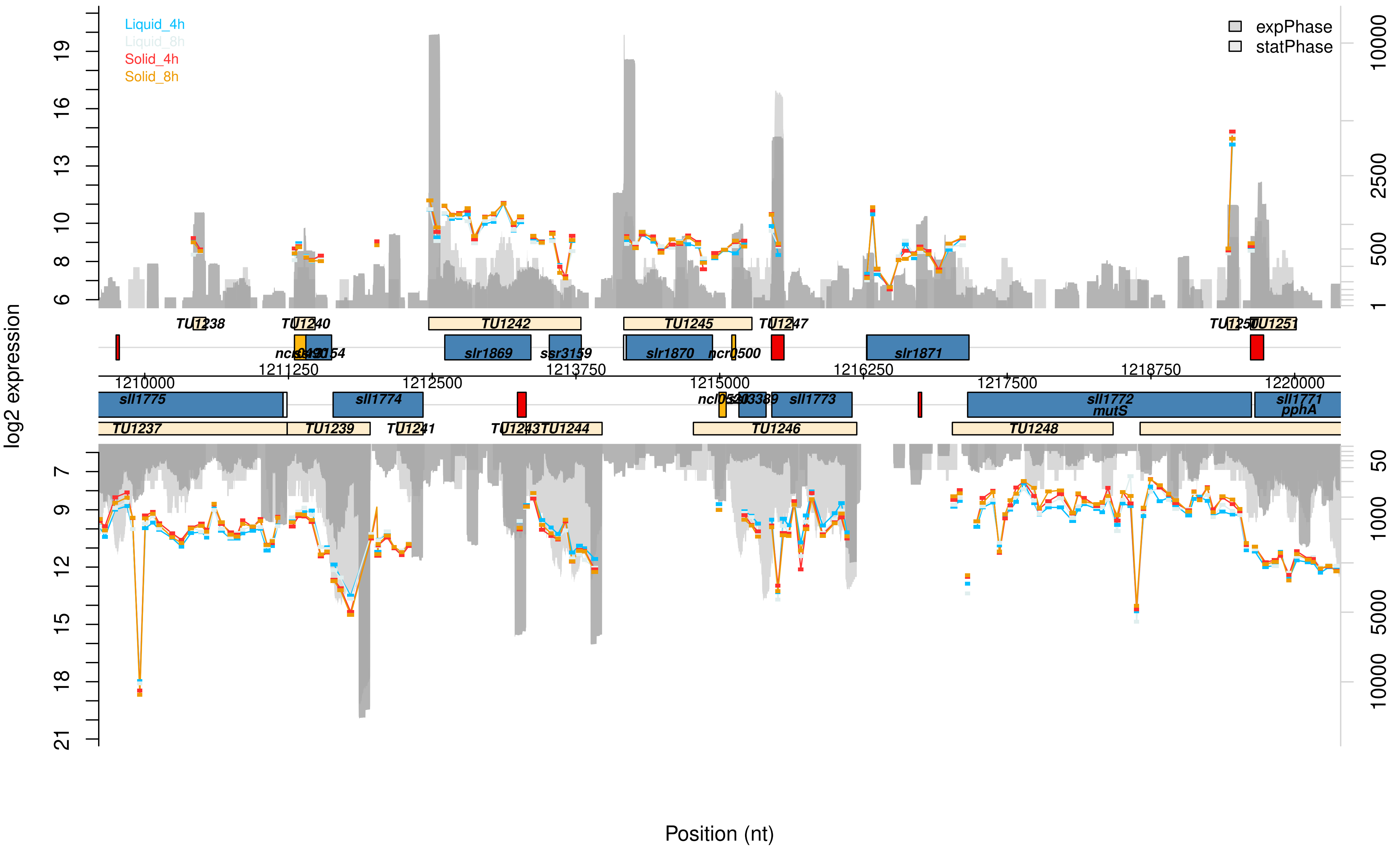
Liquid_4h
Liquid_8h
Solid_4h
Solid_8h
expPhase
statPhase
log2 expression
Position (nt)
TU1237
TU1238
TU1239
TU1240
TU1241
TU1242
TU1245
TU1246
TU1247
TU1248
TU1251
sll1775
sll1774
slr1869
ssr3159
slr1870
ncr0500
slr1871
sll1773
sll1772
mutS
sll1771
pphA
1210000
1211250
1212500
1213750
1215000
1216250
1217500
1218750
1220000

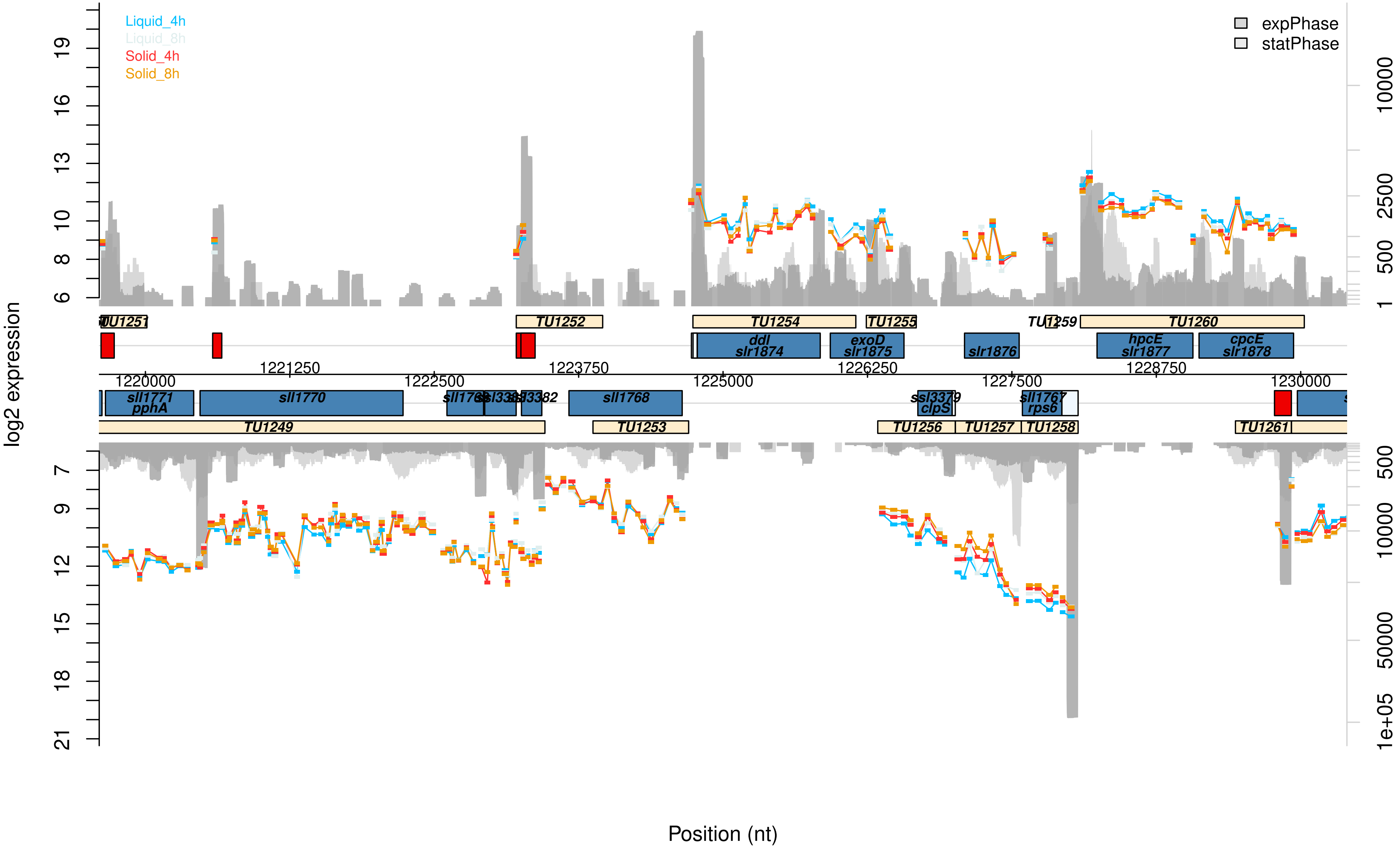

Liquid_4h
Liquid_8h
Solid_4h
Solid_8h
expPhase
statPhase
log2 expression
Position (nt)
TU1251
TU1252
TU1254
TU1255
TU1259
TU1260
ddl
slr1874
exoD
slr1875
slr1876
hpcE
slr1877
cpcE
slr1878
1220000
1221250
1222500
1223750
1225000
1226250
1227500
1228750
1230000
sll1771
pphA
sll1770
sll1768
ssl3379
clpS
sll1767
rps6
TU1249
TU1253
TU1256
TU1257
TU1258
TU1261

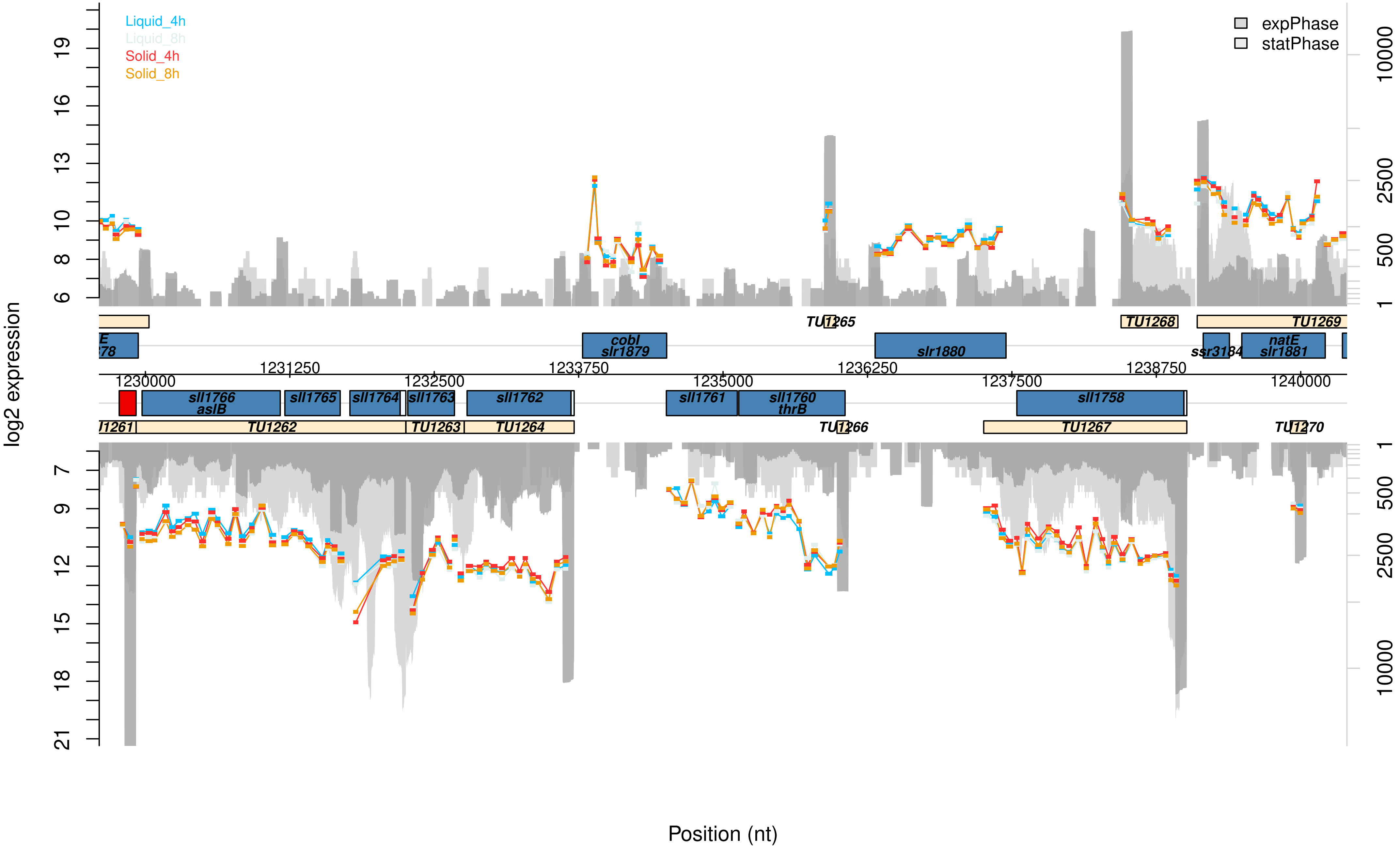

Liquid_4h
Liquid_8h
Solid_4h
Solid_8h
expPhase
statPhase
log2 expression
Position (nt)
TU1265
TU1268
TU1269
cobI
slr1879
slr1880
ssr3184
natE
slr1881
1231250
1233750
1236250
1238750
1230000
1232500
1235000
1237500
1240000
sll1766
asIB
sll1765
sll1764
sll1763
sll1762
sll1761
sll1760
thrB
sll1758
TU1262
TU1263
TU1264
TU1266
TU1267
TU1270

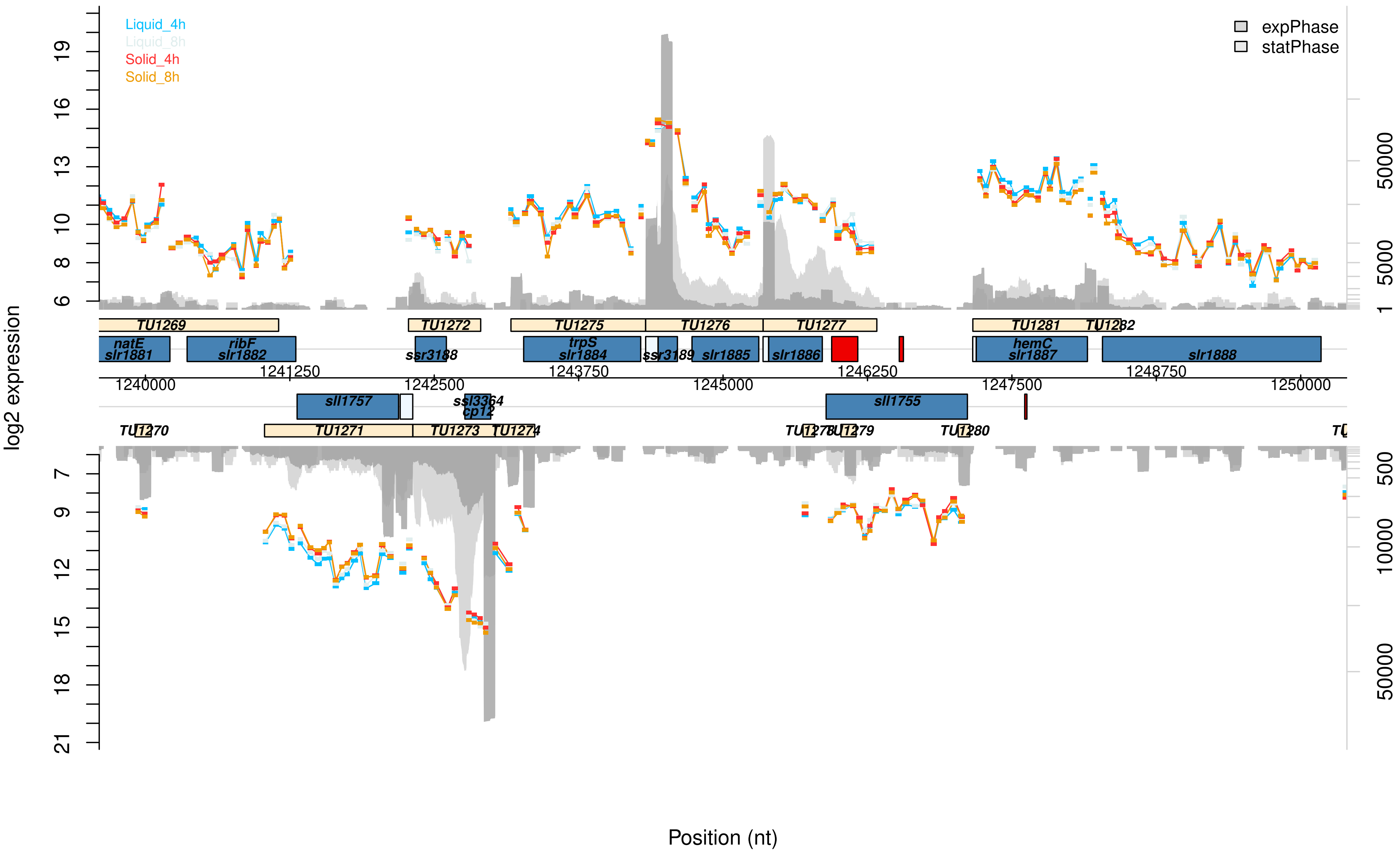
Liquid_4h
Liquid_8h
Solid_4h
Solid_8h
expPhase
statPhase
log2 expression
Position (nt)
TU1269
TU1272
TU1275
TU1276
TU1277
TU1281
TU1282
natE
slr1881
ribF
slr1882
ssr3188
trpS
slr1884
ssr3189
slr1885
slr1886
hemC
slr1887
slr1888
1240000
1241250
1242500
1243750
1245000
1246250
1247500
1248750
1250000
sll1757
ssl3364
cp12
sll1755
TU1270
TU1271
TU1273
TU1274
TU1278
TU1279
TU1280

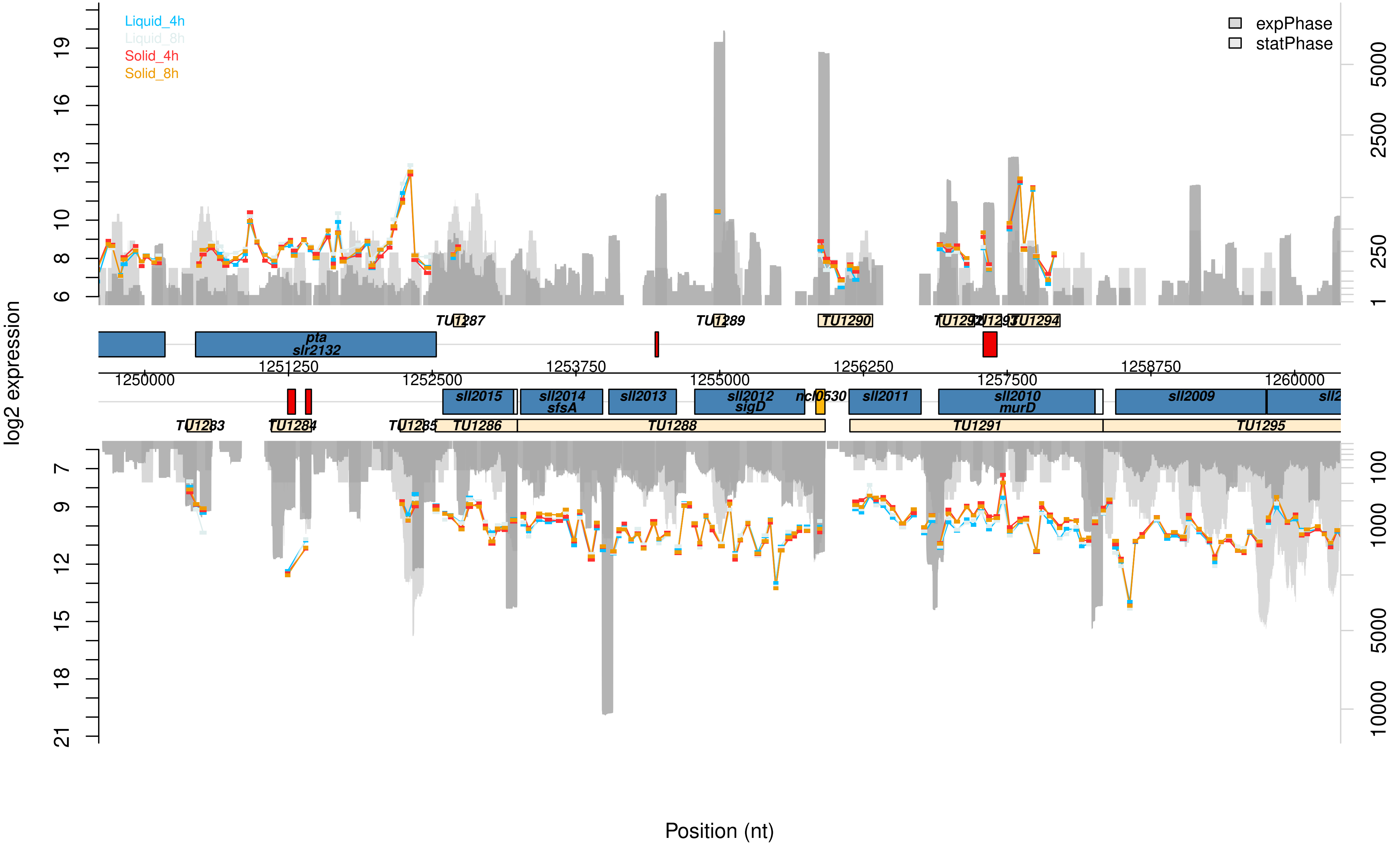

log2 expression
Liquid_4h
Liquid_8h
Solid_4h
Solid_8h
expPhase
statPhase
TU1287
TU1289
TU1290
TU1292
TU1293
TU1294
pta
slr2132
1250000
1251250
1252500
1253750
1255000
1256250
1257500
1258750
1260000
sll2015
sll2014
sfsA
sll2013
sll2012
sigD
ncl0530
sll2011
sll2010
murD
sll2009
TU1283
TU1284
TU1285
TU1286
TU1288
TU1291
TU1295
Position (nt)

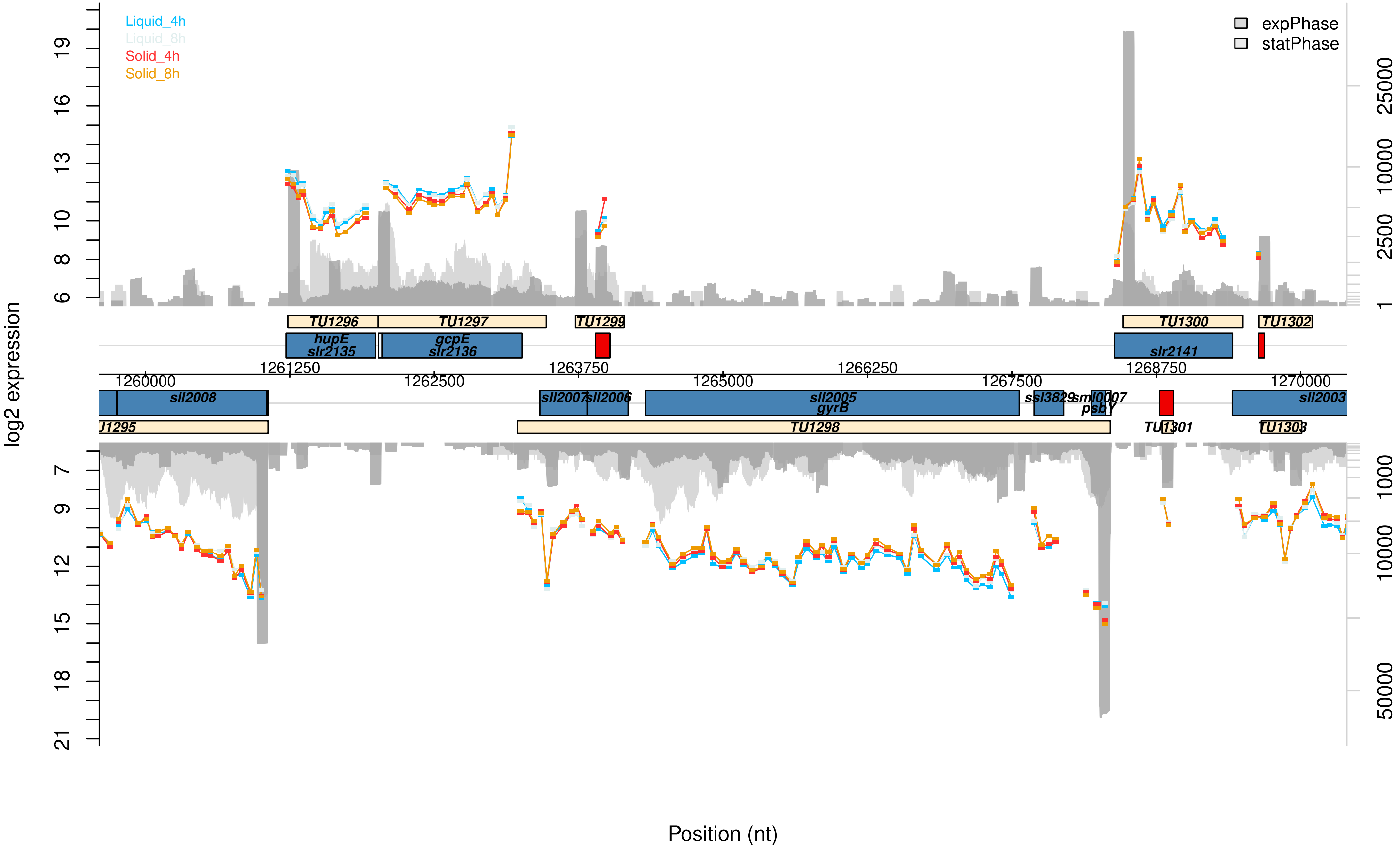

Liquid_4h
Liquid_8h
Solid_4h
Solid_8h
expPhase
statPhase
log2 expression
Position (nt)
TU1296
TU1297
TU1299
TU1300
TU1302
hupE
slr2135
gcpE
slr2136
slr2141
1261250
1263750
1266250
1268750
1260000
1262500
1265000
1267500
1270000
sll2008
sll2007
sll2006
sll2005
gyrB
ssl3829
sml0007
psbY
sll2003
J1295
TU1298
TU1301
TU1303

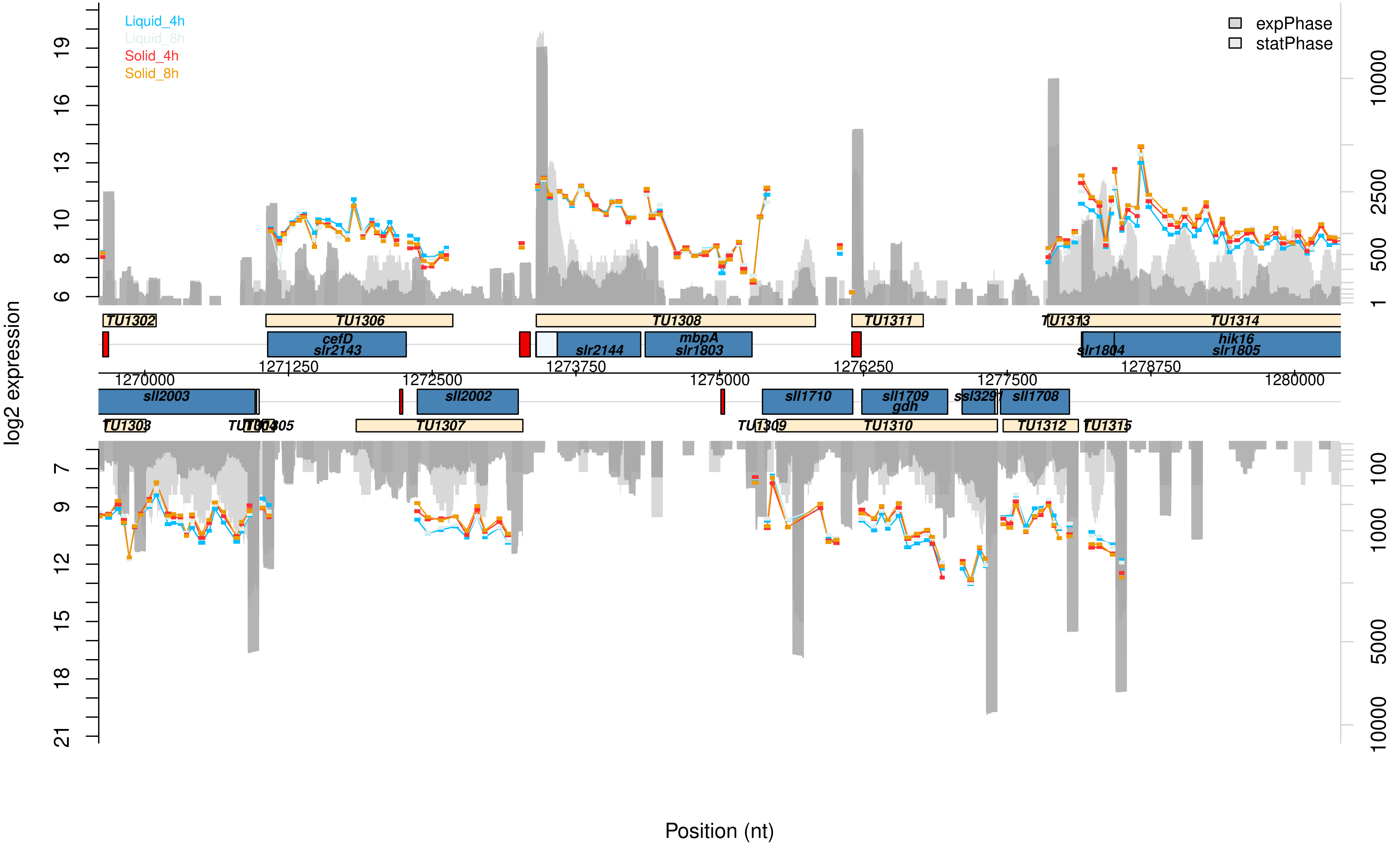

Liquid_4h
Liquid_8h
Solid_4h
Solid_8h
expPhase
statPhase
log2 expression
Position (nt)
TU1302
TU1306
TU1308
TU1311
TU1313
TU1314
cefD
slr2143
slr2144
mbpA
slr1803
slr1804
hik16
slr1805
sll2003
sll2002
sll1710
sll1709
gdh
ssl3291
sll1708
TU1303
TU1307
TU1310
TU1312
TU1315

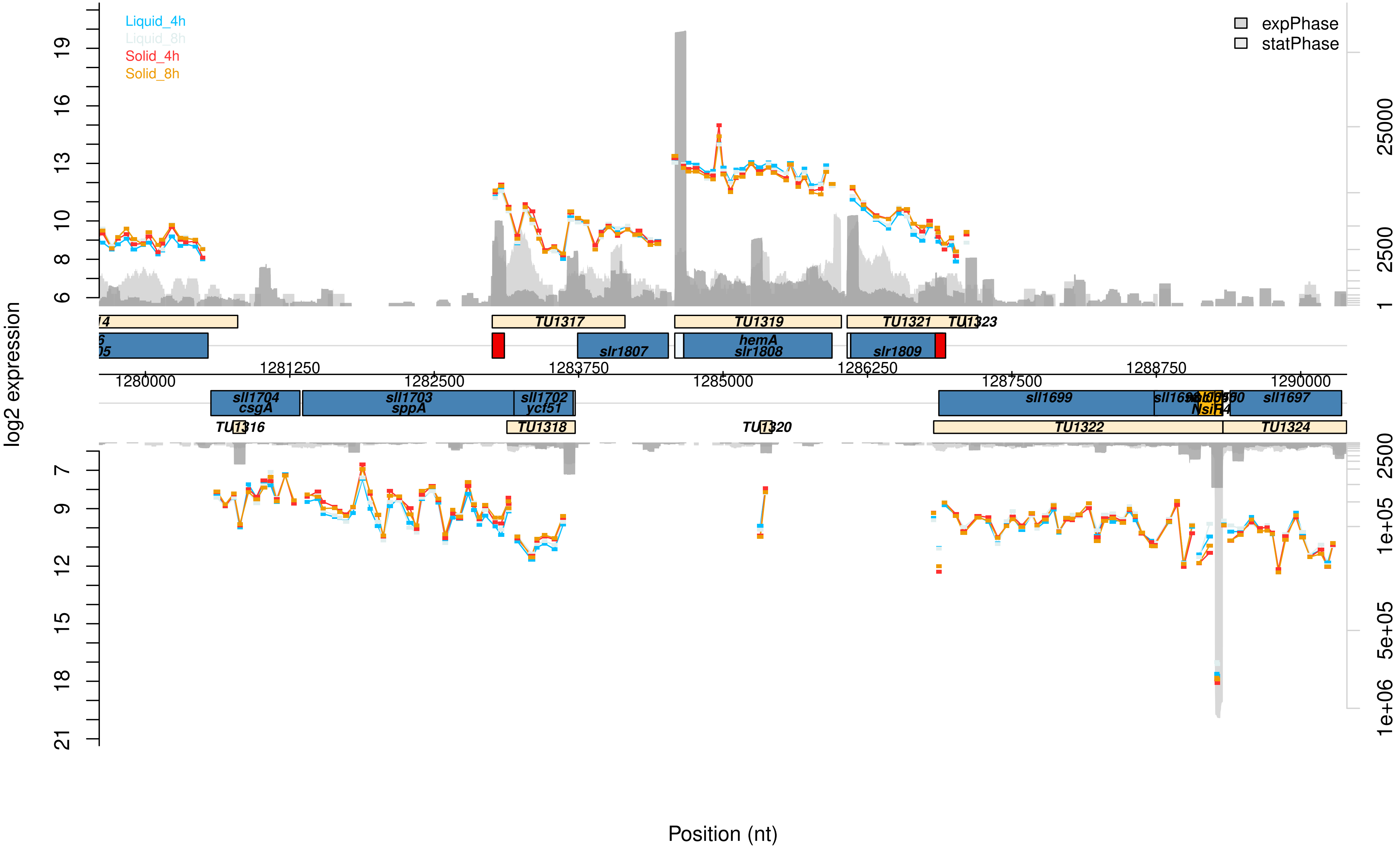

Liquid_4h
Liquid_8h
Solid_4h
Solid_8h
expPhase
statPhase
log2 expression
Position (nt)
TU1317
TU1319
TU1321
TU1323
slr1807
hemA
slr1808
slr1809
sll1704
csgA
sll1703
sppA
sll1702
ycf51
sll1699
sll1697
TU1316
TU1318
TU1320
TU1322
TU1324
1280000
1281250
1282500
1283750
1285000
1286250
1287500
1288750
1290000

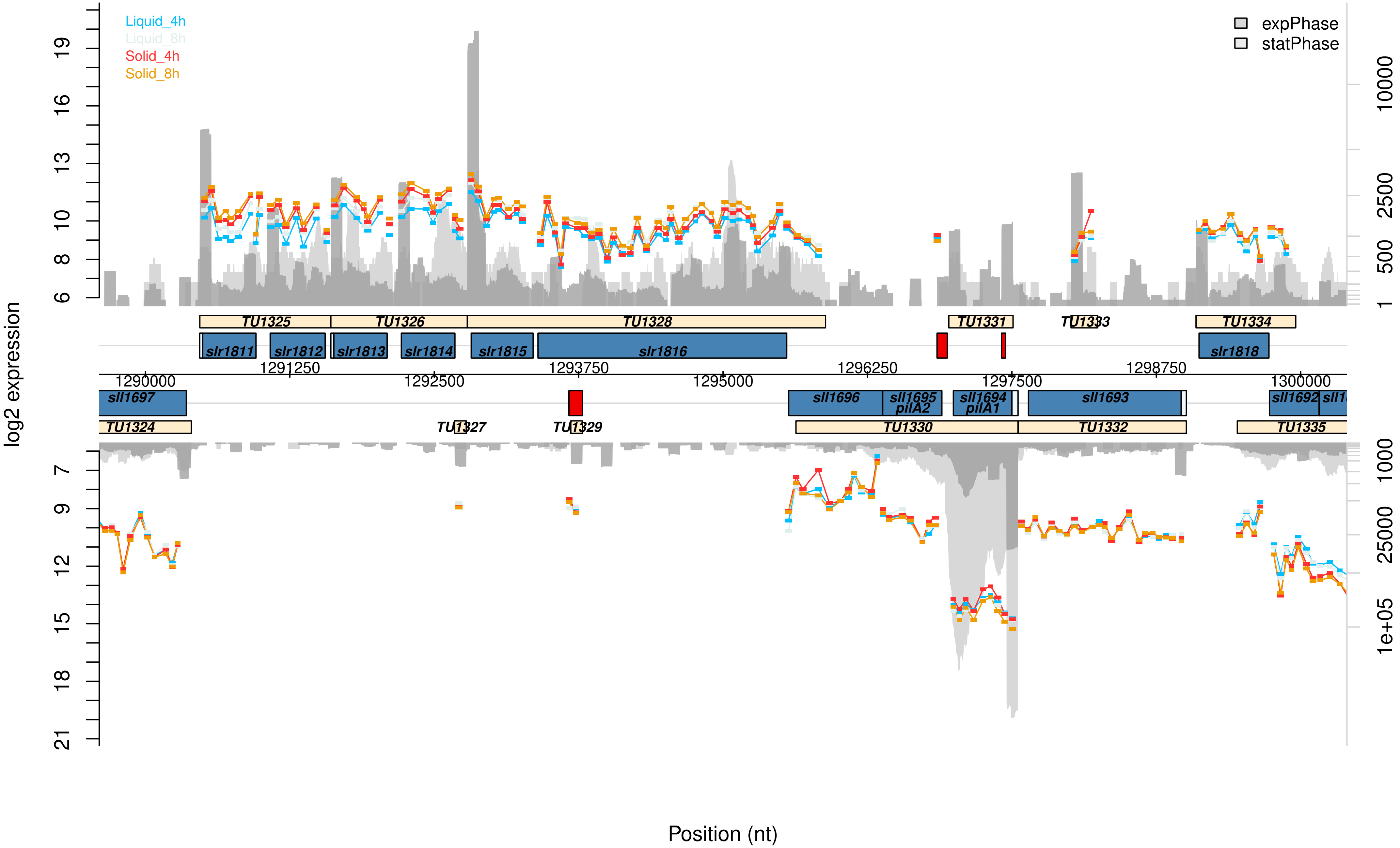

Liquid_4h
Liquid_8h
Solid_4h
Solid_8h
expPhase
statPhase
log2 expression
Position (nt)
TU1325
TU1326
TU1328
TU1331
TU1333
TU1334
slr1811
slr1812
slr1813
slr1814
slr1815
slr1816
slr1818
1290000
1291250
1292500
1293750
1295000
1296250
1297500
1298750
1300000
sll1697
sll1696
sll1695
pilA2
sll1694
pilA1
sll1693
sll1692
TU1324
TU1327
TU1329
TU1330
TU1332
TU1335

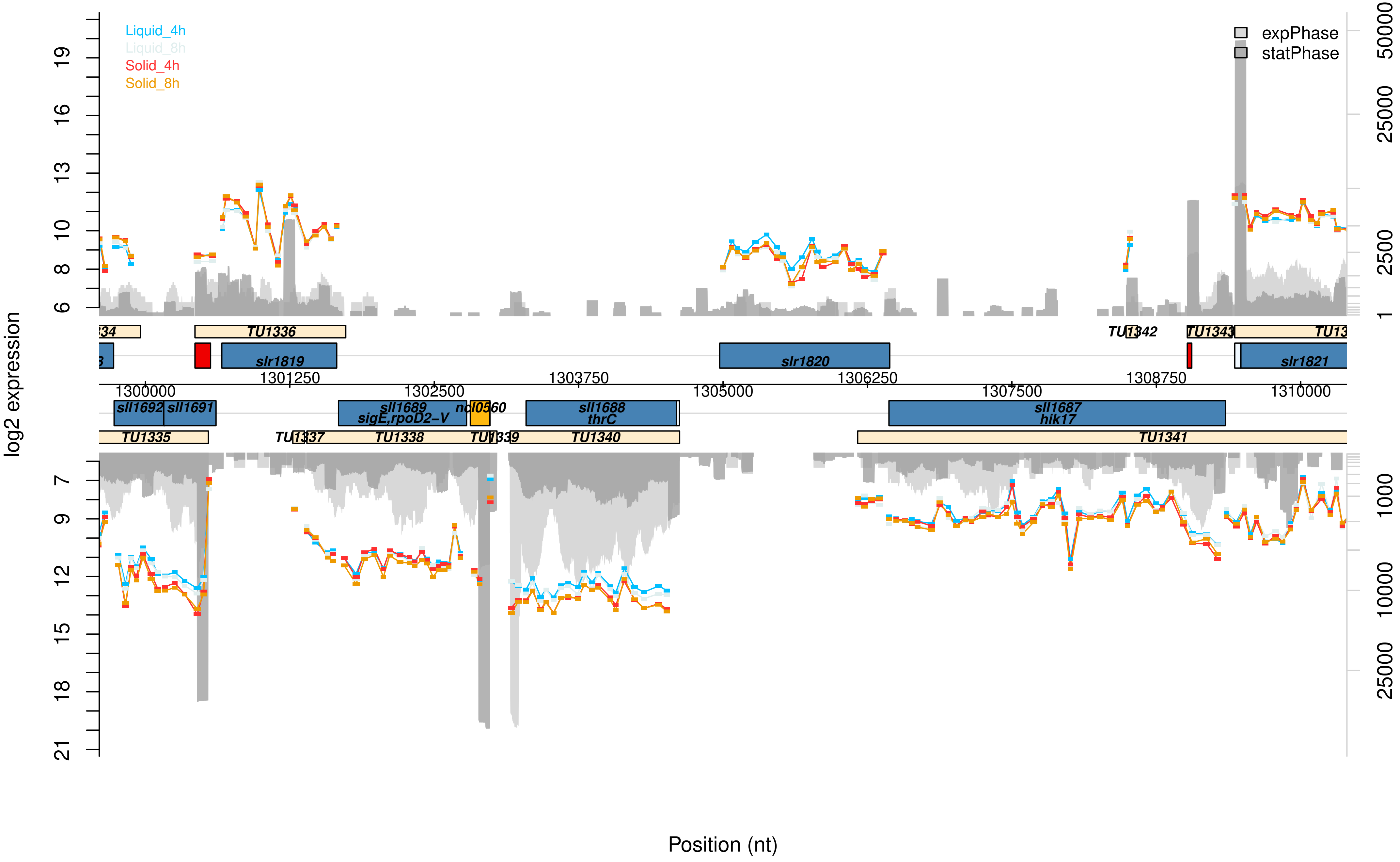

Liquid_4h
Liquid_8h
Solid_4h
Solid_8h
expPhase
statPhase
log2 expression
Position (nt)
TU1336
TU1342
TU1343
slr1819
slr1820
slr1821
sll1692
sll1691
sll1689
sigE,rpoD2−V
ncl0560
sll1688
thrC
sll1687
hik17
TU1335
TU1338
TU1340
TU1341
1300000
1301250
1302500
1303750
1305000
1306250
1307500
1308750
1310000

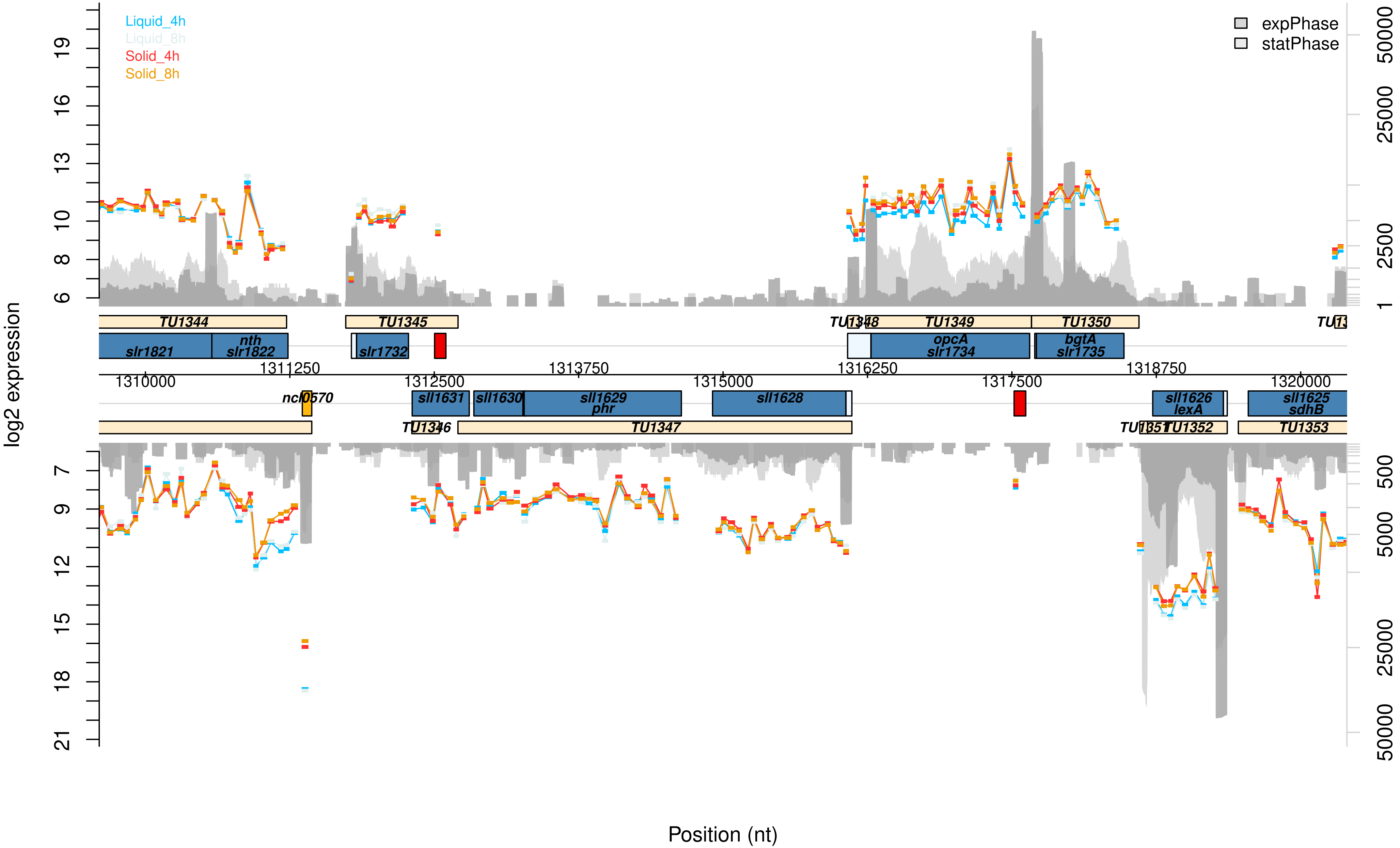

Liquid_4h
Liquid_8h
Solid_4h
Solid_8h
expPhase
statPhase
log2 expression
19
16
13
10
8
6
7
9
12
15
18
21
50000
25000
2500
1
500
5000
25000
50000
TU1344
TU1345
TU1348
TU1349
TU1350
slr1821
nth
slr1822
slr1732
opcA
slr1734
bgtA
slr1735
1311250
1312500
1313750
1315000
1316250
1317500
1318750
1310000
1320000
ncl0570
sll1631
sll1630
sll1629
phr
sll1628
sll1626
lexA
sll1625
sdhB
TU1346
TU1347
TU1351
TU1352
TU1353
Position (nt)

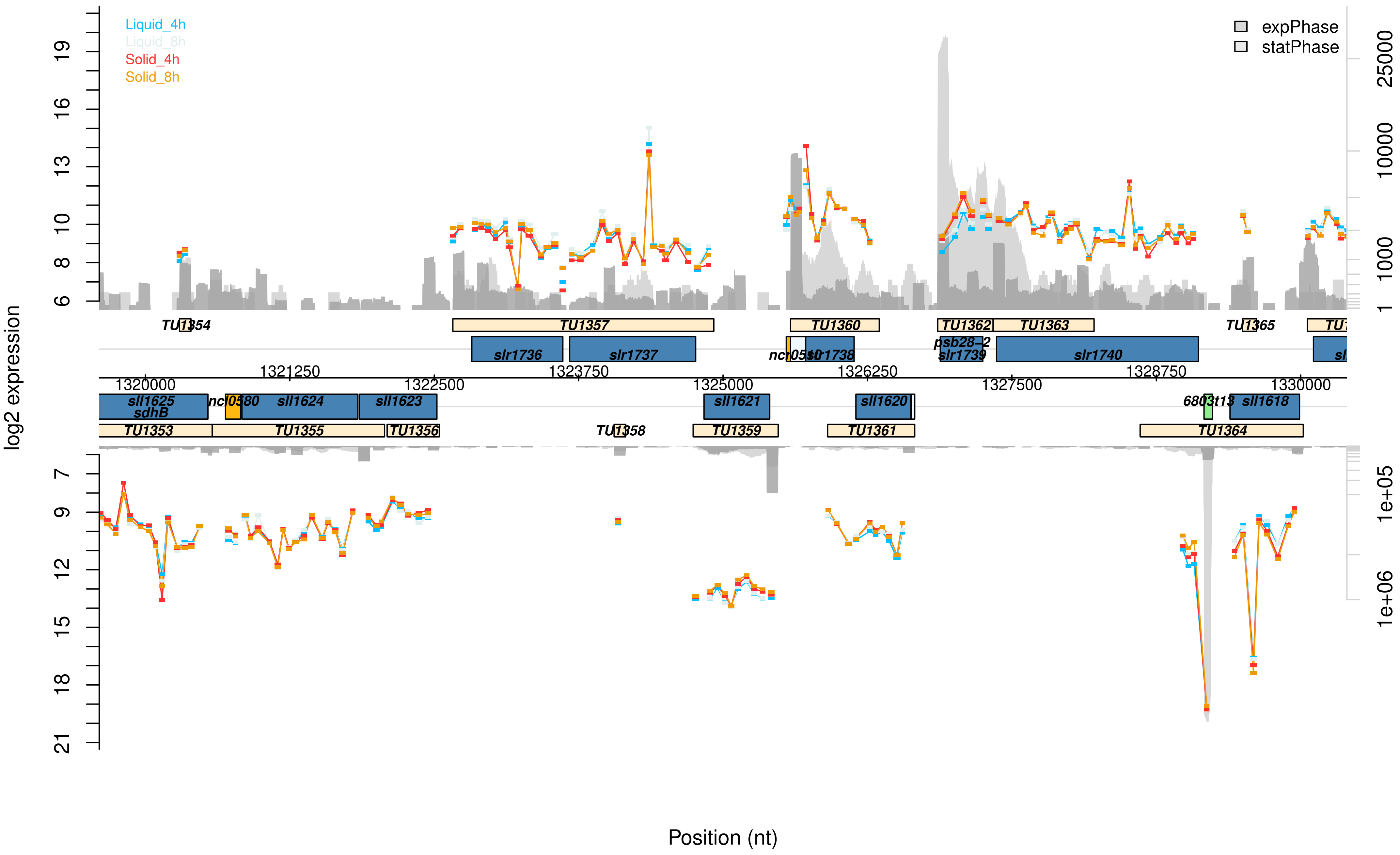

Liquid_4h
Liquid_8h
Solid_4h
Solid_8h
expPhase
statPhase
log2 expression
Position (nt)
TU1353
TU1354
TU1355
TU1356
TU1357
TU1358
TU1359
TU1360
TU1361
TU1362
TU1363
TU1364
TU1365
sll1625
sdhB
ncl0580
sll1624
sll1623
slr1736
slr1737
sll1621
slr1738
sll1620
psb28-2
slr1739
slr1740
sll1618
6803t13
1320000
1321250
1322500
1323750
1325000
1326250
1327500
1328750
1330000

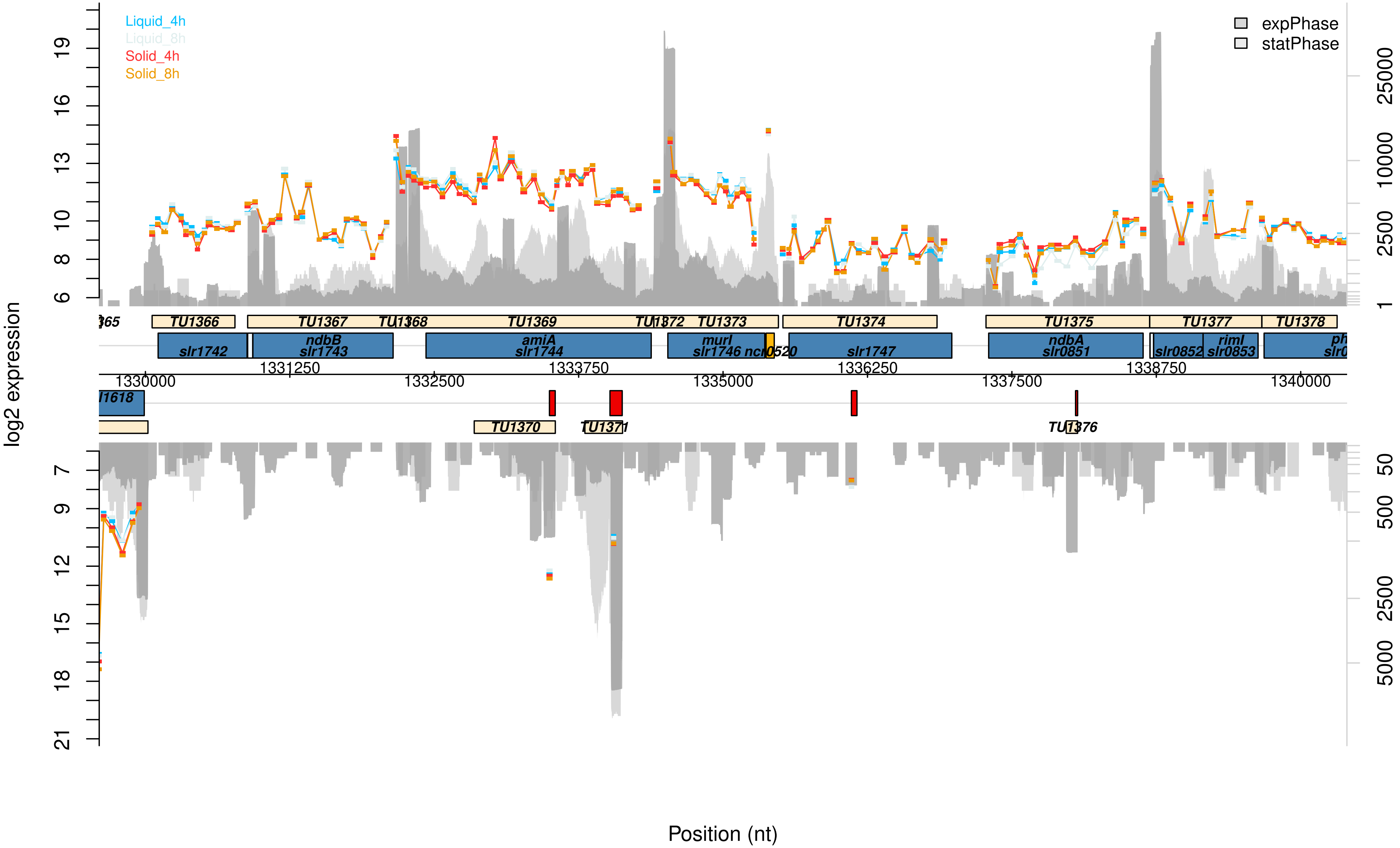

Liquid_4h
Liquid_8h
Solid_4h
Solid_8h
expPhase
statPhase
log2 expression
Position (nt)
TU1366
TU1367
TU1368
TU1369
TU1372
TU1373
TU1374
TU1375
TU1377
TU1378
slr1742
ndbB
slr1743
amiA
slr1744
murI
slr1746
slr1747
ndbA
slr0851
slr0852
rimI
slr0853
1330000
1331250
1332500
1333750
1335000
1336250
1337500
1338750
1340000
TU1370
TU1371
TU1376

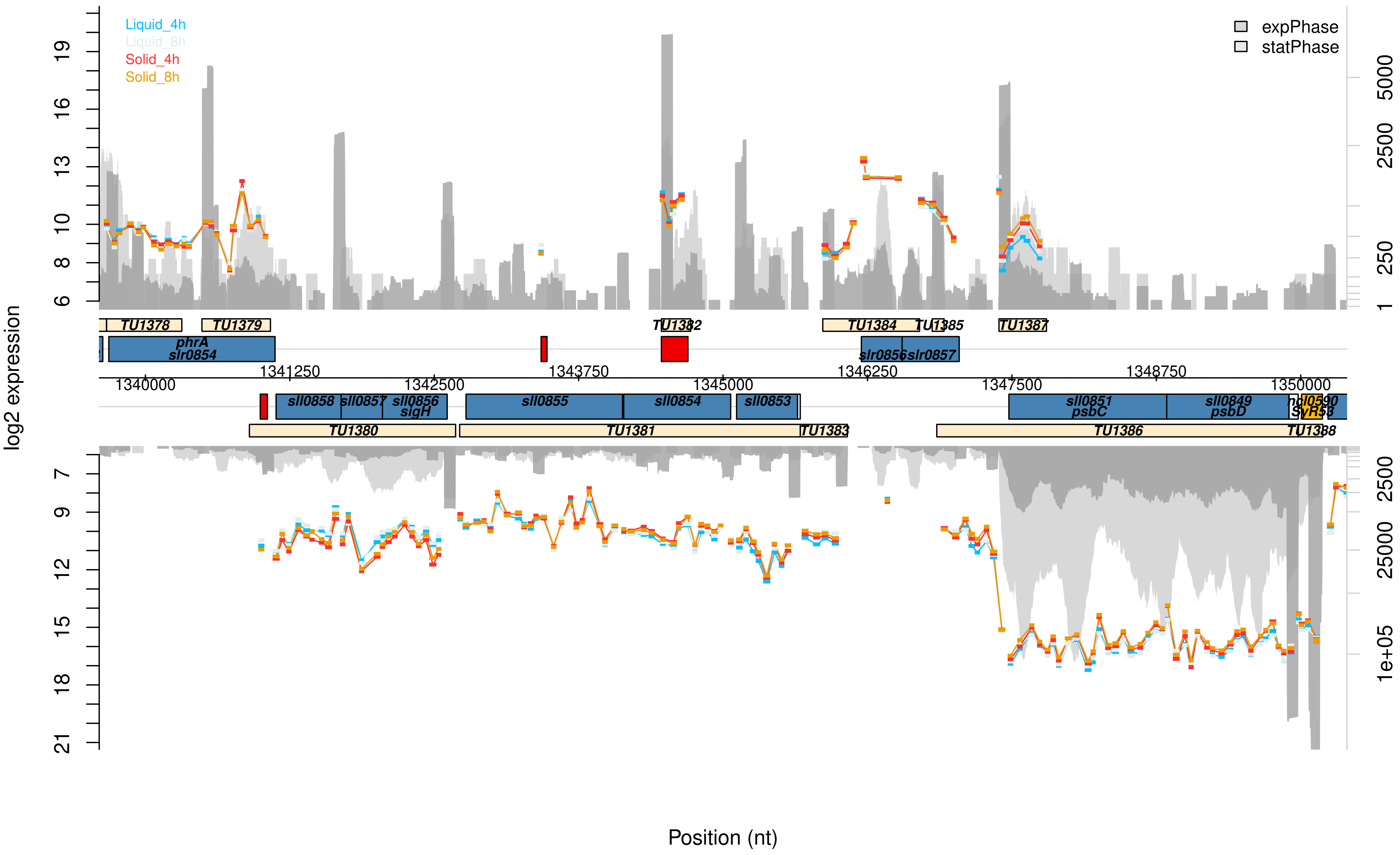

Liquid_4h
Liquid_8h
Solid_4h
Solid_8h
expPhase
statPhase
log2 expression
TU1378
TU1379
TU1382
TU1384
TU1385
TU1387
phrA
slr0854
slr0856
slr0857
1340000
1341250
1342500
1343750
1345000
1346250
1347500
1348750
1350000
sll0858
sll0857
sll0856
sigH
sll0855
sll0854
sll0853
sll0851
psbC
sll0849
psbD
SyR53
TU1380
TU1381
TU1383
TU1386
TU1388
Position (nt)

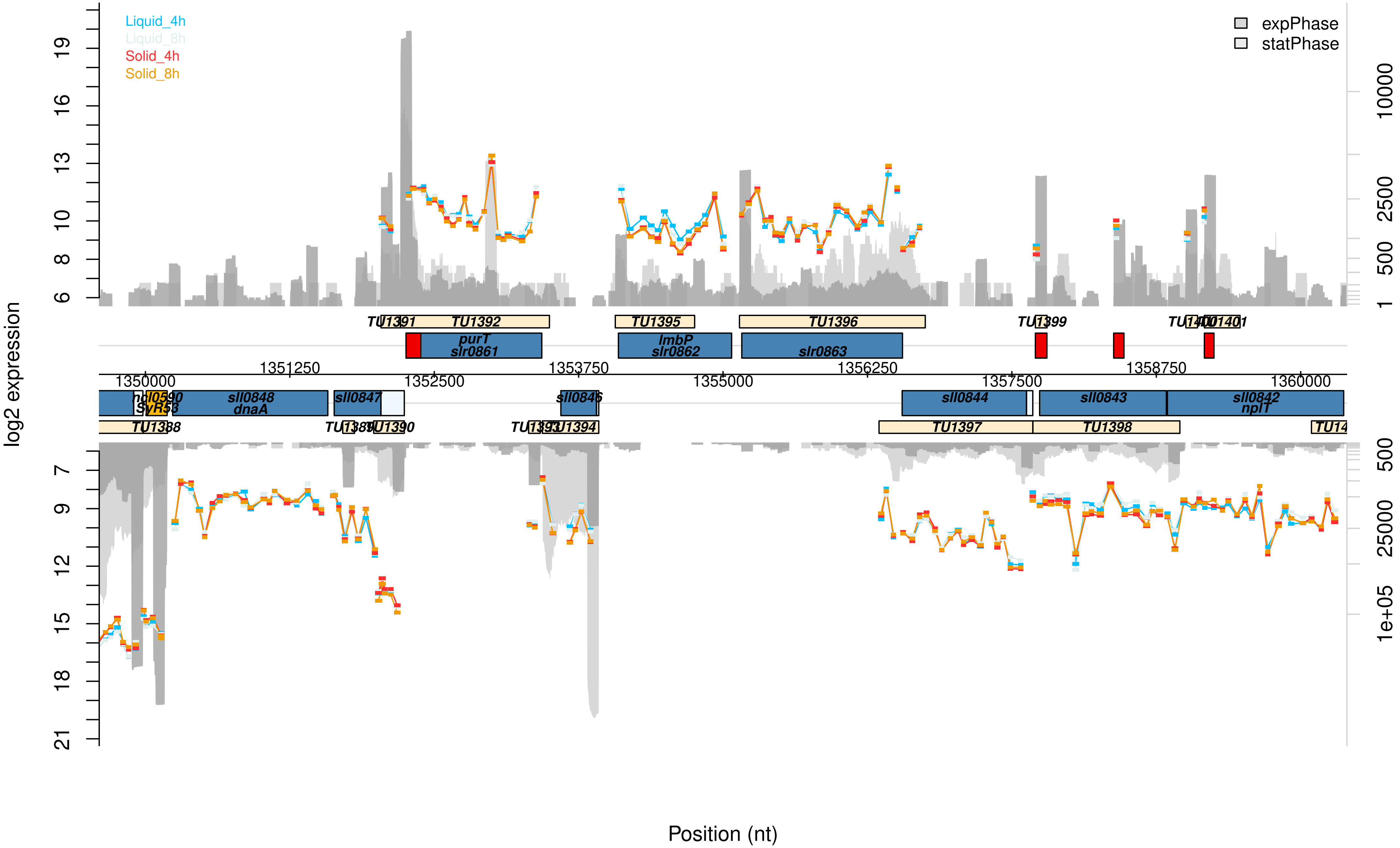

Liquid_4h
Liquid_8h
Solid_4h
Solid_8h
expPhase
statPhase
log2 expression
Position (nt)
TU1391
TU1392
TU1395
TU1396
TU1399
purT
slr0861
lmbP
slr0862
slr0863
1350000
1351250
1352500
1353750
1355000
1356250
1357500
1358750
1360000
dnaA
sll0848
sll0847
sll0846
sll0844
sll0843
sll0842
nplT
SyR53
TU1388
TU1390
TU1394
TU1397
TU1398

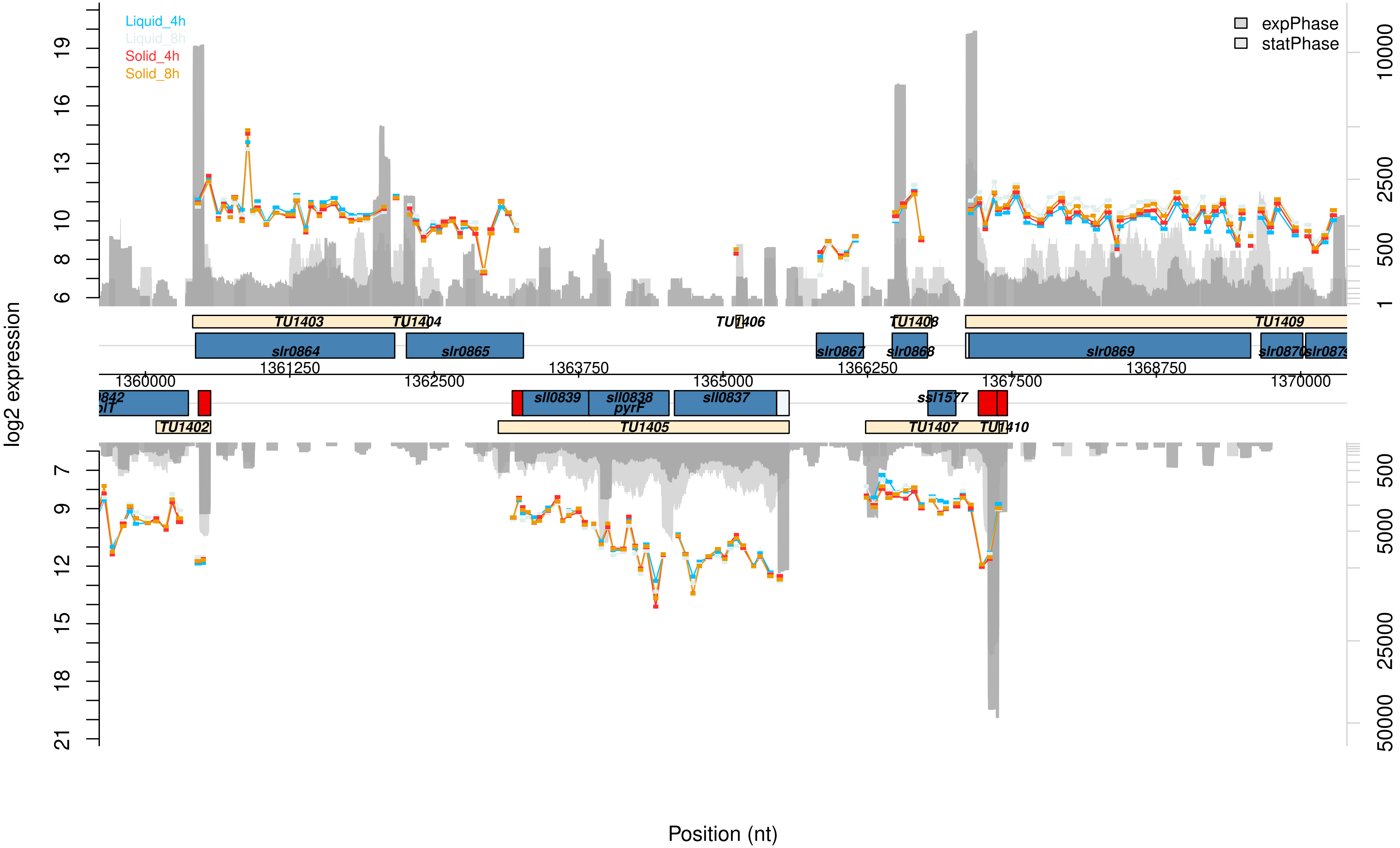

Liquid_4h
Liquid_8h
Solid_4h
Solid_8h
expPhase
statPhase
log2 expression
Position (nt)
TU1403
TU1404
TU1406
TU1408
TU1409
slr0864
slr0865
slr0867
slr0868
slr0869
slr0870
1360000
1361250
1362500
1363750
1365000
1366250
1367500
1368750
1370000
sll0839
sll0838
pyrF
sll0837
ssl1577
TU1402
TU1405
TU1407
TU1410

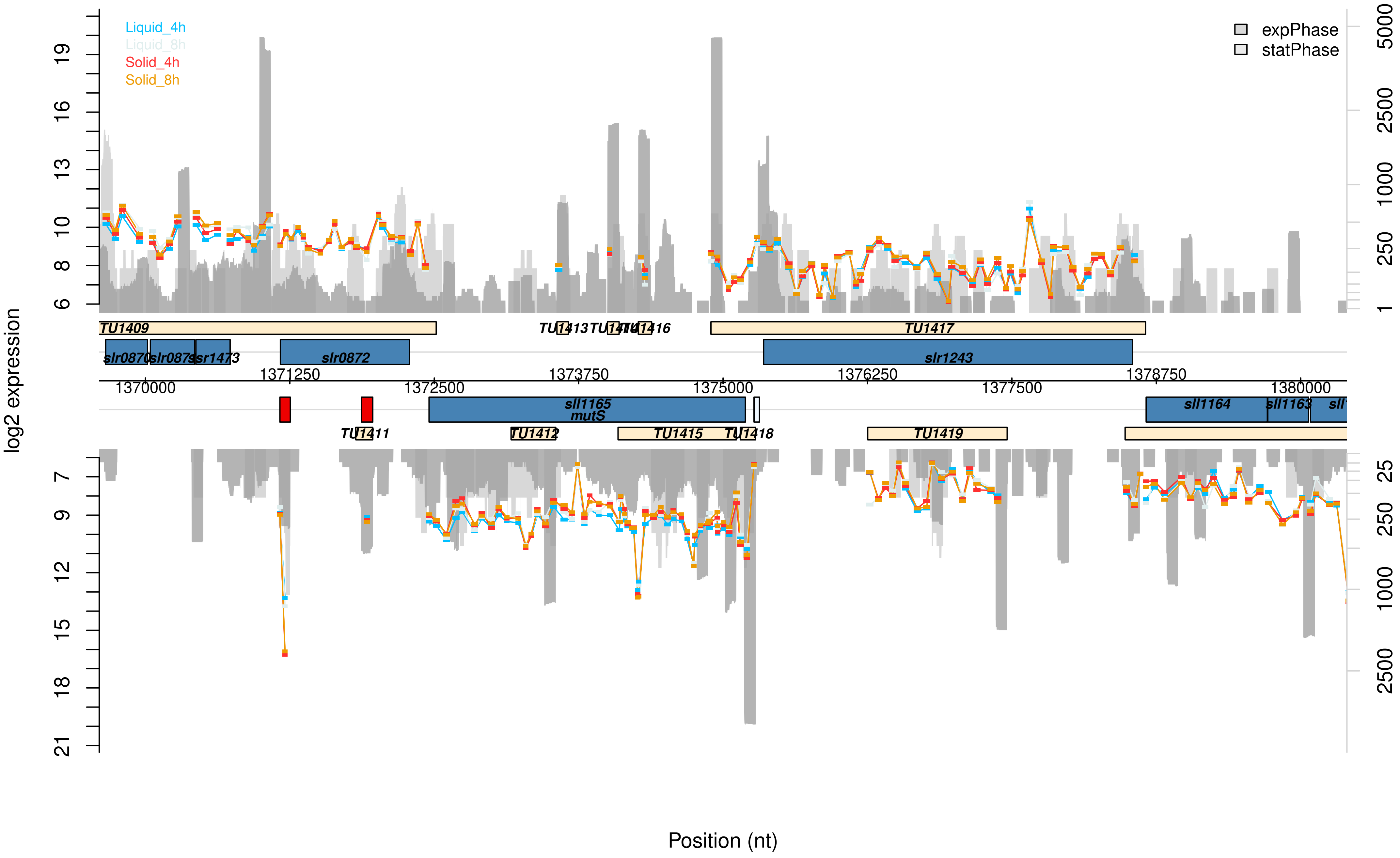

Liquid_4h
Liquid_8h
Solid_4h
Solid_8h
expPhase
statPhase
log2 expression
Position (nt)
TU1409
TU1413
TU1416
TU1417
slr0870
slr0873
ssr1473
slr0872
slr1243
1371250
1373750
1376250
1378750
1370000
1372500
1375000
1377500
1380000
sll1165
mutS
sll1164
sll1163
TU1411
TU1412
TU1415
TU1418
TU1419

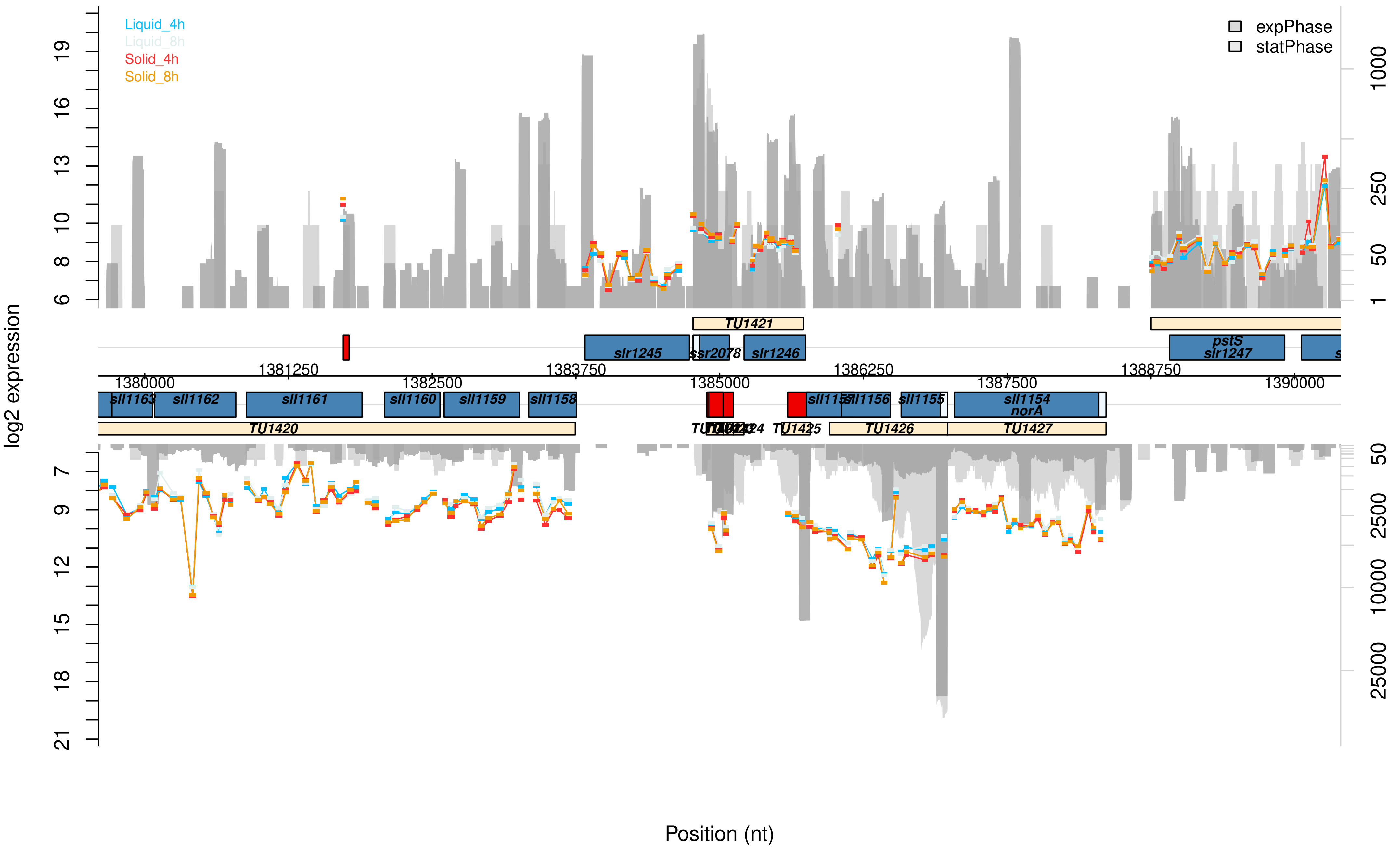

Liquid_4h
Liquid_8h
Solid_4h
Solid_8h
expPhase
statPhase
log2 expression
Position (nt)
TU1420
TU1421
TU1425
TU1426
TU1427
slr1245
ssr2078
slr1246
pstS
slr1247
sll1163
sll1162
sll1161
sll1160
sll1159
sll1158
sll1156
sll1155
sll1154
norA
1380000
1381250
1382500
1383750
1385000
1386250
1387500
1388750
1390000

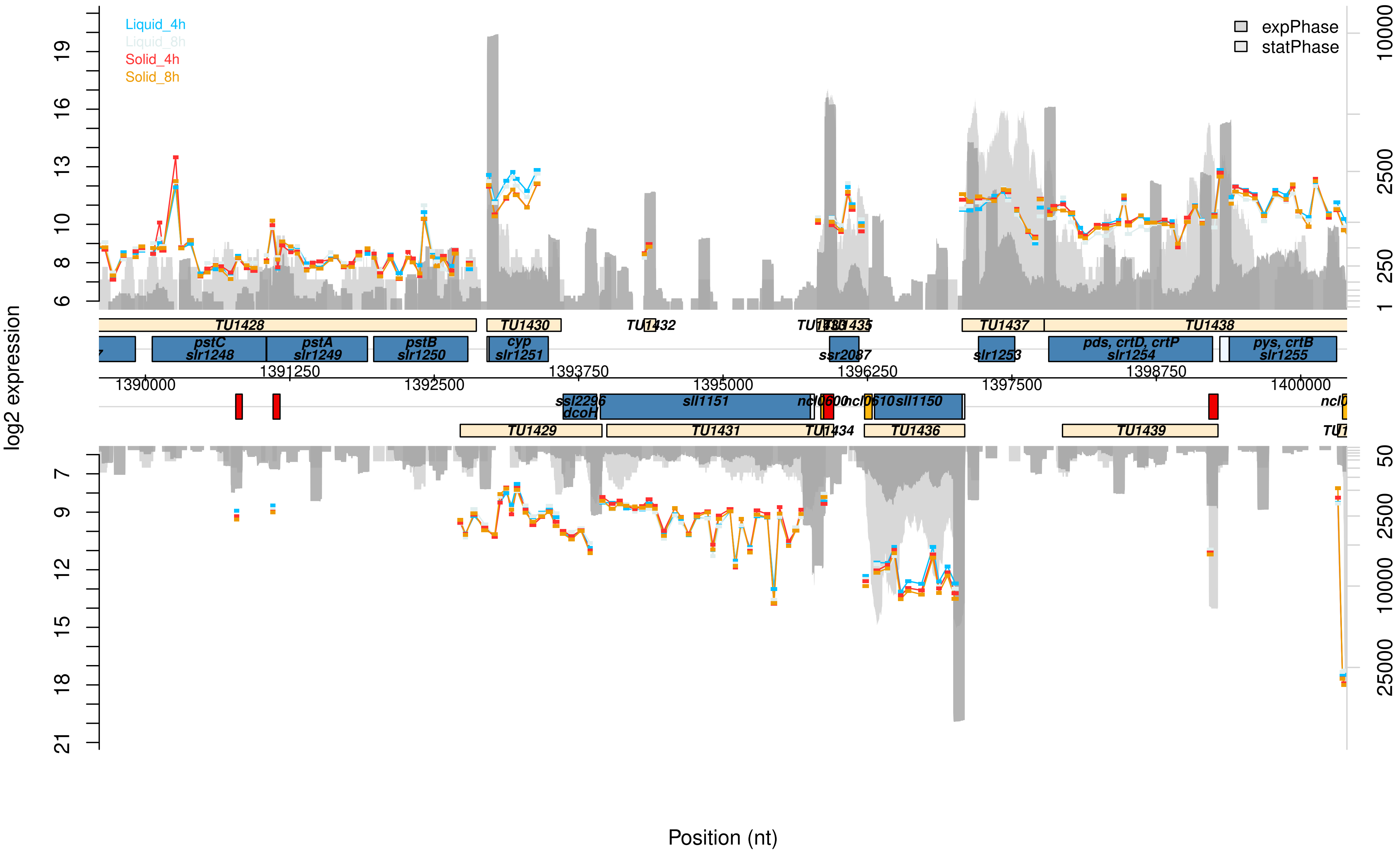
Liquid_4h
Liquid_8h
Solid_4h
Solid_8h
expPhase
statPhase
log2 expression
Position (nt)
TU1428
TU1430
TU1432
TU1437
TU1438
pstC slr1248
pstA slr1249
pstB slr1250
cyp slr1251
ssr2087
slr1253
pds, crtD, crtP slr1254
pys, crtB slr1255
1390000
1391250
1392500
1393750
1395000
1396250
1397500
1398750
1400000
ssl2296 dcoH
sll1151
sll1150
TU1429
TU1431
TU1434
TU1436
TU1439

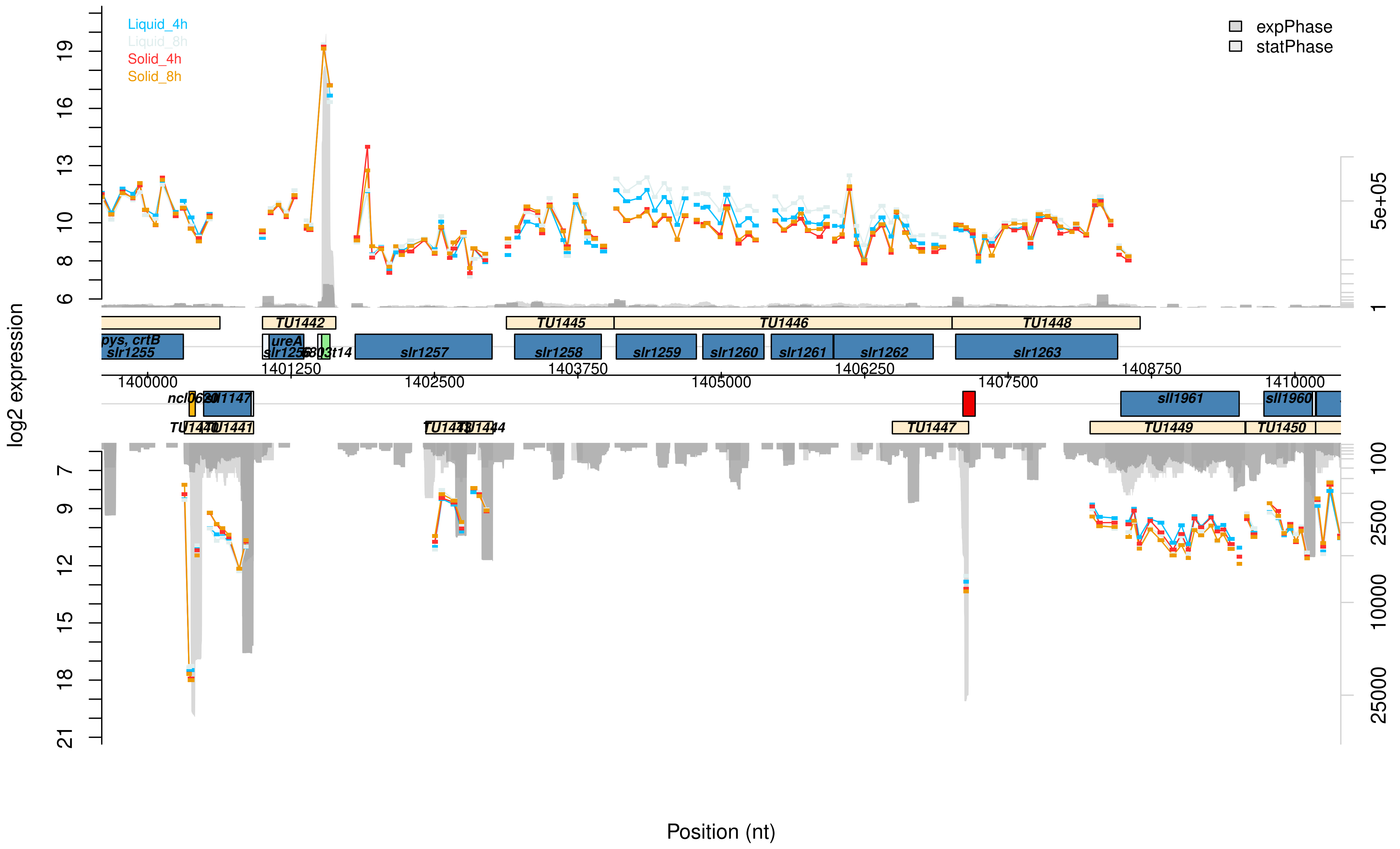

Liquid_4h
Liquid_8h
Solid_4h
Solid_8h
expPhase
statPhase
log2 expression
Position (nt)
TU1442
TU1445
TU1446
TU1448
pys, crtB
slr1255
ureA
slr1256
slr1257
slr1258
slr1259
slr1260
slr1261
slr1262
slr1263
1400000
1401250
1402500
1403750
1405000
1406250
1407500
1408750
1410000
sll1147
sll1961
sll1960
TU1441
TU1447
TU1449
TU1450
5e+05

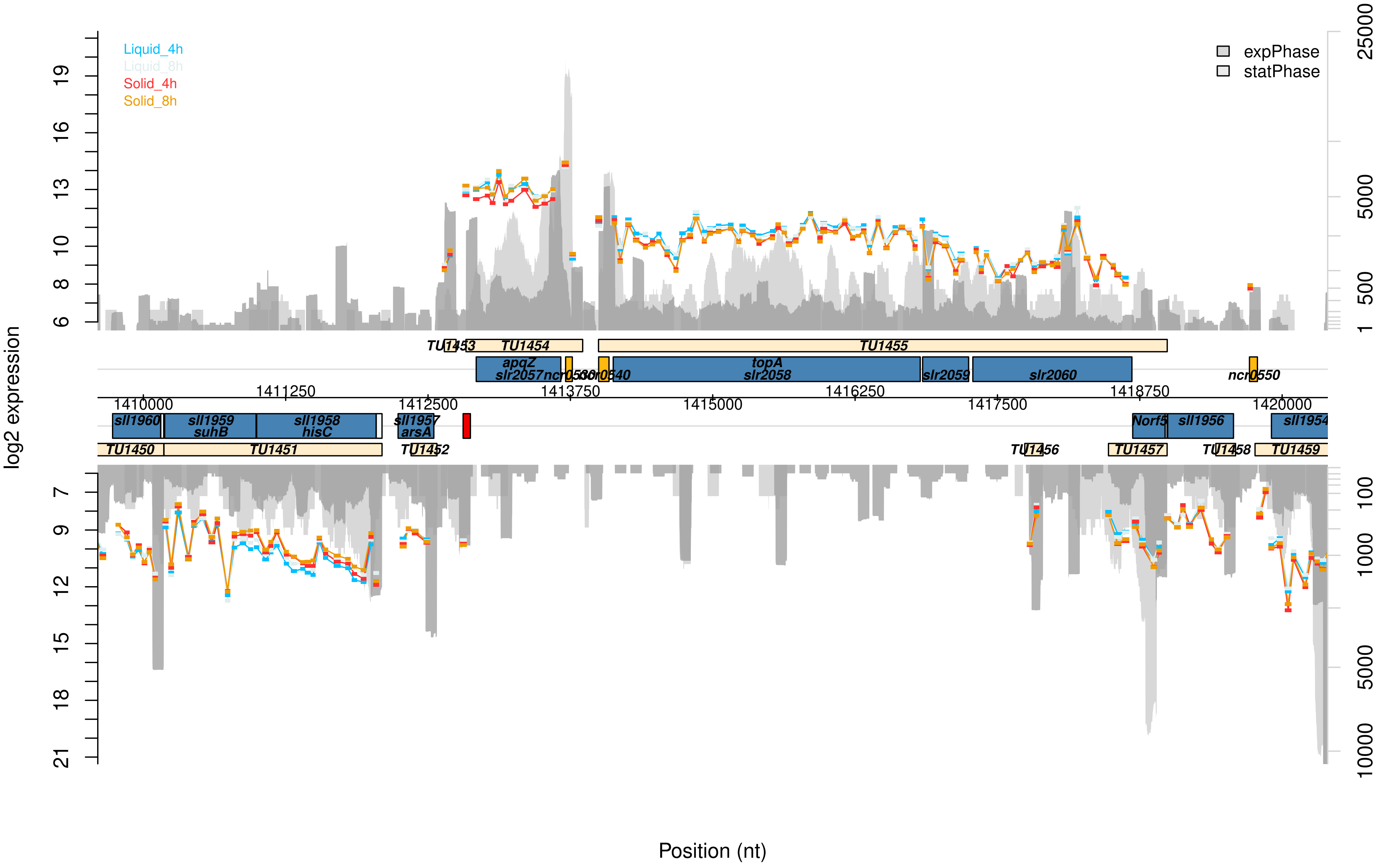

Liquid_4h
Liquid_8h
Solid_4h
Solid_8h
expPhase
statPhase
log2 expression
Position (nt)
TU1453
TU1454
TU1455
apqZ
slr2057
ncr0530
ncr0540
topA
slr2058
slr2059
slr2060
ncr0550
1410000
1411250
1412500
1413750
1415000
1416250
1417500
1418750
1420000
sll1960
sll1959
suhB
sll1958
hisC
sll1957
arsA
Norf5
sll1956
sll1954
TU1450
TU1451
TU1452
TU1456
TU1457
TU1458
TU1459

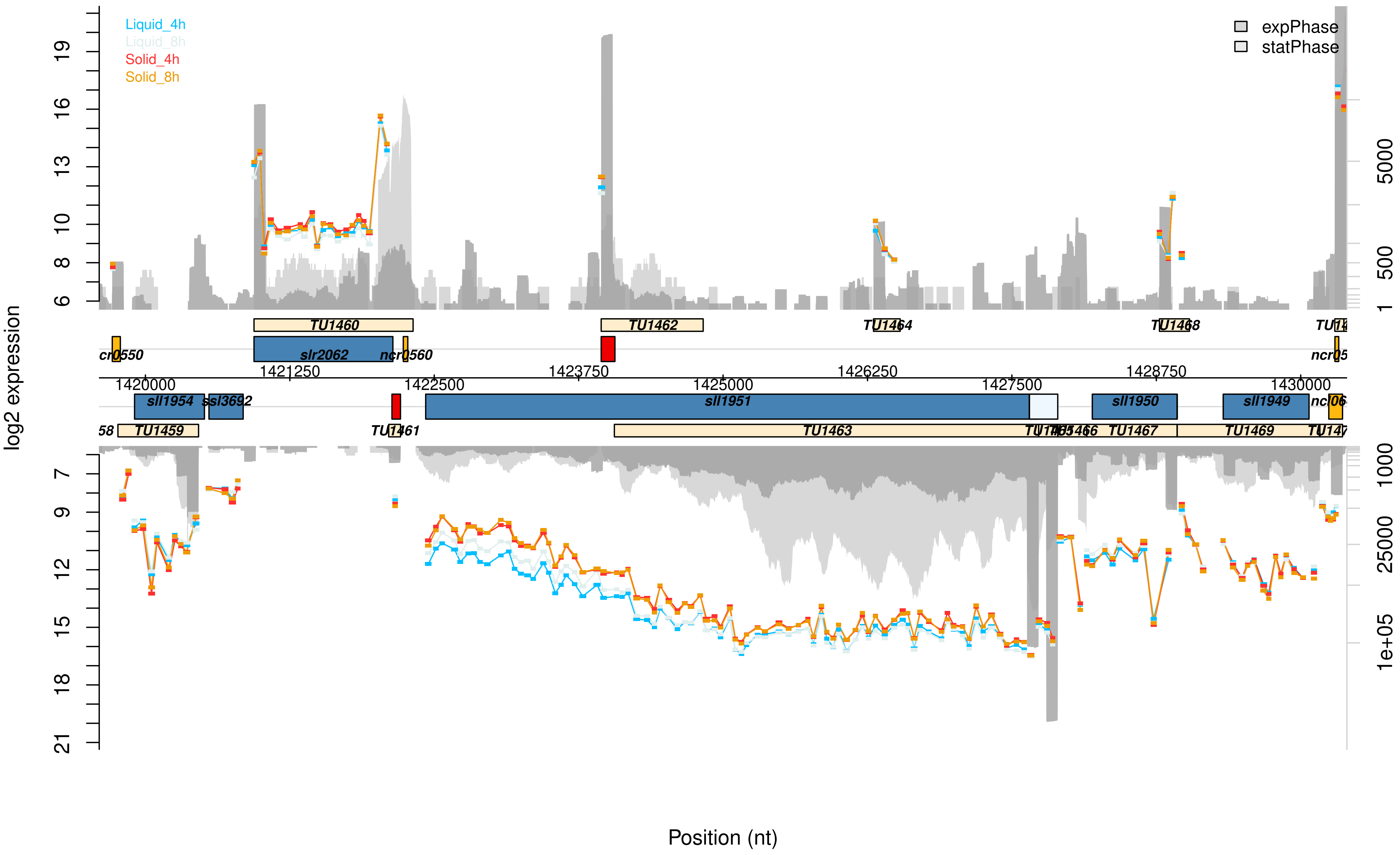

Liquid_4h
Liquid_8h
Solid_4h
Solid_8h
expPhase
statPhase
log2 expression
TU1460
TU1462
TU1464
TU1468
slr2062
ncr0560
1421250
1423750
1426250
1428750
1420000
1422500
1425000
1427500
1430000
sll1954
ssl3692
sll1951
sll1950
sll1949
TU1459
TU1461
TU1463
TU1467
TU1469
Position (nt)

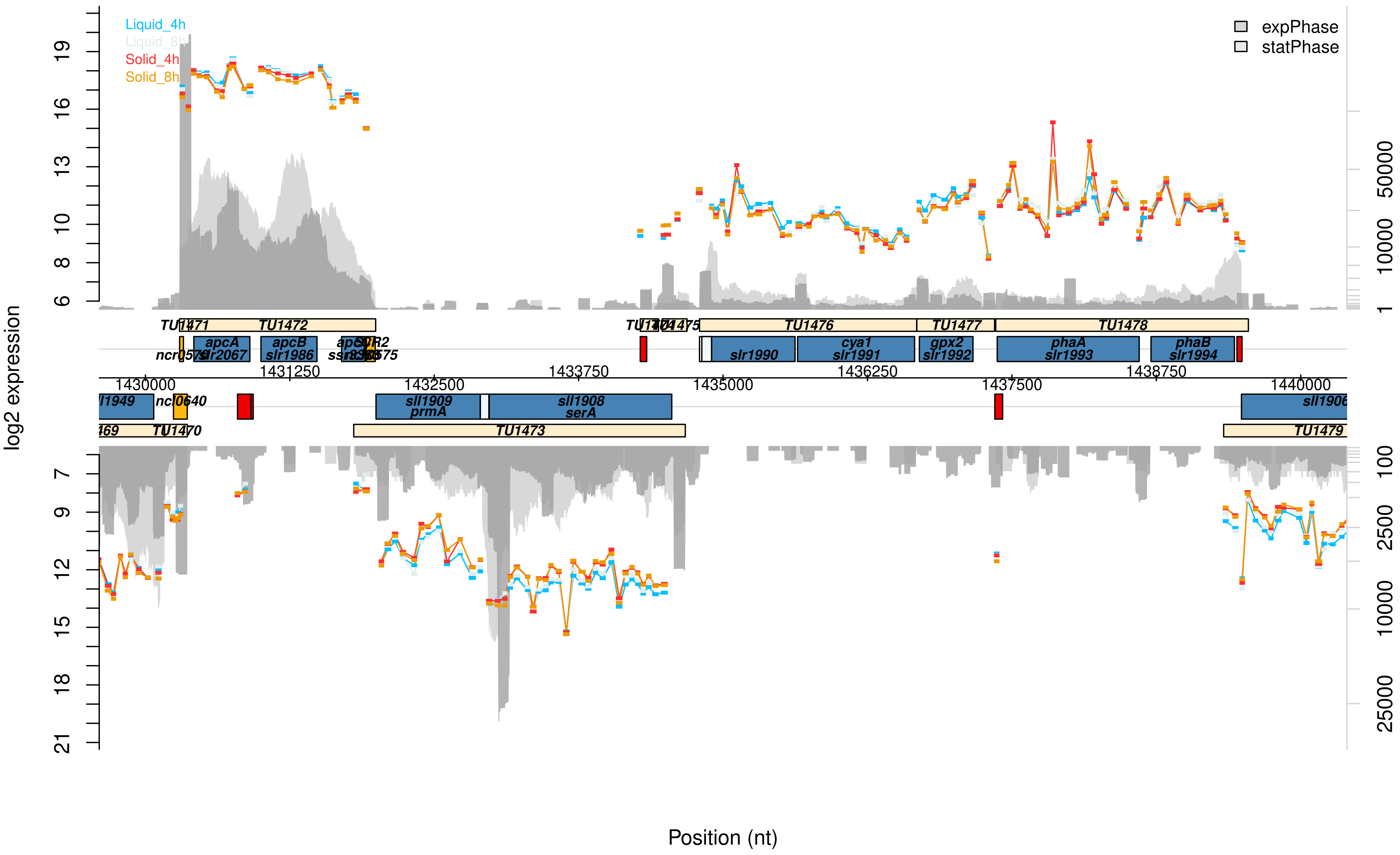
Liquid_4h
Liquid_8h
Solid_4h
Solid_8h
expPhase
statPhase
log2 expression
Position (nt)
TU1471
TU1472
apcA
slr2067
apcB
slr1986
TU1476
TU1477
TU1478
slr1990
cya1
slr1991
gpx2
slr1992
phaA
slr1993
phaB
slr1994
sll1909
prmA
sll1908
serA
TU1473
TU1470
sll1906
TU1479
1430000
1431250
1432500
1433750
1435000
1436250
1437500
1438750
1440000

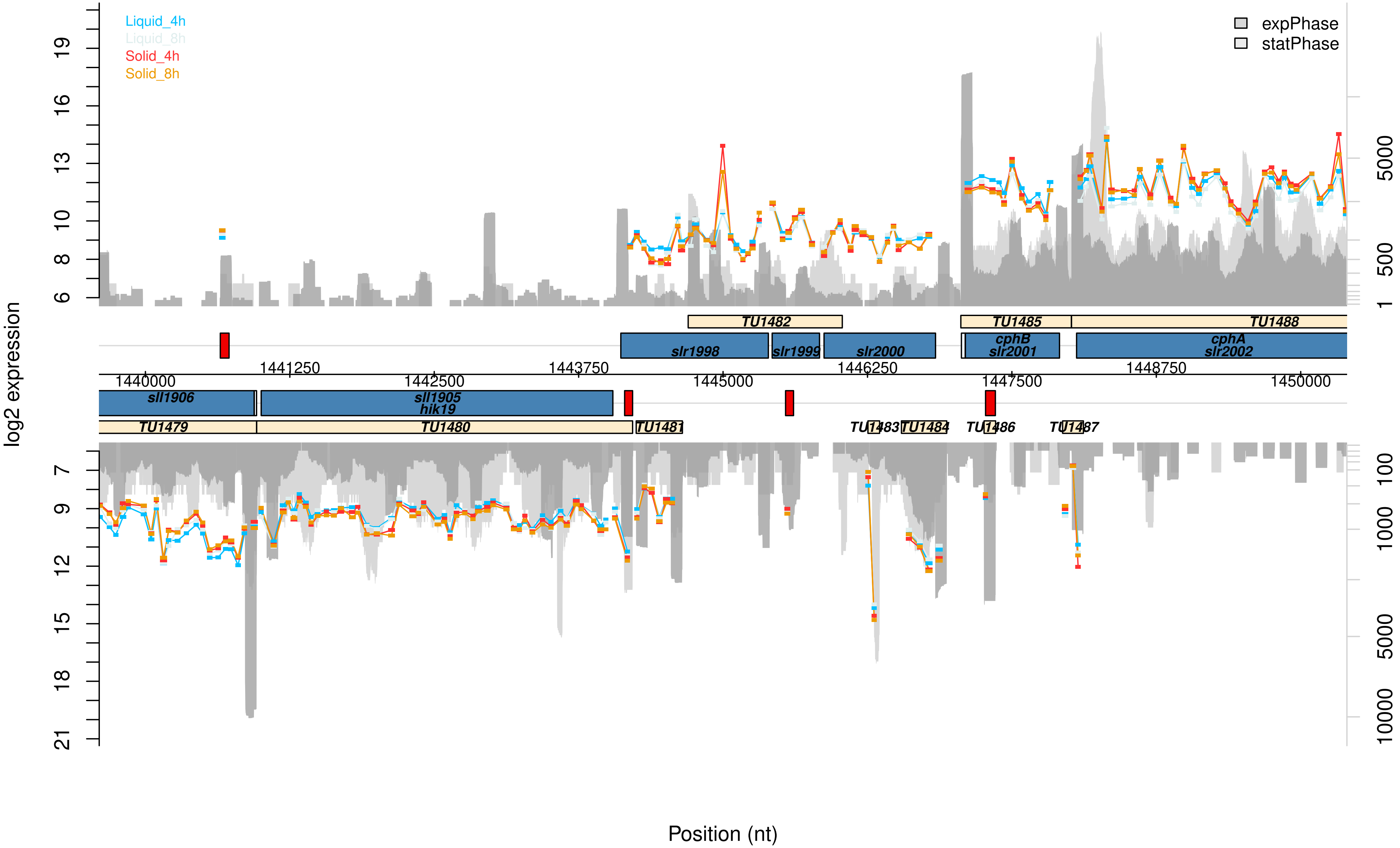

Liquid_4h
Liquid_8h
Solid_4h
Solid_8h
expPhase
statPhase
log2 expression
TU1482
TU1485
TU1488
slr1998
slr1999
slr2000
cphB
slr2001
cphA
slr2002
1441250
1443750
1446250
1448750
1440000
1442500
1445000
1447500
1450000
sll1906
sll1905
hik19
TU1479
TU1480
TU1481
TU1483
TU1484
TU1486
TU1487
Position (nt)

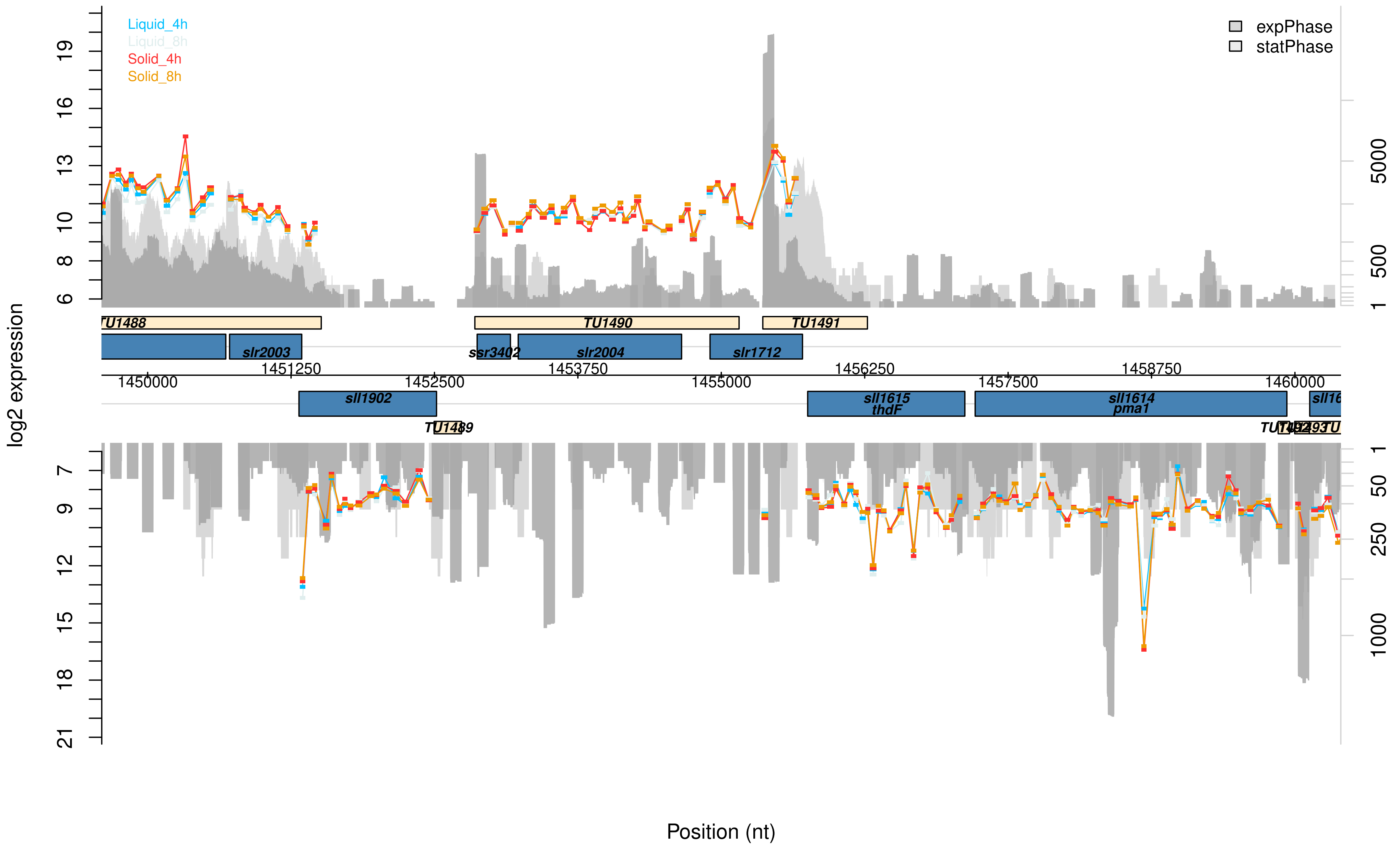

Liquid_4h
Liquid_8h
Solid_4h
Solid_8h
expPhase
statPhase
log2 expression
TU1488
TU1490
TU1491
slr2003
ssr3402
slr2004
slr1712
1451250
1453750
1456250
1458750
1450000
1452500
1455000
1457500
1460000
sll1902
sll1615
thdF
sll1614
pma1
TU1489
Position (nt)

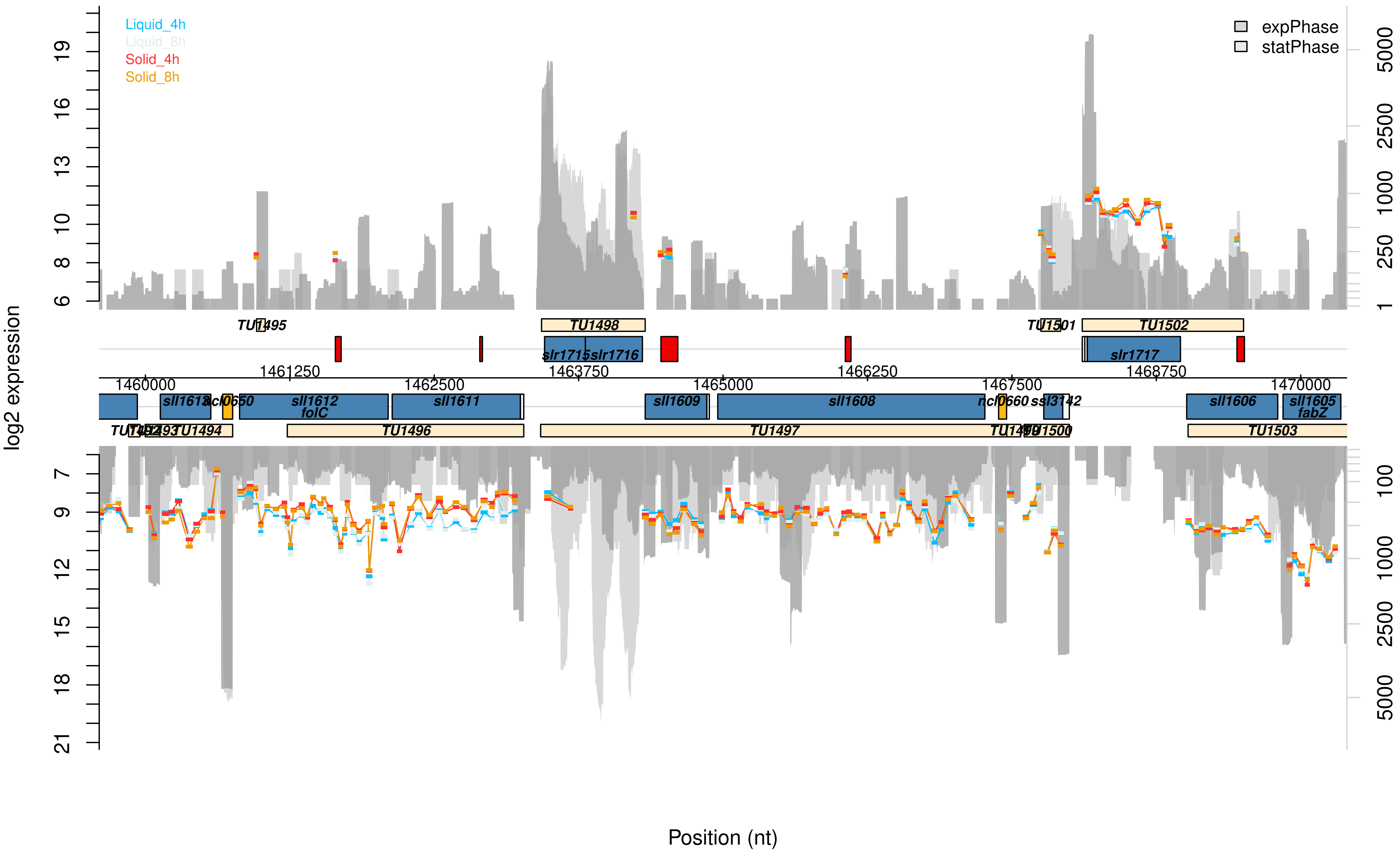

Liquid_4h
Liquid_8h
Solid_4h
Solid_8h
expPhase
statPhase
log2 expression
Position (nt)
TU1495
TU1498
TU1501
TU1502
slr1715
slr1716
slr1717
1460000
1461250
1462500
1463750
1465000
1466250
1467500
1468750
1470000
sll1613
ncl0650
sll1612
folC
sll1611
sll1609
sll1608
ncl0660
ssl3142
sll1606
sll1605
fabZ
TU1494
TU1496
TU1497
TU1500
TU1503

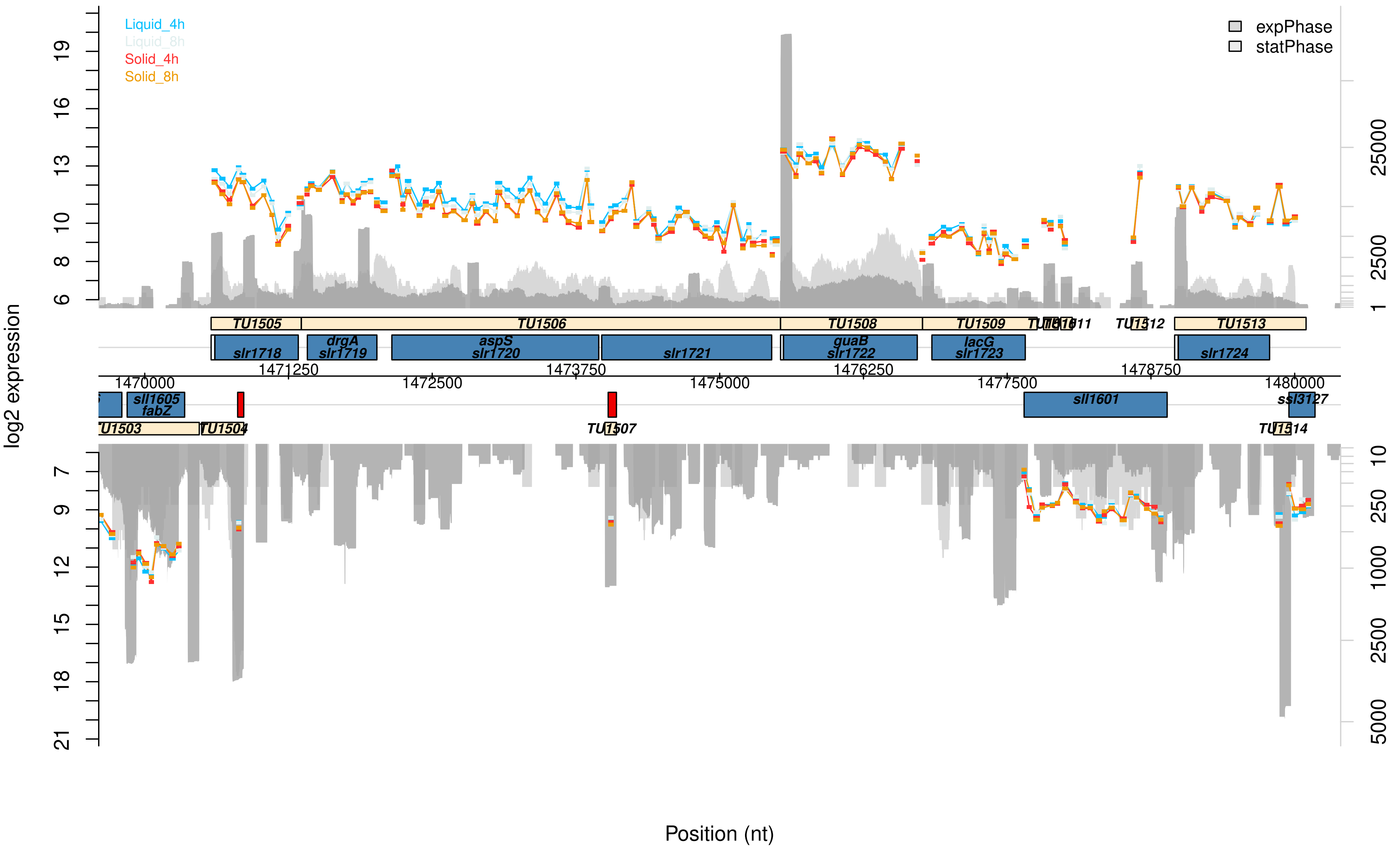

Liquid_4h
Liquid_8h
Solid_4h
Solid_8h
expPhase
statPhase
log2 expression
19
16
13
10
8
6
7
9
12
15
18
21
25000
2500
1
10
250
1000
2500
5000
TU1505
TU1506
TU1508
TU1509
TU1511
TU1512
TU1513
slr1718
drgA
slr1719
aspS
slr1720
slr1721
guaB
slr1722
lacG
slr1723
slr1724
1470000
1471250
1472500
1473750
1475000
1476250
1477500
1478750
1480000
sll1605
fabZ
sll1601
ssl3127
TU1503
TU1504
TU1507
TU1514
Position (nt)

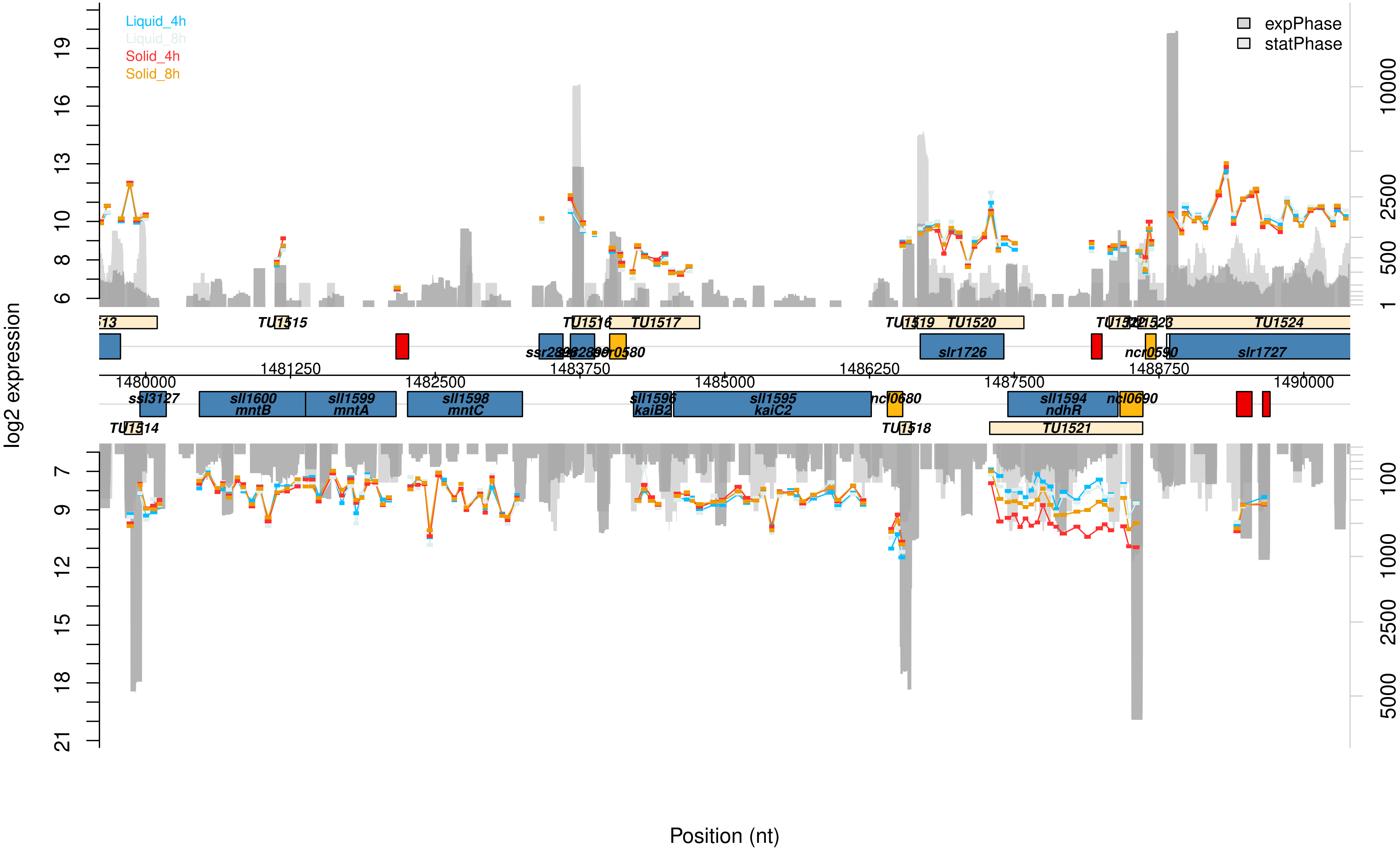

Liquid_4h
Liquid_8h
Solid_4h
Solid_8h
expPhase
statPhase
log2 expression
Position (nt)
TU1515
TU1516
TU1517
TU1519
TU1520
TU1524
ssr2848
ncr0580
slr1726
ncr0590
slr1727
1481250
1483750
1486250
1488750
1480000
1482500
1485000
1487500
1490000
ssl3127
sll1600
mntB
sll1599
mntA
sll1598
mntC
sll1596
kaiB2
sll1595
kaiC2
ncl0680
sll1594
ndhR
ncl0690
TU1514
TU1518
TU1521

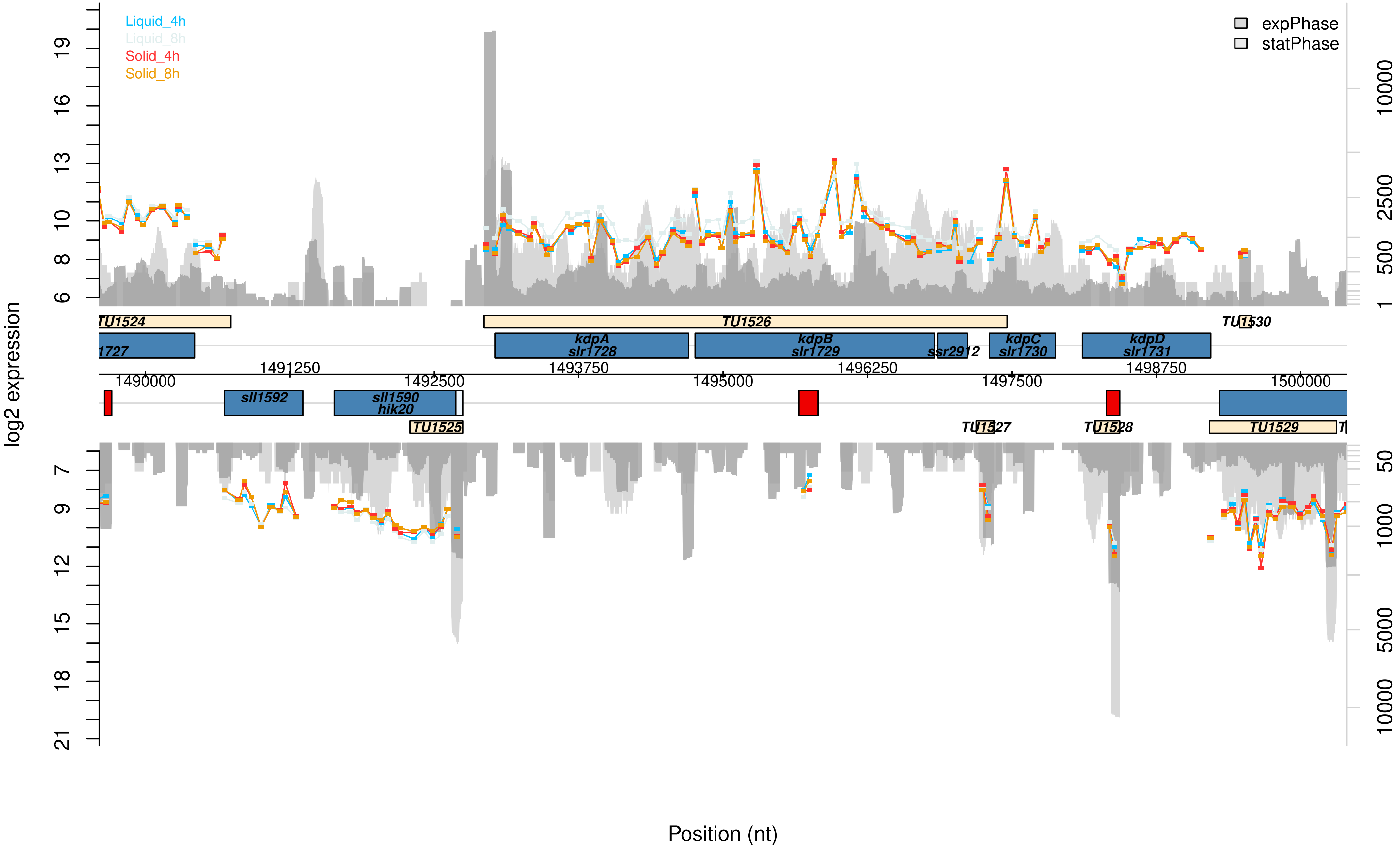

Liquid_4h
Liquid_8h
Solid_4h
Solid_8h
expPhase
statPhase
log2 expression
Position (nt)
TU1524
TU1525
TU1526
TU1527
TU1528
TU1529
TU1530
kdpA
slr1728
kdpB
slr1729
ssr2912
kdpC
slr1730
kdpD
slr1731
sll1592
sll1590
hik20
1727
1490000
1491250
1492500
1493750
1495000
1496250
1497500
1498750
1500000

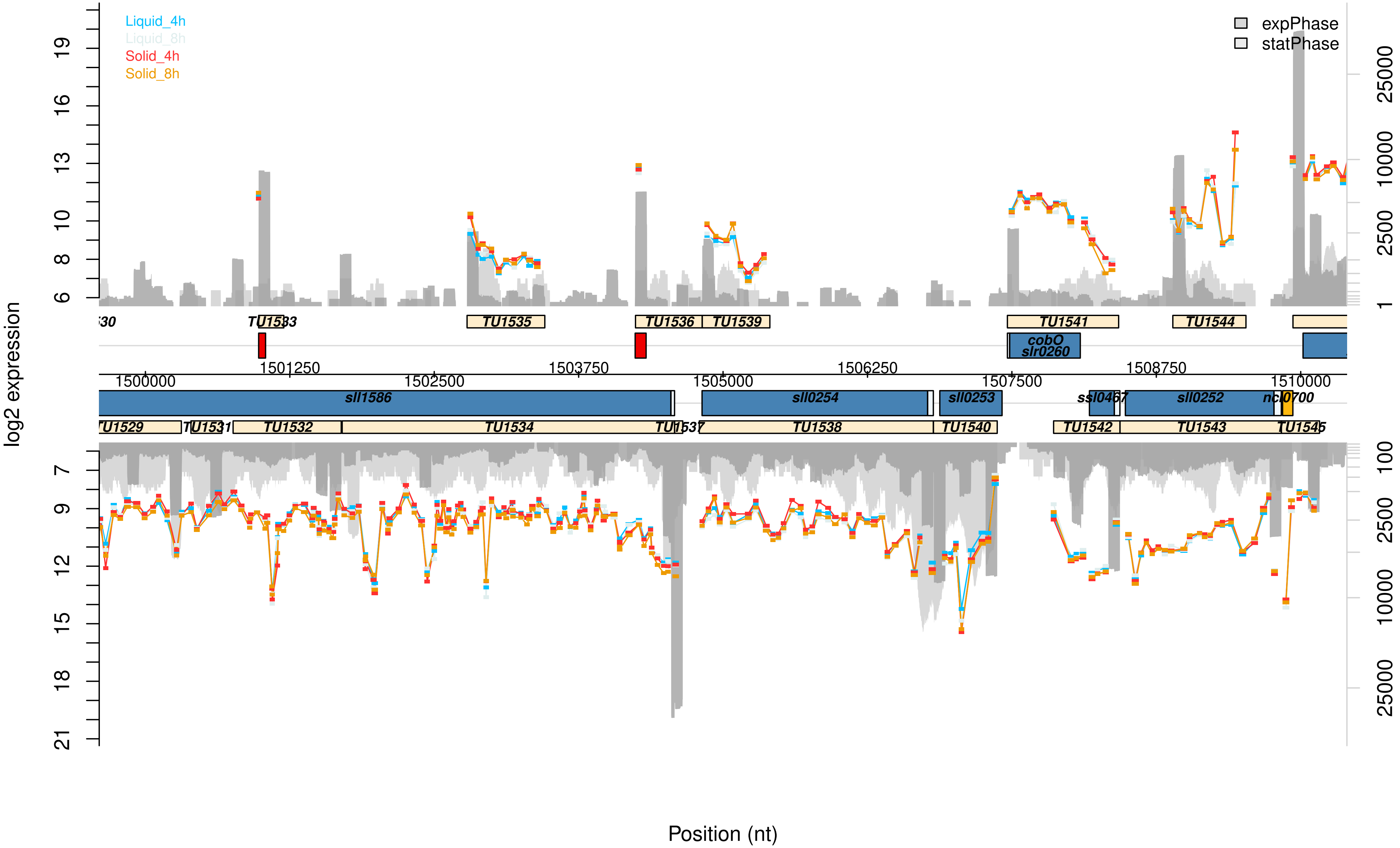

Liquid_4h
Liquid_8h
Solid_4h
Solid_8h
expPhase
statPhase
log2 expression
TU1533
TU1535
TU1536
TU1539
TU1541
TU1544
cobO
slr0260
1501250
1503750
1506250
1508750
1500000
1502500
1505000
1507500
1510000
sll1586
sll0254
sll0253
ssl0467
sll0252
ncl0700
TU1529
TU1531
TU1532
TU1534
TU1537
TU1538
TU1540
TU1542
TU1543
TU1545
Position (nt)

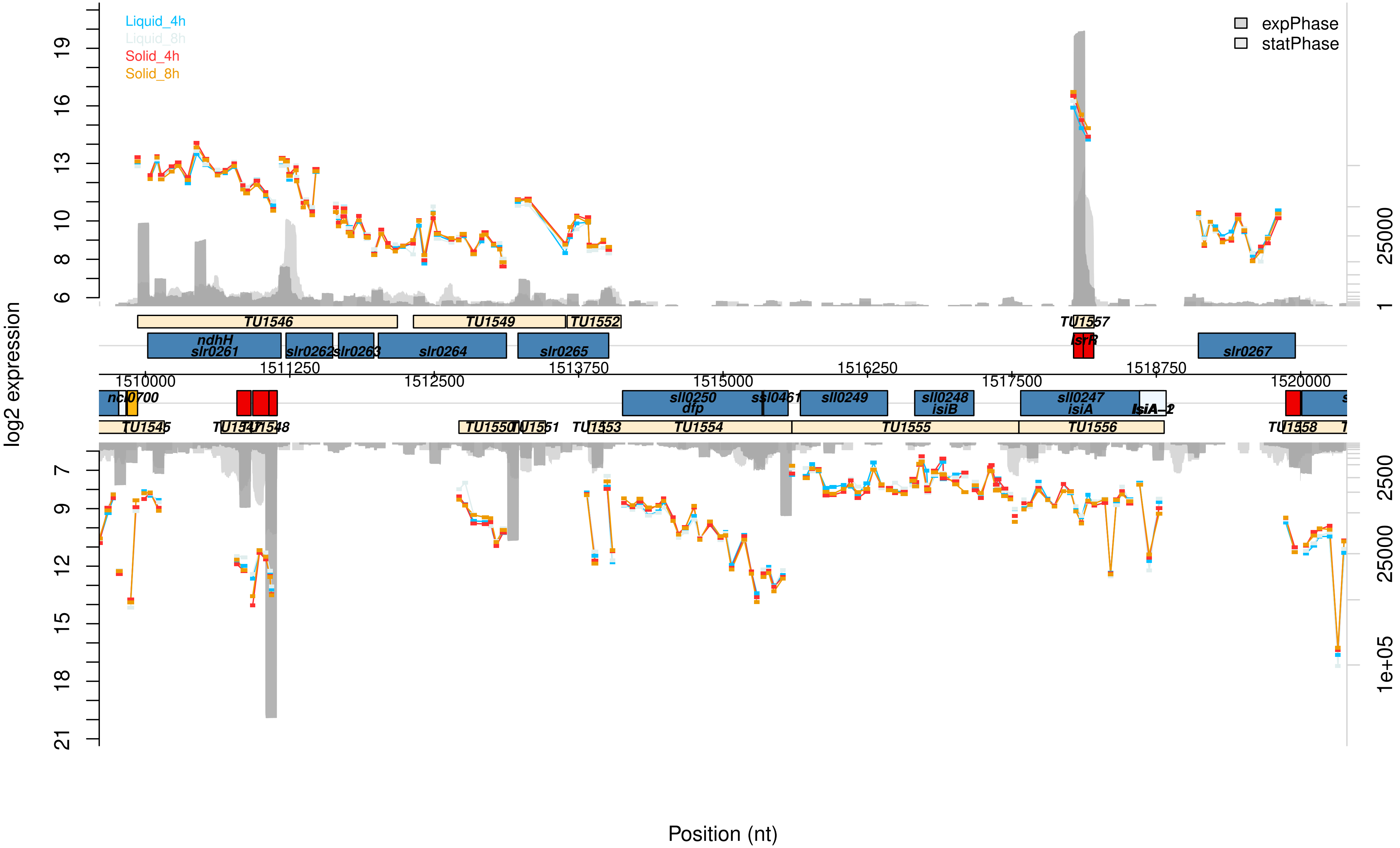

Liquid_4h
Liquid_8h
Solid_4h
Solid_8h
expPhase
statPhase
log2 expression
Position (nt)
TU1546
TU1549
TU1552
TU1557
ndhH
slr0261
slr0262
slr0263
slr0264
slr0265
IsrR
slr0267
1510000
1511250
1512500
1513750
1515000
1516250
1517500
1518750
1520000
sll0250
dfp
ssl0461
sll0249
sll0248
isiB
sll0247
isiA
IsiA-2
TU1545
TU1547
TU1548
TU1550
TU1551
TU1553
TU1554
TU1555
TU1556
TU1558

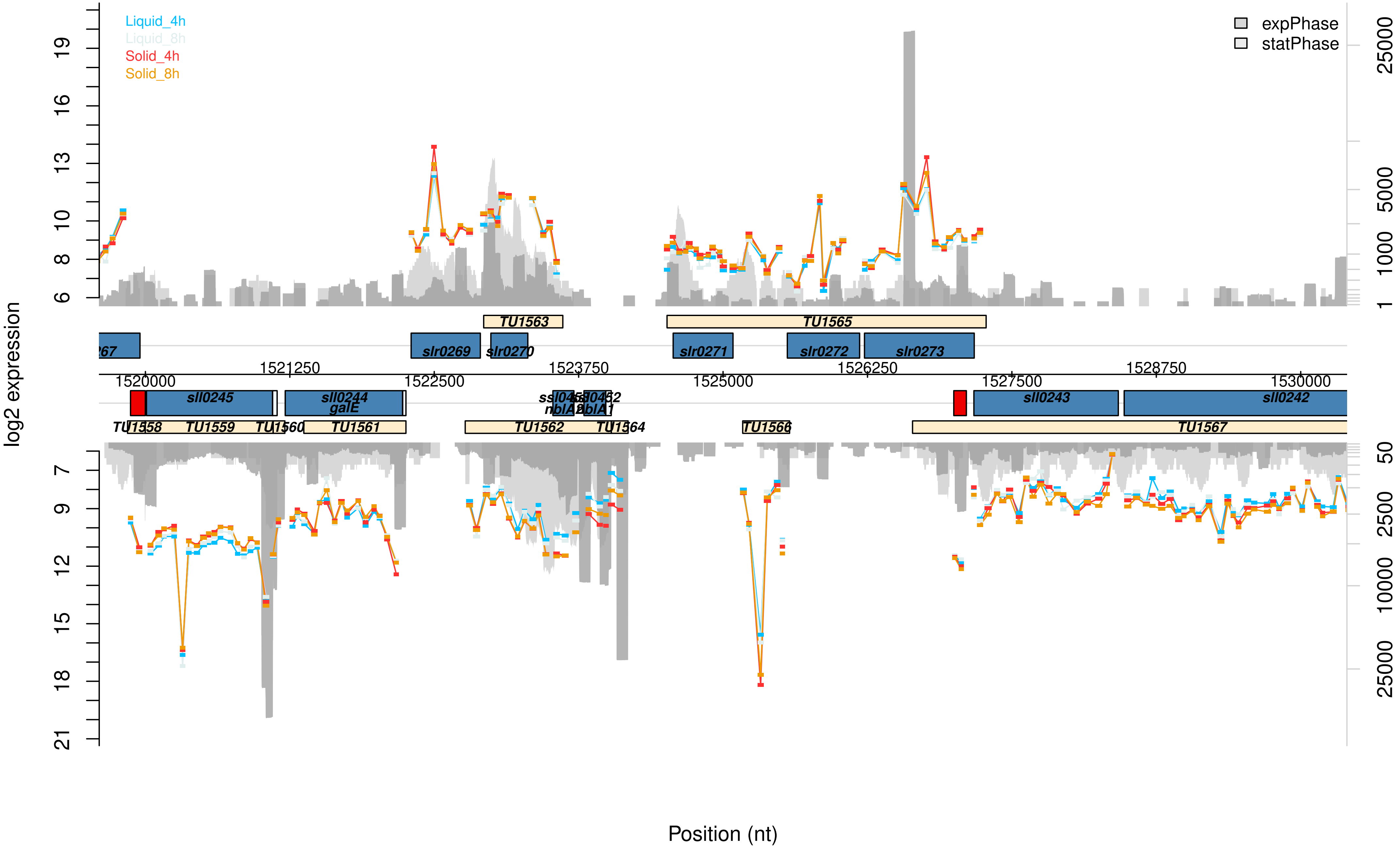

Liquid_4h
Liquid_8h
Solid_4h
Solid_8h
expPhase
statPhase
log2 expression
Position (nt)
TU1563
TU1565
slr0269
slr0270
slr0271
slr0272
slr0273
sll0245
sll0244
galE
sll0243
sll0242
TU1559
TU1561
TU1562
TU1564
TU1566
TU1567
1520000
1521250
1522500
1523750
1525000
1526250
1527500
1528750
1530000

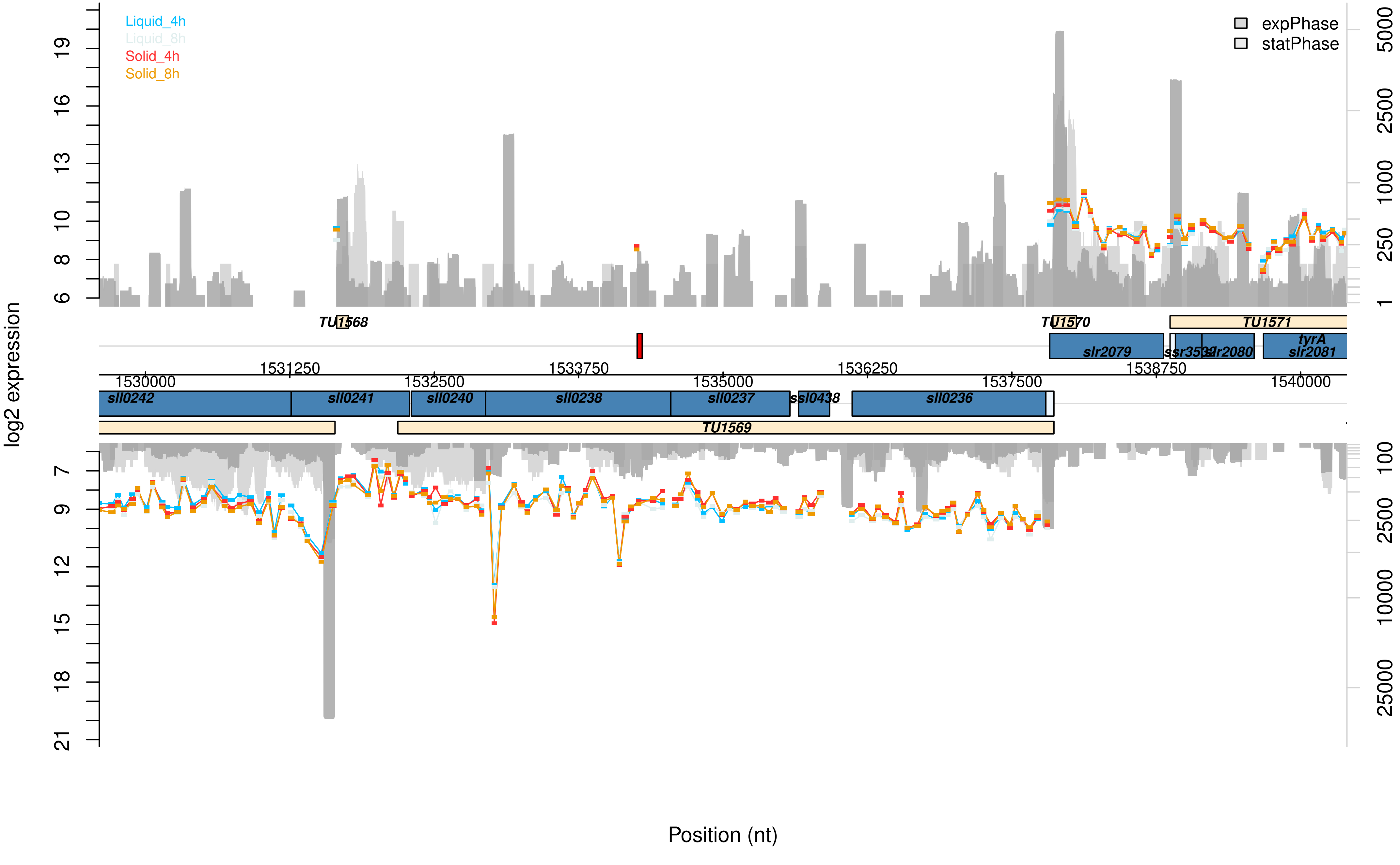
Liquid_4h
Liquid_8h
Solid_4h
Solid_8h
expPhase
statPhase
log2 expression
Position (nt)
TU1568
TU1570
TU1571
TU1569
slr2079
ssr3532
slr2080
tyrA
slr2081
sll0242
sll0241
sll0240
sll0238
sll0237
ssl0438
sll0236
1530000
1531250
1532500
1533750
1535000
1536250
1537500
1538750
1540000

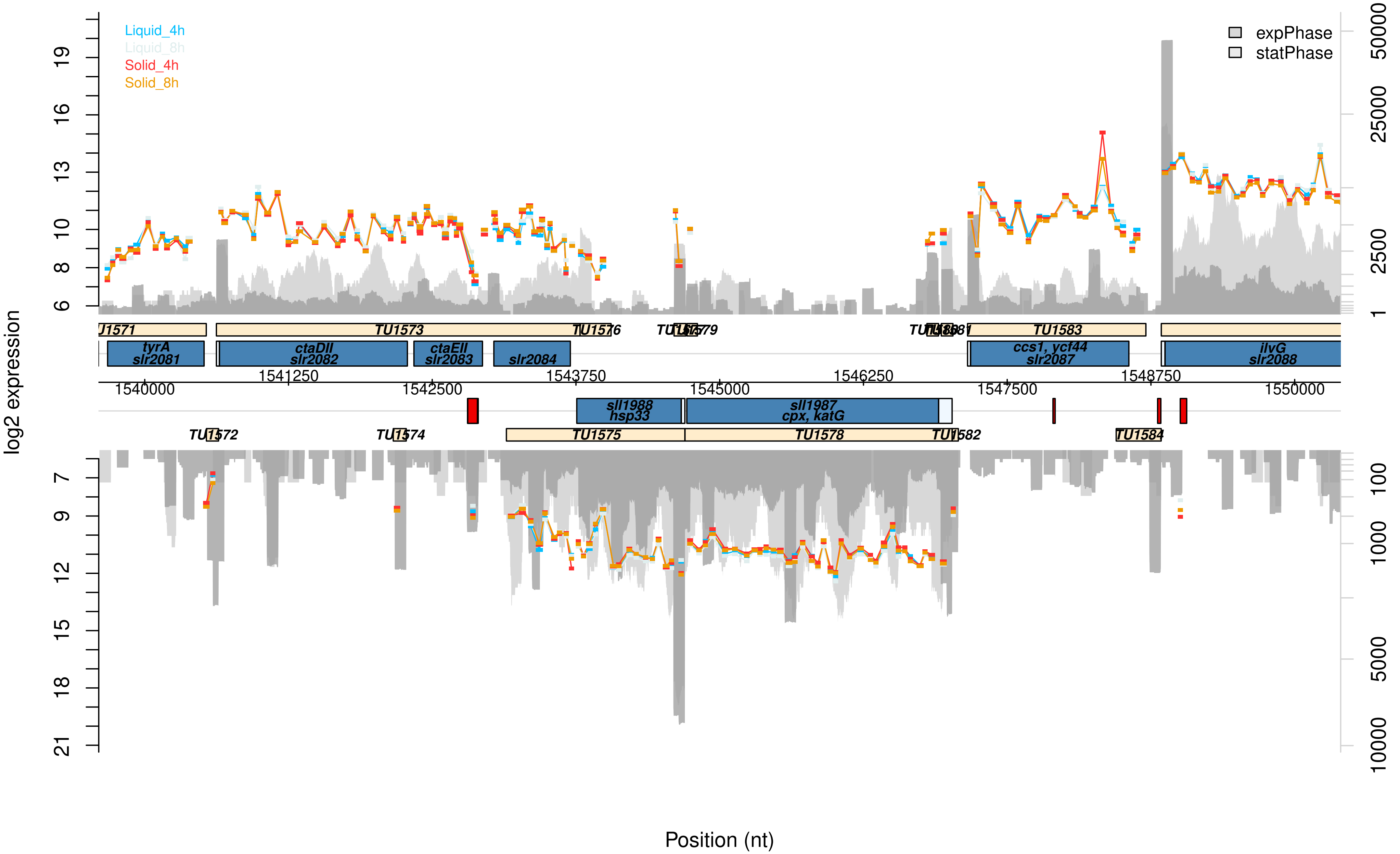

Liquid_4h
Liquid_8h
Solid_4h
Solid_8h
expPhase
statPhase
log2 expression
Position (nt)
tyrA slr2081
ctaDII slr2082
ctaEII slr2083
slr2084
ccs1, ycf44 slr2087
ilvG slr2088
sll1988 hsp33
sll1987 cpx, katG
TU1573
TU1575
TU1578
TU1583
TU1584
1540000
1541250
1542500
1543750
1545000
1546250
1547500
1548750
1550000

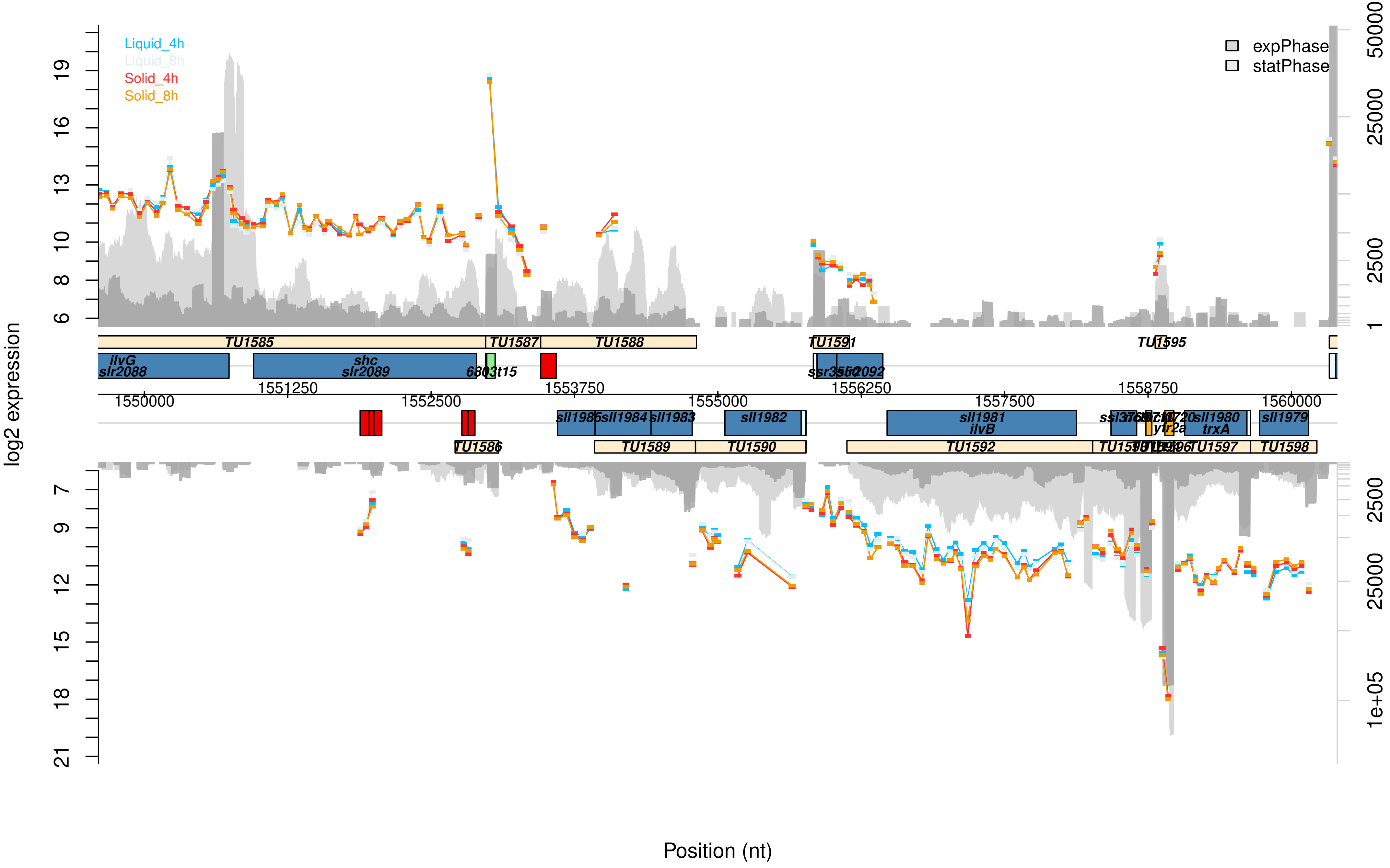

Liquid_4h
Liquid_8h
Solid_4h
Solid_8h
expPhase
statPhase
log2 expression
Position (nt)
TU1585
TU1587
TU1588
TU1591
TU1595
ilvG
slr2088
shc
slr2089
1550000
1551250
1552500
1553750
1555000
1556250
1557500
1558750
1560000
sll1985
sll1984
sll1983
sll1982
sll1981
ilvB
sll1980
trxA
sll1979
TU1586
TU1589
TU1590
TU1592
TU1597
TU1598

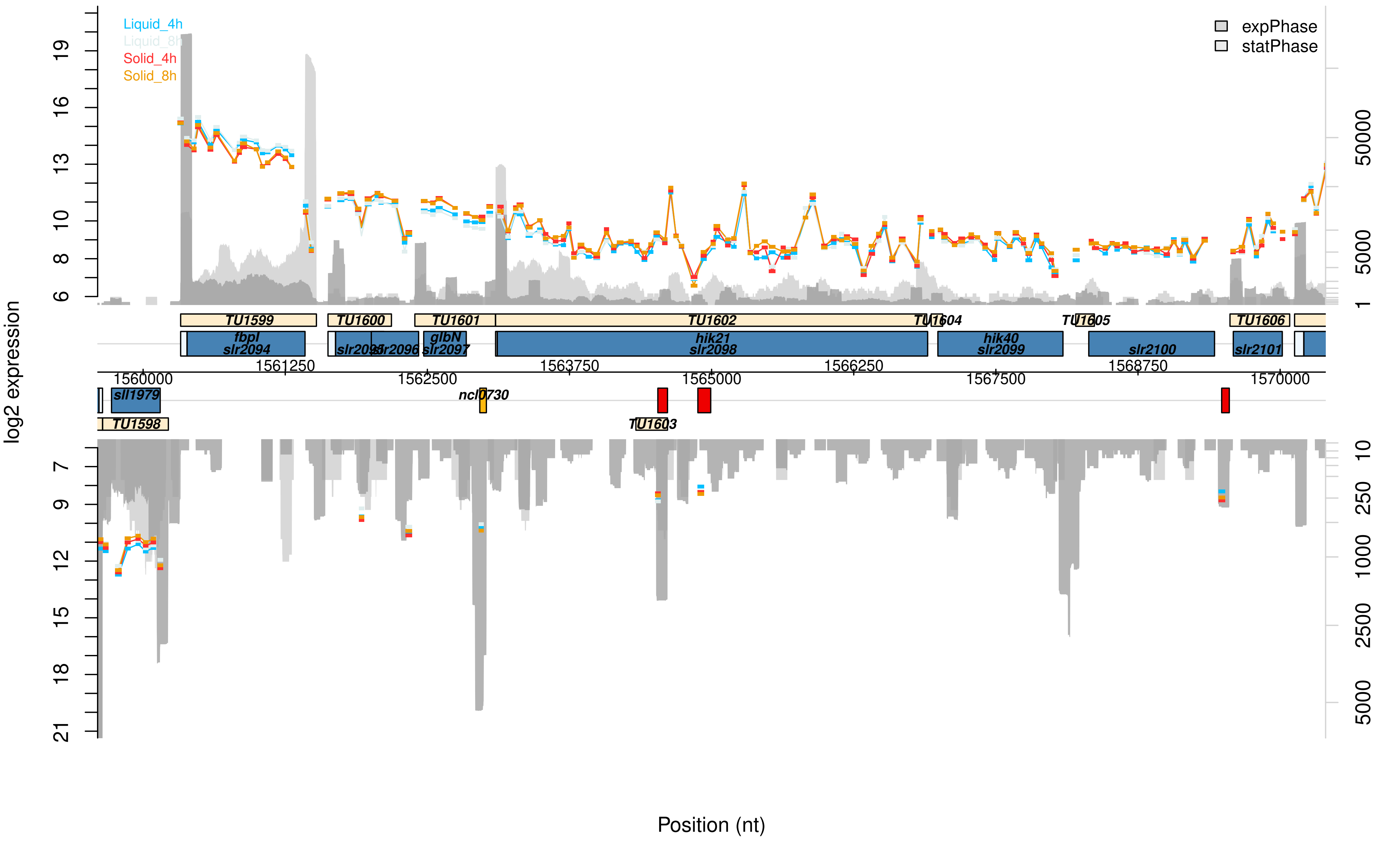

Liquid_4h
Liquid_8h
Solid_4h
Solid_8h
expPhase
statPhase
log2 expression
TU1599
TU1600
TU1601
TU1602
TU1604
TU1605
TU1606
fbpI
slr2094
slr2095
slr2096
glbN
slr2097
hik21
slr2098
hik40
slr2099
slr2100
slr2101
sll1979
ncl0730
TU1598
TU1603
1560000
1561250
1562500
1563750
1565000
1566250
1567500
1568750
1570000
Position (nt)

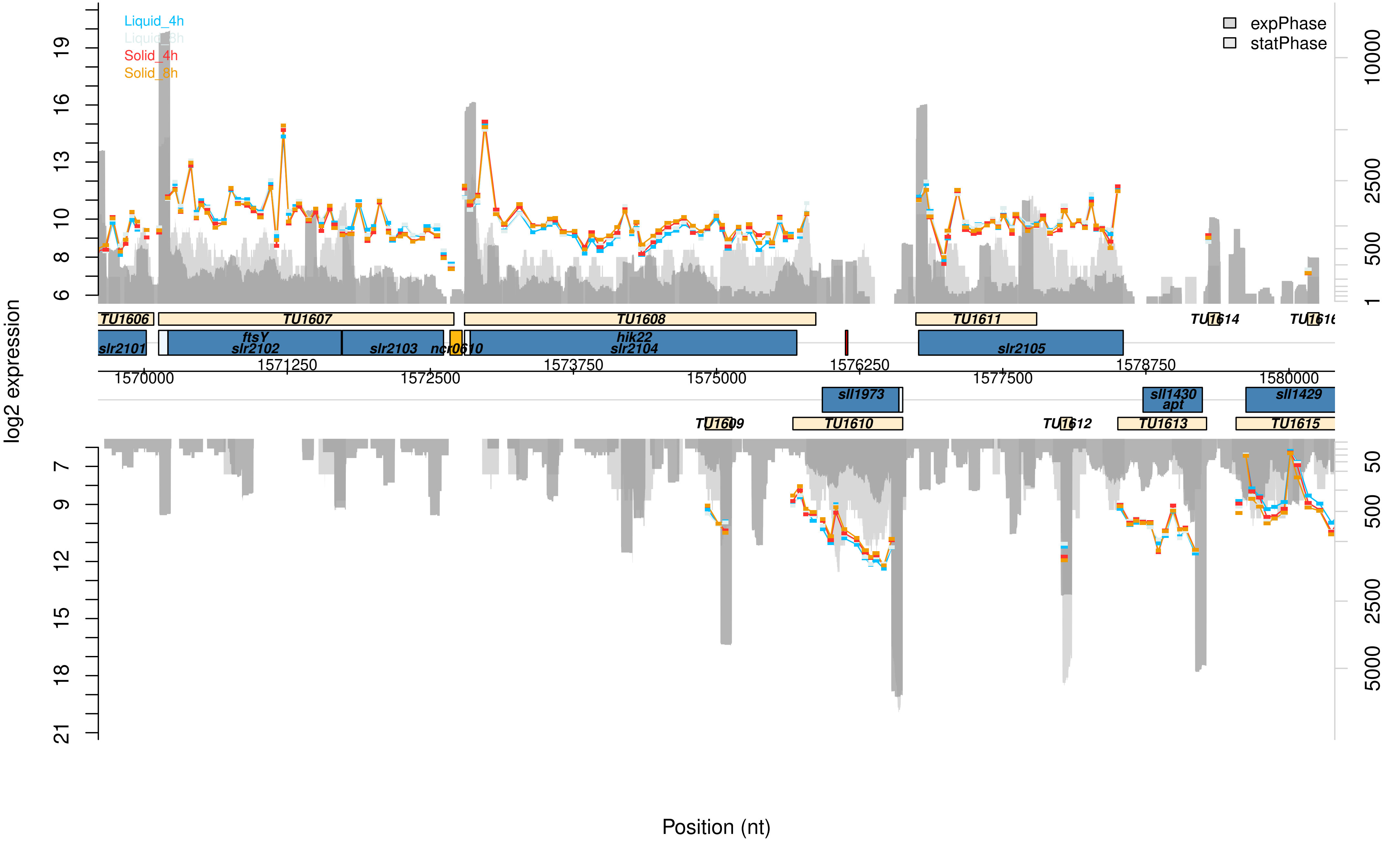

Liquid_4h
Liquid_8h
Solid_4h
Solid_8h
expPhase
statPhase
log2 expression
TU1606
TU1607
TU1608
TU1611
TU1614
slr2101
ftsY
slr2102
slr2103
ncr0610
hik22
slr2104
slr2105
1570000
1571250
1572500
1573750
1575000
1576250
1577500
1578750
1580000
sll1973
sll1430
apt
sll1429
TU1609
TU1610
TU1612
TU1613
TU1615
Position (nt)

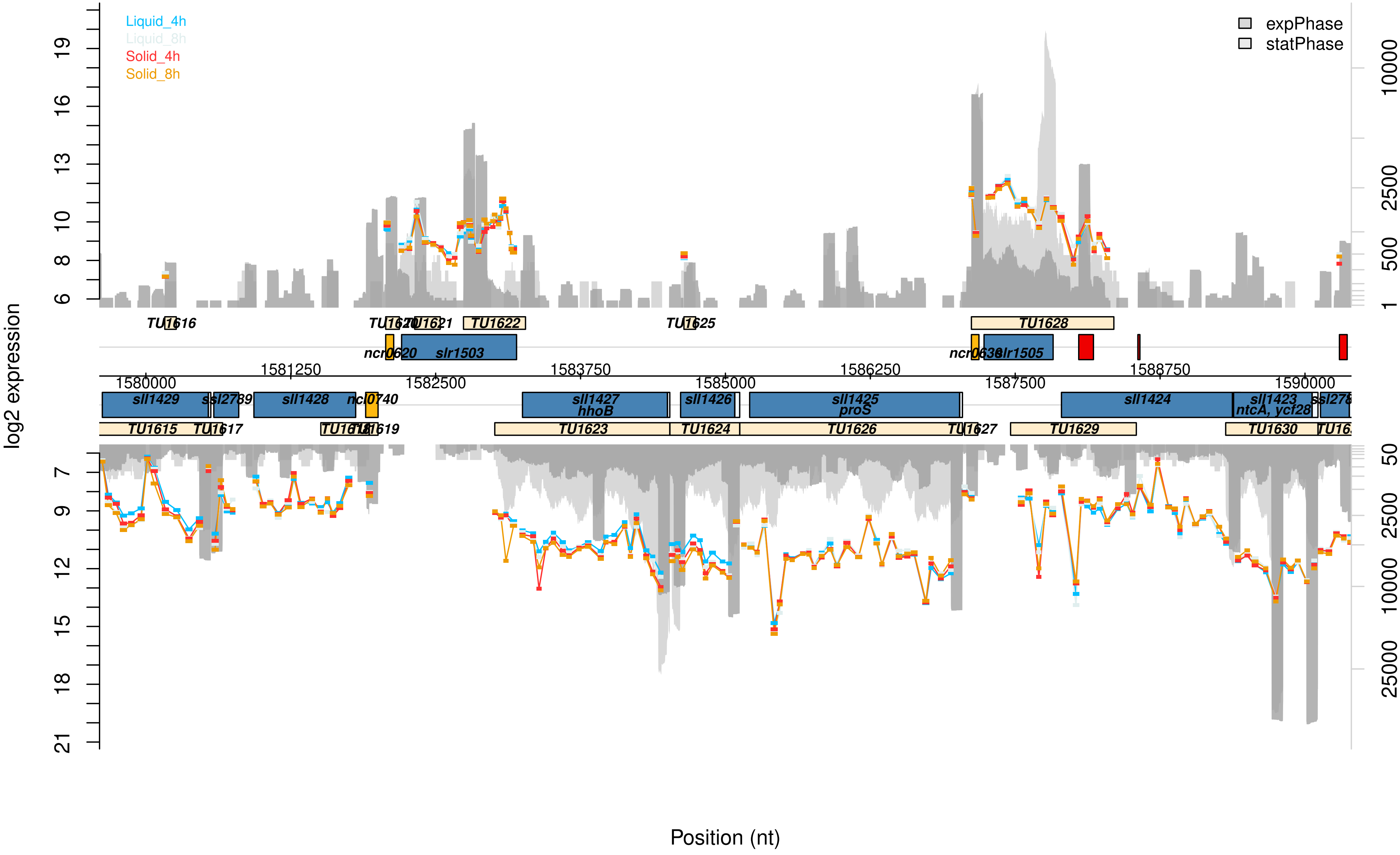
Liquid_4h
Liquid_8h
Solid_4h
Solid_8h
expPhase
statPhase
log2 expression
Position (nt)
TU1616
TU1620
TU1621
TU1622
TU1625
TU1628
ncr0620
slr1503
ncr0638
slr1505
1580000
1581250
1582500
1583750
1585000
1586250
1587500
1588750
1590000
sll1429
ssl2789
sll1428
ncl0740
sll1427
hhoB
sll1426
sll1425
proS
sll1424
sll1423
ntcA, ycf28
TU1615
TU1617
TU1618
TU1619
TU1623
TU1624
TU1626
TU1627
TU1629
TU1630

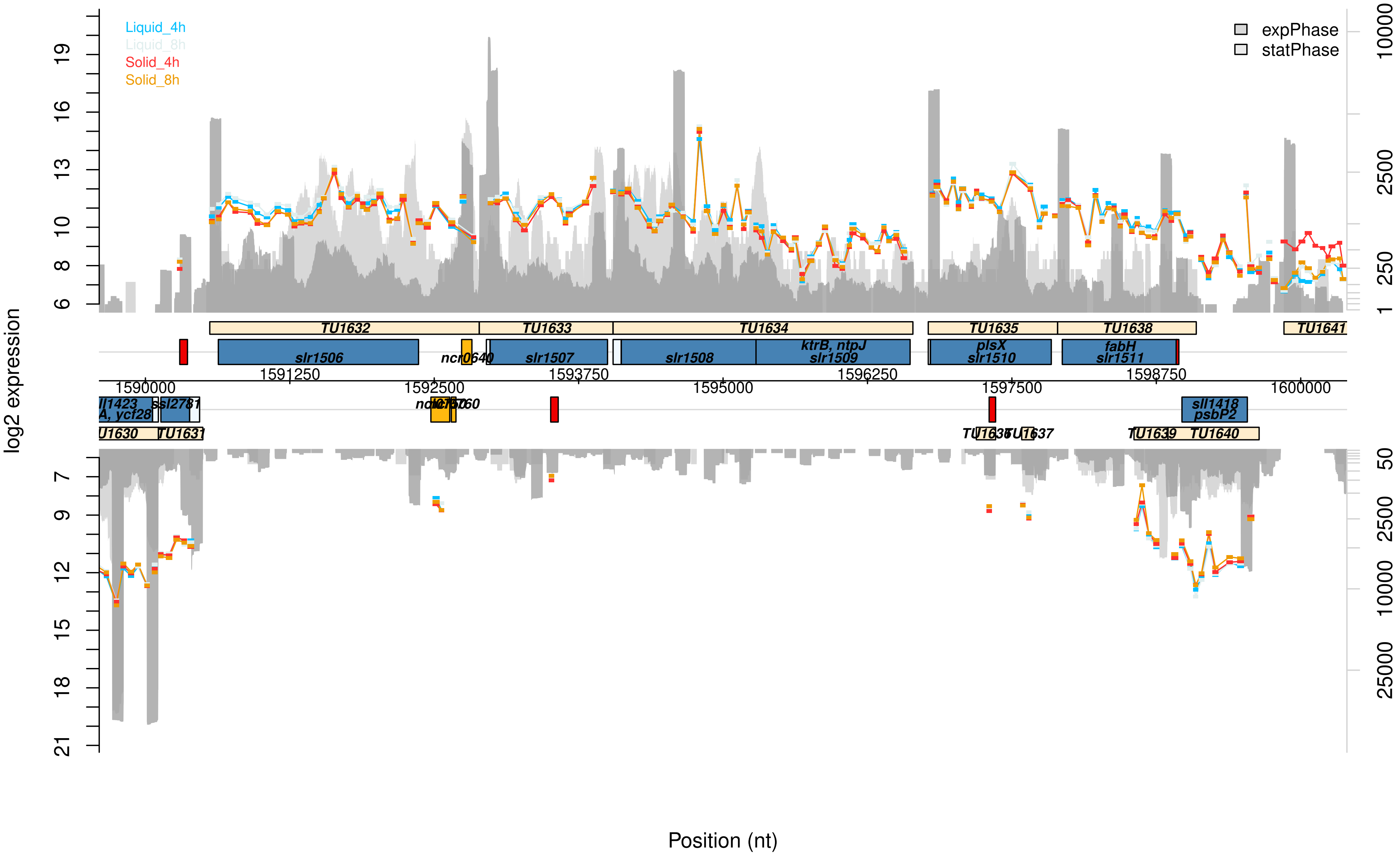

Liquid_4h
Liquid_8h
Solid_4h
Solid_8h
expPhase
statPhase
log2 expression
Position (nt)
TU1632
TU1633
TU1634
TU1635
TU1638
TU1641
slr1506
ncr0640
slr1507
slr1508
ktrB, ntpJ
slr1509
plsX
slr1510
fabH
slr1511
1590000
1591250
1592500
1593750
1595000
1596250
1597500
1598750
1600000
sll1418
psbP2
TU1630
TU1631
TU1639
TU1640

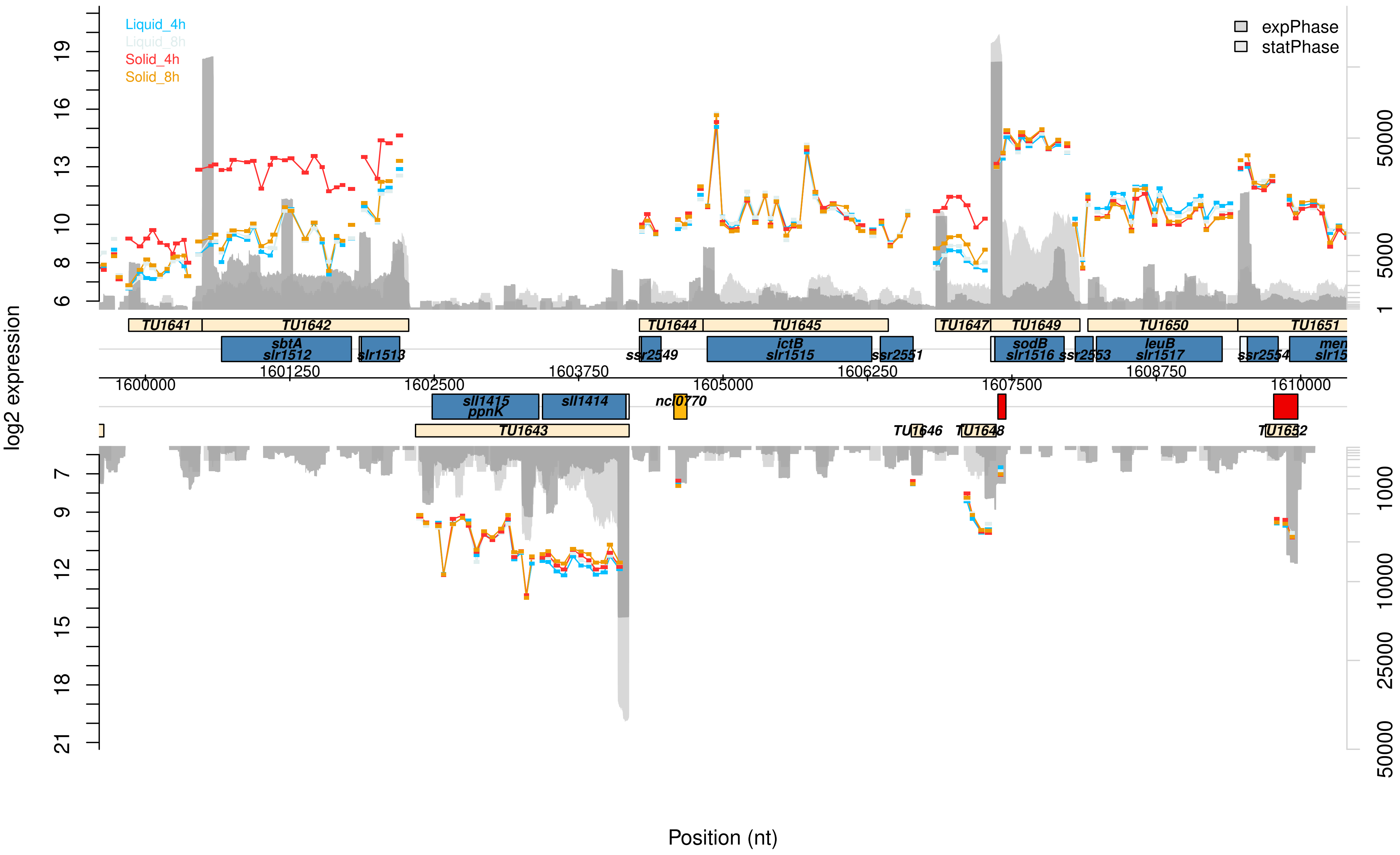
Liquid_4h
Liquid_8h
Solid_4h
Solid_8h
expPhase
statPhase
log2 expression
Position (nt)
TU1641
TU1642
TU1643
TU1644
TU1645
TU1646
TU1647
TU1648
TU1649
TU1650
TU1651
TU1652
sbtA
slr1512
slr1513
sll1415
ppnK
sll1414
ssr2549
ncl0770
ictB
slr1515
ssr2551
sodB
slr1516
ssr2553
leuB
slr1517
ssr2554
1600000
1601250
1602500
1603750
1605000
1606250
1607500
1608750
1610000

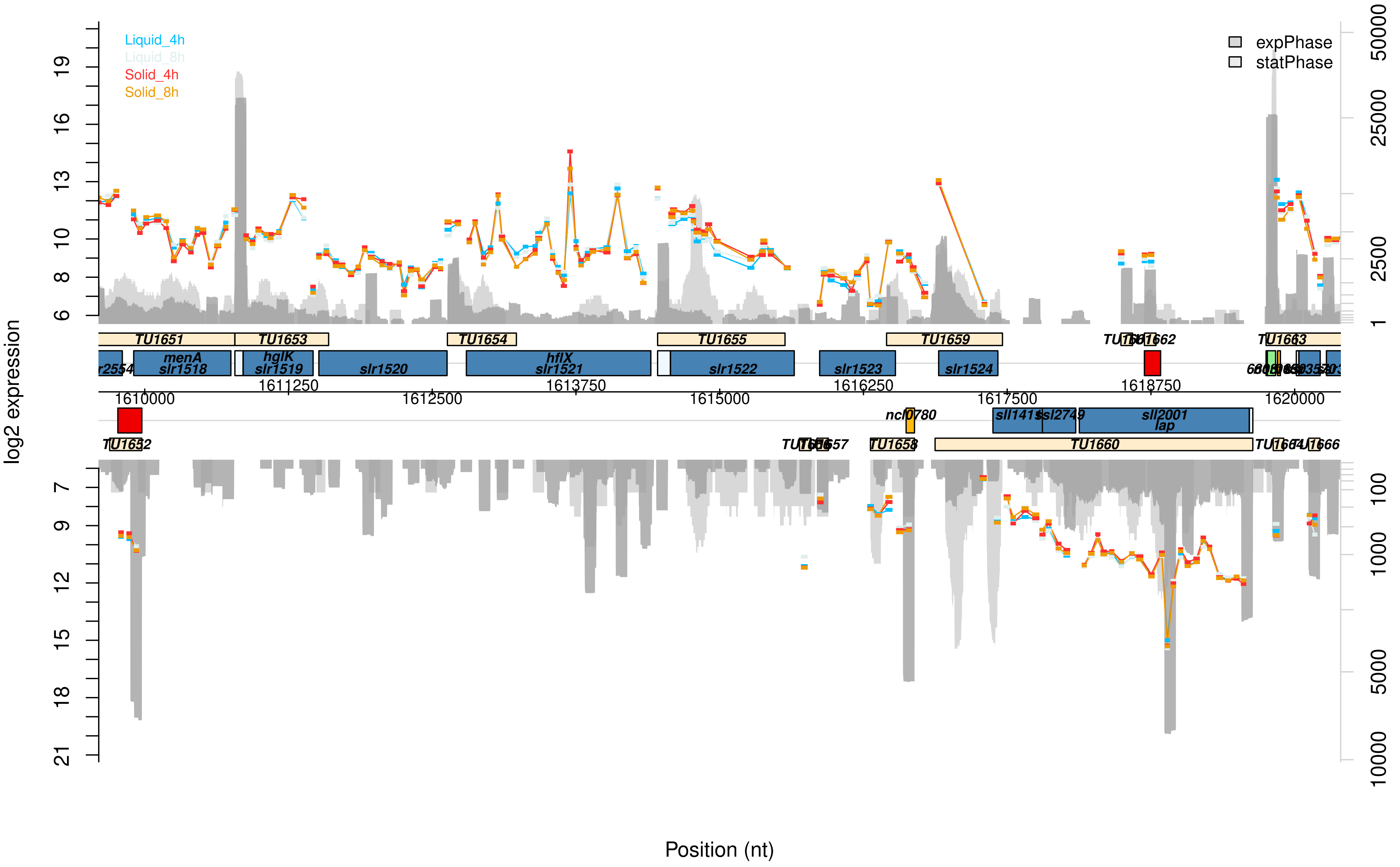
Liquid_4h
Liquid_8h
Solid_4h
Solid_8h
expPhase
statPhase
log2 expression
Position (nt)
TU1651
TU1653
TU1654
TU1655
TU1659
TU1663
menA
slr1518
hglK
slr1519
slr1520
hflX
slr1521
slr1522
slr1523
slr1524
1610000
1611250
1612500
1613750
1615000
1616250
1617500
1618750
1620000
ncl0780
sll2749
sll2001
lap
TU1652
TU1658
TU1660

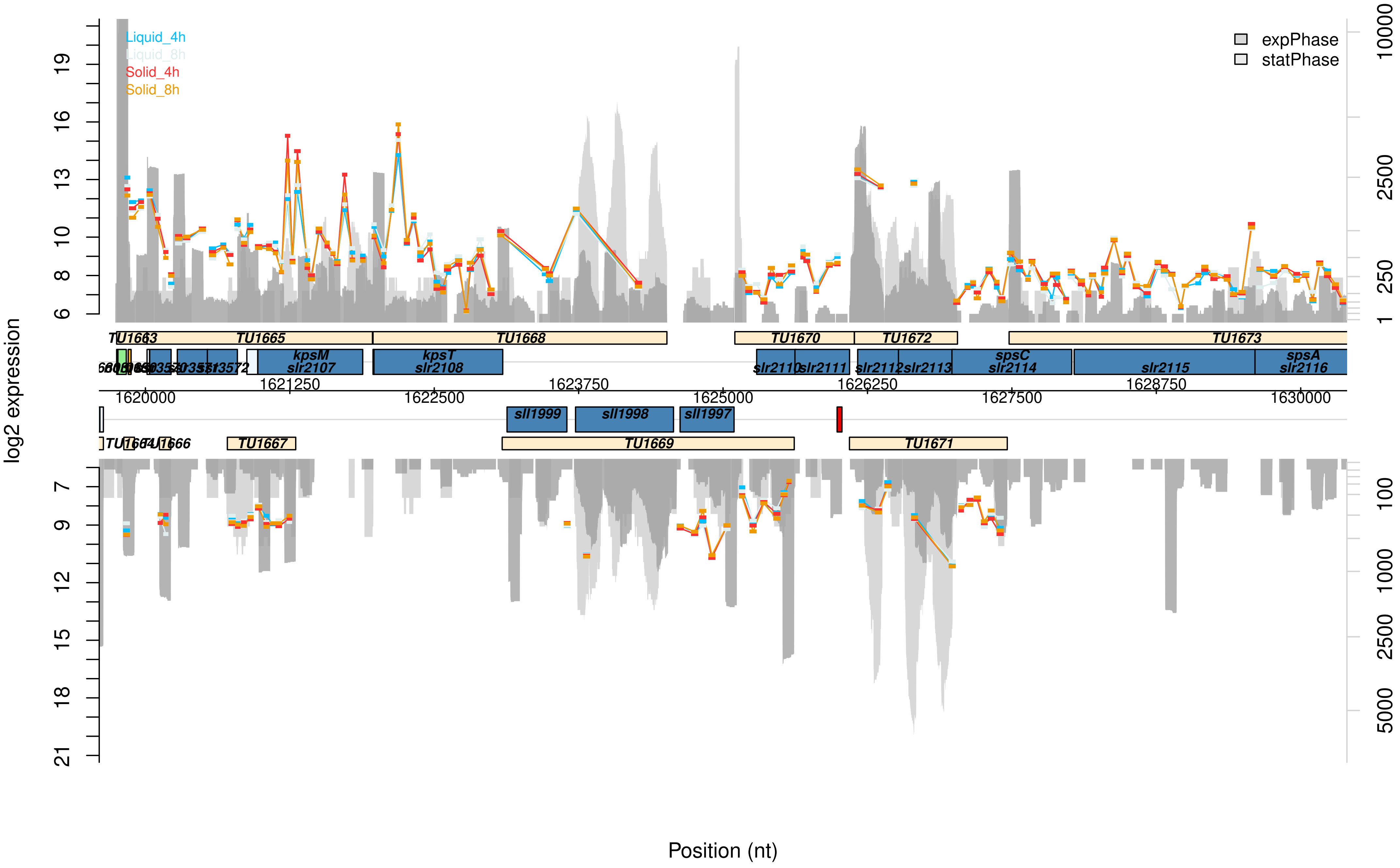

Liquid_4h
Liquid_8h
Solid_4h
Solid_8h
expPhase
statPhase
log2 expression
Position (nt)
TU1663
TU1665
TU1668
TU1670
TU1672
TU1673
kpsM
slr2107
kpsT
slr2108
slr2110
slr2111
slr2112
slr2113
spsC
slr2114
slr2115
spsA
slr2116
sll1999
sll1998
sll1997
TU1667
TU1669
TU1671
1620000
1621250
1622500
1623750
1625000
1626250
1627500
1628750
1630000

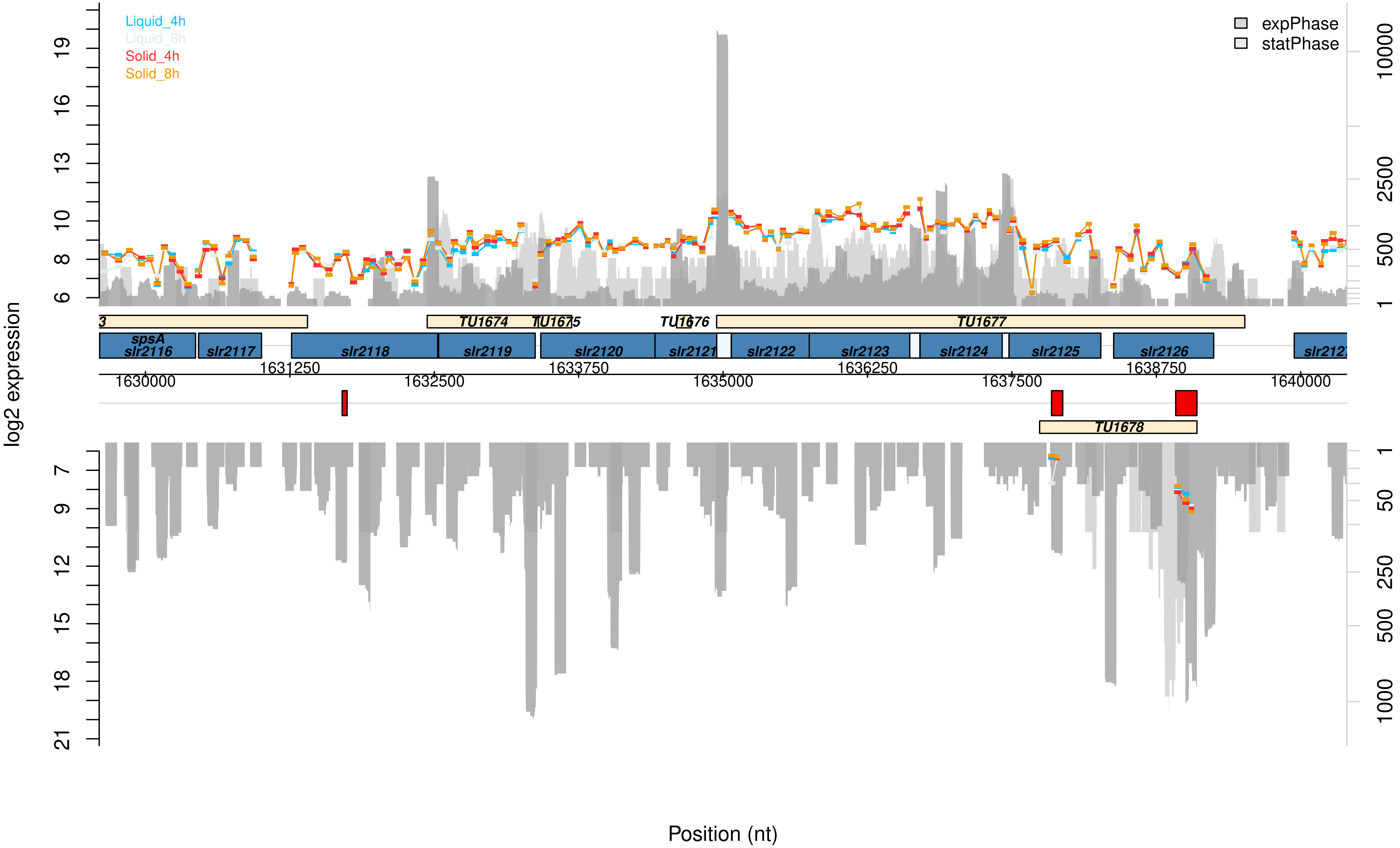
Liquid_4h
Liquid_8h
Solid_4h
Solid_8h
expPhase
statPhase
log2 expression
TU1674
TU1675
TU1676
TU1677
TU1678
spsA
slr2116
slr2117
slr2118
slr2119
slr2120
slr2121
slr2122
slr2123
slr2124
slr2125
slr2126
1630000
1631250
1632500
1633750
1635000
1636250
1637500
1638750
1640000
Position (nt)

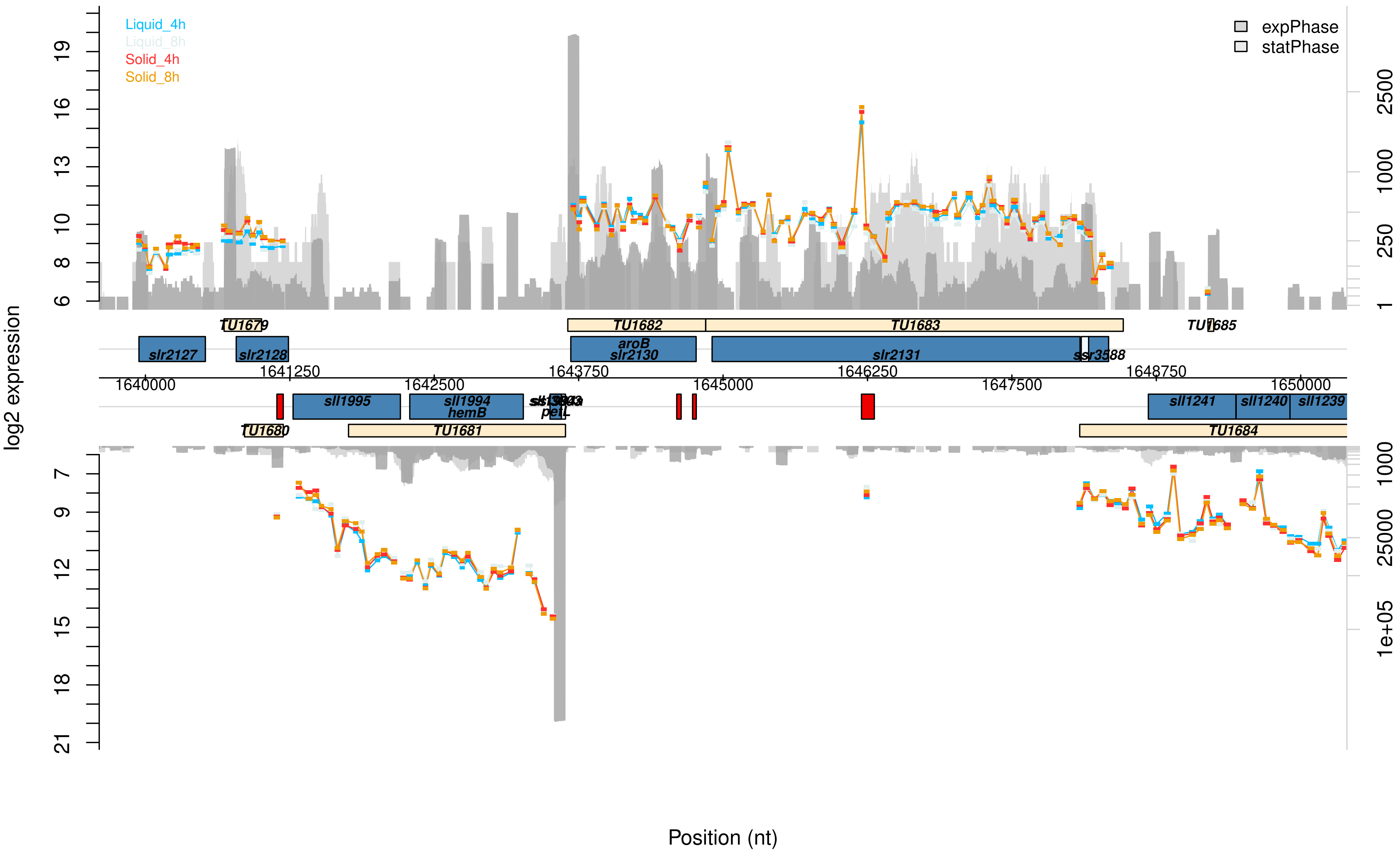

Liquid_4h
Liquid_8h
Solid_4h
Solid_8h
expPhase
statPhase
log2 expression
Position (nt)
TU1679
TU1682
TU1683
TU1685
slr2127
slr2128
aroB
slr2130
slr2131
ssr3588
1640000
1641250
1642500
1643750
1645000
1646250
1647500
1648750
1650000
sll1995
sll1994
hemB
petL
sll1241
sll1240
sll1239
TU1680
TU1681
TU1684

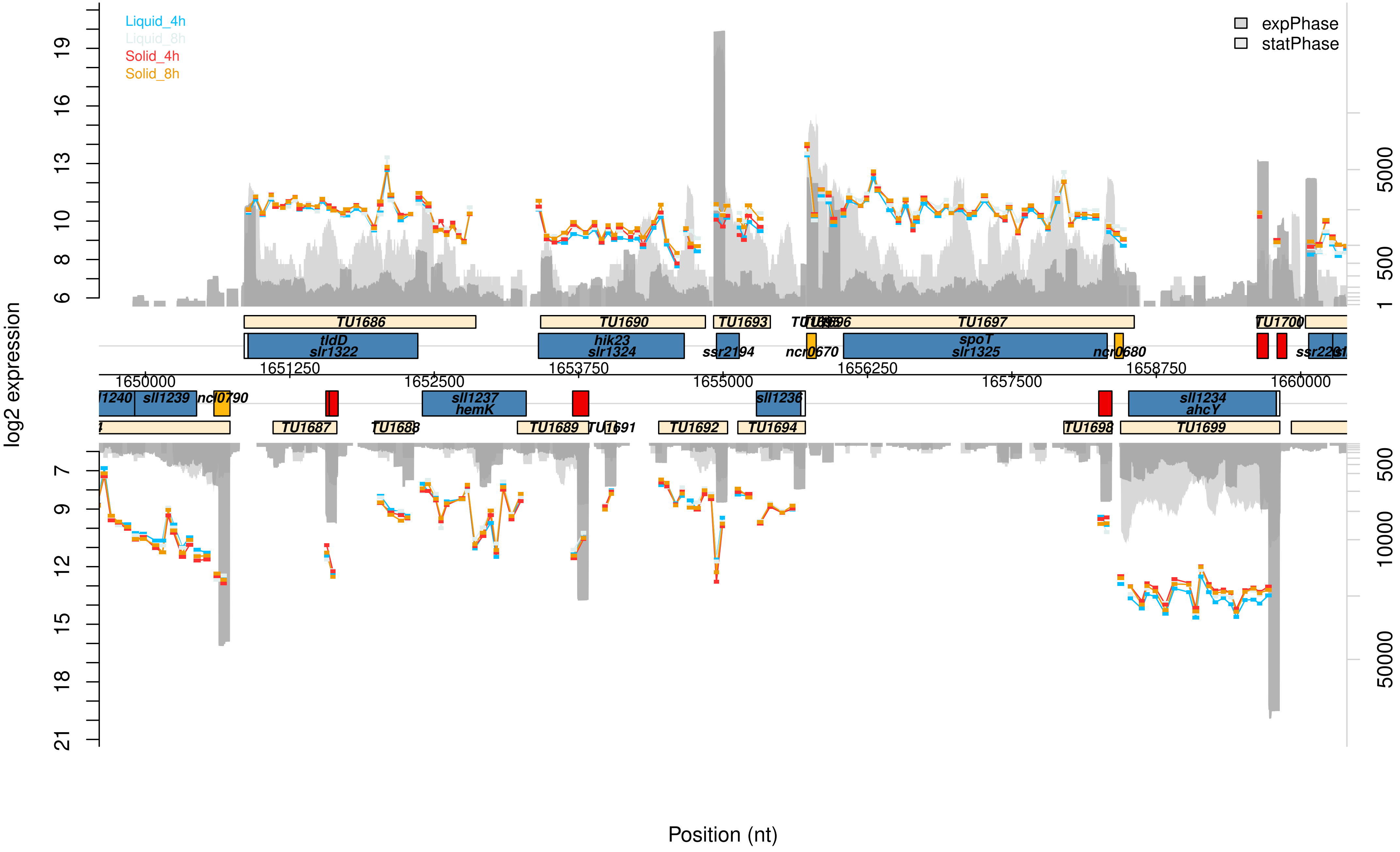
Liquid_4h
Liquid_8h
Solid_4h
Solid_8h
expPhase
statPhase
log2 expression
Position (nt)
TU1686
TU1690
TU1693
TU1697
TU1700
tldD
slr1322
hik23
slr1324
ssr2194
ncr0670
spoT
slr1325
ncr0680
1651250
1653750
1656250
1658750
1650000
1652500
1655000
1657500
1660000
sll1239
sll1237
hemK
sll1236
sll1234
ahcY
TU1687
TU1689
TU1691
TU1692
TU1694
TU1698
TU1699

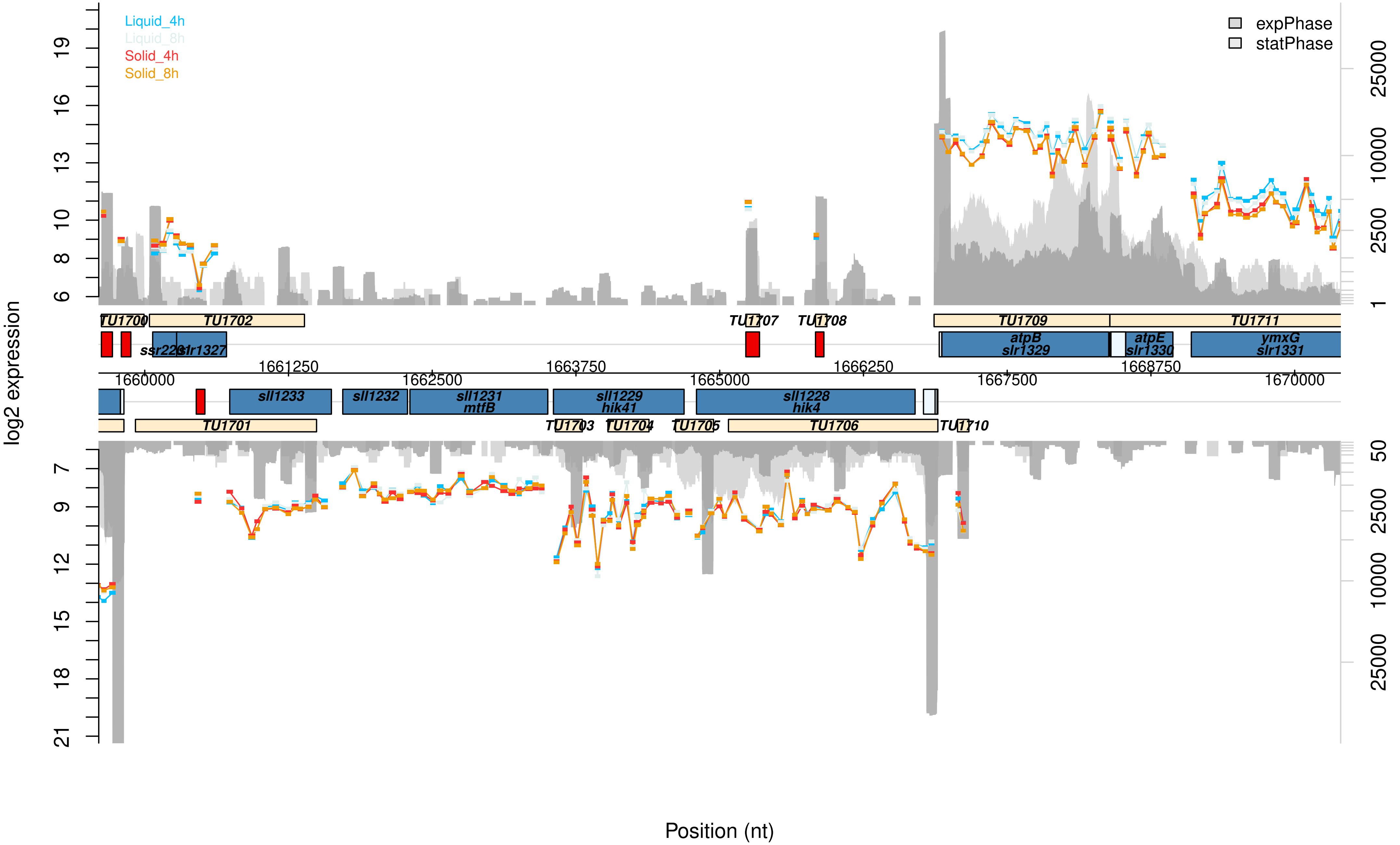

Liquid_4h
Liquid_8h
Solid_4h
Solid_8h
expPhase
statPhase
log2 expression
Position (nt)
TU1700
TU1702
TU1707
TU1708
TU1709
TU1711
ssr2281
slr1327
atpB
slr1329
atpE
slr1330
ymxG
slr1331
sll1233
sll1232
sll1231
mtfB
sll1229
hik41
sll1228
hik4
TU1701
TU1703
TU1704
TU1705
TU1706
TU1710
1660000
1661250
1662500
1663750
1665000
1666250
1667500
1668750
1670000

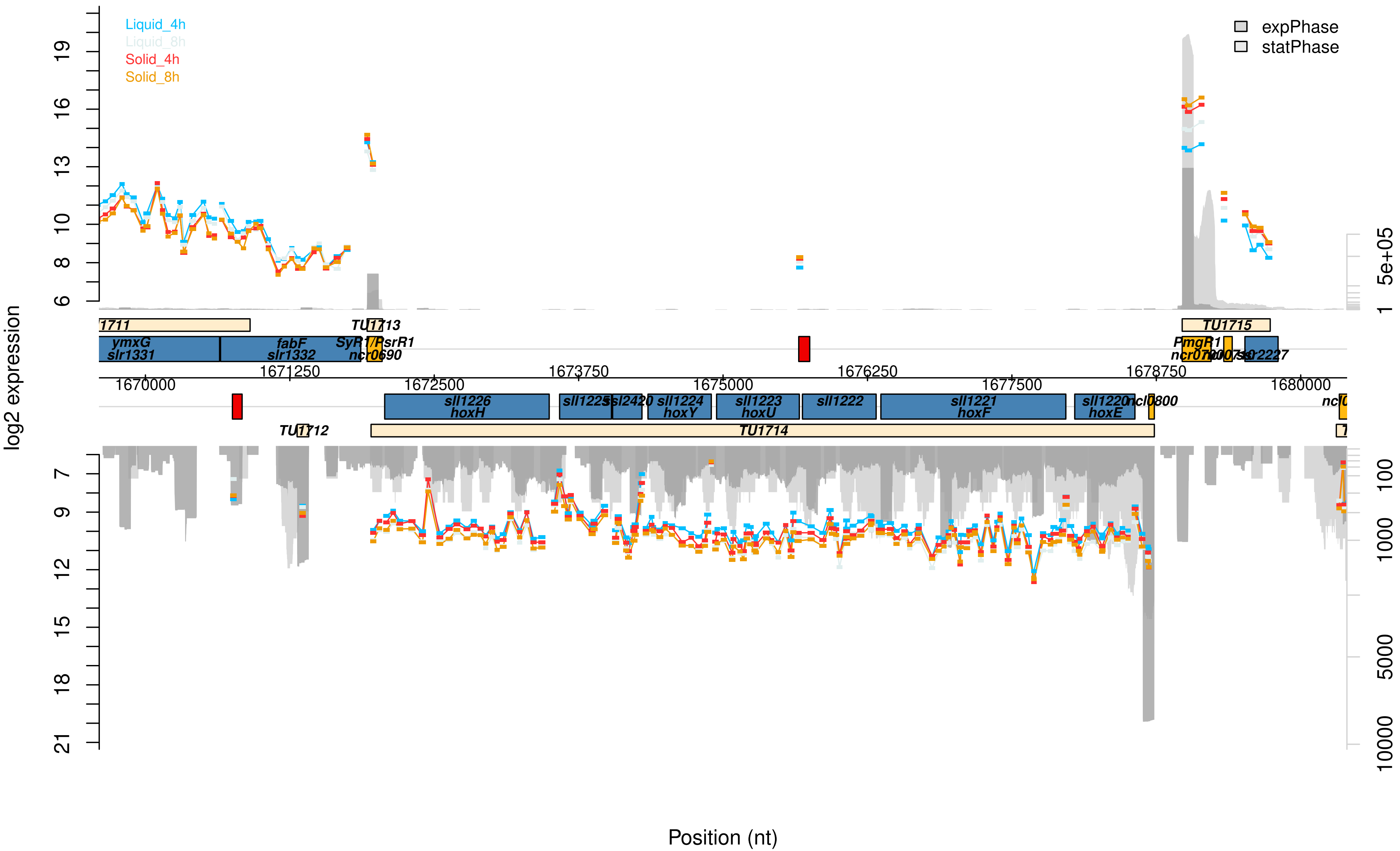

Liquid_4h
Liquid_8h
Solid_4h
Solid_8h
expPhase
statPhase
log2 expression
Position (nt)
TU1713
TU1715
ymxG
slr1331
fabF
slr1332
SyR1/PsrR1
ncr0690
PmgR1
sll1226
hoxH
sll1224
hoxY
sll1223
hoxU
sll1222
sll1221
hoxF
sll1220
hoxE
ncl0800
TU1712
TU1714
1670000
1671250
1672500
1673750
1675000
1676250
1677500
1678750
1680000

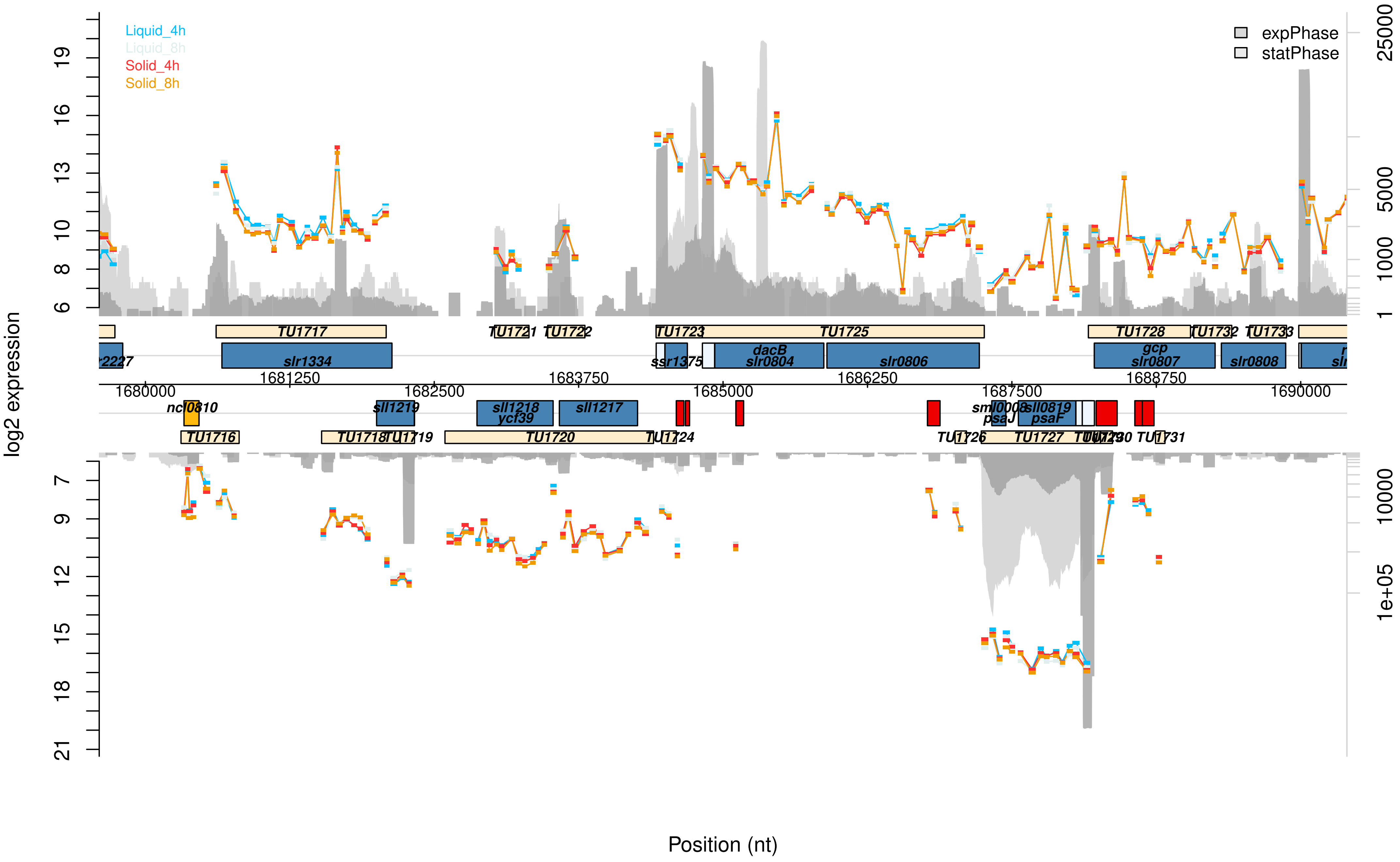

Liquid_4h
Liquid_8h
Solid_4h
Solid_8h
expPhase
statPhase
log2 expression
Position (nt)
TU1717
TU1721
TU1722
TU1723
TU1725
TU1728
TU1732
TU1733
slr1334
ssr1375
dacB
slr0804
slr0806
gcp
slr0807
slr0808
1680000
1681250
1682500
1683750
1685000
1686250
1687500
1688750
1690000
ncl0810
sll1219
sll1218
ycf39
sll1217
sml0008
psaJ
sll0819
psaF
TU1716
TU1718
TU1719
TU1720
TU1724
TU1726
TU1727
TU1731

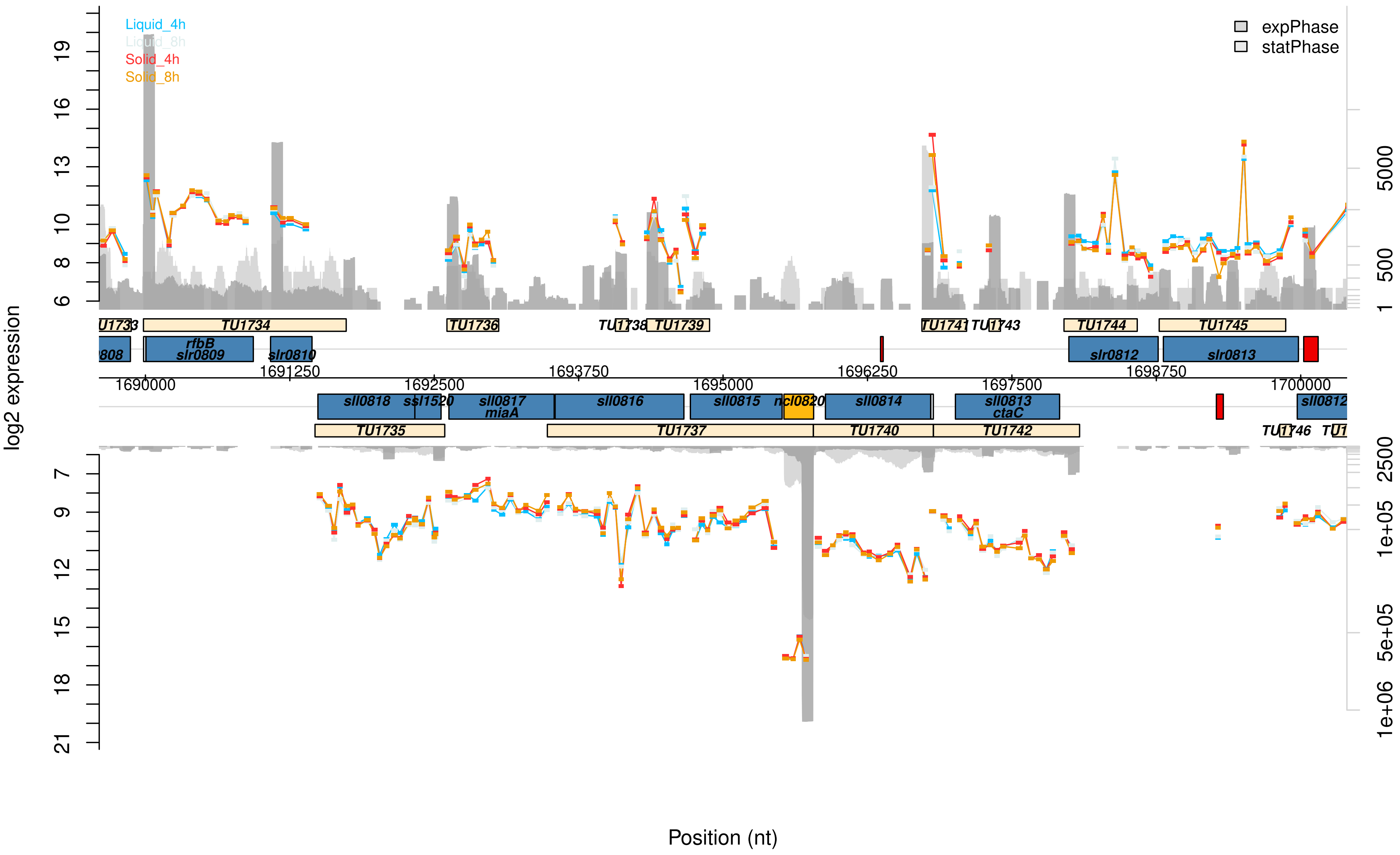

Liquid_4h
Liquid_8h
Solid_4h
Solid_8h
expPhase
statPhase
log2 expression
Position (nt)
TU1734
TU1736
TU1739
TU1741
TU1744
TU1745
rfbB
slr0809
slr0810
slr0812
slr0813
1690000
1691250
1692500
1693750
1695000
1696250
1697500
1698750
1700000
sll0818
sll0817
miaA
sll0816
sll0815
sll0814
sll0813
ctaC
TU1735
TU1737
TU1740
TU1742

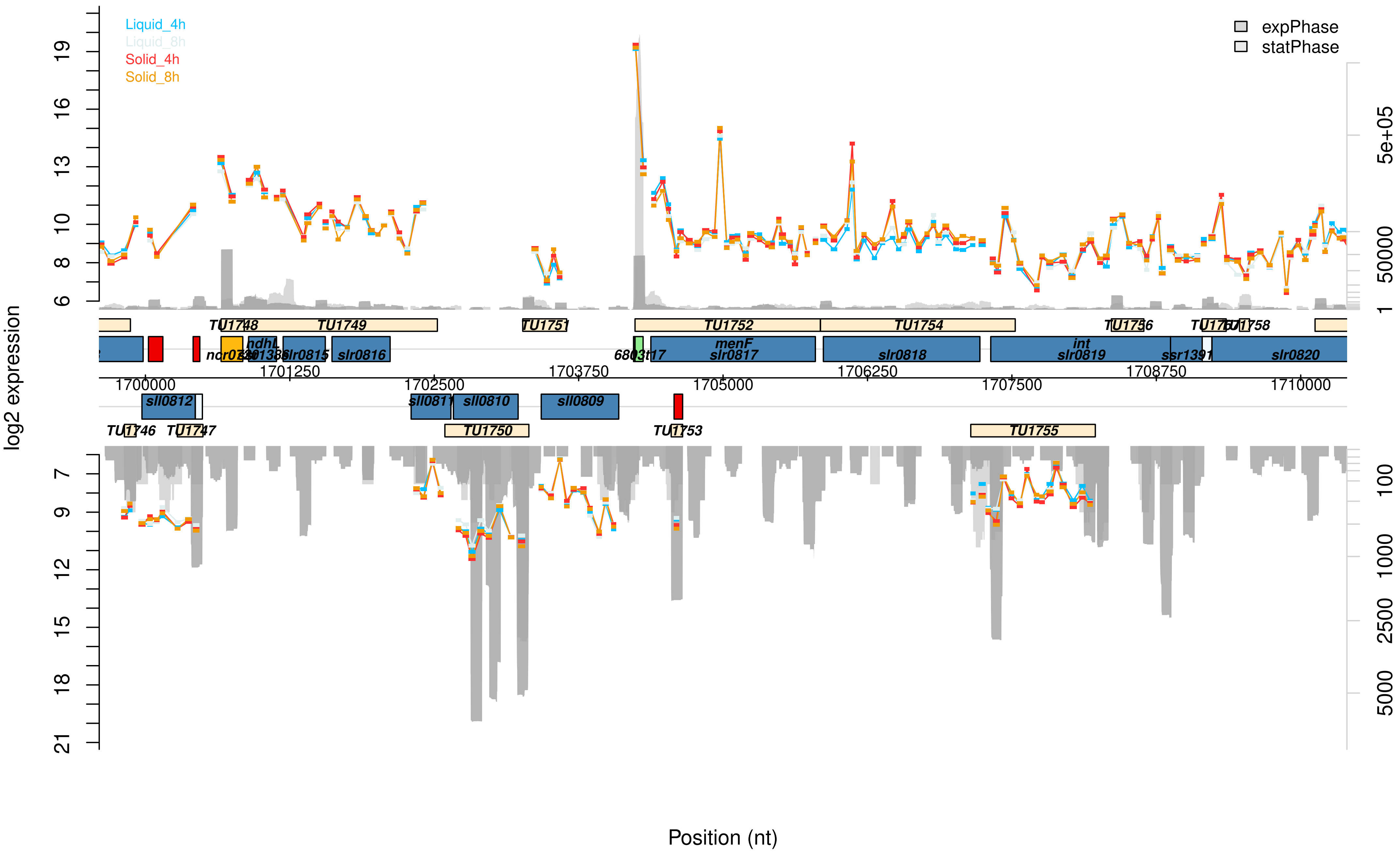
Liquid_4h
Liquid_8h
Solid_4h
Solid_8h
expPhase
statPhase
log2 expression
Position (nt)
TU1748
TU1749
TU1751
TU1752
TU1754
TU1756
TU1758
ndhL
slr0815
slr0816
menF
slr0817
slr0818
int
slr0819
ssr1391
slr0820
1700000
1701250
1702500
1703750
1705000
1706250
1707500
1708750
1710000
sll0812
sll0810
sll0809
TU1746
TU1747
TU1750
TU1753
TU1755

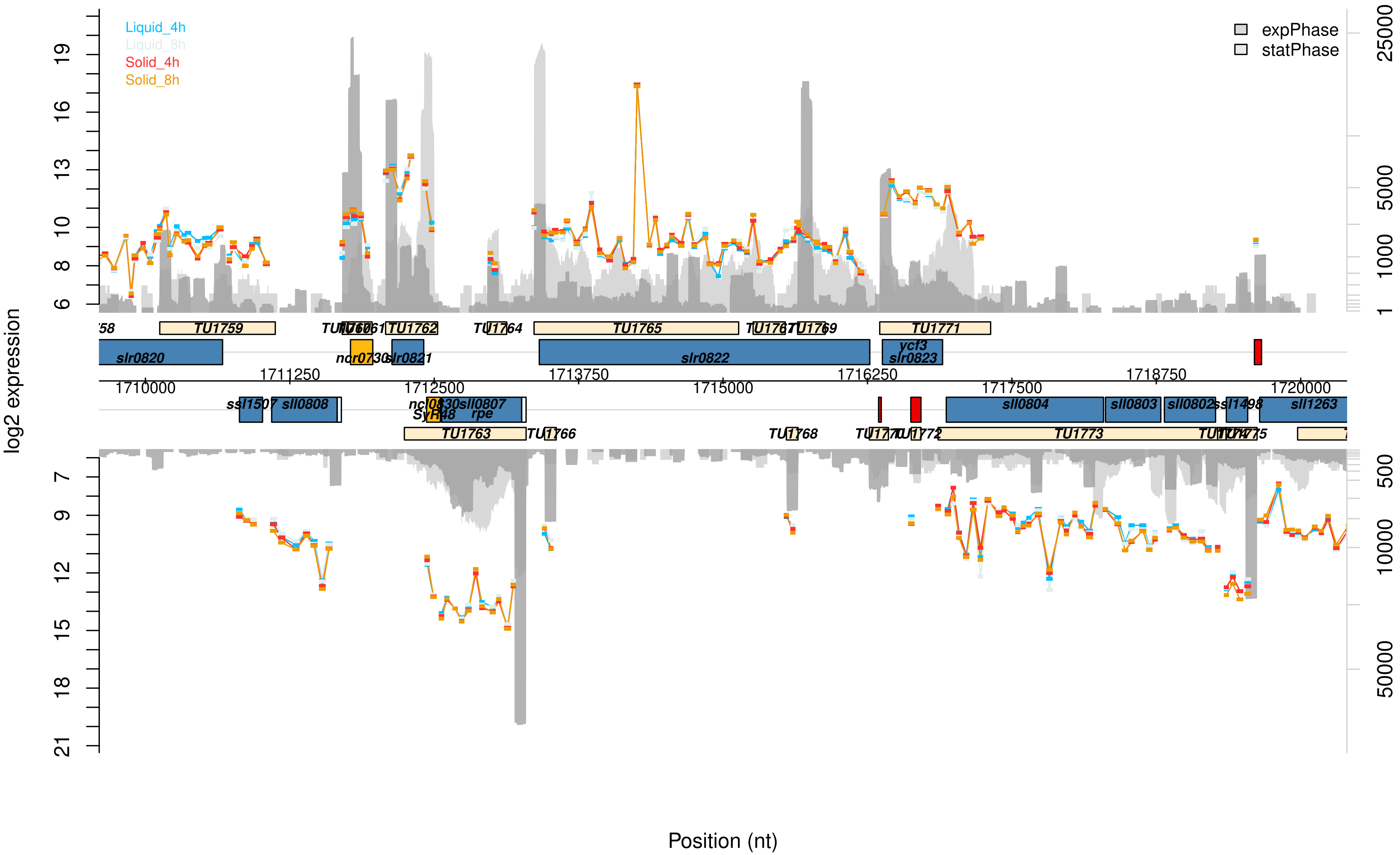

Liquid_4h
Liquid_8h
Solid_4h
Solid_8h
expPhase
statPhase
log2 expression
Position (nt)
TU1759
TU1762
TU1764
TU1765
TU1769
TU1771
slr0820
nor0730
slr0821
slr0822
ycf3
slr0823
1710000
1711250
1712500
1713750
1715000
1716250
1717500
1718750
1720000
ssl1507
sll0808
sll0807
SyR48
rpe
sll0804
sll0803
sll0802
ssl1498
sll1263
TU1763
TU1766
TU1768
TU1772
TU1773
TU1775

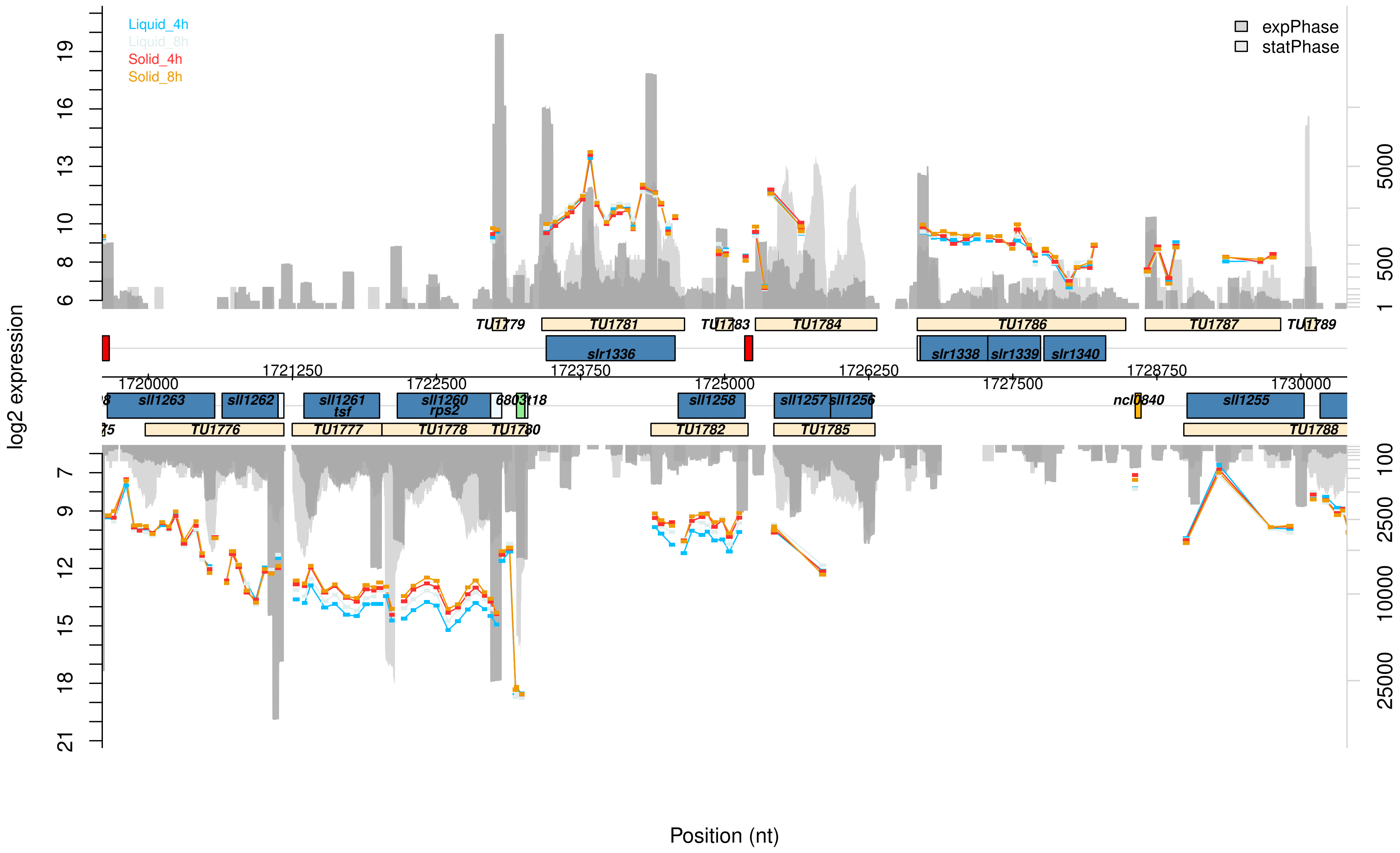

Liquid_4h
Liquid_8h
Solid_4h
Solid_8h
expPhase
statPhase
log2 expression
Position (nt)
TU1779
TU1781
TU1783
TU1784
TU1786
TU1787
TU1789
slr1336
slr1338
slr1339
slr1340
1720000
1721250
1722500
1723750
1725000
1726250
1727500
1728750
1730000
sll1263
sll1262
sll1261
tsf
sll1260
rps2
sll1258
sll1257
sll1256
ncl0840
sll1255
TU1776
TU1777
TU1778
TU1780
TU1782
TU1785
TU1788

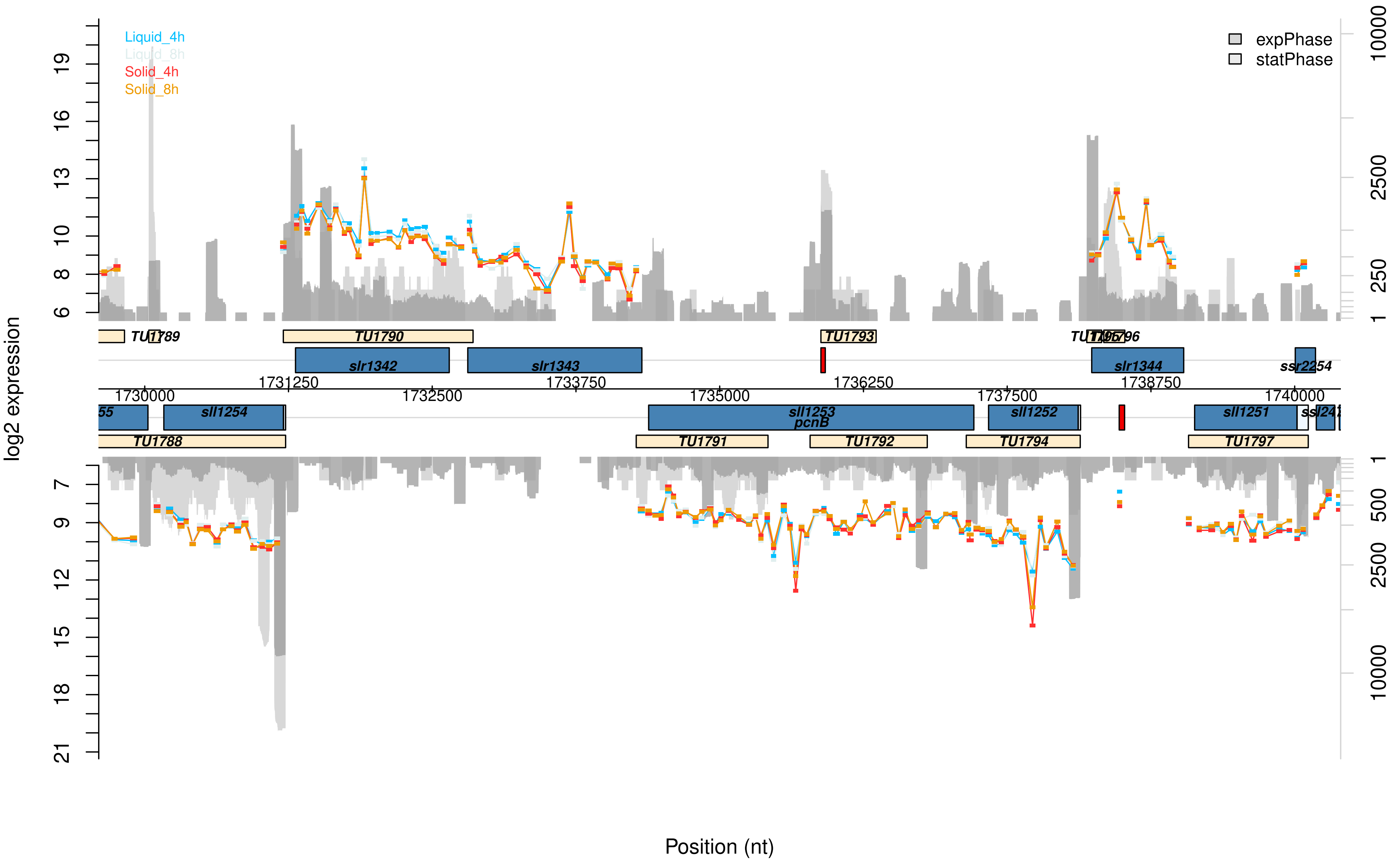

Liquid_4h
Liquid_8h
Solid_4h
Solid_8h
expPhase
statPhase
log2 expression
Position (nt)
TU1789
TU1790
TU1793
slr1342
slr1343
slr1344
ssr2254
1731250
1733750
1736250
1738750
1730000
1732500
1735000
1737500
1740000
sll1254
sll1253
pcnB
sll1252
sll1251
TU1788
TU1791
TU1792
TU1794
TU1797

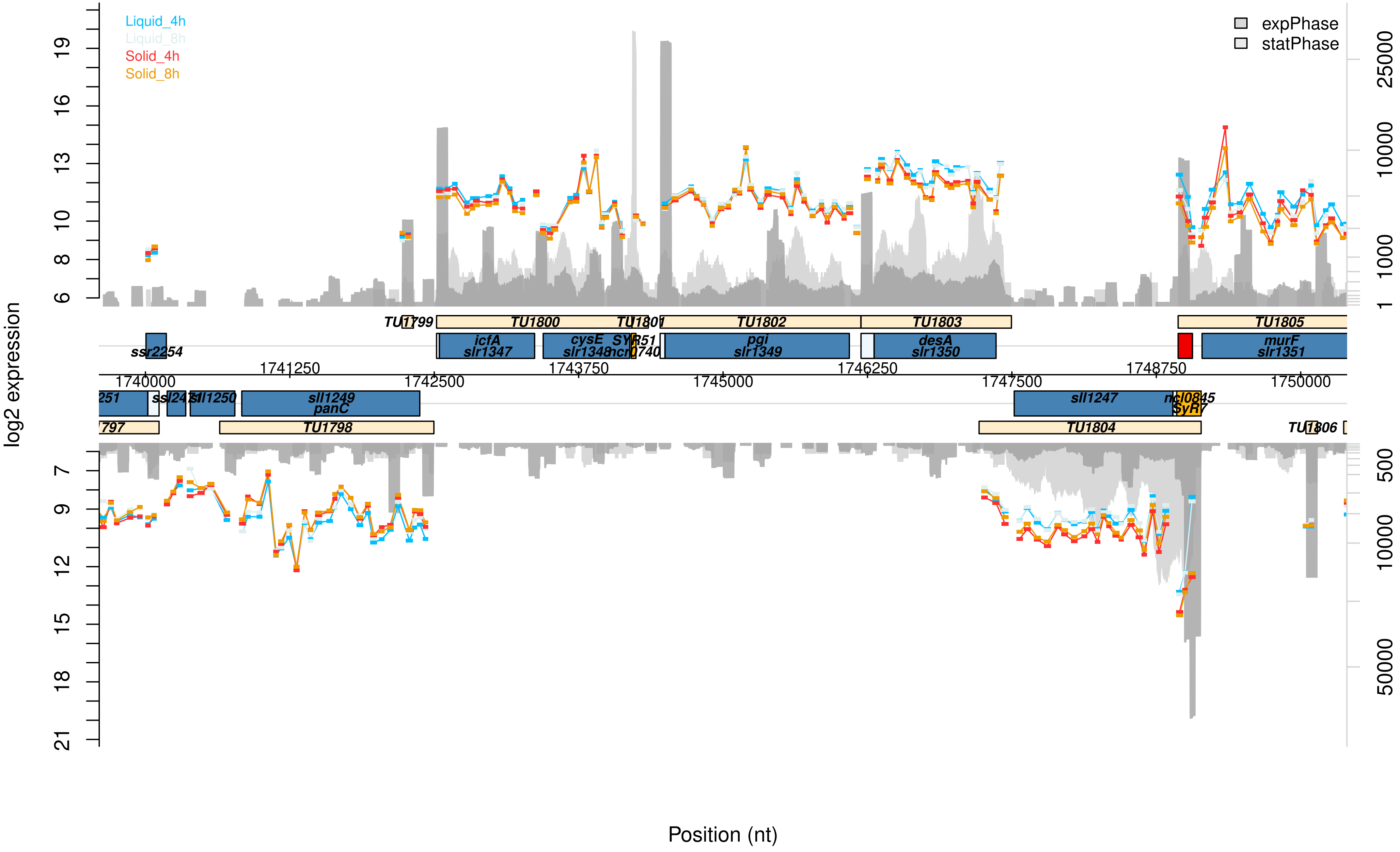

Liquid_4h
Liquid_8h
Solid_4h
Solid_8h
expPhase
statPhase
log2 expression
Position (nt)
TU1799
TU1800
TU1801
TU1802
TU1803
TU1805
ssr2254
icfA
slr1347
cysE
slr1348
pgi
slr1349
desA
slr1350
murF
slr1351
1740000
1741250
1742500
1743750
1745000
1746250
1747500
1748750
1750000
sll1250
sll1249
panC
sll1247
TU1798
TU1804
TU1806

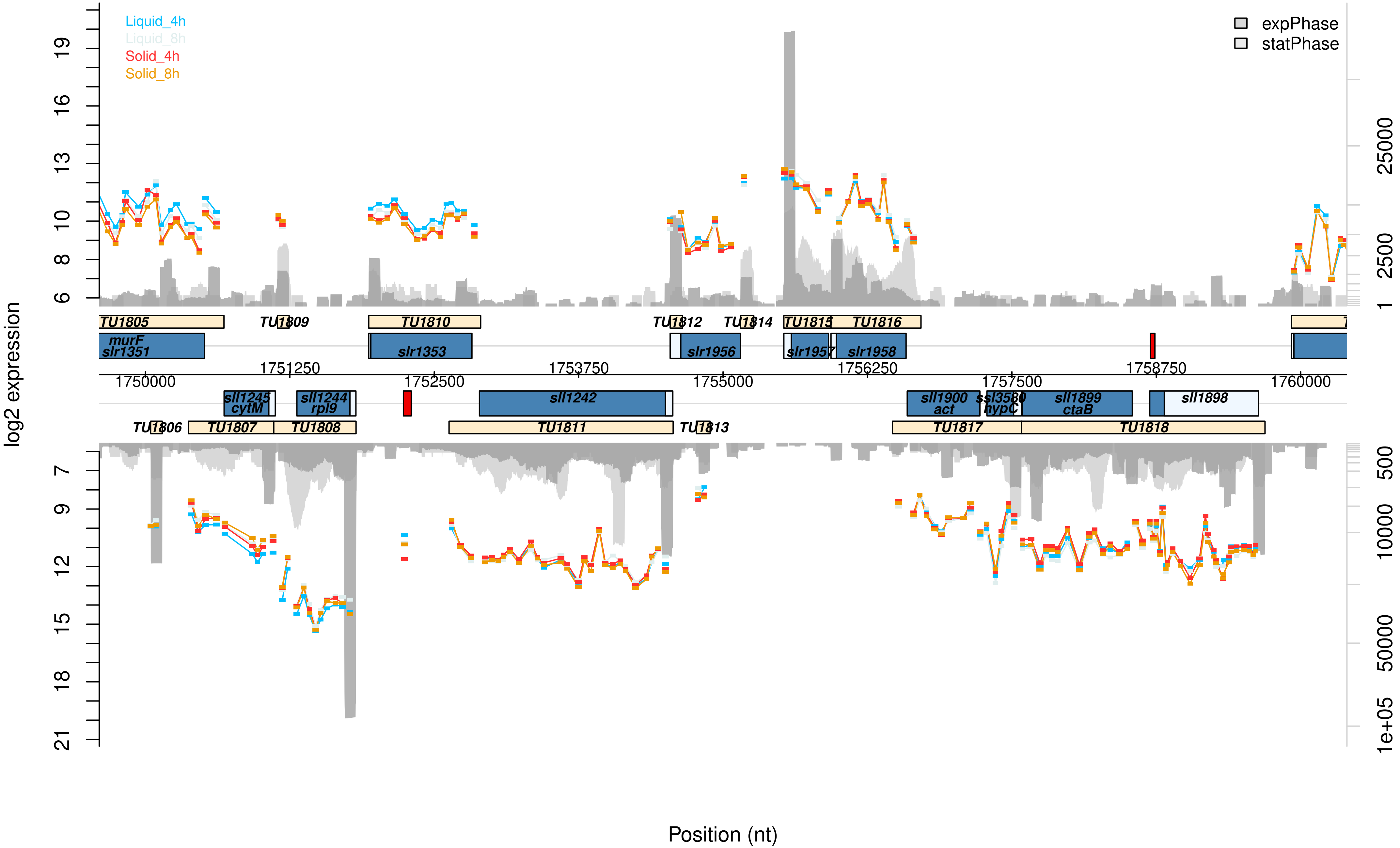

Liquid_4h
Liquid_8h
Solid_4h
Solid_8h
expPhase
statPhase
log2 expression
Position (nt)
TU1805
TU1809
TU1810
TU1812
TU1814
TU1815
TU1816
murF
slr1351
slr1353
slr1956
slr1957
slr1958
1750000
1751250
1752500
1753750
1755000
1756250
1757500
1758750
1760000
sll1245
cytM
sll1244
rpl9
sll1242
sll1900
act
ssl3580
hypC
sll1899
ctaB
sll1898
TU1806
TU1807
TU1808
TU1811
TU1813
TU1817
TU1818

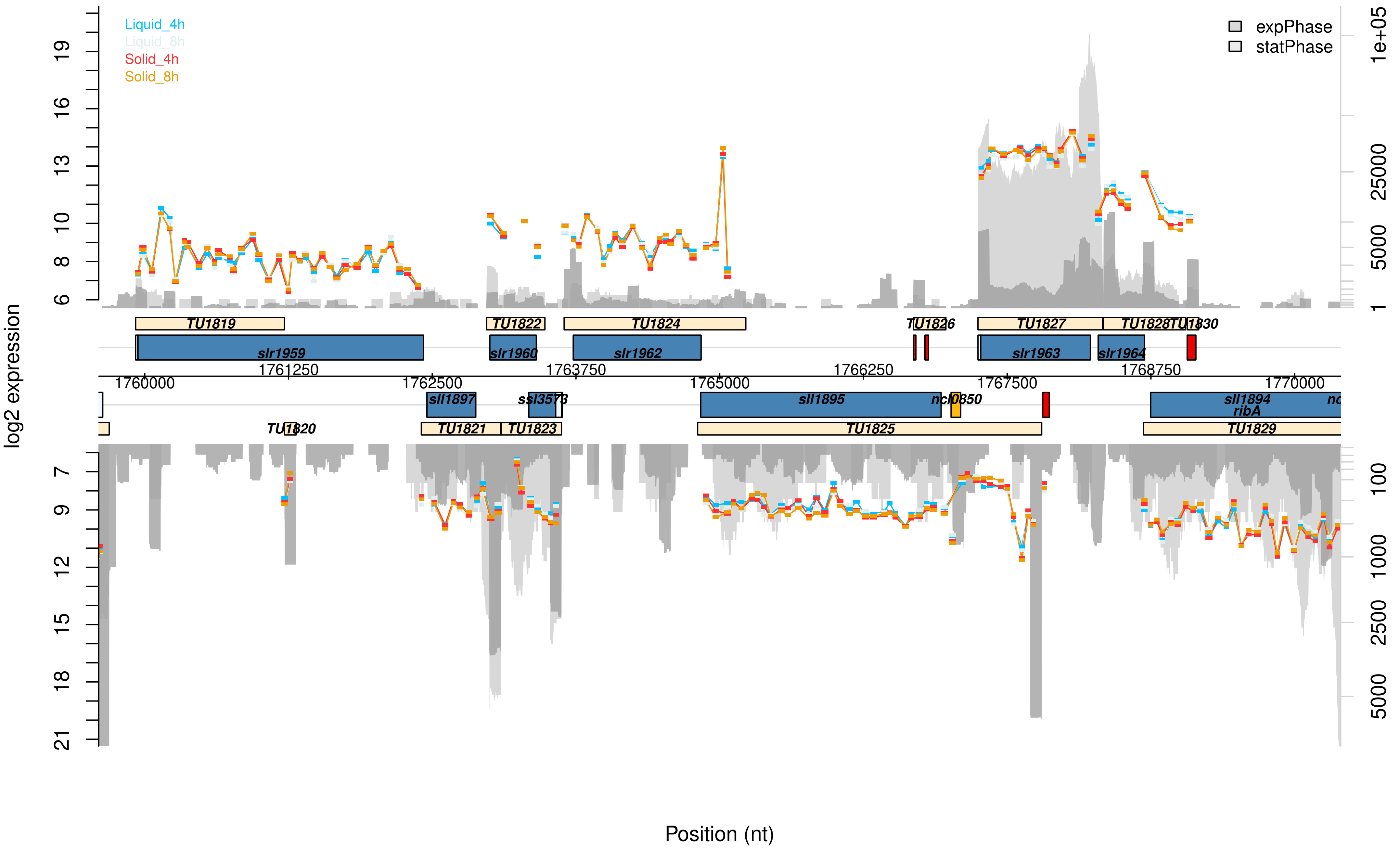

Liquid_4h
Liquid_8h
Solid_4h
Solid_8h
expPhase
statPhase
log2 expression
TU1819
TU1822
TU1824
TU1826
TU1827
TU1828
TU1830
slr1959
slr1960
slr1962
slr1963
slr1964
1761250
1763750
1766250
1768750
1760000
1762500
1765000
1767500
1770000
sll1897
ssl3573
sll1895
ncl0850
sll1894
ribA
TU1820
TU1821
TU1823
TU1825
TU1829
Position (nt)

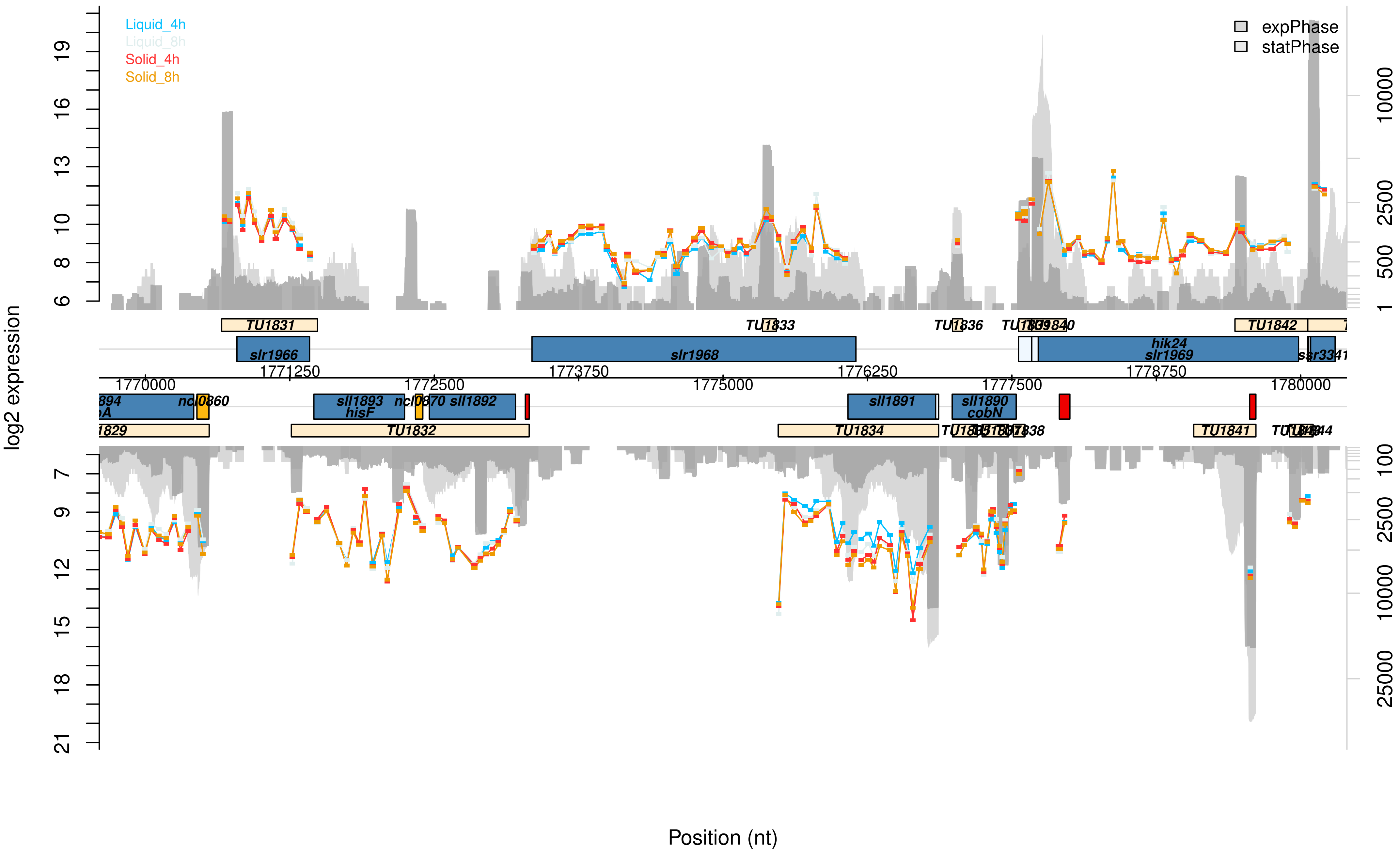
Liquid_4h
Liquid_8h
Solid_4h
Solid_8h
expPhase
statPhase
log2 expression
Position (nt)
TU1831
TU1833
TU1836
TU1842
slr1966
slr1968
hik24
slr1969
ssr3341
1770000
1771250
1772500
1773750
1775000
1776250
1777500
1778750
1780000
sll1893
hisF
sll1892
sll1891
sll1890
cobN
TU1832
TU1834
TU1841

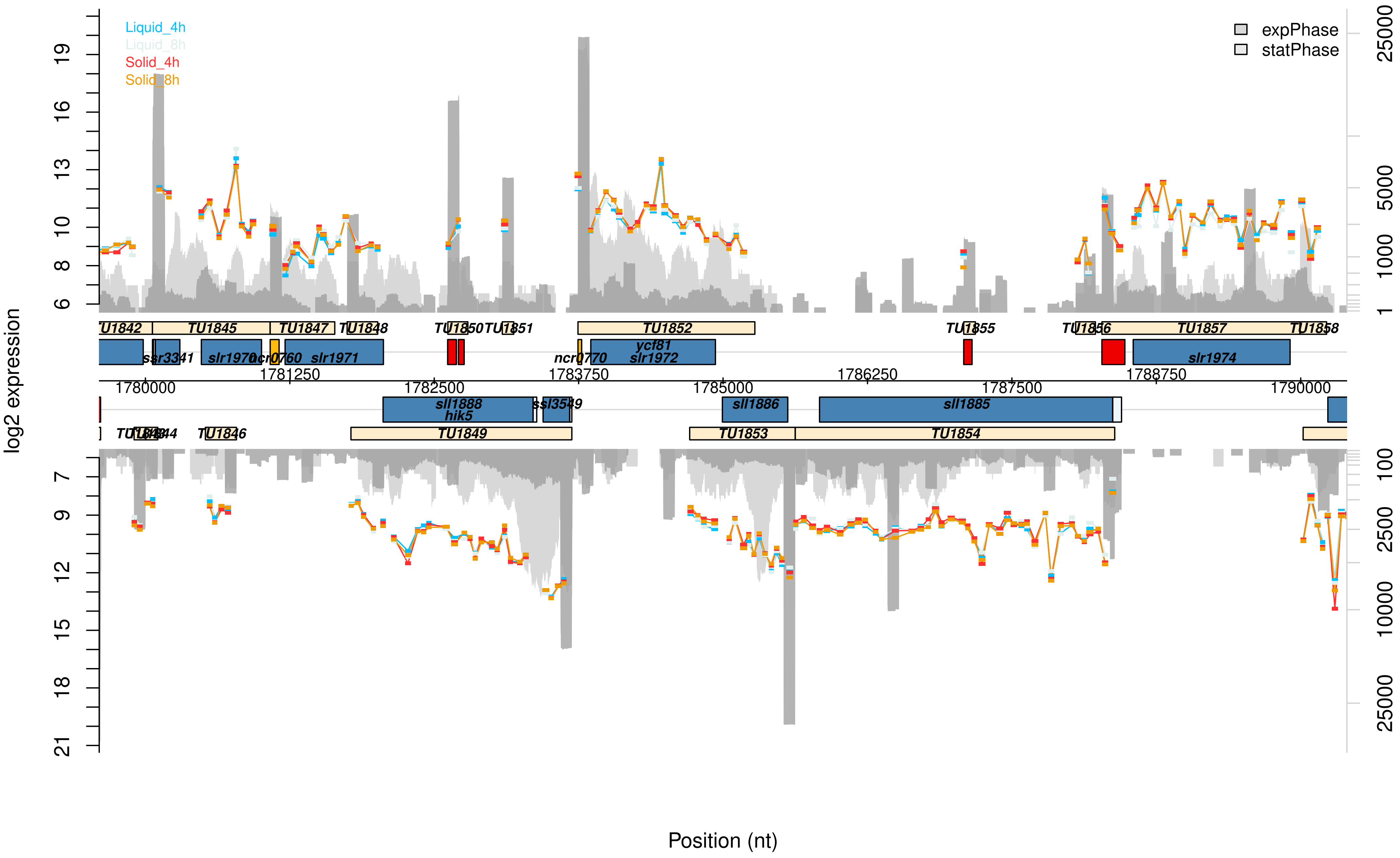

Liquid_4h
Liquid_8h
Solid_4h
Solid_8h
expPhase
statPhase
log2 expression
Position (nt)
TU1842
TU1845
TU1847
TU1848
TU1850
TU1851
TU1852
TU1855
TU1856
TU1857
TU1858
ssr3341
slr1970
ncr0760
slr1971
ncr0770
ycf81
slr1972
slr1974
sll1888
hik5
ssl3549
sll1886
sll1885
TU1846
TU1849
TU1853
TU1854
1780000
1781250
1782500
1783750
1785000
1786250
1787500
1788750
1790000

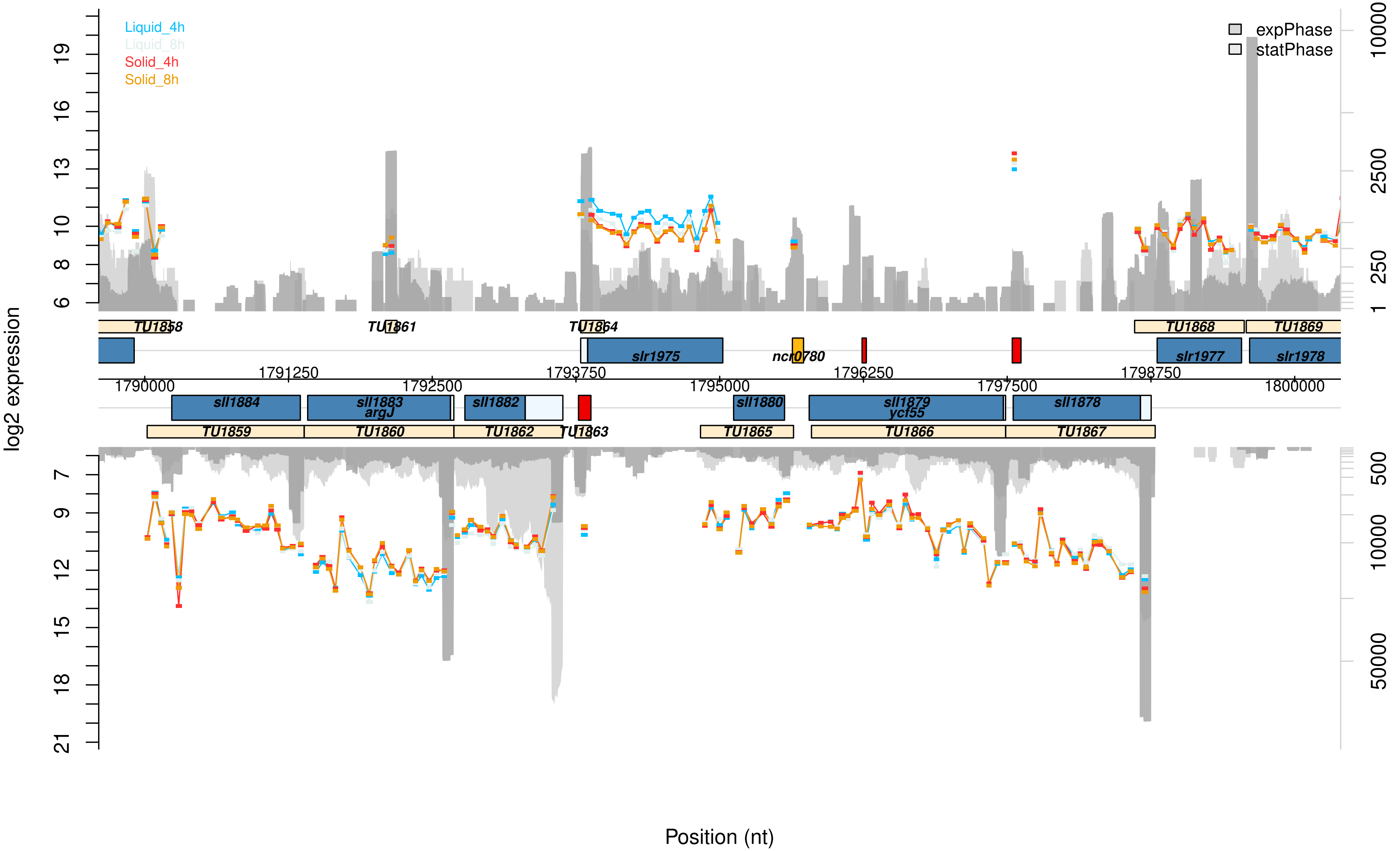

Liquid_4h
Liquid_8h
Solid_4h
Solid_8h
expPhase
statPhase
log2 expression
Position (nt)
TU1858
TU1861
TU1864
TU1868
TU1869
slr1975
ncr0780
slr1977
slr1978
1790000
1791250
1792500
1793750
1795000
1796250
1797500
1798750
1800000
sll1884
sll1883
argJ
sll1882
sll1880
sll1879
ycf55
sll1878
TU1859
TU1860
TU1862
TU1863
TU1865
TU1866
TU1867

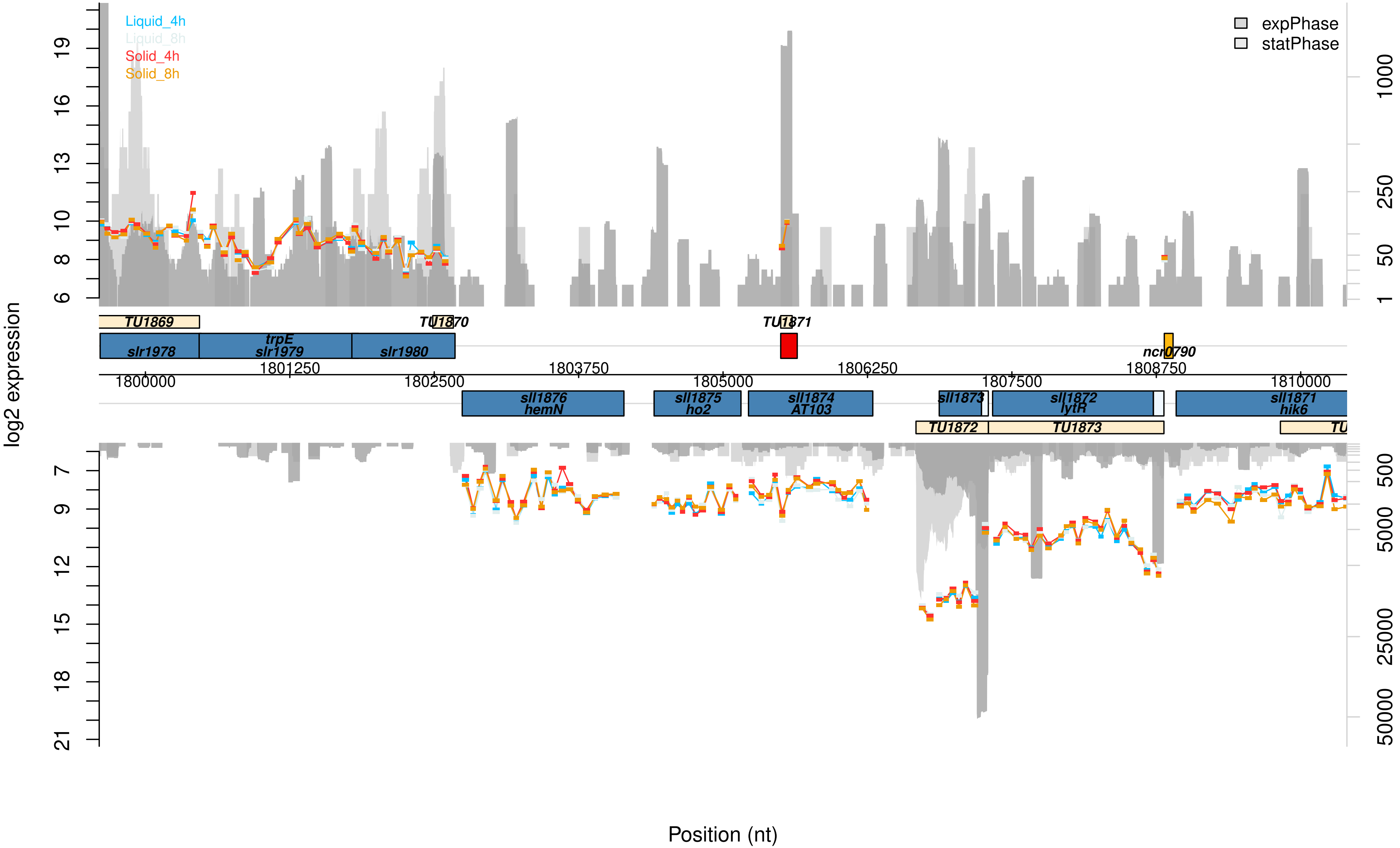

Liquid_4h
Liquid_8h
Solid_4h
Solid_8h
expPhase
statPhase
log2 expression
Position (nt)
TU1869
TU1870
TU1871
TU1872
TU1873
slr1978
trpE
slr1979
slr1980
ncr0790
sll1876
hemN
sll1875
ho2
sll1874
AT103
sll1873
sll1872
lytR
sll1871
hik6
1800000
1801250
1802500
1803750
1805000
1806250
1807500
1808750
1810000

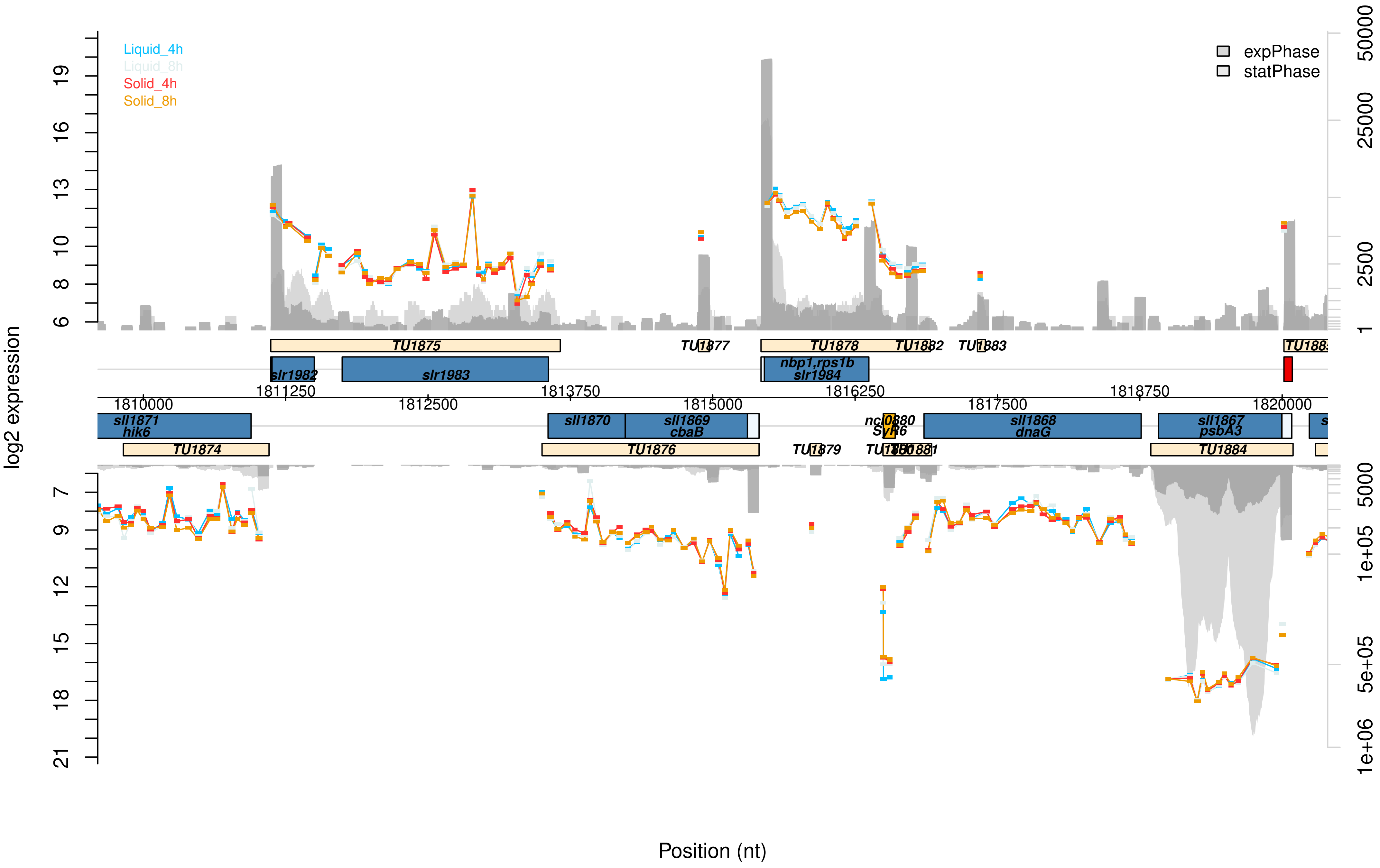

Liquid_4h
Liquid_8h
Solid_4h
Solid_8h
expPhase
statPhase
log2 expression
Position (nt)
TU1875
TU1877
TU1878
TU1882
TU1883
slr1982
slr1983
nbp1,rps1b
slr1984
sll1871
hik6
sll1870
sll1869
cbaB
ncl0880
SyR6
sll1868
dnaG
sll1867
psbA3
TU1874
TU1876
TU1879
TU1884
1810000
1811250
1812500
1813750
1815000
1816250
1817500
1818750
1820000

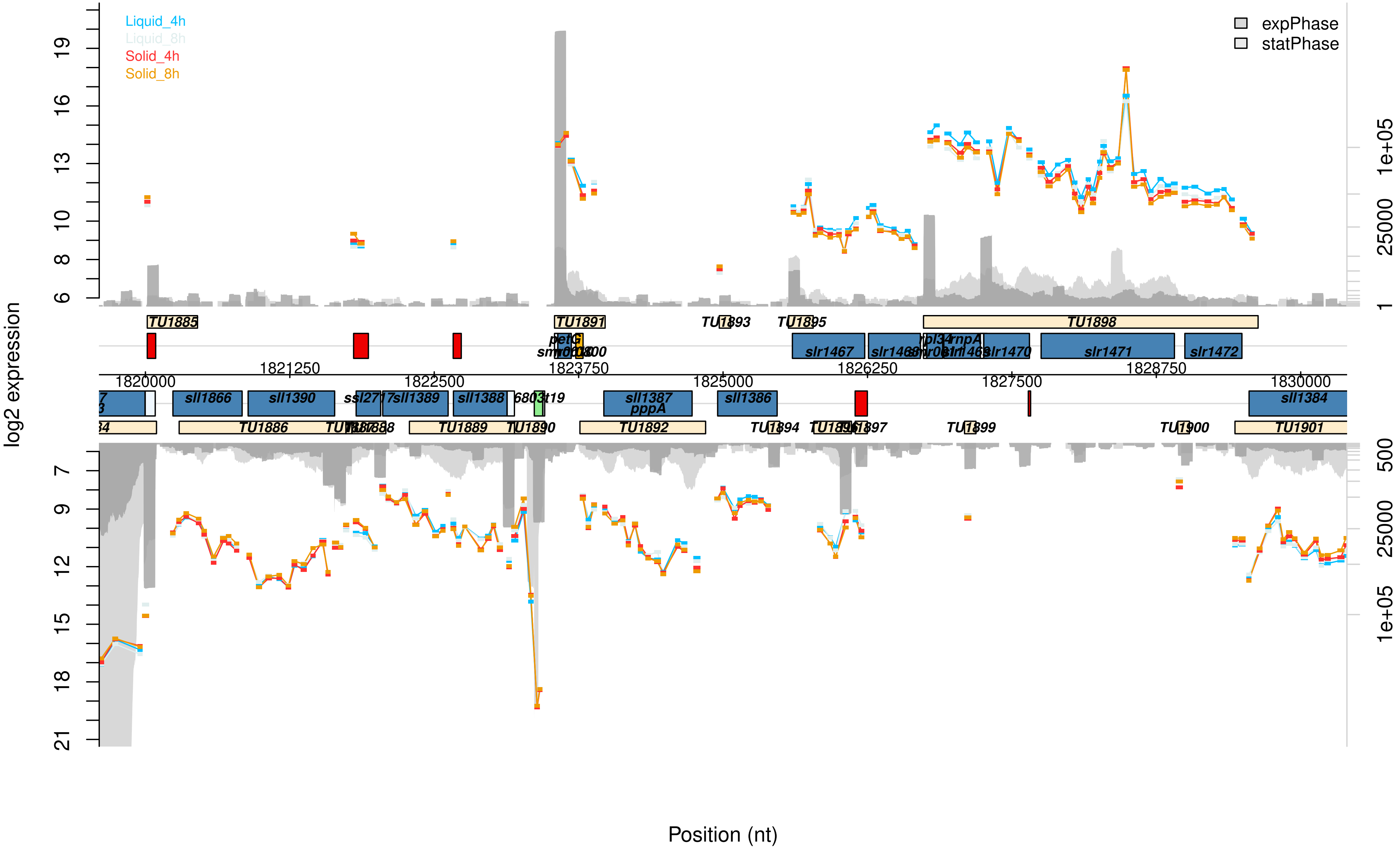
Liquid_4h
Liquid_8h
Solid_4h
Solid_8h
expPhase
statPhase
log2 expression
Position (nt)
TU1885
TU1891
TU1893
TU1895
TU1898
slr1467
slr1471
slr1472
slr1470
petG
sll1866
sll1390
sll1389
sll1388
sll1387
pppA
sll1386
sll1384
TU1886
TU1889
TU1890
TU1892
TU1894
TU1897
TU1899
TU1900
TU1901
1820000
1821250
1822500
1823750
1825000
1826250
1827500
1828750
1830000

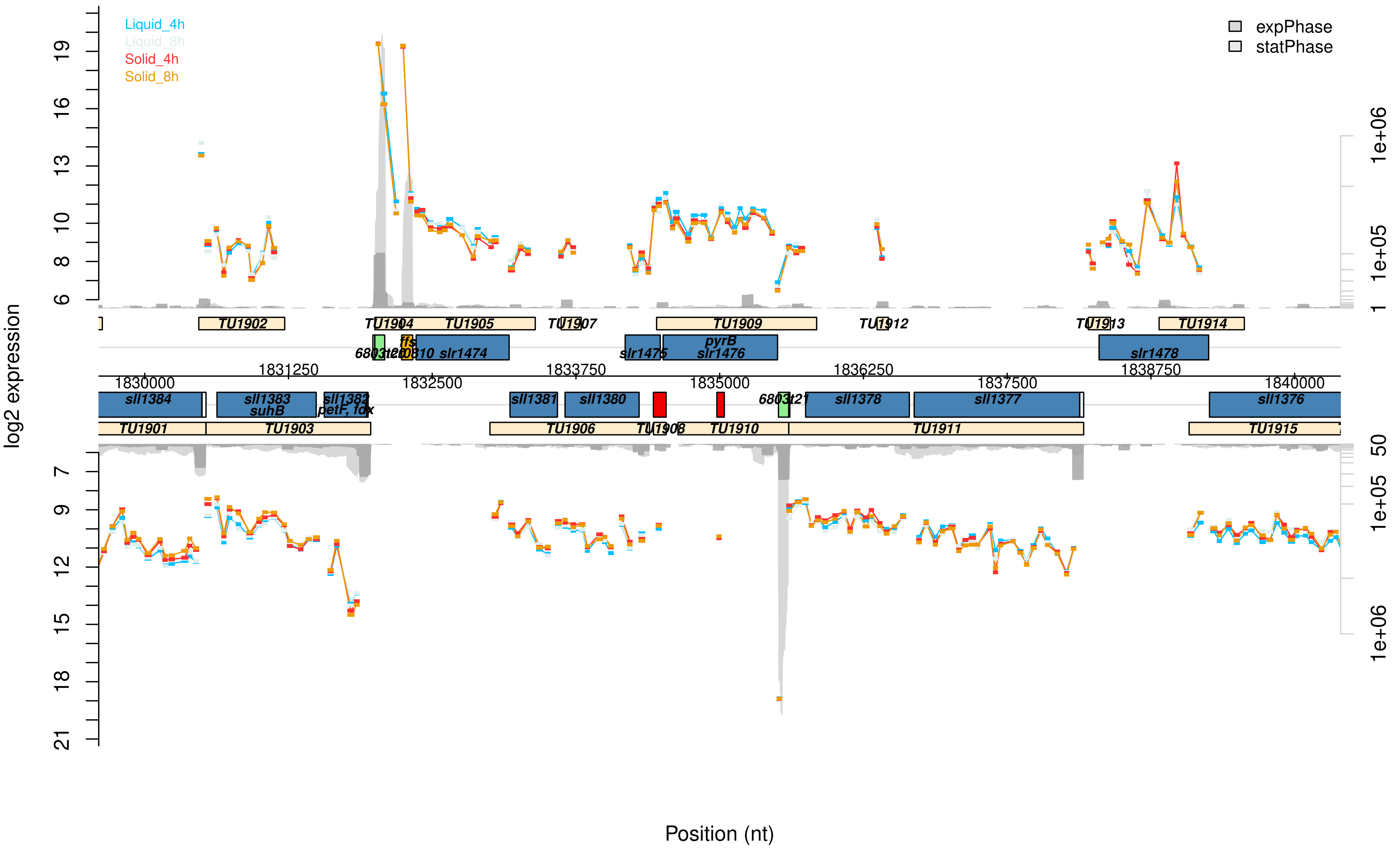

Liquid_4h
Liquid_8h
Solid_4h
Solid_8h
expPhase
statPhase
log2 expression
Position (nt)
TU1901
TU1902
TU1903
TU1904
TU1905
TU1906
TU1907
TU1908
TU1909
TU1910
TU1911
TU1912
TU1913
TU1914
TU1915
sll1384
sll1383
suhB
sll1382
petF, fdx
sll1381
sll1380
sll1378
sll1377
sll1376
slr1474
slr1475
pyrB
slr1476
slr1478
ffs
1830000
1831250
1832500
1833750
1835000
1836250
1837500
1838750
1840000

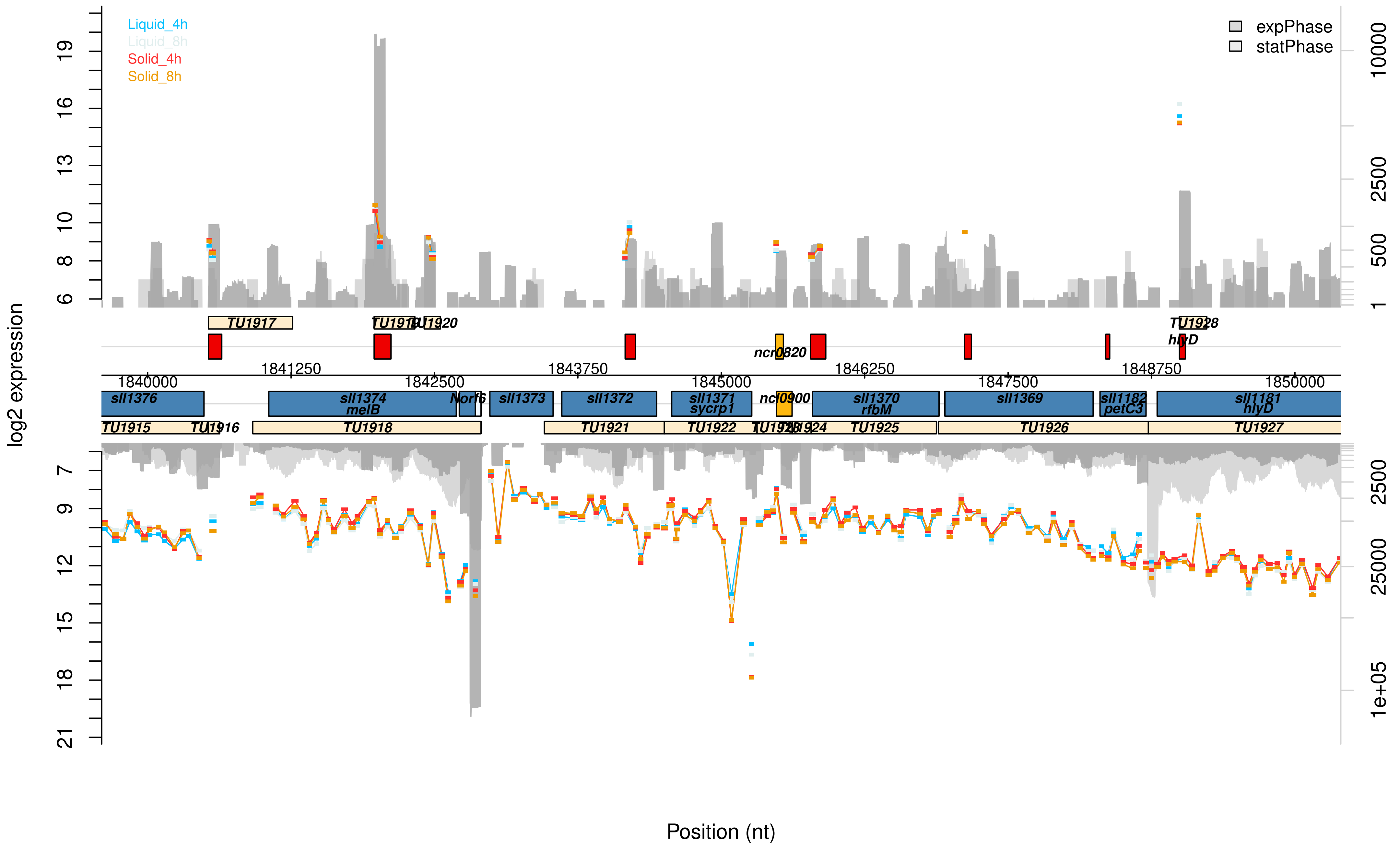

Liquid_4h
Liquid_8h
Solid_4h
Solid_8h
expPhase
statPhase
log2 expression
Position (nt)
TU1917
TU1919
TU1920
TU1928
ncr0820
hlyD
1840000
1841250
1842500
1843750
1845000
1846250
1847500
1848750
1850000
sll1376
sll1374
melB
Norf6
sll1373
sll1372
sll1371
sycrp1
ncl0900
sll1370
rfbM
sll1369
sll1182
petC3
sll1181
hlyD
TU1915
TU1916
TU1918
TU1921
TU1922
TU1923
TU1924
TU1925
TU1926
TU1927

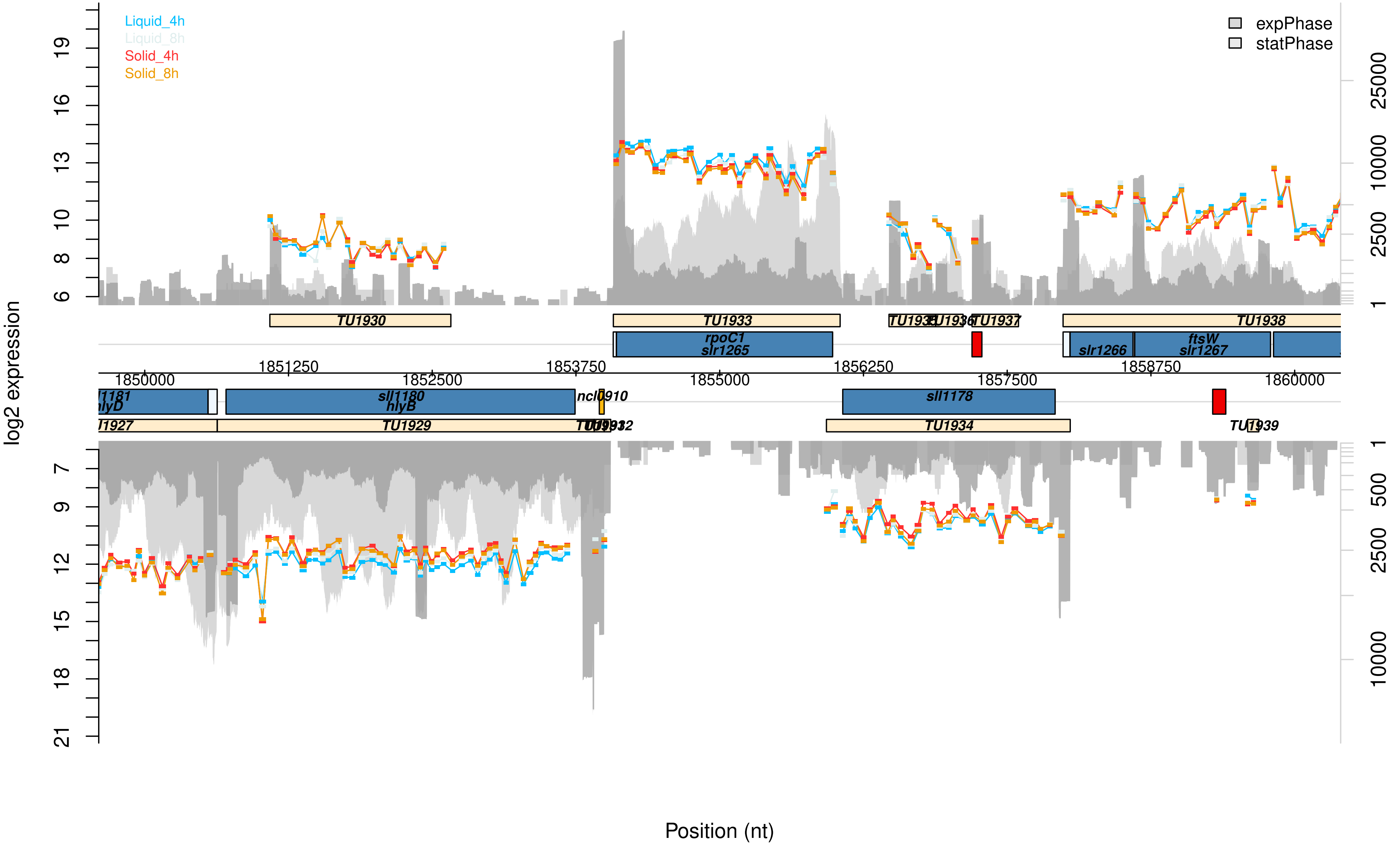

Liquid_4h
Liquid_8h
Solid_4h
Solid_8h
expPhase
statPhase
log2 expression
Position (nt)
TU1930
TU1933
TU1937
TU1938
rpoC1
slr1265
slr1266
ftsW
slr1267
1850000
1851250
1852500
1853750
1855000
1856250
1857500
1858750
1860000
hlyD
sll1180
hlyB
ncl0910
sll1178
TU1929
TU1934

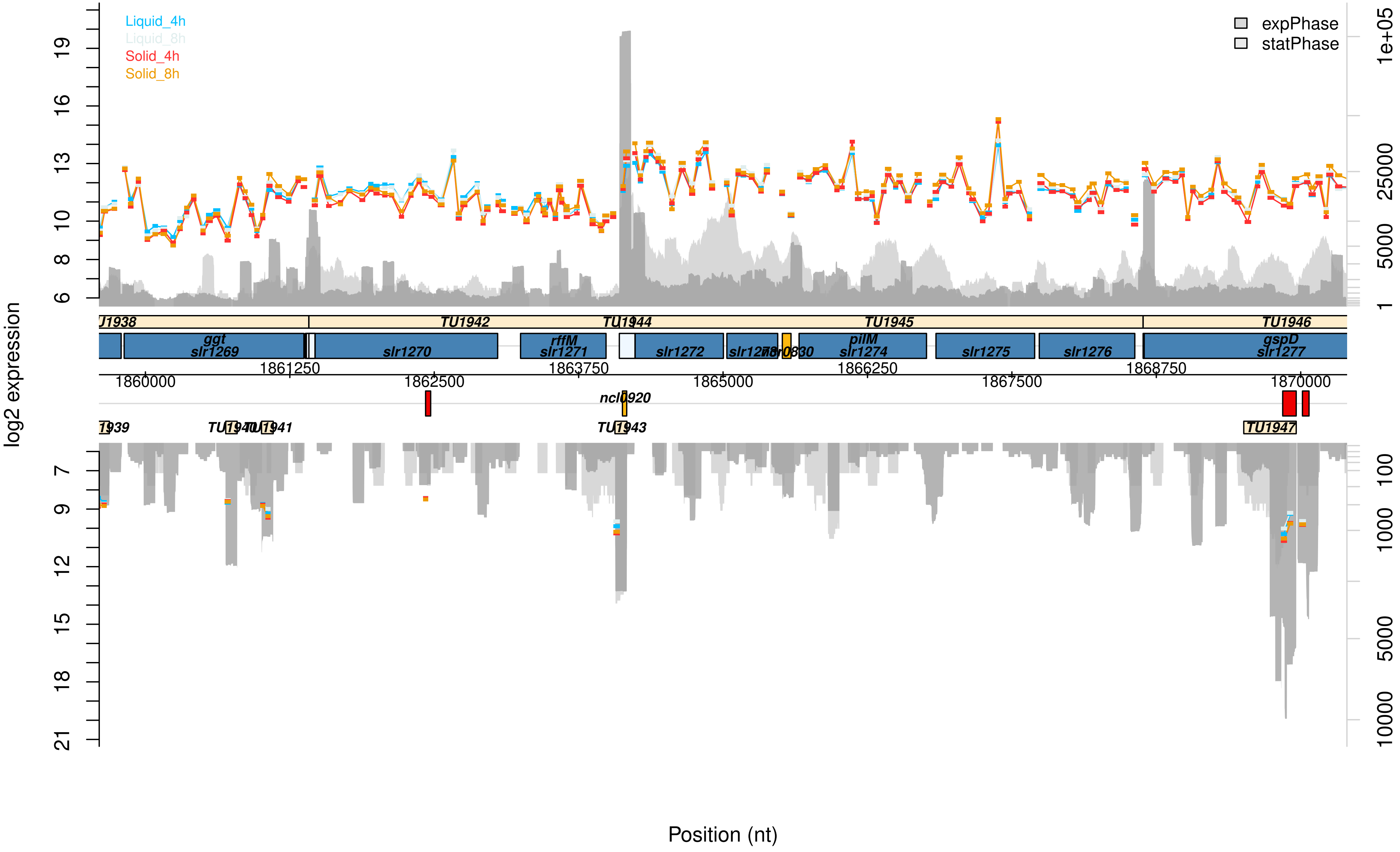

Liquid_4h
Liquid_8h
Solid_4h
Solid_8h
expPhase
statPhase
log2 expression
TU1942
TU1944
TU1945
TU1946
ggt
slr1269
slr1270
rffM
slr1271
slr1272
pilM
slr1274
slr1275
slr1276
gspD
slr1277
1860000
1861250
1862500
1863750
1865000
1866250
1867500
1868750
1870000
ncl0920
TU1943
TU1947
Position (nt)

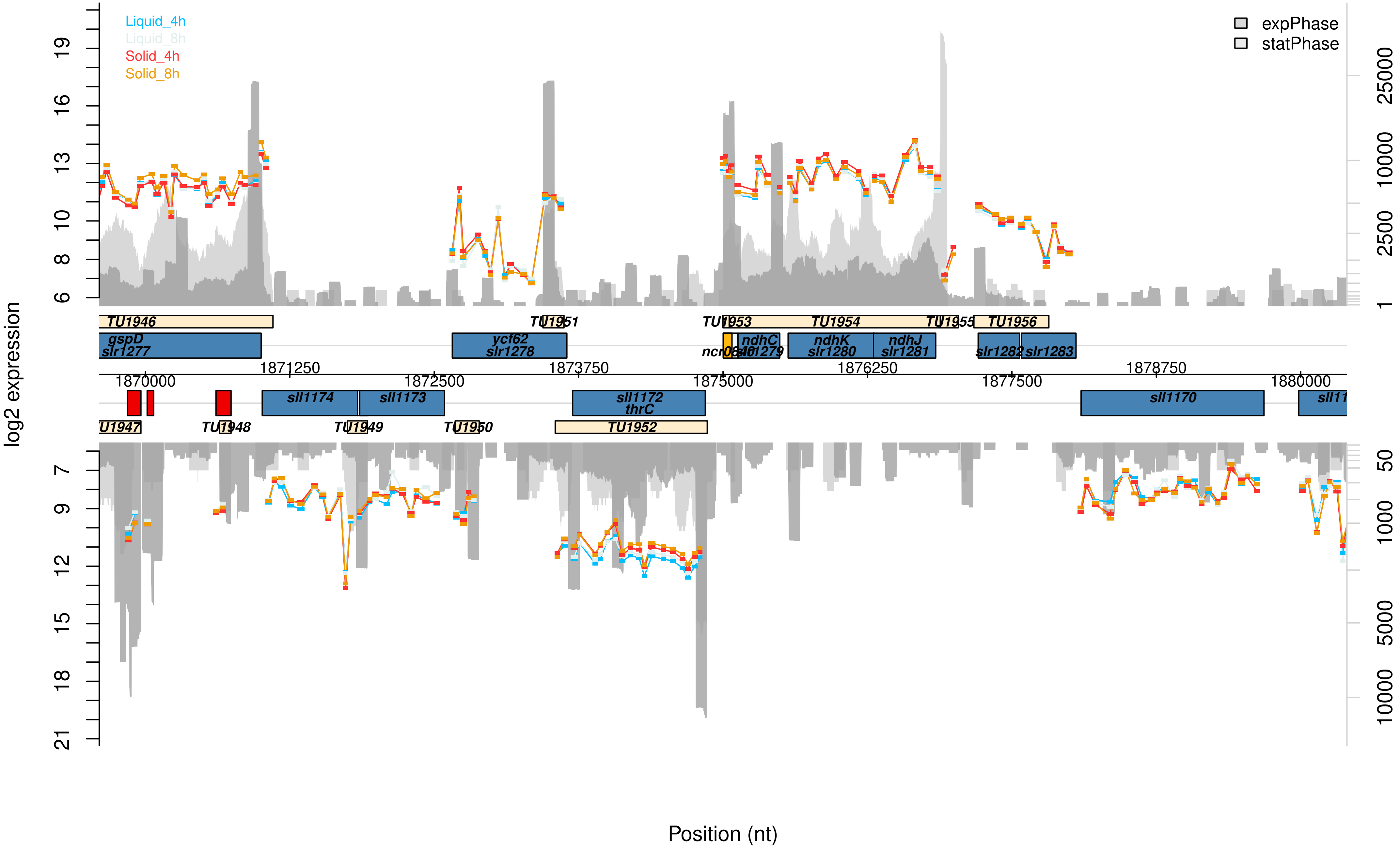

Liquid_4h
Liquid_8h
Solid_4h
Solid_8h
expPhase
statPhase
log2 expression
Position (nt)
TU1946
TU1951
TU1953
TU1954
TU1955
TU1956
gspD
slr1277
ycf62
slr1278
ndhC
slr1279
ndhK
slr1280
ndhJ
slr1281
slr1282
slr1283
1870000
1871250
1872500
1873750
1875000
1876250
1877500
1878750
1880000
sll1174
sll1173
sll1172
thrC
sll1170
TU1948
TU1949
TU1950
TU1952

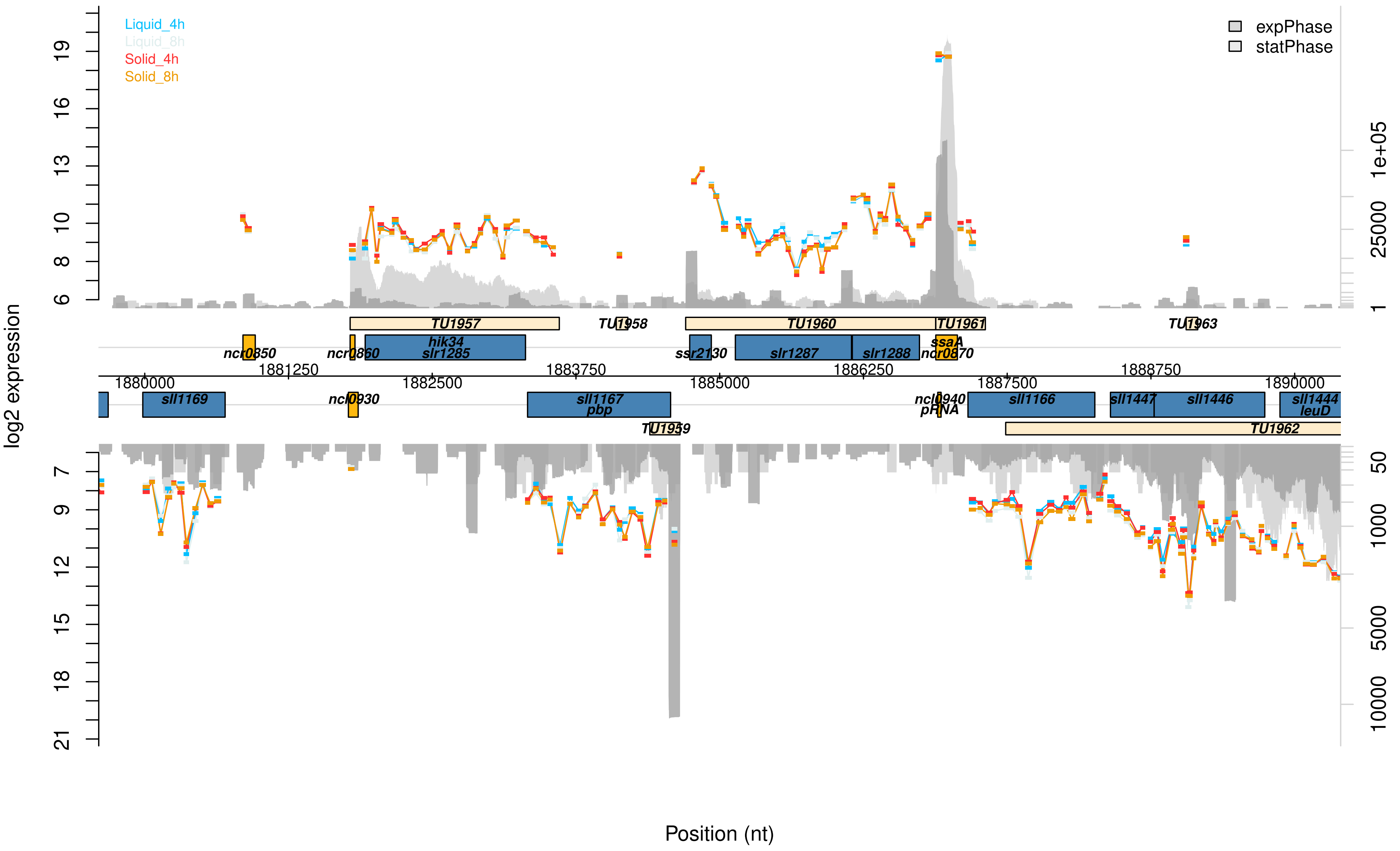

Liquid_4h
Liquid_8h
Solid_4h
Solid_8h
expPhase
statPhase
log2 expression
Position (nt)
TU1957
TU1958
TU1960
TU1961
TU1963
ncr0850
ncr0860
hik34
slr1285
ssr2130
slr1287
slr1288
ssaA
ncr0870
1880000
1881250
1882500
1883750
1885000
1886250
1887500
1888750
1890000
sll1169
ncl0930
sll1167
pbp
ncl0940
pRNA
sll1166
sll1447
sll1446
sll1444
leuD
TU1959
TU1962

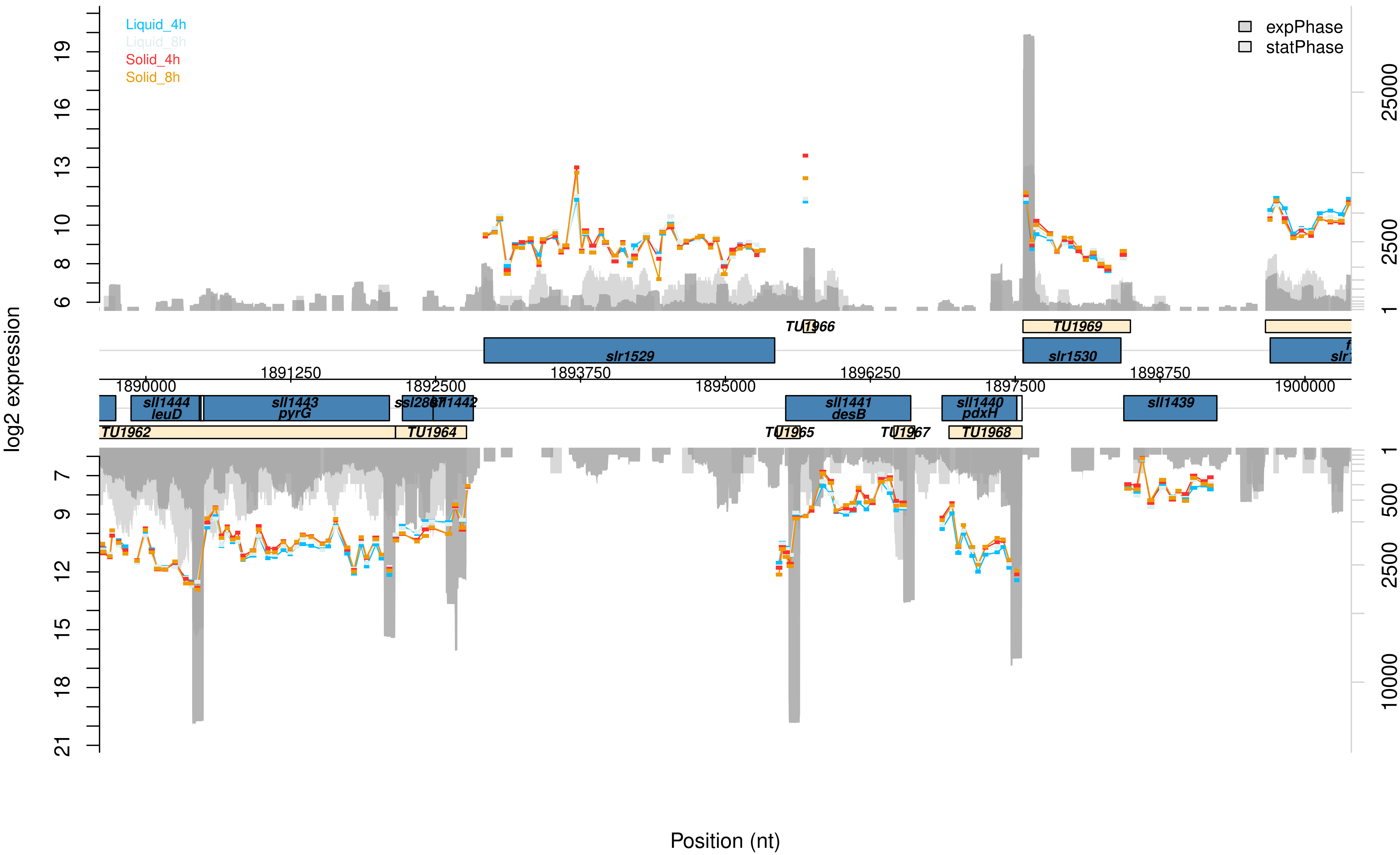

Liquid_4h
Liquid_8h
Solid_4h
Solid_8h
expPhase
statPhase
log2 expression
Position (nt)
TU1966
TU1969
slr1529
slr1530
1891250
1893750
1896250
1898750
1890000
1892500
1895000
1897500
1900000
sll1444
leuD
sll1443
pyrG
ssl2867
sll1442
sll1441
desB
sll1440
pdxH
sll1439
TU1962
TU1964
TU1965
TU1967
TU1968

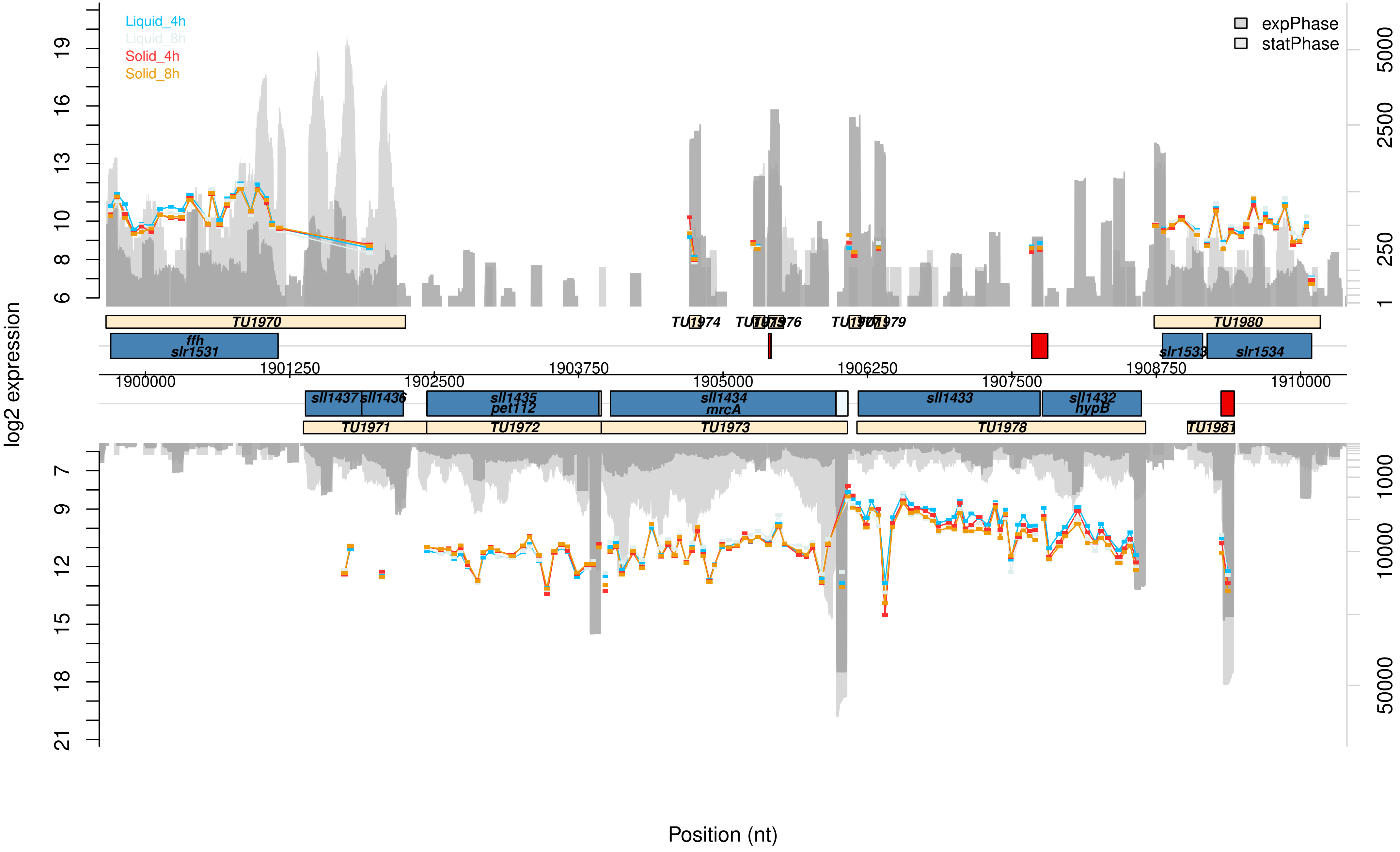

Liquid_4h
Liquid_8h
Solid_4h
Solid_8h
expPhase
statPhase
log2 expression
TU1970
TU1974
TU1979
TU1980
ffh
slr1531
slr1533
slr1534
1900000
1901250
1902500
1903750
1905000
1906250
1907500
1908750
1910000
sll1437
sll1436
sll1435
pet112
sll1434
mrcA
sll1433
sll1432
hypB
TU1971
TU1972
TU1973
TU1978
TU1981
Position (nt)

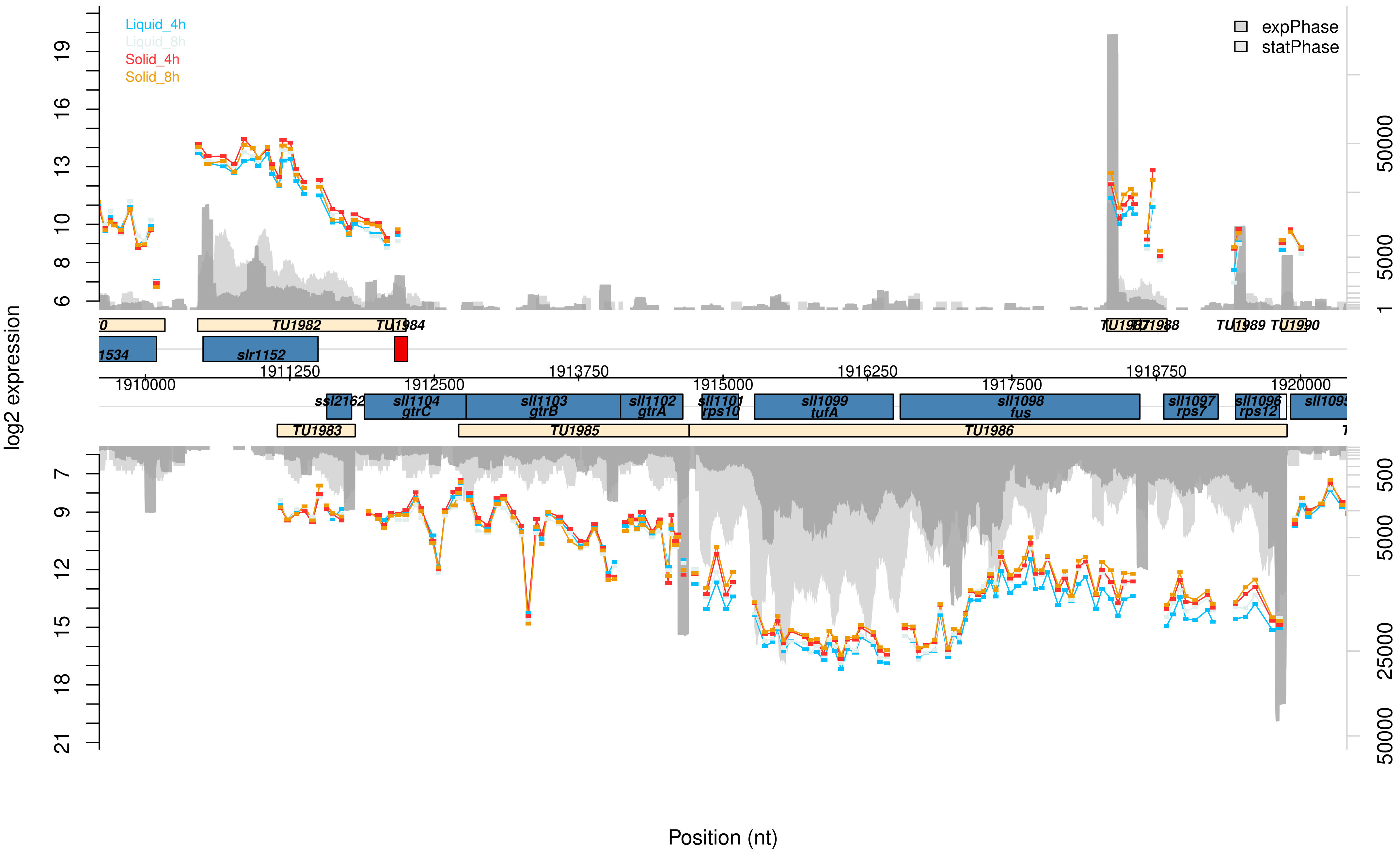

Liquid_4h
Liquid_8h
Solid_4h
Solid_8h
expPhase
statPhase
log2 expression
TU1982
TU1984
slr1152
1534
TU1987
TU1988
TU1989
TU1990
1910000
1911250
1912500
1913750
1915000
1916250
1917500
1918750
1920000
ssl2162
sll1104
gtrC
sll1103
gtrB
sll1102
gtrA
sll1101
rps10
sll1099
tufA
sll1098
fus
sll1097
rps7
sll1096
rps12
TU1983
TU1985
TU1986
Position (nt)

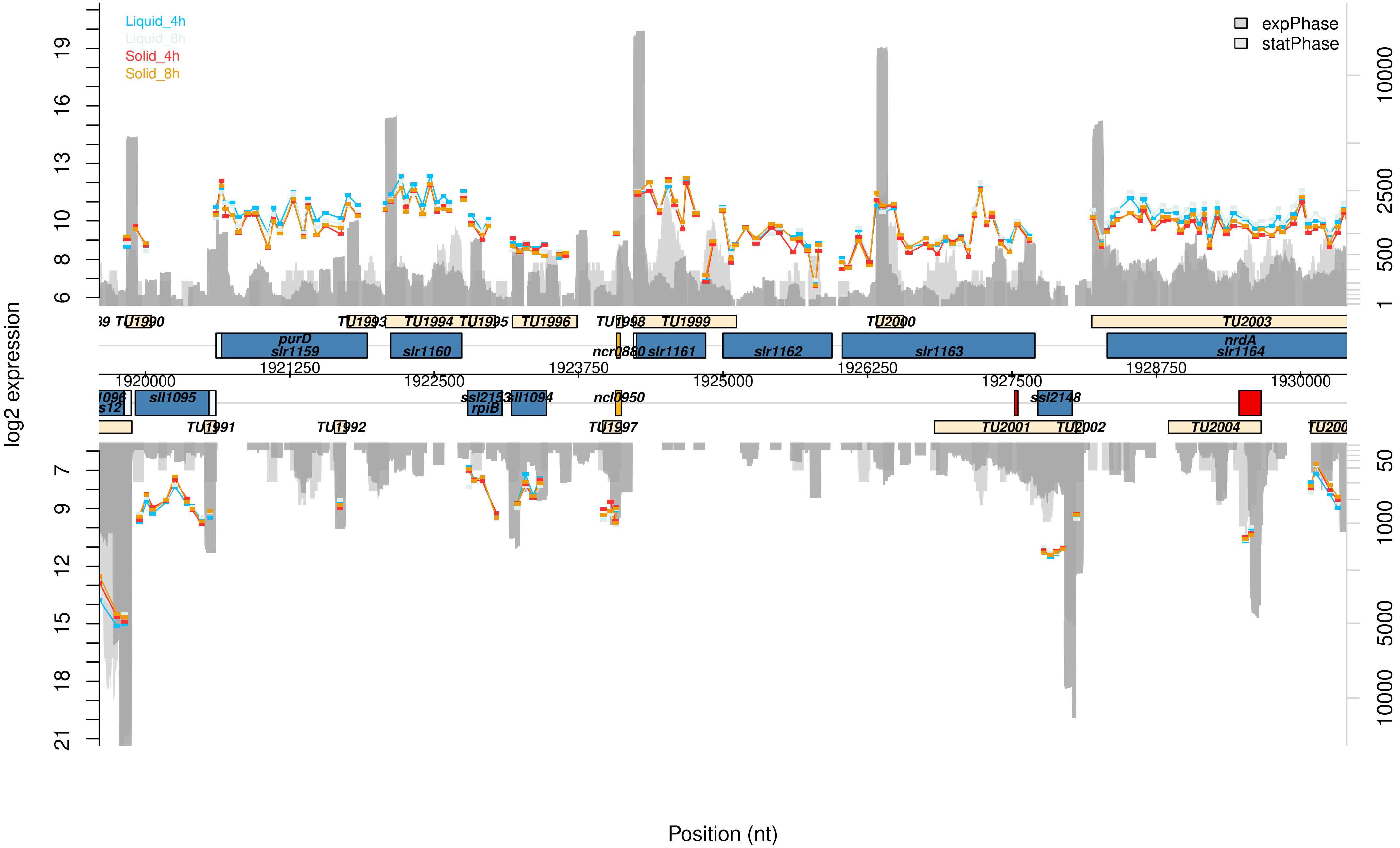

Liquid_4h
Liquid_8h
Solid_4h
Solid_8h
expPhase
statPhase
log2 expression
Position (nt)
TU1990
TU1991
TU1992
TU1993
TU1994
TU1995
TU1996
TU1997
TU1998
TU1999
TU2000
TU2001
TU2002
TU2003
TU2004
purD
slr1159
slr1160
ncr0880
slr1161
slr1162
slr1163
nrdA
slr1164
sll1095
ssl2153
rpiB
sll1094
ncl0950
ssl2148
1920000
1921250
1922500
1923750
1925000
1926250
1927500
1928750
1930000

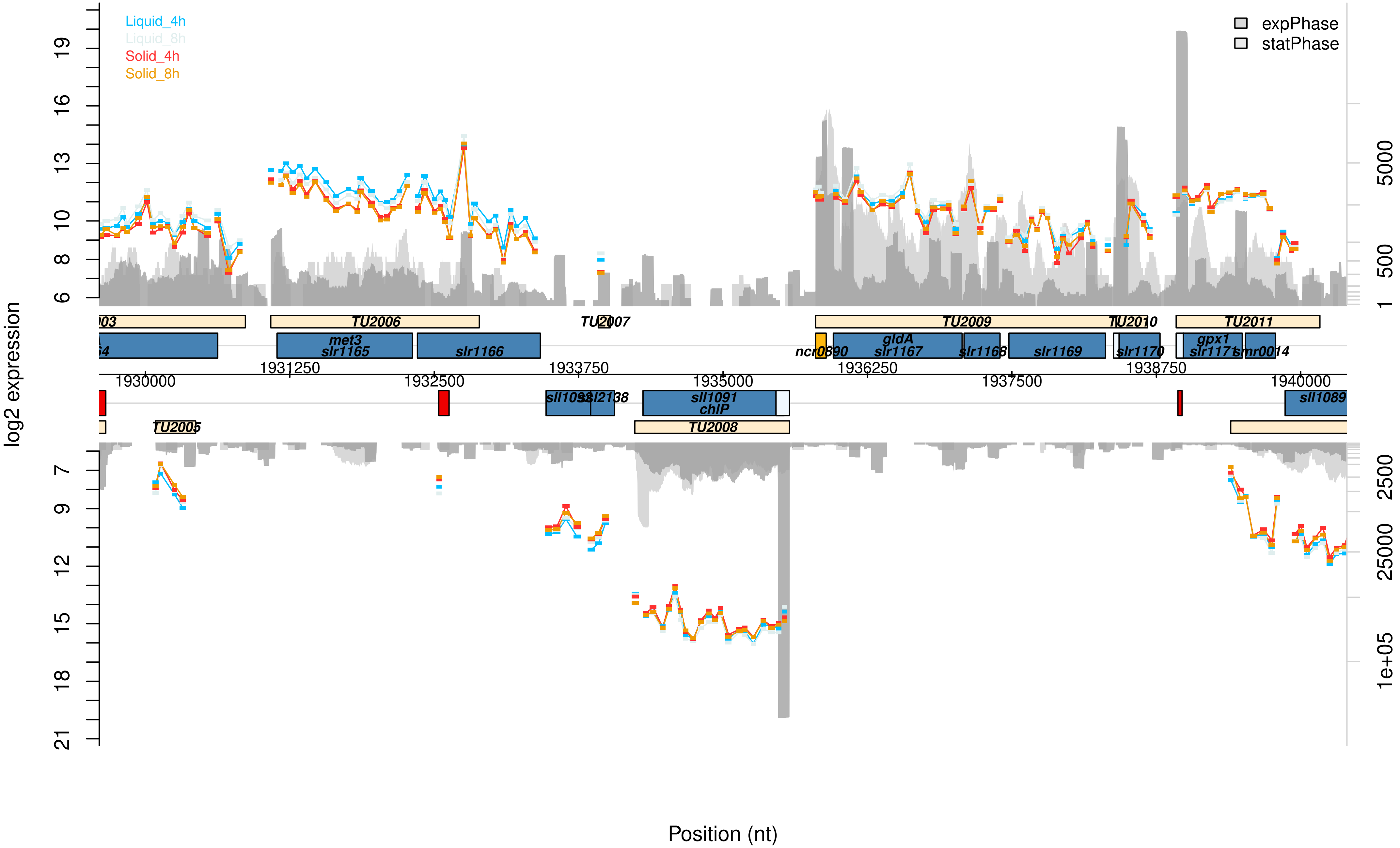

Liquid_4h
Liquid_8h
Solid_4h
Solid_8h
expPhase
statPhase
log2 expression
TU2006
TU2007
TU2009
TU2010
TU2011
met3
slr1165
slr1166
ncr0890
gldA
slr1167
slr1168
slr1169
slr1170
gpx1
slr1171
smr0014
1930000
1931250
1932500
1933750
1935000
1936250
1937500
1938750
1940000
sll1091
chlP
sll1089
TU2005
TU2008
Position (nt)

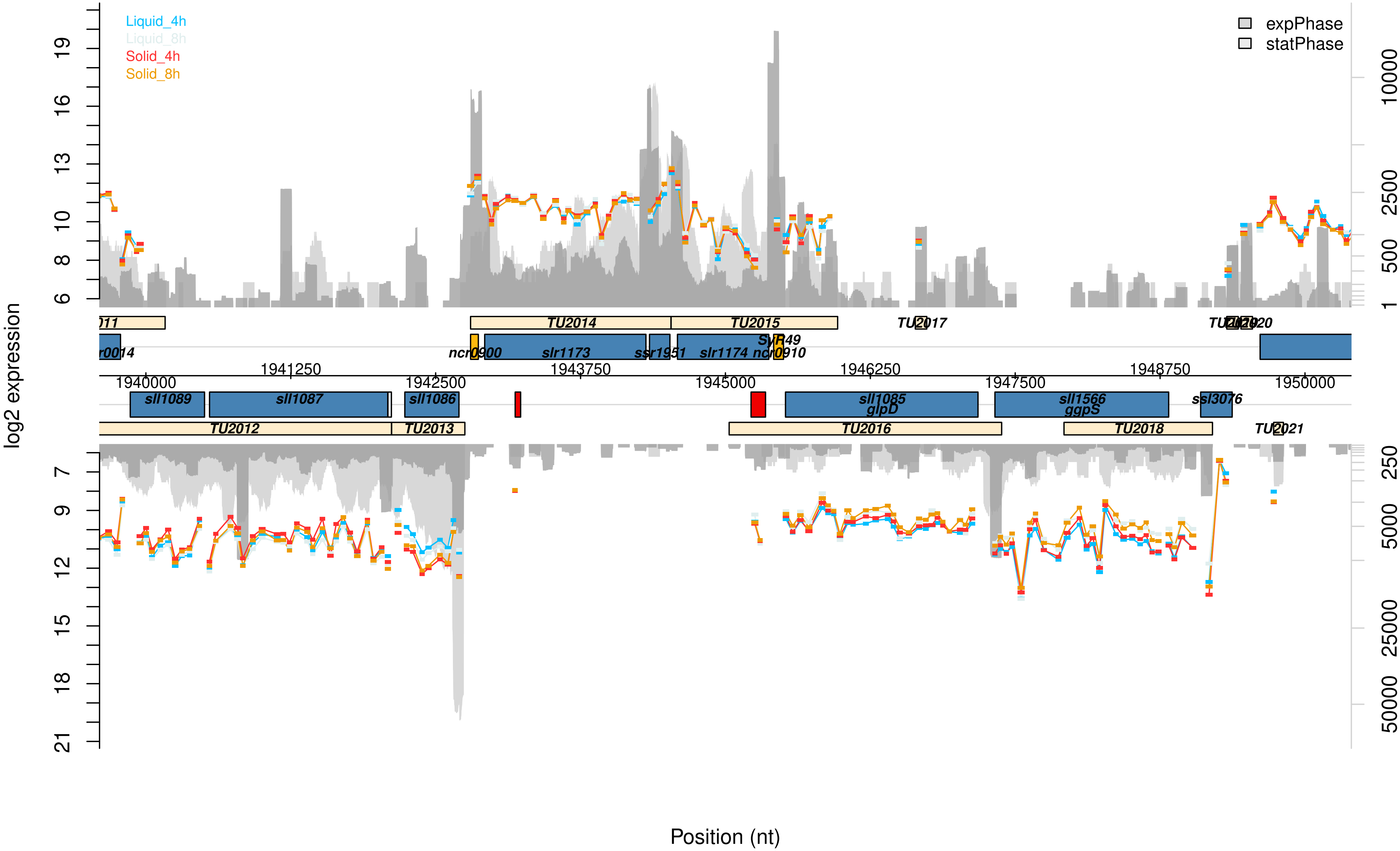

Liquid_4h
Liquid_8h
Solid_4h
Solid_8h
expPhase
statPhase
log2 expression
TU2014
TU2015
TU2017
ncr0900
slr1173
ssr1951
slr1174
ncr0910
SyR49
1941250
1943750
1946250
1948750
1940000
1942500
1945000
1947500
1950000
sll1089
sll1087
sll1086
sll1085
glpD
sll1566
ggpS
ssl3076
TU2012
TU2013
TU2016
TU2018
TU2021
Position (nt)

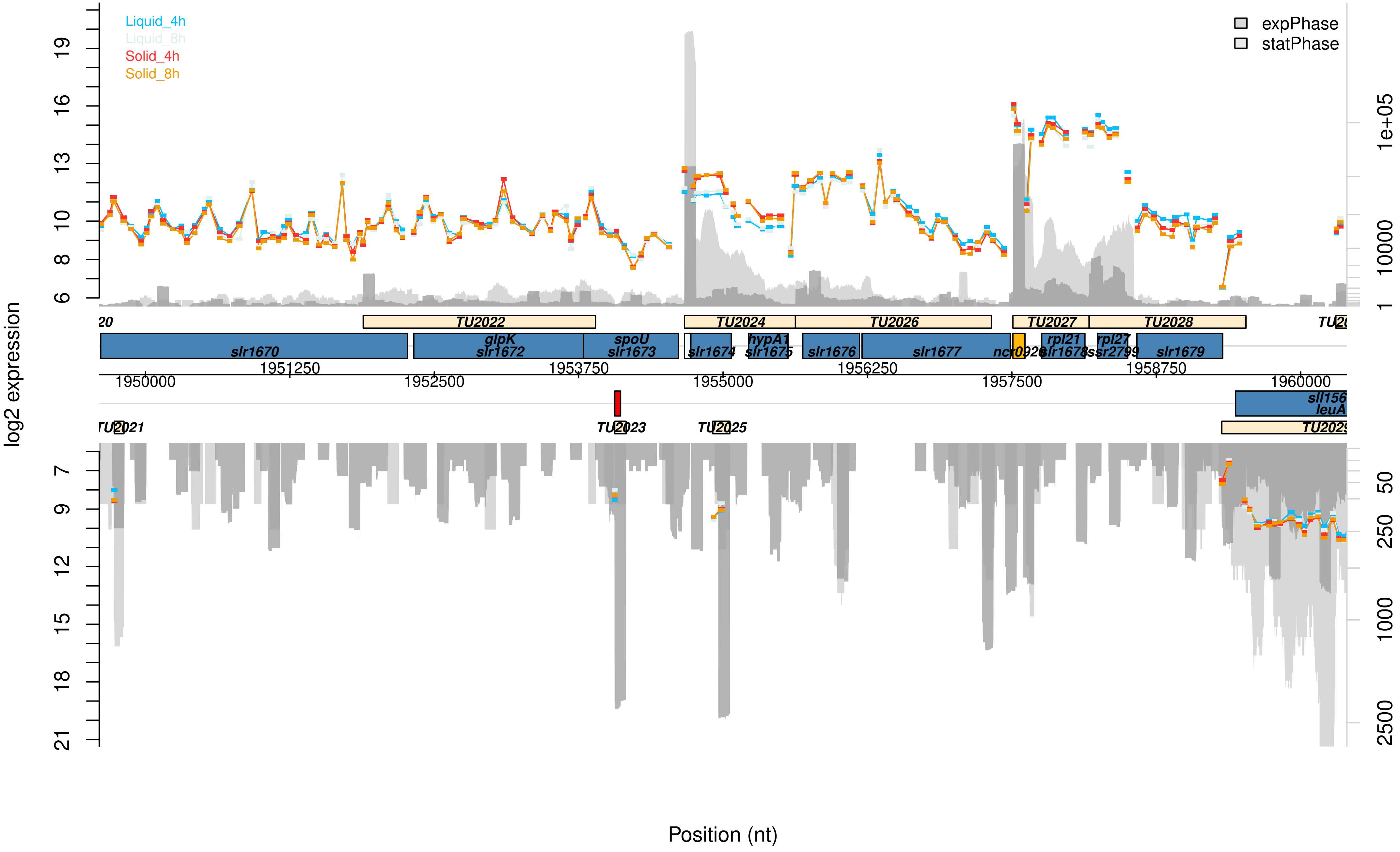
Liquid_4h
Liquid_8h
Solid_4h
Solid_8h
expPhase
statPhase
log2 expression
Position (nt)
TU2022
TU2024
TU2026
TU2027
TU2028
slr1670
glpK
slr1672
spoU
slr1673
slr1674
hypA1
slr1675
slr1676
slr1677
rpl21
slr1678
rpl27
ssr2799
slr1679
1950000
1951250
1952500
1953750
1955000
1956250
1957500
1958750
1960000
leuA
TU2023
TU2025

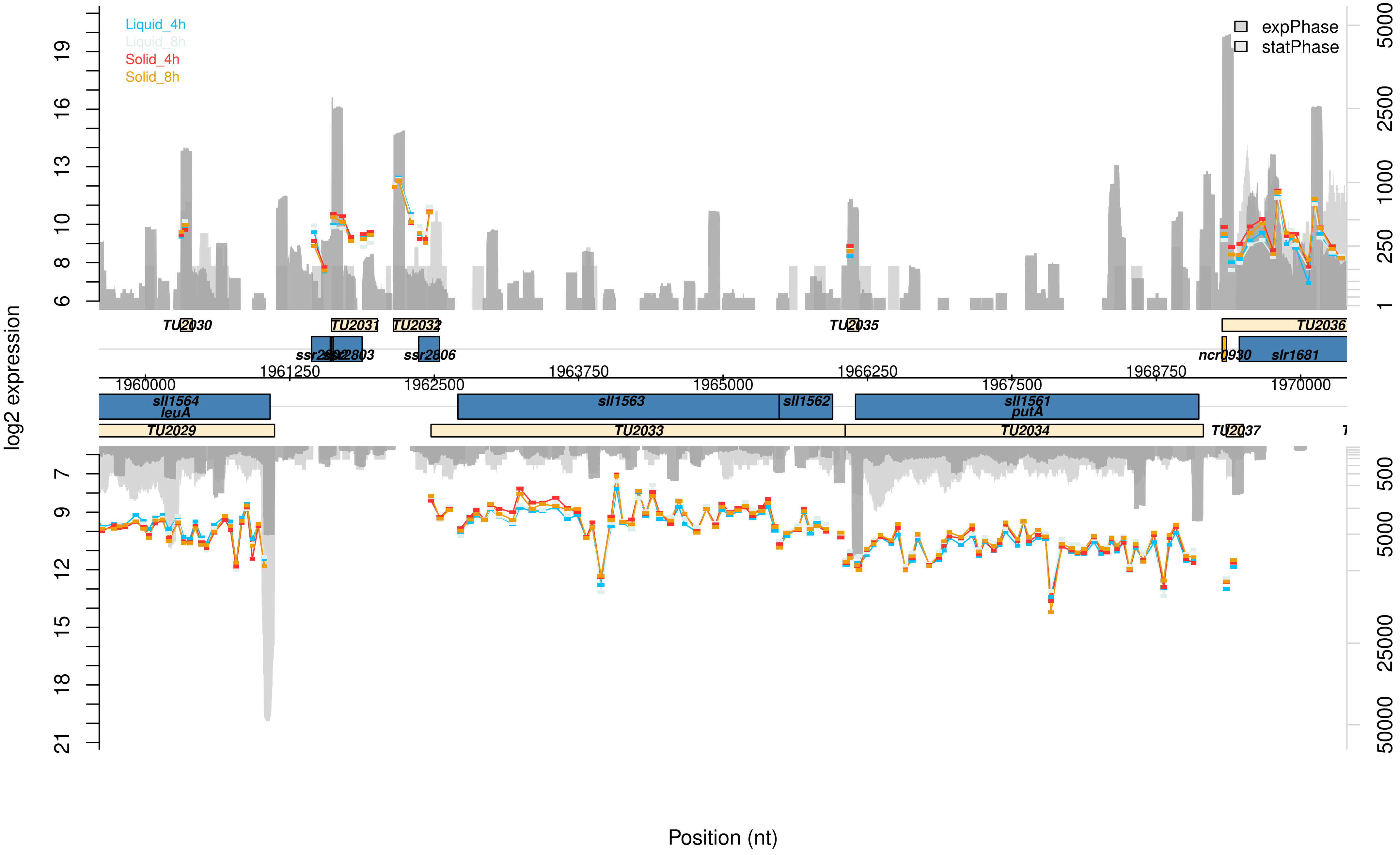

Liquid_4h
Liquid_8h
Solid_4h
Solid_8h
expPhase
statPhase
log2 expression
TU2030
TU2031
TU2032
TU2035
TU2036
ssr2803
ssr2806
ncr0930
slr1681
1960000
1961250
1962500
1963750
1965000
1966250
1967500
1968750
1970000
sll1564
leuA
sll1563
sll1562
sll1561
putA
TU2029
TU2033
TU2034
TU2037
Position (nt)

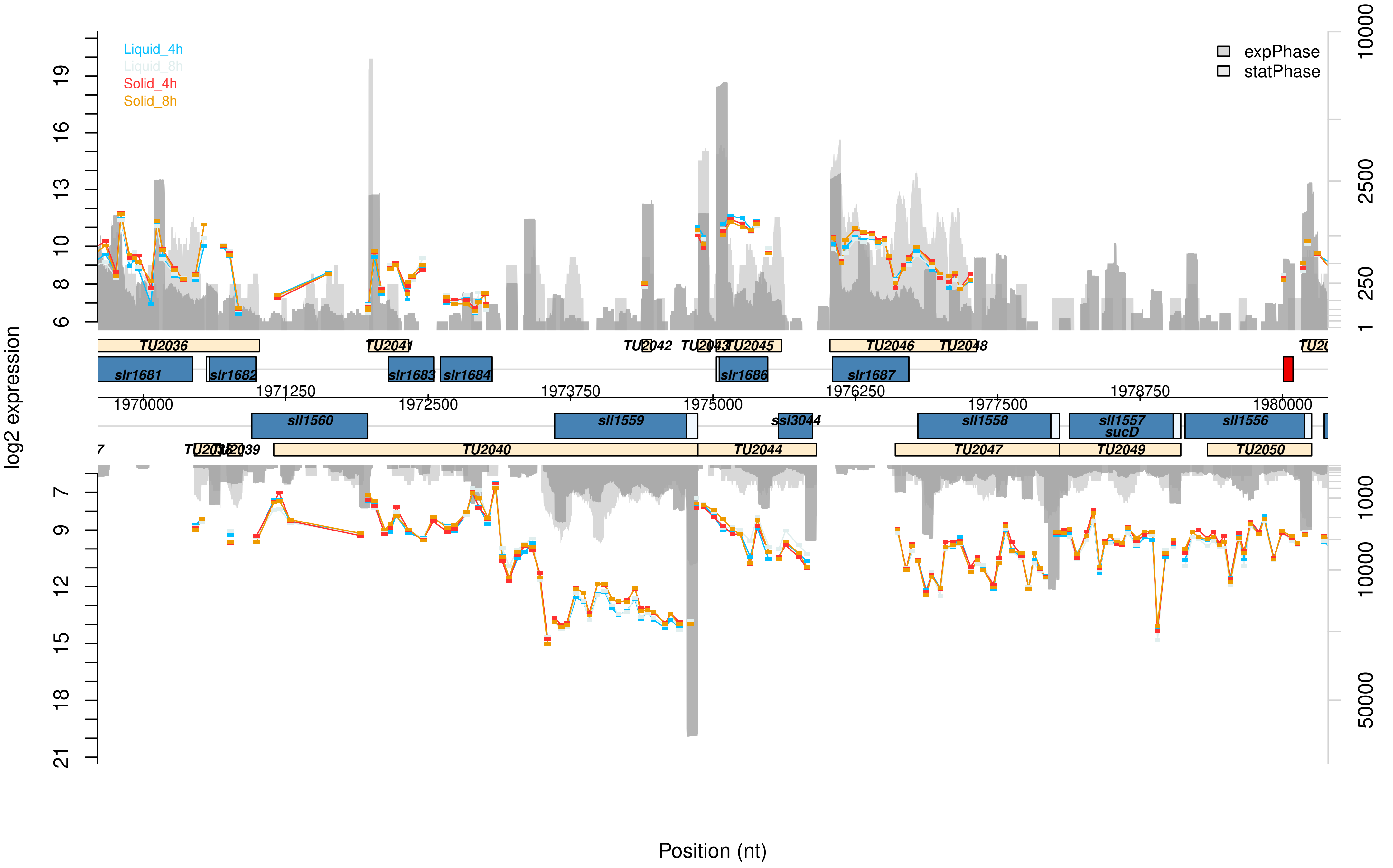
Liquid_4h
Liquid_8h
Solid_4h
Solid_8h
expPhase
statPhase
log2 expression
Position (nt)
TU2036
TU2041
TU2042
TU2043
TU2045
TU2046
TU2048
slr1681
slr1682
slr1683
slr1684
slr1686
slr1687
1970000
1971250
1972500
1973750
1975000
1976250
1977500
1978750
1980000
sll1560
sll1559
ssl3044
sll1558
sll1557
sucD
sll1556
TU2038
TU2039
TU2040
TU2044
TU2047
TU2049
TU2050

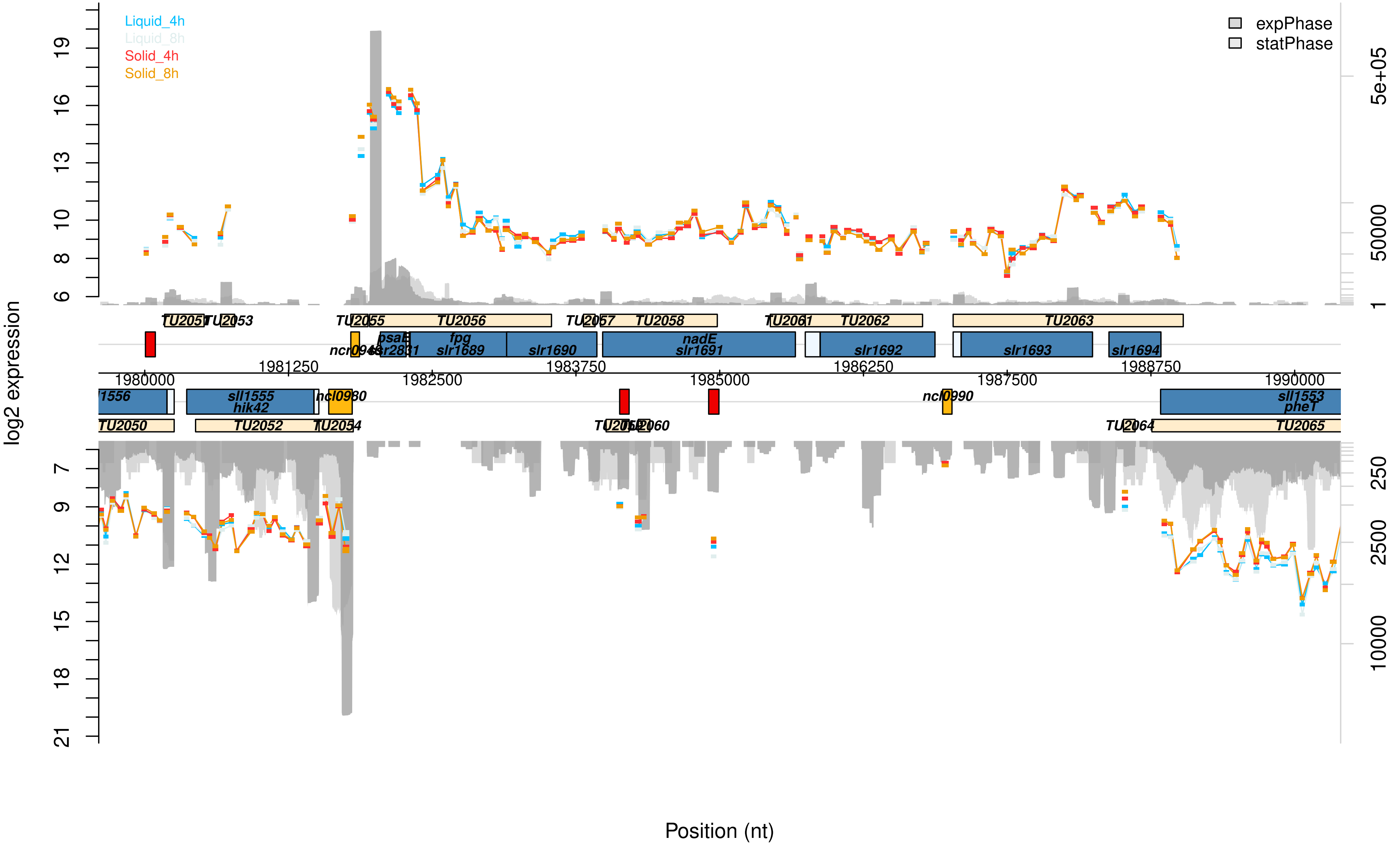

Liquid_4h
Liquid_8h
Solid_4h
Solid_8h
expPhase
statPhase
log2 expression
Position (nt)
TU2051
TU2053
TU2055
TU2056
TU2057
TU2058
TU2061
TU2062
TU2063
psaE
fpg
slr1689
slr1690
nadE
slr1691
slr1692
slr1693
slr1694
1980000
1981250
1982500
1983750
1985000
1986250
1987500
1988750
1990000
sll1555
hik42
ncl0980
ncl0990
sll1553
pheT
TU2050
TU2052
TU2054
TU2060
TU2064
TU2065

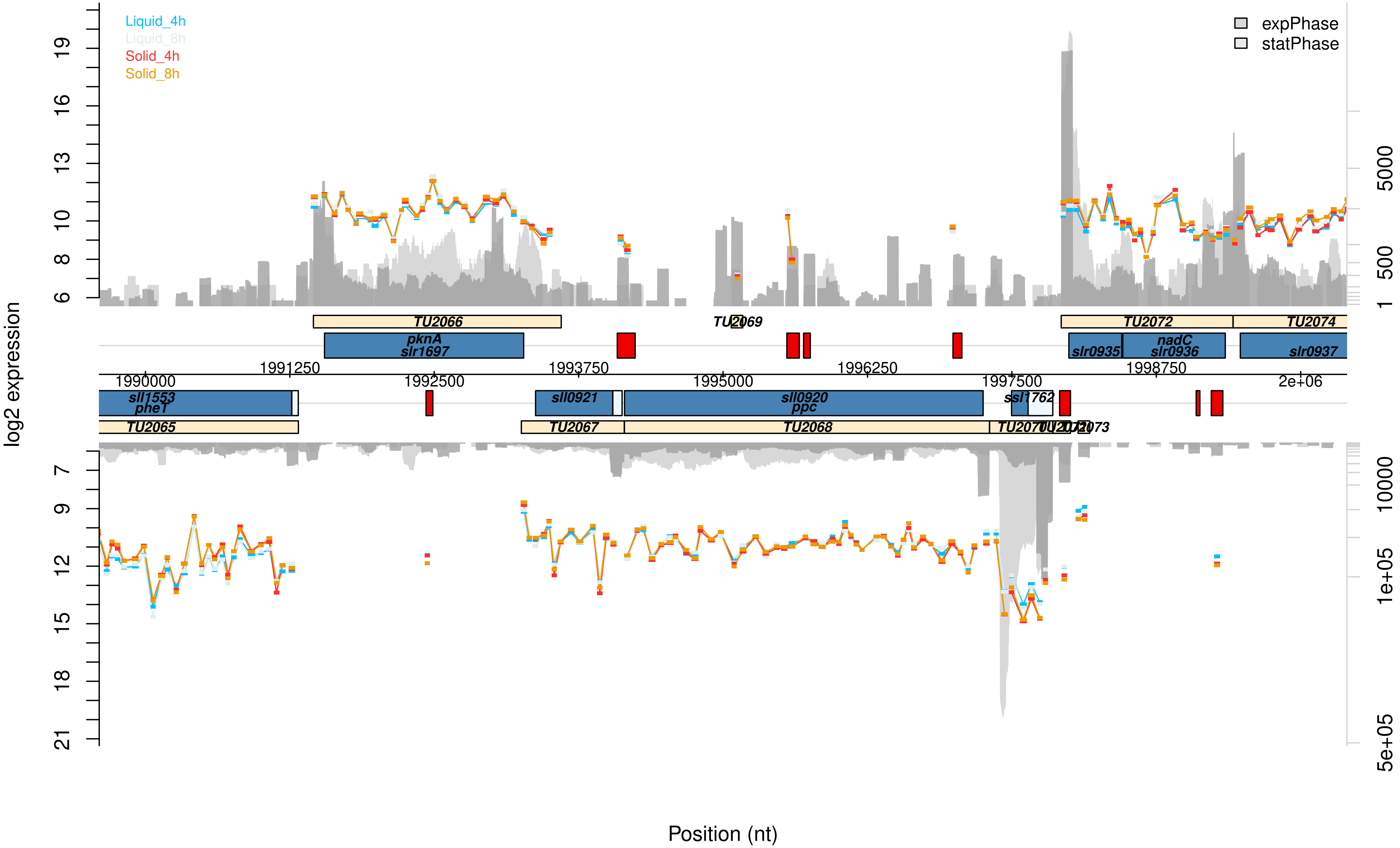

Liquid_4h
Liquid_8h
Solid_4h
Solid_8h
expPhase
statPhase
log2 expression
TU2066
TU2069
TU2072
TU2074
pknA
slr1697
slr0935
nadC
slr0936
slr0937
1990000
1991250
1992500
1993750
1995000
1996250
1997500
1998750
2e+06
sll1553
pheT
sll0921
sll0920
ppc
ssl1762
TU2065
TU2067
TU2068
Position (nt)

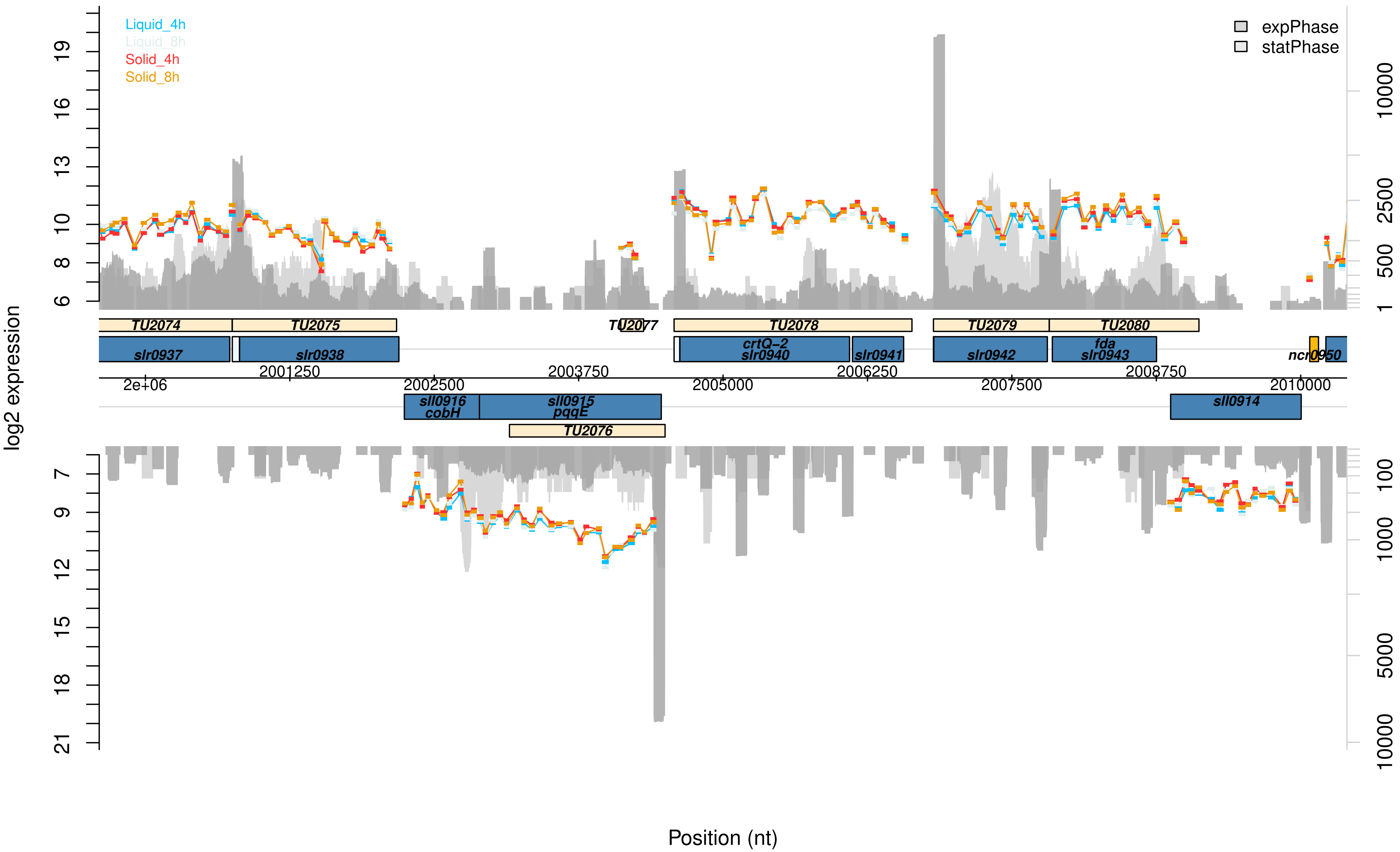

Liquid_4h
Liquid_8h
Solid_4h
Solid_8h
expPhase
statPhase
log2 expression
TU2074
TU2075
TU2077
TU2078
TU2079
TU2080
slr0937
slr0938
crtQ–2
slr0940
slr0941
slr0942
fda
slr0943
ncr0950
2e+06
2001250
2002500
2003750
2005000
2006250
2007500
2008750
2010000
sll0916
cobH
sll0915
pqqE
sll0914
TU2076
Position (nt)

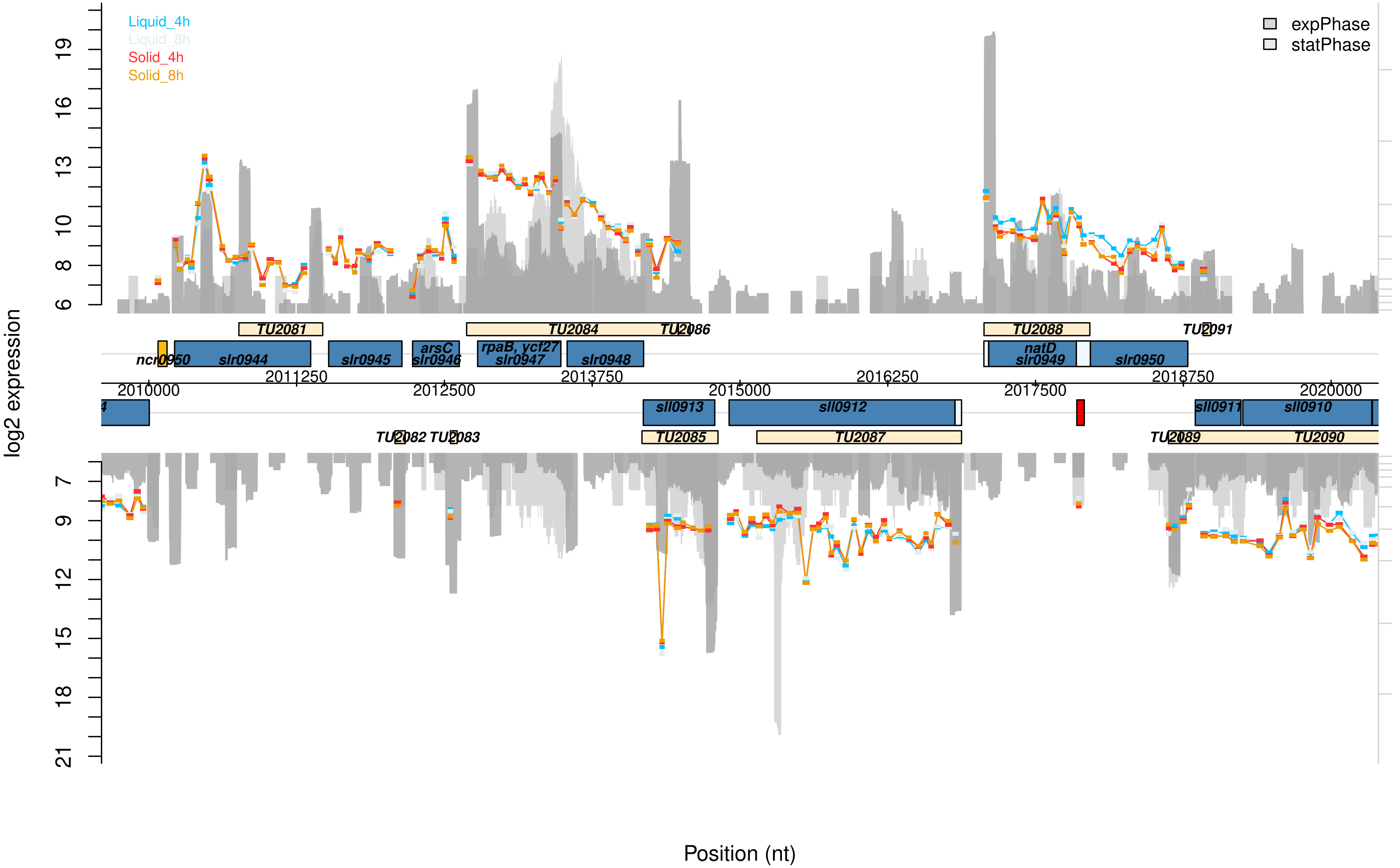

Liquid_4h
Liquid_8h
Solid_4h
Solid_8h
expPhase
statPhase
log2 expression
TU2081
TU2084
TU2086
TU2088
TU2091
ncr0950
slr0944
slr0945
arsC
slr0946
rpaB, ycf27
slr0947
slr0948
natD
slr0949
slr0950
2010000
2011250
2012500
2013750
2015000
2016250
2017500
2018750
2020000
sll0913
sll0912
sll0911
sll0910
TU2082
TU2083
TU2085
TU2087
TU2089
TU2090
Position (nt)

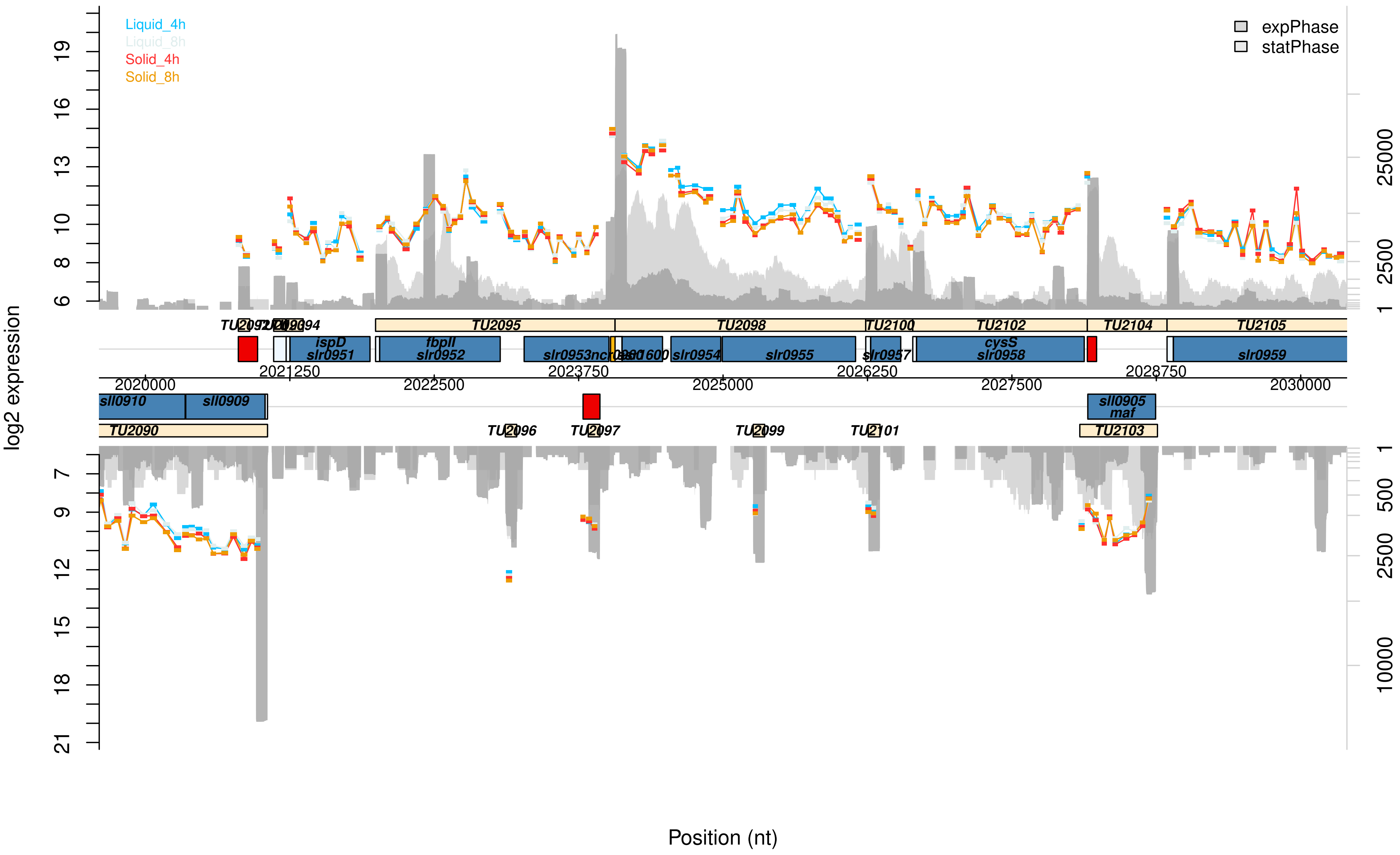

Liquid_4h
Liquid_8h
Solid_4h
Solid_8h
expPhase
statPhase
log2 expression
19
16
13
10
8
6
7
9
12
15
18
21
25000
2500
1
500
10000
TU2095
TU2098
TU2100
TU2102
TU2104
TU2105
ispD
slr0951
fbpII
slr0952
slr0954
slr0955
slr0957
cysS
slr0958
slr0959
2020000
2021250
2022500
2023750
2025000
2026250
2027500
2028750
2030000
sll0910
sll0909
sll0905
maf
TU2090
TU2096
TU2097
TU2099
TU2101
TU2103
Position (nt)

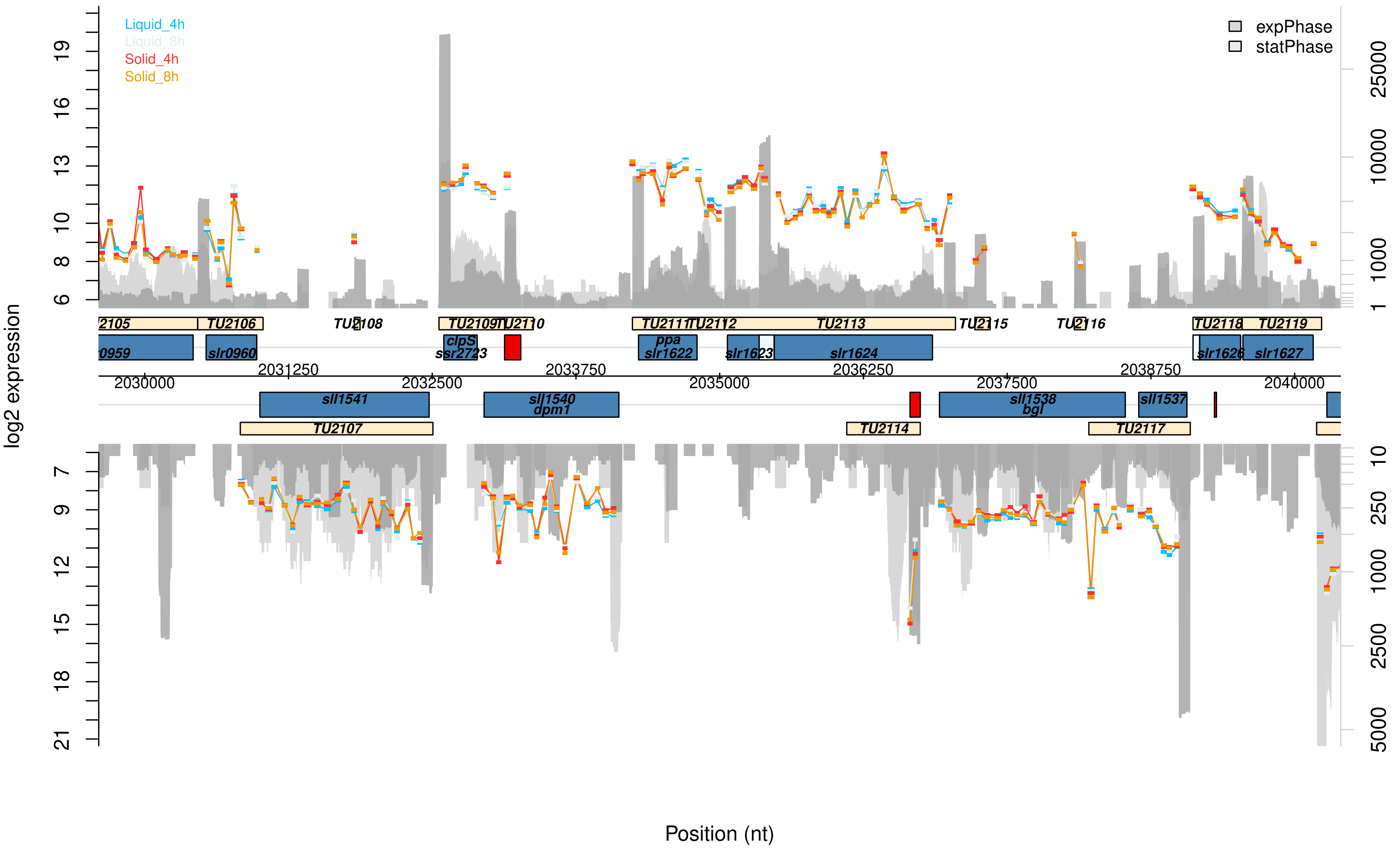

Liquid_4h
Liquid_8h
Solid_4h
Solid_8h
expPhase
statPhase
log2 expression
Position (nt)
TU2105
TU2106
TU2108
TU2109
TU2110
TU2111
TU2112
TU2113
TU2115
TU2116
TU2118
TU2119
slr0959
slr0960
clpS
ssr2723
ppa
slr1622
slr1623
slr1624
slr1626
slr1627
sll1541
sll1540
dpm1
sll1538
bgl
sll1537
TU2107
TU2114
TU2117
2030000
2031250
2032500
2033750
2035000
2036250
2037500
2038750
2040000

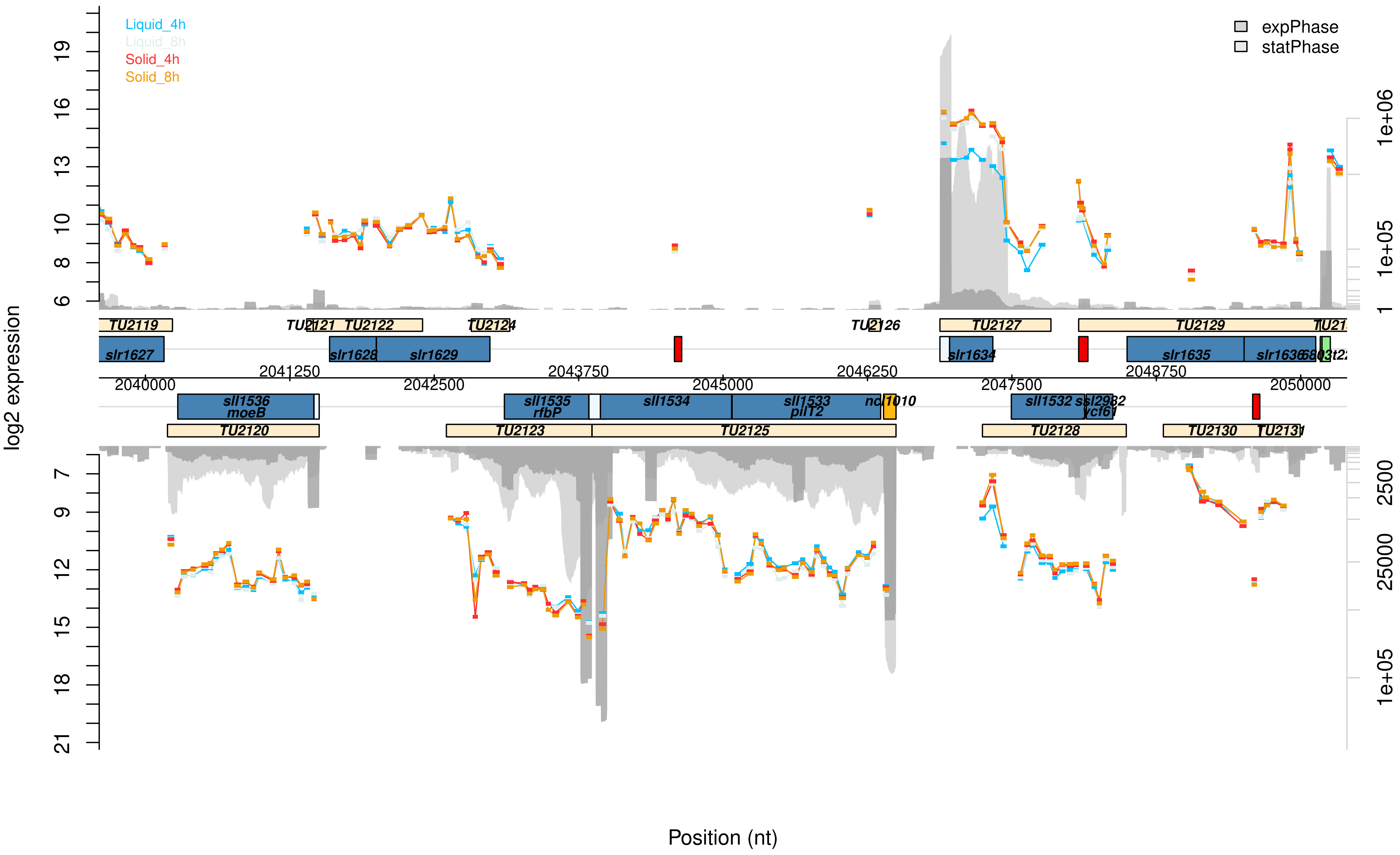

Liquid_4h
Liquid_8h
Solid_4h
Solid_8h
expPhase
statPhase
log2 expression
Position (nt)
TU2119
TU2121
TU2122
TU2124
TU2126
TU2127
TU2129
slr1627
slr1628
slr1629
slr1634
slr1635
slr1636
2040000
2041250
2042500
2043750
2045000
2046250
2047500
2048750
2050000
sll1536
moeB
sll1535
rfbP
sll1534
sll1533
pilT2
ncl1010
sll1532
ssl2982
ycf61
TU2120
TU2123
TU2125
TU2128
TU2130
TU2131

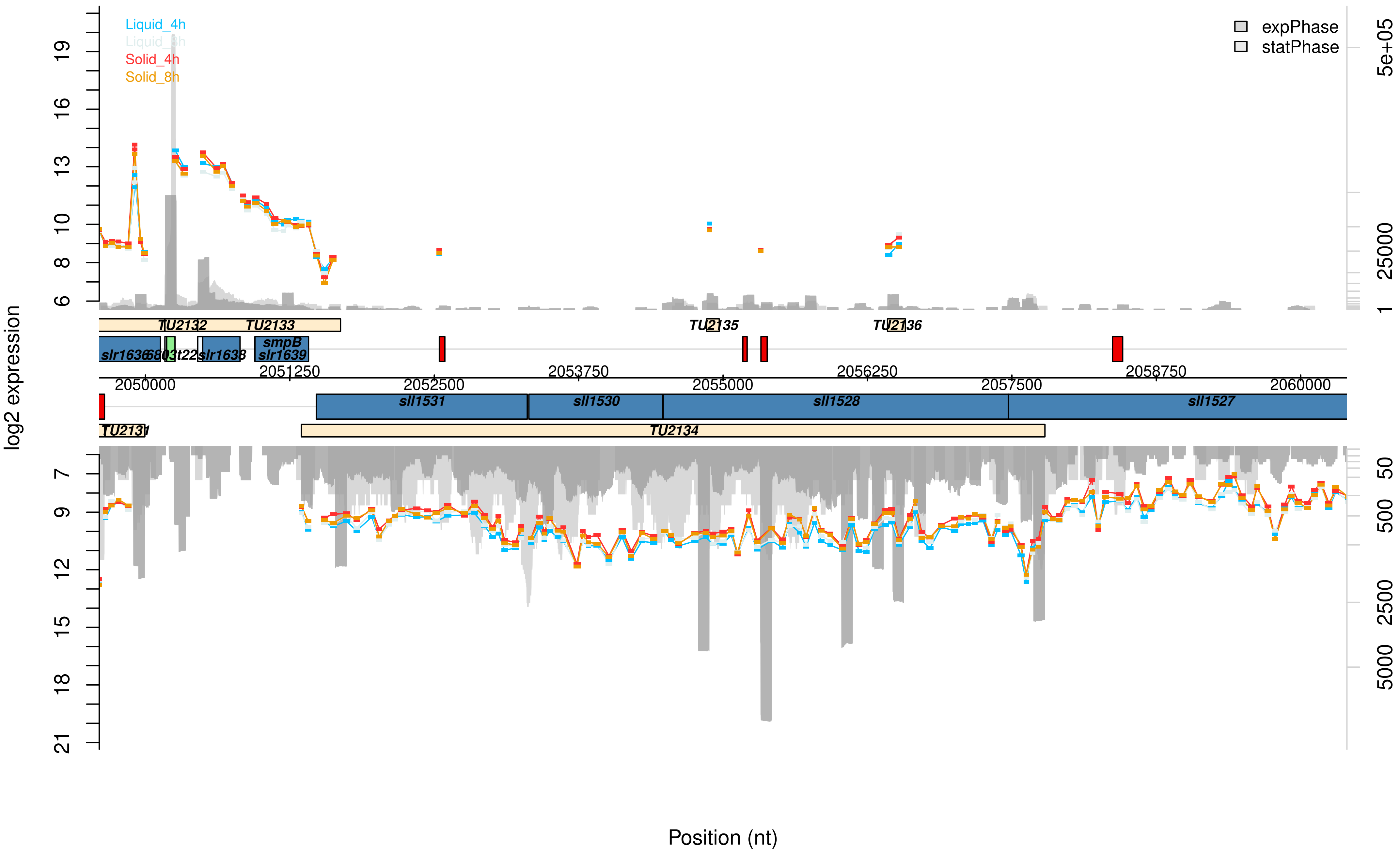

Liquid_4h
Liquid_8h
Solid_4h
Solid_8h
expPhase
statPhase
log2 expression
Position (nt)
TU2131
TU2132
TU2133
TU2134
TU2135
TU2136
slr1636
slr1638
smpB
slr1639
sll1531
sll1530
sll1528
sll1527
2050000
2051250
2052500
2053750
2055000
2056250
2057500
2058750
2060000

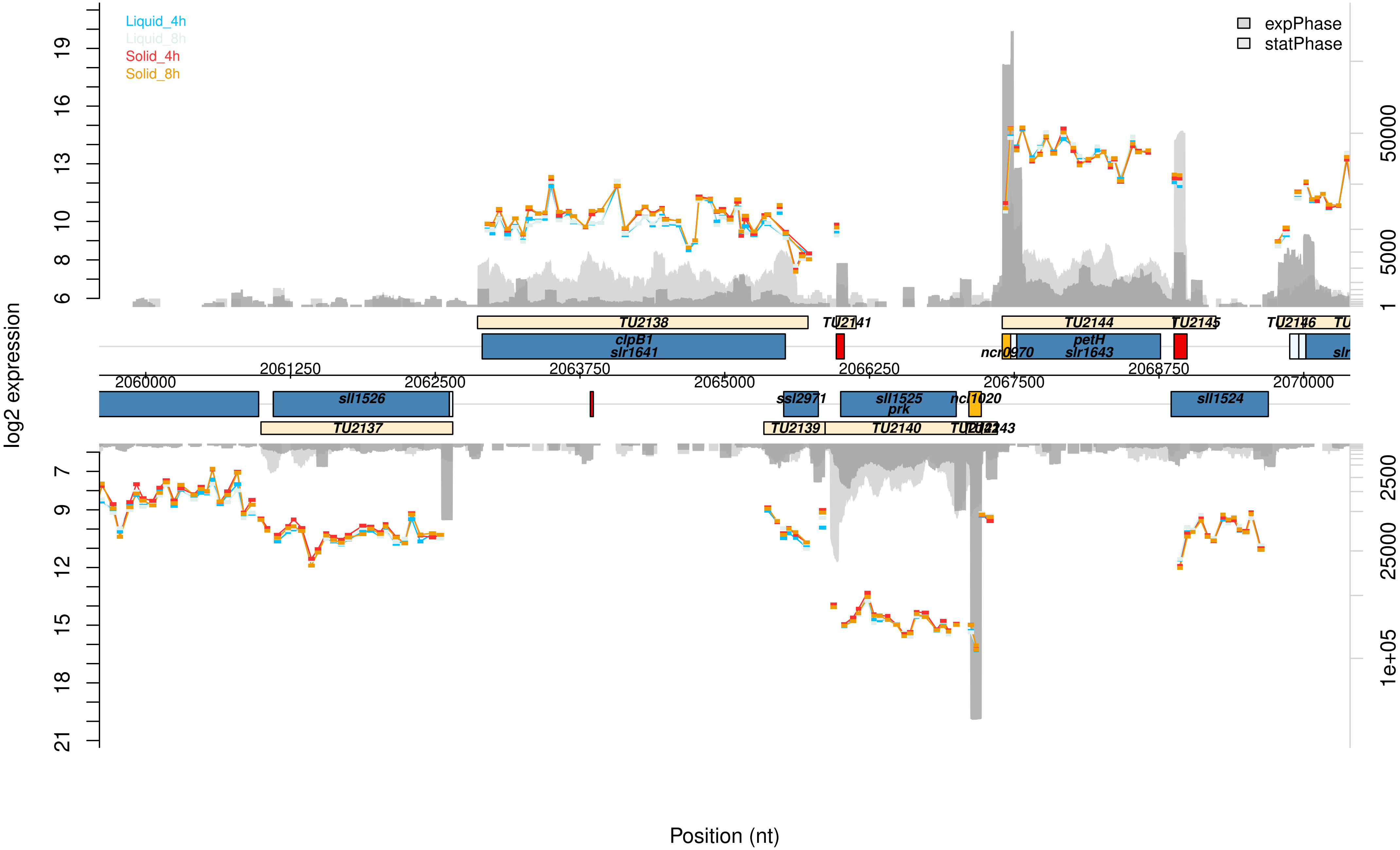

Liquid_4h
Liquid_8h
Solid_4h
Solid_8h
expPhase
statPhase
log2 expression
Position (nt)
TU2138
TU2141
TU2144
TU2145
TU2146
clpB1
slr1641
ncr0970
petH
slr1643
sll1526
sll1524
ssl2971
sll1525
prk
ncr1020
TU2137
TU2139
TU2140
2060000
2061250
2062500
2063750
2065000
2066250
2067500
2068750
2070000

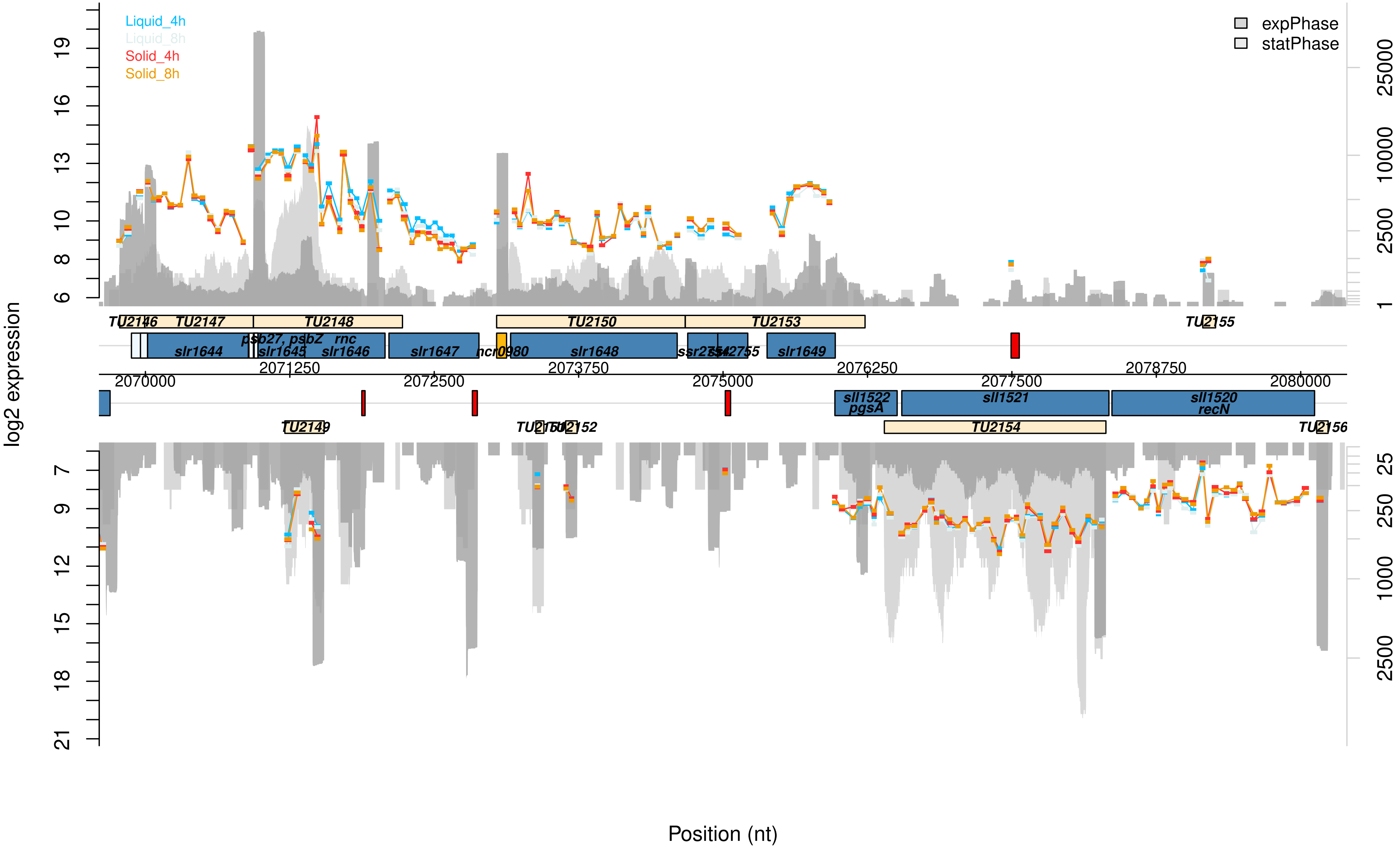

Liquid_4h
Liquid_8h
Solid_4h
Solid_8h
expPhase
statPhase
log2 expression
TU2146
TU2147
TU2148
TU2150
TU2153
TU2155
slr1644
psb27, psbZ
slr1645
rnc
slr1646
slr1647
ncr0980
slr1648
slr1649
2071250
2073750
2076250
2078750
2070000
2072500
2075000
2077500
2080000
sll1522
pgsA
sll1521
sll1520
recN
TU2149
TU2152
TU2154
TU2156
Position (nt)

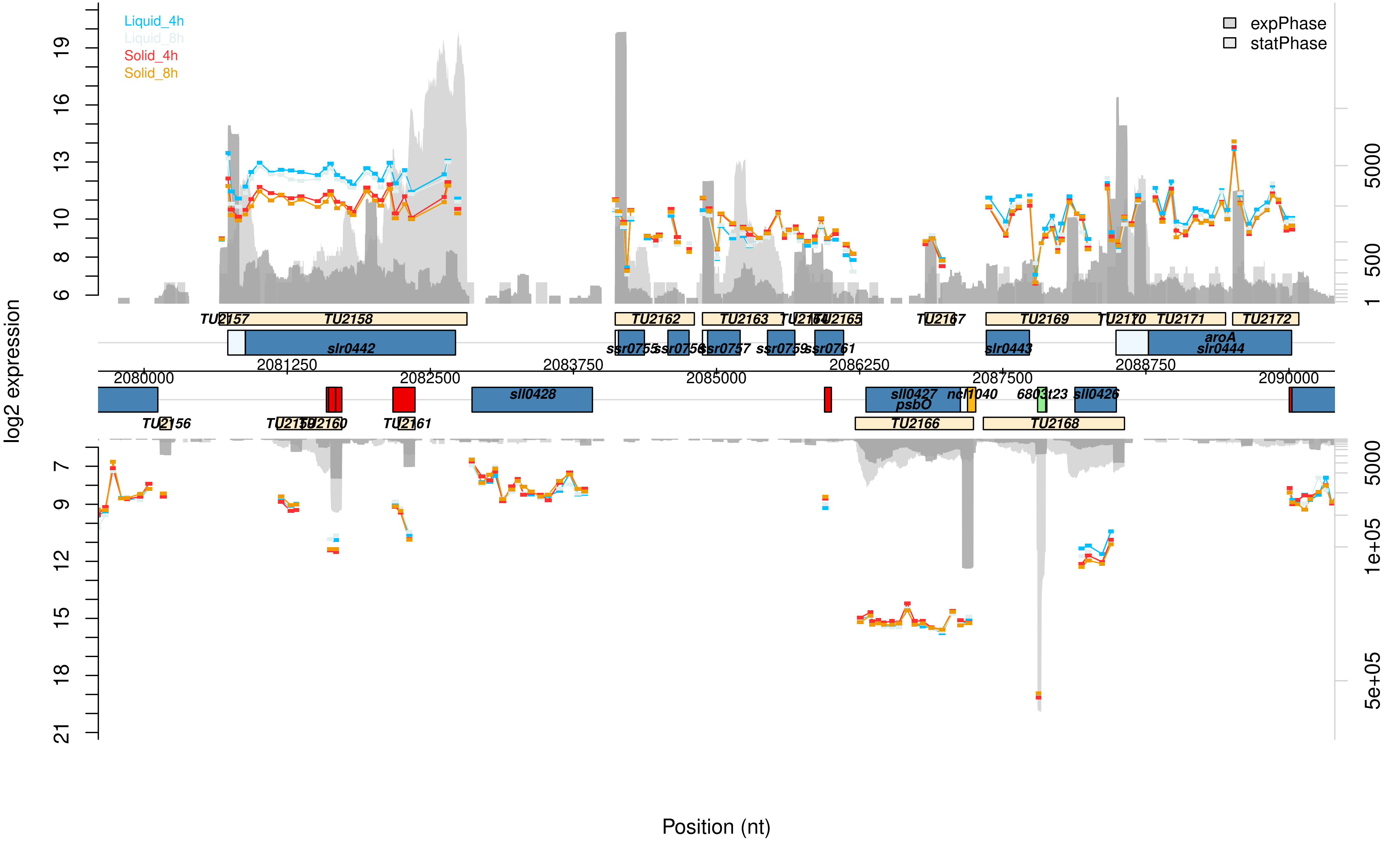

Liquid_4h
Liquid_8h
Solid_4h
Solid_8h
expPhase
statPhase
log2 expression
Position (nt)
TU2157
TU2158
TU2162
TU2163
TU2164
TU2165
TU2167
TU2169
TU2170
TU2171
TU2172
slr0442
ssr0755
ssr0756
ssr0757
ssr0759
ssr0761
slr0443
aroA
slr0444
2080000
2081250
2082500
2083750
2085000
2086250
2087500
2088750
2090000
sll0428
sll0427
psbO
ncl1040
6803t23
sll0426
TU2156
TU2159
TU2160
TU2161
TU2166
TU2168

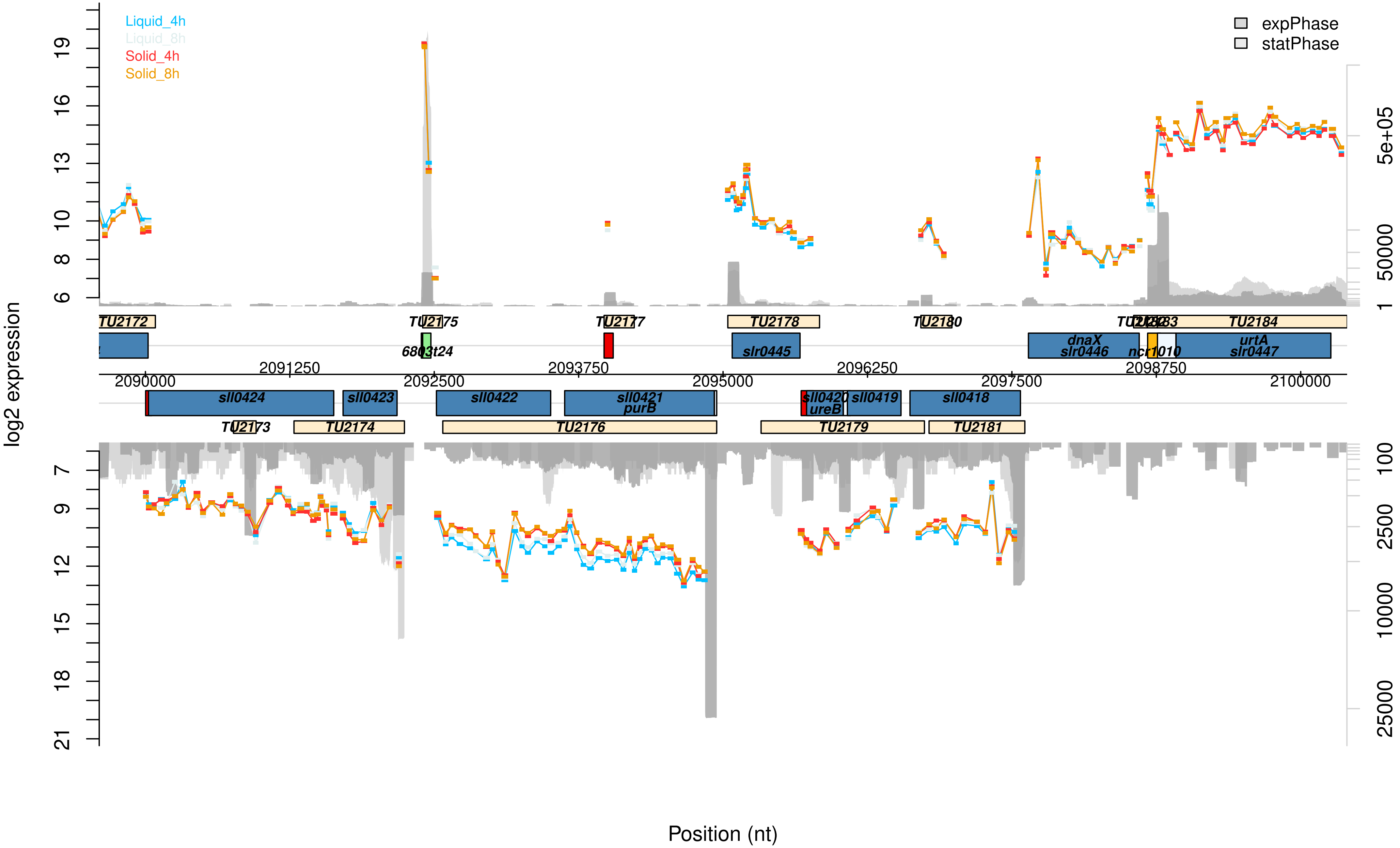

Liquid_4h
Liquid_8h
Solid_4h
Solid_8h
expPhase
statPhase
log2 expression
Position (nt)
TU2172
TU2175
TU2177
TU2178
TU2180
TU2183
TU2184
6803t24
slr0445
dnaX
slr0446
ncr1010
urtA
slr0447
2090000
2091250
2092500
2093750
2095000
2096250
2097500
2098750
2100000
sll0424
sll0423
sll0422
sll0421
purB
sll0420
ureB
sll0419
sll0418
TU2173
TU2174
TU2176
TU2179
TU2181

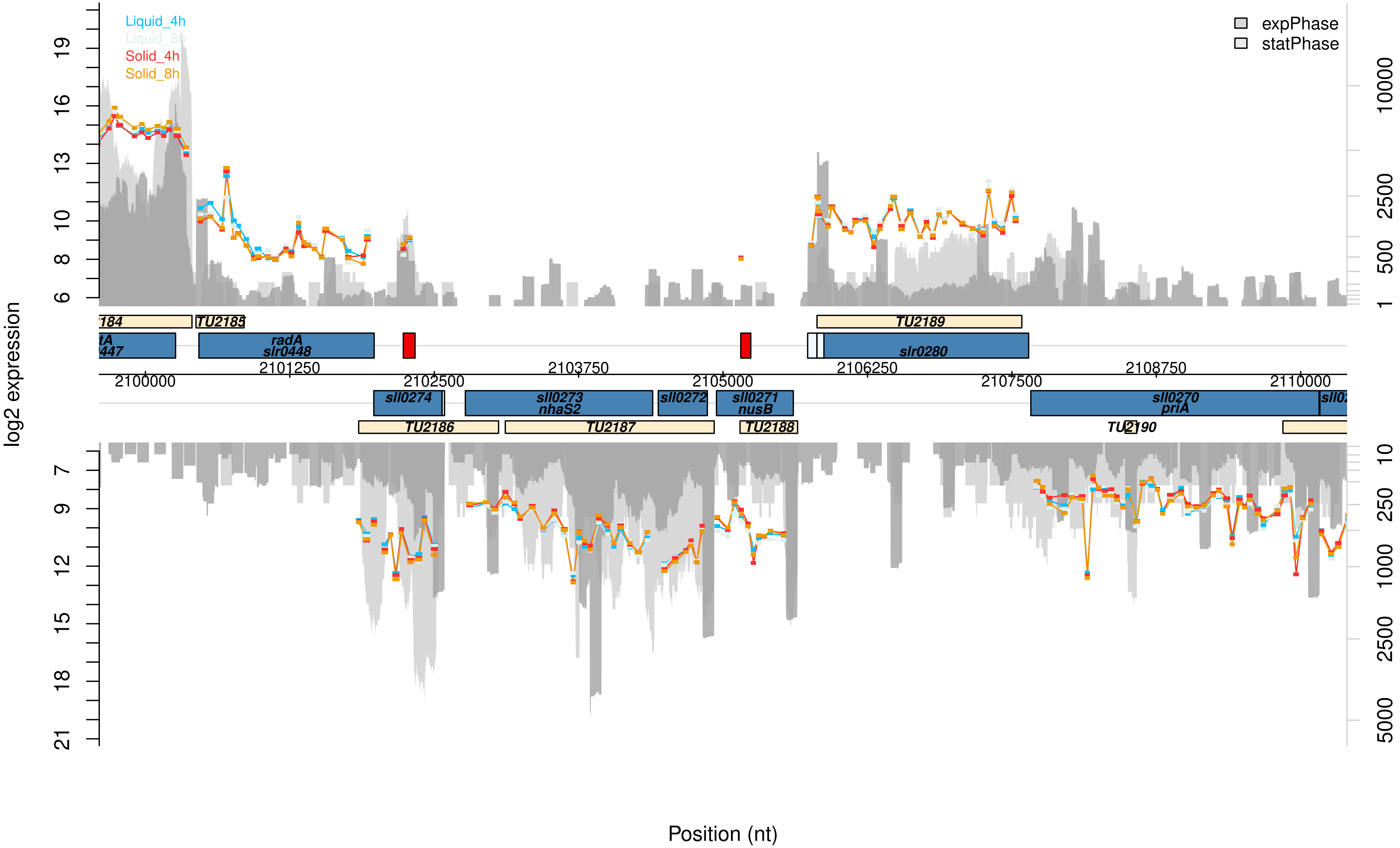

Liquid_4h
Liquid_8h
Solid_4h
Solid_8h
expPhase
statPhase
log2 expression
TU2185
TU2189
radA
slr0448
slr0280
2101250
2103750
2106250
2108750
2100000
2102500
2105000
2107500
2110000
sll0274
sll0273
nhaS2
sll0272
sll0271
nusB
sll0270
priA
TU2186
TU2187
TU2188
TU2190
Position (nt)

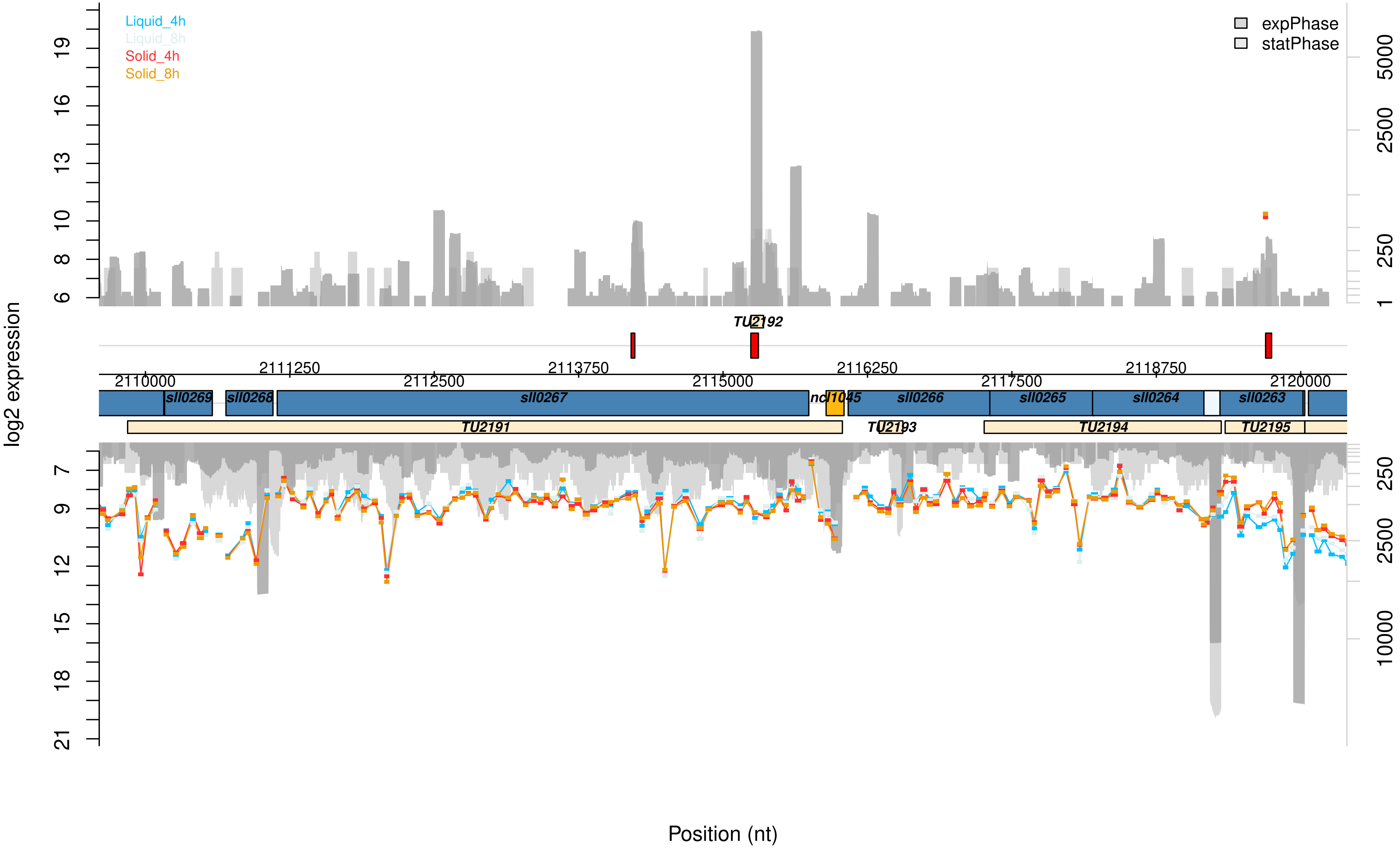

Liquid_4h
Liquid_8h
Solid_4h
Solid_8h
expPhase
statPhase
log2 expression
TU2192
2110000
2111250
2112500
2113750
2115000
2116250
2117500
2118750
2120000
sll0269
sll0268
sll0267
ncl1045
sll0266
sll0265
sll0264
sll0263
TU2191
TU2193
TU2194
TU2195
Position (nt)

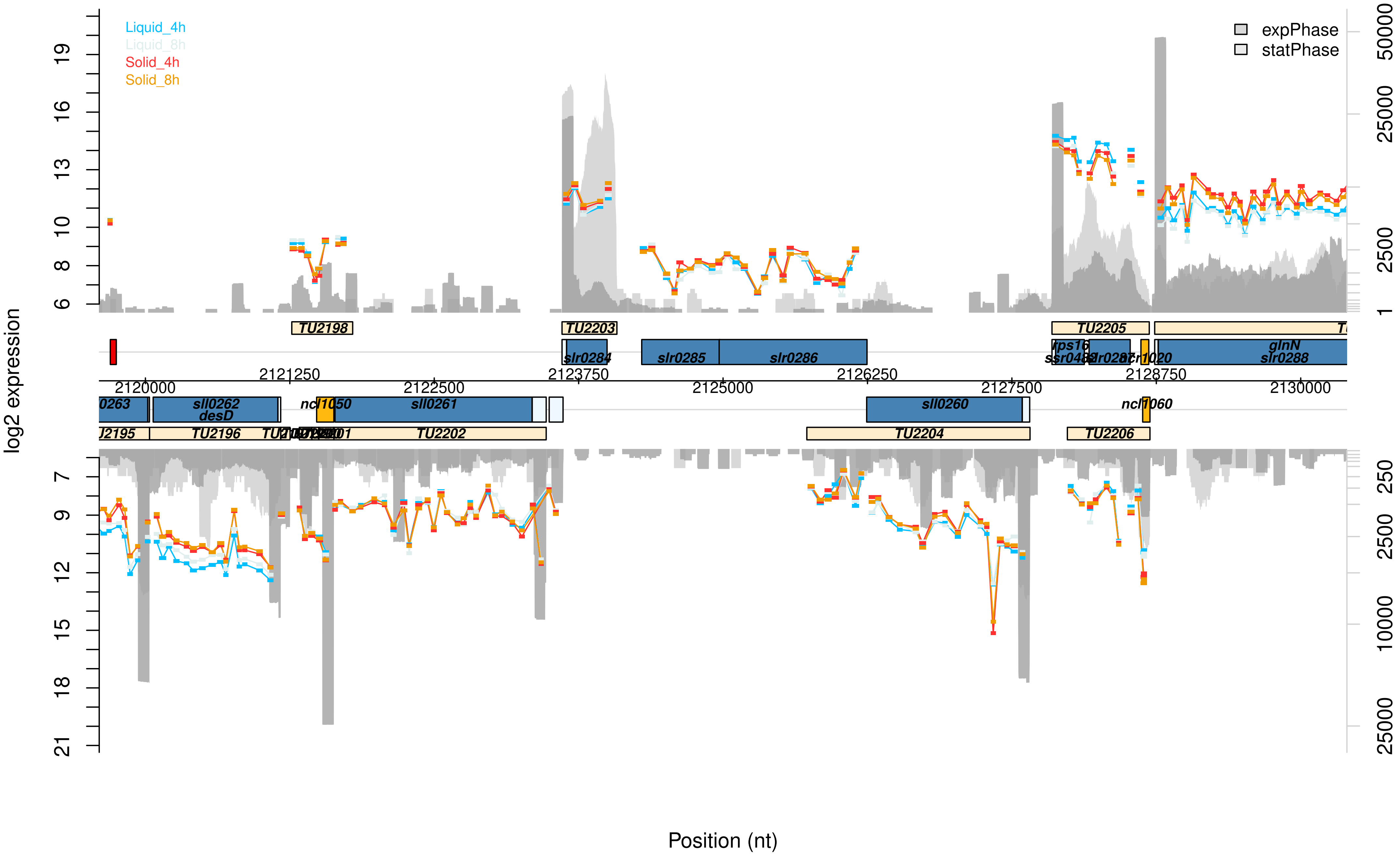

Liquid_4h
Liquid_8h
Solid_4h
Solid_8h
expPhase
statPhase
log2 expression
Position (nt)
TU2198
TU2203
TU2205
TU2204
TU2206
TU2196
TU2202
slr0284
slr0285
slr0286
sll0262
desD
ncl1050
sll0261
sll0260
ncl1060
glnN
slr0288
rps16
2120000
2121250
2122500
2123750
2125000
2126250
2127500
2128750
2130000

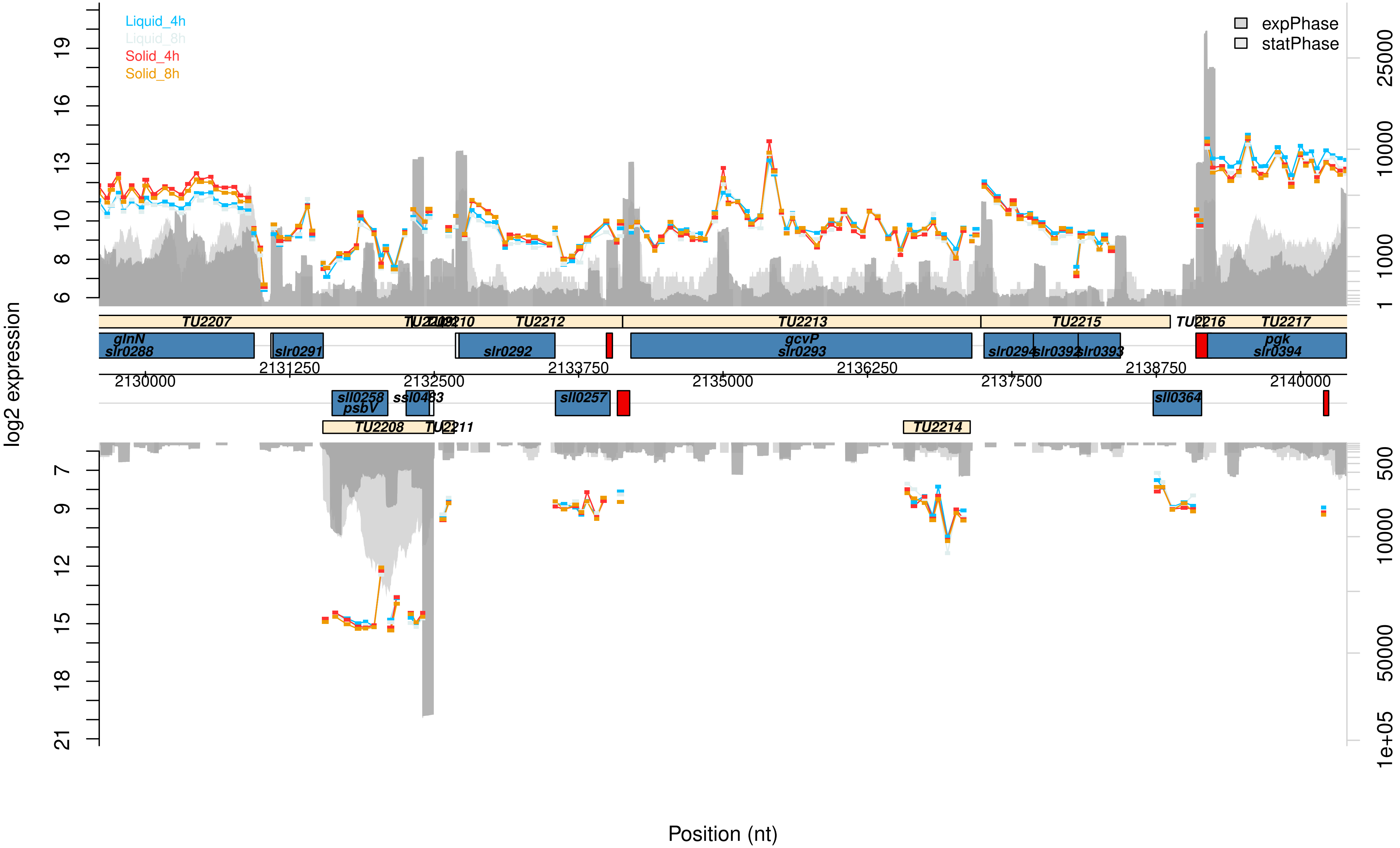

Liquid_4h
Liquid_8h
Solid_4h
Solid_8h
expPhase
statPhase
log2 expression
TU2207
TU2212
TU2213
TU2215
TU2217
TU2208
TU2214
glnN
slr0288
slr0291
slr0292
gcvP
slr0293
slr0294
slr0392
slr0393
pgk
slr0394
sll0258
psbV
sll0257
sll0364
2130000
2131250
2132500
2133750
2135000
2136250
2137500
2138750
2140000
Position (nt)

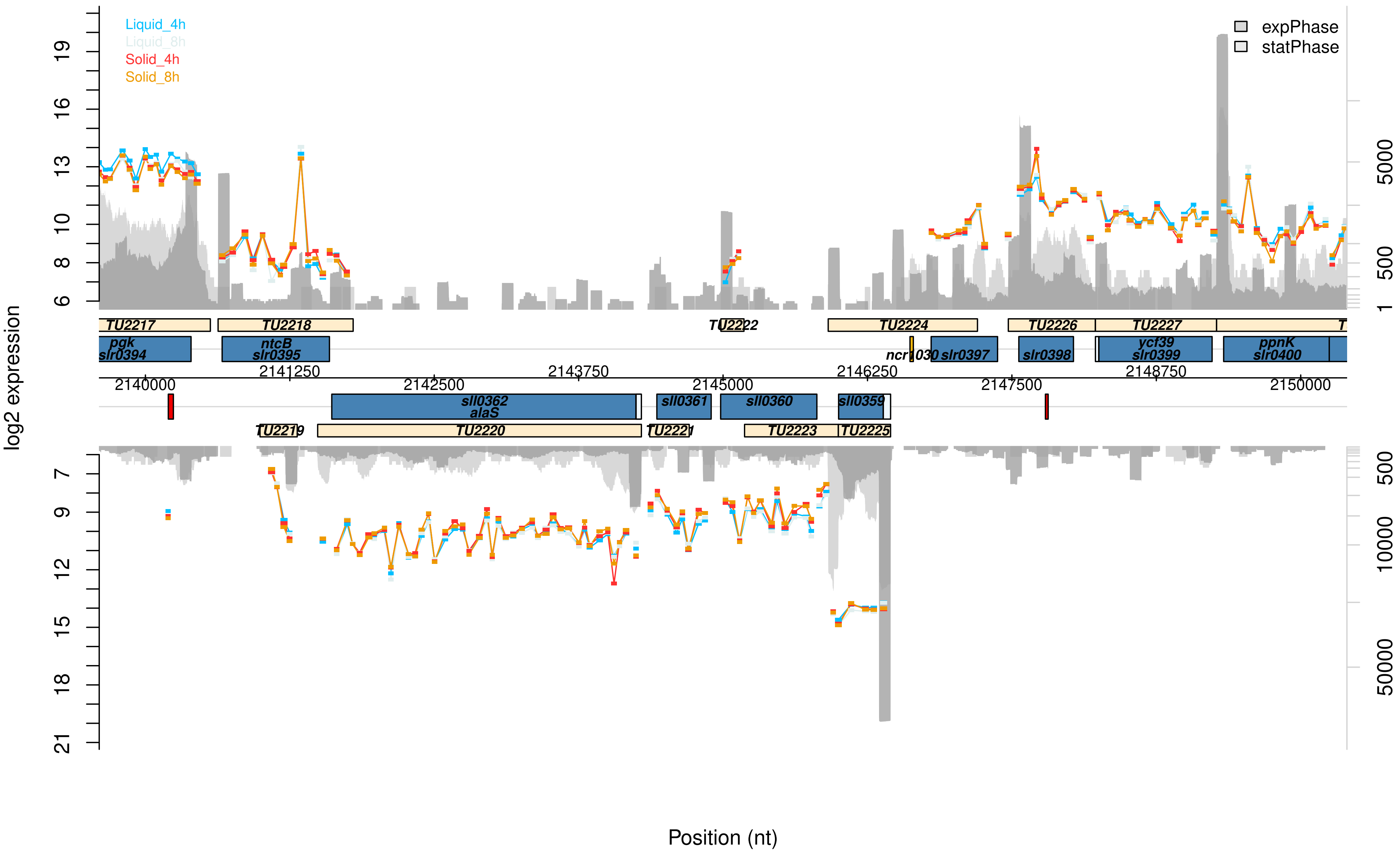

Liquid_4h
Liquid_8h
Solid_4h
Solid_8h
expPhase
statPhase
log2 expression
6
8
10
13
16
19
7
9
12
15
18
21
1
500
5000
10000
50000
TU2217
TU2218
TU2222
TU2224
TU2226
TU2227
T
pgk
slr0394
ntcB
slr0395
ncr1030
slr0397
slr0398
ycf39
slr0399
ppnK
slr0400
2140000
2141250
2142500
2143750
2145000
2146250
2147500
2148750
2150000
sll0362
alaS
sll0361
sll0360
sll0359
TU2219
TU2220
TU2221
TU2223
TU2225
Position (nt)

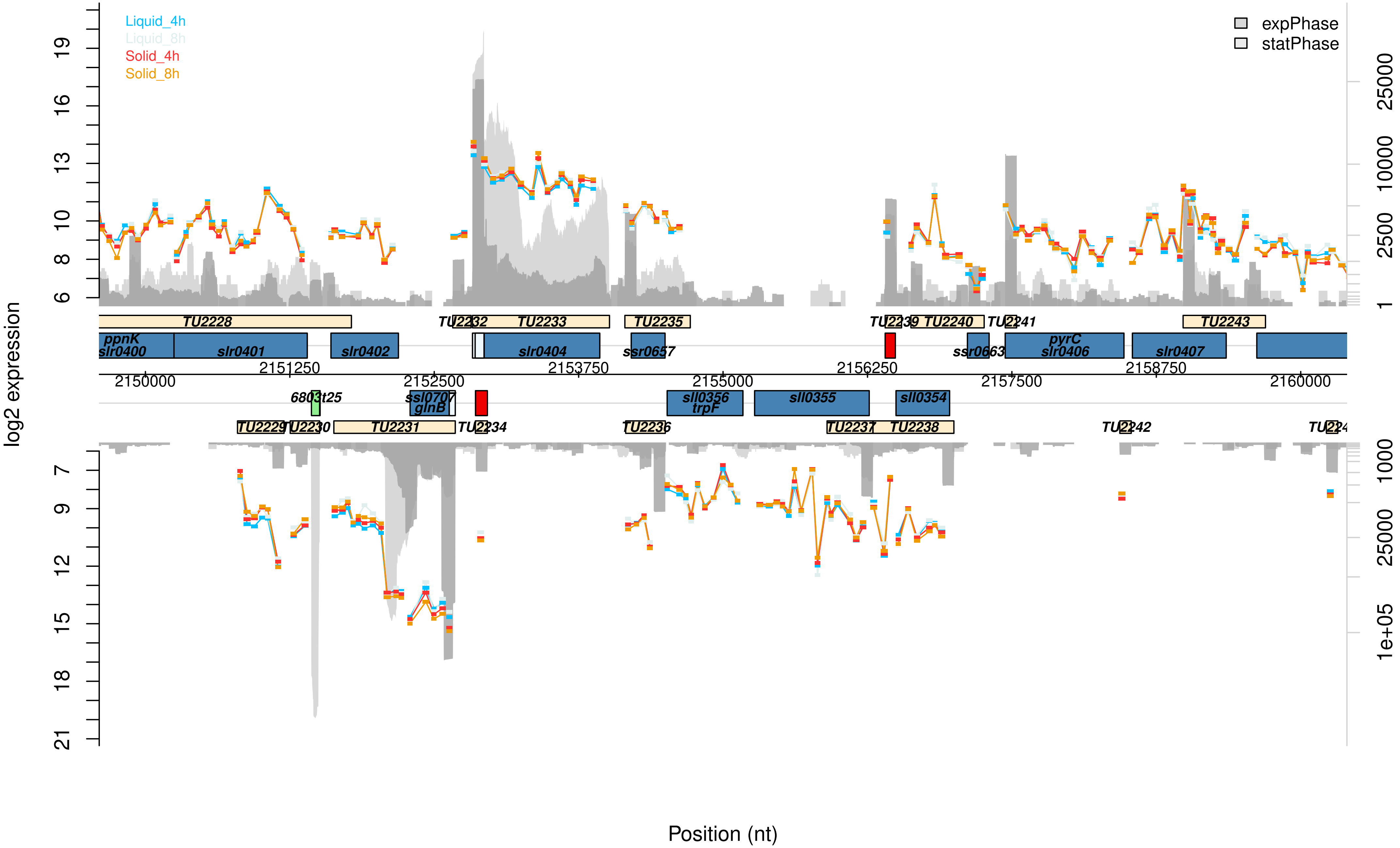

Liquid_4h
Liquid_8h
Solid_4h
Solid_8h
expPhase
statPhase
log2 expression
Position (nt)
TU2228
TU2232
TU2233
TU2235
TU2239
TU2240
TU2241
TU2243
ppnK
slr0400
slr0401
slr0402
slr0404
ssr0657
ssr0663
pyrC
slr0406
slr0407
2150000
2151250
2152500
2153750
2155000
2156250
2157500
2158750
2160000
6803t25
ssl0707
glnB
sll0356
trpF
sll0355
sll0354
TU2229
TU2230
TU2231
TU2234
TU2236
TU2237
TU2238
TU2242

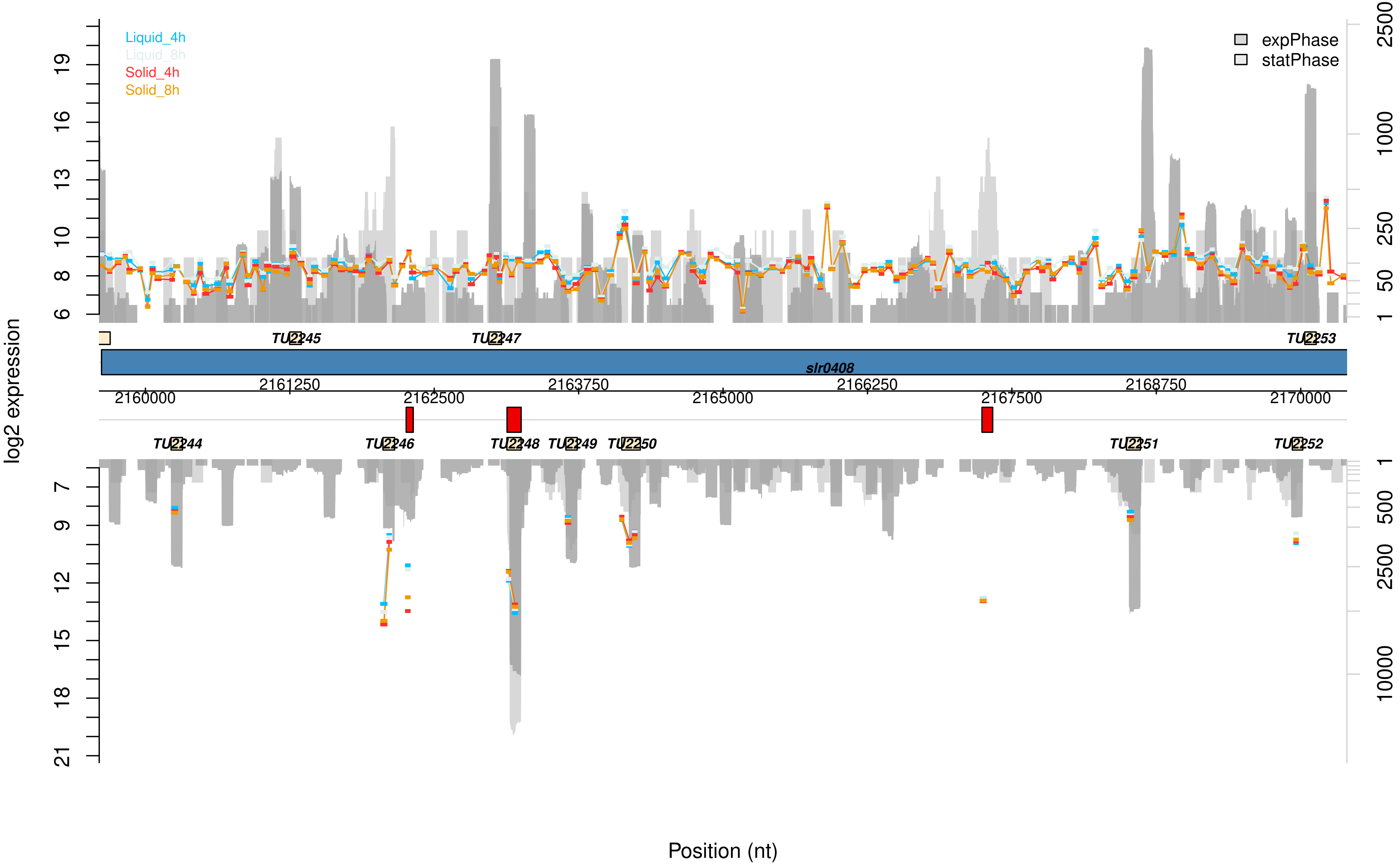

Liquid_4h
Liquid_8h
Solid_4h
Solid_8h
expPhase
statPhase
log2 expression
TU2245
TU2247
TU2253
slr0408
TU2244
TU2246
TU2248
TU2249
TU2250
TU2251
TU2252
2160000
2161250
2162500
2163750
2165000
2166250
2167500
2168750
2170000
Position (nt)

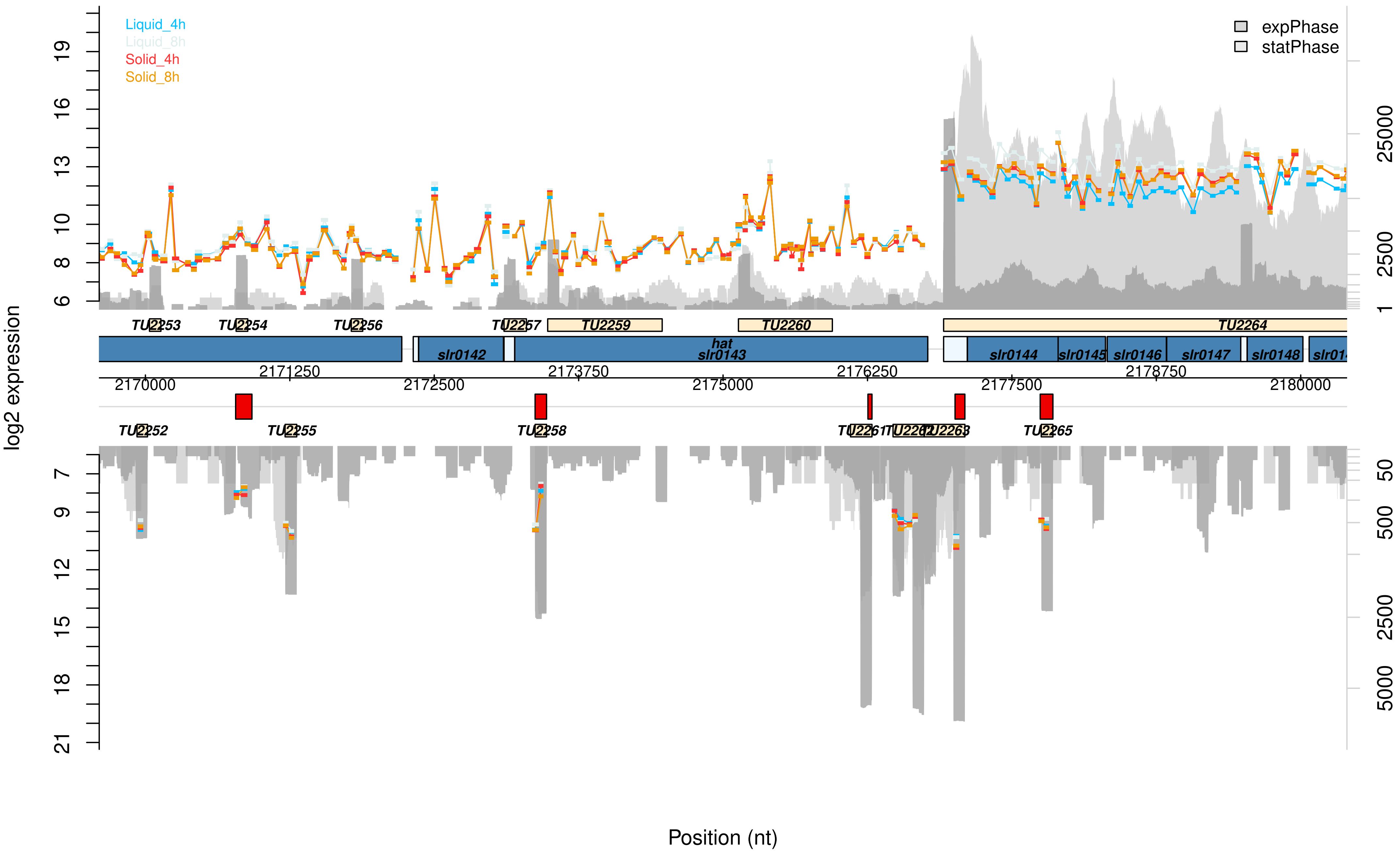

Liquid_4h
Liquid_8h
Solid_4h
Solid_8h
expPhase
statPhase
log2 expression
TU2253
TU2254
TU2256
TU2257
TU2259
TU2260
TU2264
slr0142
hat
slr0143
slr0144
slr0145
slr0146
slr0147
slr0148
2170000
2171250
2172500
2173750
2175000
2176250
2177500
2178750
2180000
TU2252
TU2255
TU2258
TU2261
TU2262
TU2263
TU2265
Position (nt)

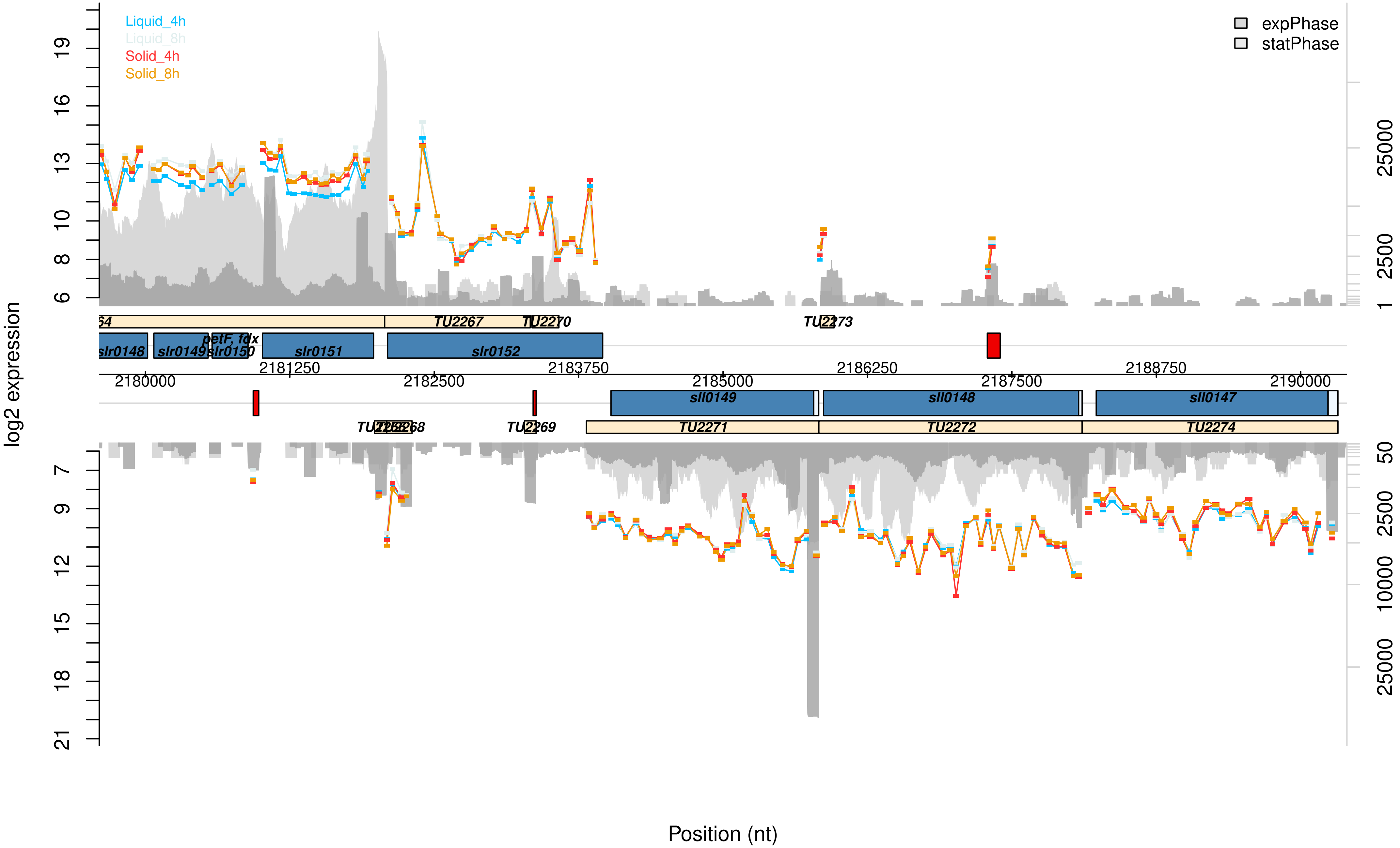
Liquid_4h
Liquid_8h
Solid_4h
Solid_8h
expPhase
statPhase
log2 expression
TU2267
TU2270
TU2273
slr0148
slr0149
petF, fdx
slr0150
slr0151
slr0152
2181250
2183750
2186250
2188750
2180000
2182500
2185000
2187500
2190000
sll0149
sll0148
sll0147
TU2268
TU2269
TU2271
TU2272
TU2274
Position (nt)

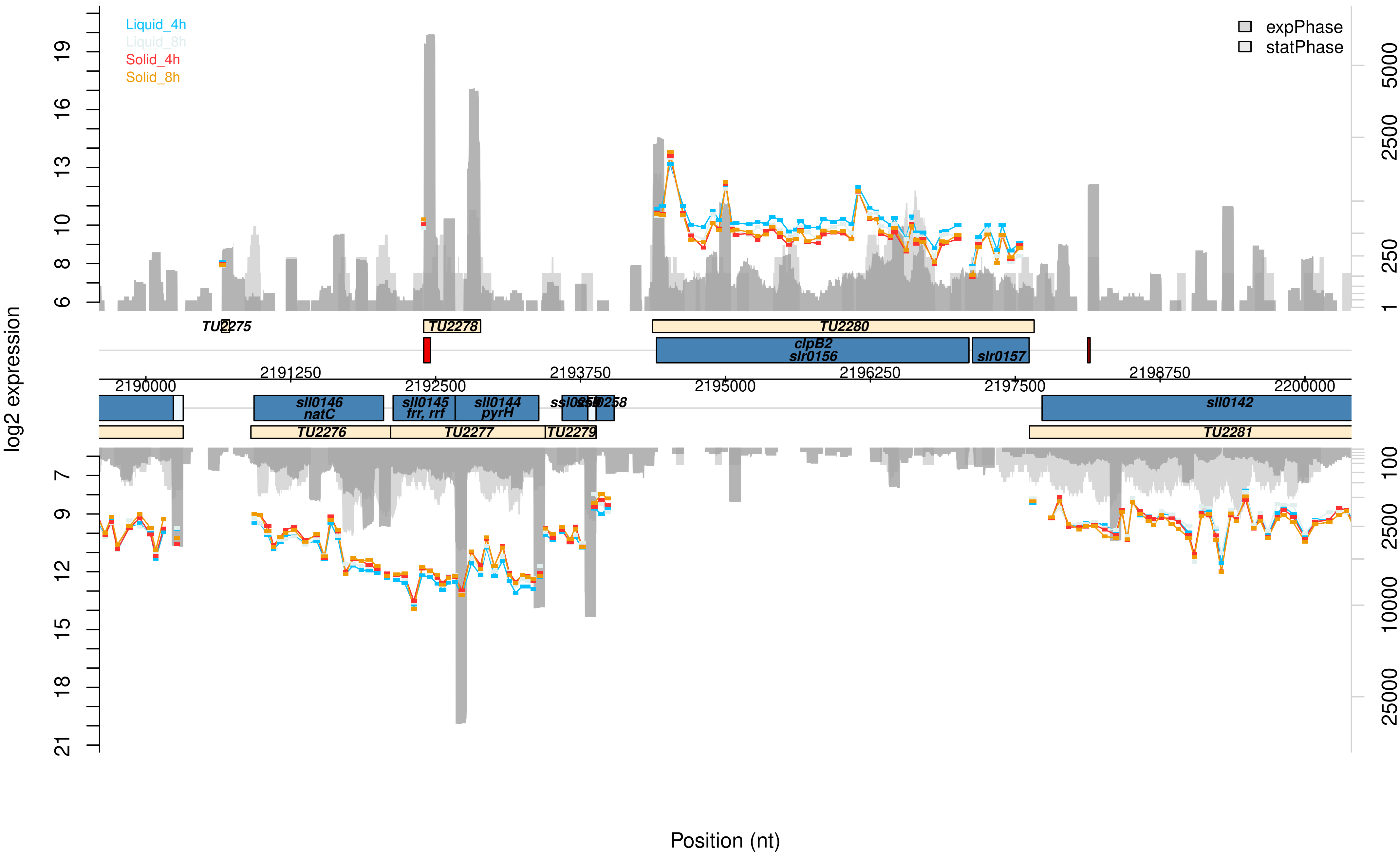

Liquid_4h
Liquid_8h
Solid_4h
Solid_8h
expPhase
statPhase
log2 expression
Position (nt)
TU2275
TU2278
TU2280
clpB2
slr0156
slr0157
sll0146
natC
sll0145
frr, rrf
sll0144
pyrH
sll0142
TU2276
TU2277
TU2279
TU2281
2190000
2191250
2192500
2193750
2195000
2196250
2197500
2198750
2200000

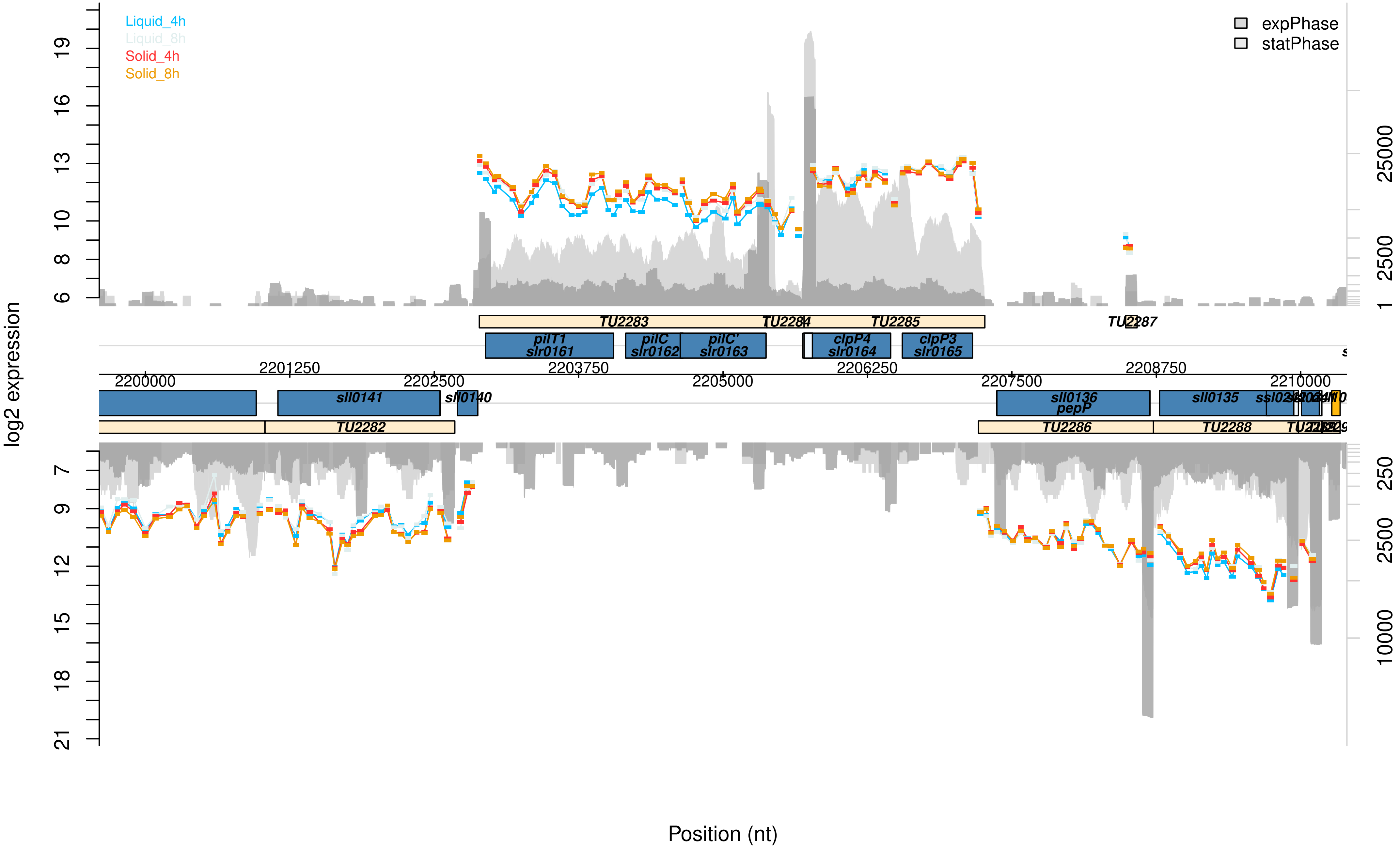

Liquid_4h
Liquid_8h
Solid_4h
Solid_8h
expPhase
statPhase
log2 expression
Position (nt)
TU2283
TU2284
TU2285
TU2287
pilT1
slr0161
pilC
slr0162
pilC'
slr0163
clpP4
slr0164
clpP3
slr0165
sll0141
sll0140
sll0136
pepP
sll0135
TU2282
TU2286
TU2288
2200000
2201250
2202500
2203750
2205000
2206250
2207500
2208750
2210000

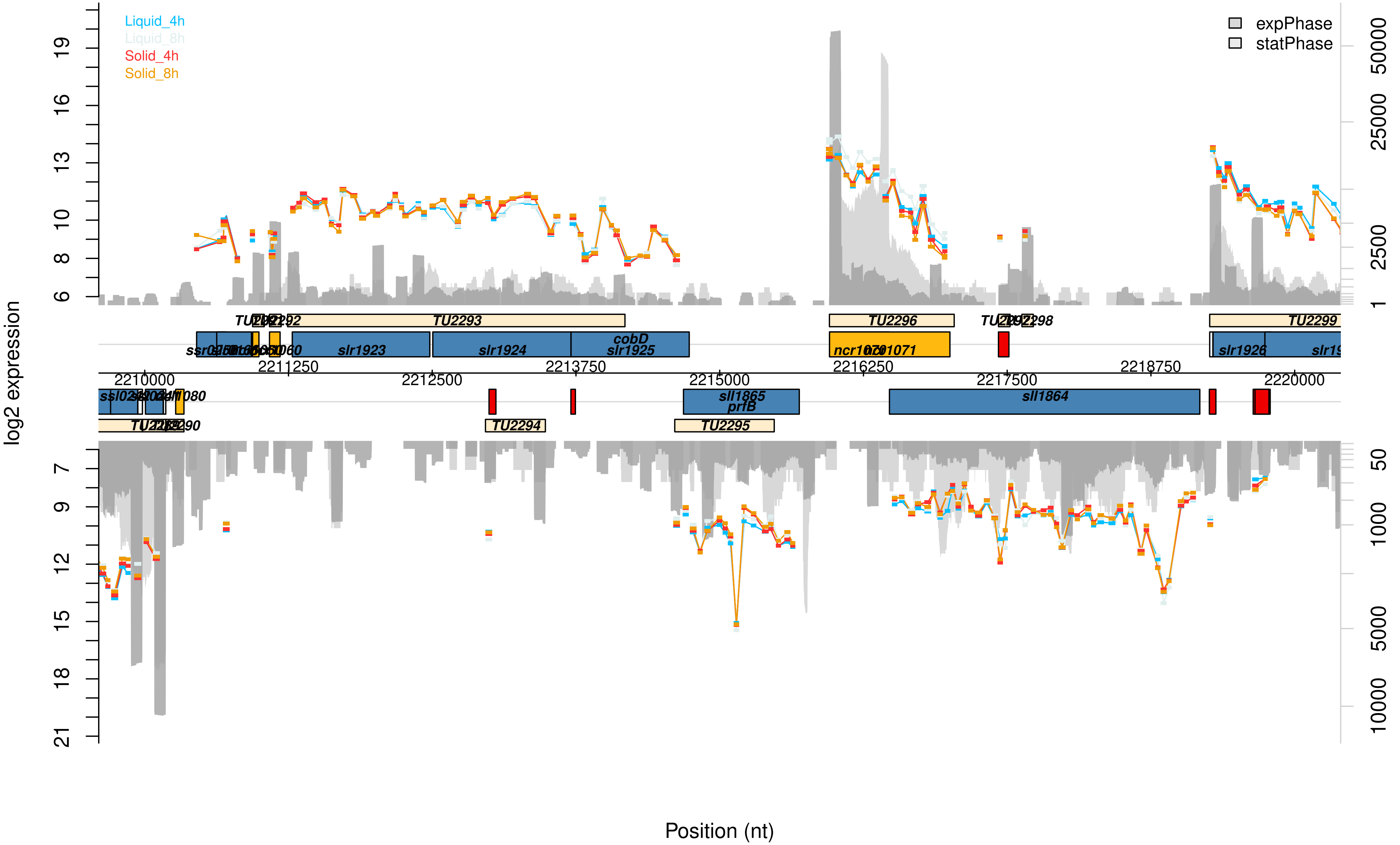

Liquid_4h
Liquid_8h
Solid_4h
Solid_8h
expPhase
statPhase
log2 expression
Position (nt)
TU2293
TU2296
TU2299
slr1923
slr1924
cobD
slr1925
slr1926
sll1865
prfB
sll1864
TU2294
TU2295
2210000
2211250
2212500
2213750
2215000
2216250
2217500
2218750
2220000

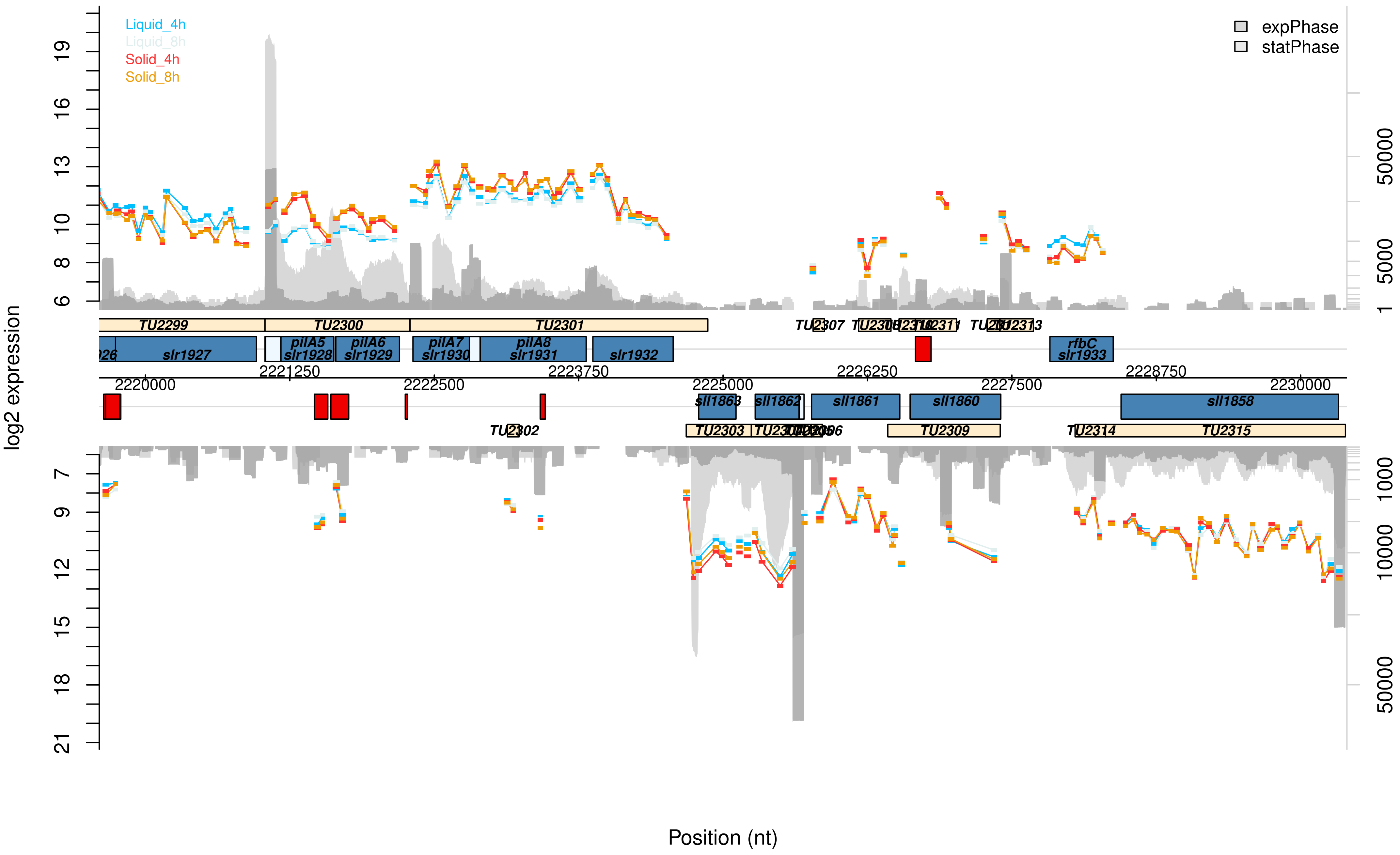

Liquid_4h
Liquid_8h
Solid_4h
Solid_8h
expPhase
statPhase
log2 expression
Position (nt)
TU2299
TU2300
TU2301
TU2302
TU2303
TU2307
TU2309
TU2311
TU2313
TU2314
TU2315
slr1927
pilA5
slr1928
pilA6
slr1929
pilA7
slr1930
pilA8
slr1931
slr1932
rfbC
slr1933
sll1863
sll1862
sll1861
sll1860
sll1858
2220000
2221250
2222500
2223750
2225000
2226250
2227500
2228750
2230000

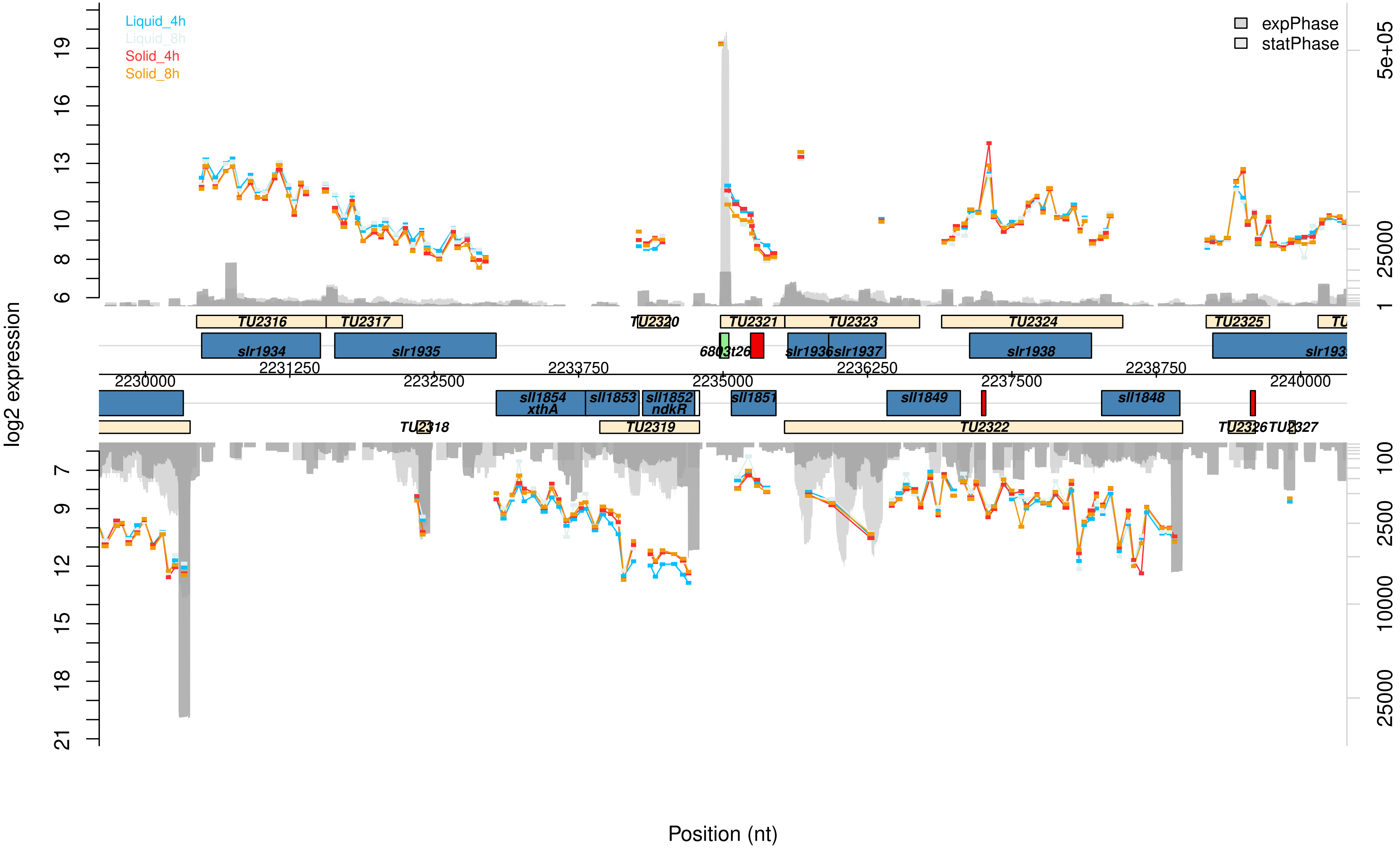

Liquid_4h
Liquid_8h
Solid_4h
Solid_8h
expPhase
statPhase
log2 expression
TU2316
TU2317
TU2320
TU2321
TU2323
TU2324
TU2325
slr1934
slr1935
6803t26
slr1936
slr1937
slr1938
2230000
2231250
2232500
2233750
2235000
2236250
2237500
2238750
2240000
sll1854
xthA
sll1853
sll1852
ndkR
sll1851
sll1849
sll1848
TU2318
TU2319
TU2322
TU2326
TU2327
Position (nt)

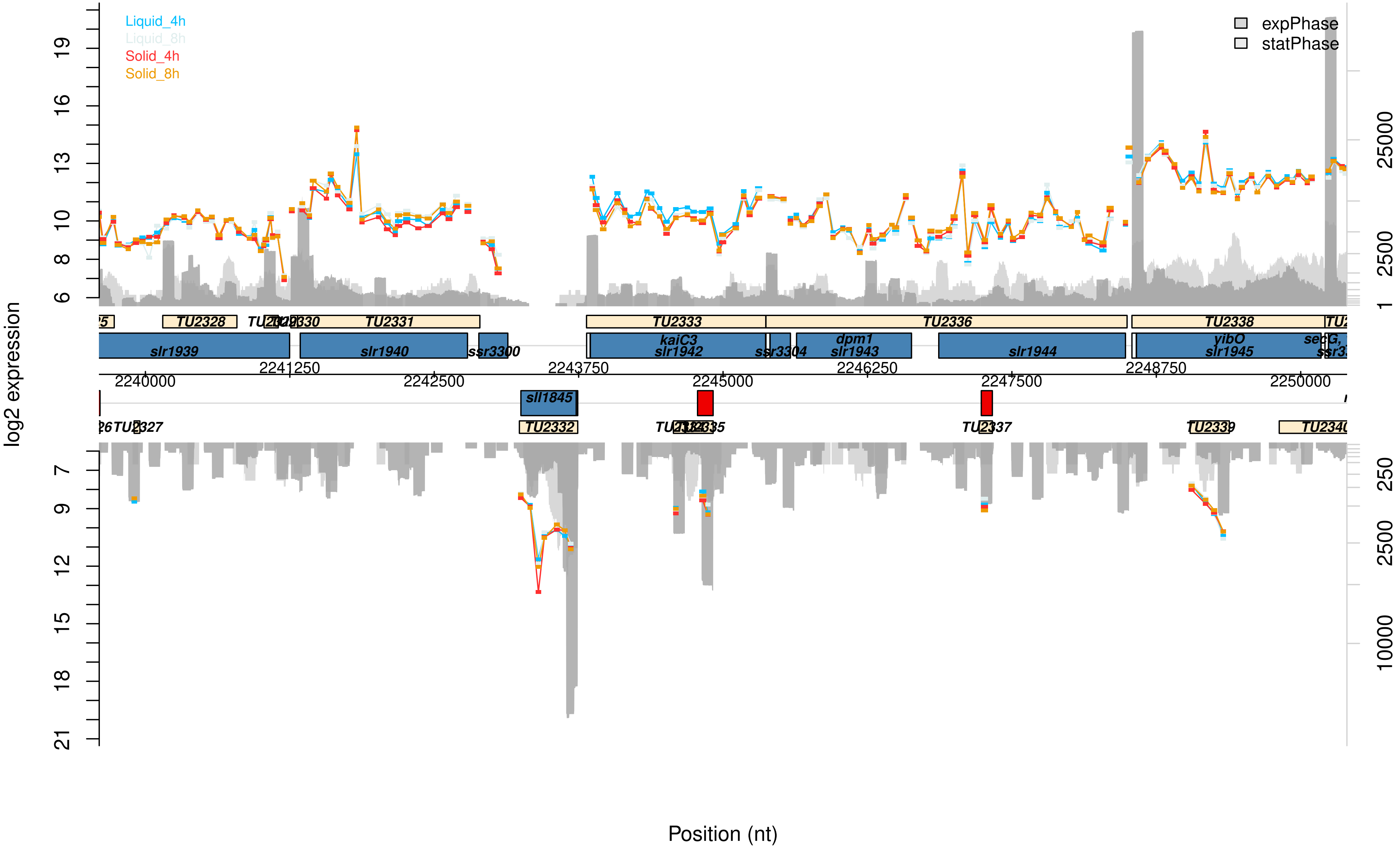

Liquid_4h
Liquid_8h
Solid_4h
Solid_8h
expPhase
statPhase
log2 expression
Position (nt)
TU2328
TU2331
TU2333
TU2336
TU2338
slr1939
slr1940
ssr3300
kaiC3
slr1942
dpm1
slr1943
slr1944
yibO
slr1945
sll1845
TU2332
TU2337
TU2339
2240000
2241250
2242500
2243750
2245000
2246250
2247500
2248750
2250000

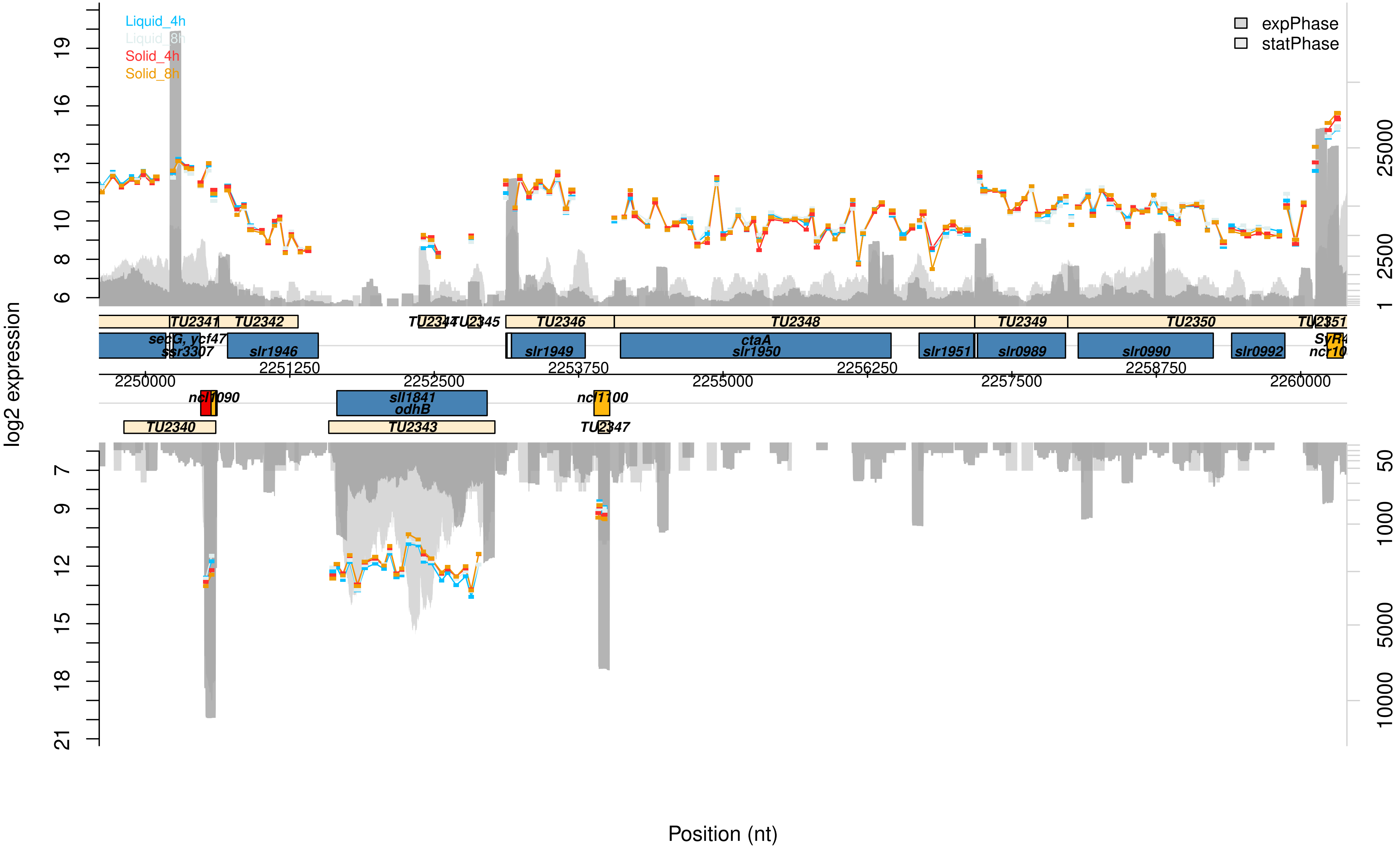
Liquid_4h
Liquid_8h
Solid_4h
Solid_8h
expPhase
statPhase
log2 expression
Position (nt)
TU2341
TU2342
TU2344
TU2345
TU2346
TU2348
TU2349
TU2350
TU2351
secG, ycf47
ssr3307
slr1946
slr1949
ctaA
slr1950
slr1951
slr0989
slr0990
slr0992
ncl1090
sll1841
odhB
ncl1100
TU2340
TU2343
TU2347
2250000
2251250
2252500
2253750
2255000
2256250
2257500
2258750
2260000

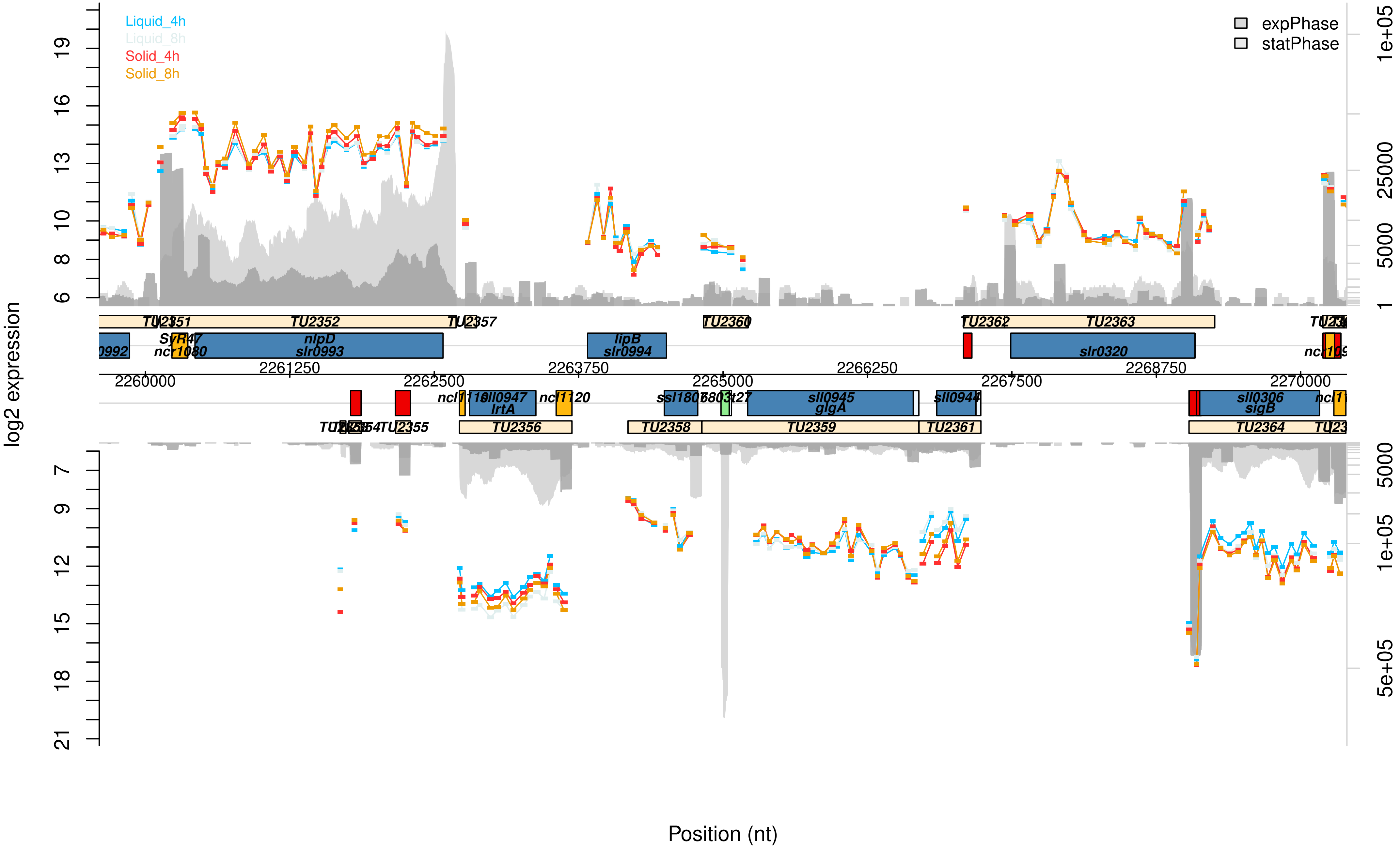

Liquid_4h
Liquid_8h
Solid_4h
Solid_8h
expPhase
statPhase
log2 expression
Position (nt)
TU2351
TU2352
TU2357
TU2360
TU2362
TU2363
SyR47
ncr1080
nlpD
slr0993
lipB
slr0994
slr0320
2261250
2263750
2266250
2268750
2260000
2262500
2265000
2267500
2270000
sll0947
lrtA
ncl1120
glgA
sll0945
sll0944
sll0306
sigB
TU2355
TU2356
TU2358
TU2359
TU2361
TU2364

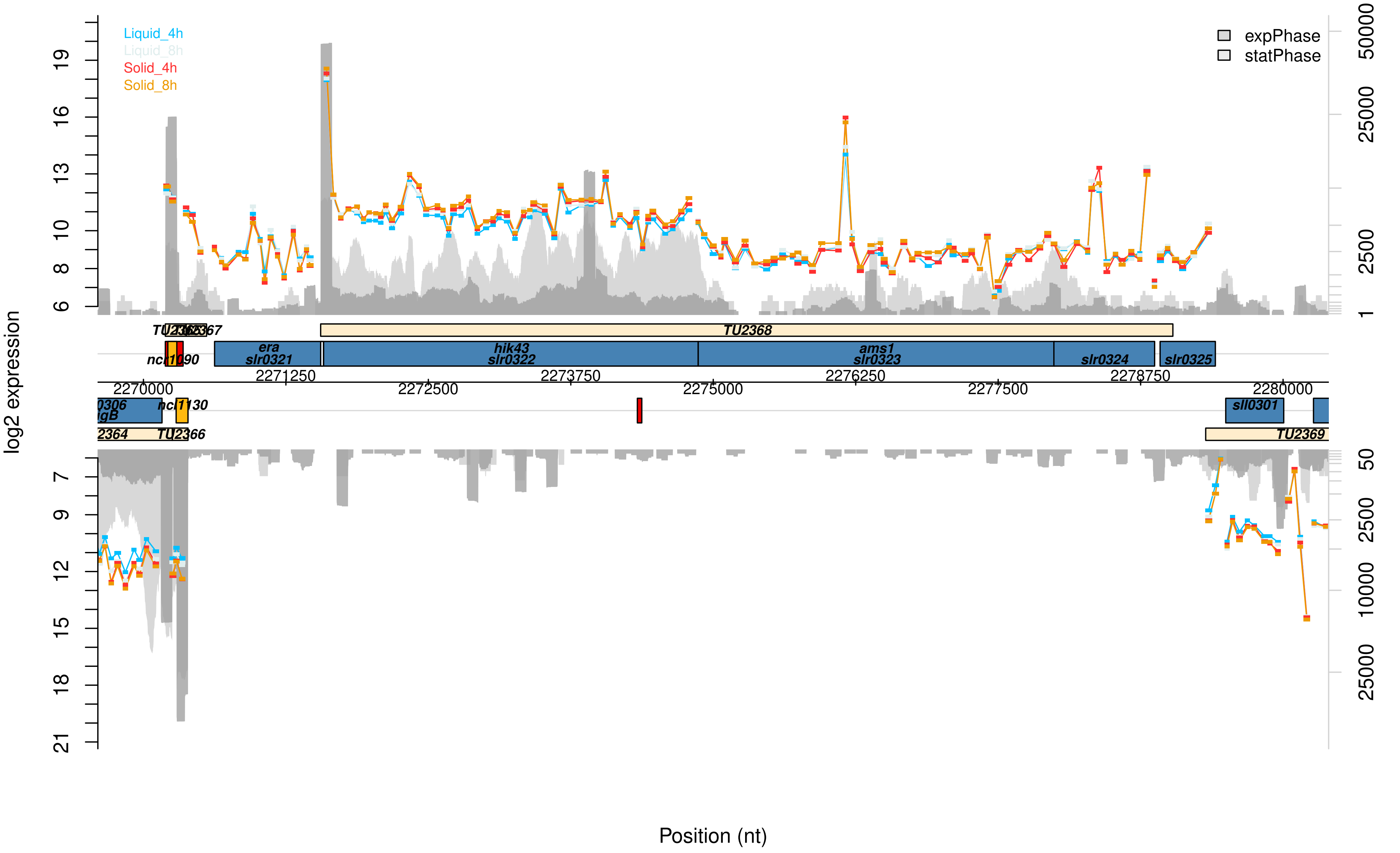
Liquid_4h
Liquid_8h
Solid_4h
Solid_8h
expPhase
statPhase
log2 expression
Position (nt)
TU2368
era
slr0321
hik43
slr0322
ams1
slr0323
slr0324
slr0325
ncl1090
ncl1130
sll0301
TU2366
TU2369
2270000
2271250
2272500
2273750
2275000
2276250
2277500
2278750
2280000

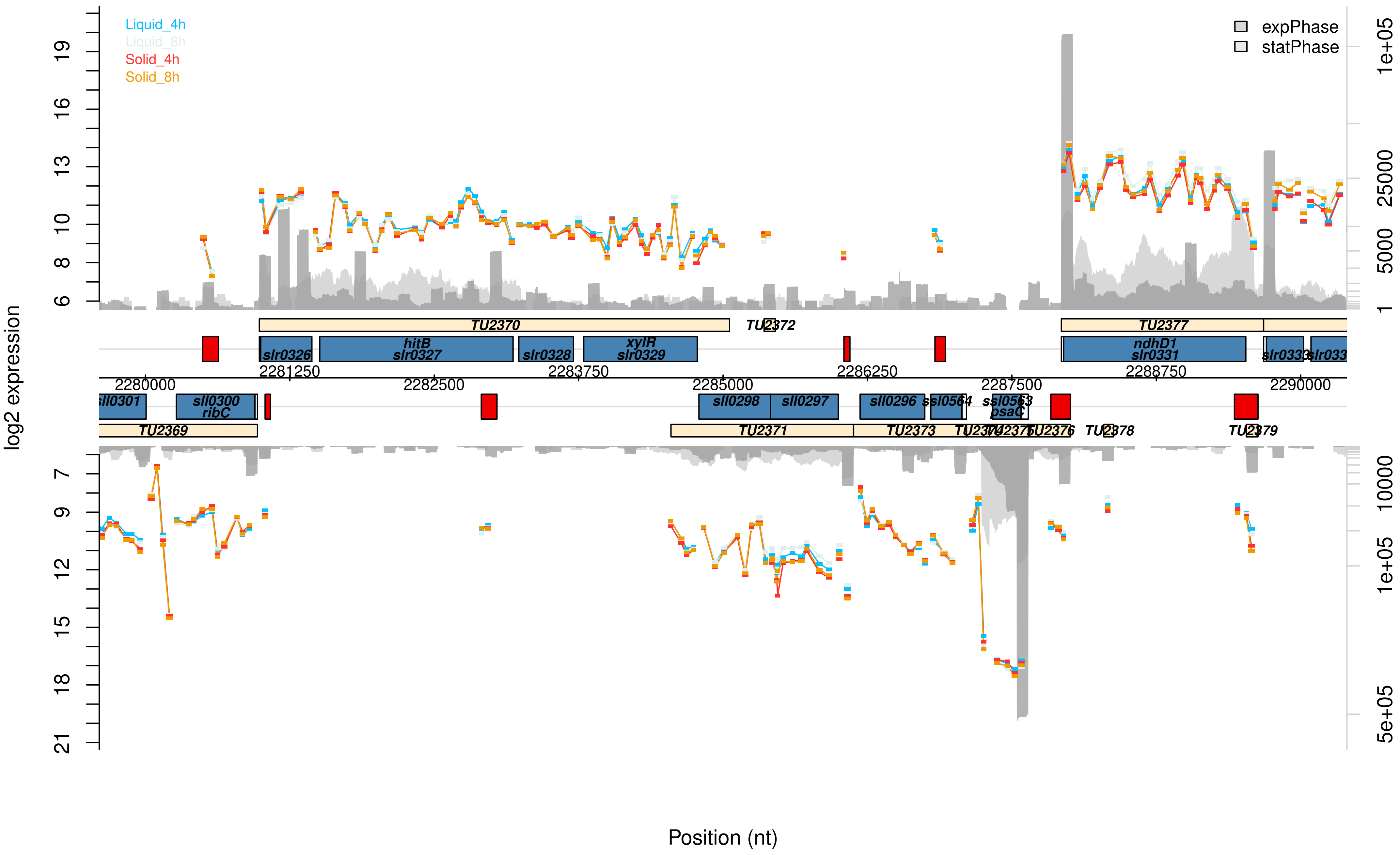

Liquid_4h
Liquid_8h
Solid_4h
Solid_8h
expPhase
statPhase
log2 expression
Position (nt)
TU2370
TU2372
TU2377
slr0326
hitB
slr0327
slr0328
xylR
slr0329
ndhD1
slr0331
sll0301
sll0300
ribC
sll0298
sll0297
sll0296
ssl0564
ssl0563
psaC
TU2369
TU2371
TU2373
TU2376
TU2378
TU2379
2280000
2281250
2282500
2283750
2285000
2286250
2287500
2288750
2290000

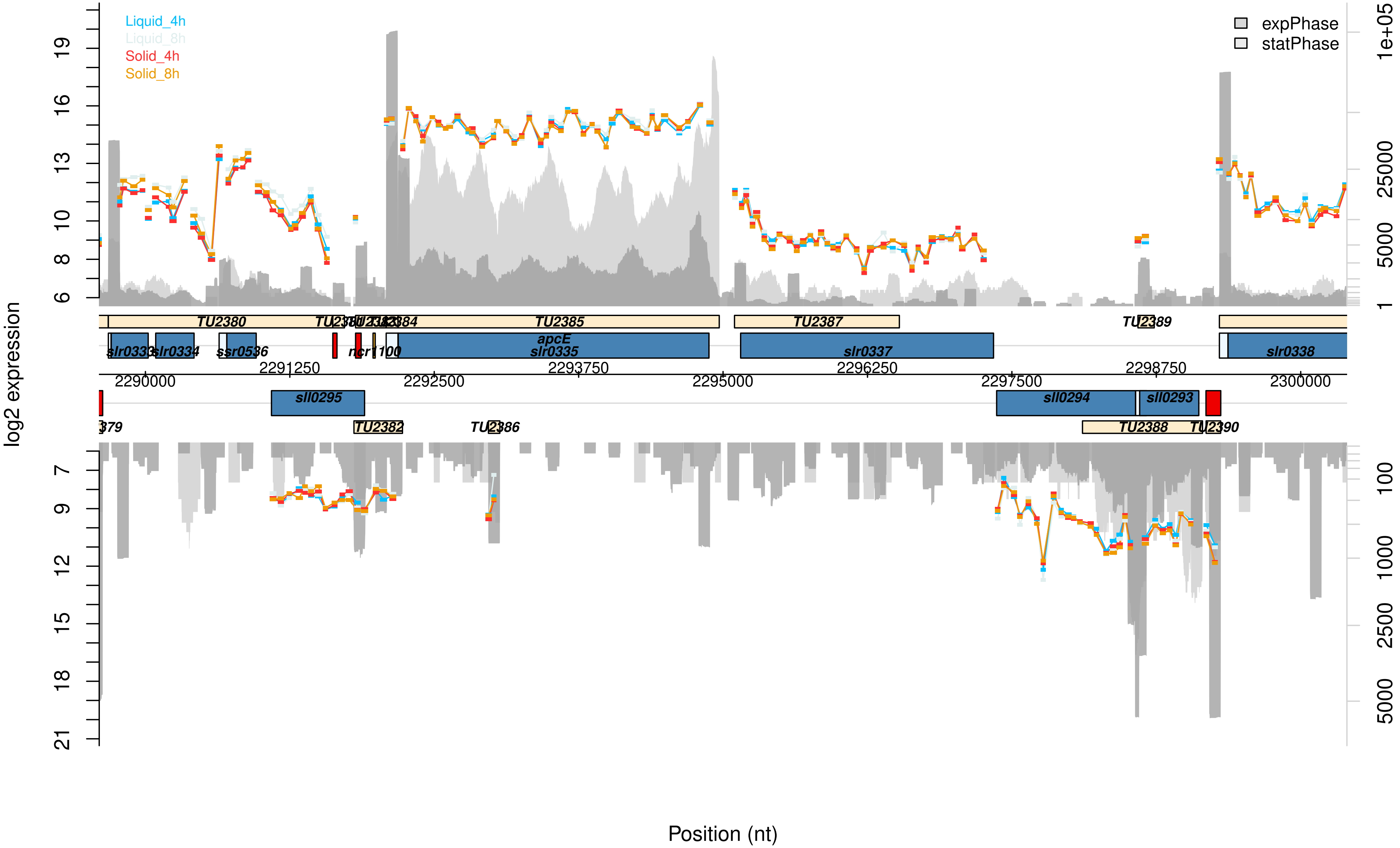

Liquid_4h
Liquid_8h
Solid_4h
Solid_8h
expPhase
statPhase
log2 expression
TU2380
TU2385
TU2387
TU2389
slr0333
slr0334
ssr0536
ncr1100
apcE
slr0335
slr0337
slr0338
2290000
2291250
2292500
2293750
2295000
2296250
2297500
2298750
2300000
sll0295
sll0294
sll0293
TU2382
TU2386
TU2388
Position (nt)

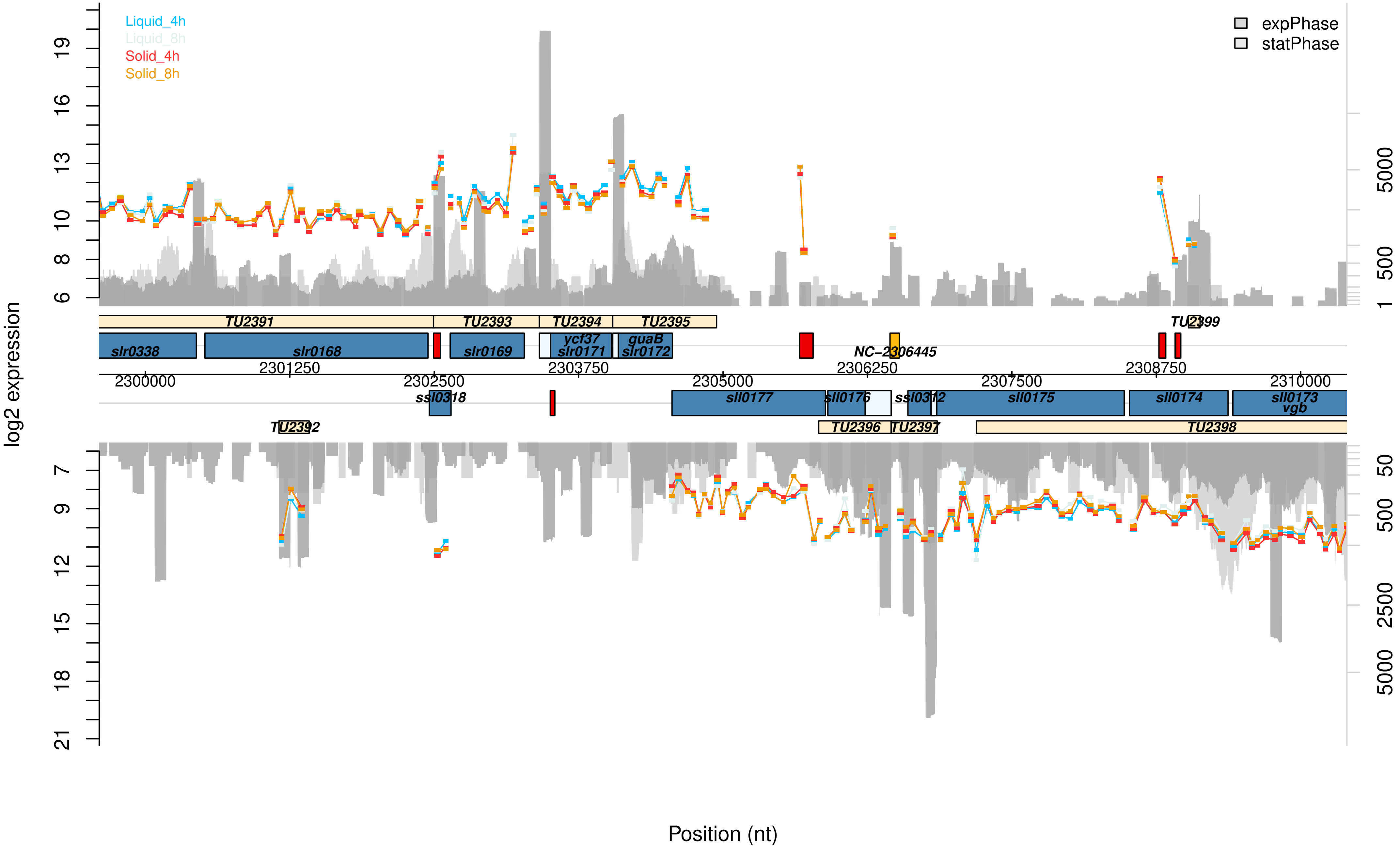

Liquid_4h
Liquid_8h
Solid_4h
Solid_8h
expPhase
statPhase
log2 expression
TU2391
TU2393
TU2394
TU2395
TU2399
slr0338
slr0168
slr0169
ycf37
slr0171
guaB
slr0172
NC-2306445
2300000
2301250
2302500
2303750
2305000
2306250
2307500
2308750
2310000
ssl0318
sll0177
sll0176
ssl0312
sll0175
sll0174
sll0173
vgb
TU2392
TU2396
TU2397
TU2398
Position (nt)

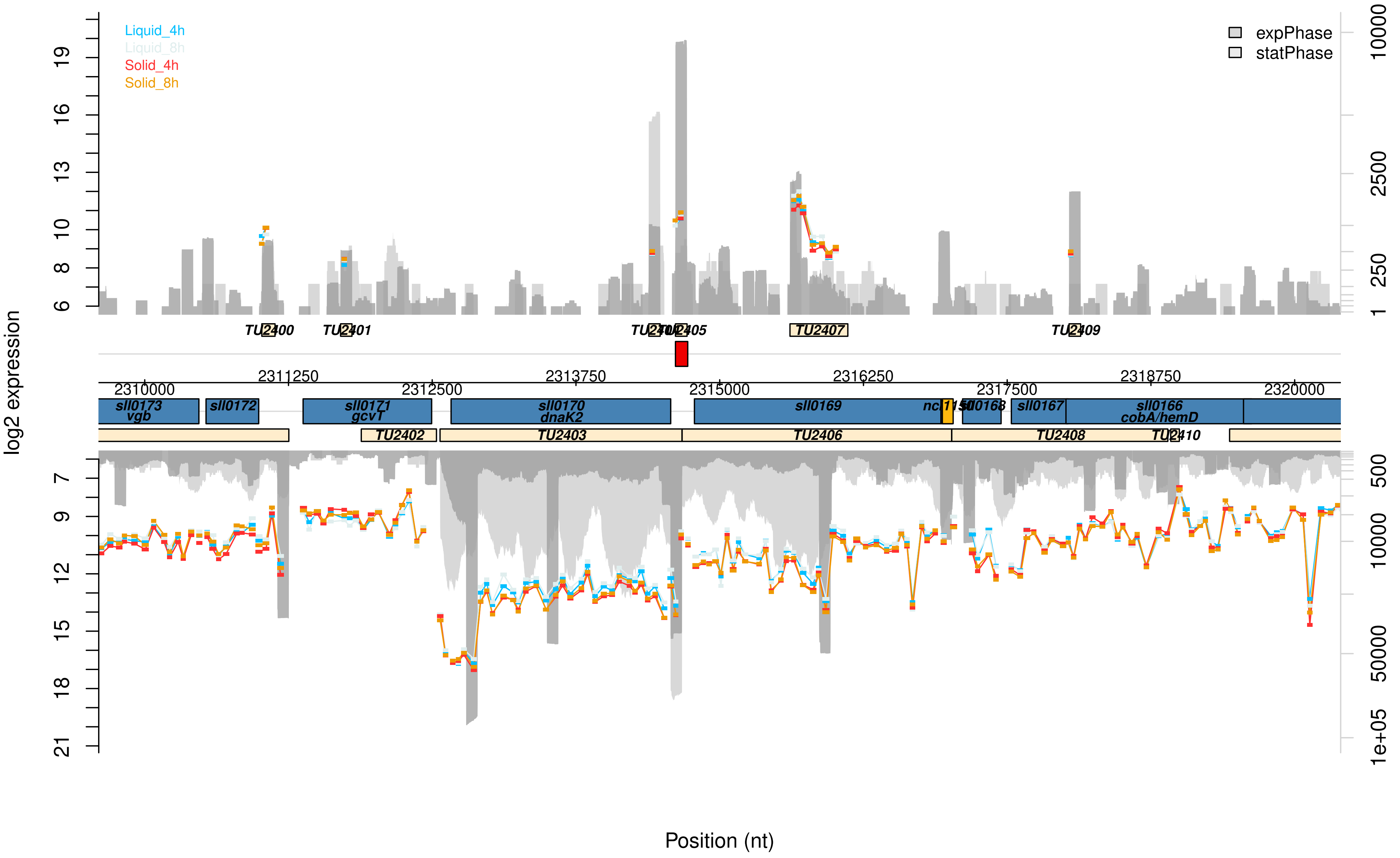
Liquid_4h
Liquid_8h
Solid_4h
Solid_8h
expPhase
statPhase
log2 expression
Position (nt)
TU2400
TU2401
TU2405
TU2407
TU2409
TU2402
TU2403
TU2406
TU2408
sll0173
vgb
sll0172
sll0171
gcvT
sll0170
dnaK2
sll0169
sll0167
sll0166
cobA/hemD
2310000
2311250
2312500
2313750
2315000
2316250
2317500
2318750
2320000

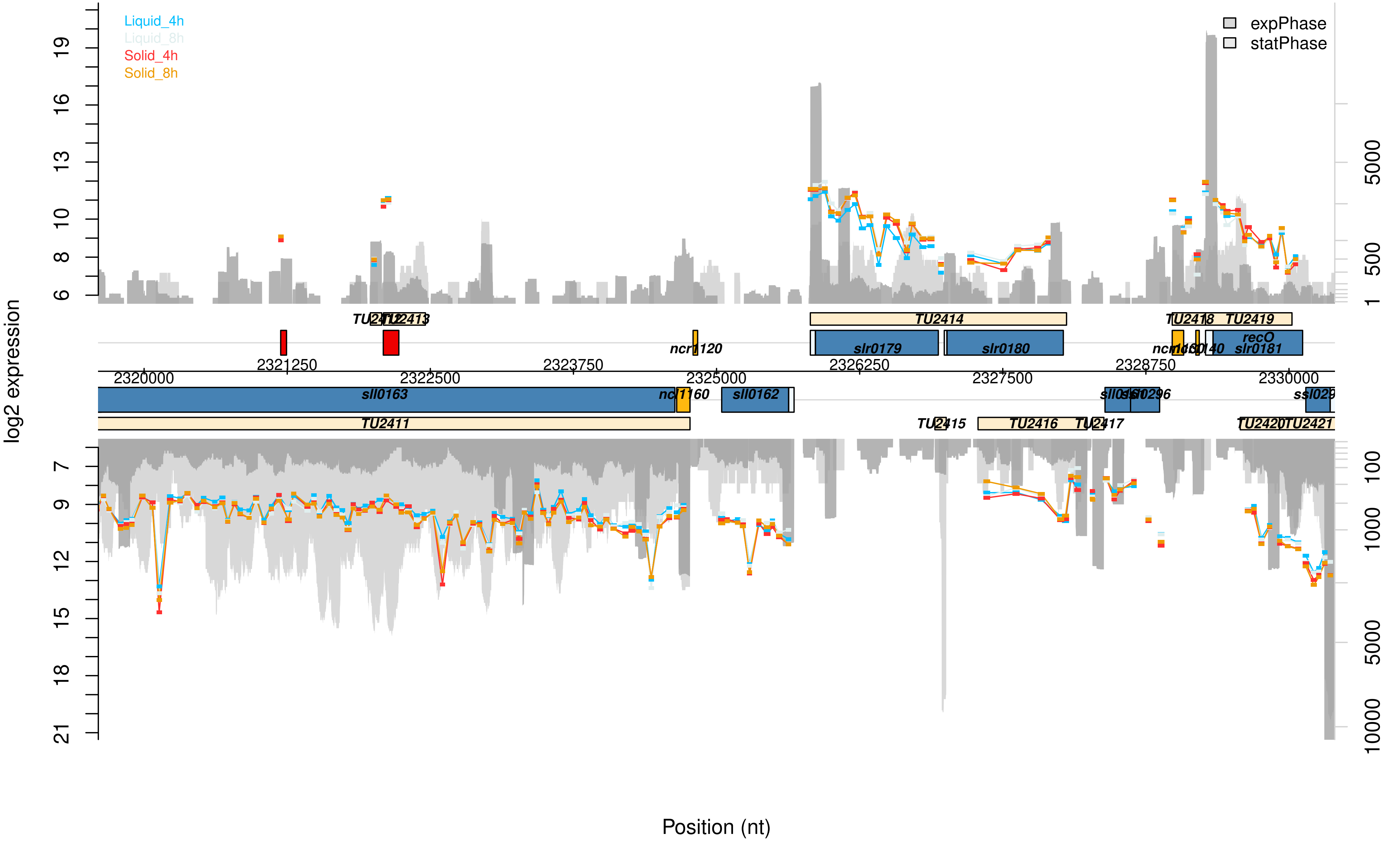
Liquid_4h
Liquid_8h
Solid_4h
Solid_8h
expPhase
statPhase
log2 expression
Position (nt)
TU2411
TU2414
TU2415
TU2416
TU2417
TU2419
sll0163
sll0162
slr0179
slr0180
recO
slr0181
ncr1120
2320000
2321250
2322500
2323750
2325000
2326250
2327500
2328750
2330000

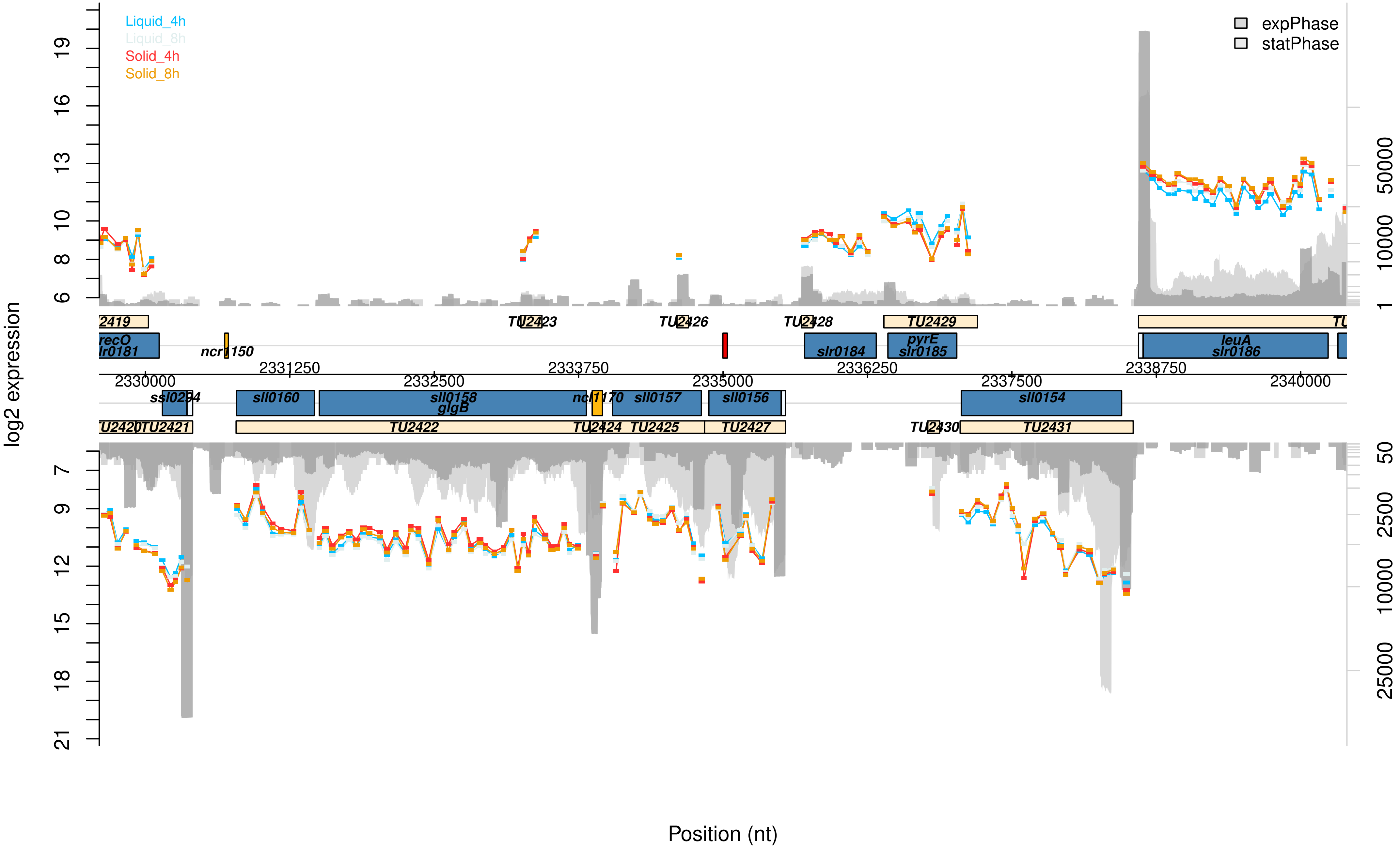

Liquid_4h
Liquid_8h
Solid_4h
Solid_8h
expPhase
statPhase
log2 expression
Position (nt)
2419
TU2423
TU2426
TU2428
TU2429
recO
slr0181
ncr1150
slr0184
pyrE
slr0185
leuA
slr0186
2330000
2331250
2332500
2333750
2335000
2336250
2337500
2338750
2340000
ssl0294
sll0160
sll0158
glgB
ncr1170
sll0157
sll0156
sll0154
TU2420
TU2421
TU2422
TU2424
TU2425
TU2427
TU2430
TU2431

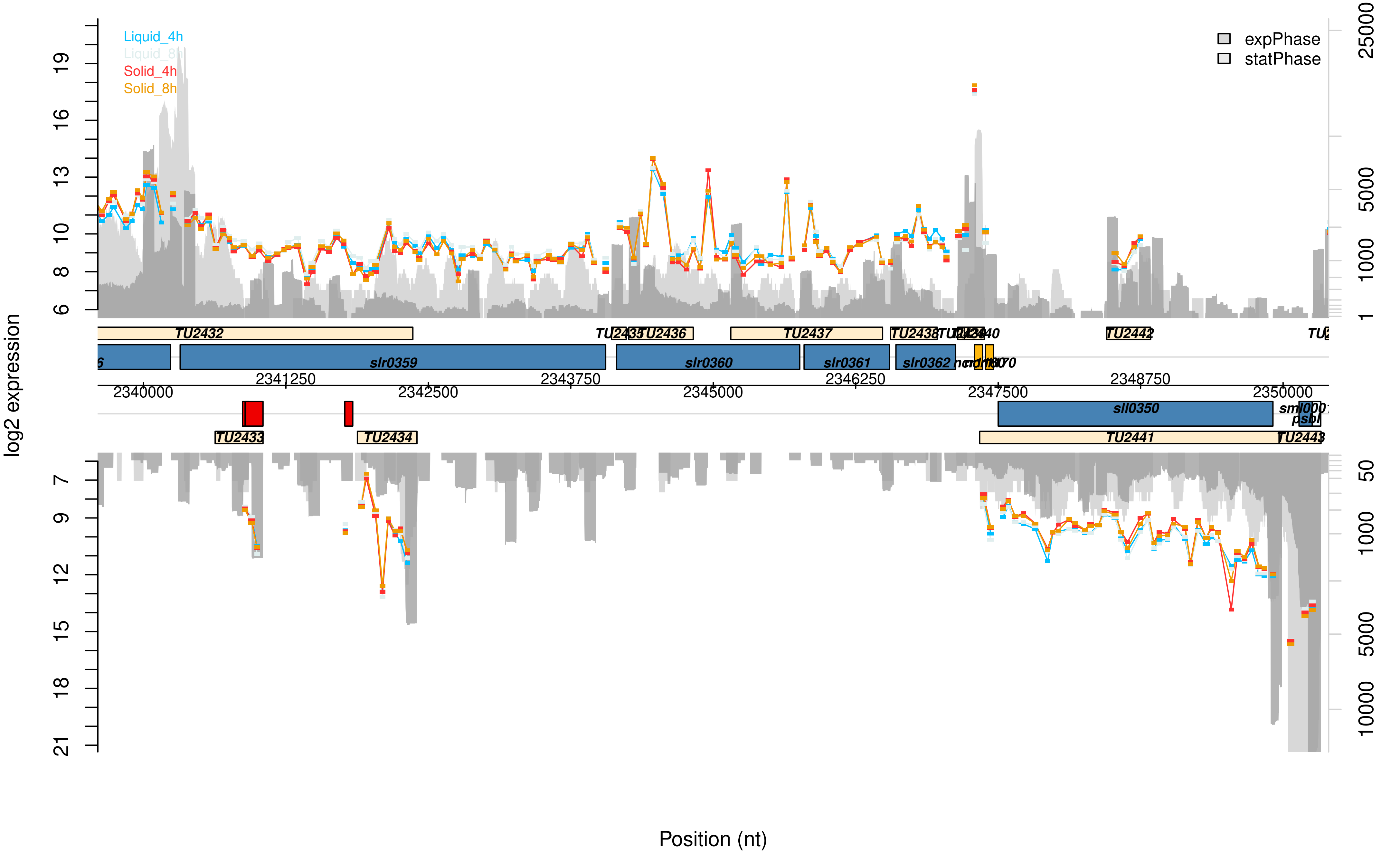

Liquid_4h
Liquid_8h
Solid_4h
Solid_8h
expPhase
statPhase
log2 expression
Position (nt)
TU2432
TU2435
TU2436
TU2437
TU2438
TU2442
slr0359
slr0360
slr0361
slr0362
sll0350
TU2433
TU2434
TU2441
TU2443
2340000
2341250
2342500
2343750
2345000
2346250
2347500
2348750
2350000

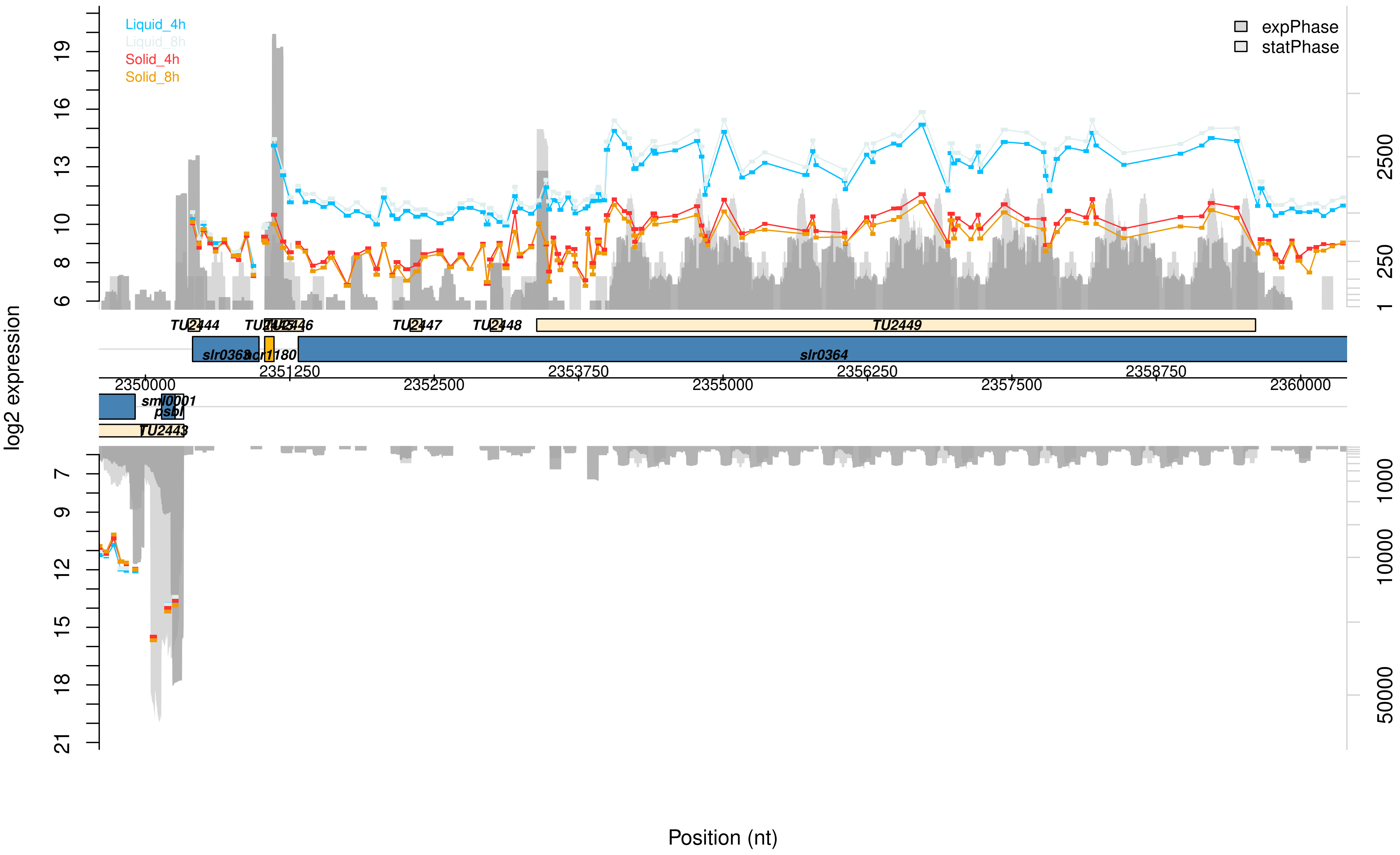

Liquid_4h
Liquid_8h
Solid_4h
Solid_8h
expPhase
statPhase
log2 expression
TU2444
TU2445
TU2446
TU2447
TU2448
TU2449
slr0363
ncr1180
slr0364
2350000
2351250
2352500
2353750
2355000
2356250
2357500
2358750
2360000
sml0001
psbI
TU2443
Position (nt)

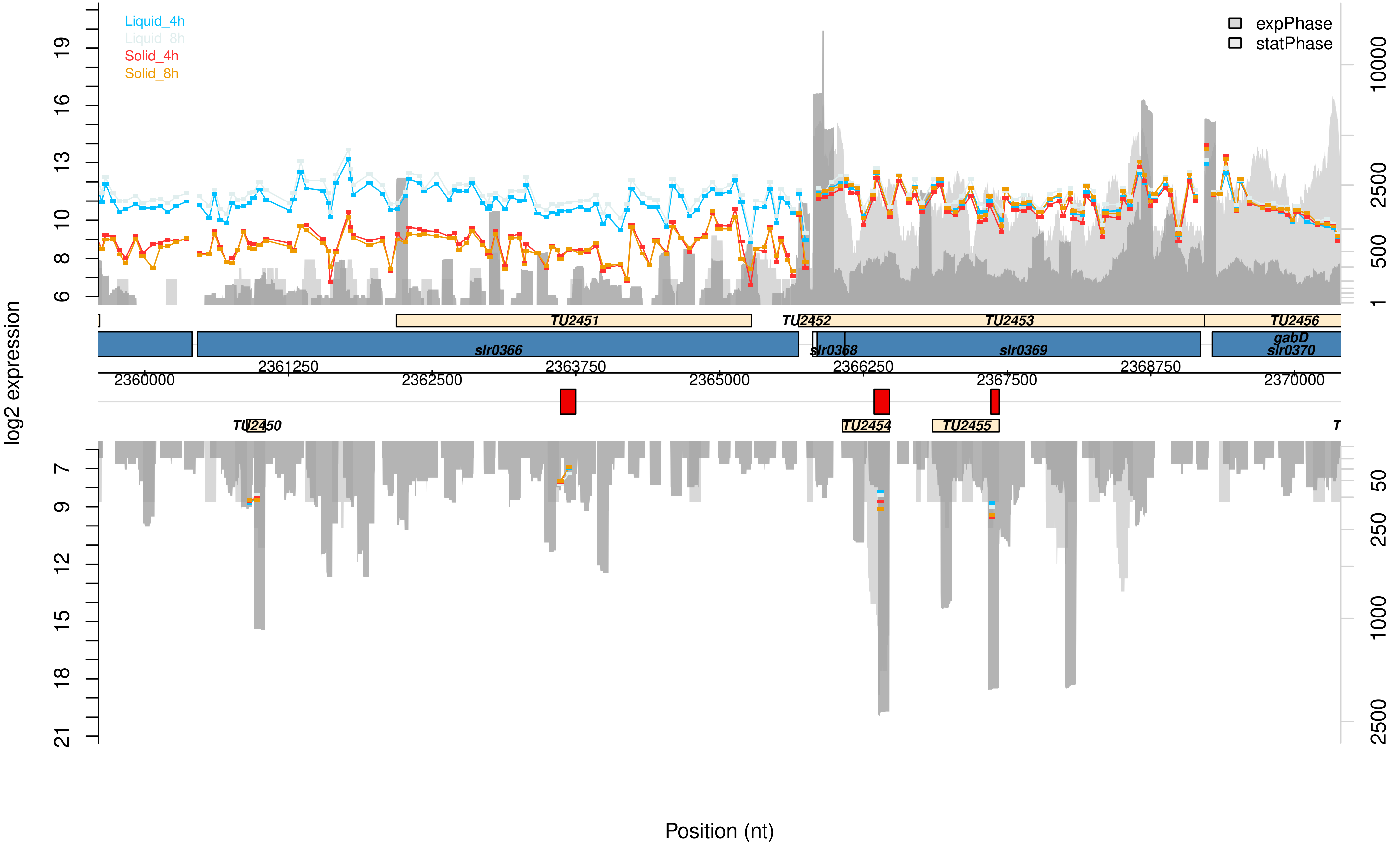

Liquid_4h
Liquid_8h
Solid_4h
Solid_8h
expPhase
statPhase
log2 expression
TU2451
TU2452
TU2453
TU2456
slr0366
slr0368
slr0369
gabD
slr0370
TU2450
TU2454
TU2455
2360000
2361250
2362500
2363750
2365000
2366250
2367500
2368750
2370000
Position (nt)

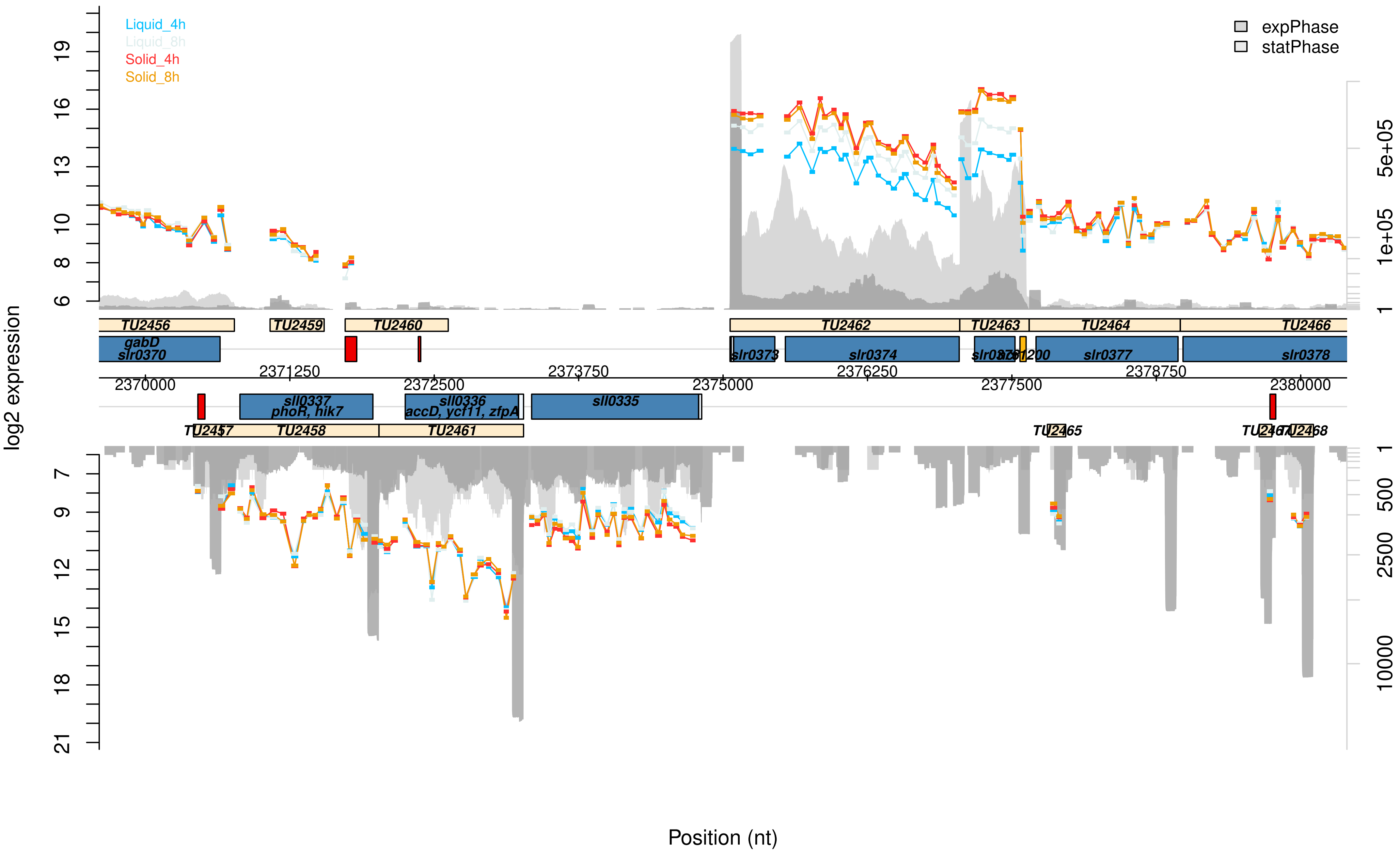

Liquid_4h
Liquid_8h
Solid_4h
Solid_8h
expPhase
statPhase
log2 expression
TU2456
TU2459
TU2460
TU2462
TU2463
TU2464
TU2466
gabD
slr0370
slr0373
slr0374
slr0377
slr0378
sll0337
phoR, hik7
sll0336
accD, ycf11, zfpA
sll0335
TU2458
TU2461
TU2465
TU2468
Position (nt)

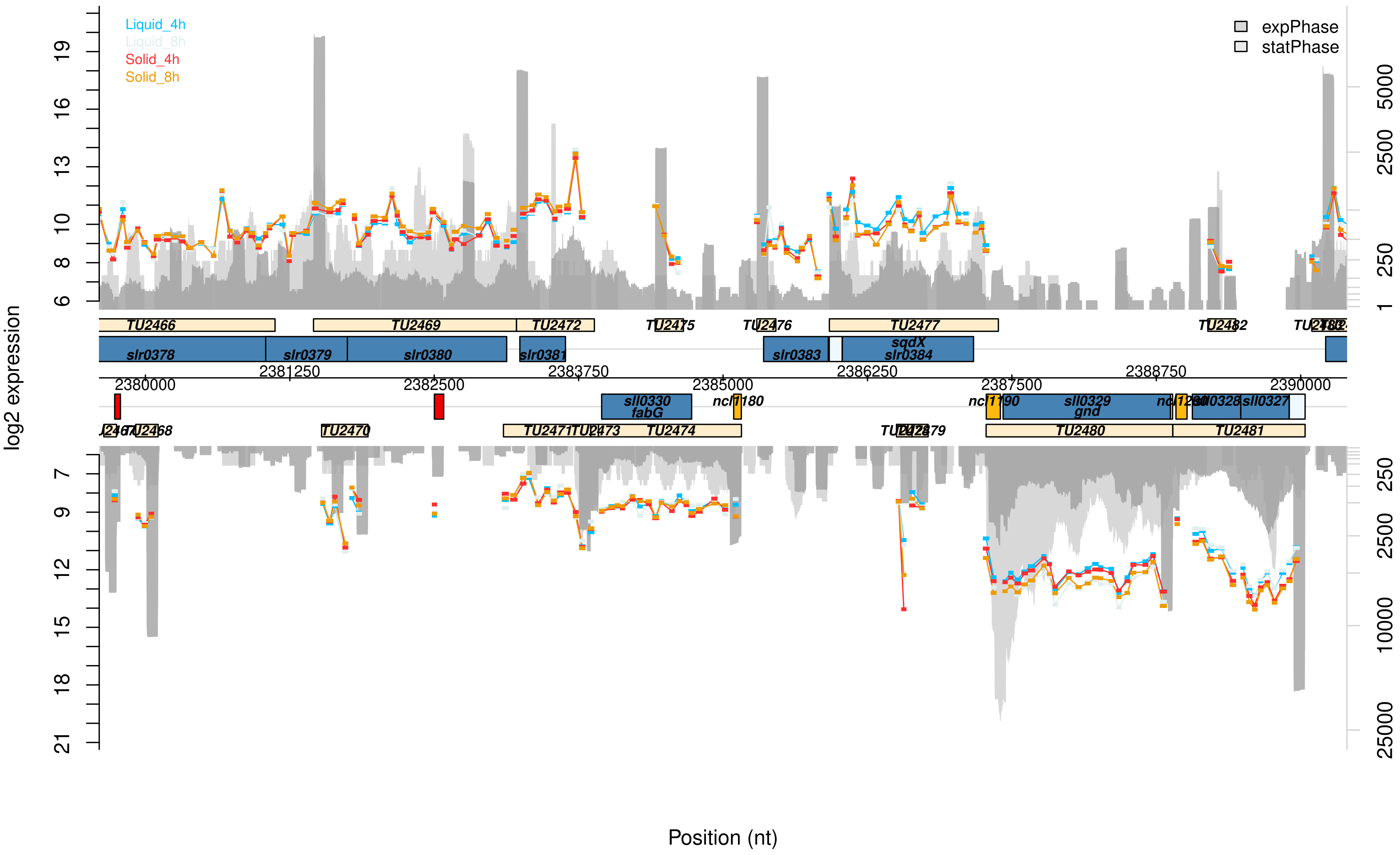
Liquid_4h
Liquid_8h
Solid_4h
Solid_8h
expPhase
statPhase
log2 expression
Position (nt)
TU2466
TU2469
TU2472
TU2475
TU2476
TU2477
TU2482
slr0378
slr0379
slr0380
slr0381
slr0383
sqdX
slr0384
2381250
2383750
2386250
2388750
2380000
2382500
2385000
2387500
2390000
sll0330
fabG
ncl1180
sll0329
gnd
sll0327
TU2468
TU2470
TU2474
TU2480
TU2481

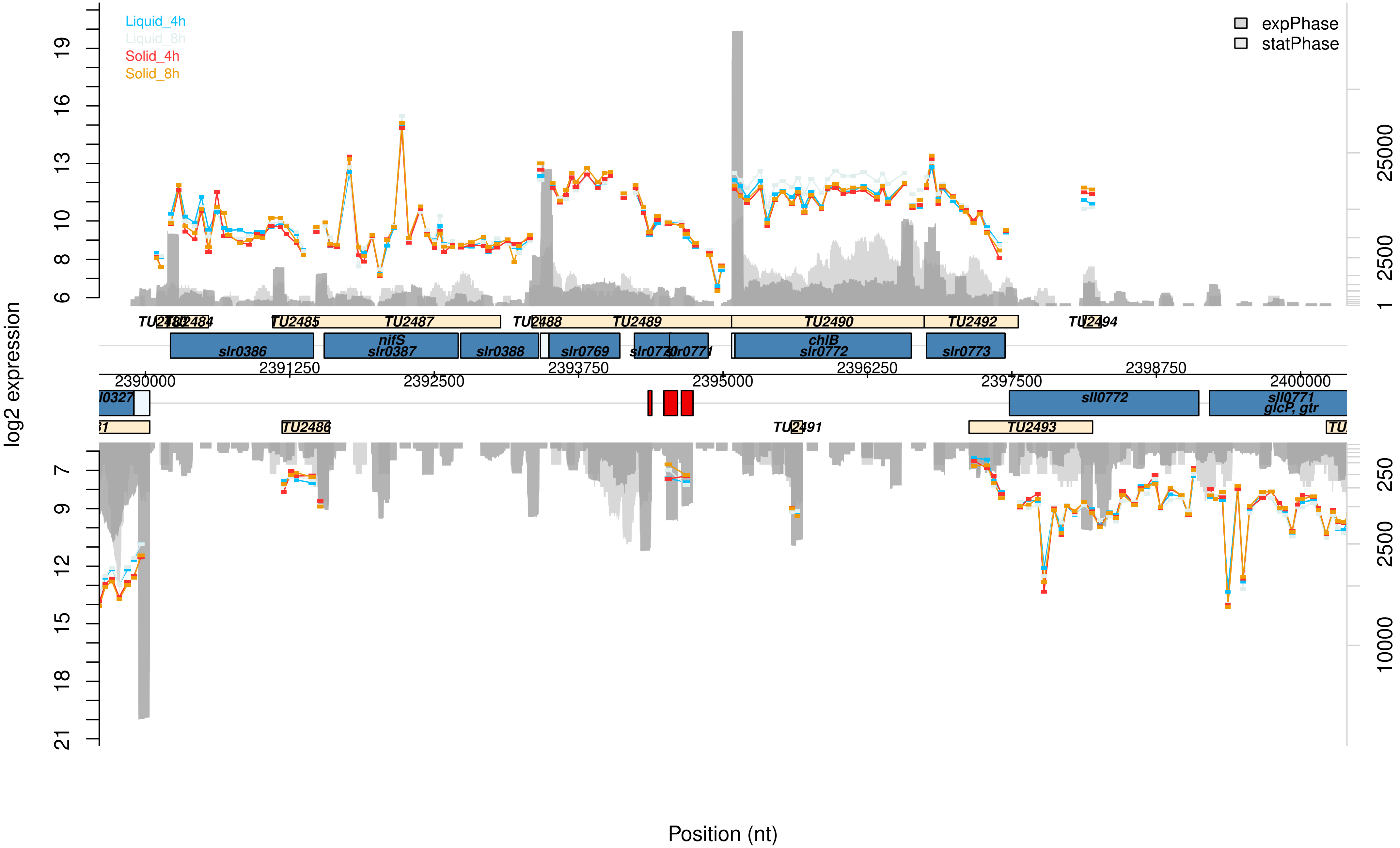
Liquid_4h
Liquid_8h
Solid_4h
Solid_8h
expPhase
statPhase
log2 expression
TU2484
TU2485
TU2487
TU2488
TU2489
TU2490
TU2492
TU2494
slr0386
nifS
slr0387
slr0388
slr0769
slr0771
chlB
slr0772
slr0773
2390000
2391250
2392500
2393750
2395000
2396250
2397500
2398750
2400000
sll0772
sll0771
glcP, gtr
TU2486
TU2491
TU2493
Position (nt)

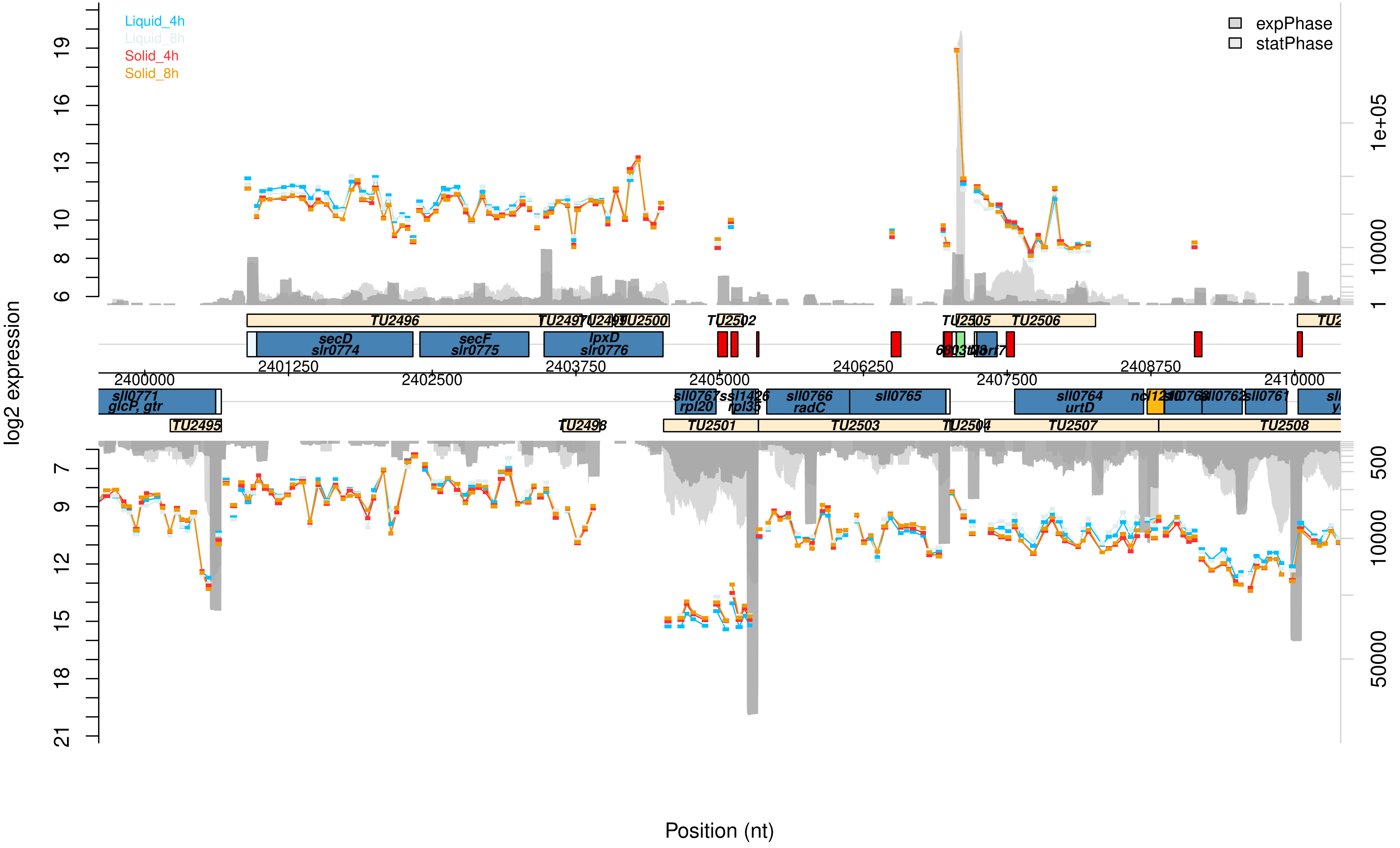

Liquid_4h
Liquid_8h
Solid_4h
Solid_8h
expPhase
statPhase
log2 expression
Position (nt)
TU2496
TU2497
TU2499
TU2500
TU2502
TU2505
TU2506
secD
slr0774
secF
slr0775
lpxD
slr0776
2400000
2401250
2402500
2403750
2405000
2406250
2407500
2408750
2410000
sll0771
glcP, gtr
sll0767
rpl20
ssl1426
rpl35
sll0766
radC
sll0765
sll0764
urtD
sll0763
sll0762
sll0761
TU2495
TU2498
TU2501
TU2503
TU2504
TU2507
TU2508

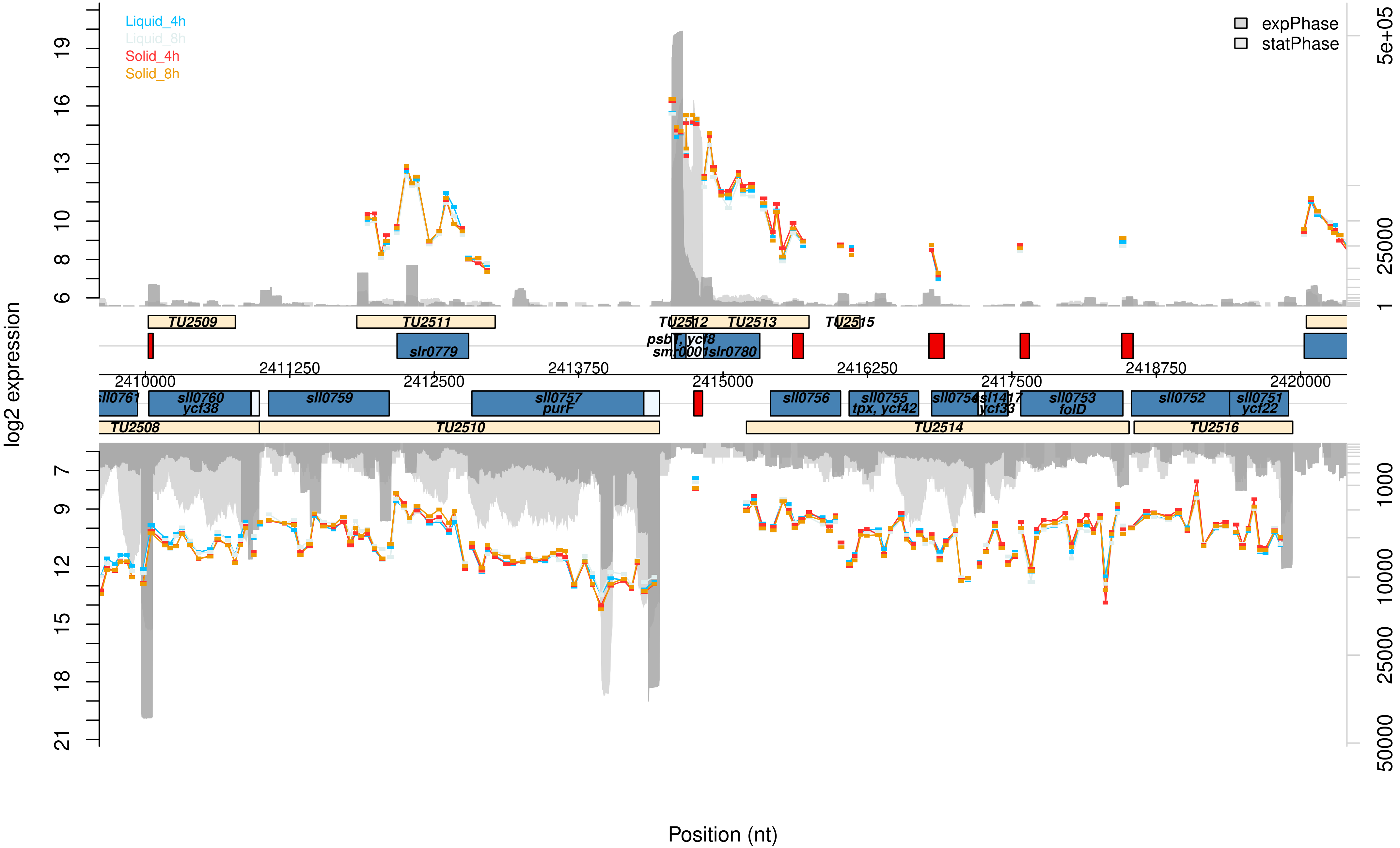

Liquid_4h
Liquid_8h
Solid_4h
Solid_8h
expPhase
statPhase
log2 expression
Position (nt)
TU2509
TU2511
TU2512
TU2513
TU2515
slr0779
psbT, ycf8
smr0001
slr0780
2410000
2411250
2412500
2413750
2415000
2416250
2417500
2418750
2420000
sll0761
sll0760
ycf38
sll0759
sll0757
purF
sll0756
sll0755
tpx, ycf42
sll0754
ssl1417
ycf33
sll0753
folD
sll0752
sll0751
ycf22
TU2508
TU2510
TU2514
TU2516

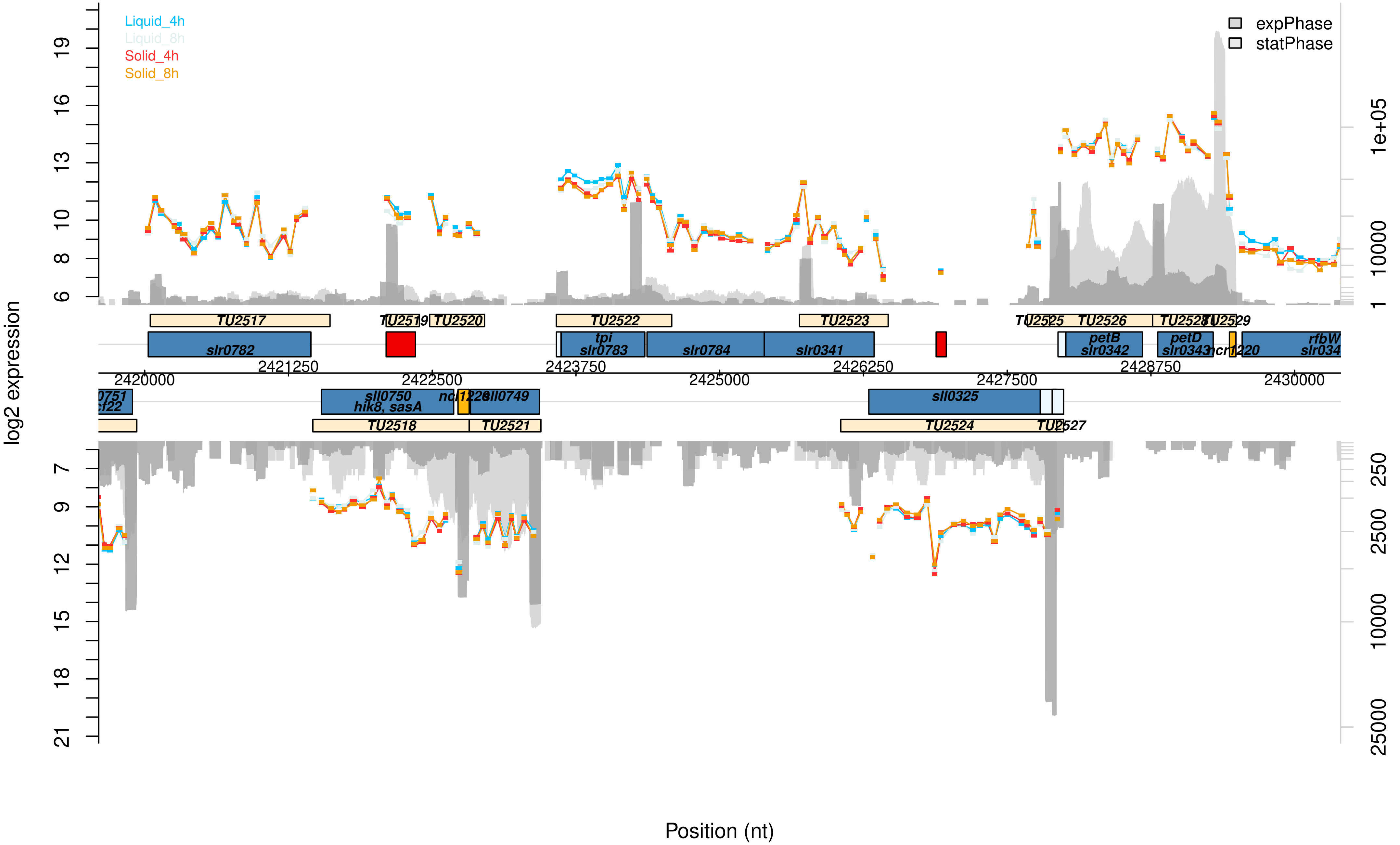

Liquid_4h
Liquid_8h
Solid_4h
Solid_8h
expPhase
statPhase
log2 expression
Position (nt)
TU2517
TU2519
TU2520
TU2522
TU2523
TU2525
TU2526
TU2528
TU2529
slr0782
tpi
slr0783
slr0784
slr0341
petB
slr0342
petD
slr0343
ncr1220
rfbW
sll0750
hik8, sasA
sll0749
sll0325
TU2518
TU2521
TU2524
TU2527
2420000
2421250
2422500
2423750
2425000
2426250
2427500
2428750
2430000

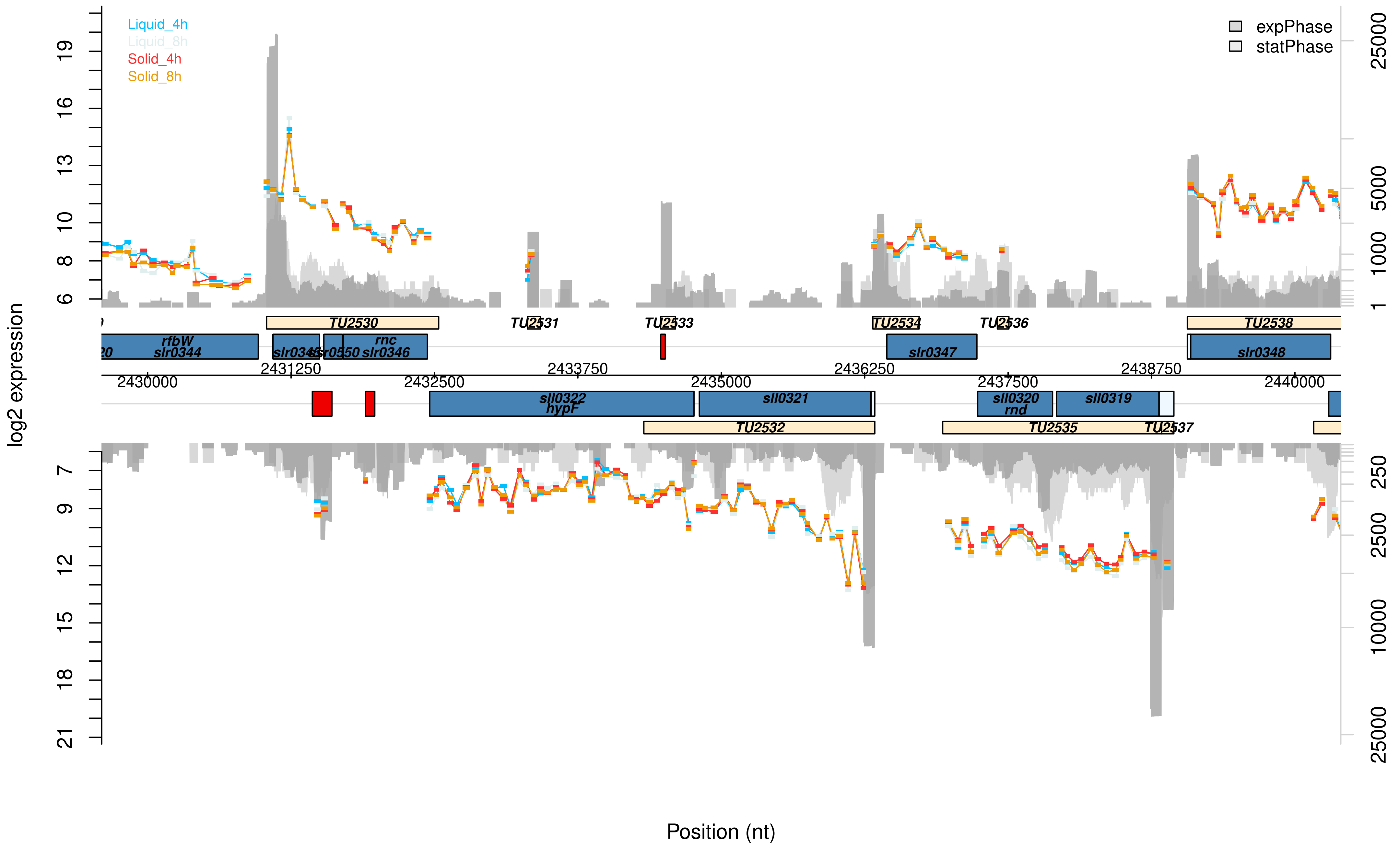

Liquid_4h
Liquid_8h
Solid_4h
Solid_8h
expPhase
statPhase
log2 expression
Position (nt)
rfbW
slr0344
slr0345
slr0350
rnc
slr0346
slr0347
slr0348
sll0322
hypF
sll0321
sll0320
rnd
sll0319
TU2530
TU2531
TU2532
TU2533
TU2534
TU2535
TU2536
TU2537
TU2538
2430000
2431250
2432500
2433750
2435000
2436250
2437500
2438750
2440000

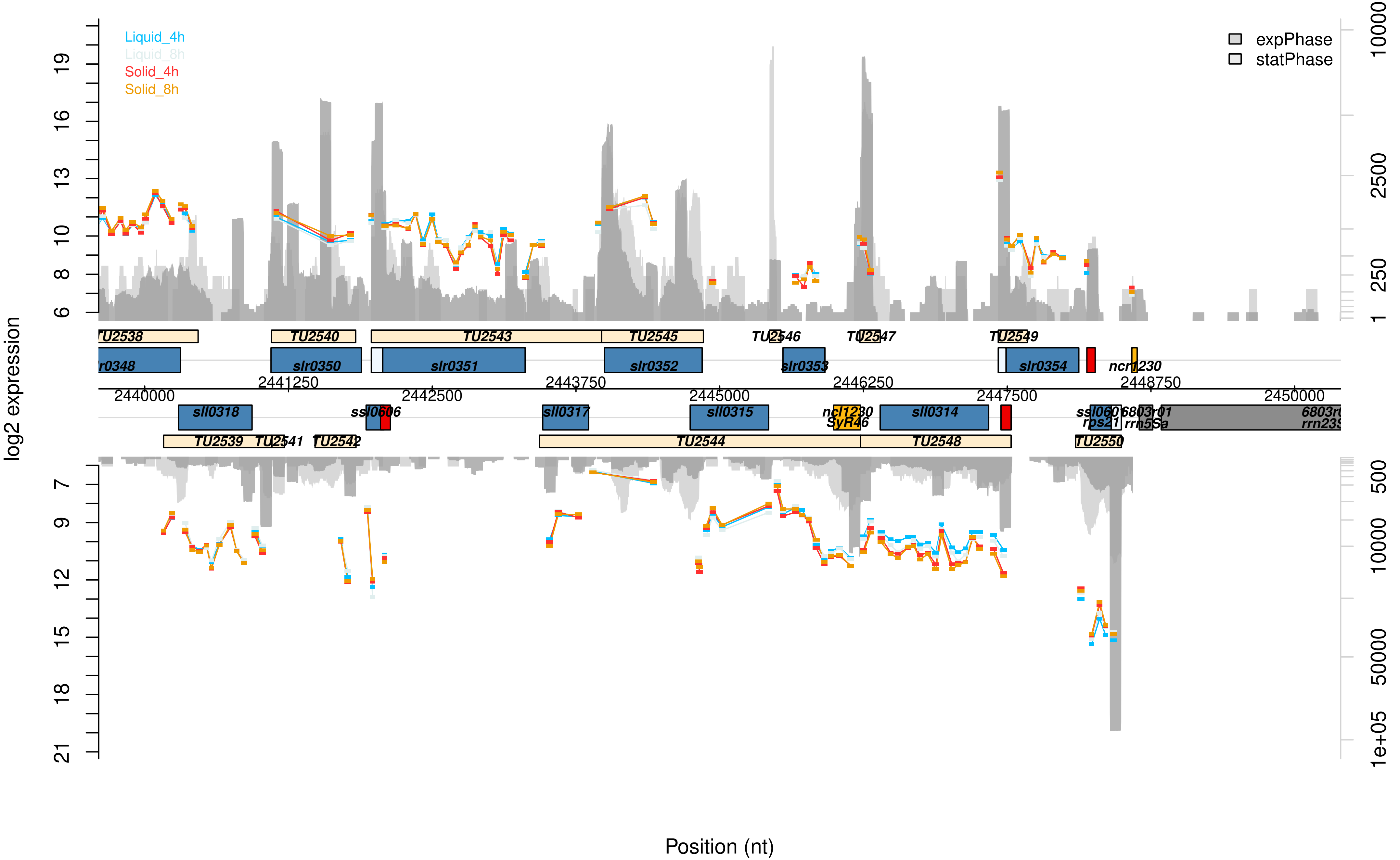

Liquid_4h
Liquid_8h
Solid_4h
Solid_8h
expPhase
statPhase
log2 expression
Position (nt)
TU2538
TU2540
TU2543
TU2545
TU2546
TU2547
TU2549
slr0350
slr0351
slr0352
slr0353
slr0354
ncr1230
2440000
2441250
2442500
2443750
2445000
2446250
2447500
2448750
2450000
sll0318
ssl0606
sll0317
sll0315
ncl1230
SyR46
sll0314
rps21
rrn5Sa
TU2539
TU2542
TU2544
TU2548
TU2550

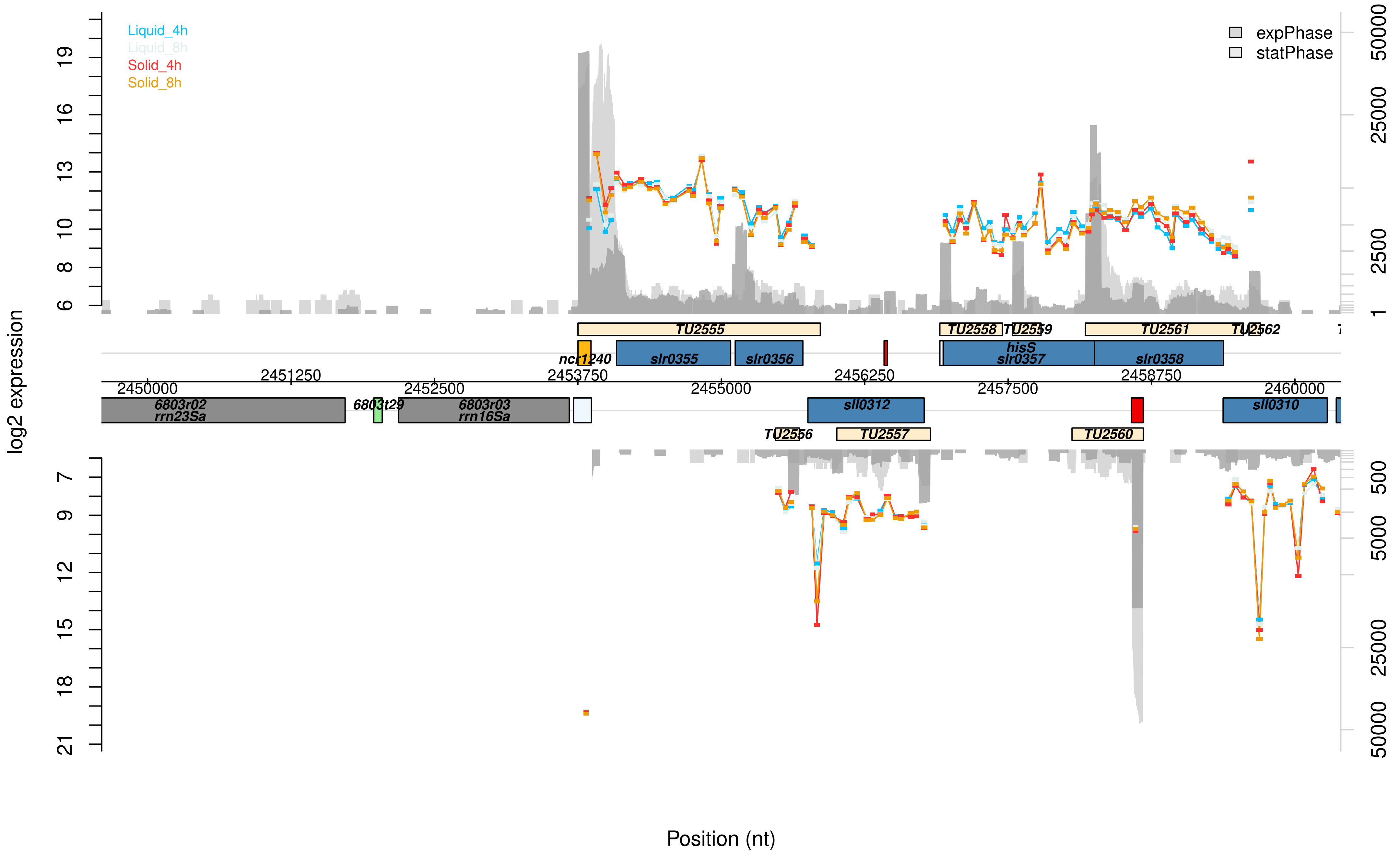

Liquid_4h
Liquid_8h
Solid_4h
Solid_8h
expPhase
statPhase
log2 expression
Position (nt)
TU2555
TU2558
TU2559
TU2561
TU2562
ncr1240
slr0355
slr0356
hisS
slr0357
slr0358
2450000
2451250
2452500
2453750
2455000
2456250
2457500
2458750
2460000
6803r02
rrn23Sa
6803t29
6803r03
rrn16Sa
sll0312
sll0310
TU2556
TU2557
TU2560

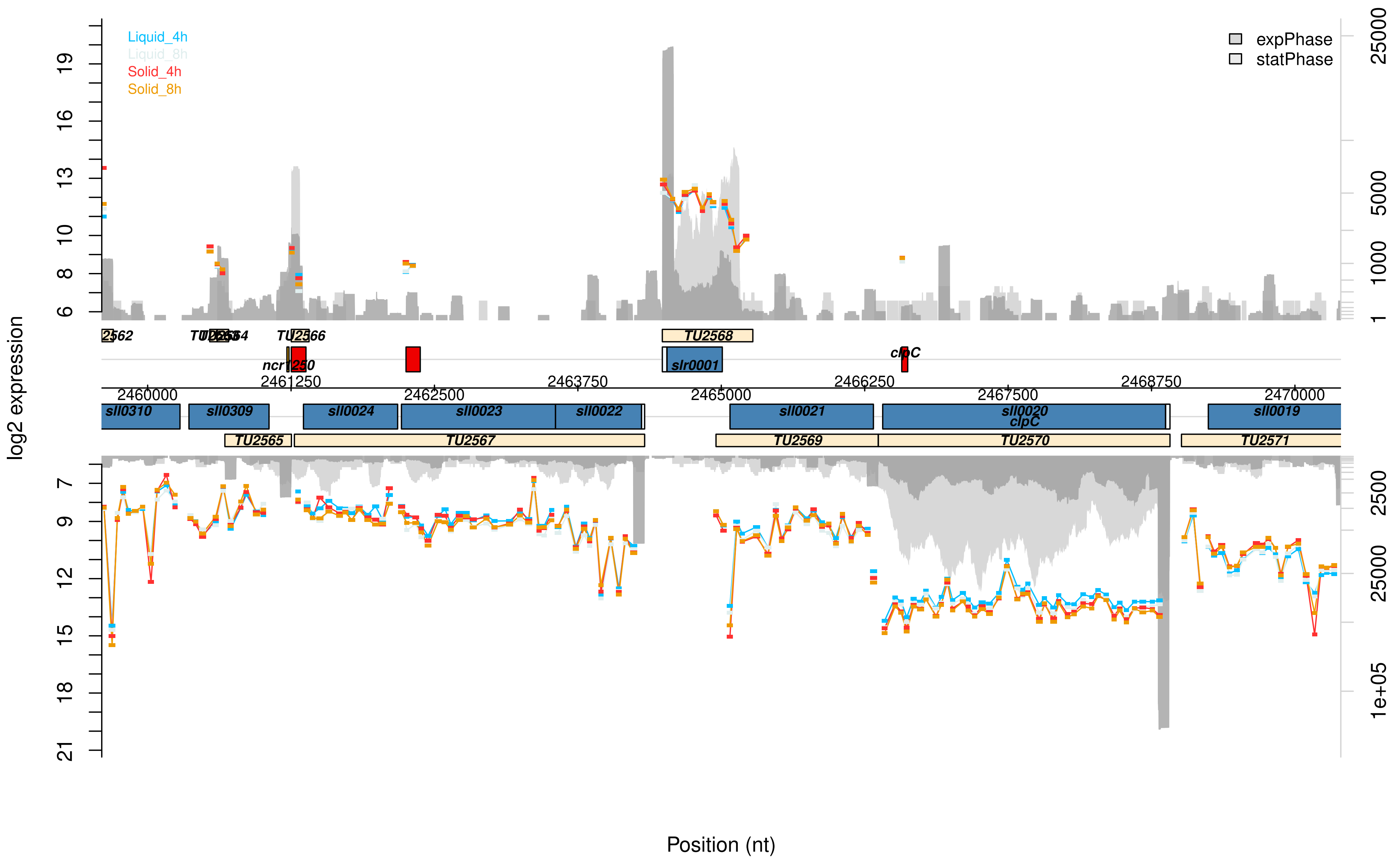

Liquid_4h
Liquid_8h
Solid_4h
Solid_8h
expPhase
statPhase
log2 expression
Position (nt)
2562
TU2564
TU2566
TU2568
ncr1250
slr0001
clpC
2461250
2463750
2466250
2468750
2460000
2462500
2465000
2467500
2470000
sll0310
sll0309
sll0024
sll0023
sll0022
sll0021
sll0020
clpC
sll0019
TU2565
TU2567
TU2569
TU2570
TU2571

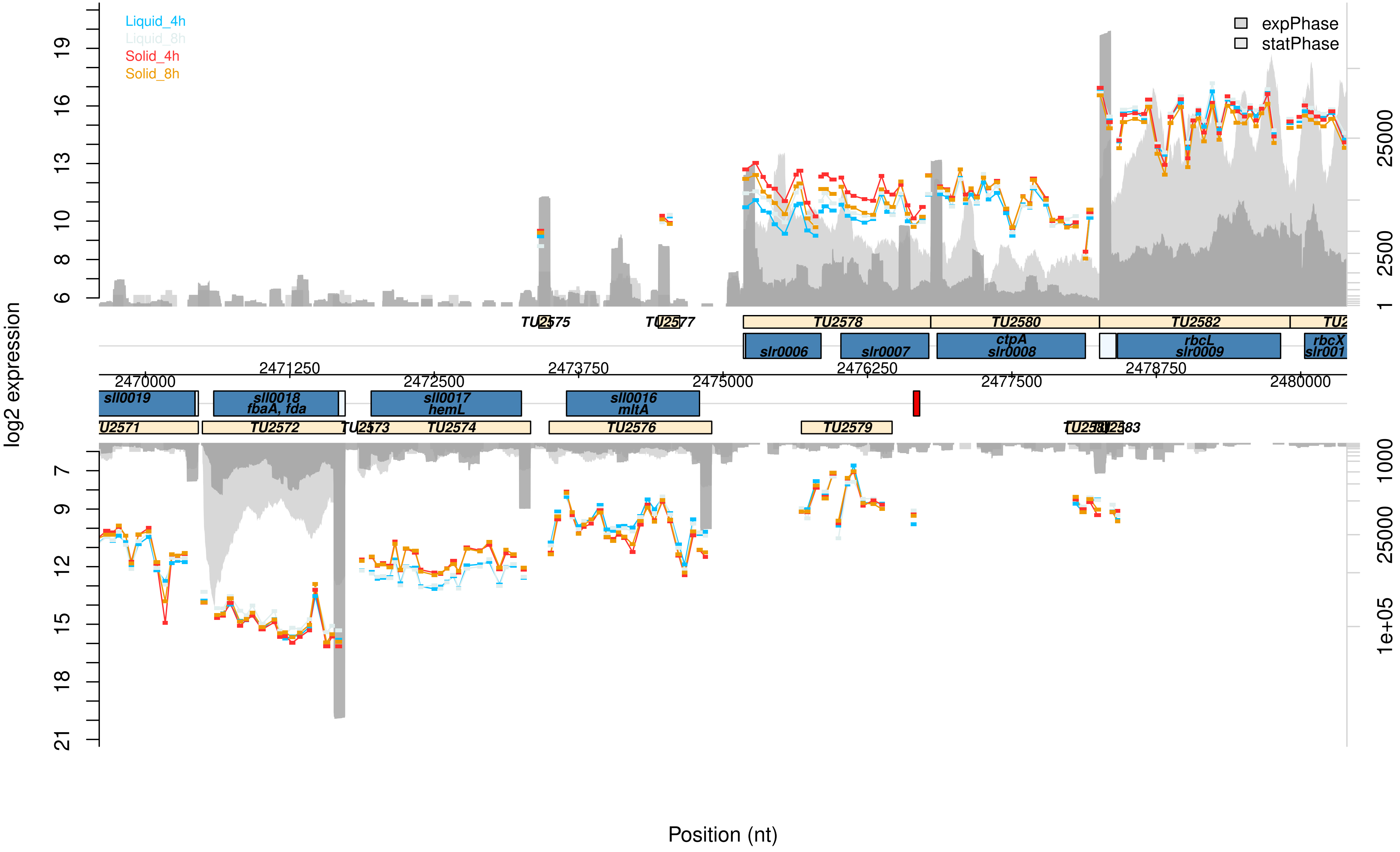

Liquid_4h
Liquid_8h
Solid_4h
Solid_8h
expPhase
statPhase
log2 expression
TU2575
TU2577
TU2578
TU2580
TU2582
slr0006
slr0007
ctpA
slr0008
rbcL
slr0009
rbcX
2470000
2471250
2472500
2473750
2475000
2476250
2477500
2478750
2480000
sll0019
sll0018
fbaA, fda
sll0017
hemL
sll0016
mltA
TU2572
TU2574
TU2576
TU2579
Position (nt)

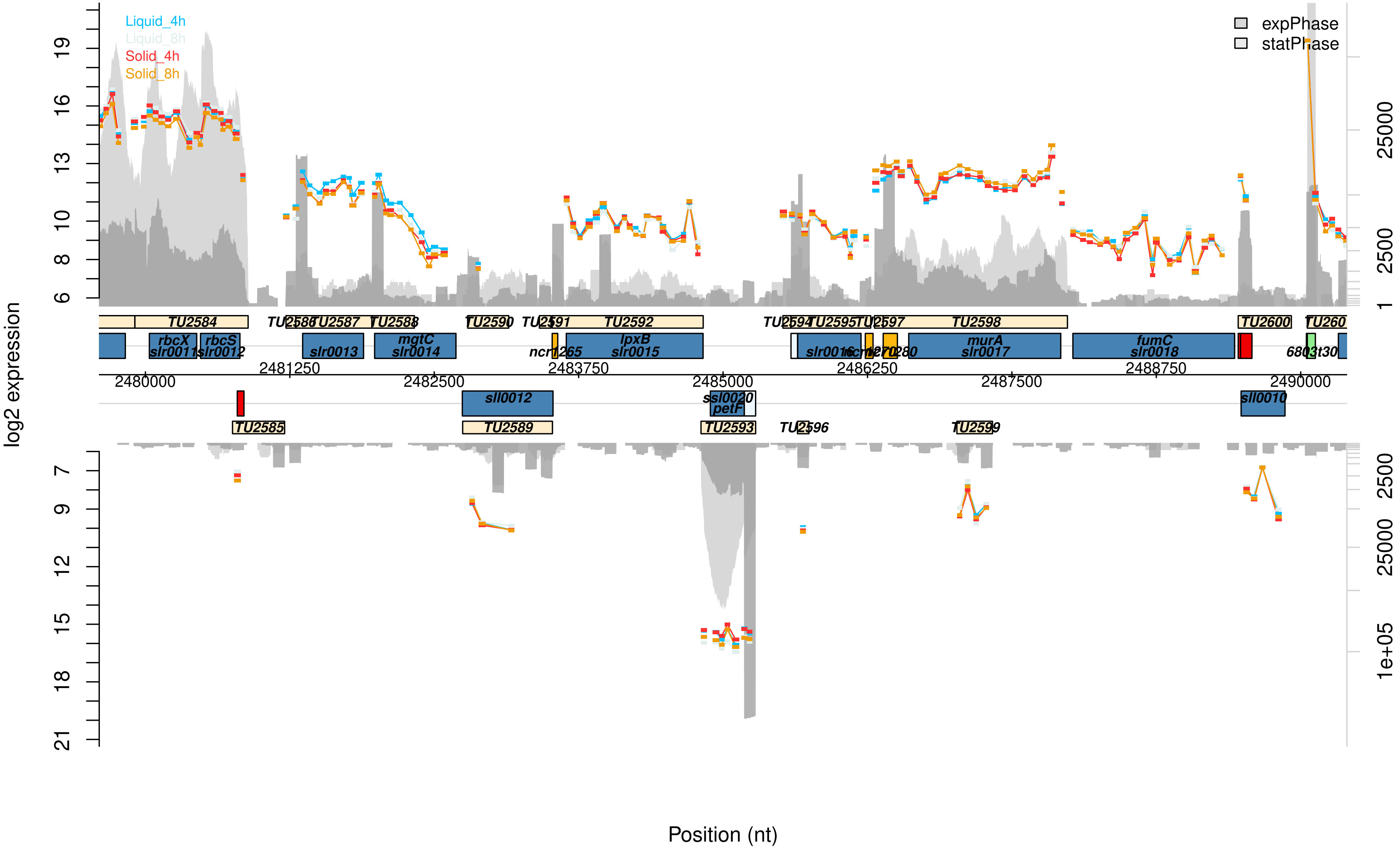

Liquid_4h
Liquid_8h
Solid_4h
Solid_8h
expPhase
statPhase
log2 expression
Position (nt)
TU2584
TU2585
TU2587
TU2588
TU2589
TU2590
TU2592
TU2593
TU2595
TU2597
TU2598
TU2599
TU2600
rbcX
slr0011
rbcS
slr0012
slr0013
mgtC
slr0014
sll0012
lpxB
slr0015
ssl0020
petF
murA
slr0017
fumC
slr0018
sll0010
2480000
2481250
2482500
2483750
2485000
2486250
2487500
2488750
2490000

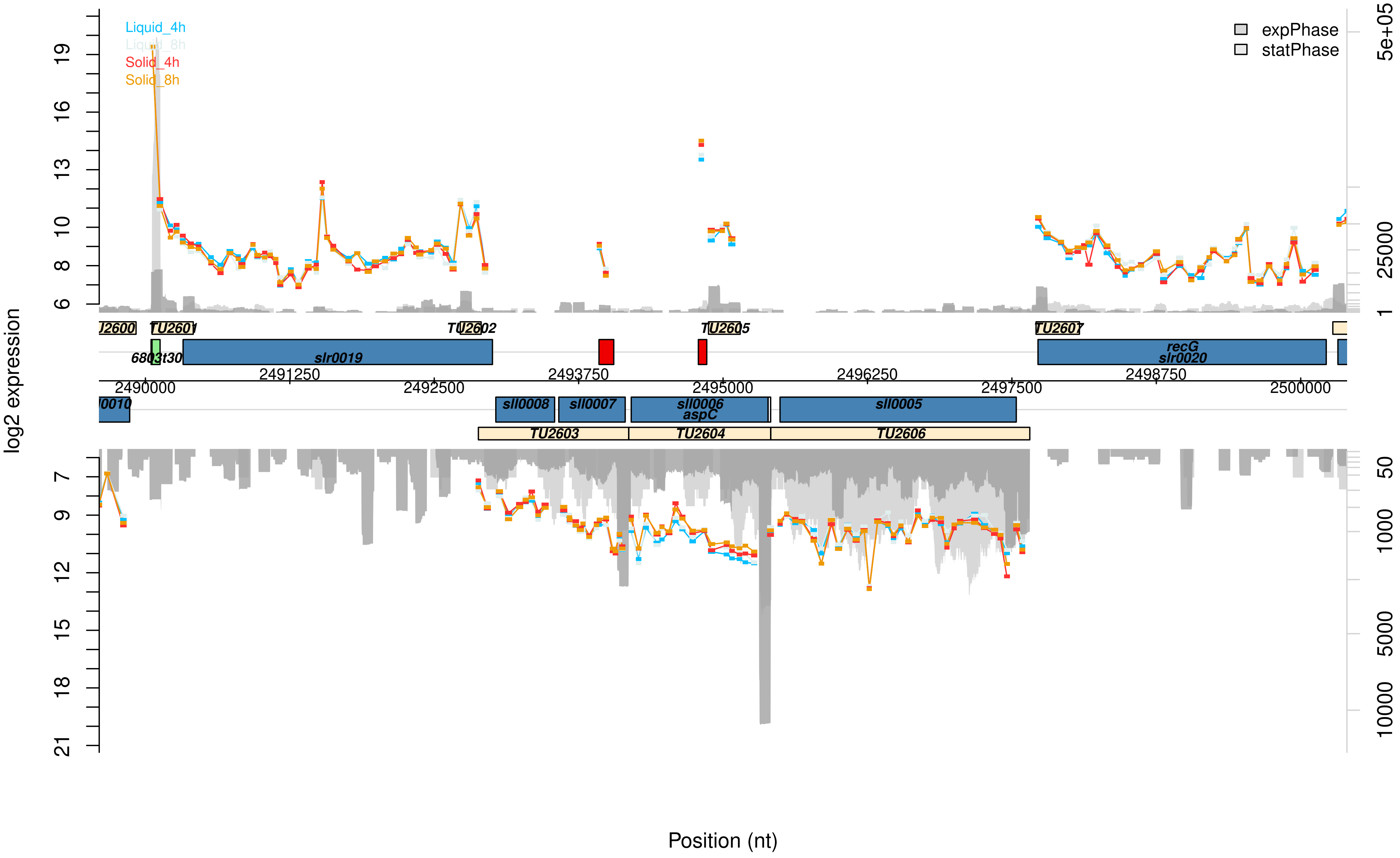

Liquid_4h
Liquid_8h
Solid_4h
Solid_8h
expPhase
statPhase
log2 expression
Position (nt)
TU2600
TU2601
TU2602
TU2605
TU2607
6803t30
slr0019
recG
slr0020
sll0010
sll0008
sll0007
sll0006
aspC
sll0005
TU2603
TU2604
TU2606
2490000
2491250
2492500
2493750
2495000
2496250
2497500
2498750
2500000
5e+05
25000
1
50
1000
5000
10000

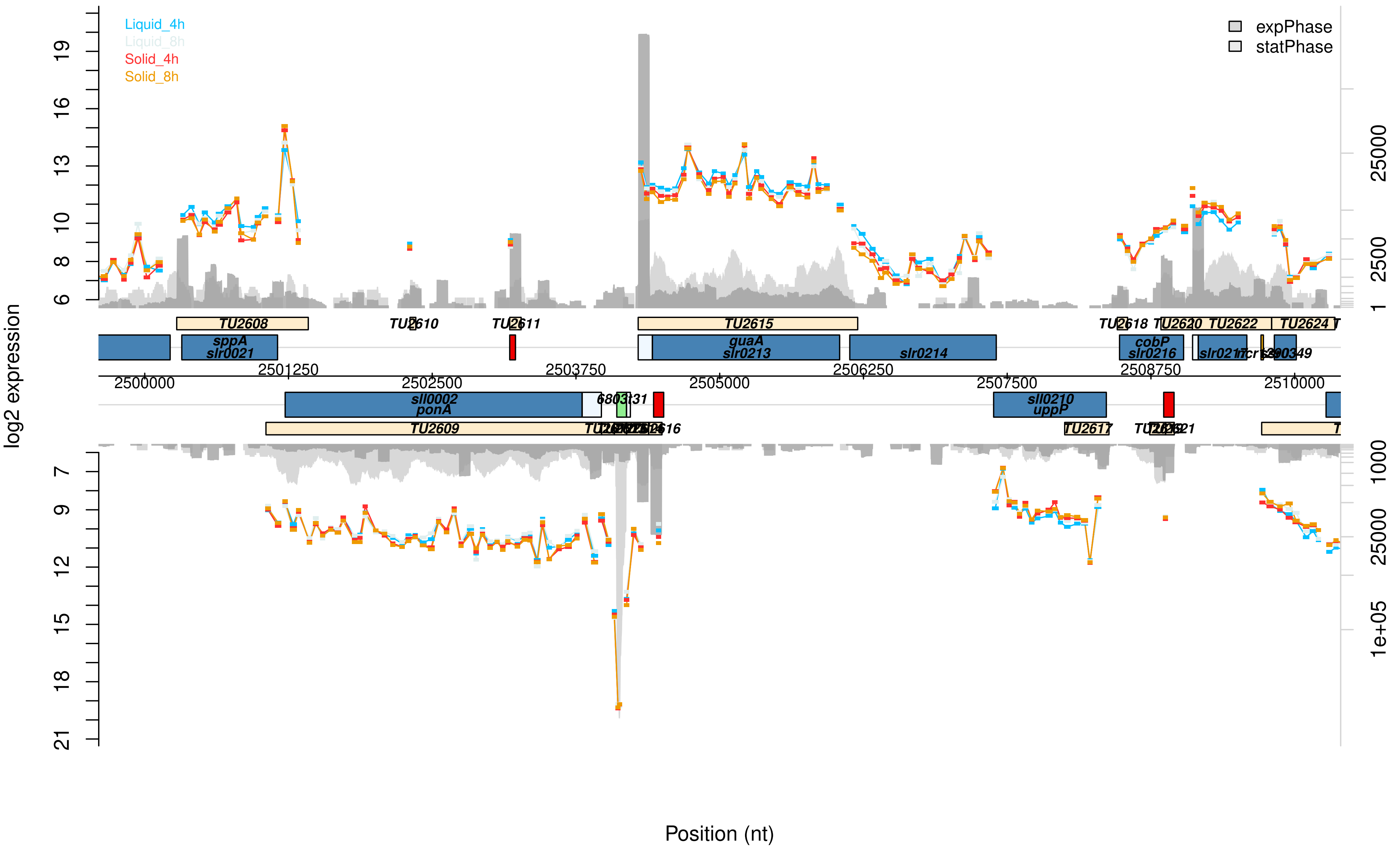

Liquid_4h
Liquid_8h
Solid_4h
Solid_8h
expPhase
statPhase
log2 expression
Position (nt)
TU2608
TU2610
TU2611
TU2615
TU2618
TU2620
TU2622
TU2624
sppA
slr0021
guaA
slr0213
slr0214
cobP
slr0216
sll0002
ponA
sll0210
uppP
TU2609
TU2617
2500000
2501250
2502500
2503750
2505000
2506250
2507500
2508750
2510000

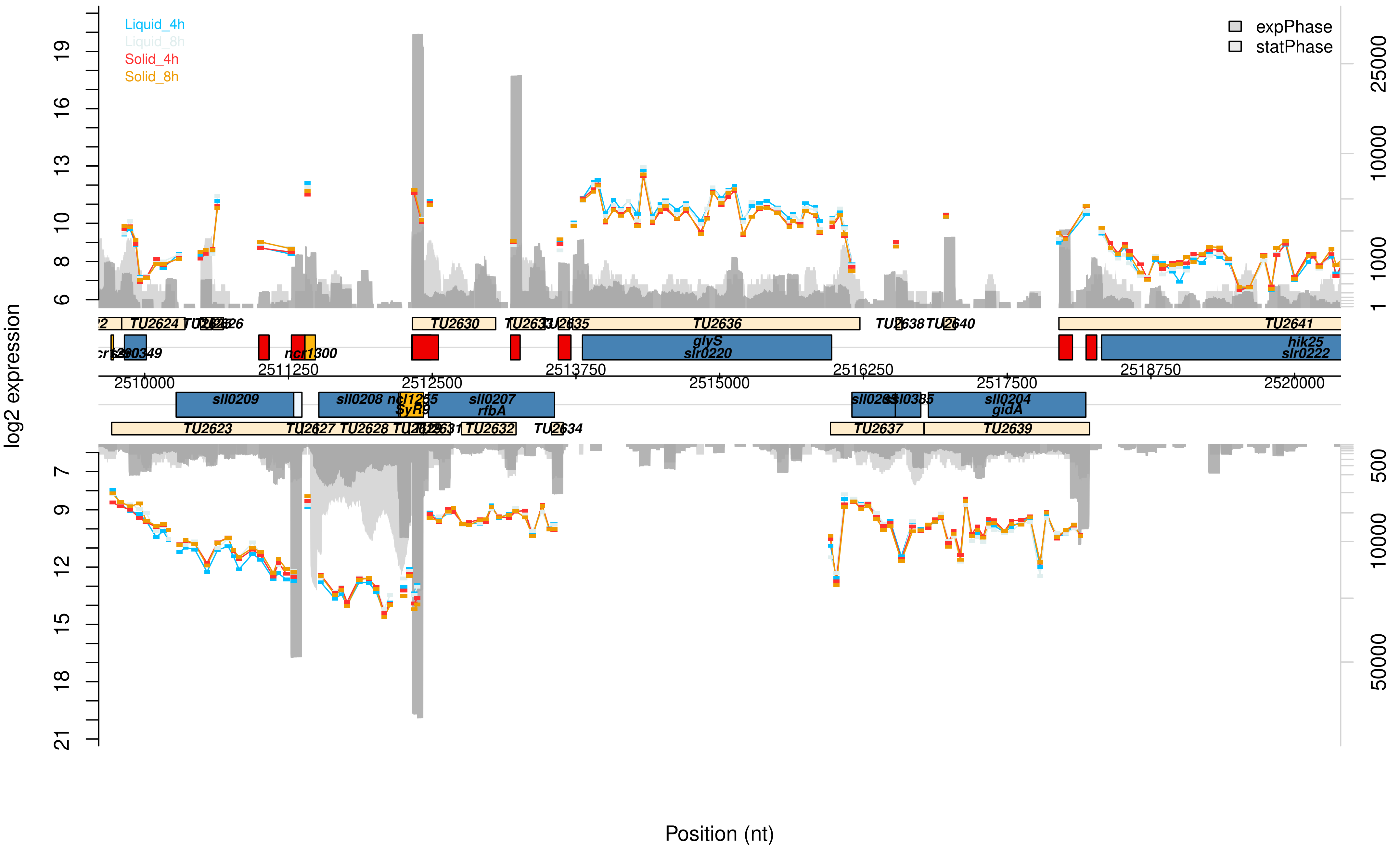

Liquid_4h
Liquid_8h
Solid_4h
Solid_8h
expPhase
statPhase
log2 expression
Position (nt)
TU2624
TU2630
TU2636
TU2641
glyS
slr0220
hik25
slr0222
ncr1300
sll0209
sll0208
sll0207
rfbA
sll0204
gidA
TU2623
TU2628
TU2632
TU2637
TU2639
2510000
2511250
2512500
2513750
2515000
2516250
2517500
2518750
2520000

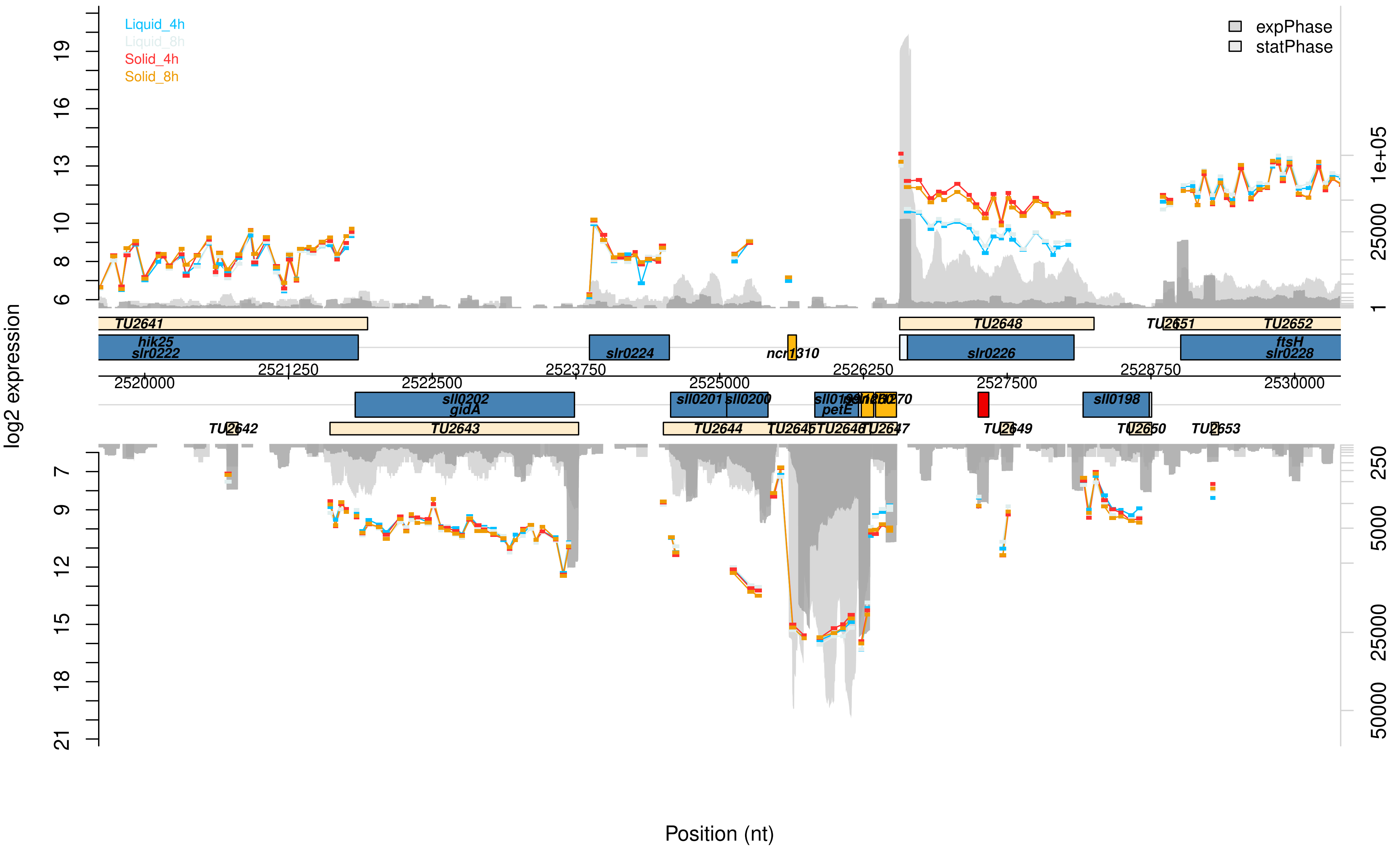

Liquid_4h
Liquid_8h
Solid_4h
Solid_8h
expPhase
statPhase
log2 expression
Position (nt)
TU2641
hik25
slr0222
slr0224
ncr1310
TU2648
slr0226
TU2651
TU2652
ftsH
slr0228
sll0202
gidA
sll0201
sll0200
petE
sll0198
TU2642
TU2643
TU2644
TU2645
TU2646
TU2647
TU2649
TU2650
TU2653

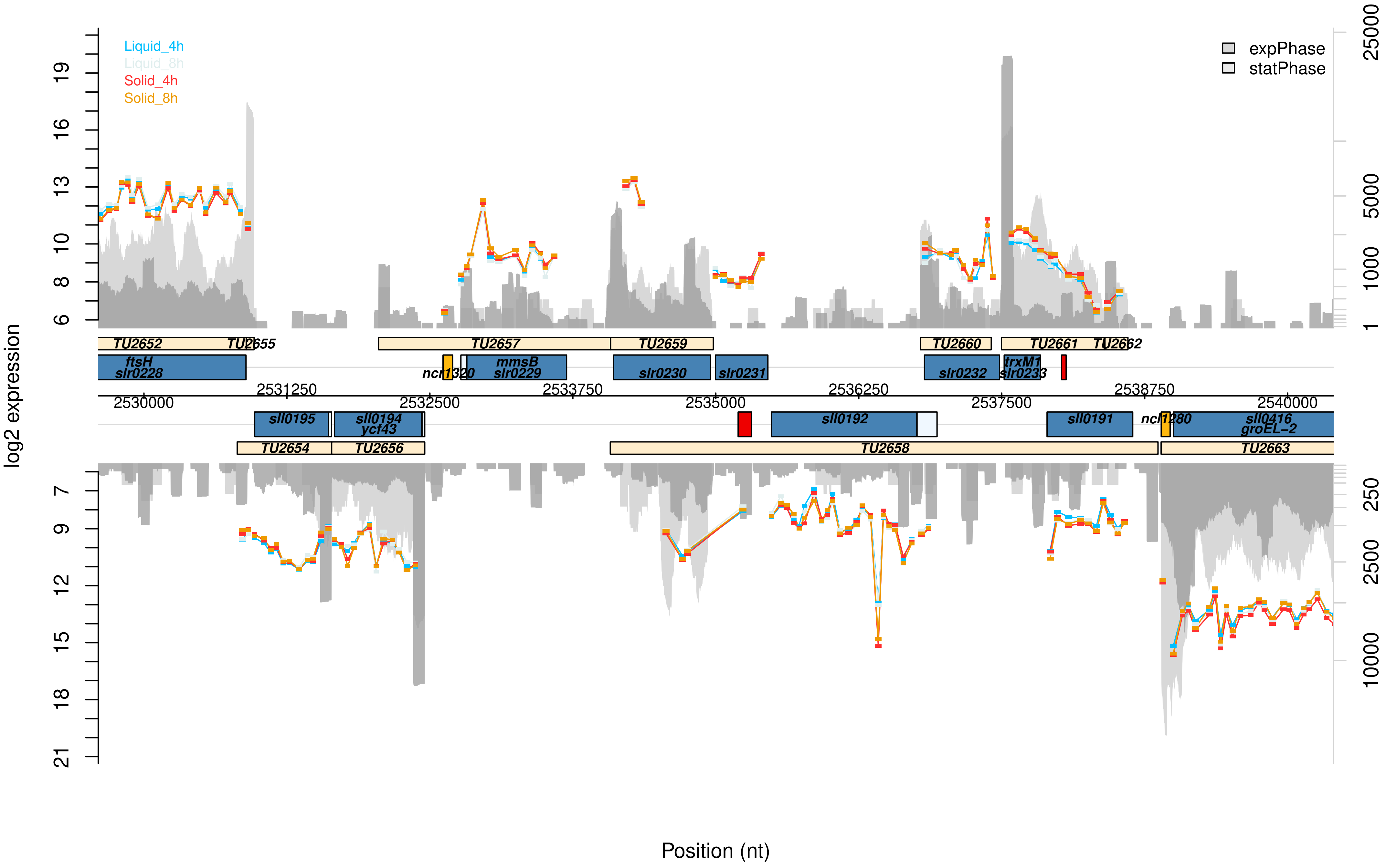

Liquid_4h
Liquid_8h
Solid_4h
Solid_8h
expPhase
statPhase
log2 expression
TU2652
TU2655
TU2657
TU2659
TU2660
TU2661
TU2662
ftsH
slr0228
ncr1320
mmsB
slr0229
slr0230
slr0231
slr0232
trxM1
slr0233
2530000
2531250
2532500
2533750
2535000
2536250
2537500
2538750
2540000
sll0195
sll0194
ycf43
sll0192
sll0191
ncl1280
sll0416
groEL−2
TU2654
TU2656
TU2658
TU2663
Position (nt)

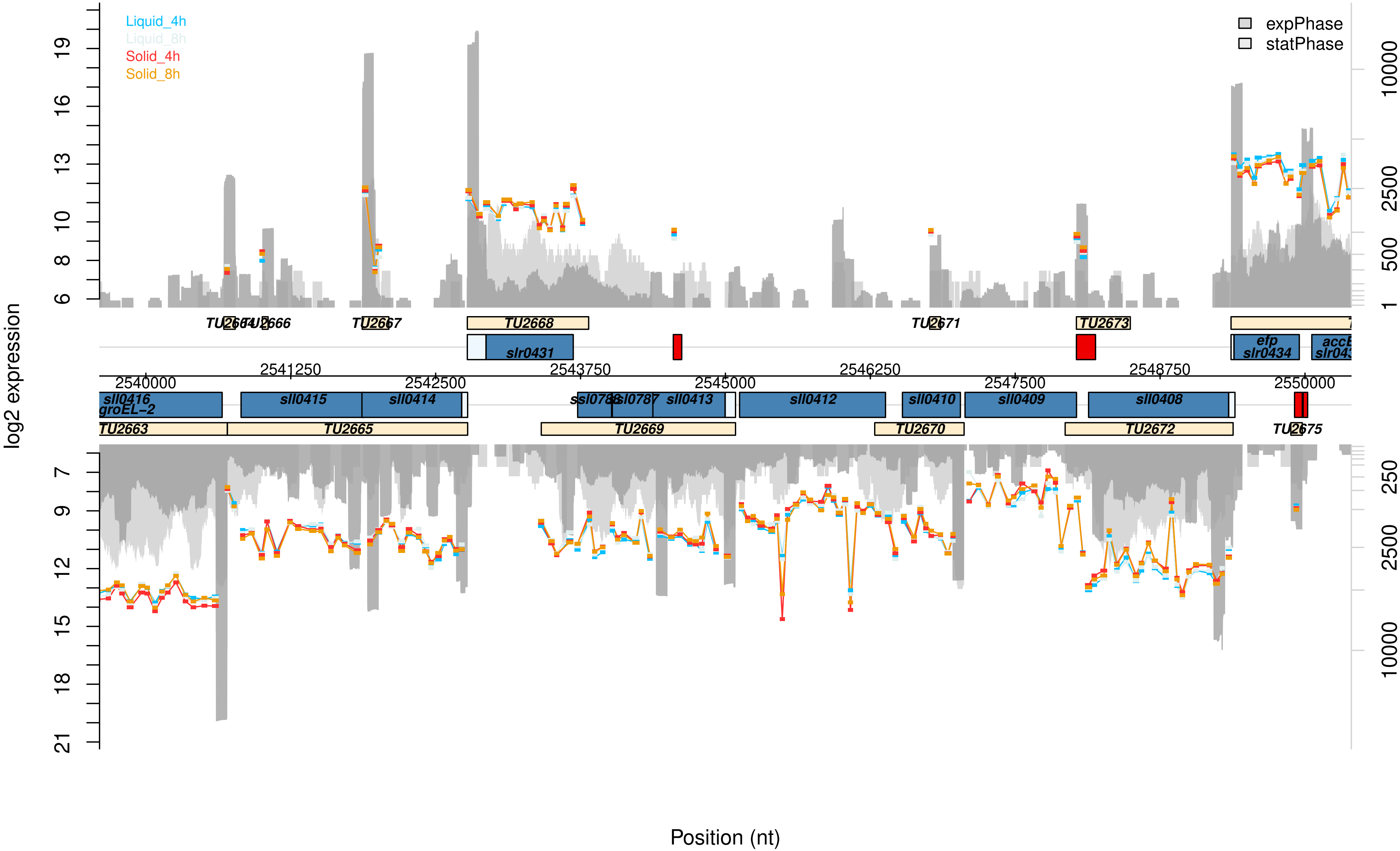

Liquid_4h
Liquid_8h
Solid_4h
Solid_8h
expPhase
statPhase
log2 expression
Position (nt)
TU2664
TU2666
TU2667
TU2668
TU2671
TU2673
slr0431
efp
slr0434
accB
2541250
2543750
2546250
2548750
2540000
2542500
2545000
2547500
2550000
sll0416
groEL-2
sll0415
sll0414
ssl0788
sll0787
sll0413
sll0412
sll0410
sll0409
sll0408
TU2663
TU2665
TU2669
TU2670
TU2672
TU2675

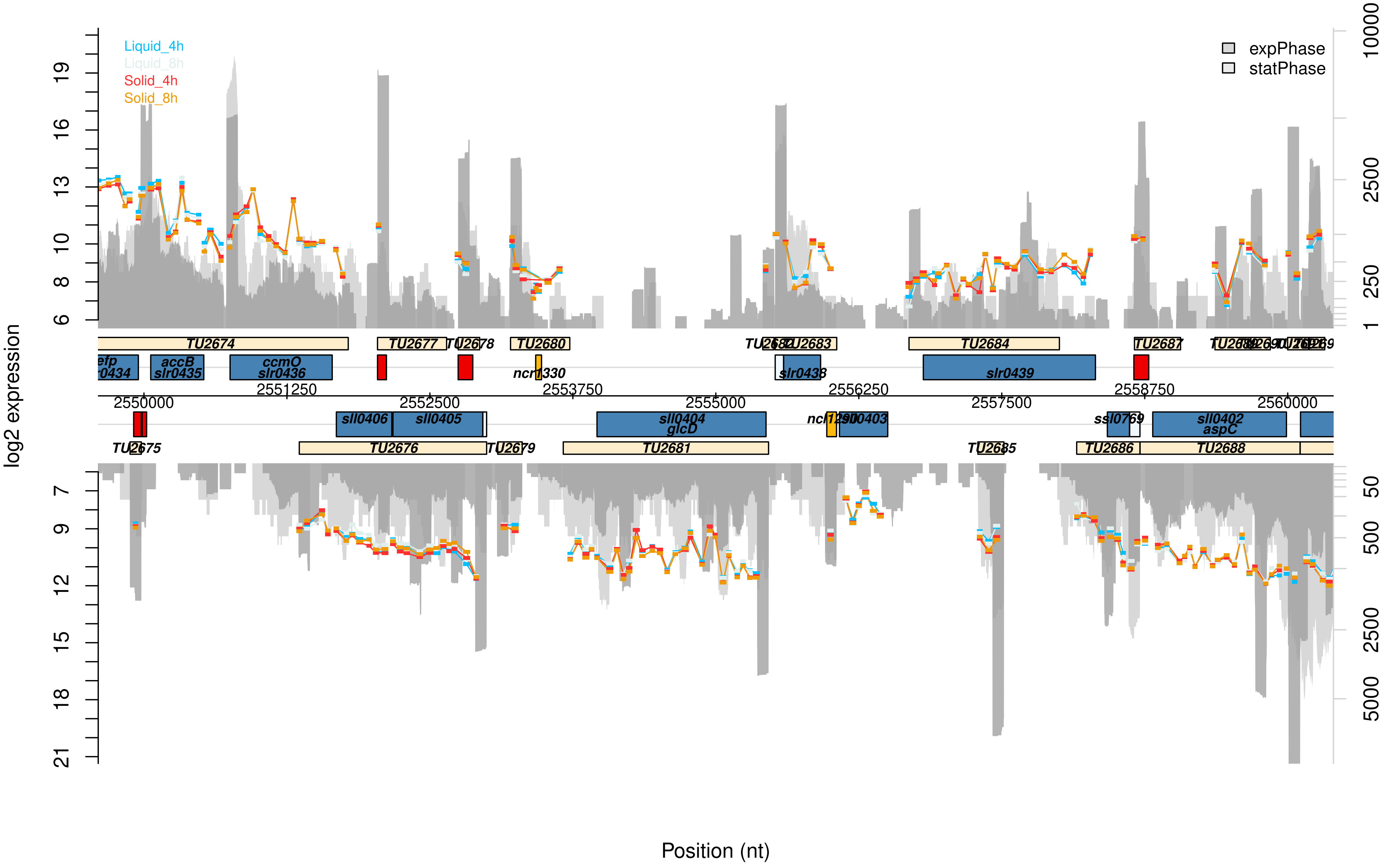

Liquid_4h
Liquid_8h
Solid_4h
Solid_8h
expPhase
statPhase
log2 expression
Position (nt)
TU2674
TU2677
TU2678
TU2680
TU2683
TU2684
TU2687
TU2675
TU2676
TU2679
TU2681
TU2685
TU2686
TU2688
accB
slr0435
ccmO
slr0436
ncr1330
slr0438
slr0439
sll0406
sll0405
sll0404
glcD
ssl0769
sll0402
aspC
2550000
2551250
2552500
2553750
2555000
2556250
2557500
2558750
2560000

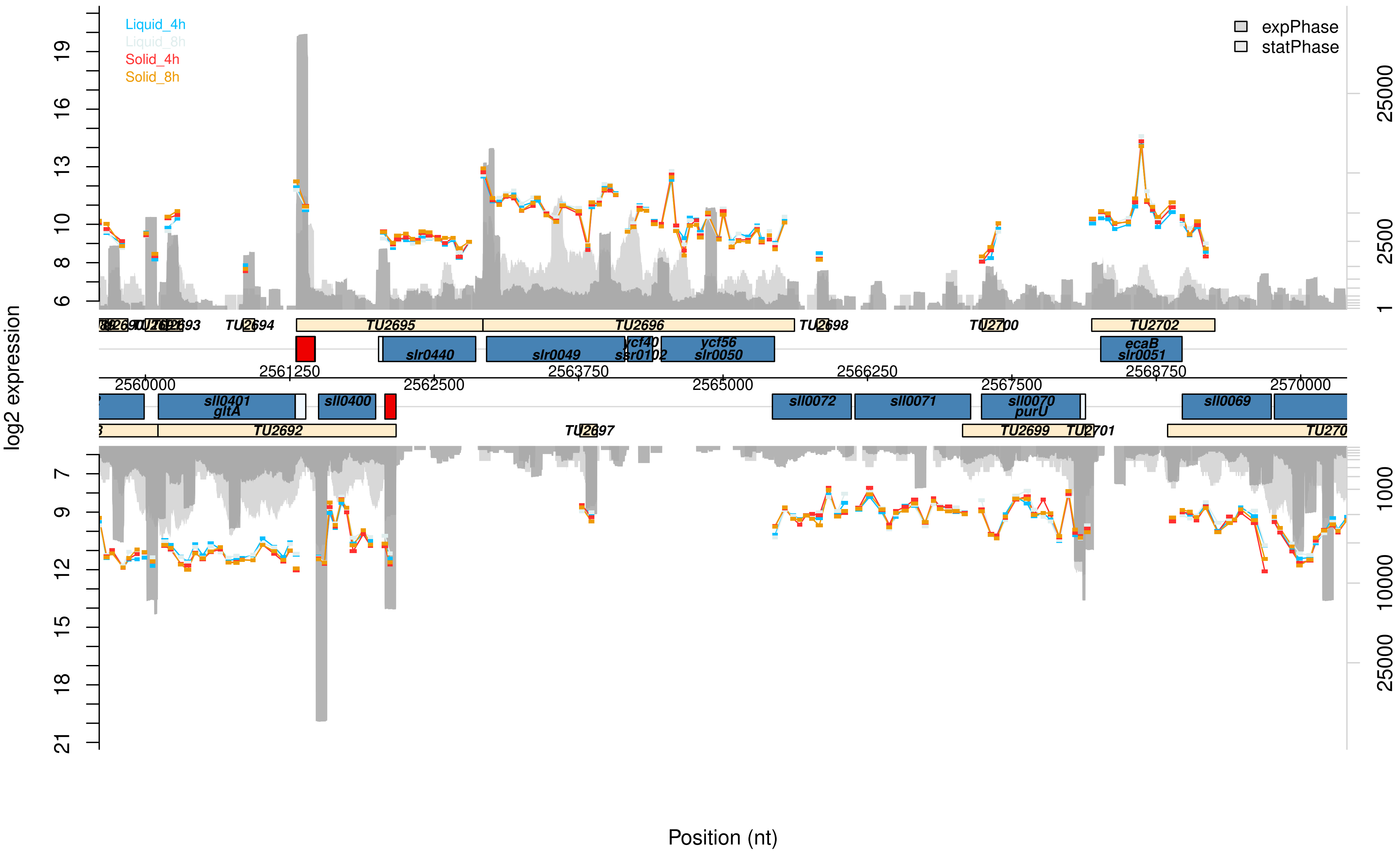

Liquid_4h
Liquid_8h
Solid_4h
Solid_8h
expPhase
statPhase
log2 expression
Position (nt)
TU2694
TU2695
TU2696
TU2698
TU2700
TU2702
slr0440
slr0049
ycf40
ssr0102
ycf56
slr0050
ecaB
slr0051
2561250
2563750
2566250
2568750
2560000
2562500
2565000
2567500
2570000
sll0401
gltA
sll0400
sll0072
sll0071
sll0070
purU
sll0069
TU2692
TU2697
TU2699
TU2701

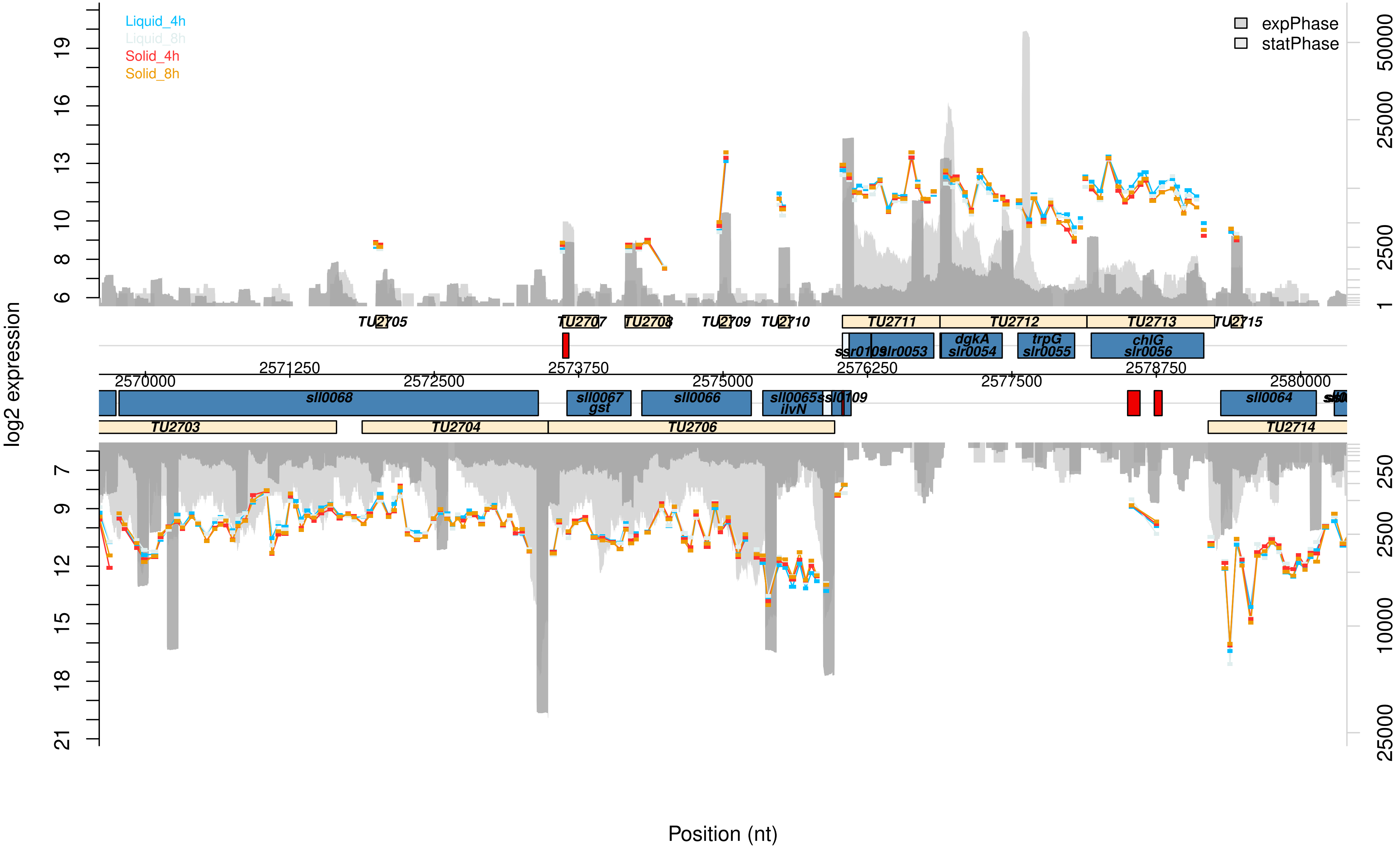

Liquid_4h
Liquid_8h
Solid_4h
Solid_8h
expPhase
statPhase
log2 expression
TU2705
TU2707
TU2708
TU2709
TU2710
TU2711
TU2712
TU2713
TU2715
ssr0109
slr0053
dgkA
slr0054
trpG
slr0055
chlG
slr0056
2570000
2571250
2572500
2573750
2575000
2576250
2577500
2578750
2580000
sll0068
sll0067
gst
sll0066
sll0065
ilvN
ssl0109
sll0064
TU2703
TU2704
TU2706
TU2714
Position (nt)

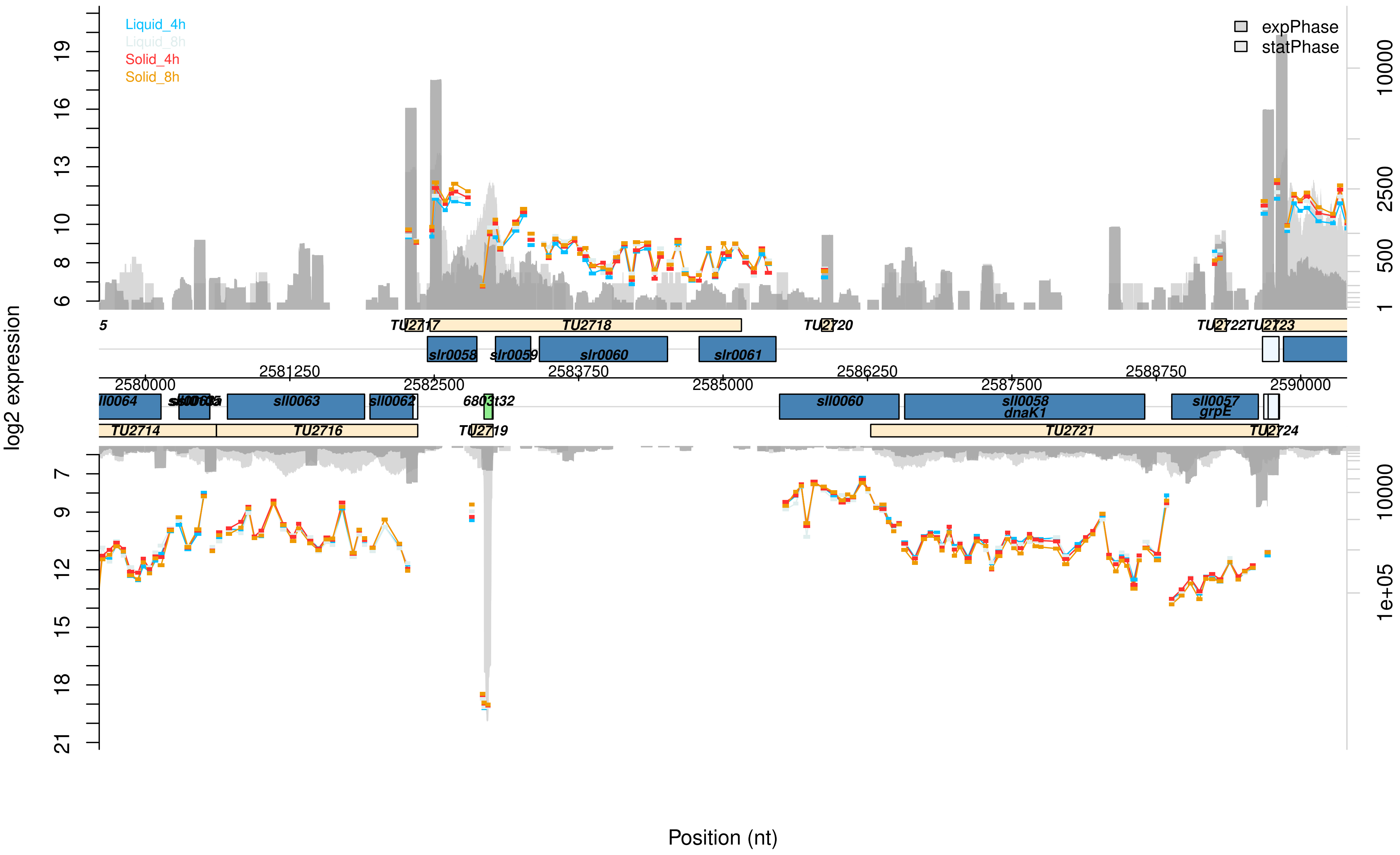

Liquid_4h
Liquid_8h
Solid_4h
Solid_8h
expPhase
statPhase
log2 expression
TU2717
TU2718
TU2720
TU2722TU2723
slr0058
slr0059
slr0060
slr0061
2580000
2581250
2582500
2583750
2585000
2586250
2587500
2588750
2590000
sll0064
sll0063
sll0062
6803t32
sll0060
sll0058
dnaK1
sll0057
grpE
TU2714
TU2716
TU2719
TU2721
TU2724
Position (nt)

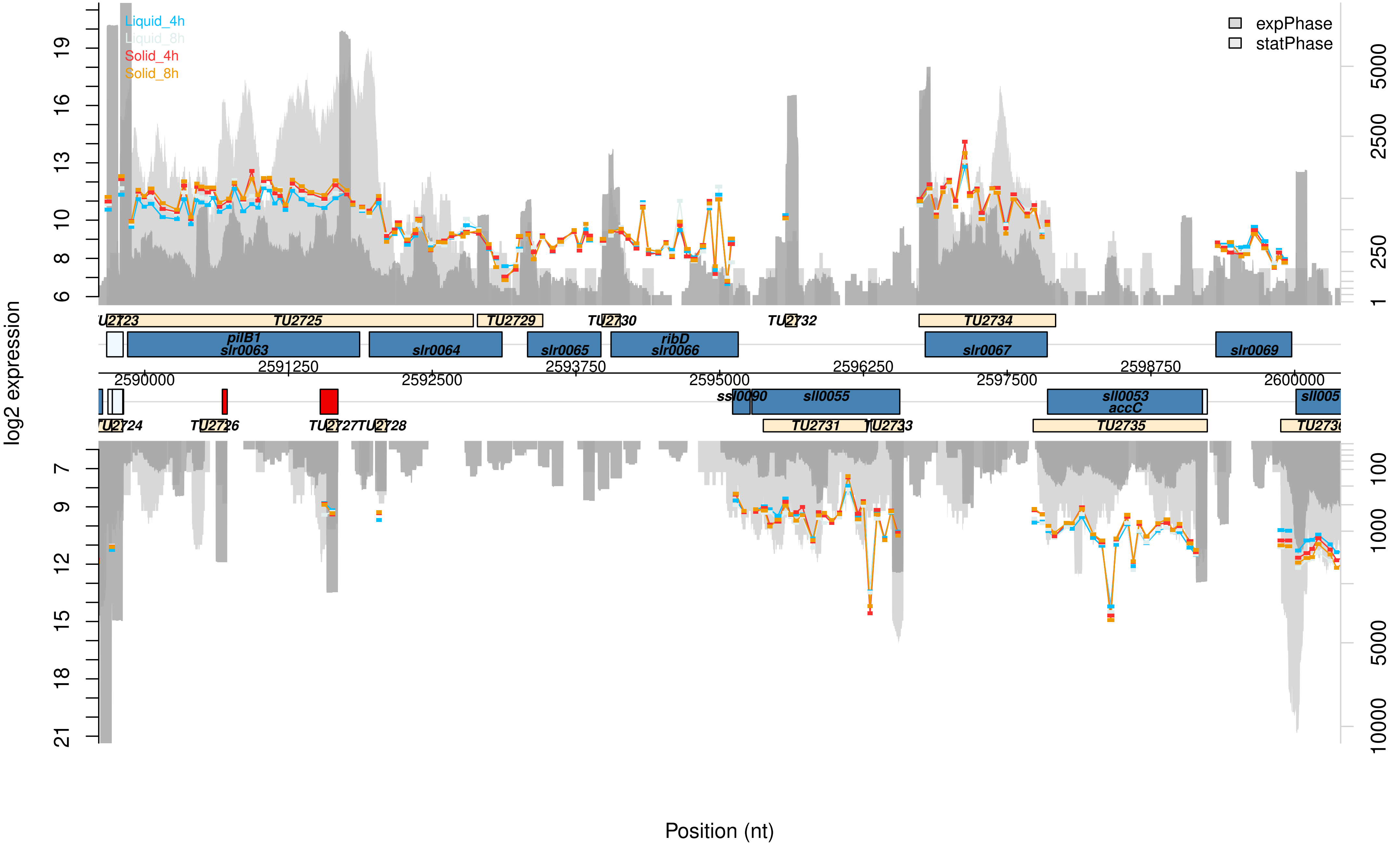

Liquid_4h
Liquid_8h
Solid_4h
Solid_8h
expPhase
statPhase
log2 expression
Position (nt)
TU2723
TU2725
TU2729
TU2730
TU2732
TU2734
pilB1
slr0063
slr0064
slr0065
ribD
slr0066
slr0067
slr0069
2590000
2591250
2592500
2593750
2595000
2596250
2597500
2598750
2600000
ssl0090
sll0055
sll0053
accC
TU2724
TU2726
TU2727
TU2728
TU2731
TU2733
TU2735

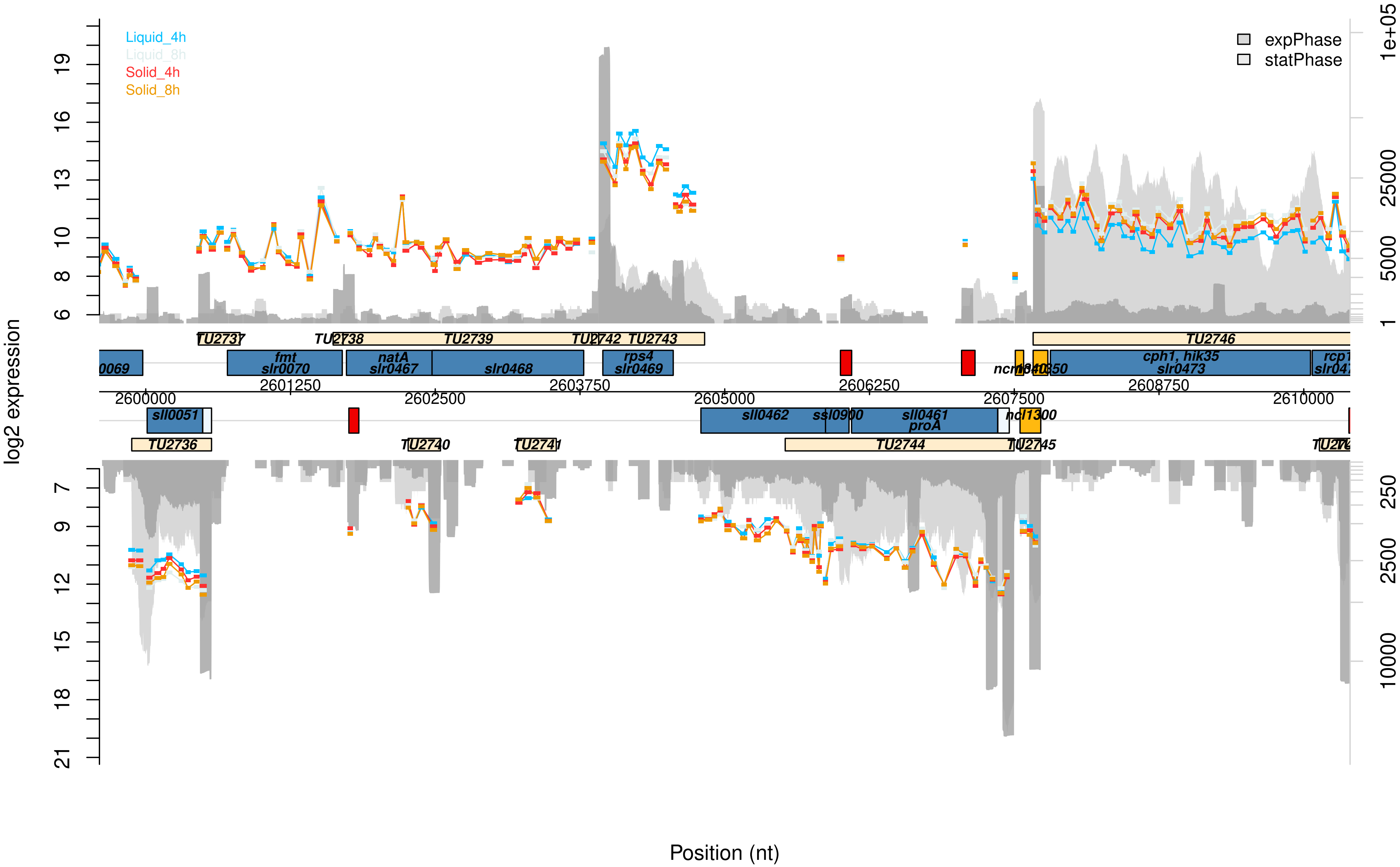

Liquid_4h
Liquid_8h
Solid_4h
Solid_8h
expPhase
statPhase
log2 expression
Position (nt)
TU2737
TU2738
TU2739
TU2742
TU2743
TU2746
fmt
slr0070
natA
slr0467
slr0468
rps4
slr0469
cph1, hik35
slr0473
sll0051
sll0462
ssl0900
sll0461
proA
TU2736
TU2740
TU2741
TU2744
TU2745
2600000
2601250
2602500
2603750
2605000
2606250
2607500
2608750
2610000

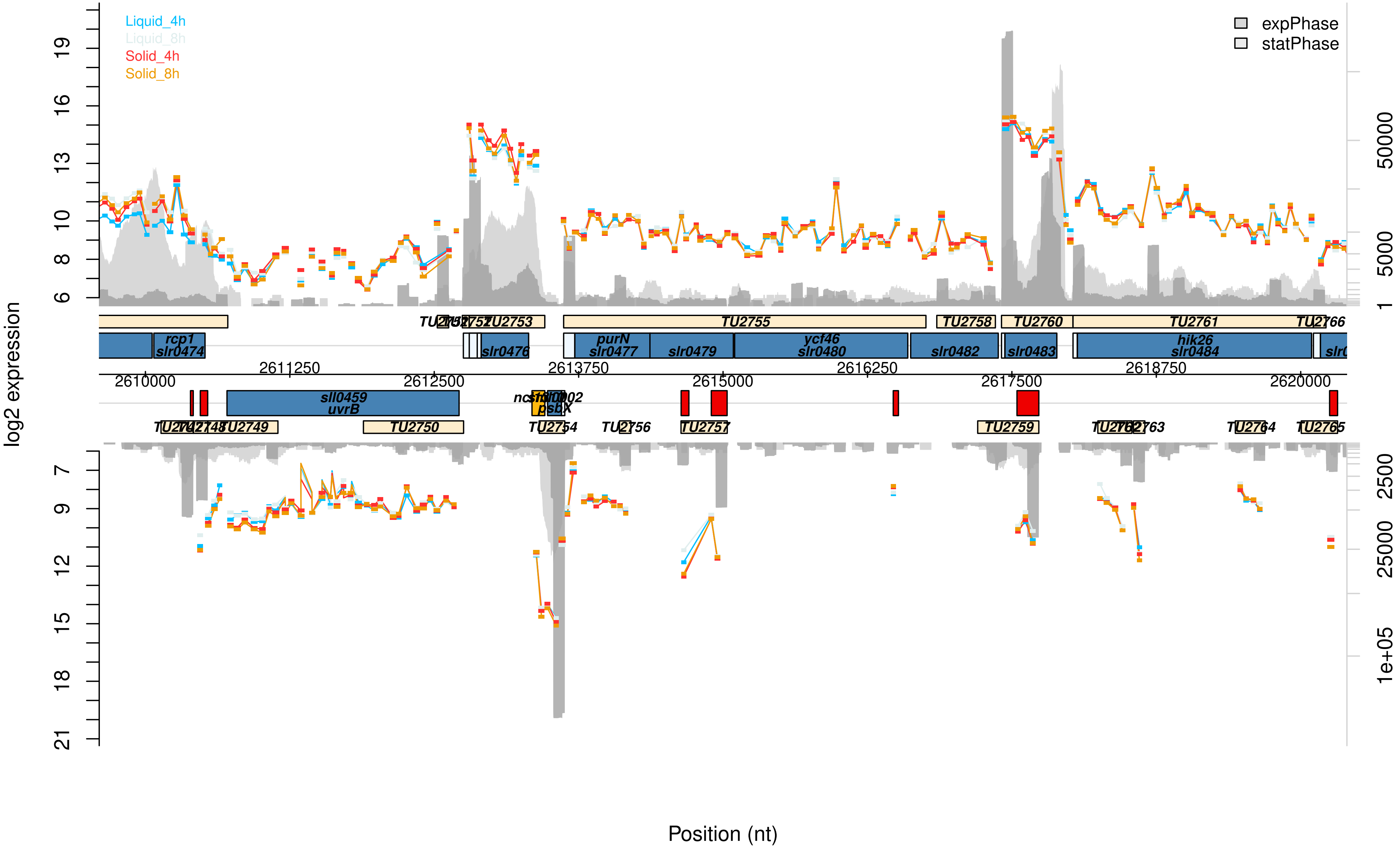
Liquid_4h
Liquid_8h
Solid_4h
Solid_8h
expPhase
statPhase
log2 expression
Position (nt)
2610000
2611250
2612500
2613750
2615000
2616250
2617500
2618750
2620000
rcp1
slr0474
slr0476
purN
slr0477
slr0479
ycf46
slr0480
slr0482
slr0483
hik26
slr0484
sll0459
uvrB
psbX
TU2753
TU2755
TU2758
TU2760
TU2761
TU2766
TU2749
TU2750
TU2754
TU2756
TU2757
TU2759
TU2763
TU2764
TU2765
50000
5000
1
2500
25000
1e+05

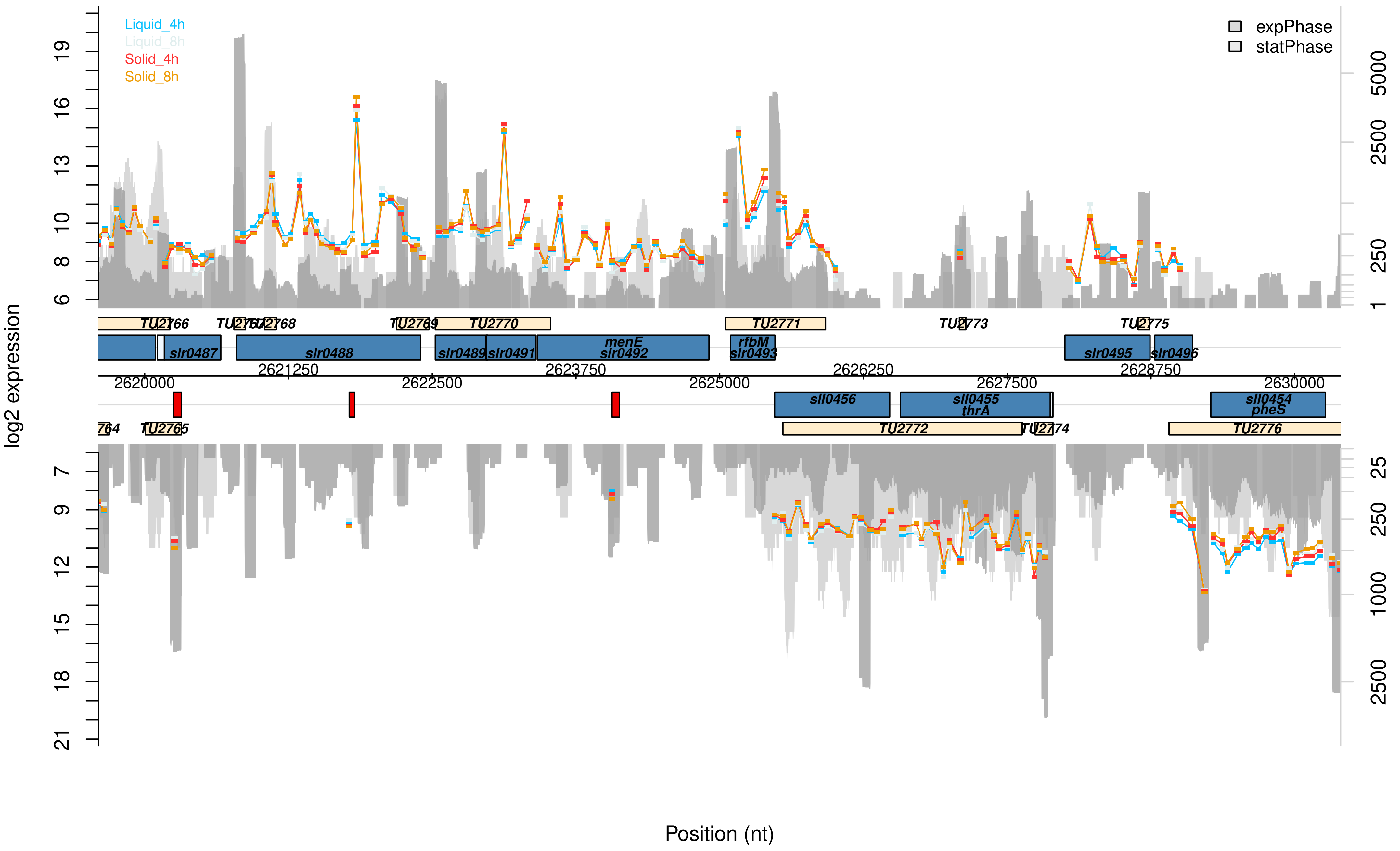

Liquid_4h
Liquid_8h
Solid_4h
Solid_8h
expPhase
statPhase
log2 expression
Position (nt)
TU2766
TU2767
TU2768
TU2769
TU2770
TU2771
TU2773
TU2775
slr0487
slr0488
slr0489
slr0491
menE
slr0492
rfbM
slr0493
slr0495
slr0496
2620000
2621250
2622500
2623750
2625000
2626250
2627500
2628750
2630000
sll0456
sll0455
thrA
sll0454
pheS
TU2765
TU2772
TU2774
TU2776

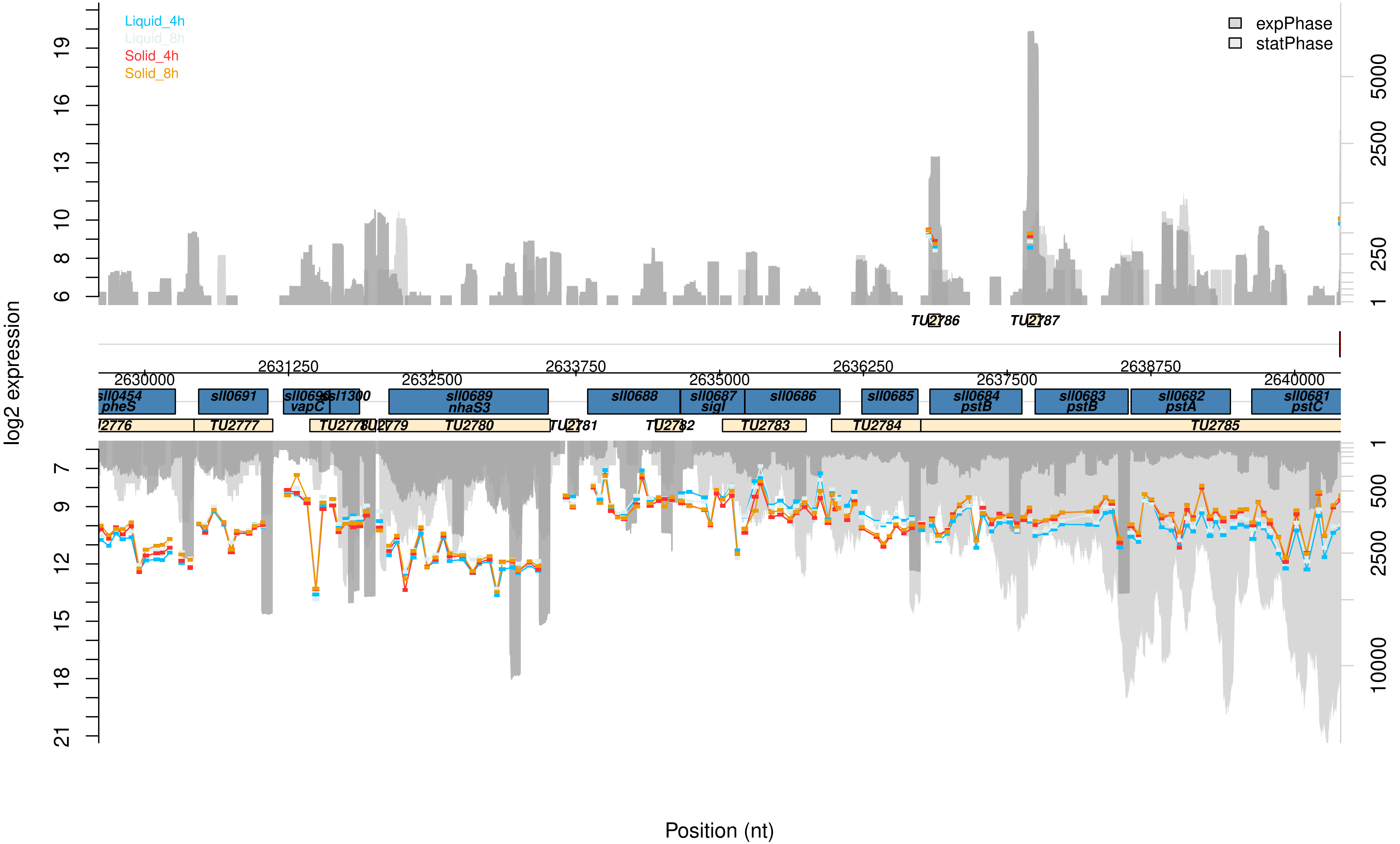

Liquid_4h
Liquid_8h
Solid_4h
Solid_8h
expPhase
statPhase
log2 expression
Position (nt)
2630000
2631250
2632500
2633750
2635000
2636250
2637500
2638750
2640000
sll0454 pheS
sll0691
sll0690 vapC
ssl1300
sll0689 nhaS3
sll0688
sll0687 sigI
sll0686
sll0685
sll0684 pstB
sll0683 pstB
sll0682 pstA
sll0681 pstC
TU2777
TU2780
TU2783
TU2784
TU2785
TU2786
TU2787

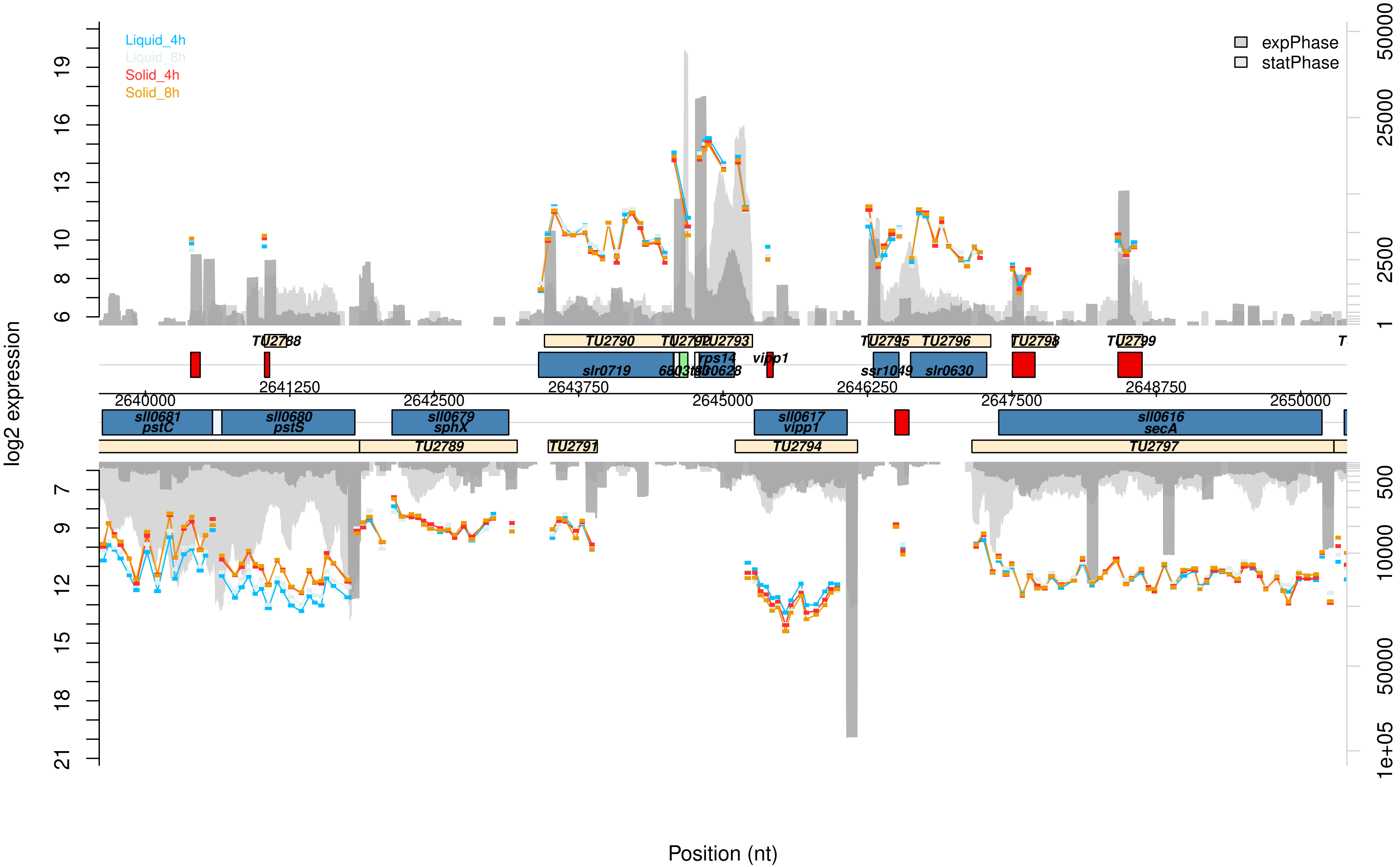

log2 expression
Position (nt)
Liquid_4h
Liquid_8h
Solid_4h
Solid_8h
expPhase
statPhase
TU2788
TU2790
TU2792
TU2793
TU2795
TU2796
TU2798
TU2799
slr0719
rps14
vipp1
ssr1049
slr0630
2641250
2643750
2646250
2648750
2640000
2642500
2645000
2647500
2650000
sll0681
pstC
sll0680
pstS
sll0679
sphX
sll0617
vipp1
sll0616
secA
TU2789
TU2791
TU2794
TU2797

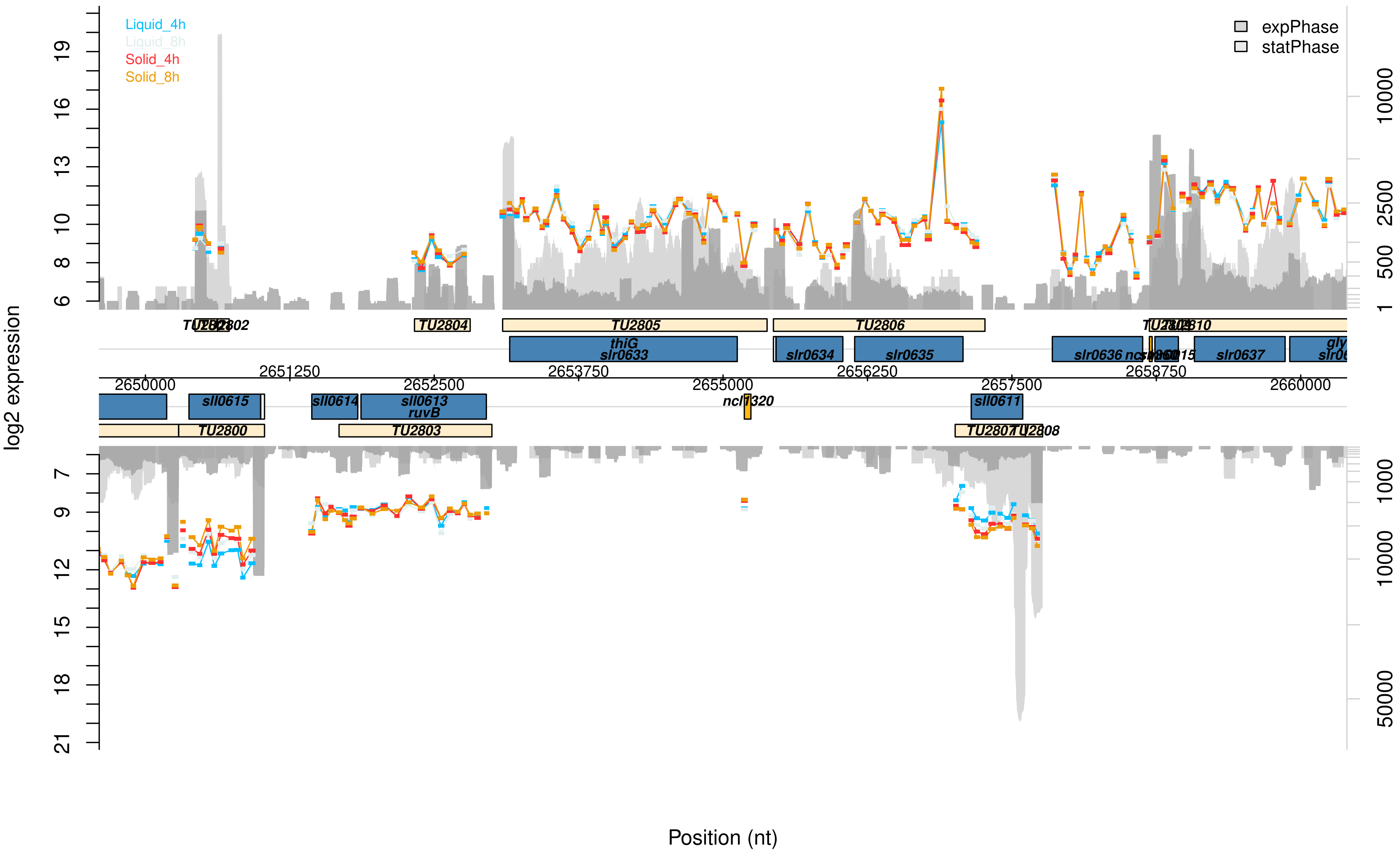

Liquid_4h
Liquid_8h
Solid_4h
Solid_8h
expPhase
statPhase
log2 expression
Position (nt)
TU2805
TU2806
TU2804
TU2810
thiG
slr0633
slr0634
slr0635
slr0636
slr0637
2650000
2651250
2652500
2653750
2655000
2656250
2657500
2658750
2660000
sll0615
sll0614
sll0613
ruvB
ncl1320
sll0611
TU2800
TU2803
TU2807
TU2808

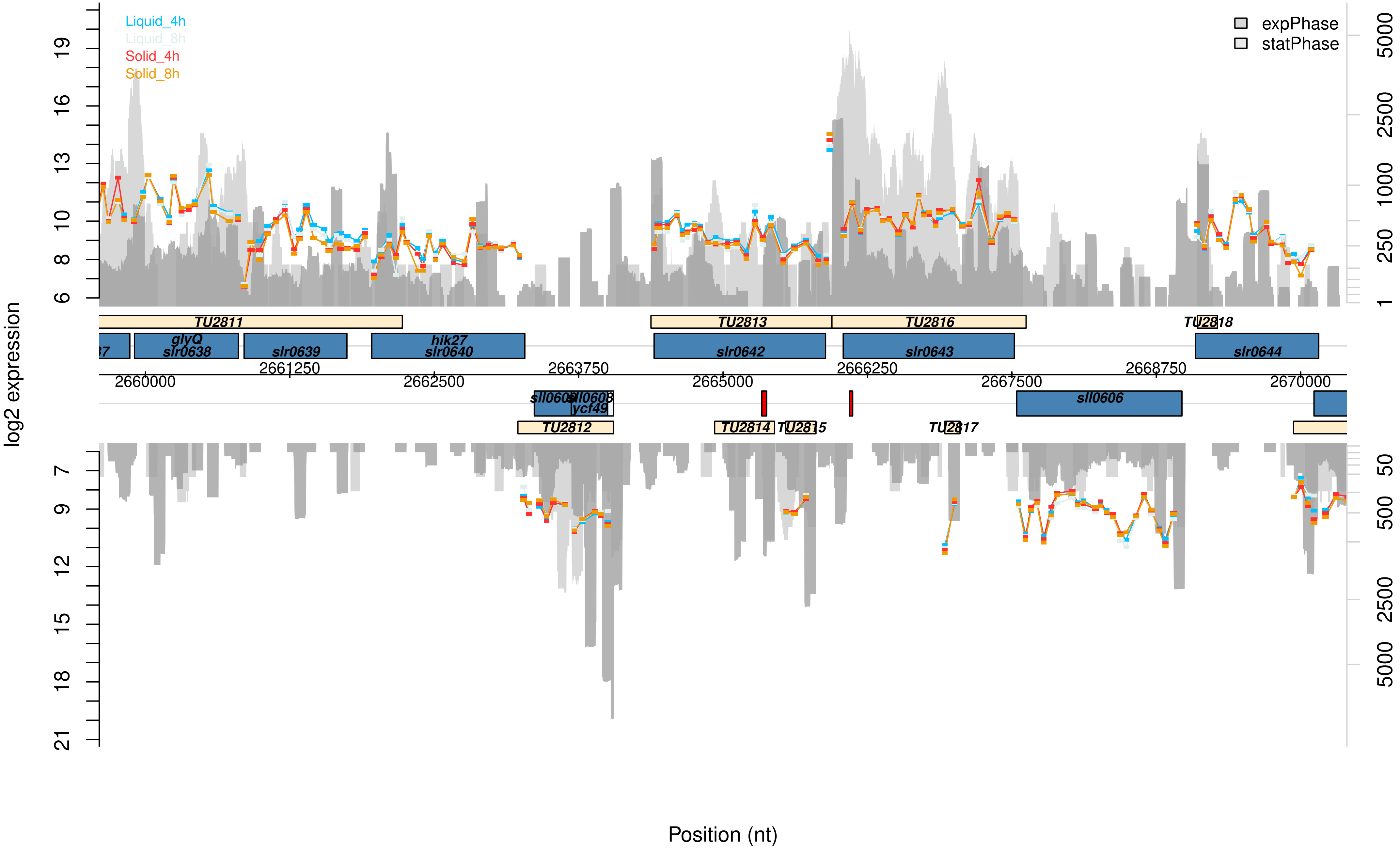
Liquid_4h
Liquid_8h
Solid_4h
Solid_8h
expPhase
statPhase
log2 expression
TU2811
TU2813
TU2816
TU2818
glyQ
slr0638
slr0639
hik27
slr0640
slr0642
slr0643
slr0644
sll0605
sll0608
ycf49
sll0606
TU2812
TU2814
TU2815
TU2817
Position (nt)

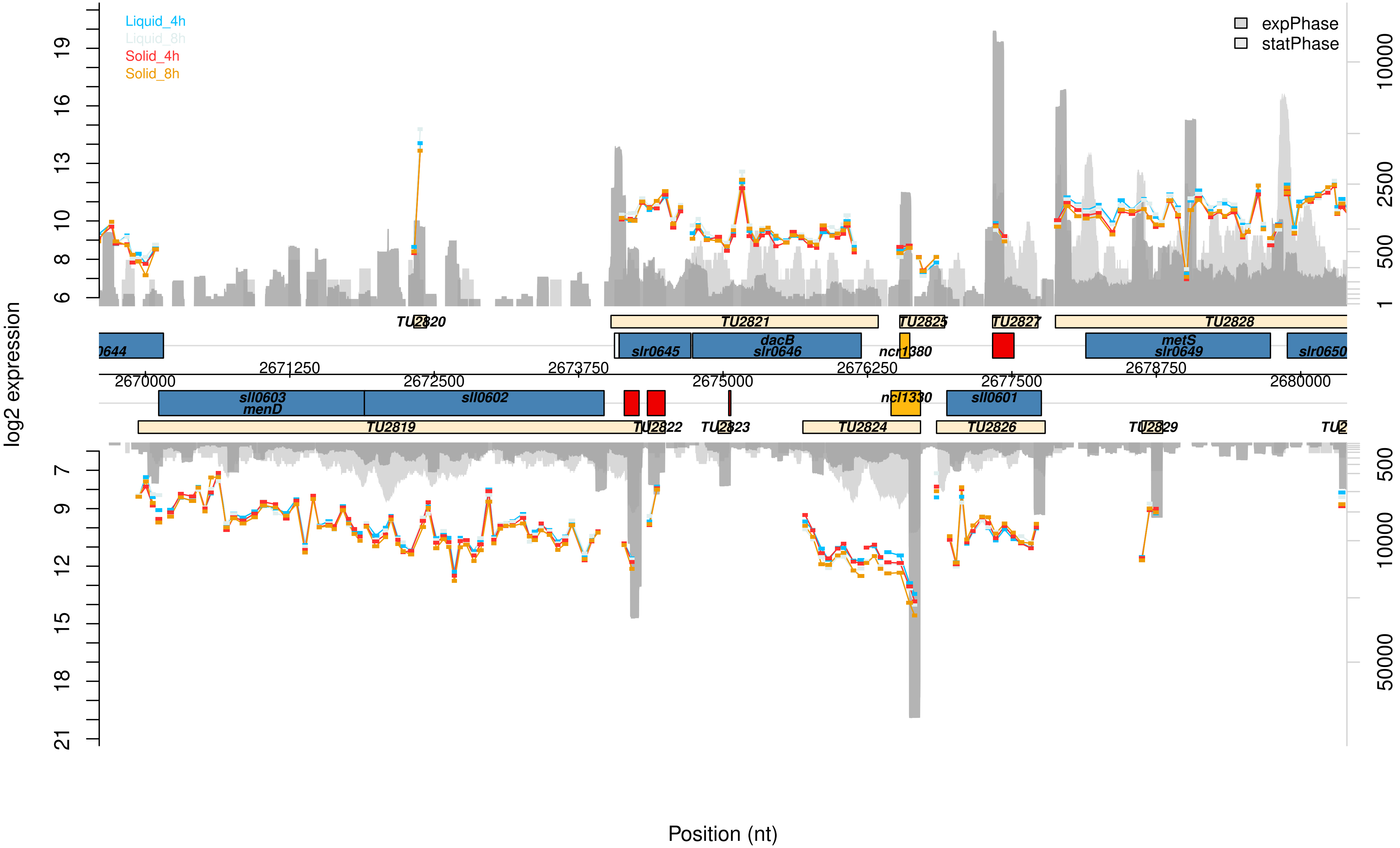

Liquid_4h
Liquid_8h
Solid_4h
Solid_8h
expPhase
statPhase
log2 expression
Position (nt)
TU2820
TU2821
TU2825
TU2827
TU2828
slr0645
dacB
slr0646
ncr1380
metS
slr0649
slr0650
sll0603
menD
sll0602
ncl1330
sll0601
TU2819
TU2822
TU2823
TU2824
TU2826
TU2829
2670000
2671250
2672500
2673750
2675000
2676250
2677500
2678750
2680000

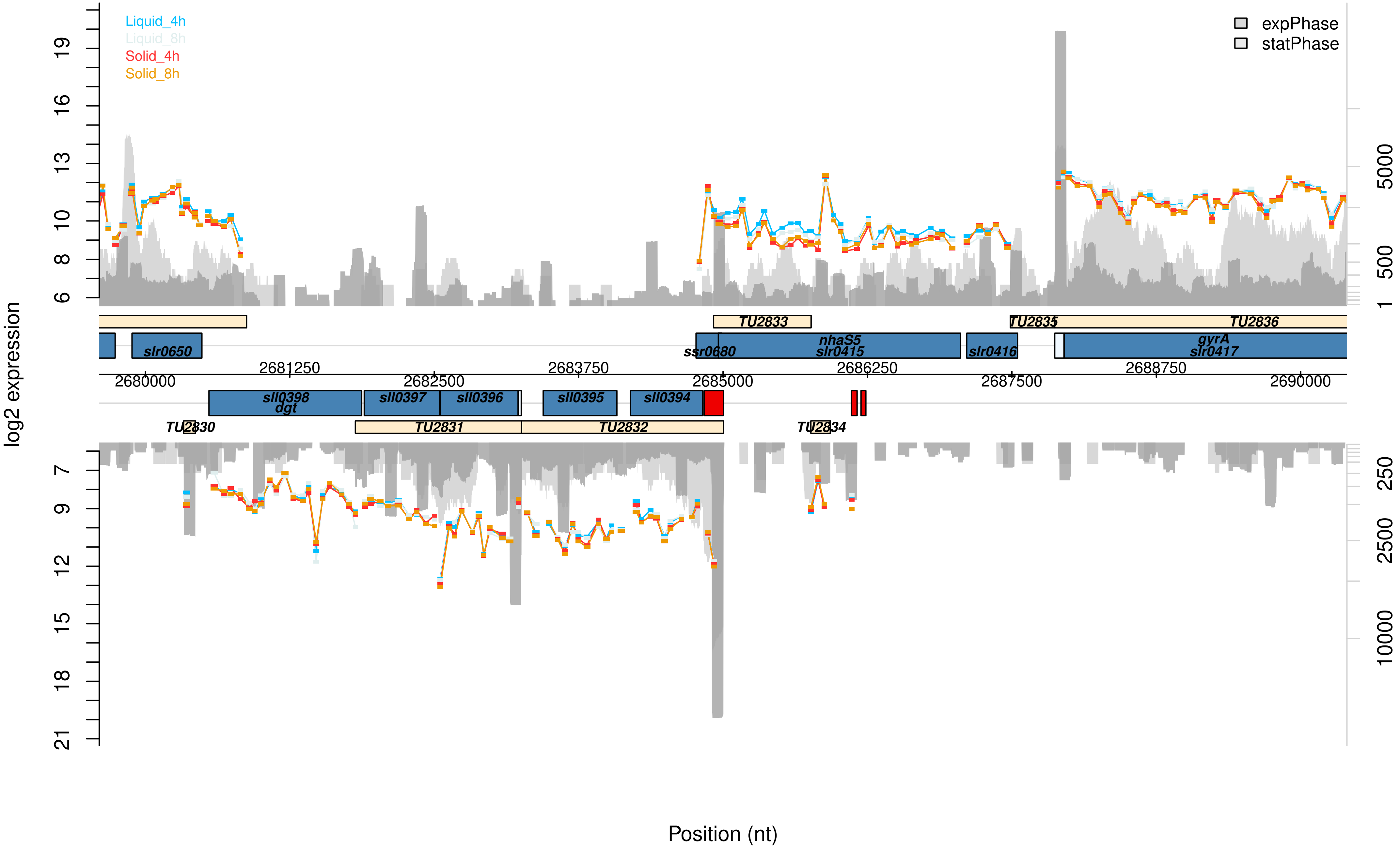

Liquid_4h
Liquid_8h
Solid_4h
Solid_8h
expPhase
statPhase
log2 expression
Position (nt)
TU2830
TU2831
TU2832
TU2833
TU2834
TU2835
TU2836
slr0650
sll0398
dgt
sll0397
sll0396
sll0395
sll0394
ssr0680
nhaS5
slr0415
slr0416
gyrA
slr0417
2680000
2681250
2682500
2683750
2685000
2686250
2687500
2688750
2690000

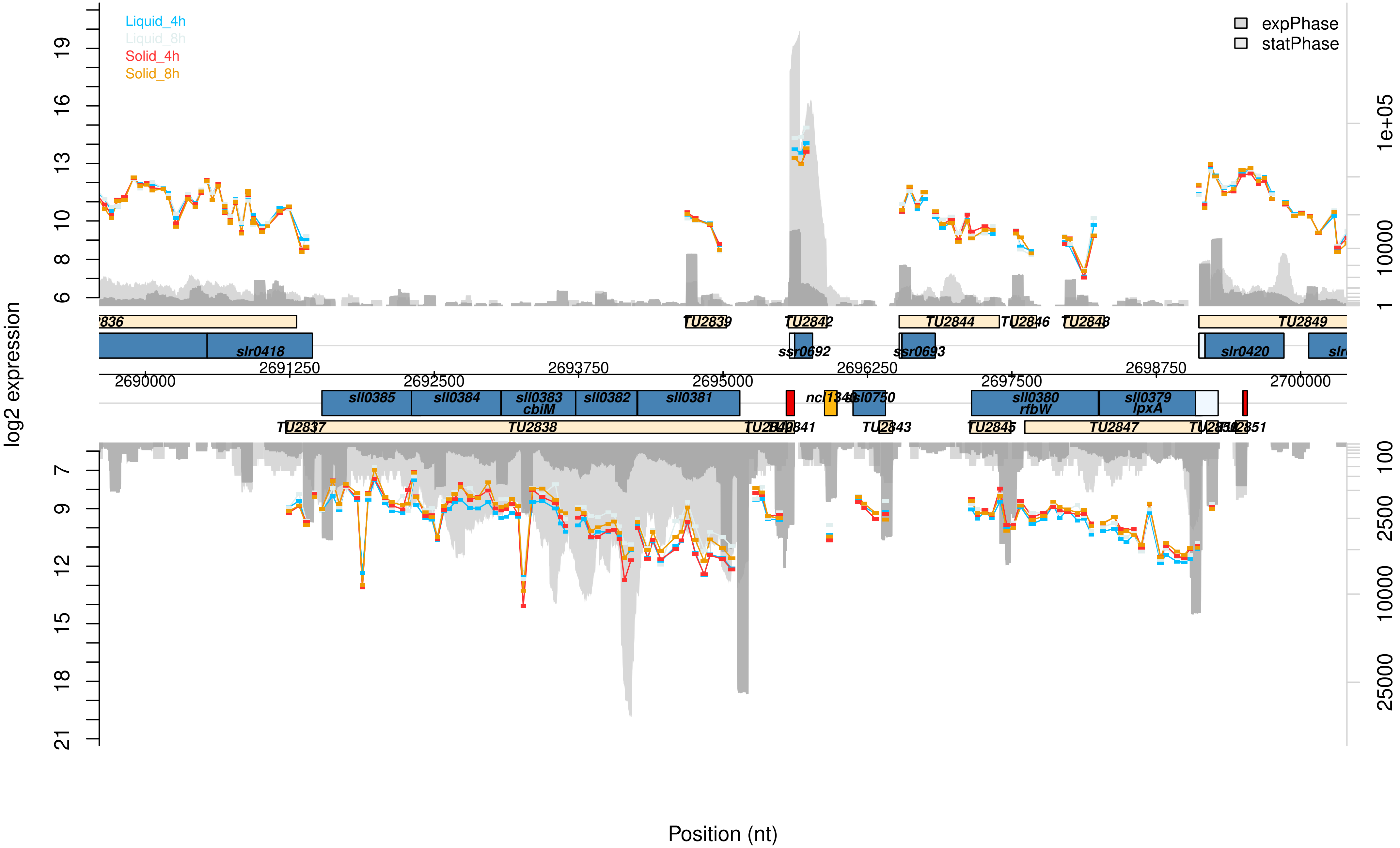

Liquid_4h
Liquid_8h
Solid_4h
Solid_8h
expPhase
statPhase
log2 expression
Position (nt)
TU2839
TU2842
TU2844
TU2846
TU2848
TU2849
slr0418
ssr0692
ssr0693
slr0420
sll0385
sll0384
sll0383
cbiM
sll0382
sll0381
sll0380
rfbW
sll0379
lpxA
TU2837
TU2838
TU2843
TU2845
TU2847
2690000
2691250
2692500
2693750
2695000
2696250
2697500
2698750
2700000

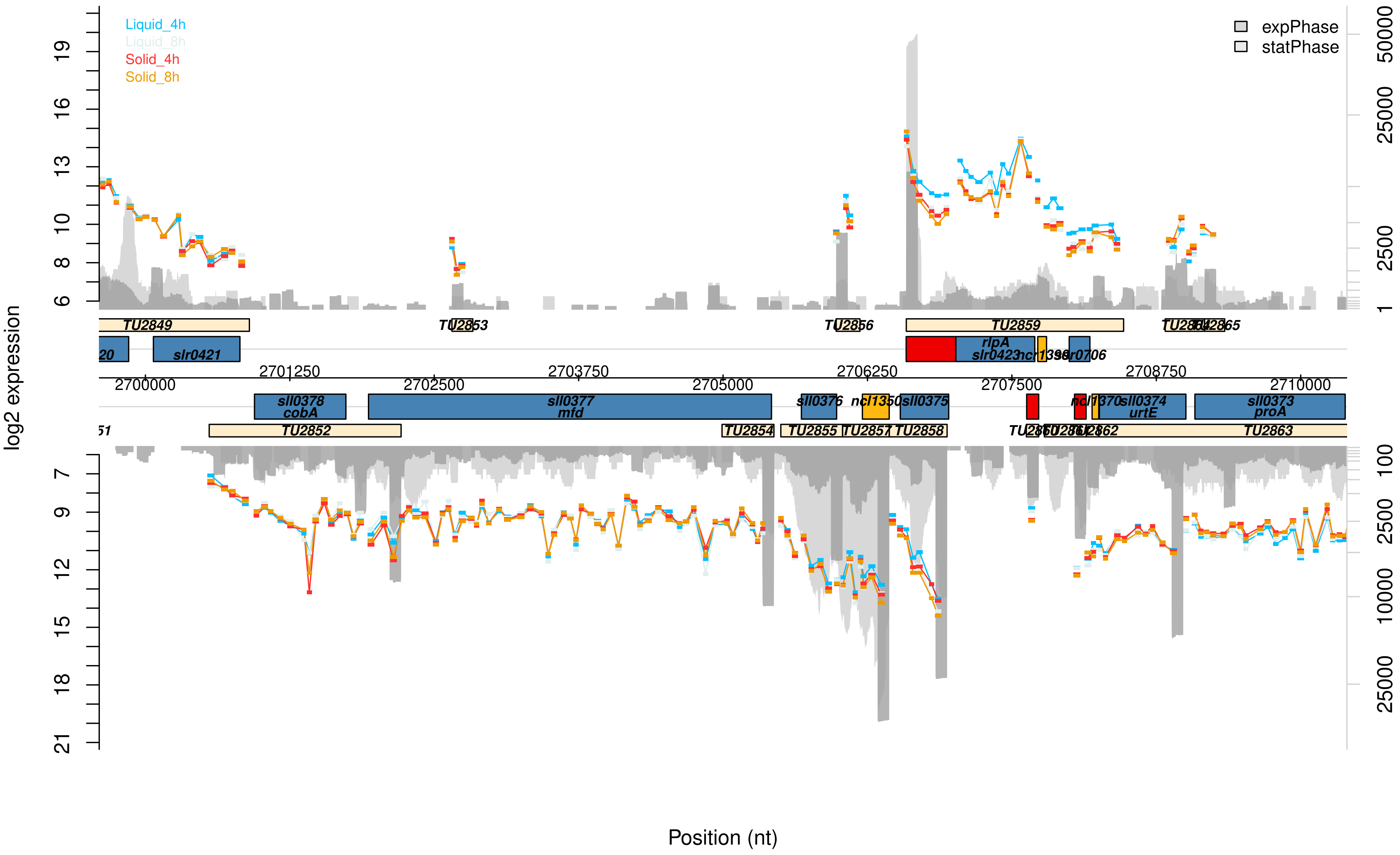

Liquid_4h
Liquid_8h
Solid_4h
Solid_8h
expPhase
statPhase
log2 expression
Position (nt)
TU2849
TU2853
TU2856
TU2859
TU2852
TU2854
TU2855
TU2857
TU2858
TU2863
slr0421
rlpA
slr0423
sll0378
cobA
sll0377
mfd
sll0376
ncl1350
sll0375
sll0374
urtE
sll0373
proA
2700000
2701250
2702500
2703750
2705000
2706250
2707500
2708750
2710000

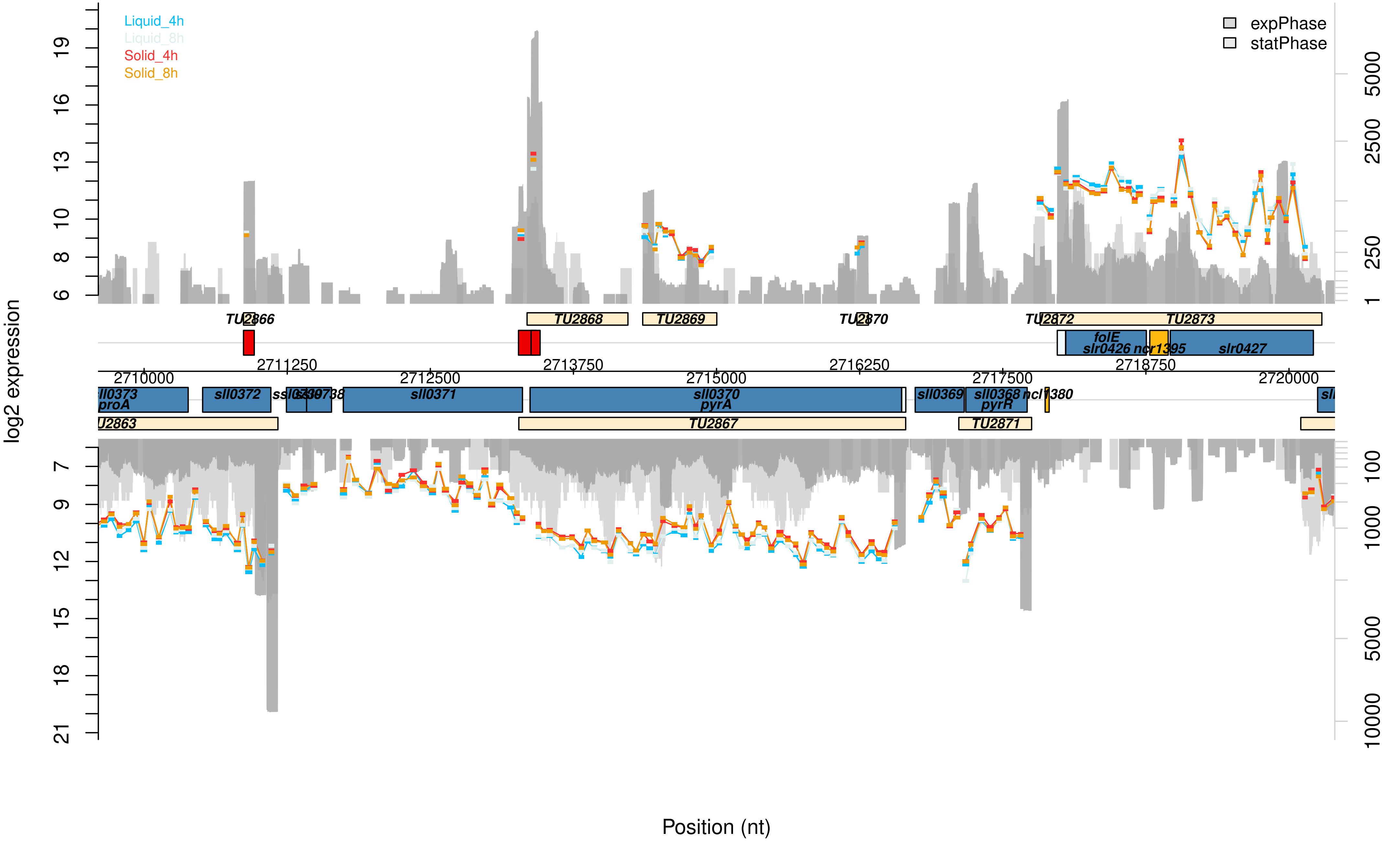

Liquid_4h
Liquid_8h
Solid_4h
Solid_8h
expPhase
statPhase
log2 expression
Position (nt)
TU2866
TU2868
TU2869
TU2870
TU2872
TU2873
folE
slr0426
ncr1395
slr0427
2710000
2711250
2712500
2713750
2715000
2716250
2717500
2718750
2720000
sll0373
proA
sll0372
sll0371
sll0370
pyrA
sll0369
sll0368
pyrR
ncl1380
TU2863
TU2867
TU2871

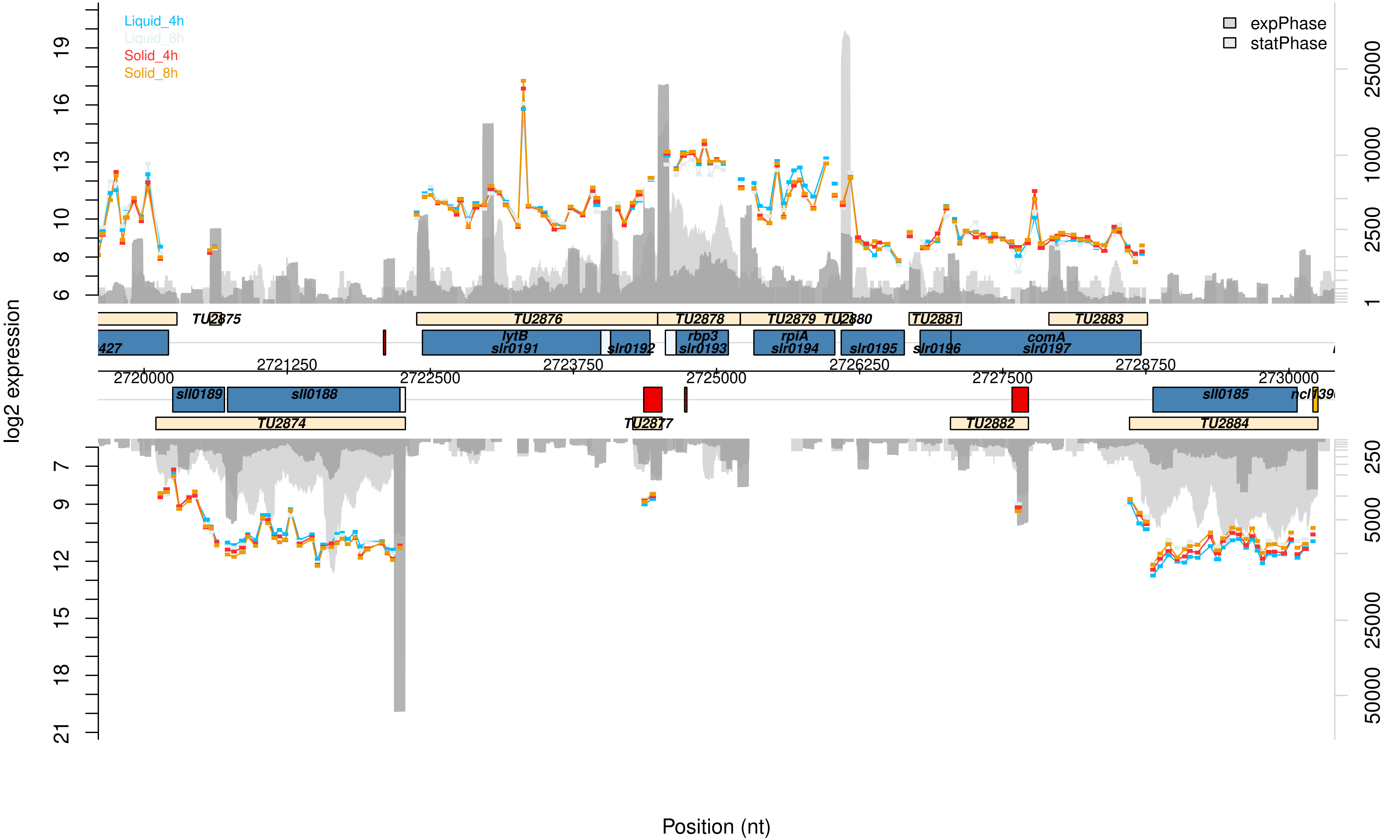

Liquid_4h
Liquid_8h
Solid_4h
Solid_8h
expPhase
statPhase
log2 expression
Position (nt)
TU2875
TU2876
TU2878
TU2879
TU2880
TU2881
TU2883
TU2874
TU2877
TU2882
TU2884
lytB
slr0191
slr0192
rbp3
slr0193
rpiA
slr0194
slr0195
slr0196
comA
slr0197
sll0189
sll0188
sll0185
2720000
2721250
2722500
2723750
2725000
2726250
2727500
2728750
2730000

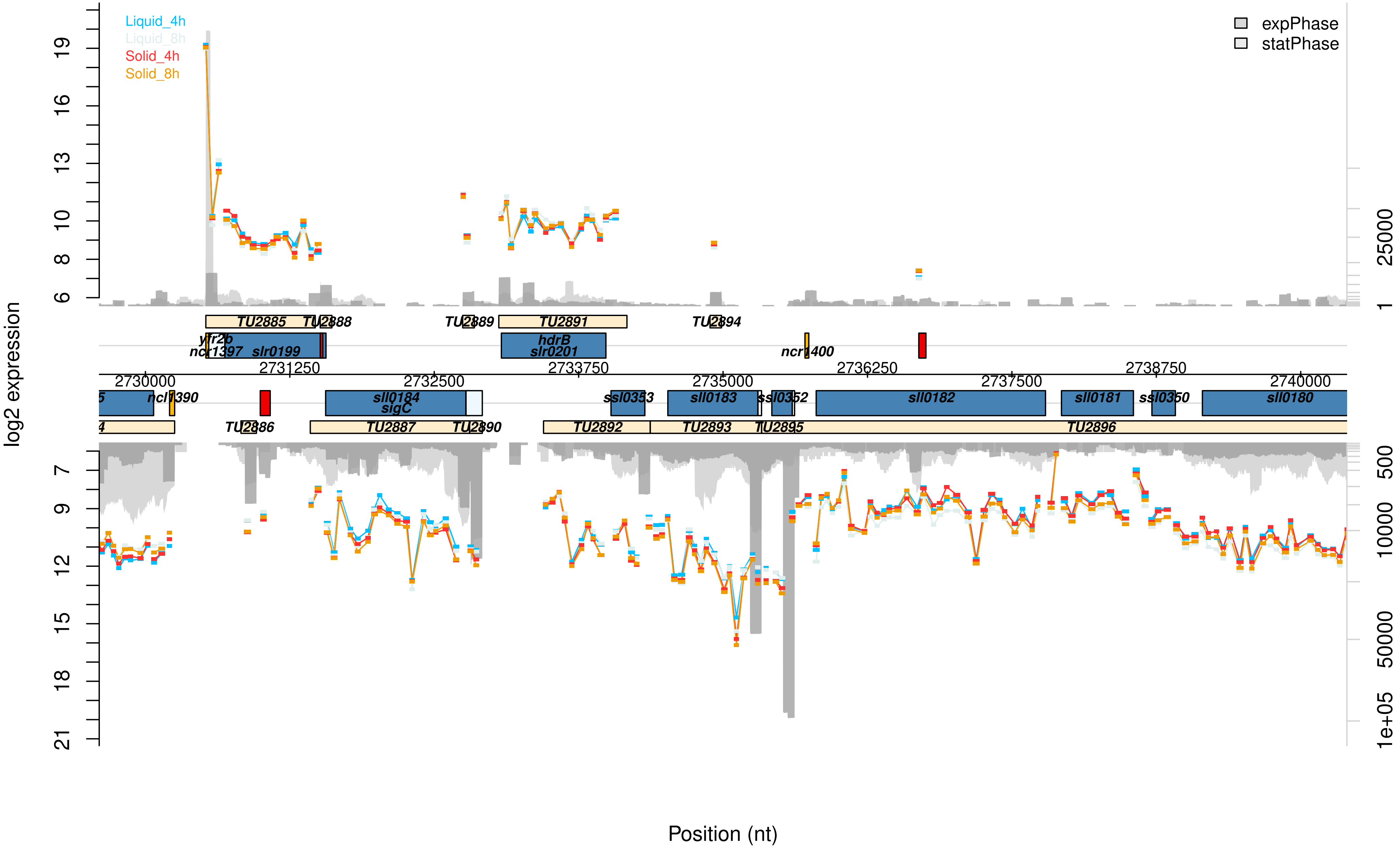

Liquid_4h
Liquid_8h
Solid_4h
Solid_8h
expPhase
statPhase
log2 expression
Position (nt)
TU2885
TU2888
TU2889
TU2891
TU2894
yfr2b
ncr1397
slr0199
hdrB
slr0201
ncr1400
ncl1390
sll0184
sigC
ssl0353
sll0183
ssl0352
sll0182
sll0181
ssl0350
sll0180
TU2886
TU2887
TU2890
TU2892
TU2893
TU2895
TU2896

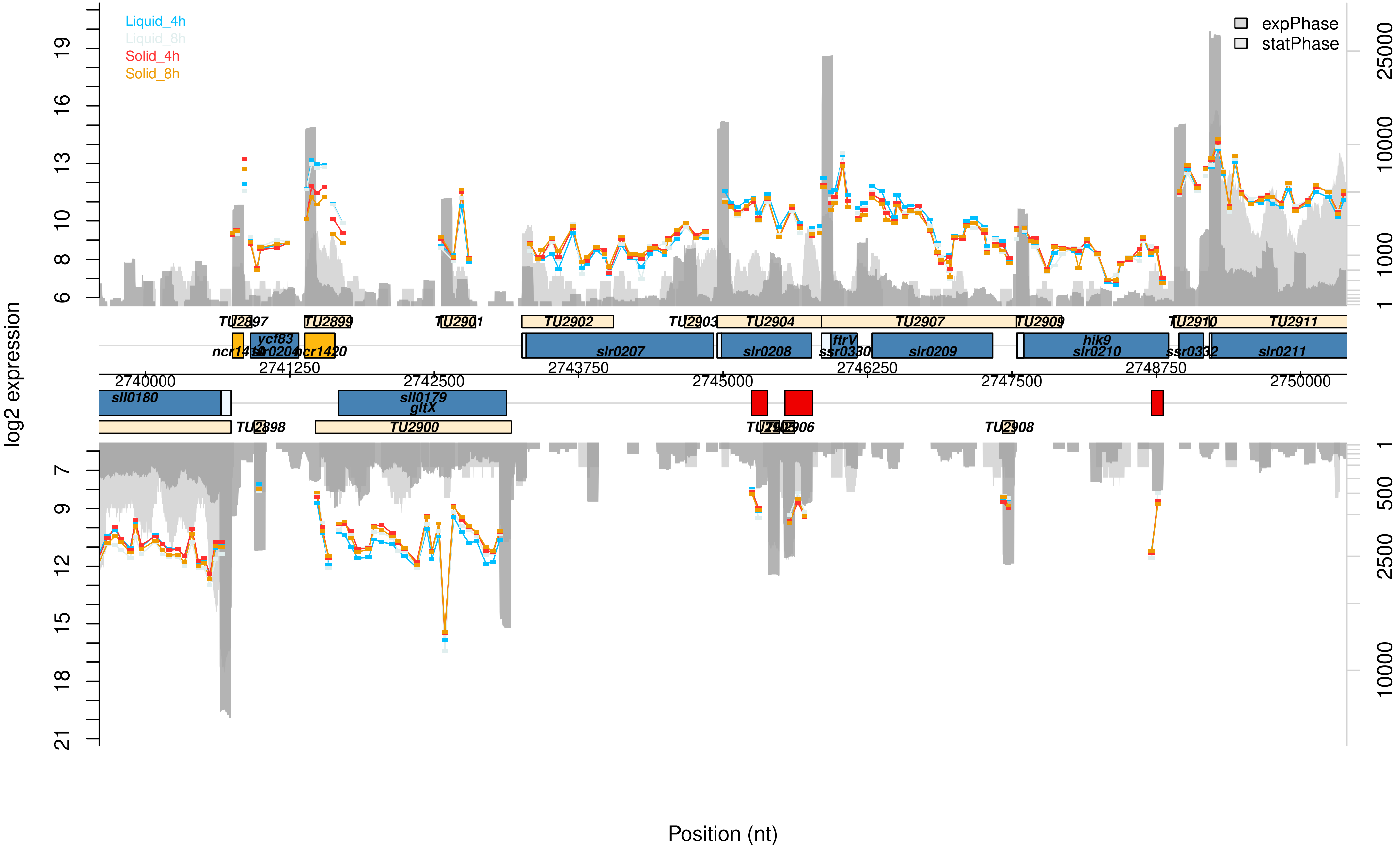
Liquid_4h
Liquid_8h
Solid_4h
Solid_8h
expPhase
statPhase
log2 expression
Position (nt)
TU2897
TU2899
TU2901
TU2902
TU2903
TU2904
TU2907
TU2909
TU2910
TU2911
ncr1410
ycf83
slr0204
ncr1420
slr0207
slr0208
ftrV
ssr0330
slr0209
hik9
slr0210
ssr0332
slr0211
sll0180
sll0179
gltX
TU2898
TU2900
TU2908
2740000
2741250
2742500
2743750
2745000
2746250
2747500
2748750
2750000

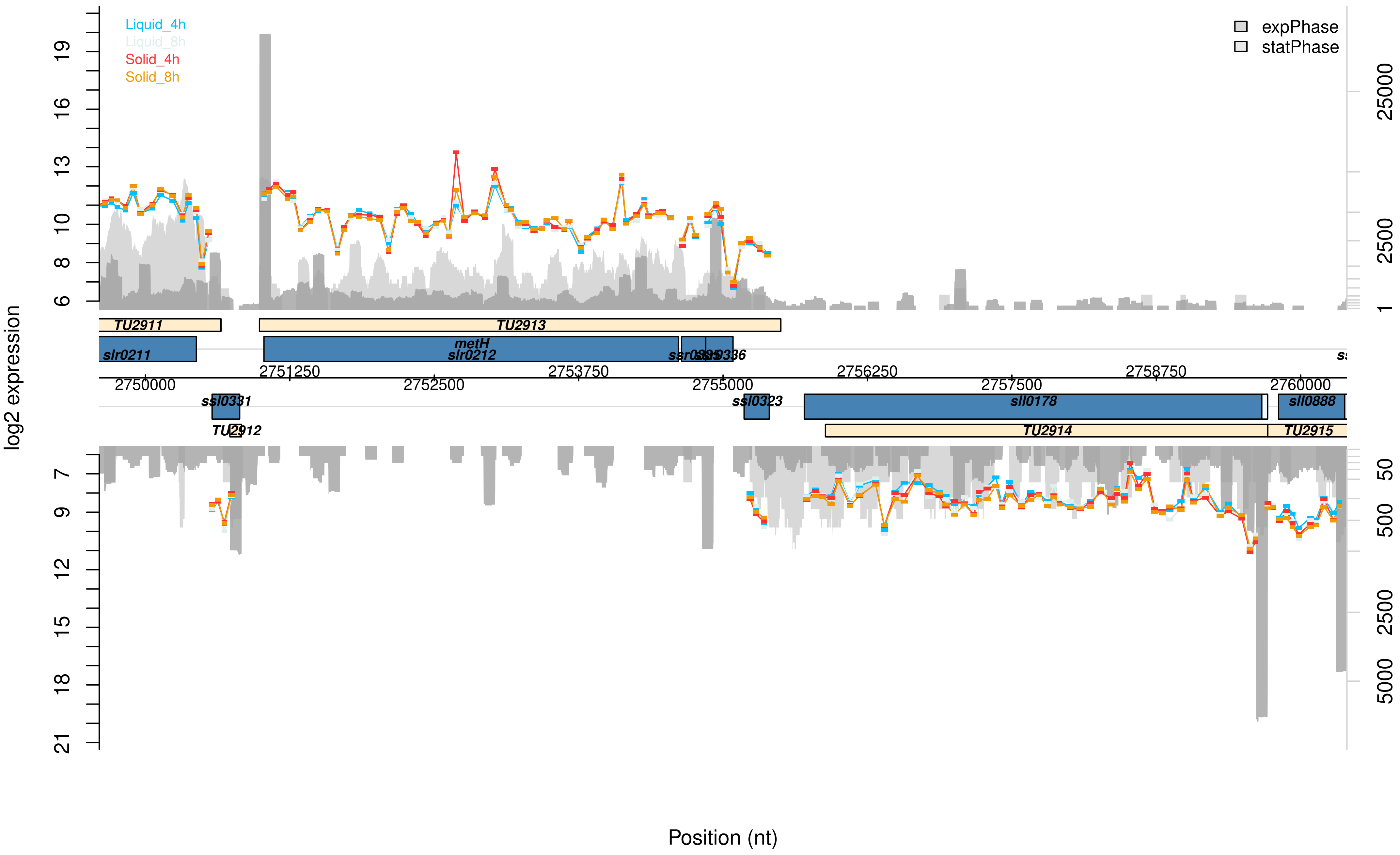

Liquid_4h
Liquid_8h
Solid_4h
Solid_8h
expPhase
statPhase
log2 expression
TU2911
TU2913
slr0211
metH
slr0212
2751250
2753750
2756250
2758750
2750000
2752500
2755000
2757500
2760000
ssl0331
ssl0323
sll0178
sll0888
TU2912
TU2914
TU2915
Position (nt)

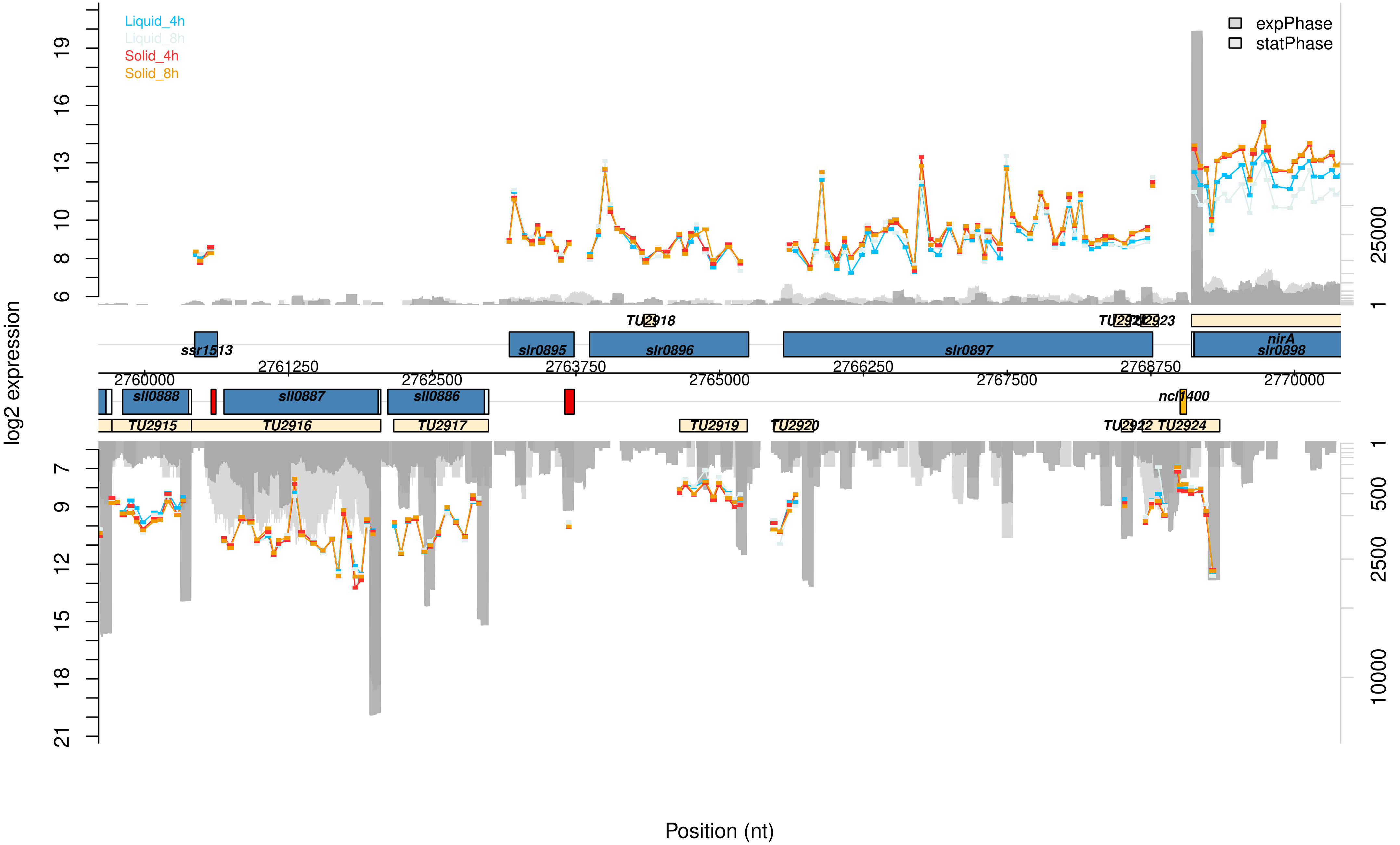

Liquid_4h
Liquid_8h
Solid_4h
Solid_8h
expPhase
statPhase
log2 expression
Position (nt)
ssr1513
slr0895
slr0896
slr0897
nirA
slr0898
TU2918
sll0888
sll0887
sll0886
ncl1400
TU2915
TU2916
TU2917
TU2919
TU2920
TU2924
2760000
2761250
2762500
2763750
2765000
2766250
2767500
2768750
2770000

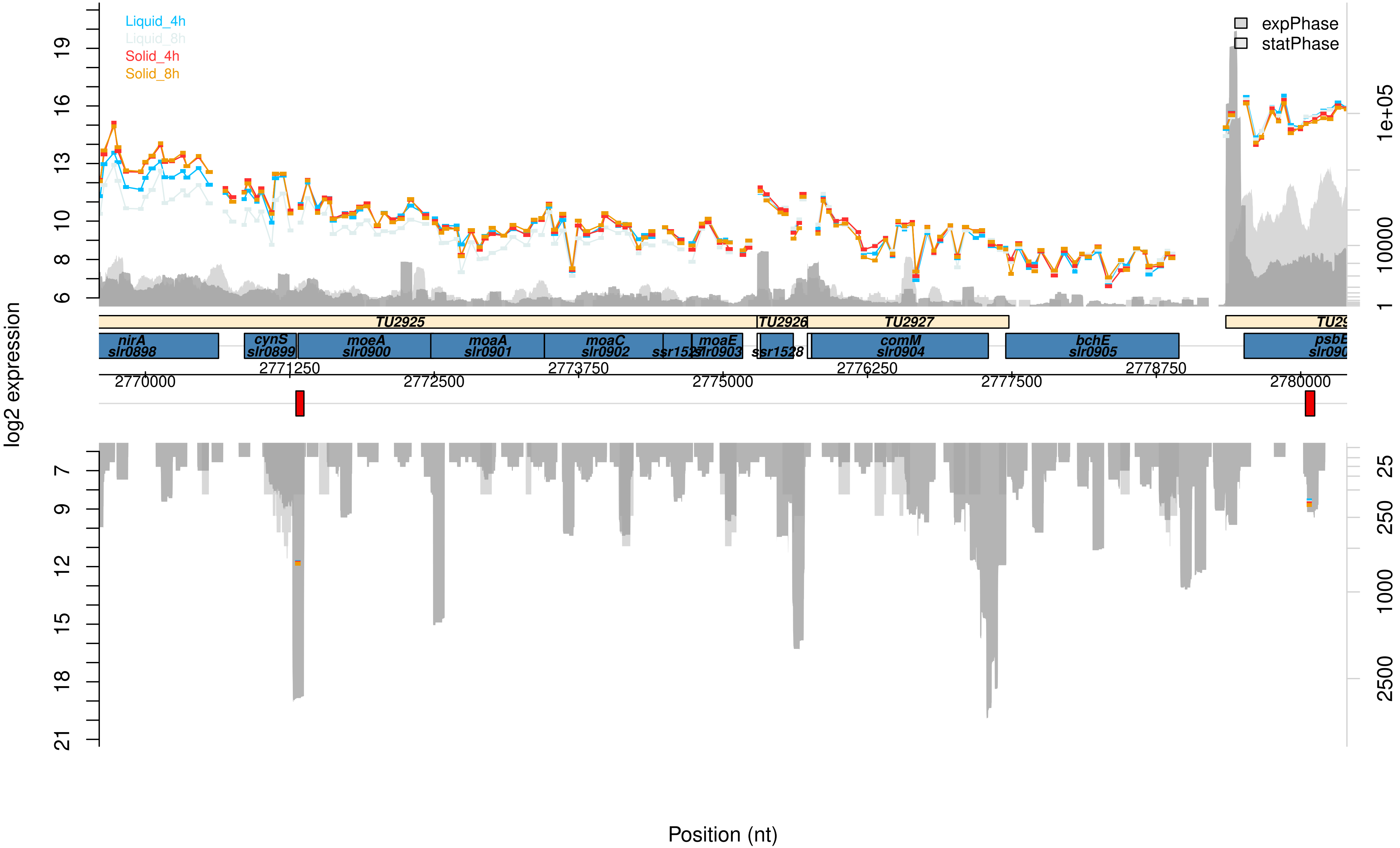

Liquid_4h
Liquid_8h
Solid_4h
Solid_8h
expPhase
statPhase
log2 expression
TU2925
TU2926
TU2927
nirA slr0898
cynS slr0899
moeA slr0900
moaA slr0901
moaC slr0902
ssr1527
moaE slr0903
ssr1528
comM slr0904
bchE slr0905
psbE
2770000
2771250
2772500
2773750
2775000
2776250
2777500
2778750
2780000
Position (nt)

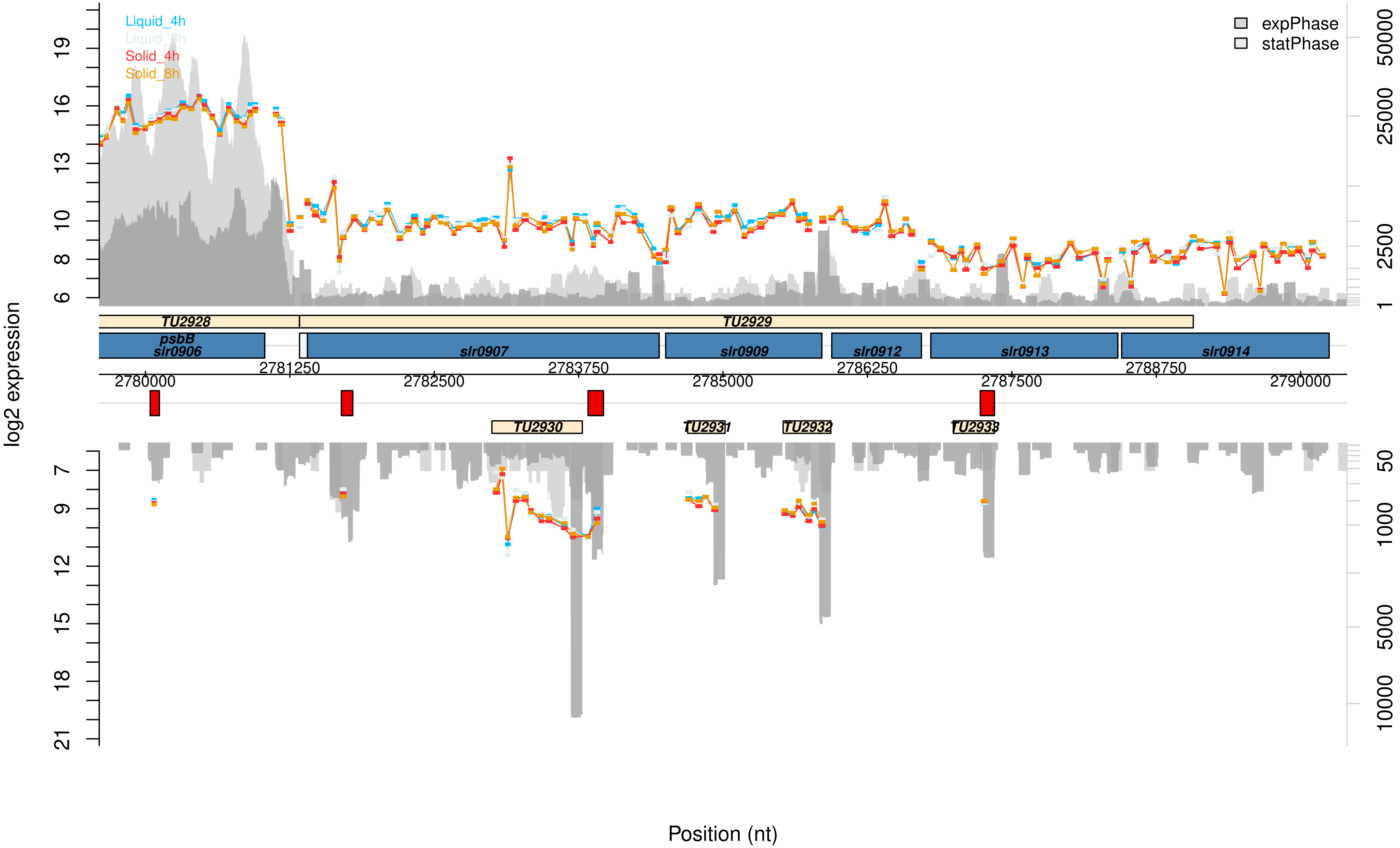

Liquid_4h
Liquid_8h
Solid_4h
Solid_8h
expPhase
statPhase
log2 expression
TU2928
TU2929
psbB
slr0906
slr0907
slr0909
slr0912
slr0913
slr0914
TU2930
TU2931
TU2932
TU2933
2780000
2781250
2782500
2783750
2785000
2786250
2787500
2788750
2790000
Position (nt)

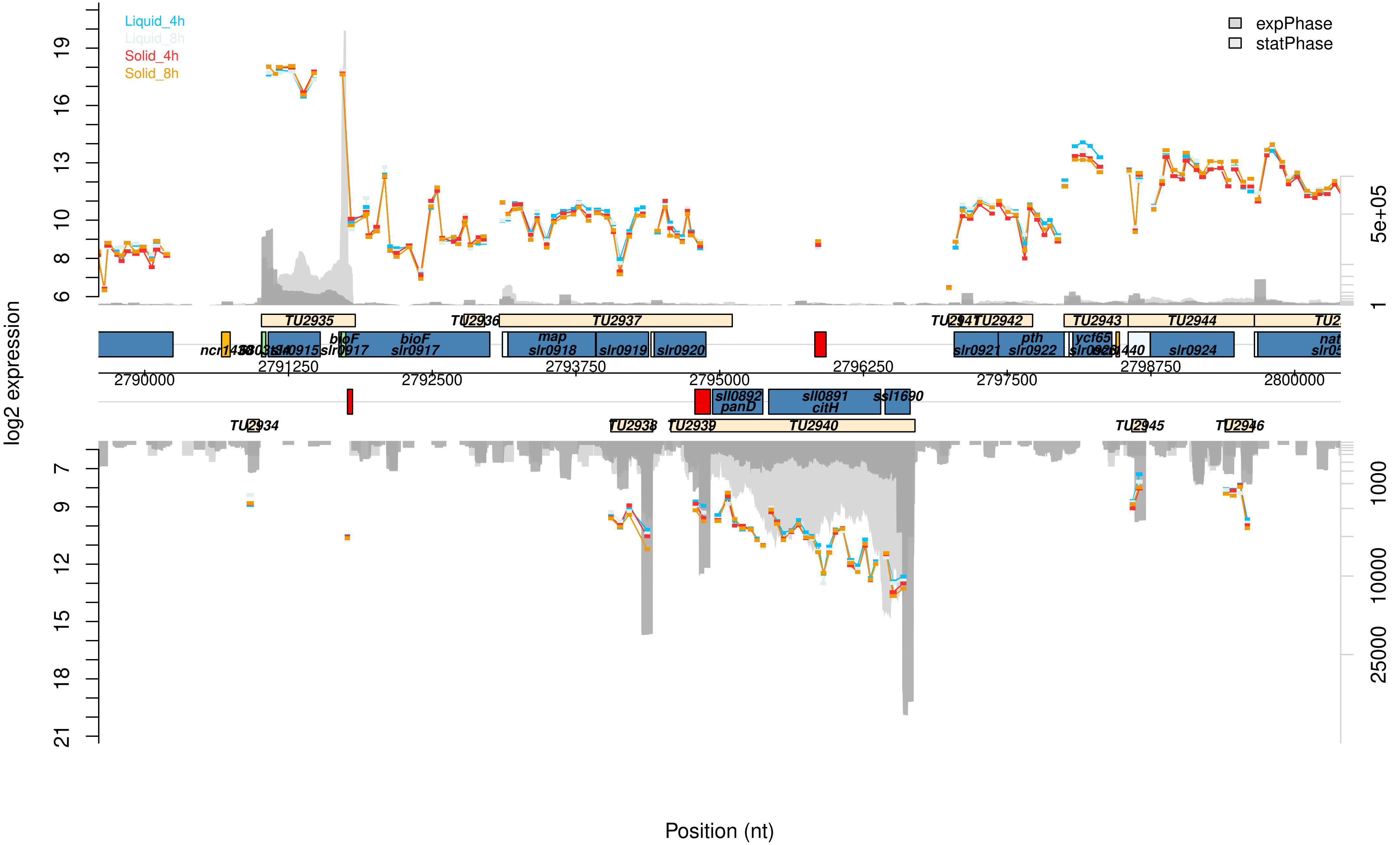

Liquid_4h
Liquid_8h
Solid_4h
Solid_8h
expPhase
statPhase
log2 expression
Position (nt)
TU2934
TU2935
TU2936
TU2937
TU2938
TU2939
TU2940
TU2942
TU2943
TU2944
TU2945
TU2946
bioF
slr0917
map
slr0918
slr0919
slr0920
slr0921
pth
slr0922
ycf65
slr0924
sll0892
panD
sll0891
citH
ssl1690
2790000
2791250
2792500
2793750
2795000
2796250
2797500
2798750
2800000
5e+05
1000
10000
25000

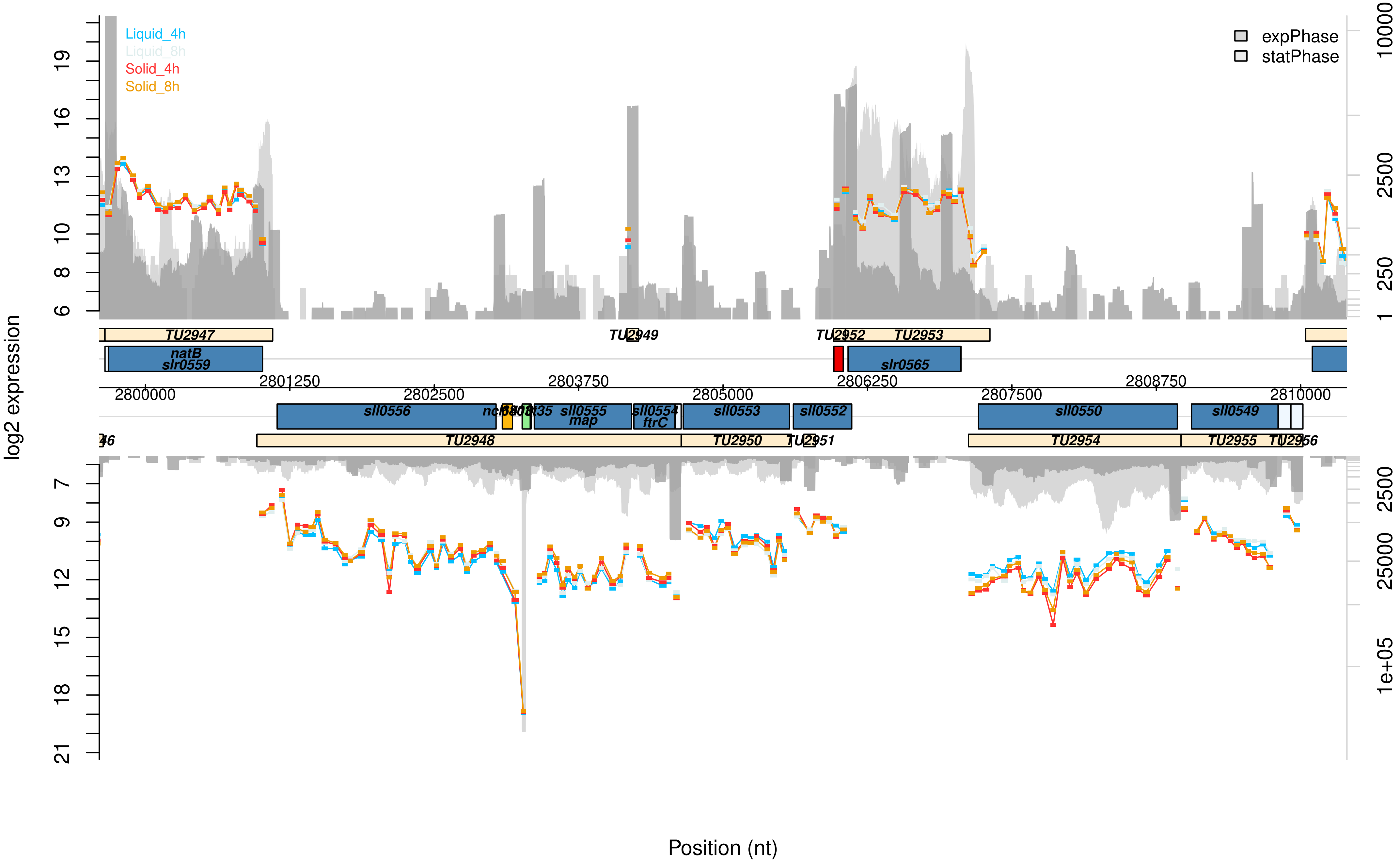

Liquid_4h
Liquid_8h
Solid_4h
Solid_8h
expPhase
statPhase
log2 expression
natB
slr0559
TU2947
TU2949
TU2952
TU2953
slr0565
sll0556
sll0555
map
sll0554
ftrC
sll0553
sll0552
sll0550
sll0549
TU2948
TU2950
TU2951
TU2954
TU2955
TU2956
2800000
2801250
2802500
2803750
2805000
2806250
2807500
2808750
2810000
Position (nt)

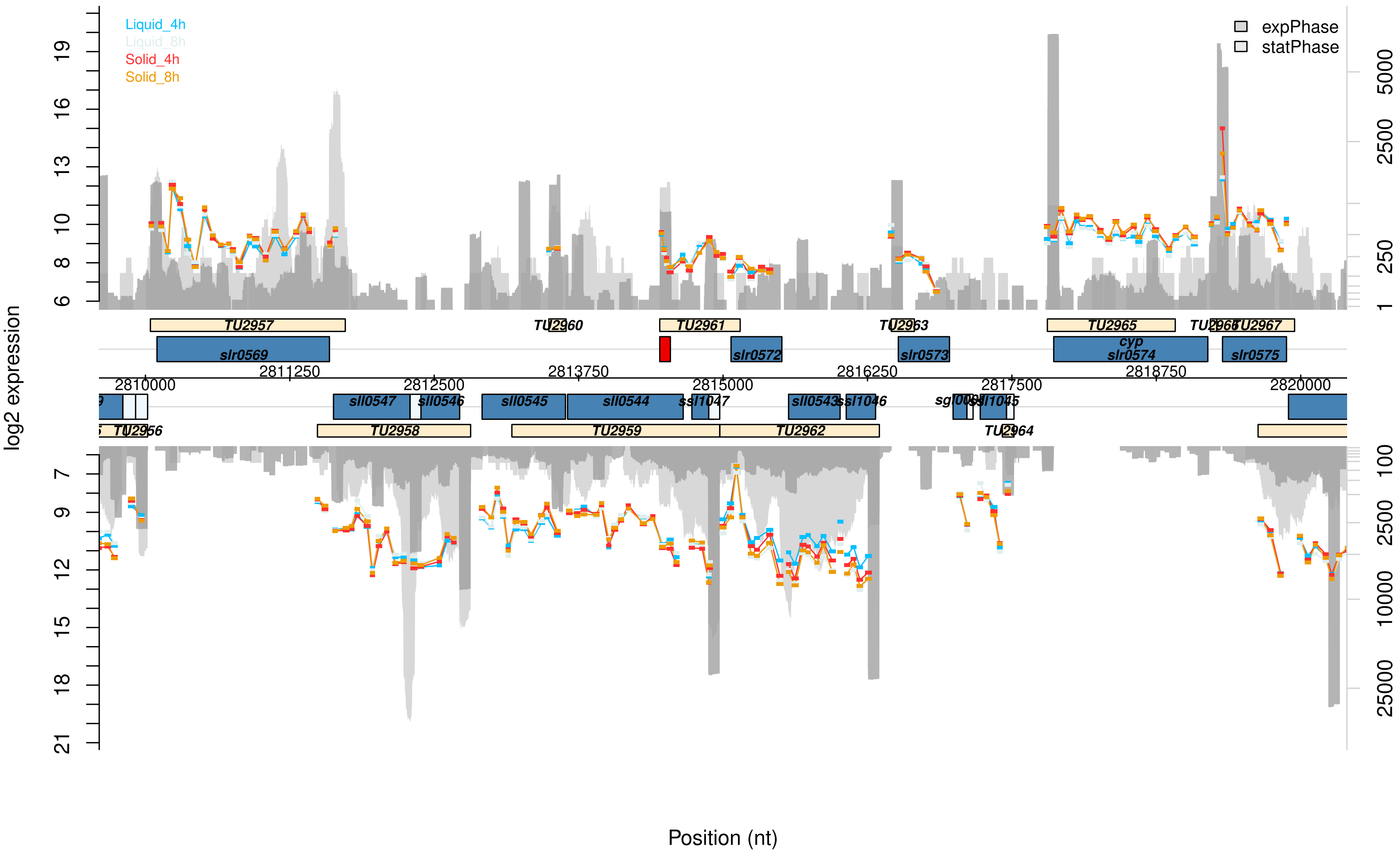

Liquid_4h
Liquid_8h
Solid_4h
Solid_8h
expPhase
statPhase
log2 expression
Position (nt)
TU2957
TU2960
TU2961
TU2963
TU2965
TU2966
TU2967
slr0569
slr0572
slr0573
cyp
slr0574
slr0575
2811250
2813750
2816250
2818750
2810000
2812500
2815000
2817500
2820000
sll0547
sll0546
sll0545
sll0544
ssl1047
sll0543
ssl1046
ssl1045
TU2956
TU2958
TU2959
TU2962
TU2964

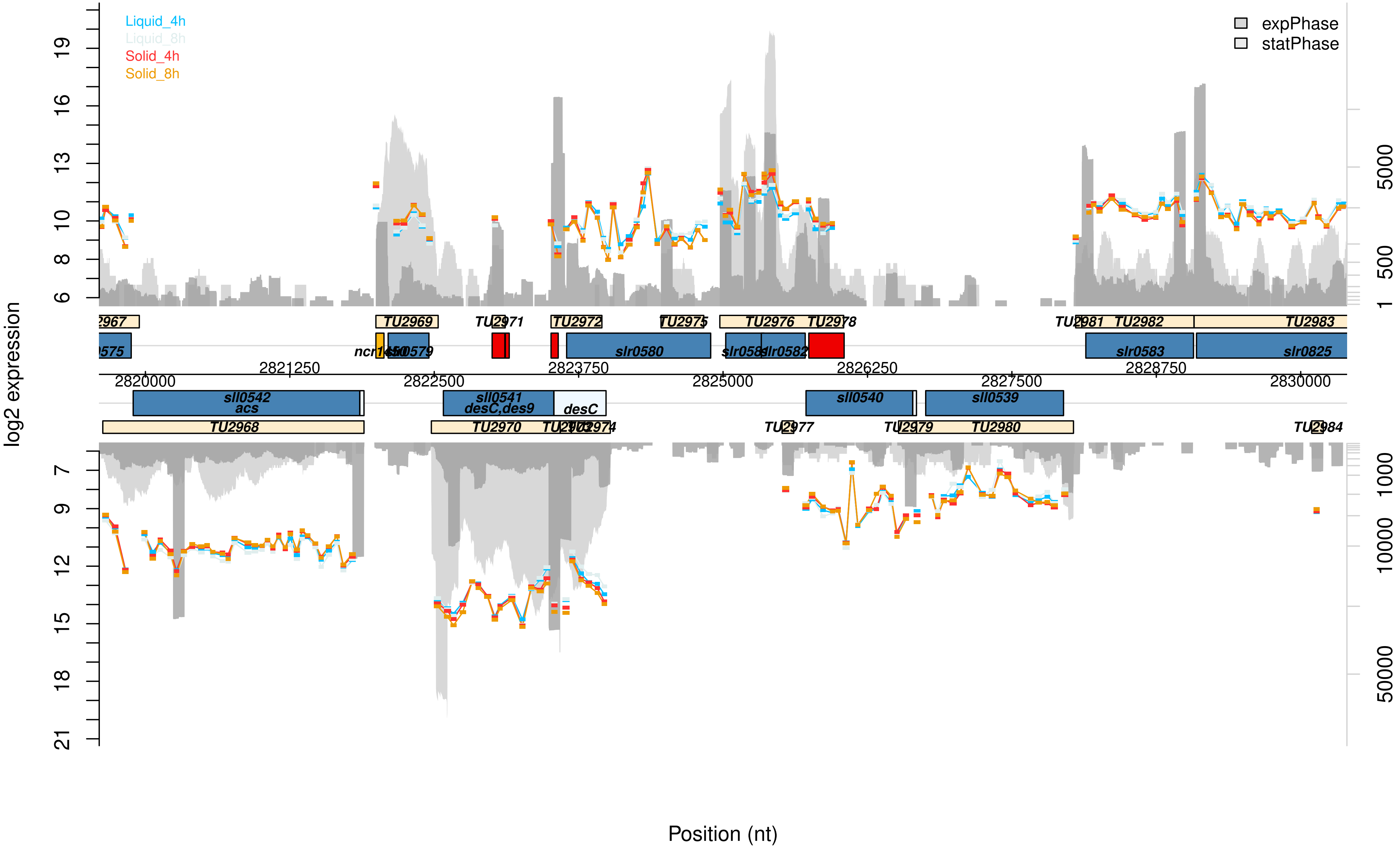

Liquid_4h
Liquid_8h
Solid_4h
Solid_8h
expPhase
statPhase
log2 expression
Position (nt)
2820000
2821250
2822500
2823750
2825000
2826250
2827500
2828750
2830000
TU2969
TU2972
TU2975
TU2976
TU2978
TU2981
TU2982
TU2983
ncr1450
slr0580
slr0583
slr0825
sll0542
acs
sll0541
desC,des9
desC
sll0540
sll0539
TU2968
TU2970
TU2980

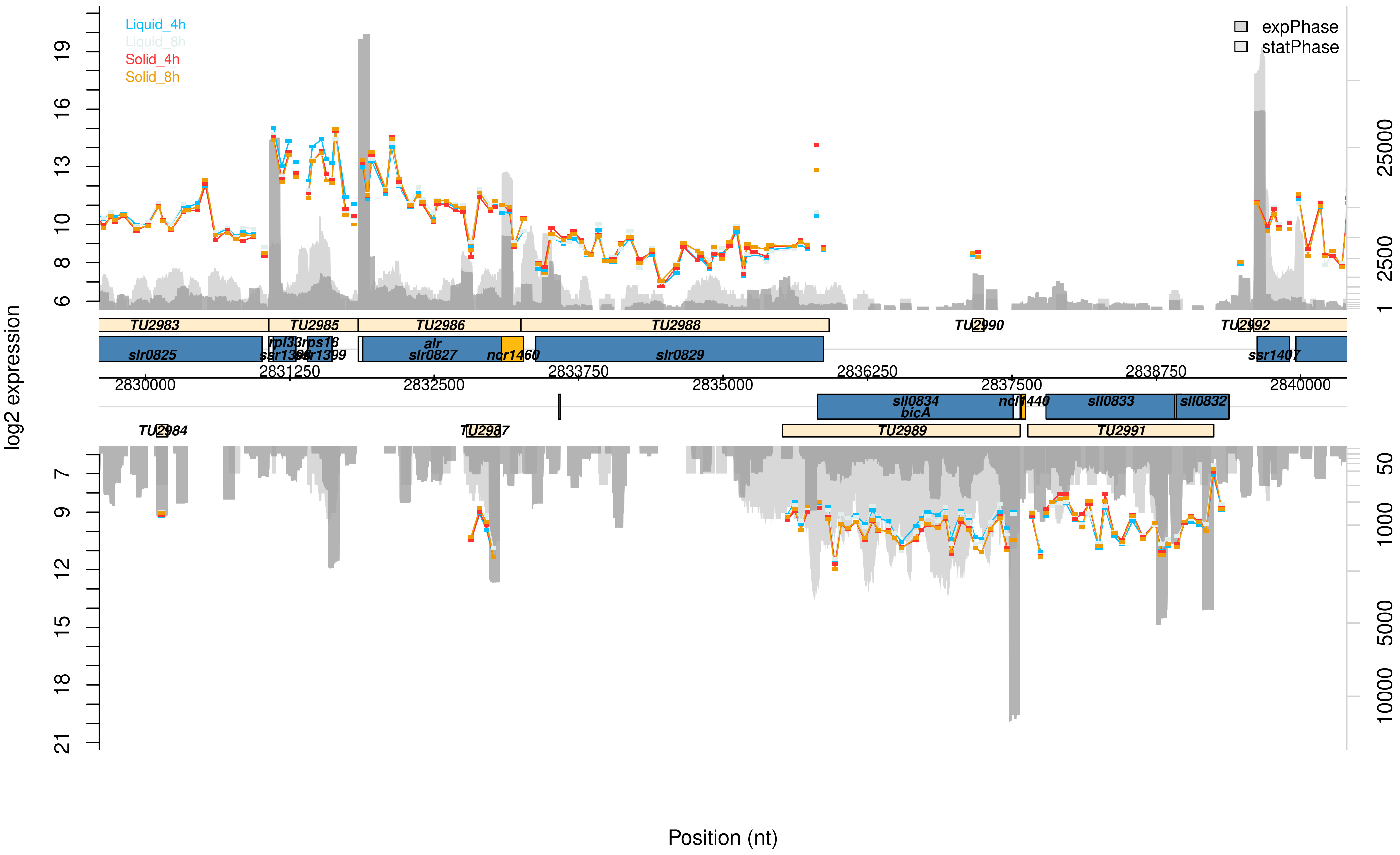

Liquid_4h
Liquid_8h
Solid_4h
Solid_8h
expPhase
statPhase
log2 expression
Position (nt)
TU2983
TU2985
TU2986
TU2988
TU2990
TU2992
slr0825
rpl33
rps18
ssr1398
ssr1399
alr
slr0827
ncr1460
slr0829
ssr1407
2830000
2831250
2832500
2833750
2835000
2836250
2837500
2838750
2840000
sll0834
bicA
ncl1440
sll0833
sll0832
TU2984
TU2987
TU2989
TU2991

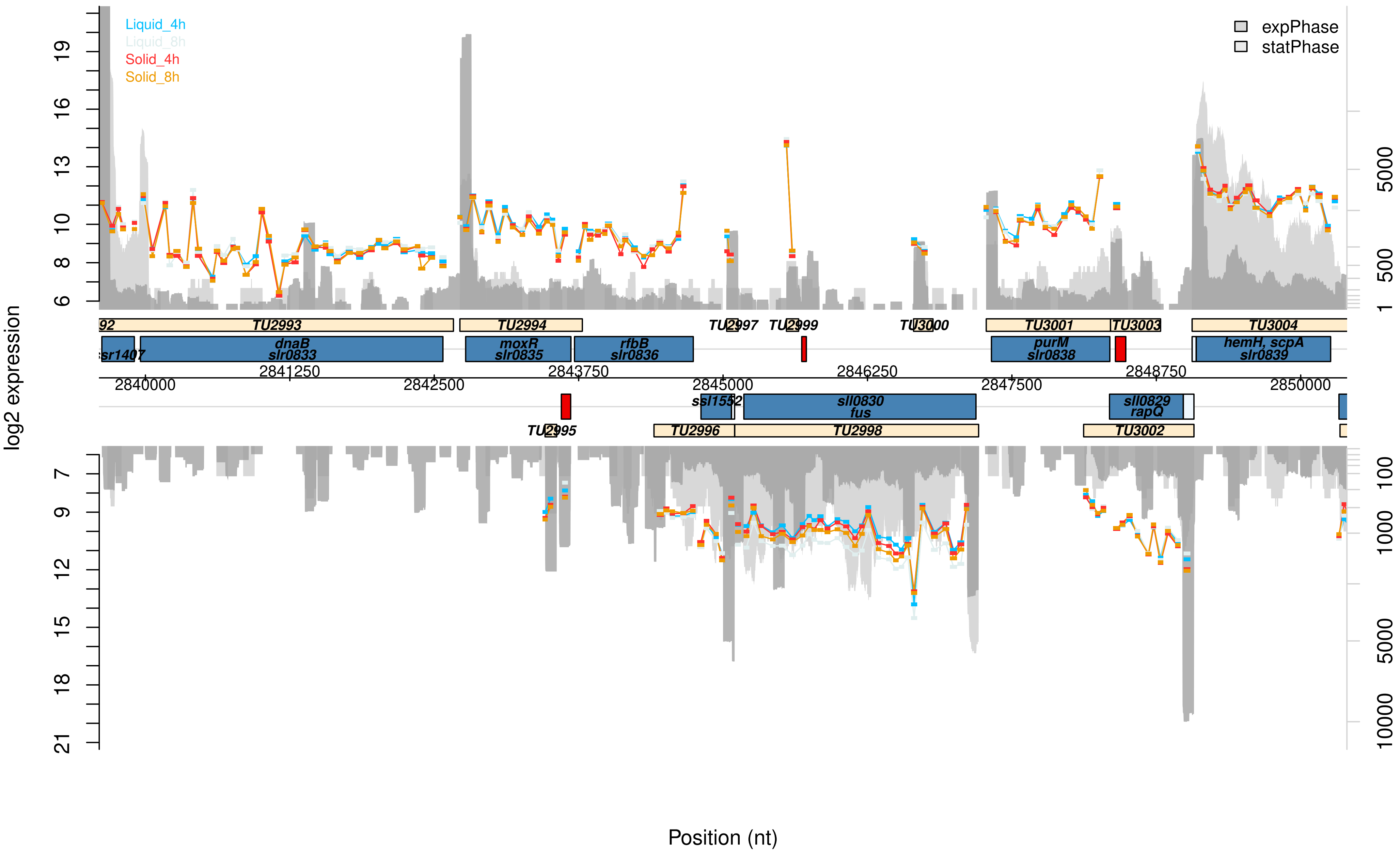

Liquid_4h
Liquid_8h
Solid_4h
Solid_8h
expPhase
statPhase
log2 expression
Position (nt)
TU2993
TU2994
TU2997
TU2999
TU3000
TU3001
TU3003
TU3004
dnaB
slr0833
moxR
slr0835
rfbB
slr0836
purM
slr0838
hemH, scpA
slr0839
ssl1552
sll0830
fus
sll0829
rapQ
TU2995
TU2996
TU2998
TU3002
2840000
2841250
2842500
2843750
2845000
2846250
2847500
2848750
2850000

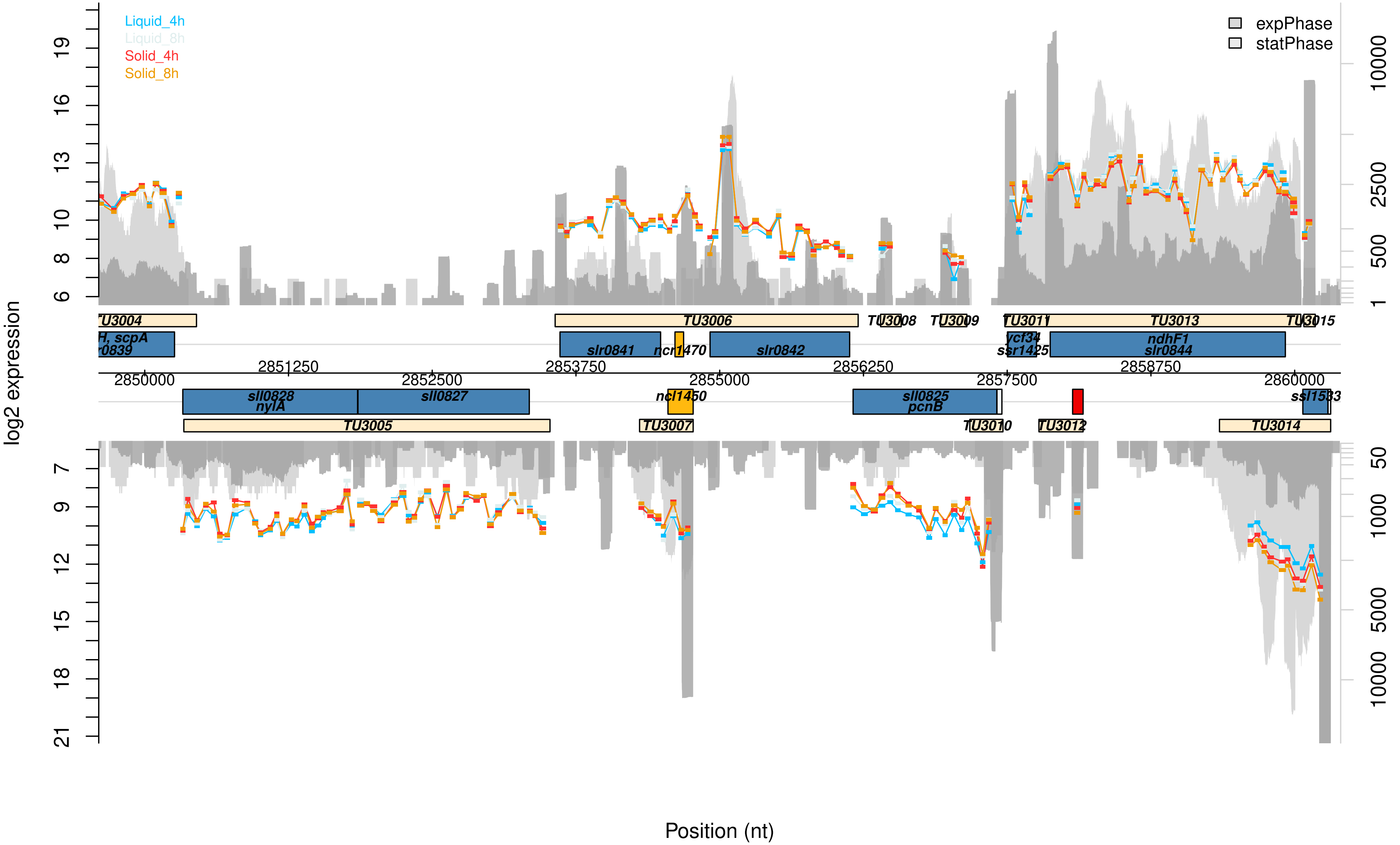

Liquid_4h
Liquid_8h
Solid_4h
Solid_8h
expPhase
statPhase
log2 expression
Position (nt)
TU3004
TU3006
TU3008
TU3009
TU3011
TU3013
TU3015
TU3005
TU3007
TU3010
TU3012
TU3014
slr0841
ncr1470
slr0842
ycf34
ssr1425
ndhF1
slr0844
sll0828
nylA
sll0827
ncl1450
sll0825
pcnB
ssl1533
2850000
2851250
2852500
2853750
2855000
2856250
2857500
2858750
2860000

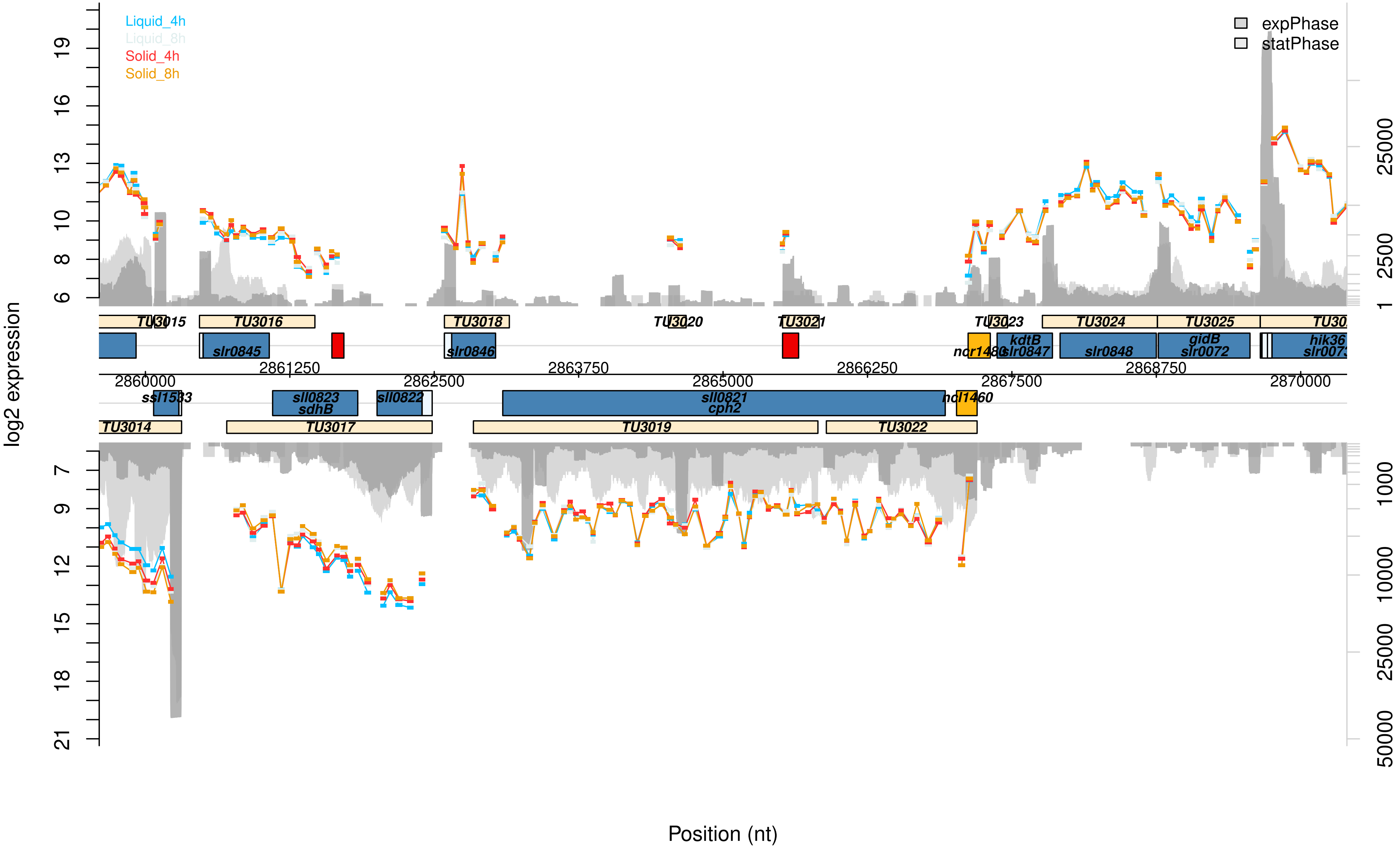

Liquid_4h
Liquid_8h
Solid_4h
Solid_8h
expPhase
statPhase
log2 expression
Position (nt)
TU3014
TU3015
TU3016
TU3017
TU3018
TU3019
TU3020
TU3021
TU3022
TU3023
TU3024
TU3025
slr0845
slr0846
ssl1533
sll0823
sdhB
sll0822
sll0821
cph2
ncl1460
ncr1480
kdtB
slr0847
slr0848
gidB
slr0072
2860000
2861250
2862500
2863750
2865000
2866250
2867500
2868750
2870000

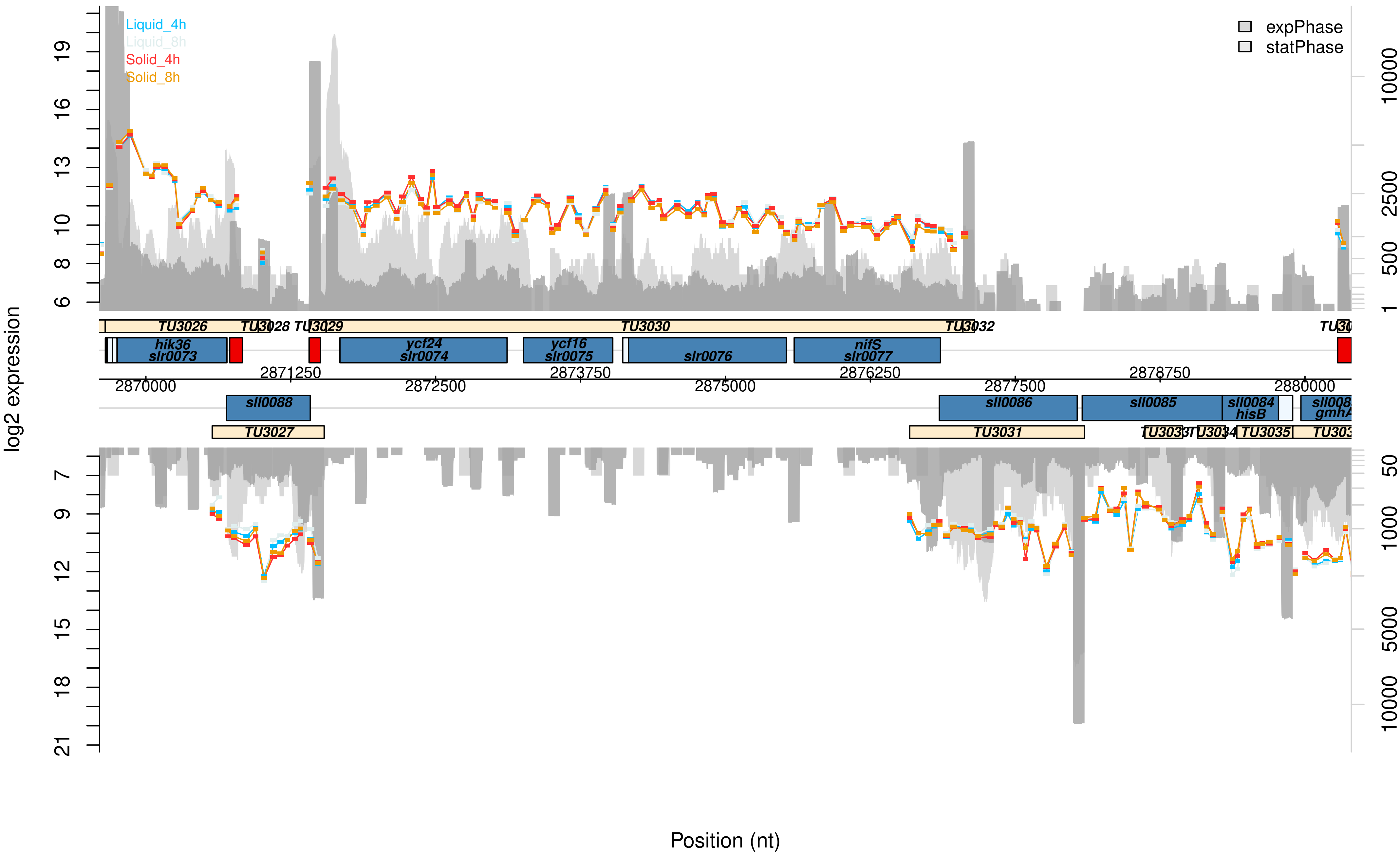

Liquid_4h
Liquid_8h
Solid_4h
Solid_8h
expPhase
statPhase
log2 expression
Position (nt)
TU3026
TU3028
TU3029
TU3030
TU3032
hik36
slr0073
ycf24
slr0074
ycf16
slr0075
slr0076
nifS
slr0077
2869000
2870000
2871250
2872500
2873750
2875000
2876250
2877500
2878750
2880000
sll0088
TU3027
sll0086
TU3031
sll0085
sll0084
hisB
TU3035

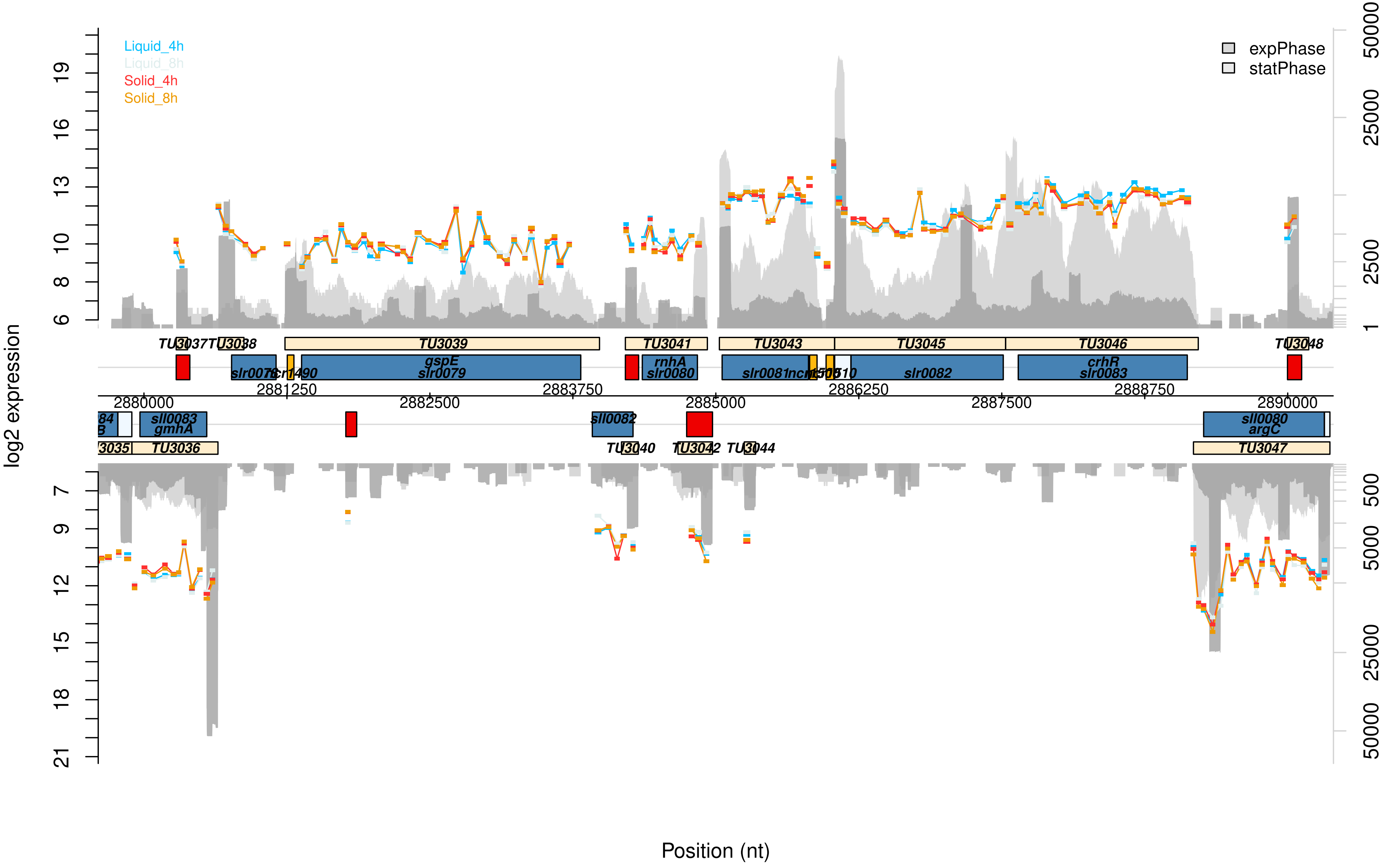
Liquid_4h
Liquid_8h
Solid_4h
Solid_8h
expPhase
statPhase
log2 expression
Position (nt)
TU3037
TU3038
TU3039
TU3041
TU3043
TU3045
TU3046
TU3048
slr0078
ncr1490
gspE
slr0079
rnhA
slr0080
slr0081
slr0082
crhR
slr0083
2880000
2881250
2882500
2883750
2885000
2886250
2887500
2888750
2890000
sll0083
gmhA
sll0082
sll0080
argC
TU3035
TU3036
TU3040
TU3042
TU3044
TU3047

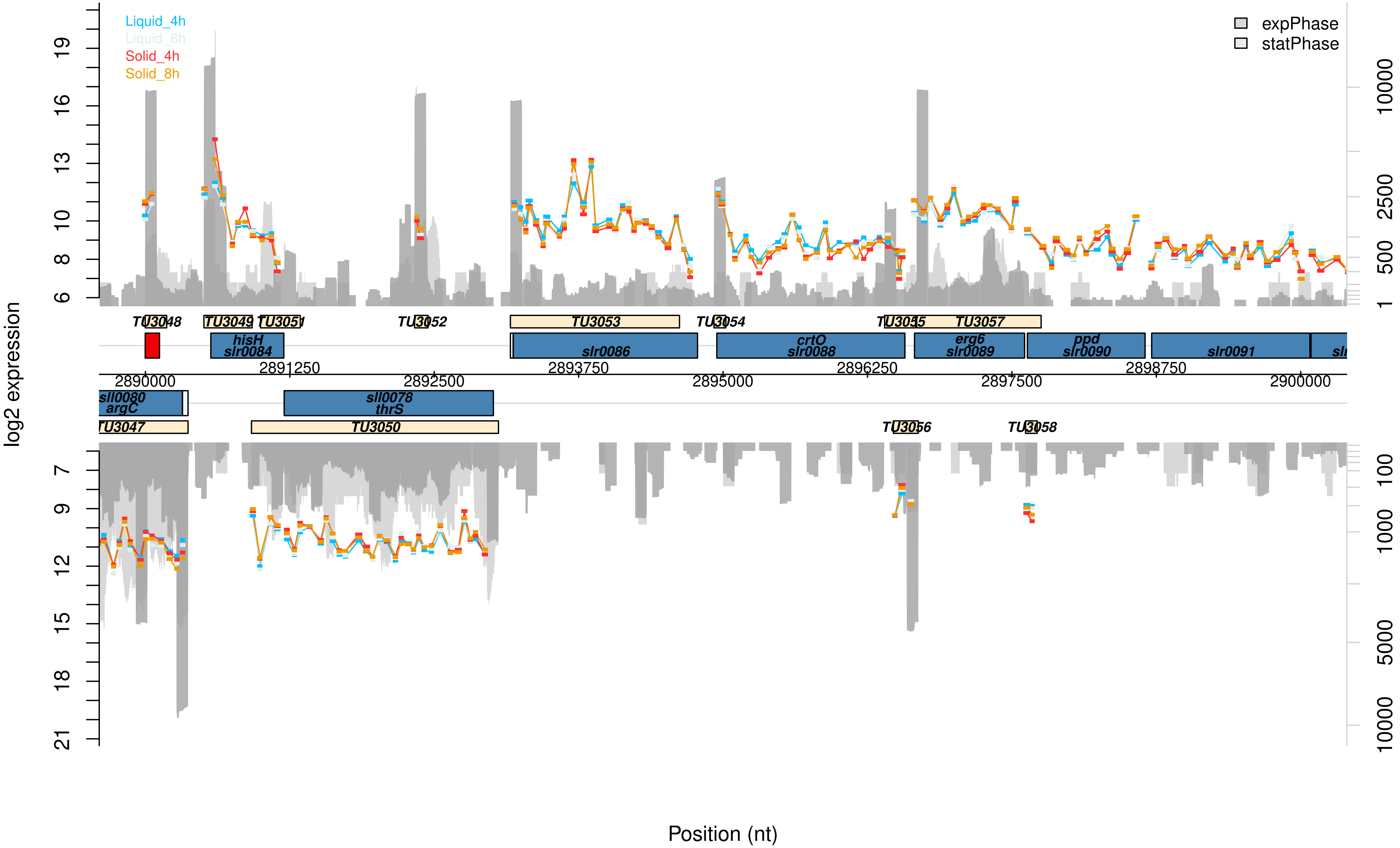

Liquid_4h
Liquid_8h
Solid_4h
Solid_8h
expPhase
statPhase
log2 expression
Position (nt)
TU3048
TU3049
TU3051
TU3052
TU3053
TU3054
TU3055
TU3057
hisH
slr0084
slr0086
crtO
slr0088
erg6
slr0089
ppd
slr0090
slr0091
2890000
2891250
2892500
2893750
2895000
2896250
2897500
2898750
2900000
sll0080
argC
sll0078
thrS
TU3047
TU3050
TU3056
TU3058

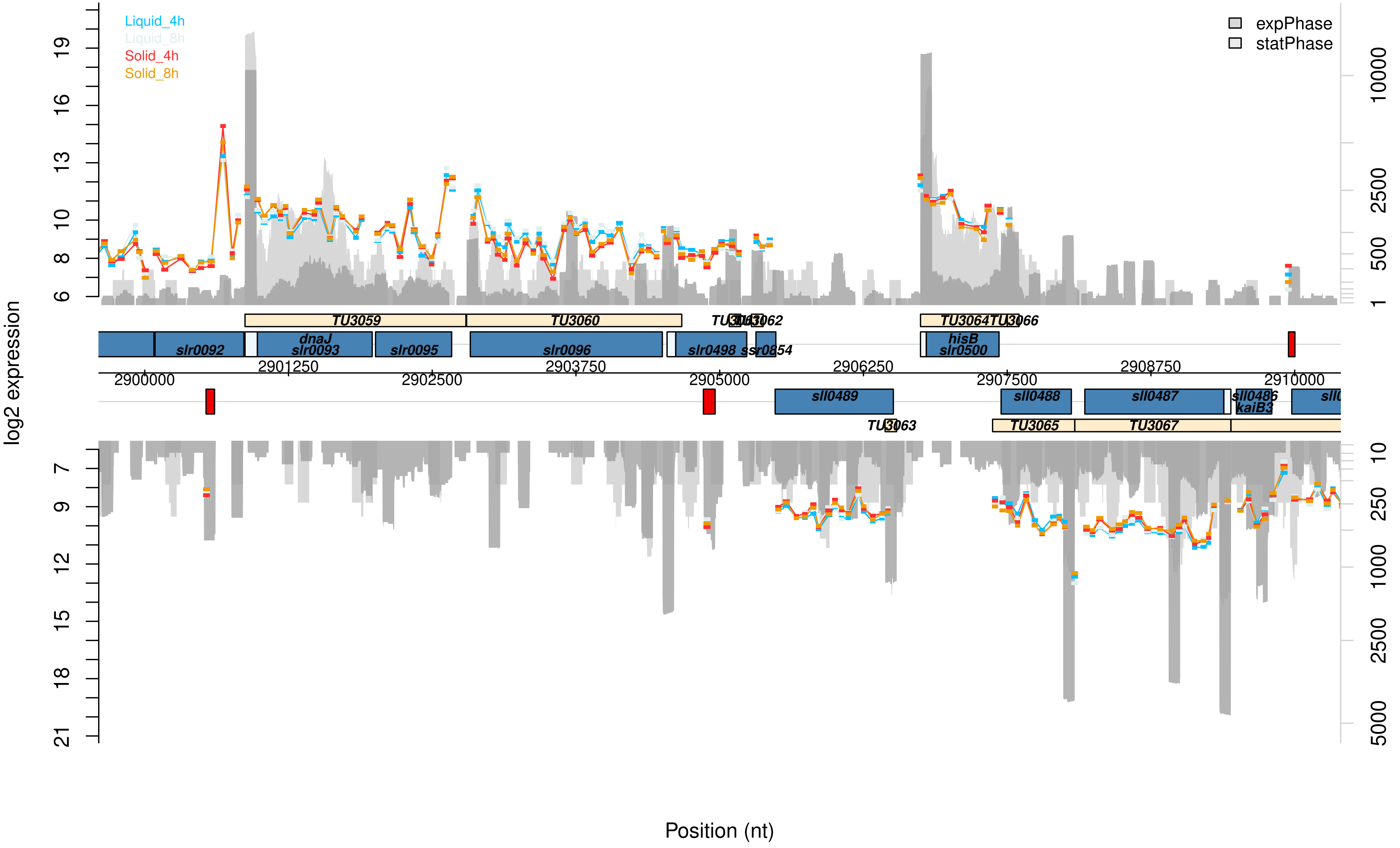

Liquid_4h
Liquid_8h
Solid_4h
Solid_8h
expPhase
statPhase
log2 expression
Position (nt)
TU3059
TU3060
TU3062
TU3064
TU3066
TU3063
TU3065
TU3067
slr0092
dnaJ
slr0093
slr0095
slr0096
slr0498
ssr0854
hisB
slr0500
sll0489
sll0488
sll0487
sll0486
kaiB3
2900000
2901250
2902500
2903750
2905000
2906250
2907500
2908750
2910000

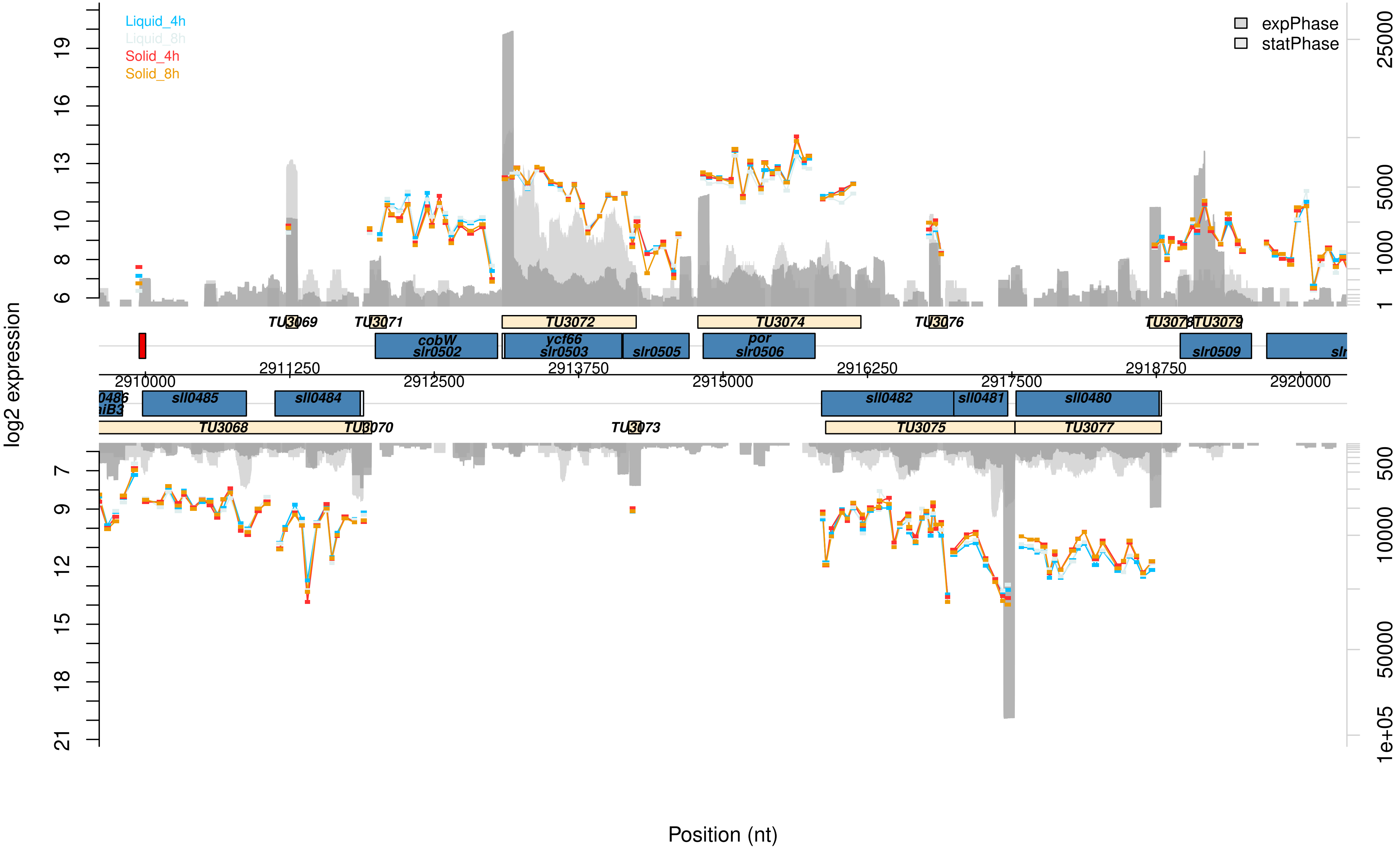

Liquid_4h
Liquid_8h
Solid_4h
Solid_8h
expPhase
statPhase
log2 expression
Position (nt)
TU3068
TU3069
TU3070
TU3071
TU3072
TU3073
TU3074
TU3075
TU3076
TU3077
TU3078
TU3079
cobW
slr0502
ycf66
slr0503
slr0505
por
slr0506
slr0509
sll0485
sll0484
sll0482
sll0481
sll0480
2910000
2911250
2912500
2913750
2915000
2916250
2917500
2918750
2920000

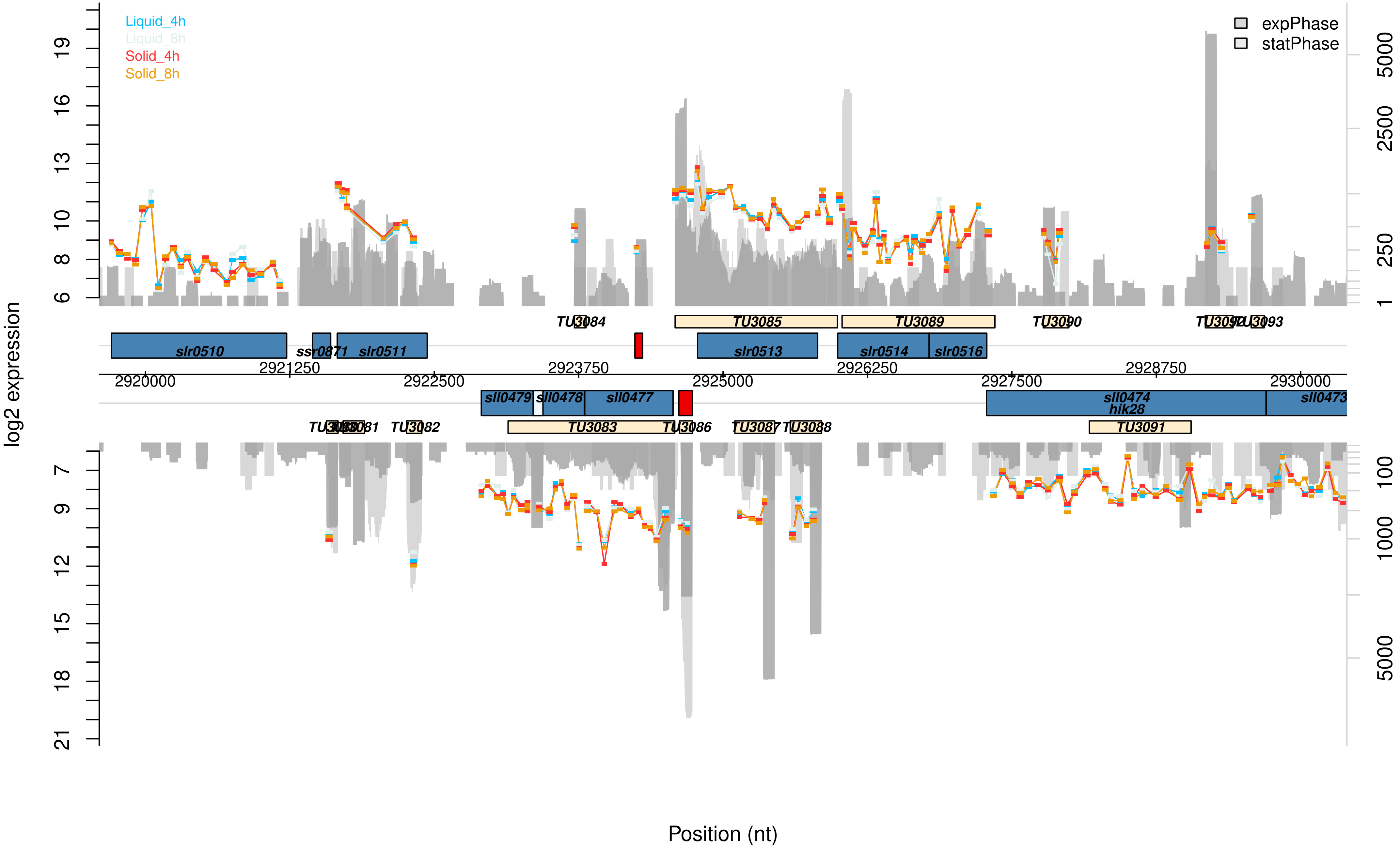

Liquid_4h
Liquid_8h
Solid_4h
Solid_8h
expPhase
statPhase
log2 expression
Position (nt)
TU3084
TU3085
TU3089
TU3090
TU3093
slr0510
ssr0871
slr0511
slr0513
slr0514
slr0516
2920000
2921250
2922500
2923750
2925000
2926250
2927500
2928750
2930000
sll0479
sll0478
sll0477
sll0474
hik28
sll0473
TU3082
TU3083
TU3086
TU3088
TU3091

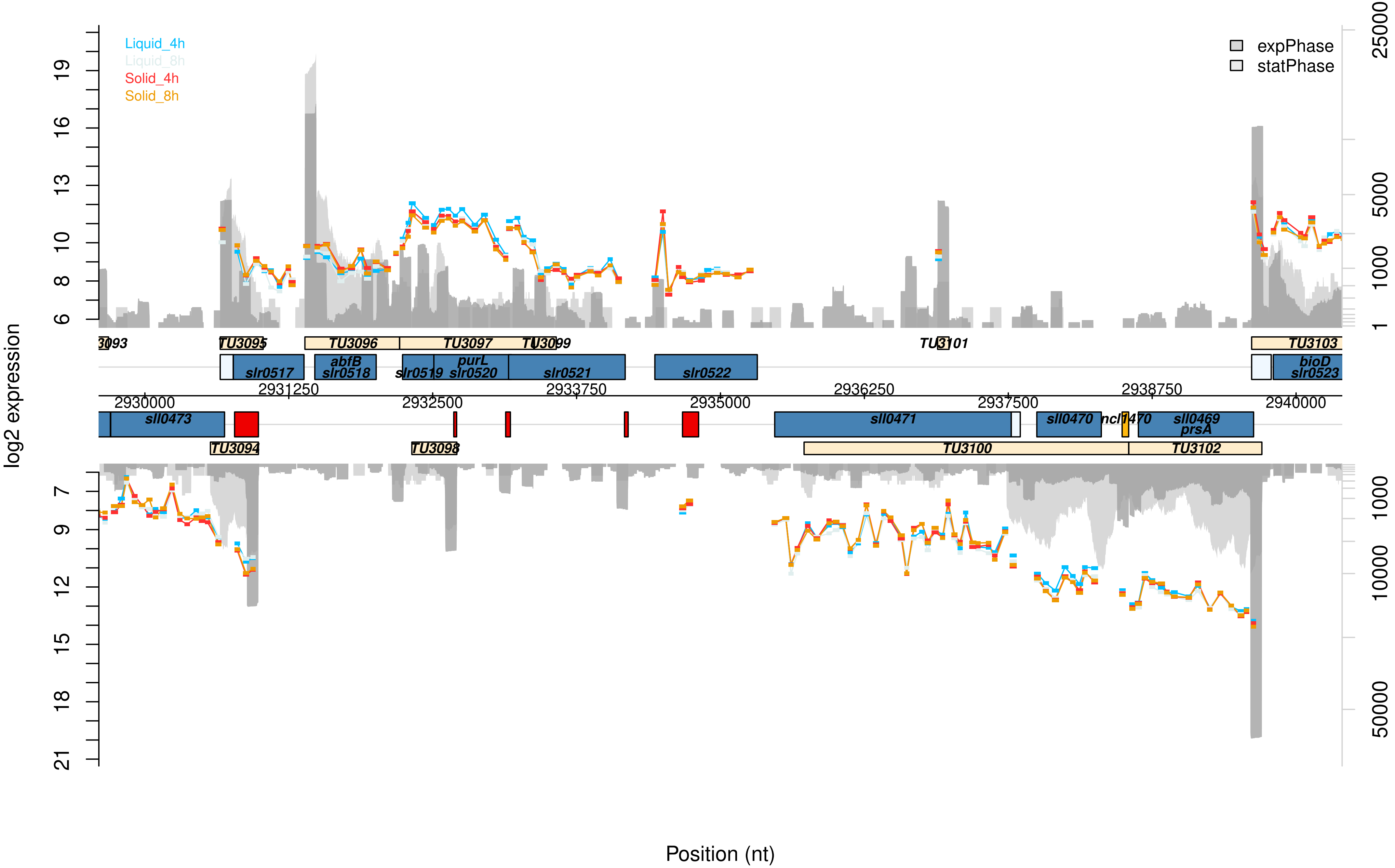

Liquid_4h
Liquid_8h
Solid_4h
Solid_8h
expPhase
statPhase
log2 expression
Position (nt)
TU3095
TU3096
TU3097
TU3099
TU3101
TU3103
slr0517
abfB
slr0518
slr0519
purL
slr0520
slr0521
slr0522
bioD
slr0523
2930000
2931250
2932500
2933750
2935000
2936250
2937500
2938750
2940000
sll0473
sll0471
sll0470
sll0469
prsA
TU3094
TU3098
TU3100
TU3102

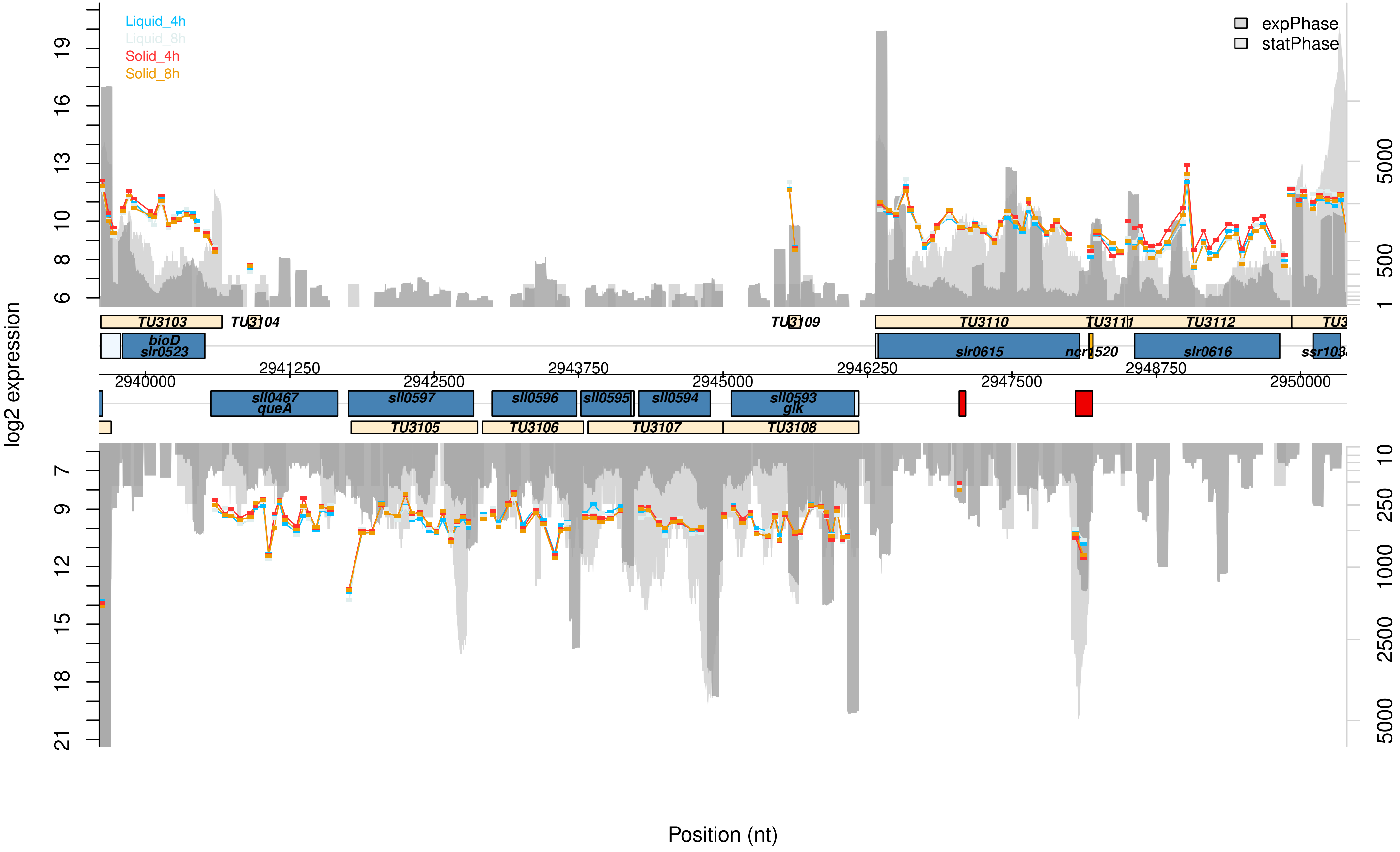

Liquid_4h
Liquid_8h
Solid_4h
Solid_8h
expPhase
statPhase
log2 expression
Position (nt)
TU3103
TU3104
TU3109
TU3110
TU3111
TU3112
bioD
slr0523
slr0615
ncr1520
slr0616
2941250
2943750
2946250
2948750
2940000
2942500
2945000
2947500
2950000
sll0467
queA
sll0597
sll0596
sll0595
sll0594
sll0593
glk
TU3105
TU3106
TU3107
TU3108

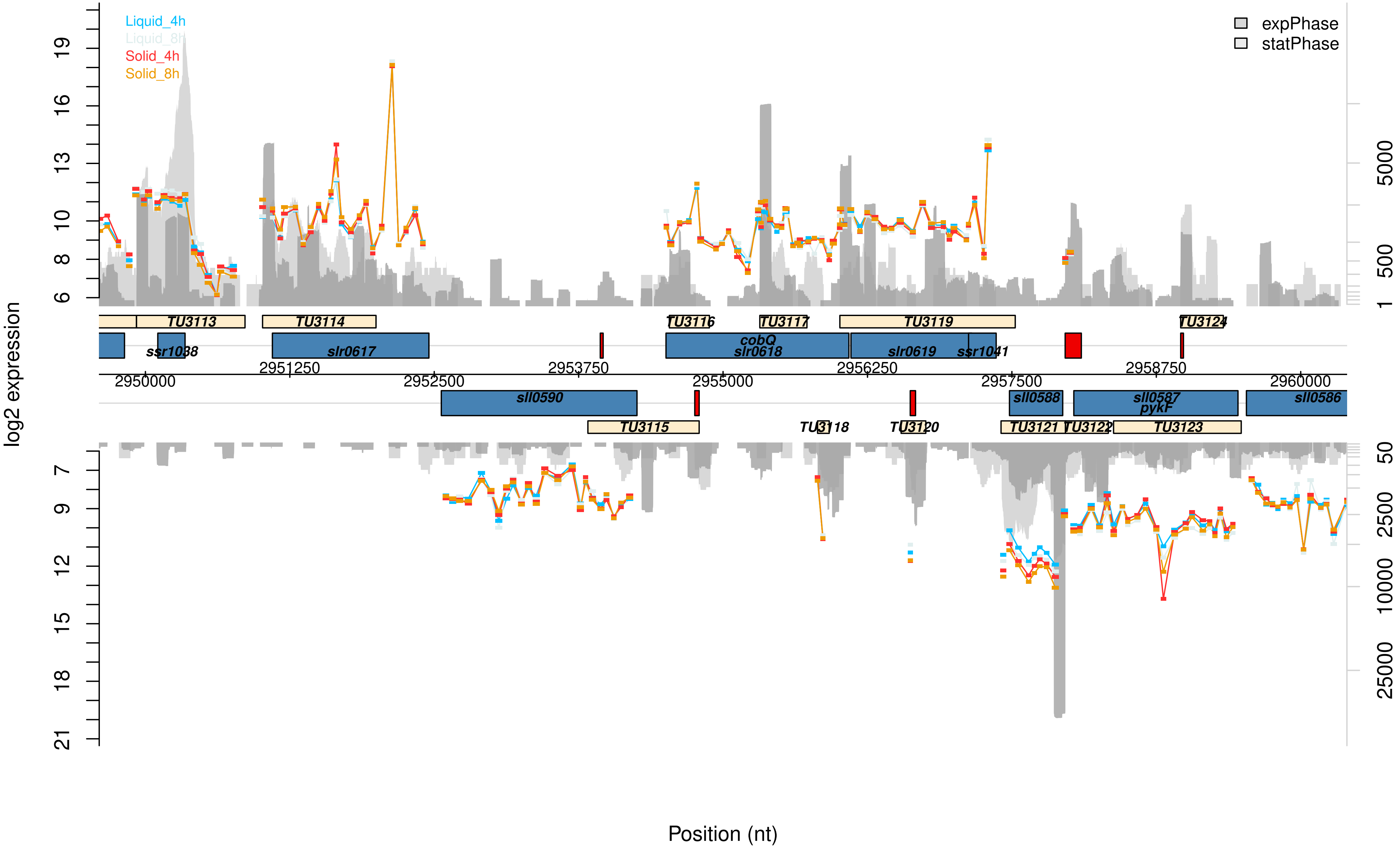
Liquid_4h
Liquid_8h
Solid_4h
Solid_8h
expPhase
statPhase
log2 expression
Position (nt)
TU3113
TU3114
TU3116
TU3117
TU3119
TU3124
ssr1038
slr0617
cobQ
slr0618
slr0619
ssr1041
2951250
2953750
2956250
2958750
2950000
2952500
2955000
2957500
2960000
sll0590
sll0588
sll0587
pykF
sll0586
TU3115
TU3118
TU3120
TU3121
TU3122
TU3123

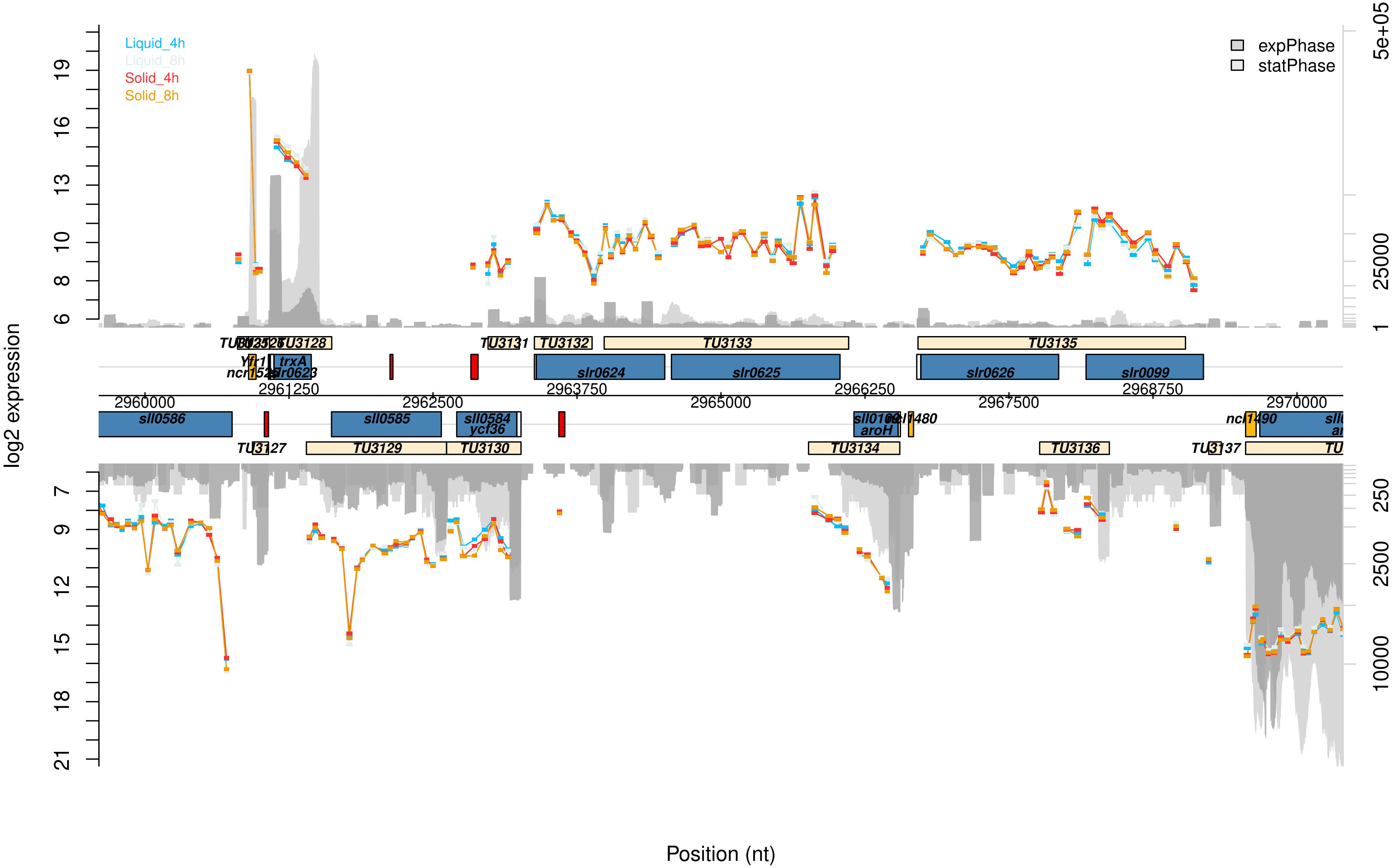

Liquid_4h
Liquid_8h
Solid_4h
Solid_8h
expPhase
statPhase
log2 expression
Position (nt)
TU3128
TU3131
TU3132
TU3133
TU3135
trxA
slr0623
slr0624
slr0625
slr0626
slr0099
sll0586
sll0585
sll0584
ycf36
aroH
TU3127
TU3129
TU3130
TU3134
TU3136
TU3137
2960000
2961250
2962500
2963750
2965000
2966250
2967500
2968750
2970000
5e+05
25000
2500
250
10000

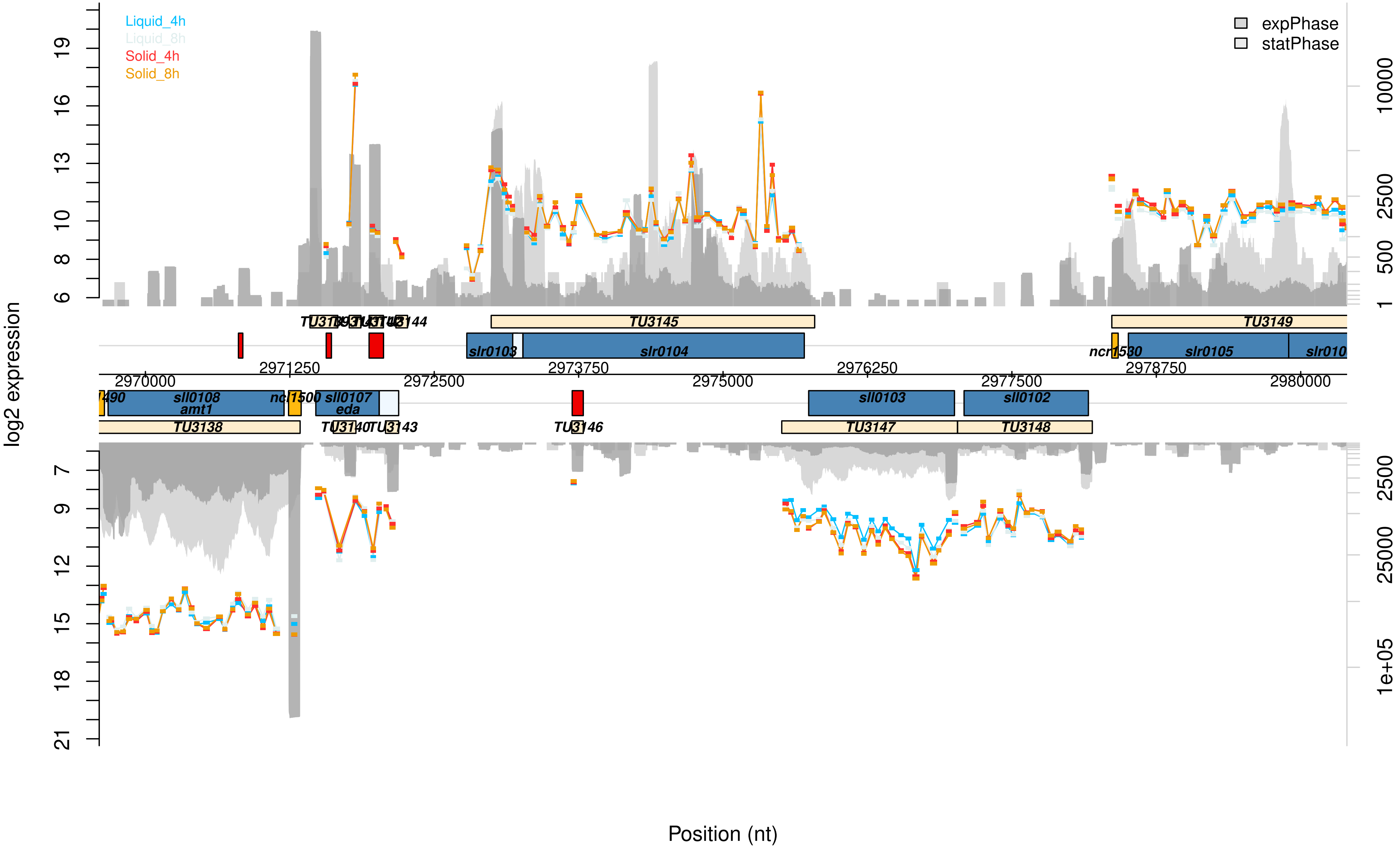

Liquid_4h
Liquid_8h
Solid_4h
Solid_8h
expPhase
statPhase
log2 expression
Position (nt)
TU3145
TU3149
TU3144
slr0103
slr0104
ncr1530
slr0105
slr010
2970000
2971250
2972500
2973750
2975000
2976250
2977500
2978750
2980000
490
sll0108
amt1
ncl1500
sll0107
eda
sll0103
sll0102
TU3138
TU3143
TU3146
TU3147
TU3148

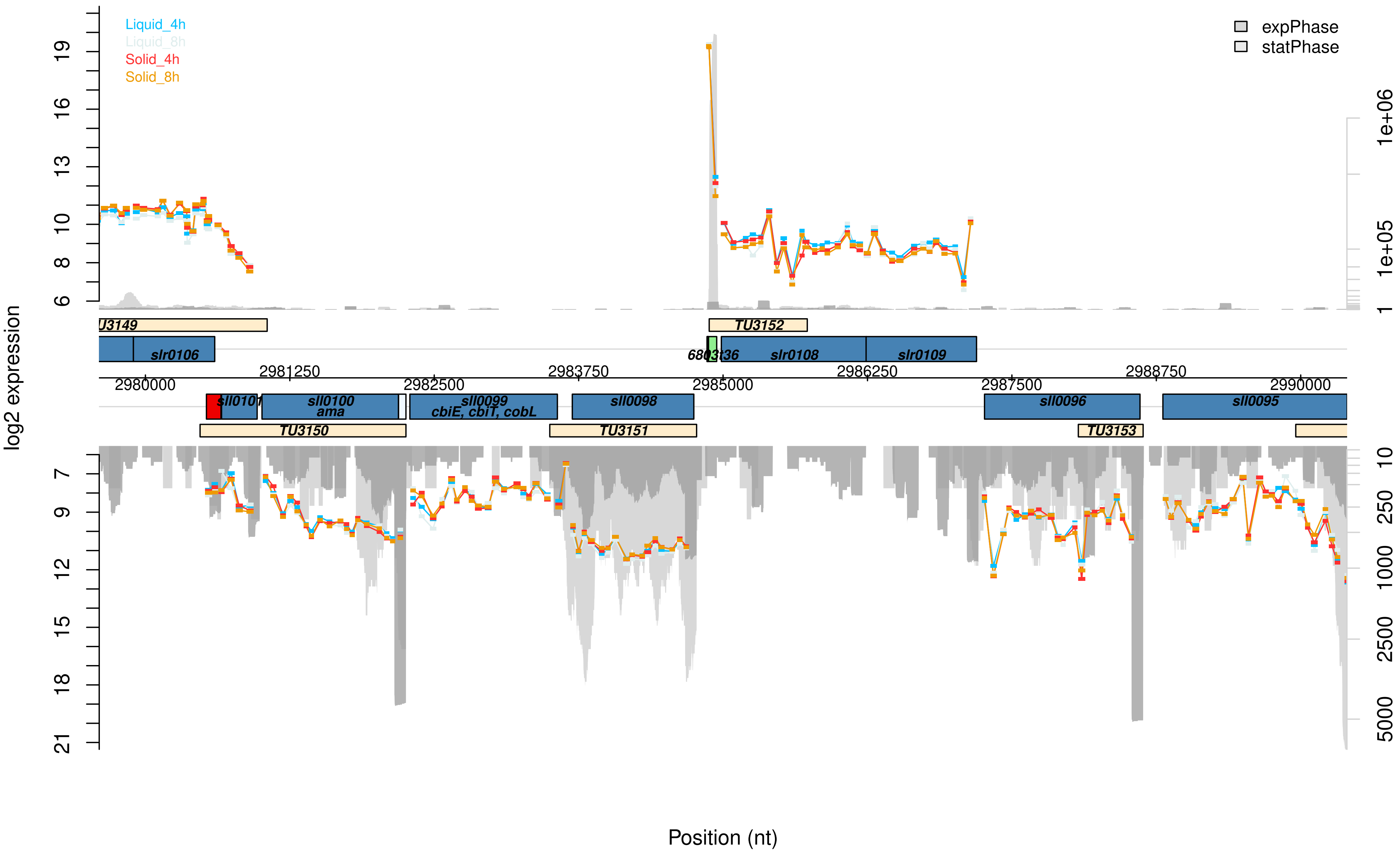

Liquid_4h
Liquid_8h
Solid_4h
Solid_8h
expPhase
statPhase
log2 expression
Position (nt)
U3149
TU3152
slr0106
6803t36
slr0108
slr0109
2980000
2981250
2982500
2983750
2985000
2986250
2987500
2988750
2990000
sll0101
sll0100
ama
sll0099
cbiE, cbiT, cobL
sll0098
sll0096
sll0095
TU3150
TU3151
TU3153

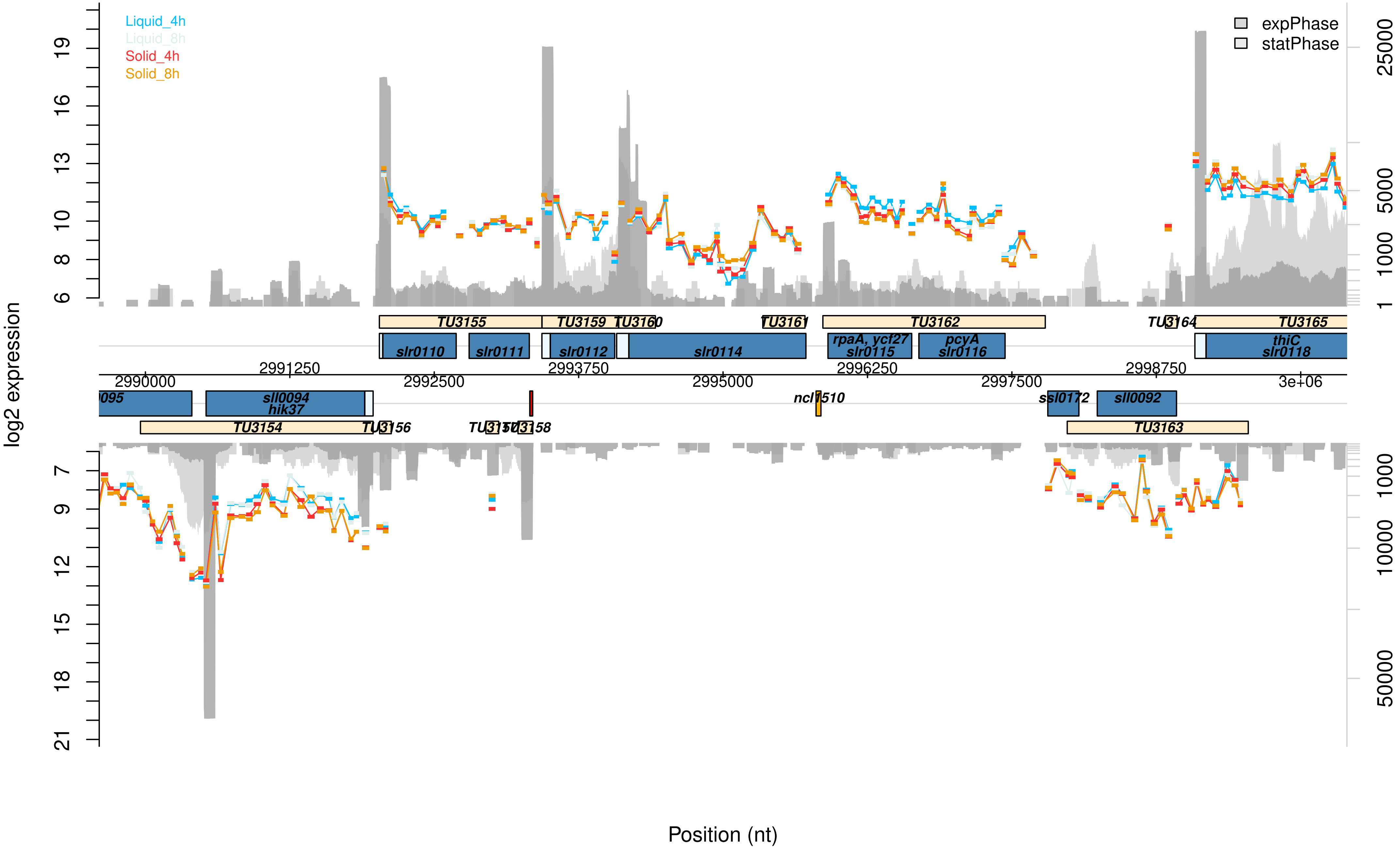

Liquid_4h
Liquid_8h
Solid_4h
Solid_8h
expPhase
statPhase
log2 expression
Position (nt)
TU3155
TU3159
TU3160
TU3161
TU3162
TU3164
TU3165
slr0110
slr0111
slr0112
slr0114
rpaA, ycf27
slr0115
pcyA
slr0116
thiC
slr0118
sll0094
hik37
ncl1510
ssl0172
sll0092
TU3154
TU3156
TU3158
TU3163
2990000
2991250
2992500
2993750
2995000
2996250
2997500
2998750
3e+06

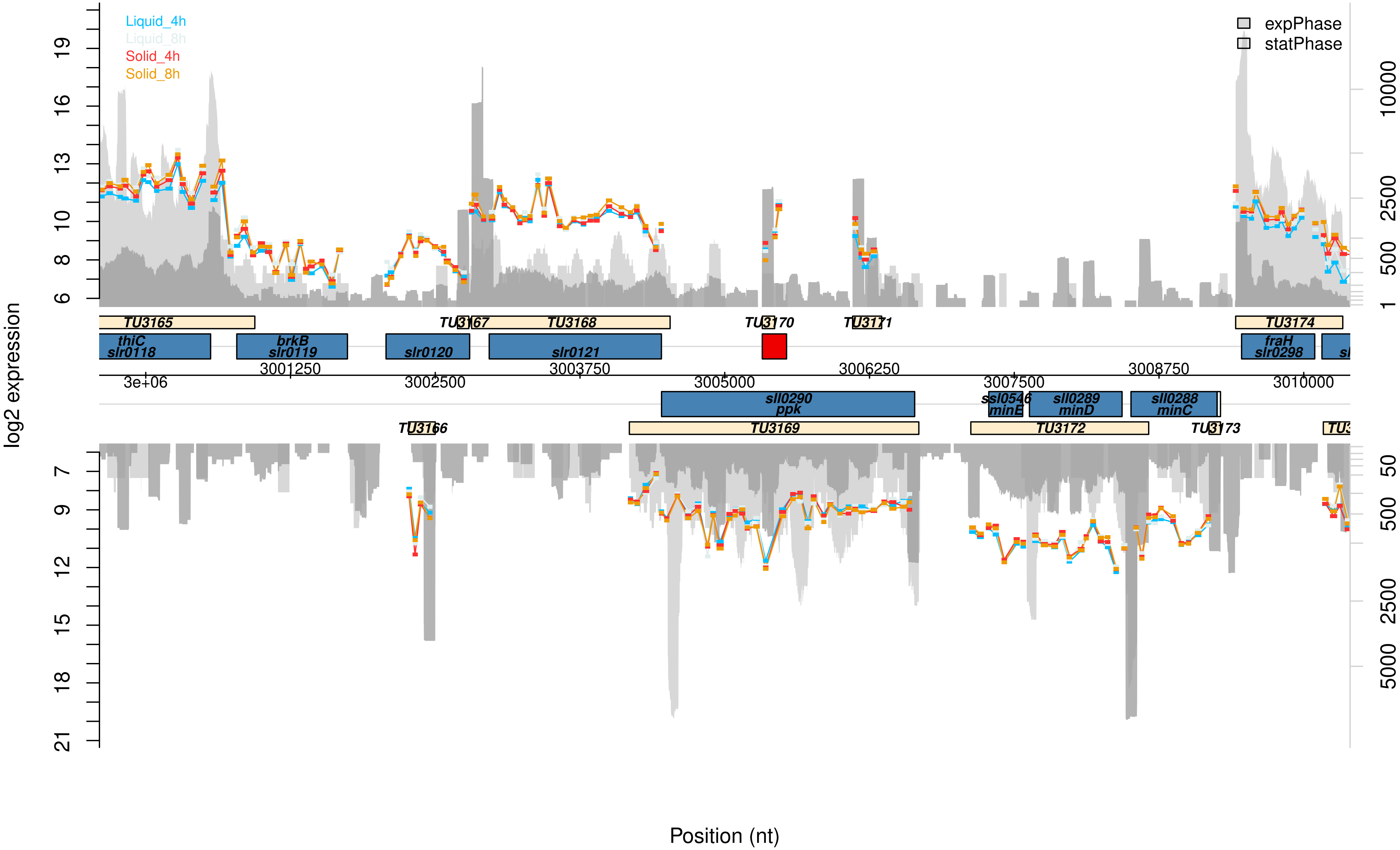

Liquid_4h
Liquid_8h
Solid_4h
Solid_8h
expPhase
statPhase
log2 expression
Position (nt)
TU3165
TU3166
TU3167
TU3168
TU3169
TU3170
TU3171
TU3172
TU3173
TU3174
thiC
slr0118
brkB
slr0119
slr0120
slr0121
sll0290
ppk
ssl0546
minE
sll0289
minD
sll0288
minC
fraH
slr0298
3e+06
3001250
3002500
3003750
3005000
3006250
3007500
3008750
3010000

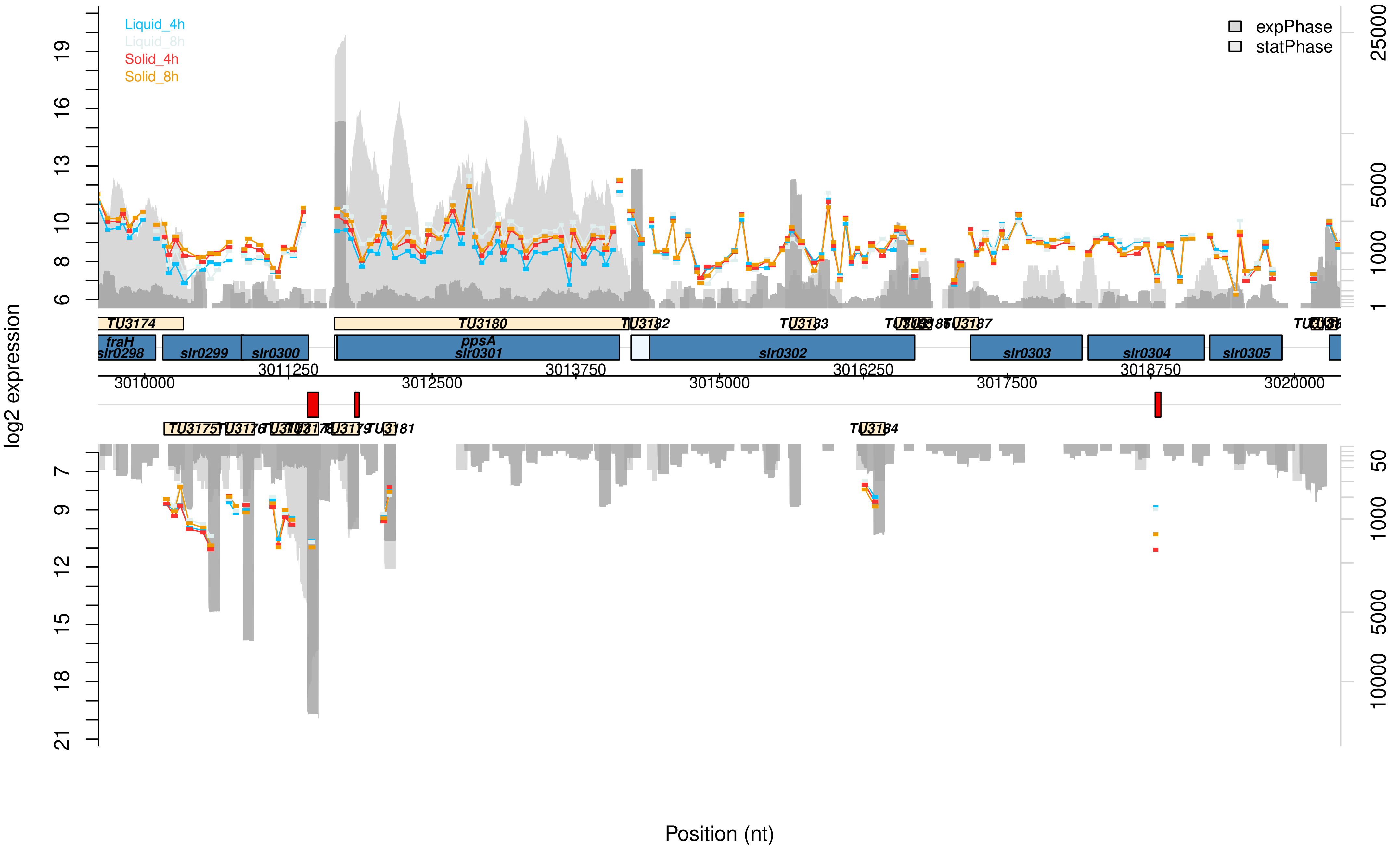

Liquid_4h
Liquid_8h
Solid_4h
Solid_8h
expPhase
statPhase
log2 expression
Position (nt)
TU3174
TU3180
TU3182
TU3183
TU3187
fraH
slr0298
slr0299
slr0300
ppsA
slr0301
slr0302
slr0303
slr0304
slr0305
3010000
3011250
3012500
3013750
3015000
3016250
3017500
3018750
3020000
TU3175
TU3181
TU3184

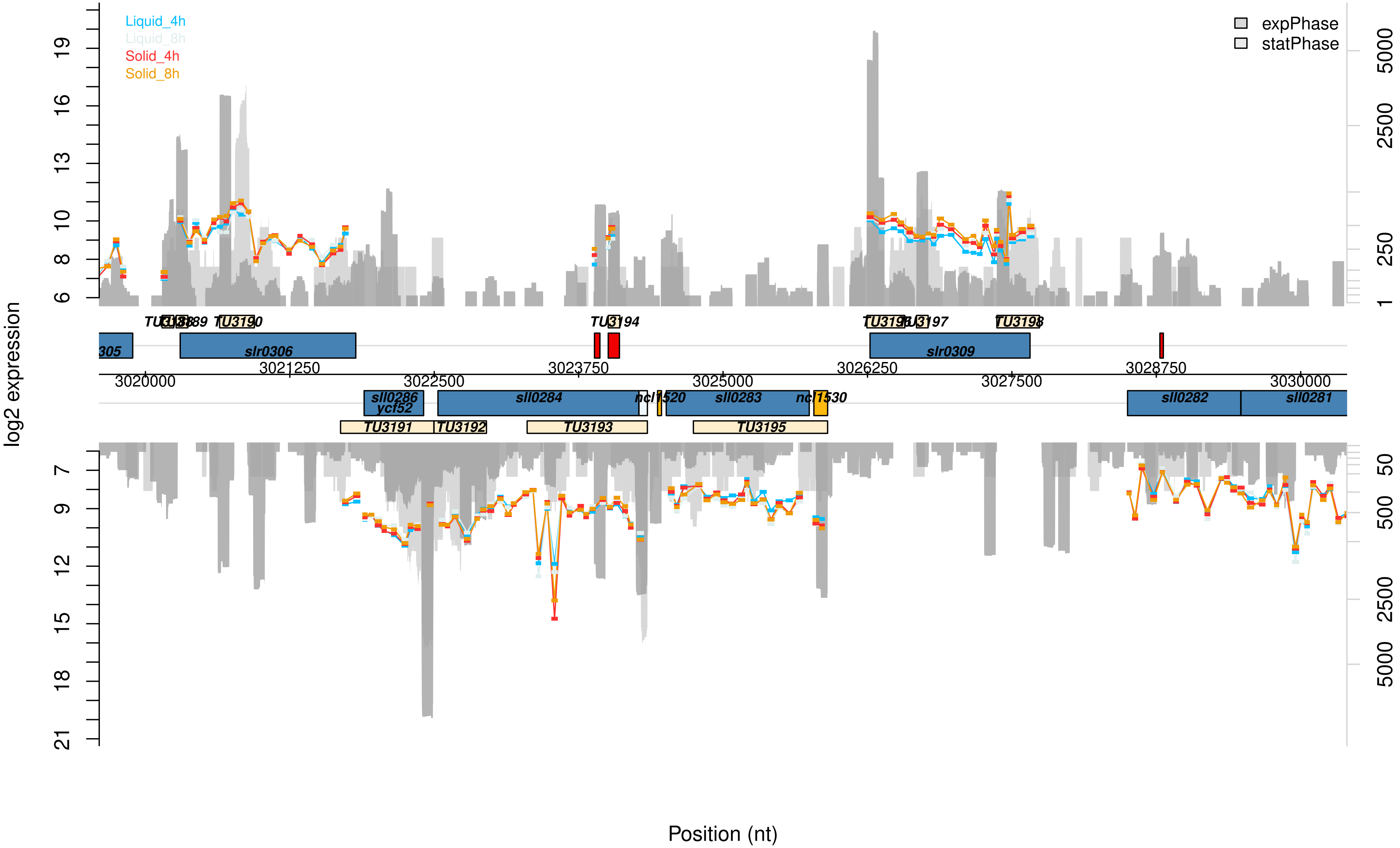
Liquid_4h
Liquid_8h
Solid_4h
Solid_8h
expPhase
statPhase
log2 expression
Position (nt)
19
16
13
10
8
6
7
9
12
15
18
21
5000
2500
250
1
50
500
2500
5000
3020000
3021250
3022500
3023750
3025000
3026250
3027500
3028750
3030000
305
slr0306
slr0309
sll0286
ycf52
sll0284
ncl1520
sll0283
ncl1530
sll0282
sll0281
TU3194
TU3197
TU3198
TU3190
TU3191
TU3192
TU3193
TU3195

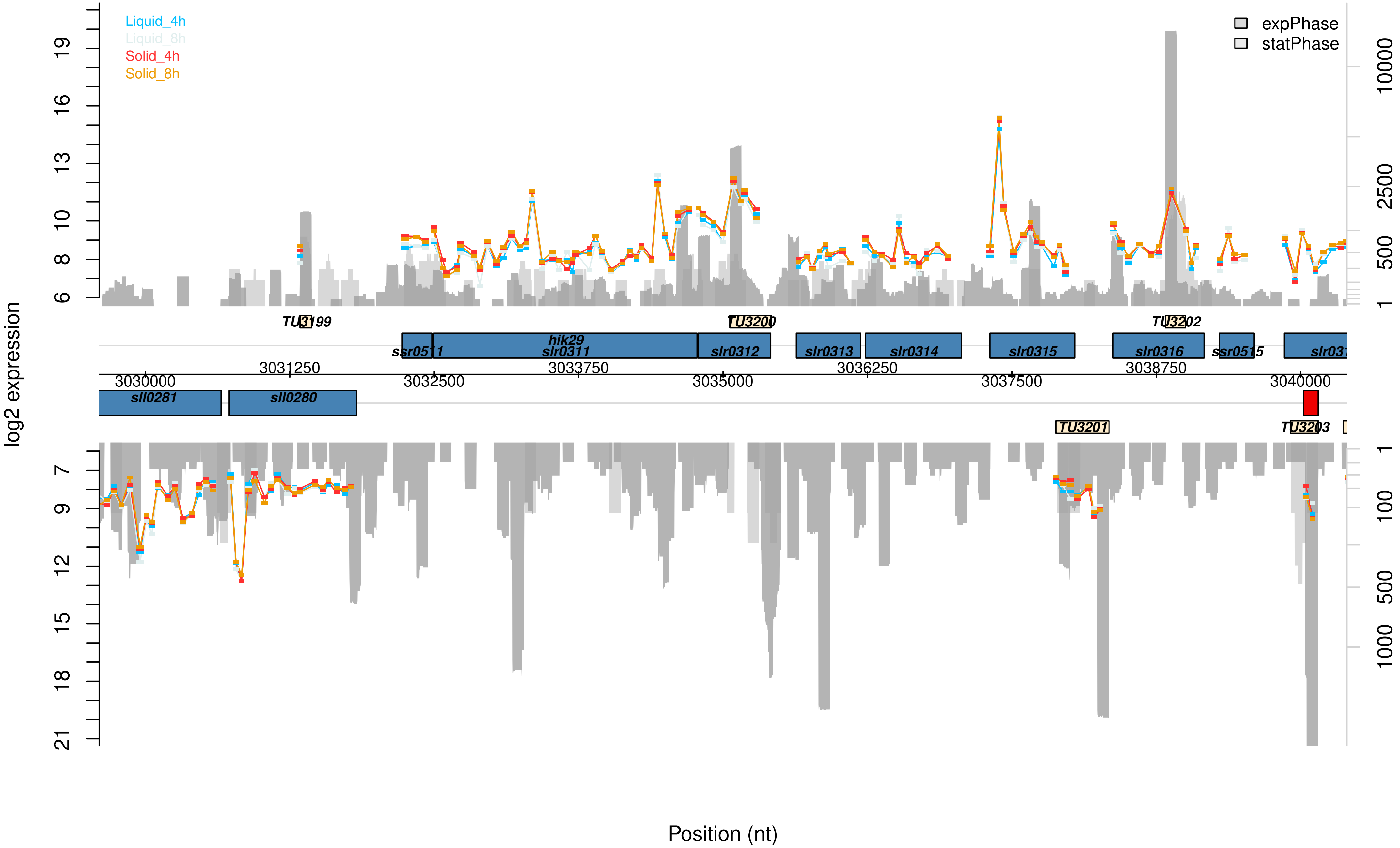
Liquid_4h
Liquid_8h
Solid_4h
Solid_8h
expPhase
statPhase
log2 expression
TU3199
TU3200
TU3202
TU3201
TU3203
ssr0511
hik29
slr0311
slr0312
slr0313
slr0314
slr0315
slr0316
ssr0515
sll0281
sll0280
3030000
3031250
3032500
3033750
3035000
3036250
3037500
3038750
3040000
Position (nt)

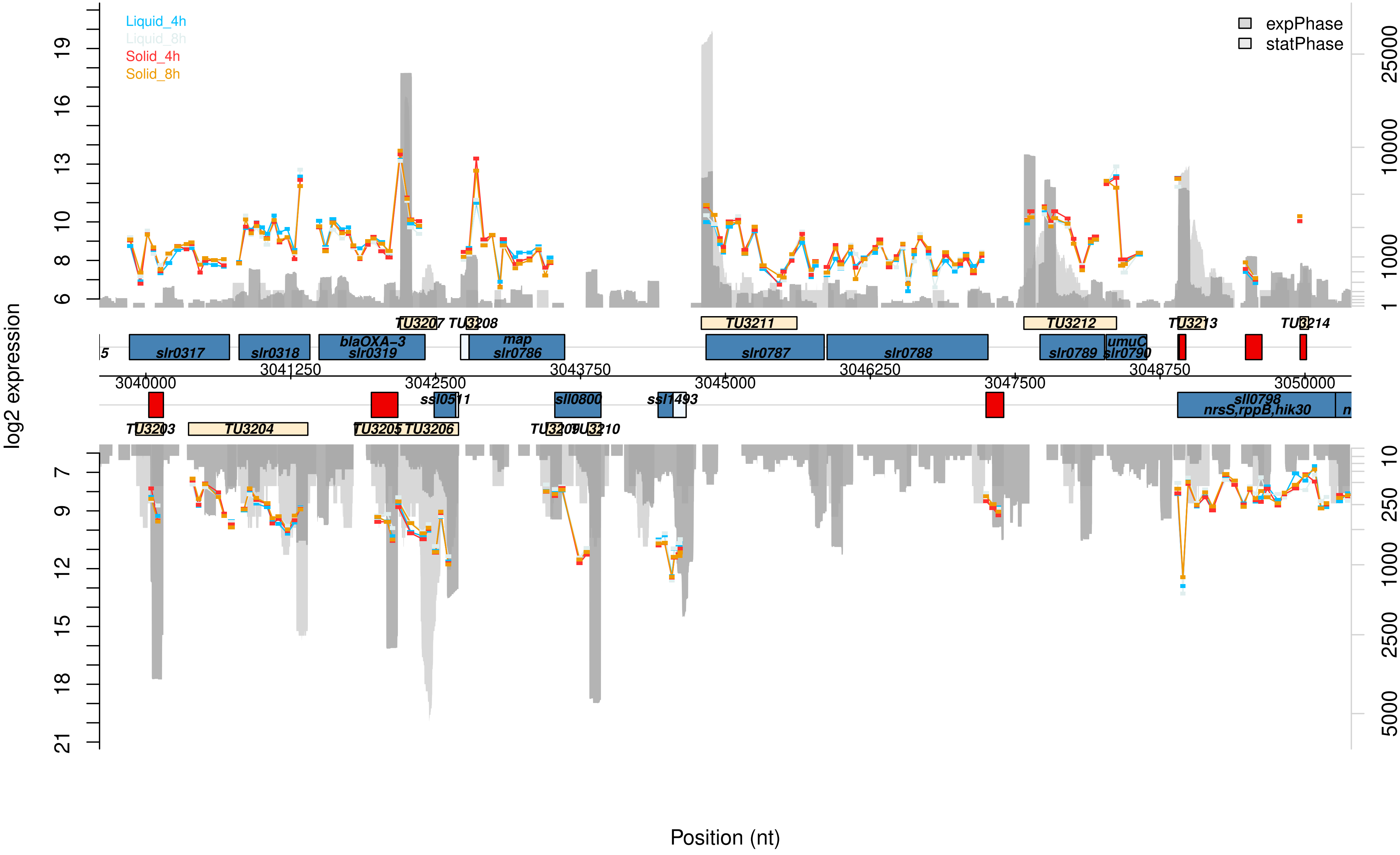

Liquid_4h
Liquid_8h
Solid_4h
Solid_8h
expPhase
statPhase
log2 expression
19
16
13
10
8
6
7
9
12
15
18
21
25000
10000
1000
1
10
250
1000
2500
5000
TU3207
TU3208
TU3211
TU3212
TU3213
TU3214
slr0317
slr0318
blaOXA-3
slr0319
map
slr0786
slr0787
slr0788
slr0789
umuC
slr0790
3041250
3043750
3046250
3048750
3040000
3042500
3045000
3047500
3050000
ssl0511
sll0800
ssl1493
sll0798
nrsS,rppB,hik30
TU3203
TU3204
TU3205
TU3206
TU3209
TU3210
Position (nt)

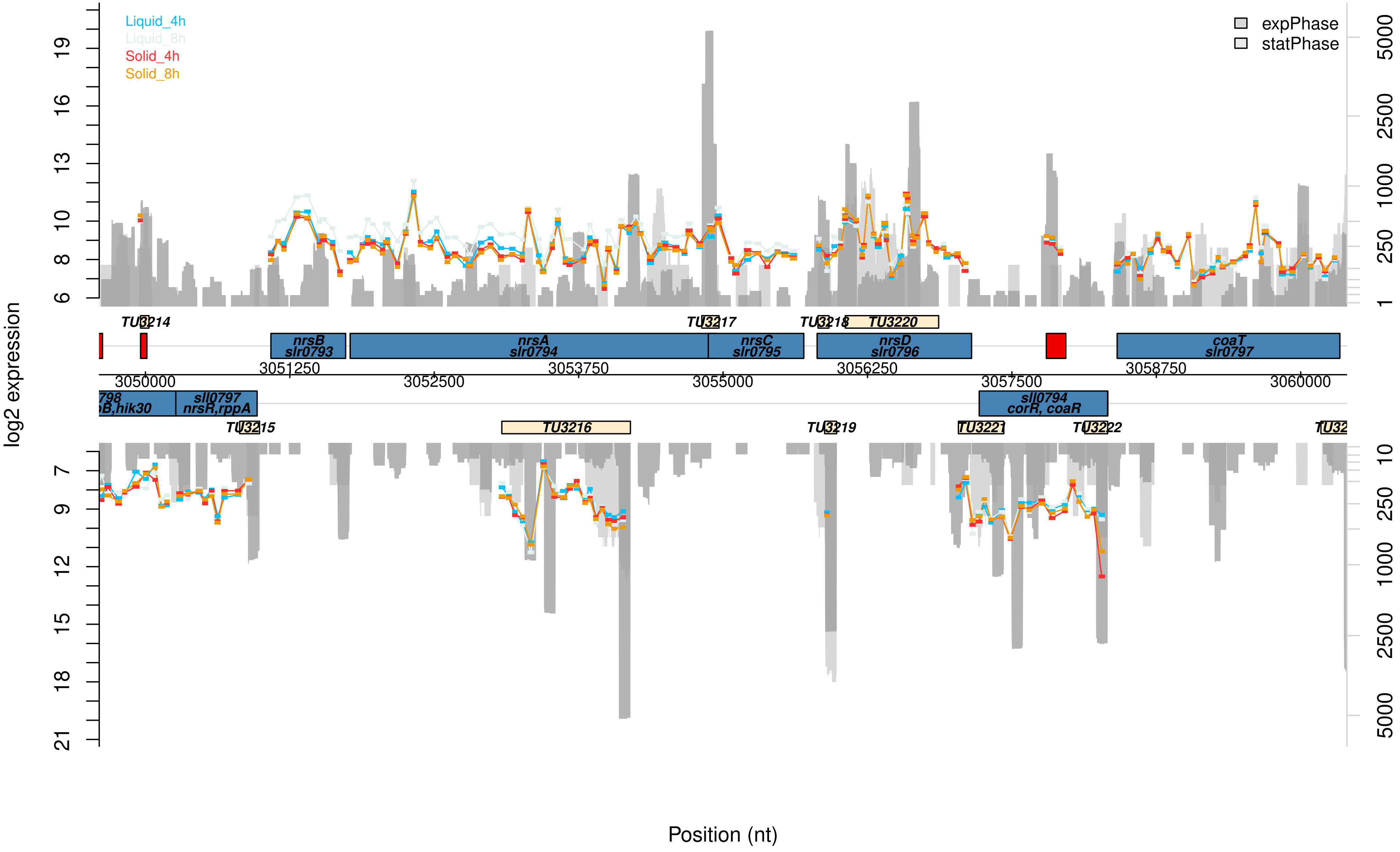

Liquid_4h
Liquid_8h
Solid_4h
Solid_8h
expPhase
statPhase
log2 expression
TU3214
TU3217
TU3218
TU3220
nrsB slr0793
nrsA slr0794
nrsC slr0795
nrsD slr0796
coaT slr0797
3051250
3053750
3056250
3058750
3050000
3052500
3055000
3057500
3060000
sll0797 nrsR,rppA
sll0794 corR, coaR
TU3215
TU3216
TU3219
TU3221
TU3222
Position (nt)

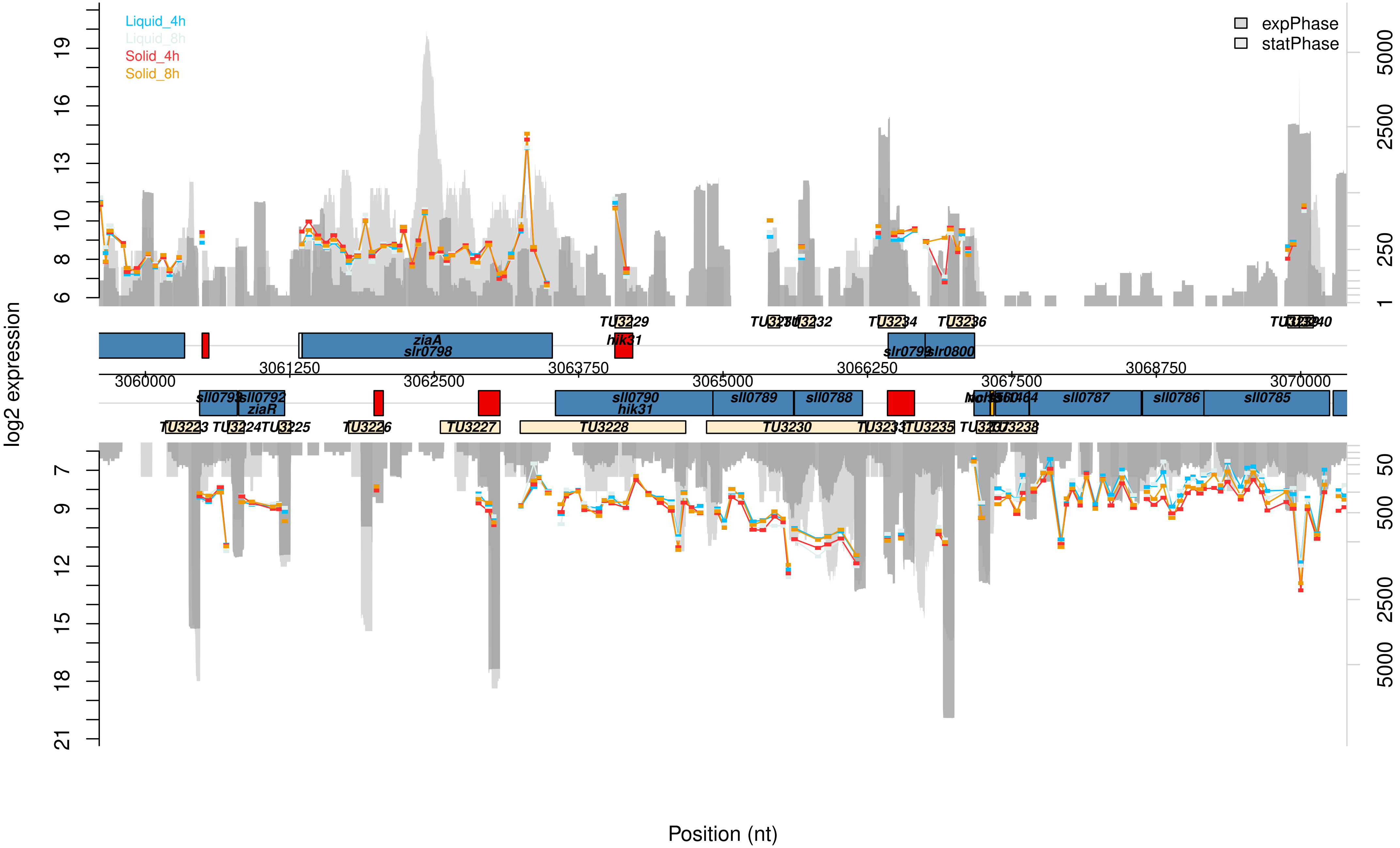
Liquid_4h
Liquid_8h
Solid_4h
Solid_8h
expPhase
statPhase
log2 expression
Position (nt)
ziaA
slr0798
hik31
slr0799
slr0800
sll0793
sll0792
ziaR
sll0790
hik31
sll0789
sll0788
sll0787
sll0786
sll0785
TU3223
TU3224
TU3225
TU3226
TU3227
TU3228
TU3229
TU3230
TU3232
TU3234
TU3235
TU3236
TU3238
3060000
3061250
3062500
3063750
3065000
3066250
3067500
3068750
3070000

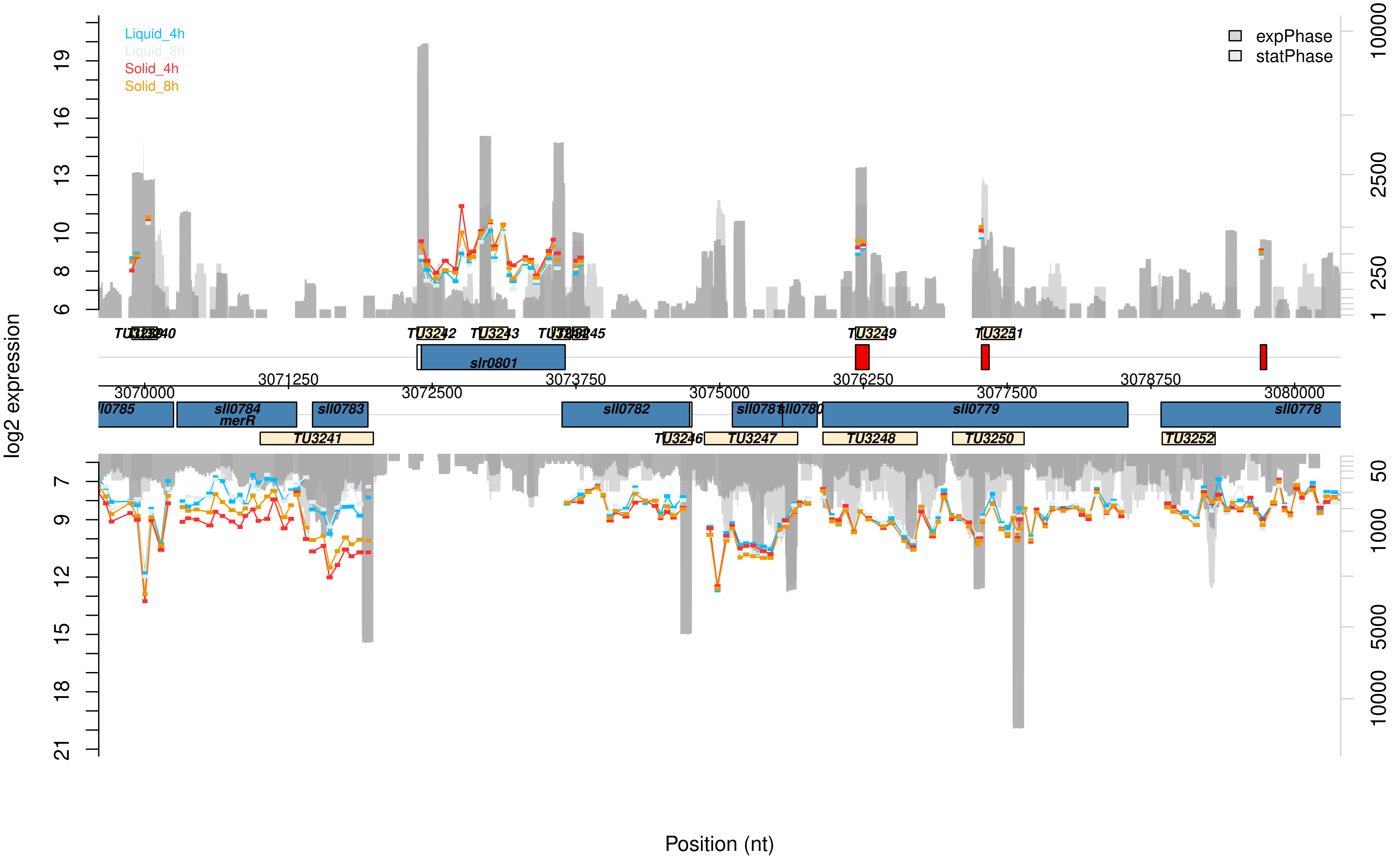

Liquid_4h
Liquid_8h
Solid_4h
Solid_8h
expPhase
statPhase
log2 expression
Position (nt)
slr0801
sll0785
sll0784
merR
sll0783
sll0782
sll0781
sll0780
sll0779
sll0778
TU3241
TU3242
TU3243
TU3245
TU3246
TU3247
TU3248
TU3249
TU3250
TU3251
TU3252
3070000
3071250
3072500
3073750
3075000
3076250
3077500
3078750
3080000

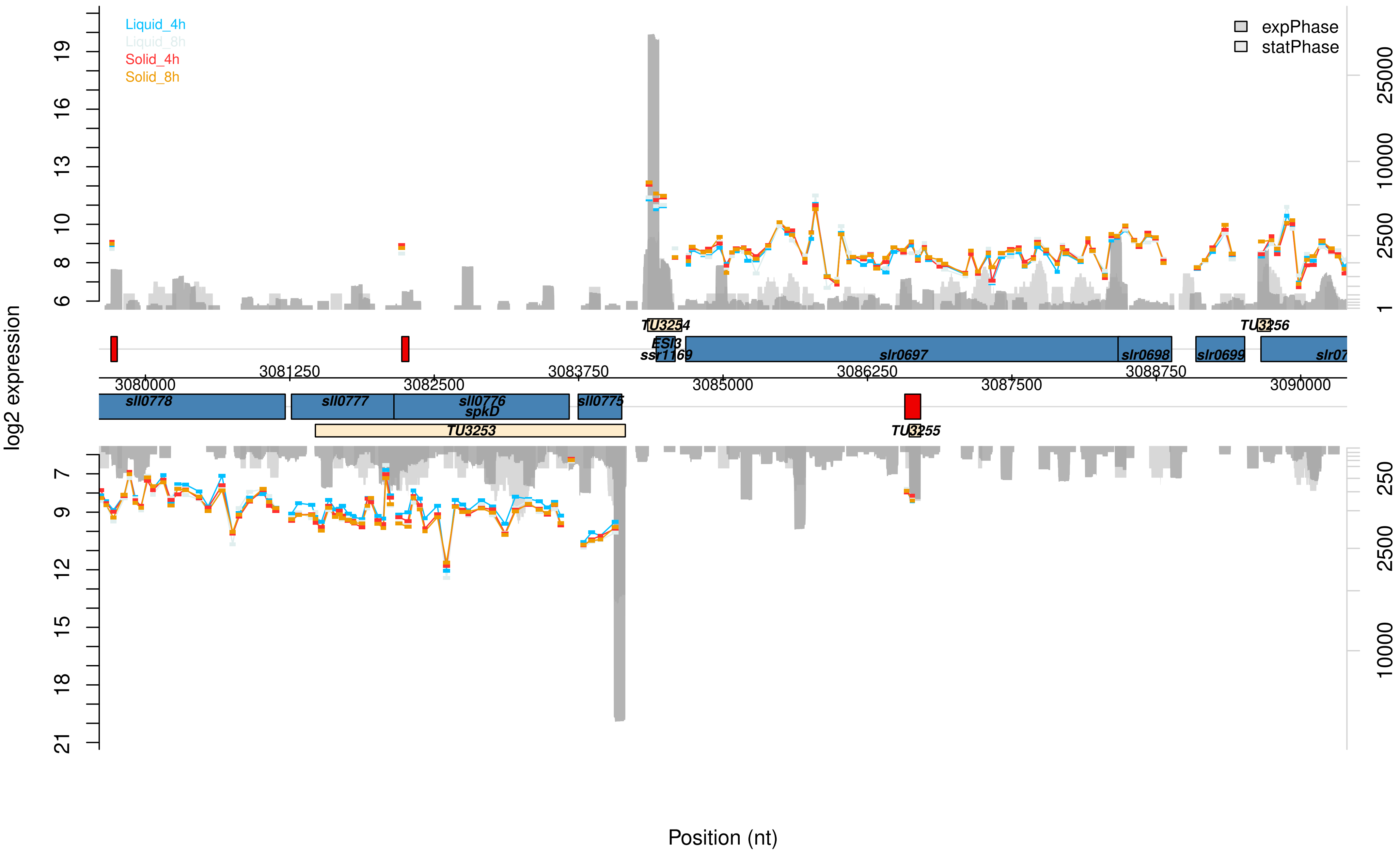

Liquid_4h
Liquid_8h
Solid_4h
Solid_8h
expPhase
statPhase
log2 expression
Position (nt)
TU3254
ESI3
ssr1169
slr0697
slr0698
slr0699
slr07
TU3256
3080000
3081250
3082500
3083750
3085000
3086250
3087500
3088750
3090000
sll0778
sll0777
sll0776
spkD
sll0775
TU3253
TU3255

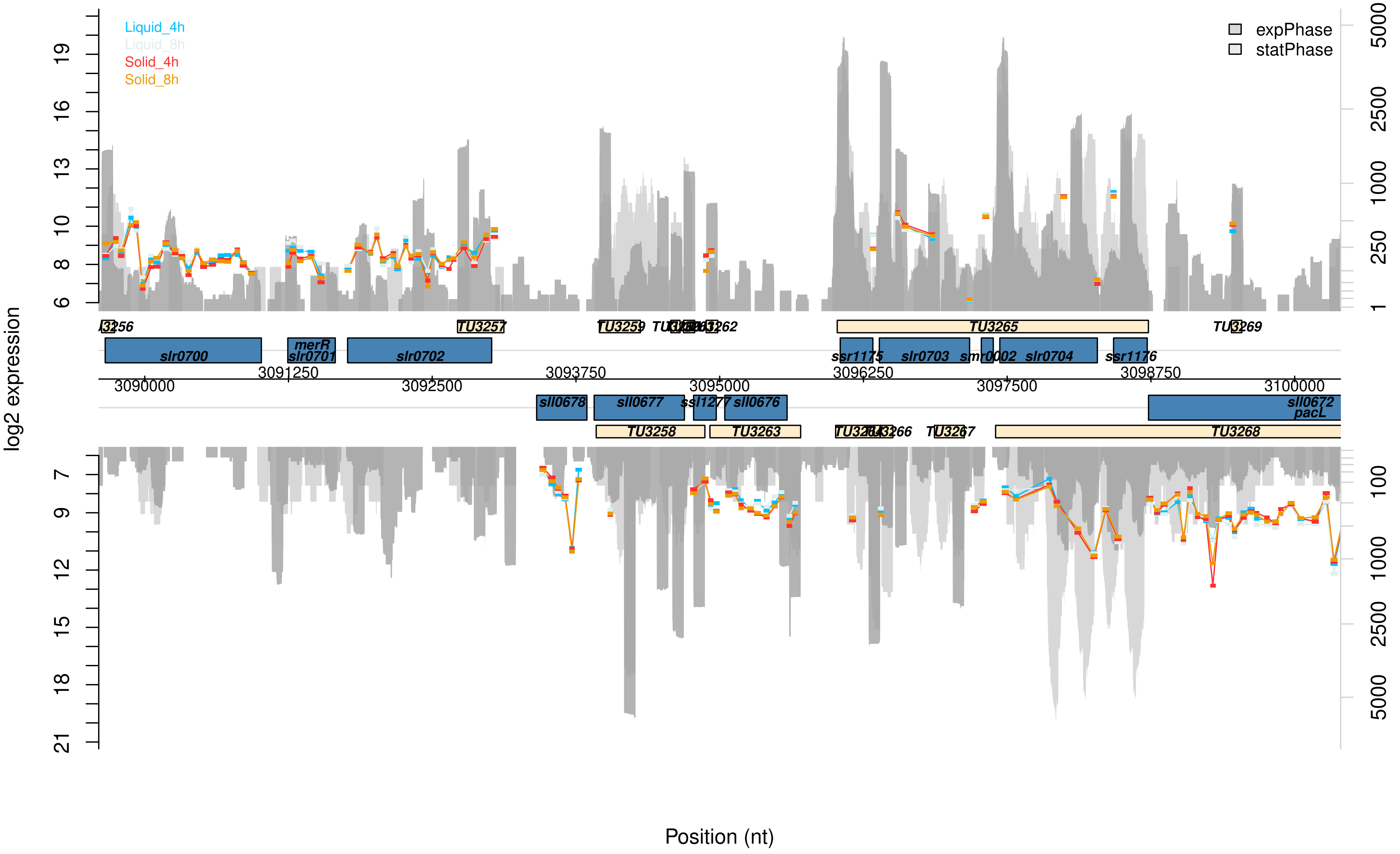

Liquid_4h
Liquid_8h
Solid_4h
Solid_8h
expPhase
statPhase
log2 expression
Position (nt)
slr0700
merR
slr0701
slr0702
TU3257
TU3259
TU3265
TU3269
ssr1175
slr0703
smr0002
slr0704
ssr1176
3090000
3091250
3092500
3093750
3095000
3096250
3097500
3098750
3100000
sll0678
sll0677
ssl1277
sll0676
sll0672
pacL
TU3258
TU3263
TU3267
TU3268

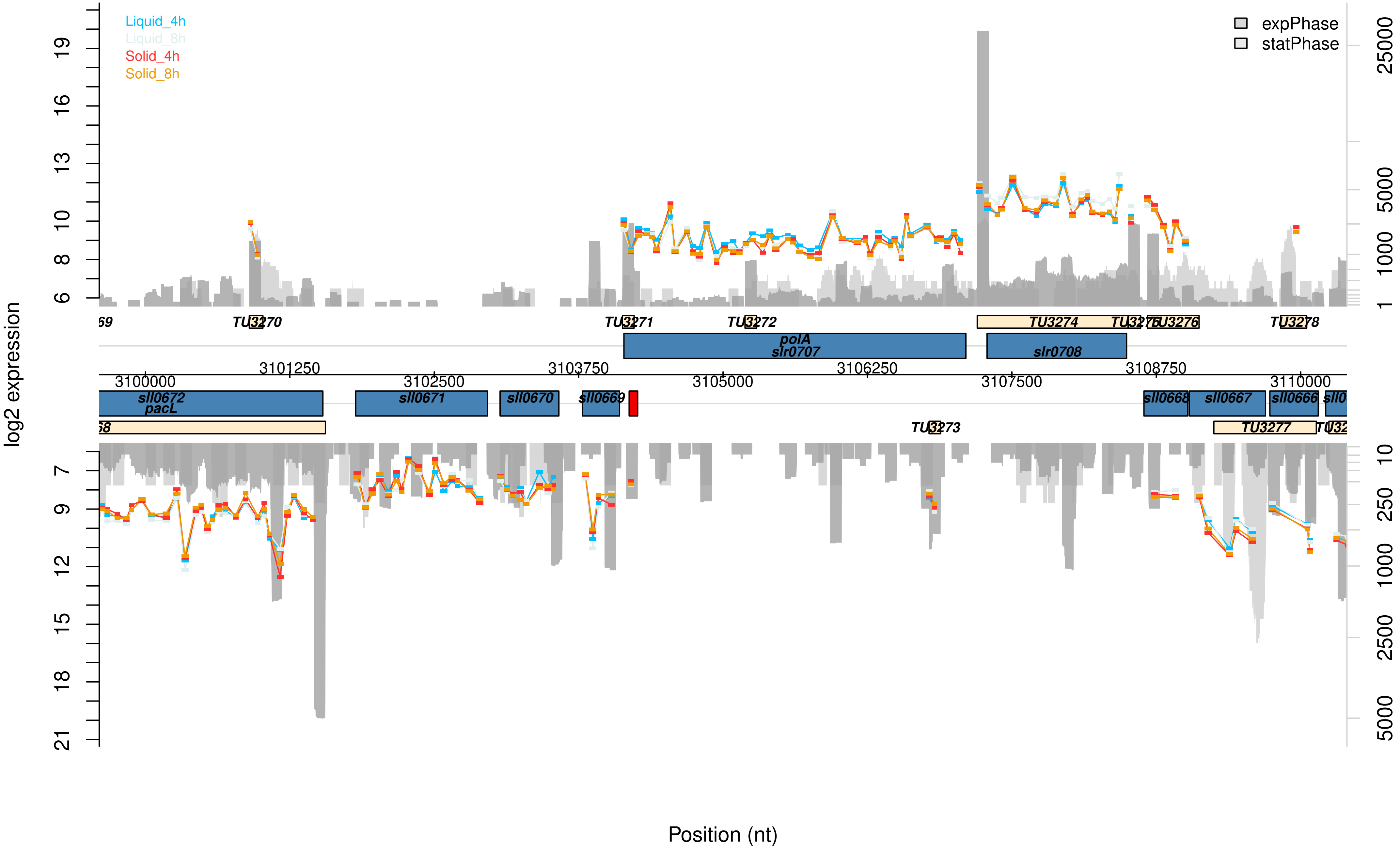
Liquid_4h
Liquid_8h
Solid_4h
Solid_8h
expPhase
statPhase
log2 expression
Position (nt)
TU3270
TU3271
TU3272
TU3273
TU3274
TU3276
TU3277
TU3278
polA
slr0707
slr0708
sll0672
pacL
sll0671
sll0670
sll0669
sll0668
sll0667
sll0666
3100000
3101250
3102500
3103750
3105000
3106250
3107500
3108750
3110000

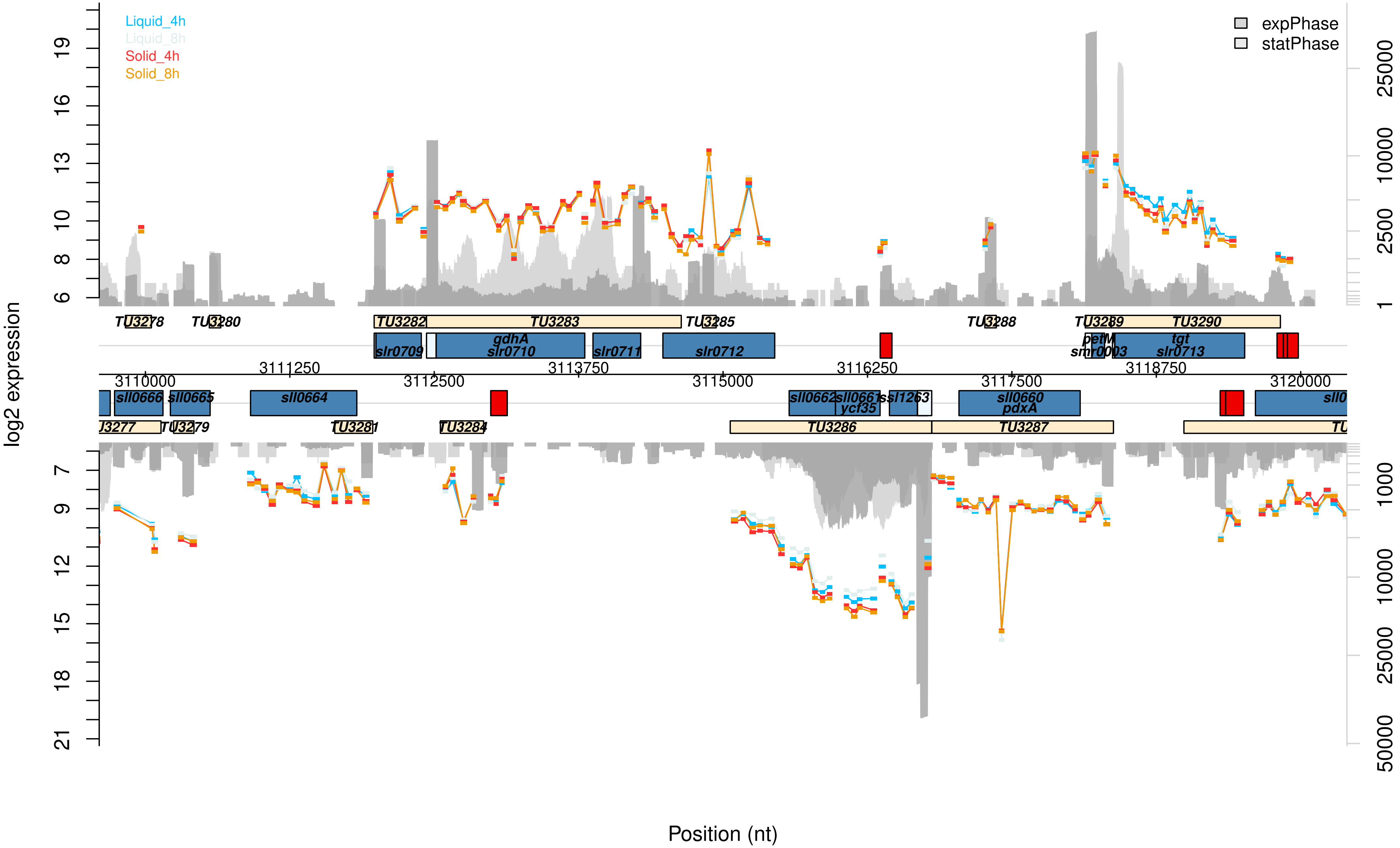

Liquid_4h
Liquid_8h
Solid_4h
Solid_8h
expPhase
statPhase
log2 expression
Position (nt)
TU3278
TU3280
TU3282
TU3283
TU3285
TU3288
TU3289
TU3290
slr0709
gdhA
slr0710
slr0711
slr0712
petM
smr0003
tgt
slr0713
3110000
3111250
3112500
3113750
3115000
3116250
3117500
3118750
3120000
sll0666
sll0665
sll0664
sll0662
sll0661
ycf35
ssl1263
sll0660
pdxA
TU3279
TU3281
TU3284
TU3286
TU3287

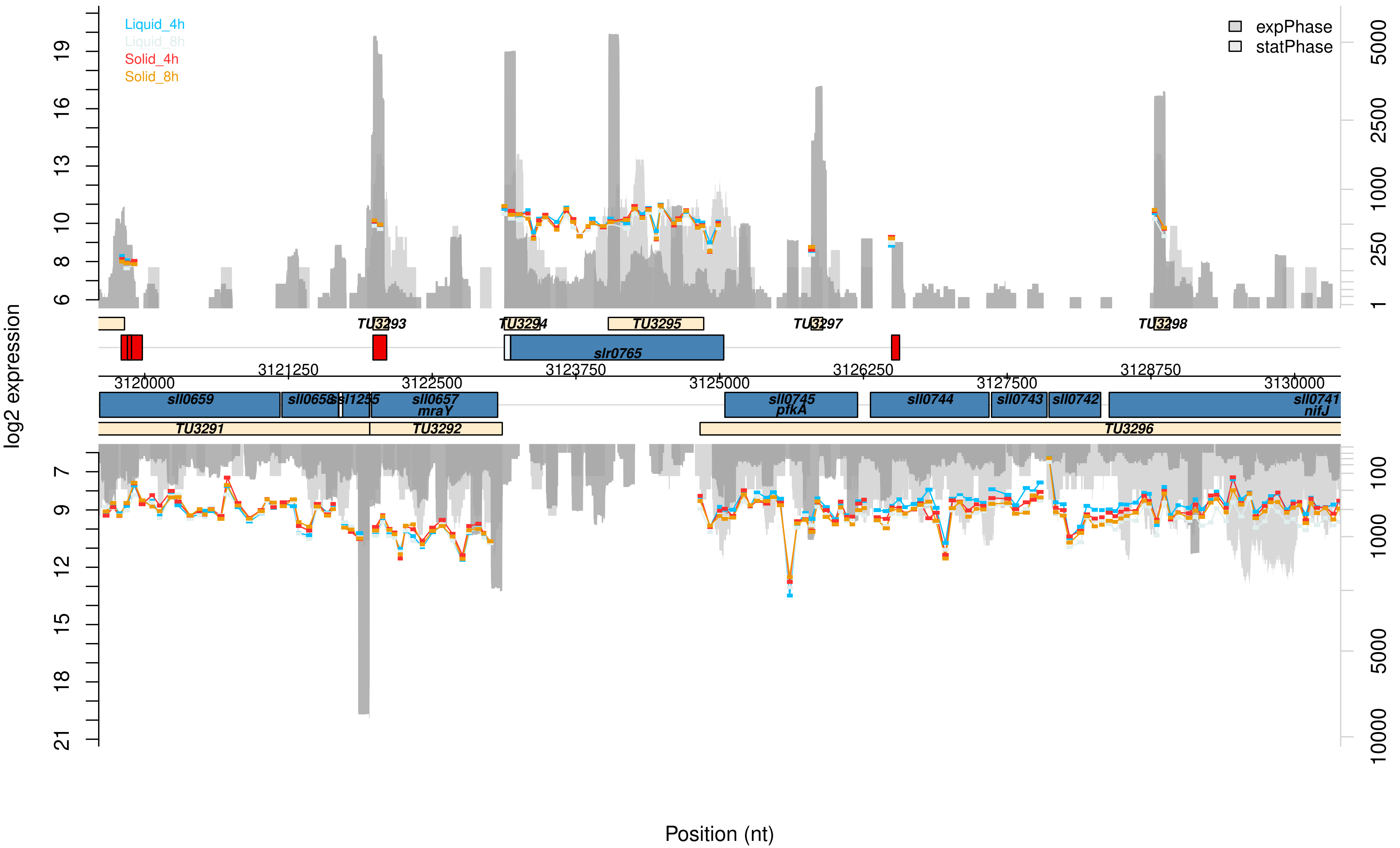

Liquid_4h
Liquid_8h
Solid_4h
Solid_8h
expPhase
statPhase
log2 expression
TU3293
TU3294
TU3295
TU3297
TU3298
slr0765
3120000
3121250
3122500
3123750
3125000
3126250
3127500
3128750
3130000
sll0659
sll0658
ssl1235
sll0657
mraY
sll0745
pfkA
sll0744
sll0743
sll0742
sll0741
nifJ
TU3291
TU3292
TU3296
Position (nt)

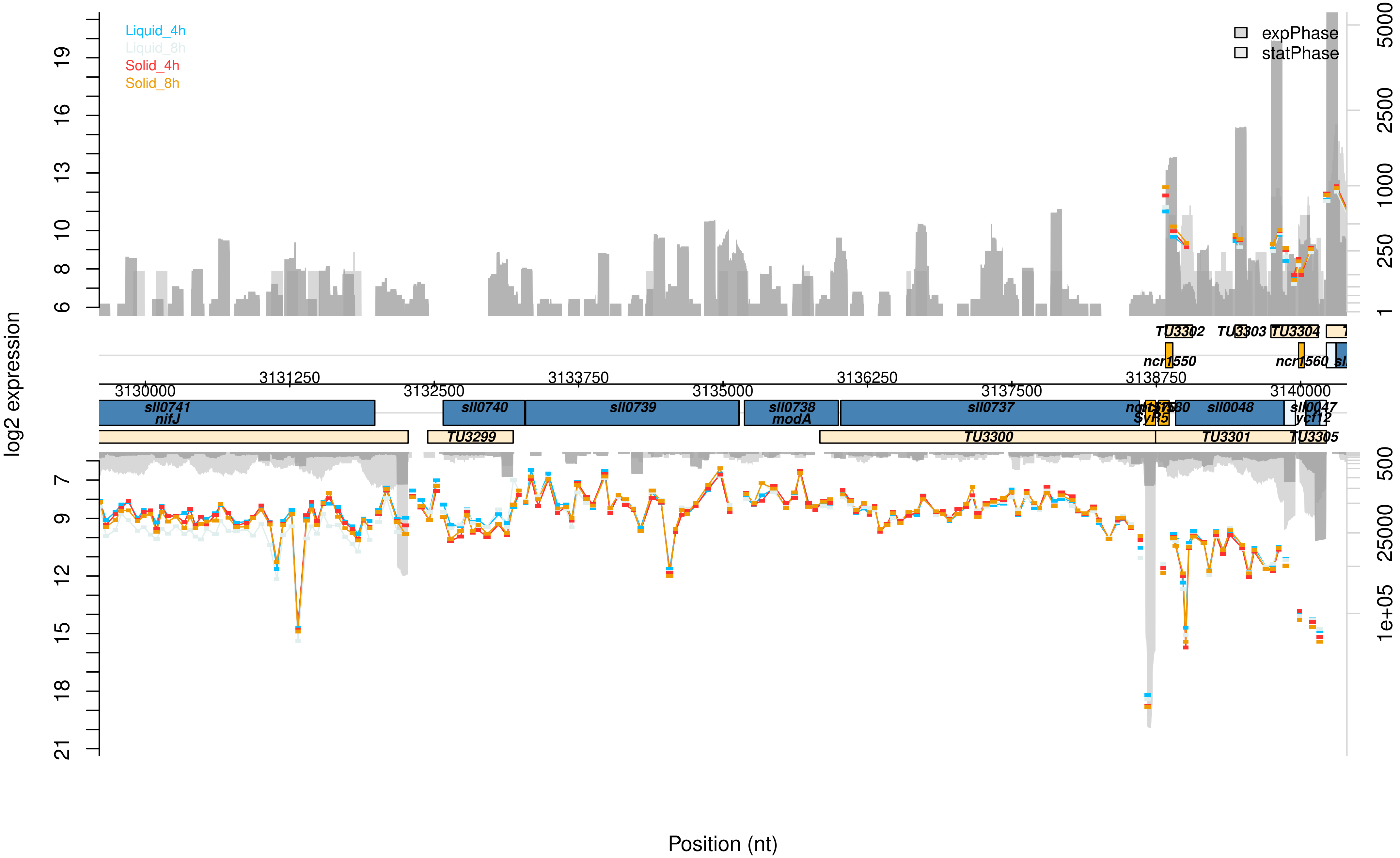

Liquid_4h
Liquid_8h
Solid_4h
Solid_8h
expPhase
statPhase
log2 expression
Position (nt)
TU3302
TU3303
TU3304
ncr1550
ncr1560
3130000
3131250
3132500
3133750
3135000
3136250
3137500
3138750
3140000
sll0741
nifJ
sll0740
sll0739
sll0738
modA
sll0737
sll0048
sll0047
ycf12
TU3299
TU3300
TU3301
TU3305

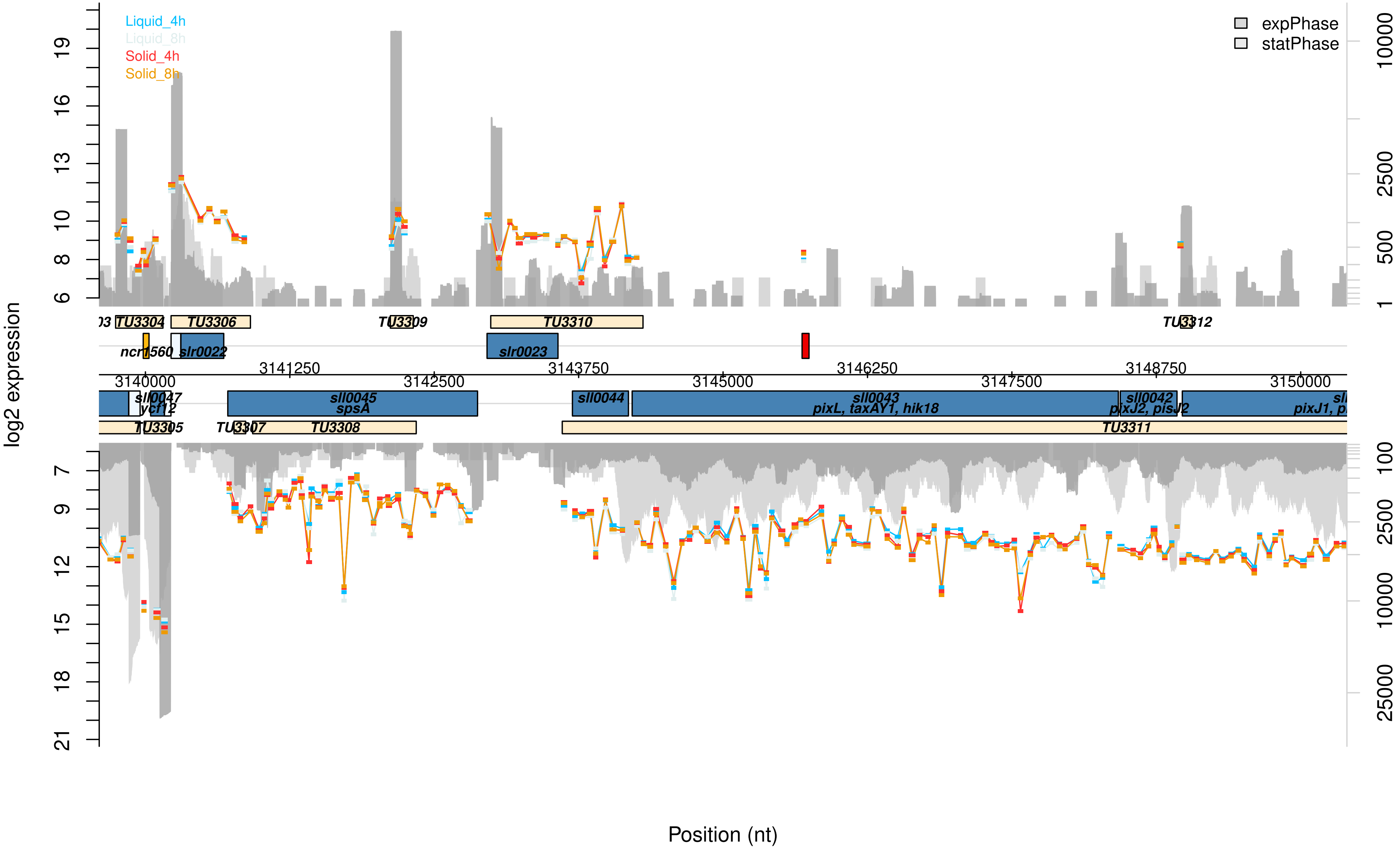

Liquid_4h
Liquid_8h
Solid_4h
Solid_8h
expPhase
statPhase
log2 expression
Position (nt)
TU3304
TU3306
TU3309
TU3310
TU3312
ncr1560
slr0022
slr0023
sll0047
ycf12
sll0045
spsA
sll0044
sll0043
pixL, taxAY1, hik18
sll0042
pixJ2, pisJ2
pixJ1
TU3305
TU3307
TU3308
TU3311
3140000
3141250
3142500
3143750
3145000
3146250
3147500
3148750
3150000

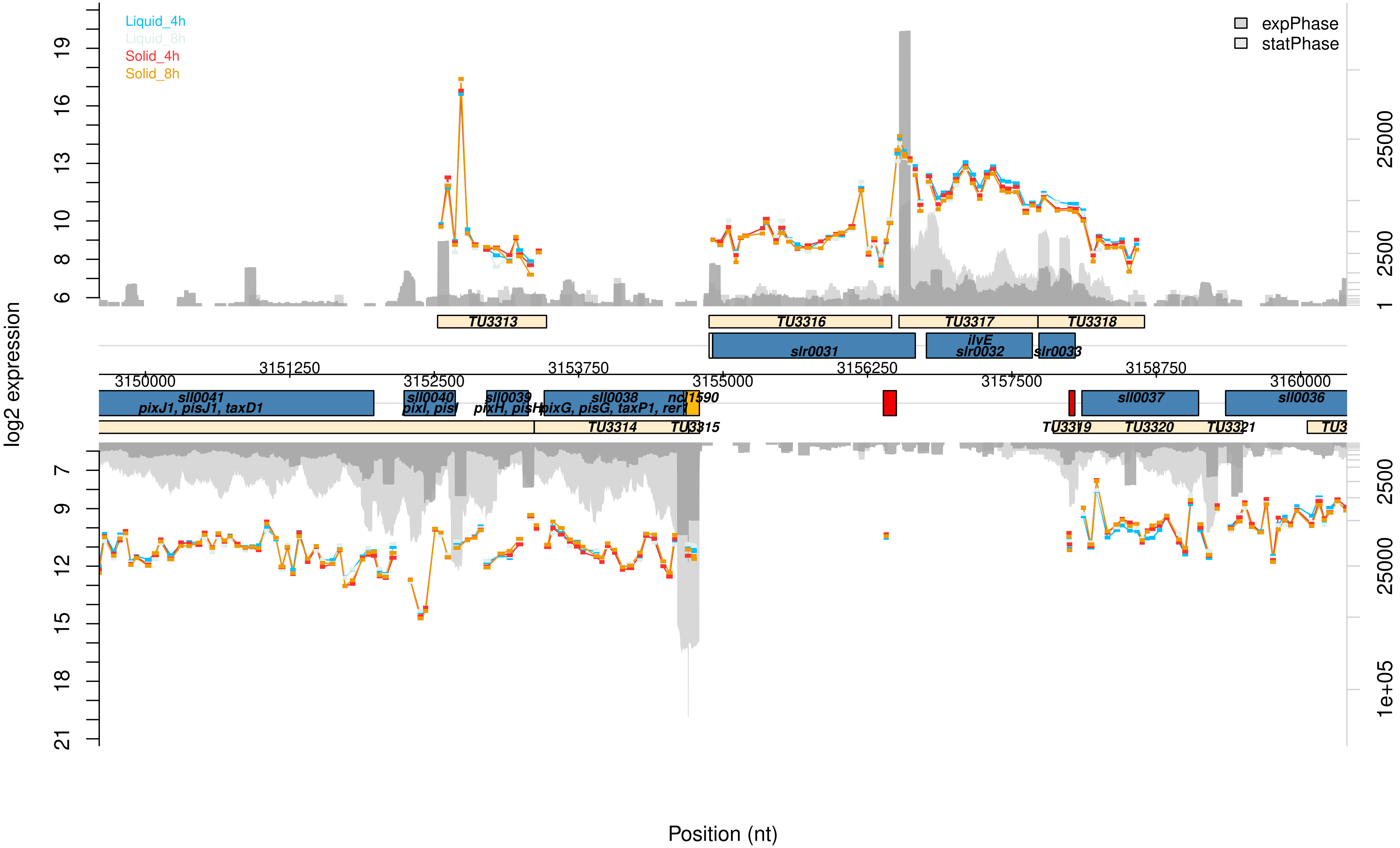

Liquid_4h
Liquid_8h
Solid_4h
Solid_8h
expPhase
statPhase
log2 expression
Position (nt)
TU3313
TU3316
TU3317
TU3318
slr0031
ilvE
slr0032
slr0033
3150000
3151250
3152500
3153750
3155000
3156250
3157500
3158750
3160000
sll0041
pixJ1, pisJ1, taxD1
sll0040
pixI, pisI
sll0039
pixH, pisH
sll0038
pixG, pisG, taxP1, rer1
ncl1590
sll0037
sll0036
TU3314
TU3315
TU3319
TU3320
TU3321

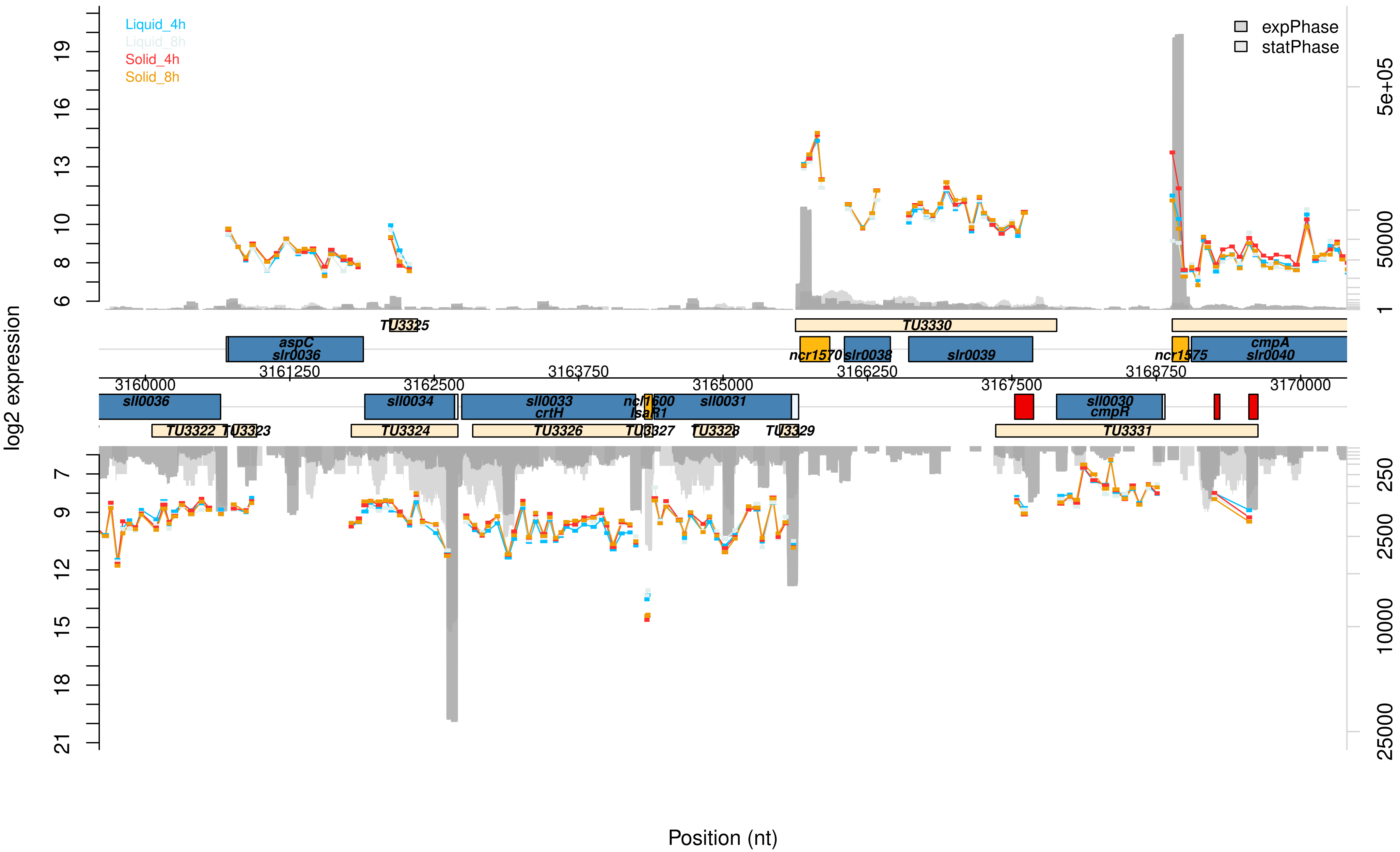

Liquid_4h
Liquid_8h
Solid_4h
Solid_8h
expPhase
statPhase
log2 expression
5e+05
50000
1
250
2500
10000
25000
TU3325
TU3330
aspC
slr0036
ncr1570
slr0038
slr0039
ncr1575
cmpA
slr0040
3161250
3163750
3166250
3168750
3160000
3162500
3165000
3167500
3170000
sll0036
sll0034
sll0033
crtH
ncl1600
IsaR1
sll0031
sll0030
cmpR
TU3322
TU3323
TU3324
TU3326
TU3327
TU3328
TU3329
TU3331
Position (nt)

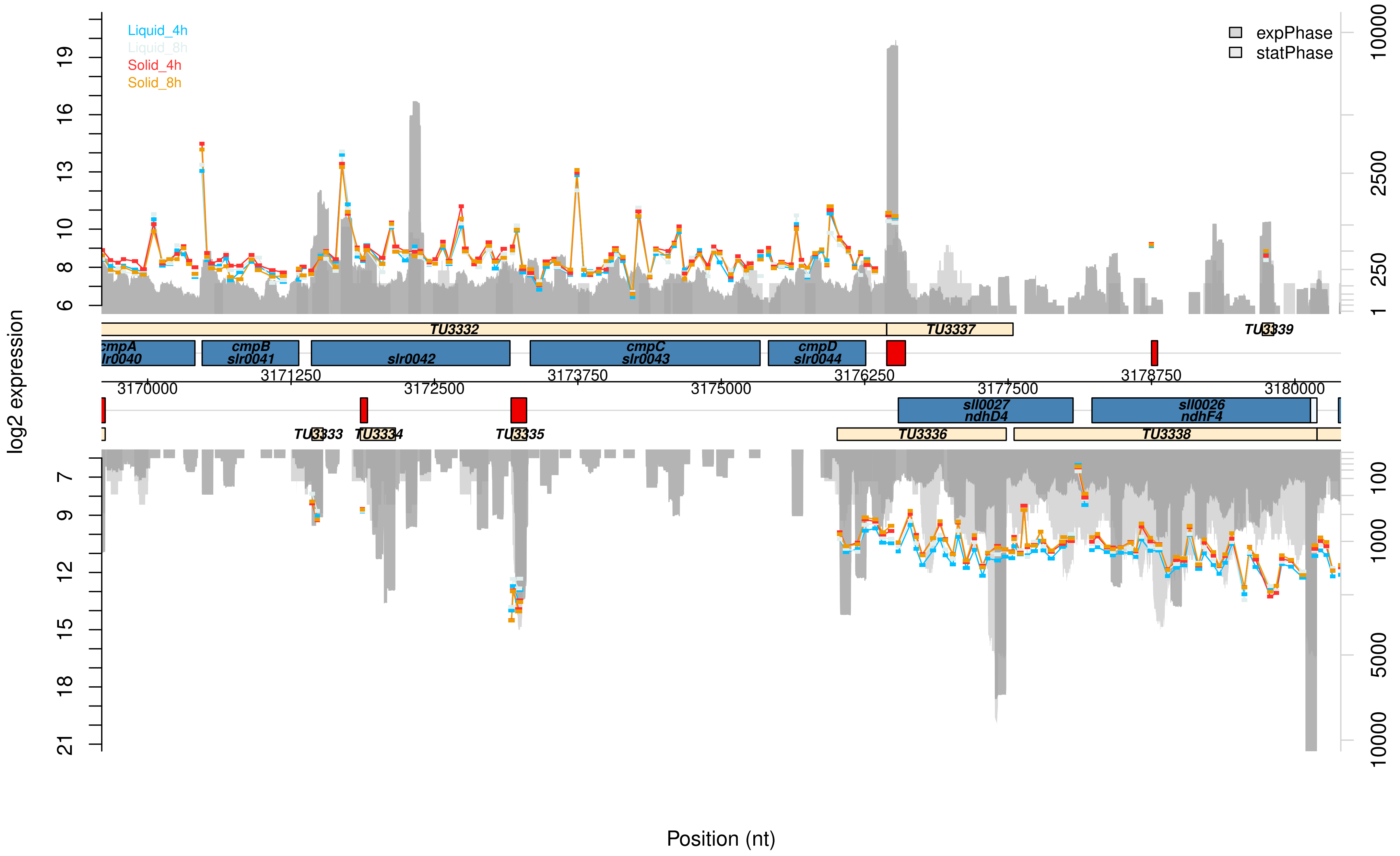

Liquid_4h
Liquid_8h
Solid_4h
Solid_8h
expPhase
statPhase
log2 expression
TU3332
TU3337
TU3339
cmpA
slr0040
cmpB
slr0041
slr0042
cmpC
slr0043
cmpD
slr0044
3170000
3171250
3172500
3173750
3175000
3176250
3177500
3178750
3180000
sll0027
ndhD4
sll0026
ndhF4
TU3333
TU3334
TU3335
TU3336
TU3338
Position (nt)

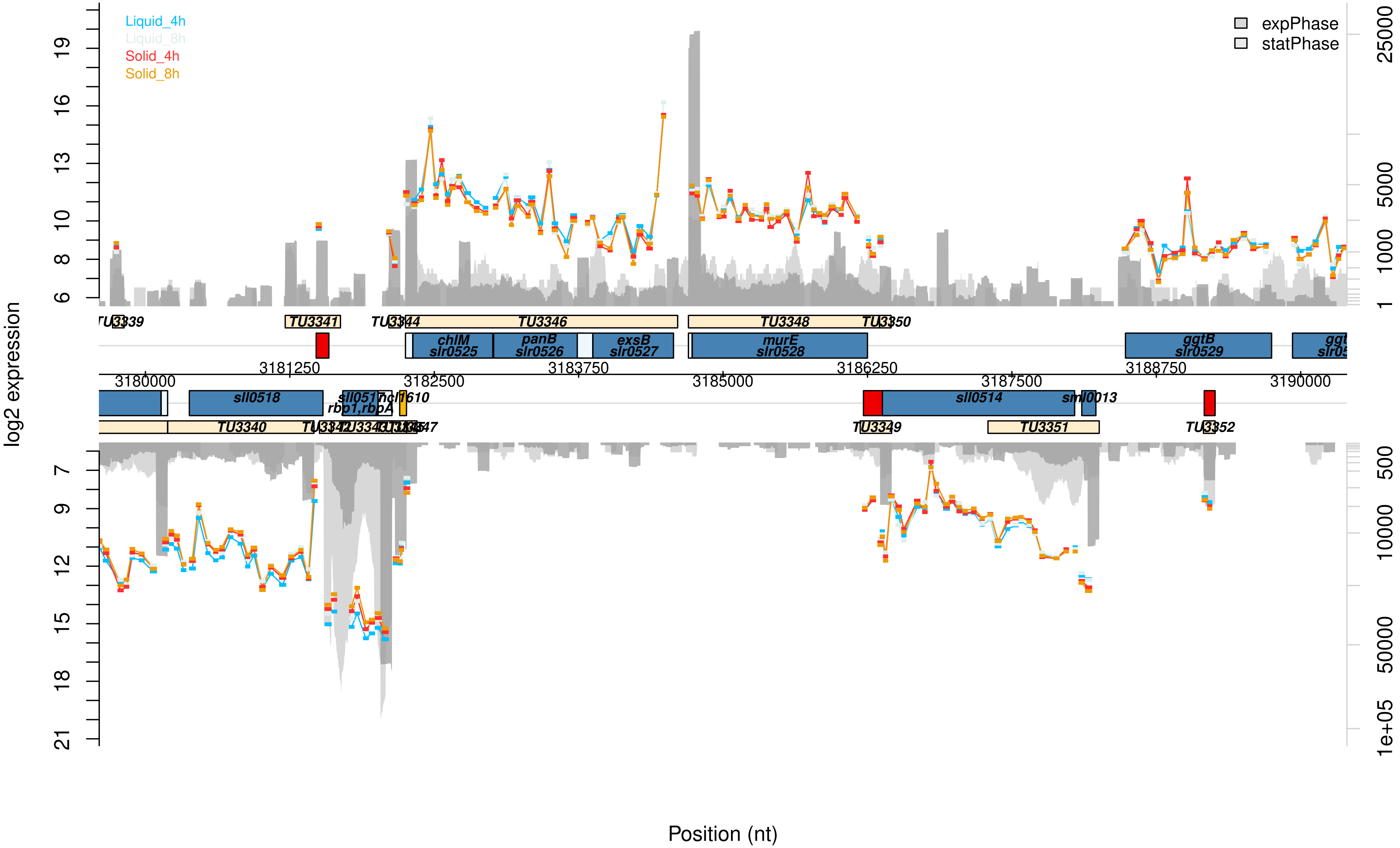

Liquid_4h
Liquid_8h
Solid_4h
Solid_8h
expPhase
statPhase
log2 expression
Position (nt)
TU3339
TU3341
TU3344
TU3346
TU3348
TU3350
chlM
slr0525
panB
slr0526
exsB
slr0527
murE
slr0528
ggtB
slr0529
3181250
3183750
3186250
3188750
3180000
3182500
3185000
3187500
3190000
sll0518
sll0517
rbp1,rbpA
ncl1610
sll0514
sml0013
TU3340
TU3349
TU3351
TU3352

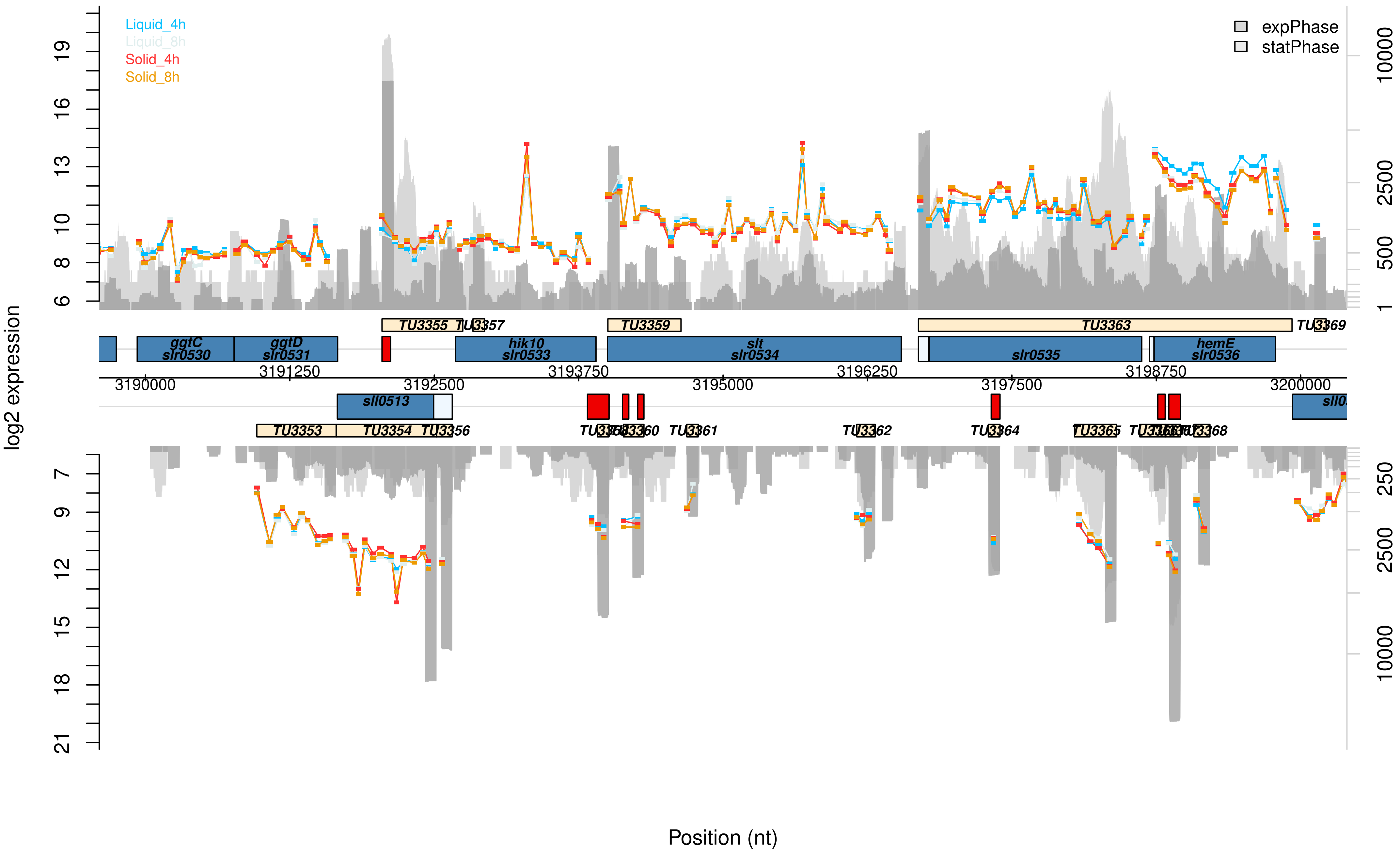

Liquid_4h
Liquid_8h
Solid_4h
Solid_8h
expPhase
statPhase
log2 expression
Position (nt)
TU3355
TU3357
TU3359
TU3363
TU3369
ggtC
slr0530
ggtD
slr0531
hik10
slr0533
slt
slr0534
slr0535
hemE
slr0536
sll0513
TU3353
TU3354
TU3356
TU3361
TU3362
TU3364
TU3365
TU3368
3190000
3191250
3192500
3193750
3195000
3196250
3197500
3198750
3200000

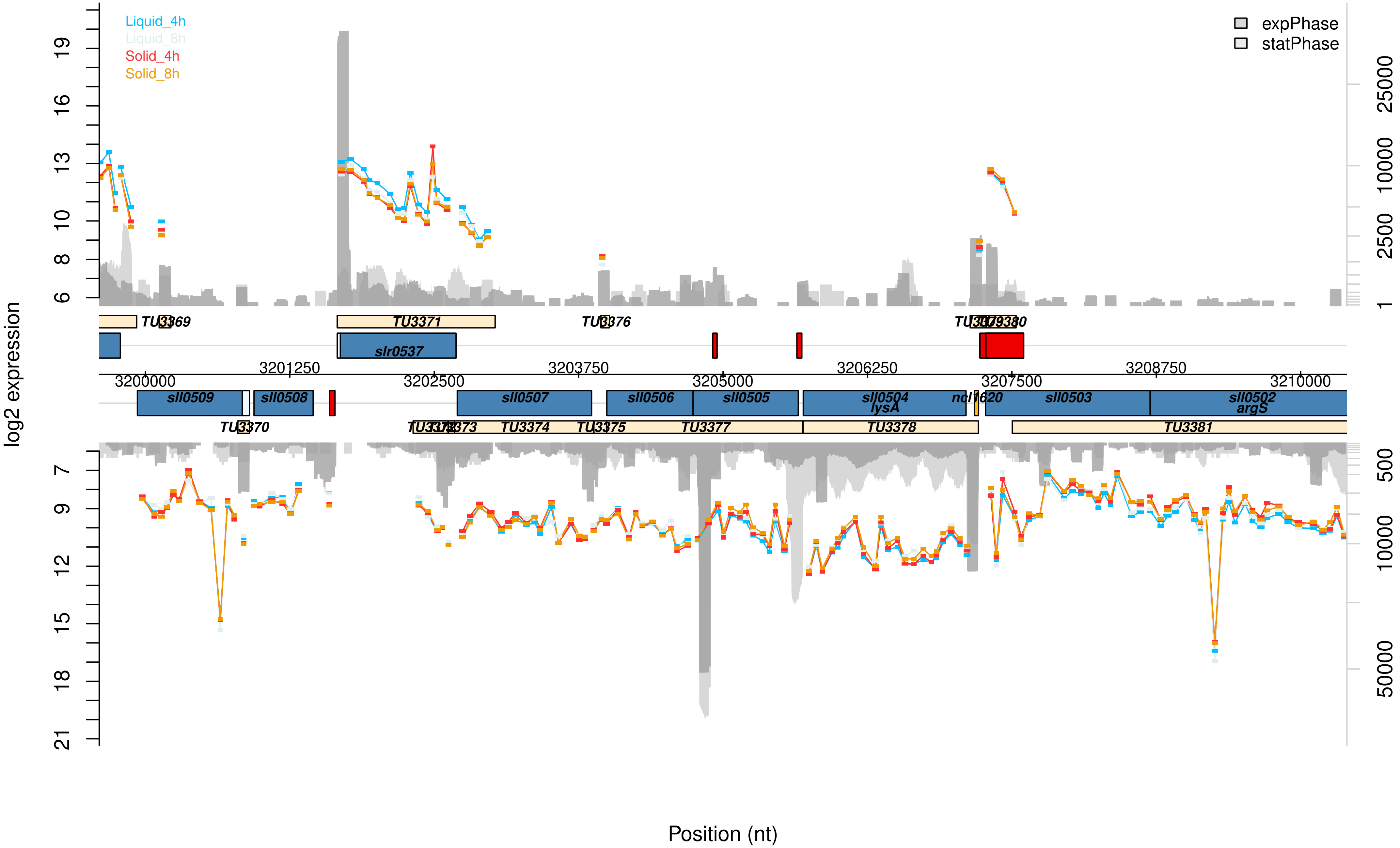

Liquid_4h
Liquid_8h
Solid_4h
Solid_8h
expPhase
statPhase
log2 expression
Position (nt)
TU3369
TU3371
TU3376
TU3380
slr0537
sll0509
sll0508
sll0507
sll0506
sll0505
sll0504
lysA
ncl1620
sll0503
sll0502
argS
TU3370
TU3373
TU3374
TU3375
TU3377
TU3378
TU3381
3200000
3201250
3202500
3203750
3205000
3206250
3207500
3208750
3210000

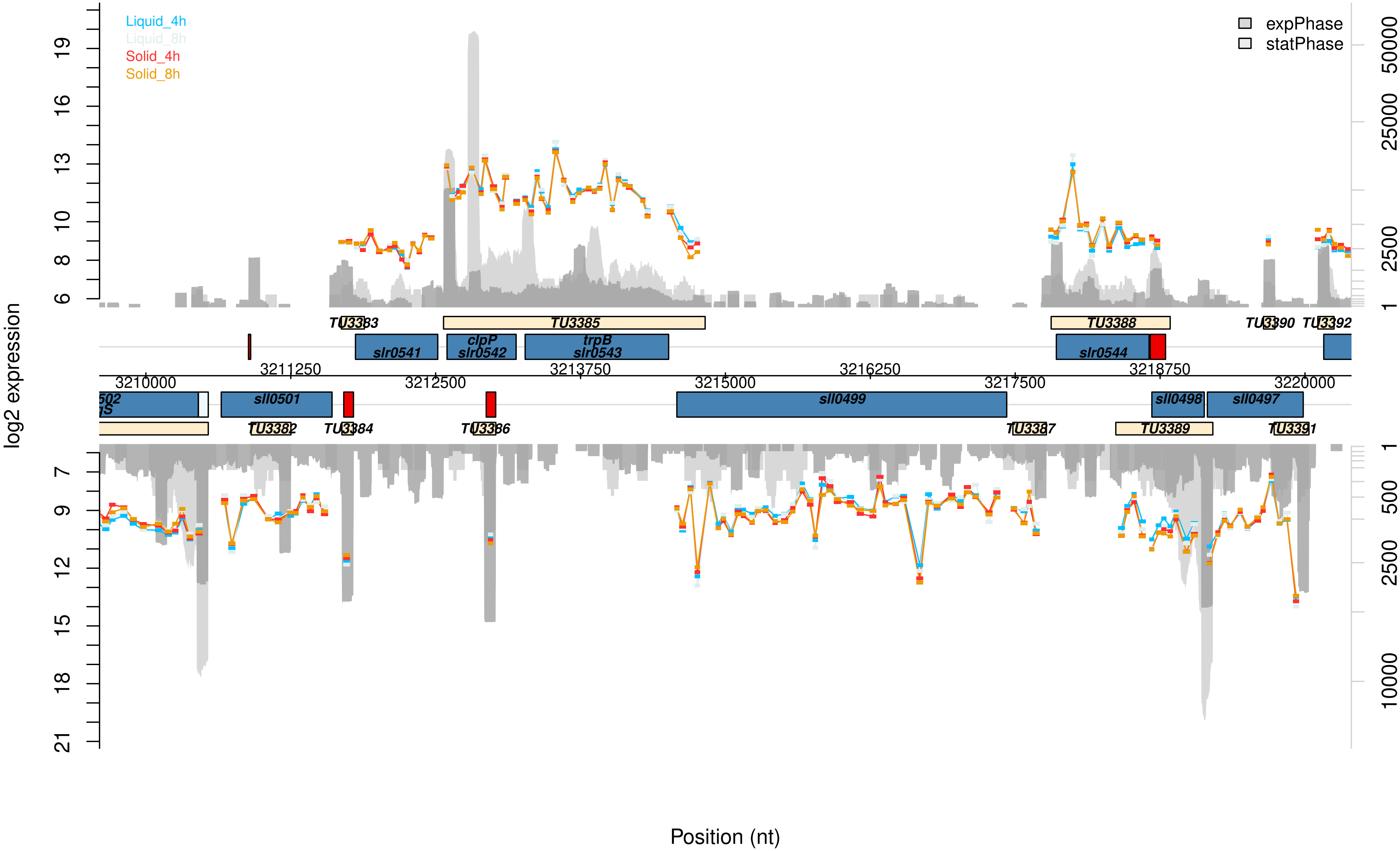

Liquid_4h
Liquid_8h
Solid_4h
Solid_8h
expPhase
statPhase
log2 expression
Position (nt)
TU3383
TU3385
TU3388
TU3390
TU3392
slr0541
clpP
slr0542
trpB
slr0543
slr0544
3210000
3211250
3212500
3213750
3215000
3216250
3217500
3218750
3220000
sll0501
sll0499
sll0498
sll0497
TU3382
TU3384
TU3386
TU3387
TU3389
TU3391

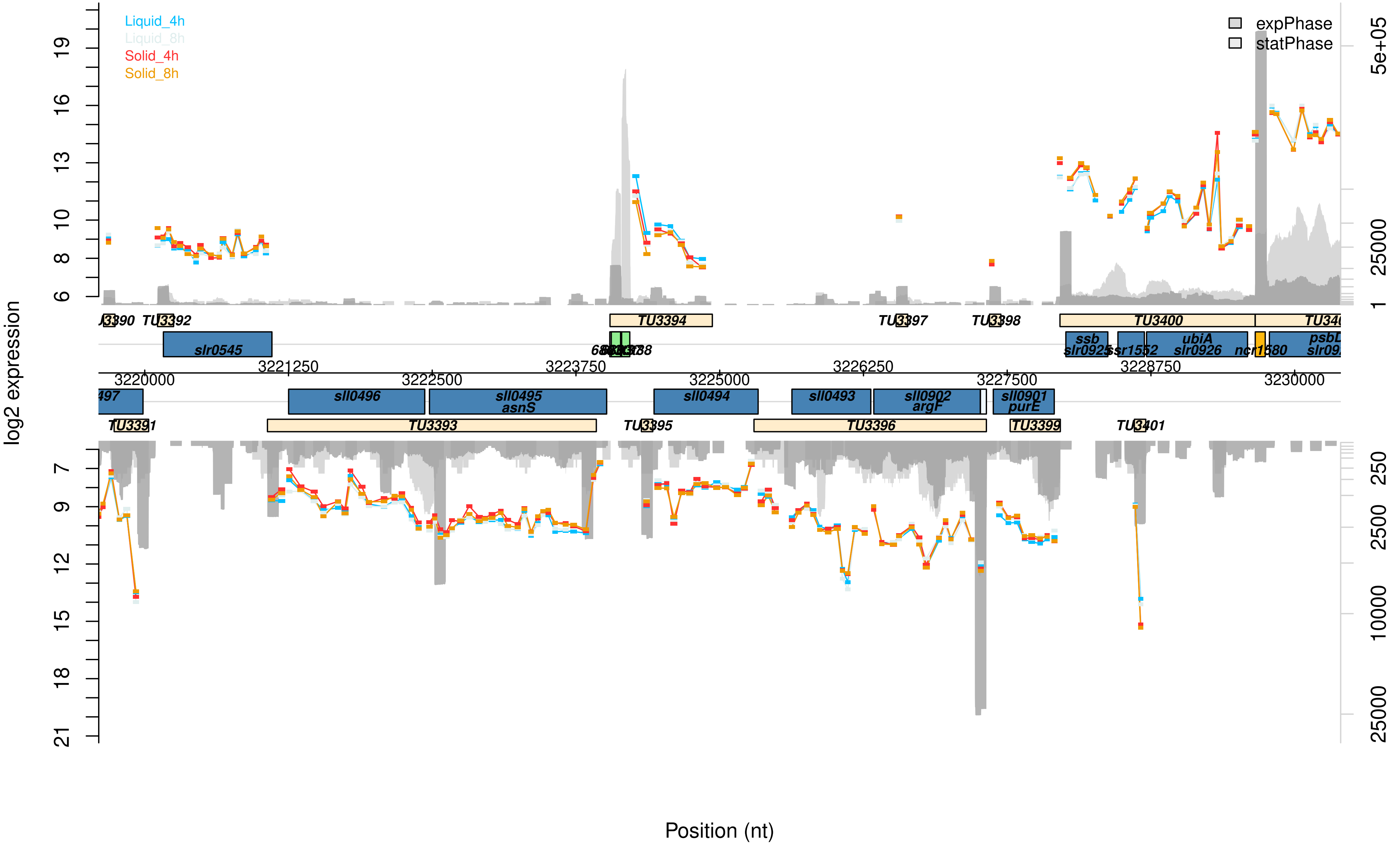

Liquid_4h
Liquid_8h
Solid_4h
Solid_8h
expPhase
statPhase
log2 expression
Position (nt)
TU3392
slr0545
TU3394
TU3397
TU3398
TU3400
ssb
slr0925
ssr1552
ubiA
slr0926
ncr1580
psbL
3220000
3221250
3222500
3223750
3225000
3226250
3227500
3228750
3230000
sll0496
sll0495
asnS
sll0494
sll0493
sll0902
argF
sll0901
purE
TU3391
TU3393
TU3395
TU3396
TU3399
TU3401
5e+05
25000
2500
250
10000

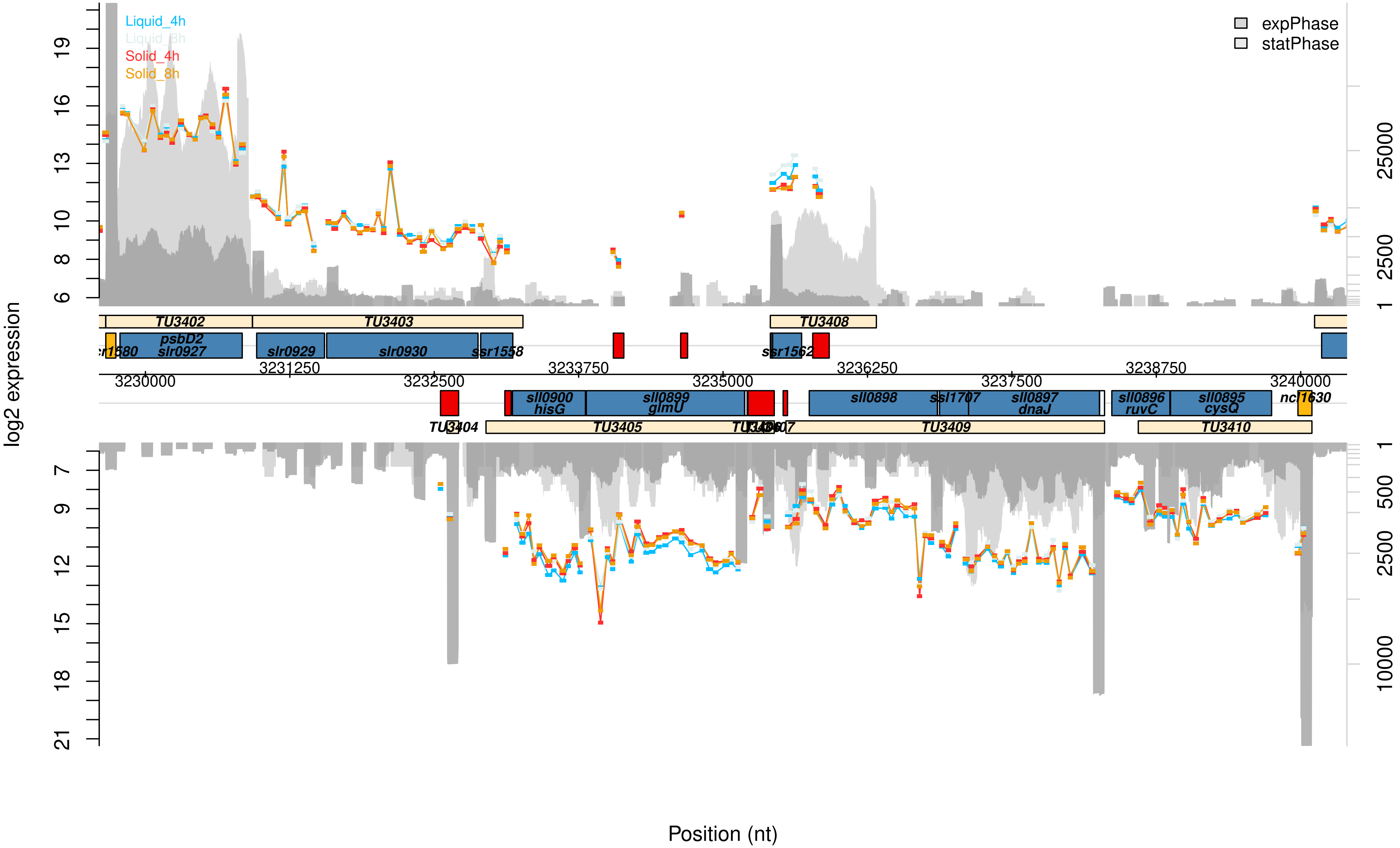

log2 expression
Liquid_4h
Liquid_8h
Solid_4h
Solid_8h
expPhase
statPhase
TU3402
TU3403
TU3408
psbD2
slr0927
slr0929
slr0930
ssr1558
ssr1562
3229000
3230000
3231250
3232500
3233750
3235000
3236250
3237500
3238750
3240000
sll0900
hisG
sll0899
glmU
sll0898
ssl1707
sll0897
dnaJ
sll0896
ruvC
sll0895
cysQ
ncl1630
TU3404
TU3405
TU3409
TU3410
Position (nt)

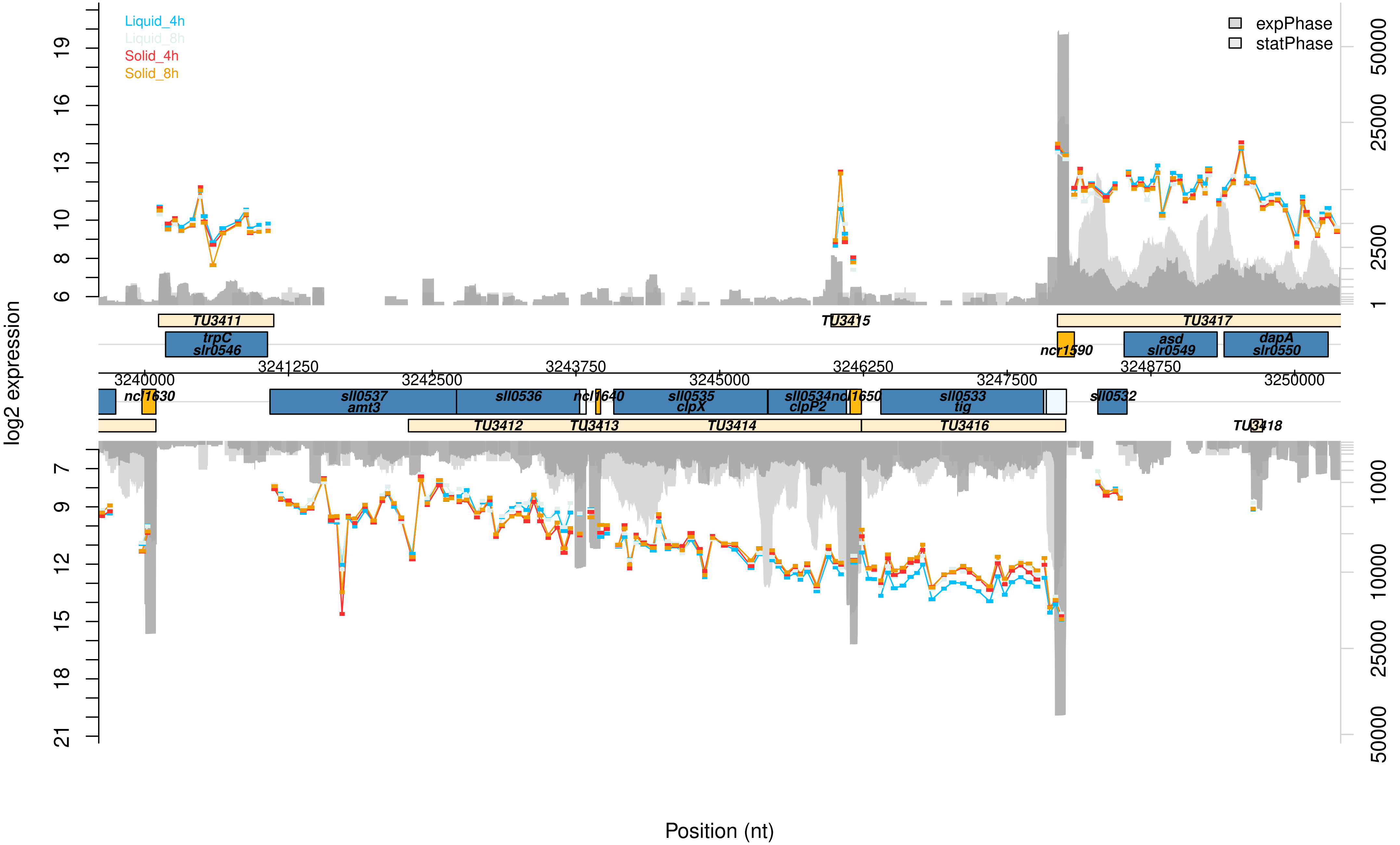

Liquid_4h
Liquid_8h
Solid_4h
Solid_8h
expPhase
statPhase
log2 expression
Position (nt)
TU3411
trpC
slr0546
TU3415
TU3417
ncr1590
asd
slr0549
dapA
slr0550
3240000
3241250
3242500
3243750
3245000
3246250
3247500
3248750
3250000
ncl1630
sll0537
amt3
sll0536
ncl1640
sll0535
clpX
sll0534
clpP2
ncl1650
sll0533
tig
sll0532
TU3412
TU3413
TU3414
TU3416
TU3418

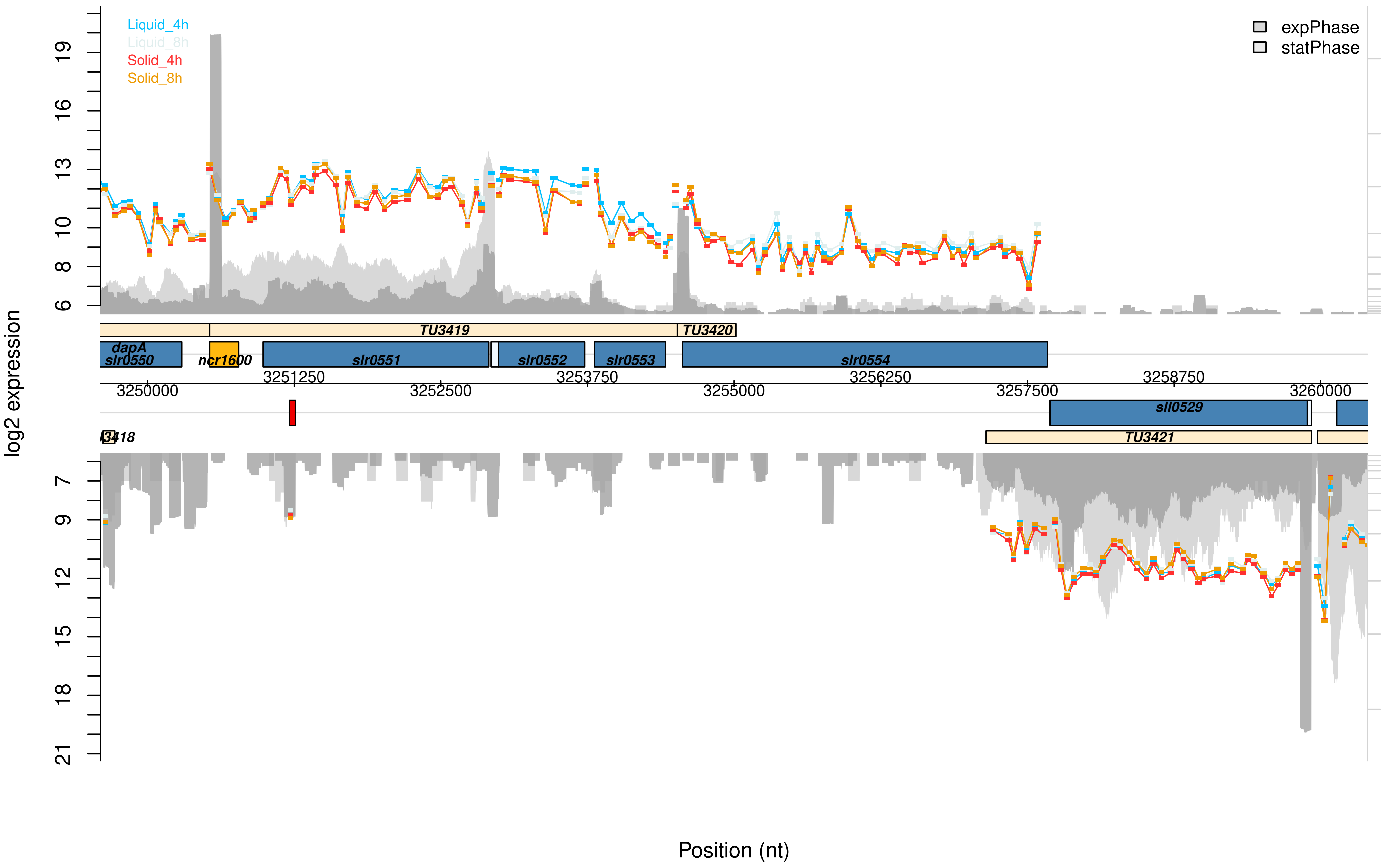

Liquid_4h
Liquid_8h
Solid_4h
Solid_8h
expPhase
statPhase
log2 expression
TU3419
TU3420
dapA
slr0550
ncr1600
slr0551
slr0552
slr0553
slr0554
3250000
3251250
3252500
3253750
3255000
3256250
3257500
3258750
3260000
sll0529
TU3421
Position (nt)

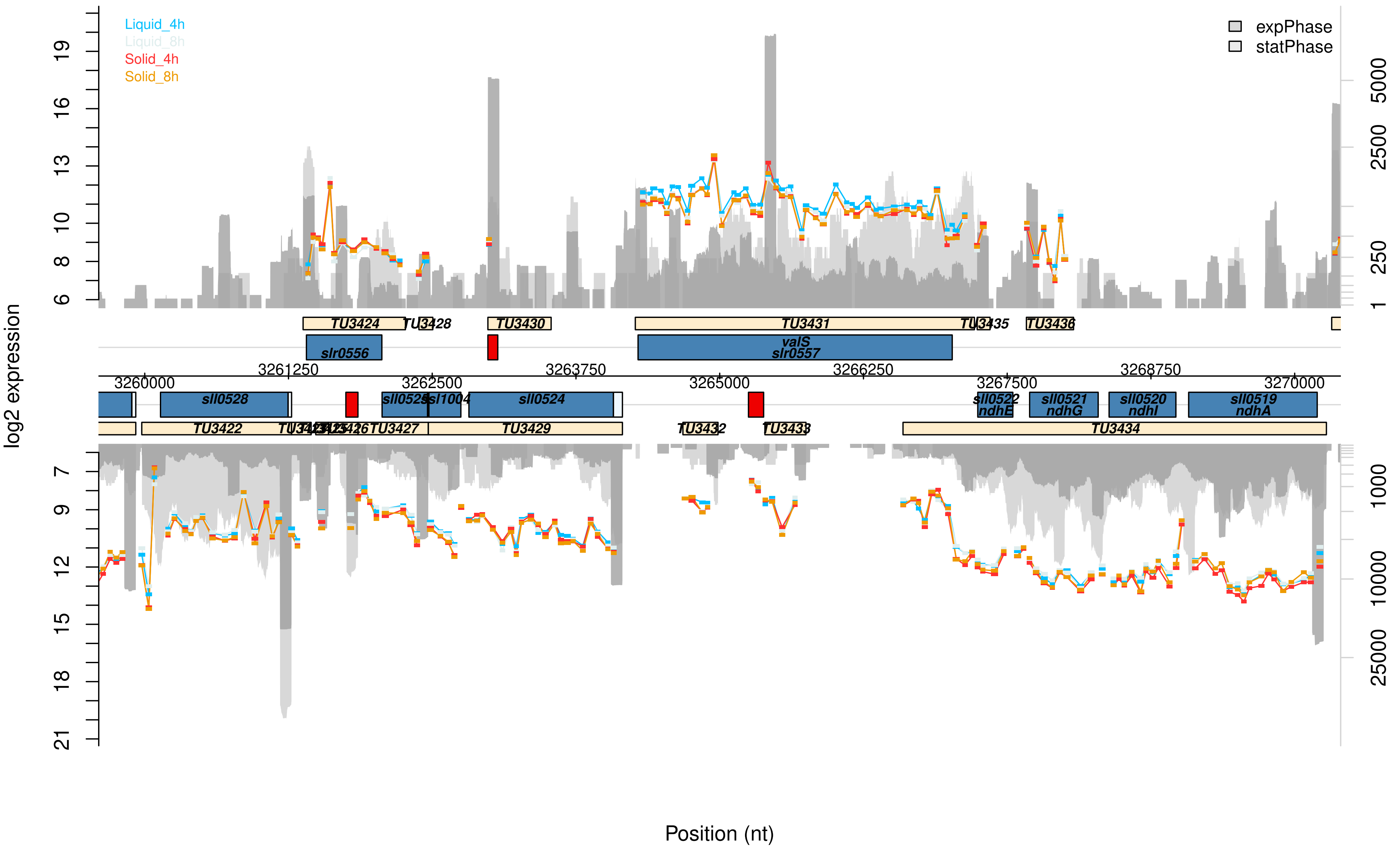

Liquid_4h
Liquid_8h
Solid_4h
Solid_8h
expPhase
statPhase
log2 expression
6
8
10
13
16
19
7
9
12
15
18
21
1
250
2500
5000
1000
10000
25000
TU3424
TU3428
TU3430
TU3431
TU3435
TU3436
slr0556
valS
slr0557
3260000
3261250
3262500
3263750
3265000
3266250
3267500
3268750
3270000
sll0528
sll0525
ssl1004
sll0524
sll0522
ndhE
sll0521
ndhG
sll0520
ndhI
sll0519
ndhA
TU3422
TU3427
TU3429
TU3432
TU3433
TU3434
Position (nt)

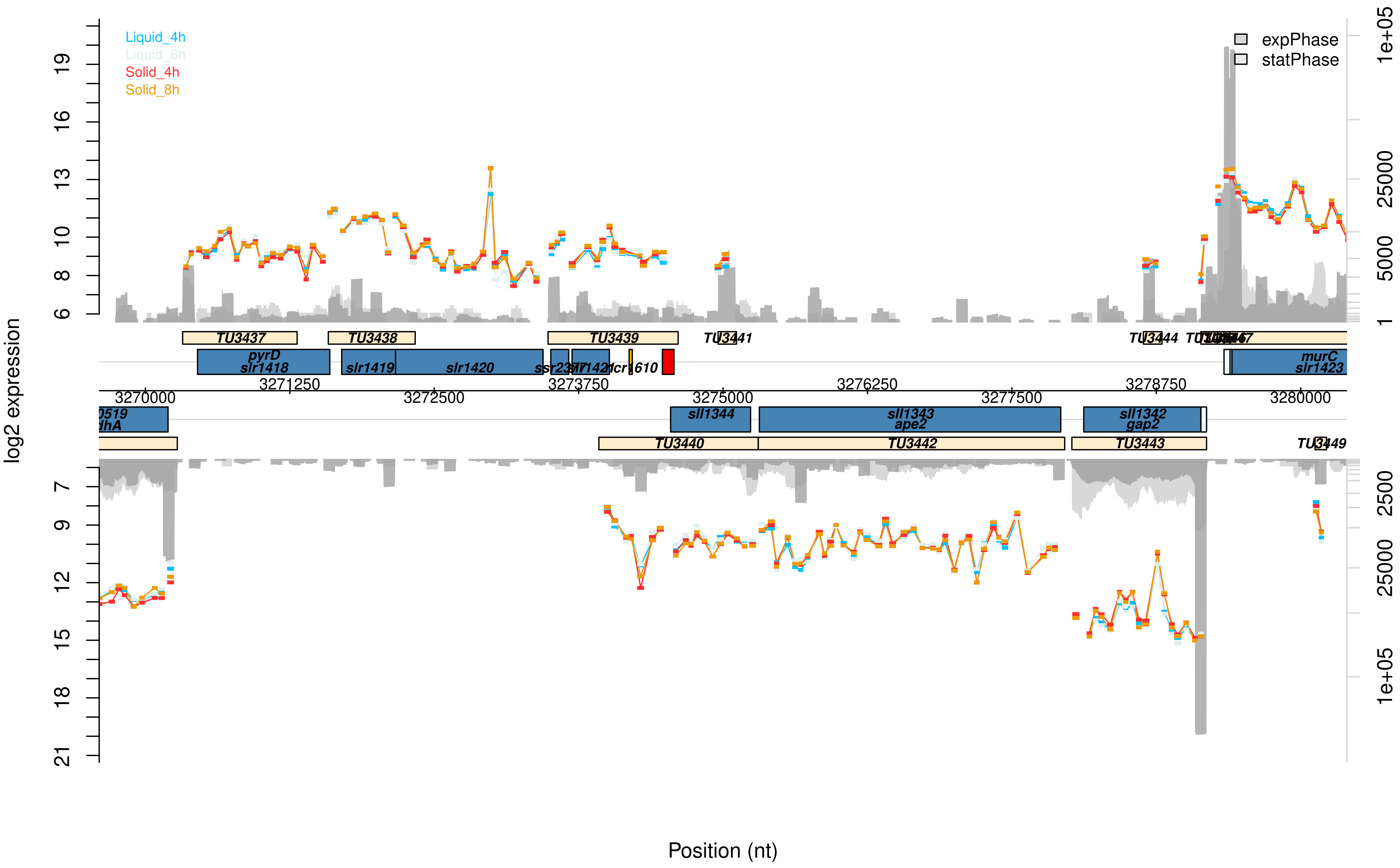

Liquid_4h
Liquid_8h
Solid_4h
Solid_8h
expPhase
statPhase
log2 expression
Position (nt)
TU3437
TU3438
TU3439
TU3441
TU3444
pyrD
slr1418
slr1419
slr1420
murC
slr1423
sll1344
sll1343
ape2
sll1342
gap2
TU3440
TU3442
TU3443
TU3449
3270000
3271250
3272500
3273750
3275000
3276250
3277500
3278750
3280000

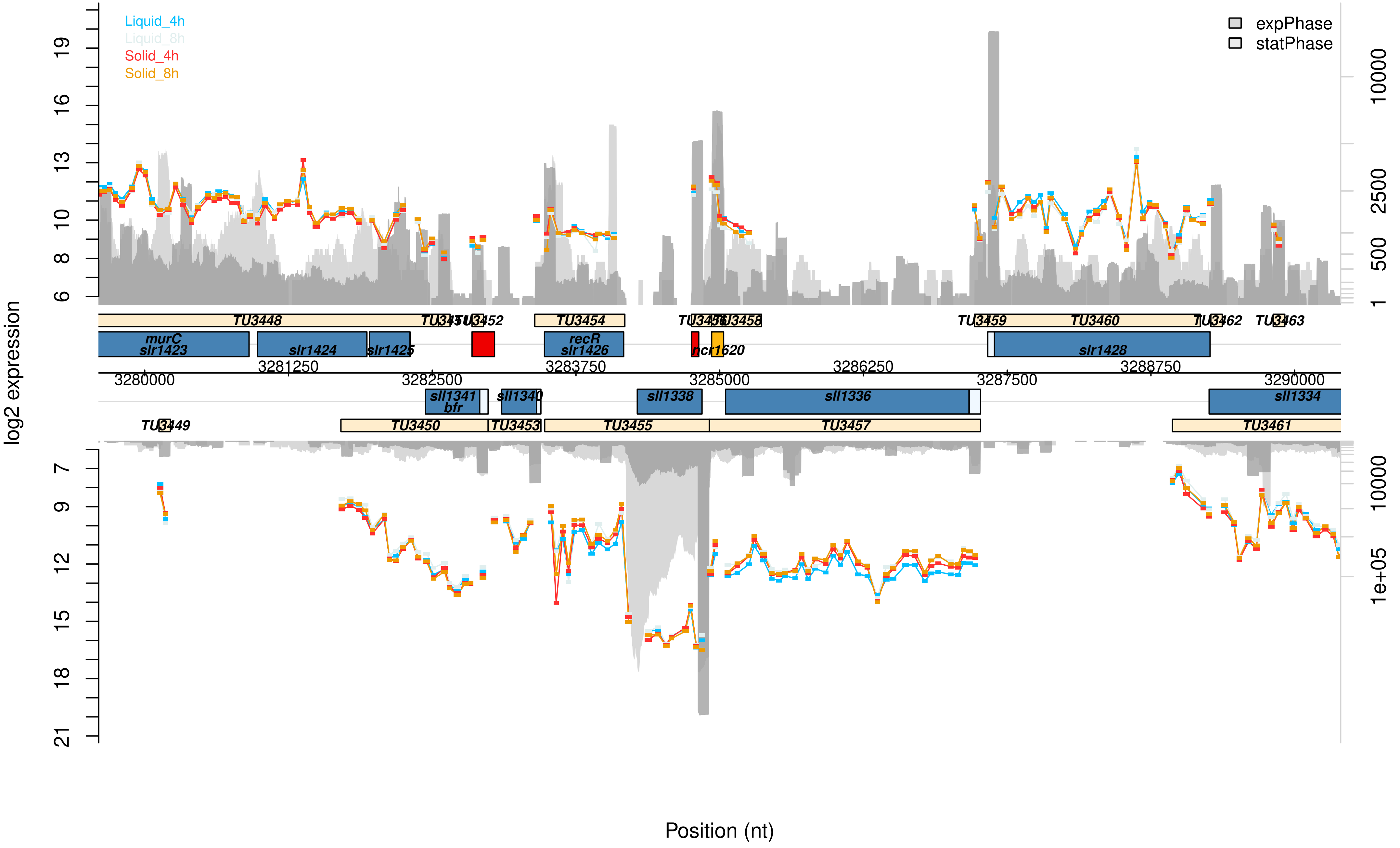
Liquid_4h
Liquid_8h
Solid_4h
Solid_8h
expPhase
statPhase
log2 expression
Position (nt)
TU3448
TU3452
TU3454
TU3459
TU3460
TU3462
TU3463
murC
slr1423
slr1424
slr1425
recR
slr1426
ncr1620
slr1428
sll1341
bfr
sll1340
sll1338
sll1336
sll1334
TU3449
TU3450
TU3453
TU3455
TU3457
TU3461
3280000
3281250
3282500
3283750
3285000
3286250
3287500
3288750
3290000

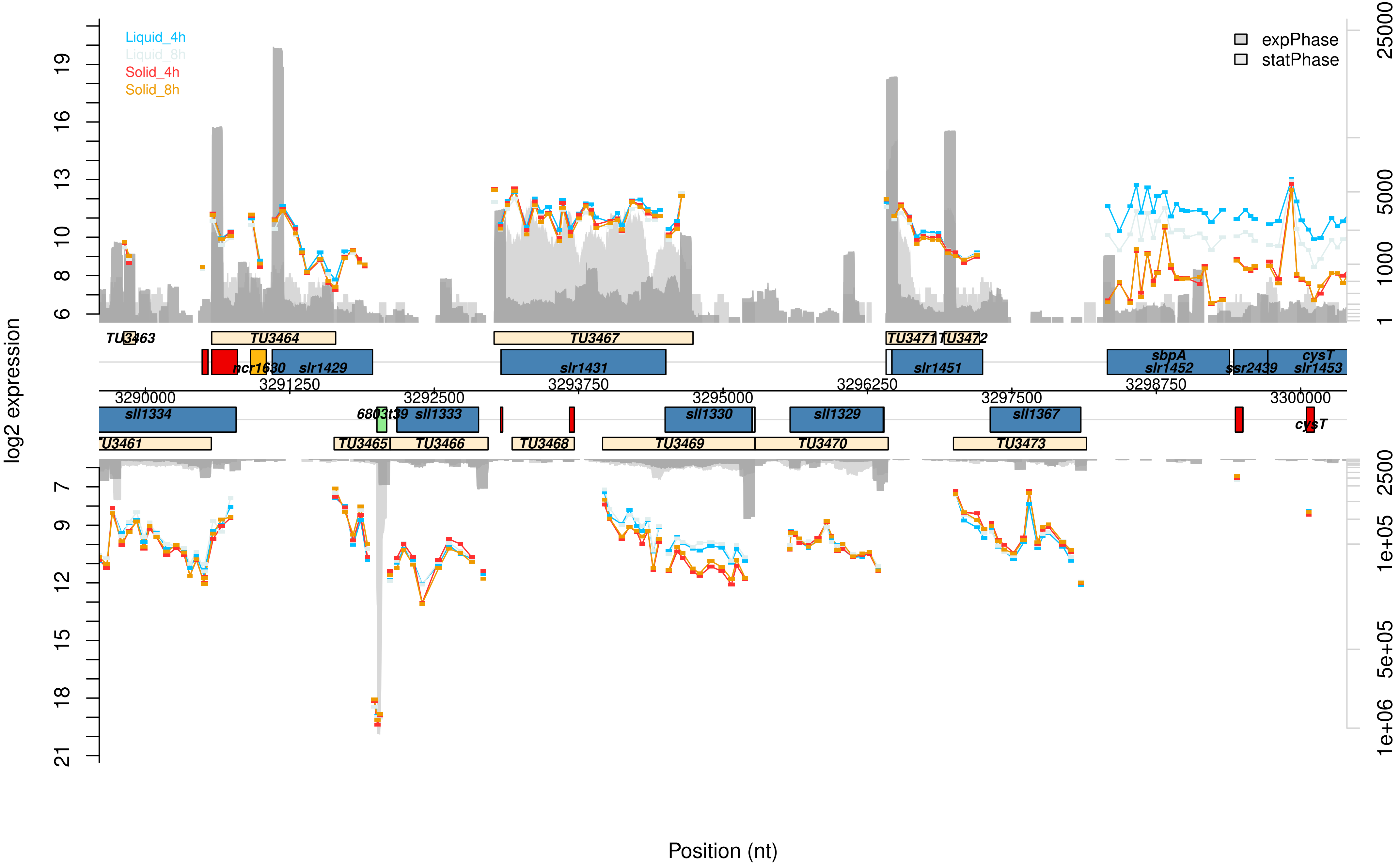

Liquid_4h
Liquid_8h
Solid_4h
Solid_8h
expPhase
statPhase
log2 expression
Position (nt)
TU3463
TU3464
TU3467
TU3471
TU3472
ncr1630
slr1429
slr1431
slr1451
sbpA
slr1452
ssr2439
cysT
slr1453
3290000
3291250
3292500
3293750
3295000
3296250
3297500
3298750
3300000
sll1334
6803t39
sll1333
sll1330
sll1329
sll1367
cysT
TU3461
TU3465
TU3466
TU3468
TU3469
TU3470
TU3473
6
8
10
13
16
19
7
9
12
15
18
21
1
1000
5000
25000
2500
1e+05
5e+05
1e+06

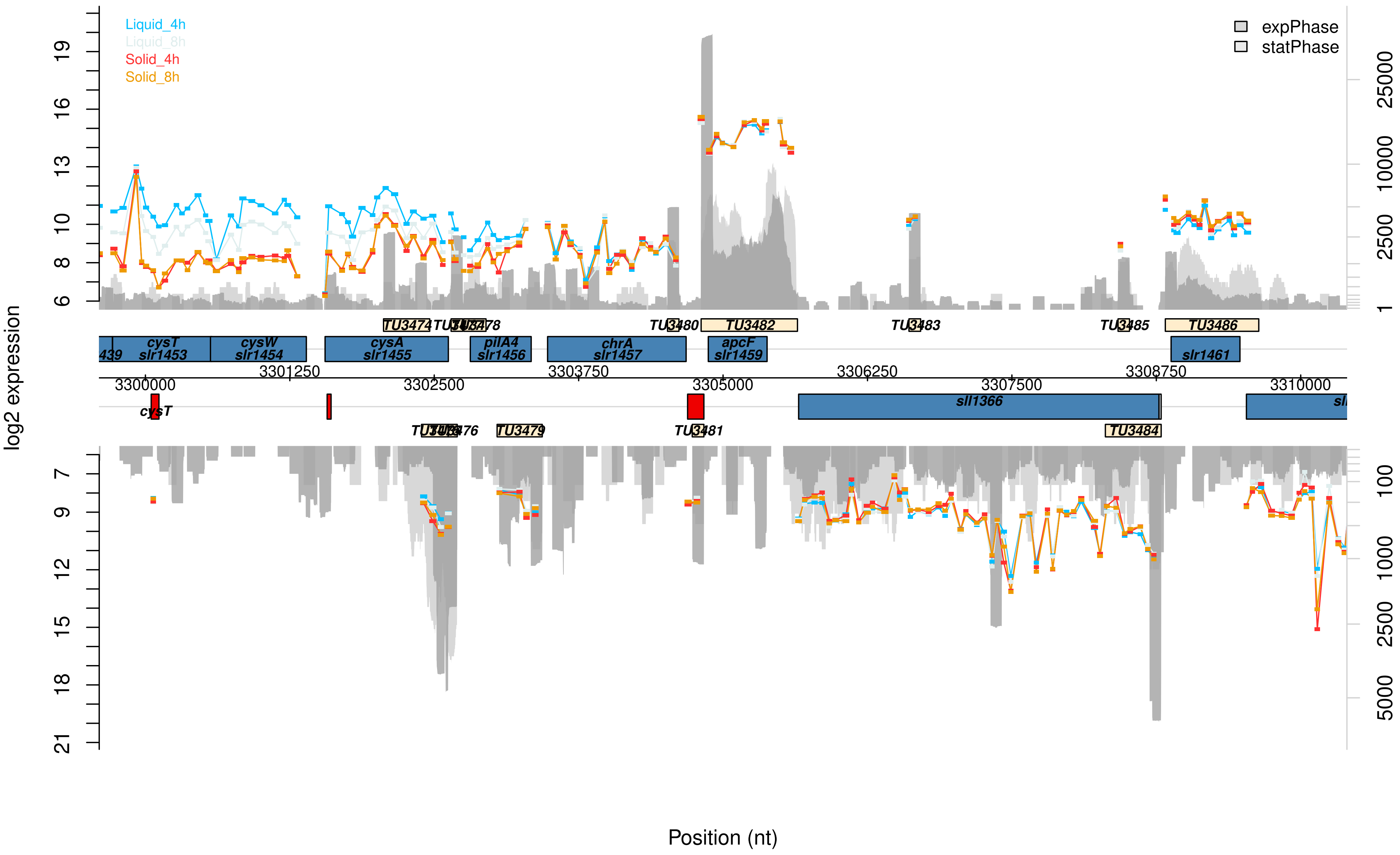

Liquid_4h
Liquid_8h
Solid_4h
Solid_8h
expPhase
statPhase
log2 expression
Position (nt)
TU3474
TU3478
TU3480
TU3482
TU3483
TU3485
TU3486
cysT
slr1453
cysW
slr1454
cysA
slr1455
pilA4
slr1456
chrA
slr1457
apcF
slr1459
slr1461
3300000
3301250
3302500
3303750
3305000
3306250
3307500
3308750
3310000
cysT
sll1366
TU3476
TU3479
TU3481
TU3484

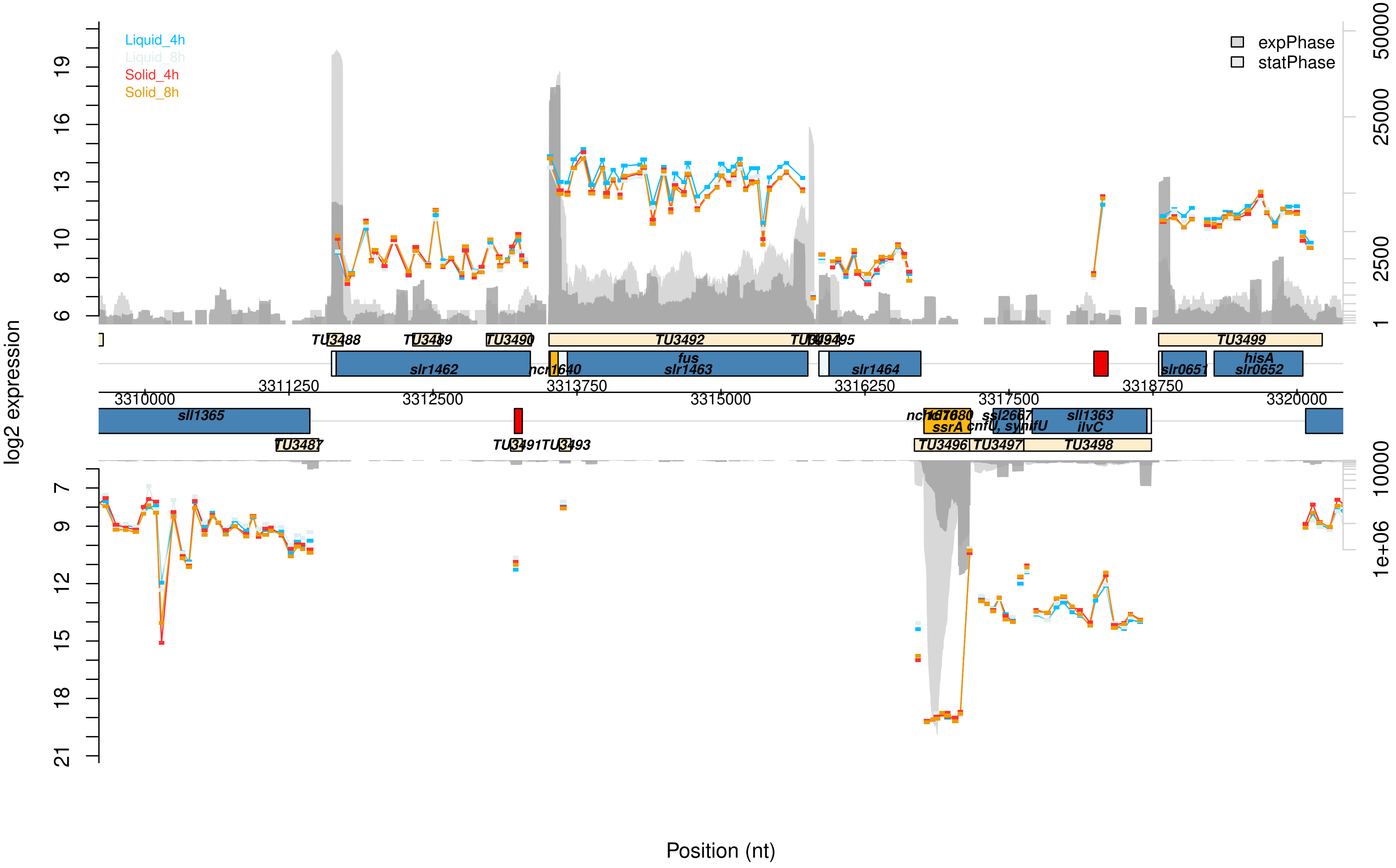
Liquid_4h
Liquid_8h
Solid_4h
Solid_8h
expPhase
statPhase
log2 expression
19
16
13
10
8
6
7
9
12
15
18
21
50000
25000
2500
1
10000
1e+06
TU3488
TU3489
TU3490
TU3492
TU3494
TU3495
TU3499
slr1462
ncl1640
fus
slr1463
slr1464
slr0651
hisA
slr0652
3310000
3311250
3312500
3313750
3315000
3316250
3317500
3318750
3320000
sll1365
ncl1680
ssrA
ssl2667
cnfU, synifU
sll1363
ilvC
TU3487
TU3491
TU3493
TU3496
TU3497
TU3498
Position (nt)

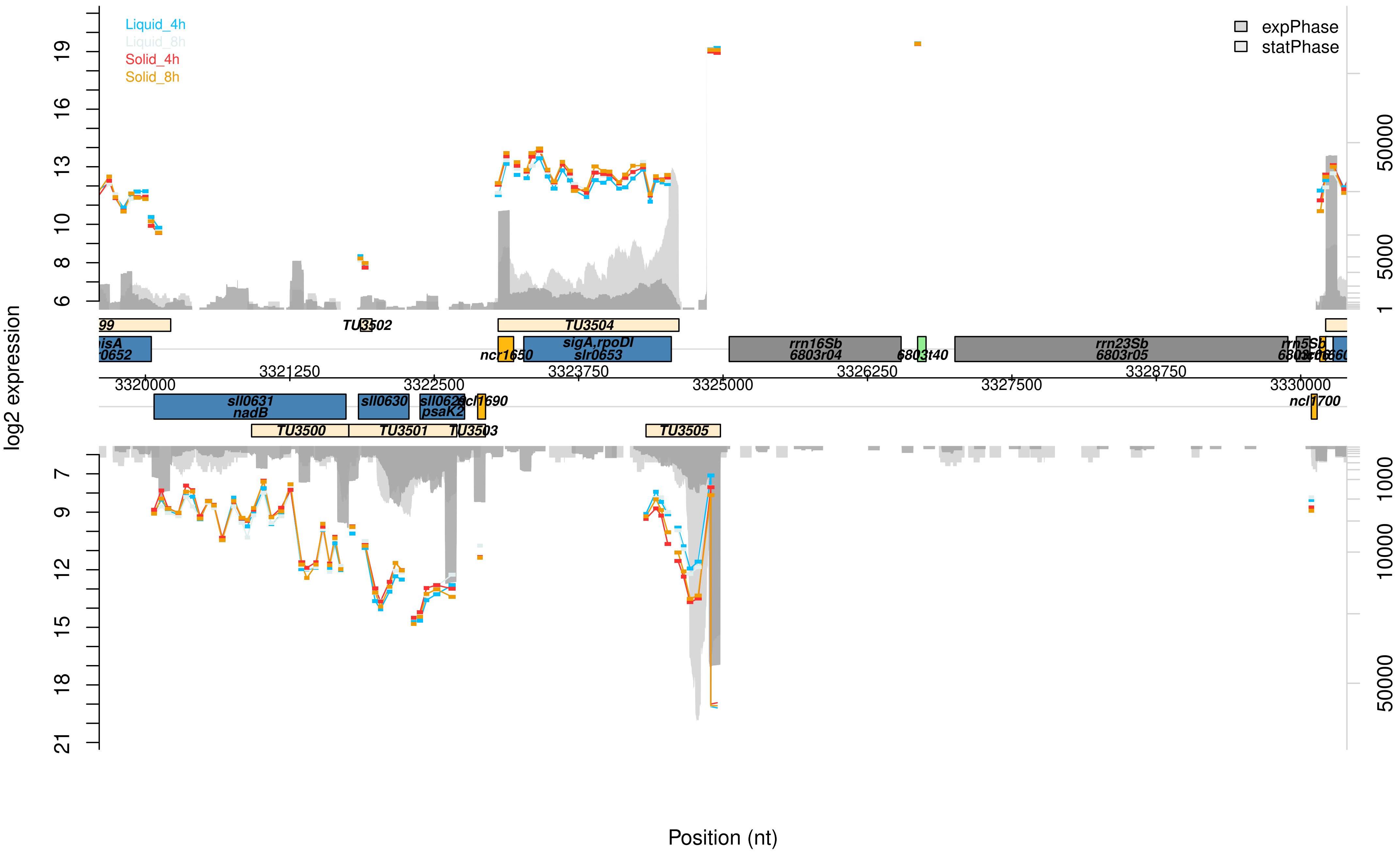

Liquid_4h
Liquid_8h
Solid_4h
Solid_8h
expPhase
statPhase
log2 expression
TU3502
TU3504
ncr1650
sigA,rpoDI
slr0653
rrn16Sb
6803r04
6803t40
rrn23Sb
6803r05
rrn5Sb
3320000
3321250
3322500
3323750
3325000
3326250
3327500
3328750
3330000
sll0631
nadB
sll0630
psaK2
ncl1690
ncl1700
TU3500
TU3501
TU3503
TU3505
Position (nt)

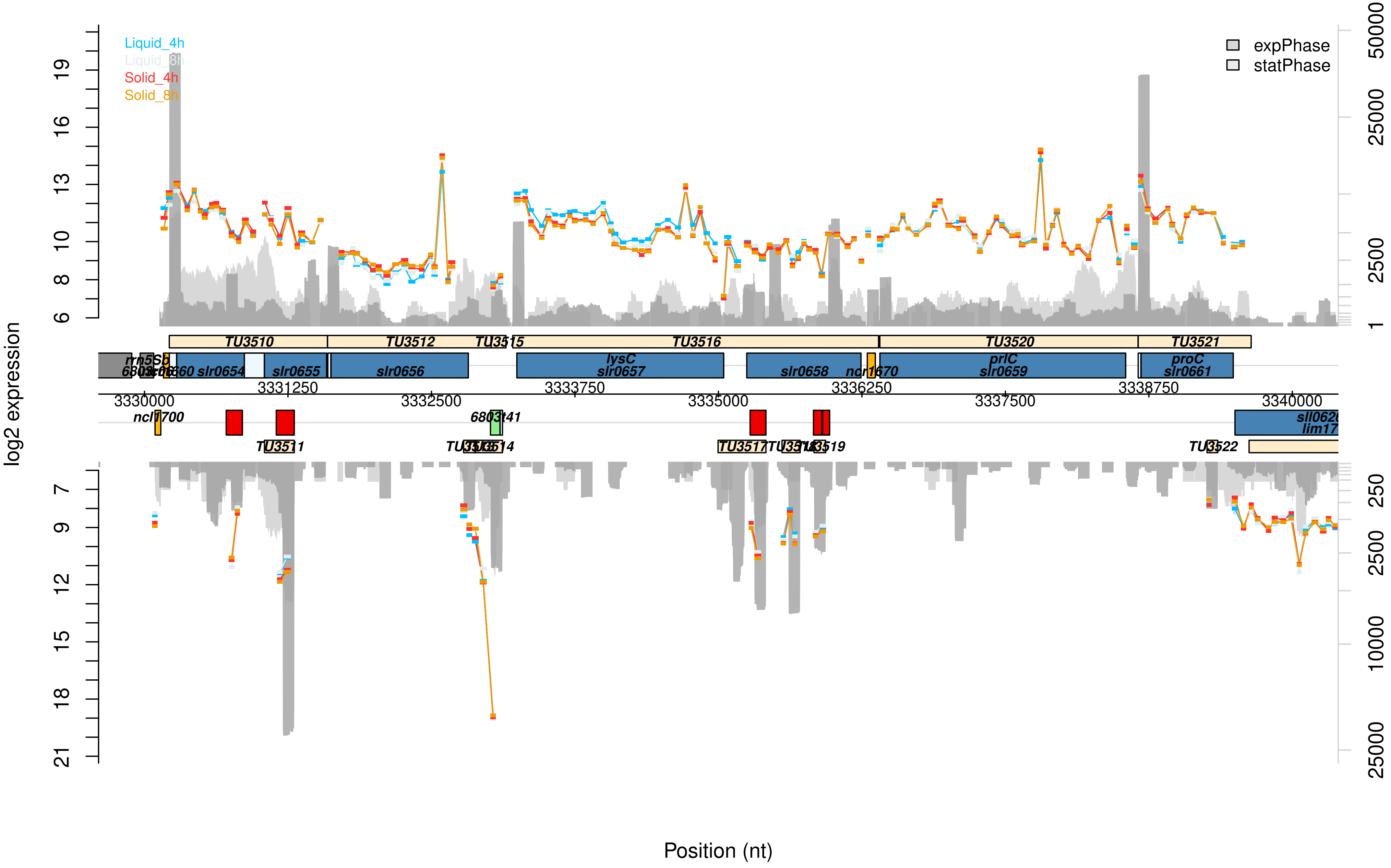

Liquid_4h
Liquid_8h
Solid_4h
Solid_8h
expPhase
statPhase
log2 expression
Position (nt)
TU3510
TU3512
TU3515
TU3516
TU3520
TU3521
TU3511
TU3514
TU3517
TU3519
TU3522
slr0654
slr0655
slr0656
lysC
slr0657
slr0658
prlC
slr0659
proC
slr0661
ncl1700
6803t41
3330000
3331250
3332500
3333750
3335000
3336250
3337500
3338750
3340000

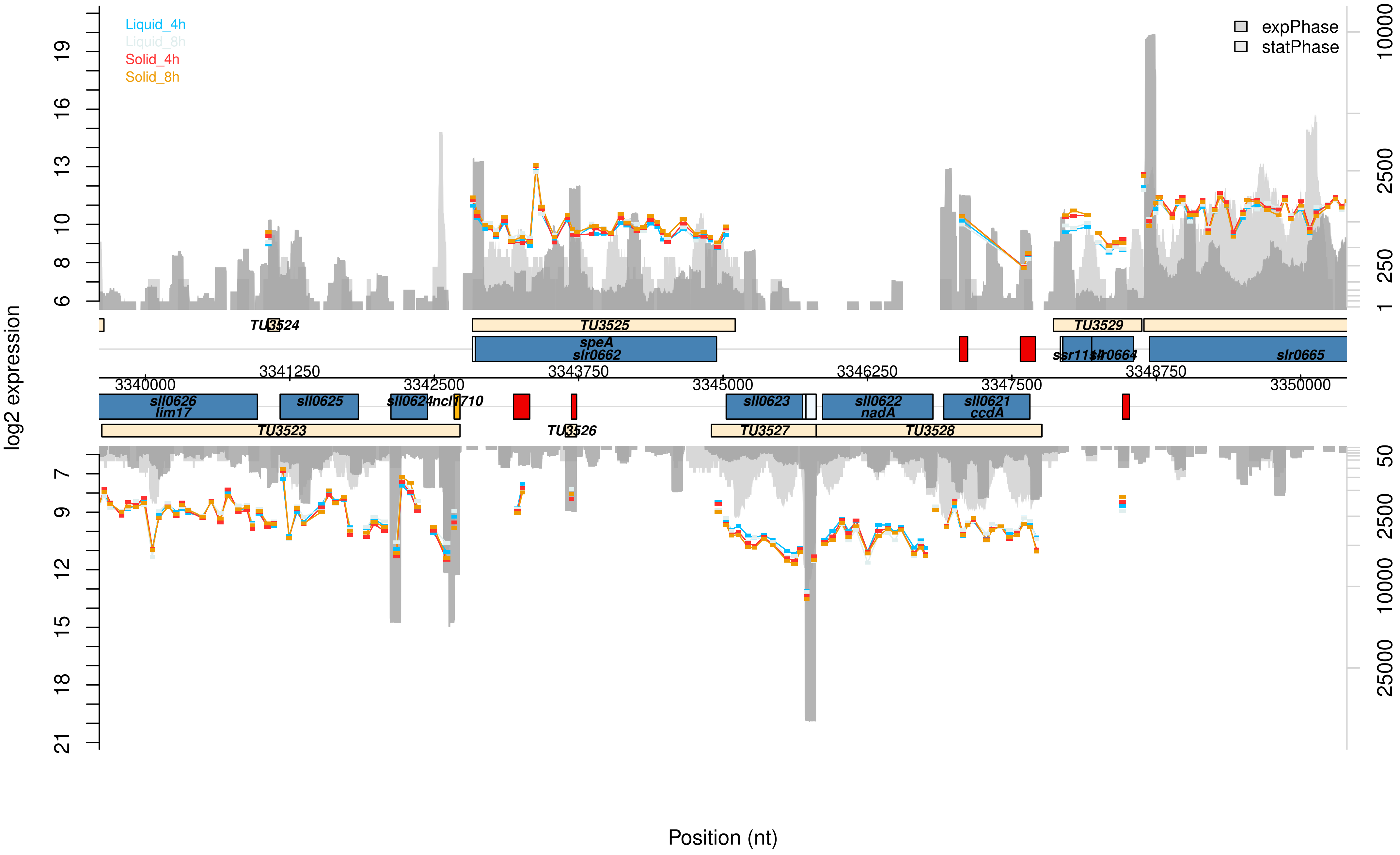
Liquid_4h
Liquid_8h
Solid_4h
Solid_8h
expPhase
statPhase
log2 expression
Position (nt)
TU3524
TU3525
TU3529
speA
slr0662
ssr1154
slr0664
slr0665
3340000
3341250
3342500
3343750
3345000
3346250
3347500
3348750
3350000
sll0626
lim17
sll0625
sll0624
ncl1710
sll0623
sll0622
nadA
sll0621
ccdA
TU3523
TU3526
TU3527
TU3528

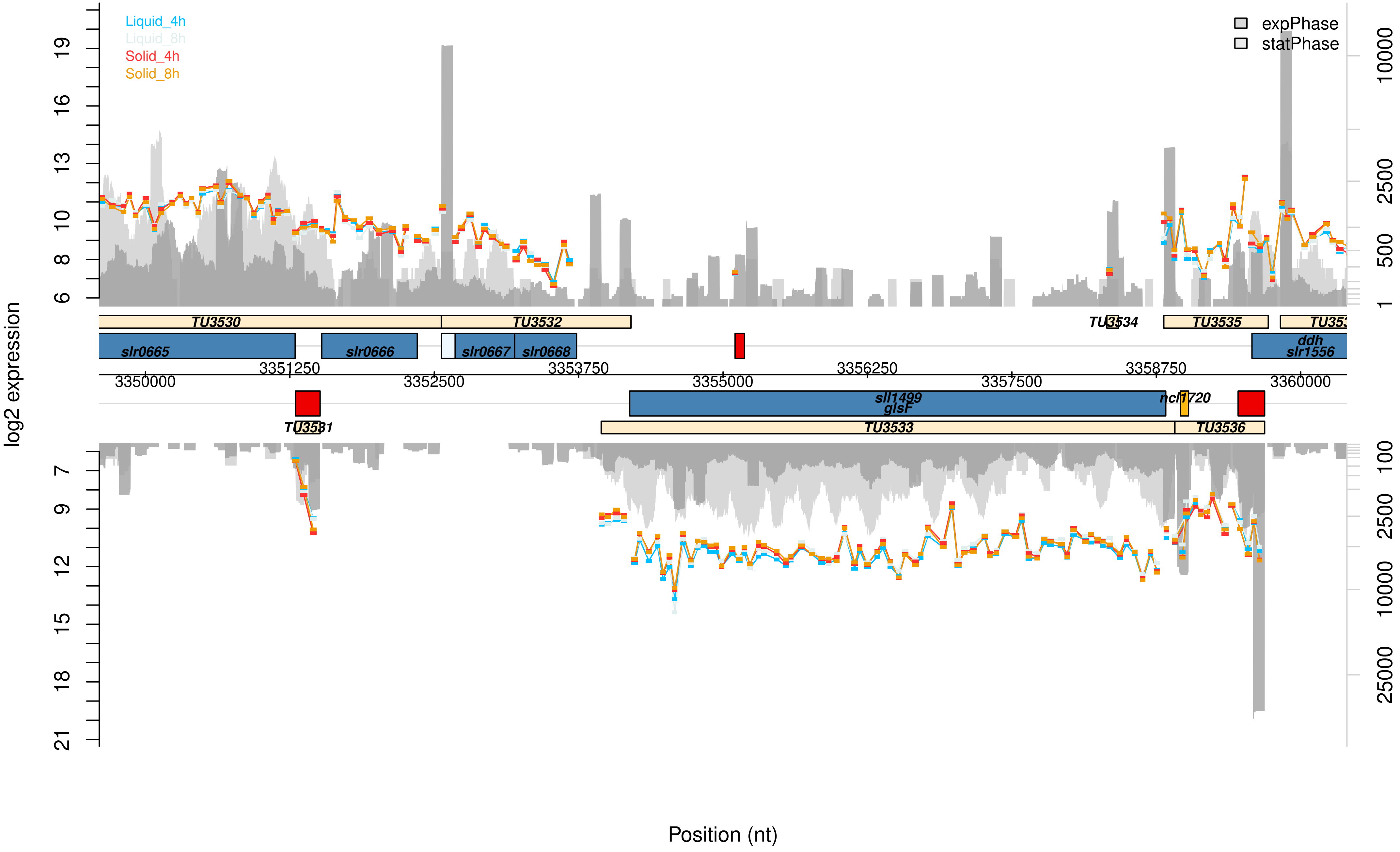

Liquid_4h
Liquid_8h
Solid_4h
Solid_8h
expPhase
statPhase
log2 expression
TU3530
TU3532
TU3534
TU3535
slr0665
slr0666
slr0667
slr0668
ddh
slr1556
3350000
3351250
3352500
3353750
3355000
3356250
3357500
3358750
3360000
sll1499
glsF
ncl1720
TU3531
TU3533
TU3536
Position (nt)

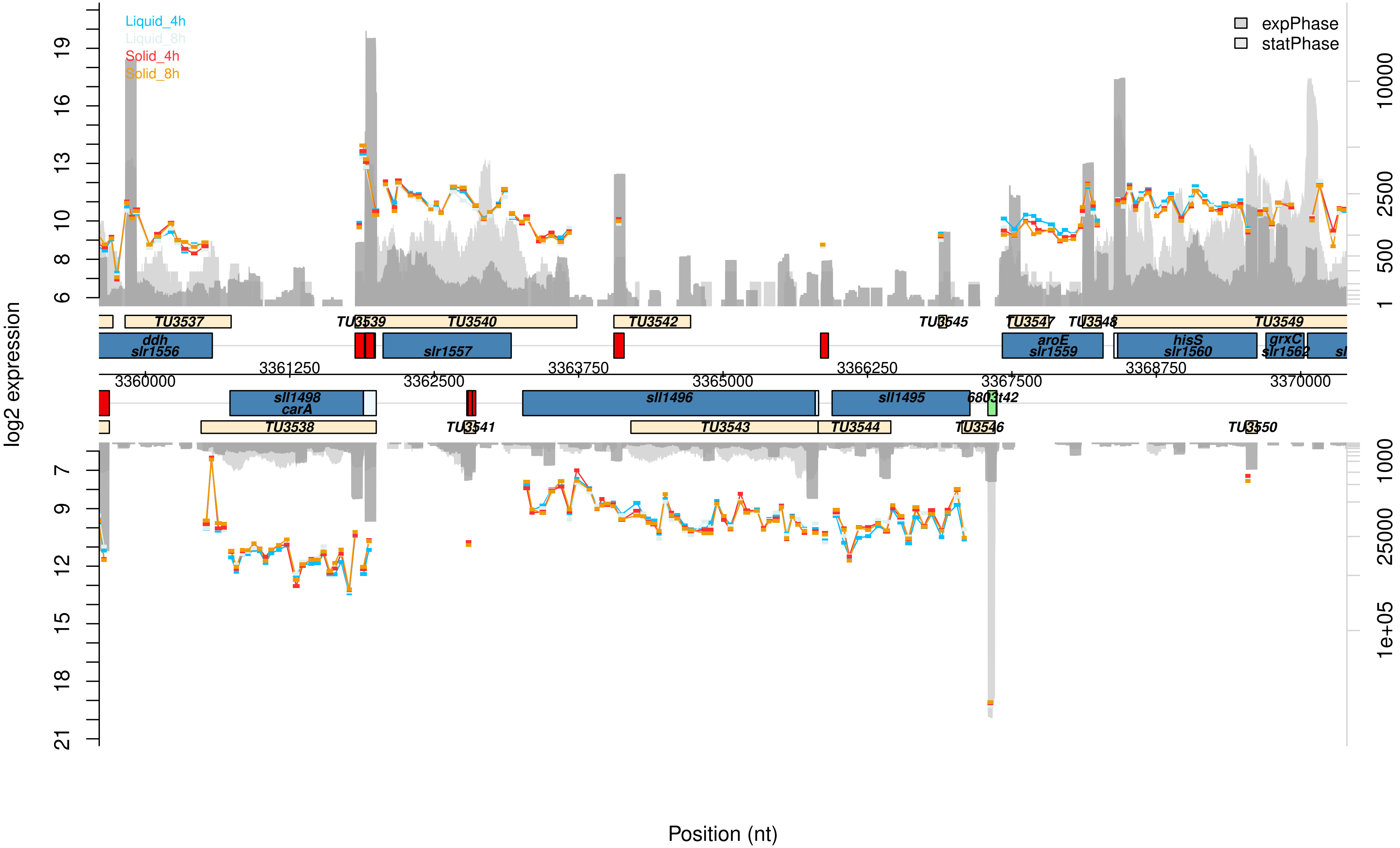

Liquid_4h
Liquid_8h
Solid_4h
Solid_8h
expPhase
statPhase
log2 expression
Position (nt)
TU3537
TU3539
TU3540
TU3542
TU3545
TU3547
TU3548
TU3549
ddh
slr1556
slr1557
aroE
slr1559
hisS
slr1560
grxC
slr1562
3361250
3363750
3366250
3368750
3360000
3362500
3365000
3367500
3370000
sll1498
carA
sll1496
sll1495
6803t42
TU3538
TU3541
TU3543
TU3544
TU3546
TU3550

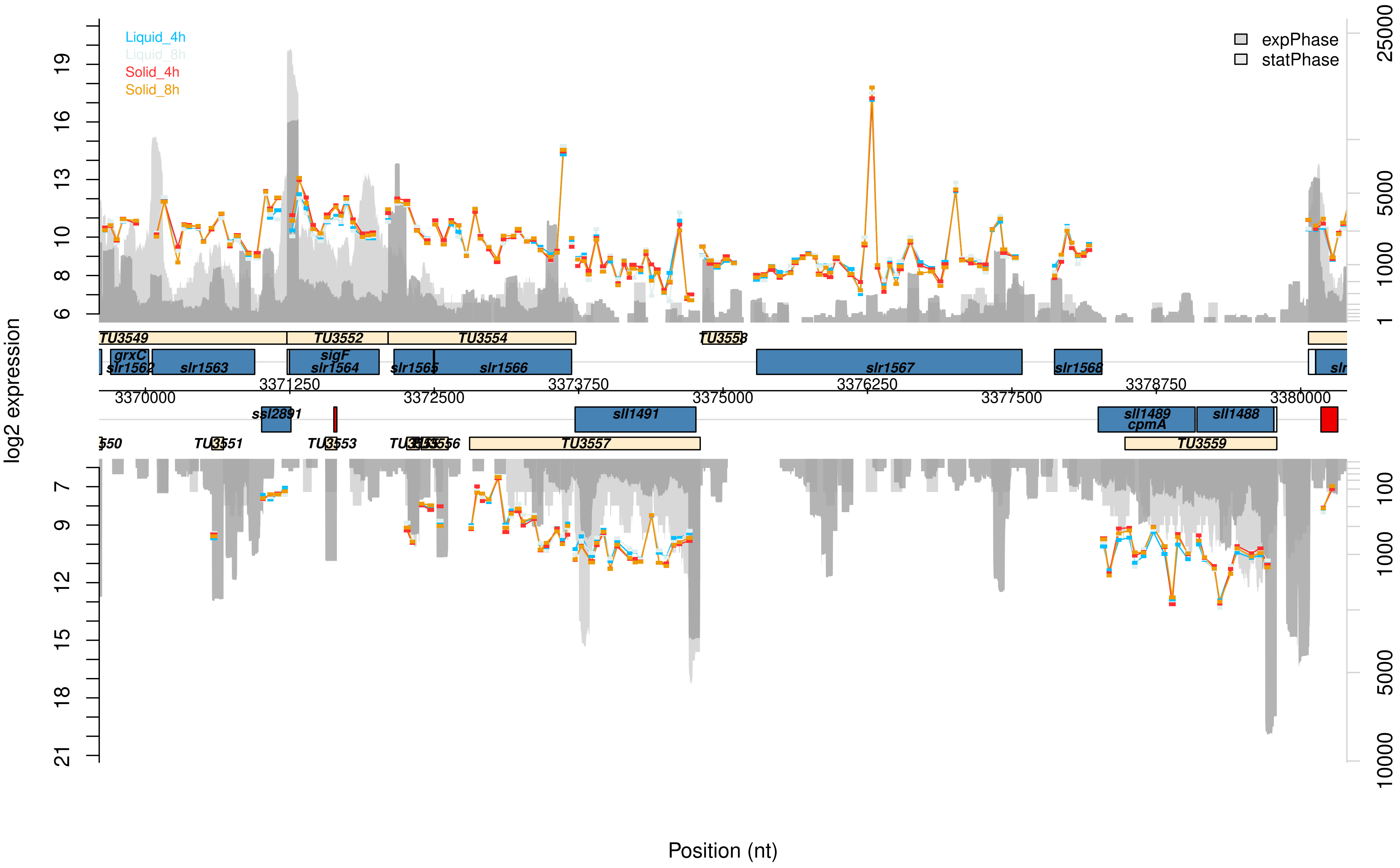

Liquid_4h
Liquid_8h
Solid_4h
Solid_8h
expPhase
statPhase
log2 expression
Position (nt)
TU3549
TU3552
TU3554
TU3558
grxC
slr1562
slr1563
sigF
slr1564
slr1565
slr1566
slr1567
slr1568
ssl2891
sll1491
sll1489
cpmA
sll1488
TU3551
TU3553
TU3557
TU3559
3370000
3371250
3372500
3373750
3375000
3376250
3377500
3378750
3380000

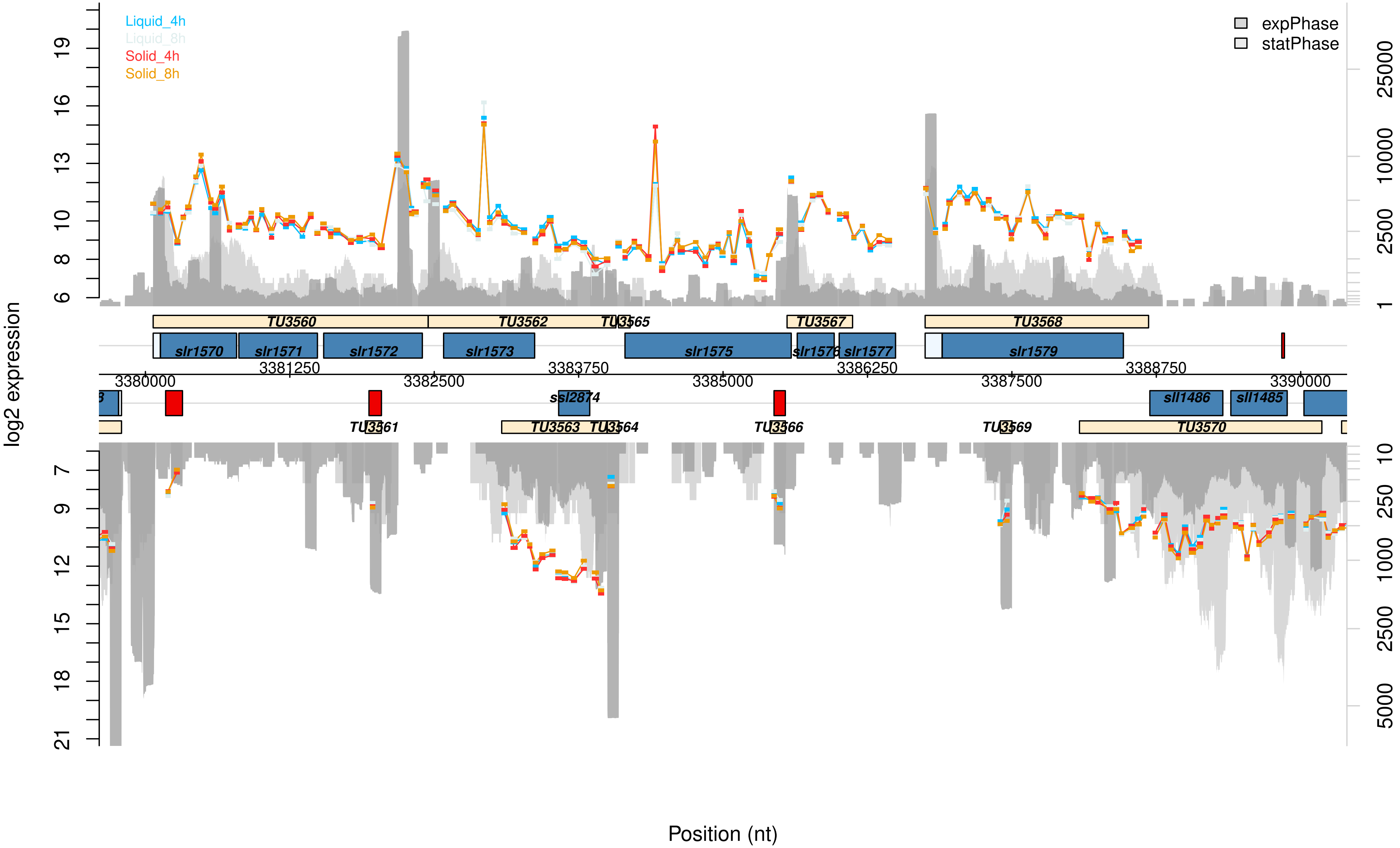

Liquid_4h
Liquid_8h
Solid_4h
Solid_8h
expPhase
statPhase
log2 expression
TU3560
TU3562
TU3565
TU3567
TU3568
slr1570
slr1571
slr1572
slr1573
slr1575
slr1576
slr1577
slr1579
3380000
3381250
3382500
3383750
3385000
3386250
3387500
3388750
3390000
ssl2874
sll1486
sll1485
TU3561
TU3563
TU3564
TU3566
TU3569
TU3570
Position (nt)

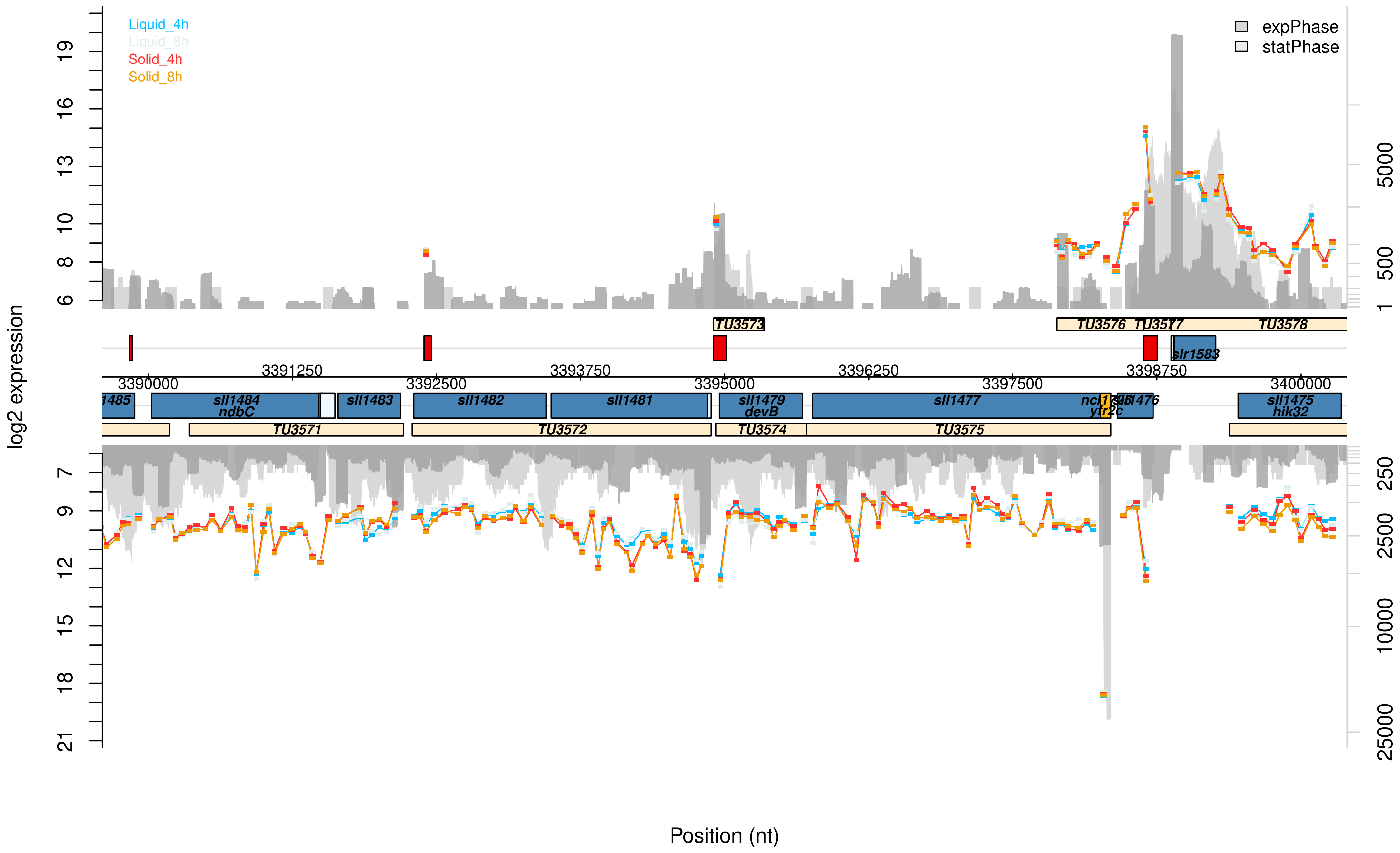

Liquid_4h
Liquid_8h
Solid_4h
Solid_8h
expPhase
statPhase
log2 expression
Position (nt)
3390000
3391250
3392500
3393750
3395000
3396250
3397500
3398750
3400000
sll1484 ndbC
sll1483
sll1482
sll1481
sll1479 devB
sll1477
sll1475 hik32
slr1583
TU3571
TU3572
TU3573
TU3574
TU3575
TU3576
TU3578

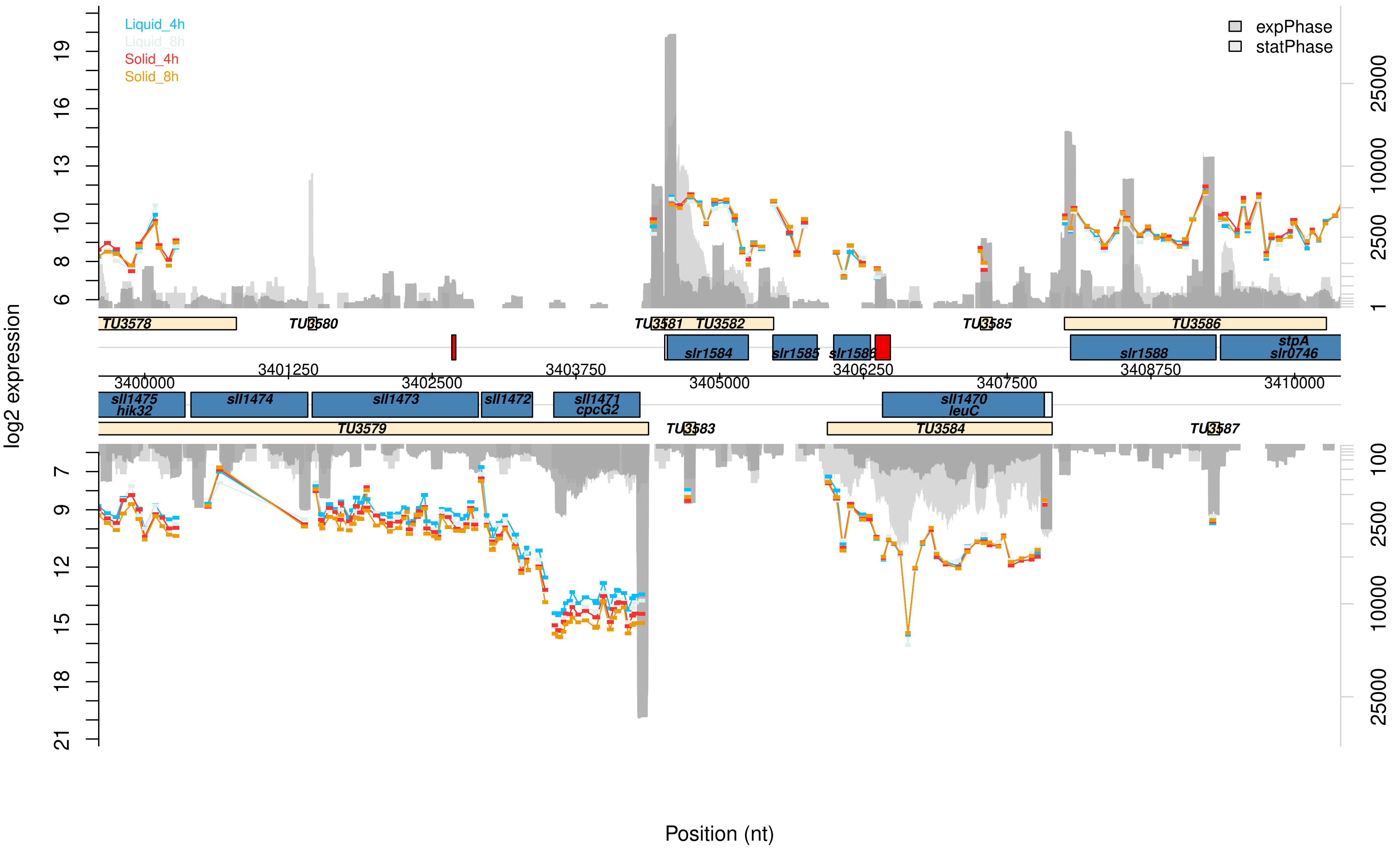

Liquid_4h
Liquid_8h
Solid_4h
Solid_8h
expPhase
statPhase
log2 expression
Position (nt)
TU3578
TU3580
TU3581
TU3582
TU3585
TU3586
slr1584
slr1585
slr1586
slr1588
stpA
slr0746
sll1475
hik32
sll1474
sll1473
sll1472
sll1471
cpcG2
sll1470
leuC
TU3579
TU3583
TU3584
TU3587
3400000
3401250
3402500
3403750
3405000
3406250
3407500
3408750
3410000

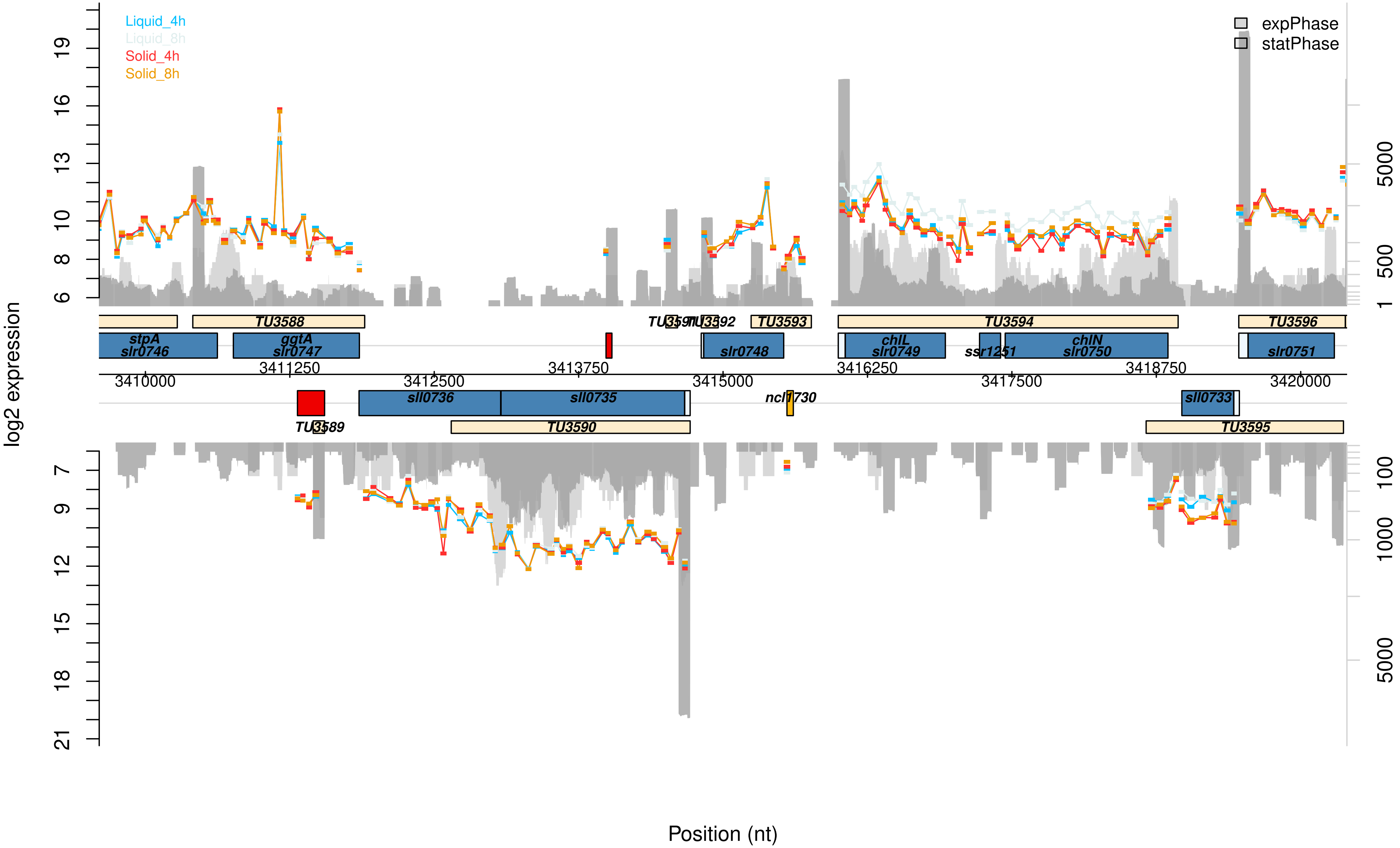

Liquid_4h
Liquid_8h
Solid_4h
Solid_8h
expPhase
statPhase
log2 expression
6
8
10
13
16
19
7
9
12
15
18
21
1
500
5000
100
1000
5000
TU3588
TU3593
TU3594
TU3596
stpA
slr0746
ggtA
slr0747
slr0748
chlL
slr0749
chlN
slr0750
slr0751
3410000
3411250
3412500
3413750
3415000
3416250
3417500
3418750
3420000
sll0736
sll0735
sll0733
TU3589
TU3590
TU3595
Position (nt)

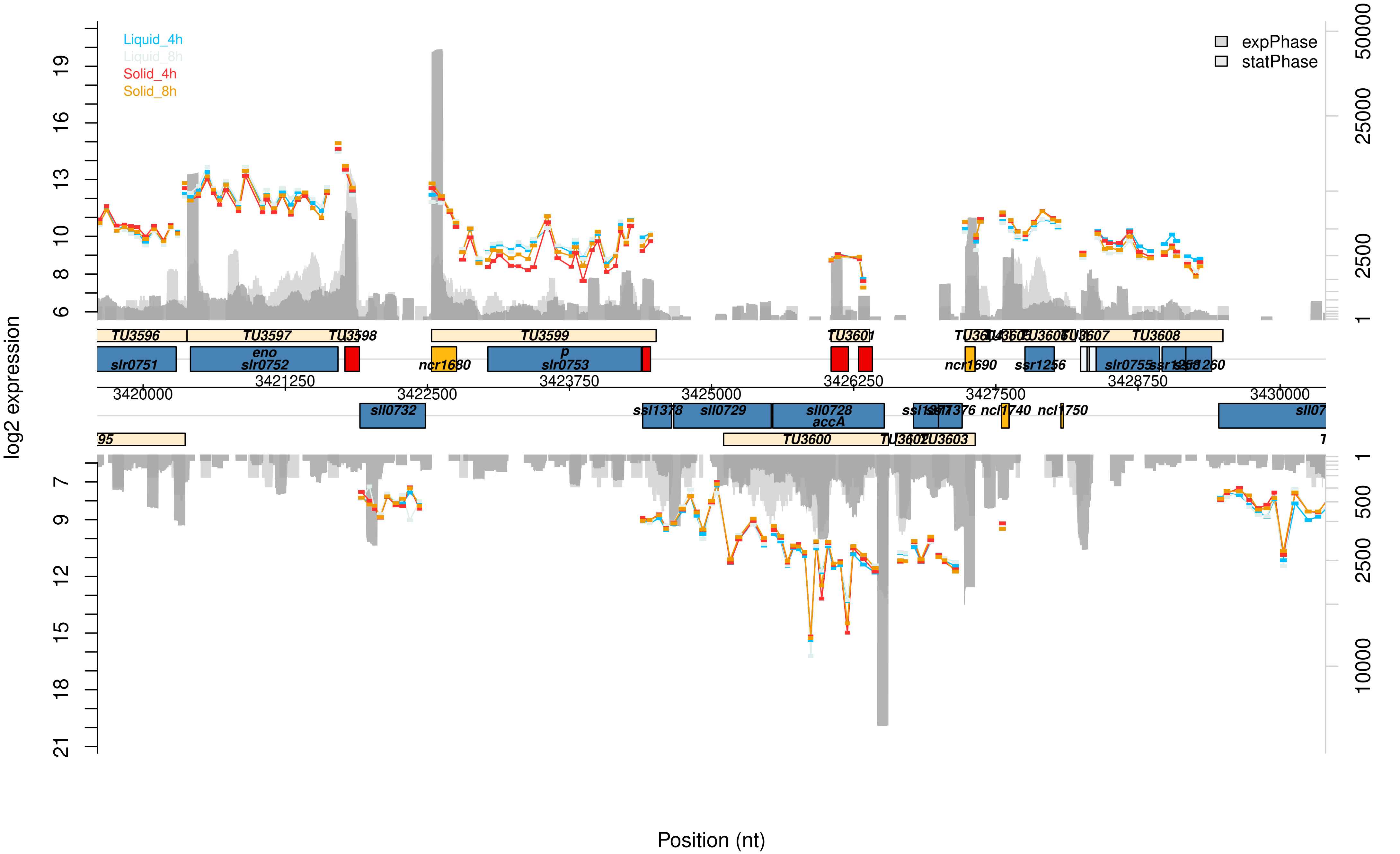

Liquid_4h
Liquid_8h
Solid_4h
Solid_8h
expPhase
statPhase
log2 expression
Position (nt)
TU3596
TU3597
TU3598
TU3599
TU3600
TU3601
TU3603
TU3608
slr0751
eno
slr0752
ncr1680
p
slr0753
ncr1690
ssr1256
slr0755
sll0732
ssl1378
sll0729
sll0728
accA
ncl1740
ncl1750
3420000
3421250
3422500
3423750
3425000
3426250
3427500
3428750
3430000

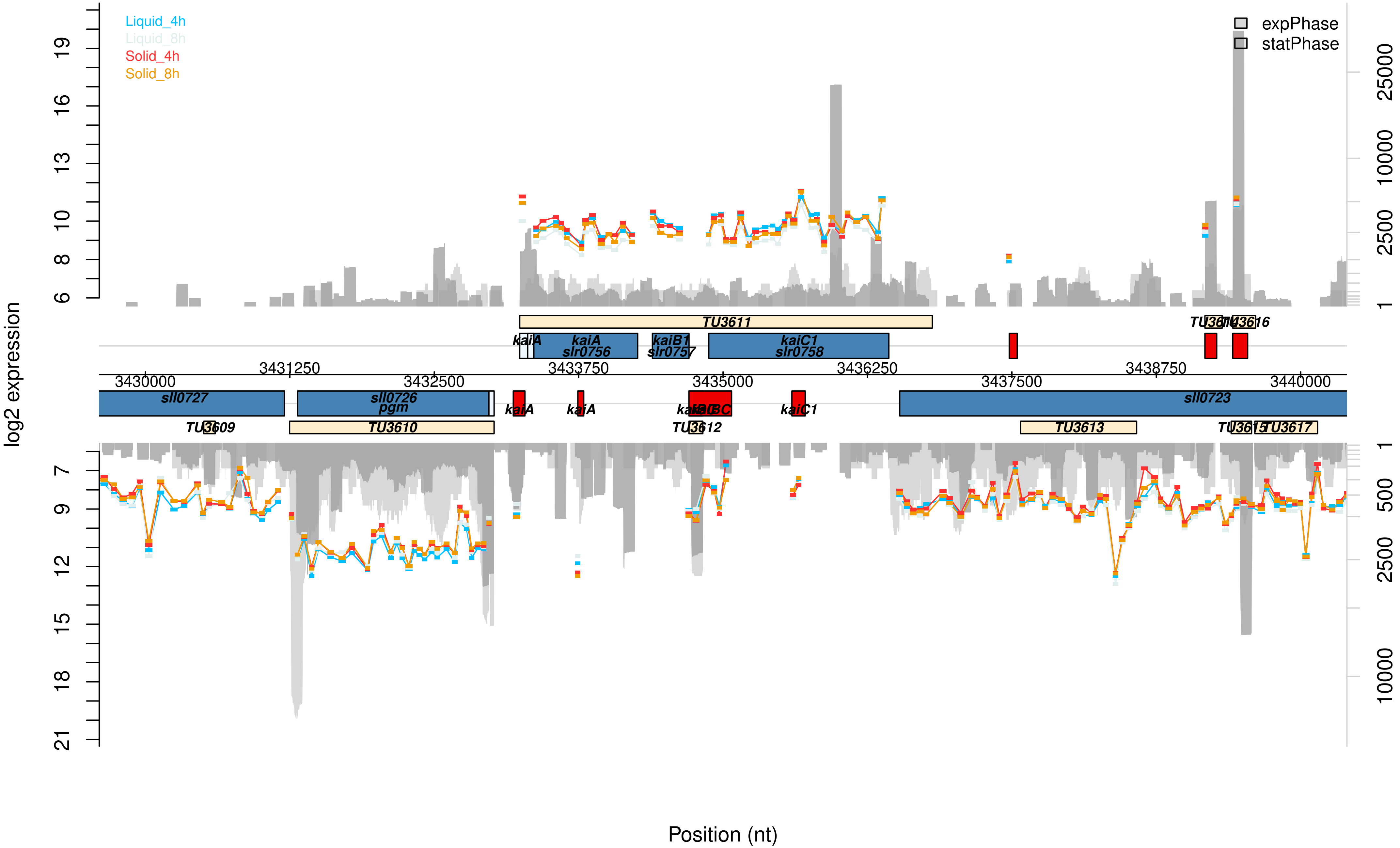

Liquid_4h
Liquid_8h
Solid_4h
Solid_8h
expPhase
statPhase
log2 expression
Position (nt)
TU3611
kaiA
kaiA slr0756
kaiB1 slr0757
kaiC1 slr0758
sll0727
sll0726 pgm
kaiA
kaiA
kaiBC
kaiC1
sll0723
TU3609
TU3610
TU3612
TU3613
TU3617
3430000
3431250
3432500
3433750
3435000
3436250
3437500
3438750
3440000

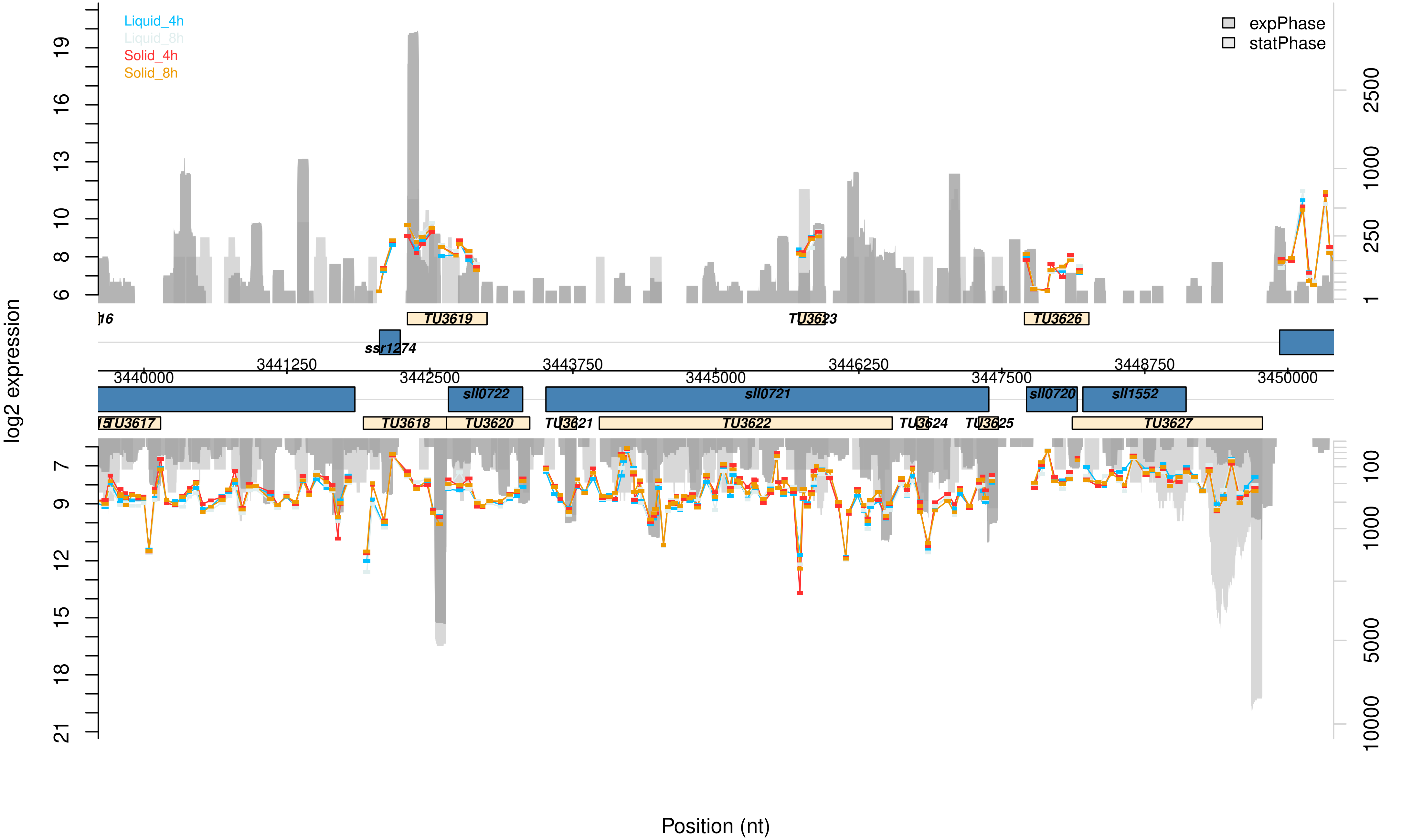

Liquid_4h
Liquid_8h
Solid_4h
Solid_8h
expPhase
statPhase
log2 expression
Position (nt)
TU3619
TU3623
TU3626
ssr1274
sll0722
sll0721
sll0720
sll1552
TU3617
TU3618
TU3620
TU3621
TU3622
TU3624
TU3625
TU3627
3440000
3441250
3442500
3443750
3445000
3446250
3447500
3448750
3450000

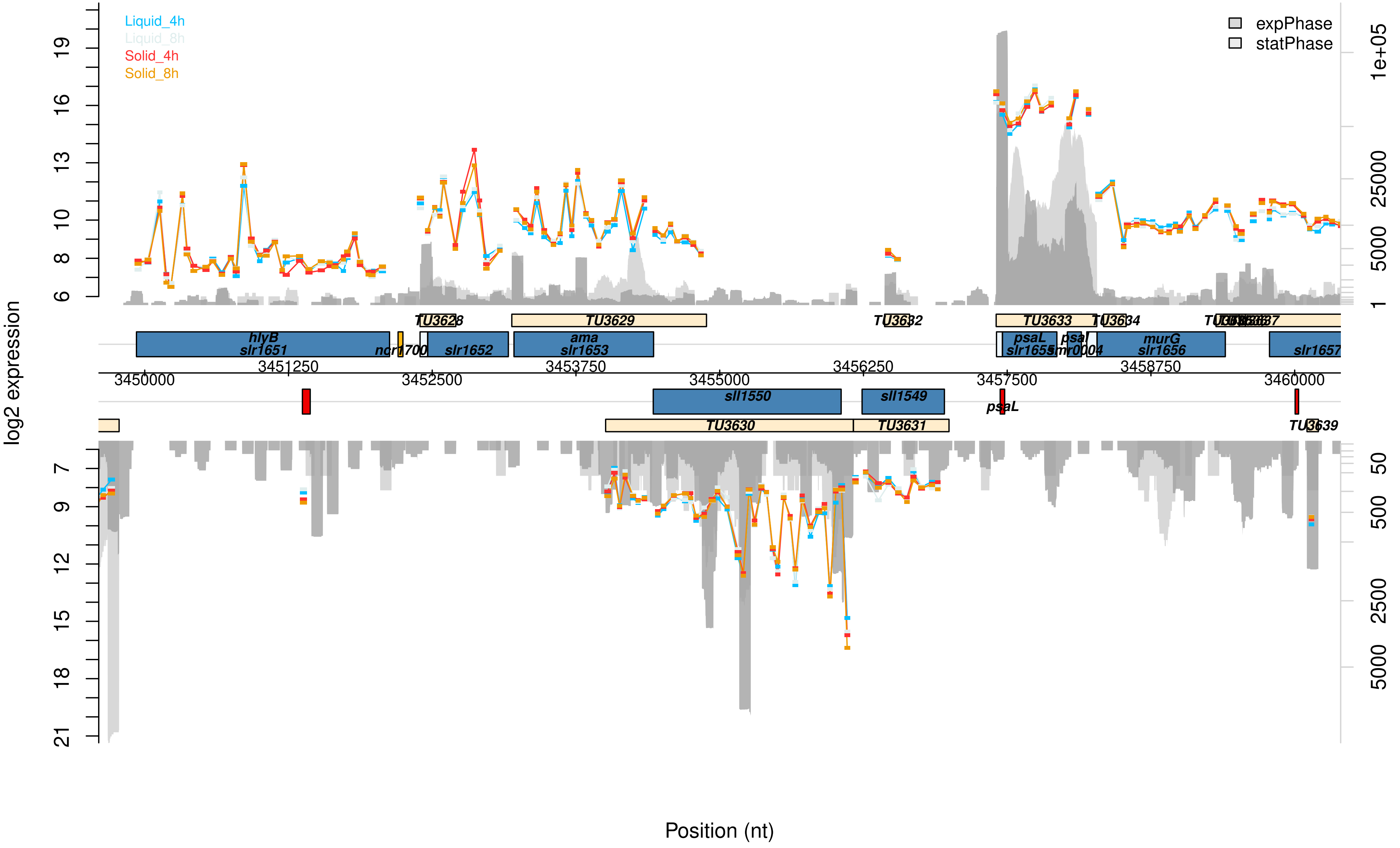
Liquid_4h
Liquid_8h
Solid_4h
Solid_8h
expPhase
statPhase
log2 expression
Position (nt)
TU3628
TU3629
TU3632
TU3633
TU3634
TU3630
TU3631
TU3639
hlyB
slr1651
ncr1700
slr1652
ama
slr1653
psaL
slr1655
psal
smr0004
murG
slr1656
slr1657
sll1550
sll1549
3450000
3451250
3452500
3453750
3455000
3456250
3457500
3458750
3460000

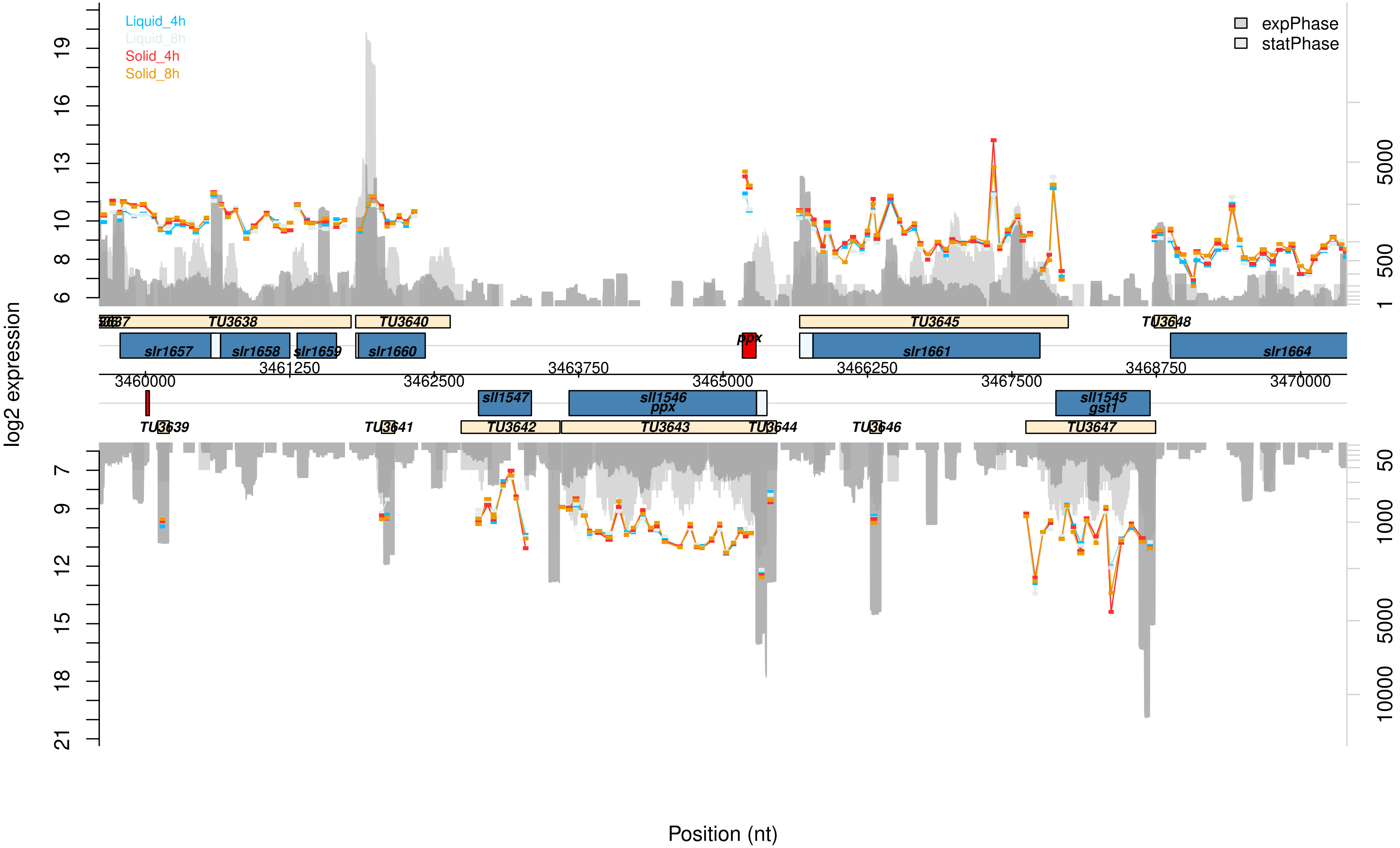

Liquid_4h
Liquid_8h
Solid_4h
Solid_8h
expPhase
statPhase
log2 expression
6
8
10
13
16
19
7
9
12
15
18
21
1
500
5000
50
1000
5000
10000
TU3638
TU3640
TU3645
TU3648
slr1657
slr1658
slr1659
slr1660
ppx
slr1661
slr1664
3460000
3461250
3462500
3463750
3465000
3466250
3467500
3468750
3470000
sll1547
sll1546
ppx
sll1545
gst1
TU3639
TU3641
TU3642
TU3643
TU3644
TU3646
TU3647
Position (nt)

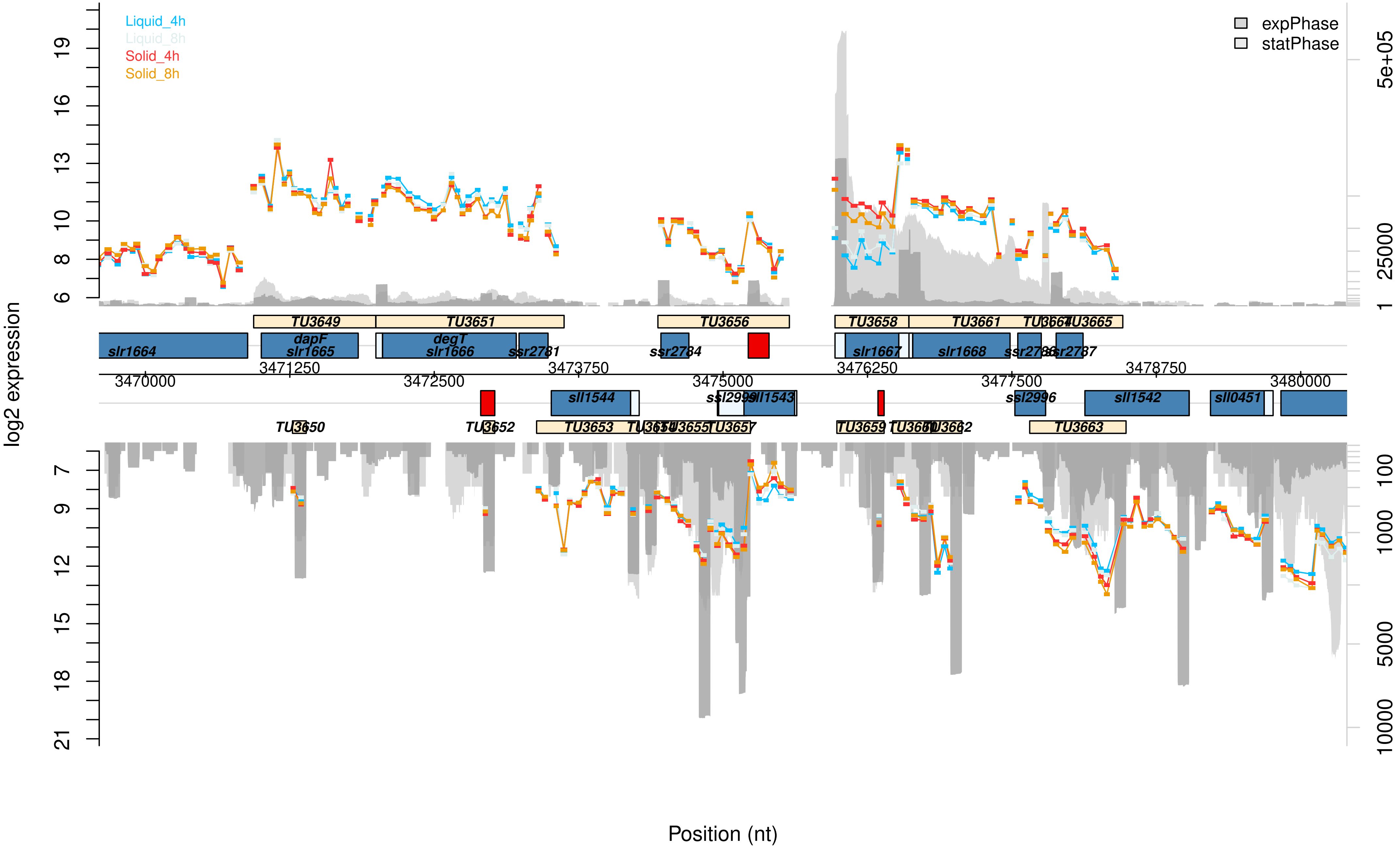
Liquid_4h
Liquid_8h
Solid_4h
Solid_8h
expPhase
statPhase
log2 expression
Position (nt)
TU3649
TU3651
TU3656
TU3658
TU3661
TU3664
TU3665
slr1664
dapF
slr1665
degT
slr1666
ssr2781
ssr2784
slr1667
slr1668
ssr2786
ssr2787
3470000
3471250
3472500
3473750
3475000
3476250
3477500
3478750
3480000
sll1544
ssl2999
sll1543
ssl2996
sll1542
sll0451
TU3650
TU3652
TU3653
TU3654
TU3655
TU3657
TU3659
TU3660
TU3662
TU3663

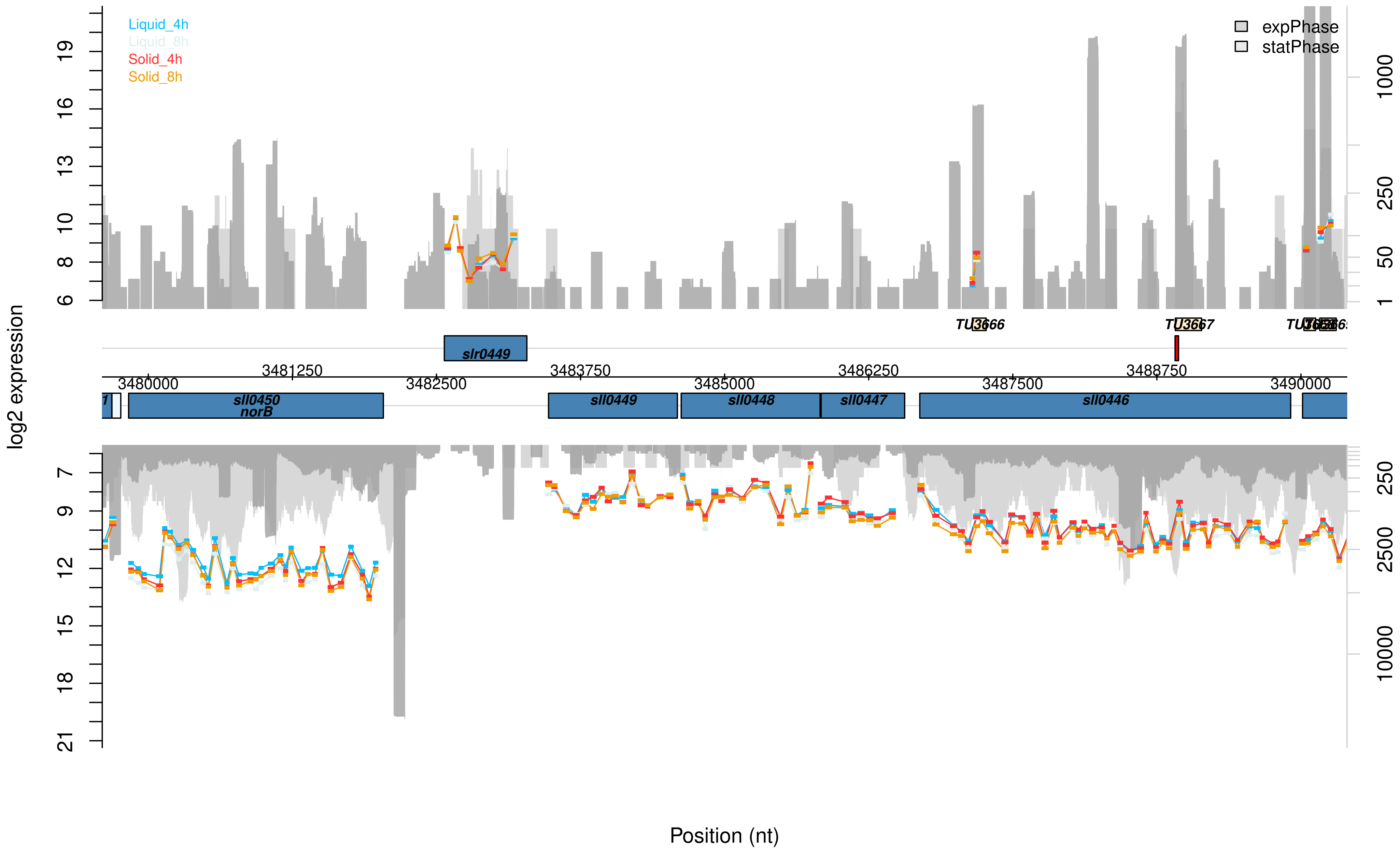

Liquid_4h
Liquid_8h
Solid_4h
Solid_8h
expPhase
statPhase
log2 expression
19
16
13
10
8
6
1000
250
50
1
TU3666
TU3667
slr0449
3480000
3481250
3482500
3483750
3485000
3486250
3487500
3488750
3490000
sll0450
norB
sll0449
sll0448
sll0447
sll0446
7
9
12
15
18
21
250
2500
10000
Position (nt)

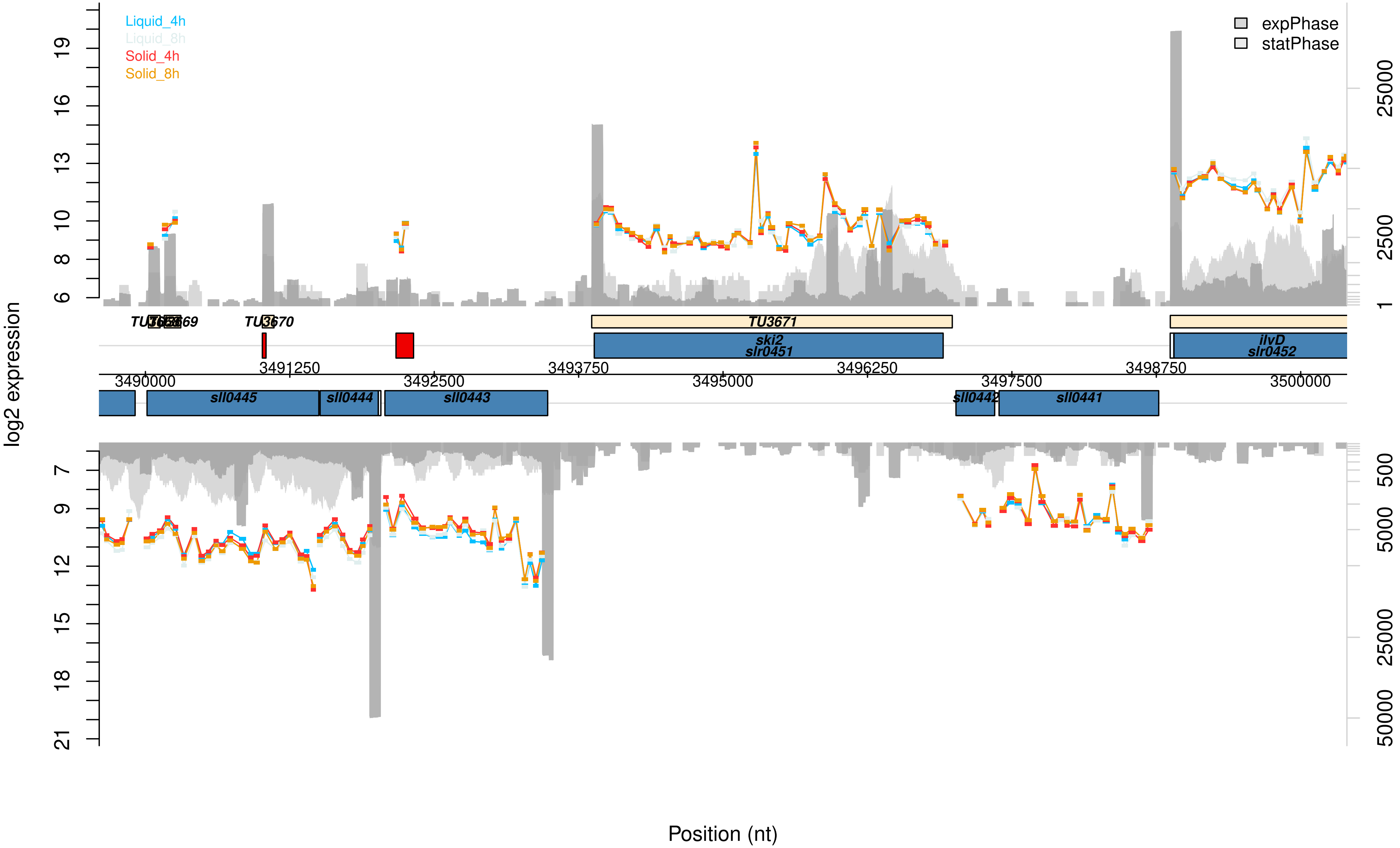

Liquid_4h
Liquid_8h
Solid_4h
Solid_8h
expPhase
statPhase
log2 expression
TU3670
TU3671
ski2
slr0451
ilvD
slr0452
sll0445
sll0444
sll0443
sll0442
sll0441
3490000
3491250
3492500
3493750
3495000
3496250
3497500
3498750
3500000
Position (nt)

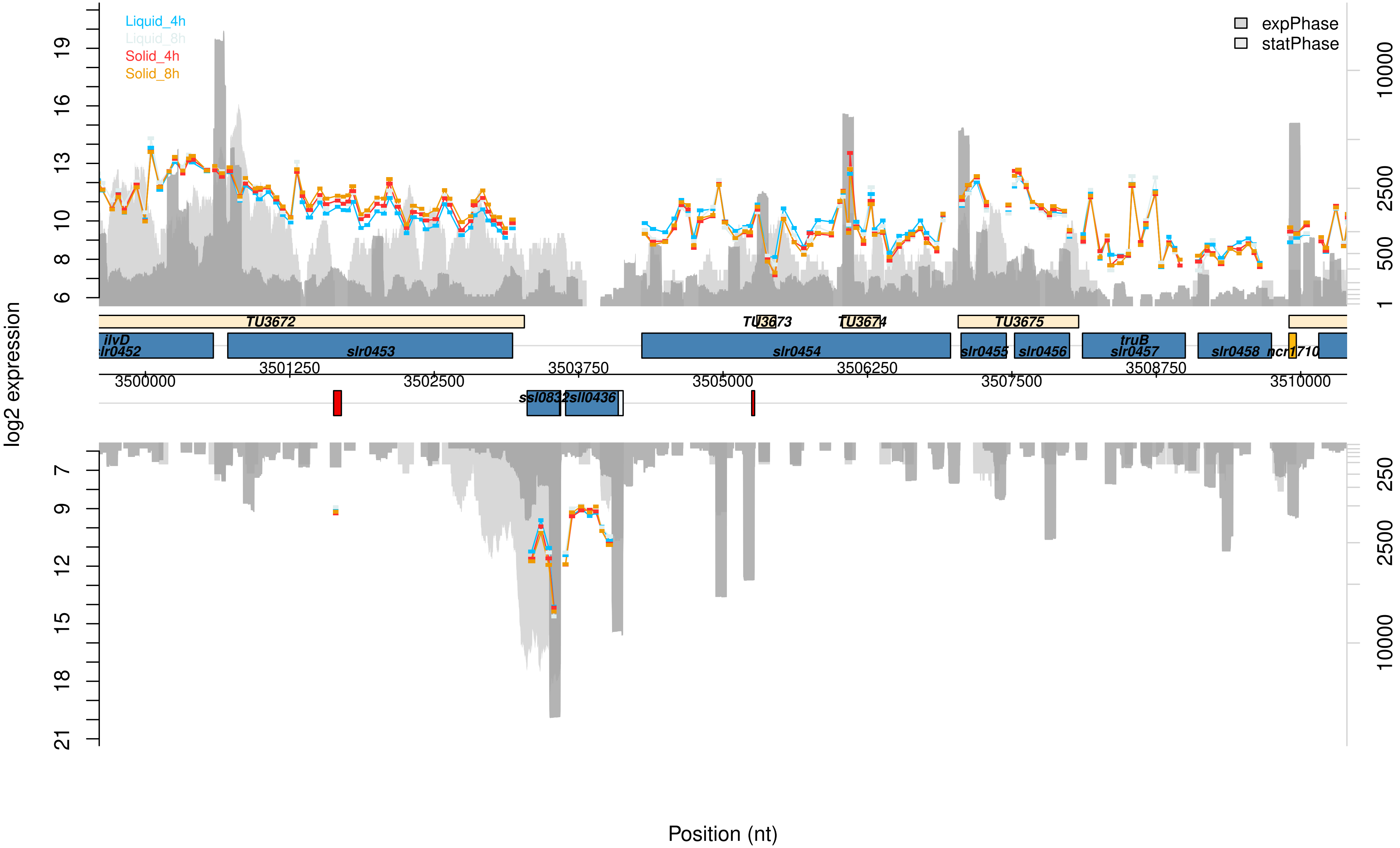

Liquid_4h
Liquid_8h
Solid_4h
Solid_8h
expPhase
statPhase
log2 expression
Position (nt)
TU3672
TU3673
TU3674
TU3675
ilvD
slr0452
slr0453
slr0454
slr0455
slr0456
truB
slr0457
slr0458
ncr1710
ssl0832
sll0436
3500000
3501250
3502500
3503750
3505000
3506250
3507500
3508750
3510000

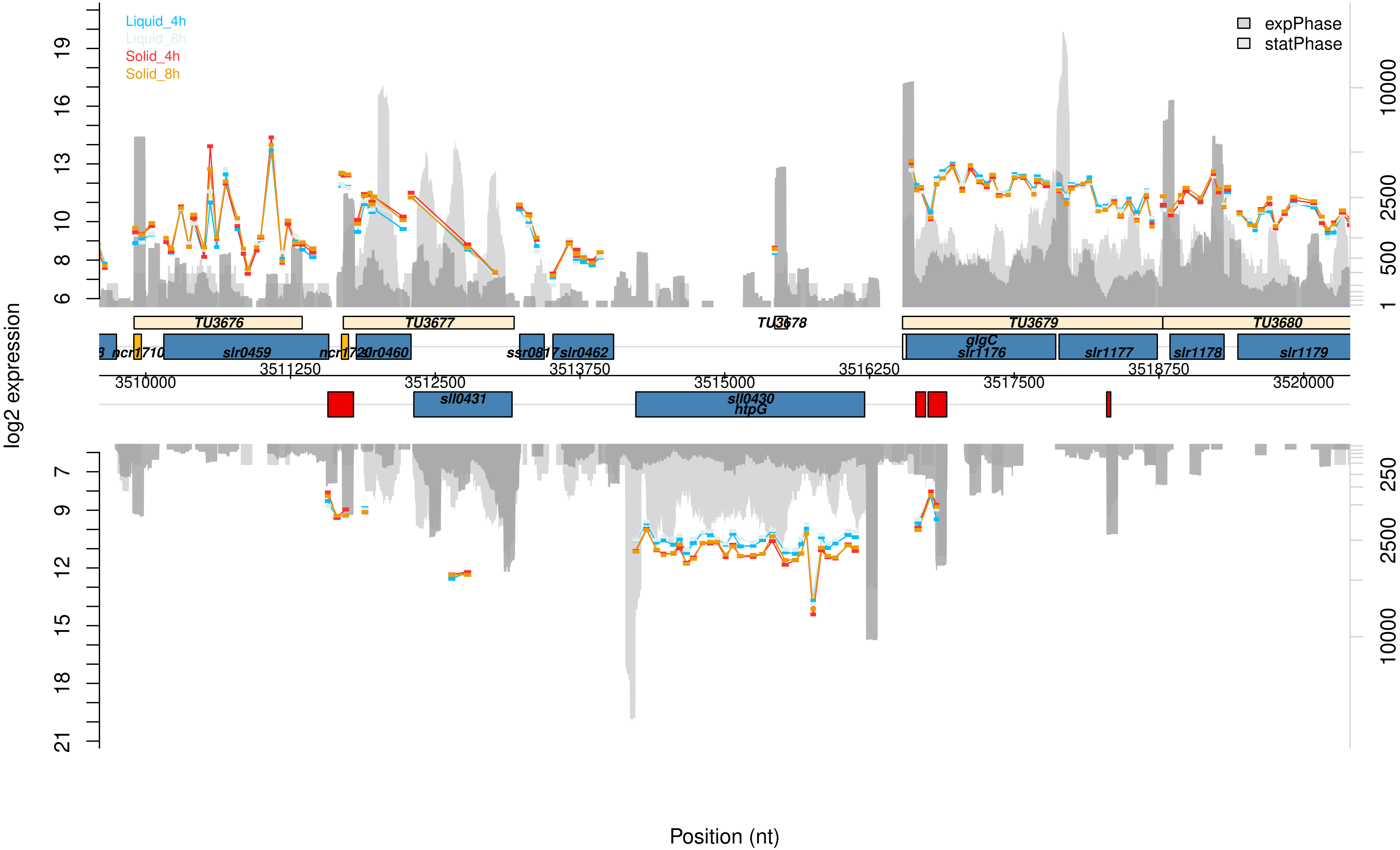

Liquid_4h
Liquid_8h
Solid_4h
Solid_8h
expPhase
statPhase
log2 expression
Position (nt)
TU3676
TU3677
TU3678
TU3679
TU3680
ncr1710
slr0459
ncr1720
slr0460
ssr0817
slr0462
glgC
slr1176
slr1177
slr1178
slr1179
sll0431
sll0430
htpG
3510000
3511250
3512500
3513750
3515000
3516250
3517500
3518750
3520000

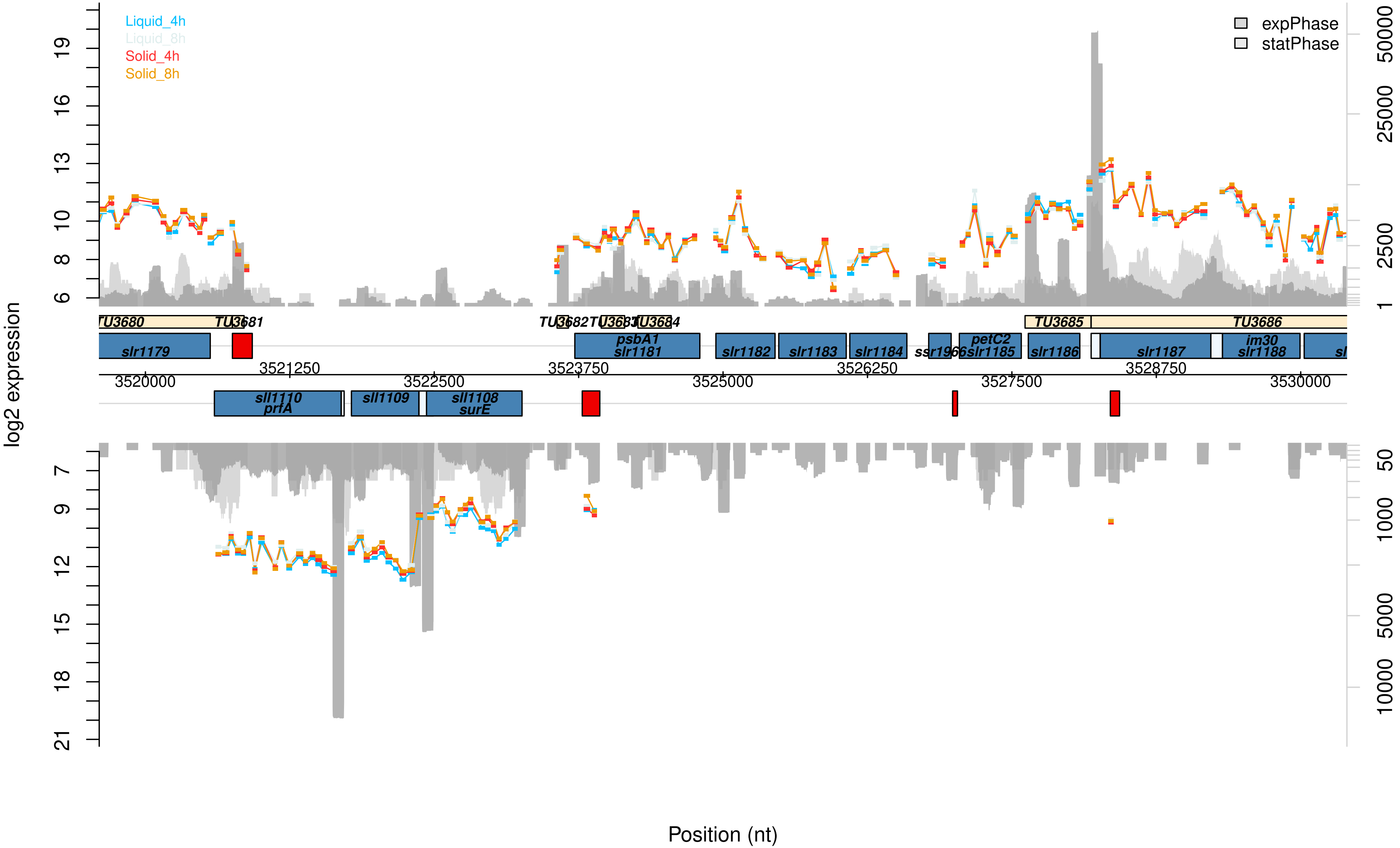

Liquid_4h
Liquid_8h
Solid_4h
Solid_8h
expPhase
statPhase
log2 expression
Position (nt)
TU3680
TU3681
TU3682
TU3683
TU3684
TU3685
TU3686
slr1179
psbA1
slr1181
slr1182
slr1183
slr1184
ssr1966
petC2
slr1185
slr1186
slr1187
im30
slr1188
sll1110
prfA
sll1109
sll1108
surE
3520000
3521250
3522500
3523750
3525000
3526250
3527500
3528750
3530000

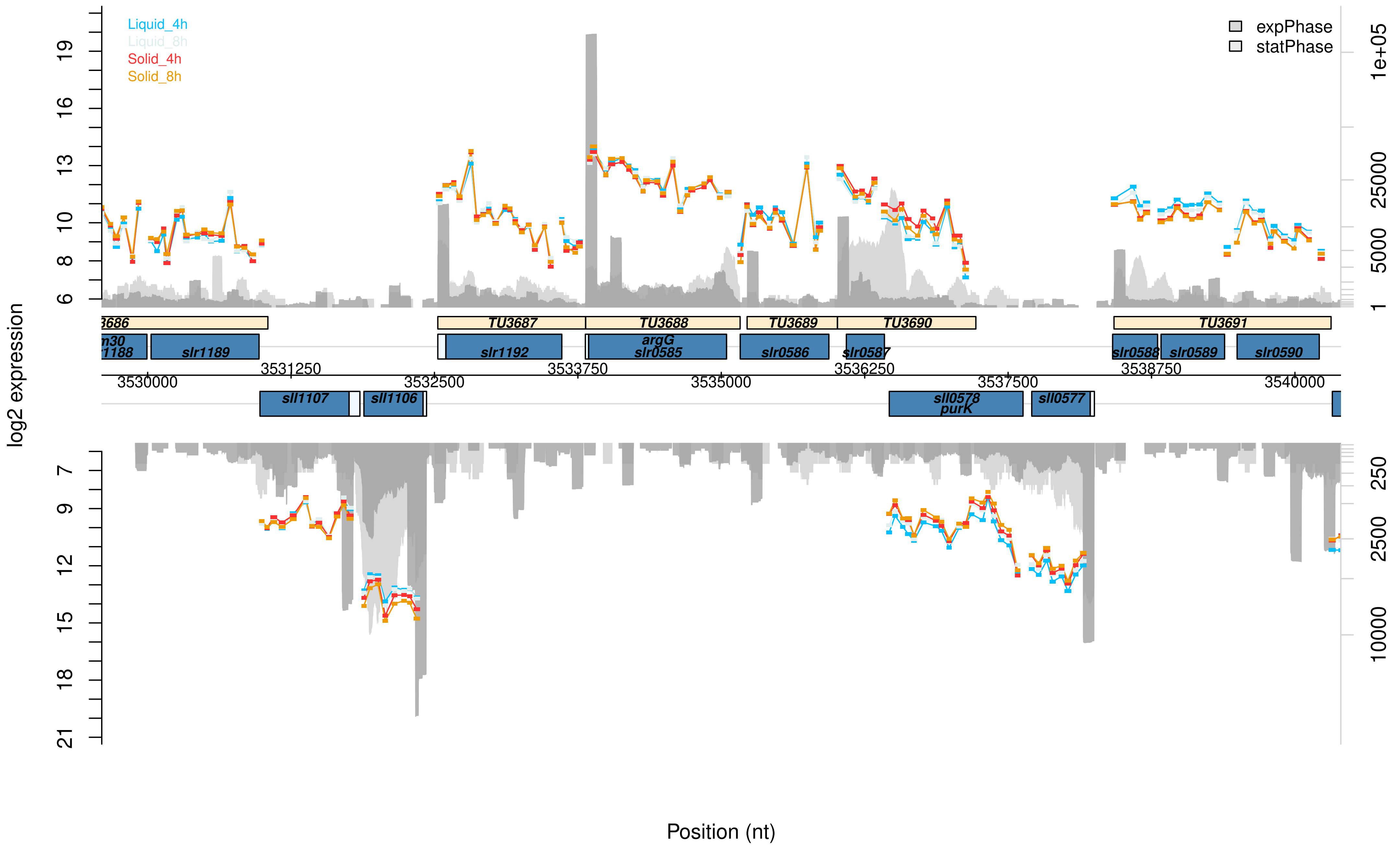

Liquid_4h
Liquid_8h
Solid_4h
Solid_8h
expPhase
statPhase
log2 expression
19
16
13
10
8
6
7
9
12
15
18
21
1e+05
25000
5000
1
250
2500
10000
3686
TU3687
TU3688
TU3689
TU3690
TU3691
m30
r1188
slr1189
slr1192
argG
slr0585
slr0586
slr0587
slr0588
slr0589
slr0590
3531250
3533750
3536250
3538750
3530000
3532500
3535000
3537500
3540000
sll1107
sll1106
sll0578
purK
sll0577
Position (nt)

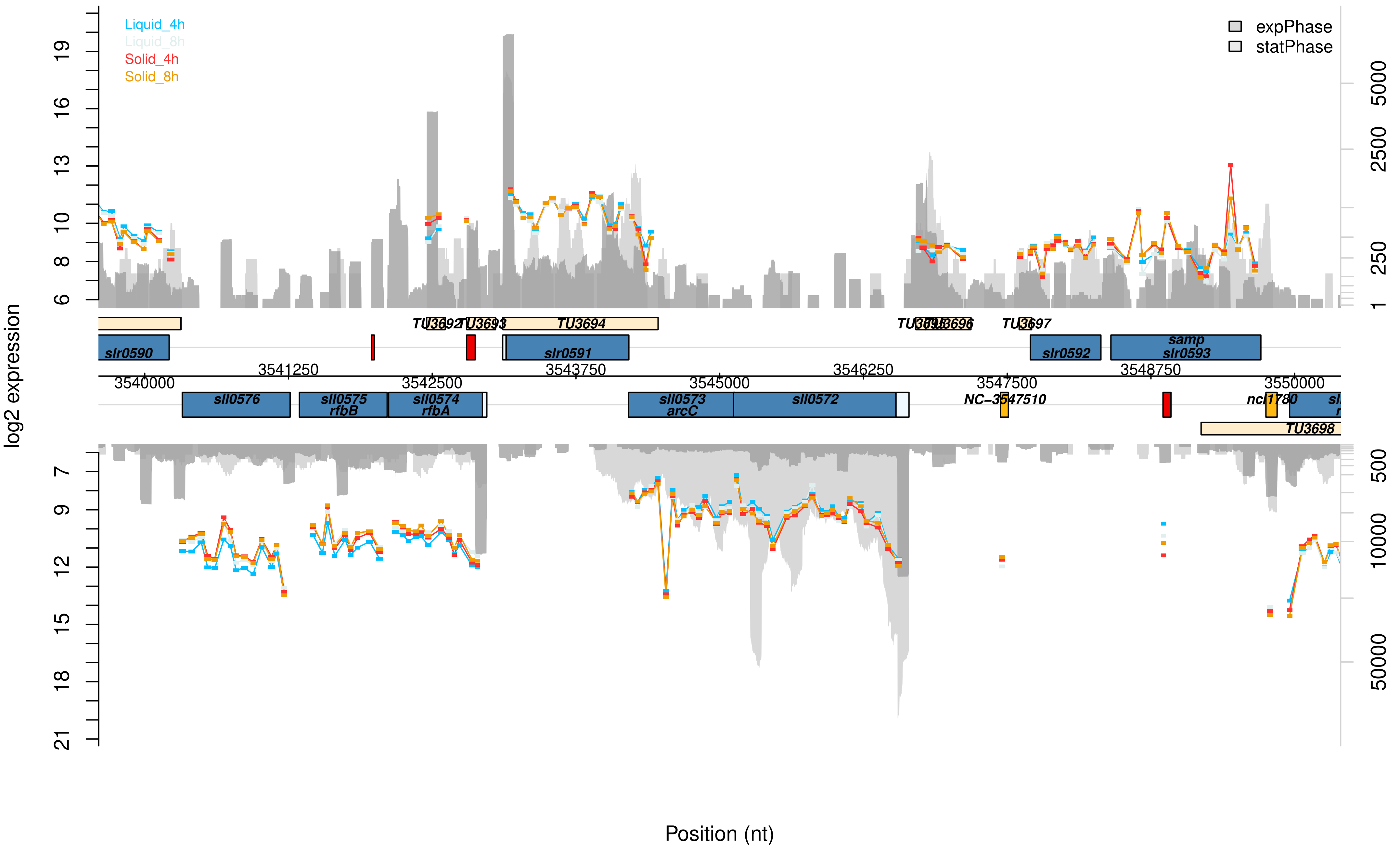

Liquid_4h
Liquid_8h
Solid_4h
Solid_8h
expPhase
statPhase
log2 expression
6
8
10
13
16
19
1
250
2500
5000
TU3692
TU3693
TU3694
TU3695
TU3696
TU3697
slr0590
slr0591
slr0592
samp
slr0593
3540000
3541250
3542500
3543750
3545000
3546250
3547500
3548750
3550000
sll0576
sll0575
rfbB
sll0574
rfbA
sll0573
arcC
sll0572
NC-3547510
ncl1780
TU3698
7
9
12
15
18
21
500
10000
50000
Position (nt)

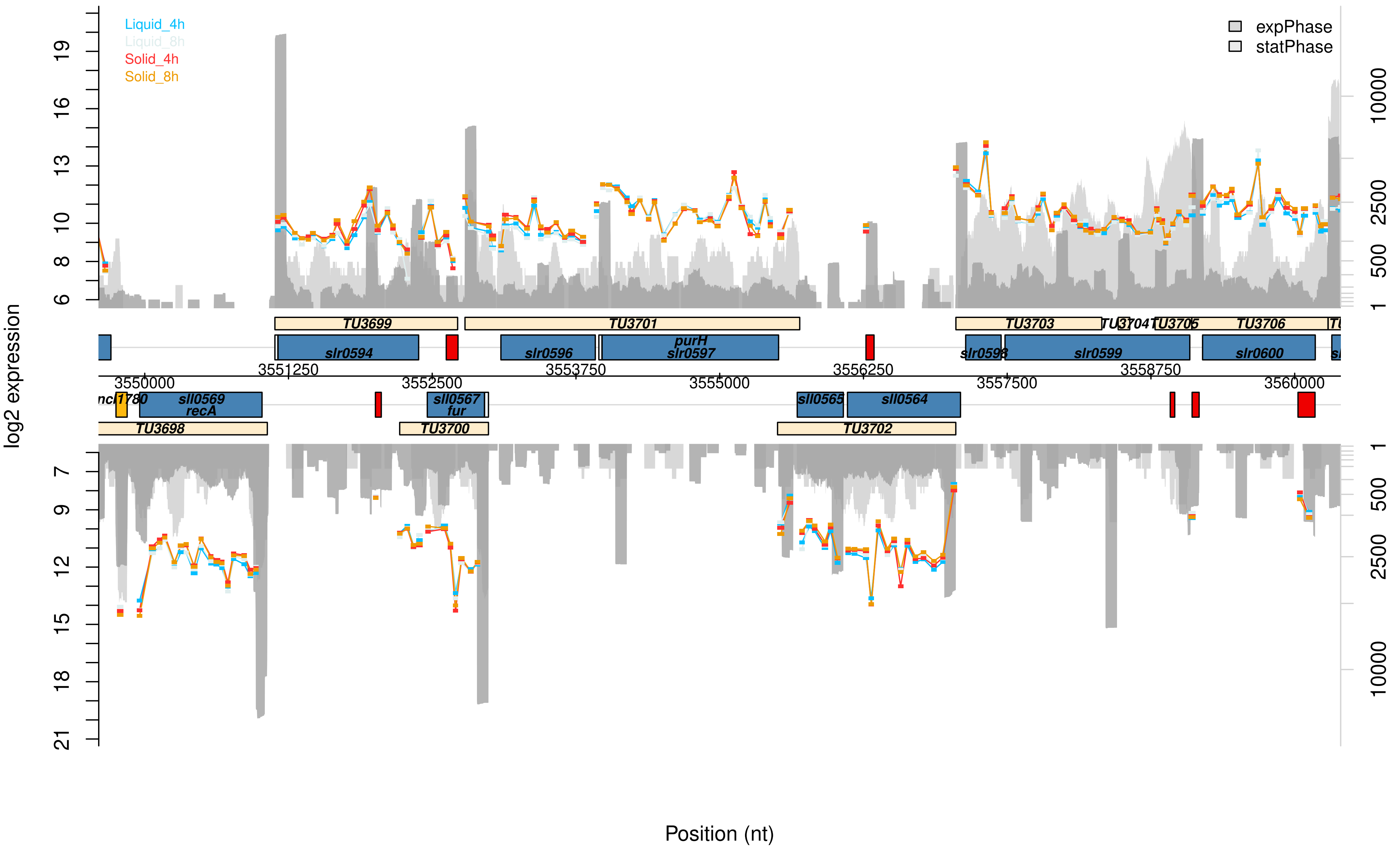

Liquid_4h
Liquid_8h
Solid_4h
Solid_8h
expPhase
statPhase
log2 expression
19
16
13
10
8
6
7
9
12
15
18
21
10000
2500
500
1
TU3699
TU3701
TU3703
TU3704
TU3705
TU3706
slr0594
slr0596
purH
slr0597
slr0598
slr0599
slr0600
3550000
3551250
3552500
3553750
3555000
3556250
3557500
3558750
3560000
ncl1780
sll0569
recA
sll0567
fur
sll0565
sll0564
TU3698
TU3700
TU3702
Position (nt)

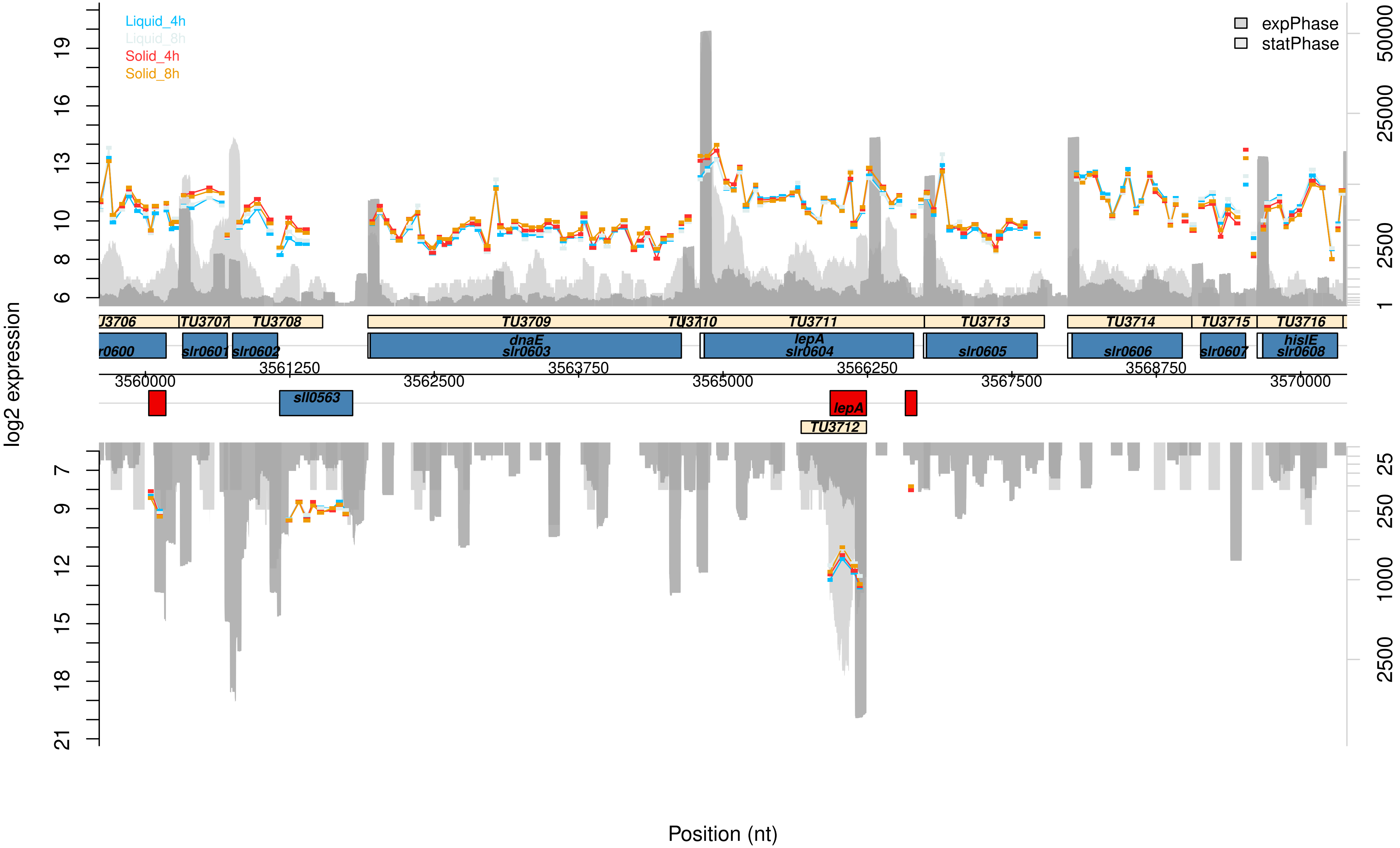

Liquid_4h
Liquid_8h
Solid_4h
Solid_8h
expPhase
statPhase
log2 expression
Position (nt)
TU3707
TU3708
TU3709
TU3711
TU3713
TU3714
TU3715
TU3716
TU3712
dnaE
slr0603
lepA
slr0604
slr0601
slr0602
slr0605
slr0606
slr0607
hisIE
slr0608
sll0563
lepA
3560000
3561250
3562500
3563750
3565000
3566250
3567500
3568750
3570000

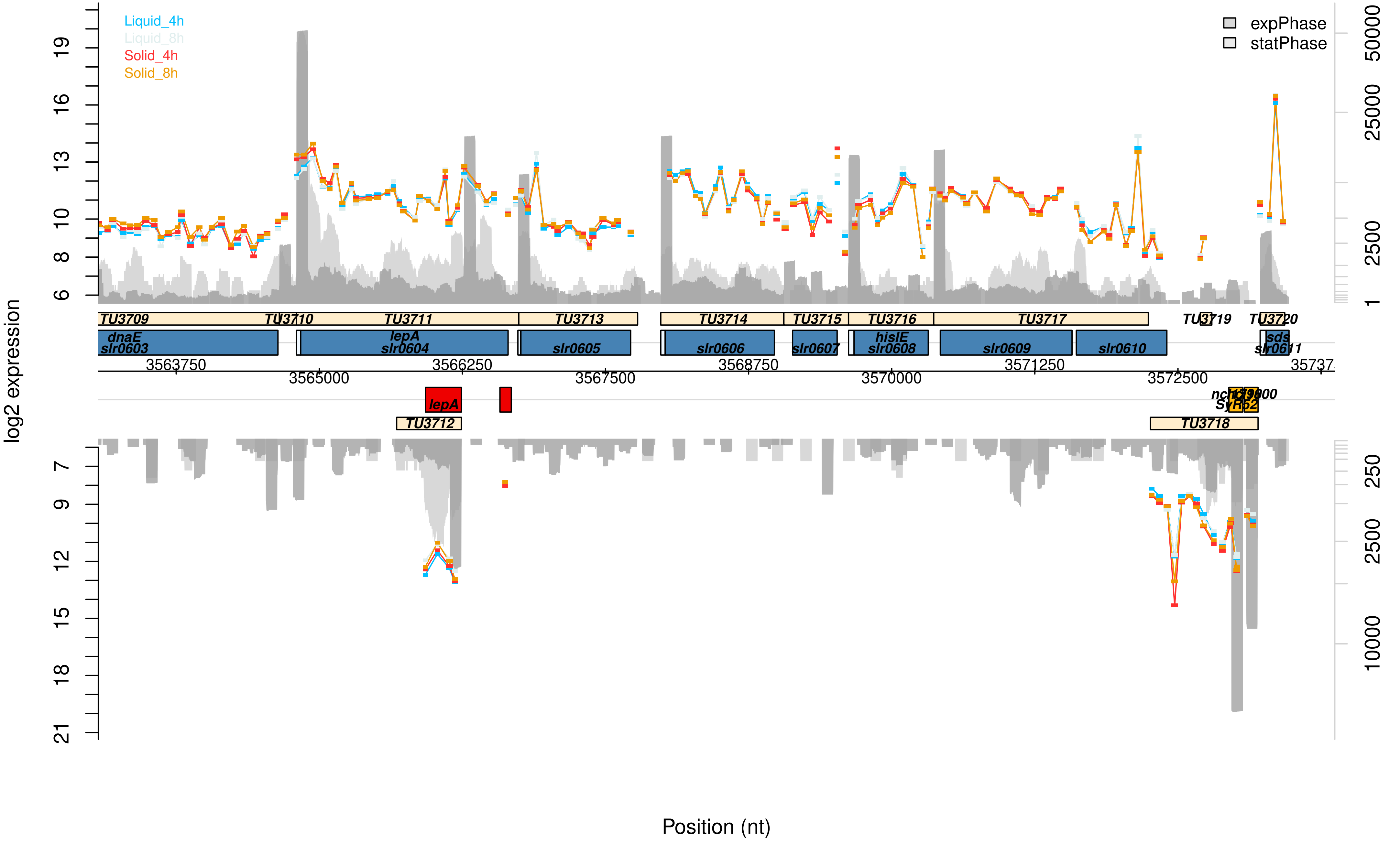

Liquid_4h
Liquid_8h
Solid_4h
Solid_8h
expPhase
statPhase
log2 expression
TU3709
TU3710
TU3711
TU3713
TU3714
TU3715
TU3716
TU3717
TU3719
TU3720
dnaE
slr0603
lepA
slr0604
slr0605
slr0606
slr0607
hisIE
slr0608
slr0609
slr0610
sds
slr0611
3563750
3565000
3566250
3567500
3568750
3570000
3571250
3572500
lepA
TU3712
TU3718
Position (nt)

Data Set S2: **Graphical visualization of the microarray results of the pSYSA plasmid.** This microarray data presents changes between Synechocystis cells of a sessile and planktonic lifestyle after 4 or 8 h, that are located on the pSYSA plasmid. The transcription values for planktonic cells after 4 h (liquid_4h, blue) or 8 h (liquid_8h, ice blue) and sessile cells after 4 h (solid_4h, red) or 8 h (solid_8h, orange) are given in log2 scale. The read numbers of exponential (dark grey) or stationary phase (light grey) grown cells are shown in log2 scale and are extracted from Kopf *et al.* (2014). Supporting data can be found in the Supplementary Data Table.

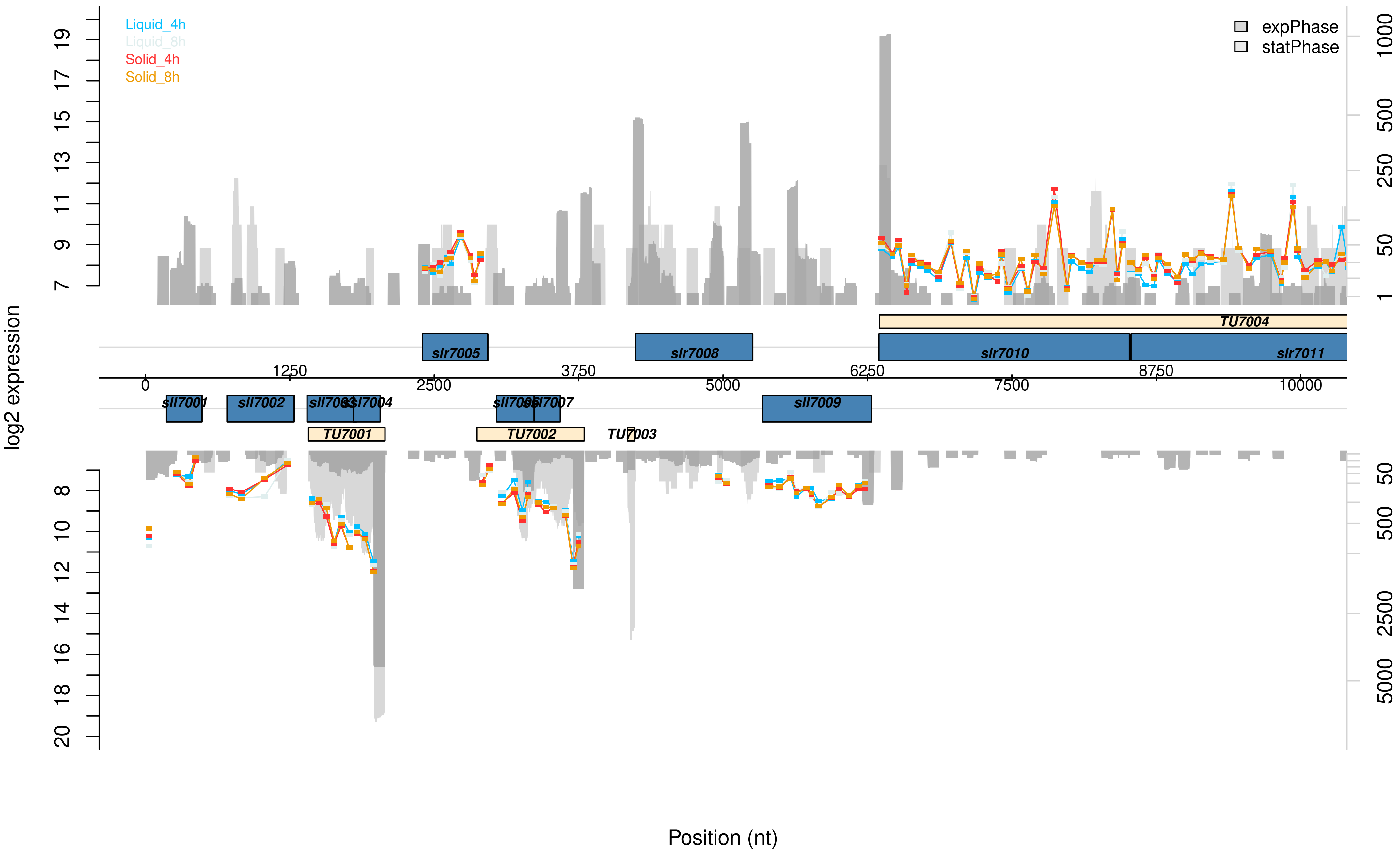

Liquid_4h
Liquid_8h
Solid_4h
Solid_8h
expPhase
statPhase
log2 expression
Position (nt)
TU7004
slr7005
slr7008
slr7010
slr7011
sll7001
sll7002
sll7003
sll7004
sll7006
sll7007
sll7009
TU7001
TU7002
TU7003
0
1250
2500
3750
5000
6250
7500
8750
10000

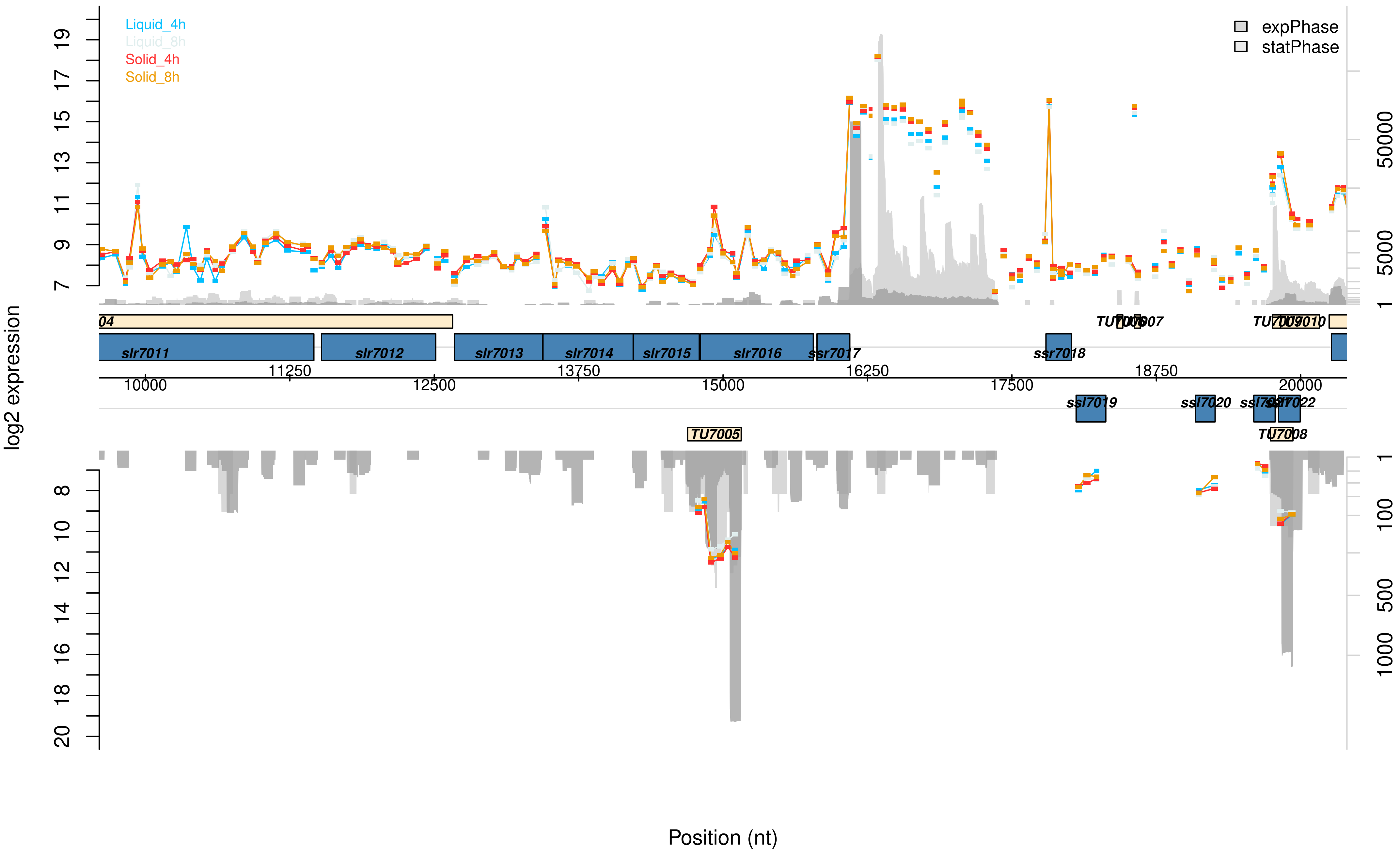

Liquid_4h
Liquid_8h
Solid_4h
Solid_8h
expPhase
statPhase
log2 expression
slr7011
slr7012
slr7013
slr7014
slr7015
slr7016
ssr7017
ssr7018
ssl7019
ssl7020
TU7005
Position (nt)

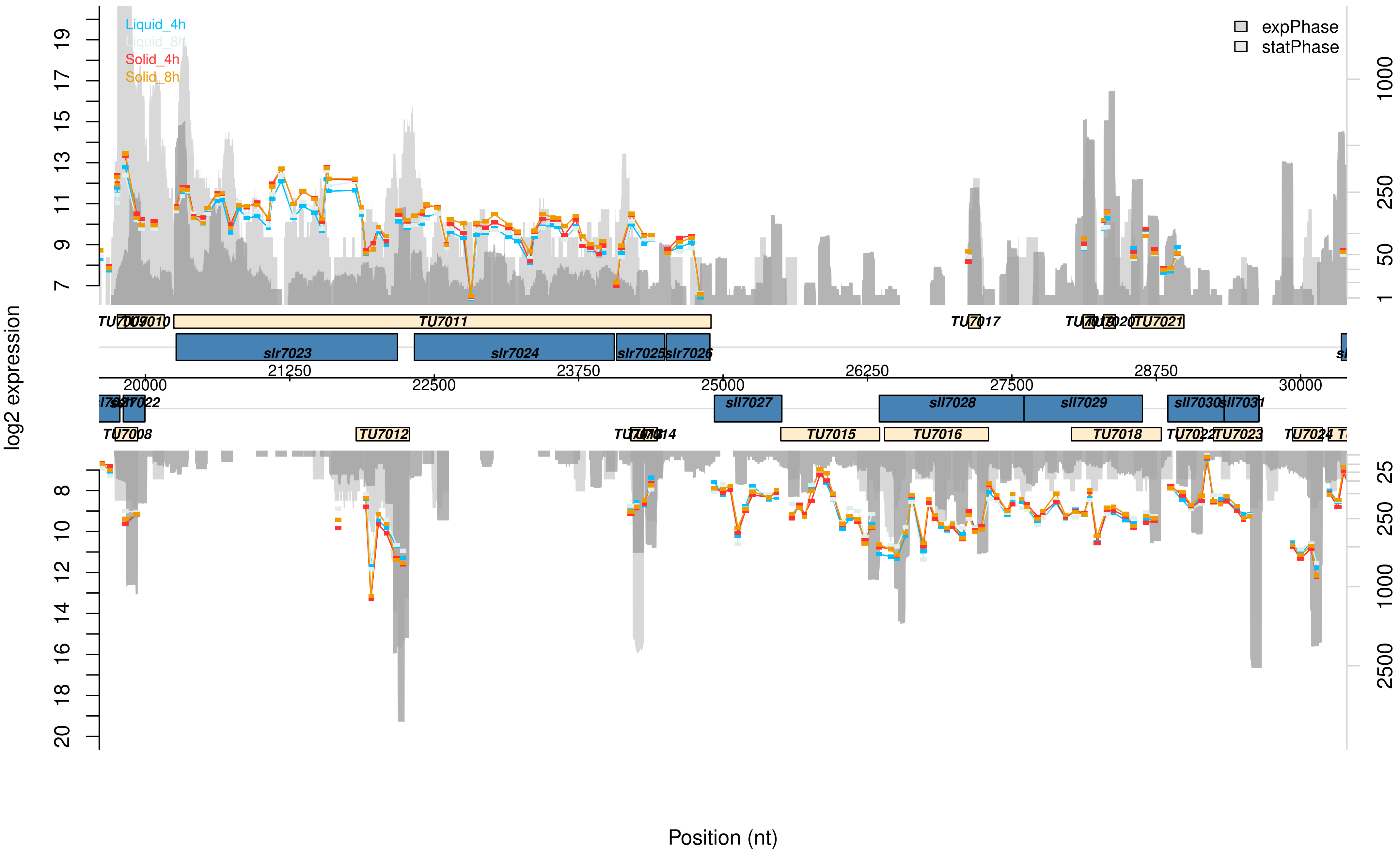

Liquid_4h
Liquid_8h
Solid_4h
Solid_8h
expPhase
statPhase
log2 expression
Position (nt)
TU7011
TU7017
TU7021
slr7023
slr7024
slr7025
slr7026
20000
21250
22500
23750
25000
26250
27500
28750
30000
sll7027
sll7028
sll7029
sll7030
sll7031
TU7008
TU7012
TU7015
TU7016
TU7018
TU7023

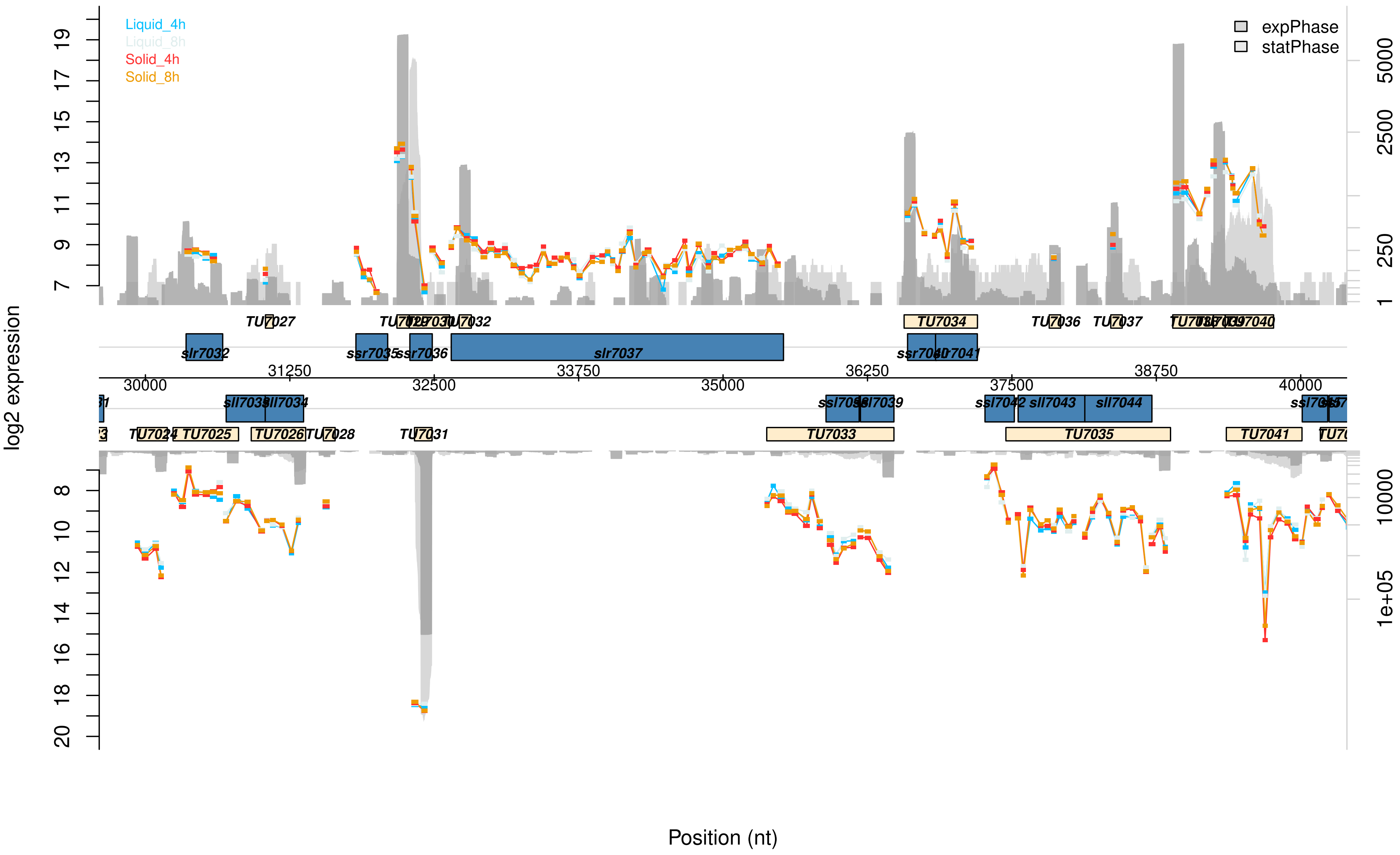

Liquid_4h
Liquid_8h
Solid_4h
Solid_8h
expPhase
statPhase
log2 expression
Position (nt)
TU7027
TU7034
TU7036
TU7037
slr7032
ssr7035
ssr7036
slr7037
sll7033
sll7034
ssl7042
sll7043
sll7044
TU7024
TU7025
TU7026
TU7028
TU7031
TU7033
TU7035
TU7041
30000
31250
32500
33750
35000
36250
37500
38750
40000

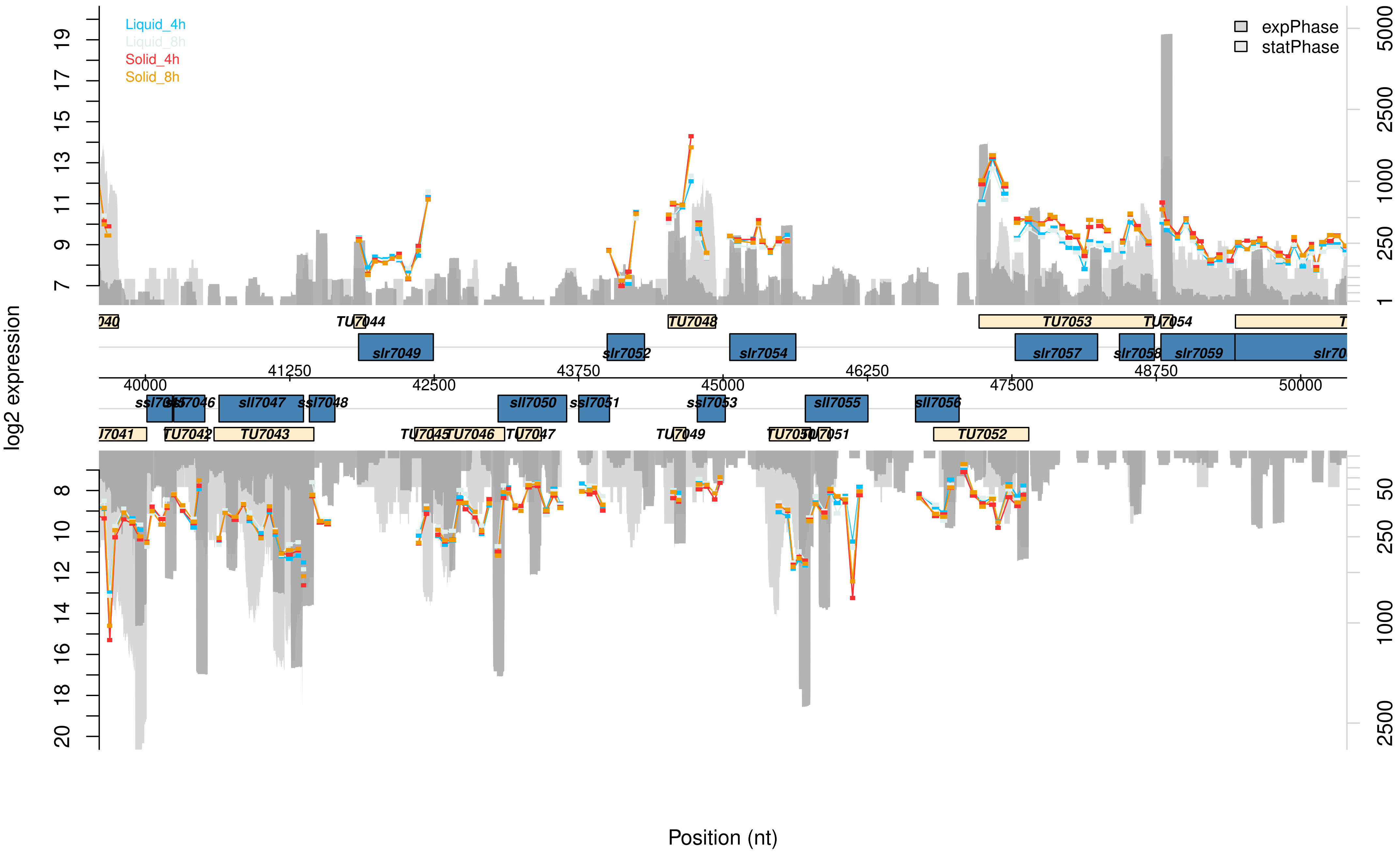

Liquid_4h
Liquid_8h
Solid_4h
Solid_8h
expPhase
statPhase
log2 expression
Position (nt)
TU7044
TU7048
TU7053
TU7054
slr7049
slr7052
slr7054
slr7057
slr7058
slr7059
40000
41250
42500
43750
45000
46250
47500
48750
50000
sll7047
ssl7048
sll7050
ssl7051
ssl7053
sll7055
sll7056
TU7042
TU7043
TU7046
TU7049
TU7052

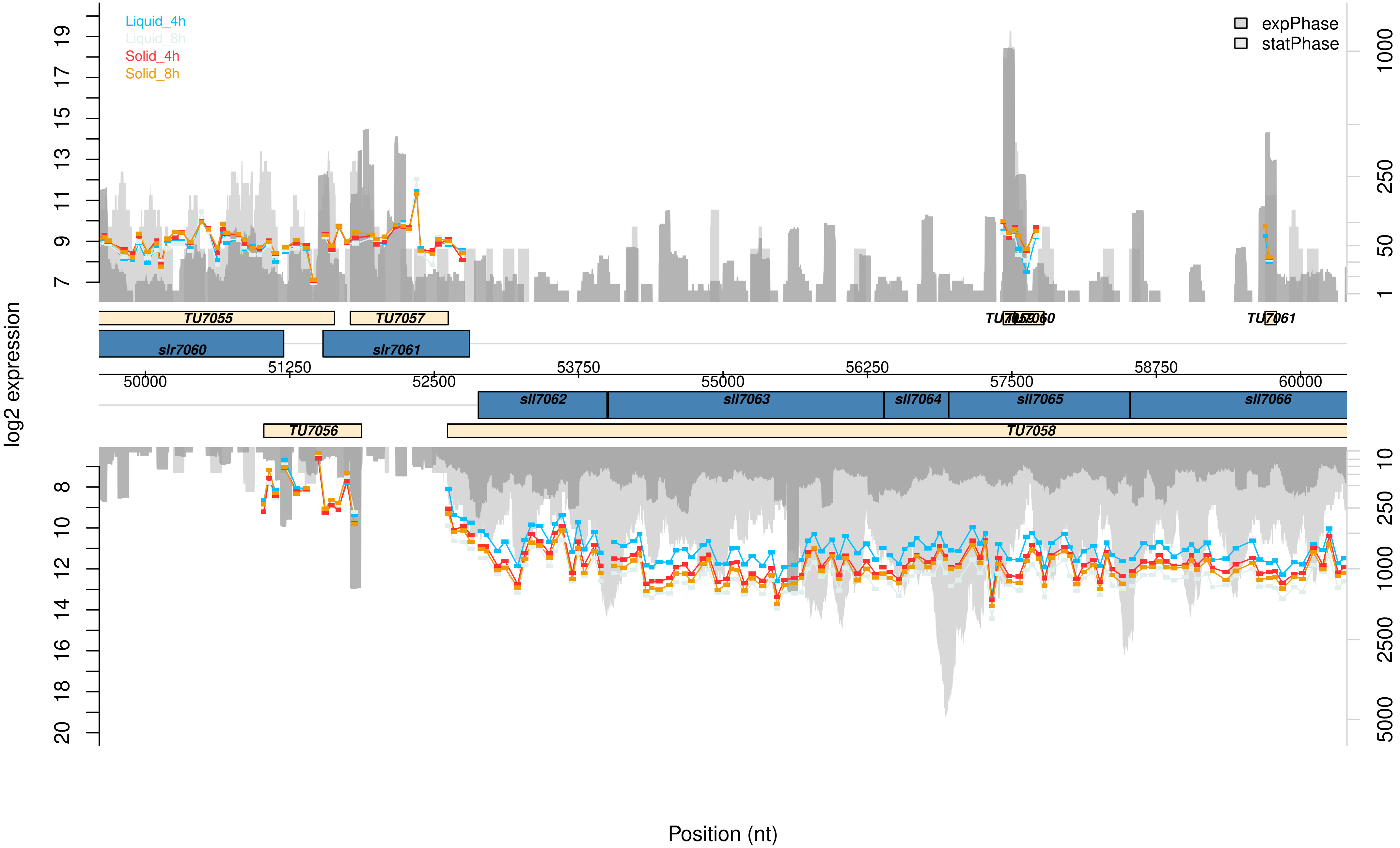

Liquid_4h
Liquid_8h
Solid_4h
Solid_8h
expPhase
statPhase
log2 expression
TU7055
TU7057
TU7059
TU7060
TU7061
slr7060
slr7061
50000
51250
52500
53750
55000
56250
57500
58750
60000
sll7062
sll7063
sll7064
sll7065
sll7066
TU7056
TU7058
Position (nt)

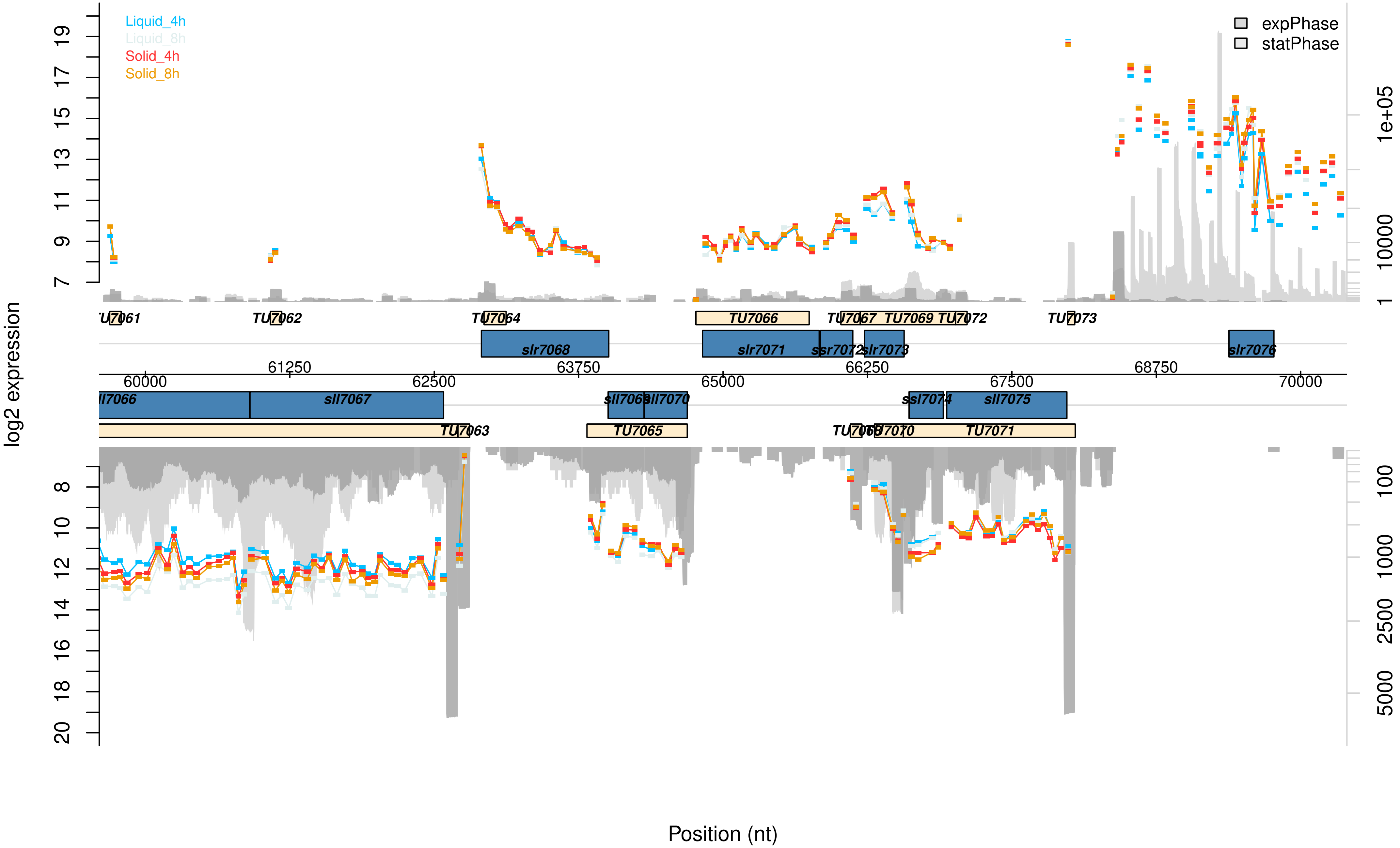

Liquid_4h
Liquid_8h
Solid_4h
Solid_8h
expPhase
statPhase
log2 expression
Position (nt)
TU7061
TU7062
TU7064
TU7066
TU7067
TU7069
TU7072
TU7073
slr7068
slr7071
ssr7072
slr7073
slr7076
60000
61250
62500
63750
65000
66250
67500
68750
70000
sll7067
sll7069
sll7070
ssl7074
sll7075
TU7063
TU7065
TU7068
TU7070
TU7071

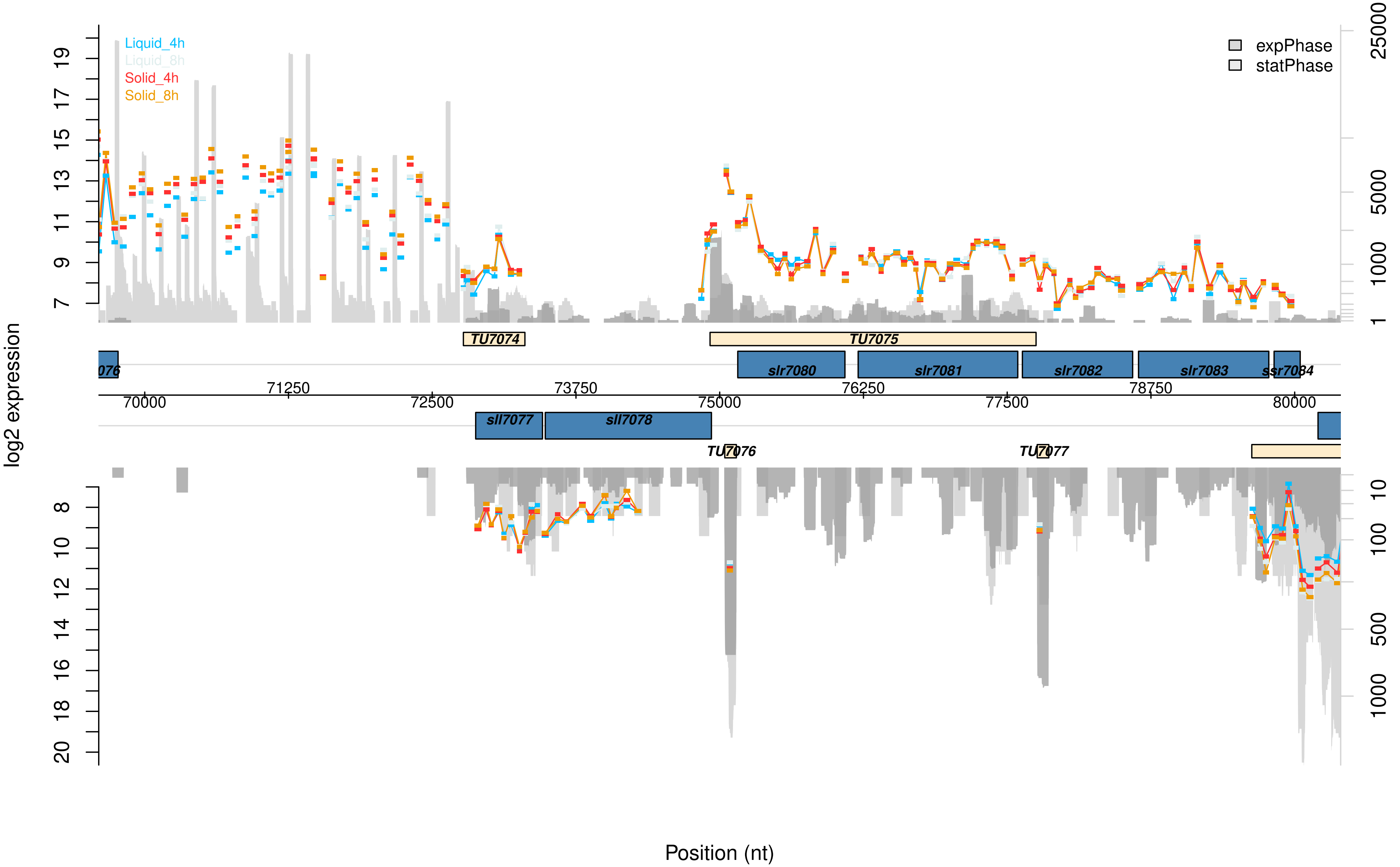

Liquid_4h
Liquid_8h
Solid_4h
Solid_8h
expPhase
statPhase
log2 expression
TU7074
TU7075
076
slr7080
slr7081
slr7082
slr7083
ssr7084
sll7077
sll7078
TU7076
TU7077
70000
71250
72500
73750
75000
76250
77500
78750
80000
Position (nt)

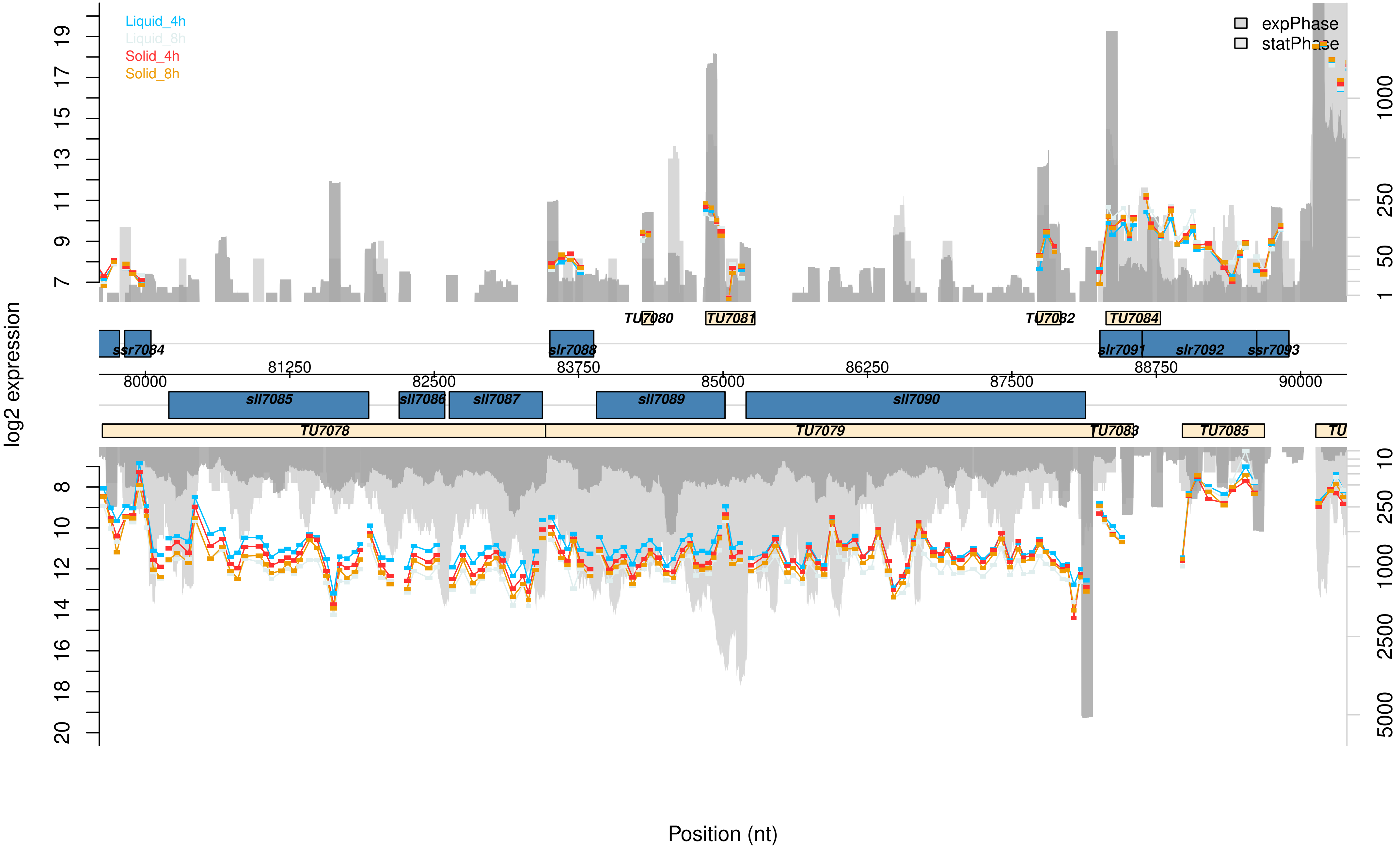

Liquid_4h
Liquid_8h
Solid_4h
Solid_8h
expPhase
statPhase
log2 expression
TU7080
TU7081
TU7082
TU7084
ssr7084
slr7088
slr7091
slr7092
ssr7093
81250
83750
86250
88750
80000
82500
85000
87500
90000
sll7085
sll7086
sll7087
sll7089
sll7090
TU7078
TU7079
TU7083
TU7085
Position (nt)

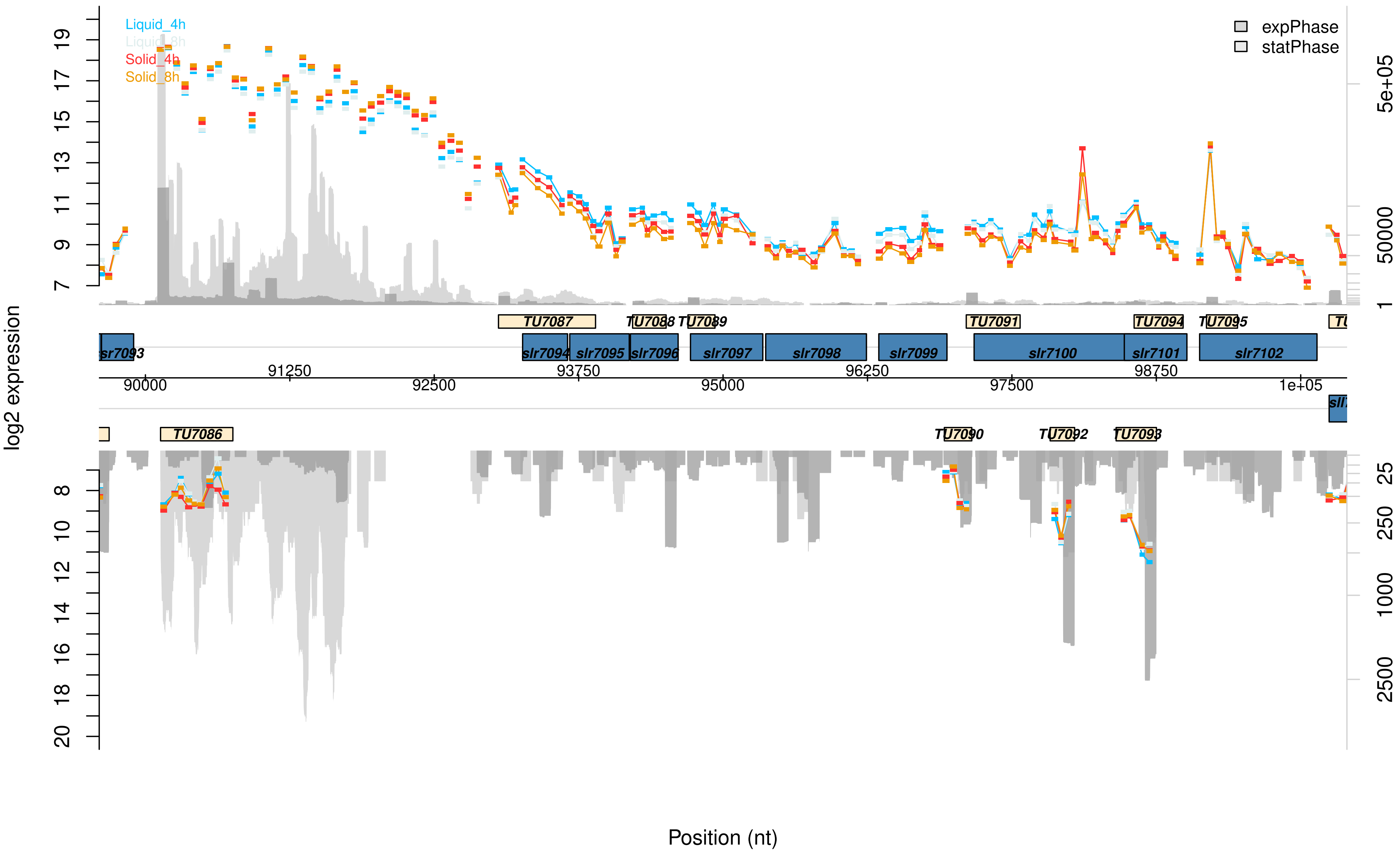

Liquid_4h
Liquid_8h
Solid_4h
Solid_8h
expPhase
statPhase
log2 expression
Position (nt)
TU7086
TU7087
TU7088
TU7089
TU7090
TU7091
TU7092
TU7093
TU7094
TU7095
sr7093
slr7094
slr7095
slr7096
slr7097
slr7098
slr7099
slr7100
slr7101
slr7102
90000
91250
92500
93750
95000
96250
97500
98750
1e+05
5e+05
50000
1
25
250
1000
2500

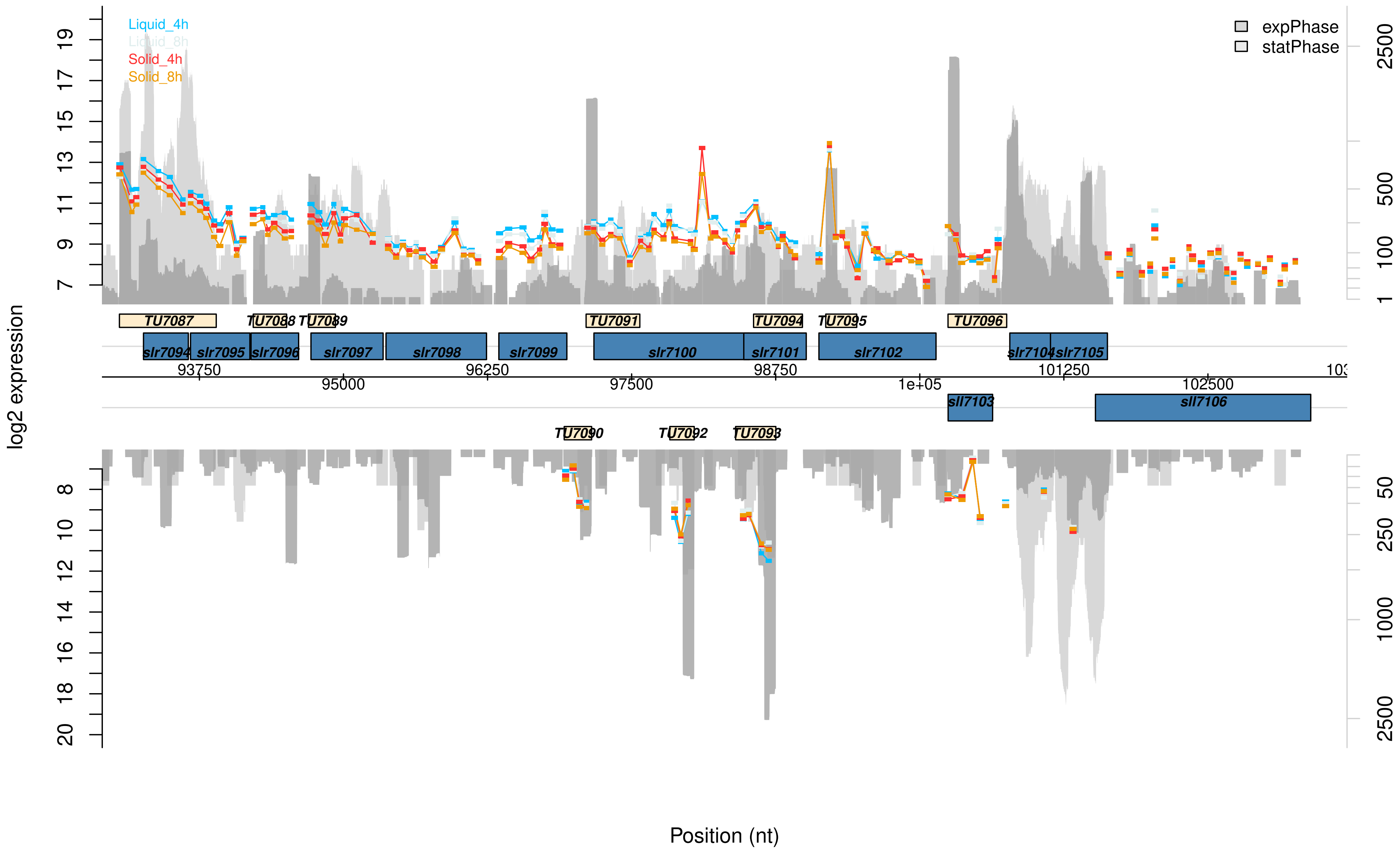

Liquid_4h
Liquid_8h
Solid_4h
Solid_8h
expPhase
statPhase
log2 expression
TU7087
TU7088
TU7089
TU7091
TU7094
TU7095
TU7096
slr7094
slr7095
slr7096
slr7097
slr7098
slr7099
slr7100
slr7101
slr7102
slr7104
slr7105
sll7103
sll7106
TU7090
TU7092
TU7093
93750
95000
96250
97500
98750
1e+05
101250
102500
Position (nt)

Data Set S3: **Graphical visualization of the microarray results of the pSYSG plasmid.** This microarray data presents changes between Synechocystis cells of a sessile and planktonic lifestyle after 4 or 8 h, that are located on the pSYSG plasmid. The transcription values for planktonic cells after 4 h (liquid_4h, blue) or 8 h (liquid_8h, ice blue) and sessile cells after 4 h (solid_4h, red) or 8 h (solid_8h, orange) are given in log2 scale. The read numbers of exponential (dark grey) or stationary phase (light grey) grown cells are shown in log2 scale and are extracted from Kopf *et al.* (2014). Supporting data can be found in the Supplementary Data Table.

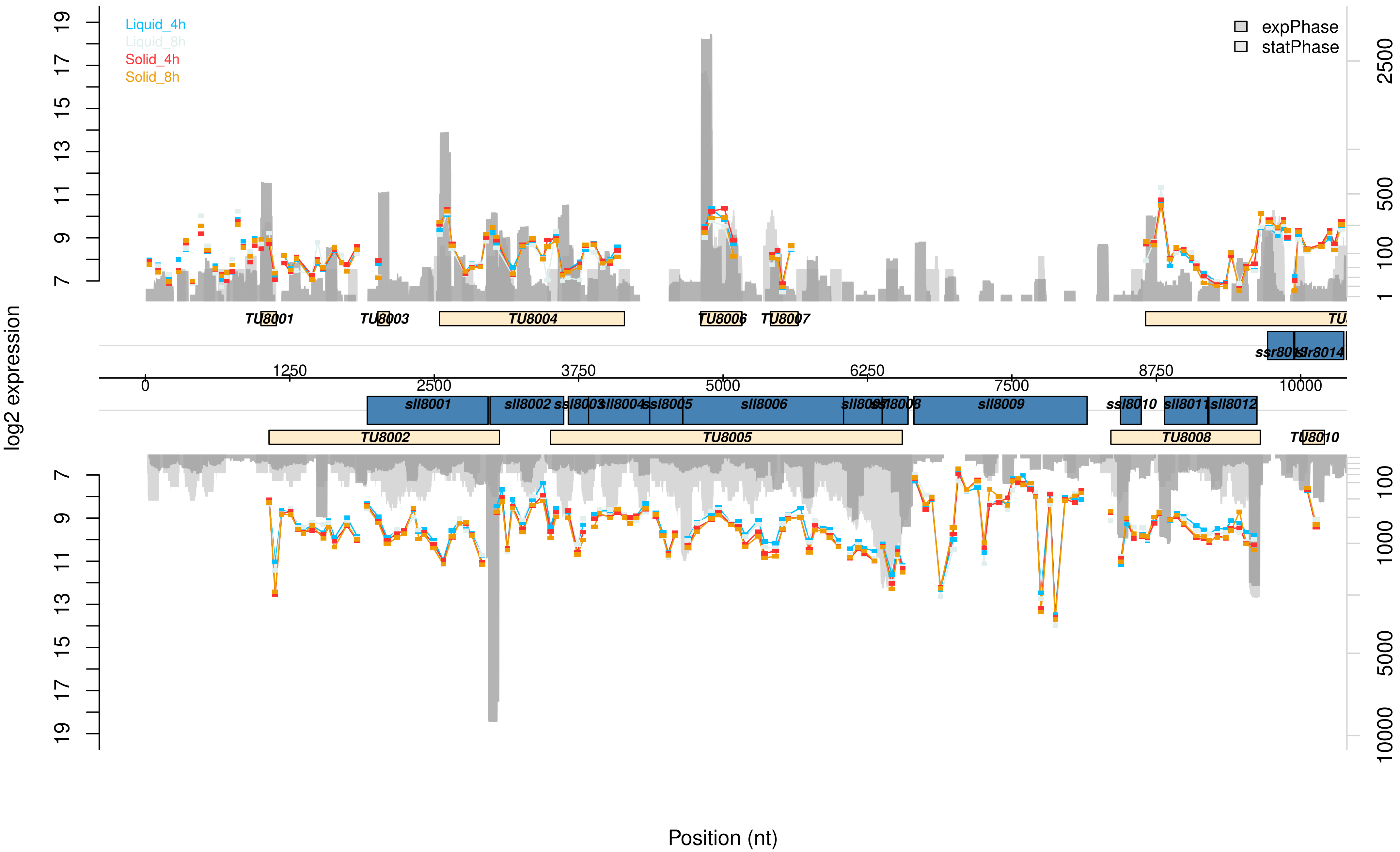

Liquid_4h
Liquid_8h
Solid_4h
Solid_8h
expPhase
statPhase
log2 expression
Position (nt)
TU8001
TU8002
TU8003
TU8004
TU8005
TU8006
TU8007
TU8008
TU8010
sll8001
sll8002
ssl8003
sll8004
ssl8005
sll8006
sll8009
ssl8010
sll8011
sll8012
ssr8014
0
1250
2500
3750
5000
6250
7500
8750
10000

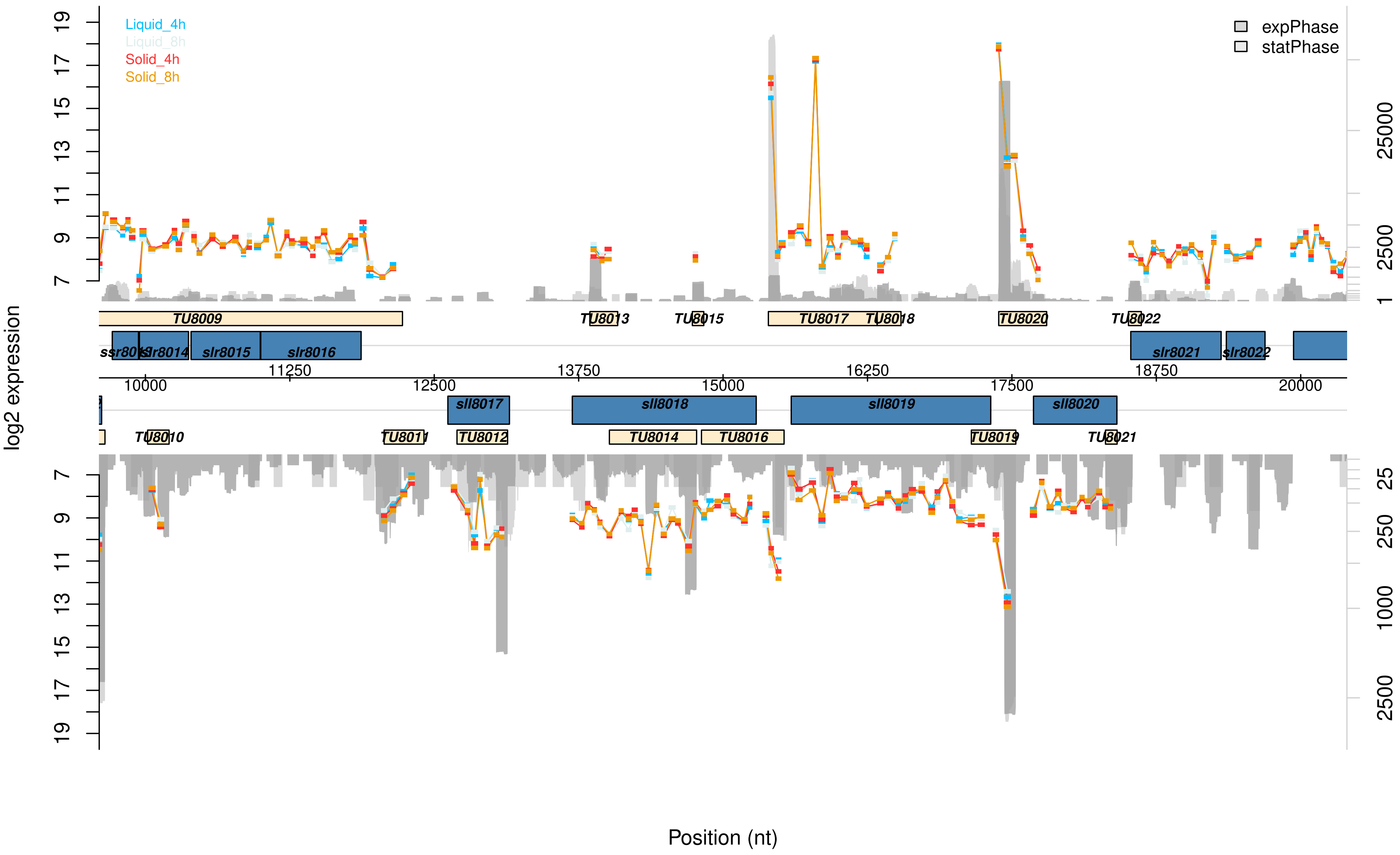

Liquid_4h
Liquid_8h
Solid_4h
Solid_8h
expPhase
statPhase
log2 expression
TU8009
TU8013
TU8015
TU8017
TU8018
TU8020
TU8022
ssr8013
slr8014
slr8015
slr8016
slr8021
slr8022
11250
13750
16250
18750
10000
12500
15000
17500
20000
sll8017
sll8018
sll8019
sll8020
TU8010
TU8011
TU8012
TU8014
TU8016
TU8019
TU8021
Position (nt)

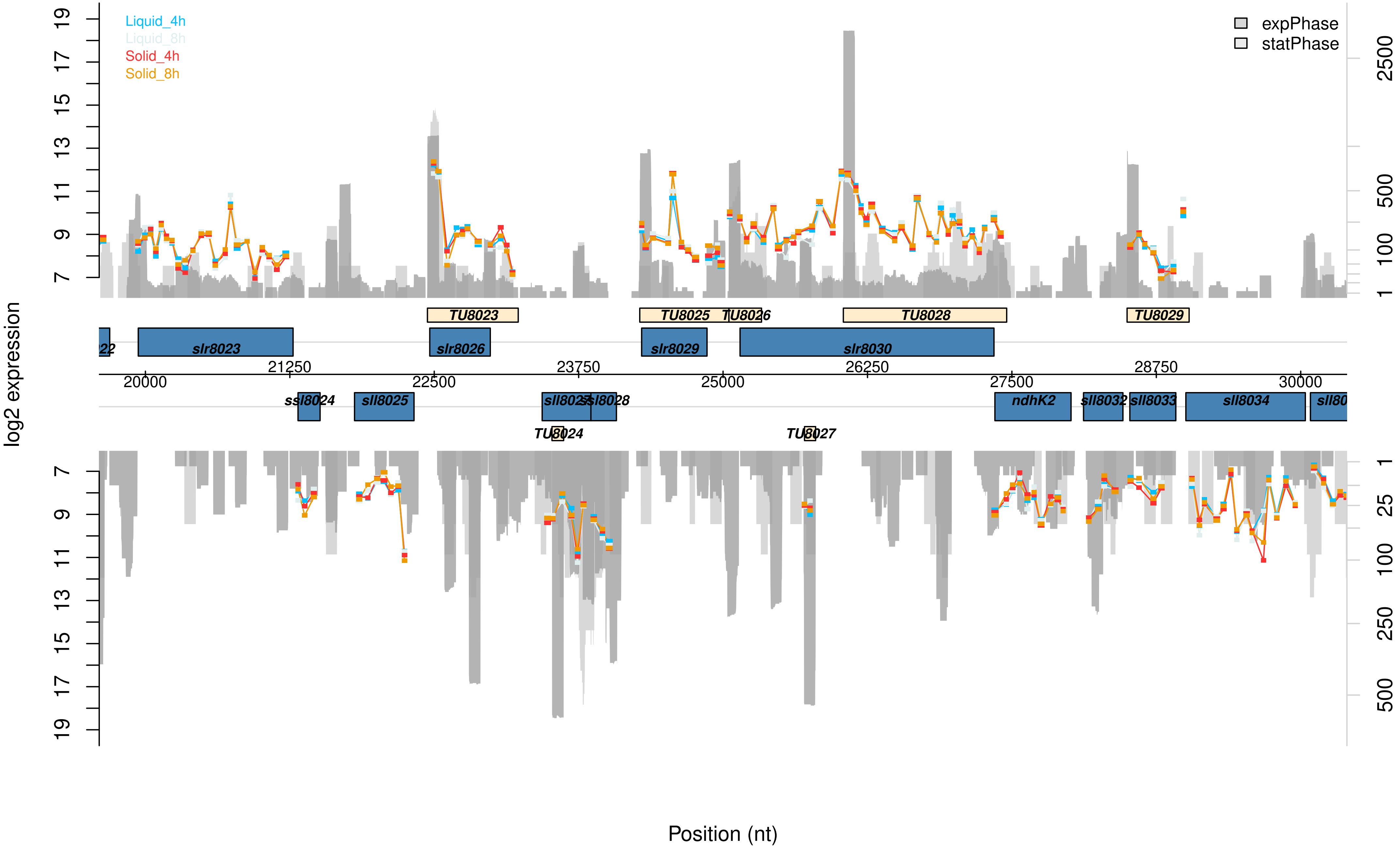

Liquid_4h
Liquid_8h
Solid_4h
Solid_8h
expPhase
statPhase
log2 expression
TU8023
TU8025
TU8026
TU8028
TU8029
slr8023
slr8026
slr8029
slr8030
21250
23750
26250
28750
20000
22500
25000
27500
30000
ssl8024
sll8025
sll8028
ndhK2
sll8032
sll8033
sll8034
TU8024
TU8027
Position (nt)

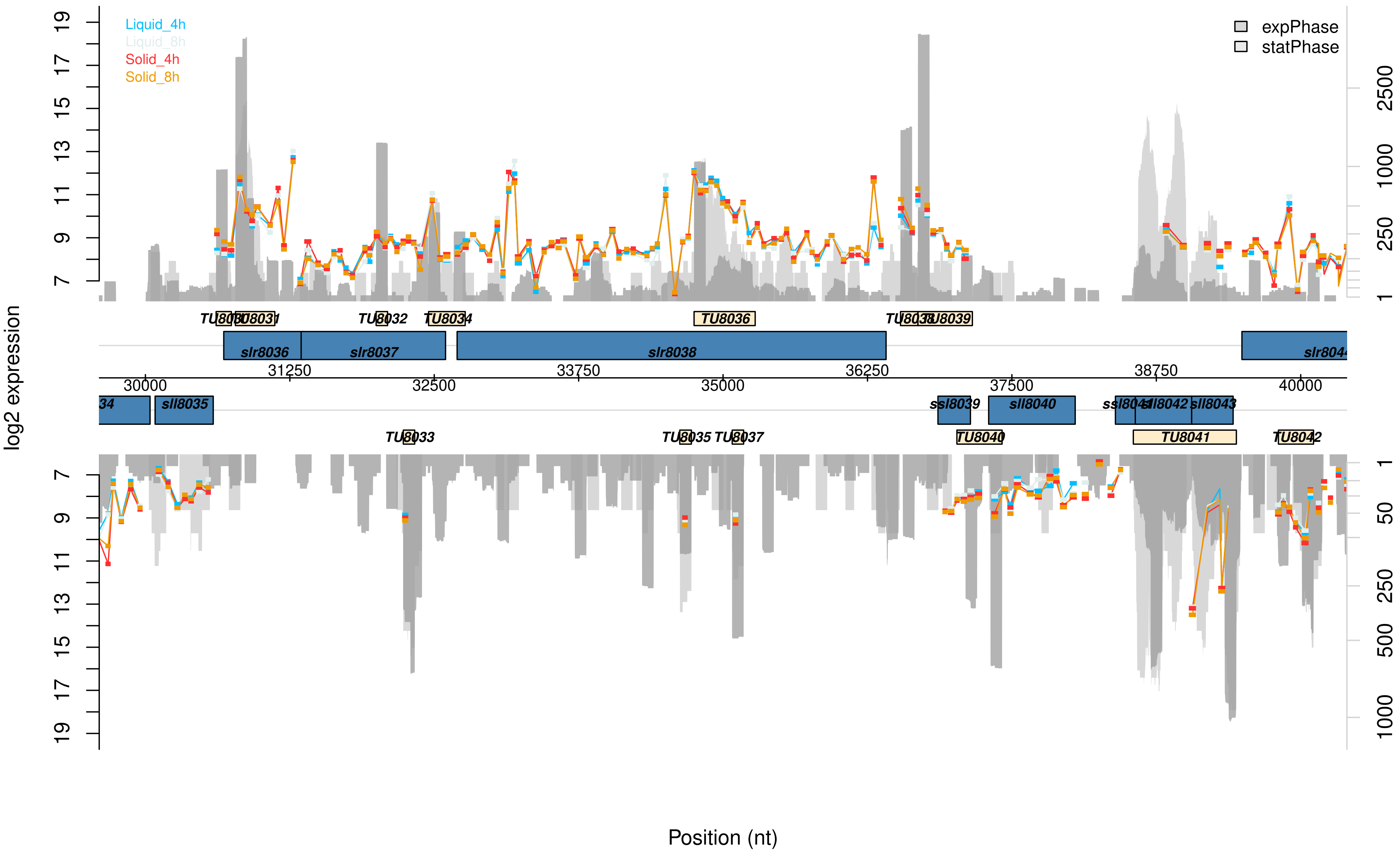

Liquid_4h
Liquid_8h
Solid_4h
Solid_8h
expPhase
statPhase
log2 expression
Position (nt)
TU8031
TU8032
TU8034
TU8036
TU8039
slr8036
slr8037
slr8038
sll8035
ssl8039
sll8040
sll8042
sll8043
TU8033
TU8035
TU8037
TU8040
TU8041
TU8042
30000
31250
32500
33750
35000
36250
37500
38750
40000

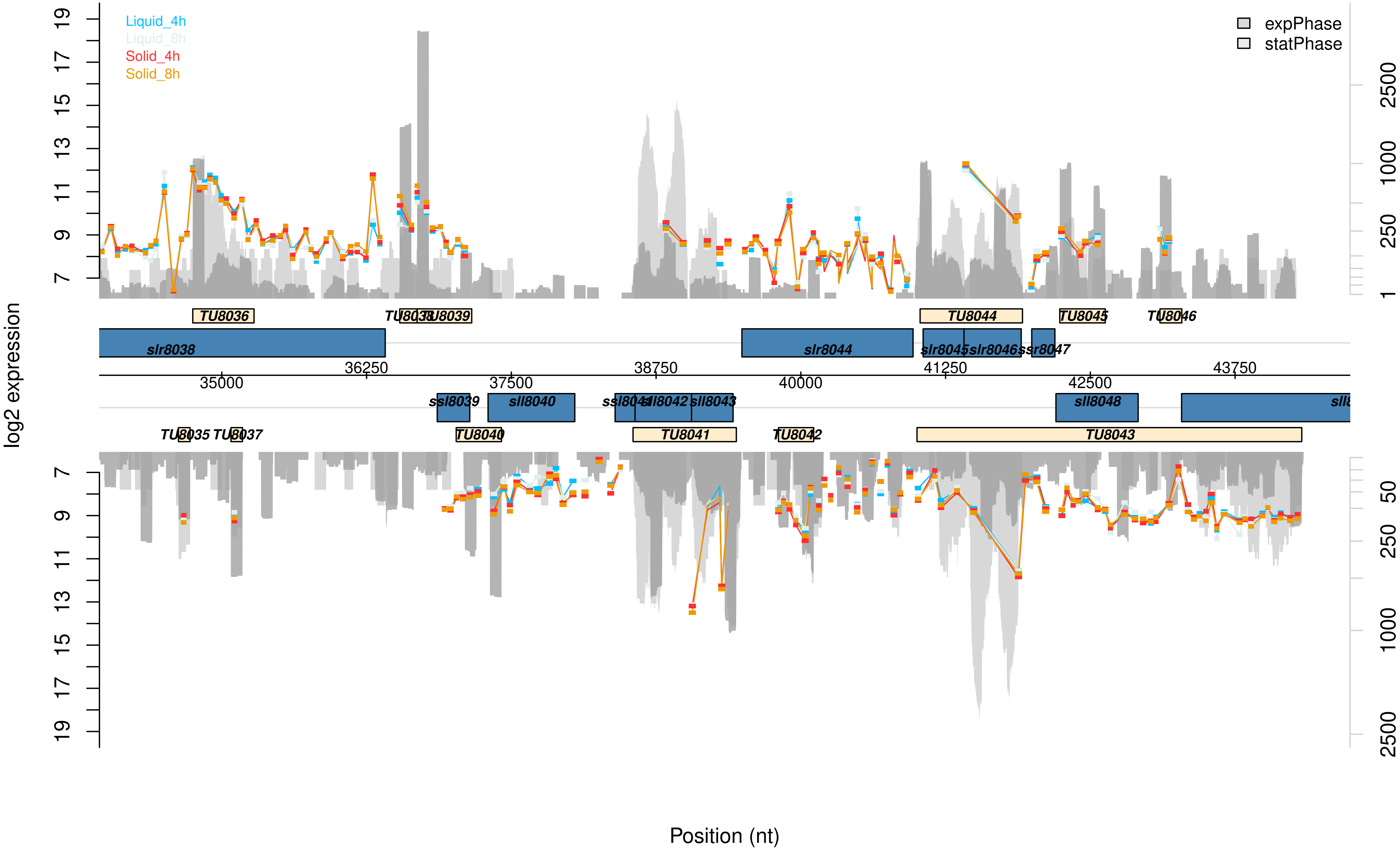

Liquid_4h
Liquid_8h
Solid_4h
Solid_8h
expPhase
statPhase
log2 expression
TU8036
TU8038
TU8039
TU8044
TU8045
TU8046
slr8038
slr8044
slr8045
slr8046
ssr8047
35000
36250
37500
38750
40000
41250
42500
43750
ssl8039
sll8040
ssl8041
sll8042
sll8043
sll8048
TU8035
TU8037
TU8040
TU8041
TU8042
TU8043
Position (nt)

Data Set S4: **Graphical visualization of the microarray results of the pSYSM plasmid.** This microarray data presents changes between Synechocystis cells of a sessile and planktonic lifestyle after 4 or 8 h, that are located on the pSYSM plasmid. The transcription values for planktonic cells after 4 h (liquid_4h, blue) or 8 h (liquid_8h, ice blue) and sessile cells after 4 h (solid_4h, red) or 8 h (solid_8h, orange) are given in log2 scale. The read numbers of exponential (dark grey) or stationary phase (light grey) grown cells are shown in log2 scale and are extracted from Kopf *et al.* (2014). Supporting data can be found in the Supplementary Data Table.

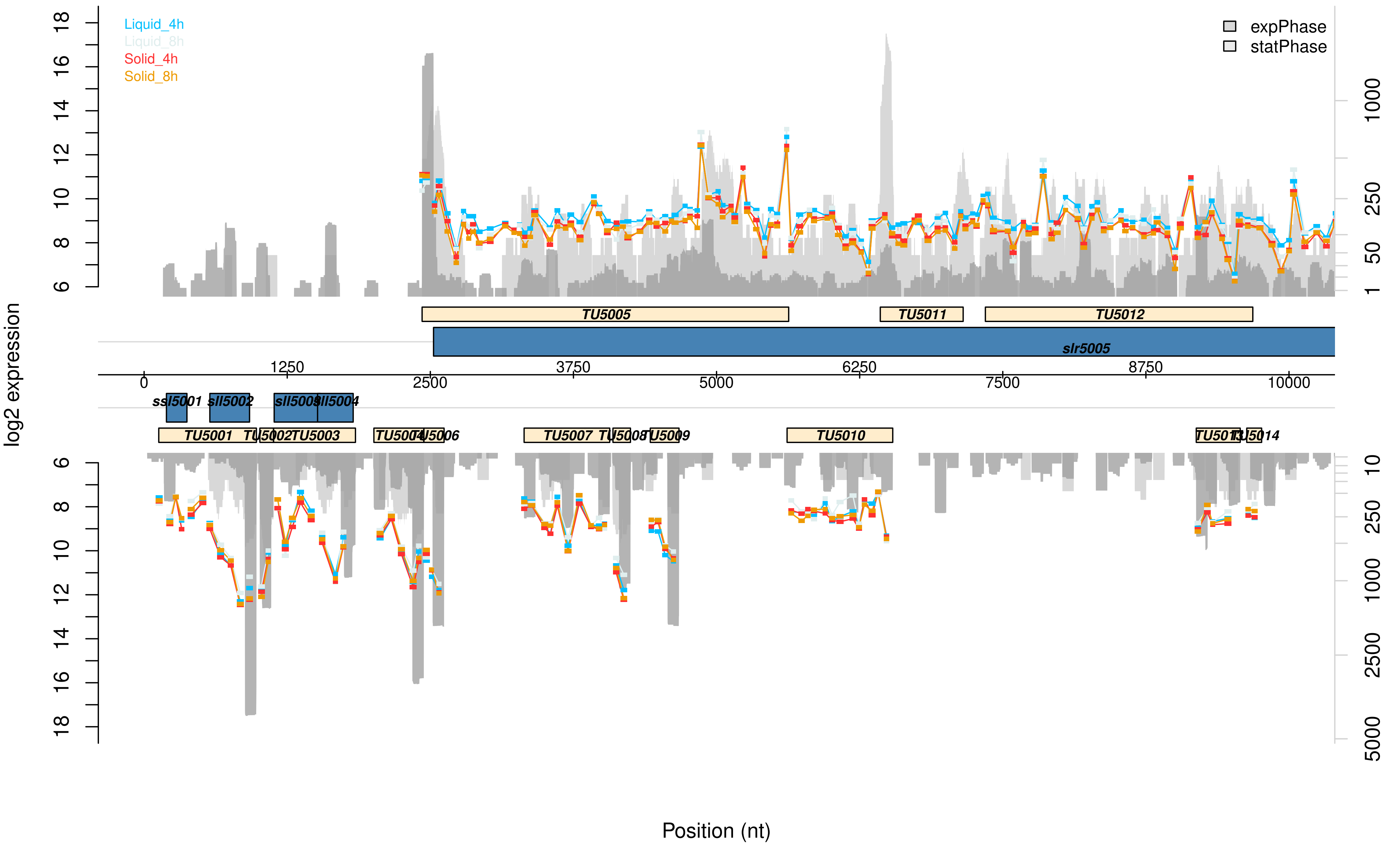
Liquid_4h
Liquid_8h
Solid_4h
Solid_8h
expPhase
statPhase
log2 expression
TU5005
TU5011
TU5012
slr5005
ssl5001
sll5002
sll5003
sll5004
TU5001
TU5002
TU5003
TU5004
TU5006
TU5007
TU5008
TU5009
TU5010
TU5014
Position (nt)

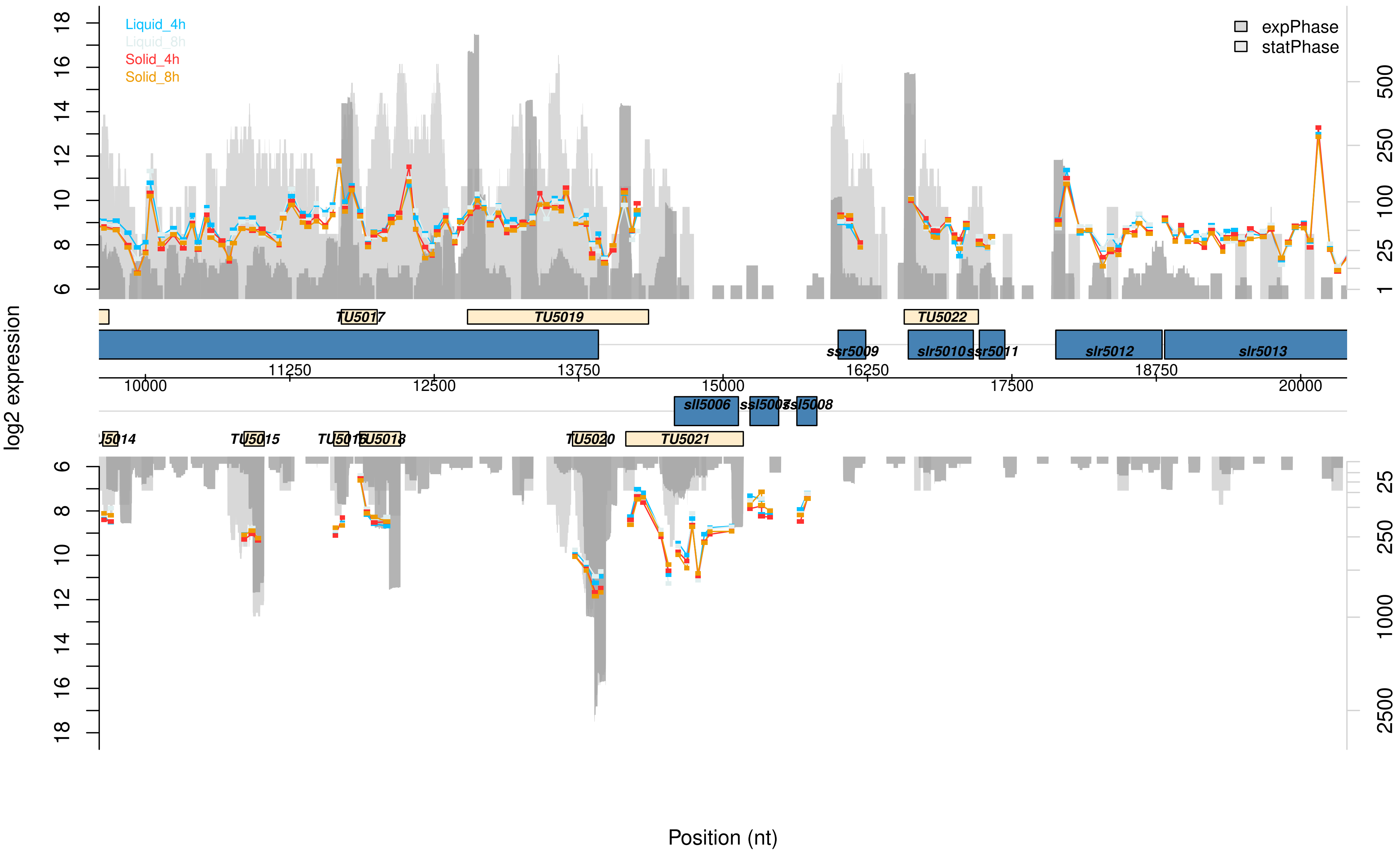

Liquid_4h
Liquid_8h
Solid_4h
Solid_8h
expPhase
statPhase
log2 expression
Position (nt)
TU5017
TU5019
TU5022
ssr5009
slr5010
ssr5011
slr5012
slr5013
sll5006
ssl5007
ssl5008
TU5014
TU5015
TU5016
TU5018
TU5020
TU5021
10000
11250
12500
13750
15000
16250
17500
18750
20000

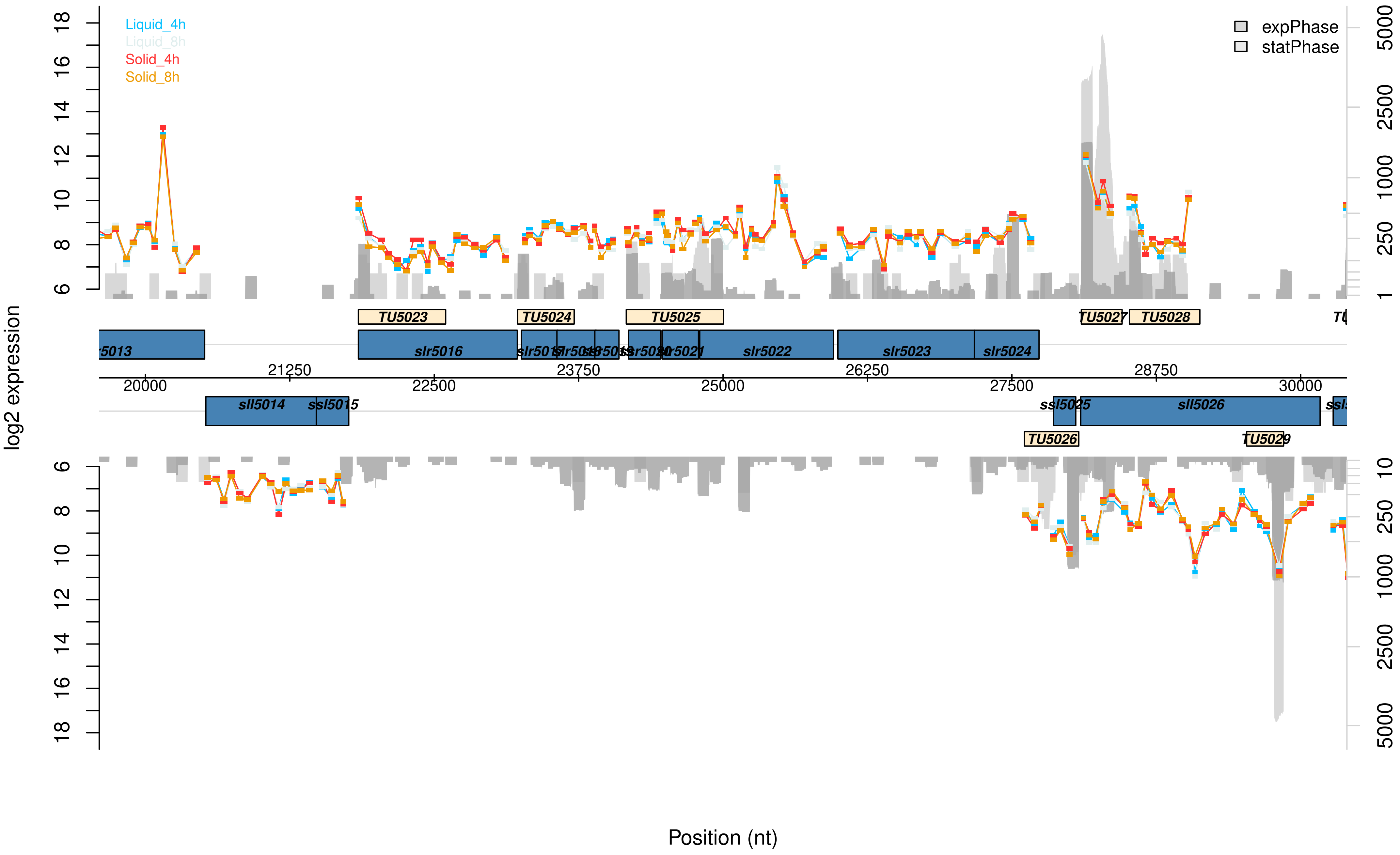
Liquid_4h
Liquid_8h
Solid_4h
Solid_8h
expPhase
statPhase
log2 expression
Position (nt)
TU5023
TU5024
TU5025
TU5027
TU5028
TU5026
TU5029
slr5016
slr5022
slr5023
slr5024
sll5014
ssl5015
sll5026
20000
21250
22500
23750
25000
26250
27500
28750
30000

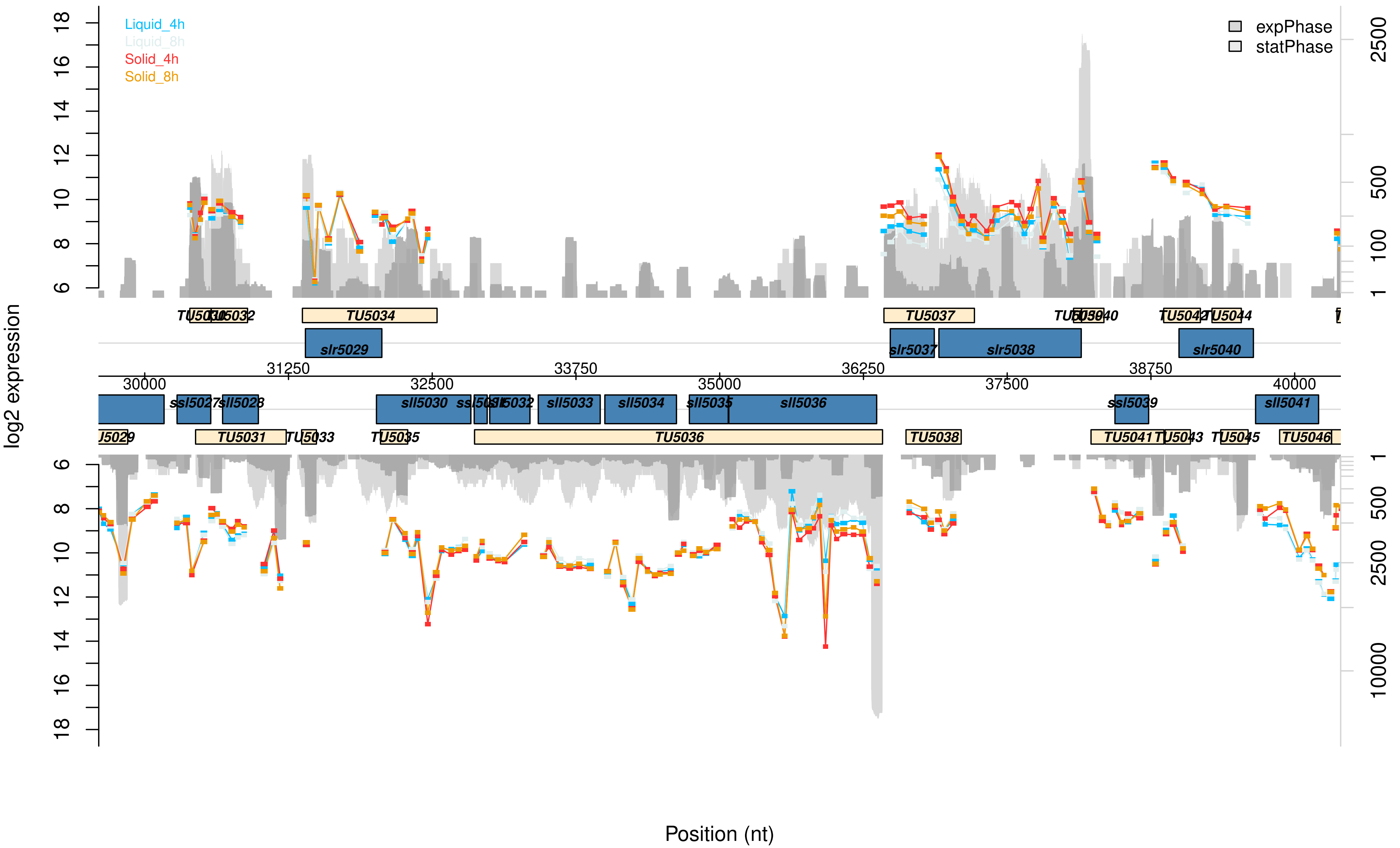
Liquid_4h
Liquid_8h
Solid_4h
Solid_8h
expPhase
statPhase
log2 expression
Position (nt)
TU5034
TU5037
TU5044
slr5029
slr5037
slr5038
slr5040
ssl5027
sll5028
sll5030
sll5032
sll5033
sll5034
sll5035
sll5036
ssl5039
sll5041
TU5031
TU5033
TU5035
TU5036
TU5038
TU5041
TU5043
TU5045
TU5046

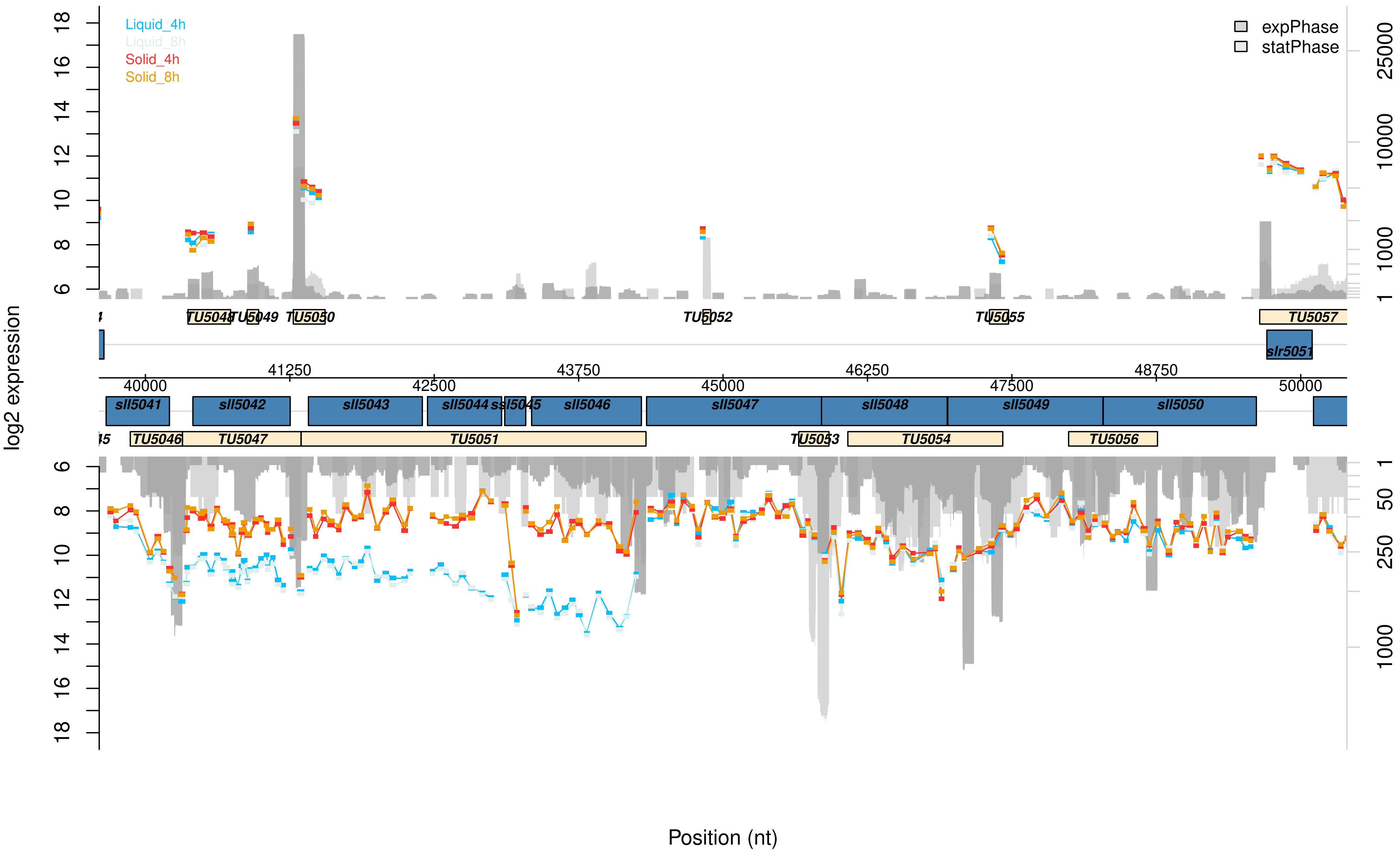
Liquid_4h
Liquid_8h
Solid_4h
Solid_8h
expPhase
statPhase
log2 expression
TU5048
TU5049
TU5050
TU5052
TU5055
TU5057
slr5051
sll5041
sll5042
sll5043
sll5044
sll5045
sll5046
sll5047
sll5048
sll5049
sll5050
TU5046
TU5047
TU5051
TU5053
TU5054
TU5056
Position (nt)

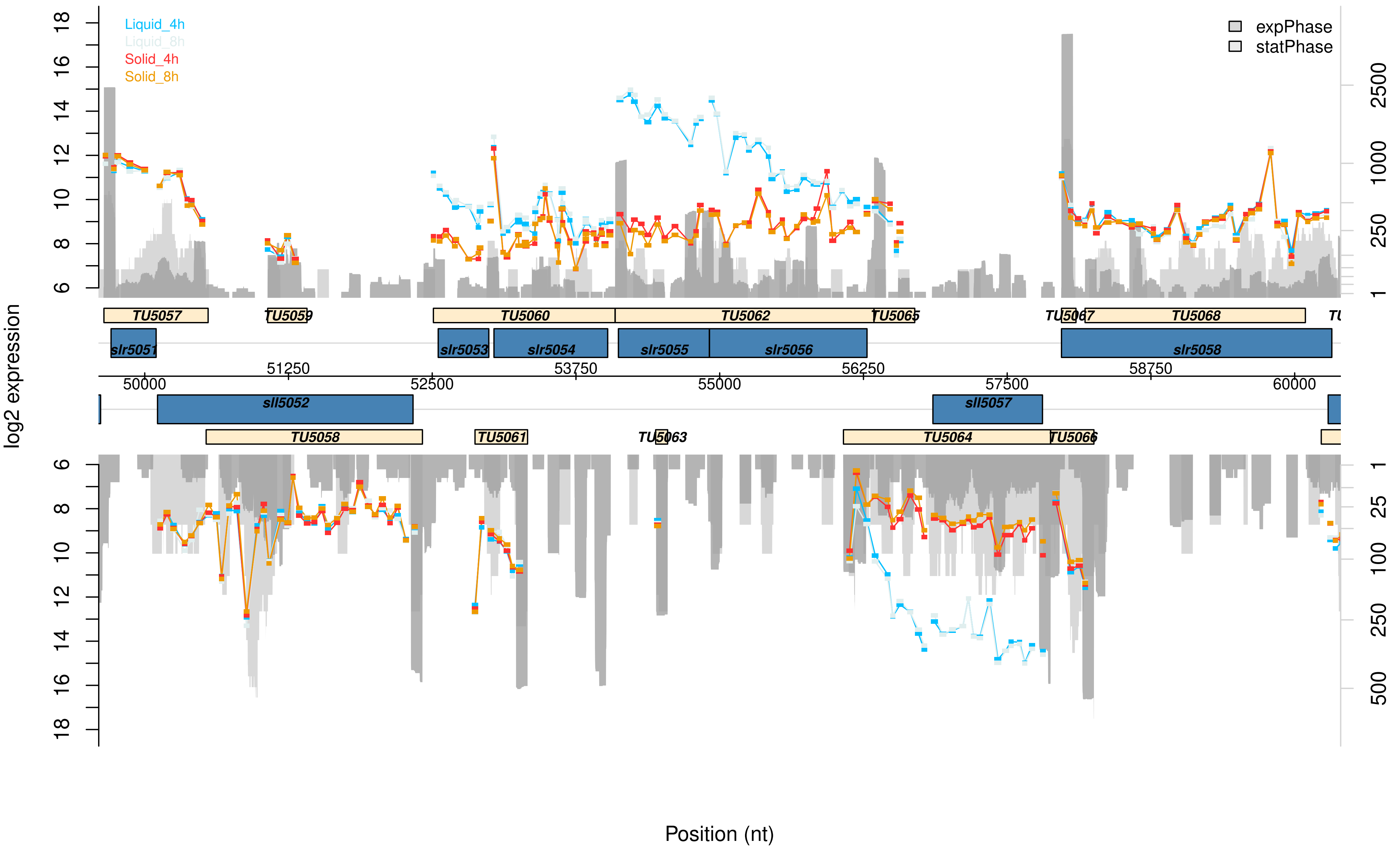

Liquid_4h
Liquid_8h
Solid_4h
Solid_8h
expPhase
statPhase
log2 expression
Position (nt)
TU5057
TU5059
TU5060
TU5062
TU5065
TU5067
TU5068
slr5051
slr5053
slr5054
slr5055
slr5056
slr5058
sll5052
sll5057
TU5058
TU5061
TU5063
TU5064
TU5066
50000
51250
52500
53750
55000
56250
57500
58750
60000

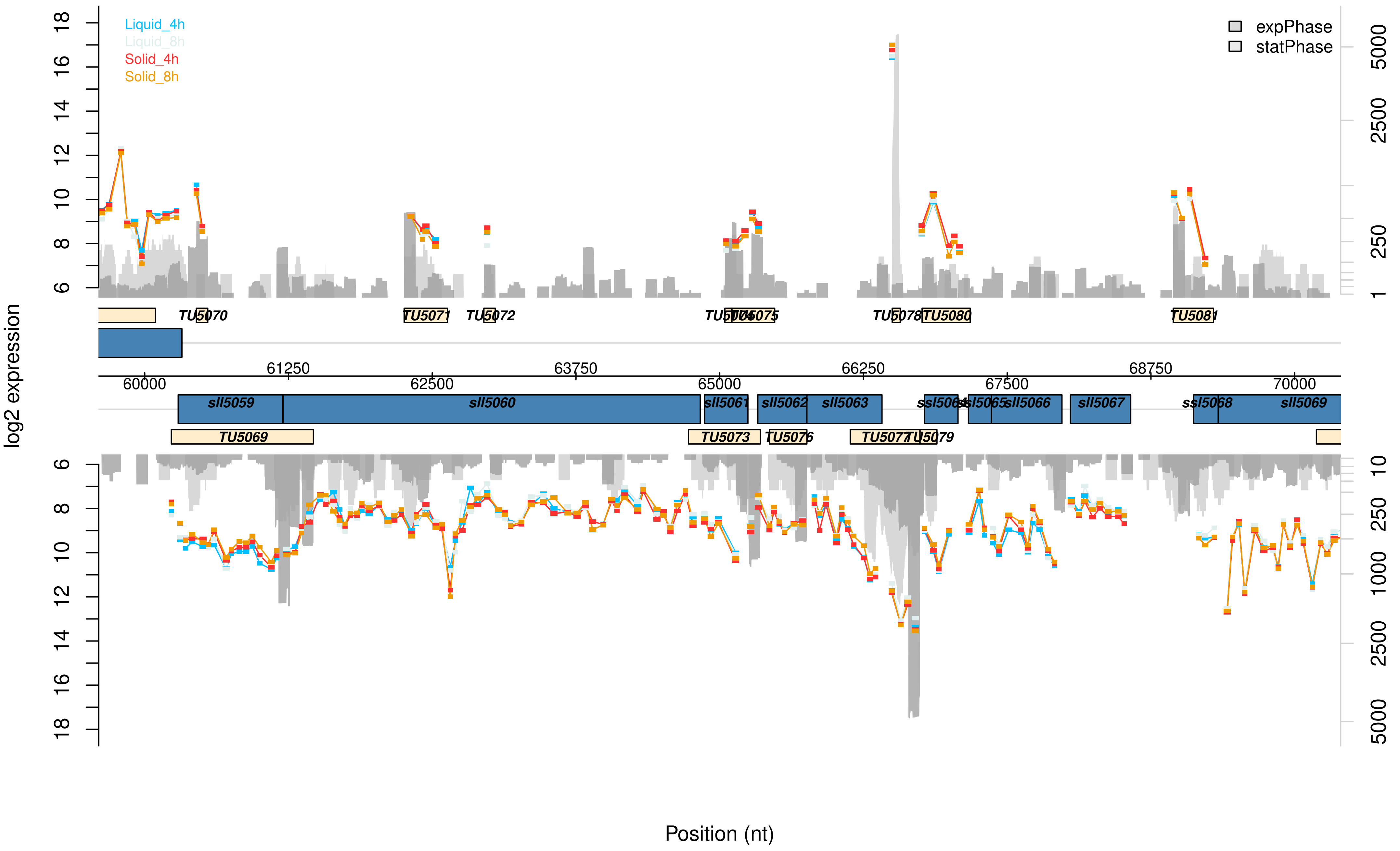

Liquid_4h
Liquid_8h
Solid_4h
Solid_8h
expPhase
statPhase
log2 expression
Position (nt)
TU5069
TU5070
TU5071
TU5072
TU5073
TU5075
TU5076
TU5077
TU5078
TU5079
TU5080
TU5081
sll5059
sll5060
sll5061
sll5062
sll5063
sll5066
sll5067
sll5069
60000
61250
62500
63750
65000
66250
67500
68750
70000

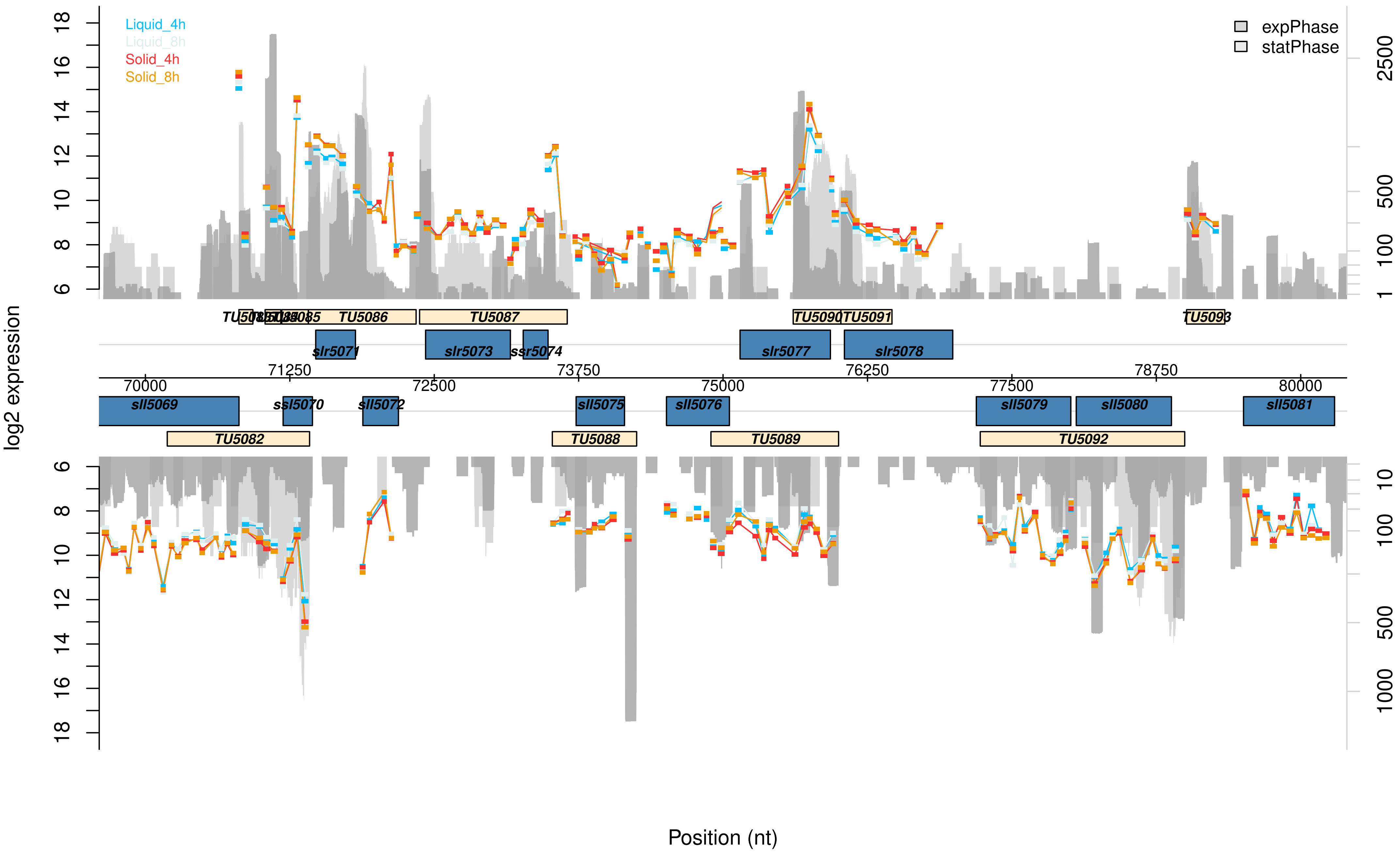

Liquid_4h
Liquid_8h
Solid_4h
Solid_8h
expPhase
statPhase
log2 expression
Position (nt)
TU5086
TU5087
TU5090
TU5091
slr5071
slr5073
ssr5074
slr5077
slr5078
70000
71250
72500
73750
75000
76250
77500
78750
80000
sll5069
ssl5070
sll5072
sll5075
sll5076
sll5079
sll5080
sll5081
TU5082
TU5088
TU5089
TU5092

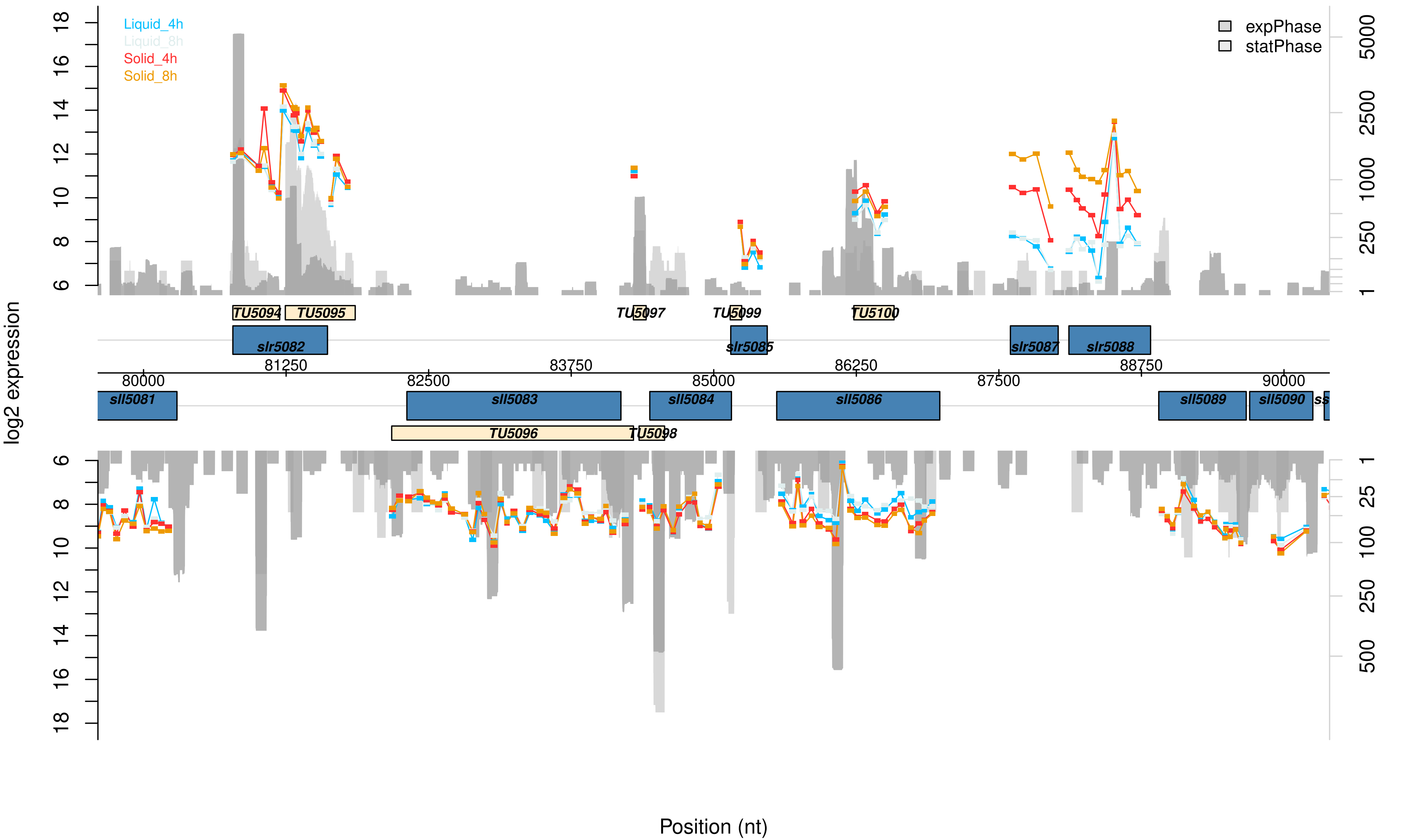

Liquid_4h
Liquid_8h
Solid_4h
Solid_8h
expPhase
statPhase
log2 expression
Position (nt)
TU5094
TU5095
TU5097
TU5099
TU5100
slr5082
slr5085
slr5087
slr5088
sll5081
sll5083
sll5084
sll5086
sll5089
sll5090
TU5096
TU5098
80000
81250
82500
83750
85000
86250
87500
88750
90000

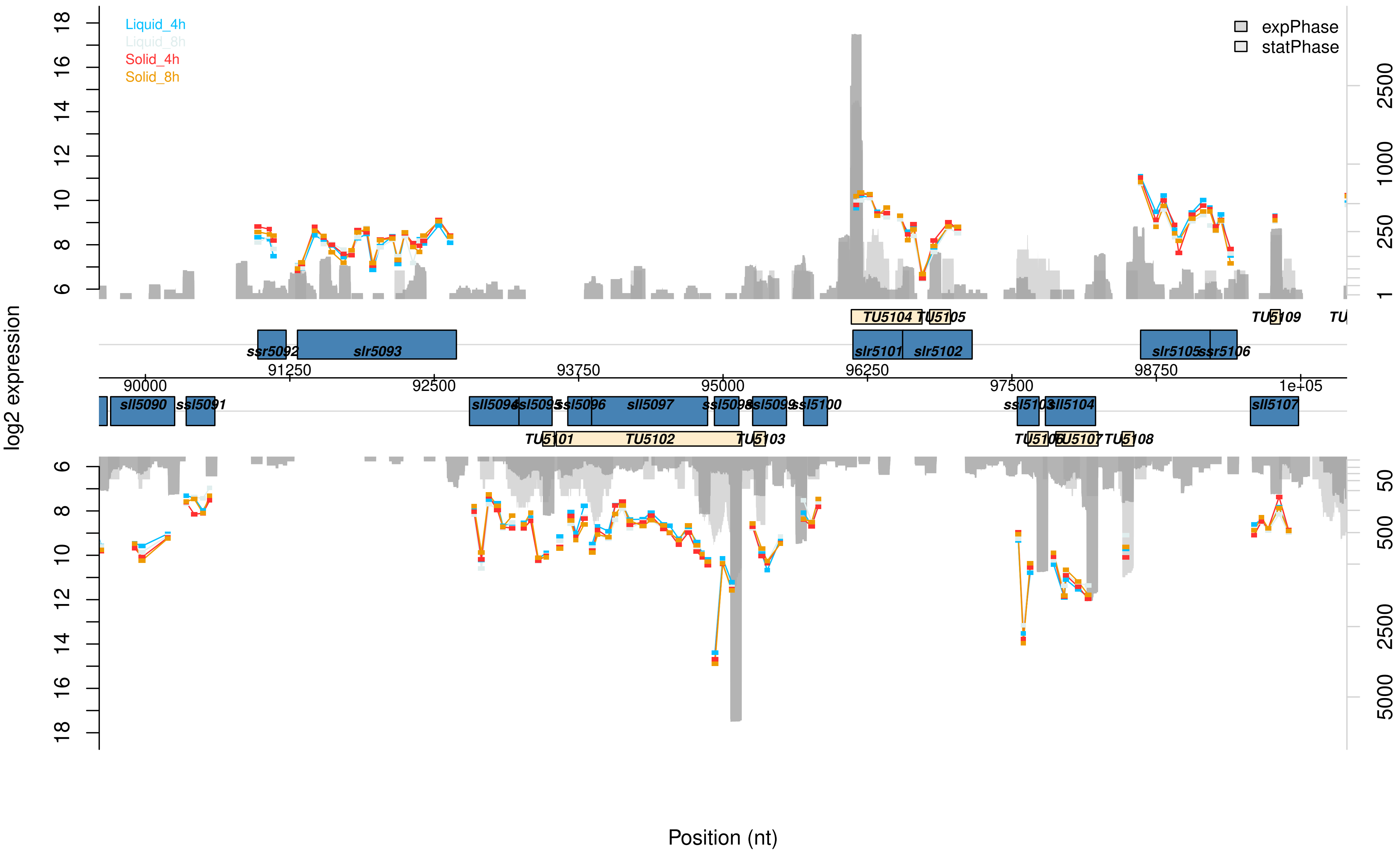

Liquid_4h
Liquid_8h
Solid_4h
Solid_8h
expPhase
statPhase
log2 expression
Position (nt)
TU5104
TU5105
TU5109
ssr5092
slr5093
slr5101
slr5102
slr5105
ssr5106
91250
93750
96250
98750
90000
92500
95000
97500
1e+05
sll5090
ssl5091
sll5094
ssl5095
ssl5096
sll5097
ssl5098
ssl5099
ssl5100
ssl5103
sll5104
sll5107
TU5101
TU5102
TU5103
TU5106
TU5107
TU5108

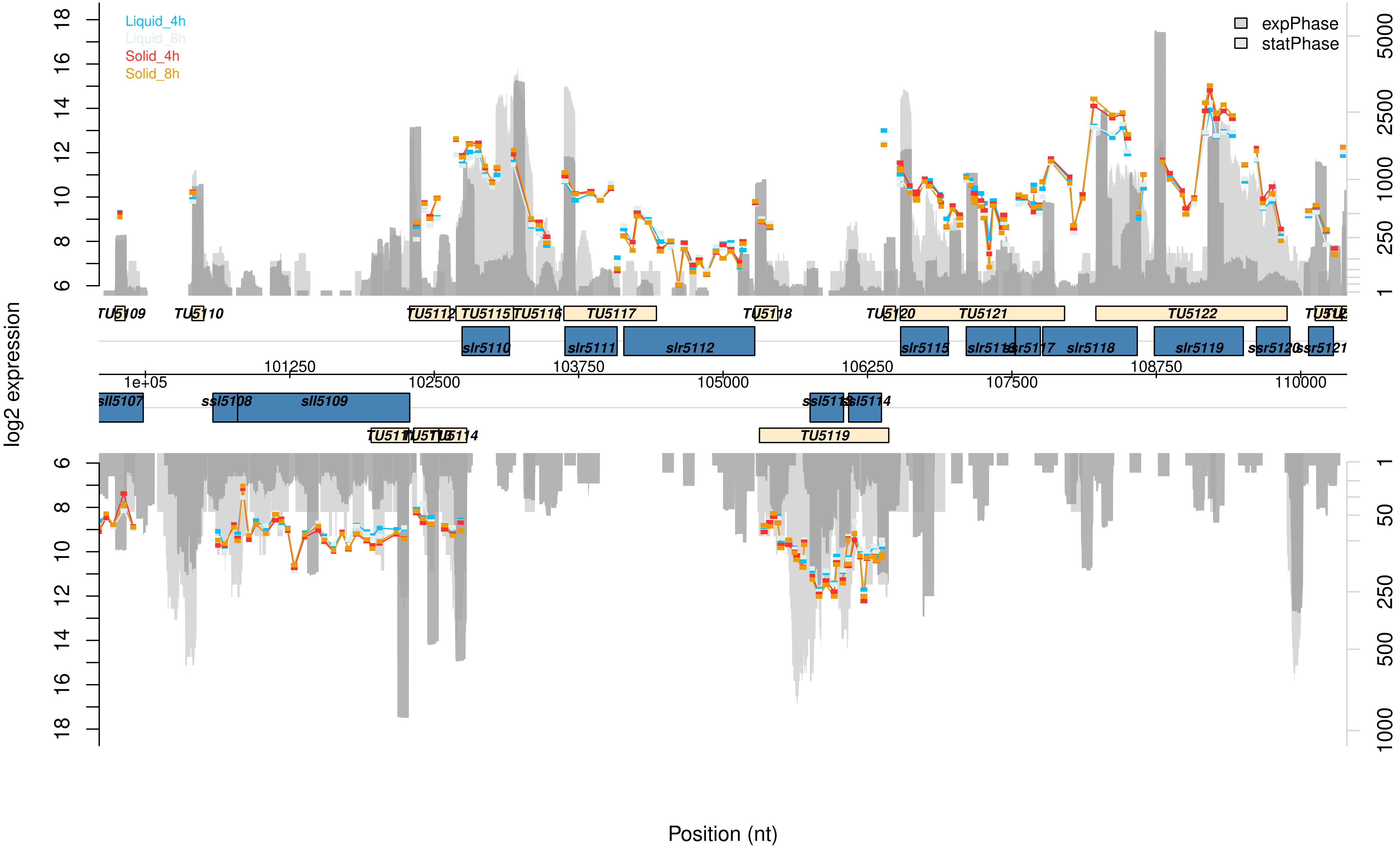

Liquid_4h
Liquid_8h
Solid_4h
Solid_8h
expPhase
statPhase
log2 expression
Position (nt)
TU5109
TU5110
TU5112
TU5115
TU5116
TU5117
TU5118
TU5120
TU5121
TU5122
slr5110
slr5111
slr5112
slr5115
slr5118
slr5119
ssr5121
101250
102500
103750
105000
106250
107500
108750
110000
1e+05
sll5107
ssl5108
sll5109
TU5119

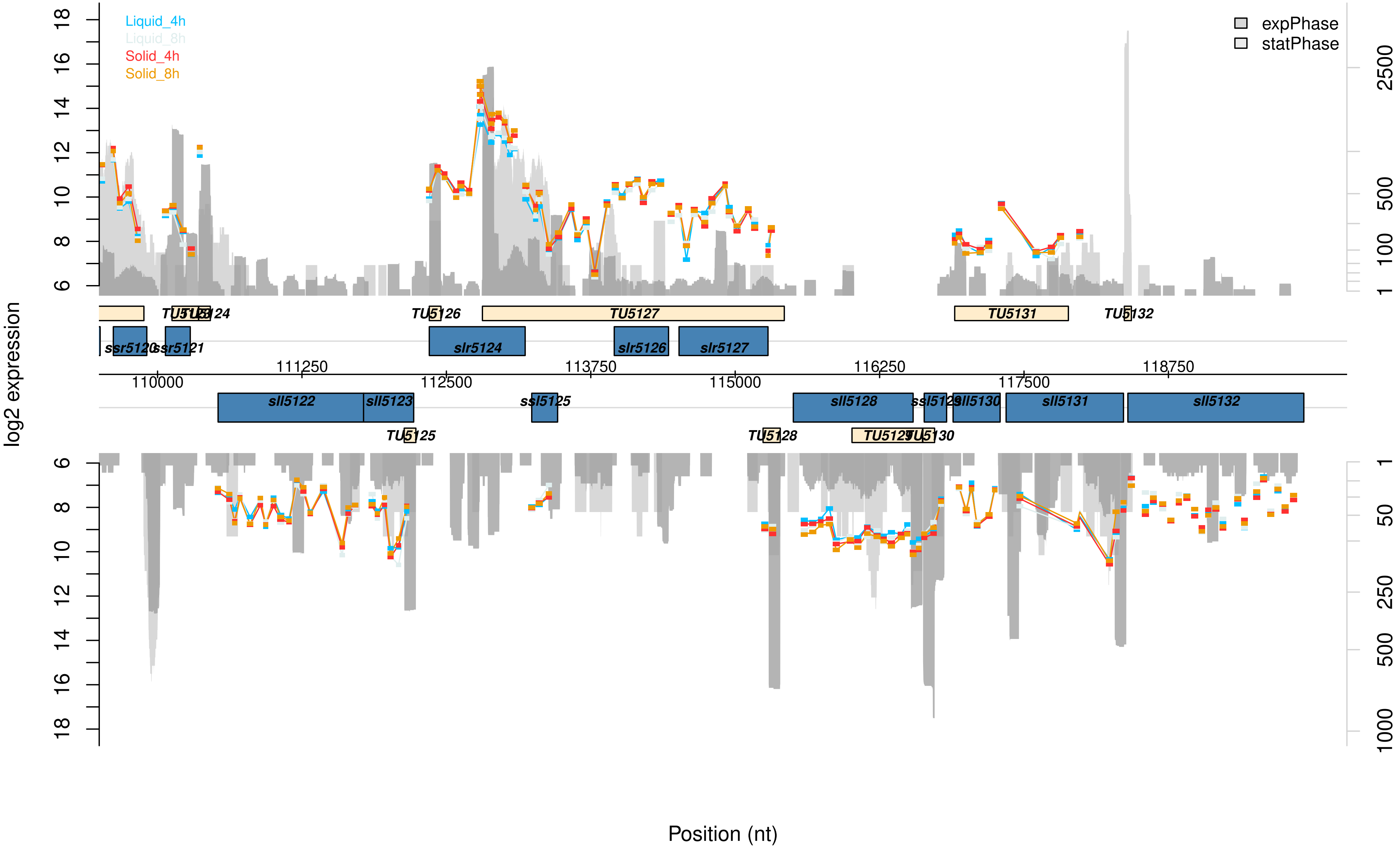

Liquid_4h
Liquid_8h
Solid_4h
Solid_8h
expPhase
statPhase
log2 expression
Position (nt)
18
16
14
12
10
8
6
2500
500
100
1
TU5124
TU5126
TU5127
TU5131
TU5132
ssr5120
ssr5121
slr5124
slr5126
slr5127
111250
113750
116250
118750
110000
112500
115000
117500
sll5122
sll5123
ssl5125
sll5128
ssl5129
sll5130
sll5131
sll5132
TU5125
TU5128
TU5129
TU5130
50
250
1000

Data Set S5: **Graphical visualization of the microarray results of the pSYSX plasmid.** This microarray data presents changes between Synechocystis cells of a sessile and planktonic lifestyle after 4 or 8 h, that are located on the pSYSX plasmid. The transcription values for planktonic cells after 4 h (liquid_4h, blue) or 8 h (liquid_8h, ice blue) and sessile cells after 4 h (solid_4h, red) or 8 h (solid_8h, orange) are given in log2 scale. The read numbers of exponential (dark grey) or stationary phase (light grey) grown cells are shown in log2 scale and are extracted from Kopf *et al.* (2014). Supporting data can be found in the Supplementary Data Table.

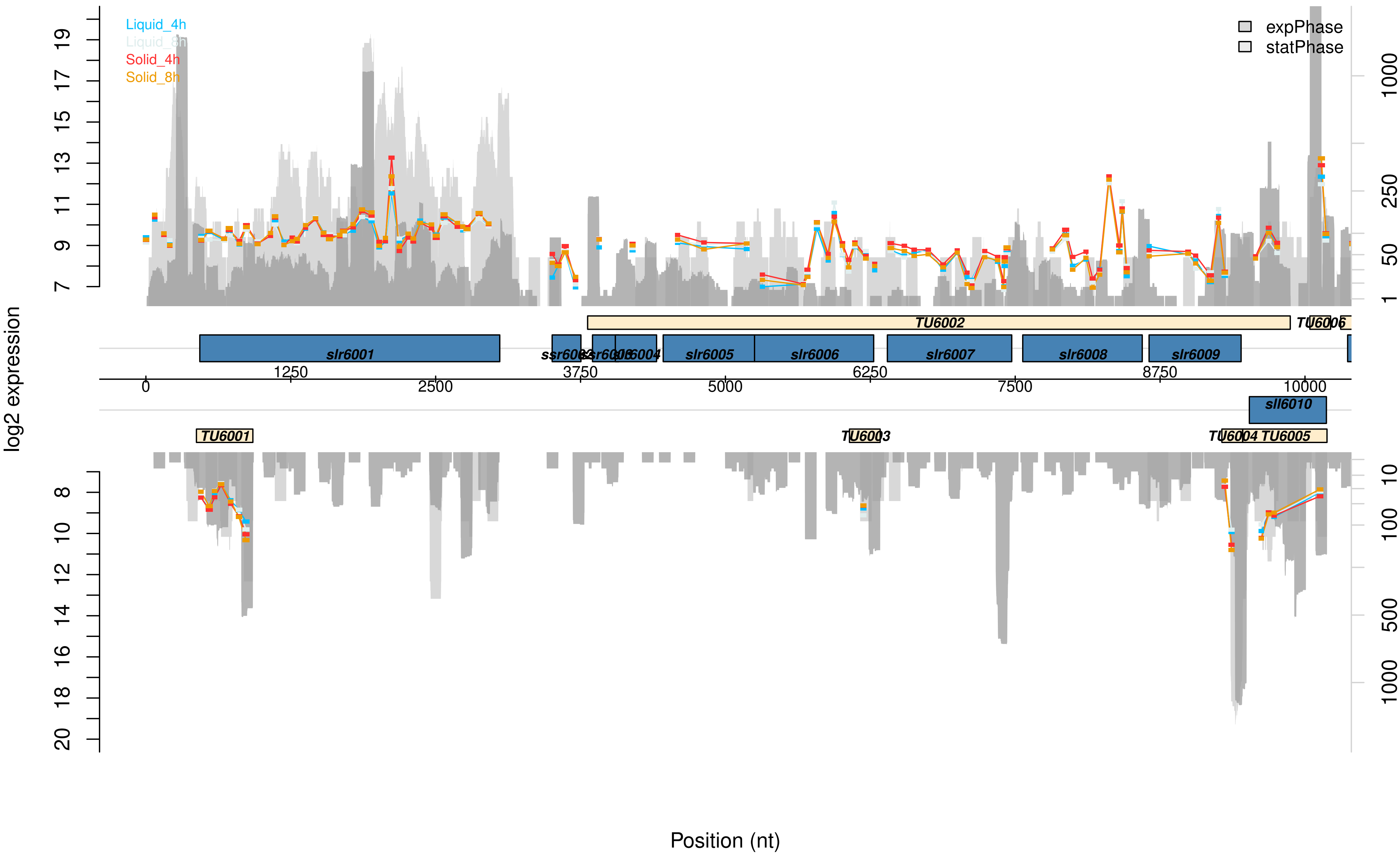

Liquid_4h
Liquid_8h
Solid_4h
Solid_8h
expPhase
statPhase
log2 expression
7
9
11
13
15
17
19
1
50
250
1000
TU6002
slr6001
slr6005
slr6006
slr6007
slr6008
slr6009
sll6010
TU6001
TU6005
0
1250
2500
3750
5000
6250
7500
8750
10000
8
10
12
14
16
18
20
10
100
500
1000
Position (nt)

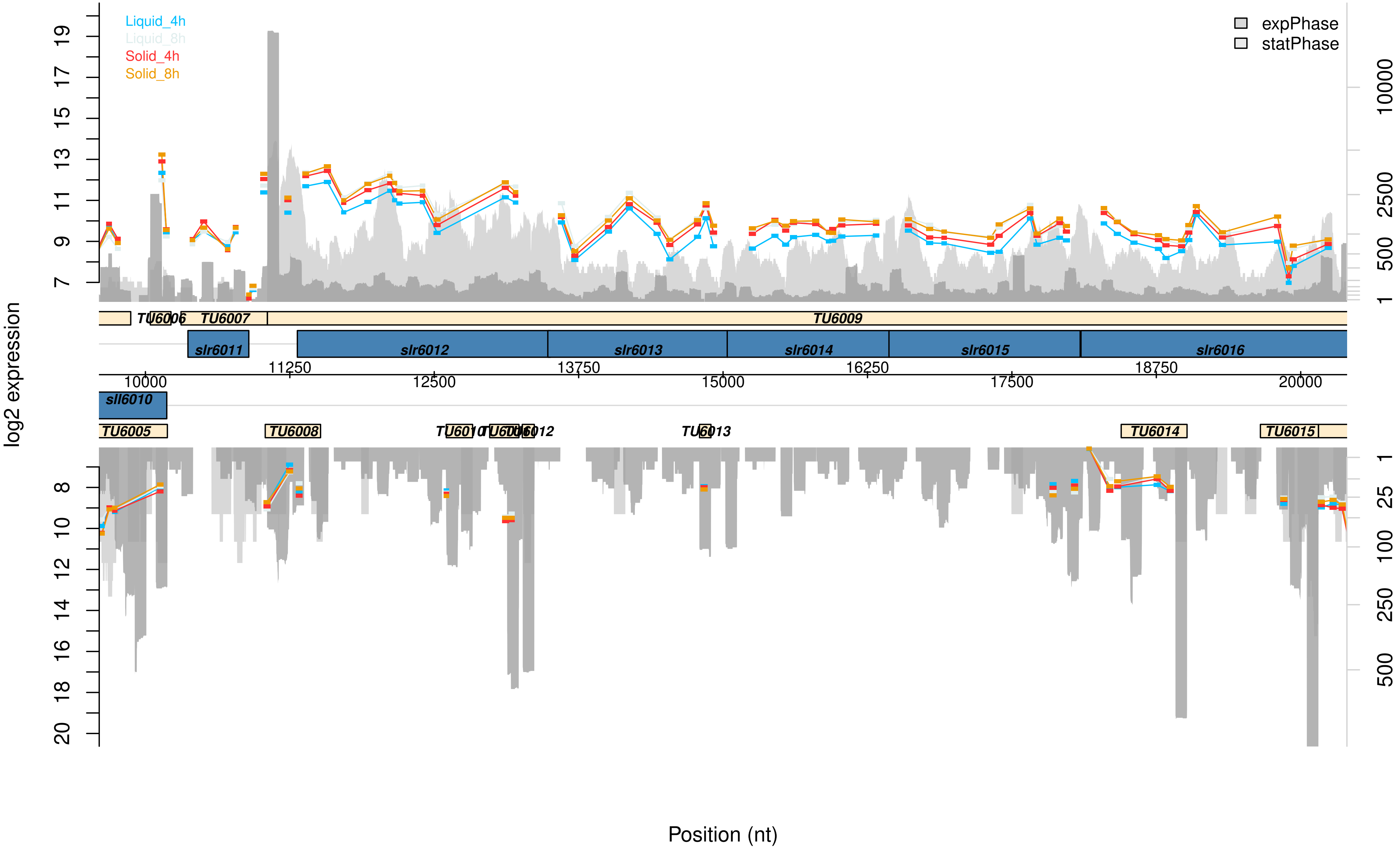

Liquid_4h
Liquid_8h
Solid_4h
Solid_8h
expPhase
statPhase
log2 expression
TU6005
TU6006
TU6007
TU6008
TU6009
TU6013
TU6014
TU6015
sll6010
slr6011
slr6012
slr6013
slr6014
slr6015
slr6016
10000
11250
12500
13750
15000
16250
17500
18750
20000
Position (nt)

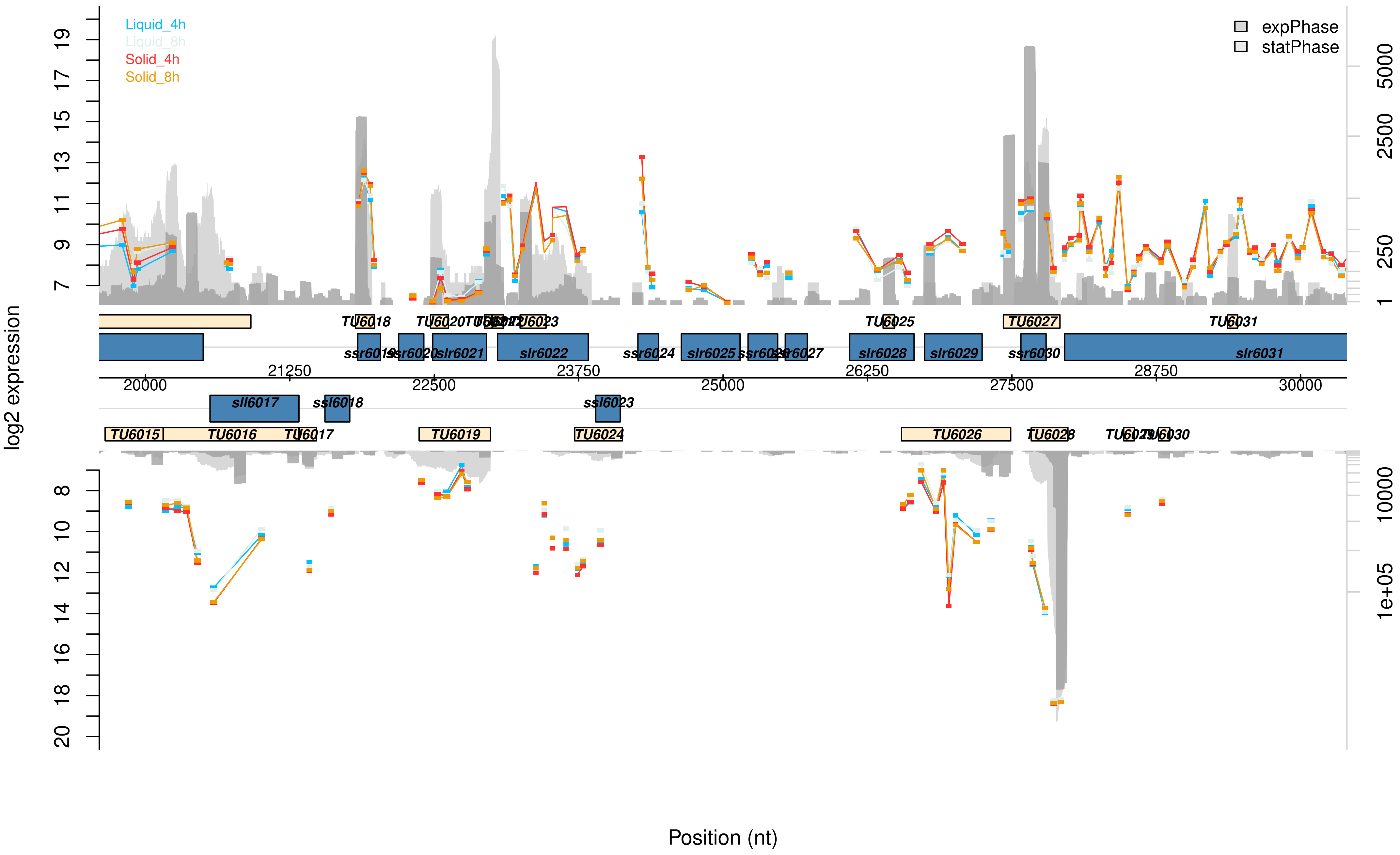

Liquid_4h
Liquid_8h
Solid_4h
Solid_8h
expPhase
statPhase
log2 expression
Position (nt)
TU6015
TU6016
TU6017
TU6018
TU6019
TU6023
TU6024
TU6025
TU6026
TU6027
TU6031
ssr6019
slr6021
slr6022
ssr6024
slr6025
slr6027
slr6028
slr6029
ssr6030
slr6031
sll6017
ssl6018
ssl6023
20000
21250
22500
23750
25000
26250
27500
28750
30000
5000
2500
250
1
10000
1e+05

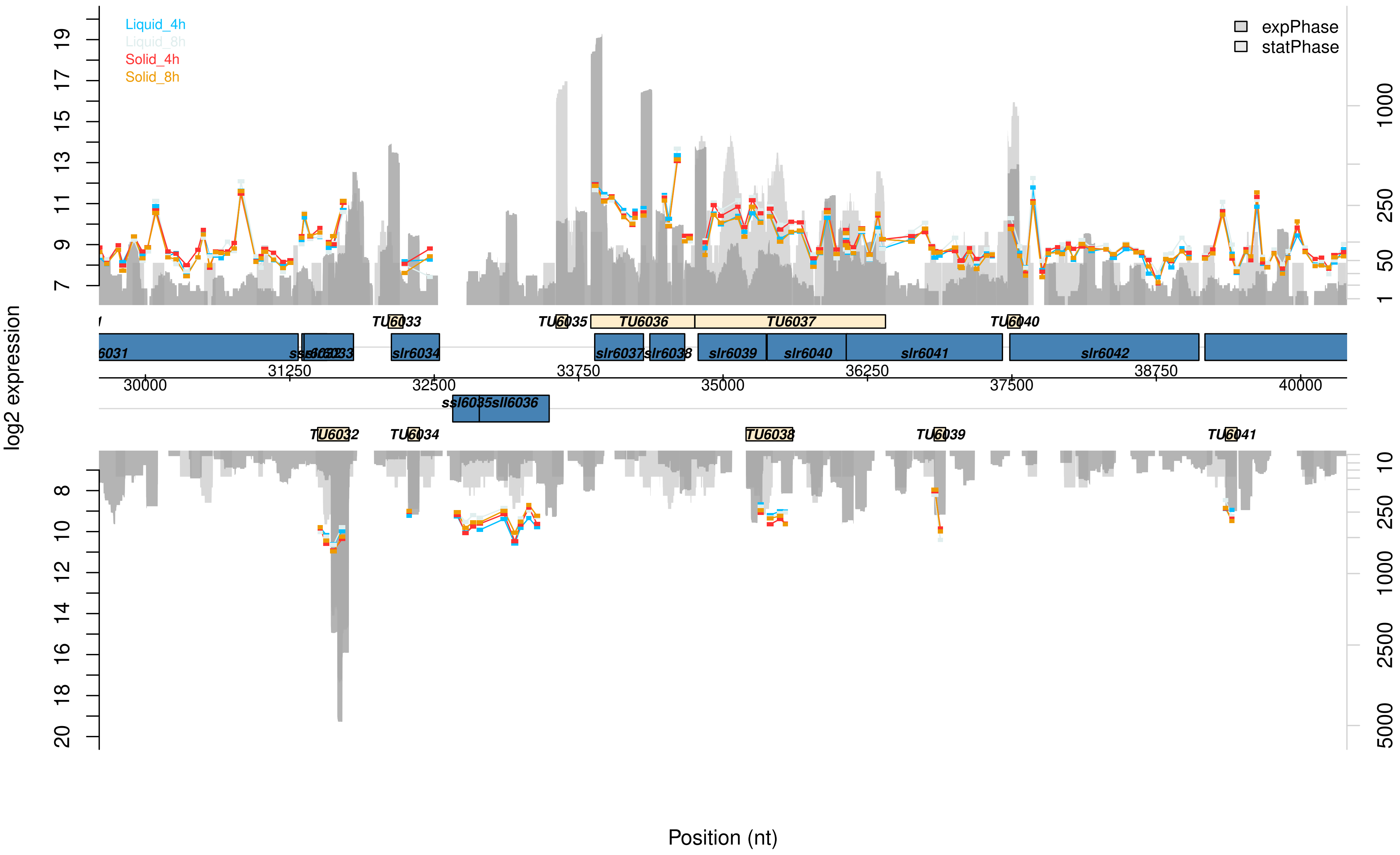

Liquid_4h
Liquid_8h
Solid_4h
Solid_8h
expPhase
statPhase
log2 expression
Position (nt)
TU6033
TU6035
TU6036
TU6037
TU6040
slr6034
slr6037
slr6038
slr6039
slr6040
slr6041
slr6042
ssl6035
sll6036
TU6032
TU6034
TU6038
TU6039
TU6041
30000
31250
32500
33750
35000
36250
37500
38750
40000

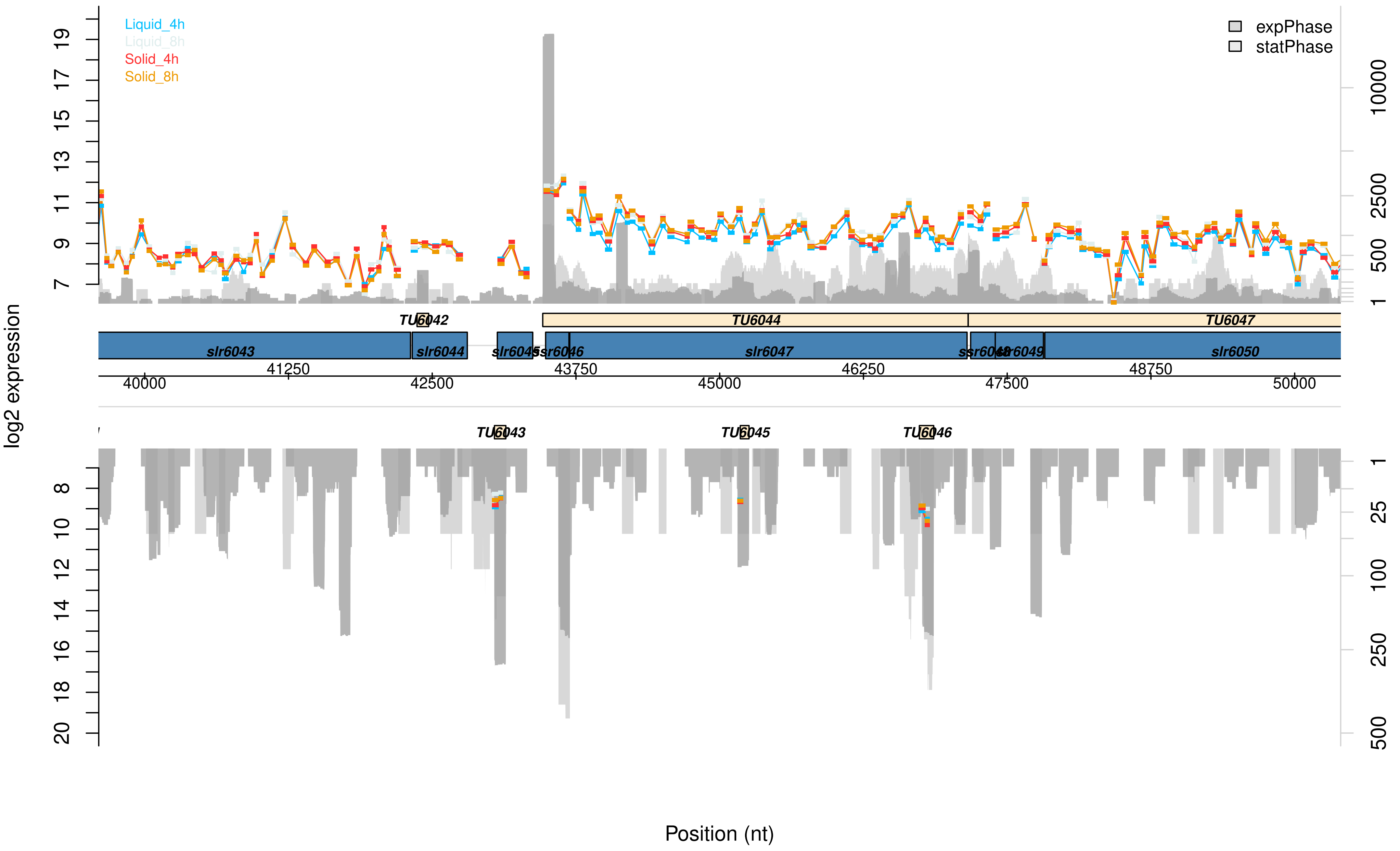

Liquid_4h
Liquid_8h
Solid_4h
Solid_8h
expPhase
statPhase
log2 expression
TU6042
TU6044
TU6047
slr6043
slr6044
slr6045
ssr6046
slr6047
ssr6048
slr6049
slr6050
41250
43750
46250
48750
40000
42500
45000
47500
50000
TU6043
TU6045
TU6046
Position (nt)

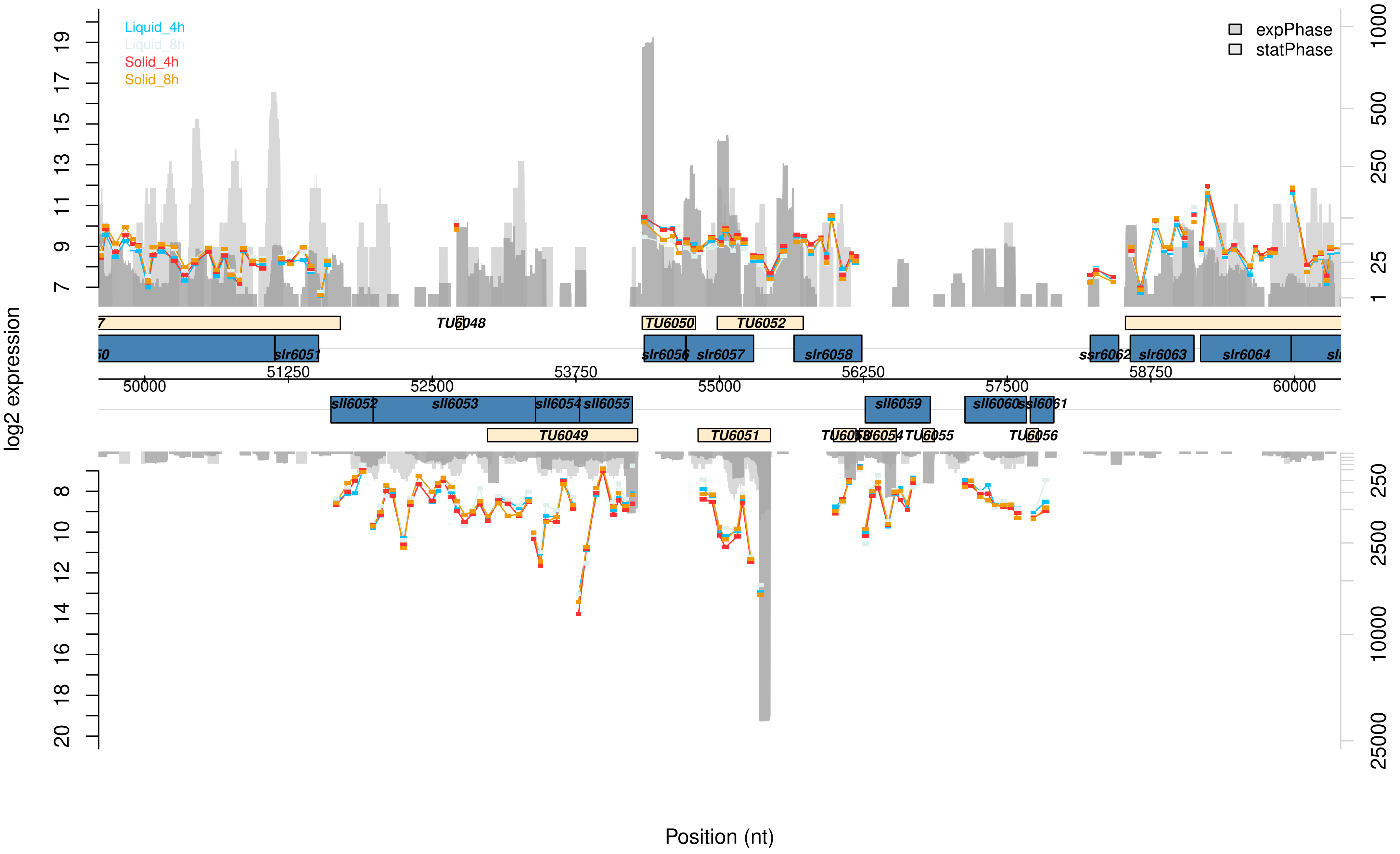
Liquid_4h
Liquid_8h
Solid_4h
Solid_8h
expPhase
statPhase
log2 expression
Position (nt)
TU6048
TU6050
TU6052
slr6051
slr6056
slr6057
slr6058
ssr6062
slr6063
slr6064
sll6052
sll6053
sll6054
sll6055
sll6059
sll6060
ssl6061
TU6049
TU6051
TU6055
TU6056
50000
51250
52500
53750
55000
56250
57500
58750
60000

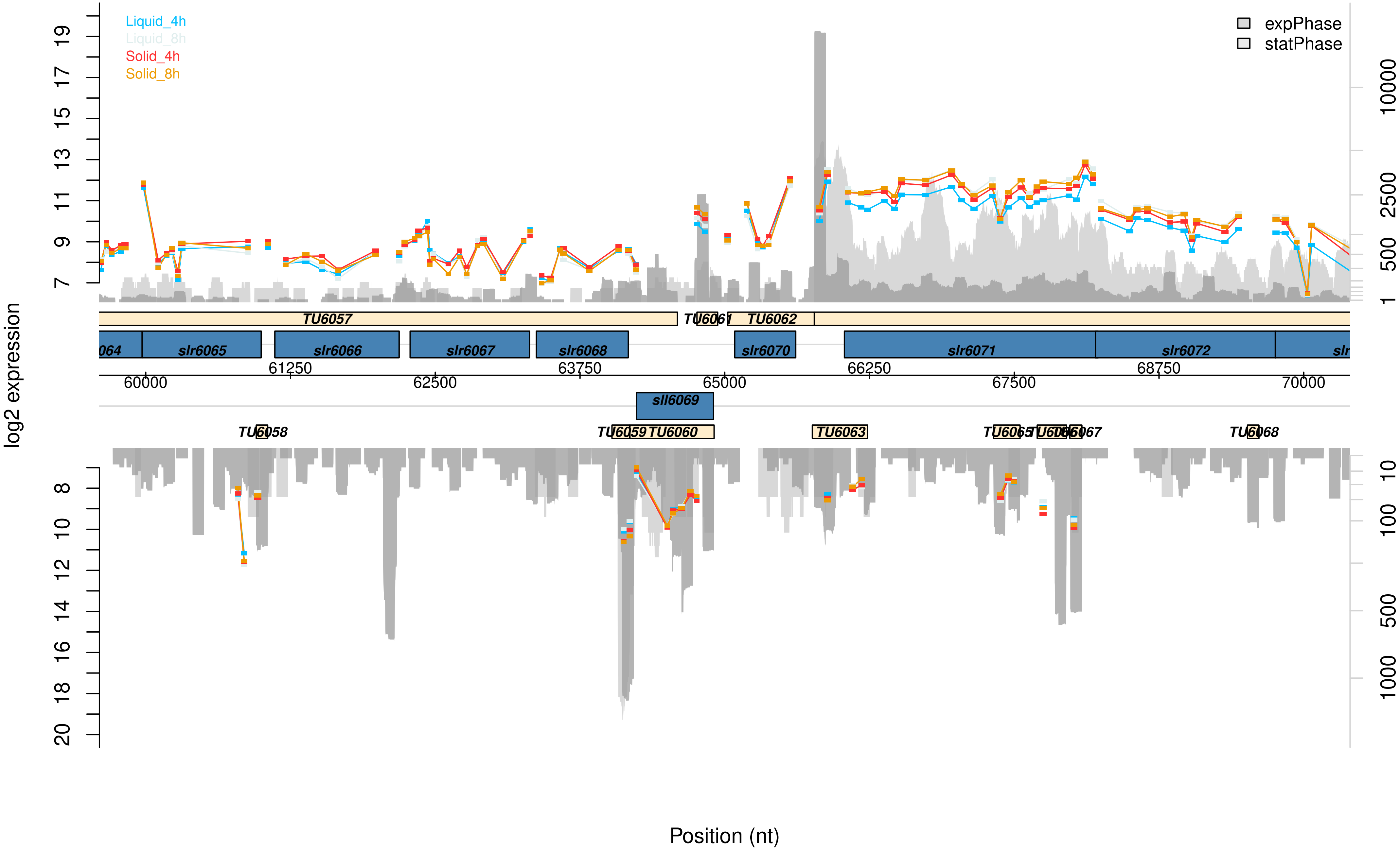

Liquid_4h
Liquid_8h
Solid_4h
Solid_8h
expPhase
statPhase
log2 expression
TU6057
TU6061
TU6062
064
slr6065
slr6066
slr6067
slr6068
slr6070
slr6071
slr6072
slr
sll6069
TU6058
TU6059
TU6060
TU6063
TU6065
TU6066
TU6067
TU6068
60000
61250
62500
63750
65000
66250
67500
68750
70000
Position (nt)

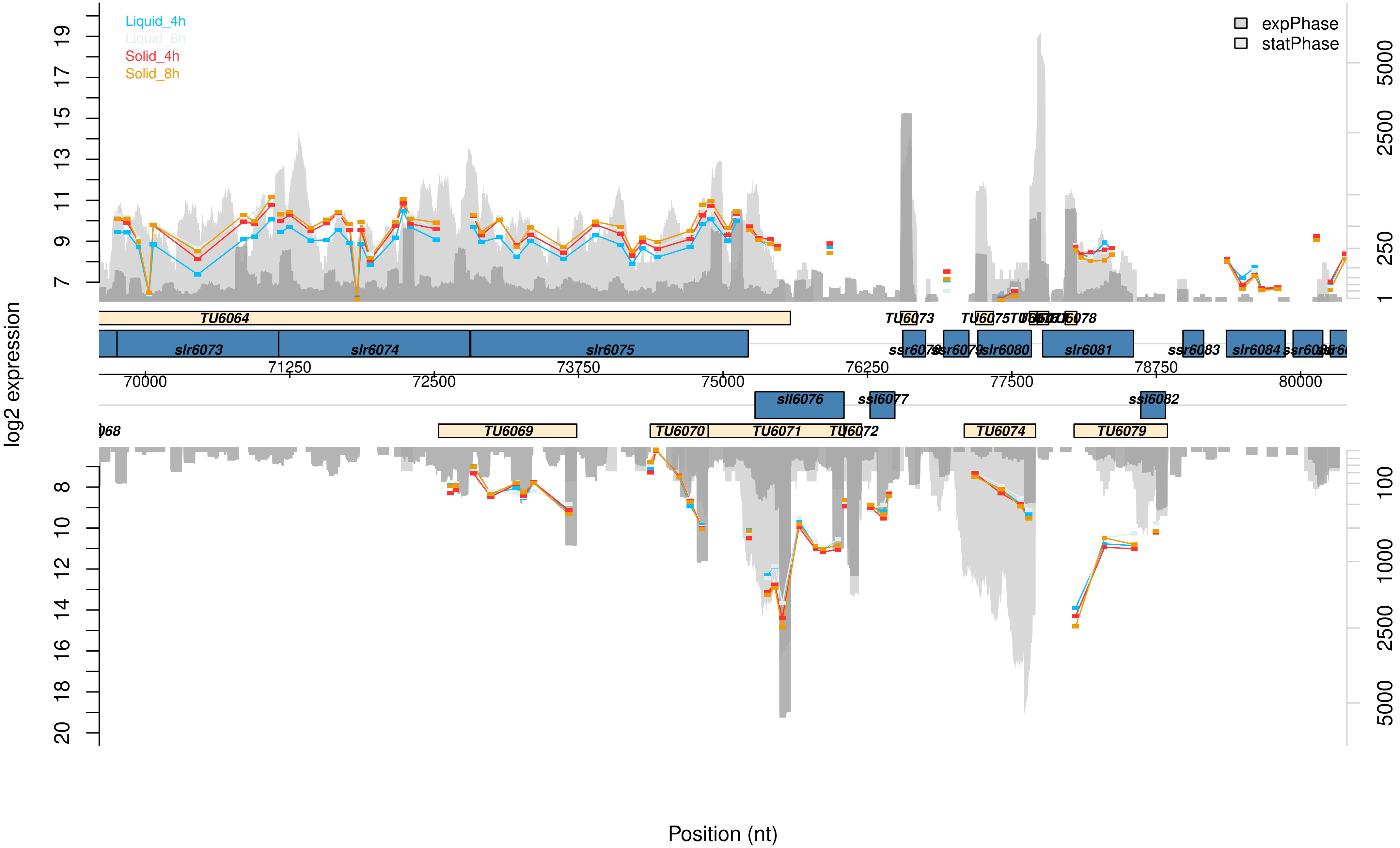

Liquid_4h
Liquid_8h
Solid_4h
Solid_8h
expPhase
statPhase
log2 expression
Position (nt)
TU6064
TU6073
TU6079
slr6073
slr6074
slr6075
slr6080
slr6081
ssr6083
slr6084
sll6076
ssl6077
ssl6082
TU6069
TU6070
TU6071
TU6072
TU6074
70000
71250
72500
73750
75000
76250
77500
78750
80000

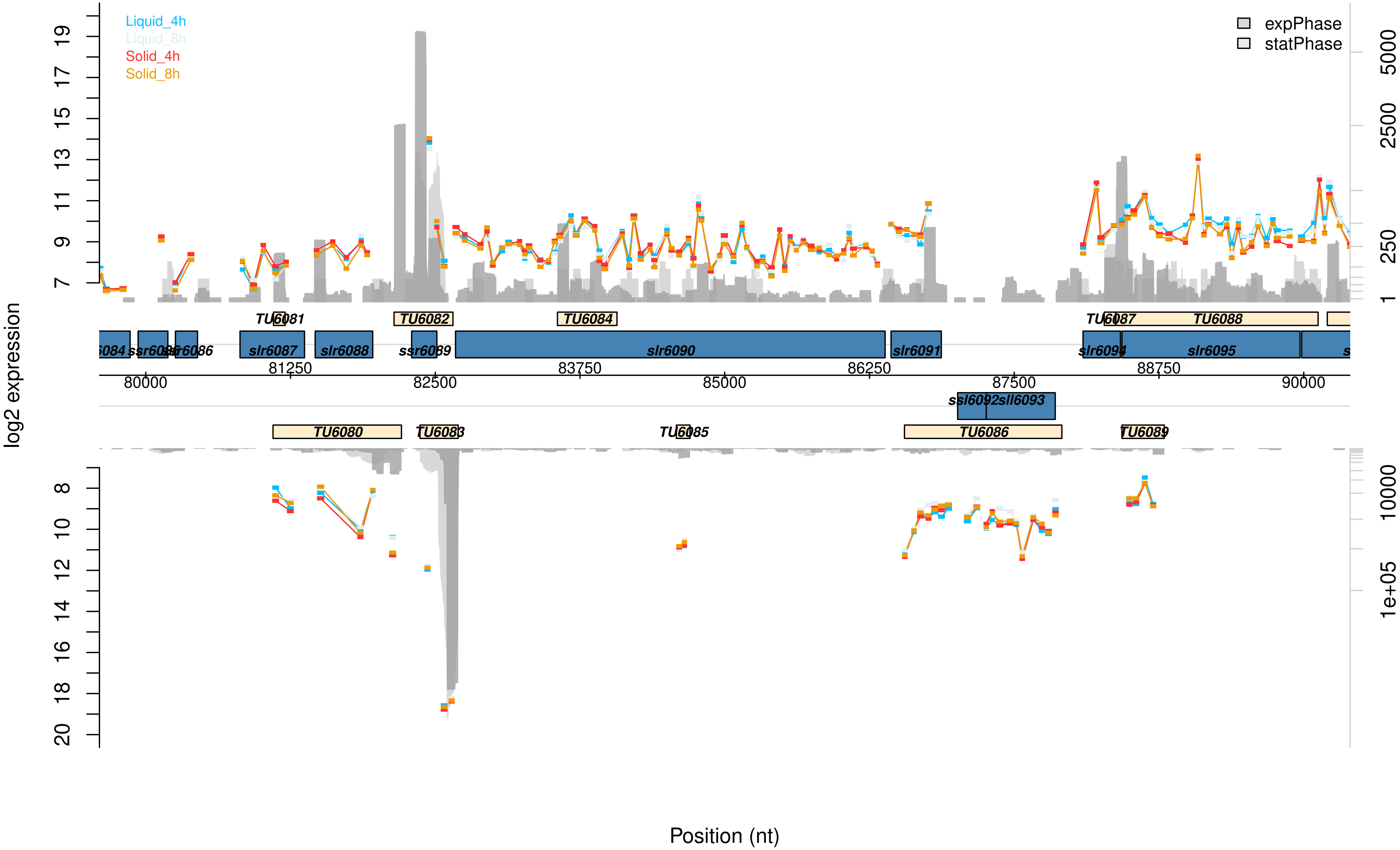

Liquid_4h
Liquid_8h
Solid_4h
Solid_8h
expPhase
statPhase
log2 expression
Position (nt)
TU6080
TU6081
TU6082
TU6083
TU6084
TU6085
TU6086
TU6087
TU6088
TU6089
ssr6086
slr6087
slr6088
ssr6089
slr6090
slr6091
ssl6092
sll6093
slr6094
slr6095
80000
81250
82500
83750
85000
86250
87500
88750
90000

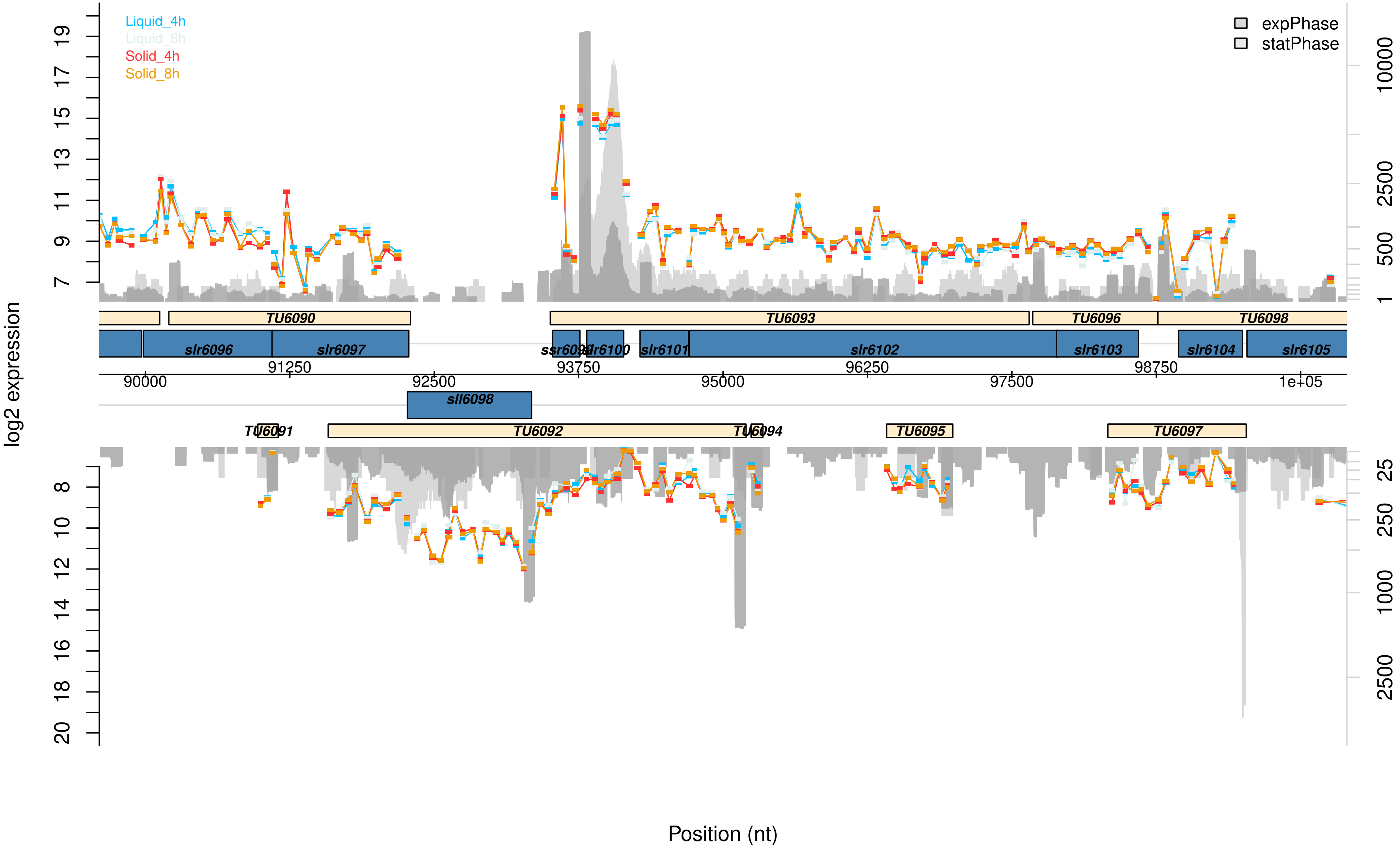

Liquid_4h
Liquid_8h
Solid_4h
Solid_8h
expPhase
statPhase
log2 expression
TU6090
TU6093
TU6096
TU6098
slr6096
slr6097
ssr6099
slr6100
slr6101
slr6102
slr6103
slr6104
slr6105
sll6098
TU6091
TU6092
TU6094
TU6095
TU6097
90000
91250
92500
93750
95000
96250
97500
98750
1e+05
Position (nt)

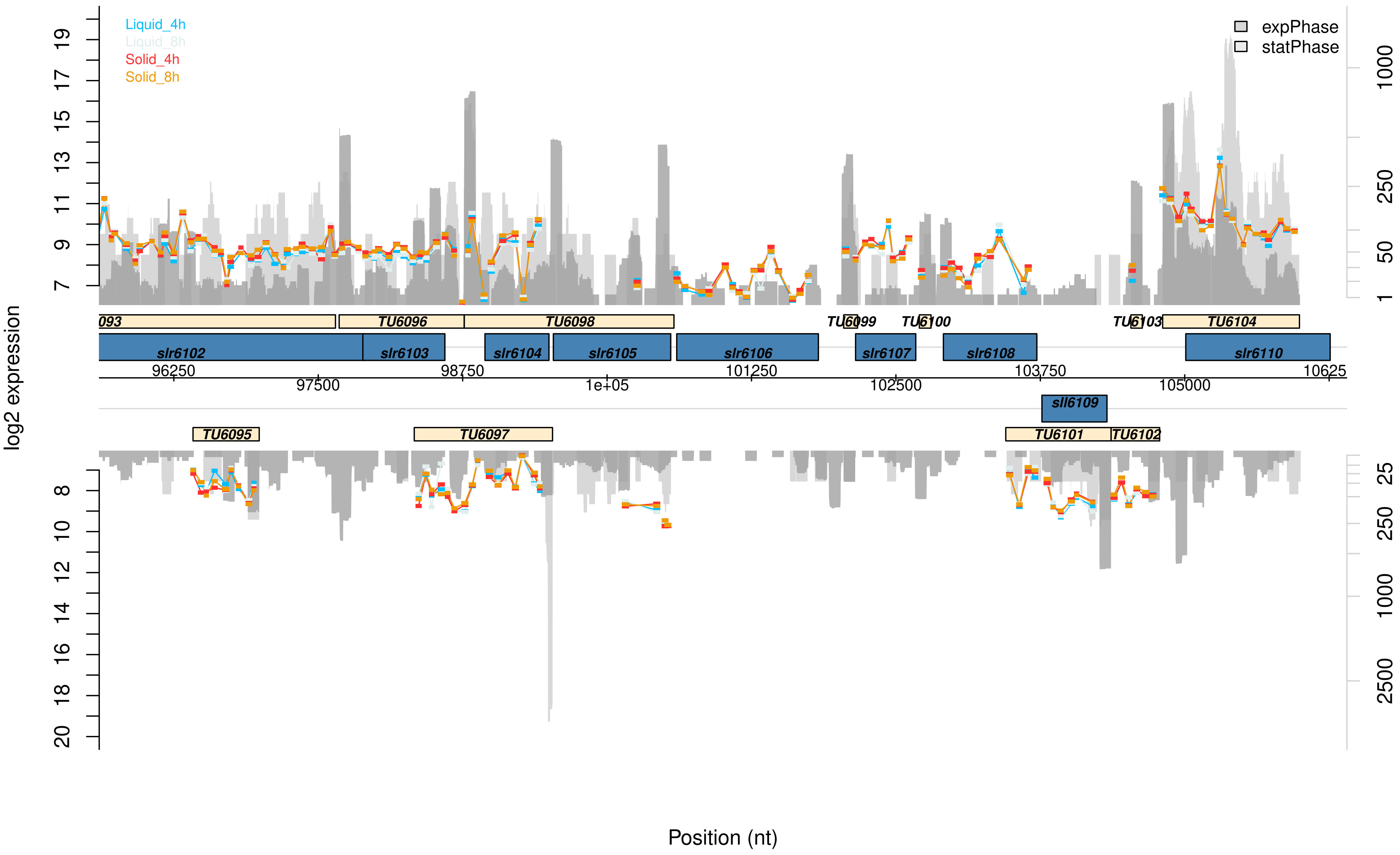
Liquid_4h
Liquid_8h
Solid_4h
Solid_8h
expPhase
statPhase
log2 expression
093
TU6096
TU6098
TU6099
TU6100
TU6103
TU6104
slr6102
slr6103
slr6104
slr6105
slr6106
slr6107
slr6108
slr6110
96250
97500
98750
1e+05
101250
102500
103750
105000
10625
sll6109
TU6095
TU6097
TU6101
TU6102
Position (nt)
