## Supplementary information for "Minor pilin genes are involved in motility and natural competence in *Synechocystis* sp. PCC 6803"

**Running Title**: Minor Pilins of *Synechocystis* sp. PCC 6803

**Keywords**

minor pilin, natural competence, surface acclimation, type IV pili, second messenger, cyanobacteria

**Figure S1**: Schematic representation and verification of different mutants of the *pilA5-pilA6* operon.
**Figure S2**: Pili diameter of minor pilin deletion mutants.
**Figure S3**: Volcano blots of Fig. 6 with data labels.
**Figure S4**: Synteny analyses of the *Synechocystis* minor pilin PilA5 (A), and the PilX homologs Slr0226 (B), Slr0442 (C) and Sll1268 (D) using the FlaG tool (Table S6).
**Figure S5**: Conserved Protein Domains in PilX homologs.
**Figure S6**: Predicted three dimensional models of pilins.
**Figure S7**: Secundary structure predictions of major and minor Pilins of *Synechocystis*.

**Table S1 (excel file)**: Supplementary protein data for evaluation of synteny data for the proteins PilA5, Slr0226, Slr0442 and Sll1268 of Fig. S4.
**Table S2**: BLASTP analysis of proteins from *Synechococcus elongatus* involved in natural transformation (Taton *et al*. 2020) to *Synechocystis* genome.
**Table S3**: Oligonucleotides used in this study
**Table S4**: Plasmids used in this study
**Table S5**: Mutant strains used in this study
**Table S6**: Bioinformatic tools

**Supplementary Data Table (excel-file):** Microarray dataset for *Synechocystis* cells grown in sessile or planktonic conditions for either 4 or 8 h.

**Data Set S1 (pdf-file):** Graphical visualization of the microarray results of the chromosome.
**Data Set S2 (pdf-file):** Graphical visualization of the microarray results of the plasmid pSYSA.

**Data Set S3 (pdf-file):** Graphical visualization of the microarray results of the plasmid pSYSG.
**Data Set S4 (pdf-file):** Graphical visualization of the microarray results of the plasmid pSYSM.
**Data Set S5 (pdf-file):** Graphical visualization of the microarray results of the plasmid pSYSX.

**Video S1 (gif file)**: Bead Assay WT
**Video S2 (gif file)**: Bead Assay Δ*pilA9-slr2019*
**Video S3 (gif file)**: Bead Assay Δ*pilA5-6*

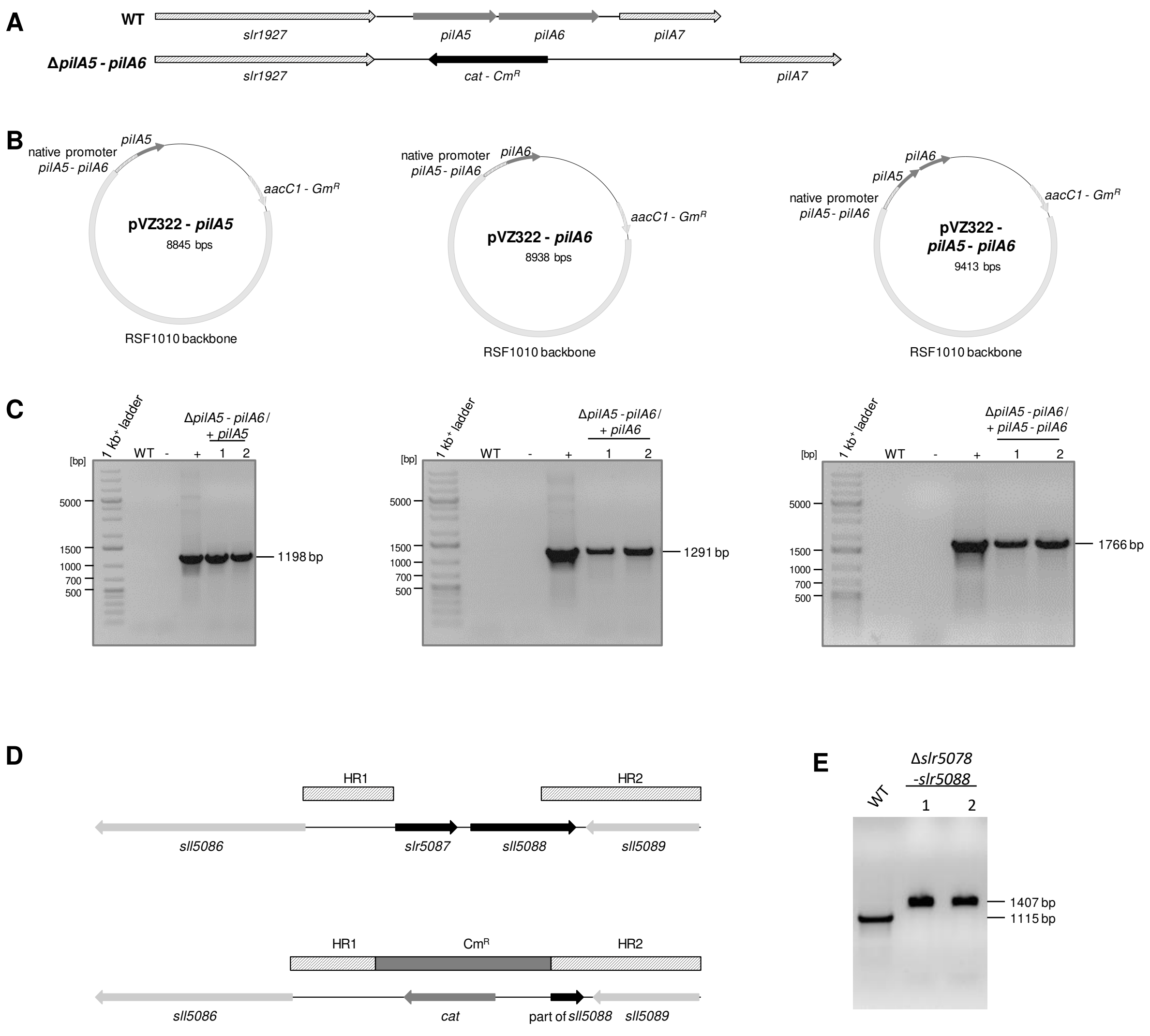


Figure S1: **Schematic representation and verification of different mutants of the *pilA5-pilA6* operon. (A)** Gene arrangement of the *pilA5-pilA6* operon in WT and the deletion mutant Δ*pilA5-pilA6* (described in Wallner *et al*. 2020). **(B)** The genes *pilA5*, *pilA6* or the combination of *pilA5* and *pilA6* have been cloned into the pVZ322 vector under the control of their natural promotor. The gene for aminoglycosidase-3'-phosphotransferase was inactivated during that process. These derivates of the pVZ322 plasmid (NCBI accession number AF100175) have been genetically transferred via three-parental mating into the deletion mutant Δ *pilA5-pilA6*. aacC1 – gentamicin 3-N-acetyltransferase **(C)** Verification of successful plasmid transfer into the deletion mutant Δ*pilA5 - pilA6* via PCR. The chromosomal DNA of the mutants, the wild type (WT), the plasmid used for conjugation as positive control (+) and water as a negative control (-) were used for PCR.

**Schematic representation and verification of the *slr5087-slr5088* operon knock out mutant. (D)** Gene arrangement of the *slr5087-slr5088* operon in WT and the deletion mutant. HR – homologous region; cat - chloramphenicol acetyltransferase; $Cm^R$ – chloramphenicol resistance cassette **(E)** Verification of successful genetic transformation and genetic insertion into *Synechocystis* pSYSM plasmid was tested via PCR. Results for WT and two different clones of Δ*slr5087-slr5088* are shown here. For more details see material and methods section. Marker – 1 kb plus DNA ladder (ThermoScientific, Germany)

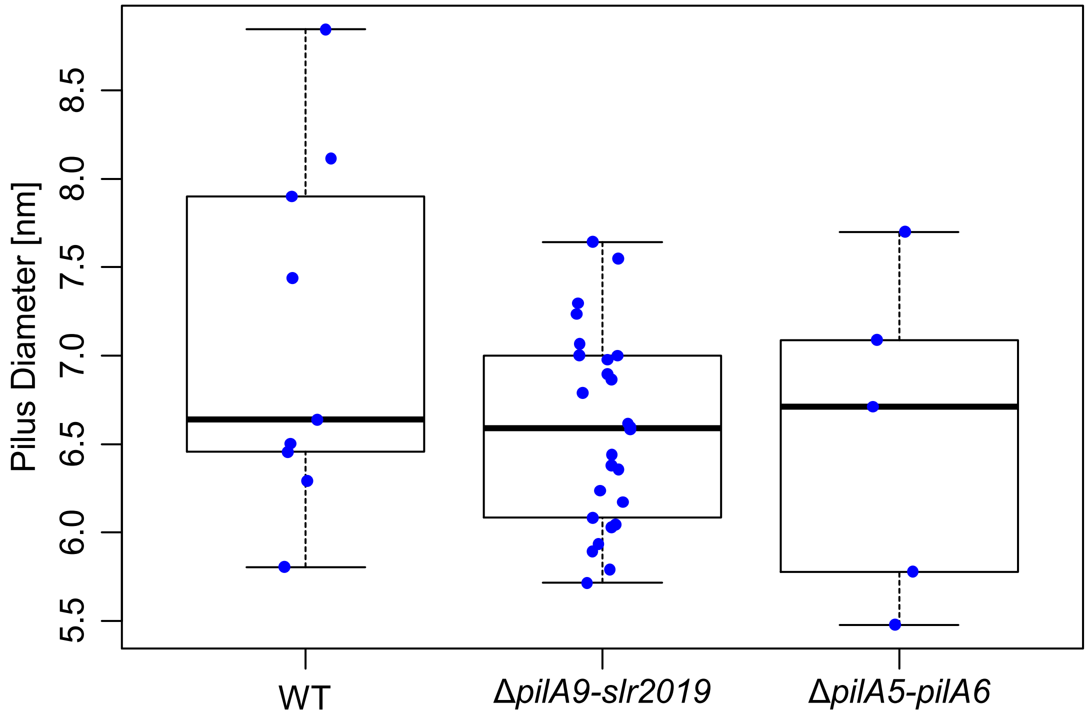


Figure S2: **Pili diameter of minor pilin deletion mutants.** Shown are averaged pili diameters of the WT, Δ*pilA9-slr2019* and Δ*pilA5-pilA6* mutants. Each data point represents the mean of several measurements on one pilus. Measurements have been taken on different TEM pictures of different experiments.

Figure S3: **Volcano blots of Fig. 6 with data labels.**

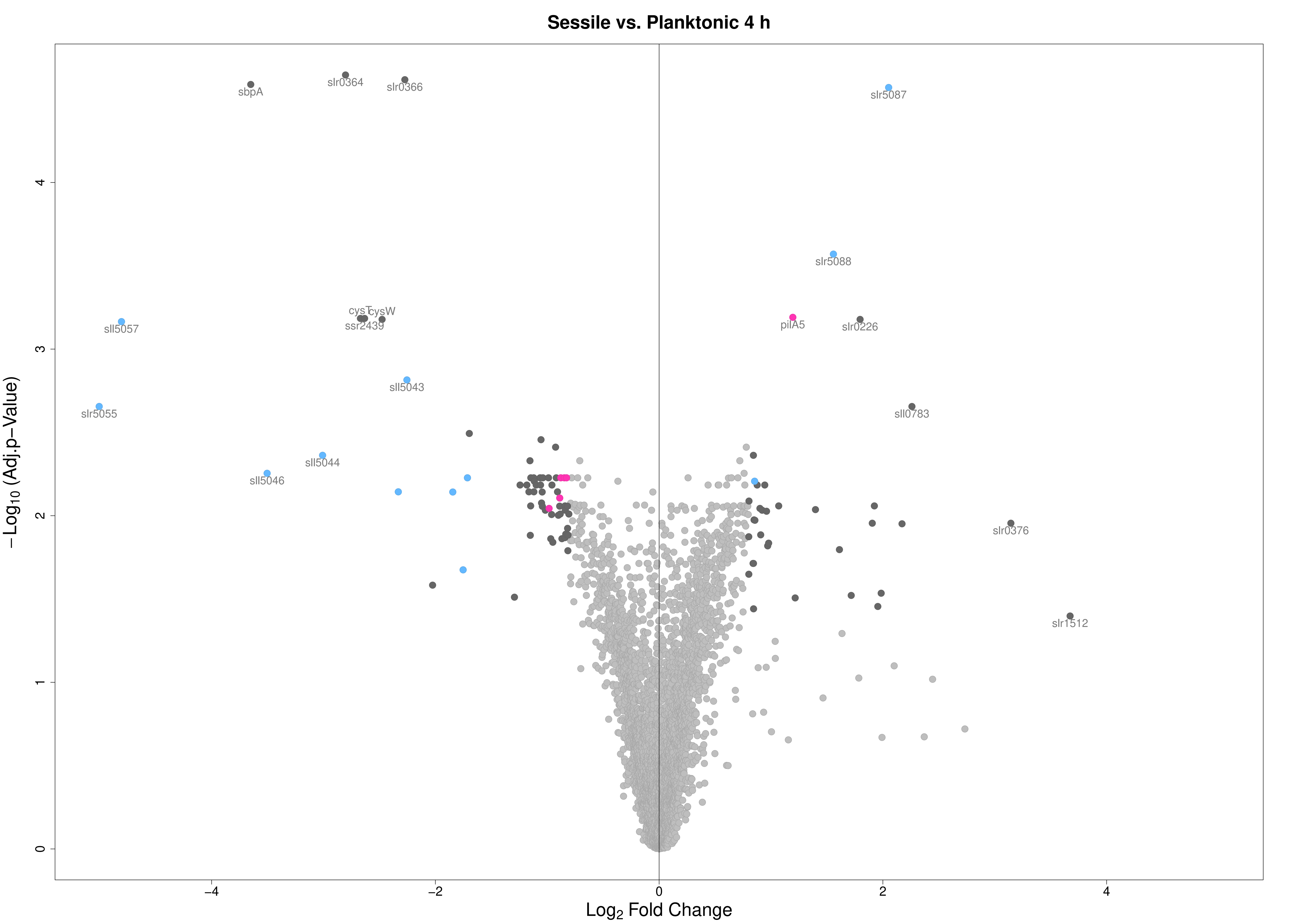
Sessile vs. Planktonic 4 h
−Log10 (Adj.p−Value)
Log2 Fold Change
sbpA
slr0364
slr0366
slr5087
slr5088
sll5057
cysT
cysW
ssr2439
pilA5
slr0226
sll5043
slr5055
sll0783
sll5044
sll5046
slr0376
slr1512
0
1
2
3
4
−4
−2
0
2
4

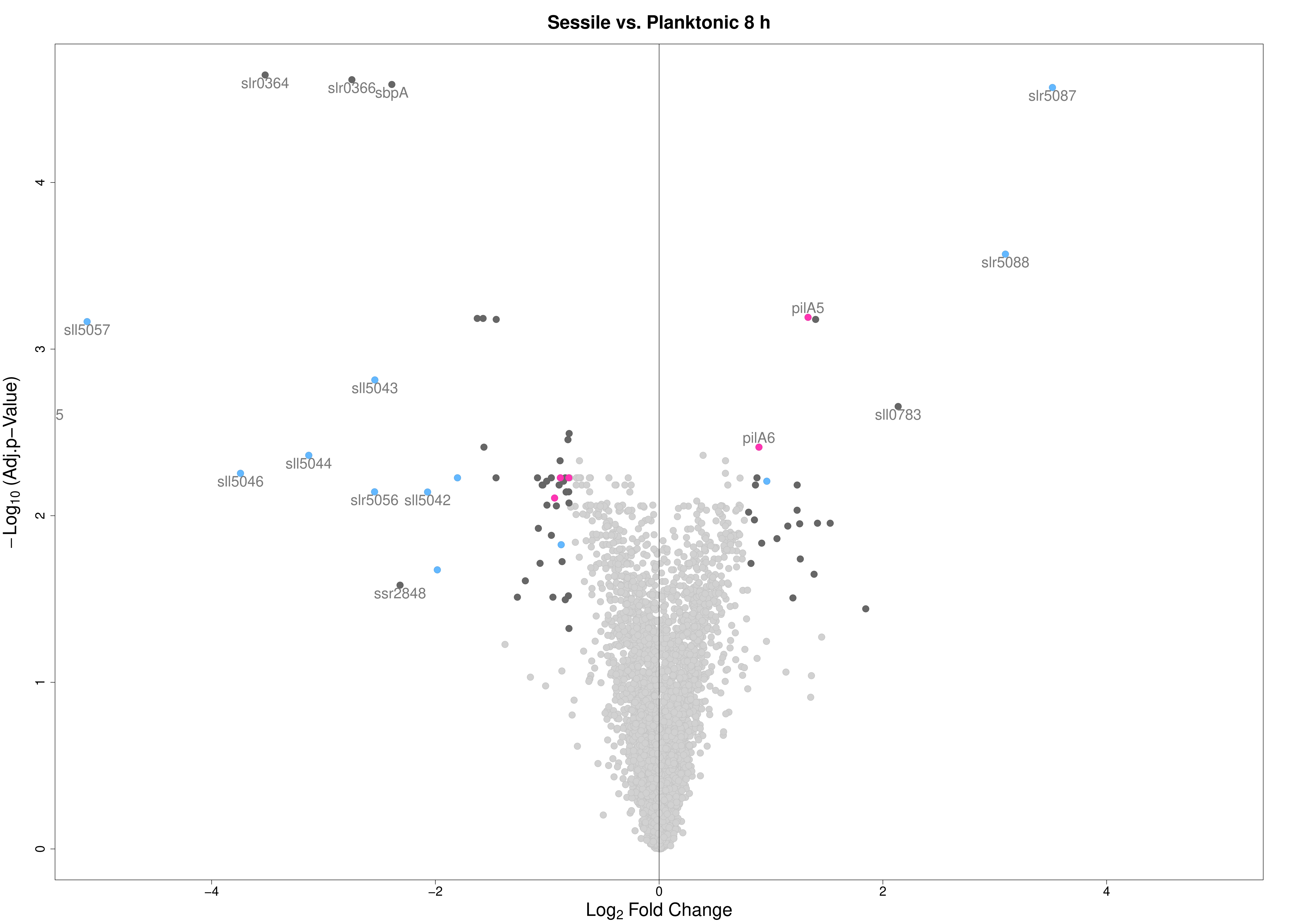
Sessile vs. Planktonic 8 h
slr0364
slr0366
sbpA
slr5087
slr5088
pilA5
sll5057
sll5043
sll0783
5
pilA6
sll5044
sll5046
slr5056
sll5042
ssr2848
−Log10 (Adj.p−Value)
Log2 Fold Change
0
1
2
3
4
−4
−2
0
2
4

Figure S4: **Synteny analyses of the *Synechocystis* minor pilin PilA5 (A), and the PilX homologs Slr0226 (B), Slr0442 (C) and Sll1268 (D) using the FlaGs tool (Table S6).** Data labeling can be found in Table S1.

A

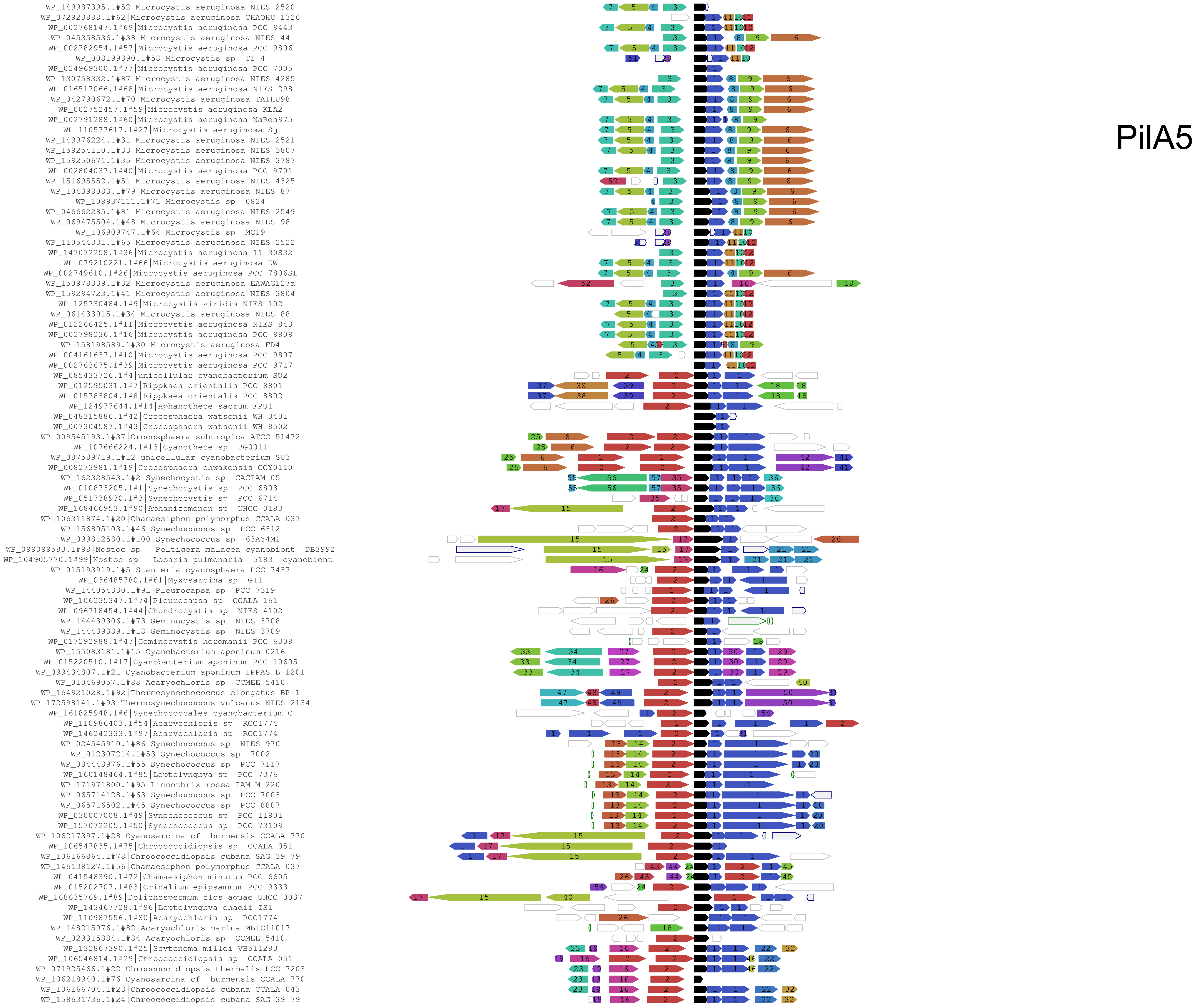
PilA5
WP_149987395.1#52|Microcystis aeruginosa NIES 2520
WP_072923888.1#62|Microcystis aeruginosa CHAOHU 1326
WP_002768147.1#69|Microcystis aeruginosa PCC 9443
WP_045358536.1#38|Microcystis aeruginosa NIES 44
WP_002782954.1#57|Microcystis aeruginosa PCC 9806
WP_008199390.1#58|Microcystis sp T1 4
WP_024969300.1#77|Microcystis aeruginosa PCC 7005
WP_130758332.1#87|Microcystis aeruginosa NIES 4285
WP_016517066.1#68|Microcystis aeruginosa NIES 298
WP_042790672.1#70|Microcystis aeruginosa TAIHU98
WP_002752457.1#59|Microcystis aeruginosa KLA2
WP_002791288.1#60|Microcystis aeruginosa NaRes975
WP_110577617.1#27|Microcystis aeruginosa Sj
WP_149976224.1#31|Microcystis aeruginosa NIES 2521
WP_159254110.1#33|Microcystis aeruginosa NIES 3807
WP_159250671.1#35|Microcystis aeruginosa NIES 3787
WP_002804037.1#40|Microcystis aeruginosa PCC 9701
WP_151695552.1#51|Microcystis aeruginosa NIES 4325
WP_104398083.1#79|Microcystis aeruginosa NIES 87
WP_108937111.1#71|Microcystis sp 0824
WP_046662285.1#81|Microcystis aeruginosa NIES 2549
WP_069475504.1#48|Microcystis aeruginosa NIES 98
WP_106909747.1#64|Microcystis sp MC19
WP_110544331.1#65|Microcystis aeruginosa NIES 2522
WP_147072258.1#36|Microcystis aeruginosa 11 30S32
WP_079210221.1#66|Microcystis aeruginosa KW
WP_002749610.1#26|Microcystis aeruginosa PCC 7806SL
WP_150978339.1#32|Microcystis aeruginosa EAWAG127a
WP_159294723.1#41|Microcystis aeruginosa NIES 3804
WP_125730484.1#9|Microcystis viridis NIES 102
WP_061433015.1#34|Microcystis aeruginosa NIES 88
WP_012266425.1#11|Microcystis aeruginosa NIES 843
WP_002798236.1#16|Microcystis aeruginosa PCC 9809
WP_158198589.1#30|Microcystis aeruginosa FD4
WP_004161637.1#10|Microcystis aeruginosa PCC 9807
WP_002763675.1#39|Microcystis aeruginosa PCC 9717
WP_085433726.1#4|unicellular cyanobacterium SU2
WP_012595031.1#7|Rippkaea orientalis PCC 8801
WP_015783804.1#8|Rippkaea orientalis PCC 8802
WP_124977644.1#14|Aphanothece sacrum FPU1
WP_048315886.1#42|Crocosphaera watsonii WH 0401
WP_007304587.1#43|Crocosphaera watsonii WH 8502
WP_009545193.1#37|Crocosphaera subtropica ATCC 51472
WP_107666224.1#13|Cyanothece sp BG0011
WP_087589719.1#12|unicellular cyanobacterium SU3
WP_008273981.1#19|Crocosphaera chwakensis CCY0110
WP_162328543.1#2|Synechocystis sp CACIAM 05
WP_010873205.1#1|Synechocystis sp PCC 6803
WP_051738930.1#3|Synechocystis sp PCC 6714
WP_168466953.1#90|Aphanizomenon sp UHCC 0183
WP_106311874.1#20|Chamaesiphon polymorphus CCALA 037
WP_156805103.1#46|Synechococcus sp PCC 6312
WP_099812580.1#100|Synechococcus sp 63AY4M1
WP_099099583.1#98|Nostoc sp Peltigera malacea cyanobiont DB3992
WP_104905770.1#99|Nostoc sp Lobaria pulmonaria 5183 cyanobiont
WP_015193919.1#5|Stanieria cyanosphaera PCC 7437
WP_036485780.1#61|Myxosarcina sp GI1
WP_144054330.1#91|Pleurocapsa sp PCC 7319
WP_106235347.1#74|Pleurocapsa sp CCALA 161
WP_096718454.1#44|Chondrocystis sp NIES 4102
WP_144439306.1#73|Geminocystis sp NIES 3708
WP_144439389.1#18|Geminocystis sp NIES 3709
WP_017292988.1#47|Geminocystis herdmanii PCC 6308
WP_155083181.1#15|Cyanobacterium aponinum 0216
WP_015220510.1#17|Cyanobacterium aponinum PCC 10605
WP_099434807.1#21|Cyanobacterium aponinum IPPAS B 1201
WP_010469057.1#88|Acaryochloris sp CCMEE 5410
WP_164921028.1#92|Thermosynechococcus elongatus BP 1
WP_172598141.1#93|Thermosynechococcus vulcanus NIES 2134
WP_161825948.1#6|Synechococcales cyanobacterium C
WP_110986403.1#54|Acaryochloris sp RCC1774
WP_146242333.1#97|Acaryochloris sp RCC1774
WP_024545910.1#86|Synechococcus sp NIES 970
WP_012307214.1#53|Synechococcus sp 7002
WP_084448976.1#55|Synechococcus sp PCC 7117
WP_160148464.1#85|Leptolyngbya sp PCC 7376
WP_171971800.1#95|Limnothrix rosea IAM M 220
WP_065714128.1#63|Synechococcus sp PCC 7003
WP_065716502.1#45|Synechococcus sp PCC 8807
WP_030007008.1#49|Synechococcus sp PCC 11901
WP_157072205.1#50|Synechococcus sp PCC 73109
WP_106217397.1#28|Cyanosarcina cf burmensis CCALA 770
WP_106547835.1#75|Chroococcidiopsis sp CCALA 051
WP_106166864.1#78|Chroococcidiopsis cubana SAG 39 79
WP_146138127.1#56|Chamaesiphon polymorphus CCALA 037
WP_041548390.1#72|Chamaesiphon minutus PCC 6605
WP_015202707.1#83|Crinalium epipsammum PCC 9333
WP_168635769.1#89|Dolichospermum flos aquae UHCC 0037
WP_143467728.1#96|Leptolyngbya ohadii IS1
WP_110987556.1#80|Acaryochloris sp RCC1774
WP_148215976.1#82|Acaryochloris marina MBIC11017
WP_029315884.1#84|Acaryochloris sp CCMEE 5410
WP_132867390.1#25|Scytonema millei VB511283
WP_106546814.1#29|Chroococcidiopsis sp CCALA 051
WP_071925466.1#22|Chroococcidiopsis thermalis PCC 7203
WP_106218940.1#76|Cyanosarcina cf burmensis CCALA 770
WP_106166704.1#23|Chroococcidiopsis cubana CCALA 043
WP_158631736.1#24|Chroococcidiopsis cubana SAG 39 79

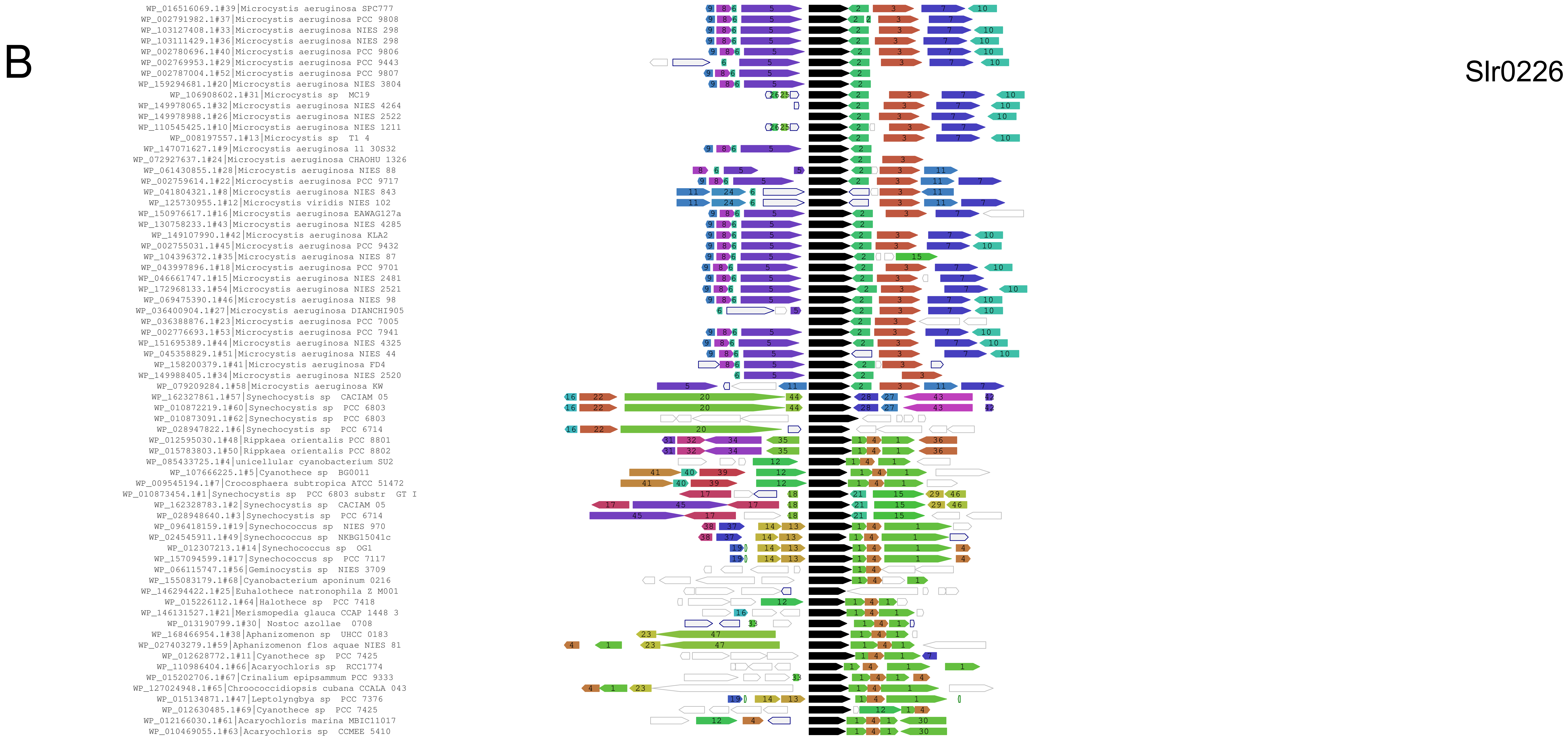
B
Slr0226
WP_016516069.1#39|Microcystis aeruginosa SPC777
WP_002791982.1#37|Microcystis aeruginosa PCC 9808
WP_103127408.1#33|Microcystis aeruginosa NIES 298
WP_103111429.1#36|Microcystis aeruginosa NIES 298
WP_002780696.1#40|Microcystis aeruginosa PCC 9806
WP_002769953.1#29|Microcystis aeruginosa PCC 9443
WP_002787004.1#52|Microcystis aeruginosa PCC 9807
WP_159294681.1#20|Microcystis aeruginosa NIES 3804
WP_106908602.1#31|Microcystis sp MC19
WP_149978065.1#32|Microcystis aeruginosa NIES 4264
WP_149978988.1#26|Microcystis aeruginosa NIES 2522
WP_110545425.1#10|Microcystis aeruginosa NIES 1211
WP_008197557.1#13|Microcystis sp T1 4
WP_147071627.1#9|Microcystis aeruginosa 11 30S32
WP_072927637.1#24|Microcystis aeruginosa CHAOHU 1326
WP_061430855.1#28|Microcystis aeruginosa NIES 88
WP_002759614.1#22|Microcystis aeruginosa PCC 9717
WP_041804321.1#8|Microcystis aeruginosa NIES 843
WP_125730955.1#12|Microcystis viridis NIES 102
WP_150976617.1#16|Microcystis aeruginosa EAWAG127a
WP_130758233.1#43|Microcystis aeruginosa NIES 4285
WP_149107990.1#42|Microcystis aeruginosa KLA2
WP_002755031.1#45|Microcystis aeruginosa PCC 9432
WP_104396372.1#35|Microcystis aeruginosa NIES 87
WP_043997896.1#18|Microcystis aeruginosa PCC 9701
WP_046661747.1#15|Microcystis aeruginosa NIES 2481
WP_172968133.1#54|Microcystis aeruginosa NIES 2521
WP_069475390.1#46|Microcystis aeruginosa NIES 98
WP_036400904.1#27|Microcystis aeruginosa DIANCHI905
WP_036388876.1#23|Microcystis aeruginosa PCC 7005
WP_002776693.1#53|Microcystis aeruginosa PCC 7941
WP_151695389.1#44|Microcystis aeruginosa NIES 4325
WP_045358829.1#51|Microcystis aeruginosa NIES 44
WP_158200379.1#41|Microcystis aeruginosa FD4
WP_149988405.1#34|Microcystis aeruginosa NIES 2520
WP_079209284.1#58|Microcystis aeruginosa KW
WP_162327861.1#57|Synechocystis sp CACIAM 05
WP_010872219.1#60|Synechocystis sp PCC 6803
WP_010873091.1#62|Synechocystis sp PCC 6803
WP_028947822.1#6|Synechocystis sp PCC 6714
WP_012595030.1#48|Rippkaea orientalis PCC 8801
WP_015783803.1#50|Rippkaea orientalis PCC 8802
WP_085433725.1#4|unicellular cyanobacterium SU2
WP_107666225.1#5|Cyanothece sp BG0011
WP_009545194.1#7|Crocosphaera subtropica ATCC 51472
WP_010873454.1#1|Synechocystis sp PCC 6803 substr GT I
WP_162328783.1#2|Synechocystis sp CACIAM 05
WP_028948640.1#3|Synechocystis sp PCC 6714
WP_096418159.1#19|Synechococcus sp NIES 970
WP_024545911.1#49|Synechococcus sp NKBG15041c
WP_012307213.1#14|Synechococcus sp OG1
WP_157094599.1#17|Synechococcus sp PCC 7117
WP_066115747.1#56|Geminocystis sp NIES 3709
WP_155083179.1#68|Cyanobacterium aponinum 0216
WP_146294422.1#25|Euhalothece natronophila Z M001
WP_015226112.1#64|Halothece sp PCC 7418
WP_146131527.1#21|Merismopedia glauca CCAP 1448 3
WP_013190799.1#30| Nostoc azollae 0708
WP_168466954.1#38|Aphanizomenon sp UHCC 0183
WP_027403279.1#59|Aphanizomenon flos aquae NIES 81
WP_012628772.1#11|Cyanothece sp PCC 7425
WP_110986404.1#66|Acaryochloris sp RCC1774
WP_015202706.1#67|Crinalium epipsammum PCC 9333
WP_127024948.1#65|Chroococcidiopsis cubana CCALA 043
WP_015134871.1#47|Leptolyngbya sp PCC 7376
WP_012630485.1#69|Cyanothece sp PCC 7425
WP_012166030.1#61|Acaryochloris marina MBIC11017
WP_010469055.1#63|Acaryochloris sp CCMEE 5410

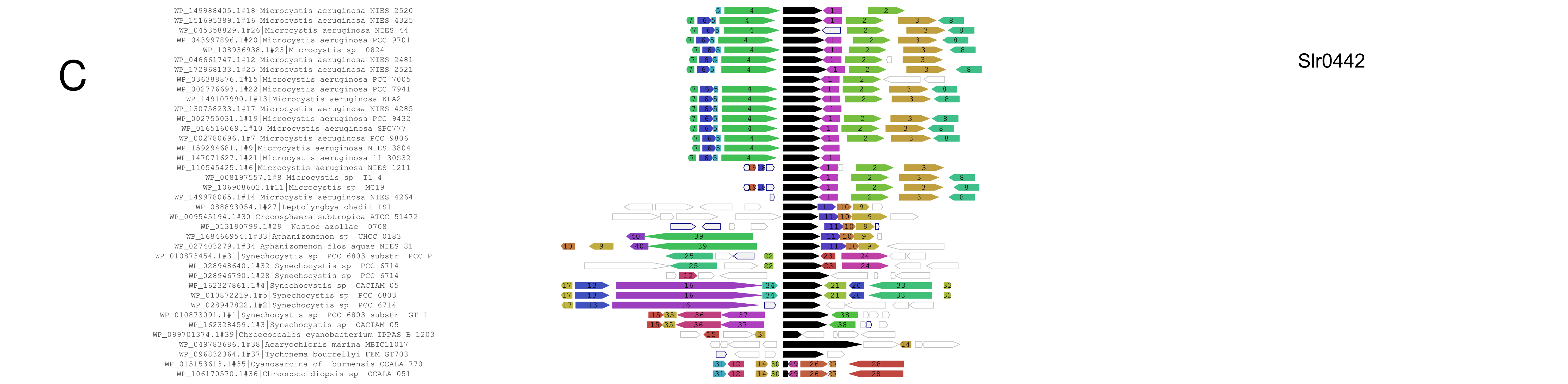
C
Slr0442
WP_149988405.1#18|Microcystis aeruginosa NIES 2520
WP_151695389.1#16|Microcystis aeruginosa NIES 4325
WP_045358829.1#26|Microcystis aeruginosa NIES 44
WP_043997896.1#20|Microcystis aeruginosa PCC 9701
WP_108936938.1#23|Microcystis sp 0824
WP_046661747.1#12|Microcystis aeruginosa NIES 2481
WP_172968133.1#25|Microcystis aeruginosa NIES 2521
WP_036388876.1#15|Microcystis aeruginosa PCC 7005
WP_002776693.1#22|Microcystis aeruginosa PCC 7941
WP_149107990.1#13|Microcystis aeruginosa KLA2
WP_130758233.1#17|Microcystis aeruginosa NIES 4285
WP_002755031.1#19|Microcystis aeruginosa PCC 9432
WP_016516069.1#10|Microcystis aeruginosa SPC777
WP_002780696.1#7|Microcystis aeruginosa PCC 9806
WP_159294681.1#9|Microcystis aeruginosa NIES 3804
WP_147071627.1#21|Microcystis aeruginosa 11 30S32
WP_110545425.1#6|Microcystis aeruginosa NIES 1211
WP_008197557.1#8|Microcystis sp T1 4
WP_106908602.1#11|Microcystis sp MC19
WP_149978065.1#14|Microcystis aeruginosa NIES 4264
WP_088893054.1#27|Leptolyngbya ohadii IS1
WP_009545194.1#30|Crocosphaera subtropica ATCC 51472
WP_013190799.1#29| Nostoc azollae 0708
WP_168466954.1#33|Aphanizomenon sp UHCC 0183
WP_027403279.1#34|Aphanizomenon flos aquae NIES 81
WP_010873454.1#31|Synechocystis sp PCC 6803 substr PCC P
WP_028948640.1#32|Synechocystis sp PCC 6714
WP_028946790.1#28|Synechocystis sp PCC 6714
WP_162327861.1#4|Synechocystis sp CACIAM 05
WP_010872219.1#5|Synechocystis sp PCC 6803
WP_028947822.1#2|Synechocystis sp PCC 6714
WP_010873091.1#1|Synechocystis sp PCC 6803 substr GT I
WP_162328459.1#3|Synechocystis sp CACIAM 05
WP_099701374.1#39|Chroococcales cyanobacterium IPPAS B 1203
WP_049783686.1#38|Acaryochloris marina MBIC11017
WP_096832364.1#37|Tychonema bourrellyi FEM GT703
WP_015153613.1#35|Cyanosarcina cf burmensis CCALA 770
WP_106170570.1#36|Chroococcidiopsis sp CCALA 051

D

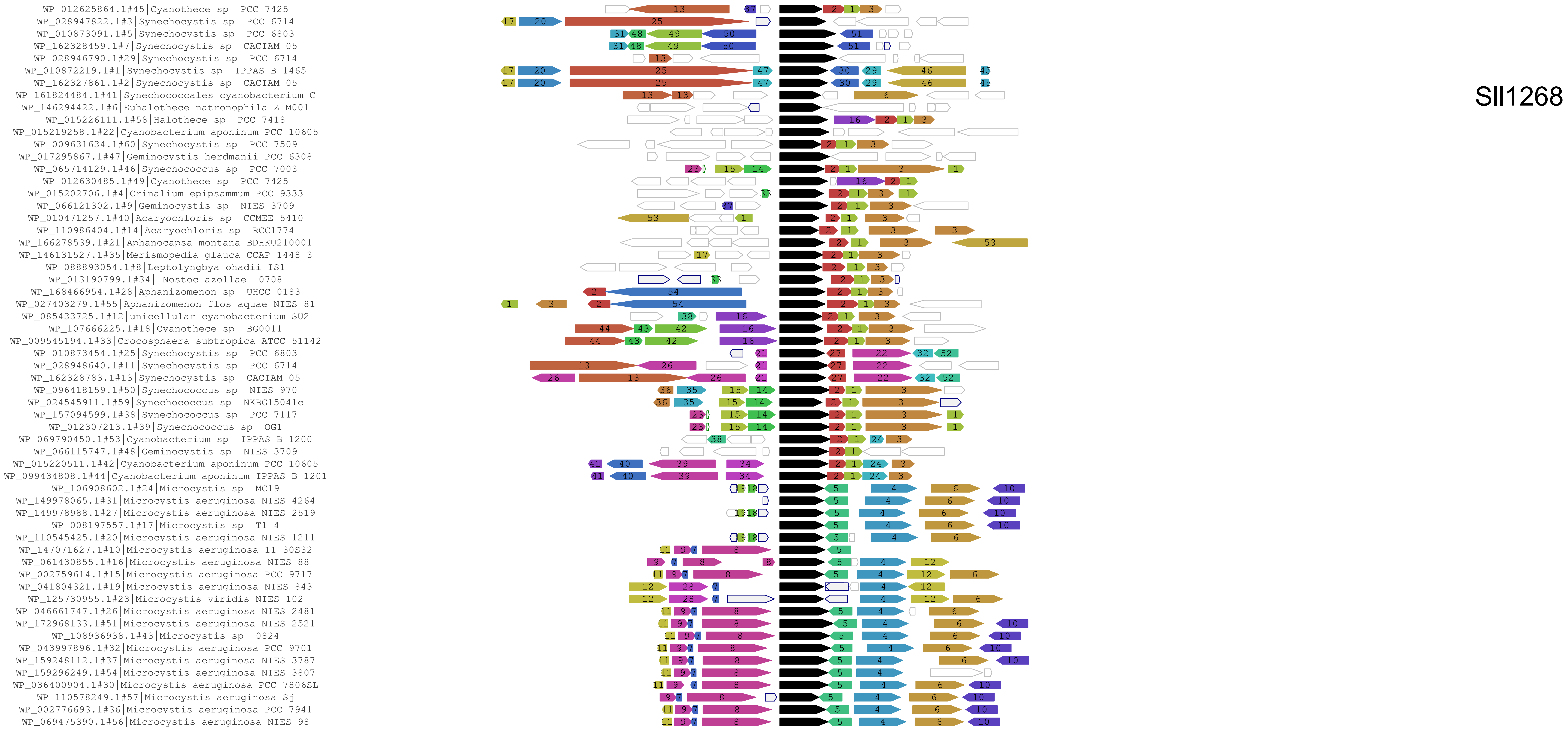
Sll1268
WP_012625864.1#45|Cyanothece sp PCC 7425
WP_028947822.1#3|Synechocystis sp PCC 6714
WP_010873091.1#5|Synechocystis sp PCC 6803
WP_162328459.1#7|Synechocystis sp CACIAM 05
WP_028946790.1#29|Synechocystis sp PCC 6714
WP_010872219.1#1|Synechocystis sp IPPAS B 1465
WP_162327861.1#2|Synechocystis sp CACIAM 05
WP_161824484.1#41|Synechococcales cyanobacterium C
WP_146294422.1#6|Euhalothece natronophila Z M001
WP_015226111.1#58|Halothece sp PCC 7418
WP_015219258.1#22|Cyanobacterium aponinum PCC 10605
WP_009631634.1#60|Synechocystis sp PCC 7509
WP_017295867.1#47|Geminocystis herdmanii PCC 6308
WP_065714129.1#46|Synechococcus sp PCC 7003
WP_012630485.1#49|Cyanothece sp PCC 7425
WP_015202706.1#4|Crinalium epipsammum PCC 9333
WP_066121302.1#9|Geminocystis sp NIES 3709
WP_010471257.1#40|Acaryochloris sp CCMEE 5410
WP_110986404.1#14|Acaryochloris sp RCC1774
WP_166278539.1#21|Aphanocapsa montana BDHKU210001
WP_146131527.1#35|Merismopedia glauca CCAP 1448 3
WP_088893054.1#8|Leptolyngbya ohadii IS1
WP_013190799.1#34| Nostoc azollae 0708
WP_168466954.1#28|Aphanizomenon sp UHCC 0183
WP_027403279.1#55|Aphanizomenon flos aquae NIES 81
WP_085433725.1#12|unicellular cyanobacterium SU2
WP_107666225.1#18|Cyanothece sp BG0011
WP_009545194.1#33|Crocosphaera subtropica ATCC 51142
WP_010873454.1#25|Synechocystis sp PCC 6803
WP_028948640.1#11|Synechocystis sp PCC 6714
WP_162328783.1#13|Synechocystis sp CACIAM 05
WP_096418159.1#50|Synechococcus sp NIES 970
WP_024545911.1#59|Synechococcus sp NKBG15041c
WP_157094599.1#38|Synechococcus sp PCC 7117
WP_012307213.1#39|Synechococcus sp OG1
WP_069790450.1#53|Cyanobacterium sp IPPAS B 1200
WP_066115747.1#48|Geminocystis sp NIES 3709
WP_015220511.1#42|Cyanobacterium aponinum PCC 10605
WP_099434808.1#44|Cyanobacterium aponinum IPPAS B 1201
WP_106908602.1#24|Microcystis sp MC19
WP_149978065.1#31|Microcystis aeruginosa NIES 4264
WP_149978988.1#27|Microcystis aeruginosa NIES 2519
WP_008197557.1#17|Microcystis sp T1 4
WP_110545425.1#20|Microcystis aeruginosa NIES 1211
WP_147071627.1#10|Microcystis aeruginosa 11 30S32
WP_061430855.1#16|Microcystis aeruginosa NIES 88
WP_002759614.1#15|Microcystis aeruginosa PCC 9717
WP_041804321.1#19|Microcystis aeruginosa NIES 843
WP_125730955.1#23|Microcystis viridis NIES 102
WP_046661747.1#26|Microcystis aeruginosa NIES 2481
WP_172968133.1#51|Microcystis aeruginosa NIES 2521
WP_108936938.1#43|Microcystis sp 0824
WP_043997896.1#32|Microcystis aeruginosa PCC 9701
WP_159248112.1#37|Microcystis aeruginosa NIES 3787
WP_159296249.1#54|Microcystis aeruginosa NIES 3807
WP_036400904.1#30|Microcystis aeruginosa PCC 7806SL
WP_110578249.1#57|Microcystis aeruginosa Sj
WP_002776693.1#36|Microcystis aeruginosa PCC 7941
WP_069475390.1#56|Microcystis aeruginosa NIES 98

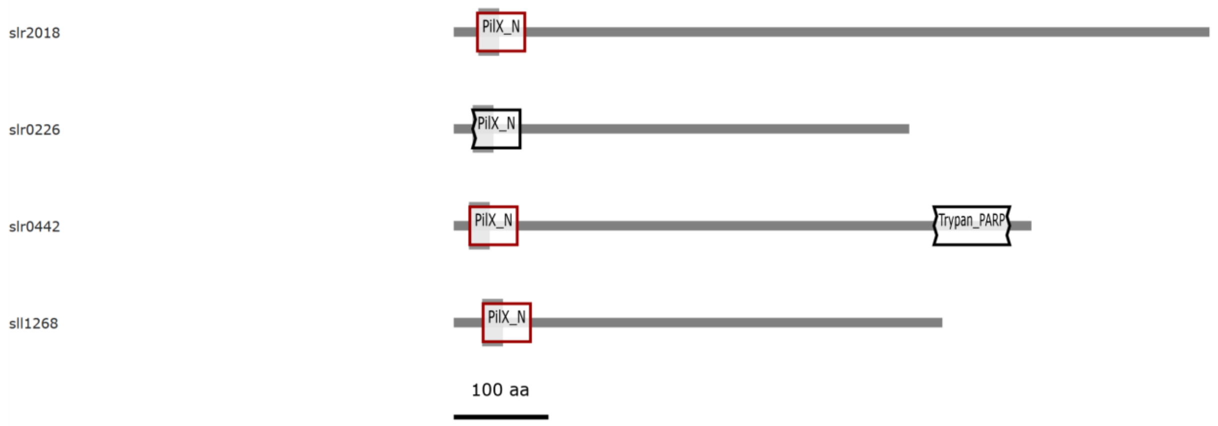


Figure S5: **Conserved Protein Domains in PilX homologs.** Using the bioinformatic online tool CDvist (Table S6) we detected an N-terminal PilX domain (PilX_N). HHsearch probability >95% (red boxes) and 100% (black box).

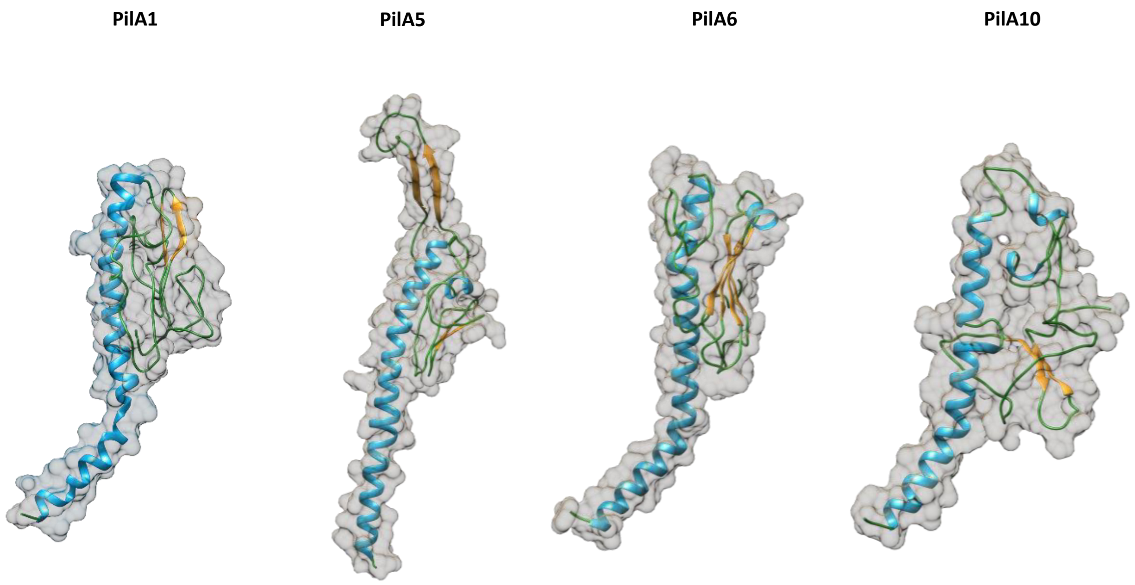


Figure S6: **Predicted three dimensional models of pilins.** Models of the mature major pilin PilA1 and the mature minor pilins PilA5, PilA6 and PilA10 were calculated with Phyre2 and visualized with Chimera (Table S6). The other minor pilins PilA9, PilA11, Slr2018, Slr0226, Slr2442 and Sll1268 showed too low homology to known tertiary structures, thus, only the helix depicted convincing conservation and the predicted models are not representative and therefore not shown here.

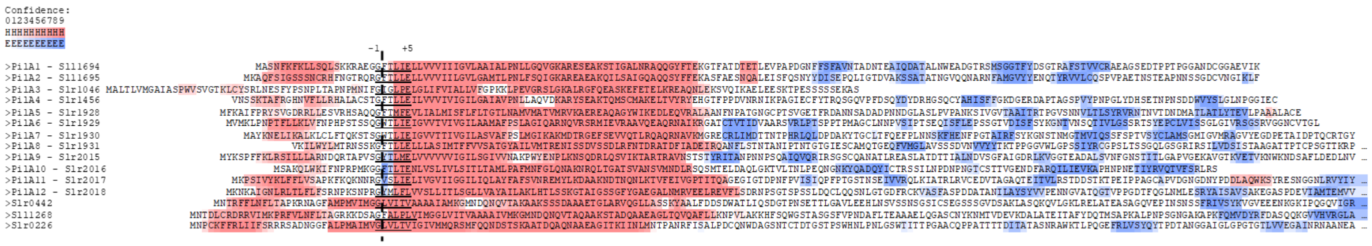

Figure S7: **Secundary structure predictions of major and minor Pilins of *Synechocystis*.** Secondary structure prediction of pilins with Ali2D. α-helices are highlighted in red and β-sheets in blue. Proposed minor pilins PilA12, Slr0442, sll1268 and Slr0226 have more predicted β-sheets further downstream, which are not shown here. The potential cleavage site by PilD is marked with the dotted line and an arrow. PilA3 was previously shown to be no minor pilin (see introduction).
