## Supplementary figures and images for "Minor pilin genes are involved in motility and natural competence in *Synechocystis* sp. PCC 6803"

### Video S1

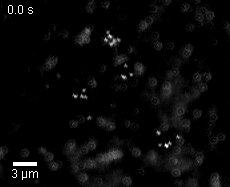

### Video S2

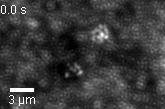

### Video S3

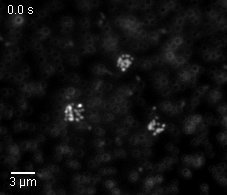
